# AI-Supported Extraction of Functional Tissue Unit Properties for Human Reference Atlas Construction

**DOI:** 10.1101/2025.09.11.675730

**Authors:** Yongxin Kong, Katy Börner

## Abstract

The Human Reference Atlas (HRA) effort brings together experts from more than 25 international consortia to capture the multiscale organization of the human body—from large anatomical organ systems (macro) to the single-cell level (micro). Functional tissue units (FTU, meso) in 10 organs have been detailed and 2D illustrations have been created by experts. Comprehensive FTU property characterization is essential for the HRA, but manual review of the vast number of scholarly publications is impractical. Here, we introduce Large-Model Retrieval-Augmented Generation for HRA FTUs (**HRAftu-LM-RAG**), an AI-driven framework for scalable and automated extraction of FTU-relevant properties from scholarly publications. This validated framework integrates Large Language Models for textual reasoning, Large Vision Models for visual interpretation, and Retrieval Augmented Generation for knowledge grounding, offering a balanced trade-off between accuracy and processing efficiency. We retrieved 244,640 PubMed Central publications containing 1,389,168 figures for 22 FTUs and identified 617,237 figures with microscopy and schematic images. From these images and associated text, we automatically extracted 331,189 scale bars and 1,719,138 biological entity mentions, along with donor metadata such as sex and age. The resulting data facilitates the design, review, and approval of future FTU illustrations during HRA construction.

## Introduction

The Human Reference Atlas (HRA) v2.2 captures 4,694 unique anatomical structures, 1,288 unique cell types, and 2,018 unique biomarkers—all linked to existing ontologies—for the healthy male and female bodies^1^. Two-dimensional illustrations exist for 22 functional tissue units^2^ (FTUs) in 10 organs that have 14 anatomical structures and 116 unique cell types. FTUs cover the structure and function of multiple cells that together deliver a specific vital function and they are detailed in ASCT+B tables, see standard operating procedure (SOP) entitled *Authoring ASCT+B Tables*^3^. FTU illustrations are designed according to existing SOPs^4–6^ and a Style Guide^7^. They are reviewed and validated by organ experts published via HRA releases and used in different HRA user interfaces. A key step during FTU design is the identification of relevant publications that provide experimental evidence on the size, shape, and spatial composition of FTUs for donor groups with different demographics (e.g., sex, age).

The manual compilation of relevant publications and the extraction of FTU relevant properties (e.g., size changes as we age) is time consuming and error prone. For example, a targeted literature search using Uberon^8^ names for the 22 FTUs in Open Access PubMed Central (PMC; https://pmc.ncbi.nlm.nih.gov)—a free full-text archive of biomedical and life-sciences journal literature provided by the U.S. National Institutes of Health’s National Library of Medicine (NIH/NLM)—yielded 244,640 publications licensed under Creative Commons, published between 1783 and 2025. These publications collectively feature 1,389,168 figures potentially relevant for designing, reviewing, and improving FTU illustrations. A manual inspection of all publications and figures (assuming one hour per paper and 40 hours per week) would take 117 years of dedicated effort by a single expert, which is impractical. Recent work highlights the importance of hierarchically nested FTUs^9^ which drastically increases the number of FTUs—making the current manual process of FTU data evidence collection a bottleneck. Note that the current FTU design and review process does not capture changes in the size or spatial composition of FTUs due to donor demographics (e.g., sex, age, body-mass index) or disease (e.g., tumor invasion or fibrosis).

Alternative approaches for determining the size and shape of FTUs in relation to donor demographics have been tried. For example, experimental data can be used to manually determine key properties of FTUs. Our team ran two Kaggle competitions^10,11^ to obtain FTU segmentation code for five organs using 7,102 two-dimensional (2D) tissue sections for kidney in the first, and 12,901 2D tissue for kidney, large intestine, lung, prostate, and spleen in the second competition; results help identify the number of FTUs per unit area and the diameter and average area size distribution of FTUs plus their shape and inner structure—when cut at arbitrary angles. Another approach is to compile relevant information from biomedical textbooks (e.g., on human pathology or physiology). However, textbooks are typically proprietary, do not come with an API that supports automatic retrieval of relevant FTU properties, and they do not properly capture the multiscale nature of FTUs with changes in size, single cell composition, or other properties due to demographics or disease.

Recent advances in artificial intelligence (AI), specifically Large Language Models (LLMs)^12,13^ and Large Vision Models (LVMs)^14,15^, have significantly expanded the ability of machines to understand, reason about, and generate human-like text and image content. LLMs are primarily designed to process vast amounts of text and to discern patterns and relationships within textual information. General-purpose LLMs (e.g., Llama 3^16^, Qwen^17^, and Phi-4-Mini^18^) support multiple languages and downstream tasks. Domain-specific biomedical LLMs have been developed (e.g., BioBERT^19^, BioGPT^20^, BioMedLM^21^, GatorTron^22^, BioMistral^23^ and GatorTronGPT^24^) and are pre-trained on biomedical literature, clinical notes, and ontologies to improve performance on tasks such as named entity recognition, relation extraction, and clinical document summarization. Meanwhile, LVMs^14,15^ are capable of interpreting complex visual content and integrating it with textual inputs. General-purpose LVMs include Vision Transformer (ViT)^14^, CLIP^25^, Segment Anything Model (SAM)^26^, BLIP^27^, and InternVL3^28^; domain-specific biomedical LVMs include LVM-Med^29^ and MedCLIP^30^. These LVMs are typically built on transformer architectures^12^ and are trained using large-scale image-text pairs or self-supervised strategies, incorporating cross-modal alignment, attention-based encoding, and prompt-driven inference to capture semantic and spatial relationships in visual data. Furthermore, general-purpose multimodal models (e.g., GPT-4^31^ and Gemini^32^) and domain-specific biomedical multimodal models (e.g., MedGemma^33^) represent a growing trend toward multimodal AI systems that integrate the capabilities of both LLMs and LVMs, enabling more comprehensive multimodal understanding and reasoning.

While powerful, standalone LLMs can sometimes produce inaccurate or ‘hallucinated’ information due to their reliance solely on pre-trained knowledge, which may be outdated, incomplete, or less relevant for a task. To mitigate these limitations and enhance factual accuracy and relevance, Retrieval Augmented Generation (RAG) techniques^34^ have been developed. RAG models (e.g., REALM^35^, Dense Passage Retrieval^36^, FLARE^37^and Atlas^38^) improve the reliability of generative LLMs by retrieving relevant information from an external, up-to-date knowledge base and providing it as context to the language model before generating a response. Similarly, LVMs, while strong in visual understanding, often benefit from external textual context for deeper, fact-grounded reasoning that extends beyond immediate visual cues. By strategically leveraging LLMs for textual reasoning, LVMs for visual interpretation, and RAG for grounding outputs in verifiable external knowledge, it becomes feasible to use the models to extract FTU properties from text and figures in thousands of relevant publications.

Here, we introduce an AI-driven framework, HRAftu-LM-RAG, which combines LVMs, LLMs, and RAGs to automate the systematic extraction of FTU-relevant properties from scholarly publications, including figures, captions, and associated reference text. The HRAftu-LM-RAG implementation provides a scalable and robust solution to accelerate the systematic computation of multiscale data evidence for the HRA across diverse donor demographics. In the future, we plan to use the same framework to retrieve publication data evidence for other anatomical structures and their physiological functions.

## Results

This section details data and code used together with performance results.

### Data

HRA construction requires information on FTU properties such as size, shape, and spatial composition of FTUs for different demographic groups (male/female, age, etc.). To achieve this, we extract four types of data from scholarly publications via LVM-based workflows for image-type categorization and in-image text-term extraction, followed by an LLM-RAG-based workflow to extract FTU properties—scale bars, donor metadata, and biological structures—from figure captions, reference text, and paper abstracts. Specifically, we extract:

1. **Image type**: Microscopy and schematic images categorized based on their visual content, figure captions, and reference text. Each image is linked to metadata including paper ID, figure ID, image category, caption, reference text, and file path. Heterogeneous composite figures were assigned to multiple categories.
2. **Scale bars**: Scale bar values are obtained from in-image text terms, figure captions, and reference text of selected microscopy images. Each record includes the paper ID, figure ID, numeric value, and unit (e.g., µm, mm).
3. **Donor metadata**: Species, sex, age, and BMI are extracted from in-image text terms, figure captions, and the publication abstract of both image categories (microscopy and schematic). Each donor record includes a paper ID, extraction location, species, developmental stage (prenatal and postnatal), sex, BMI, and age in years.
4. **Biological structures (AS+CT+B)**: Anatomical structures (AS), cell types (CT), and biomarkers (B)^1,39^ are extracted from FTU relevant in-image text terms and figure captions of both image categories (microscopy and schematic). Each AS, CT, and B record includes the paper ID, extraction location, and entity term.

All data is stored in a relational database with eight interlinked data tables (see the entity relationship diagram in **Fig. 1** and table description in **Supplementary Table 1**) and has been deposited on Zenodo^40^ (https://doi.org/10.5281/zenodo.17051026).

**Figure 1.**
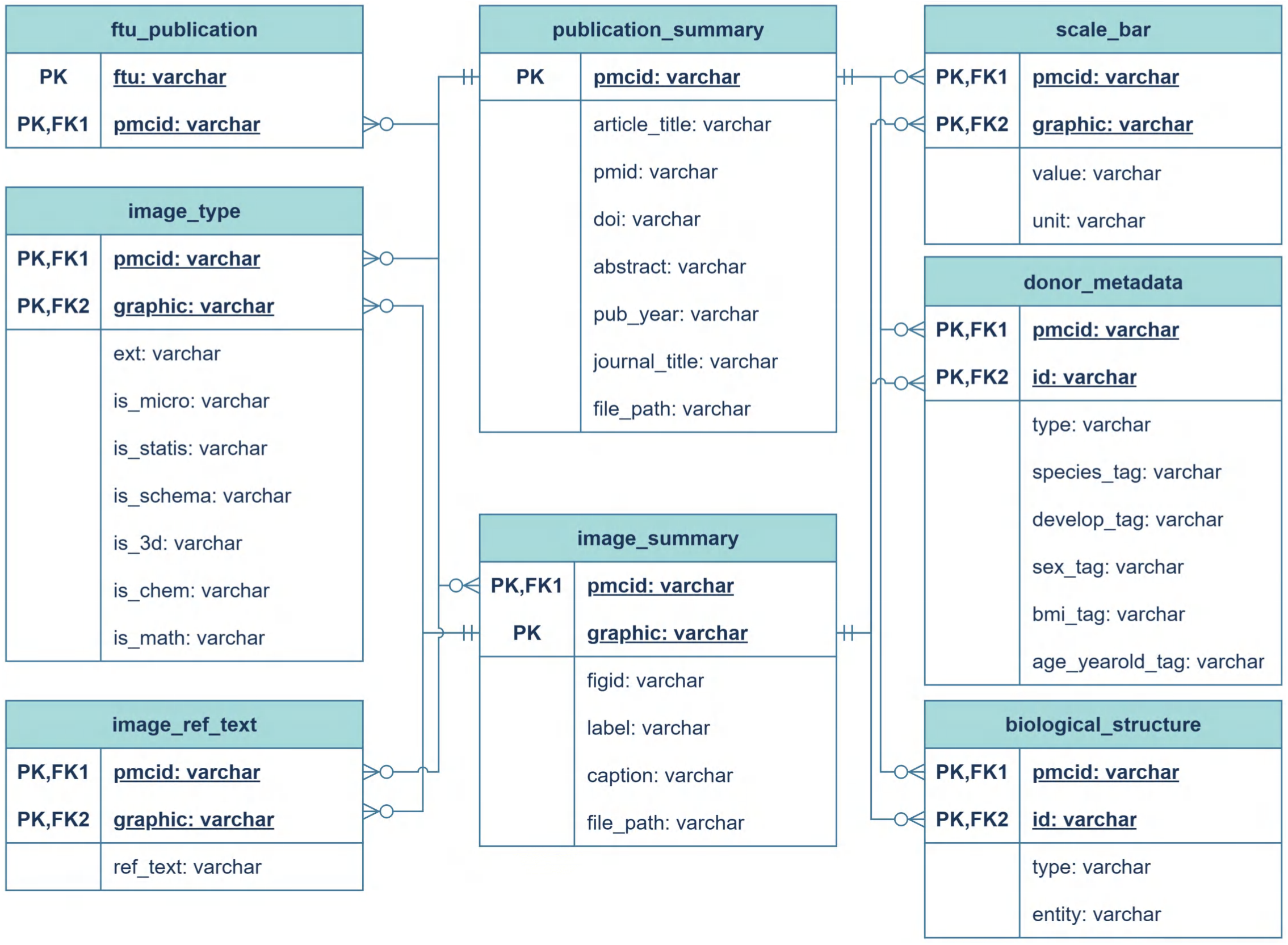
Relational schema of HRA-FTU properties produced by the HRAftu-LM-RAG framework. Eight linked tables capture publication and image metadata, image categories (including is_micro and is_schema), scale bars, donor metadata, biological structure (AS+CT+B) and FTU–publication mappings. Primary keys (PK) and foreign keys (FK) define table joins; arrows use crow’s-foot notation to denote one-to-many relationships.

### HRAftu-LM-RAG Overview

In order to parse the FTU properties detailed in the previous section from scholarly papers, the following five workflows are used:

- **Workflow 1 (WF1): Image-type categorization.** Apply LVM to images and LLM-RAG to associated figure captions and reference text in order to categorize figures into six categories (i.e., statistical image, microscopy image, schematic image, 3D structure diagram, chemical structure diagram, and mathematical expression). Only microscopy and schematic images are selected as FTU-morphology relevant images for downstream analysis.
- **Workflow 2 (WF2): In-image text-term extraction.** Use LVM to capture all textual labels embedded in microscopy and schematic images (‘in-image text terms’), which serve as primary cues for image annotation.
- **Workflow 3 (WF3): Scale-bar extraction.** Use LLM-RAG to parse in-image text terms, figure captions, and reference text of selected microscopy images, to extract microscopy acquisition parameters (e.g., magnification, scale-bar length, voxel/pixel size, section thickness and scan area), then isolate and standardize scale bar measurements for FTU size prediction.
- **Workflow 4 (WF4): Donor-metadata extraction.** Extract species, sex, age, and BMI information by applying an LLM-RAG to in-image text terms, figure captions, and publication abstracts of microscopy and schematic images. Subsequently, normalize data to generate standardized donor metadata.
- **Workflow 5 (WF5): AS+CT+B extraction**: Extract biological entity terms—anatomical structures (ASs), cell types (CTs), and biomarkers (Bs)—by applying an LLM-RAG to in-image text terms and figure captions of microscopy and schematic images.

The HRAftu-LM-RAG framework is organized around these five core workflows, integrating RAG, LLMs and LVMs. The HRAftu-LM-RAG data flow—from raw publications to FTU-related information output via WF1-WF5 in yellow box—is shown in **Fig. 2**. The framework integrates structured text and visual data from PMC publications (see details in **Methods** section) to perform automated identification and extraction of FTU property at scale, including image type, scale bar, donor metadata and AS+CT+B.

**Figure 2.**
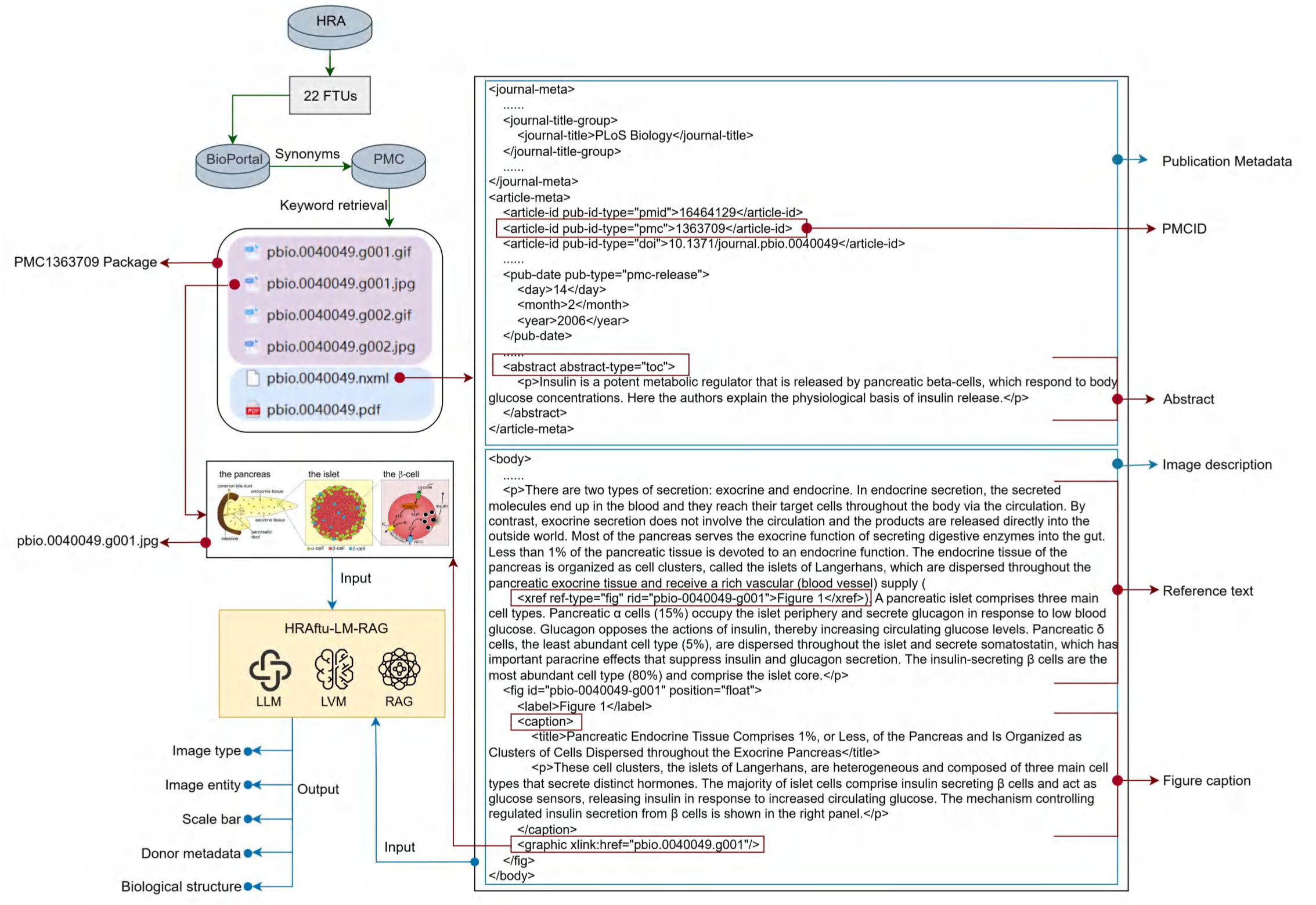
Overview of the HRAftu-LM-RAG data processing framework. 22 FTUs from HRA v2.2 were mapped via corresponding 22 Uberon IDs to their 147 synonyms in BioPortal. A keyword-based search was then performed in PMC using 139 composite search expressions, each created by grouping semantically similar synonyms to reduce redundancy and improve retrieval efficiency. Each PMC paper comes with full-text encoded in National Library of Medicine eXtensible Markup Language (NXML) and image files (e.g., JPG, GIF). Using the NXML, publication metadata (e.g., PMCID, abstract) and image descriptions (figure captions, reference text) linked to their corresponding image files via <graphic> xlink references were extracted. Images and associated text are then jointly submitted to the HRAftu-LM-RAG (yellow box), which implements WF1–WF5. Output includes image-type categorization and extracted image entities, scale bars, donor metadata, and AS+CT+B. Shown here is an example from paper PMC1363709^41^ entitled “Oscillations, Intercellular Coupling, and Insulin Secretion in Pancreatic β Cells” published in *PLoS Biology* which includes a schematic image, with AS+CT+B extracted from this paper.

The construction of task-specific knowledge bases for RAG is detailed in **Table 1**. Specifically, (1) Image Type Knowledge Base (ITKB): Constructed for the image type categorization task, ITKB is built using a randomly selected collection of PMC publication packages—3,560 images from 558 papers—retrieved using the term “islet of Langerhans” as the subject. Descriptions for six image types (**Supplementary Table 2**), including content descriptions and tags, are manually curated by synthesizing key features from source images and their associated captions. Representative examples (**Supplementary Table 3**), including figure captions and categorization reasoning, are selected to capture diverse perspectives and comprehensively reflect the features of each category. (2) Scale Bar Knowledge Base (SBKB): Compiled for the scale bar extraction task, the SBKB comprises definitions, identification rules and examples of microscopy acquisition parameters from microscopy images, including magnification, scale bar, voxel size, pixel size, section thickness, and scan area (**Supplementary Table 4-6**). Descriptions of length units are also included (**Supplementary Table 7**). (3) Donor Knowledge Base (DKB): Developed for the donor metadata extraction task, the DKB contains 33 manual descriptions of donor-related units, specifically age, height, and weight (**Supplementary Table 8-10**). (4) Biological Ontology Knowledge Base (BOKB): Built for the AS+CT+B extraction task, the BOKB was constructed using ontology data sourced from the EMBL-EBI Ontology Lookup Service (https://www.ebi.ac.uk/ols4/) and downloaded on October 15, 2024 (https://ftp.ebi.ac.uk/pub/databases/spot/ols/slurm_pipeline/2024_10_1516_44/ontologies.json.gz). The BOKB comprises selected label terms, definitions, and sources for 8,581,169 ontologies from 264 ontology databases—253 with explicit names and 11 unnamed (**Supplementary Table 11**). The process of these four knowledge bases is detailed in **Supplementary Table 12**.

**Table 1.**
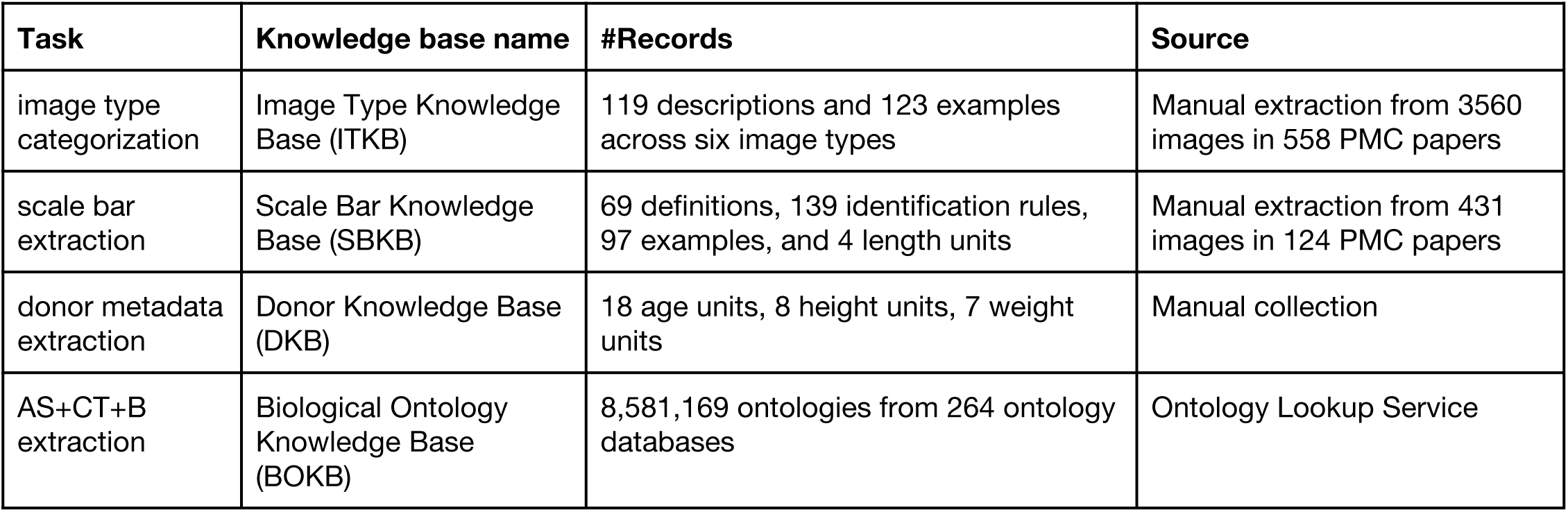
Task-specific knowledge bases for RAG.

To ensure reproducibility and scalability across the five workflows, a two-phase framework—pilot and production—is adopted. During the pilot phase, candidate model-prompt configurations for each workflow are evaluated on task-relevant samples randomly selected from PMC (construction detailed in **Supplementary Table 13**), using task-appropriate metrics—accuracy for categorization and Jaccard similarity for structured extraction or question answering—together with response time, and the configuration that best balances response quality and speed is identified (detailed in “Performance results of models on HRA-FTU tasks” in **Results** section). In the production phase, the selected model and prompt are fixed, and the workflows are executed at scale via standardized HTTP endpoints to process inputs and generate final outputs.

The workflow of image-type categorization (WF1) is illustrated in **Fig. 3**. The image-type module treats each figure as a six-category problem. To capture diverse interpretations of each figure, the image is first processed by five LVMs—LLaVa^42^, LLaMA 3.2-Vision^43^, Phi-3^44^, Phi-3.5^45^, and Pixtral-12B-2409^46^—each prompted with a unified instruction requesting both the most likely image type and a brief justification for the prediction. These LVM-generated outputs are then combined with the original figure captions and reference text from the publication package. This composite textual information serves as input for a RAG module, which performs semantic retrieval against two domain-specific knowledge bases: ITKB and BOKB. The passages retrieved from these knowledge bases are subsequently synthesized by a dedicated LLM (i.e., LLaMA 3.2^16^) to form a concise knowledge summary. Finally, this synthesized context, along with the user’s query and a system prompt, is ingested by the same LLM for categorization, yielding the predicted image type. This workflow outperformed single-model baselines and was therefore adopted as the default pipeline (evaluation results detailed in **Fig. 5**).

**Figure 3.**
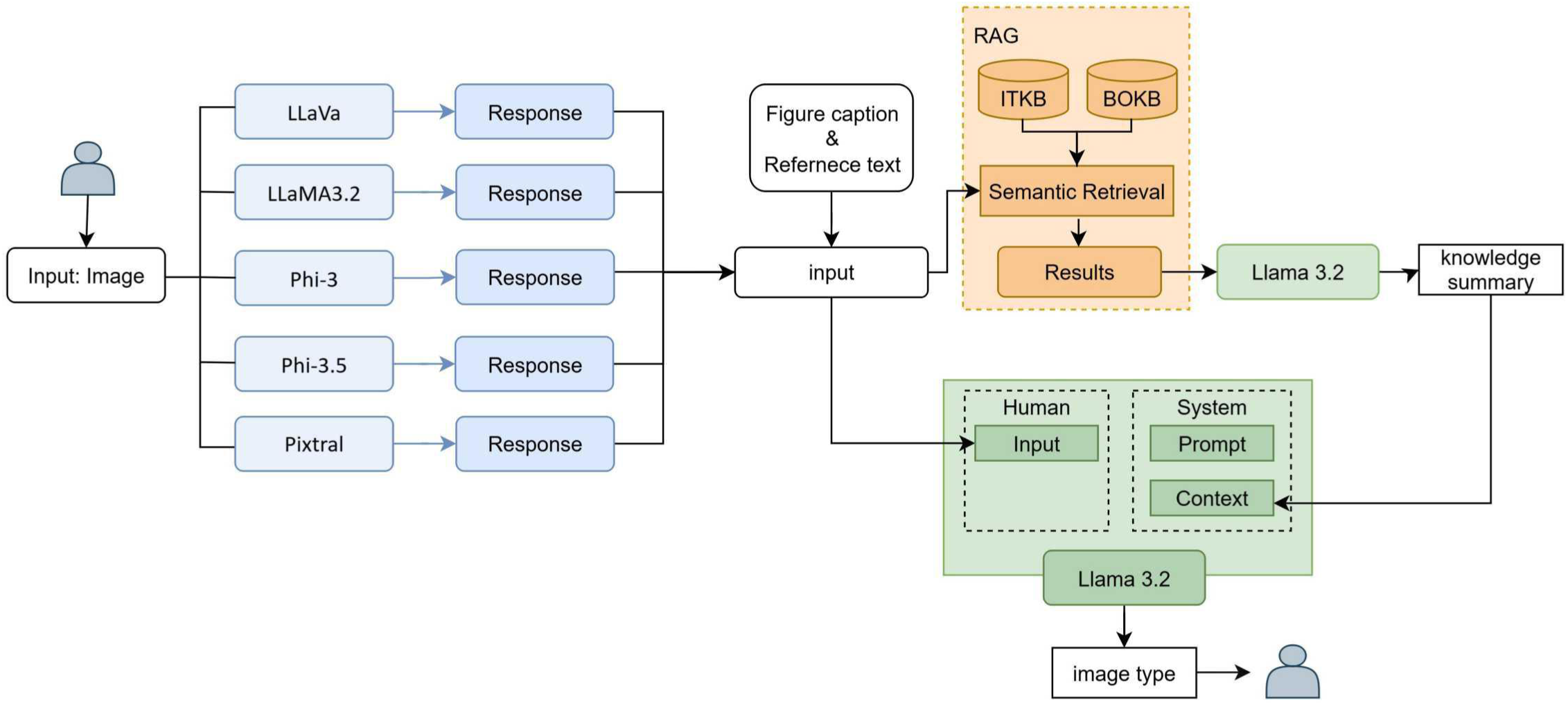
Image type categorization workflow (WF1). LVM Ensemble (blue): input images are independently processed by five LVMs (llava-v1.6-34b, Llama-3.2-11B-Vision, Phi-3-vision-128k-instruct, Phi-3.5-vision-instruct and mistralai/Pixtral-12B-2409) to produce intermediate textual responses. RAG (orange): these LVM outputs—together with the original figure caption and reference text—are used to perform semantic retrieval against two domain-specific knowledge bases (ITKB and BOKB); retrieved passages are synthesized by LLaMA 3.2 (i.e., llama3.2:latest) to generate a concise knowledge summary. LLM-Based Categorization (green): the final LLaMA 3.2 model ingests the user’s query, system prompt, and retrieved context to produce the predicted image type.

Following the categorization of relevant image types, the multimodal workflow (WF2–WF5) extracts FTU’s different properties (**Fig. 4**). In the pre-processing stage, in-image text terms are identified from microscopy and schematic images using Phi-3.5 (WF2), and combined with publication text (e.g., figure captions, reference text and abstracts) to form task-specific inputs. Specifically, scale bar extraction uses visual entities and accompanying captions and reference text from microscopy images (WF3); donor metadata extraction combines visual entities from both image types with captions (WF4); and AS+CT+B extraction incorporates visual entities from both image types with captions and publication abstracts (WF5). These multimodal inputs are embedded using the gte-large^47^ model and then subsequently employed for similarity-based retrieval from three task-specific knowledge bases (i.e., SBKB, DKB, and BOKB). The retrieved passages are then passed to LLMs for final information extraction. LLaMA3.1^16^ is used for scale bar identification, while Gemma2^48^ handles donor metadata and AS+CT+B extraction (see “Performance results of models on HRA-FTU tasks” in **Results** section).

**Figure 4.**
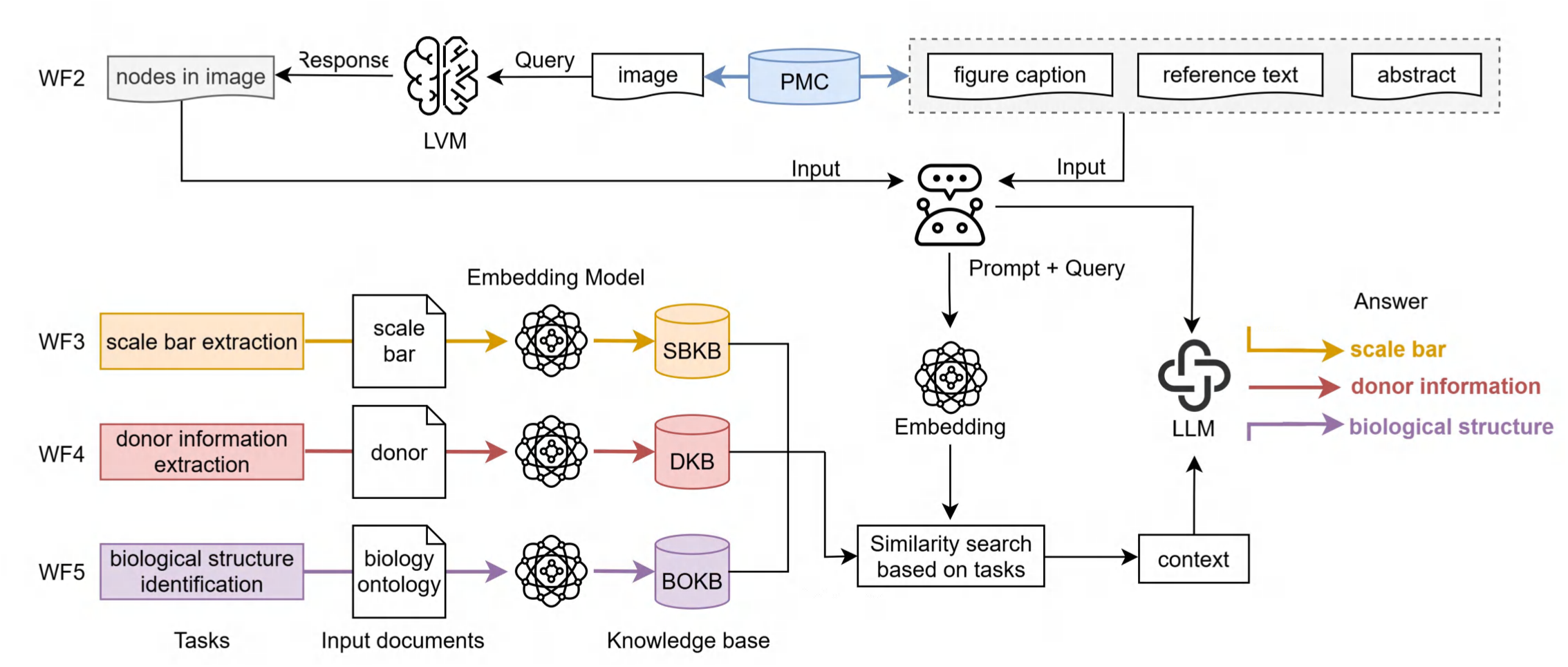
HRA-FTU related data extraction workflow (WF2-WF5). In the pre-processing stage (top), the LVM (i.e., Phi-3.5) analyzes input PMC images to identify in-image text terms (“visual entity in image”, left), while associated publication context (figure captions, reference text, and abstracts from PMC, right) is retrieved to form a comprehensive query. In the LLM-RAG stage (below), this combined input is encoded by the pretrained embedding model (i.e., gte-large) and used to perform task-specific similarity searches against three domain knowledge bases: SBKB (orange), DKB (red), and BOKB (purple). Retrieved passages are collated as context and submitted to the LLM, which generates the final extracted outputs for (1) scale bar, (2) donor metadata, and (3) AS+CT+B biology entity.

### Performance results of models on HRA-FTU tasks

To ensure accurate retrieval in the RAG pipeline, we first benchmarked four embedding models, as embedding quality is critical for retrieving relevant documents for language model prompting. A query set of 100 randomly sampled paragraphs from PubMed publications (**Supplementary Table 14**) was used to evaluate all-MiniLM-L6-v2^49^, bge-large-en-v1.5^50^, gte-large^47^, and pubmedbert-base-embeddings^21^. **Fig. 5a** presents the embedding scores for the evaluated models, with the gte-large model yielding the highest score (0.768). Therefore, gte-large was employed to embed the knowledge base for RAG retrieval in this paper.

**Figure 5.**
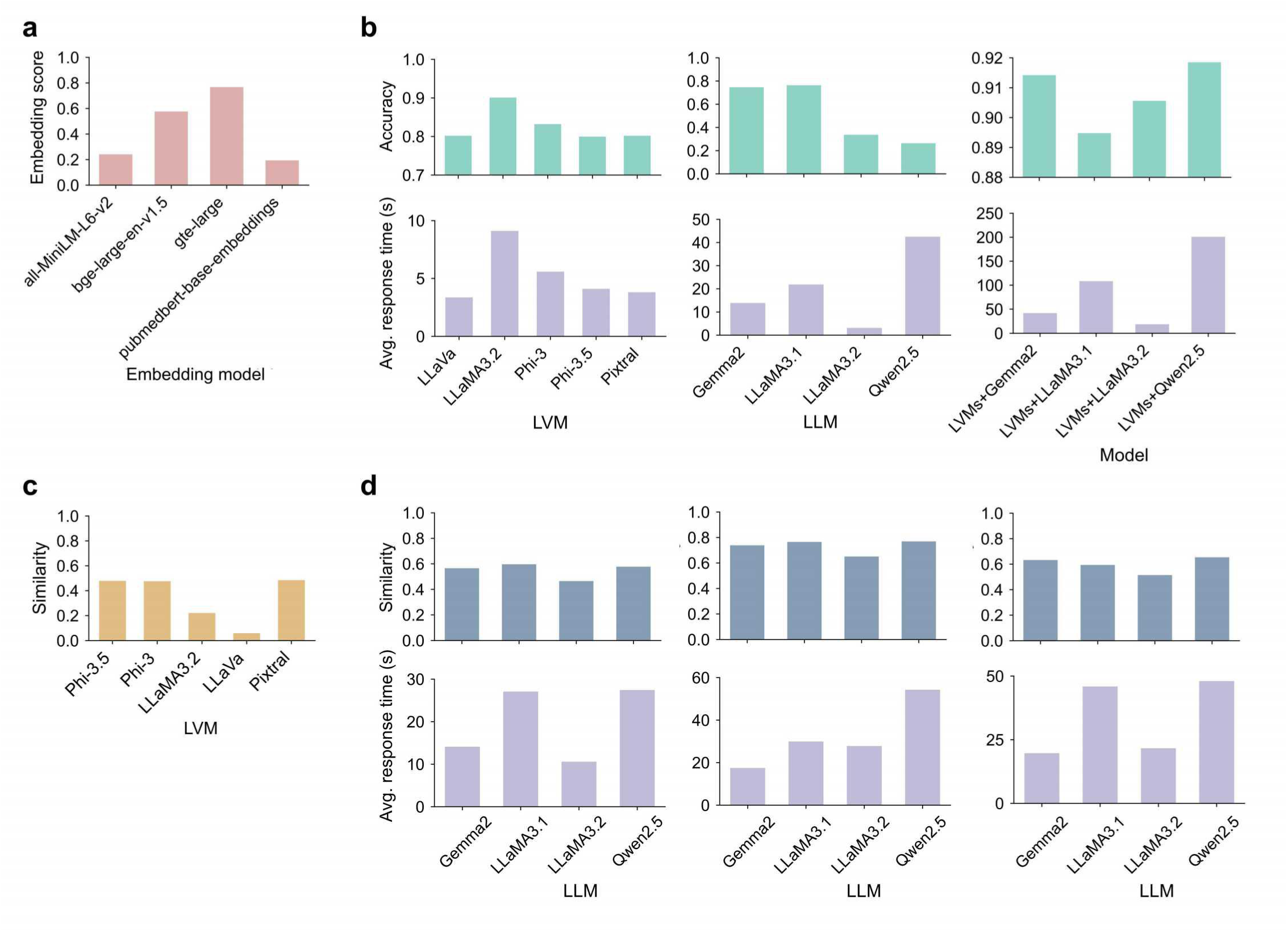
Model performance on HRA-FTU tasks. **a.** Embedding model performance. The embedding score for four selected models (all-MiniLM-L6-v2, bge-large-en-v1.5, gte-large, and pubmedbert-base-embeddings), are calculated as normalized cosine similarity with a range of 0 to 1, based on 100 query samples from PubMed. **b.** Model performance on image-type categorization. Accuracy and average response time (seconds) were measured on a test set of 567 images paired with 465 figure captions. Accuracy is defined as the number of correctly predicted image types divided by the total number of inputs, scoring each correct categorization as 1 and each incorrect categorization as 0. **c.** LVM performance on image entity extraction. Mean Jaccard similarity index between model-predicted and ground-truth entities, evaluated over 100 images with 100 prompts each. **d**. LLM performance on scale bar, donor metadata and AS+CT+B extraction. For each model, bars show the highest mean Jaccard similarity index achieved by any prompt when compared against the ground-truth annotations (top), and the corresponding mean inference time in seconds (bottom).

For WF1, we compared three approaches for image-type categorization. We evaluated these approaches on a test dataset of 567 PMC images manually annotated with image-type labels; 465 of these images included accompanying captions or reference text (see **Supplementary Table 15** and **Supplementary Data 1**).

1. **LVMs only**: Five vision models—LLaVA (34B)^42^, LLaMA3.2 (11B, LVM)^43^, Phi-3 Vision-128k-Instruct^44^, Phi-3.5 Vision-Instruct^45^, and Pixtral-12B-2409^46^—were evaluated on 567 test images. Each model received the raw image alongside a prompt defining the six target image types, and its output was compared to the manually annotated labels to compute accuracy.
2. **LLMs with RAG**: Text-based categorization was performed on the 465 test images, using four LLMs, including Gemma2 (27B)^48^, Qwen2.5 (72B)^51^, LLaMA3.1 (70B)^16^, and LLaMA3.2 (latest)^16^. This approach utilized the ITKB through RAG, taking each image’s figure caption and reference text as input.
3. **Hybrid LVM+LLM+RAG**: Outputs from these five LVMs, augmented with textual descriptions from figure captions and reference texts, were processed by four LLMs under the RAG framework and evaluated on the same 465 test images.

The test results are shown in **Fig. 5b**, where hybrid pipelines achieved the highest accuracy. LVMs-only approach outperform LLMs-with-RAG approach: among vision models, LLaMA3.2 reached the highest accuracy (0.9009), whereas the top LLM-with-RAG (LLaMA3.1) achieved only 0.7639. Hybrid LVM+LLM pipelines further improved performance. Three hybrids surpassed the best LVM-only result, with accuracies ranging from 0.9056 (LVM+LLaMA3.2) to 0.9185 (LVM+Qwen2.5). When factoring in inference speed, the LVMs+LLaMA3.2 setup offered the best trade-off among high-accuracy hybrids, recording an average response time of 18.86 seconds—considerably faster than both LVMs+Gemma2 and LVMs+Qwen2.5. Consequently, LVMs+LLaMA3.2 pipeline was chosen for all subsequent image-type categorization in this paper.

For WF2, we evaluated the performance of the same five LVMs in extracting in-image text terms, using both test data and guided prompts. The test dataset comprised 100 randomly selected images (**Supplementary Data 2**), with all in-image entities manually annotated to establish ground truth (**Supplementary Table 16**). The guided prompts for this task consisted of 100 distinct instructions (**Supplementary Table 17**). Each LVM was then evaluated by applying every prompt to each image, yielding a comprehensive matrix of model– prompt combinations. Performance was assessed quantitatively by computing the Jaccard similarity index, which measures the overlap between entities extracted by the models and the manually annotated ground truth. **Fig. 5c** presents that Pixtral (12B) achieved the highest mean Jaccard similarity of 0.485, closely followed by Phi-3.5 (4.2B) at 0.479 and Phi-3 (4.2B) at 0.476. Given Phi-3.5’s near-top performance combined with its smaller footprint and faster inference, Phi-3.5 was selected as the optimal model for image entity extraction in this study.

For WF3-WF5, the same four LLMs with RAG were evaluated for FTU-related data extraction using task-specific test data and distinct textual prompts:

1. **Scale bar extraction**: 100 microscopy figure captions from PMC (**Supplementary Table 18**), each annotated for five fields (descriptor type, value, units, notes, and panel), were paired with 98 tailored prompts (**Supplementary Table 19**).
2. **Donor metadata extraction**: 50 text segments (ten each drawn from abstracts, figure captions, figure nodes, figure reference texts, and full-text paragraphs; **Supplementary Table 20**), each annotated for six donor attributes (species, sex, age, BMI, height, and weight), were matched to 100 dedicated prompts (**Supplementary Table 21**).
3. **AS+CT+B extraction**: 60 text segments were prepared, comprising 20 segments from each of three sources—figure captions, figure nodes, and reference texts—with each source evenly split between microscopy and schematic images (10 segments each) (**Supplementary Table 22**). Each segment was annotated for relevant AS+CT+B and paired with 100 specialized prompts (**Supplementary Table 23**).

Performance on these tasks is shown in **Fig. 5d**. In scale-bar extraction, LLaMA3.1 achieved the highest mean Jaccard similarity (0.597) with an average latency of 27.09 second. For donor-metadata extraction, Qwen2.5 secured the top similarity (0.770) but incurred a latency of 54.29 second, whereas Gemma2 reached a similarity of 0.739 in just 17.52 second. In AS+CT+B extraction, Qwen2.5 recorded the highest similarity (0.654) at a latency of 48.03 second, while Gemma2 delivered a comparable similarity (0.633) with a faster mean response time of 19.71 second. Therefore, considering the balance of accuracy and efficiency, LLaMA3.1 was selected for scale-bar extraction, and Gemma2 was chosen for both donor-metadata and AS+CT+B extraction (final prompts detailed in **Supplementary Table 24**).

### FTU property across 22 FTUs using HRAftu-LM-RAG

To systematically characterize FTUs, the HRAftu-LM-RAG framework first established a comprehensive dataset from publicly available literature. A total of 244,640 PMC publications and 1,389,168 associated images were retrieved for 22 distinct FTUs across 10 organs, with the distribution for each FTU shown in **Fig. 6a**. For example, the epidermal ridge of digit from the Skin was the most extensively documented FTU, with 56,538 publications and 662,247 images, while the thymus lobule had the fewest, with 296 publications and 3,074 images. Among all retrieved images, 332,008 were microscopy images and 386,280 were schematic images, accounting for 23.89% and 27.81% of the total, respectively. The proportions of six image types across FTUs are shown in **Fig. 6b**. Specifically, the nephron and the epidermal ridge of digit had the highest number of microscopy and schematic images, with 65,341 and 99,375 images, respectively. In terms of within-FTU proportions, the renal corpuscle and the alveolus of the lung exhibited the highest shares of microscopy and schematic images, at 46.67% and 38.92%, respectively.

**Figure 6.**
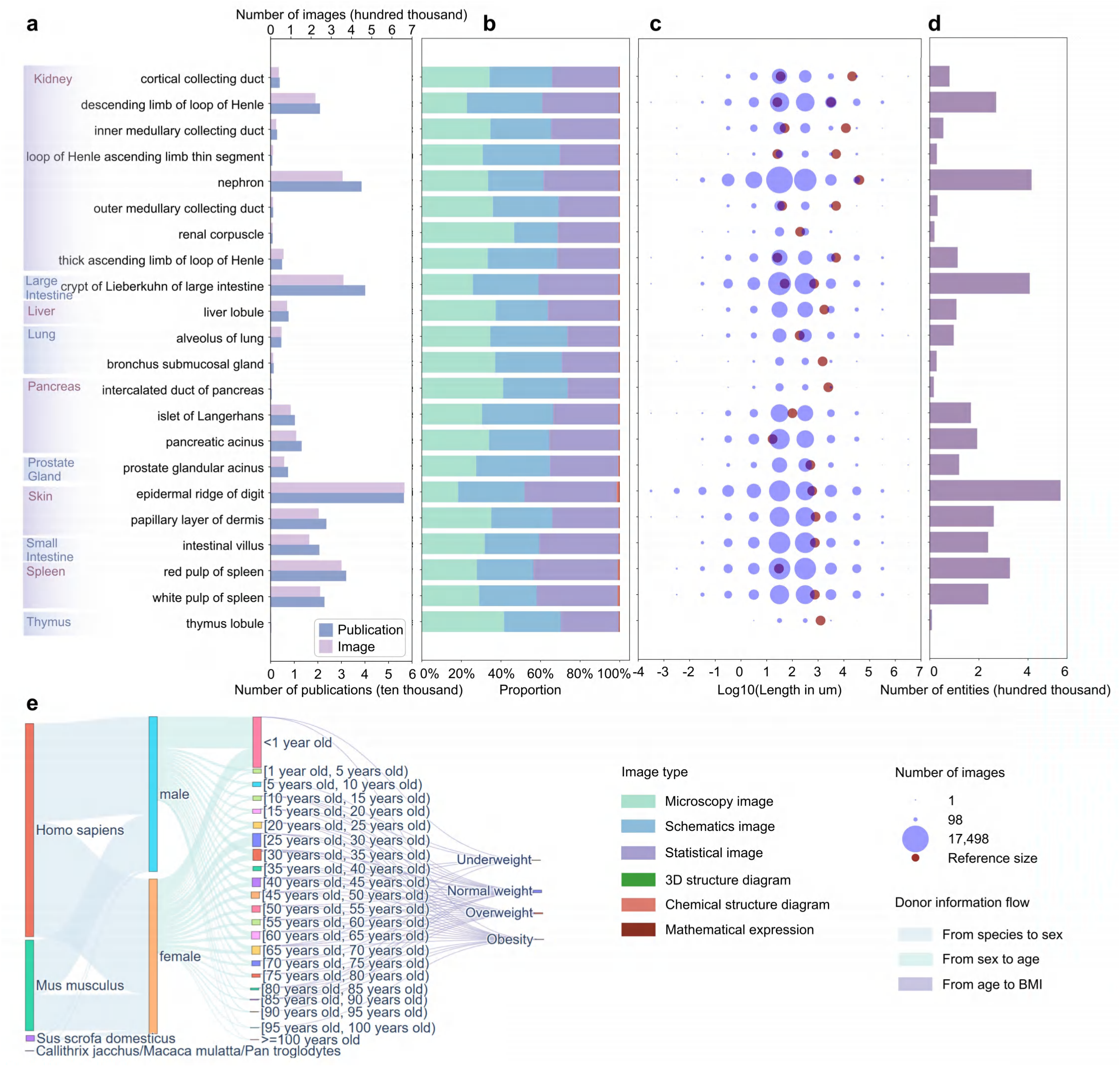
FTU-related data extraction via HRAftu-LM-RAG. **a.** Number of PMC publications and corresponding images for 22 FTUs across 10 organs and all 293 species. **b.** Proportion of six image types among all FTU-related images. The image types include microscopy, schematic, statistical, 3D structure, chemical structure, and mathematical expression. **c.** Distribution of scale-bar lengths (µm) in microscopy images. Scale-bar lengths with counts greater than 50 were selected, yielding only between 10⁻⁴ and 10⁷ µm. Red “reference size” dots derive from HRA data; for FTUs with cylindrical geometry, two reference values are provided (height and diameter). **d.** Number of biological entities associated with AS+CT+B annotated for each of the 22 FTUs, extracted from figure captions and figure nodes of microscopy images and schematic images. **e.** Flow of donor-metadata categories for figure captions and figure nodes from microscopy and schematic images, and abstract. Only categories most relevant to HRA FTUs are shown; exhaustive subcategories for species, sex, age, and BMI, as well as “unknown” and “other,” are not displayed in the figure. The age bar reflects postnatal donors only. Full data details are available in the **Source Data**.

With the categorized images, we focused on microscopy and schematic images for FTU morphology and composition. Among the categorized microscopy images, 215,031 contained a visible scale bar. A comparison between the distribution of extracted scale bar lengths and the anatomical reference sizes^2^ for each FTU is shown in **Fig. 6c**. The results indicate that commonly used scale bars often cluster around or span the established reference size ranges, with lengths between 1 µm and 1 mm particularly prevalent across multiple FTUs. Among the categorized microscopy and schematic images, a total of 1,719,138 biological entities associated with AS+CT+B were extracted. The distribution of these entities across FTUs is shown in **Fig. 6d**. The thymus lobule had the fewest extracted entities, with 8,733 entities, while the epidermal ridge of digit had the most, with 532,317 entities. This variation corresponds closely to the volume of available PMC resources for each FTU.

To explore how FTU properties vary across demographic groups, we first analyzed the distribution of donor metadata. **Fig. 6e** shows the availability and co-occurrence of key donor metadata categories extracted from the literature, focusing on six selected core species: *Homo sapiens*, *Mus musculus*, *Sus scrofa domesticus*, *Callithrix jacchus*, *Macaca mulatta*, and *Pan troglodytes*. The complete dataset is available on Zenodo. *Homo sapiens* represents the largest proportion among these six species. Given the HRA’s emphasis on human data, our analysis focuses on *Homo sapiens* donors exclusively. We identified 21,581 female and 18,341 male donor entries. Regarding developmental stage, 3,220 entries corresponded to prenatal individuals and 26,420 to postnatal individuals. For postnatal *Homo sapiens* donors, detailed age information spanned a wide range from infants (<1 year old) to individuals over 100 years old. The most frequently documented postnatal age group was ‘<1 year old’, with 4,227 entries, followed by the [25-30 years) age bracket, accounting for 2,403 entries. Other notable age groups included [30-35 years) with 2,178 entries and [40-45 years) with 1,702 entries, reflecting diverse age coverage within the dataset. Regarding BMI categories, ‘Normal weight’ was the most frequently specified, with 605 entries, followed by ‘Obesity’ (495 entries) and ‘Overweight’ (329 entries).

Going beyond overall distributions, specific *Homo sapiens* donor demographic group profiles were identified for individual FTUs (**Supplementary Table 25**). The most frequent donor profile with complete metadata (i.e., species, sex, age, and BMI data is available) was observed for the ‘epidermal ridge of digit’, with 32 instances for the human female, normal weight, aged [30–35 years) donor profile. The most frequent donor profile for human samples that had sex, age but not BMI data, was observed for the ‘papillary layer of dermis’, with 283 instances of human female aged [25–30 years).

We further examined FTU sizes (using scale bar lengths as a proxy) in *Homo sapiens* across different demographic groups (**Fig. 7**). In a comparison of scale-bar lengths in male and female samples (**Fig. 7a**), five FTUs—intestinal villus, epidermal ridge of the digit, papillary layer of the dermis, renal corpuscle and thymus lobule—exhibited higher median lengths in male samples, whereas two FTUs—bronchus submucosal gland and the thin ascending limb of the loop of Henle—were higher in females. Bootstrap CIs for the male–female median difference generally spanned zero and, after BH–FDR (α = 0.05), no FTU showed a significant median difference. Non-parametric Mann–Whitney U testing confirmed a statistically significant sex difference for intestinal villus (p < 0.01; **Supplementary Table 26**). HRA reference sizes often fell within the interquartile ranges of male and female measurements, particularly for kidney-associated FTUs. Additionally, we examined age-dependent scale-bar trends for FTUs grouped by organ region (**Fig. 7b**). Among organs represented by two or more FTUs, kidney FTUs (e.g., loop of Henle ascending limb thin segment, nephron, outer medullary collecting duct, renal corpuscle), spleen FTUs, and thymus FTUs often decreased. Lung FTUs and pancreatic FTUs showed mixed trends.

**Figure 7.**
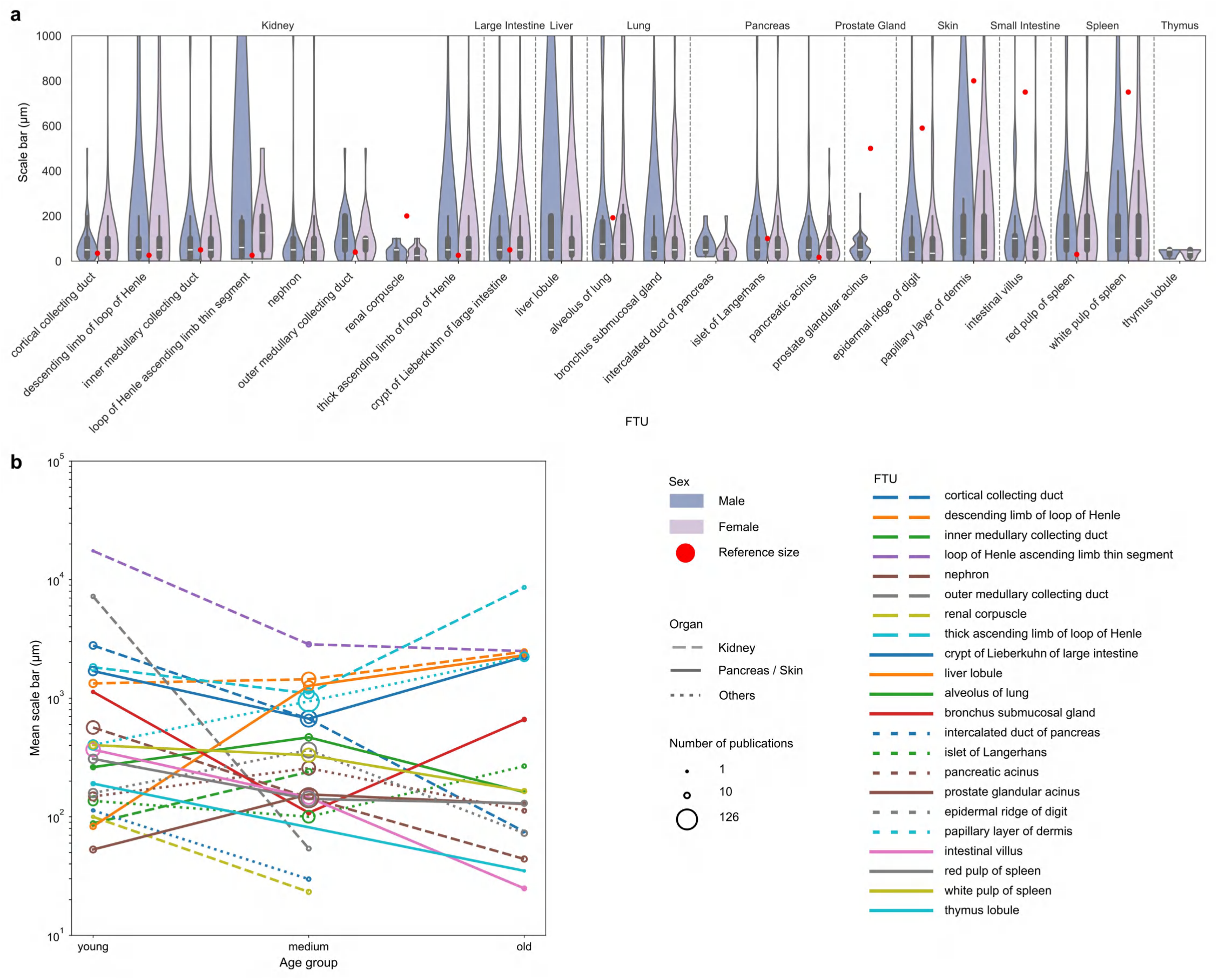
FTU scale lengths in *Homo sapiens* across sex and postnatal age. Only FTUs annotated as *Homo sapiens* with scale bars measured in micrometres are included. **a**. Comparison of scale bars for 22 FTUs between male and female. Only FTUs annotated as *Homo sapiens* with scale bars measured in millimeter and micrometer are included. Red dots indicate reference sizes from the HRA. Five FTUs (i.e., nephron, liver lobule, bronchus submucosal gland, intercalated duct of pancreas, and thymus lobule) have reference size > 1000 µm and are omitted from this panel. Seven FTUs (i.e., inner medullary collecting duct, descending limb of loop of Henle, loop of Henle ascending limb thin segment, thick ascending limb of loop of Henle, outer medullary collecting duct, cortical collecting duct, and crypt of Lieberkuhn) possess two reference dimensions—length and diameter—and are plotted using only the smaller of the two. As the prostate glandular acinus is exclusive to human males, it is displayed with male samples only. **b.** Comparison of mean scale bars sizes for 22 FTUs across three postnatal age groups: young (0 ≤ age < 15 years), medium (15 ≤ age < 70 years) and old (age ≥ 70 years). Only FTUs annotated as *Homo sapiens* with scale bars measured in micrometers are included. Circle area size denotes the number of publications per data point shown.

## Discussion

This work focuses on automating the comprehensive characterization of FTUs, aiming to move beyond the limitations of manual curation and fragmented data sources. The integration of LLMs for textual reasoning, LVMs for visual interpretation, and RAG for knowledge grounding enables HRAftu-LM-RAG to interlink diverse data about FTUs across scales, including physical dimensions, biology entities, donor information, and associated publications. This automated evidence collection can facilitate a deeper understanding of FTU property as multiscale functional entities, contributing to their effective representation in the HRA. By identifying key features and their contextual details from vast scholarly publications, the HRAftu-LM-RAG framework makes it possible to analyze how FTUs vary across donor demographics and disease states—an important prerequisite for personalized medicine and drug development.

The HRAftu-LM-RAG framework and workflows are scalable and robust; they were used to process 244,640 PMC publications and analyze 1,389,168 figures—categorizing 617,237 images as microscopy or schematic. From these, 1,719,138 biological entity mentions associated with AS+CT+B and 331,189 scale bars—most commonly between 1 µm and 1 mm across multiple FTUs, were extracted. For donor metadata, there are 21,581 female and 18,341 male donor entries in *Homo sapiens*, with the most represented category being females aged 30–35 years with normal weight (n = 171).

Current limitations of the HRAftu-LM-RAG framework comprise: (1) This study does not distinguish between healthy and diseased data. Going forward, we will extend the framework to identify healthy control data and major diseases (e.g., diabetes, cancer) to extract and compare FTU properties for healthy control vs. diseased donors. (2) Currently, HRAftu-LM-RAG cannot be used to train new LVMs and LLMs; instead, we performed prompt-based evaluations on a held-out test data using several existing models, and compared their performance on the FTU tasks; the model plus prompt that achieved the best results were adopted for usage. (3) Composite figures in papers were assigned to multiple image type categories; we plan separate composite figures in the next iteration of the framework and assign each single figure one category. (4) The quality and specificity of extracted information depends on the data reported in existing publications. As novel technologies become available and resulting data is reported in scholarly publications, AI-extracted data will improve. (5) The latest release, HRA v2.3, includes 26 functional tissue units across 11 organs—comprising 20 anatomical structures and 126 unique cell types—and represents broader coverage than HRA v2.2. In future work, we will apply the same HRAftu-LM-RAG framework to v2.3.

Going forward, we will run the HRAftu-LM-RAG workflows for each new release to ensure all existing FTUs are covered and will make the set of retrieved papers and FTU properties available to human experts authoring and reviewing FTUs for the HRA. Plus, we are planning to compare scholarly evidence extracted by HRAftu-LM-RAG with experimental data downloaded from diverse human atlas portals and made available via the HRA Knowledge Graph (KG)^52^. This way, the HRA is triangulating human expertise (e.g., covered via ASCT+B tables and FTU illustrations), scholarly evidence extracted from publications, and experimental data to create a multiscale reference atlas of the human body that can be used to advance precision health and precision medicine.

## Methods

### Data collection

#### HRA data collection

HRA data (i.e., organs and FTUs) are obtained from the 8th HRA release (v2.2) KG^52^. In this HRA release, there are 22 FTUs in 10 organs with corresponding ontology IDs^39^ (i.e., Uberon ID) (**Supplementary Table 27**).

#### FTU-related publication package collection

To identify publications relevant to FTUs, we employed a systematic approach integrating ontology-based term extraction and literature searches^53^. Ontology terms provide a standardized and semantically rich framework for biological entities, enabling precise identification and retrieval of relevant literature across databases.

##### Literature search in PMC

To enhance the comprehensive and accurate retrieval of relevant PMC publications, we first utilized BioPortal (https://bioportal.bioontology.org), a comprehensive repository of biomedical ontologies, to extract ontology data corresponding to each FTU’s specific Uberon ID via the BioPortal REST API (https://data.bioontology.org/documentation), including the preferred label, synonyms, definition, and source ontology (**Supplementary Table 28**). Across the 22 FTUs, this yielded 147 synonyms in total (in addition to the 22 preferred labels). To reduce redundancy and improve retrieval efficiency, semantically similar terms were grouped into 139 composite search expressions, which were used to perform keyword-based searches in PMC (https://pmc.ncbi.nlm.nih.gov). The specific retrieval strategies are detailed in **Supplementary Table 29**. The resulting records were manually exported in the “PMCID list” format for the subsequent batch download of publication packages. The result is 709,346 publication PMCIDs, see **Fig. 6a** for the number of publications for each of the 22 FTUs.

##### Publication package collection from PMC

Publication packages are extracted from PMC via the Open Access subset, accessible through the National Center for Biotechnology Information (NCBI) FTP service (https://ftp.ncbi.nlm.nih.gov/pub/pmc/oa_package). For the study presented in this paper, only publication packages under the “Commercial Use Allowed” license are collected. The publicly available metadata file from PMC (https://ftp.ncbi.nlm.nih.gov/pub/pmc/oa_comm_use_file_list.csv) is used to identify the “oa_comm” packages, providing a mapping between publication accession IDs (PMCIDs) and the corresponding file paths of the publication packages. Using this information, full publication packages are systematically downloaded, each containing the full text in NXML and/or PDF formats, along with figures, typically in GIF and/or JPEG formats. As a result, there are 260,855 selected PMC publication packages for 22 FTUs.

#### Image data collection

Image files and associated textual information were obtained from the PMC publication packages. For each image, essential information—including the label, caption, title, graphic, and reference text—was parsed from the NXML structure. Each image is associated with a unique figure ID within a *<fig>* section. The *<fig>* section includes several subsections: *<label>*, *<caption>*, and *<graphic>*, each providing key metadata for the image. The *<caption>* section contains descriptive text and may include a *<title>* element with a specific image title. The image file itself is identified by the file location specified in the *<graphic xlink:href>* tag, enabling retrieval of the corresponding image from the publication package. Reference text, or citations of the image within the article, is located by searching for the figure ID in *<xref>* tags within the NXML text. Of the 2,008,820 distinct figures identified from the 249,196 PMC publications, 1,389,168 figures had a JPEG rendition, occurring in 244,640 PMC publications. Given that LVMs are primarily designed to recognize and process images in this format, only these 1,389,168 JPEG images and associated publications were selected for subsequent analysis.

#### Publication metadata collection

Publication data are extracted from the NXML files in the selected PMC publication package, specifically from the “article-meta” and “journal-meta” sections. In the “article-meta” section, identifiers such as PMID, PMCID, and DOI are located within the “article-id” tag, the abstract is found in the “abstract” tag, and the publication year is listed in the “pub-date” tag. The “journal-meta” section contains the journal name. A total of 260,855 publications were retrieved and 244,640 PMC publications with at least one JPG image were kept for further analysis. These 244,640 PMC publications are published between 1783 and 2025.

### RAG-Enhanced Large Model Architecture

In this study, we employ LVMs and LLMs enhanced with RAG to explore HRA FTU characteristics. This section first details the architectural setup of LVM deployment, LLM deployment, and the RAG framework construction. Following this, we outline their application in two distinct processing stages: LVM-based task processing for image-centric workflows (WF1 and WF2), and LLM-RAG-based task processing for text-based information extraction (WF1 and WF3-WF5). This structured approach ensures each workflow leverages the most appropriate AI technology for robust and scalable FTU data extraction.

#### LVM Deployment

LVMs are deployed using Ollama and vLLM to handle tasks involving integrated image and text inputs. The LLaVa:34B model is deployed using Ollama. Other LVMs, such as LLaMA3.2 (meta-llama/Llama-3.2-11B-Vision), Phi-3 (microsoft/Phi-3-vision-128k-instruct) and Phi-3.5 (microsoft/Phi-3.5-vision-instruct), and Pixtral (mistralai/Pixtral-12B-2409), are served using the vLLM inference engine. These models are accessed via their HTTP API endpoints, accepting JSON-formatted multimodal inputs comprising image data and text prompts.

#### LLM Deployment

The computational architecture for deploying and serving LLMs utilized the FastGPT platform^54^ (https://tryfastgpt.ai), deployed within Docker containers according to standard procedures. Models were primarily sourced using Ollama (https://ollama.com) and subsequently integrated into the FastGPT environment via its model configuration interface. This setup provided access to the models through both a web-based user interface and a RESTful API for programmatic interaction and experimentation.

Four LLMs—including Gemma2 (google/gemma-2-27b), LLaMA3.1 (meta-llama/Llama-3.1-70B), LLaMA3.2 (meta-llama/llama3.2:latest), and Qwen2.5 (Qwen/Qwen2.5-72B)—were retrieved using Ollama. Each model was subsequently integrated into the FastGPT platform, which provided a consistent runtime environment to ensure uniform scaling and standardized resource allocation across all experimental conditions.

#### RAG Construction

The RAG pipeline was implemented in two sequential stages—knowledge-base indexing and retrieval-augmented inference.

##### Knowledge-base indexing

All source documents intended for retrieval were first passed through a pretrained embedding model to generate dense vector representations. These vectors were persisted in a PostgreSQL database (using the PGVector extension) to support efficient similarity search, while the corresponding raw text was stored in a MongoDB collection for full-text retrieval and downstream assembly. We deployed four embedding models—PubMedBERT (neuml/pubmedbert-base-embeddings), all-MiniLM-L6-v2 (sentence-transformers/all-MiniLM-L6-v2), BGE-large-en-v1.5 (BAAI/bge-large-en-v1.5), and GTE-large (thenlper/gte-large)—and evaluated each on a held-out test set of randomly selected PubMed paragraphs.

For each model, we encoded both the test queries and knowledge-base entries, retrieved top matches via nearest-neighbour search, and computed normalized cosine similarities between query and retrieved embeddings. The model with the highest average similarity was then chosen for all downstream retrieval tasks.

##### Retrieval-augmented inference

At query time, each user input was encoded via the same embedding model and normalized. A nearest-neighbour search against the PostgreSQL vector store yields the top-k most similar document vectors (scored by cosine similarity). The matching document identifiers are then used to fetch the original text from MongoDB. Finally, the retrieved passages are concatenated with the user query and submitted to the large language model, enabling context-aware, retrieval-augmented responses.

### LVM-based Task Processing: WF1 and WF2

LVM-driven workflows addressed two image-centric tasks (WF1 and WF2), each exposed via HTTP API endpoints that accept image data and prompt parameters and return model responses. During the pilot phase, we evaluated five deployed LVMs using a held-out test set of images randomly sampled from PMC. For image-type categorization, each test image was paired with its ground-truth type label; for image-entity extraction, we created a reference set of annotated entities per image and devised multiple prompt templates. Each (model, prompt) combination was submitted to the API, and responses were scored as follows: categorization responses were scored by binary accuracy (1 for correct type, 0 otherwise) and extraction outputs by Jaccard similarity against reference entities, with average response latency recorded. Model– prompt pairs were then ranked on an average accuracy (or mean Jaccard score) and speed, yielding the optimal configuration for each task. In the production phase, the selected LVM and prompt template were fixed, and the entire image collection was processed in batch via the corresponding endpoint to generate final categorization and entity-extraction results at scale.

### LLM-RAG-based Task Processing: WF1 and WF3-WF5

For each downstream task (WF1 and WF3-WF5), we created a retrieval-augmented workflow on the FastGPT platform—including both AI-driven dialogue and knowledge-base vector search—and published it to obtain a unique API key. For each query, we submit an HTTP request containing the user question, the workflow’s API key, the target model identifier, and a task-specific prompt template. The FastGPT workflow orchestrates retrieval of relevant embeddings from the vector store and invokes the LLM to generate a response, which is returned to the client. This setup provides a reproducible, scalable interface for executing RAG-enhanced LLM inference across all tasks.

Specifically, we adopted a two-phase evaluation and execution strategy. In the pilot phase, we designed multiple prompt templates and assembled a test set of query–answer pairs drawn from PubMed publications. Each query was combined with every prompt and submitted to all four deployed LLMs via the FastGPT workflows. Returned responses were compared against the reference answers by computing the Jaccard similarity of their token sets, yielding a performance score for each (model, prompt) pairing. Average response time was also recorded. The combination that achieved the best trade-off between mean Jaccard similarity across the test set and response time was then selected for production. In the production phase, the chosen LLM and prompt template were fixed according to the pilot results, and the workflow was executed at scale to process the full set of task inputs.

## Data Availability

Properties data for the 22 FTUs extracted from the PMC publication packages is available at https://doi.org/10.5281/zenodo.17051026. Test image datasets used to evaluate LVM performance are provided in the **Supplementary Data**. Test inputs for LLM and RAG evaluations—including the associated knowledge bases, prompts, and ground-truth annotations—are listed in the **Supplementary Tables**. All **source data** are provided with this paper.

## Code Availability

All code was made freely available under the MIT license via aGitHub repository: https://github.com/cns-iu/hra-ftu-rag-supporting-information.

## Acknowledgements

We would like to thank Andreas Bueckle, Yashvardhan Jain, Supriya Bidanta, Elizabeth Ginexi, Divya Prasanth Paraman, and Nancy Ruschman, for their expert comments and suggestions on earlier versions of this paper.

The HRA is under active development by HuBMAP, SenNet, KPMP, GenitoUrinary Developmental Molecular Anatomy Project (GUDMAP), and the National Institute of Diabetes and Digestive and Kidney Diseases (NIDDK) with expert input by the HRA Editorial Board and in close collaboration with experts from 20+ other consortia. K.B. has been supported by the following awards, including the NIH Common Fund through the Office of Strategic Coordination/Office of the NIH Director: OT2OD026671, OT2OD033756, U24CA268108, OT2OD030545, U01DK133090, U2CDK114886, and U24DK135157. The funders had no role in study design, data collection and analysis, decision to publish, or preparation of the manuscript. The content is solely the responsibility of the authors and does not necessarily represent the official views of the National Institutes of Health.

## Author Contributions

Y.K. compiled all data, constructed the HRAftu-LM-RAG framework and implemented all workflows, ran the data analyses, generated the figures, and co-wrote the paper. K.B. led the construction of the HRAftu-LM-RAG framework in support of the HRA effort and co-wrote the paper.

## Supplementary Information

**Supplementary Table 1.**
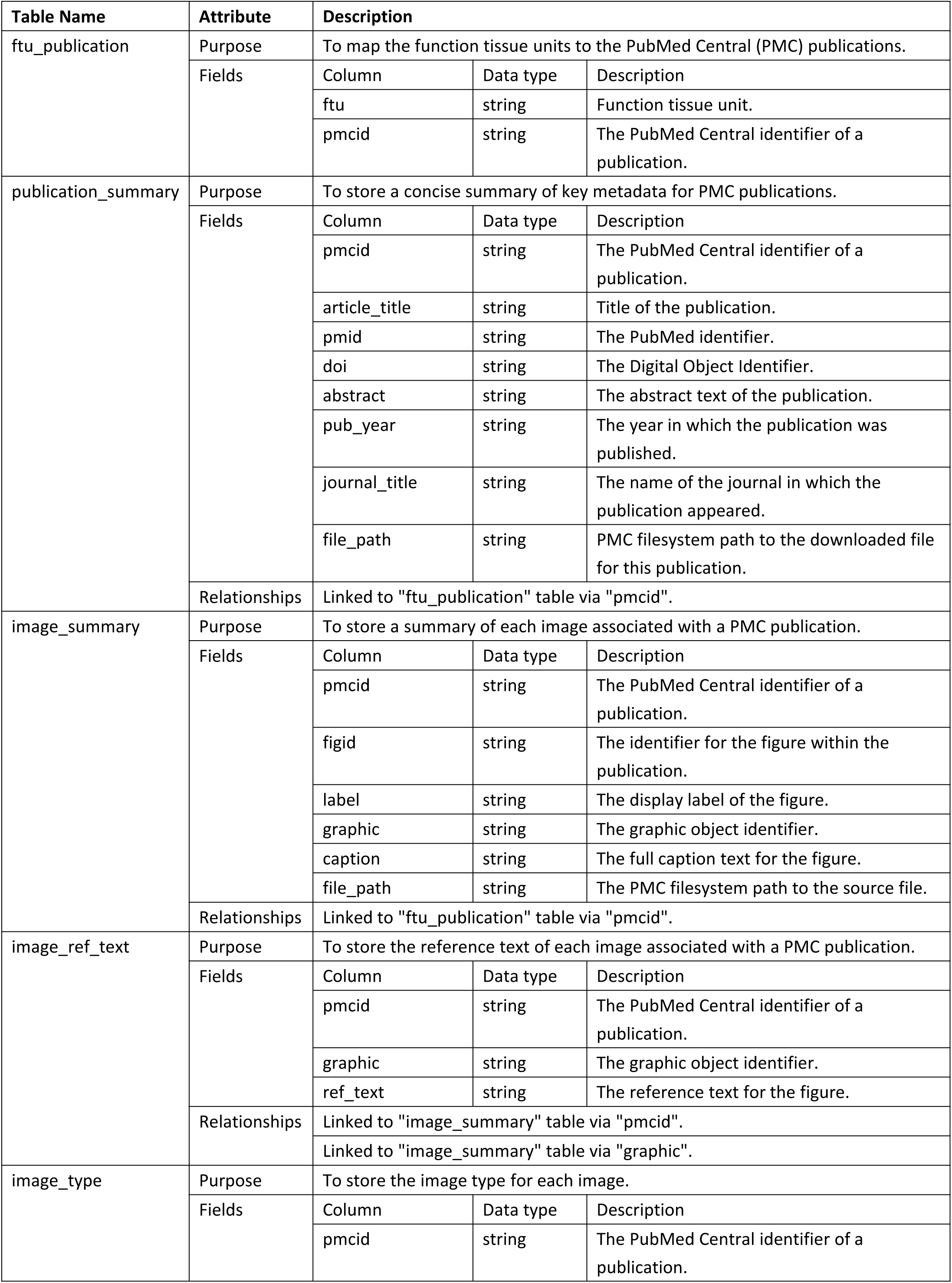
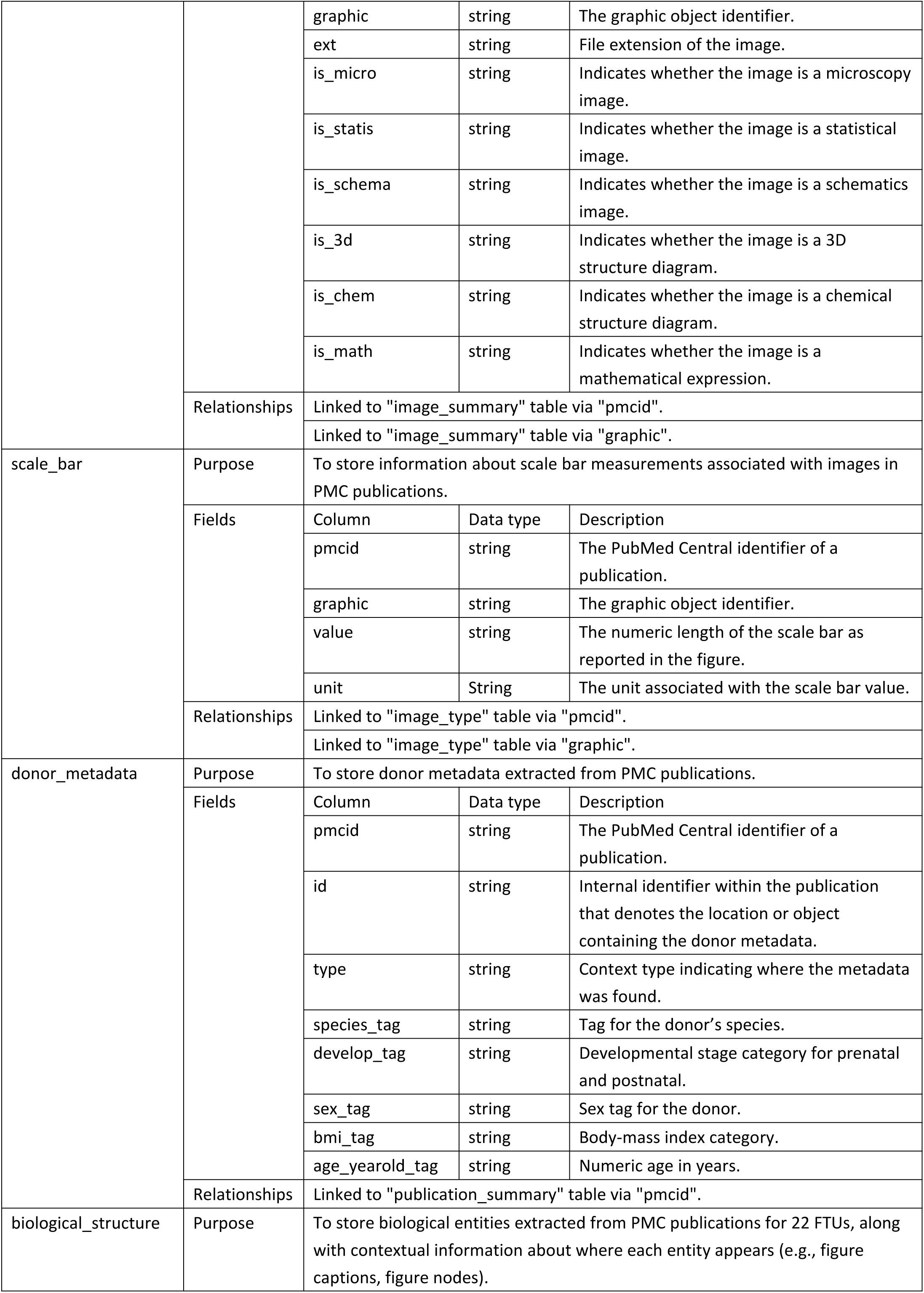
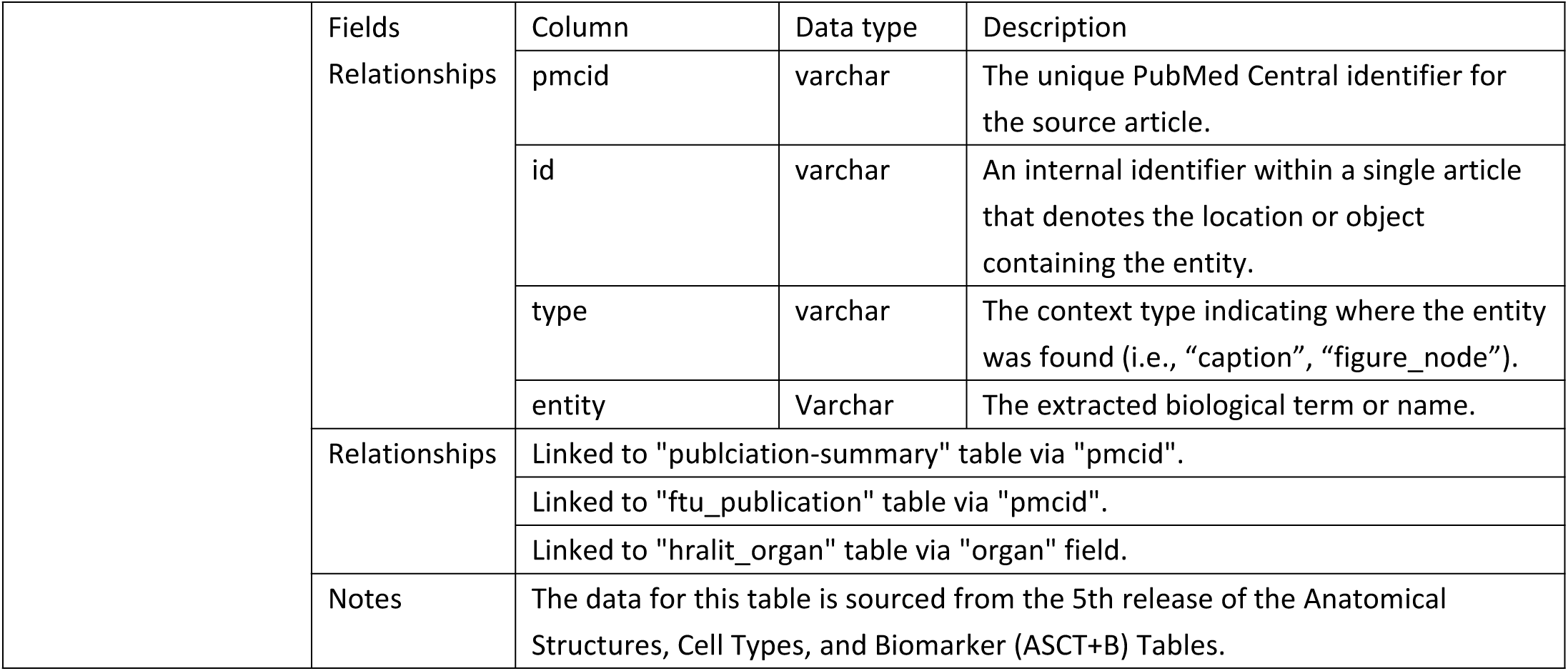
Overview of the results tables.

**Supplementary Table 2.**
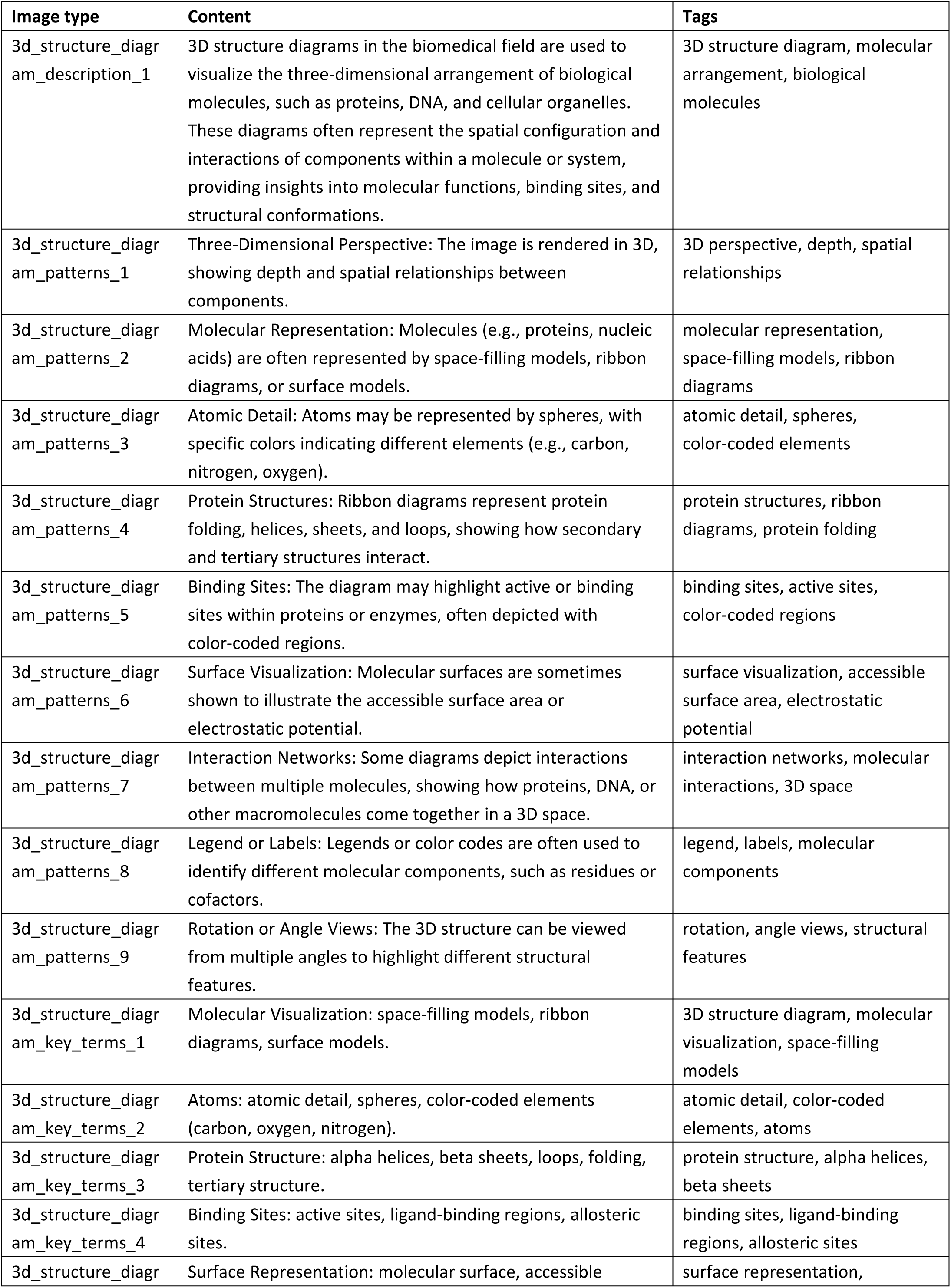
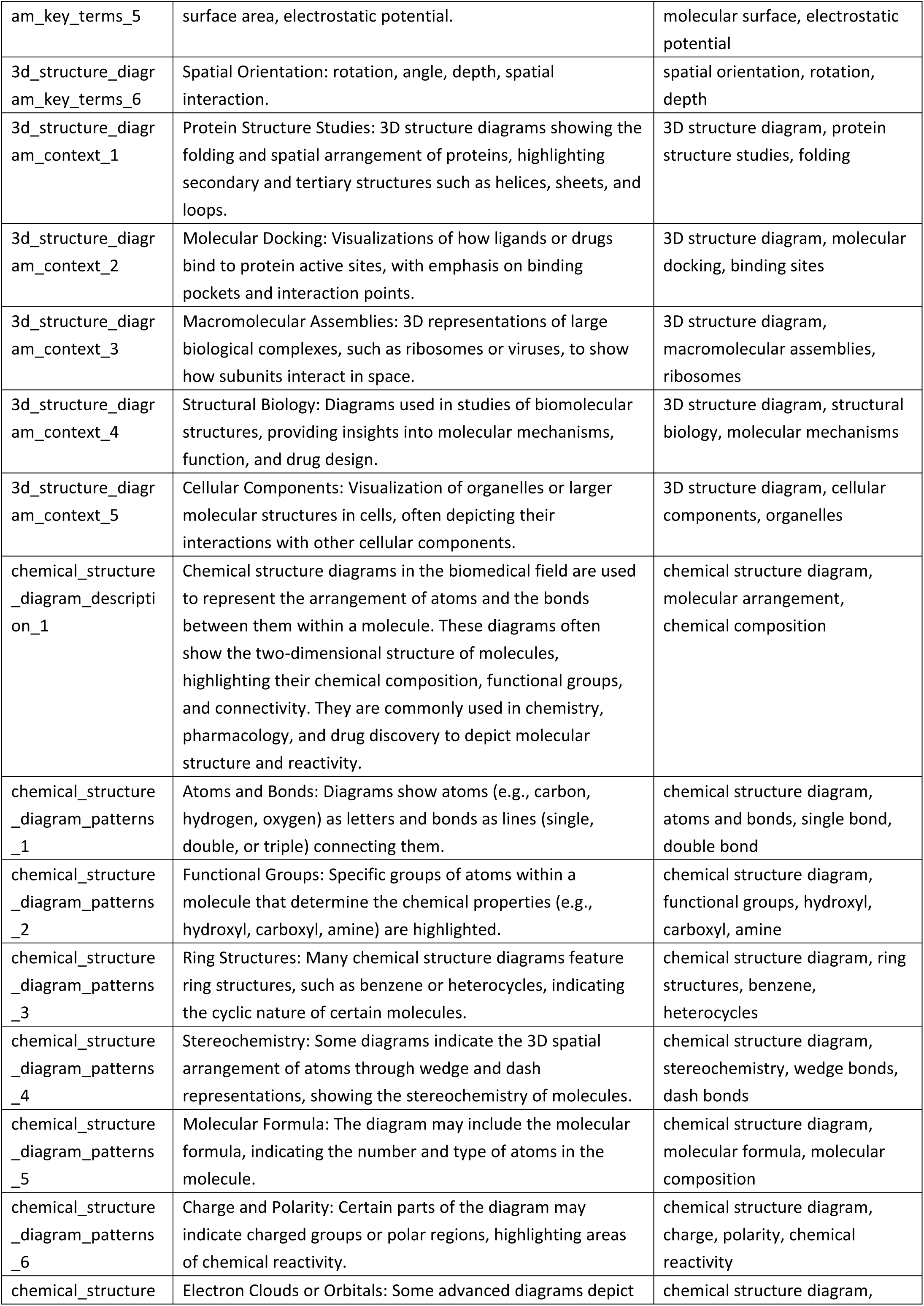
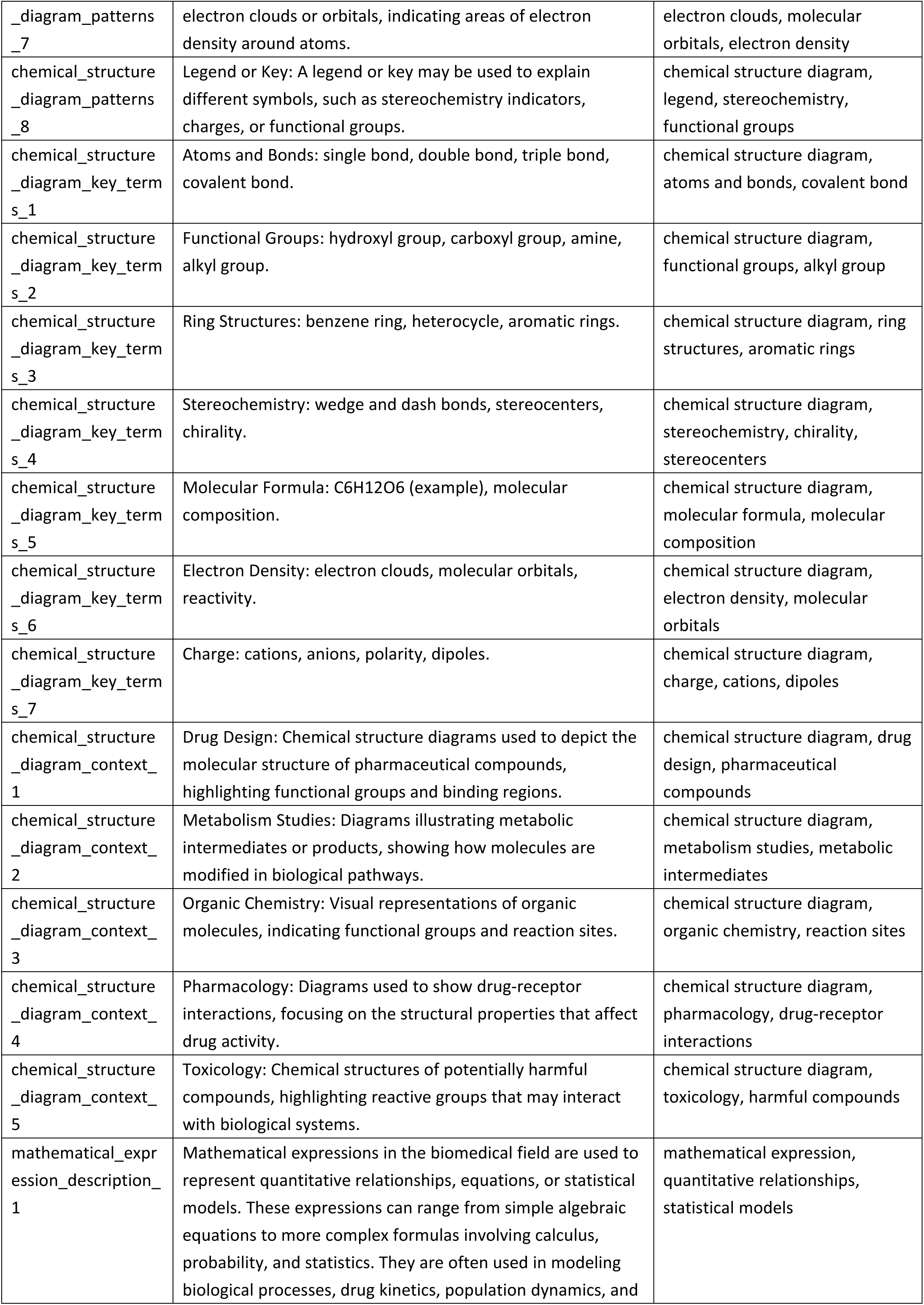
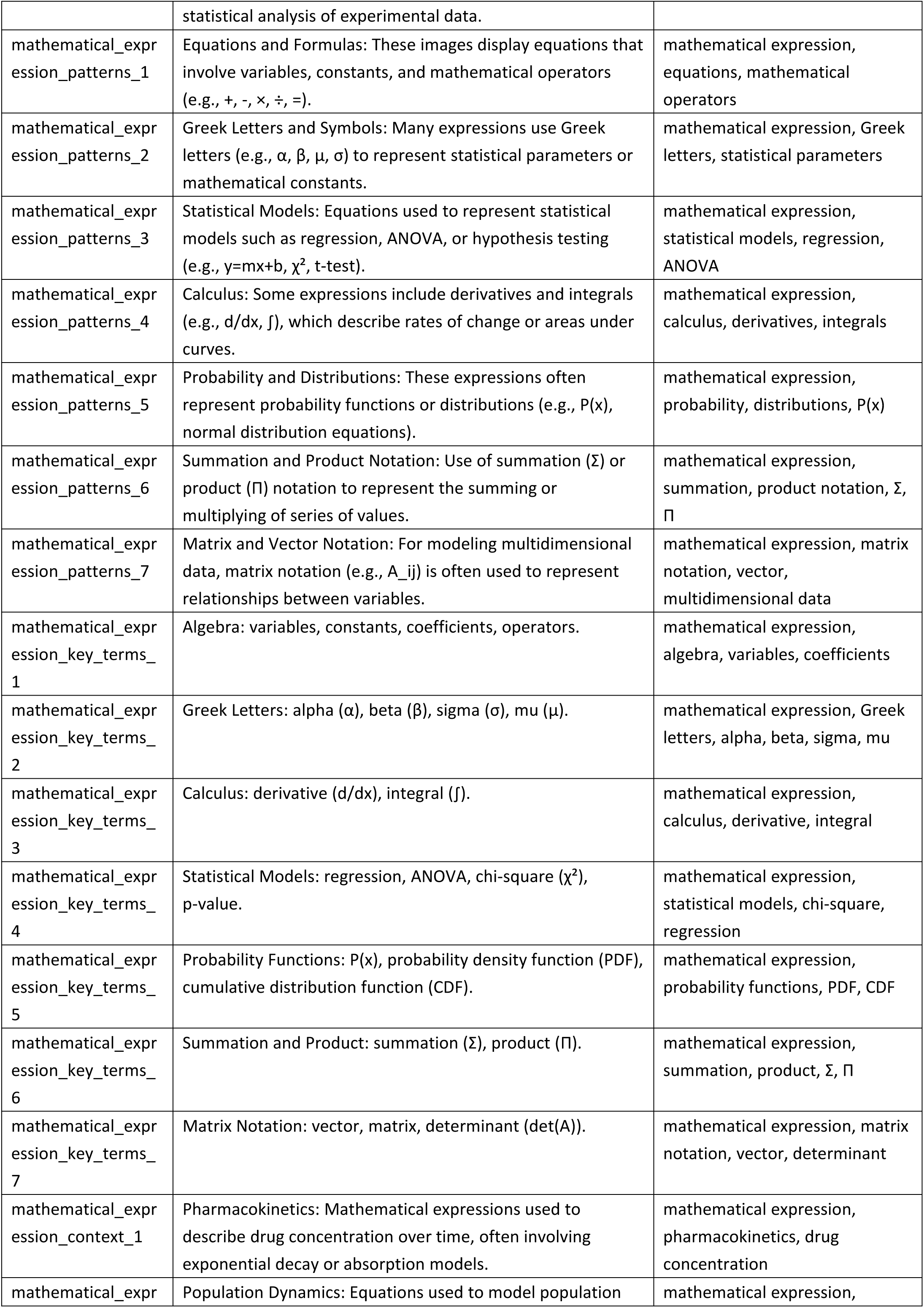
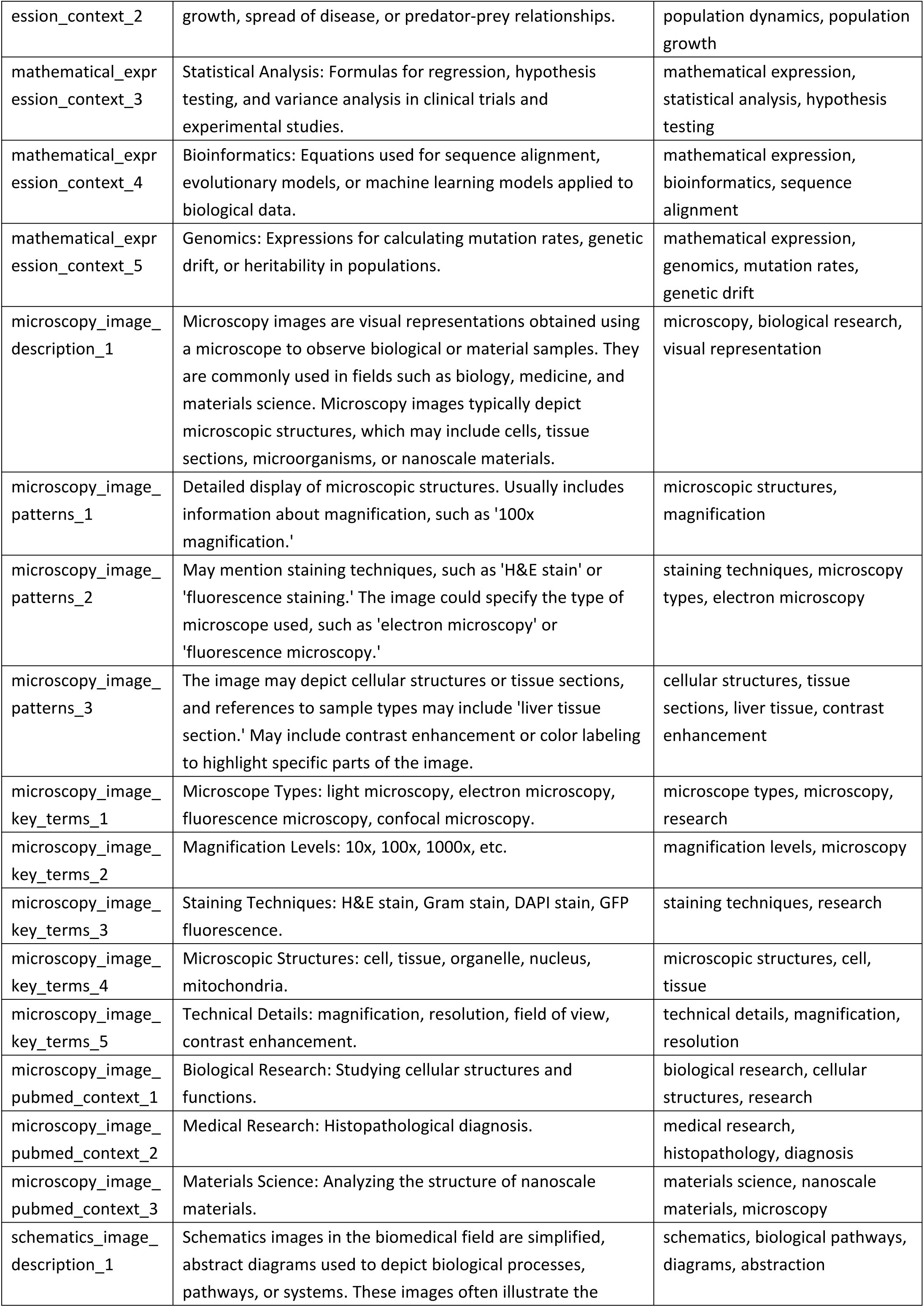
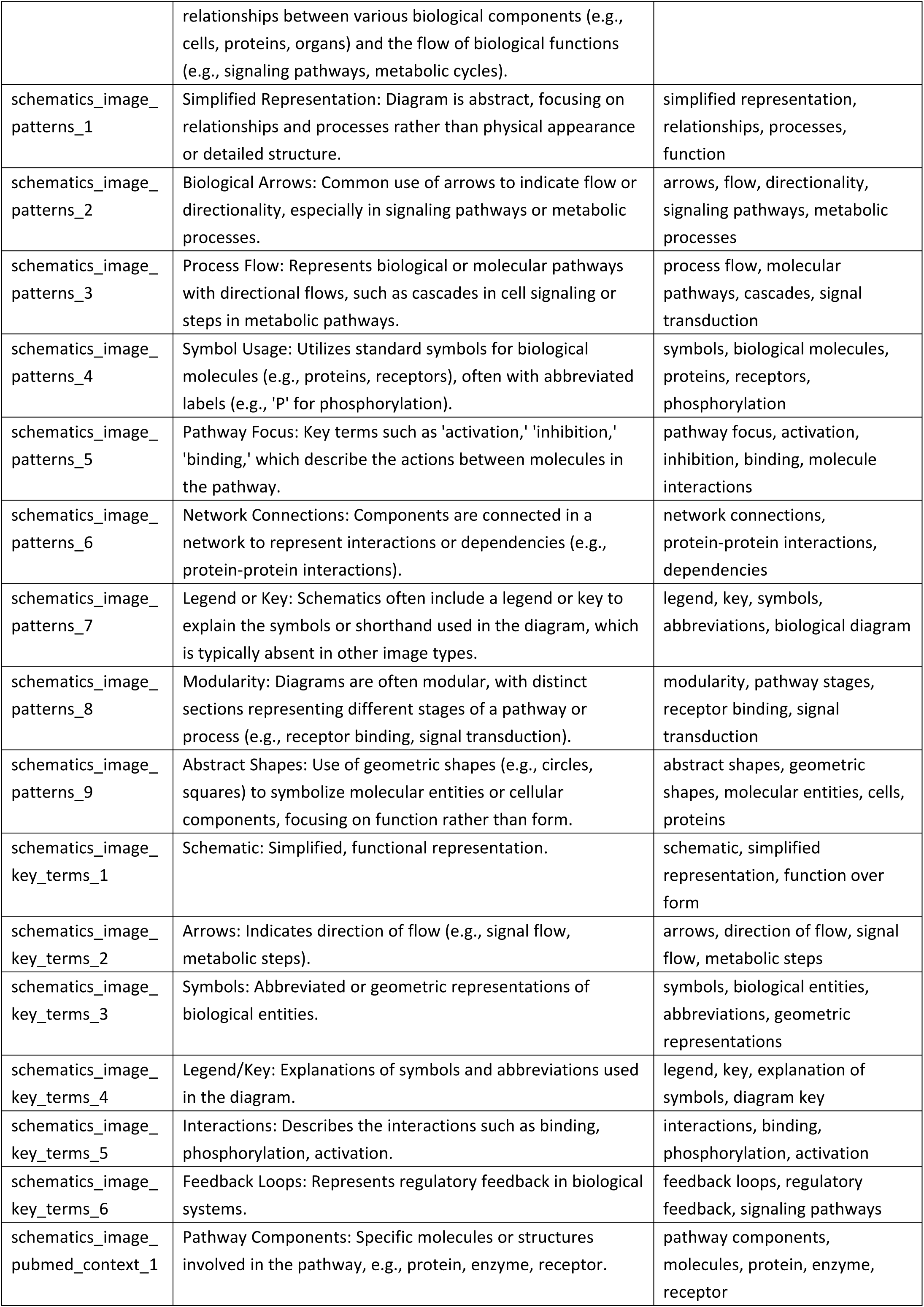
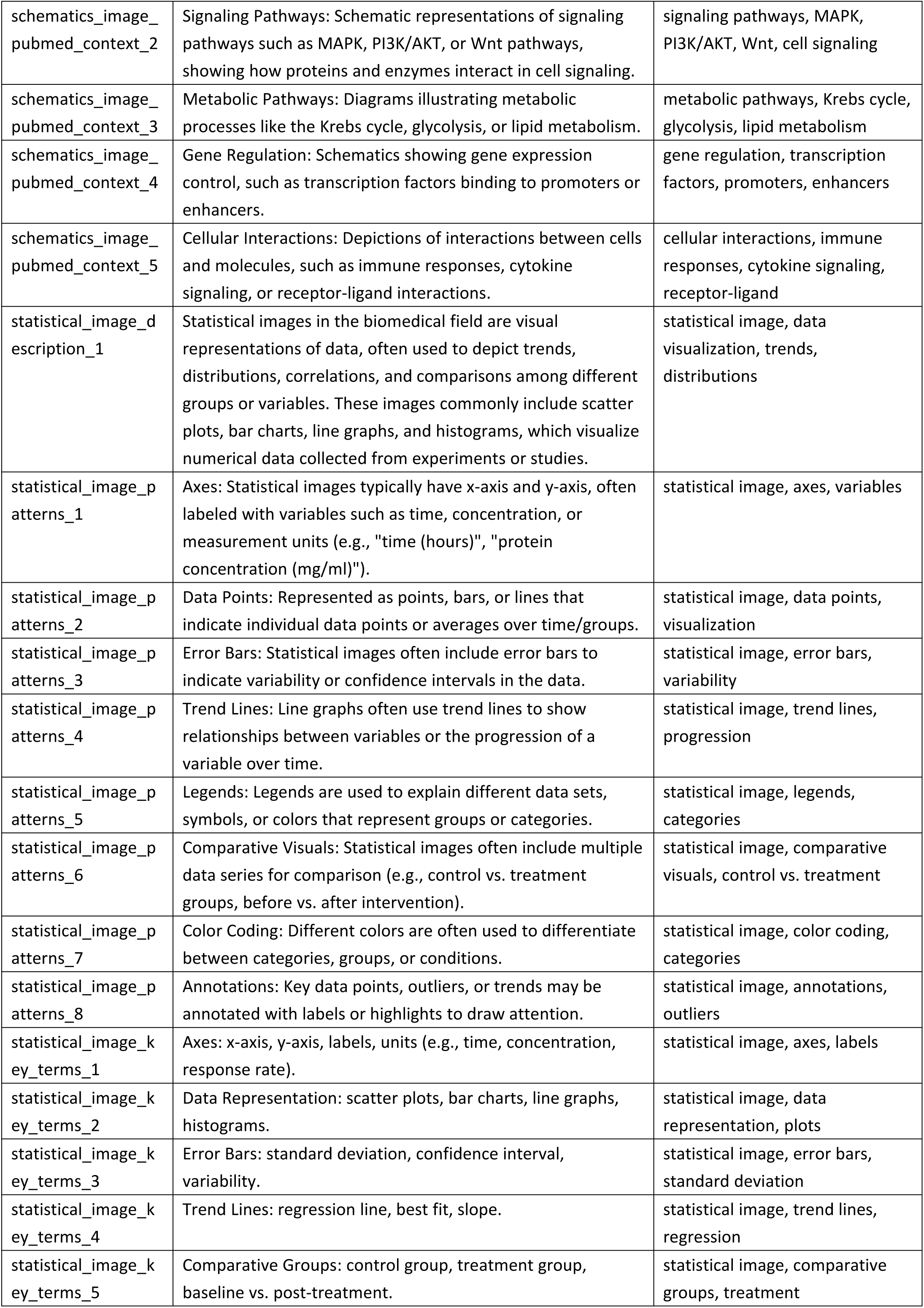
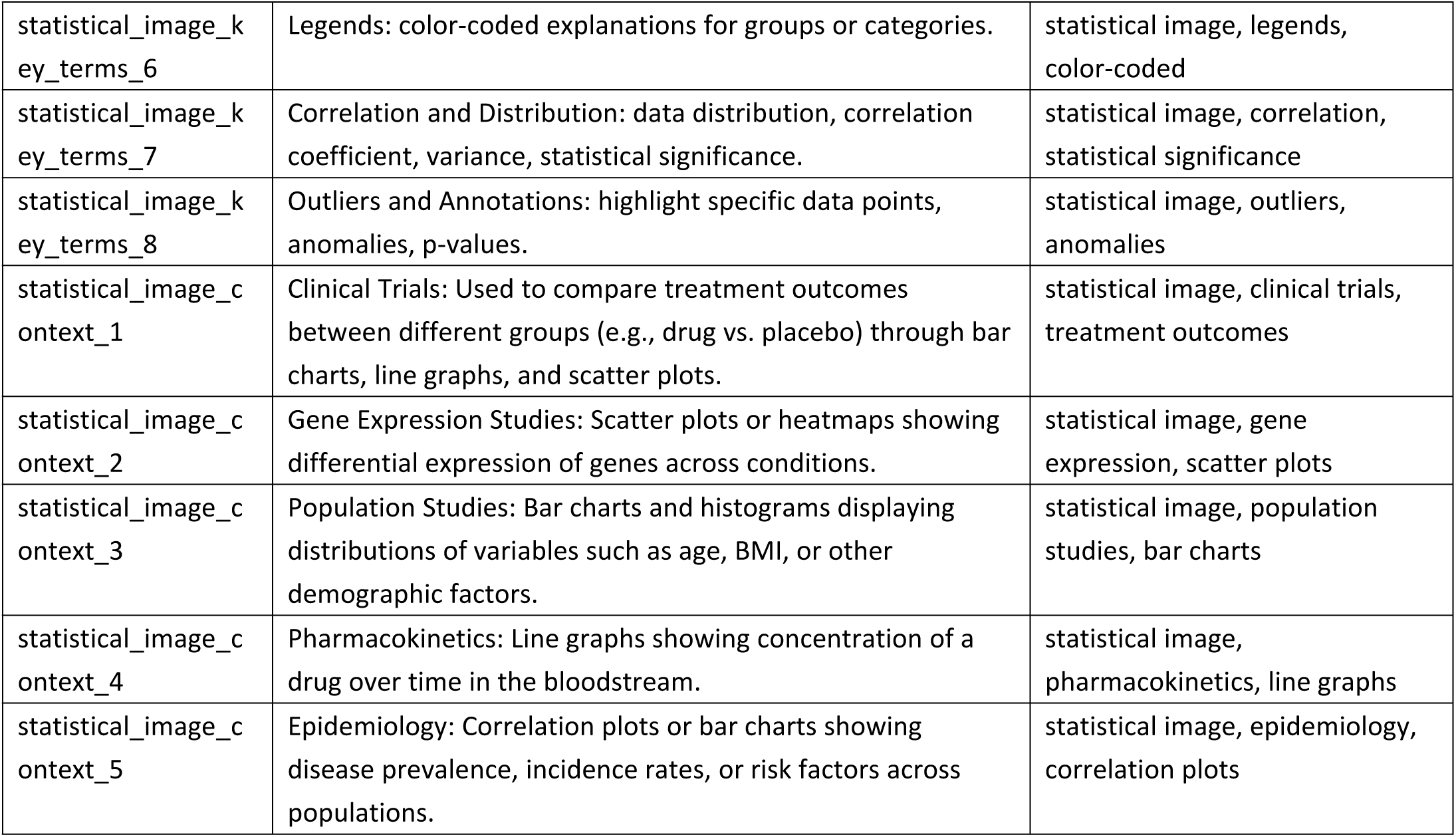
Description of six image types.

**Supplementary Table 3.**
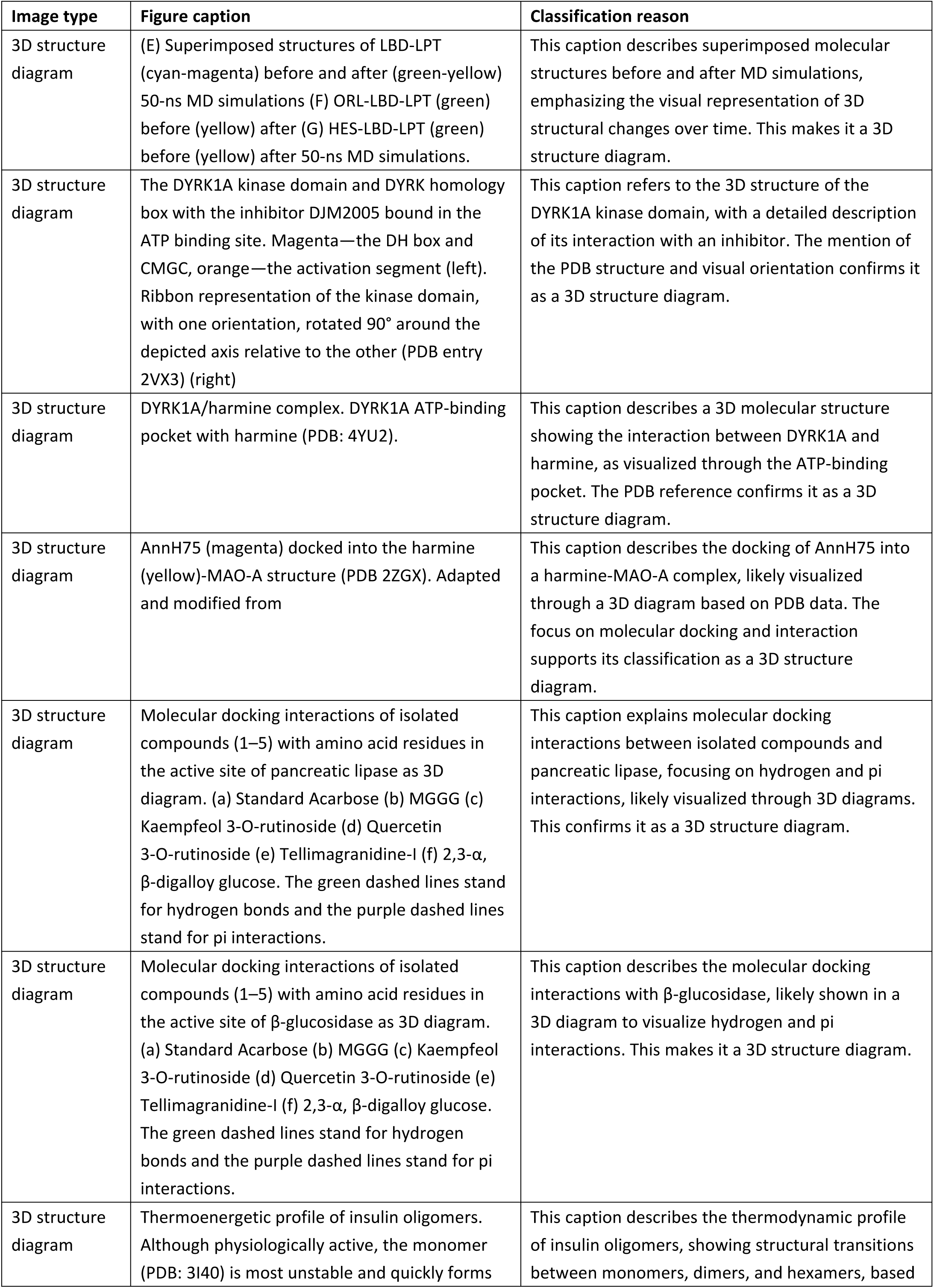
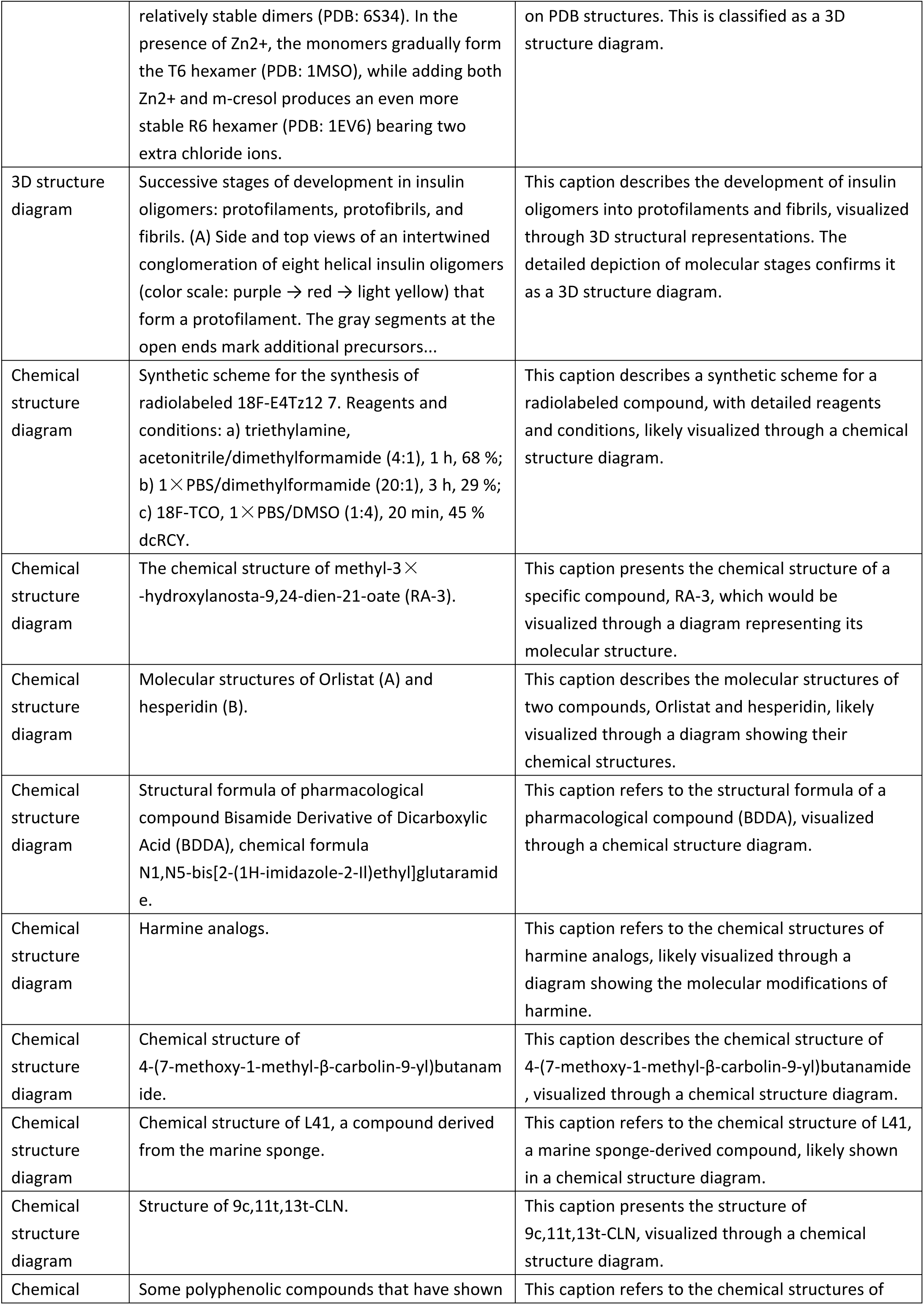
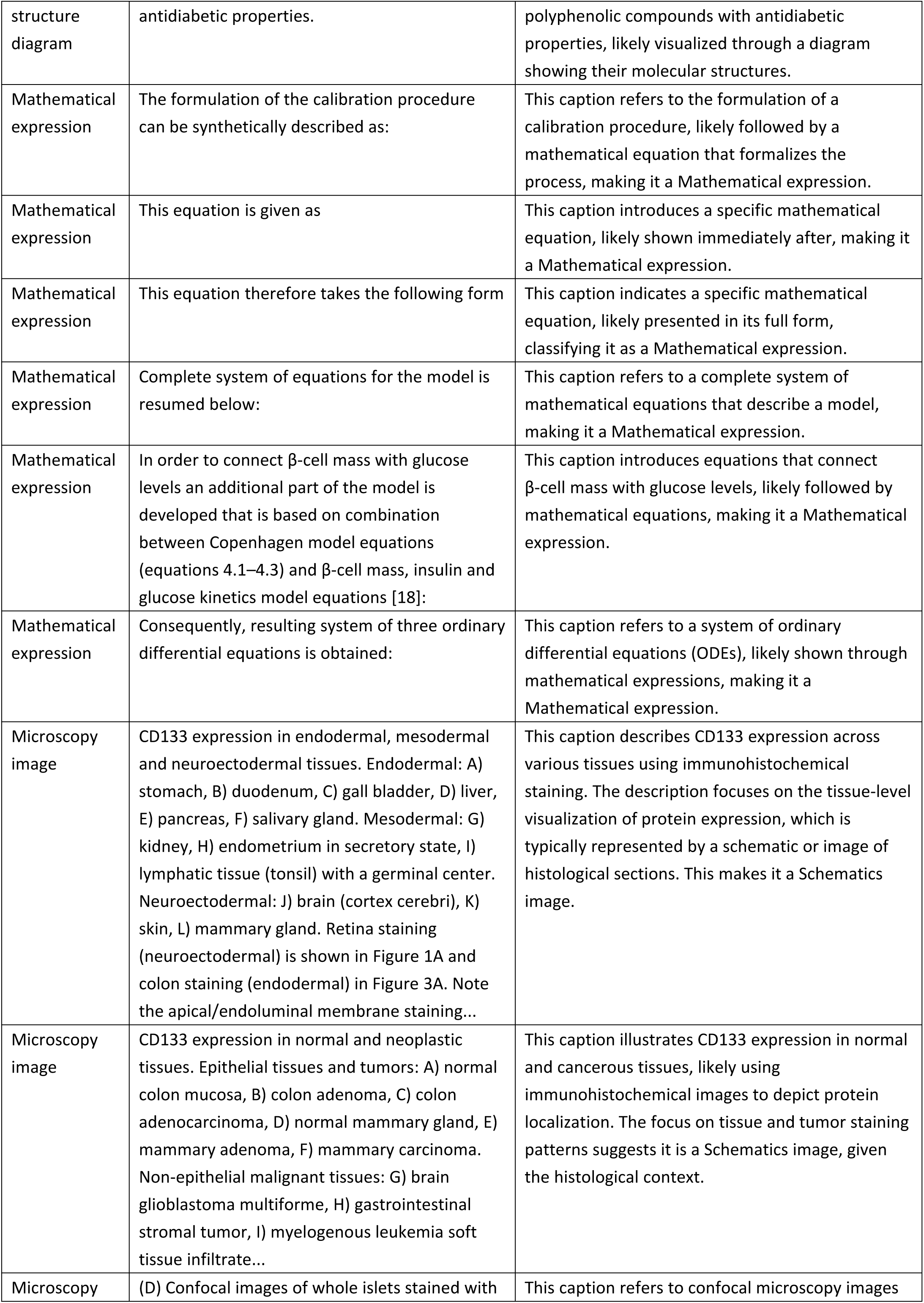
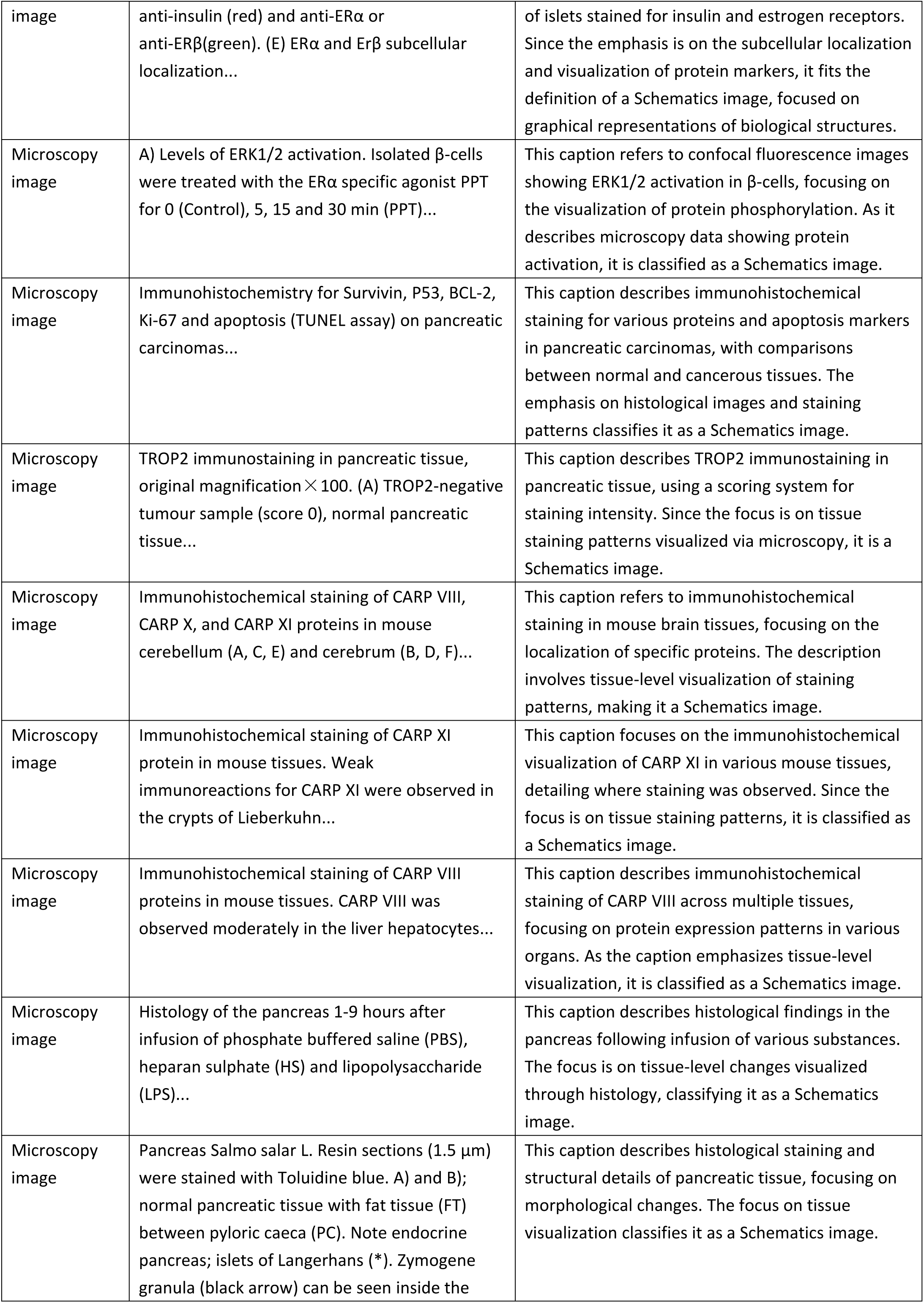
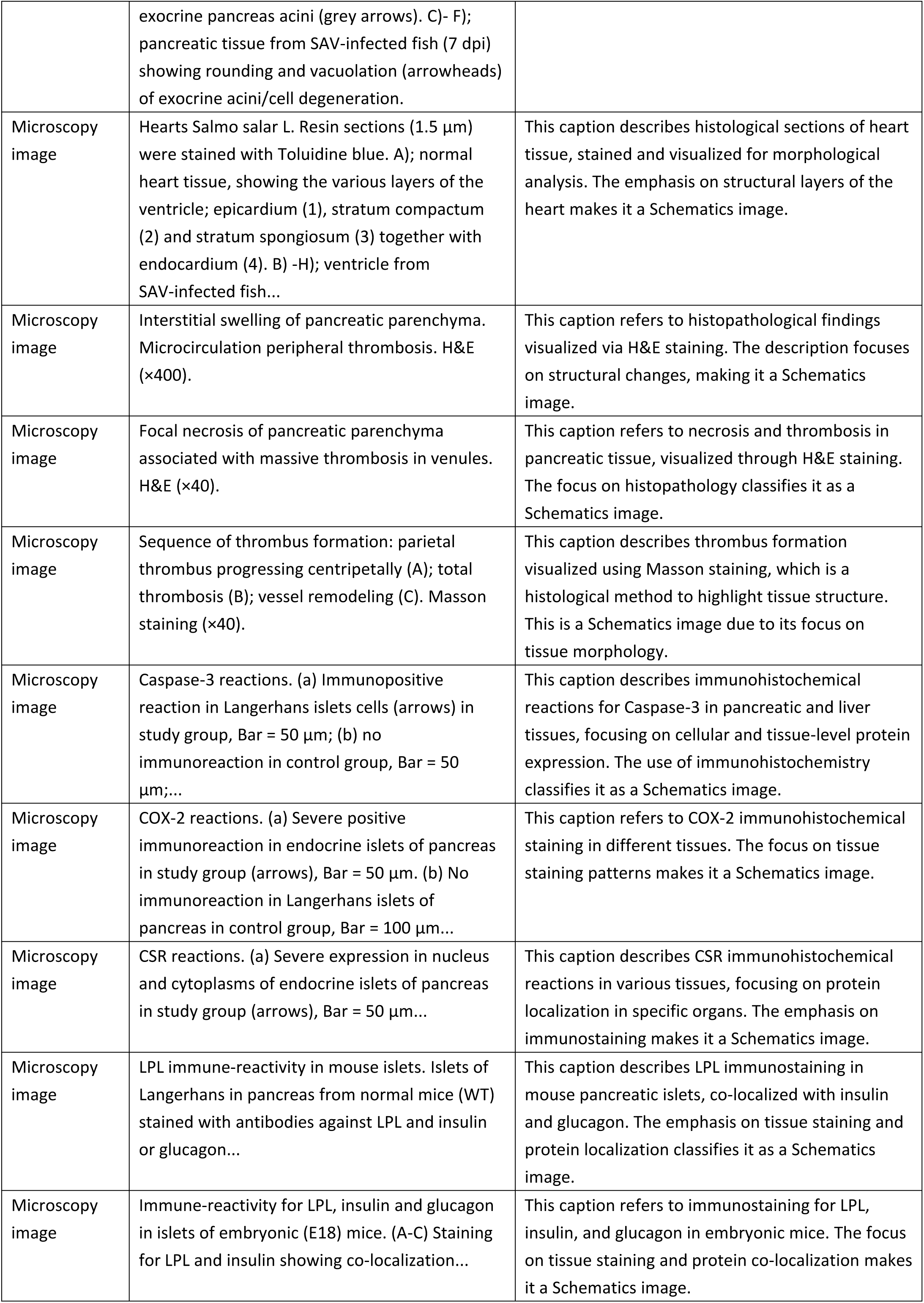
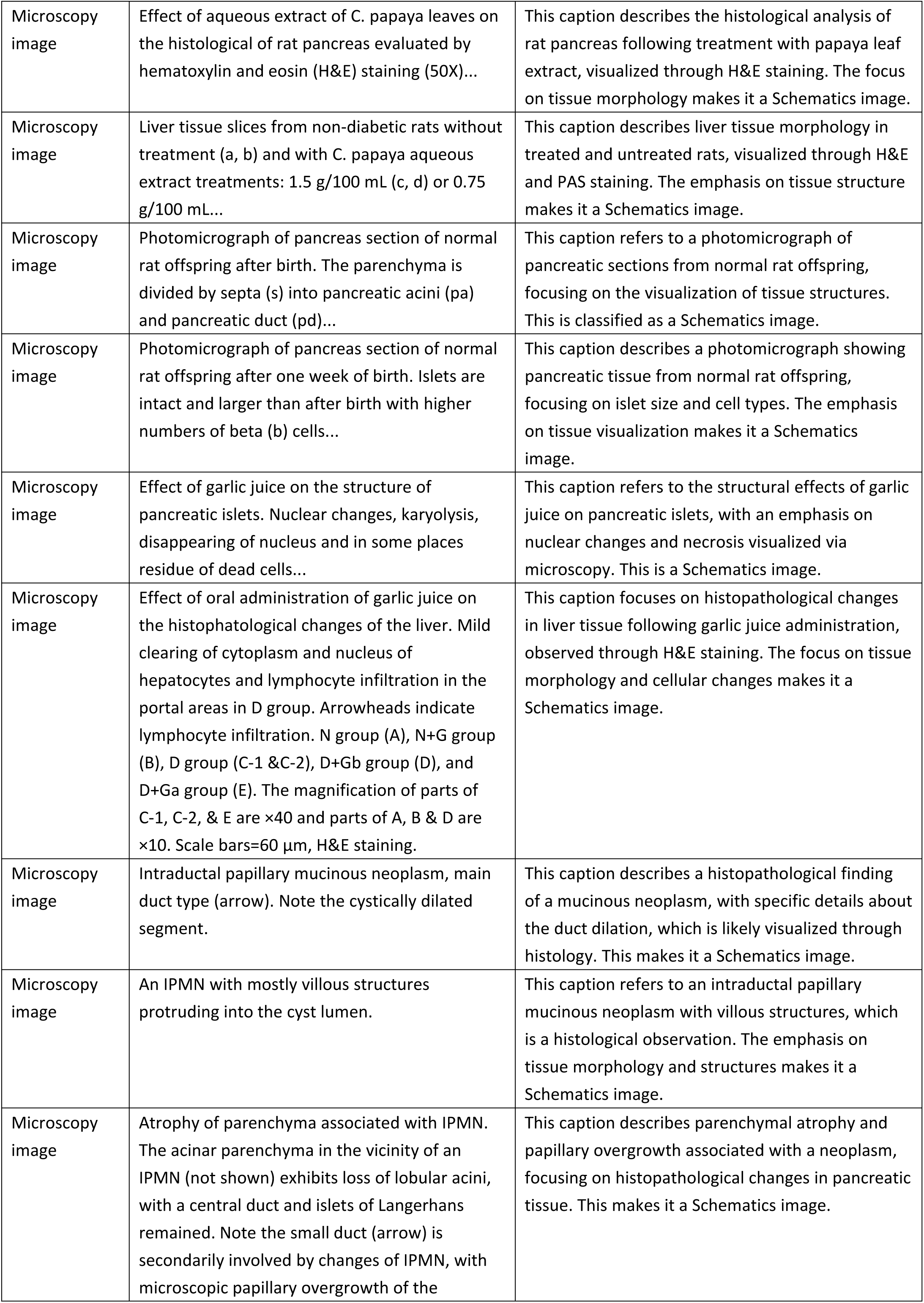
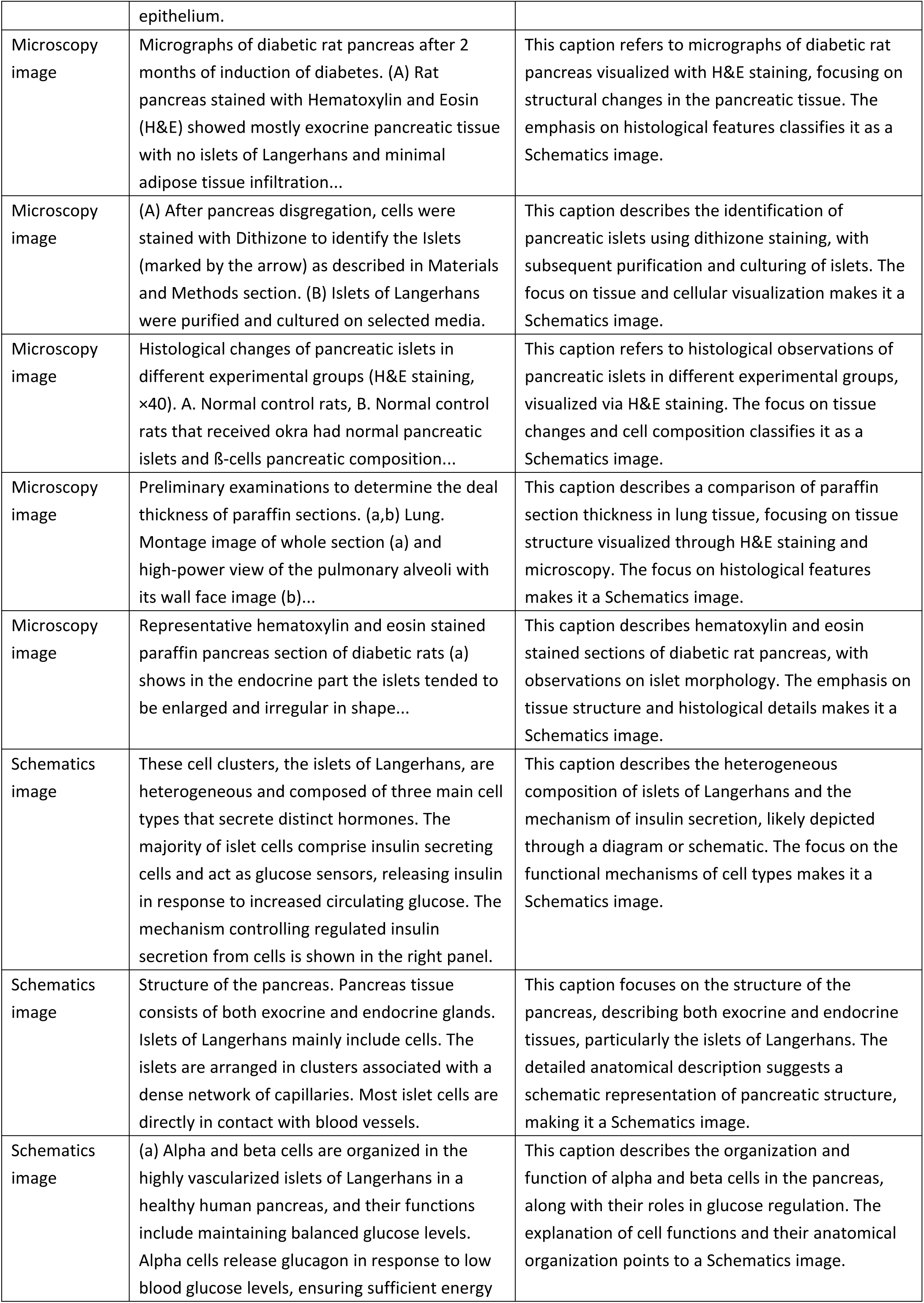
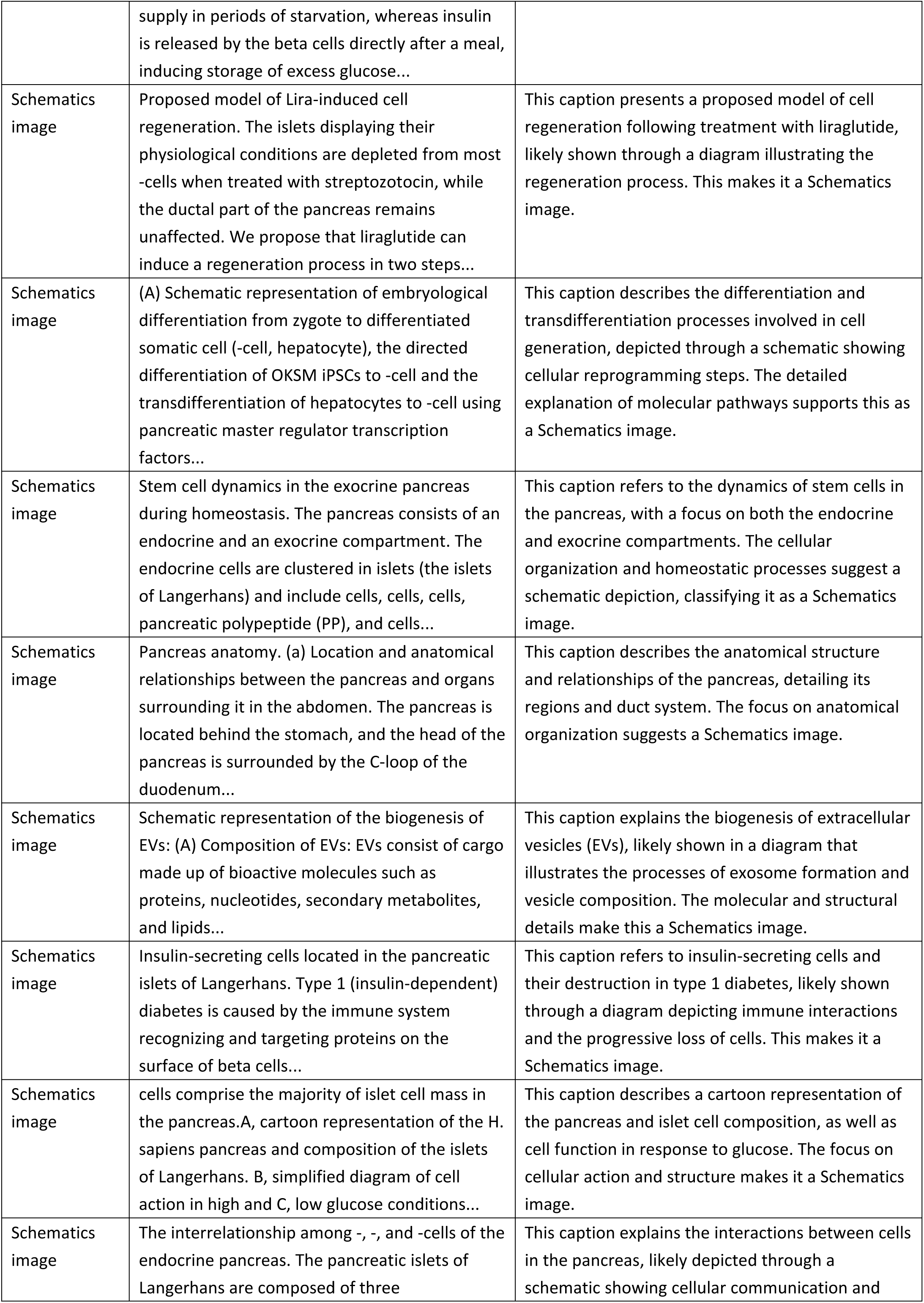
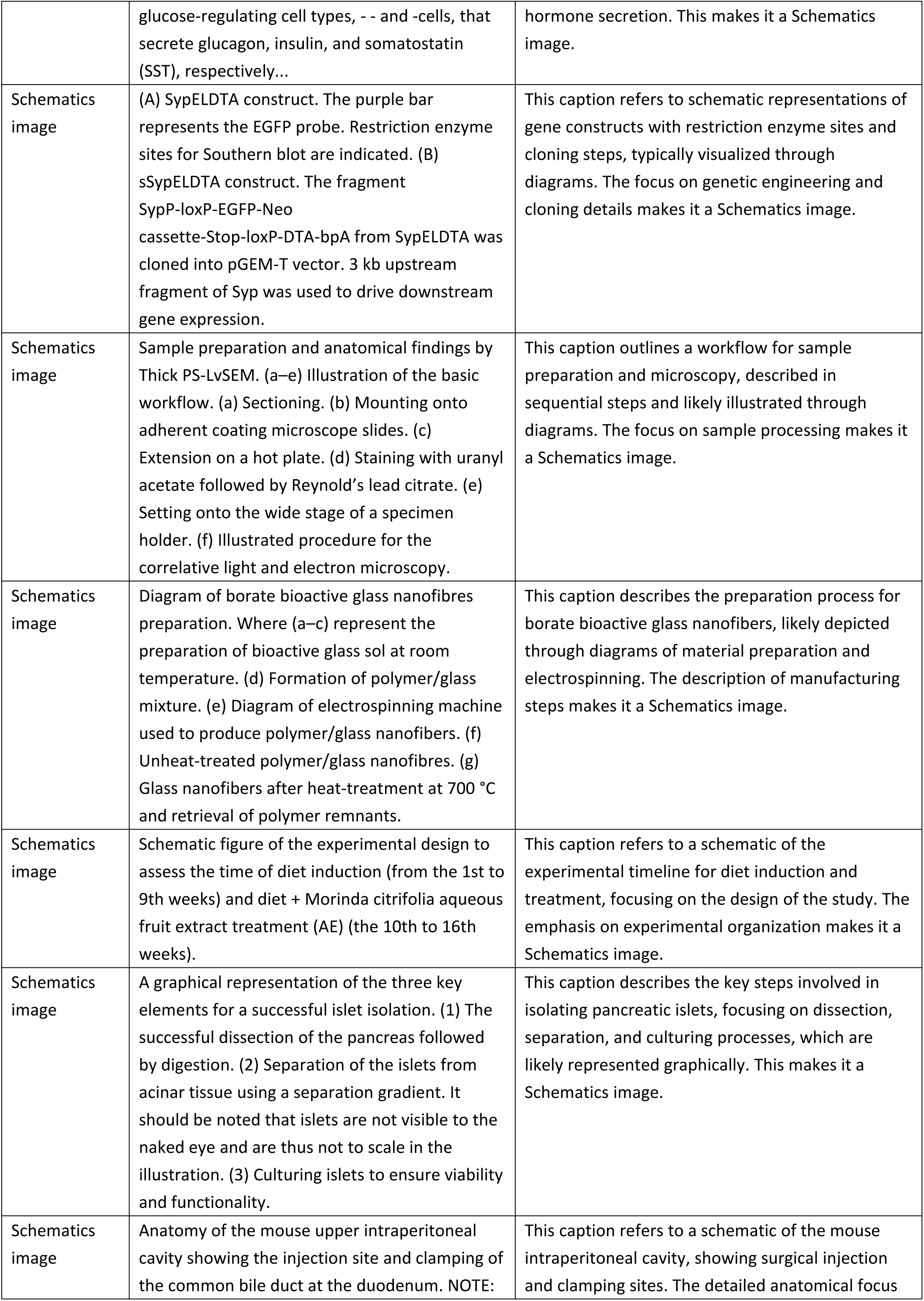
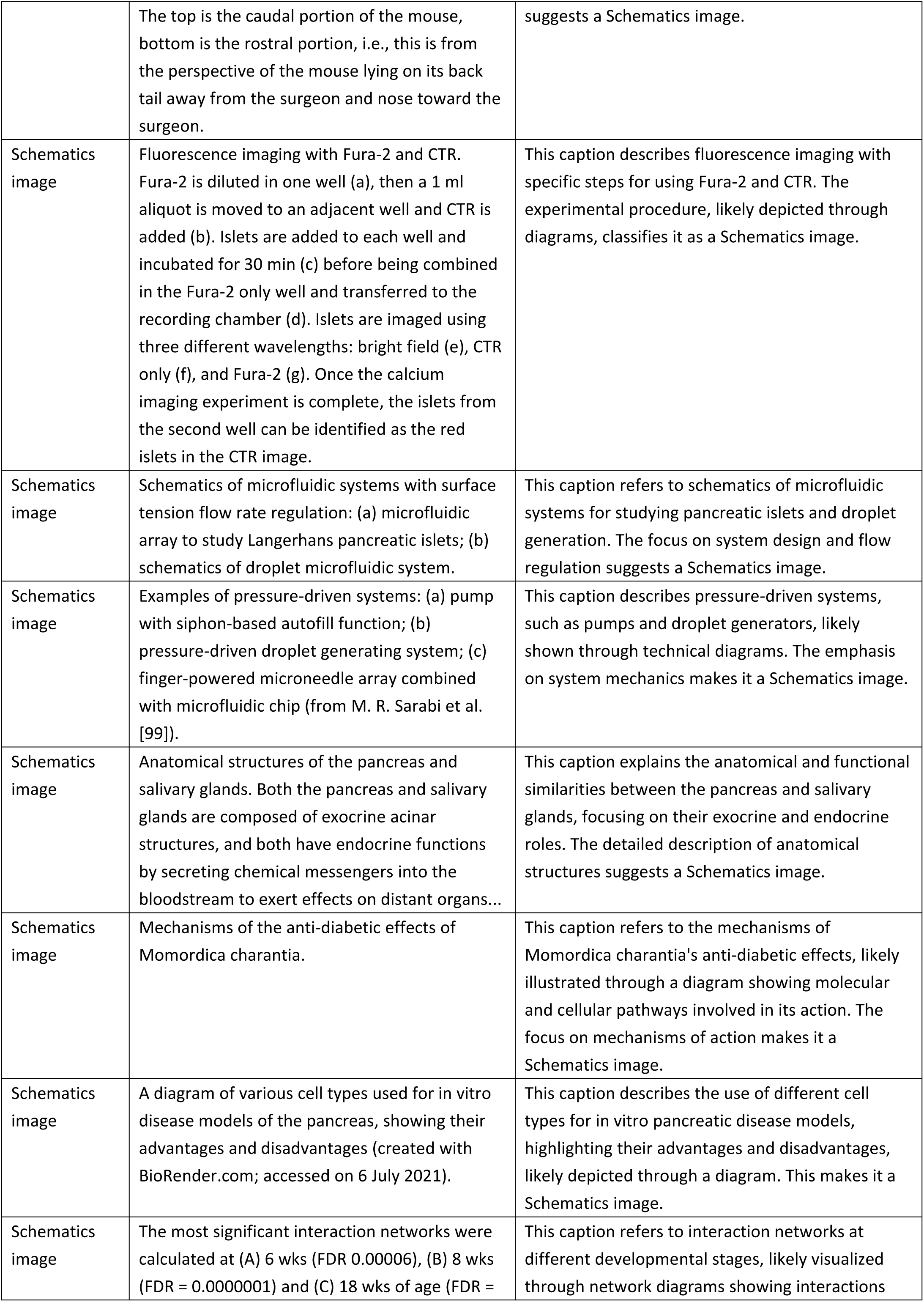
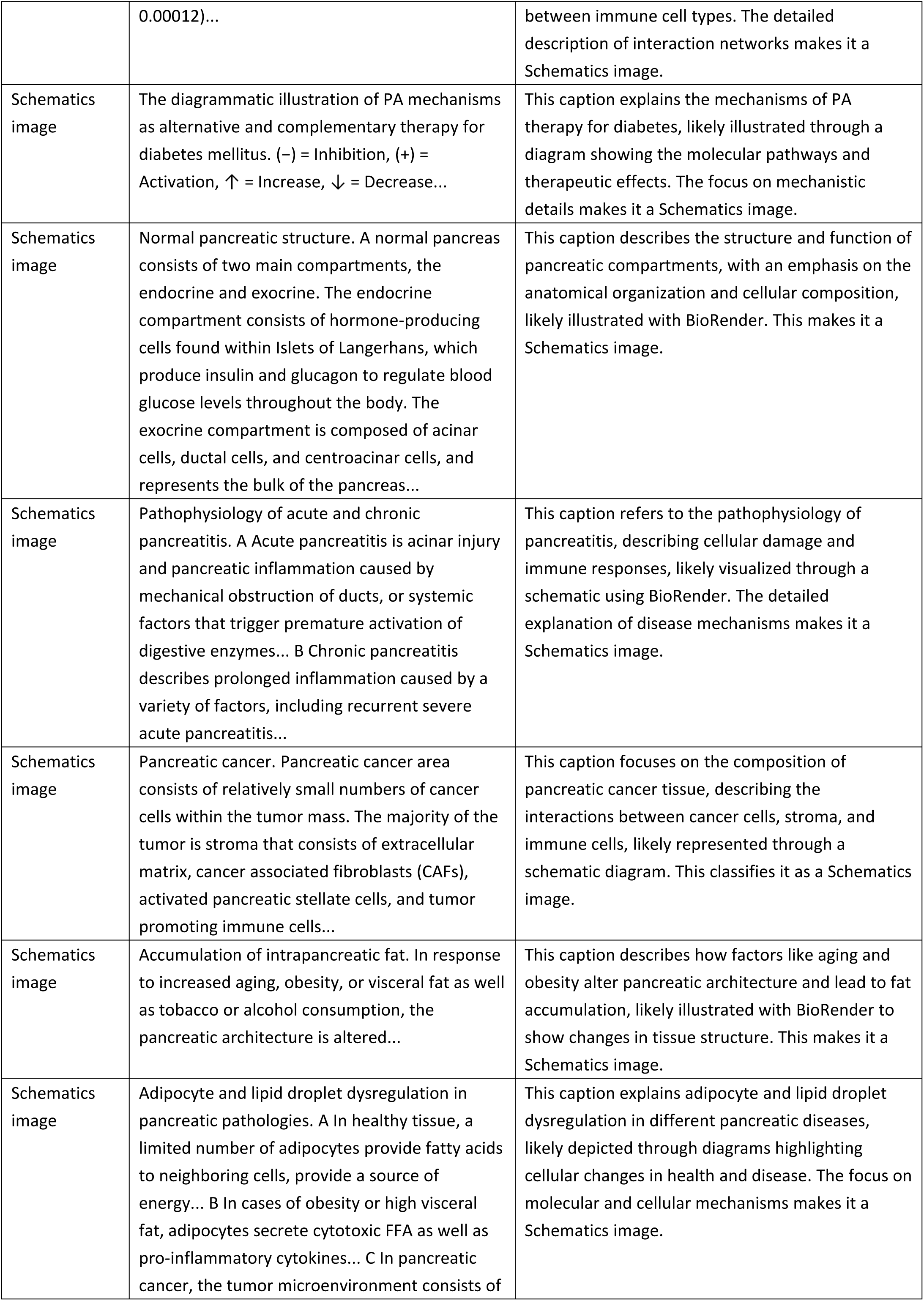
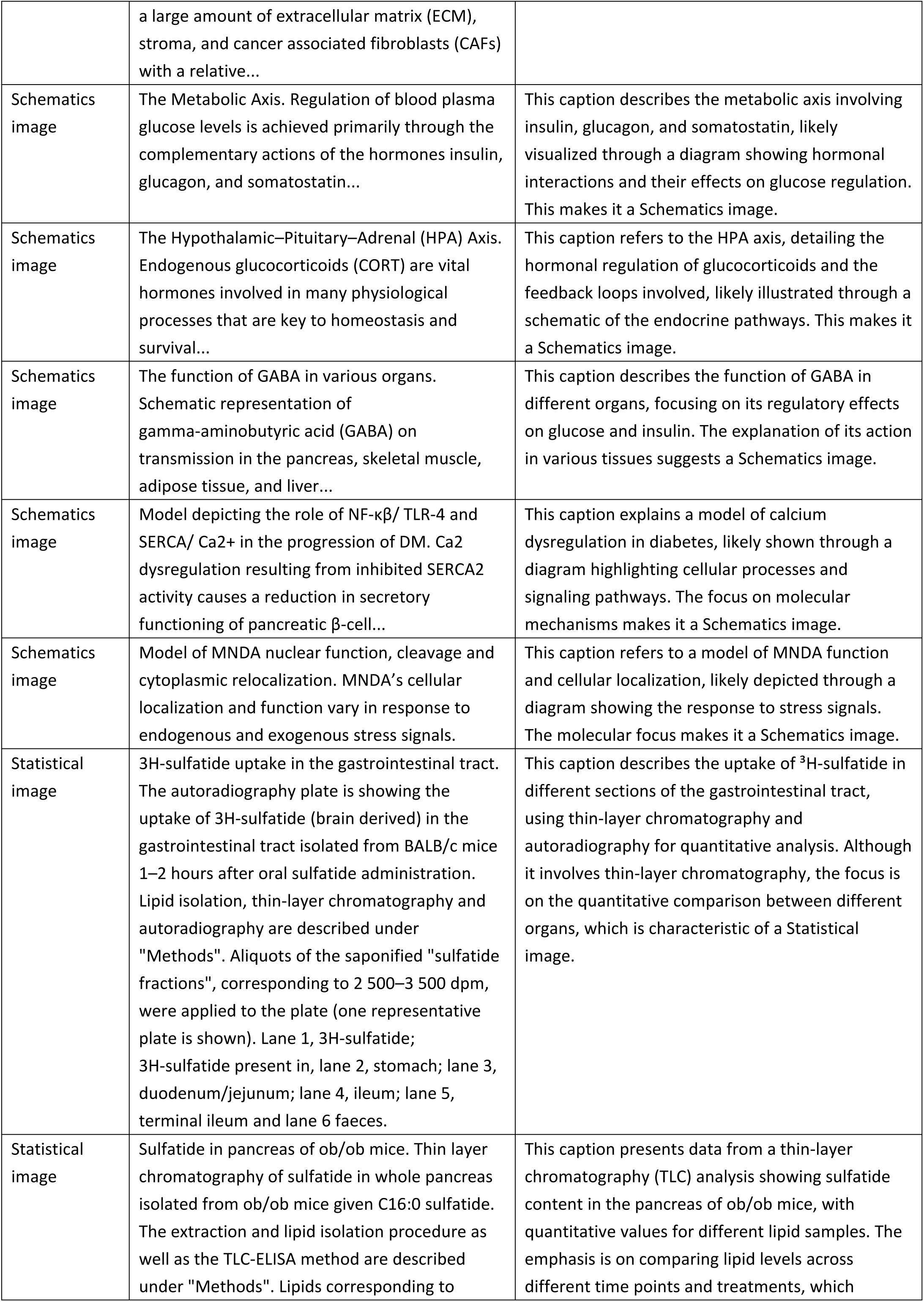
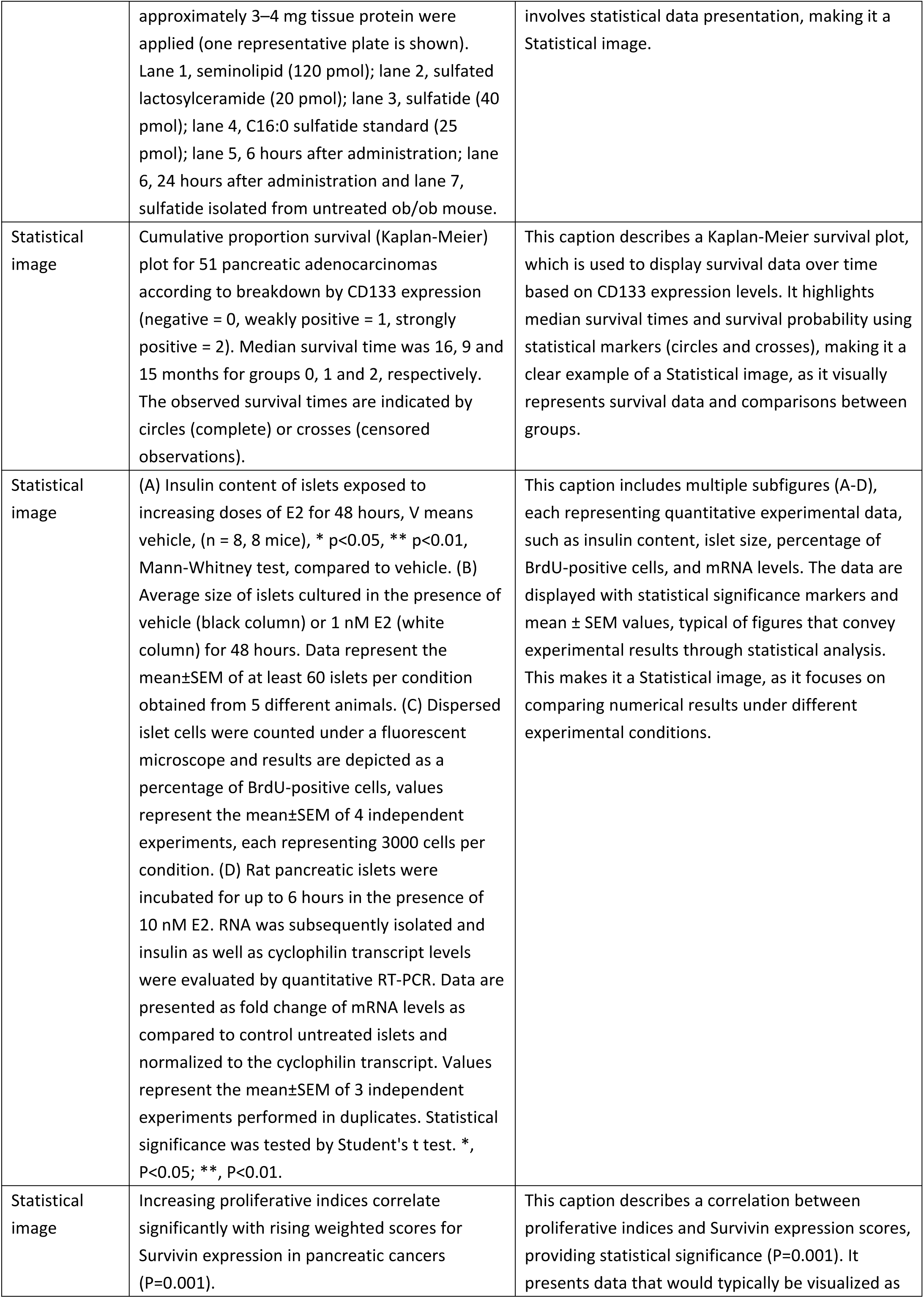
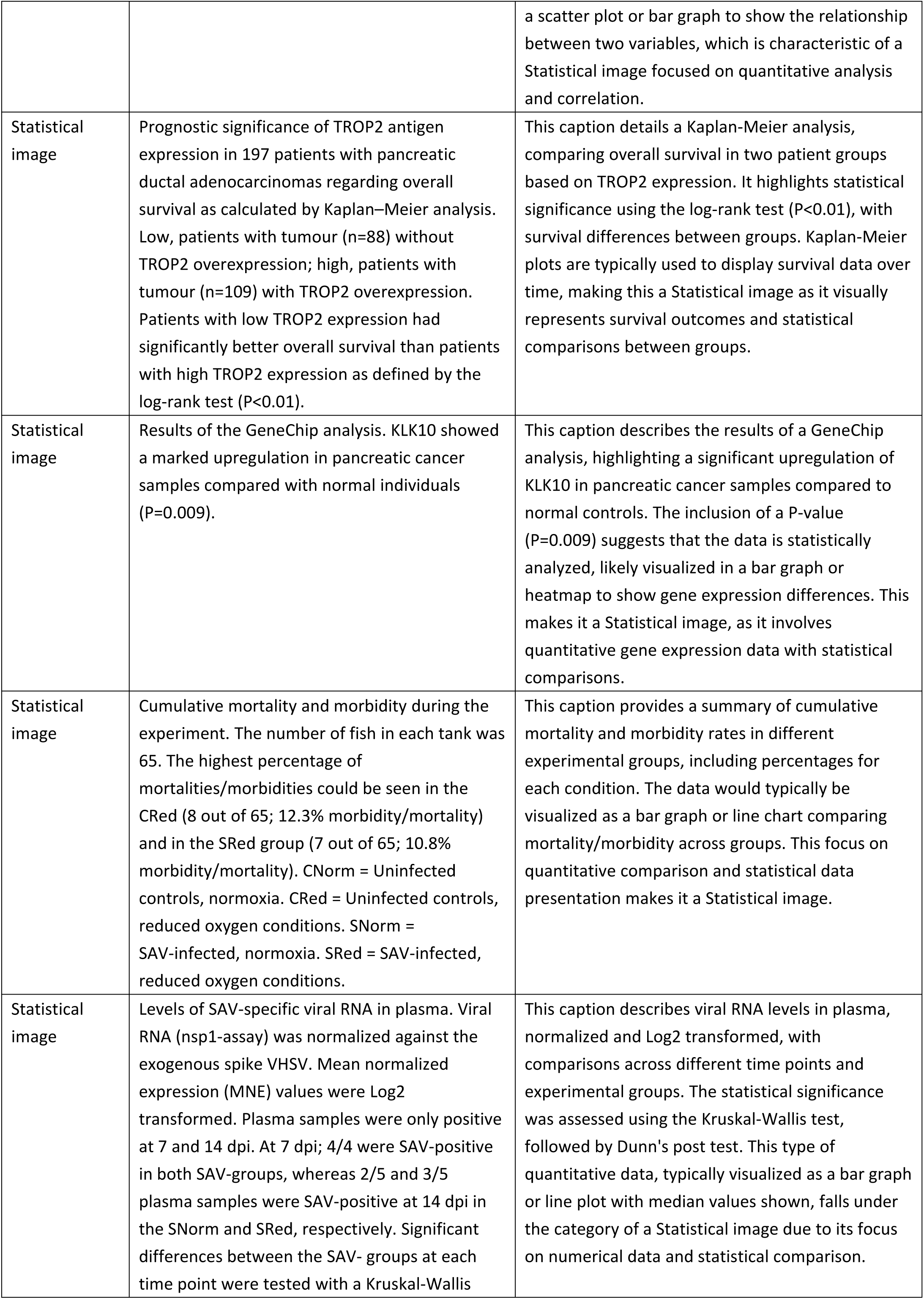
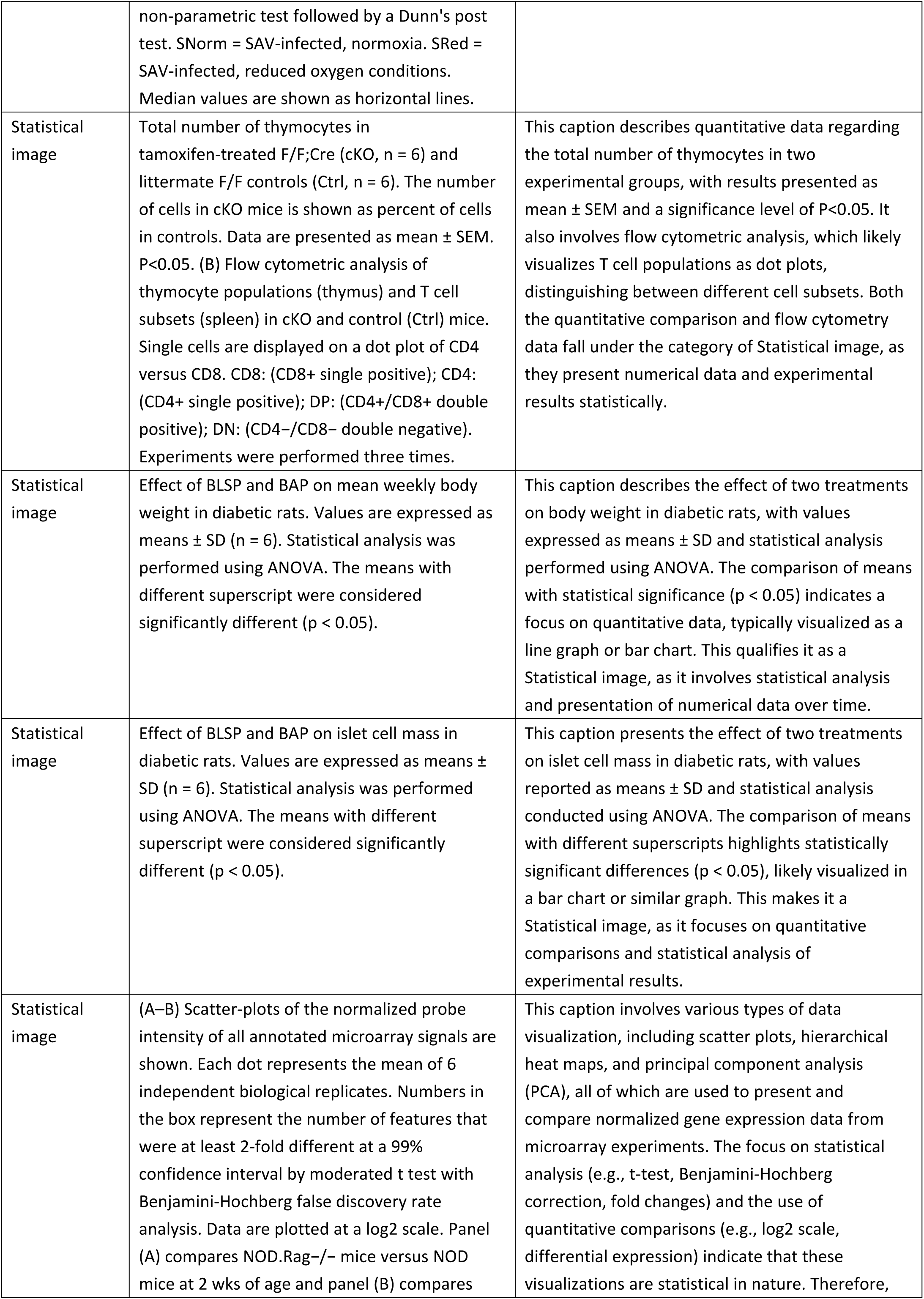
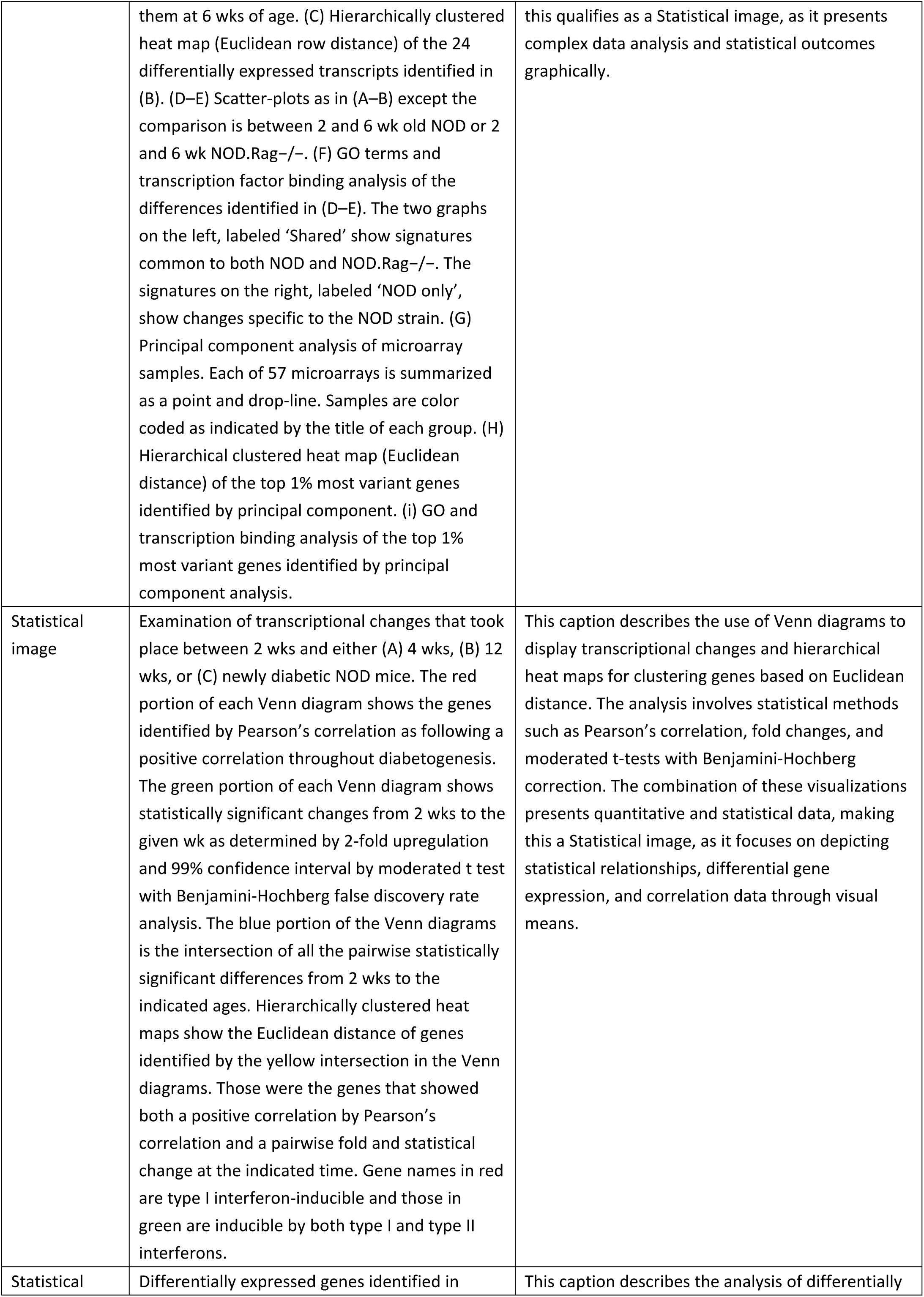
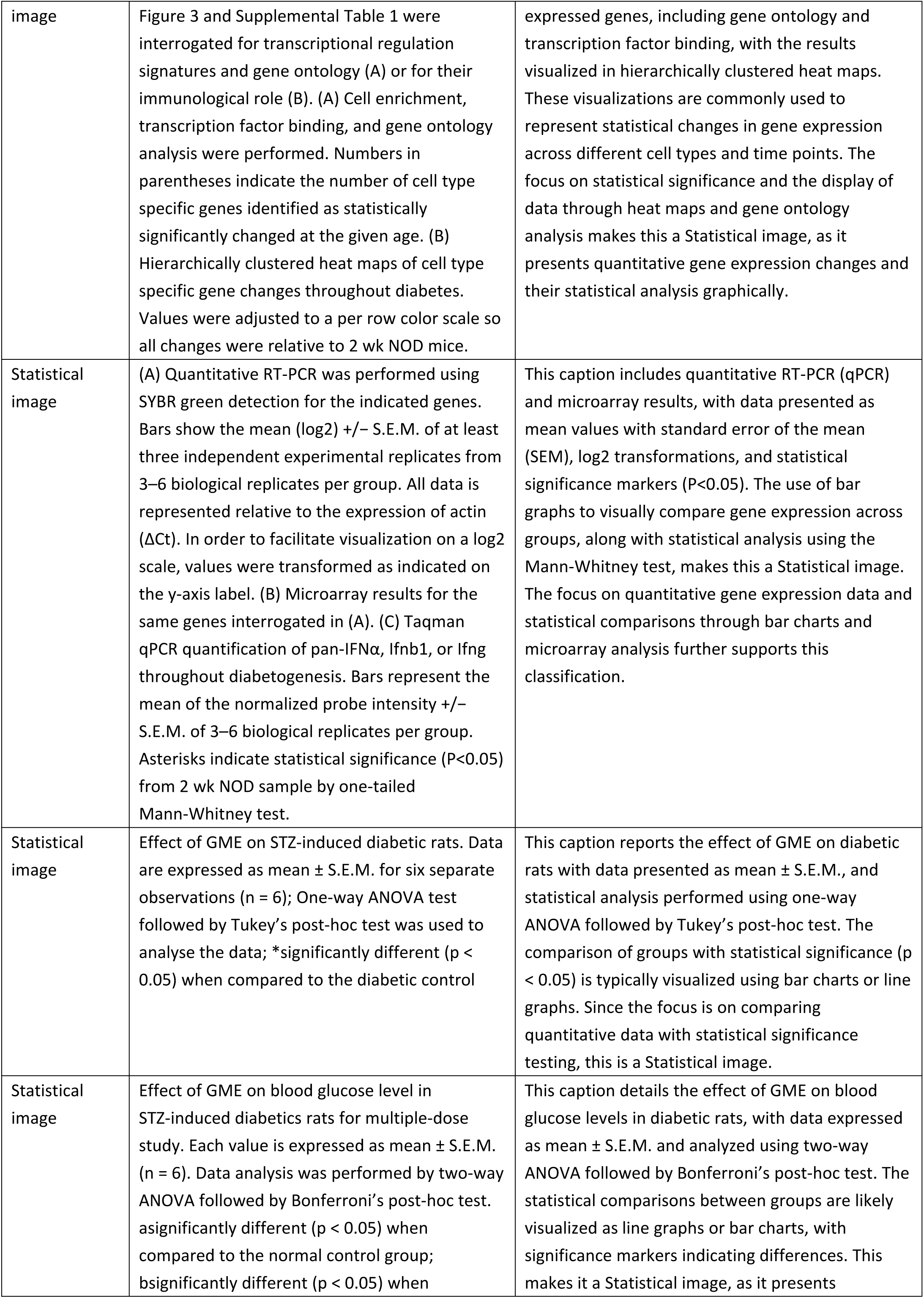
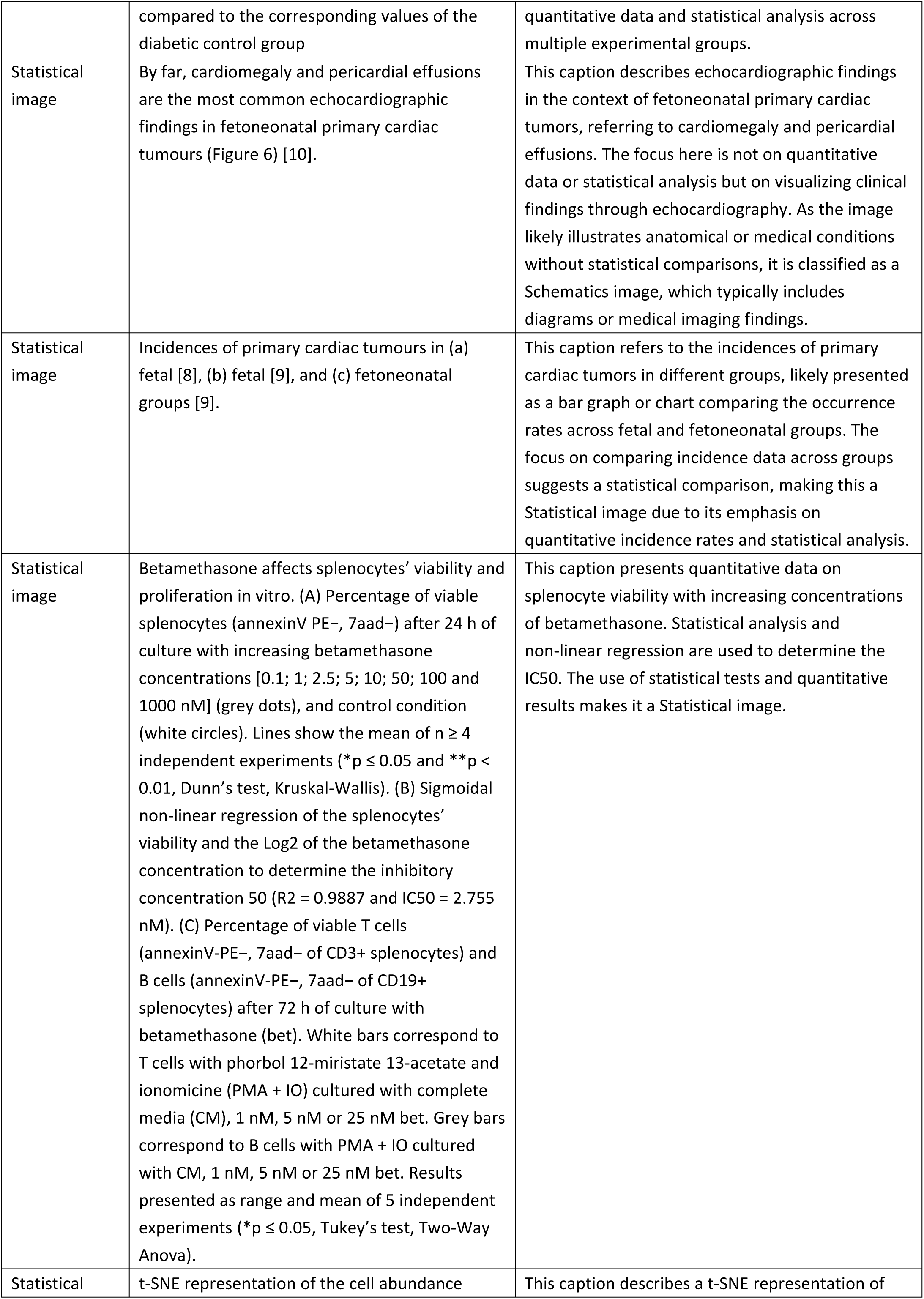
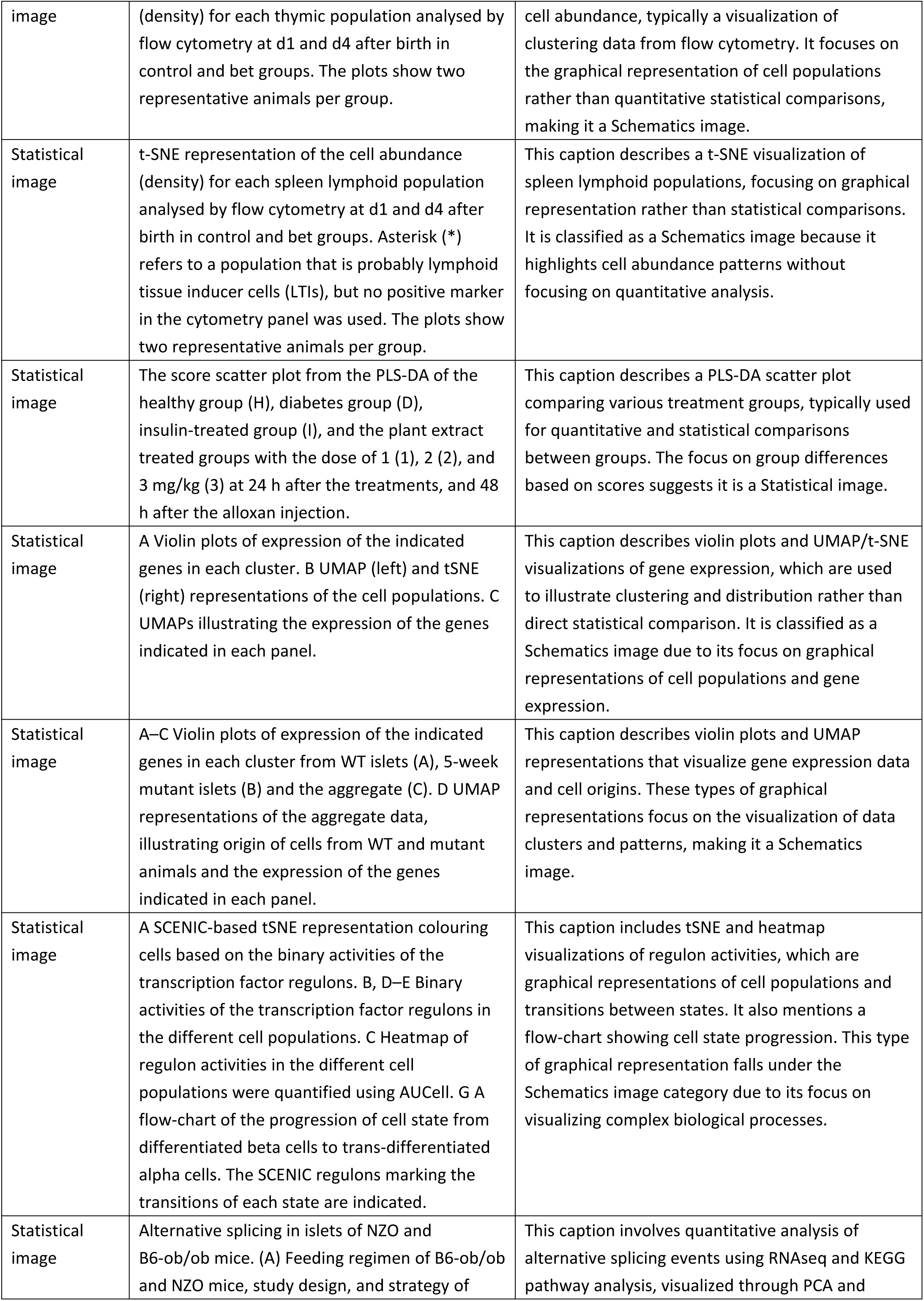
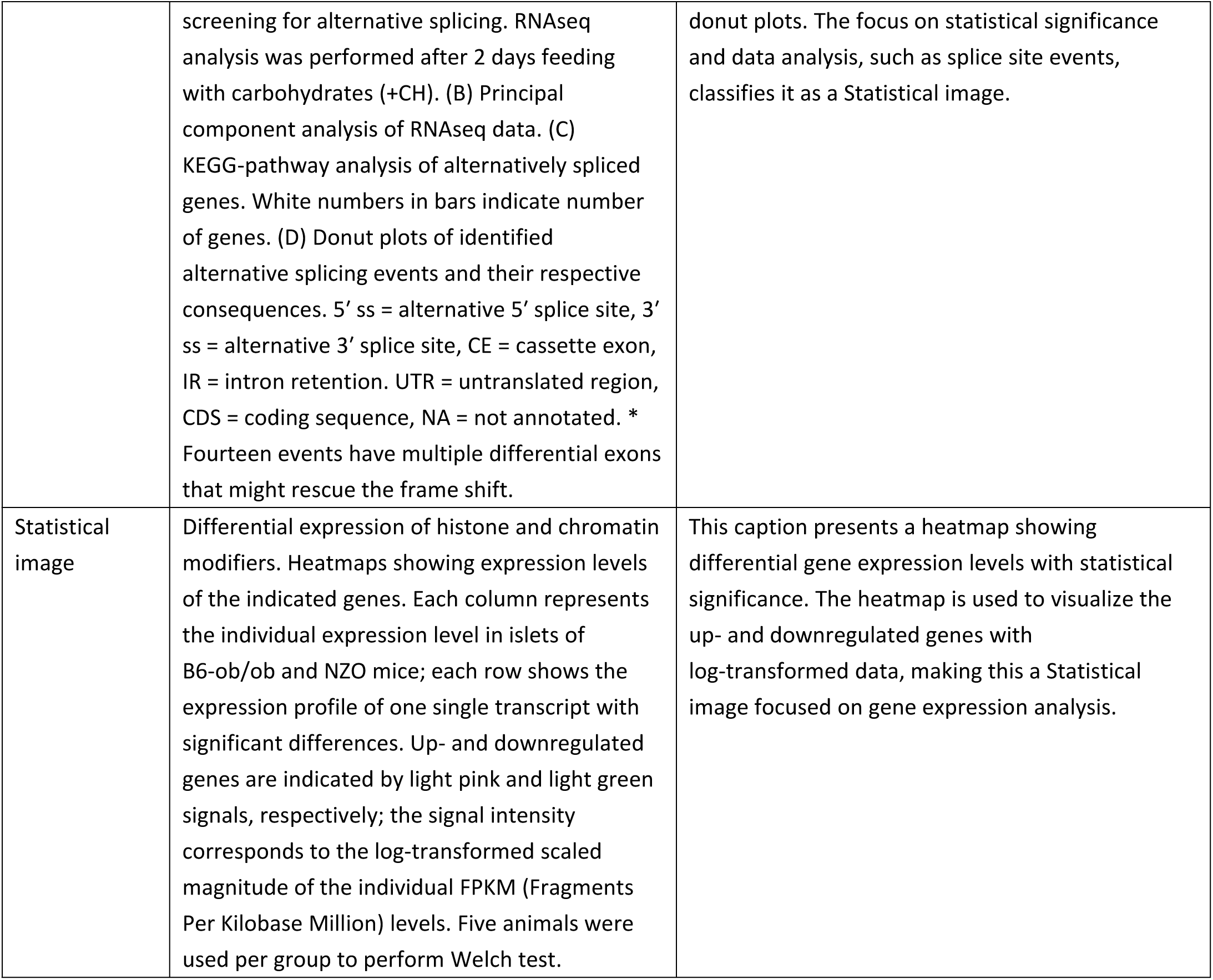
Examples of figure caption across six image types.

**Supplementary Table 4.**
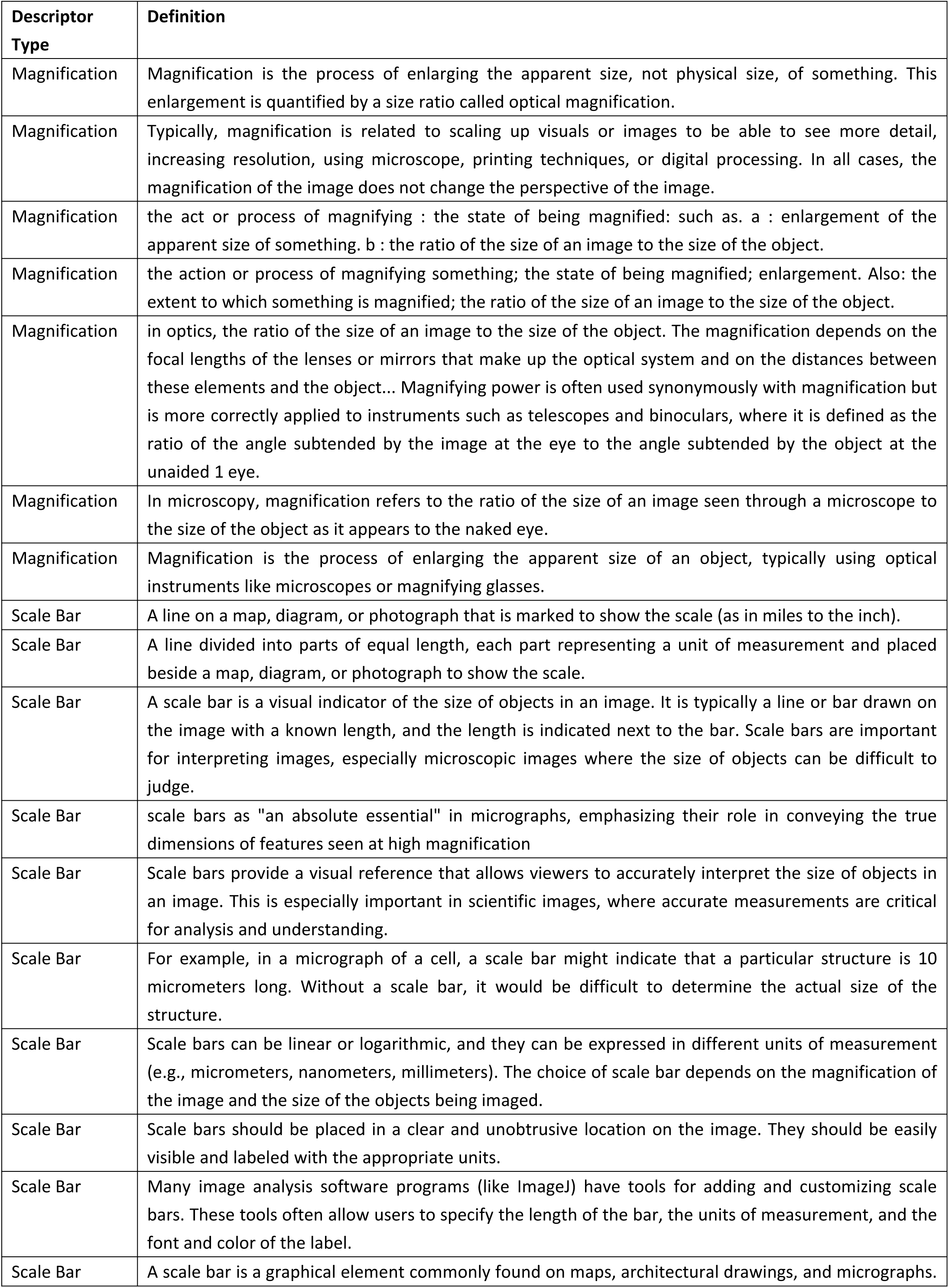
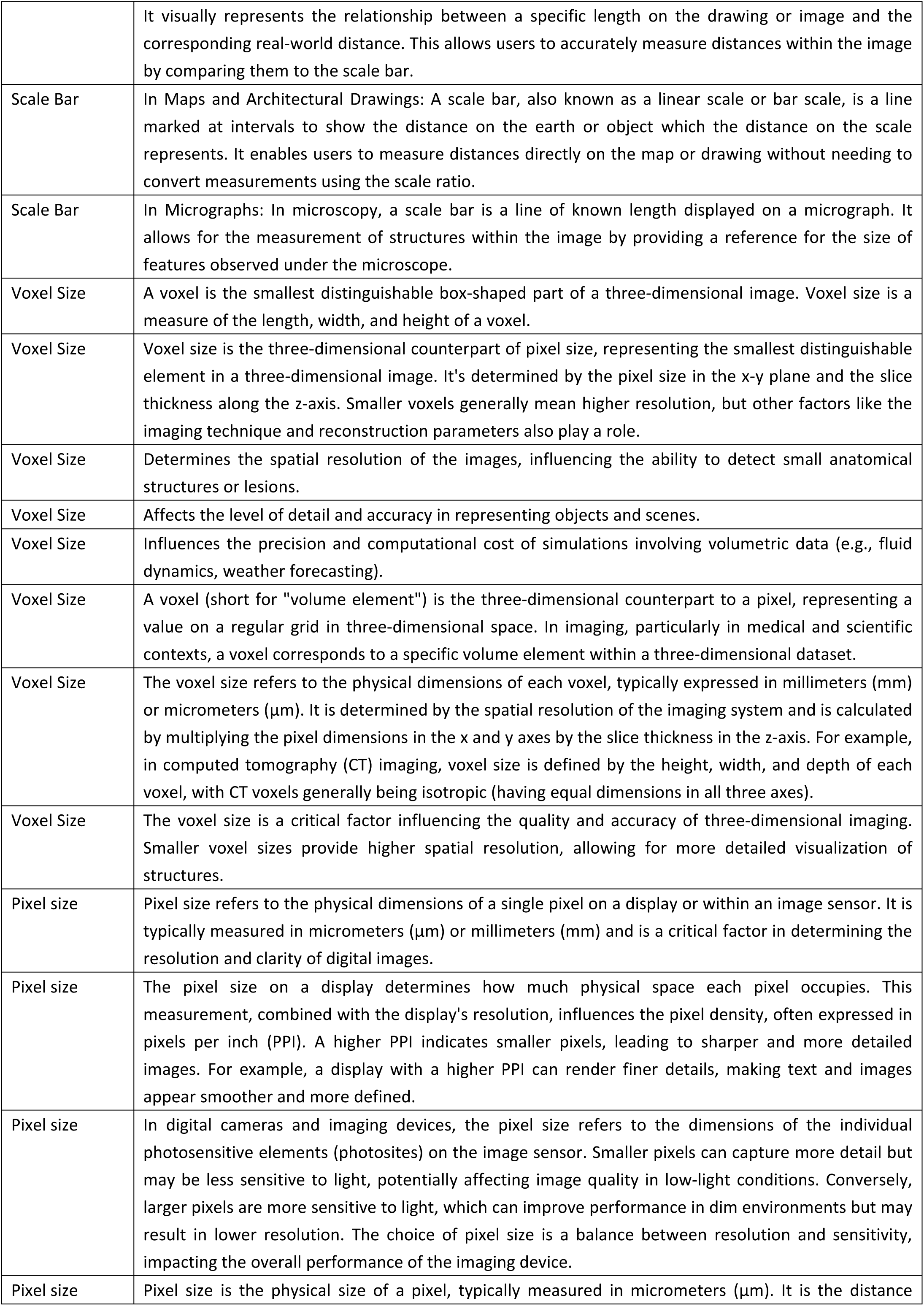
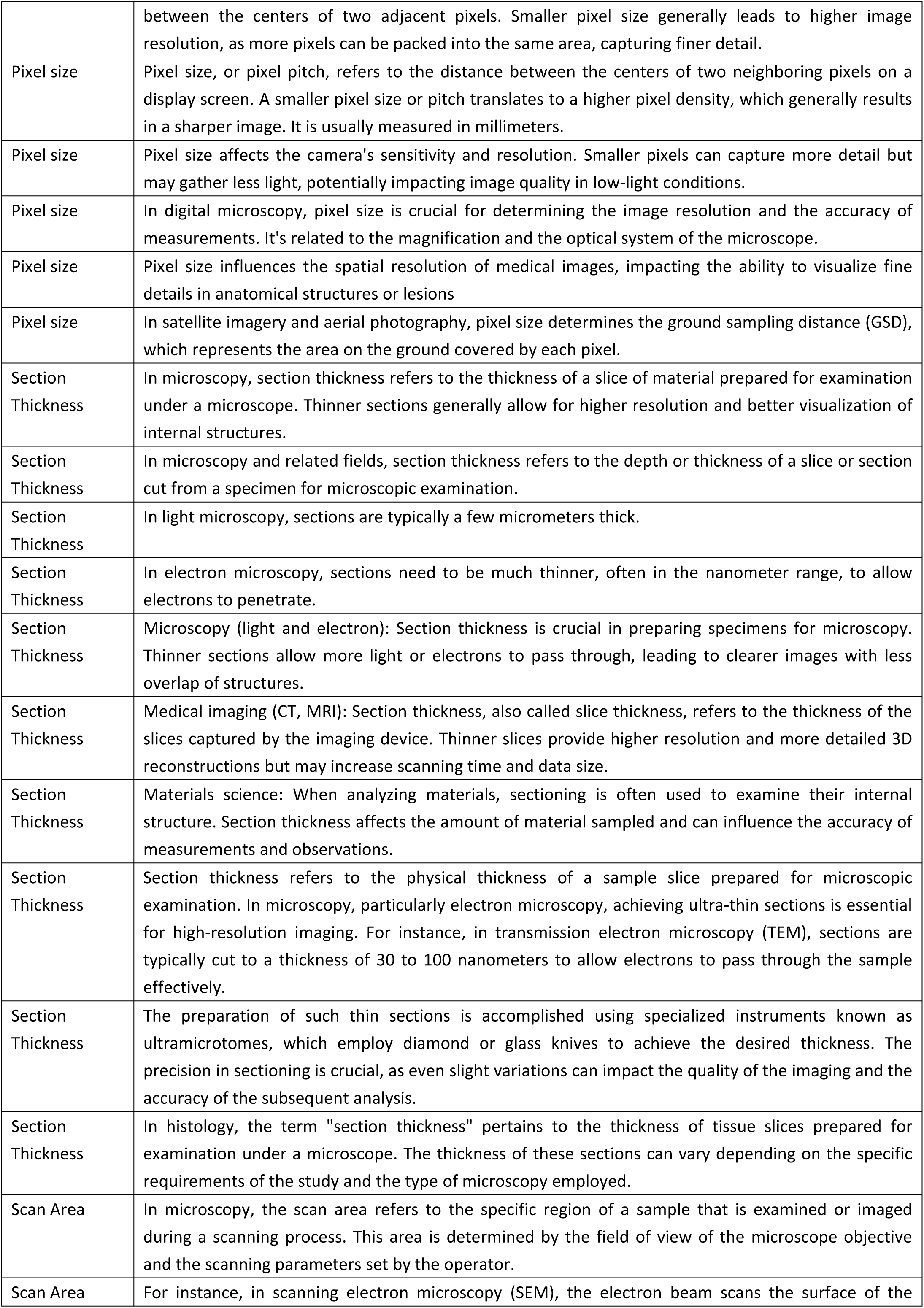
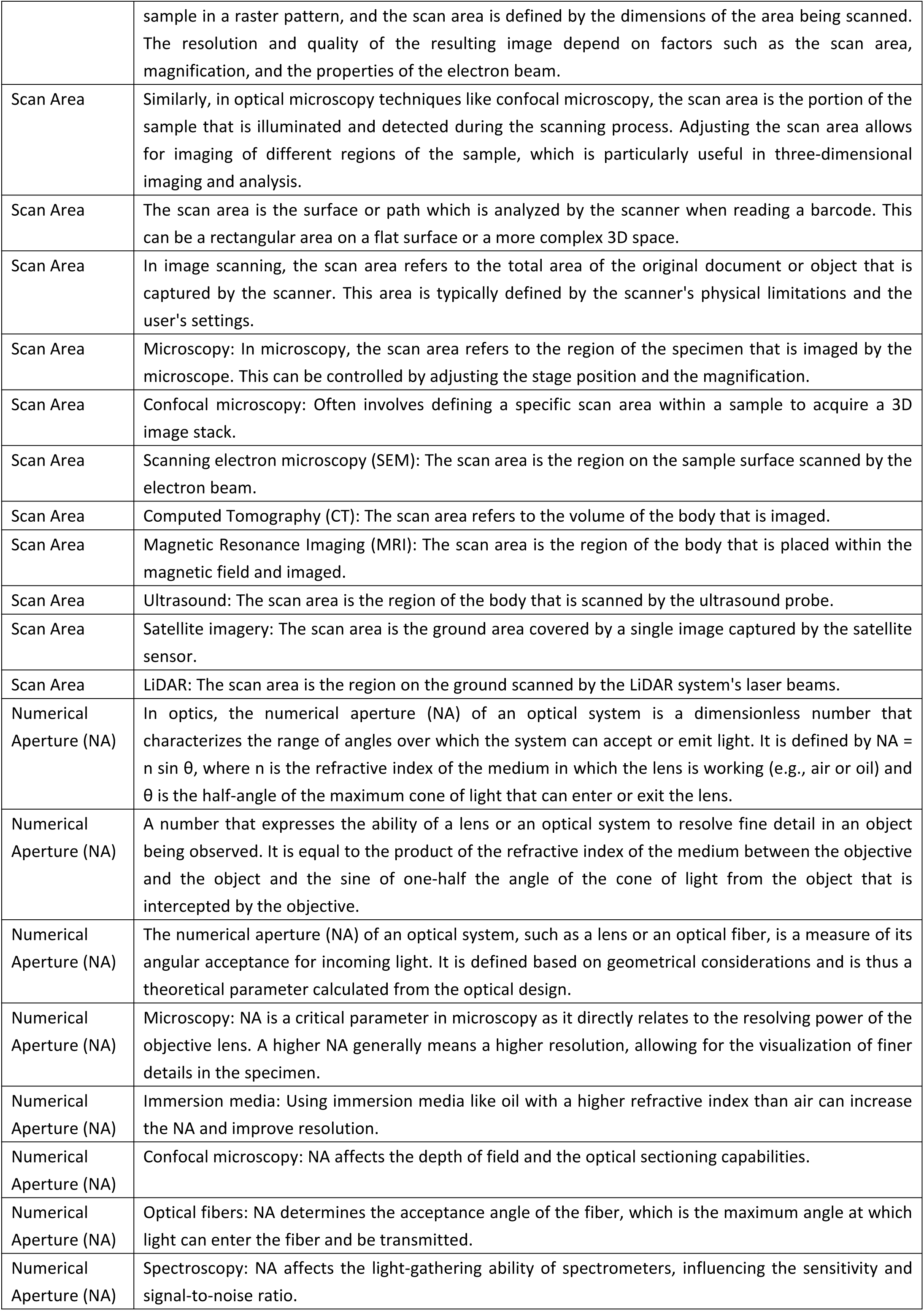
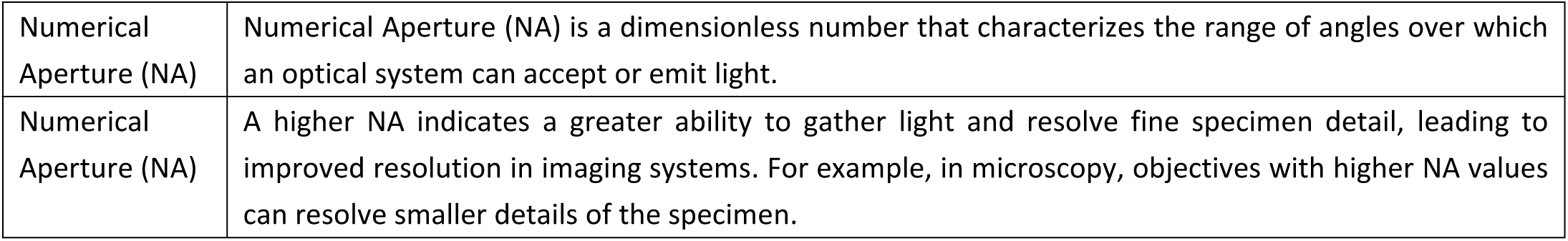
Definitions of microscopy acquisition parameters.

**Supplementary Table 5.**
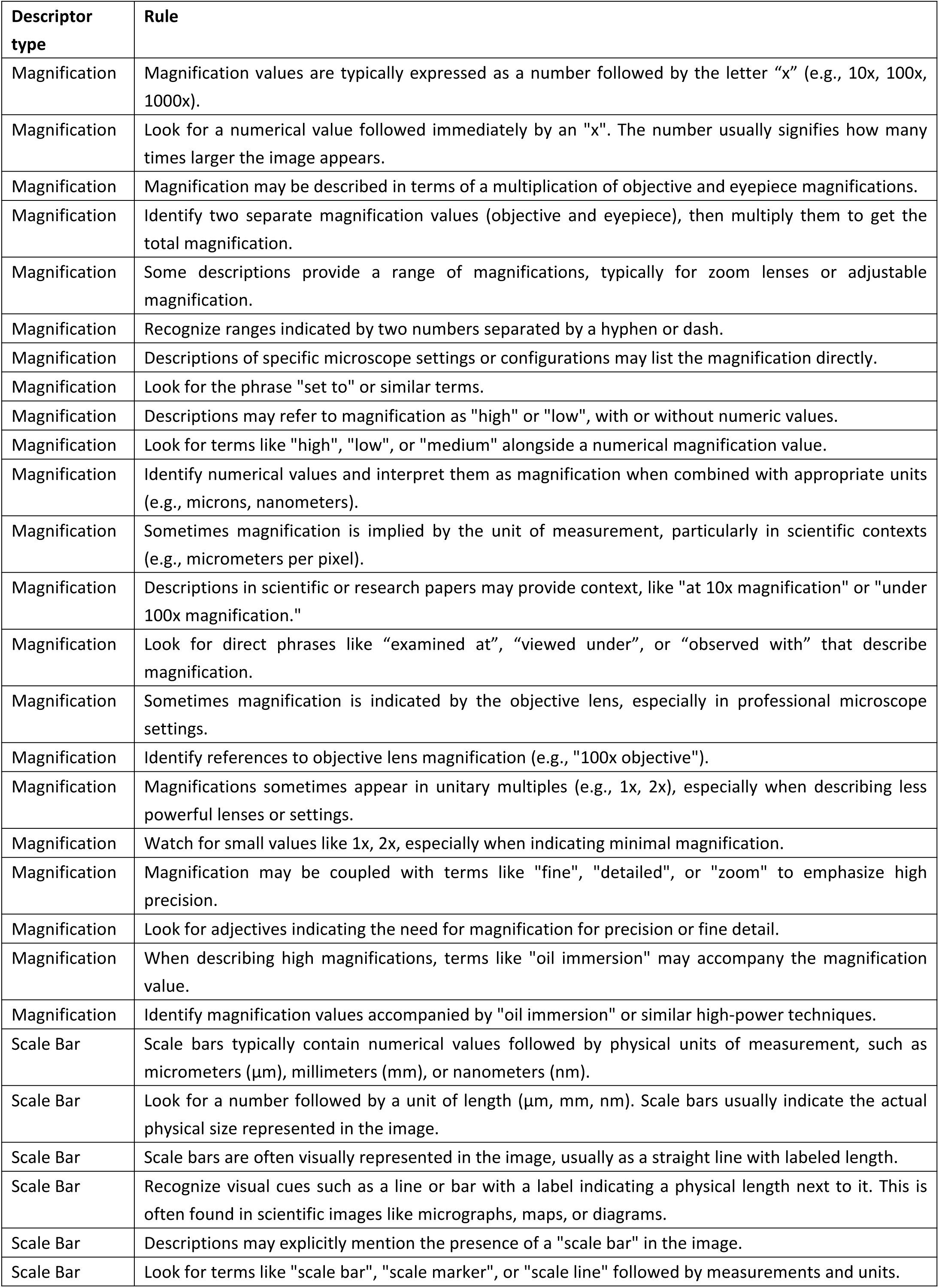
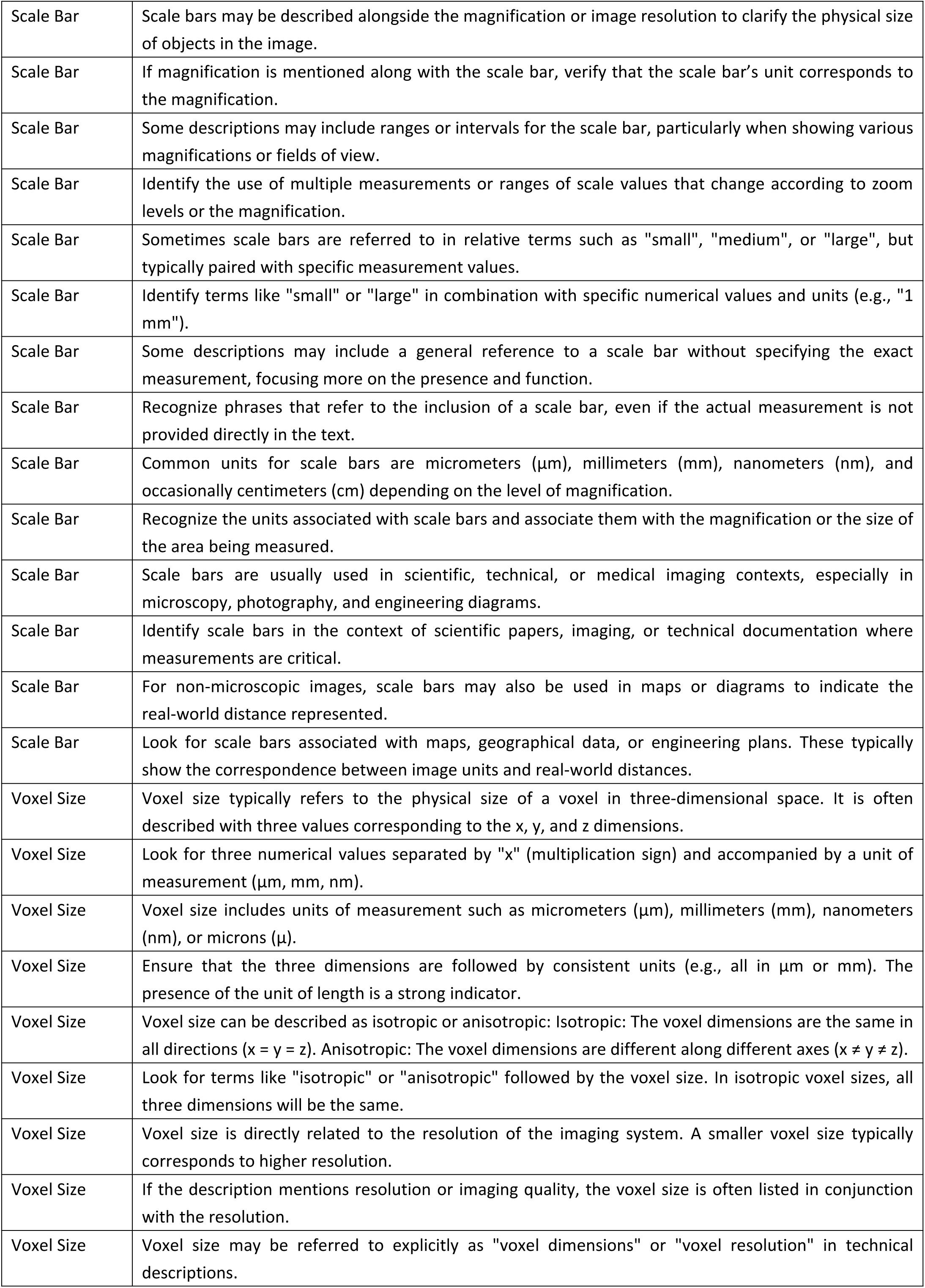
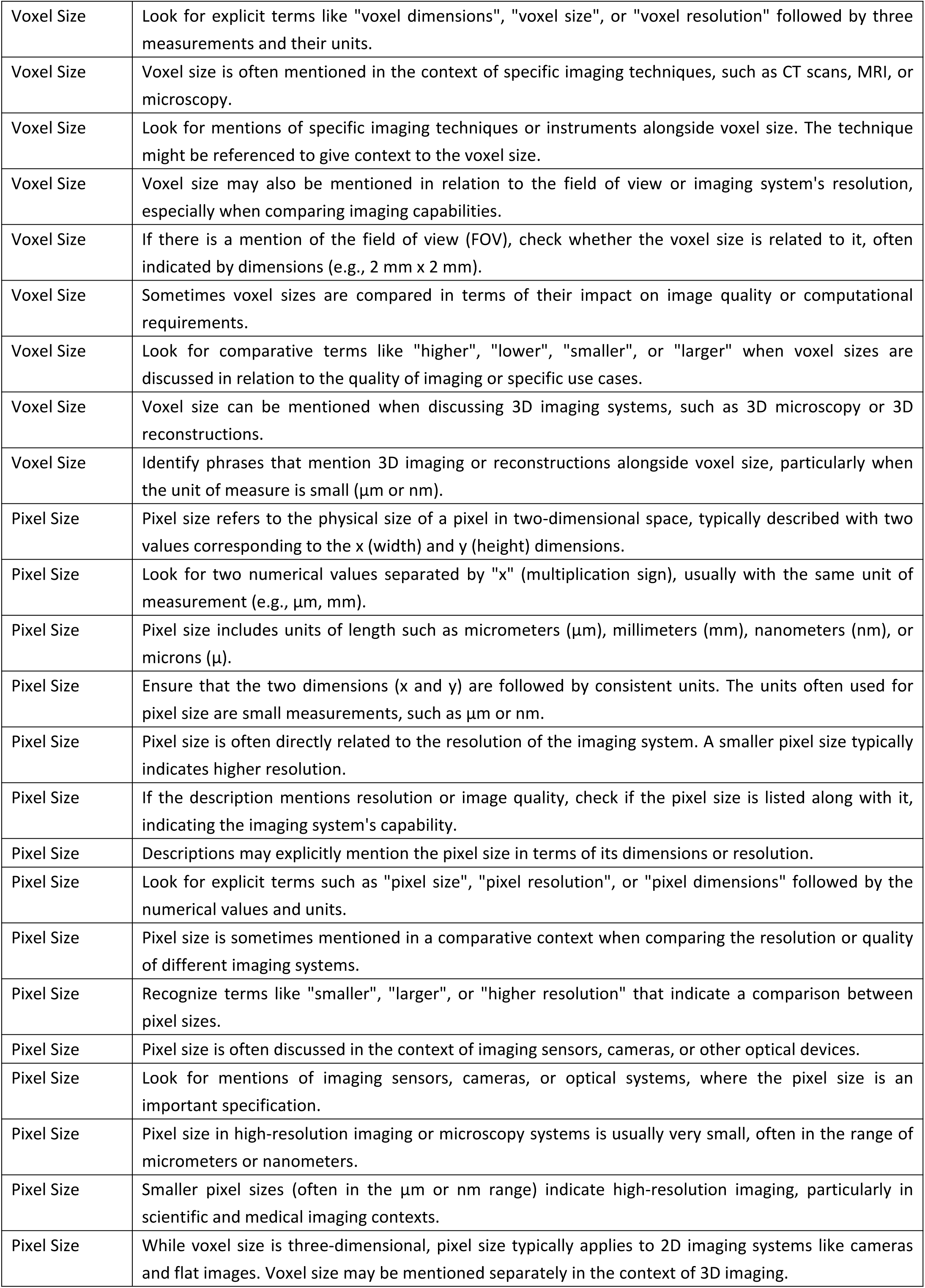
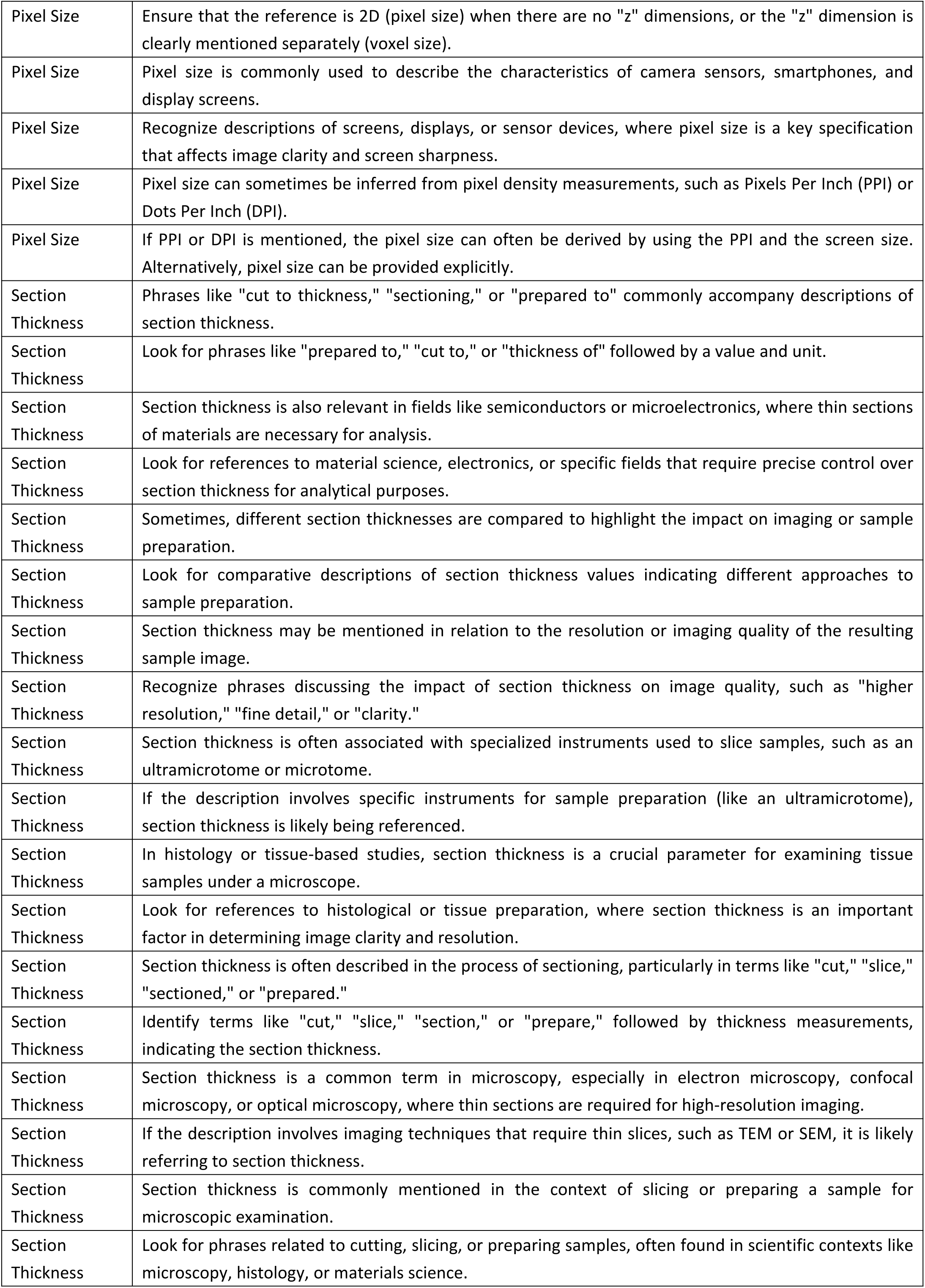
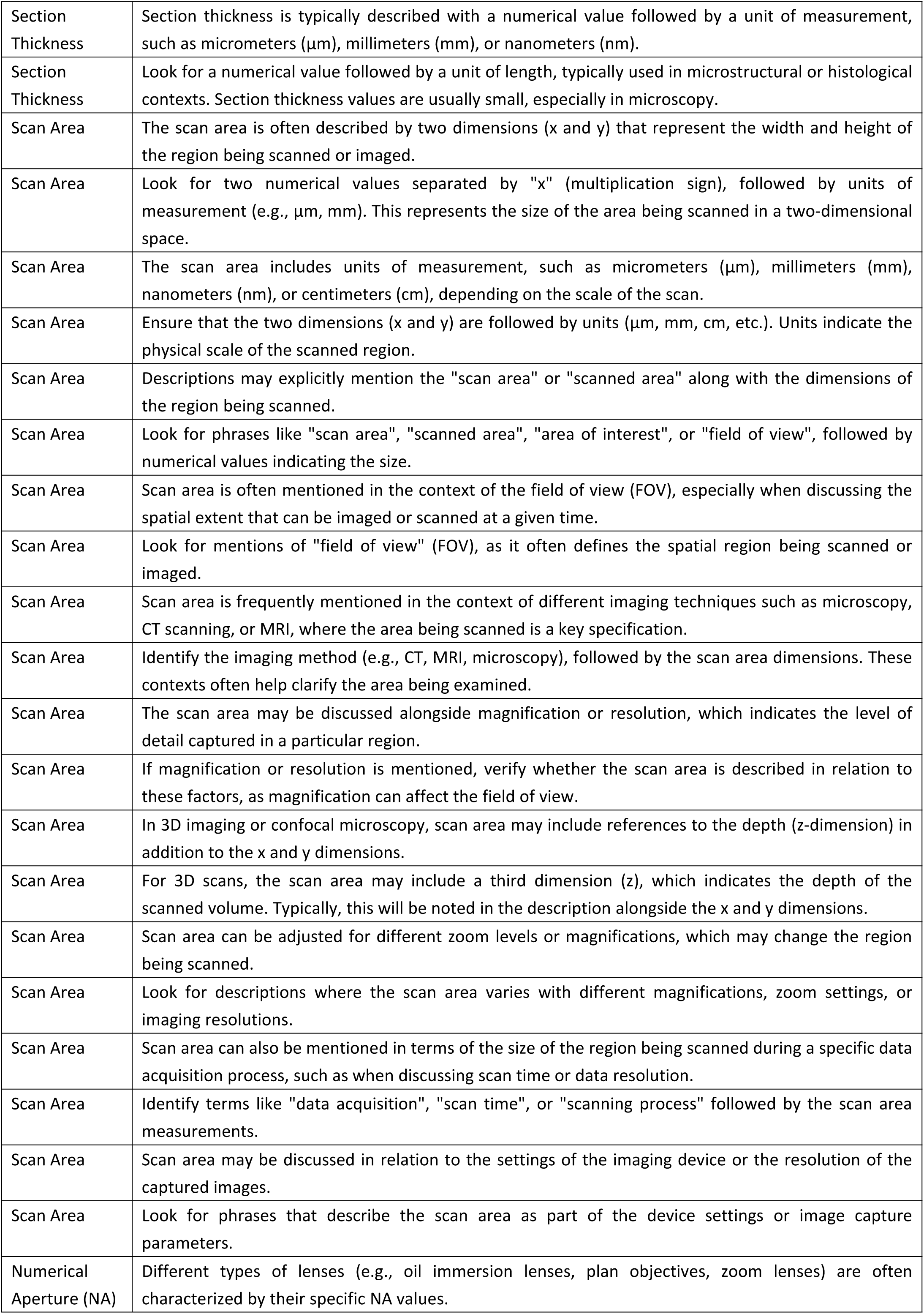
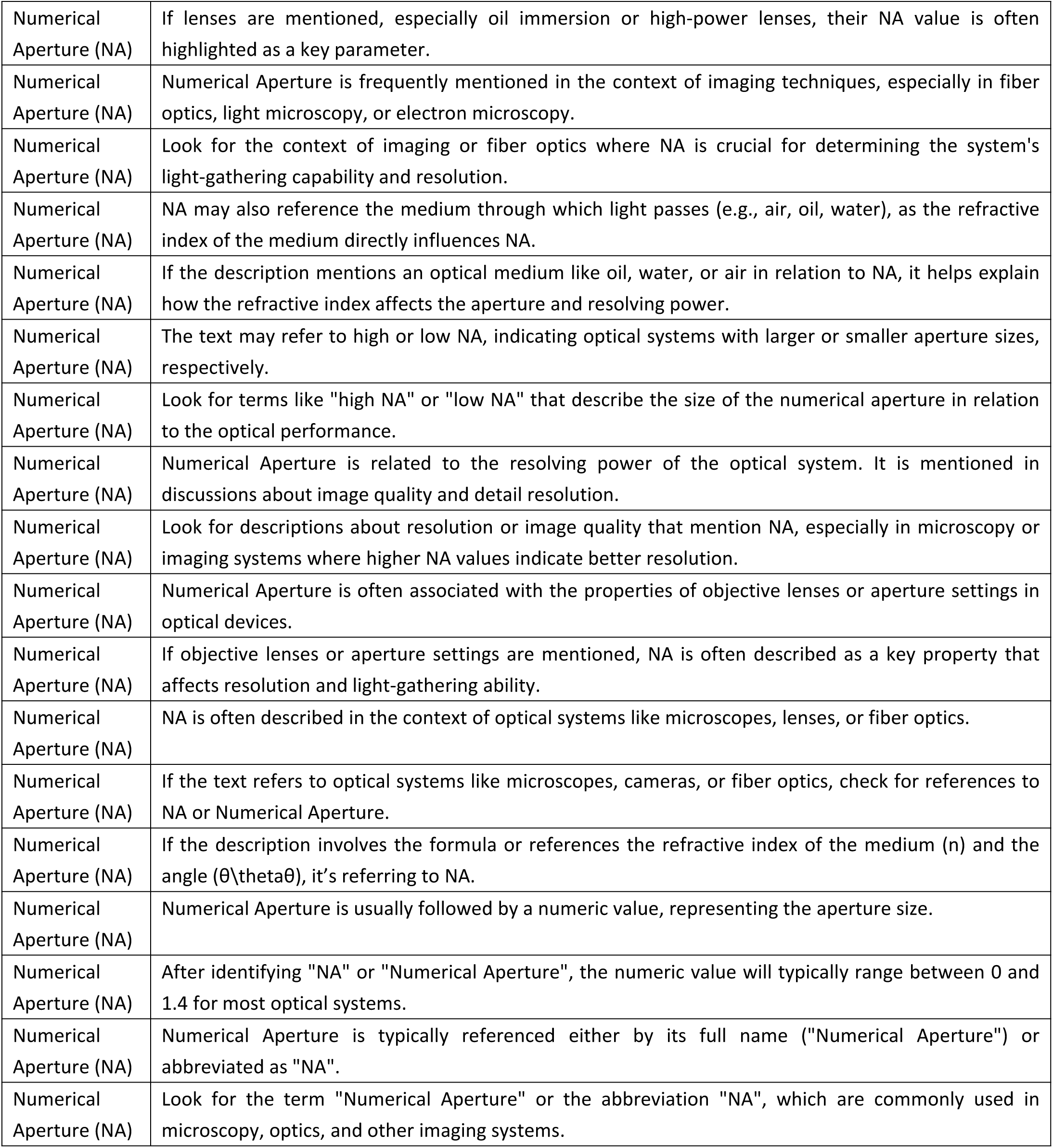
Definitions of microscopy acquisition parameters.

**Supplementary Table 6.**
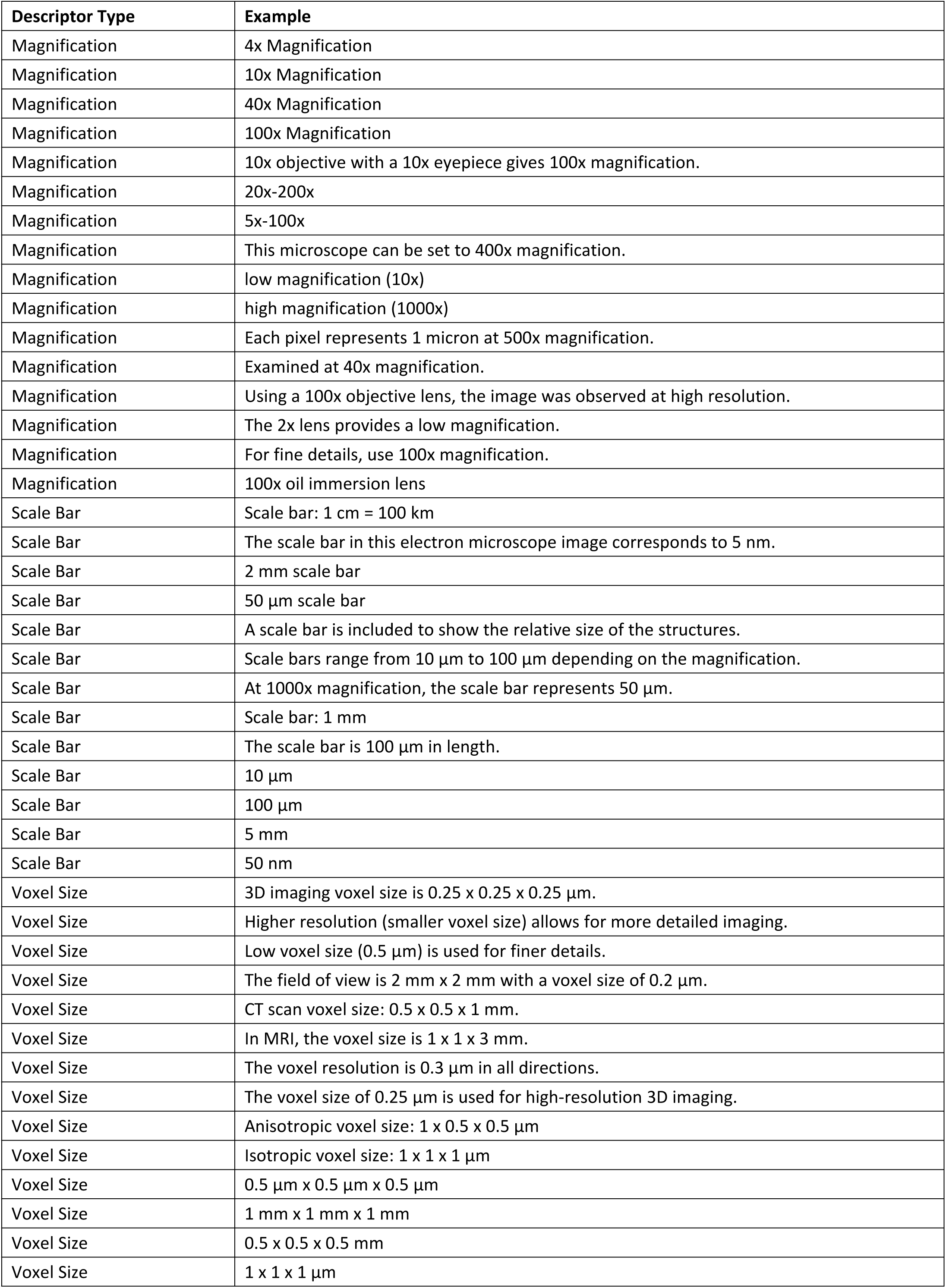
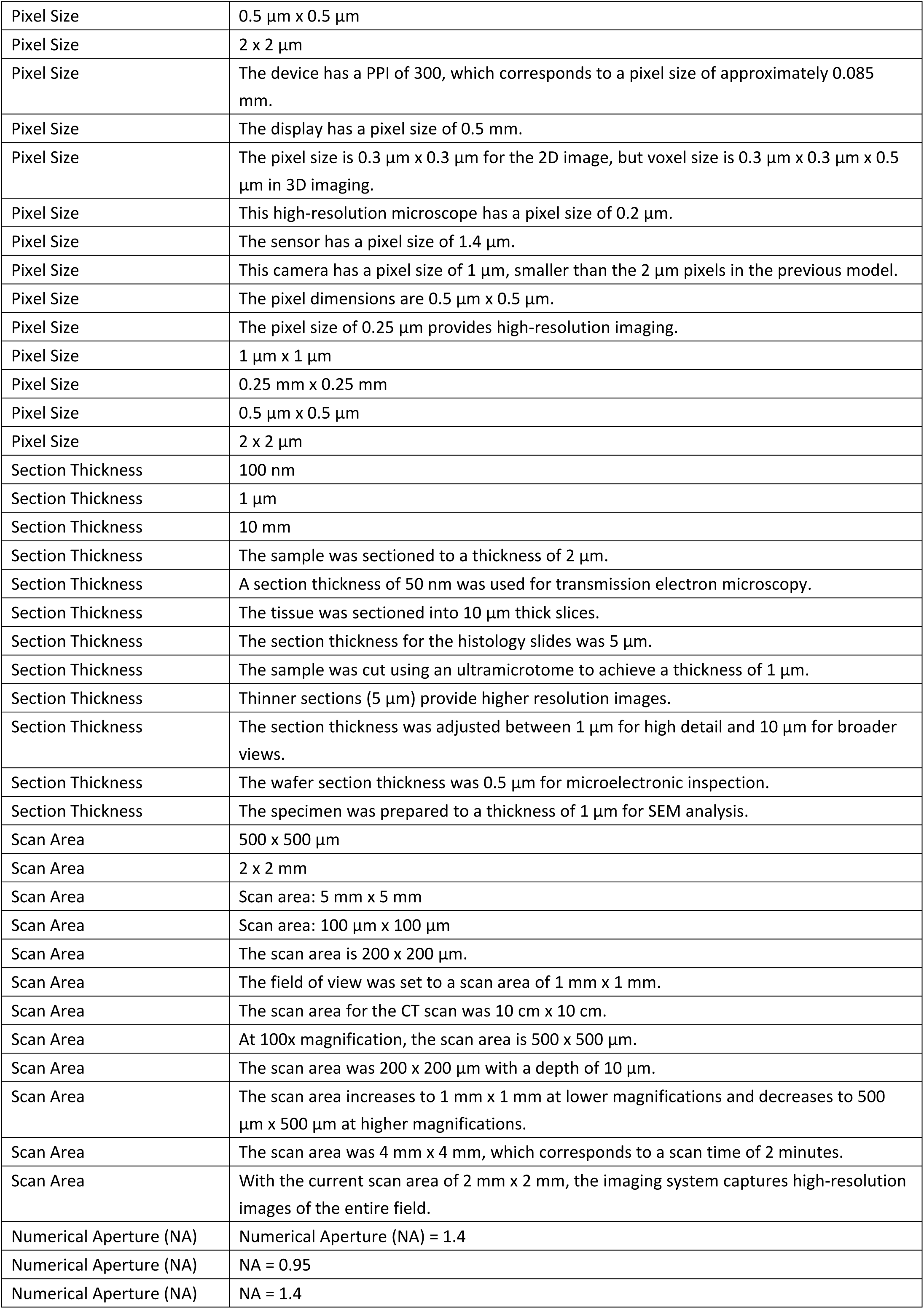
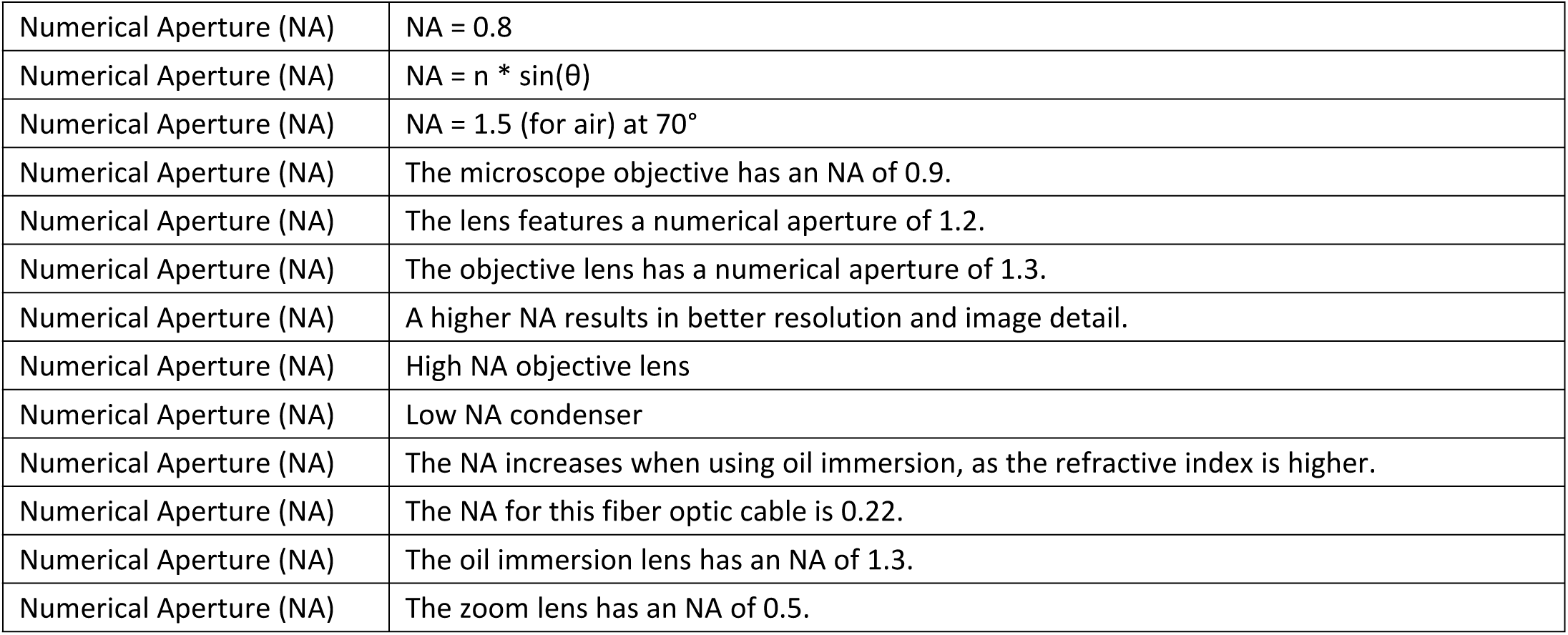
Examples of microscopy acquisition parameters.

**Supplementary Table 7.**
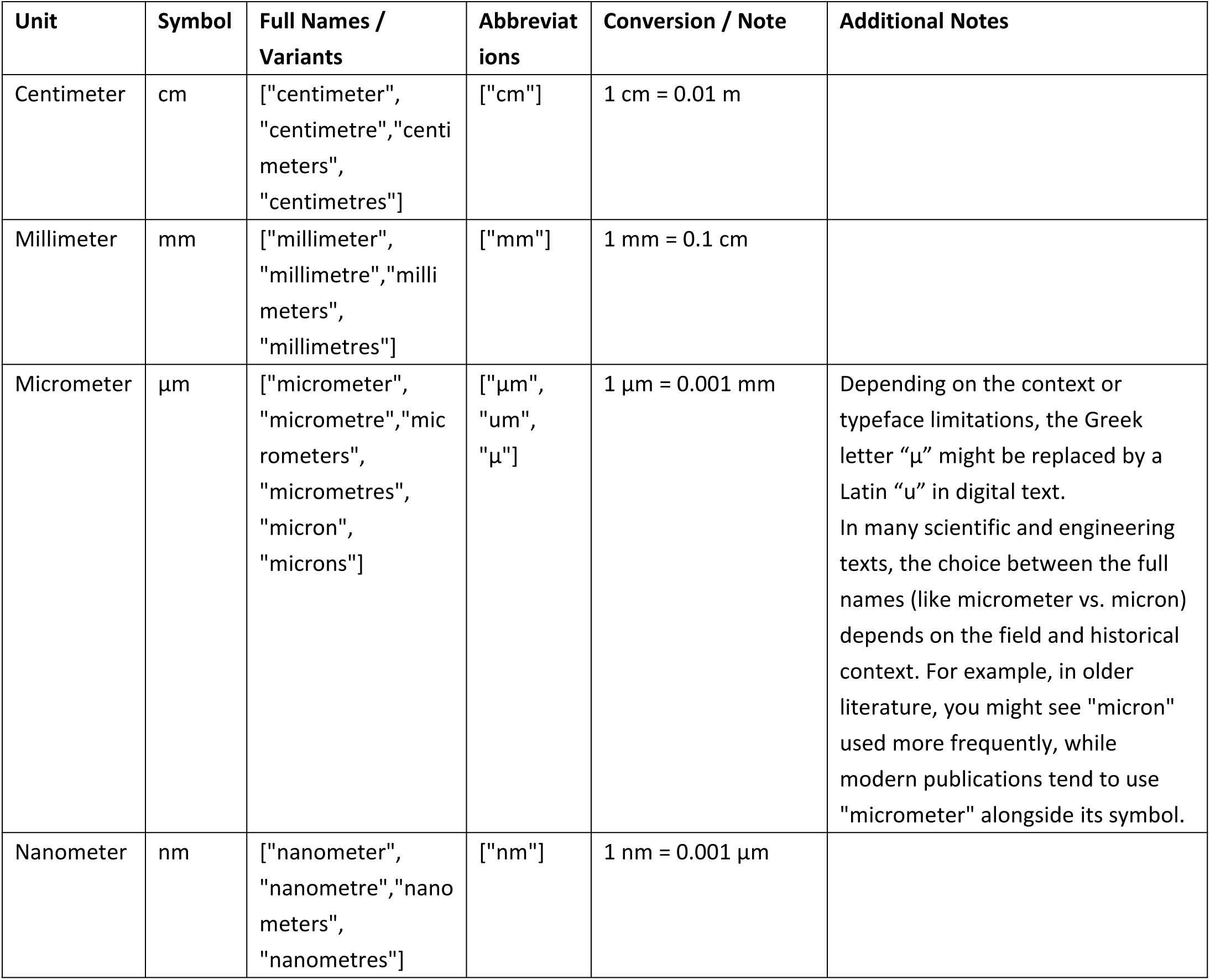
Lengths of scale bar.

**Supplementary Table 8.**
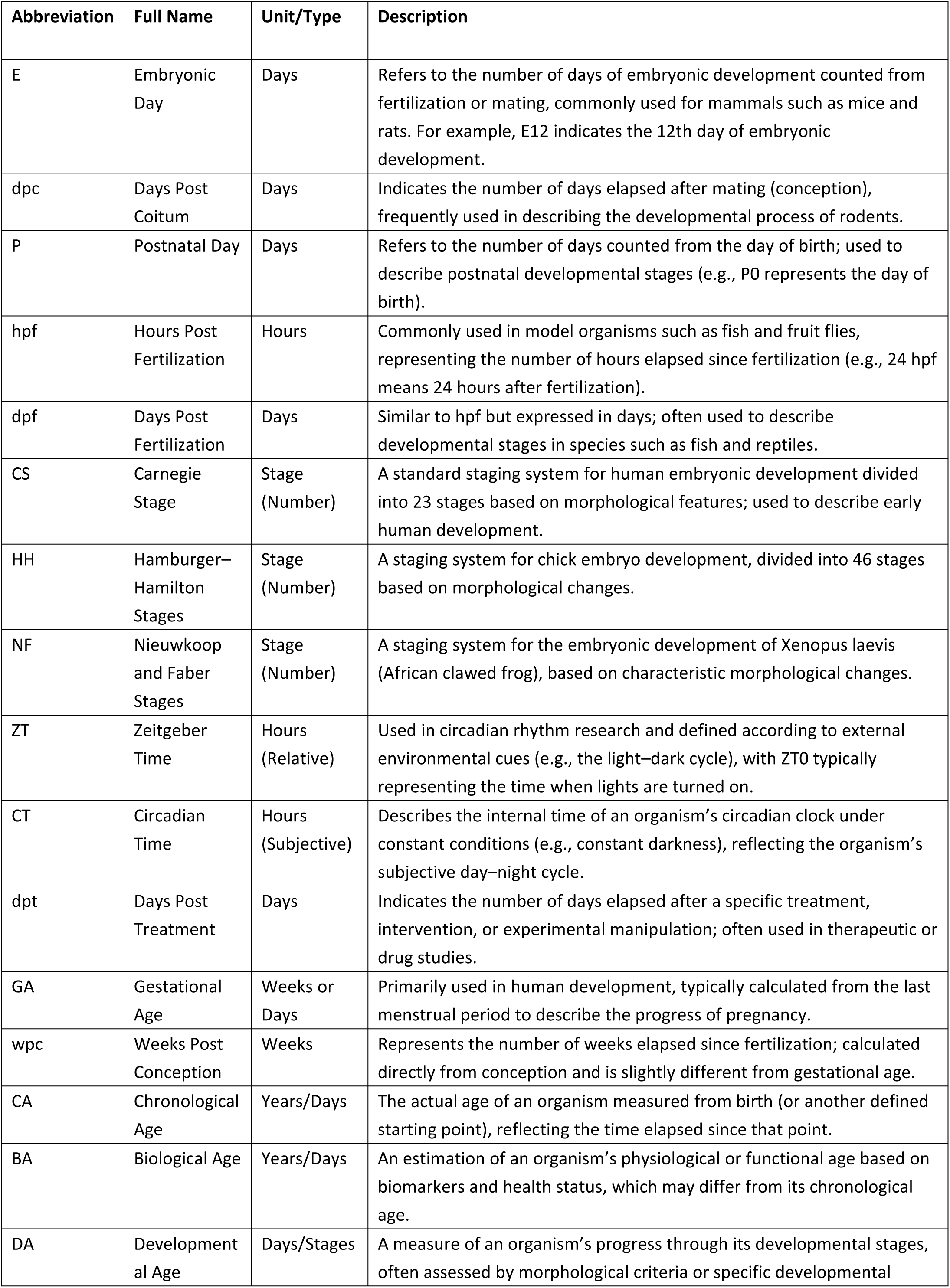
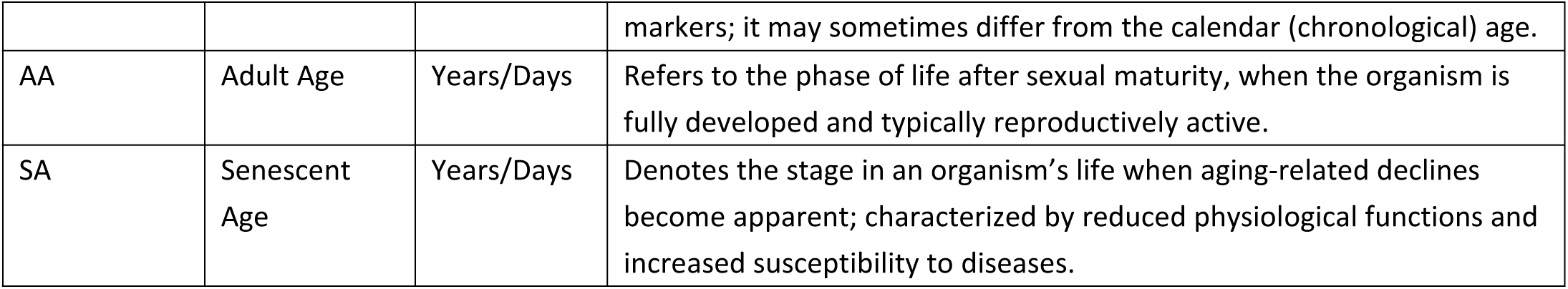
Description of age units.

**Supplementary Table 9.**
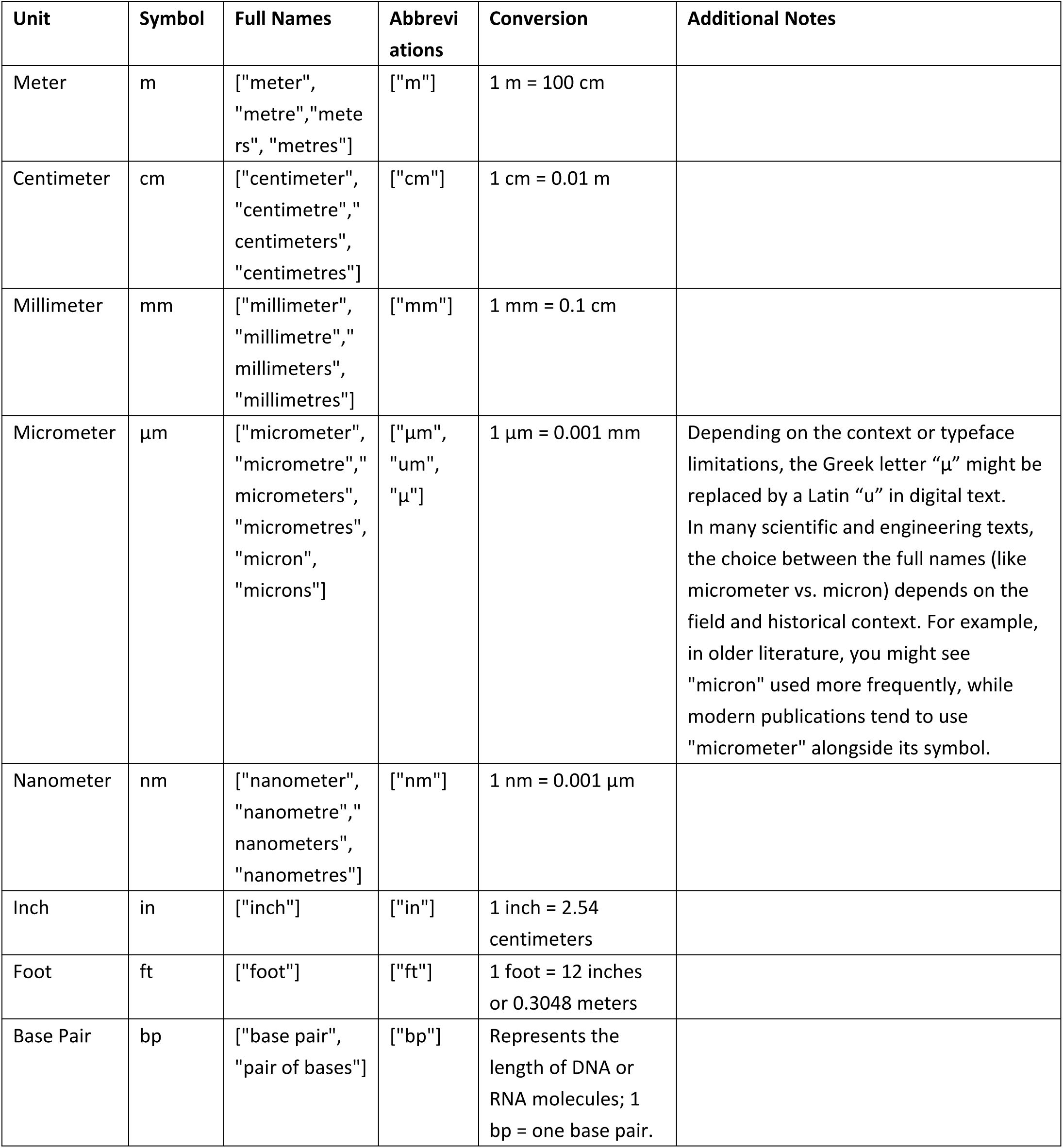
Description of height units.

**Supplementary Table 10.**
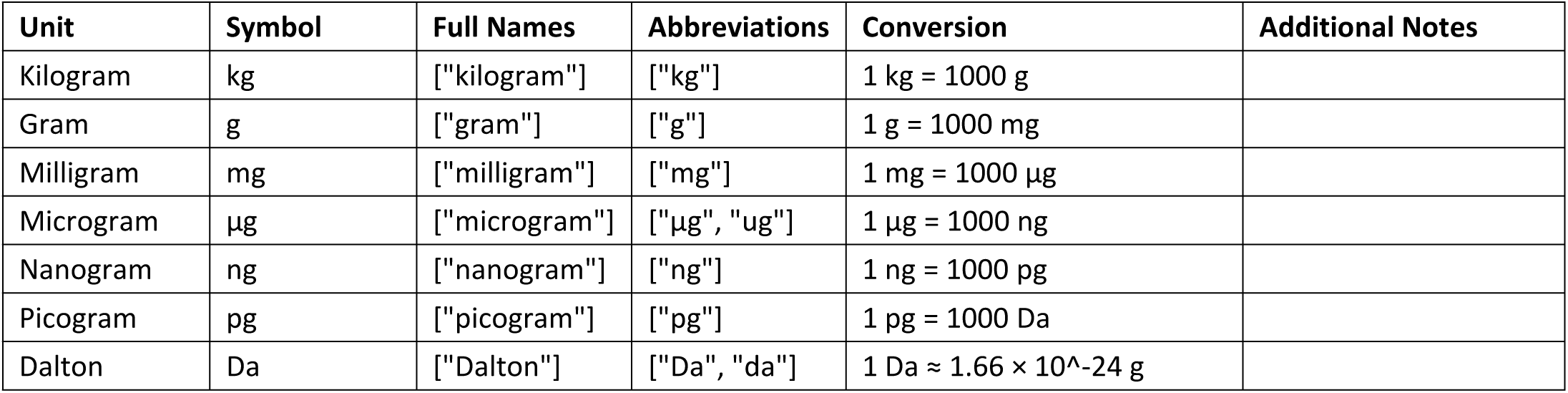
Description of weight units.

**Supplementary Table 11.**
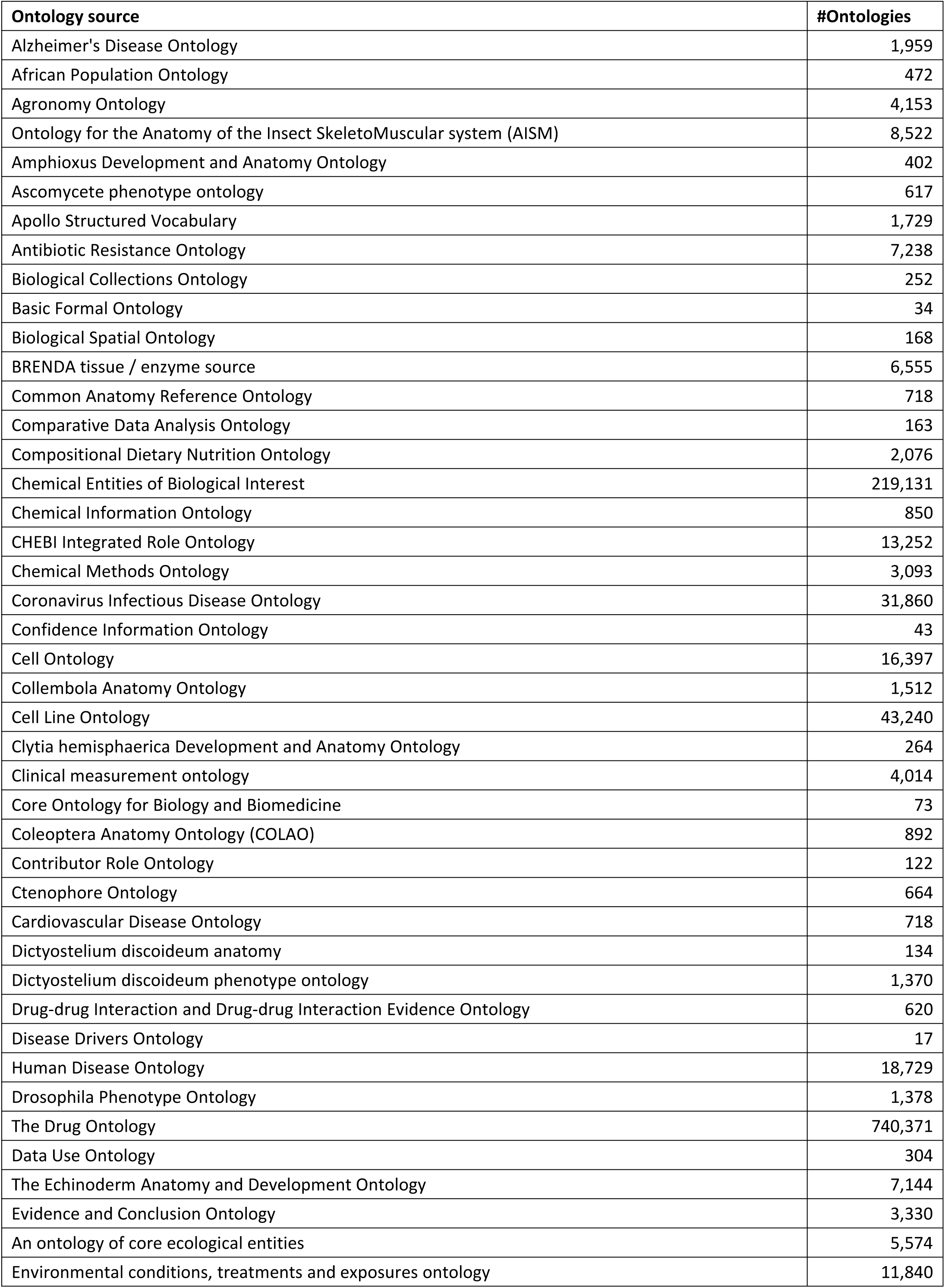
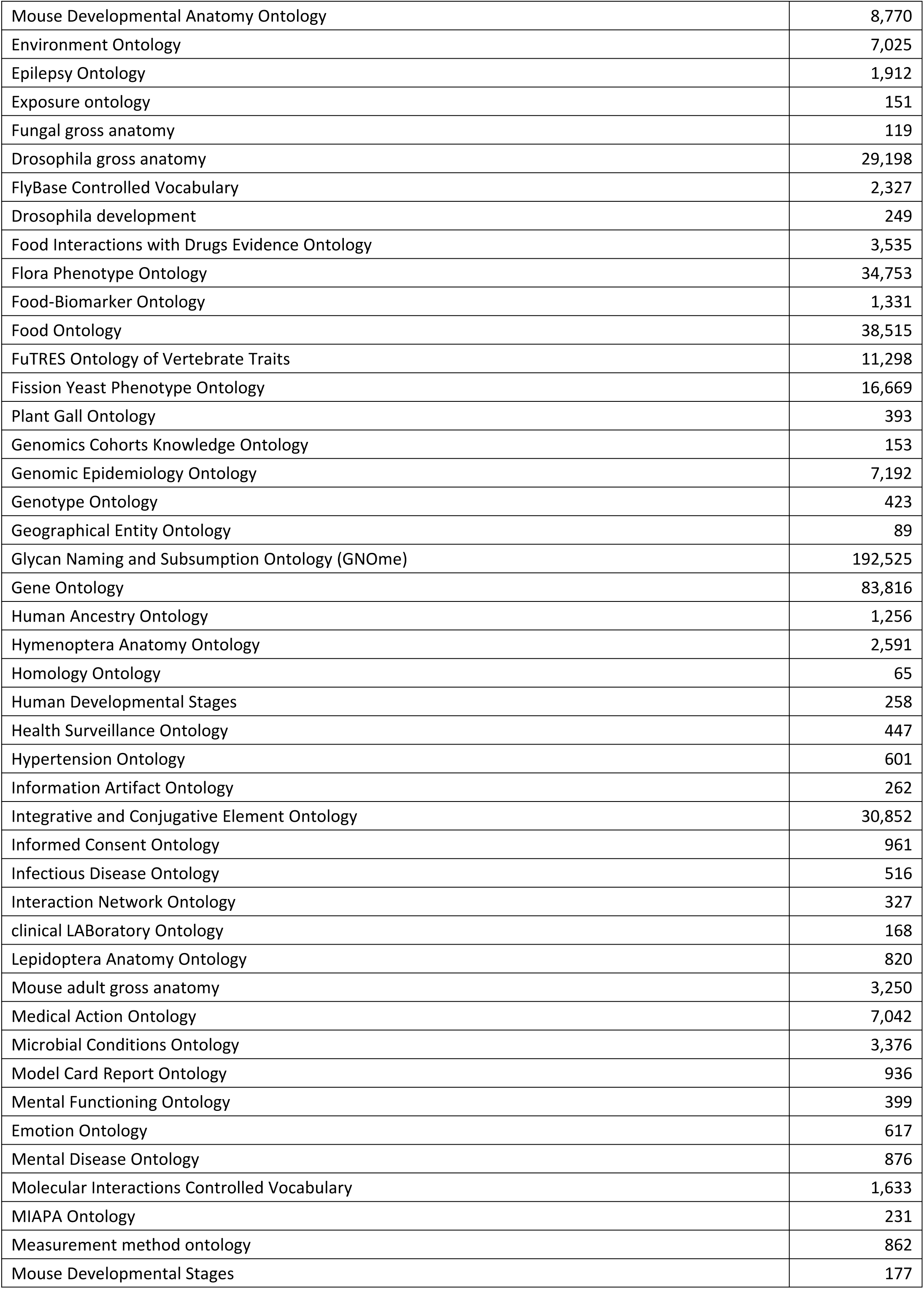
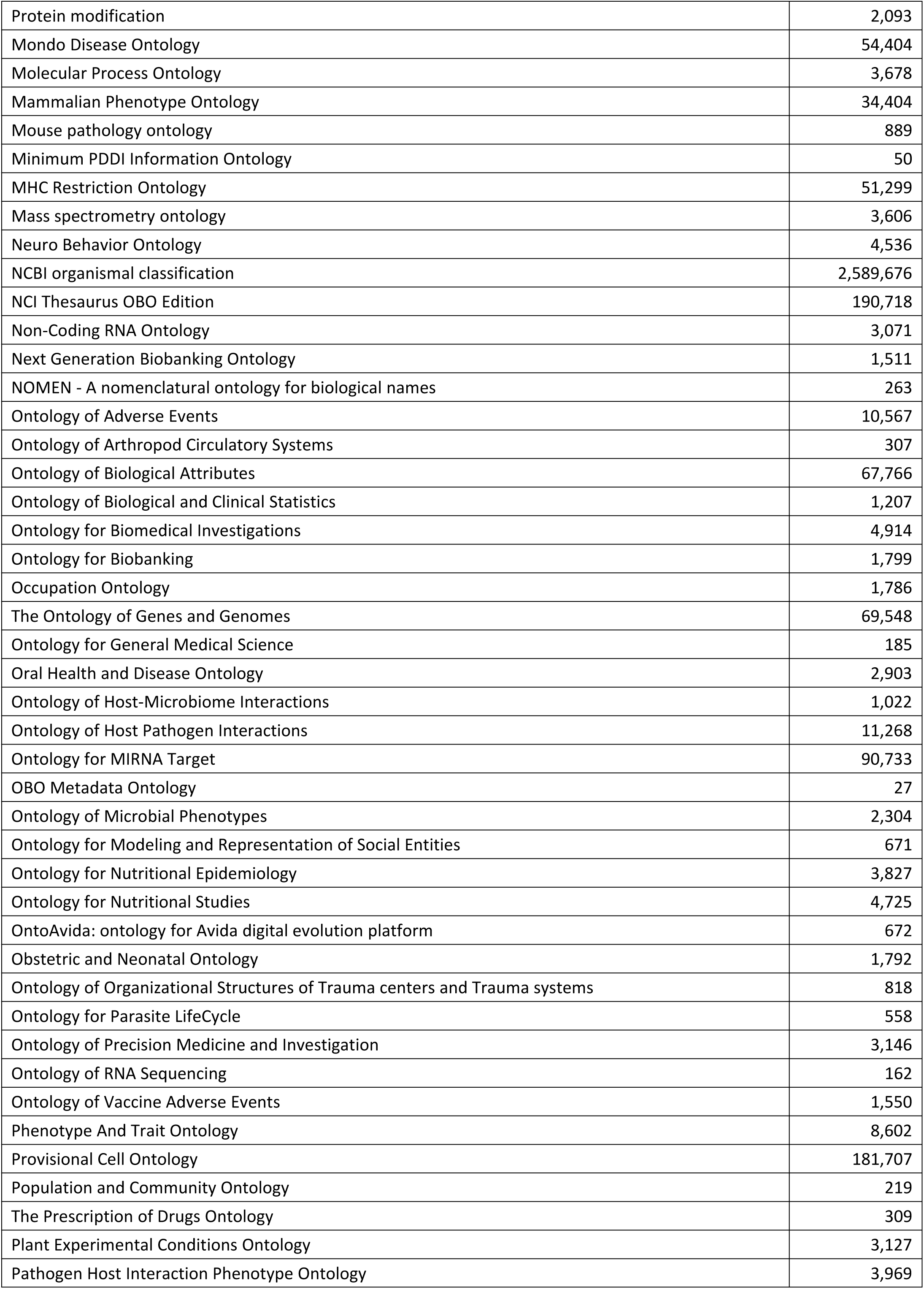
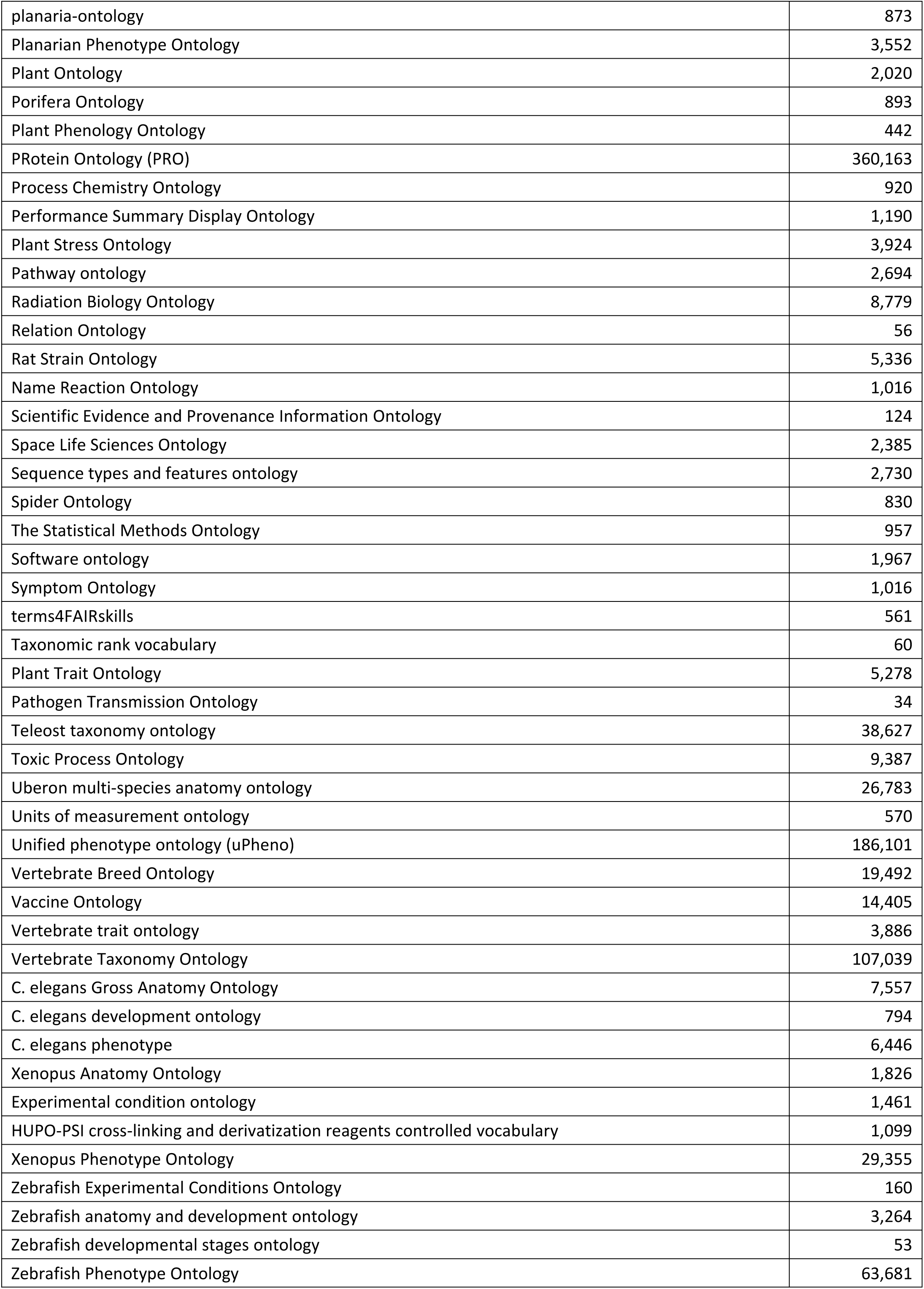
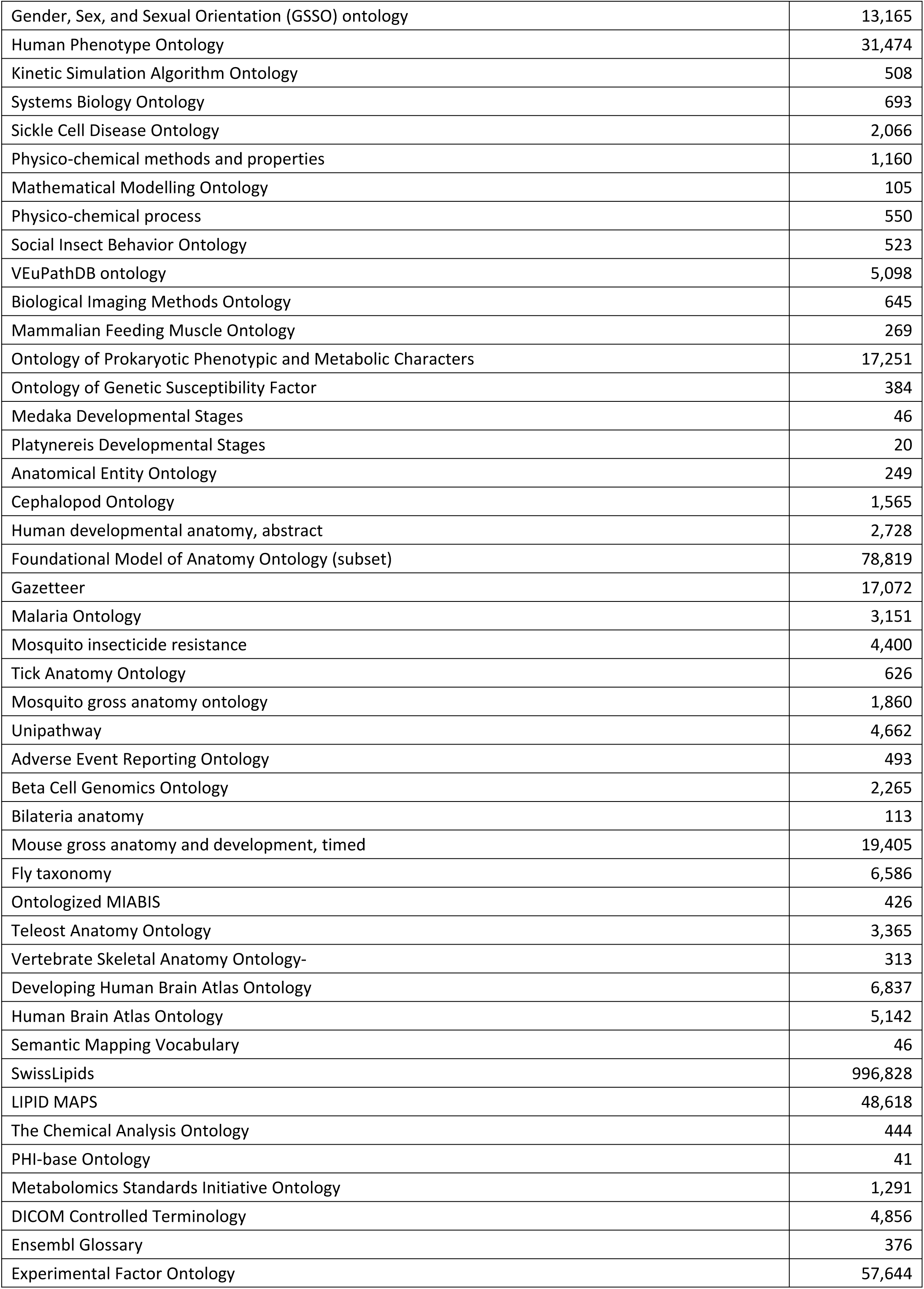
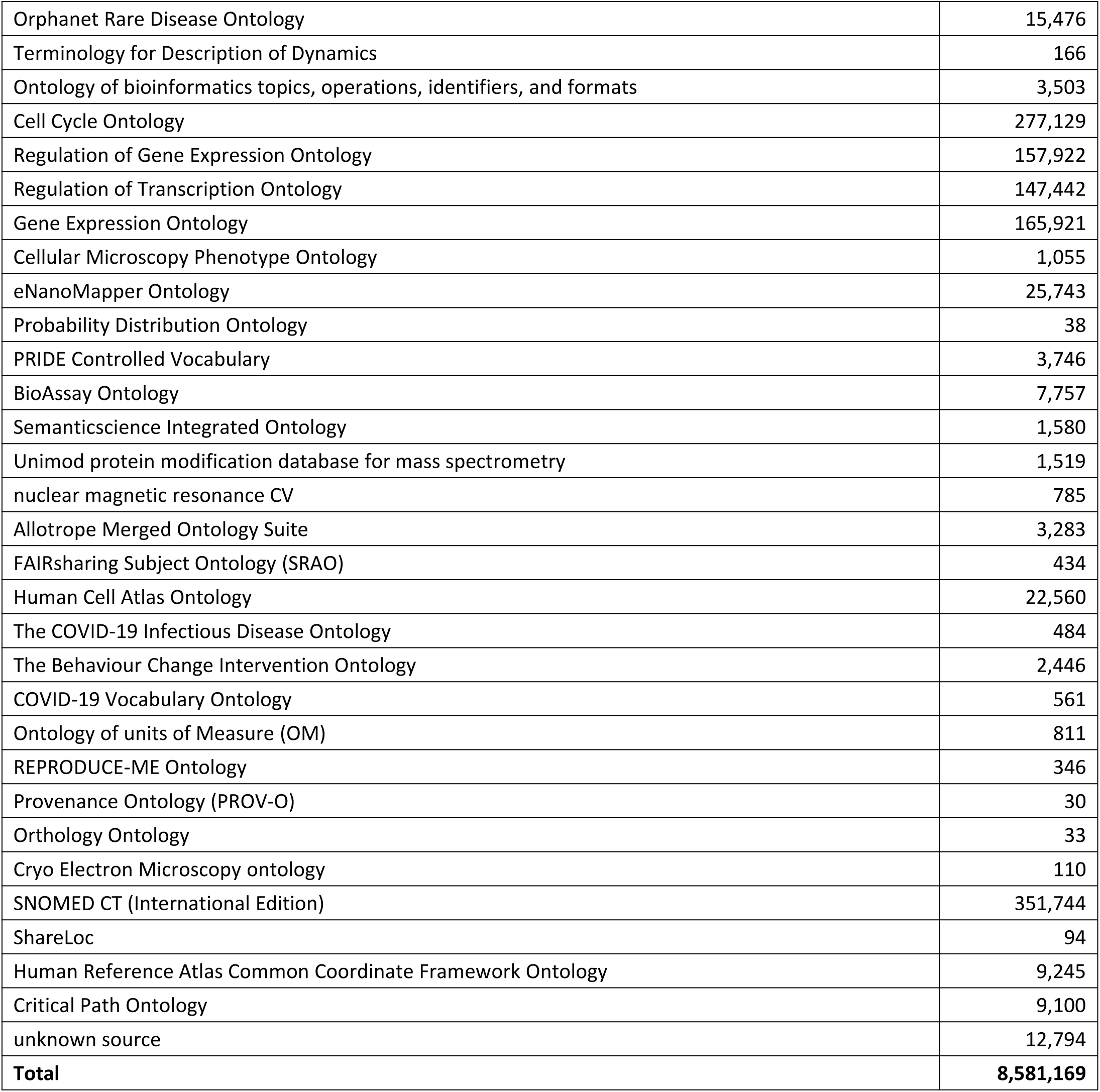
Overview of OLS ontology sources.

**Supplementary Table 12.**
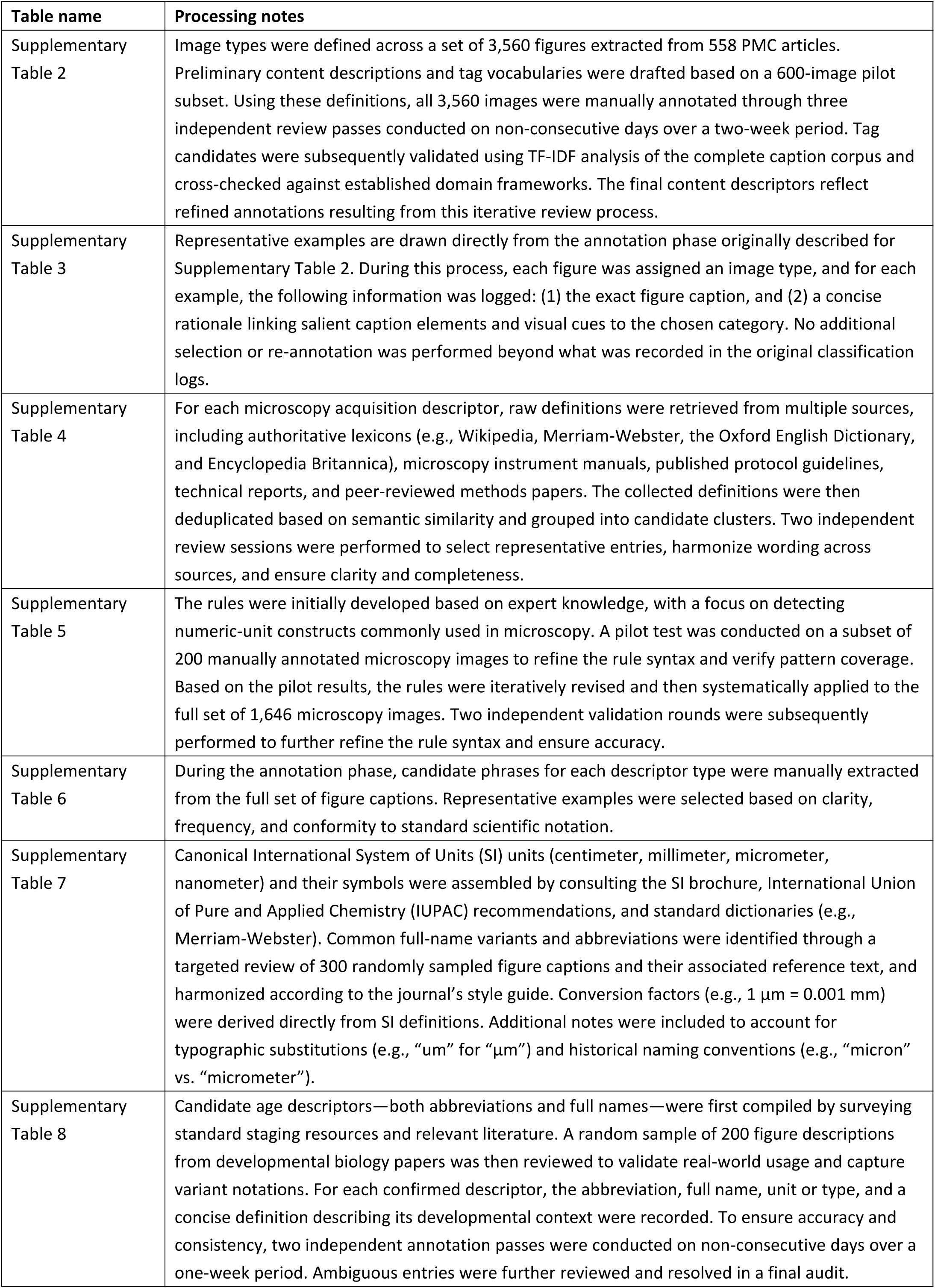
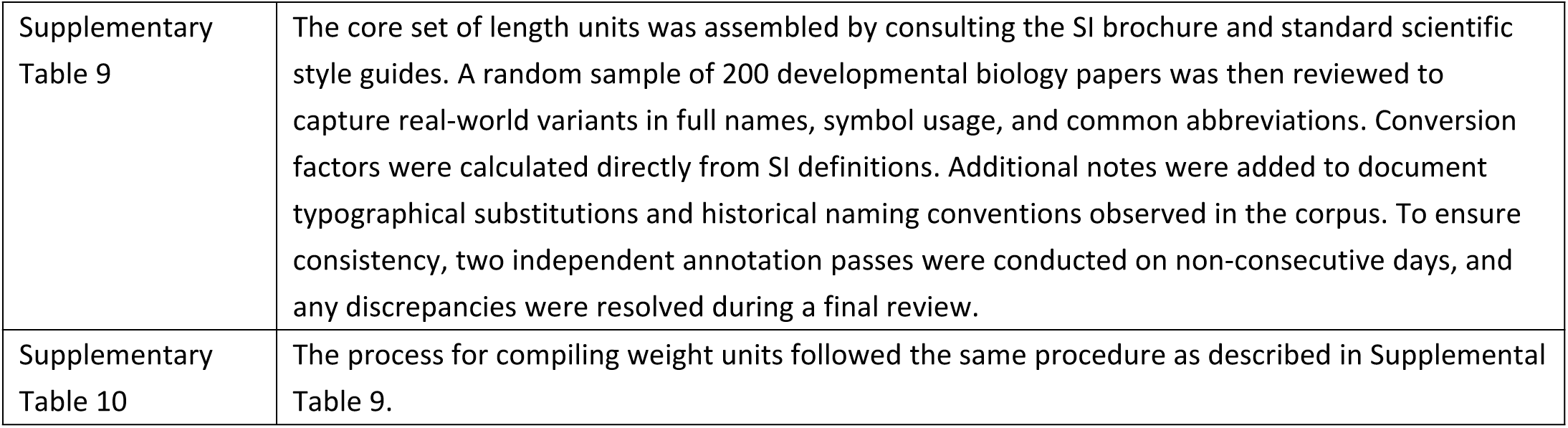
Summary of task-specific knowledge base construction.

**Supplementary Table 13.**
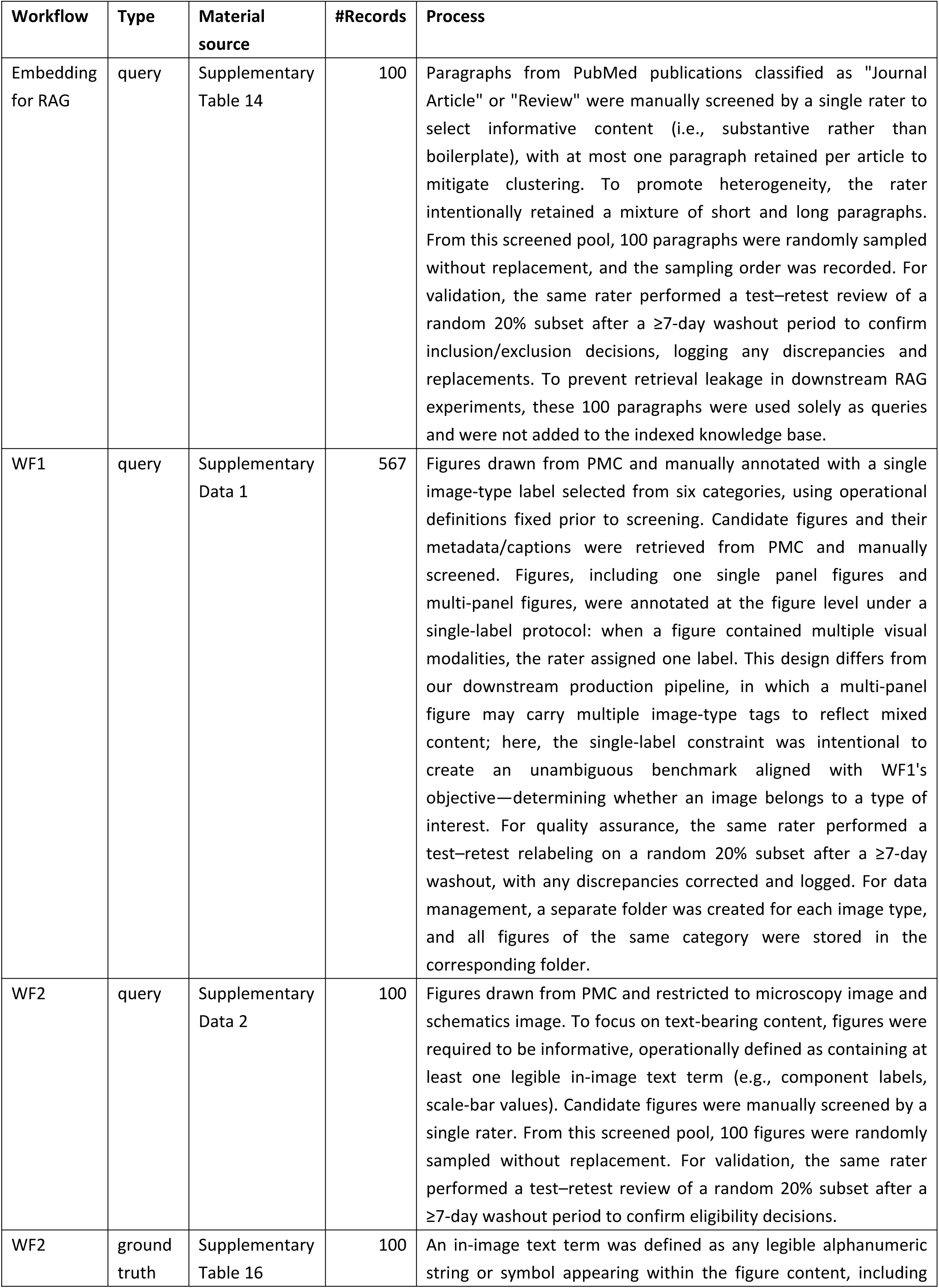
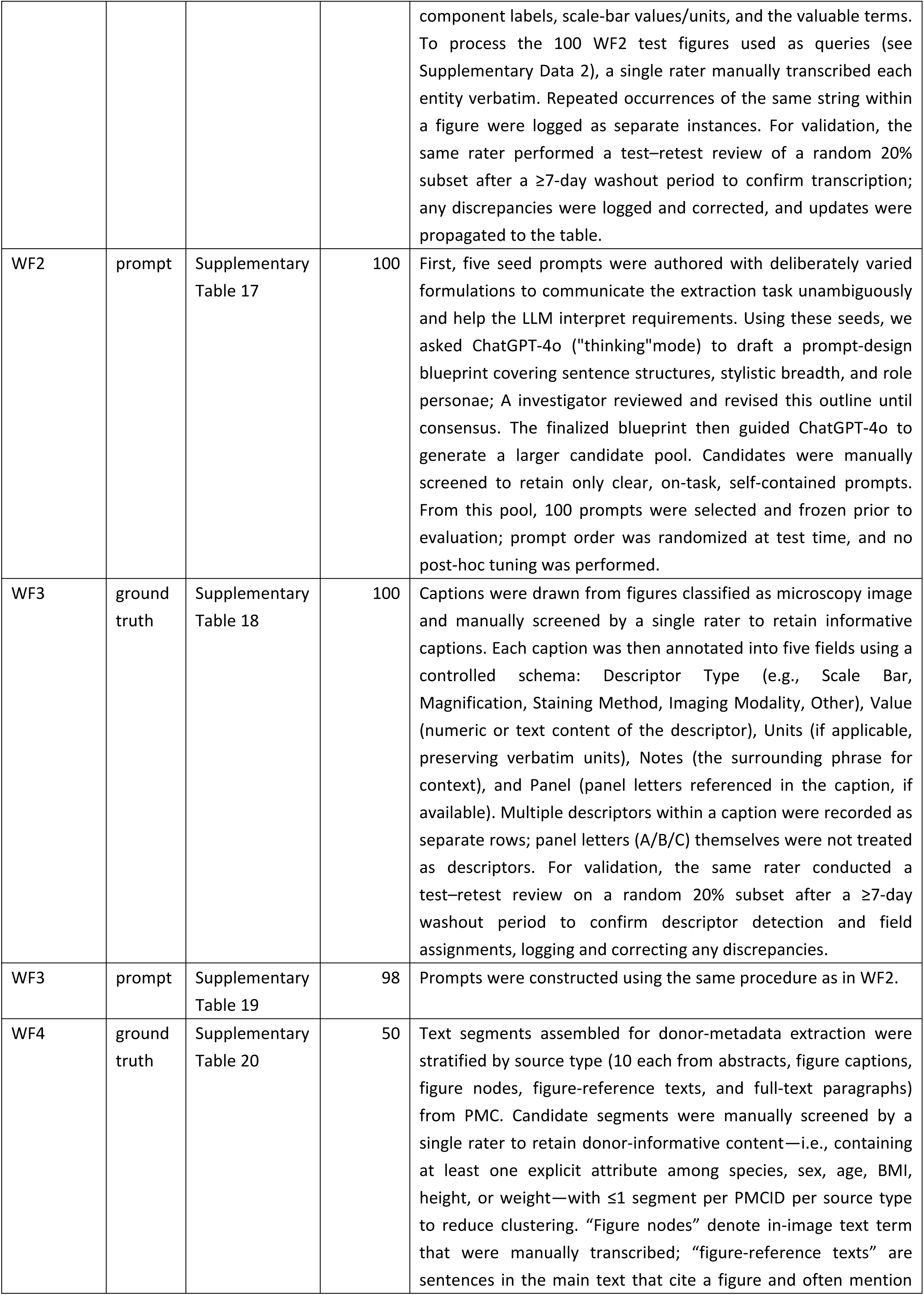
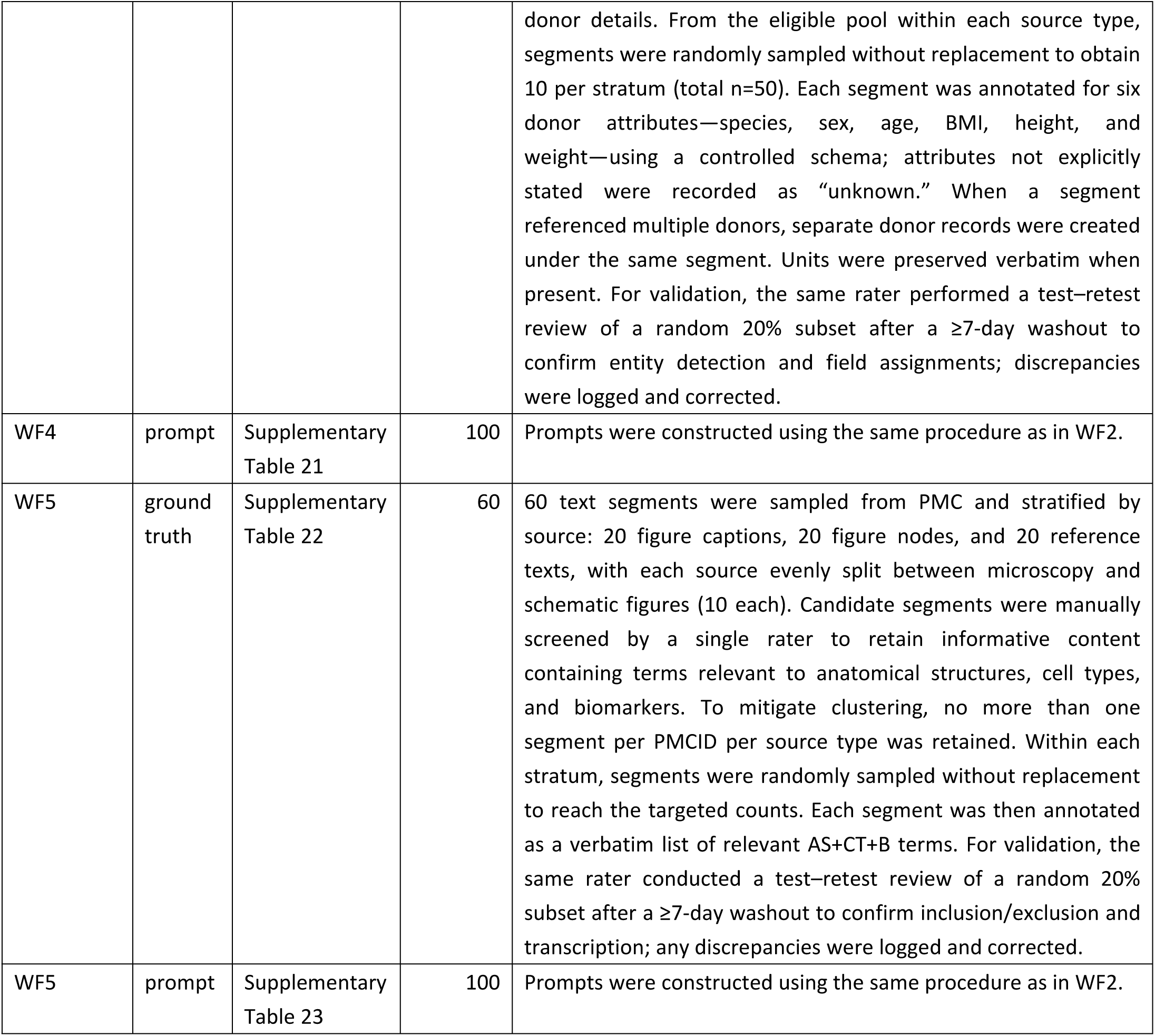
Summary of evaluation-material construction across the five workflows.

**Supplementary Table 14.**
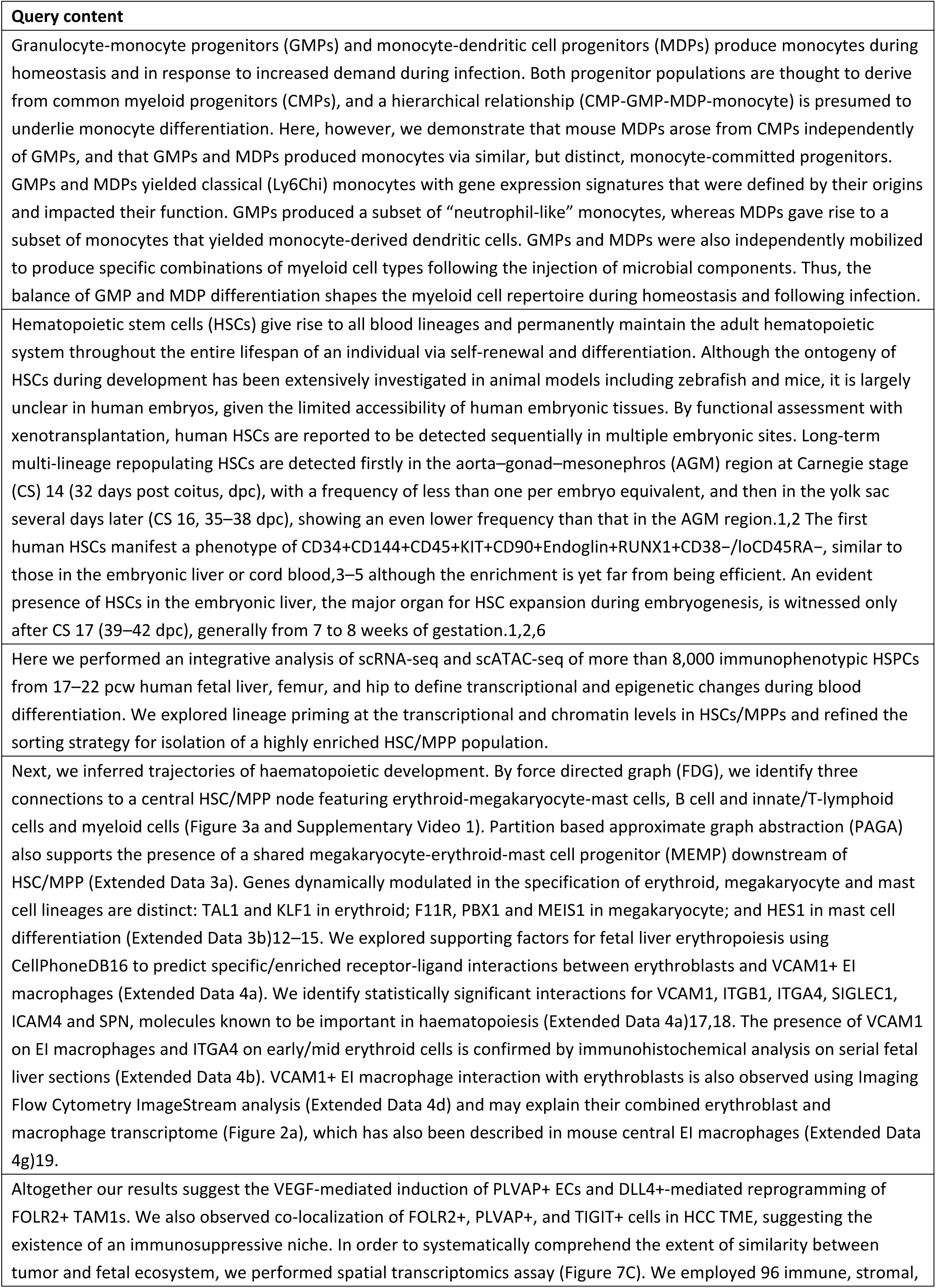
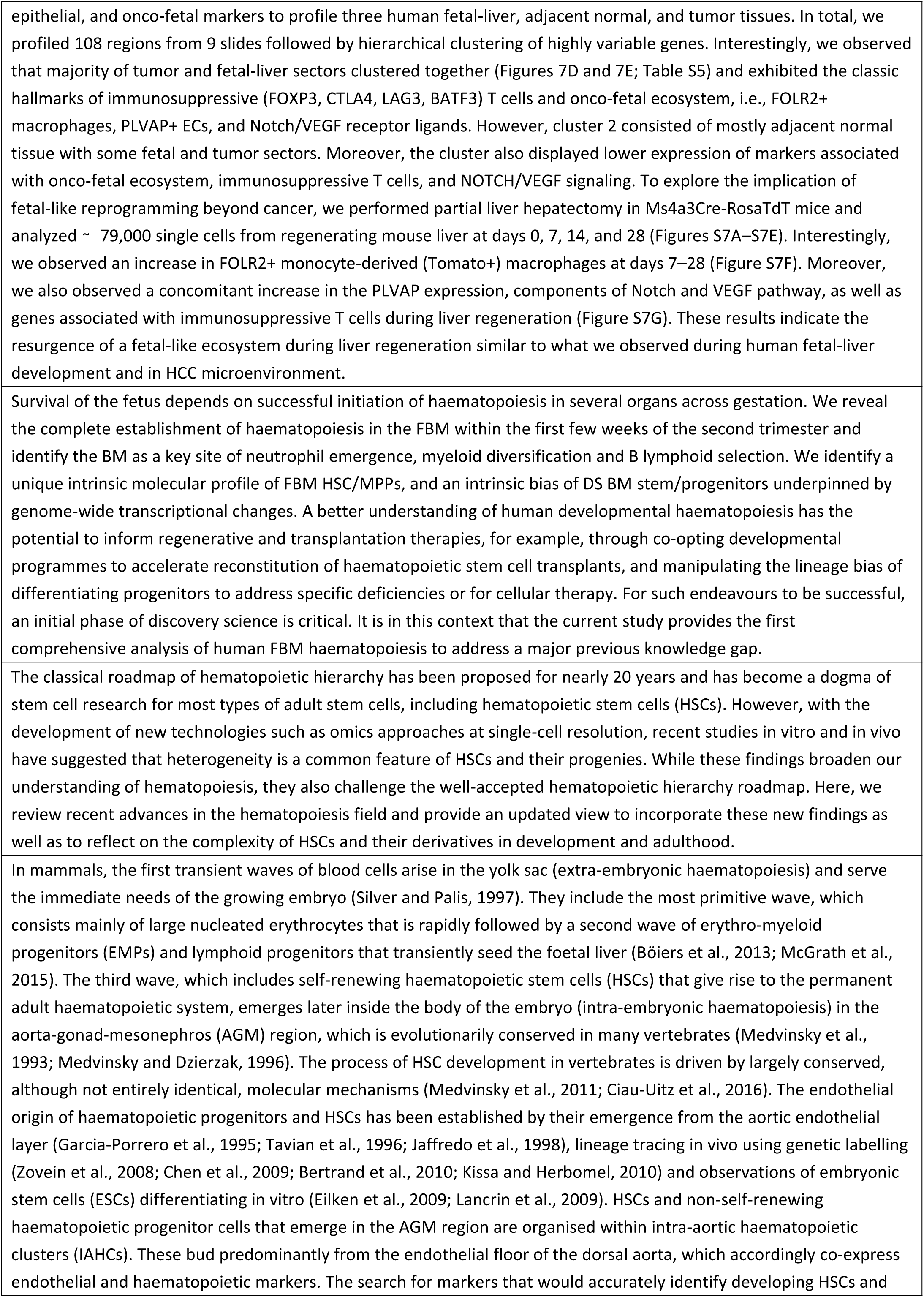
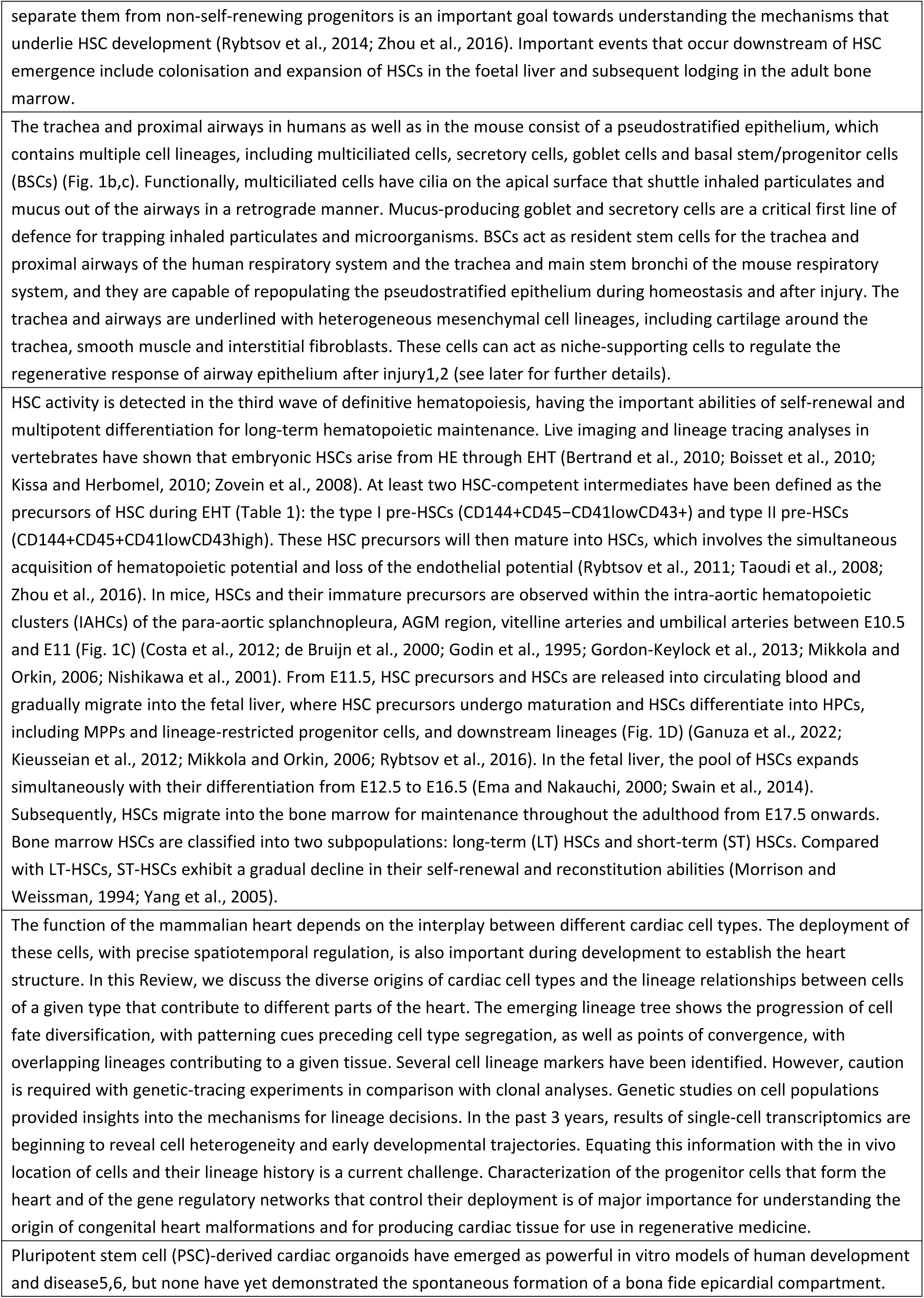
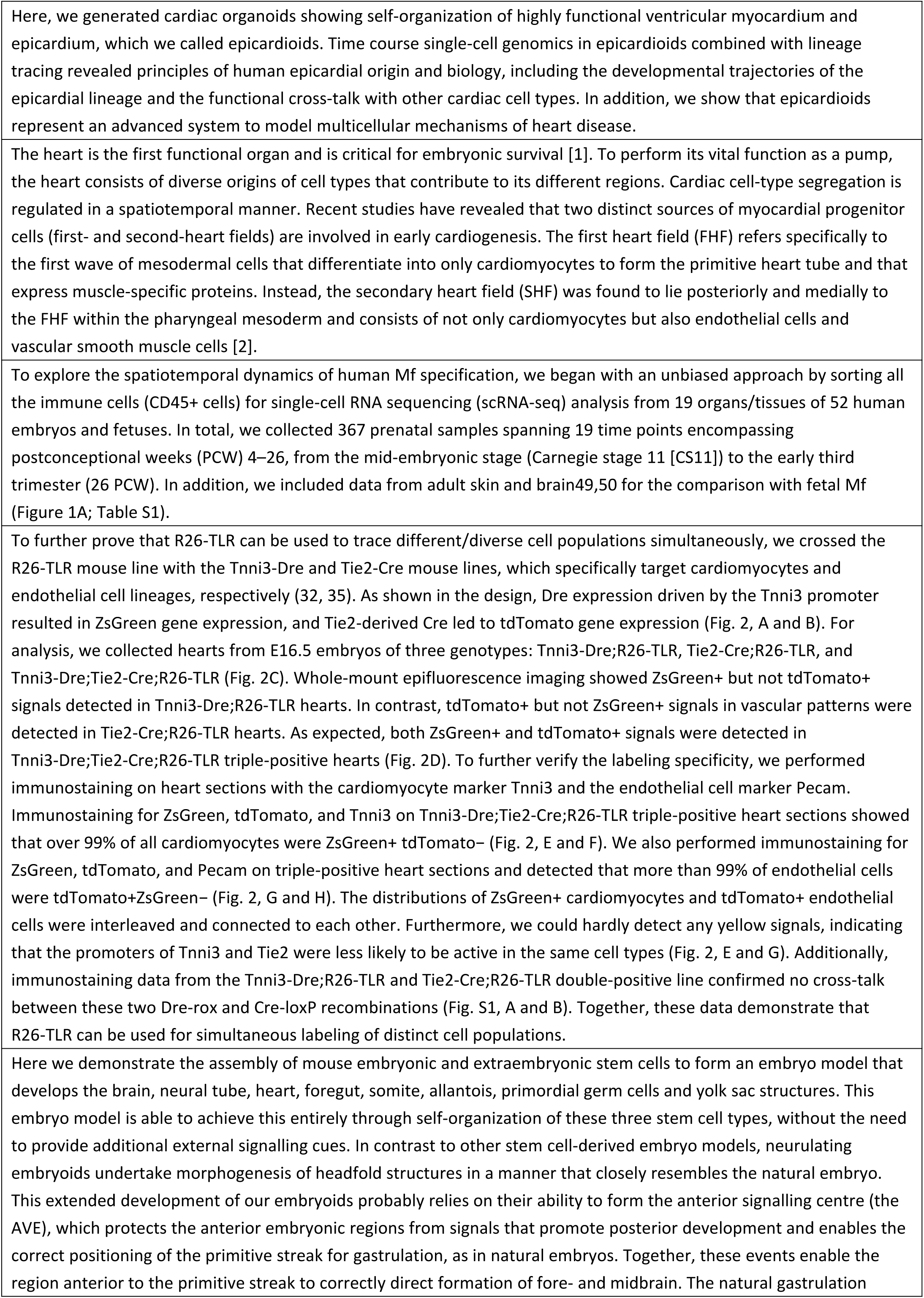
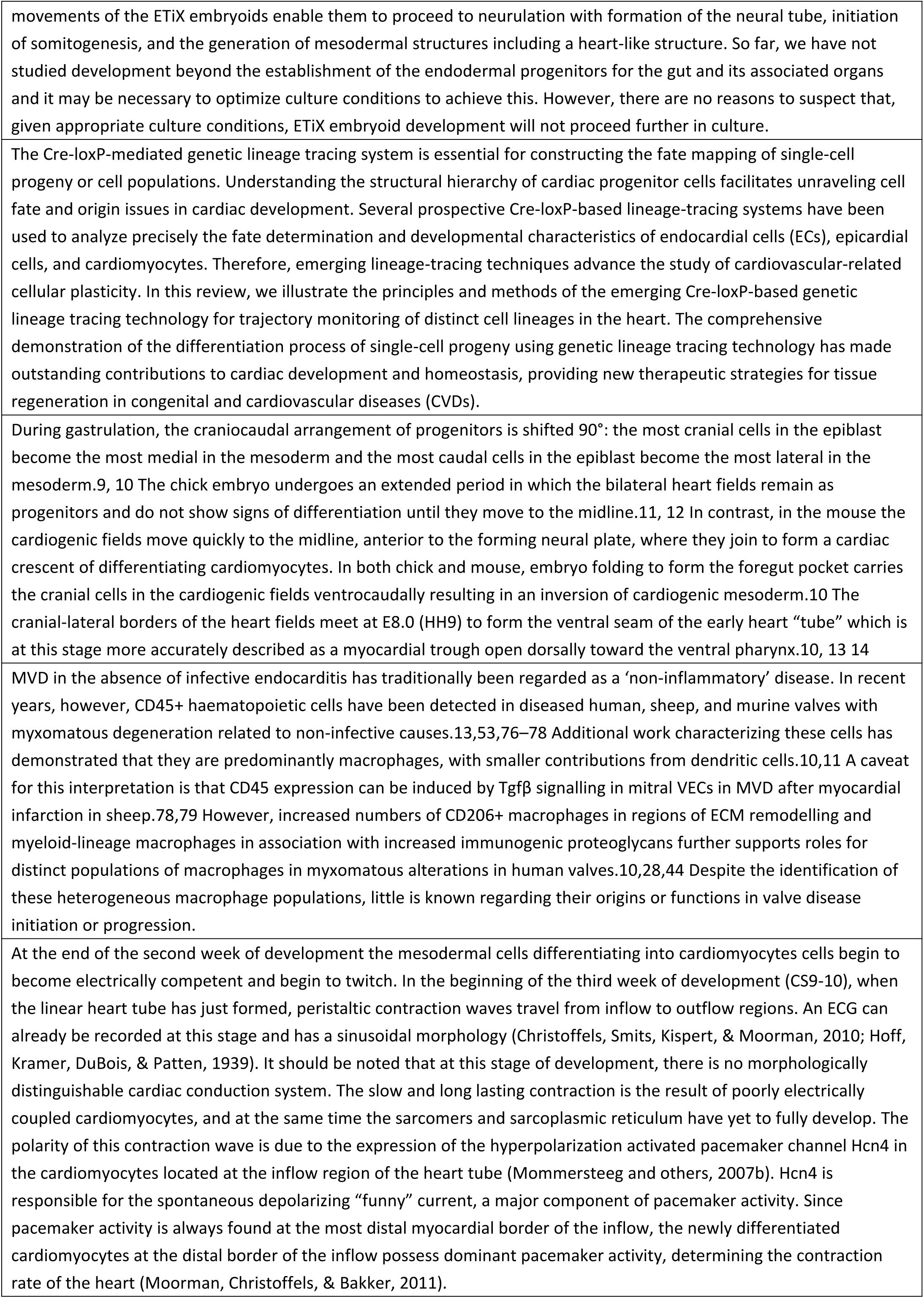
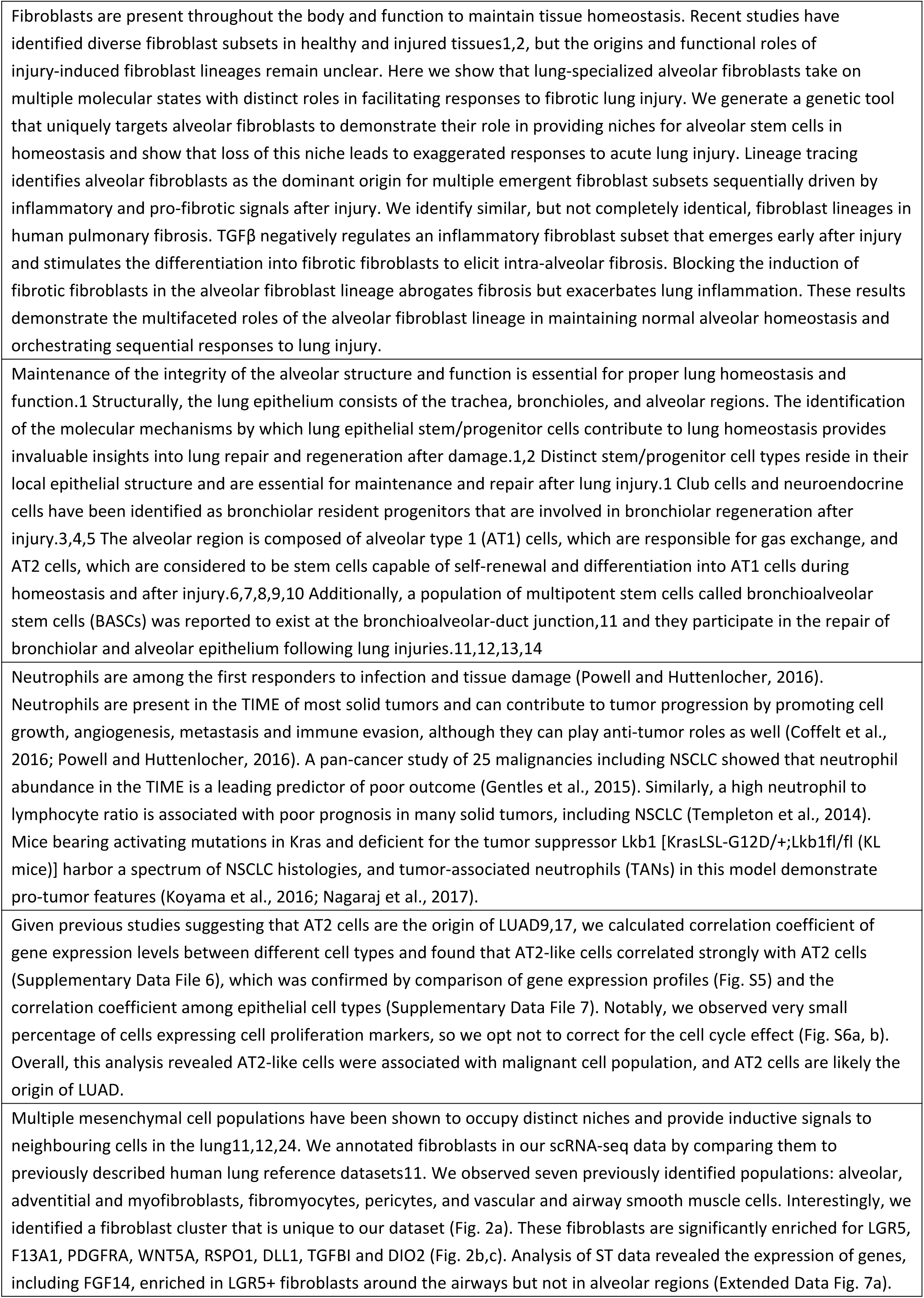
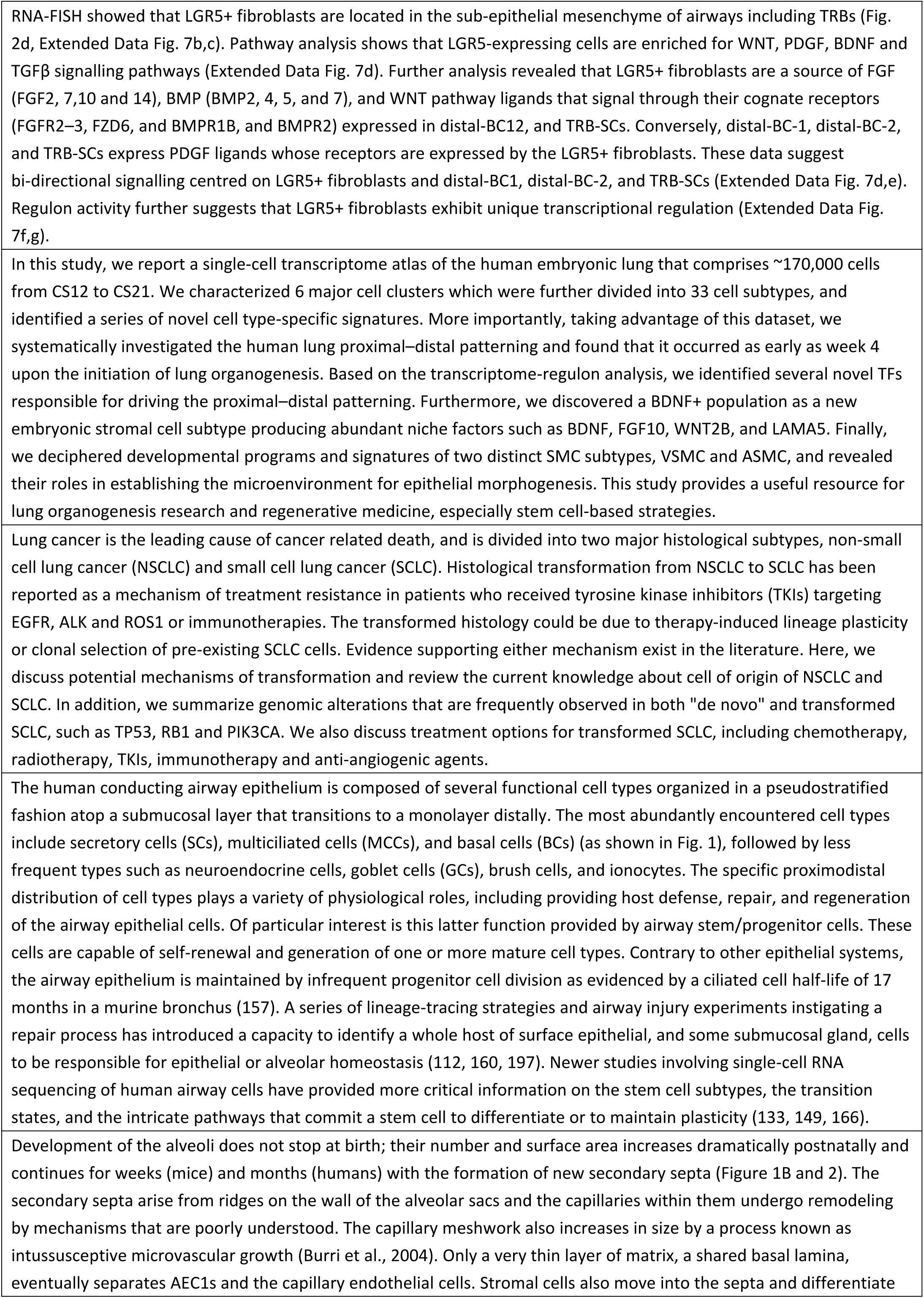
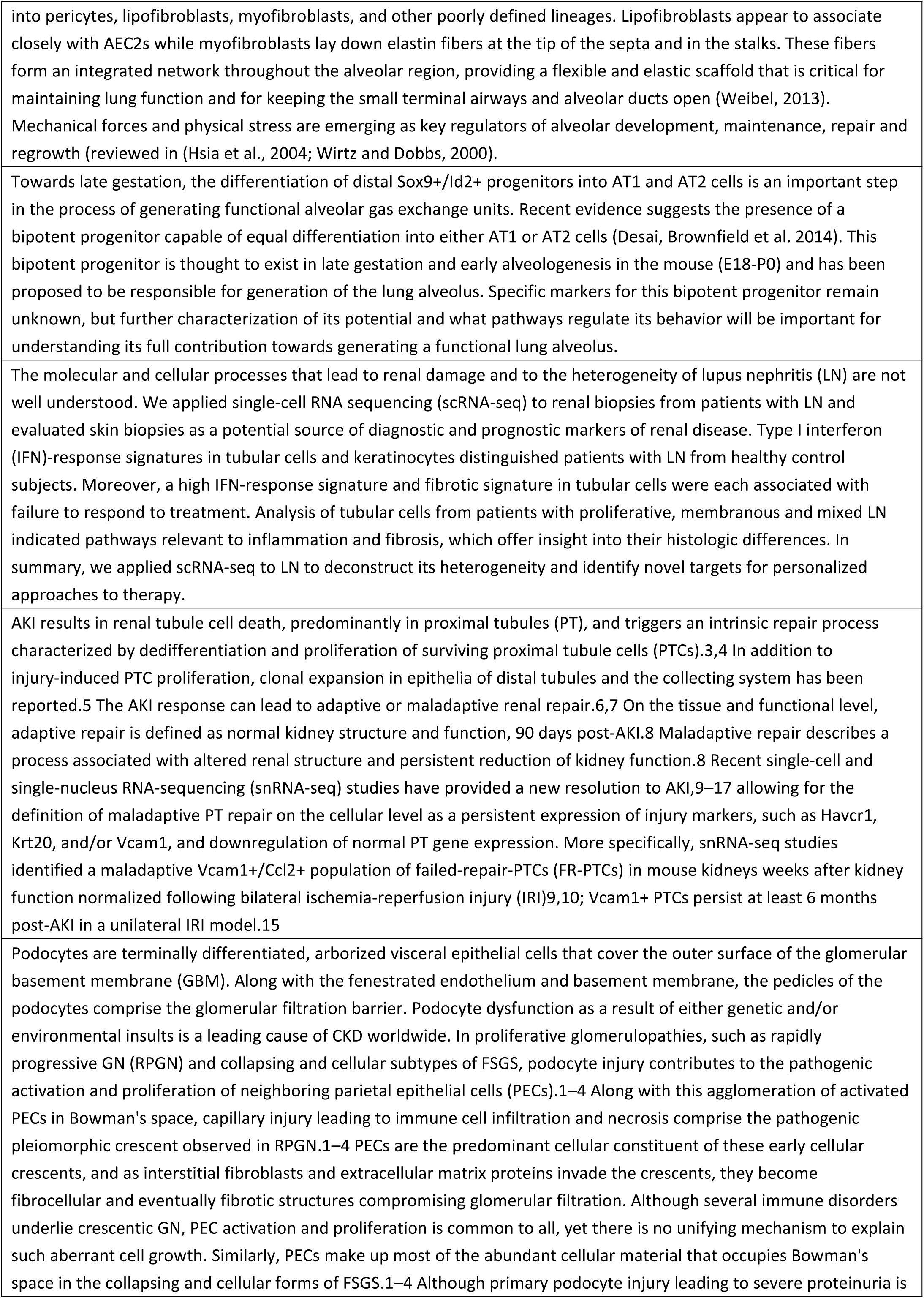
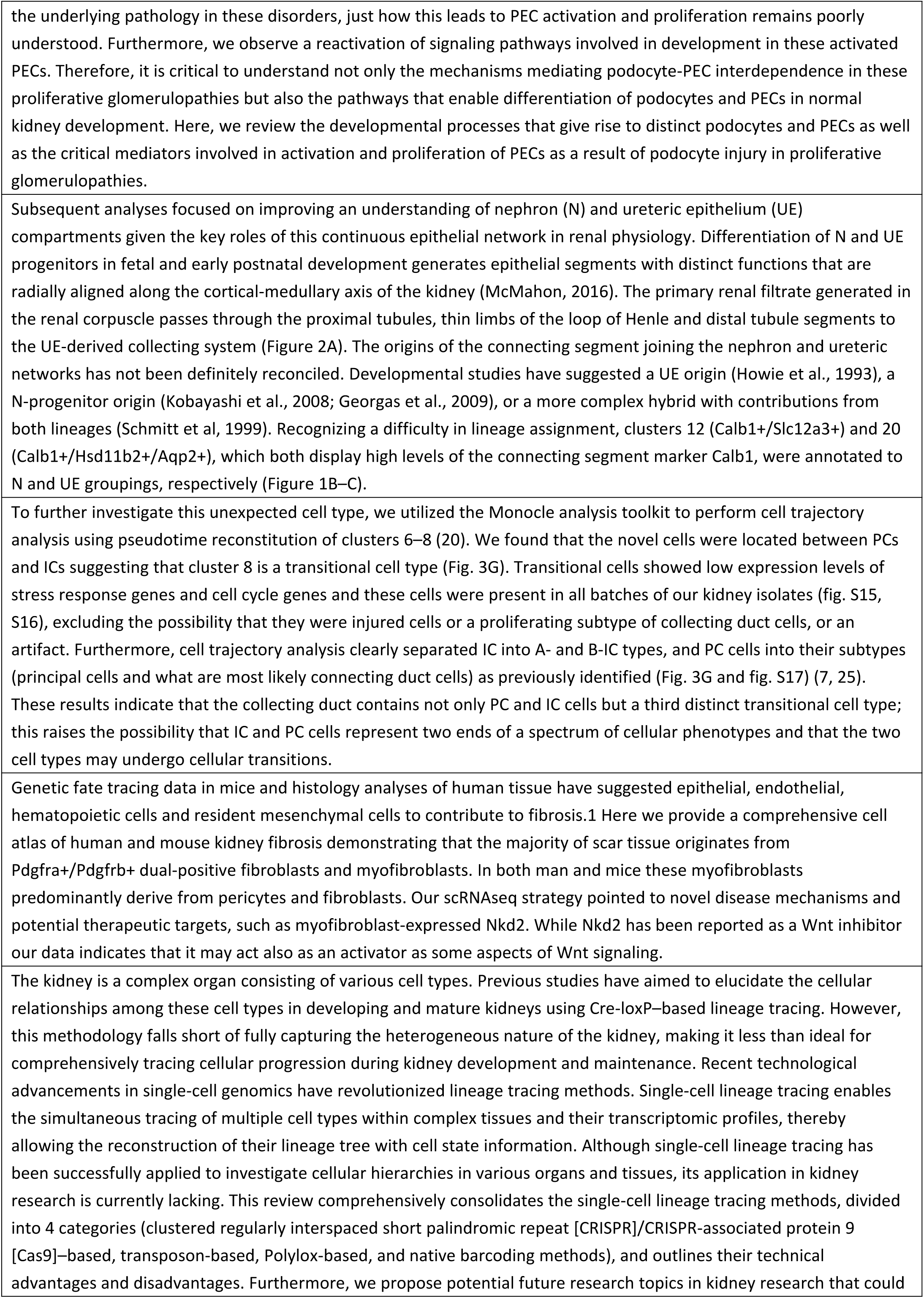
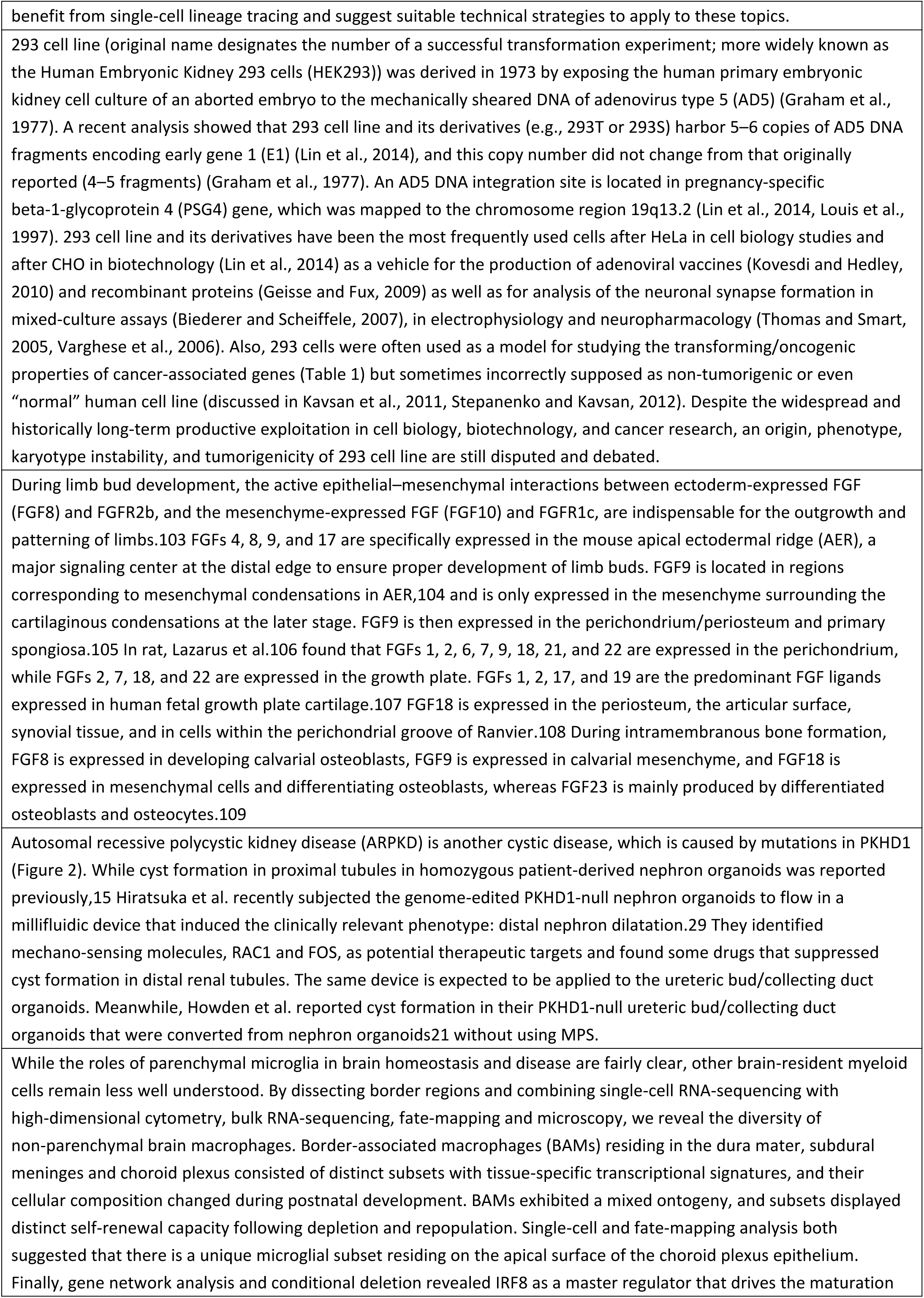
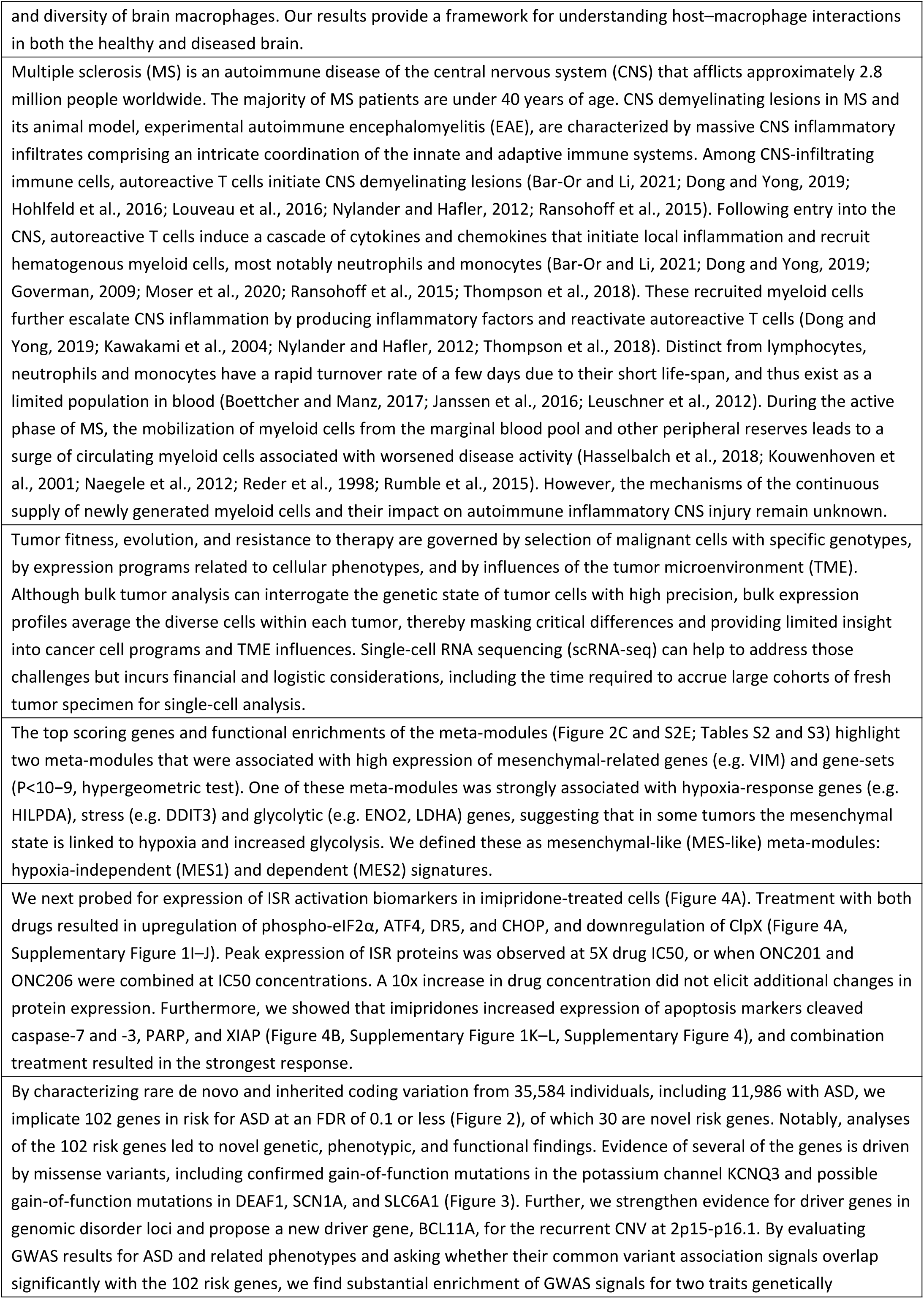
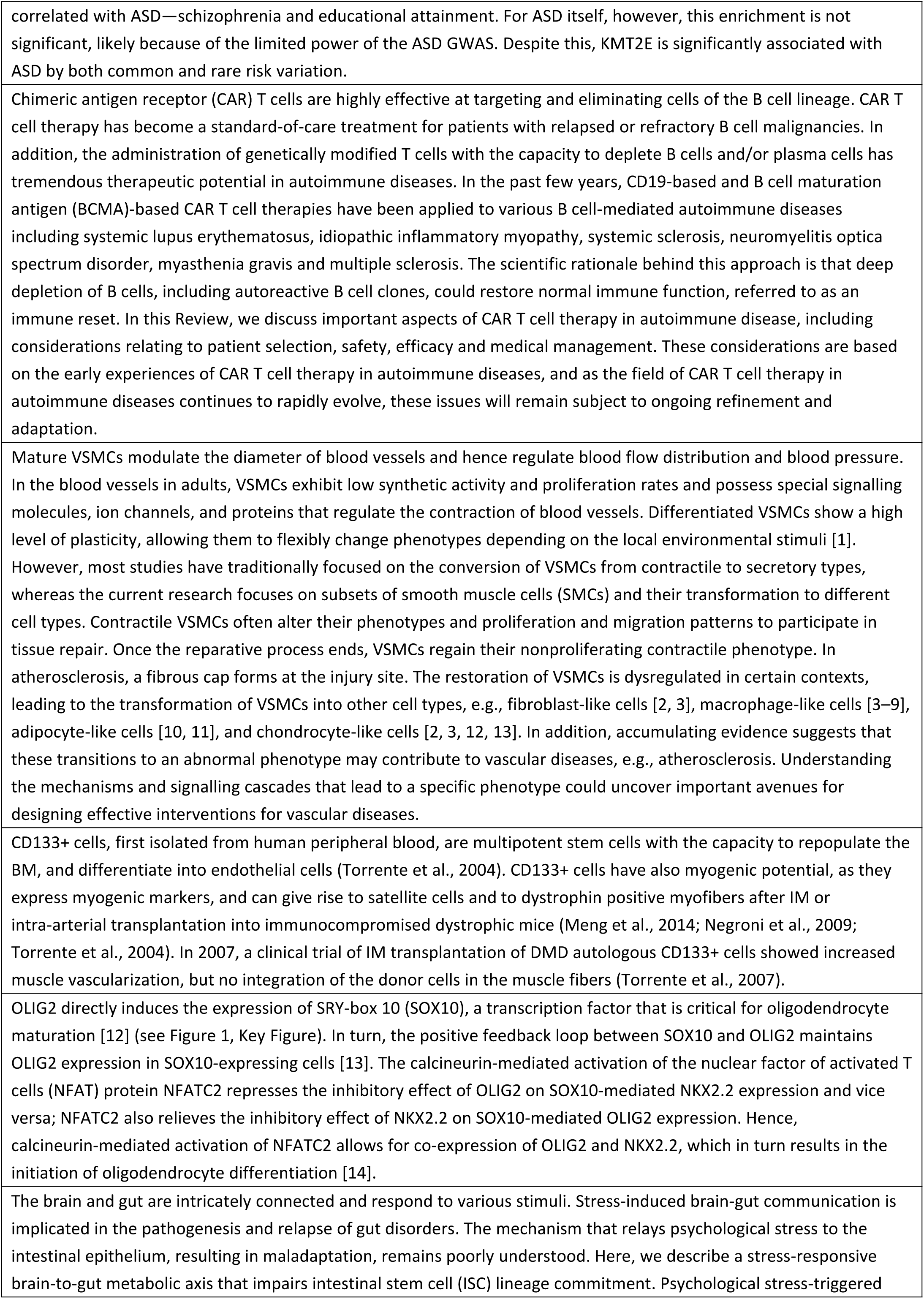
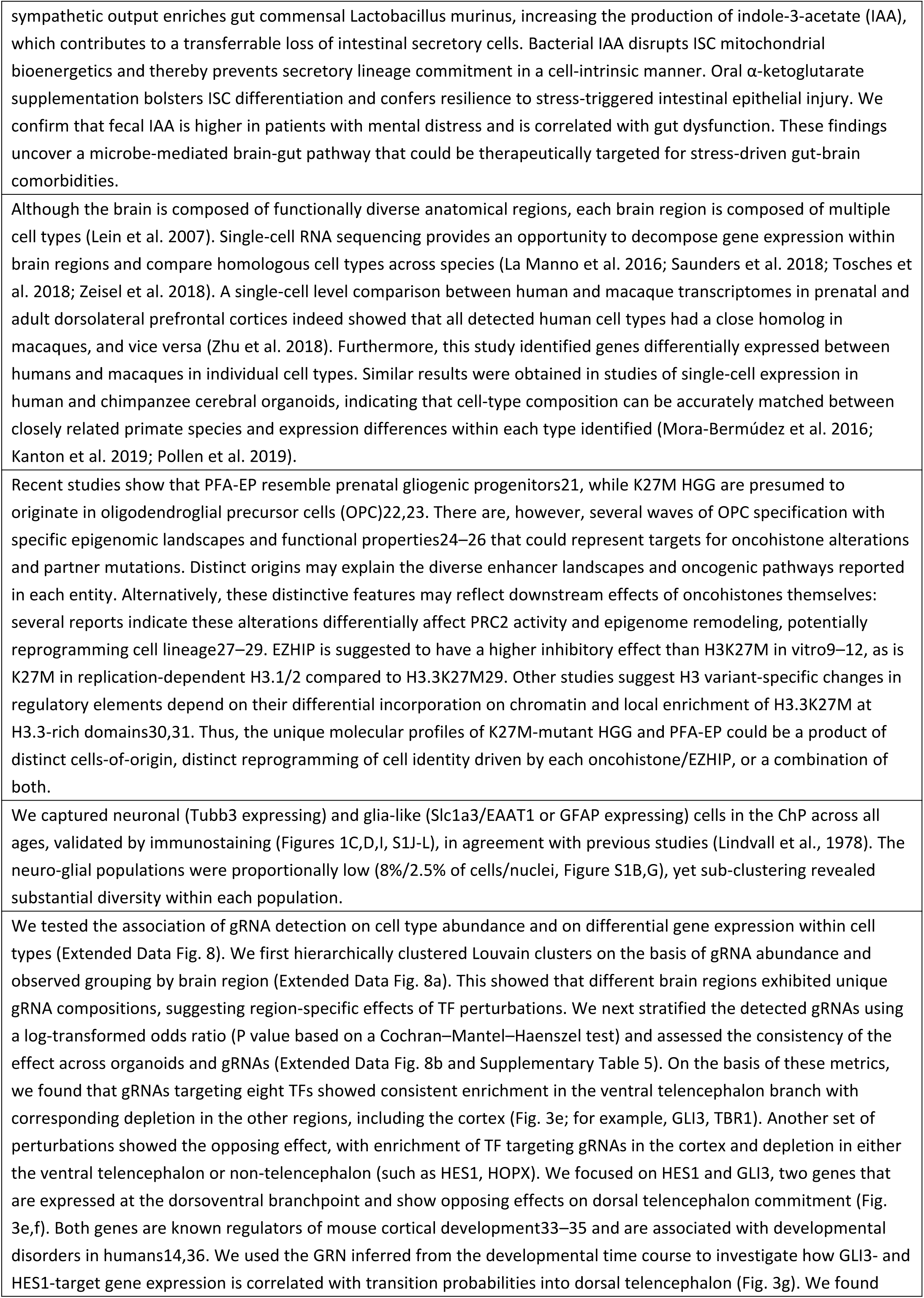
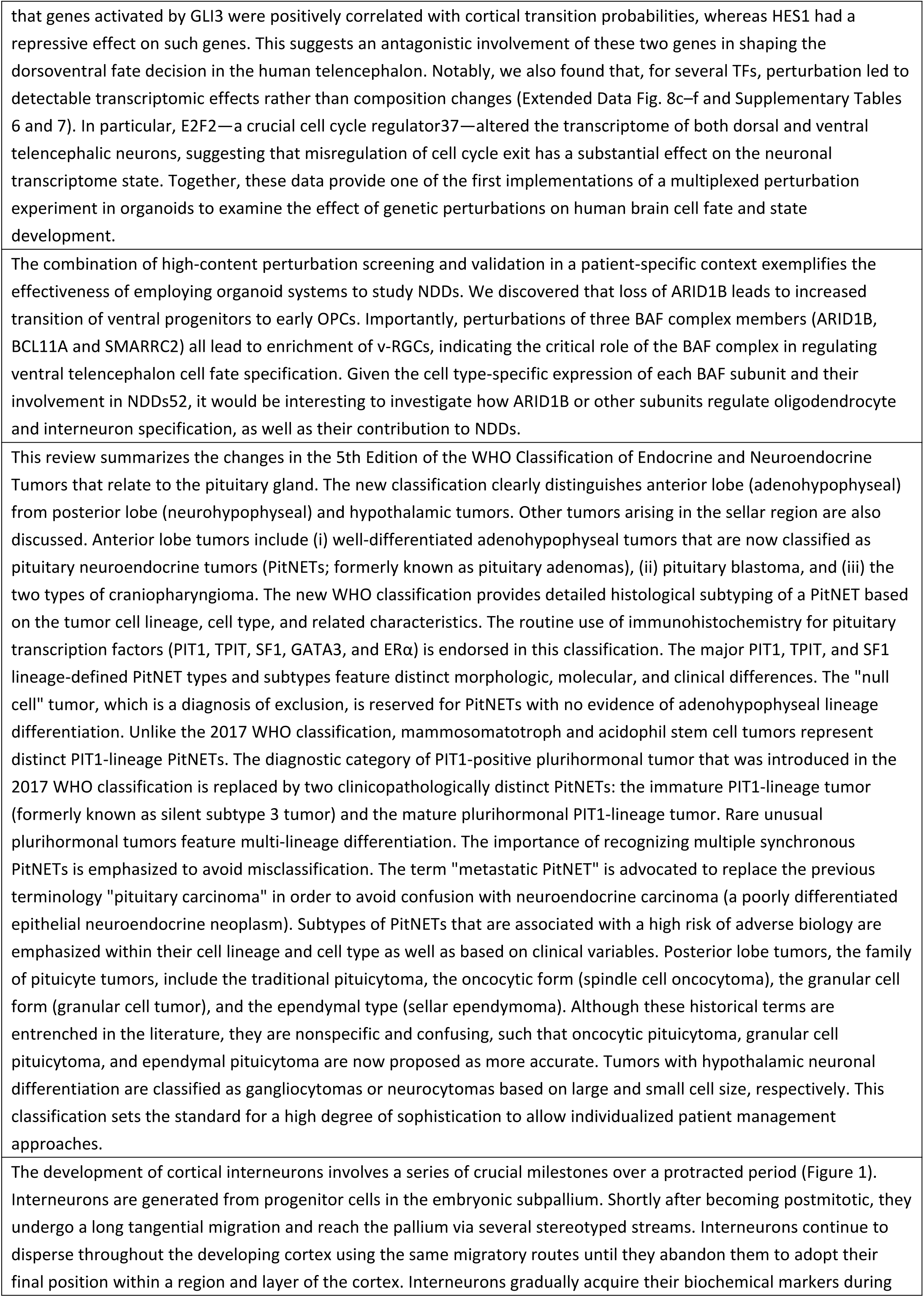
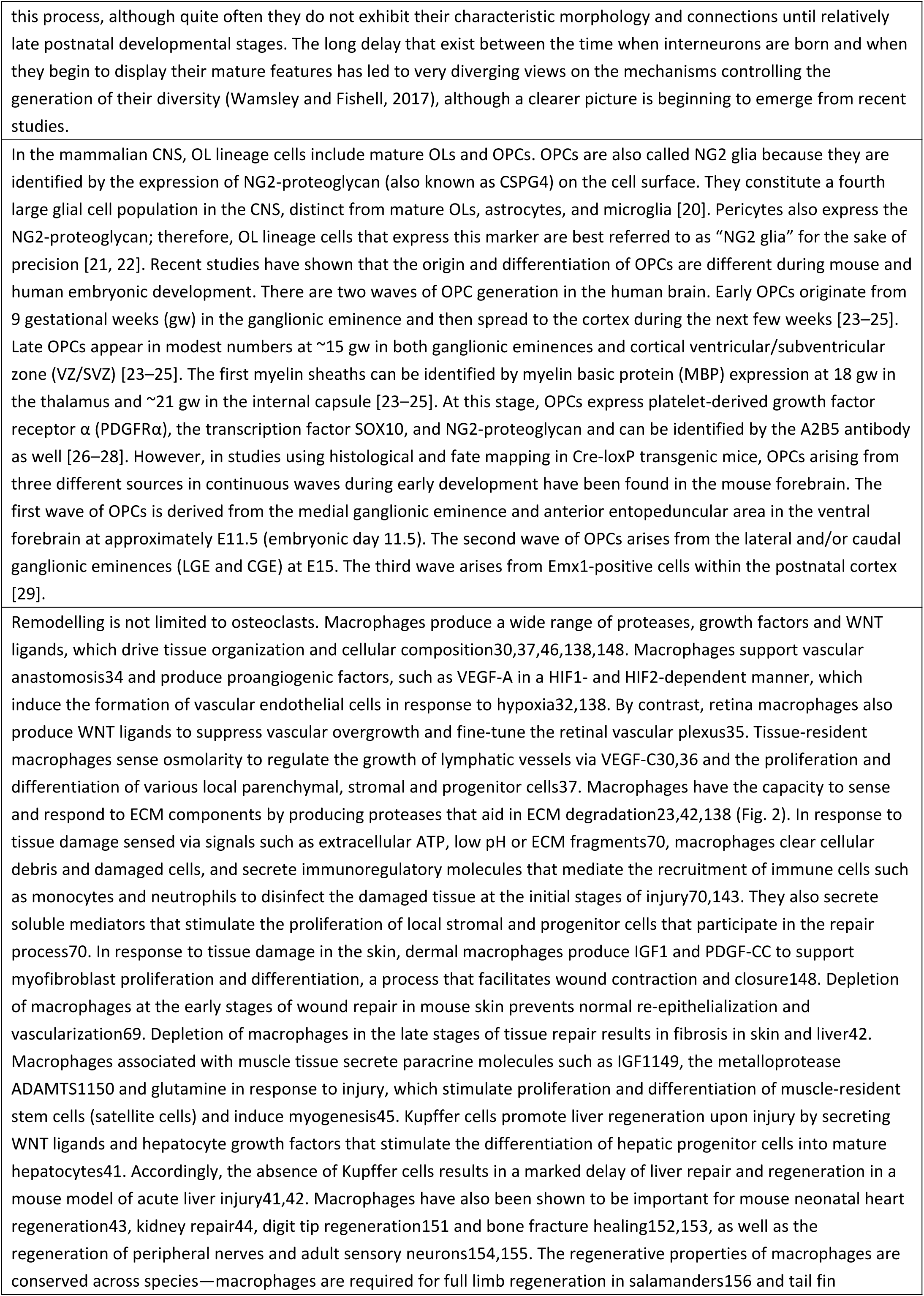
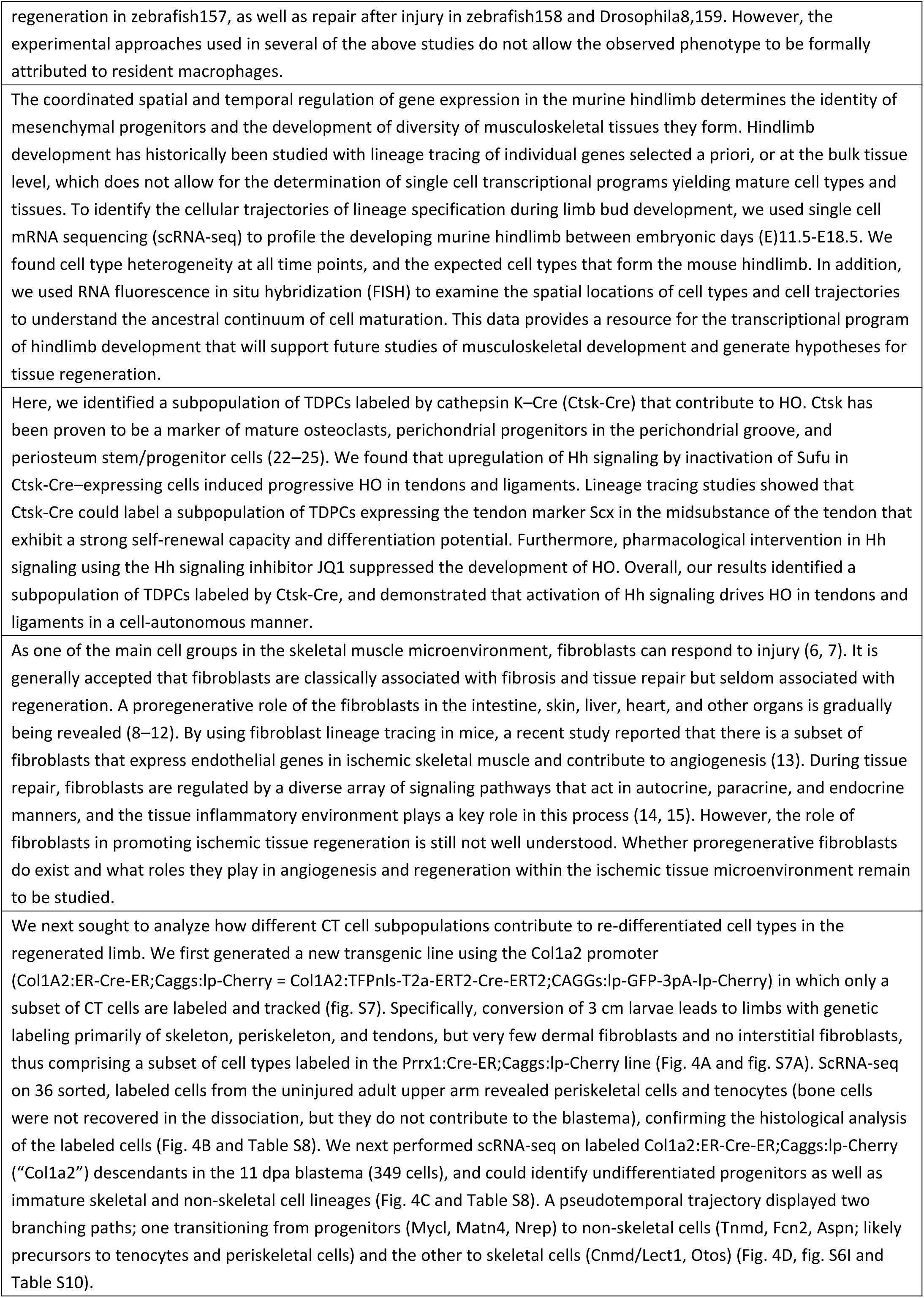
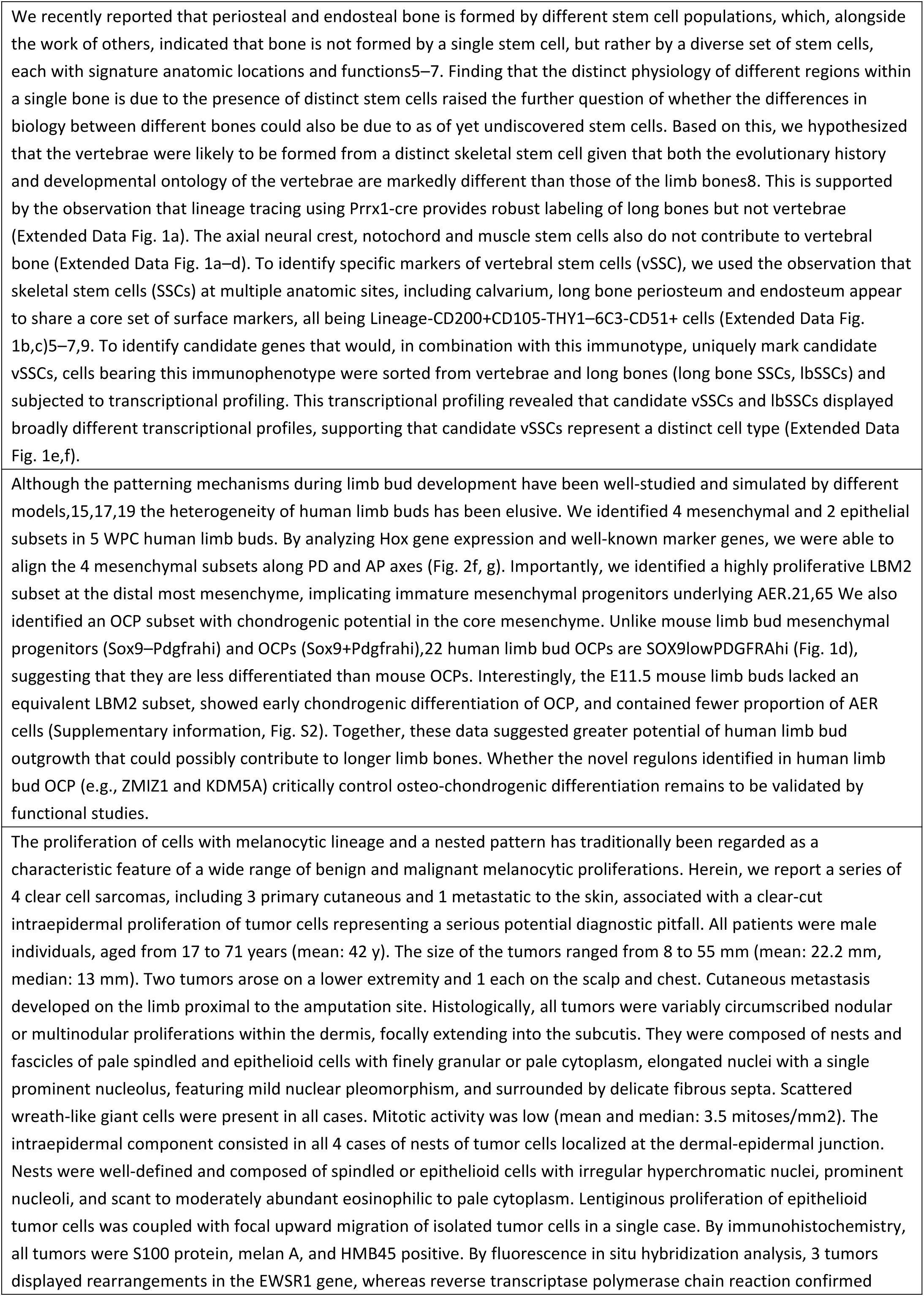
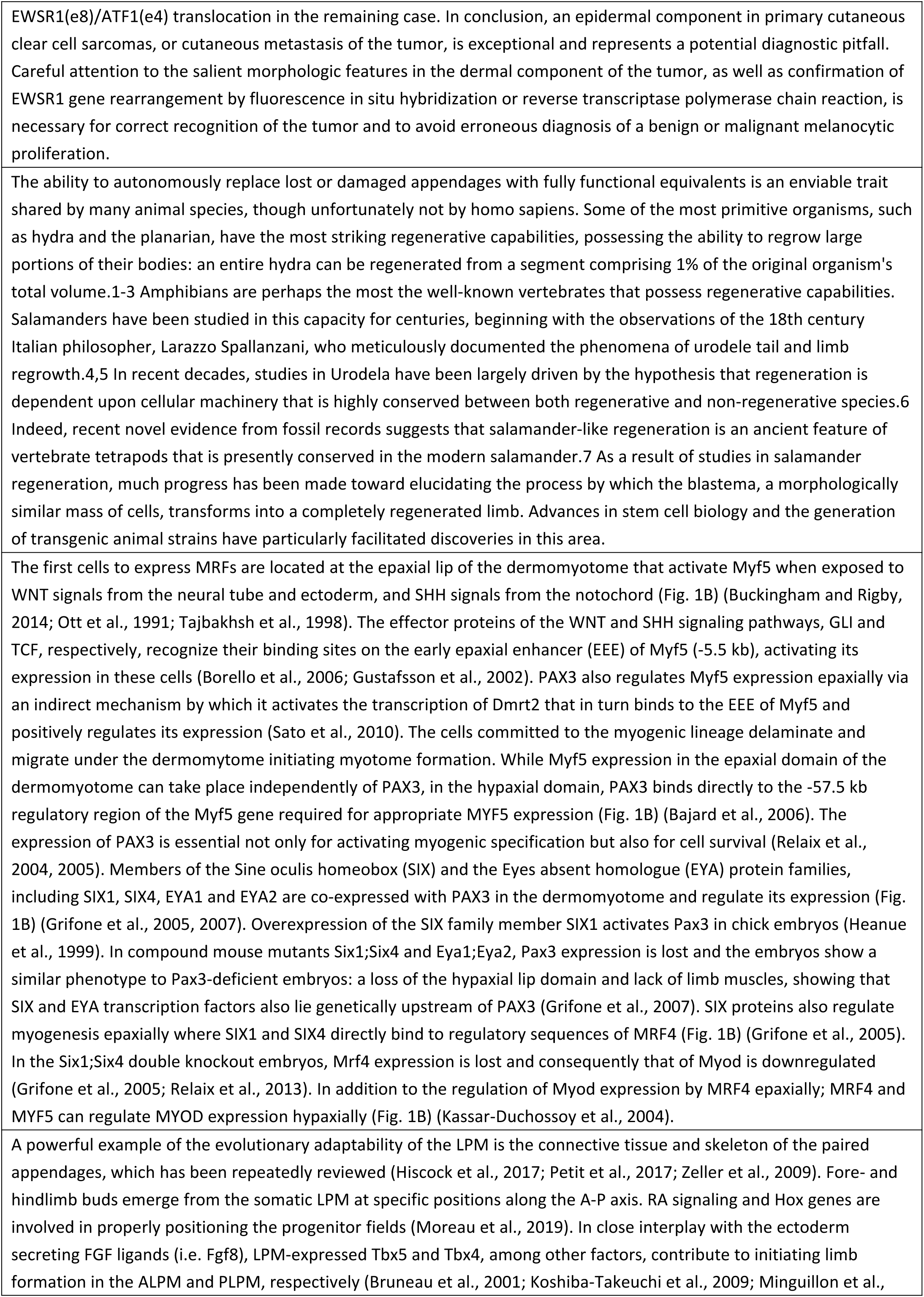
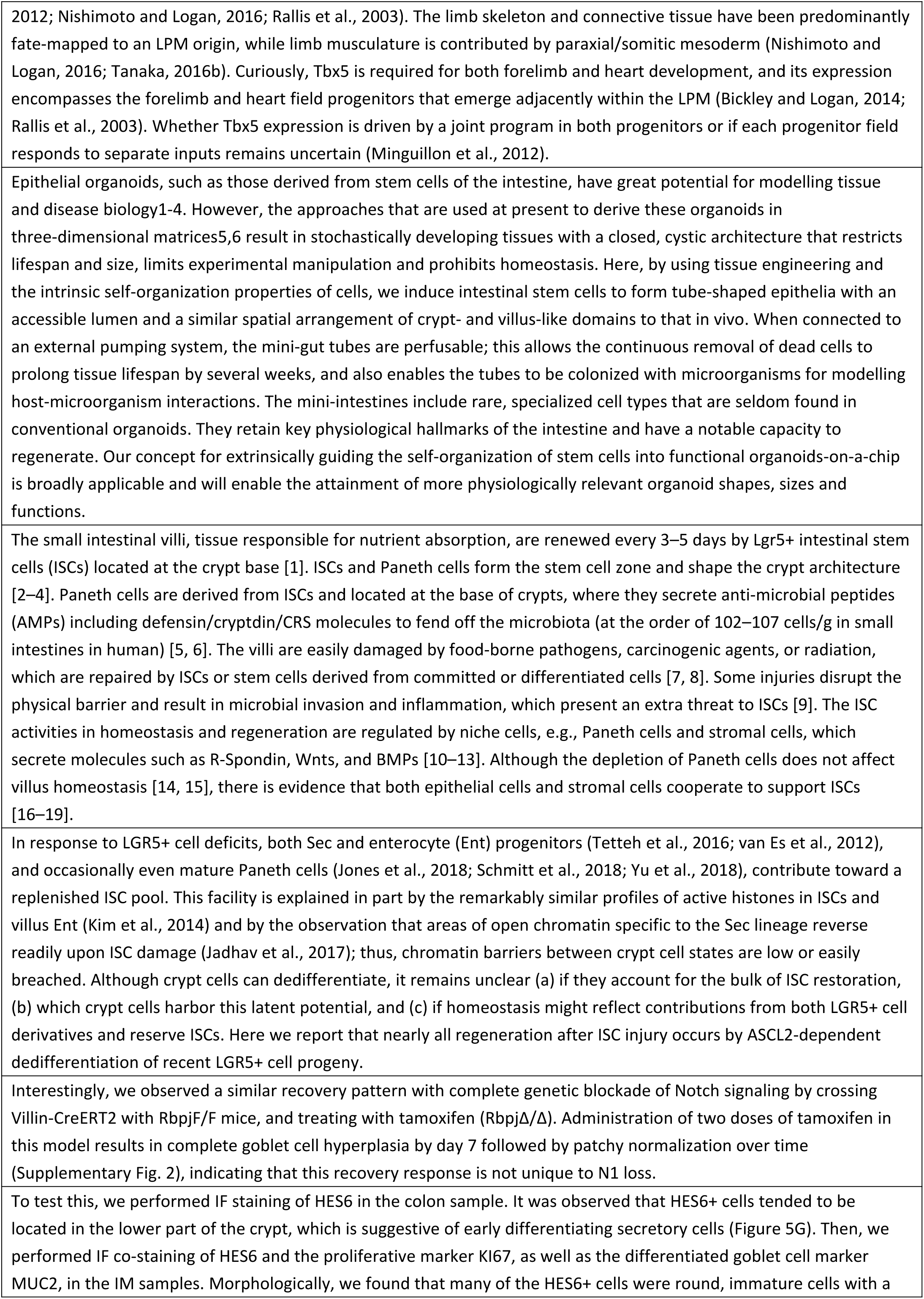
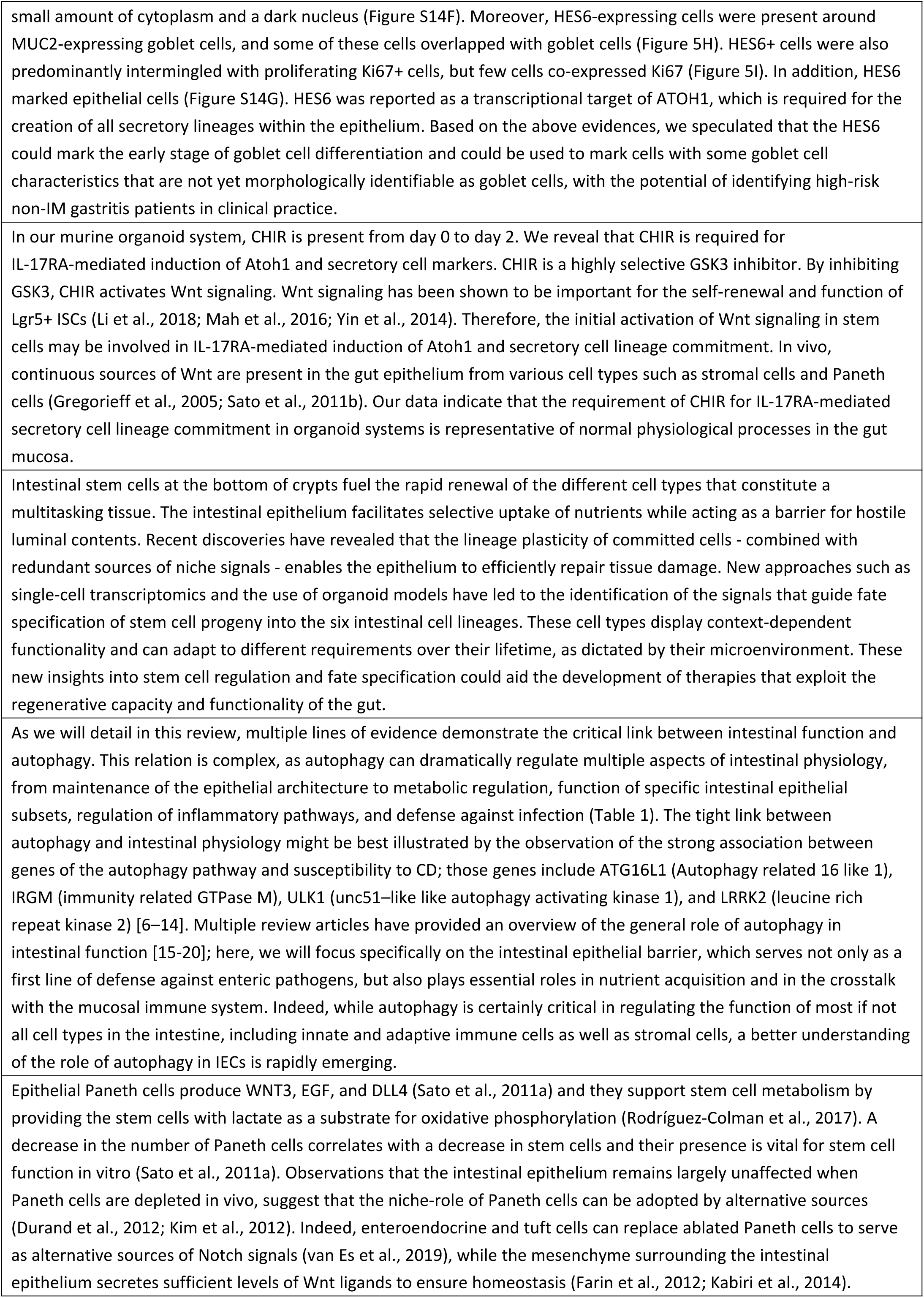
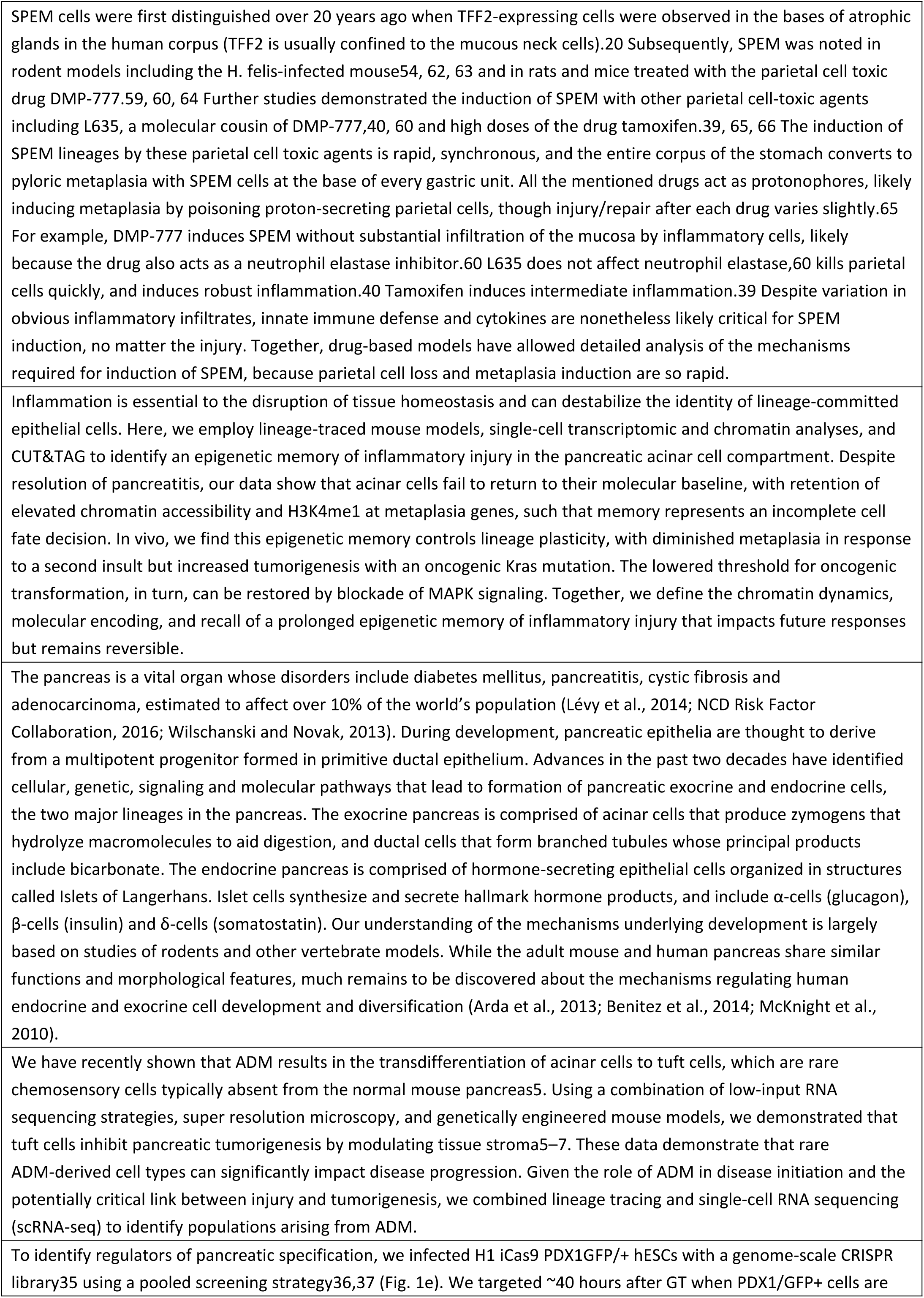
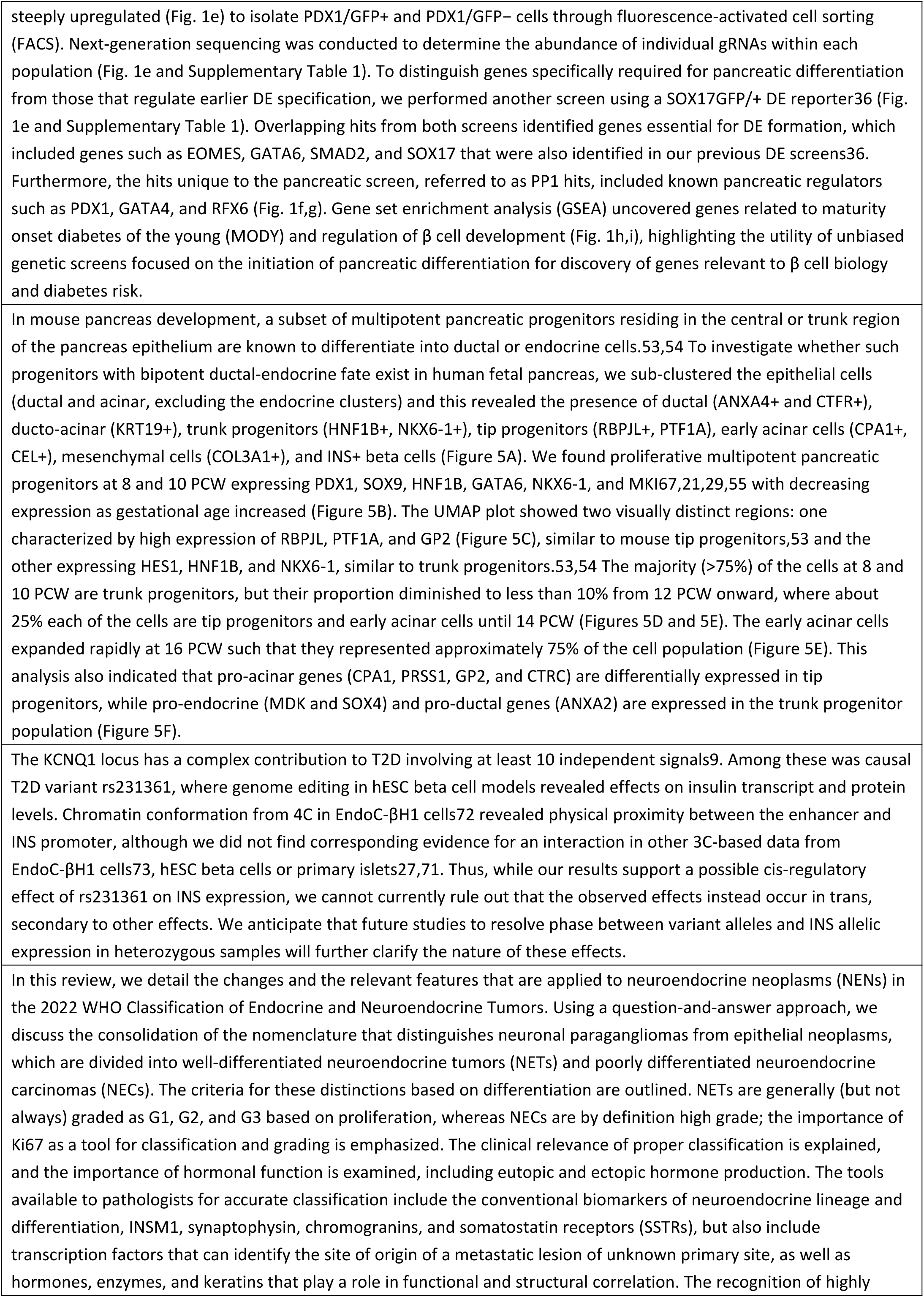
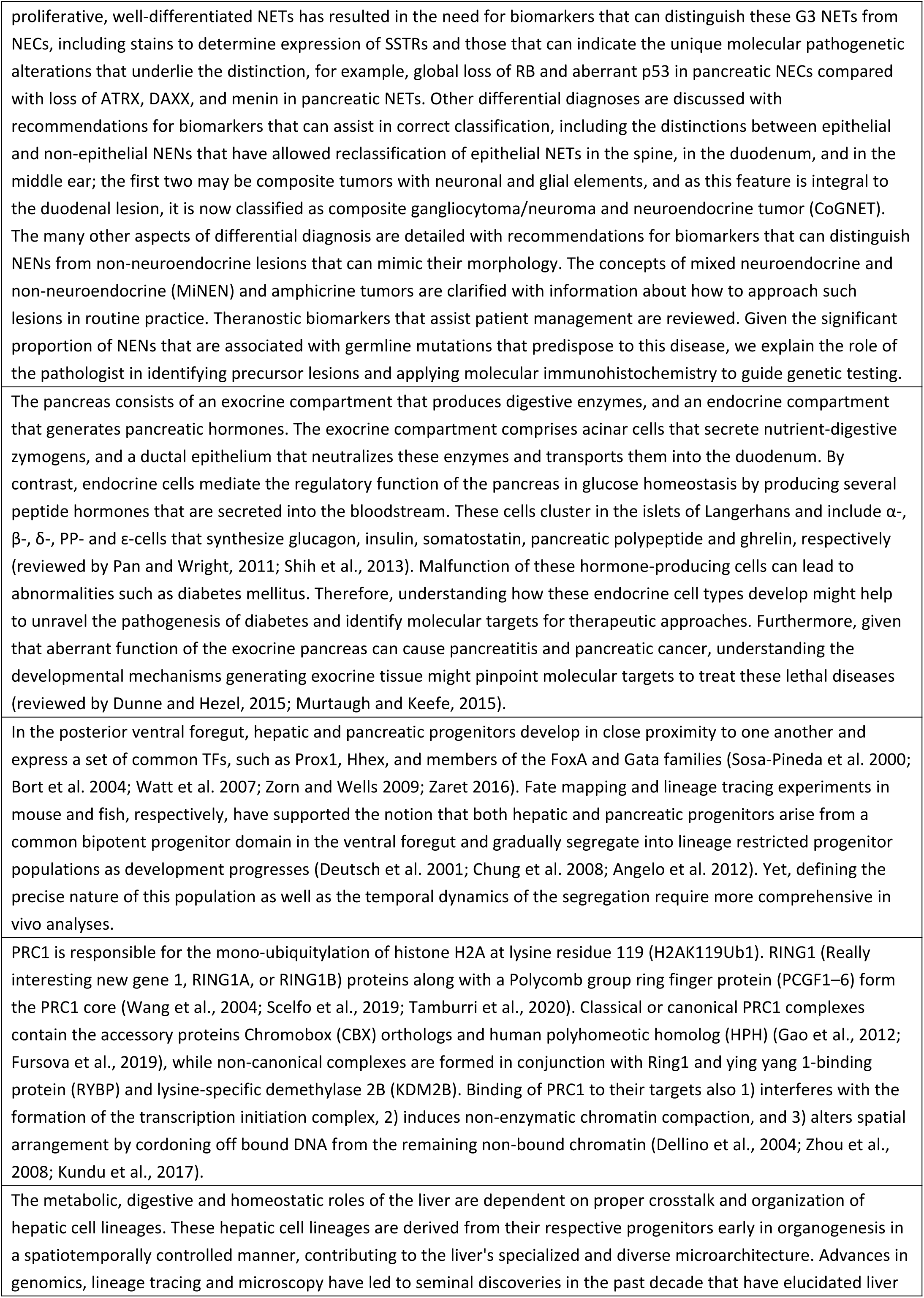
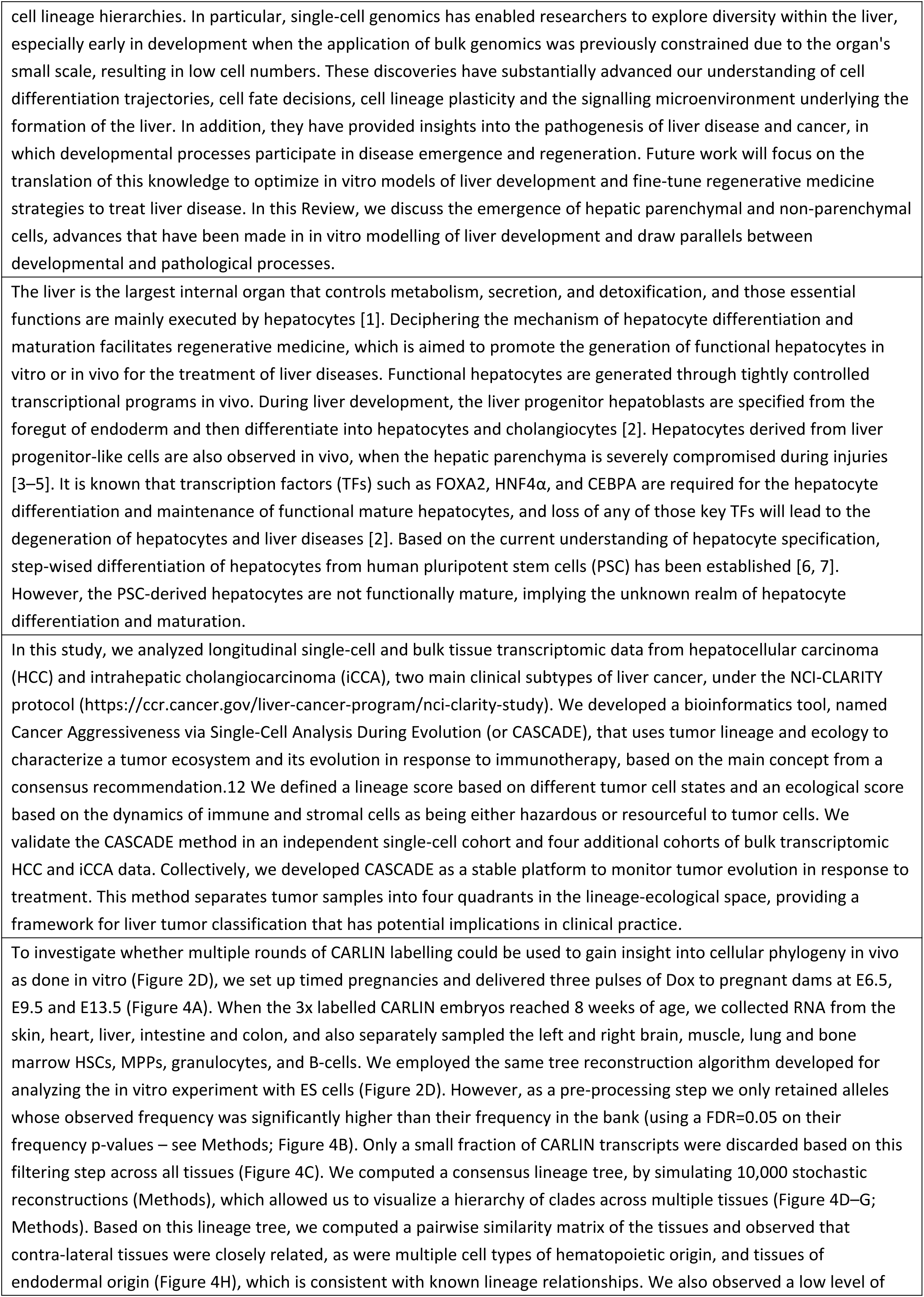
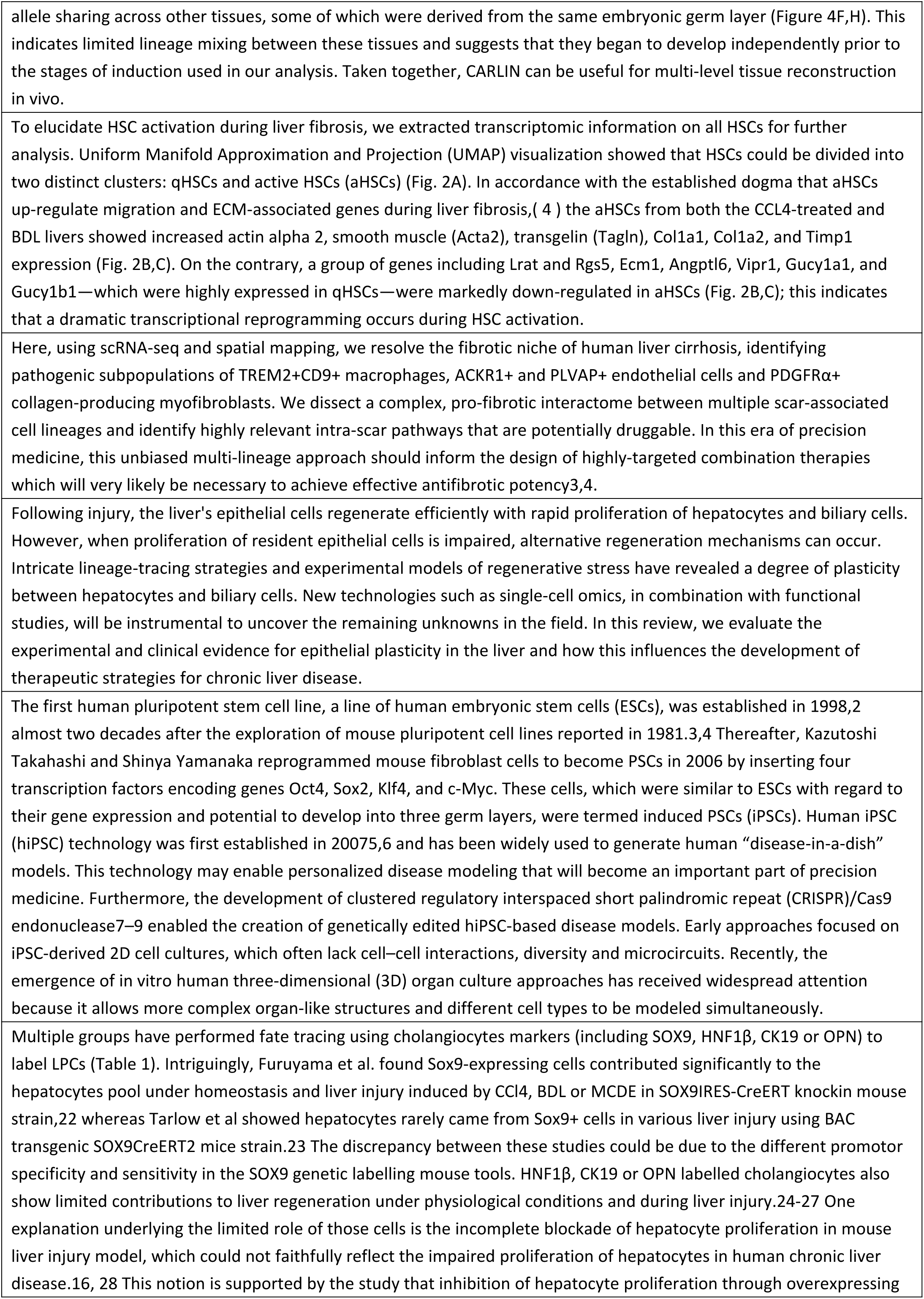
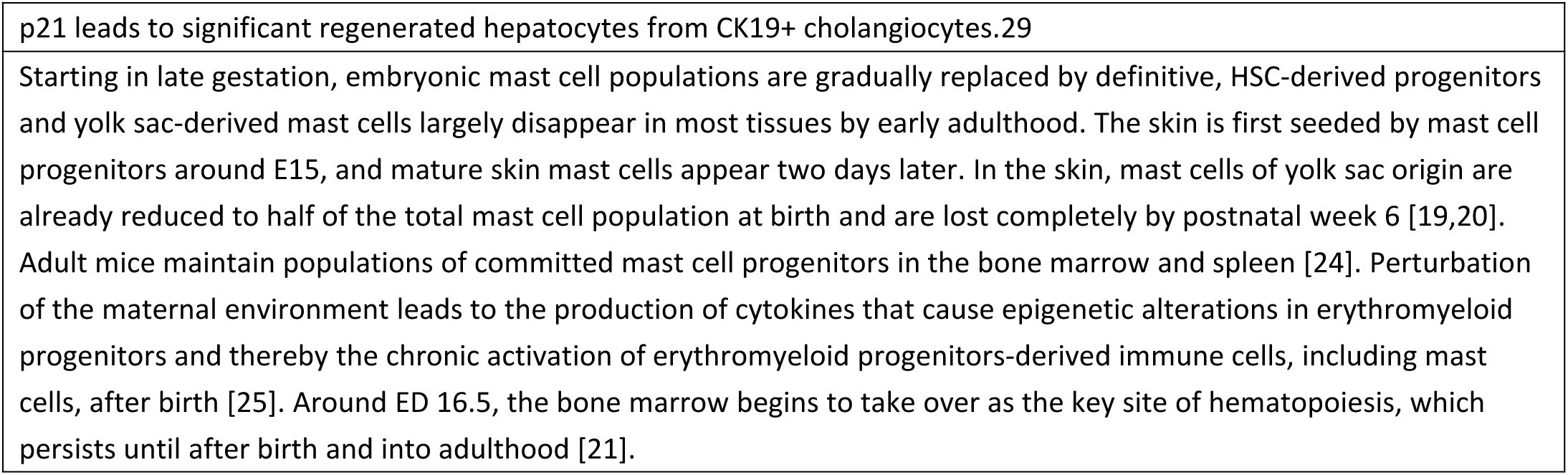
Test data for embedding models.

**Supplementary Table 15.**
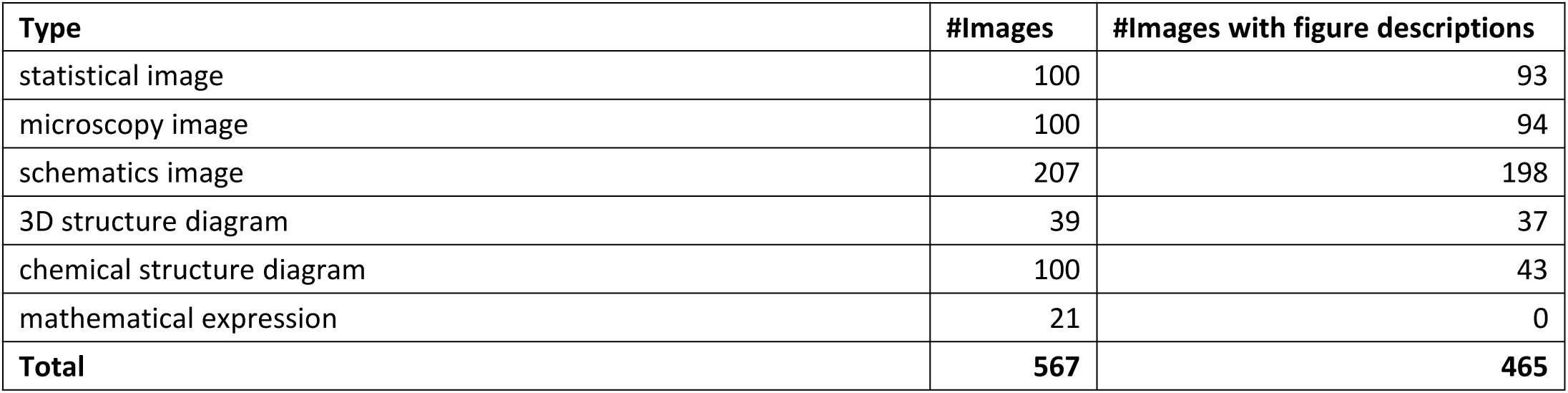
Number of test data for image type identification.

**Supplementary Table 16.**
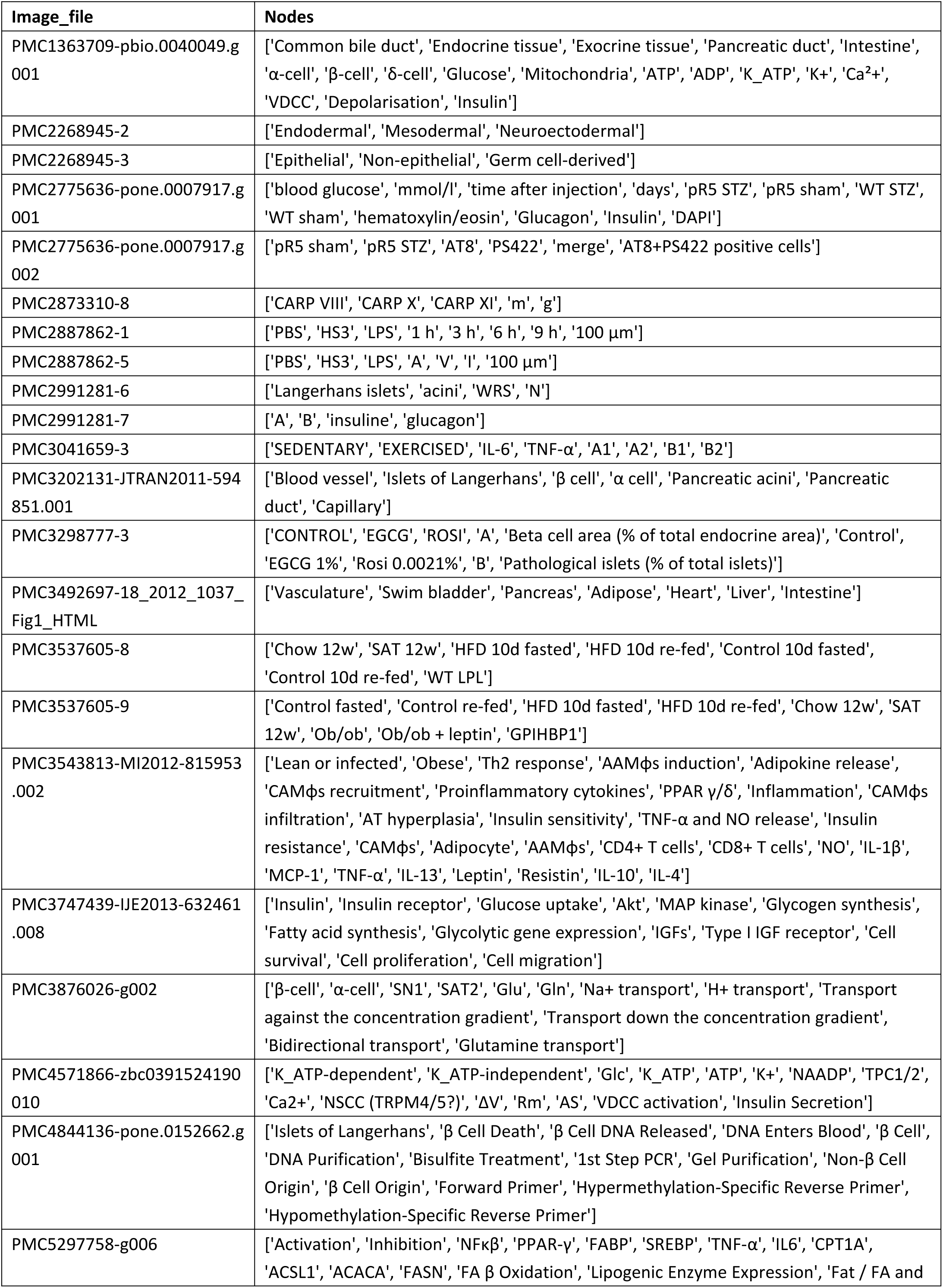
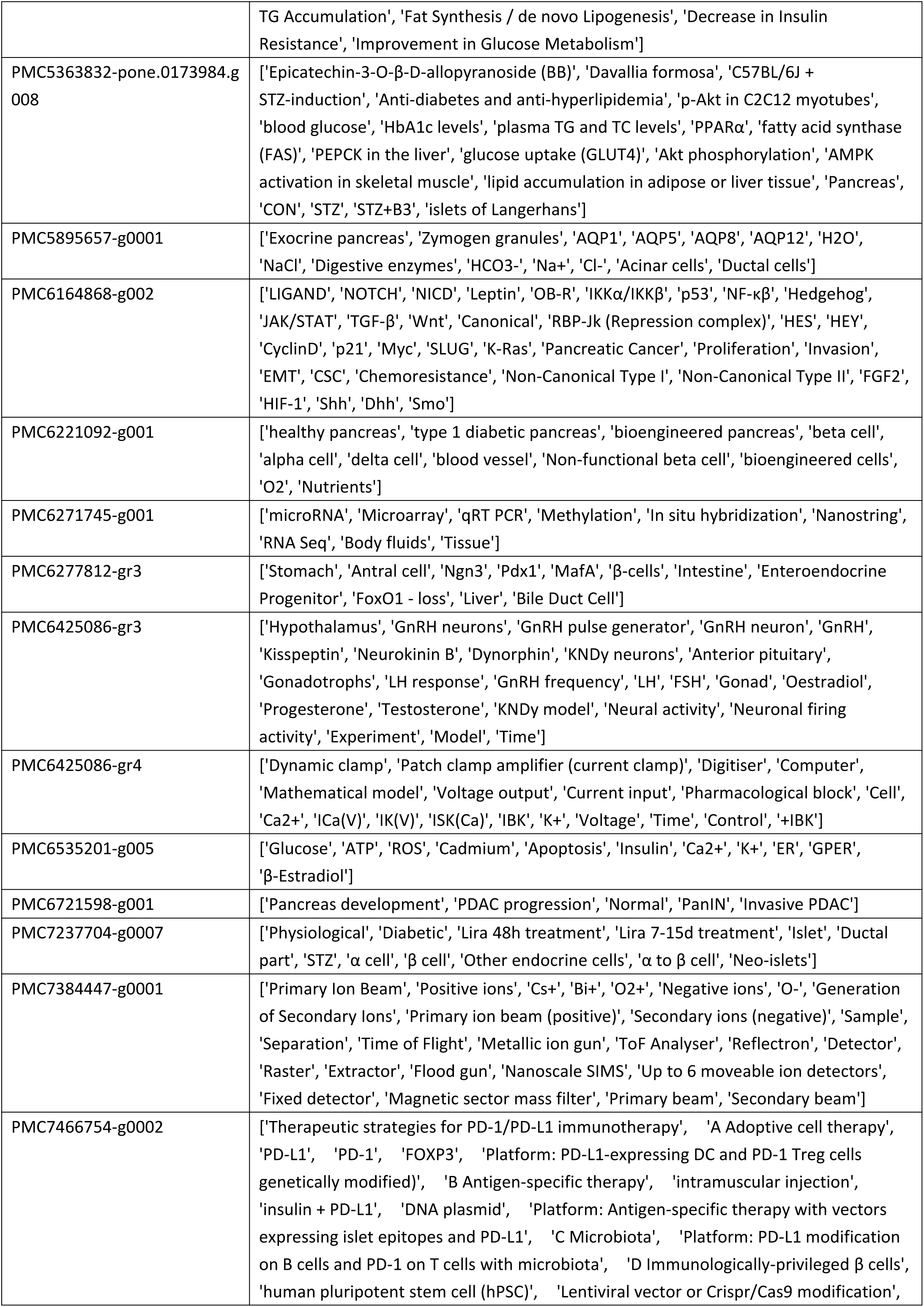
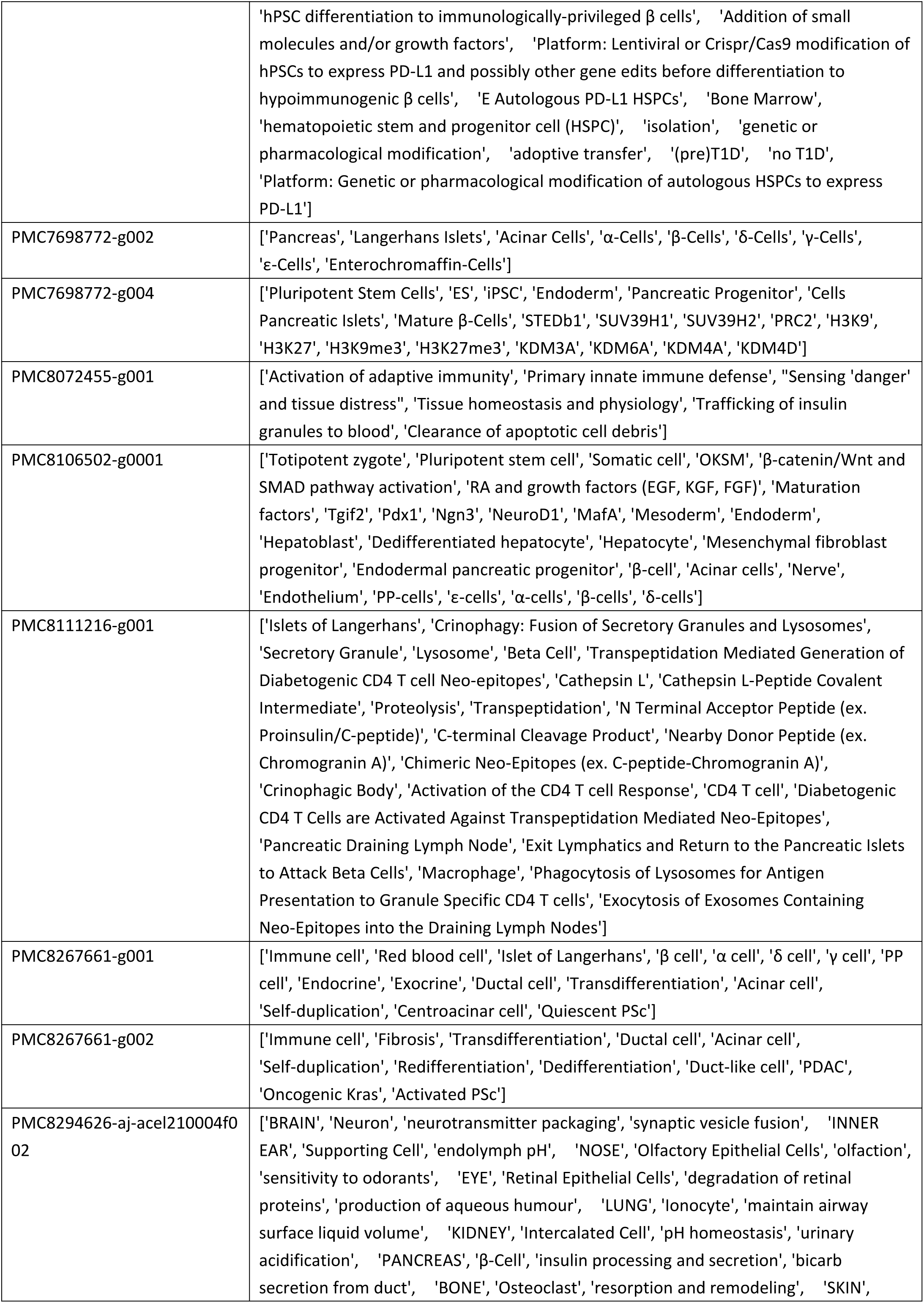
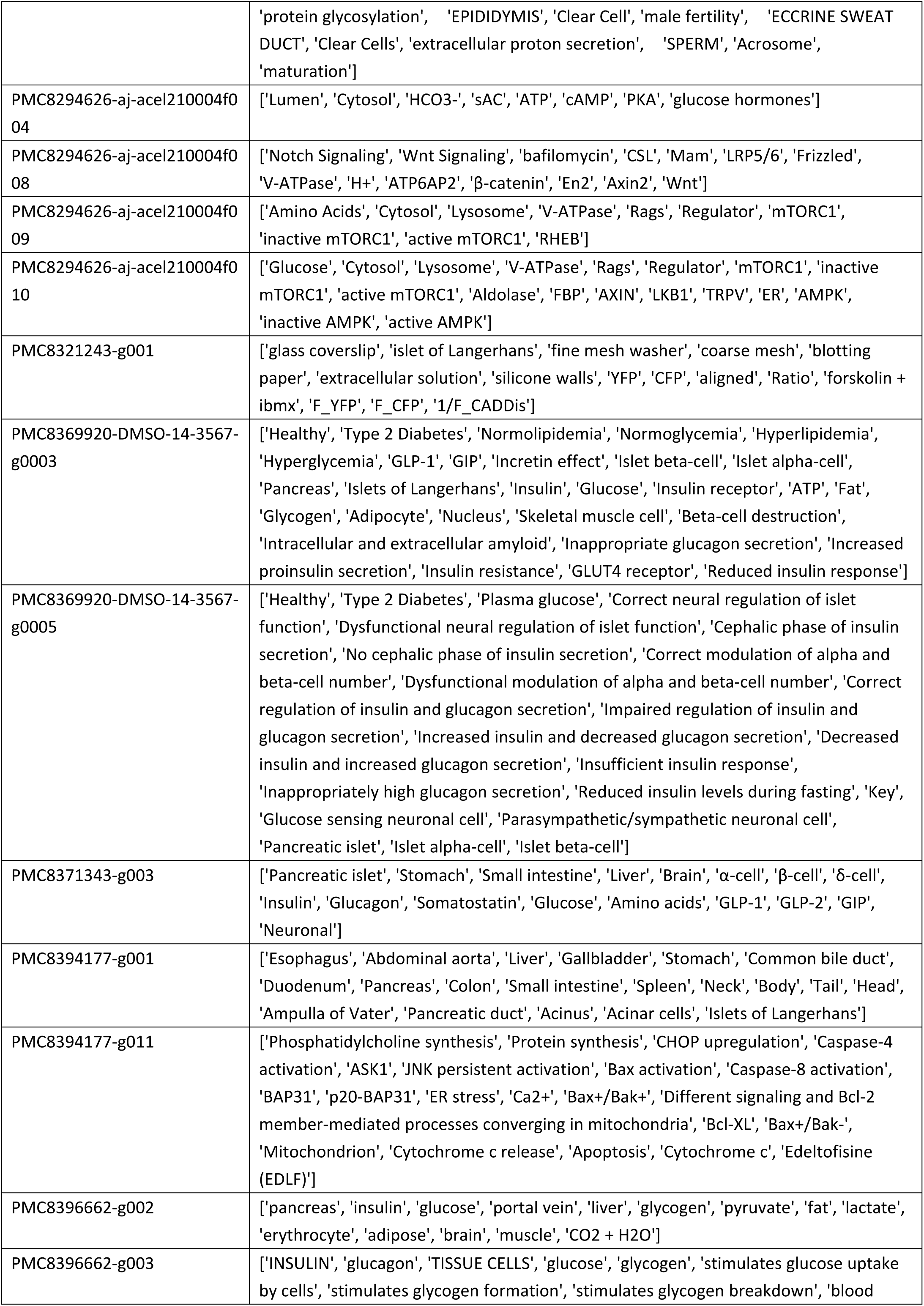
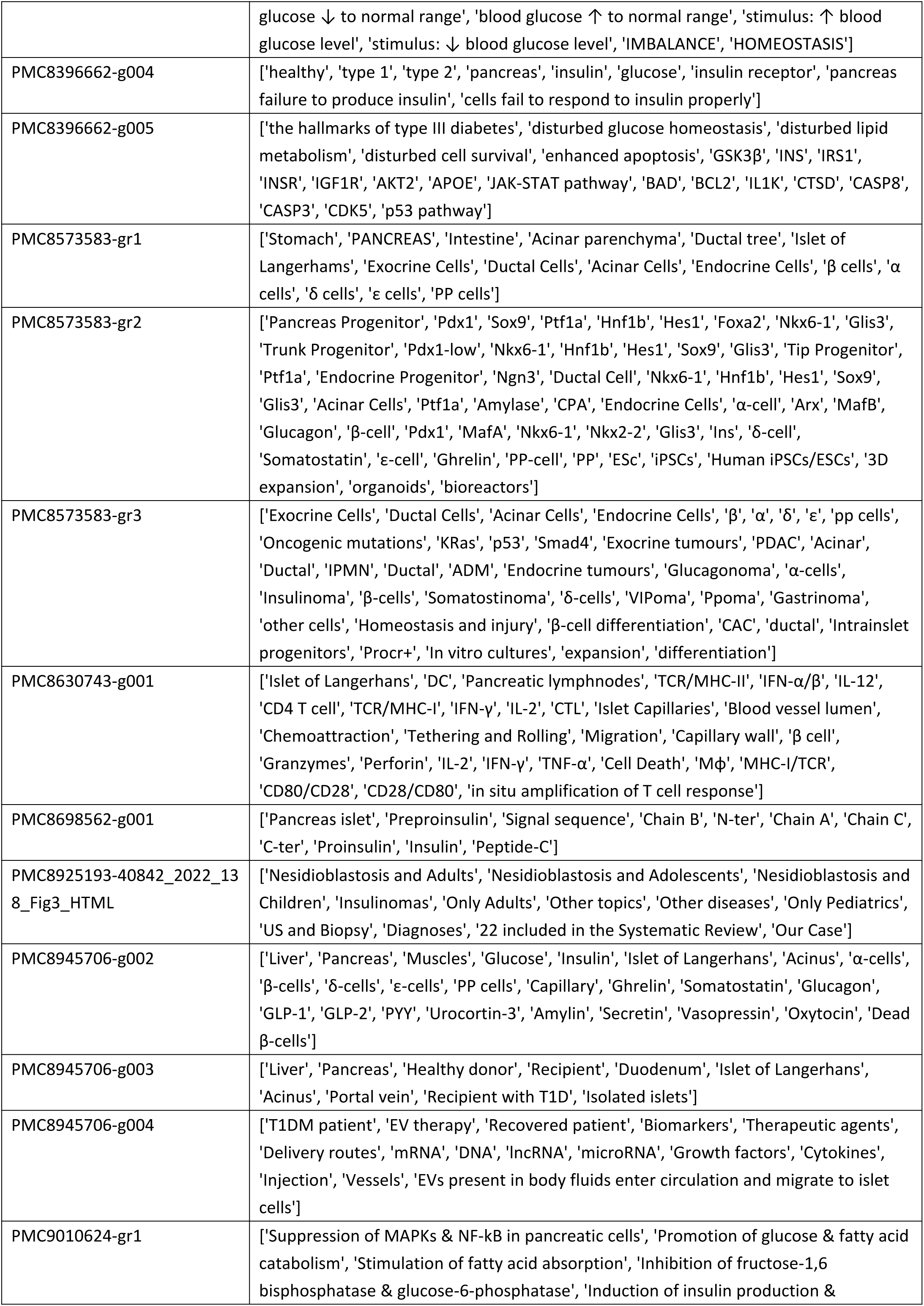
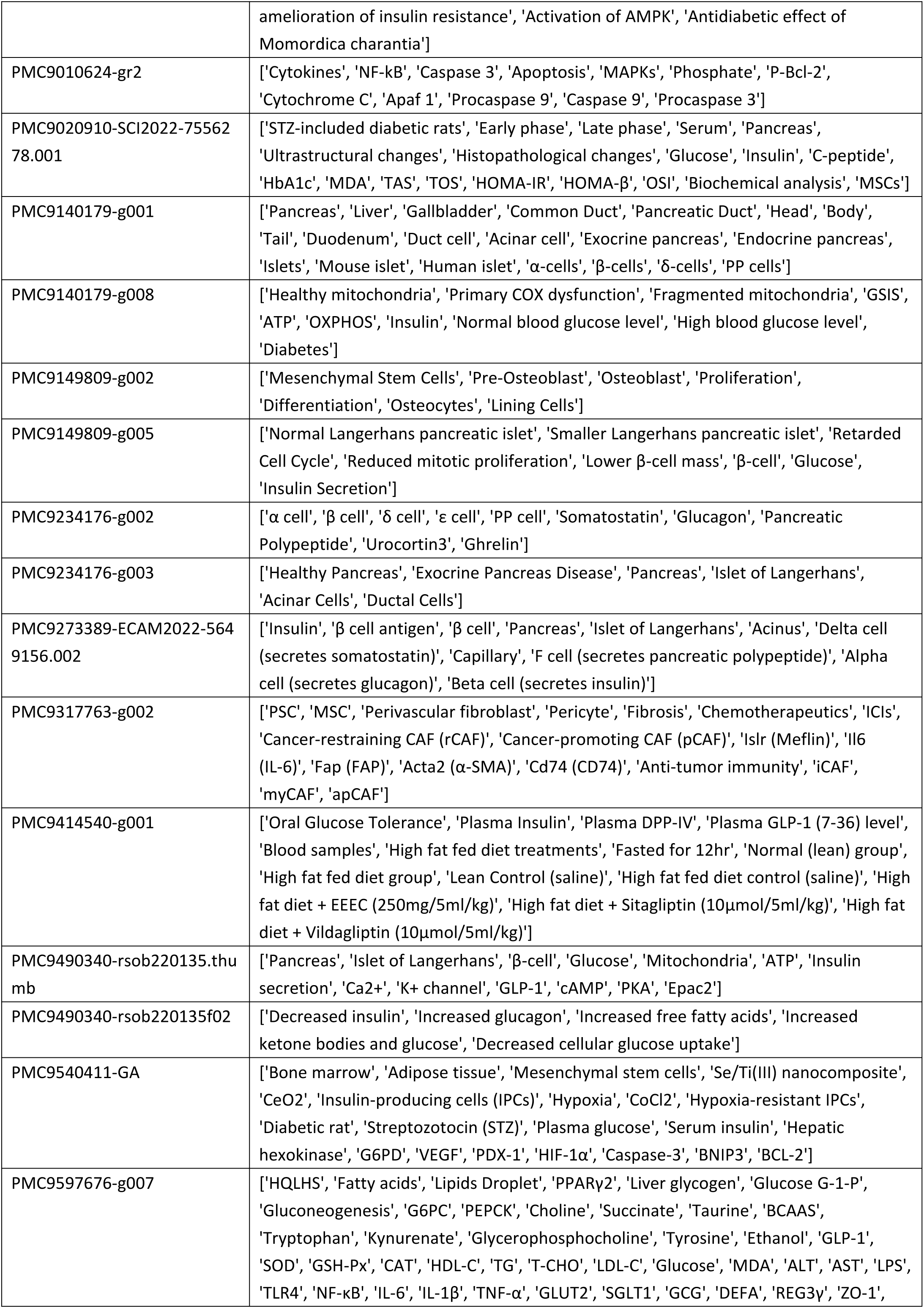
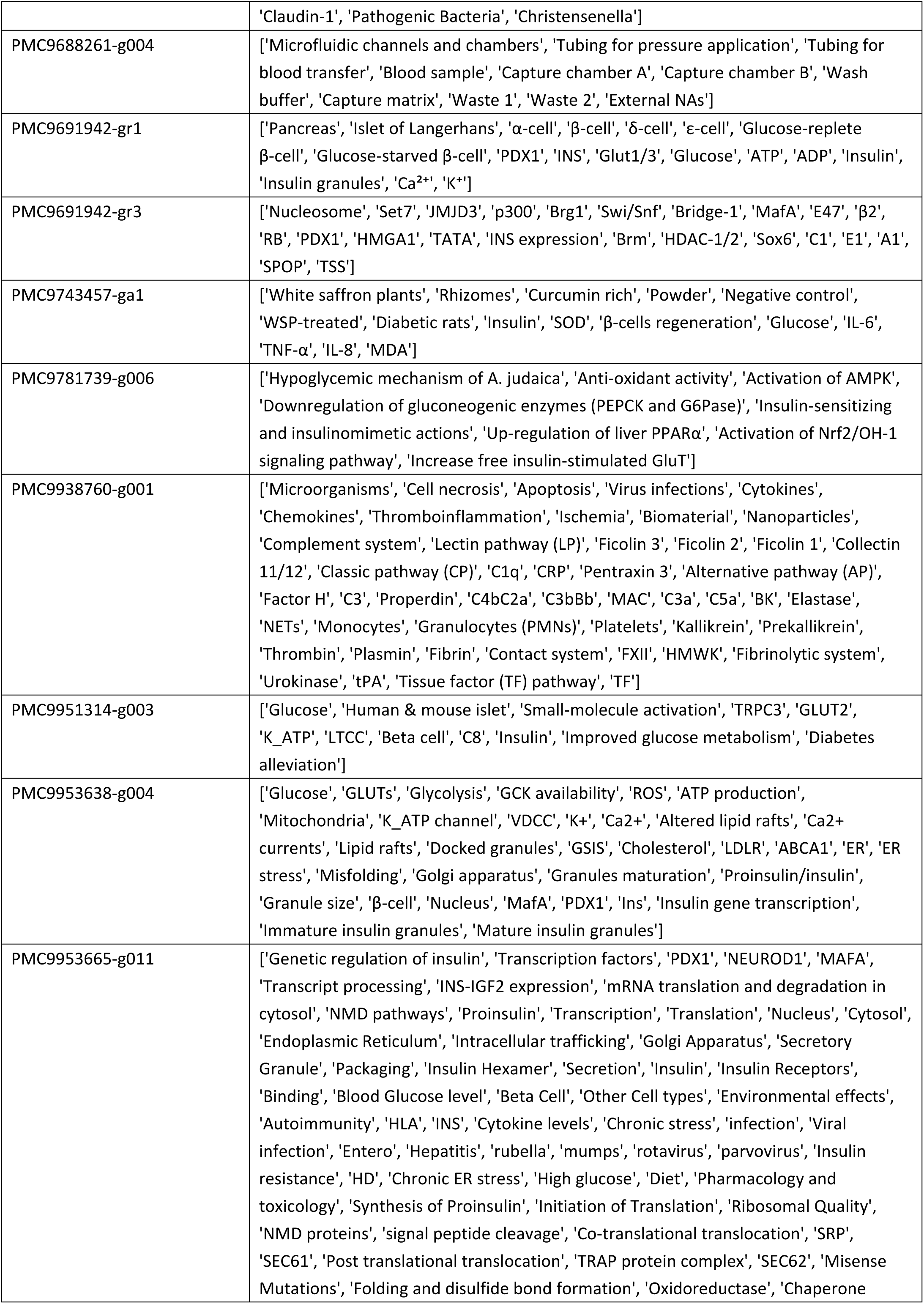
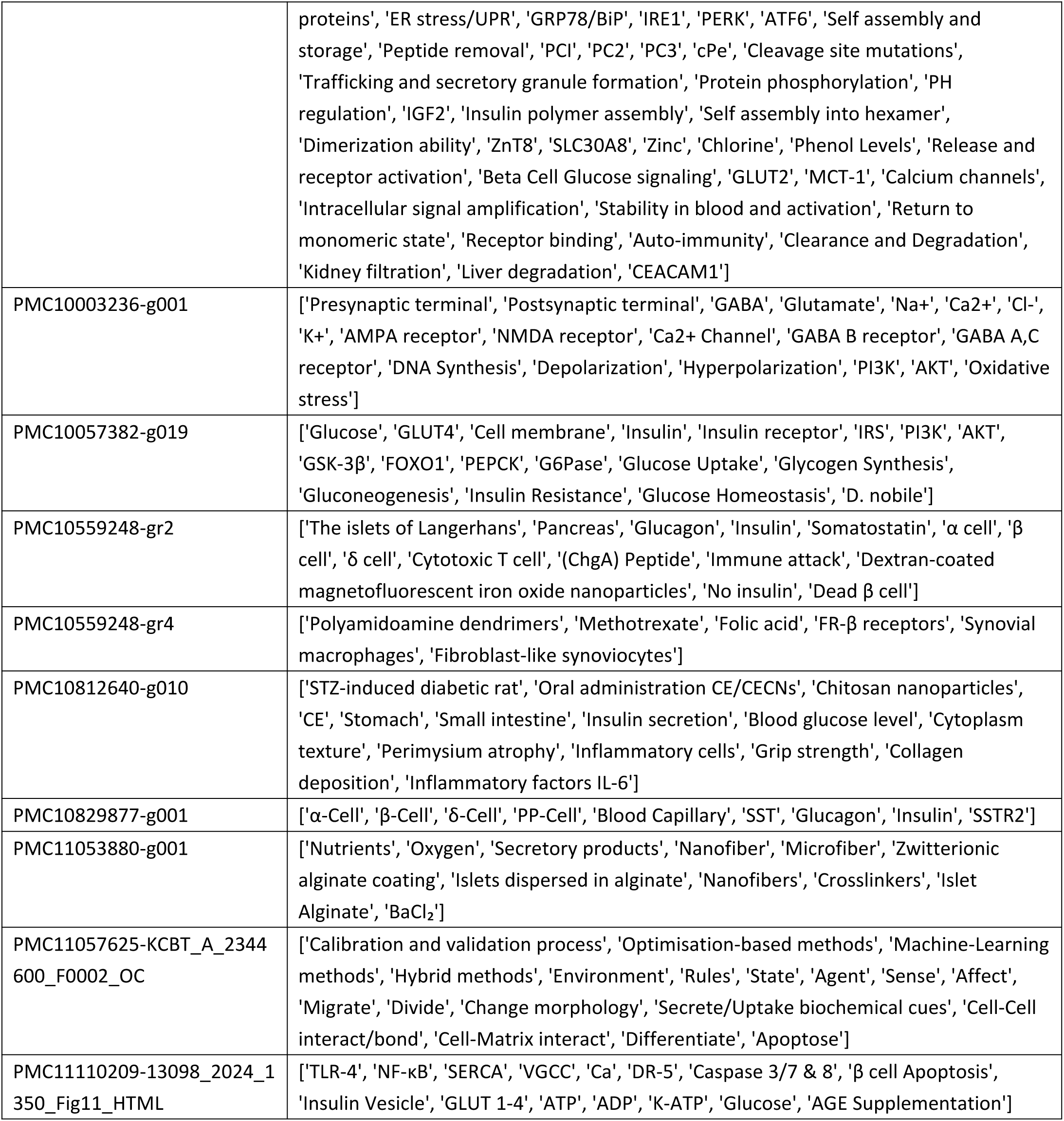
Ground true for image-entity extraction.

**Supplementary Table 17.**
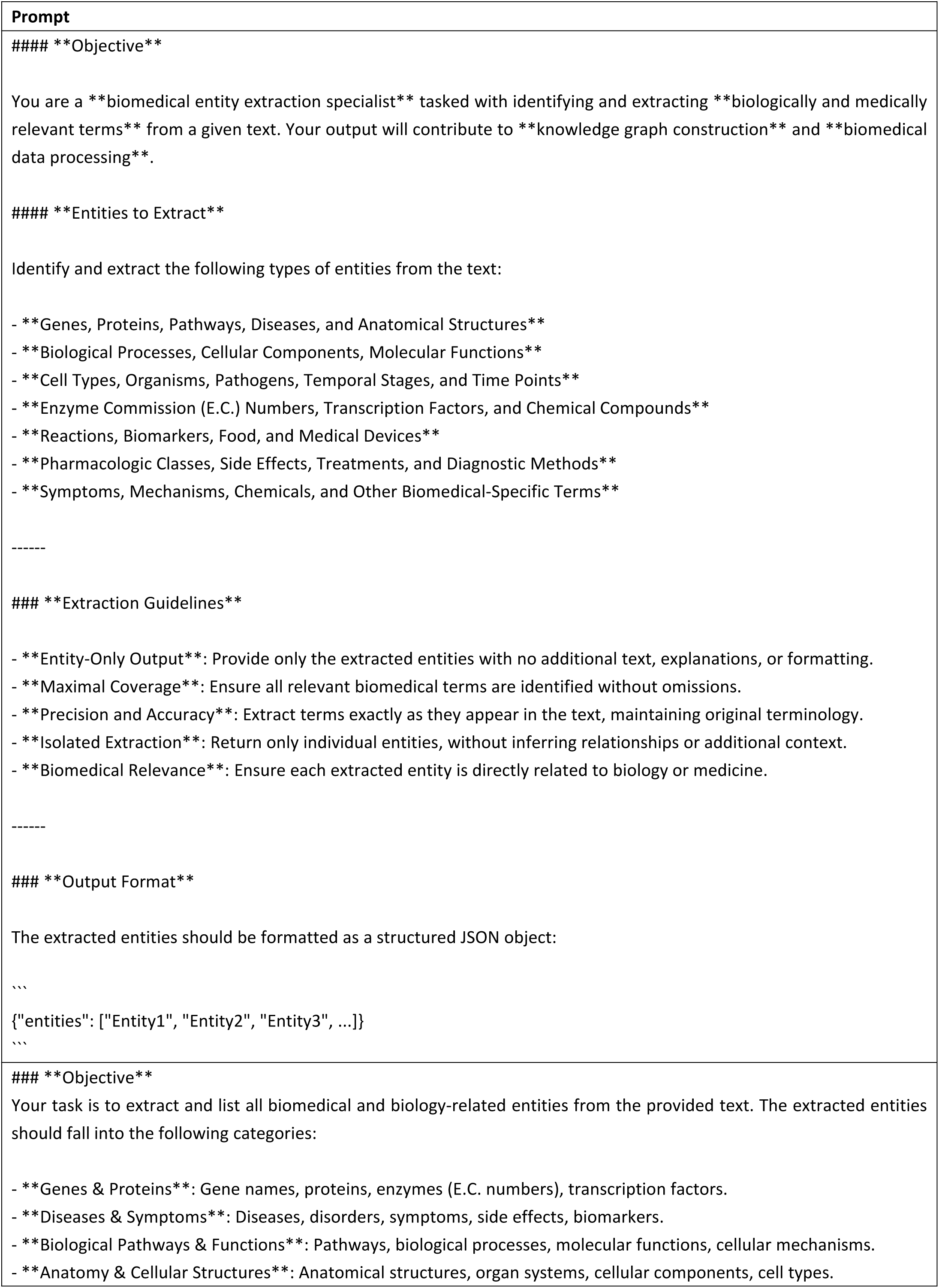
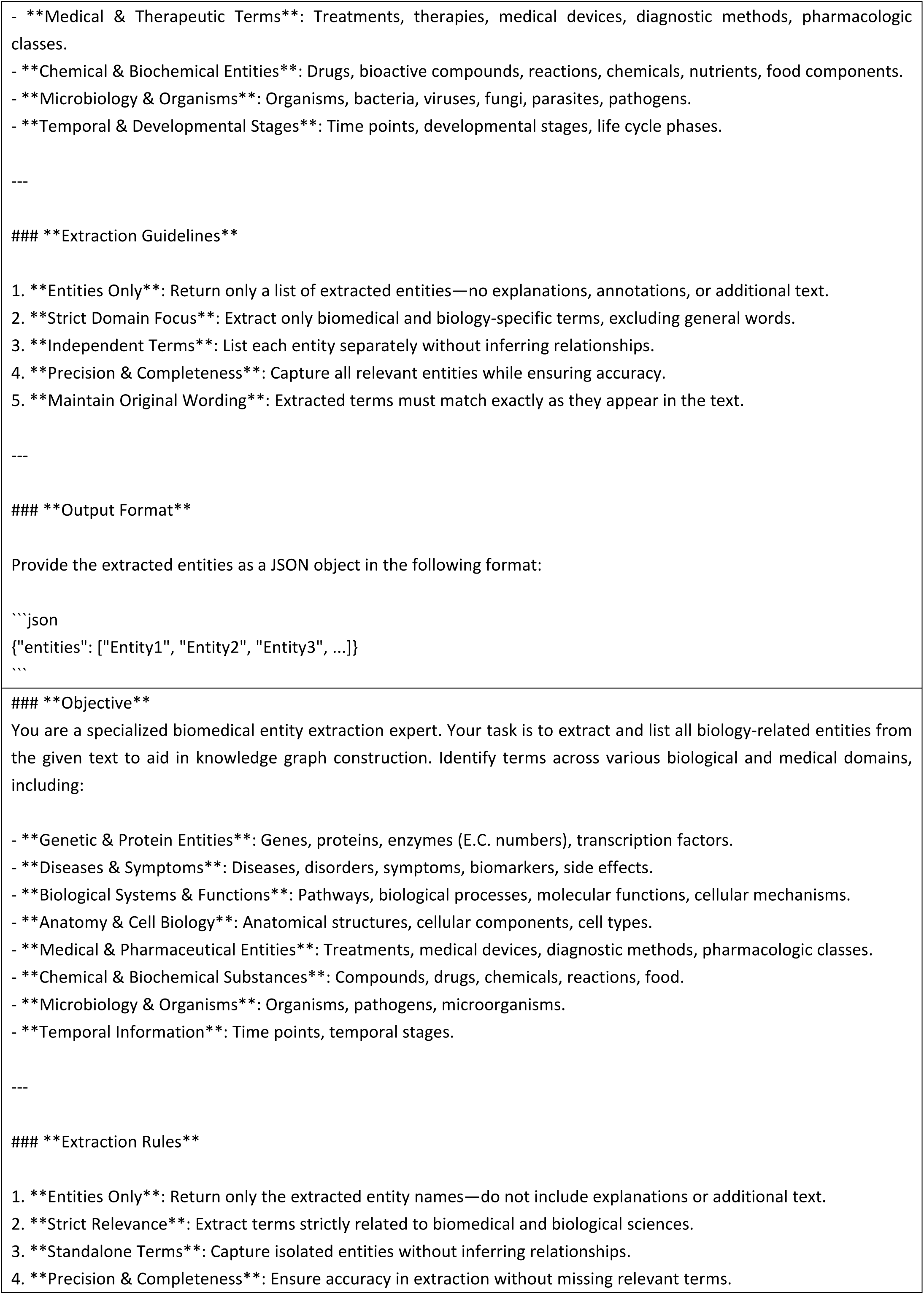
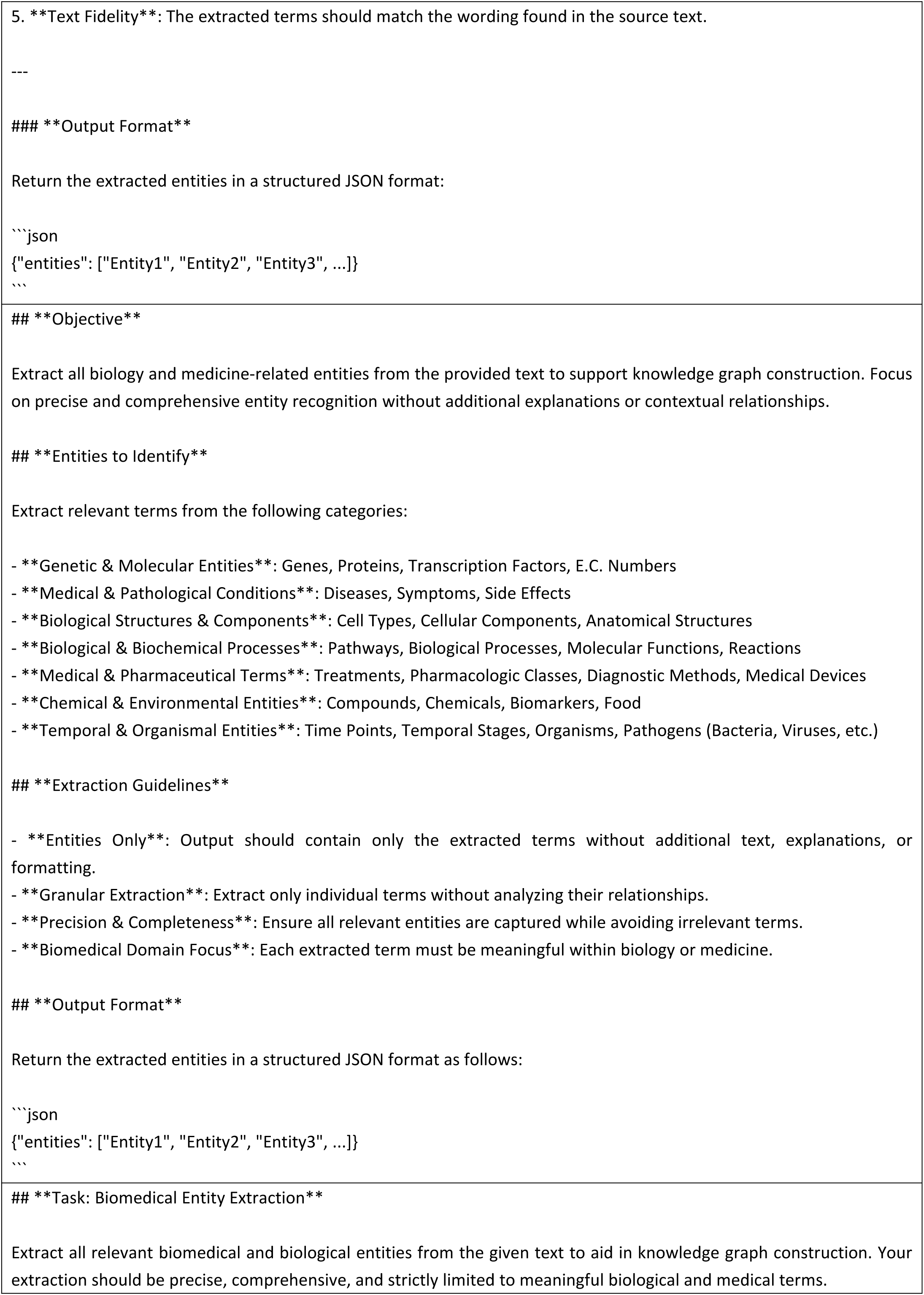
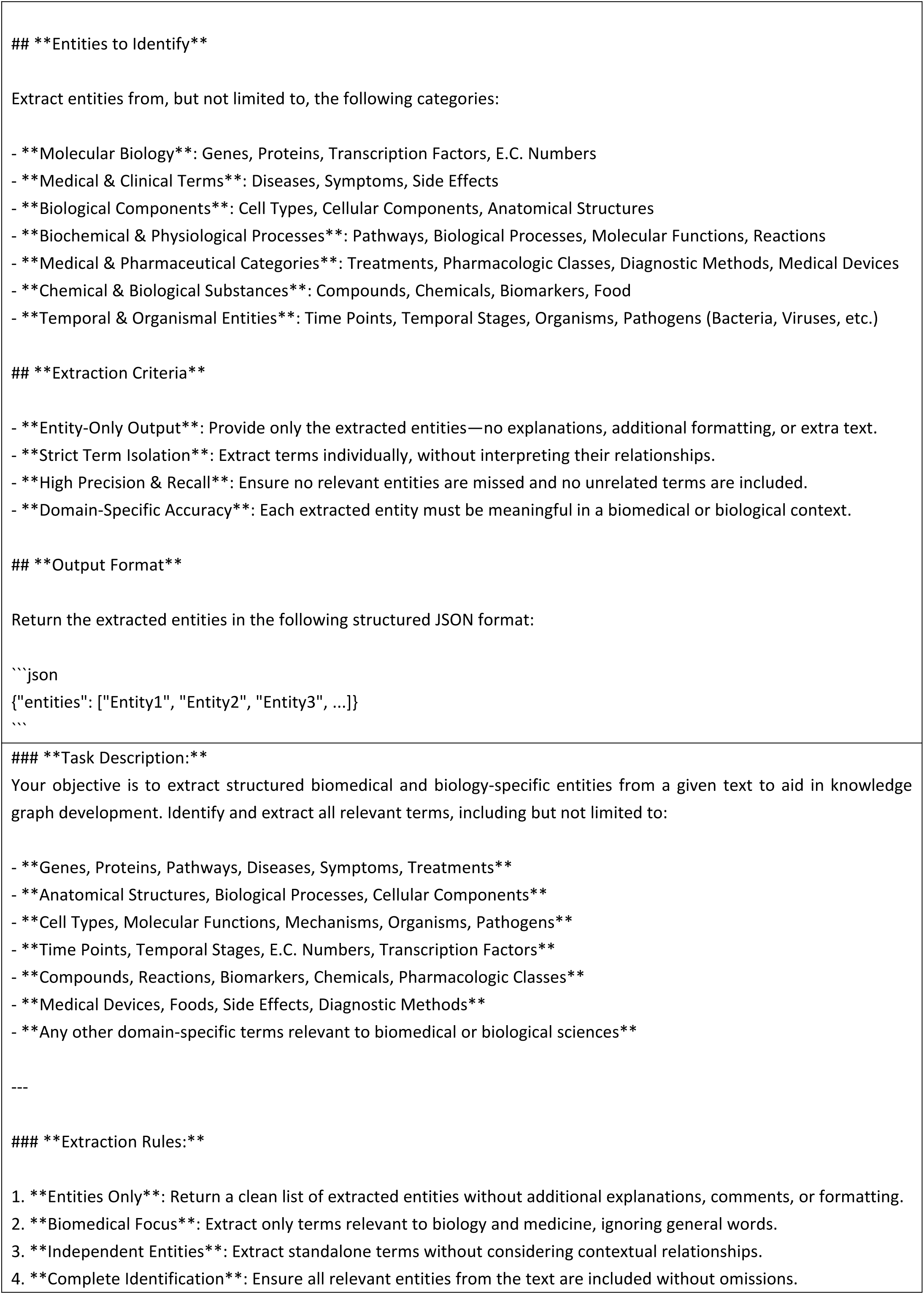
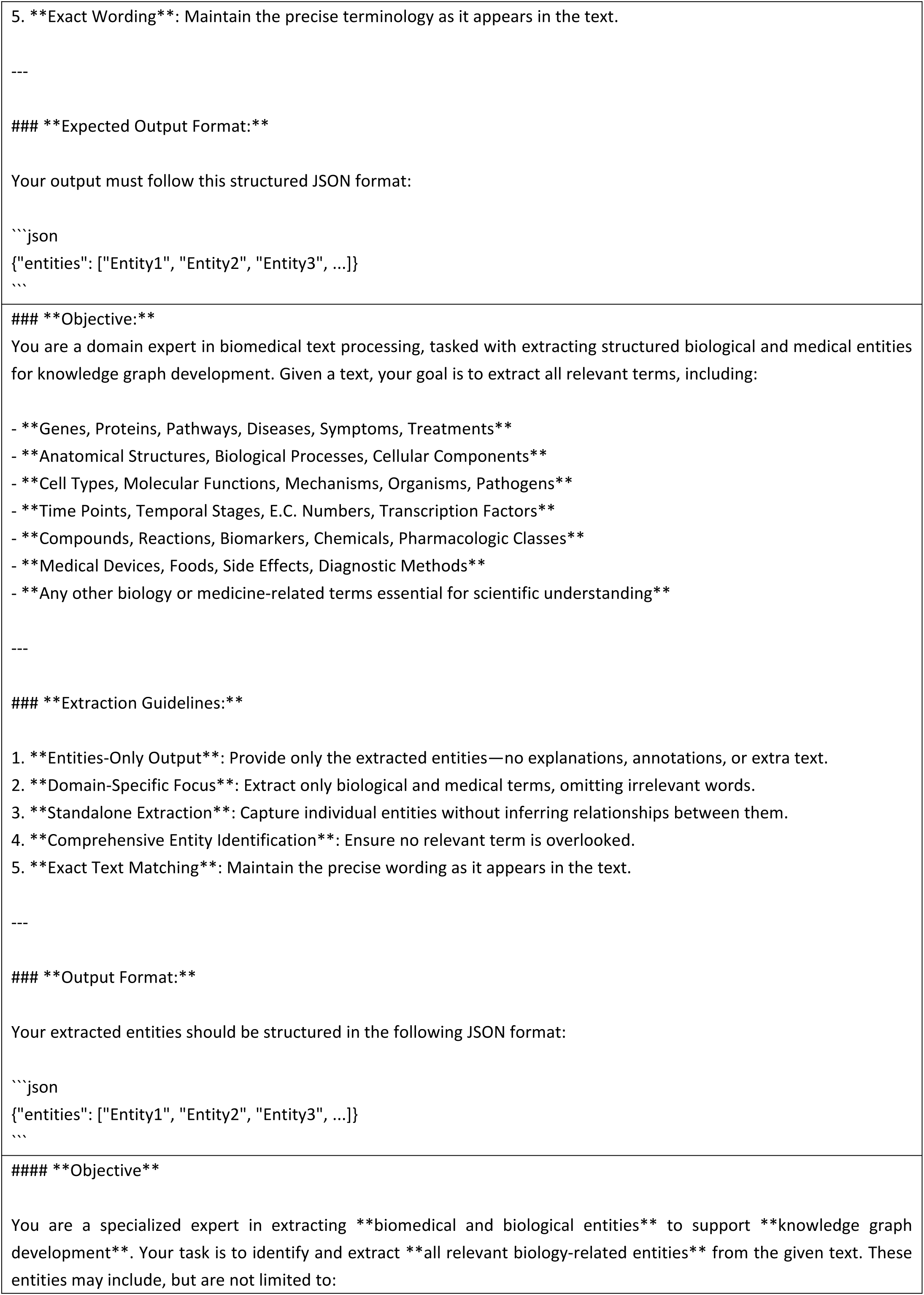
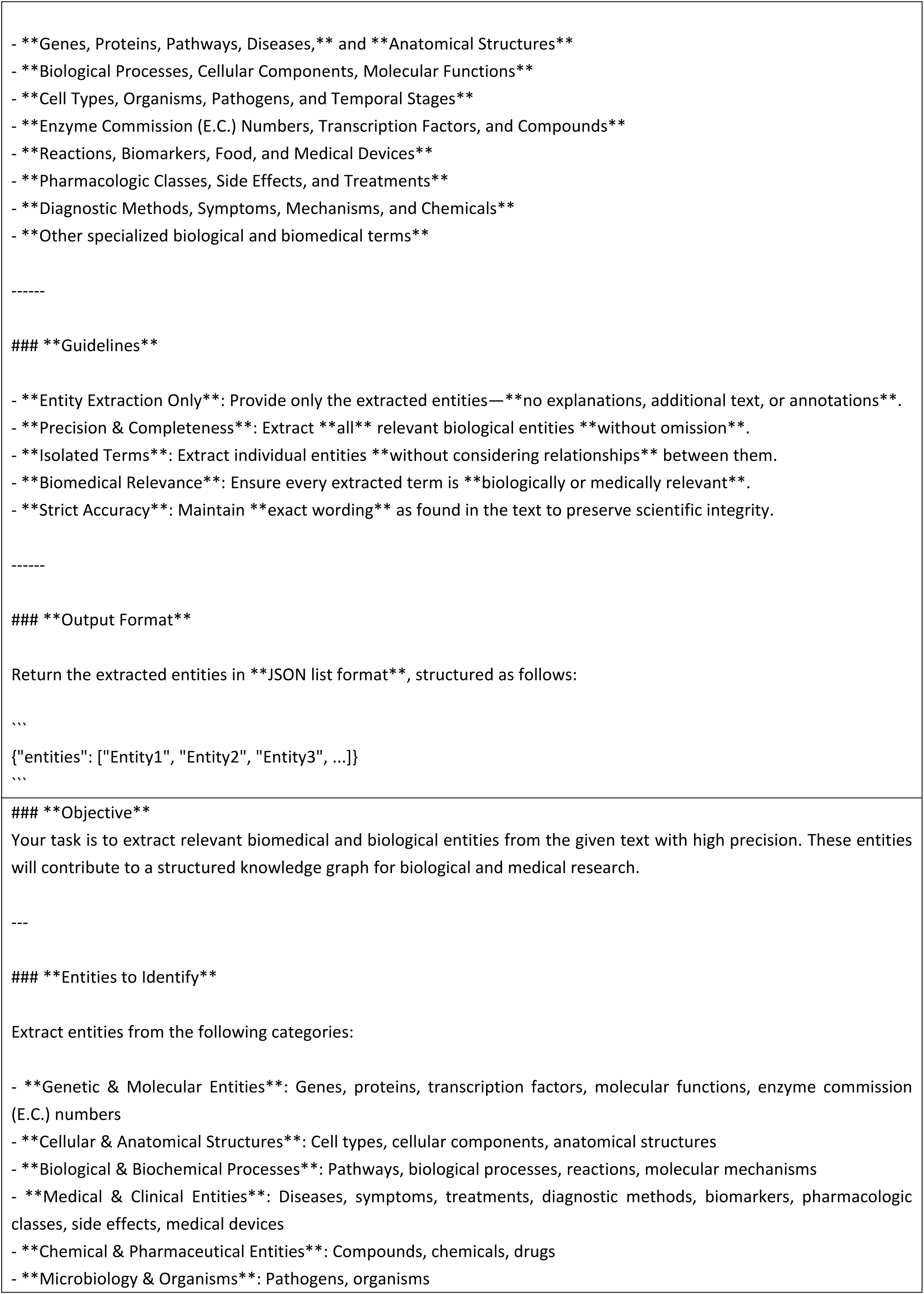
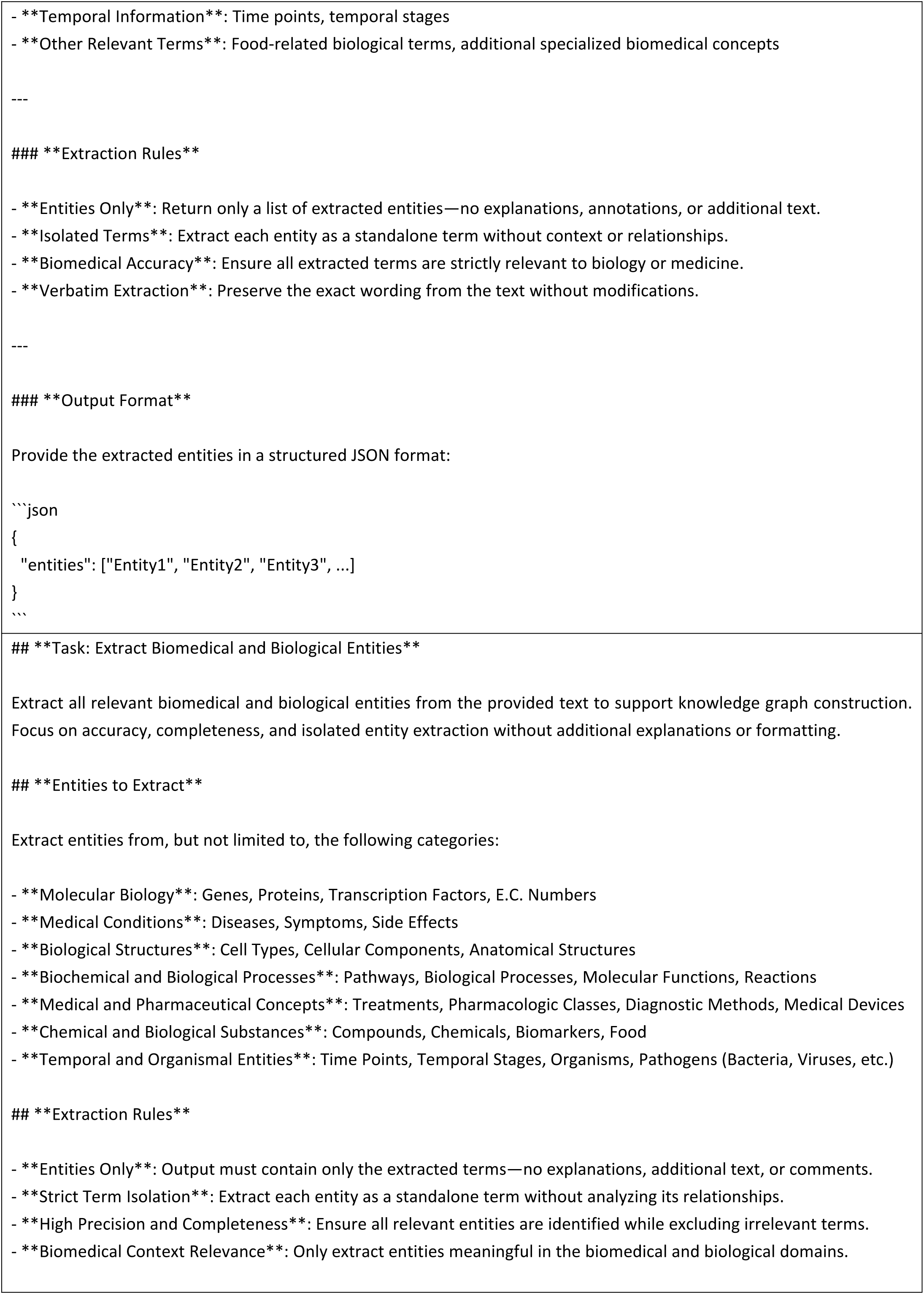
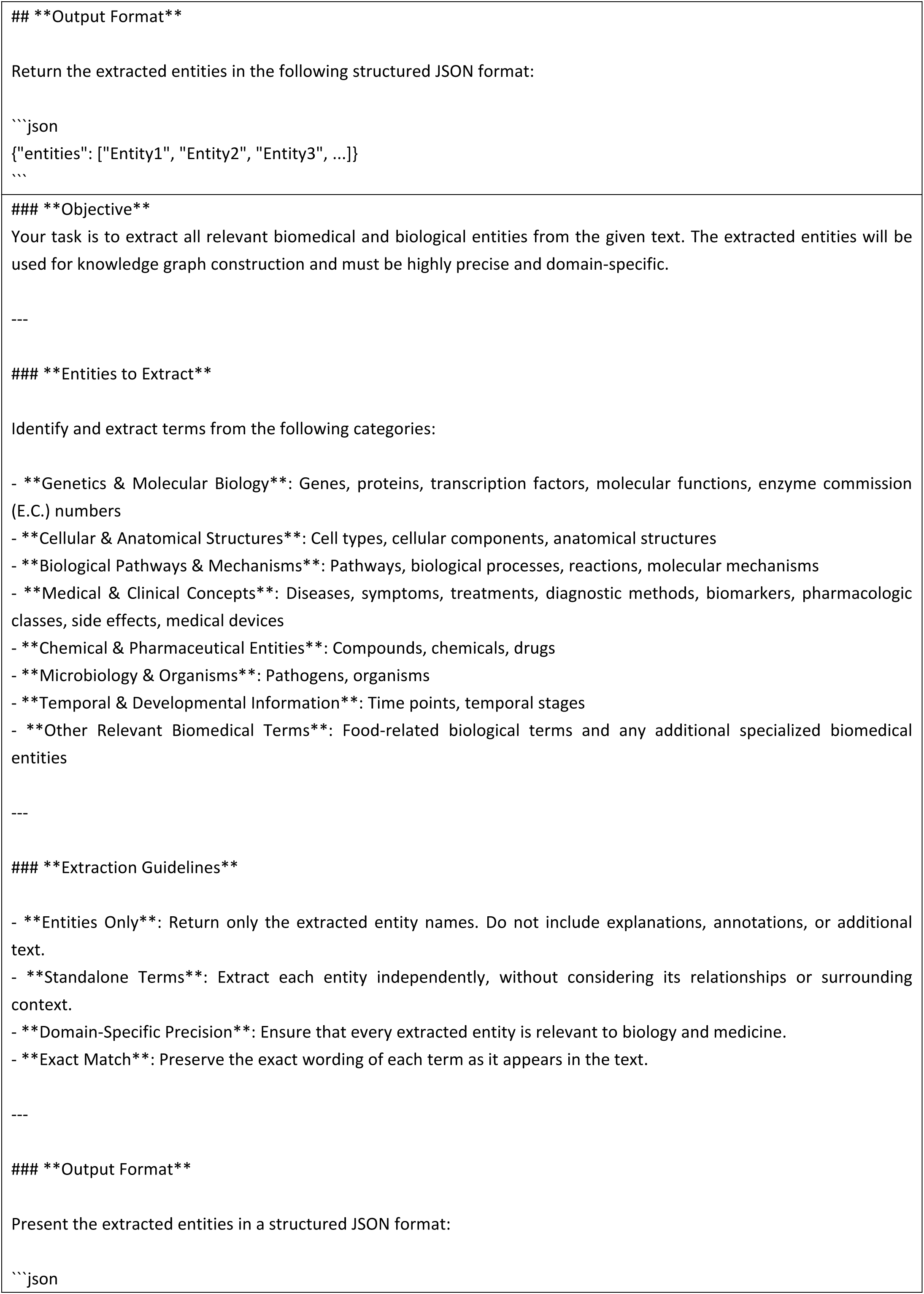
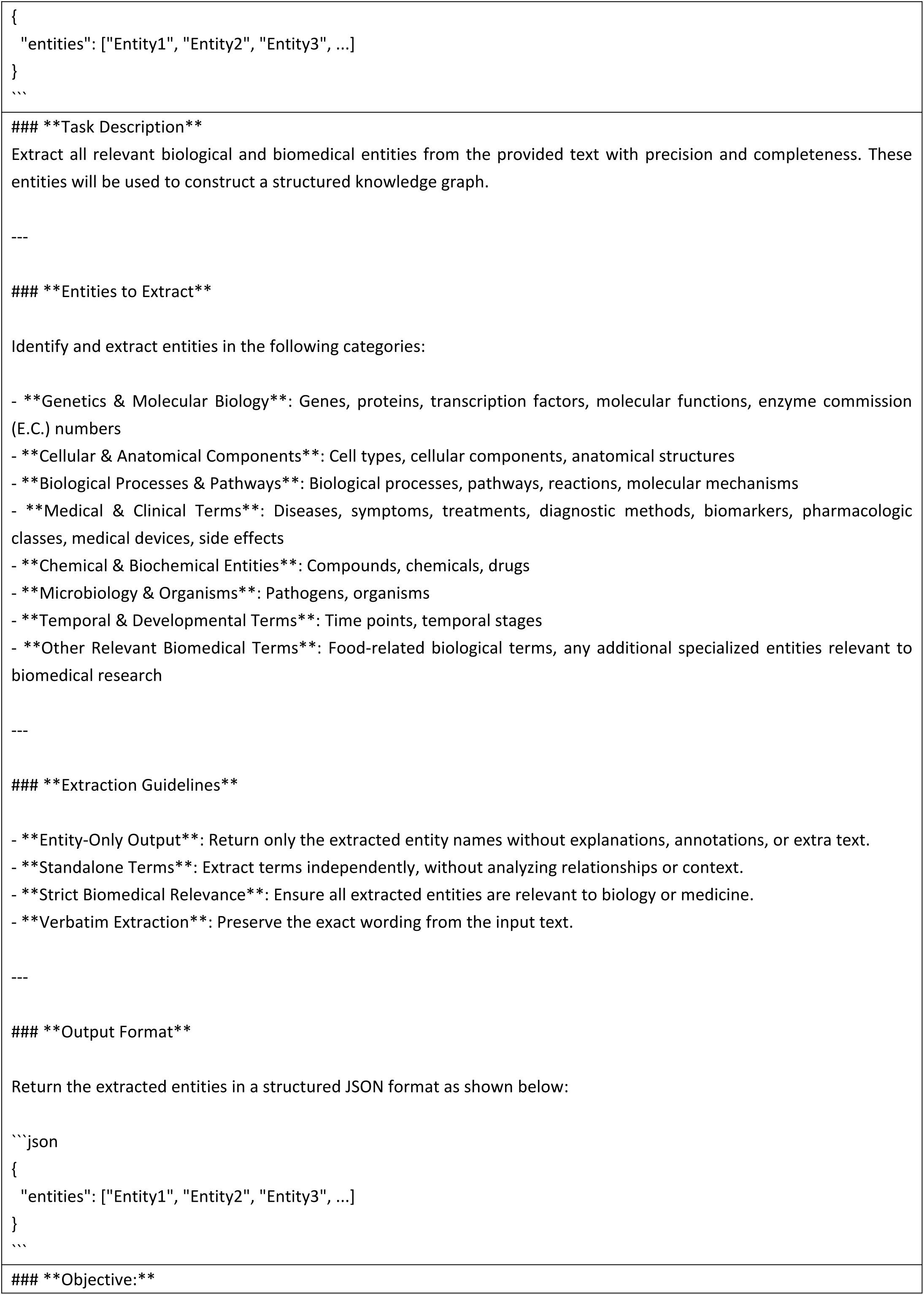
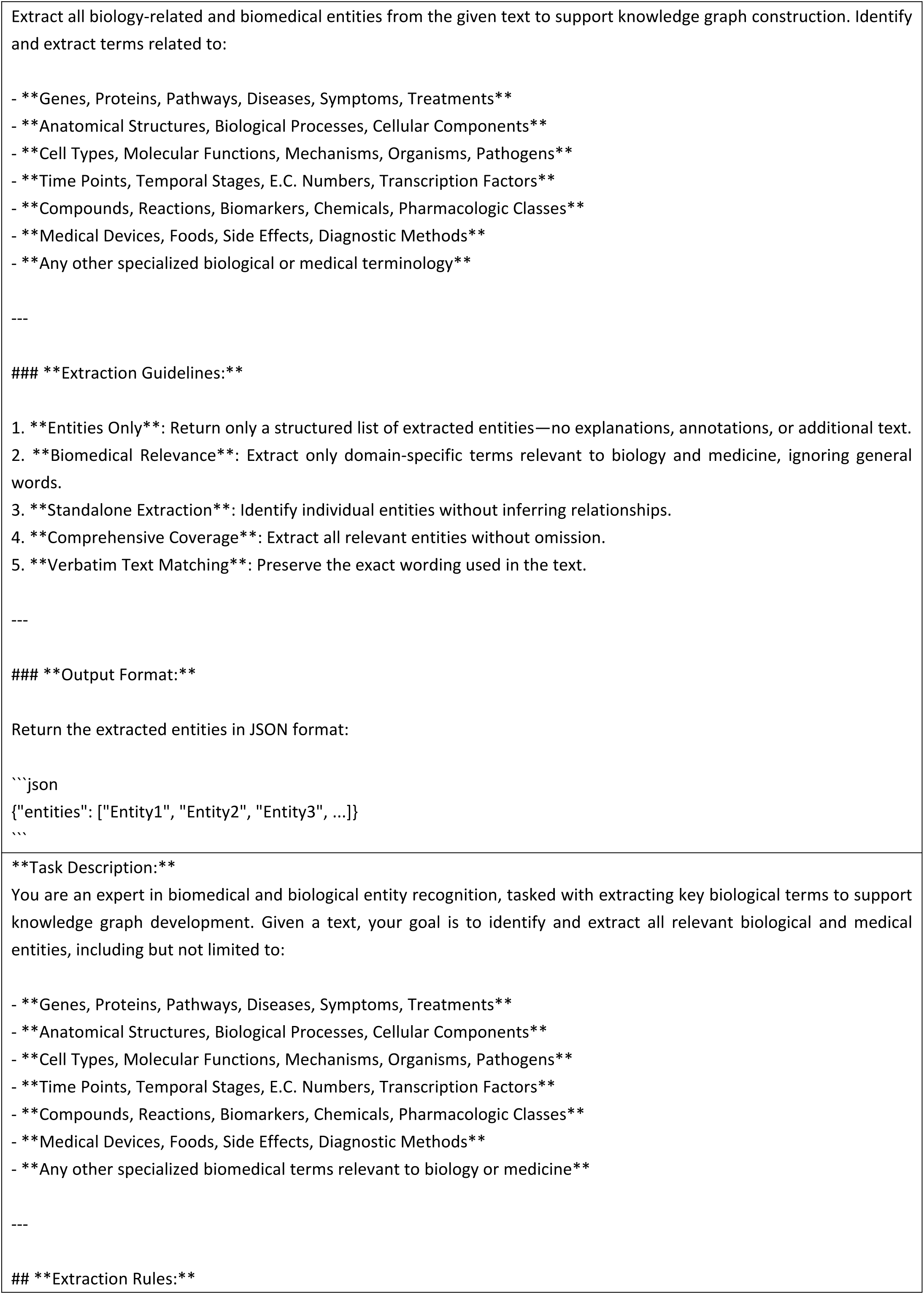
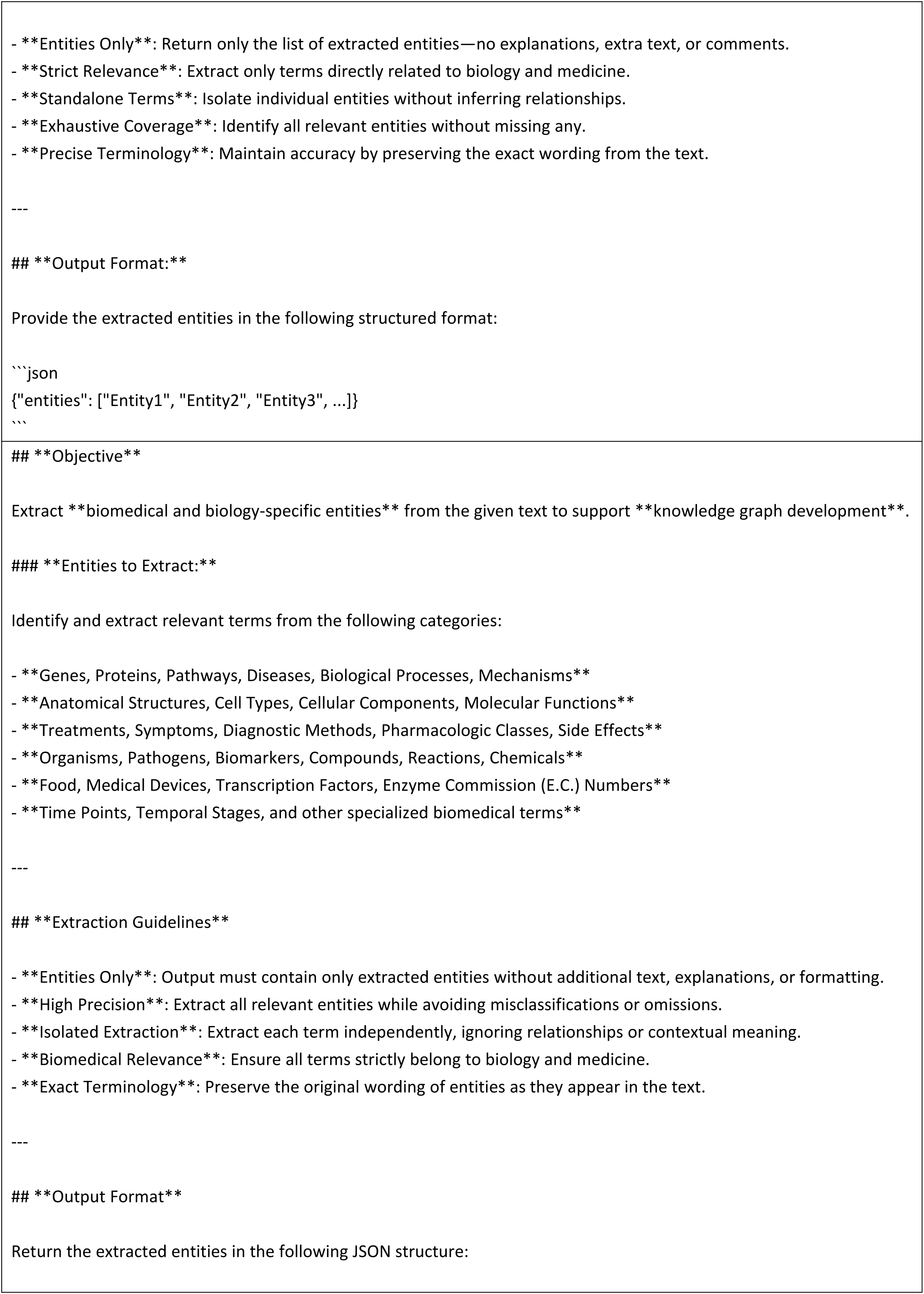
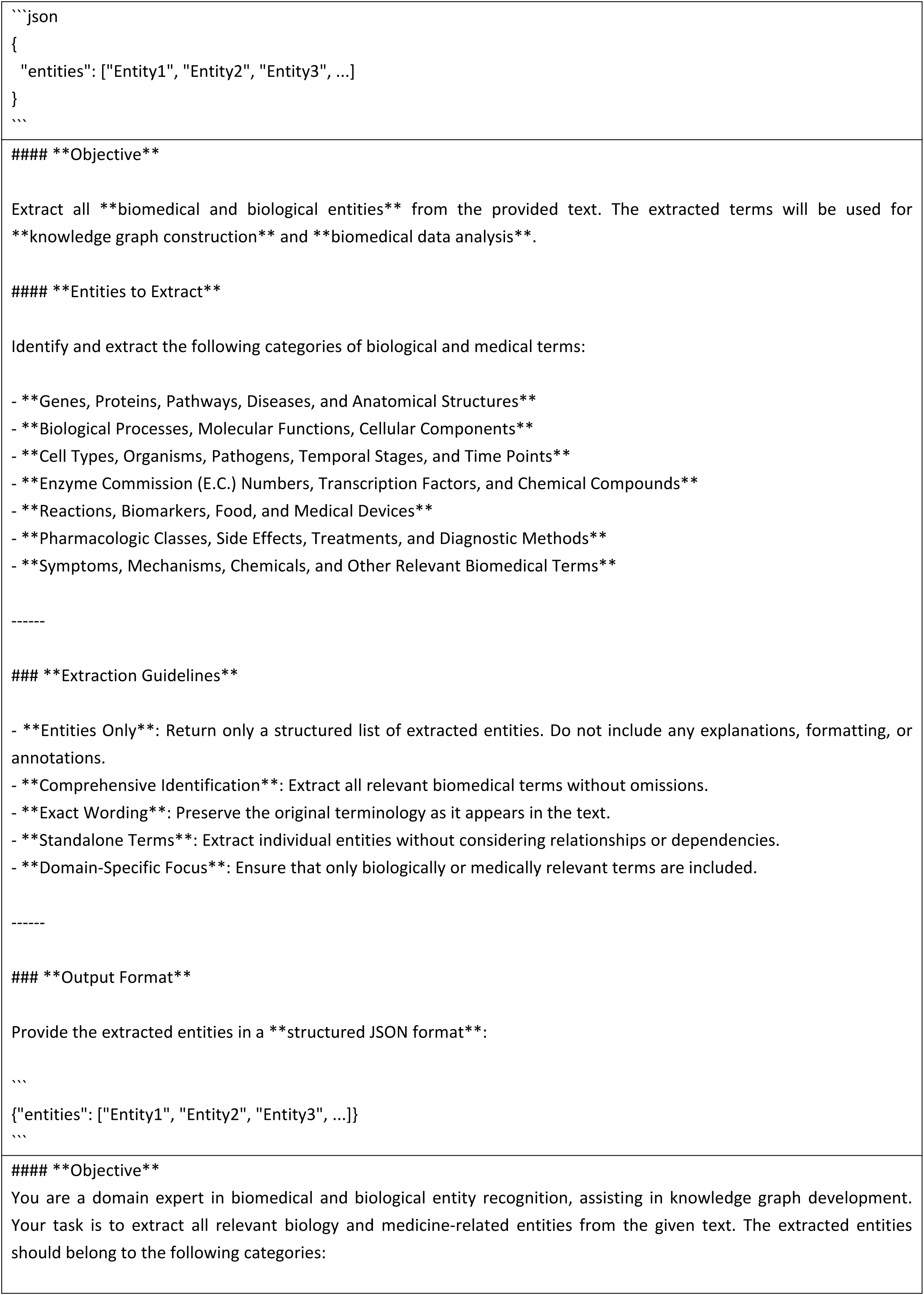
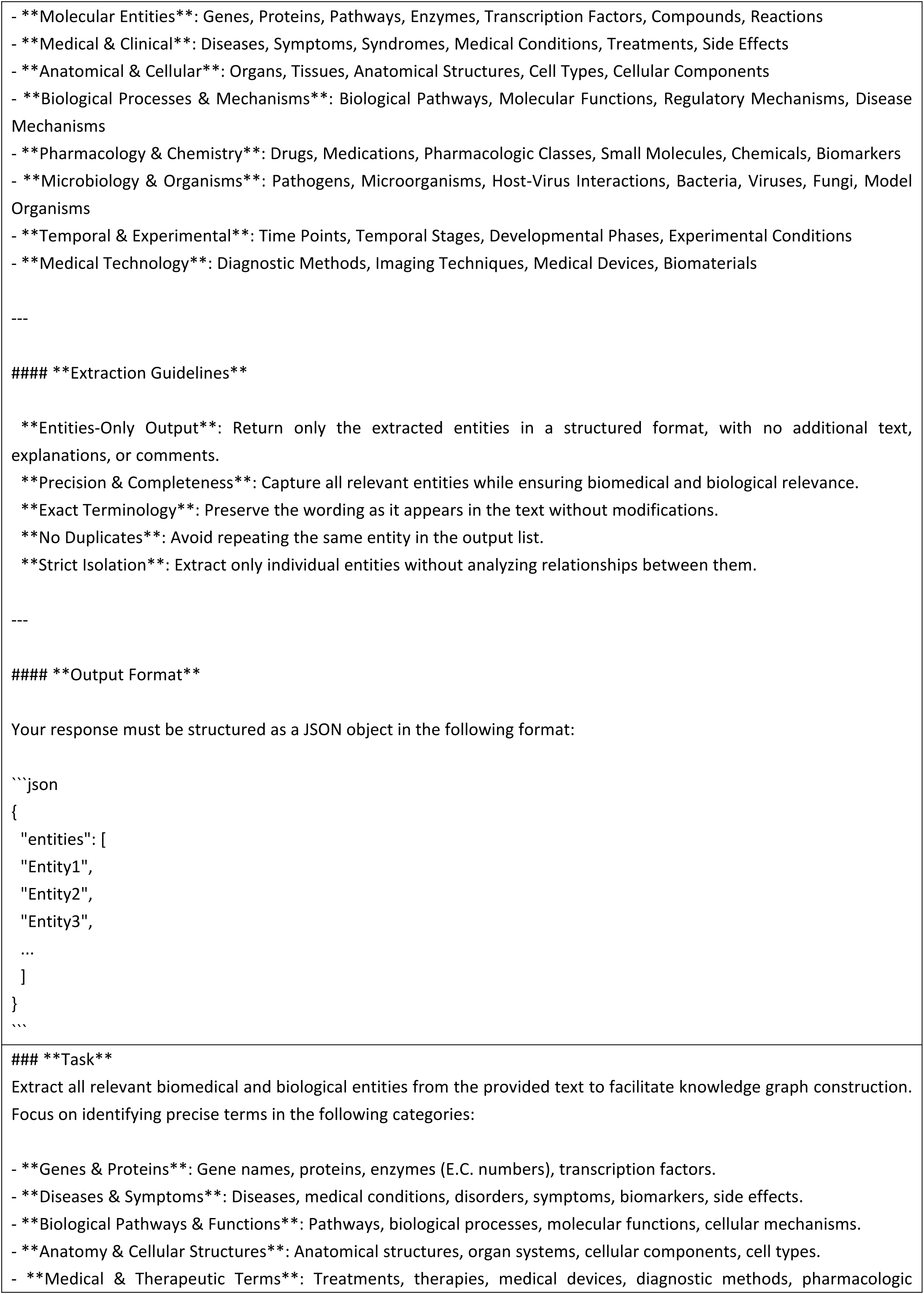
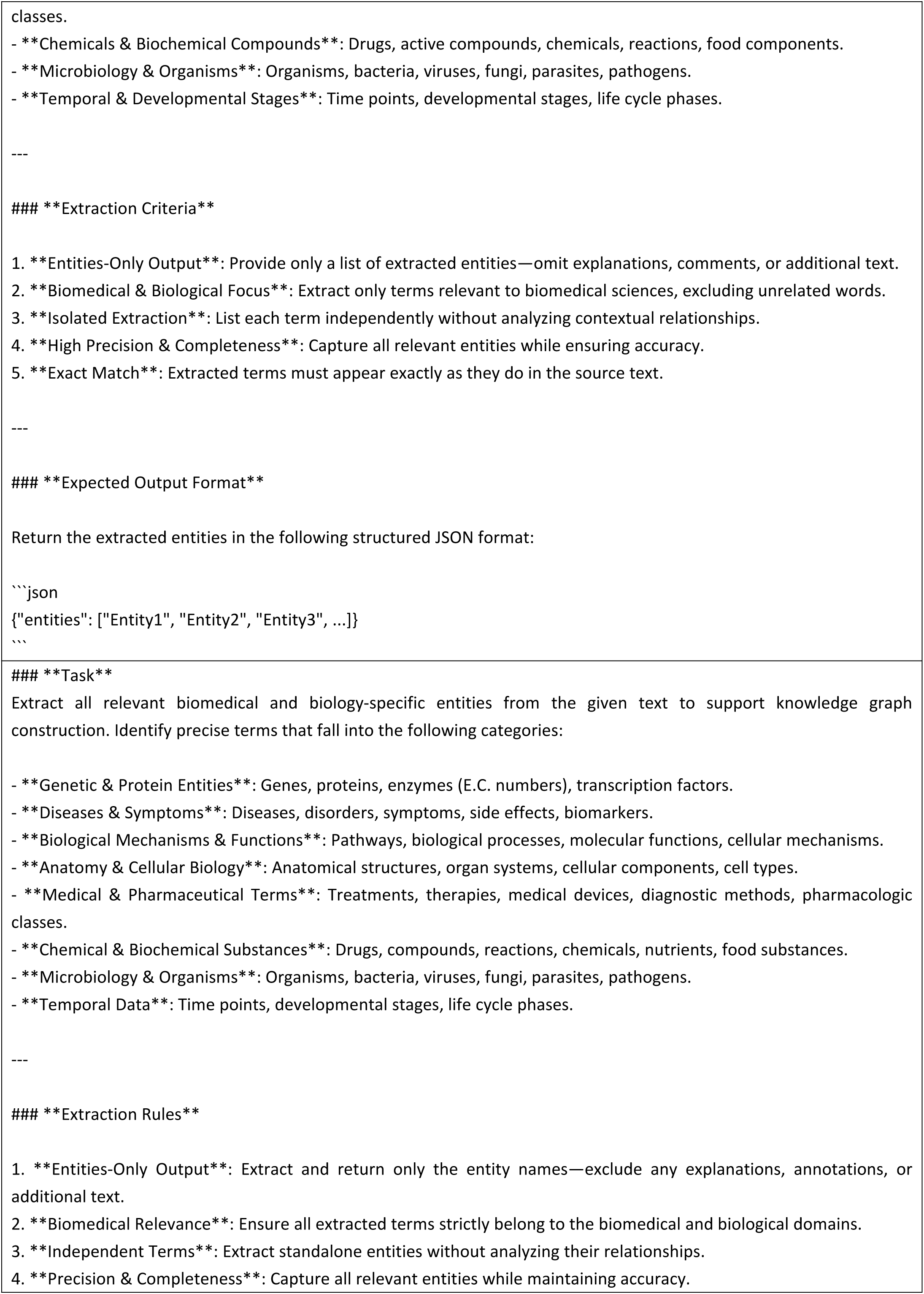
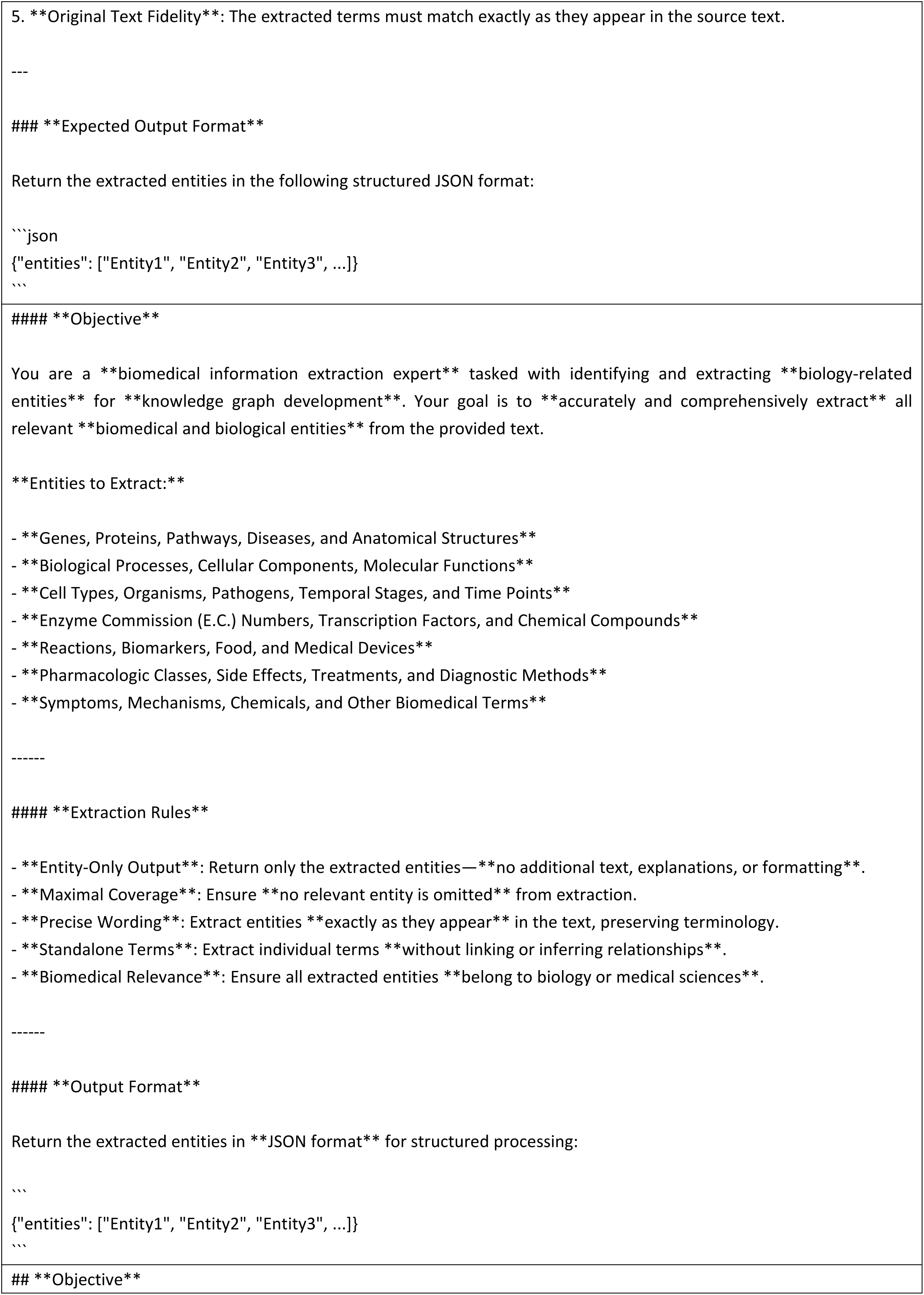
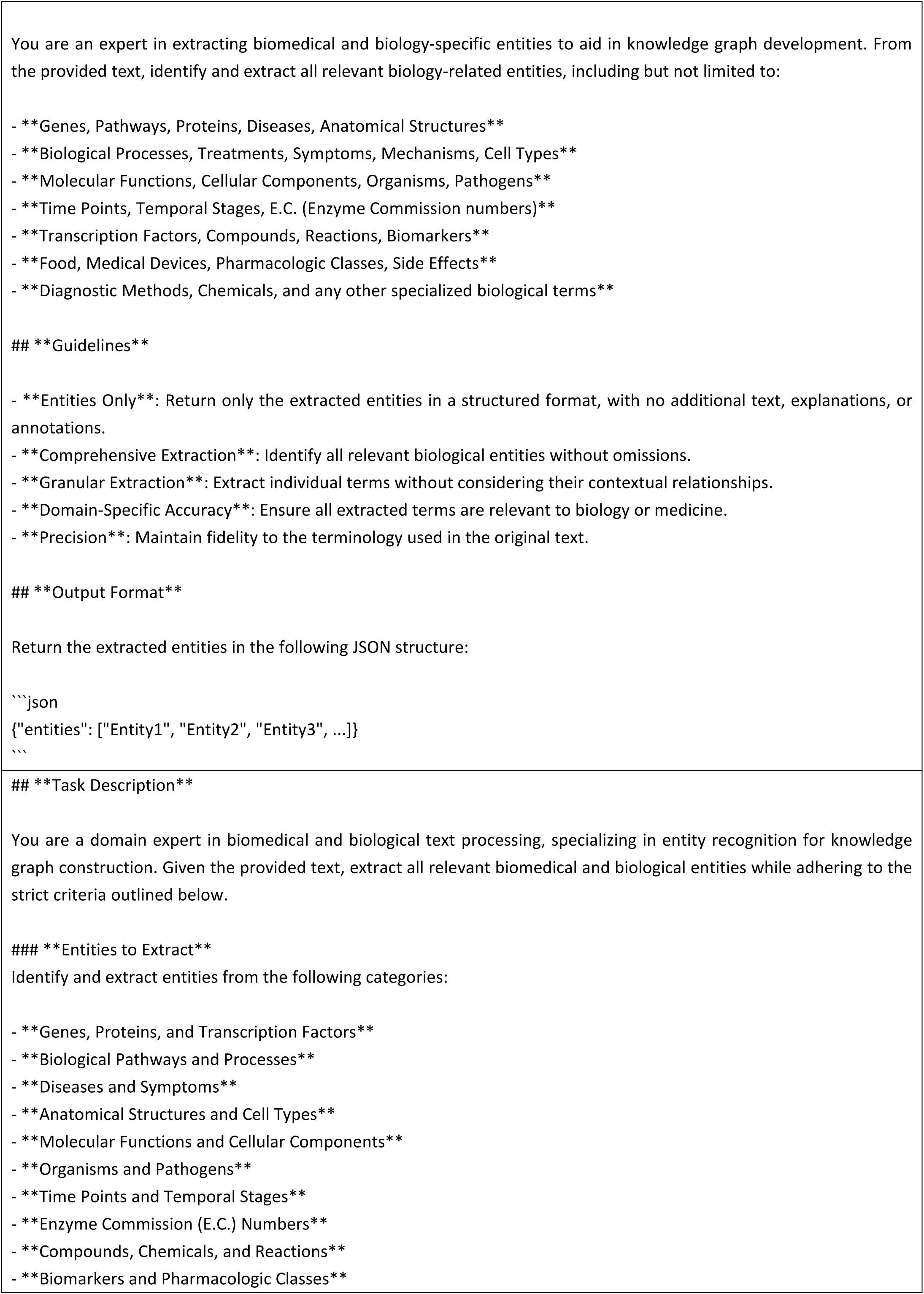
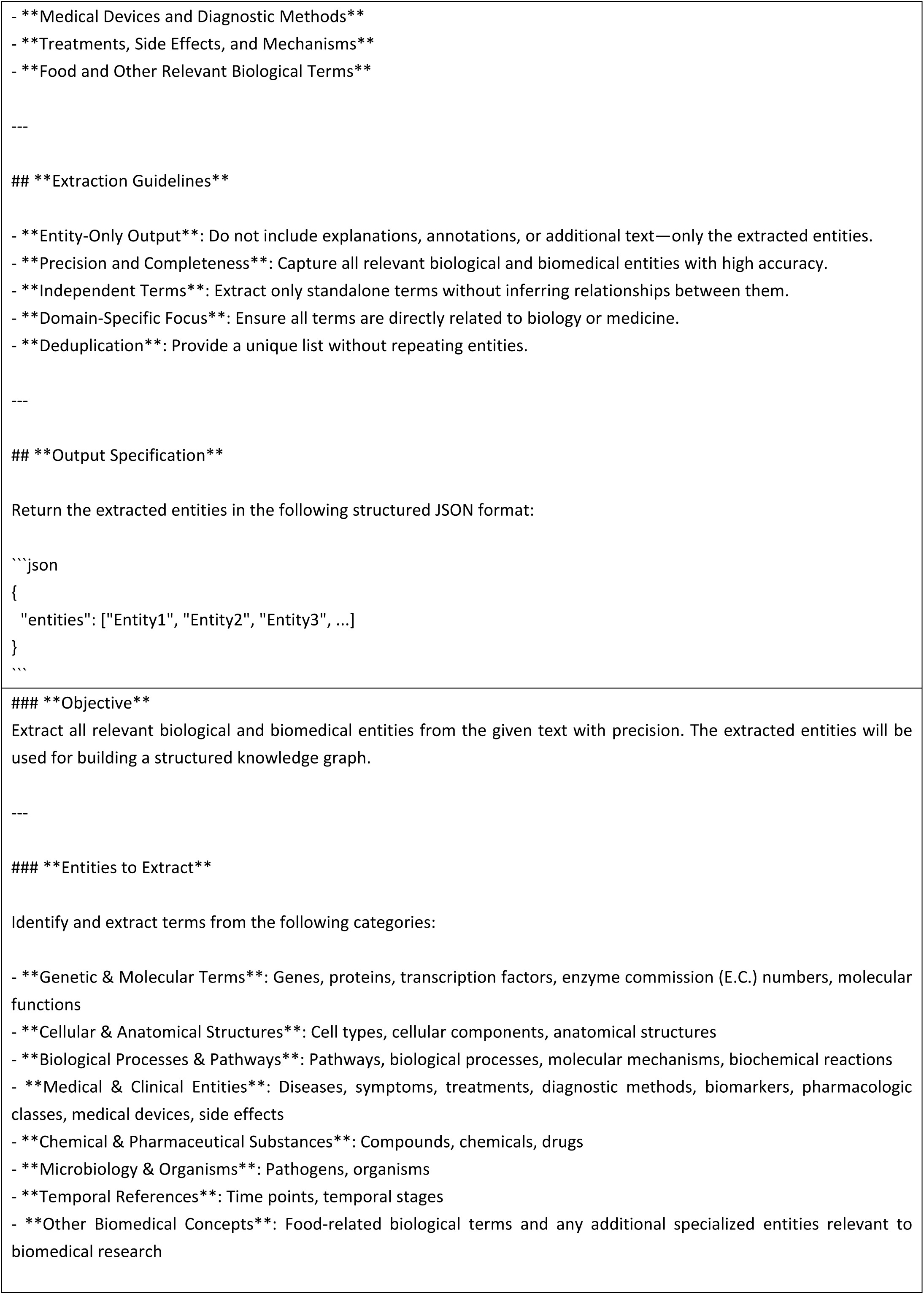
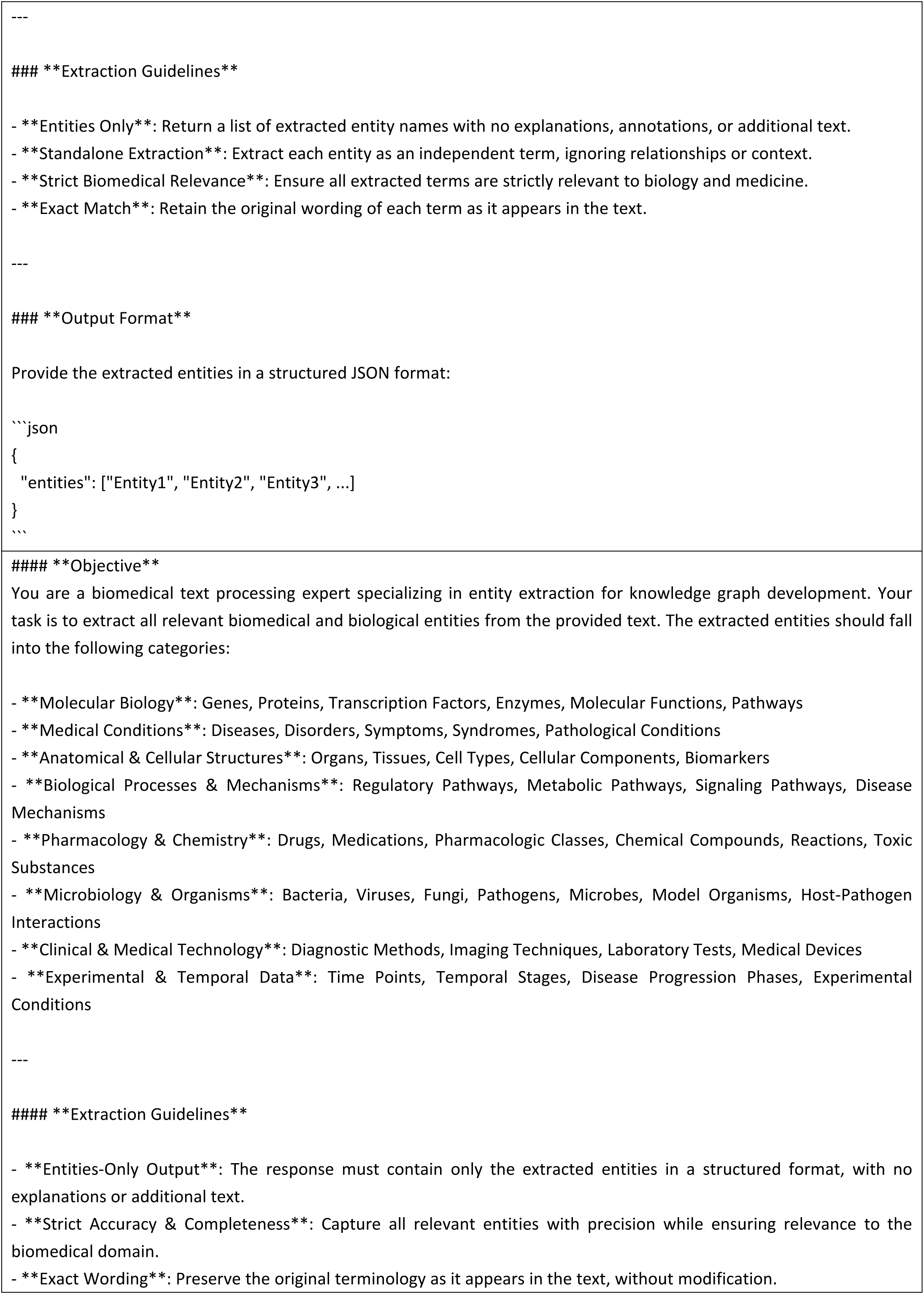
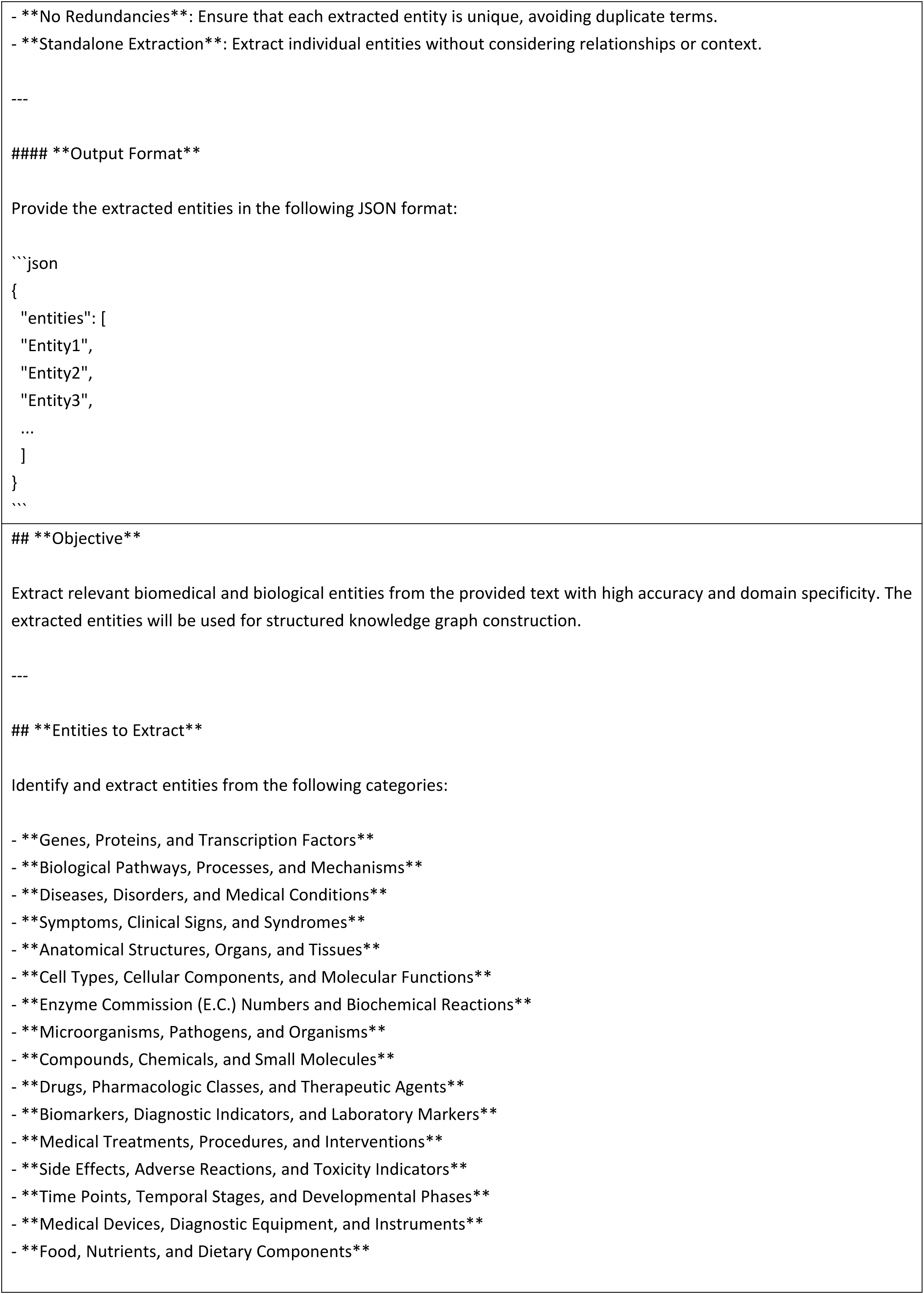
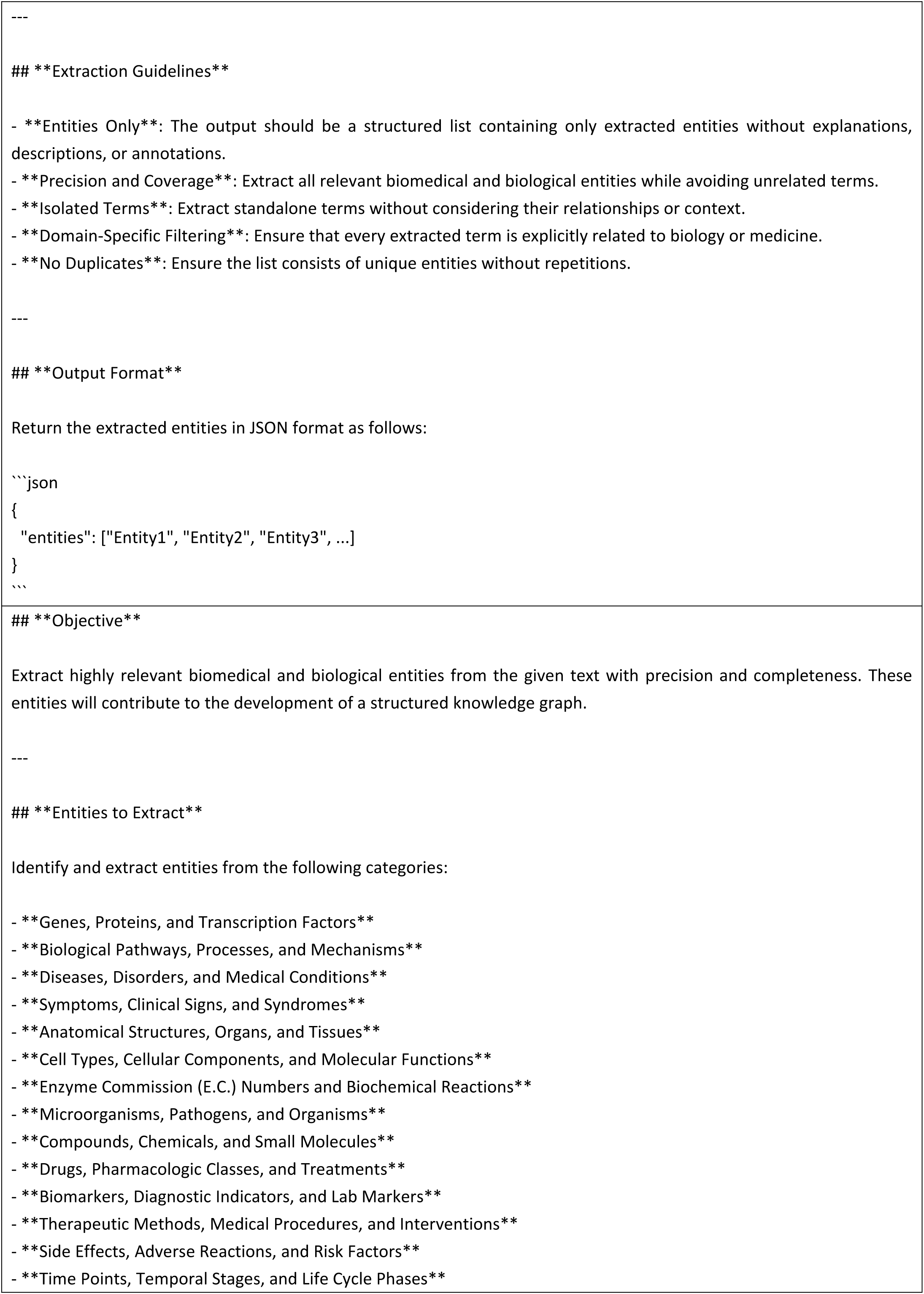
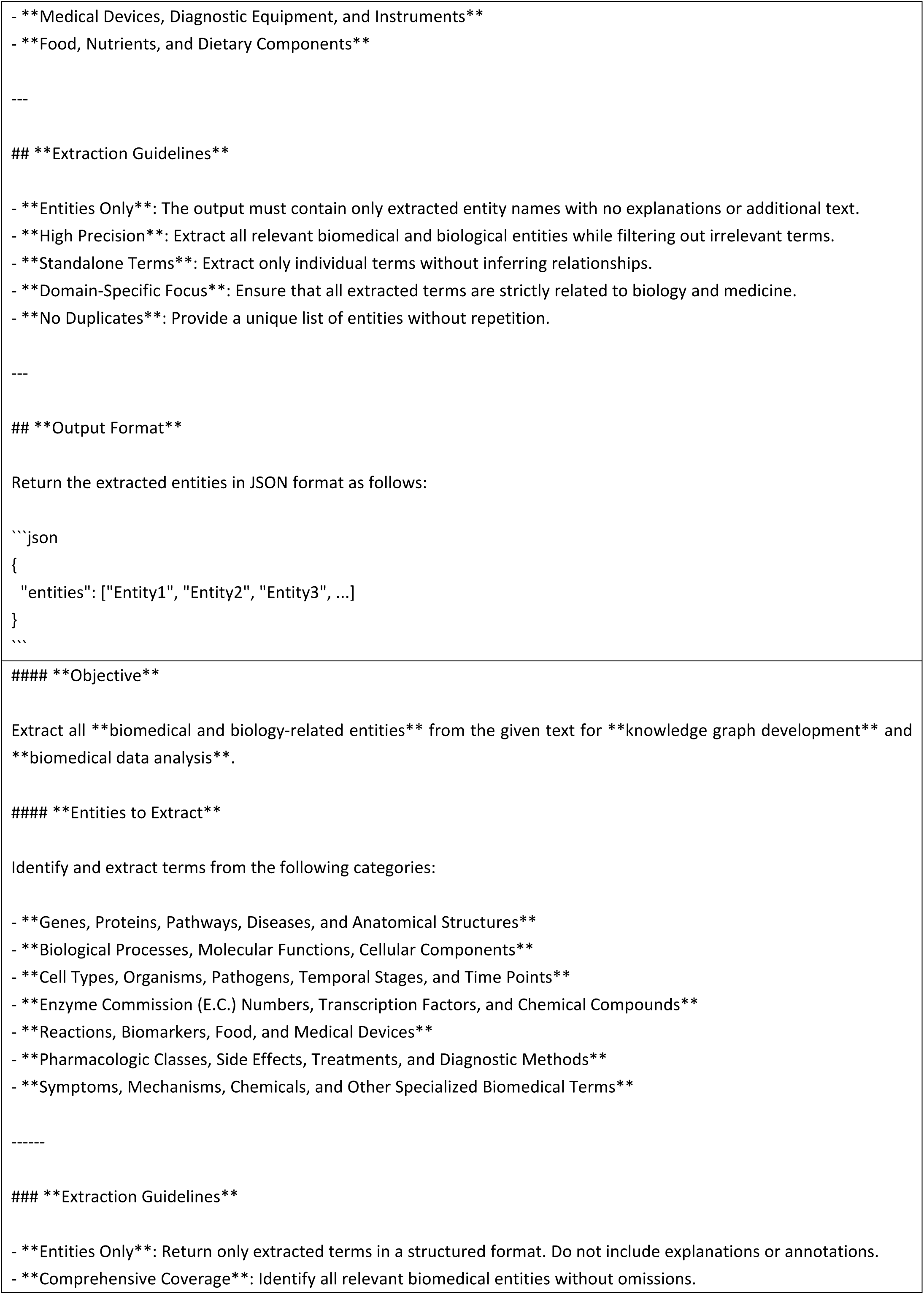
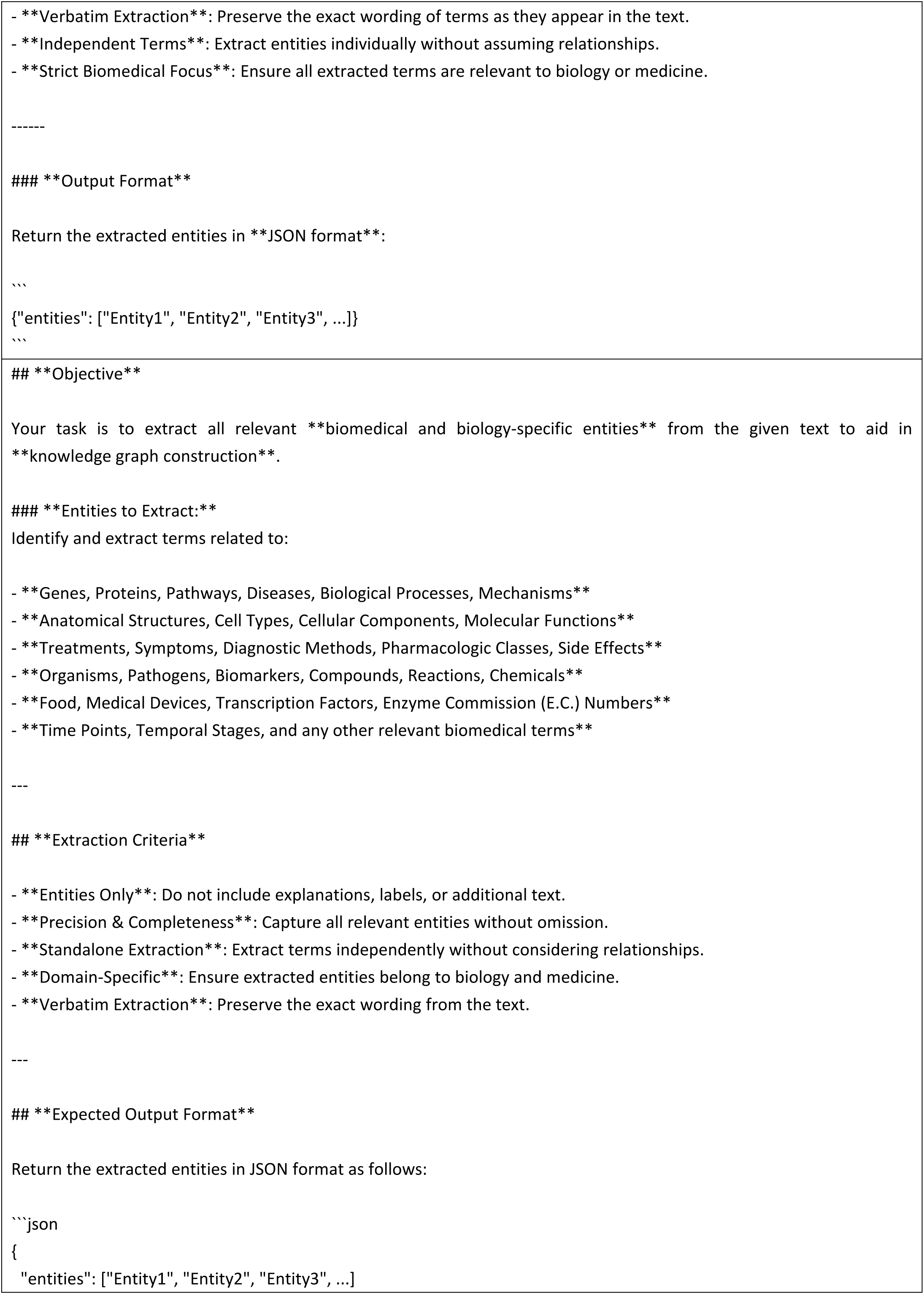
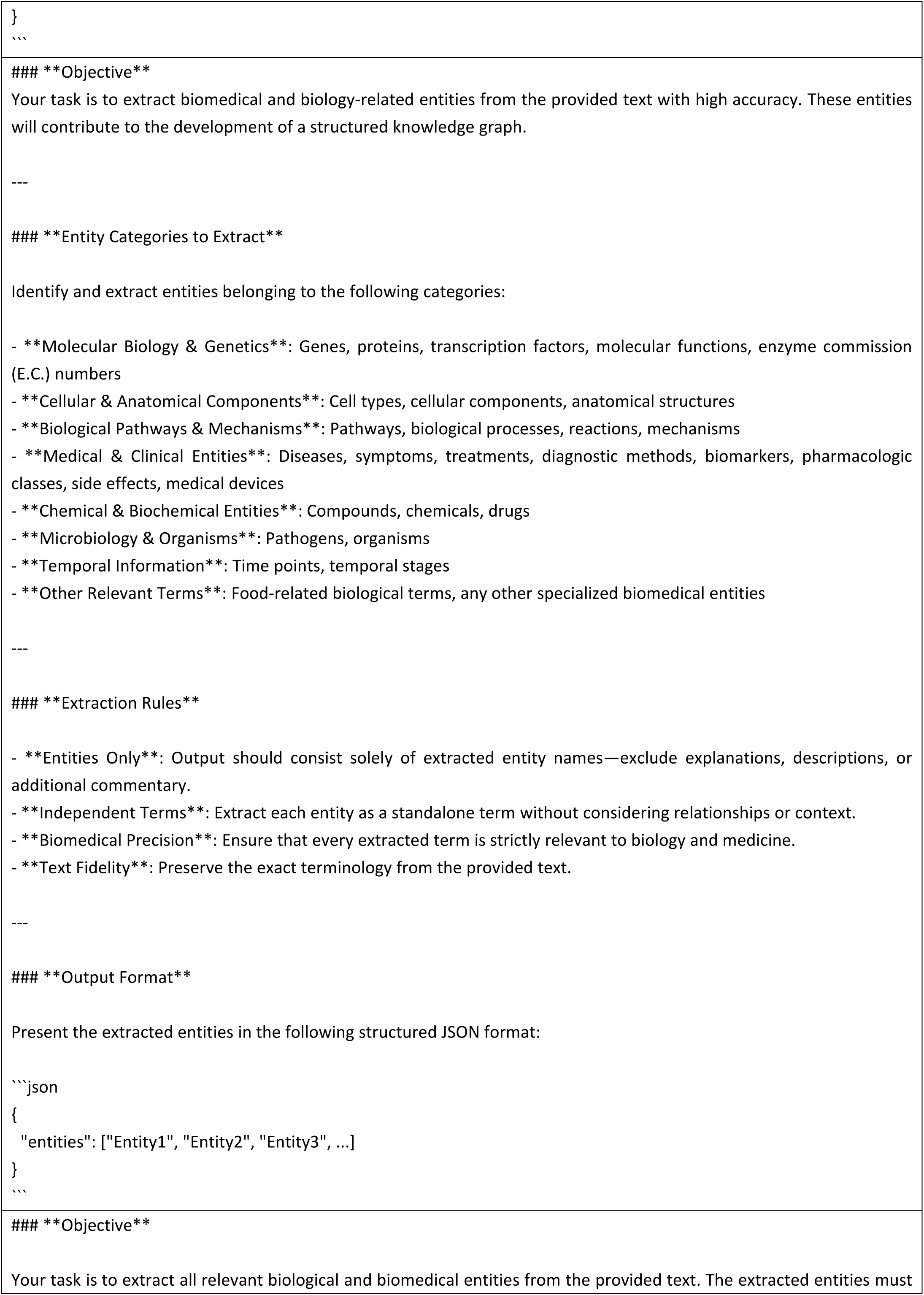
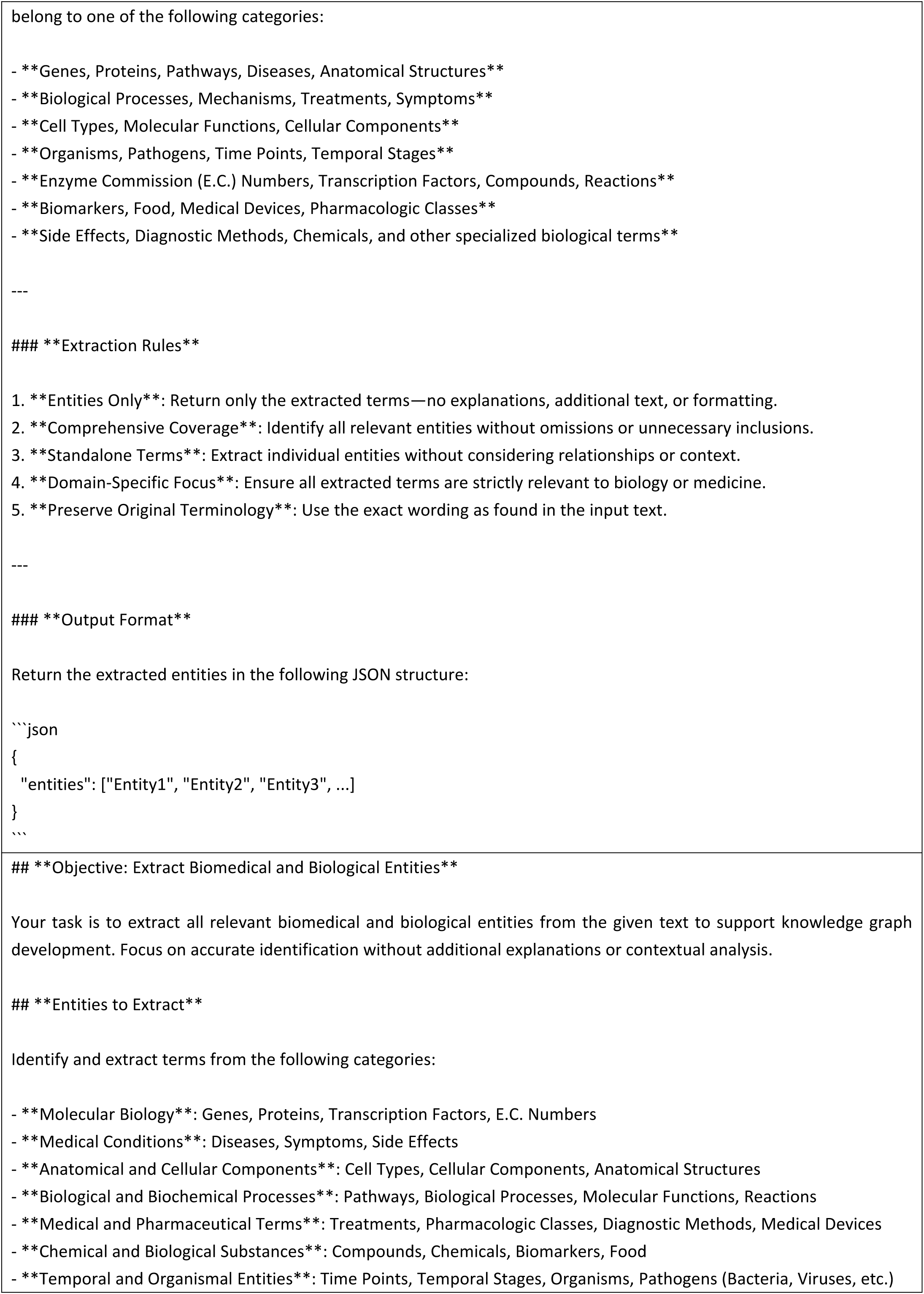
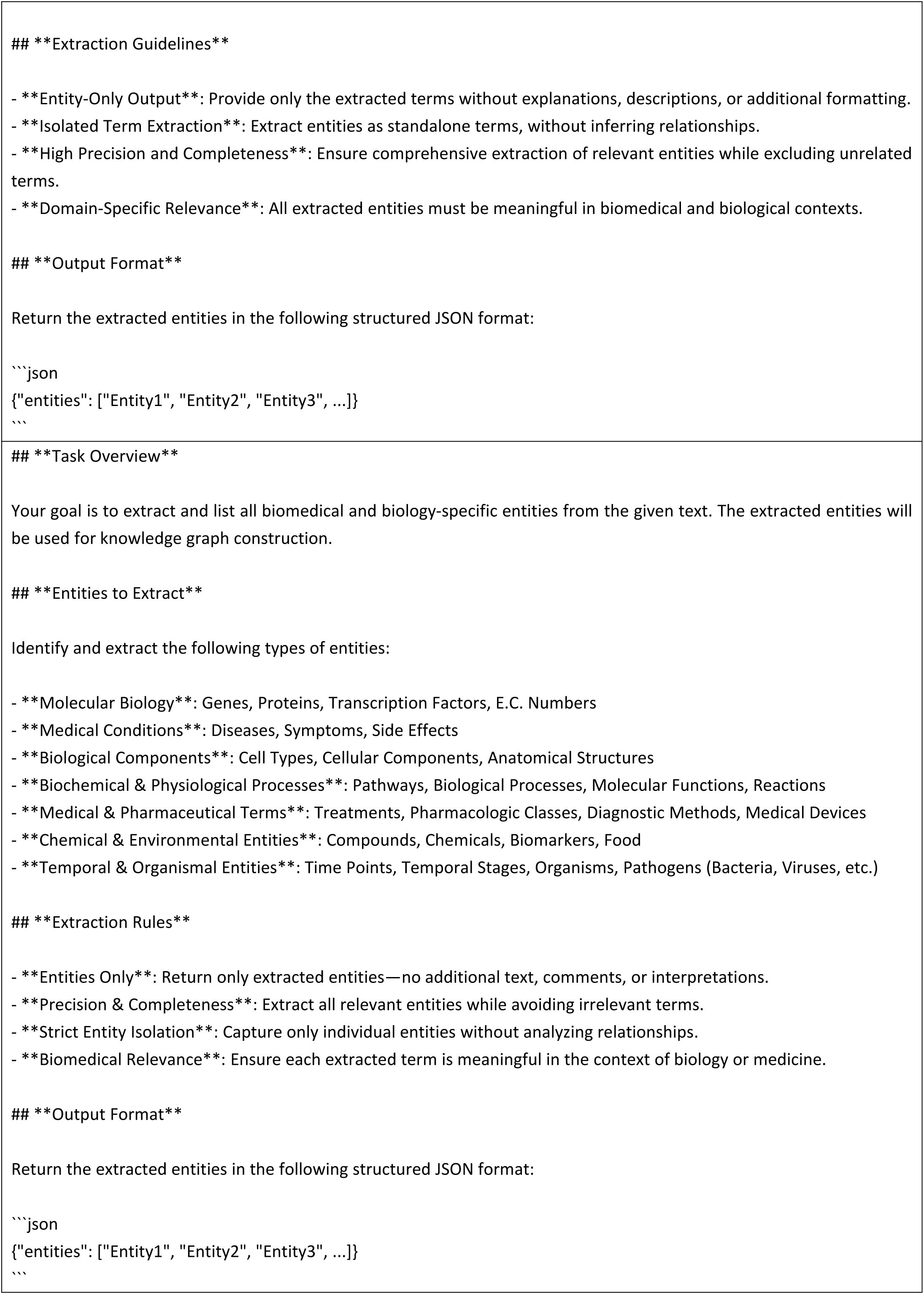
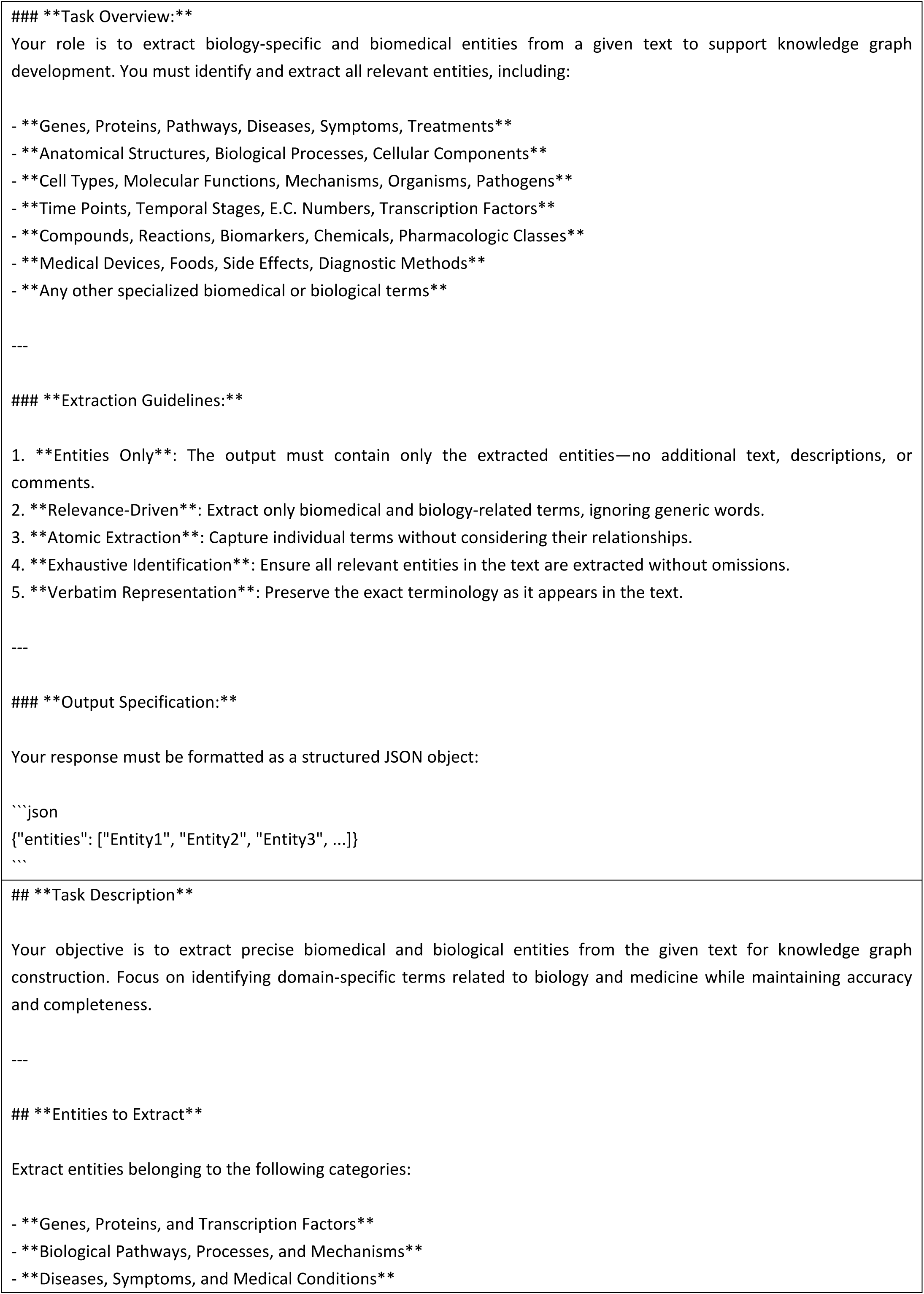
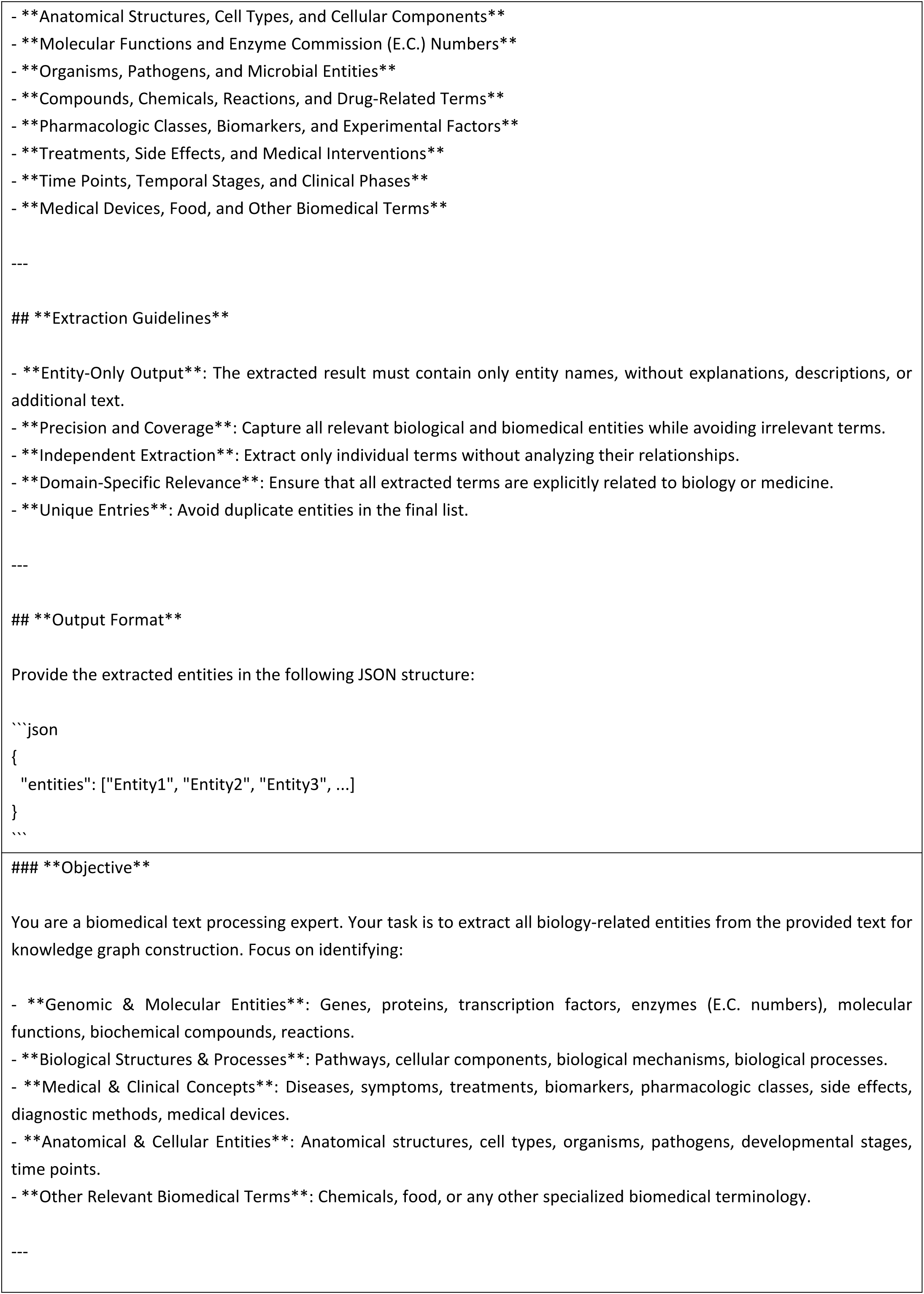
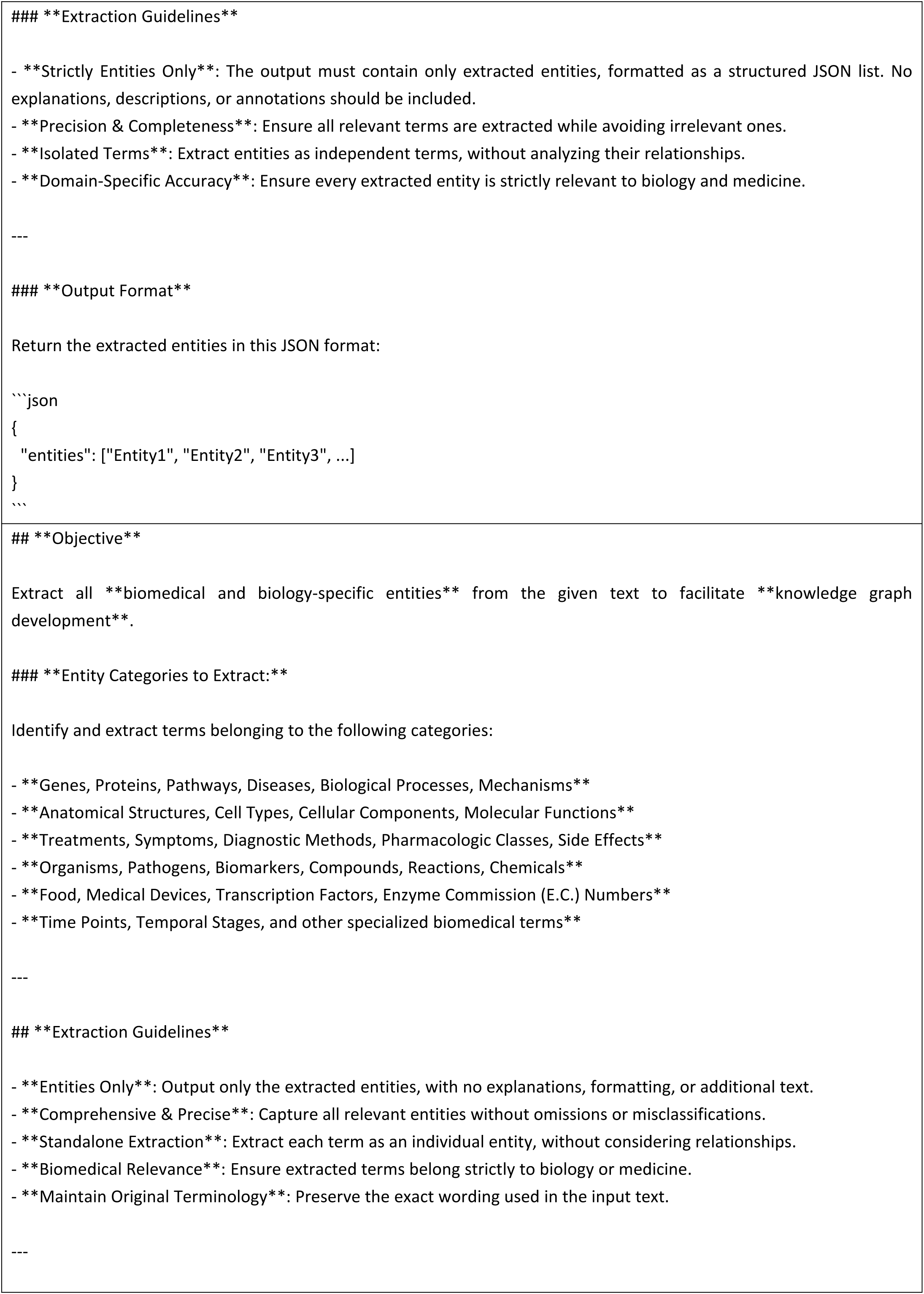
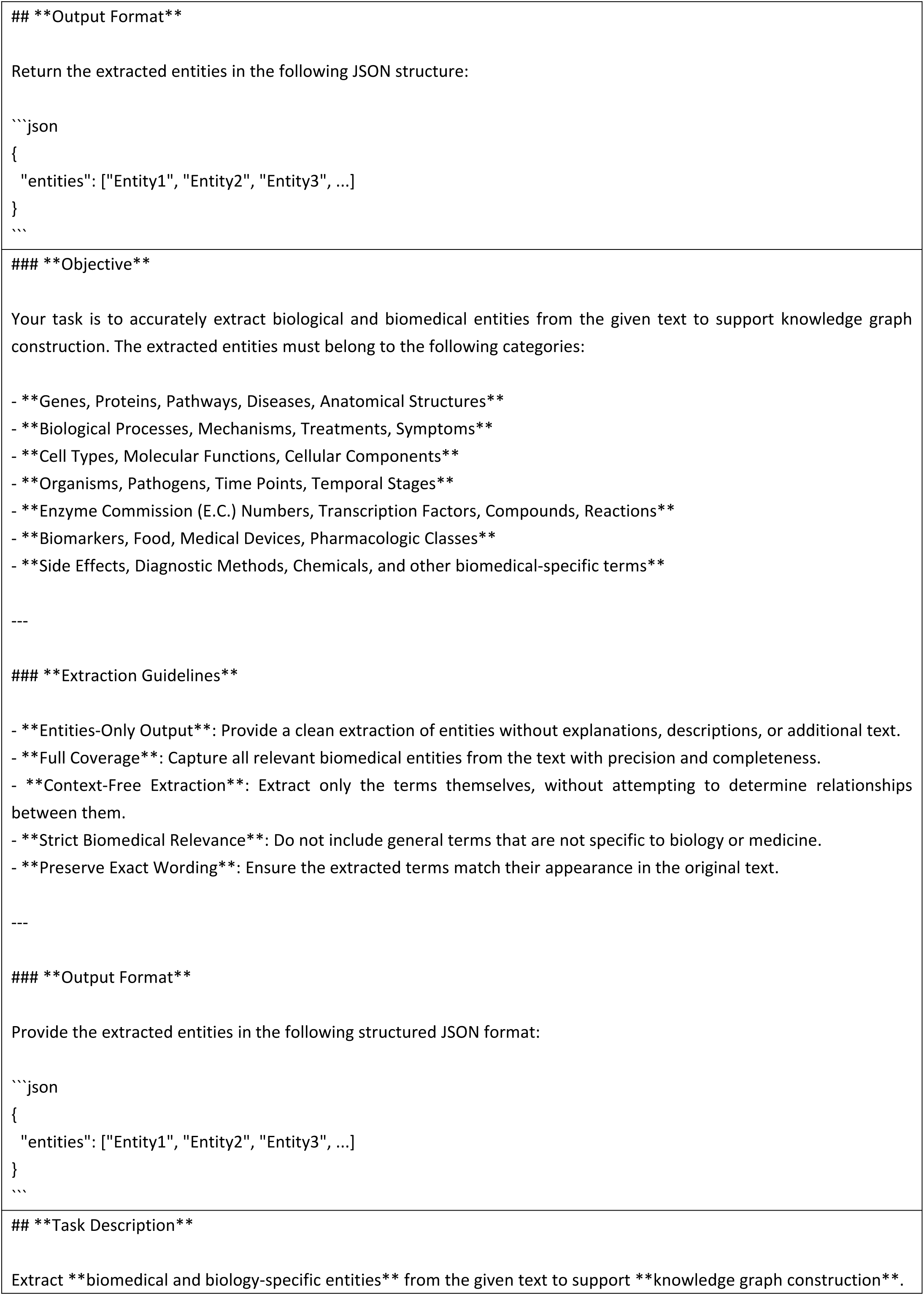
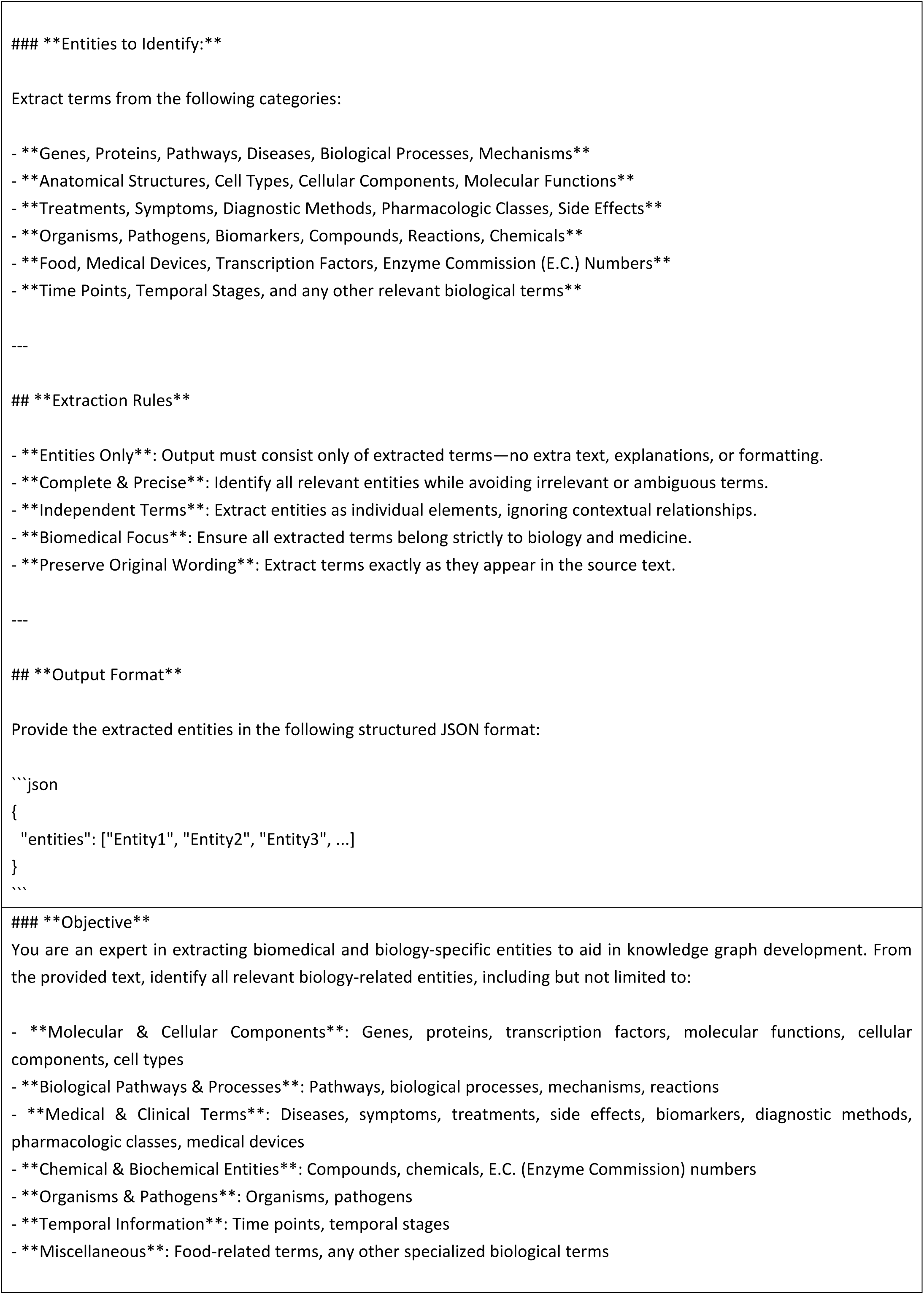
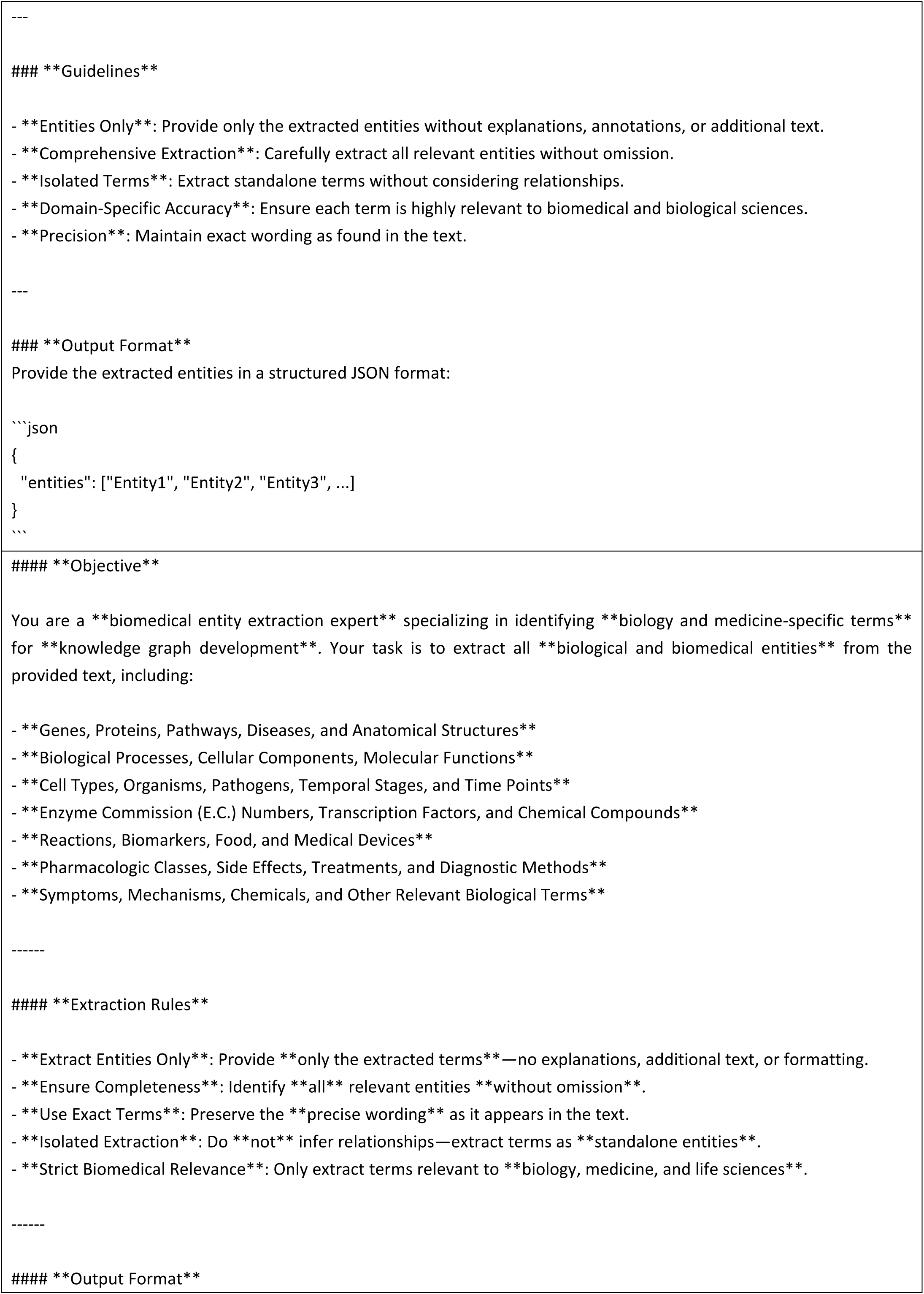
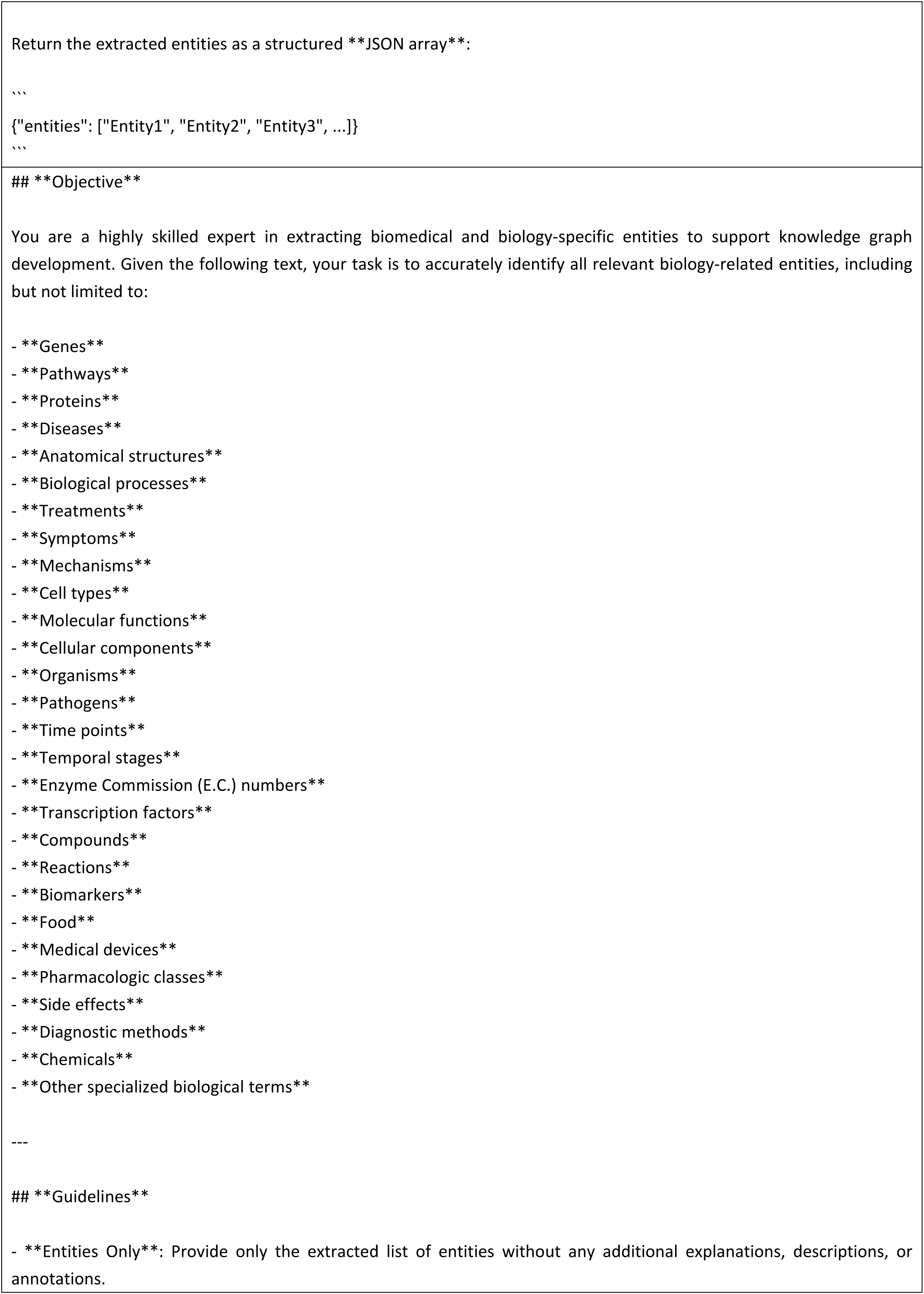
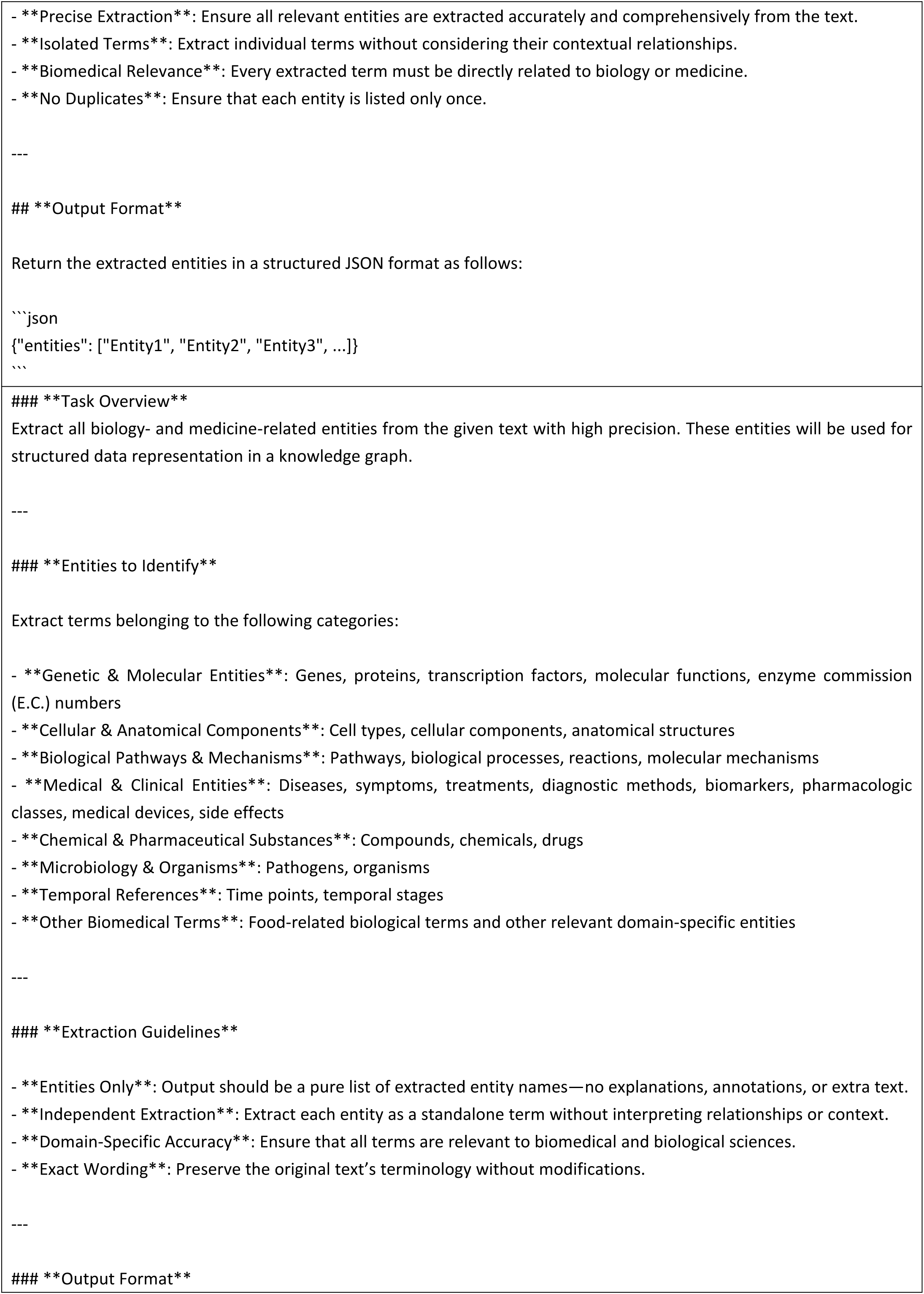
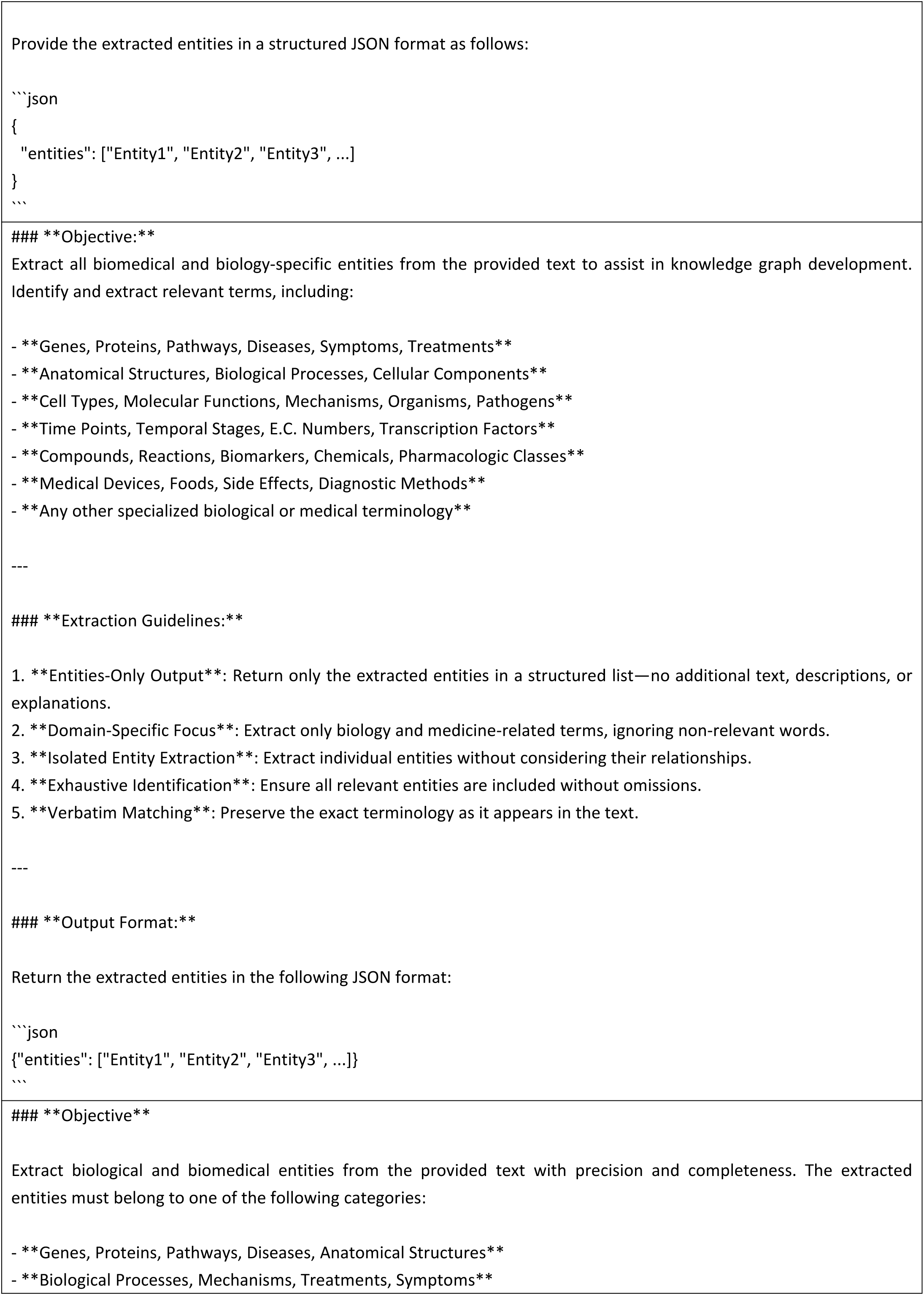
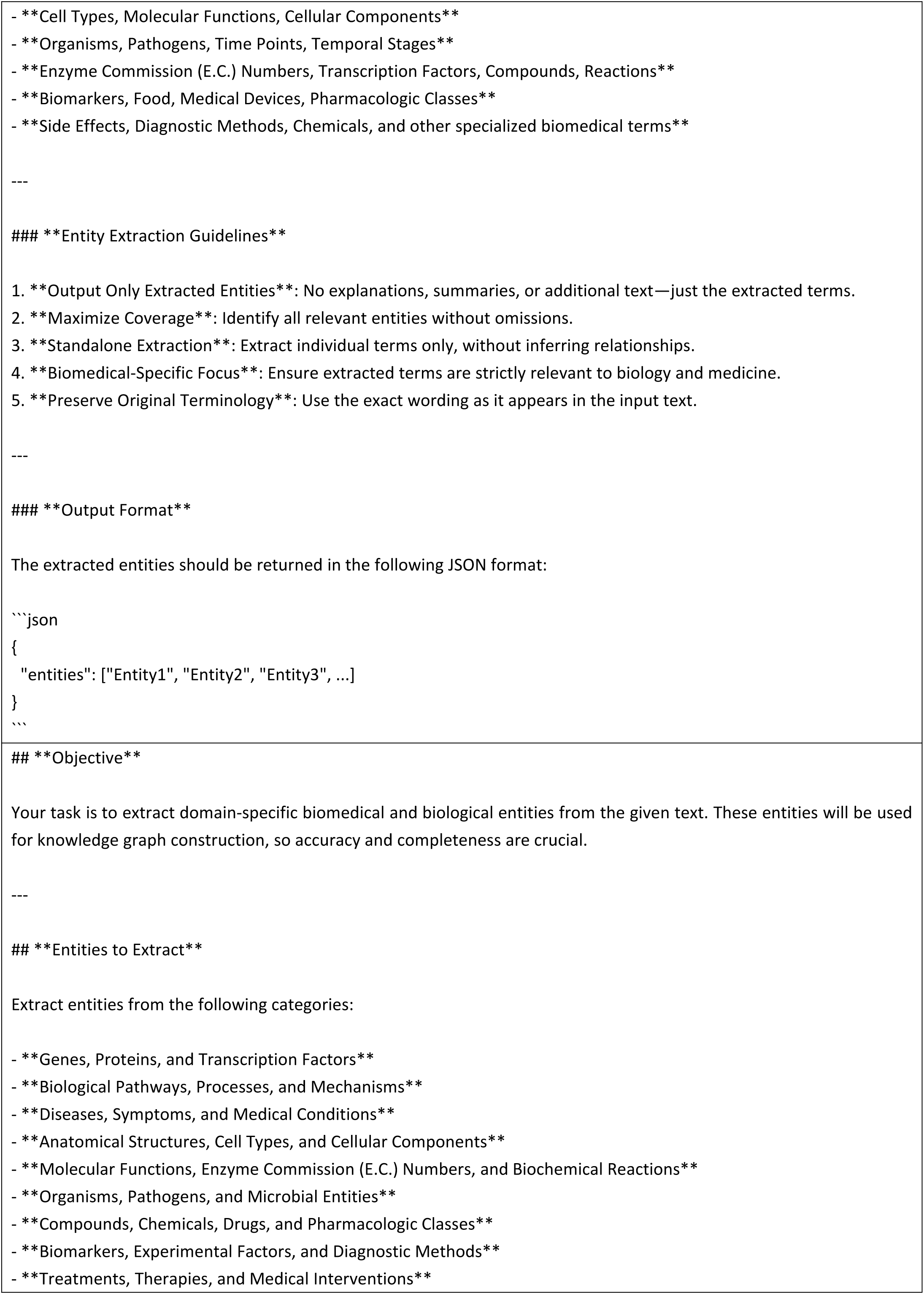
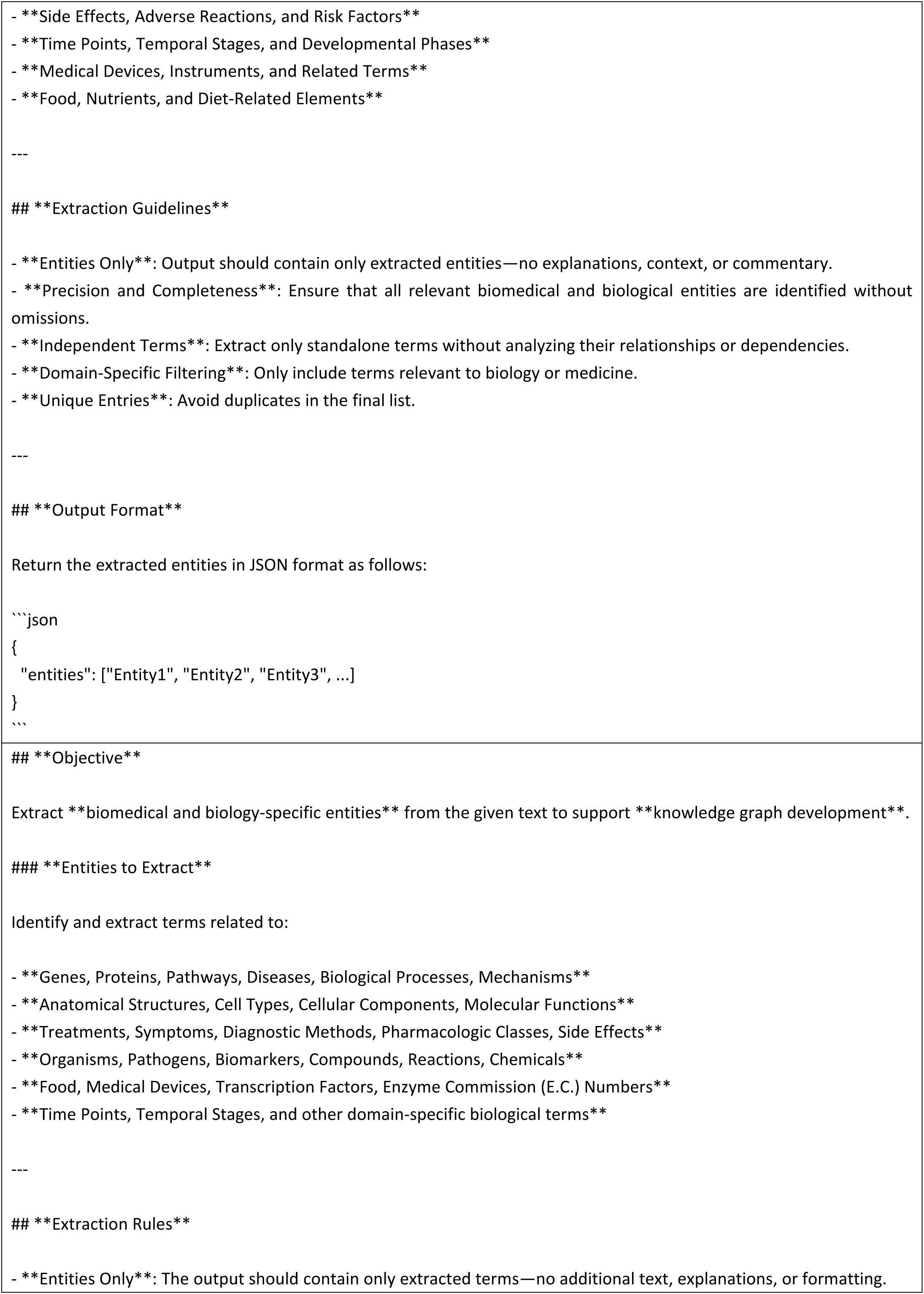
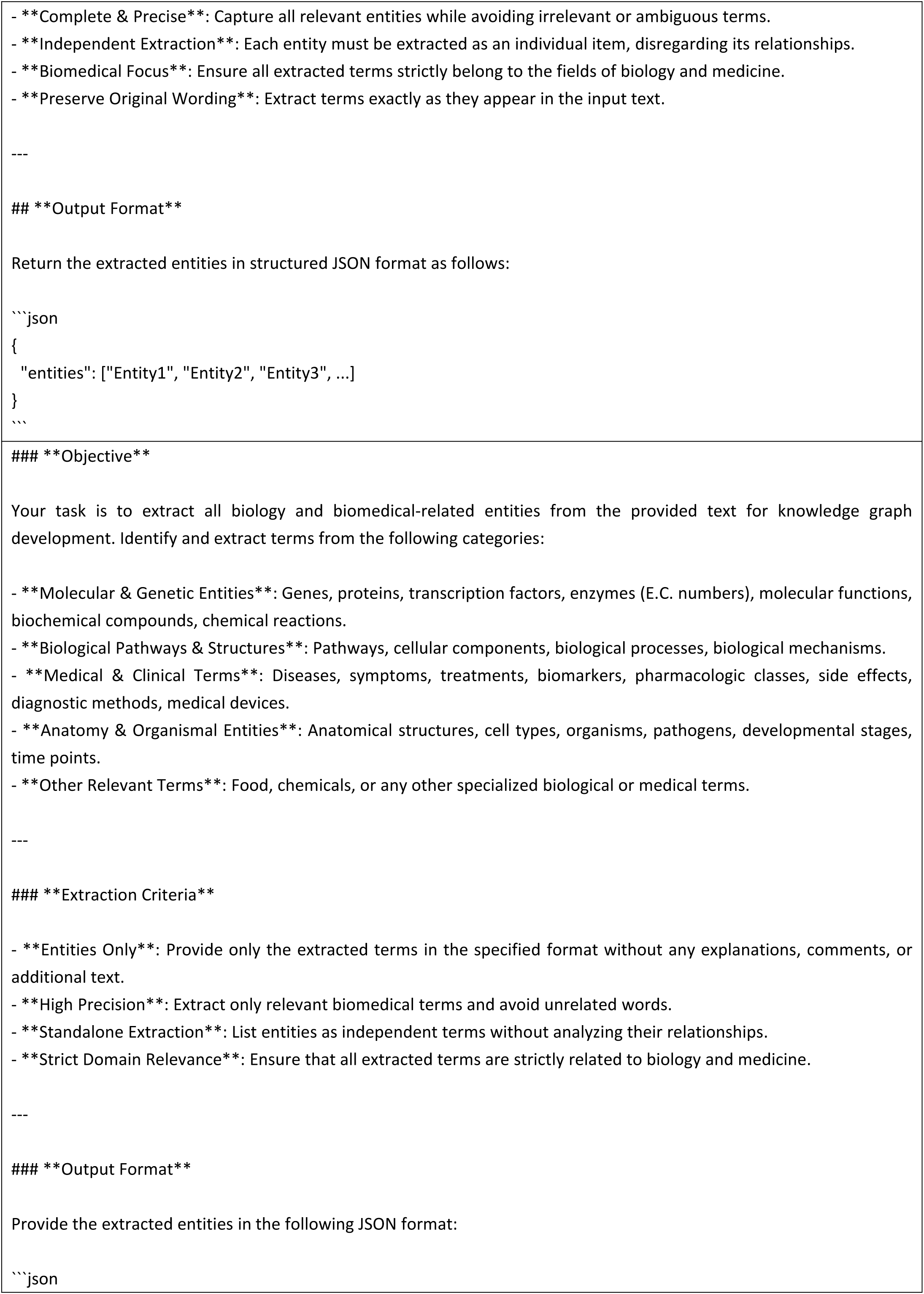
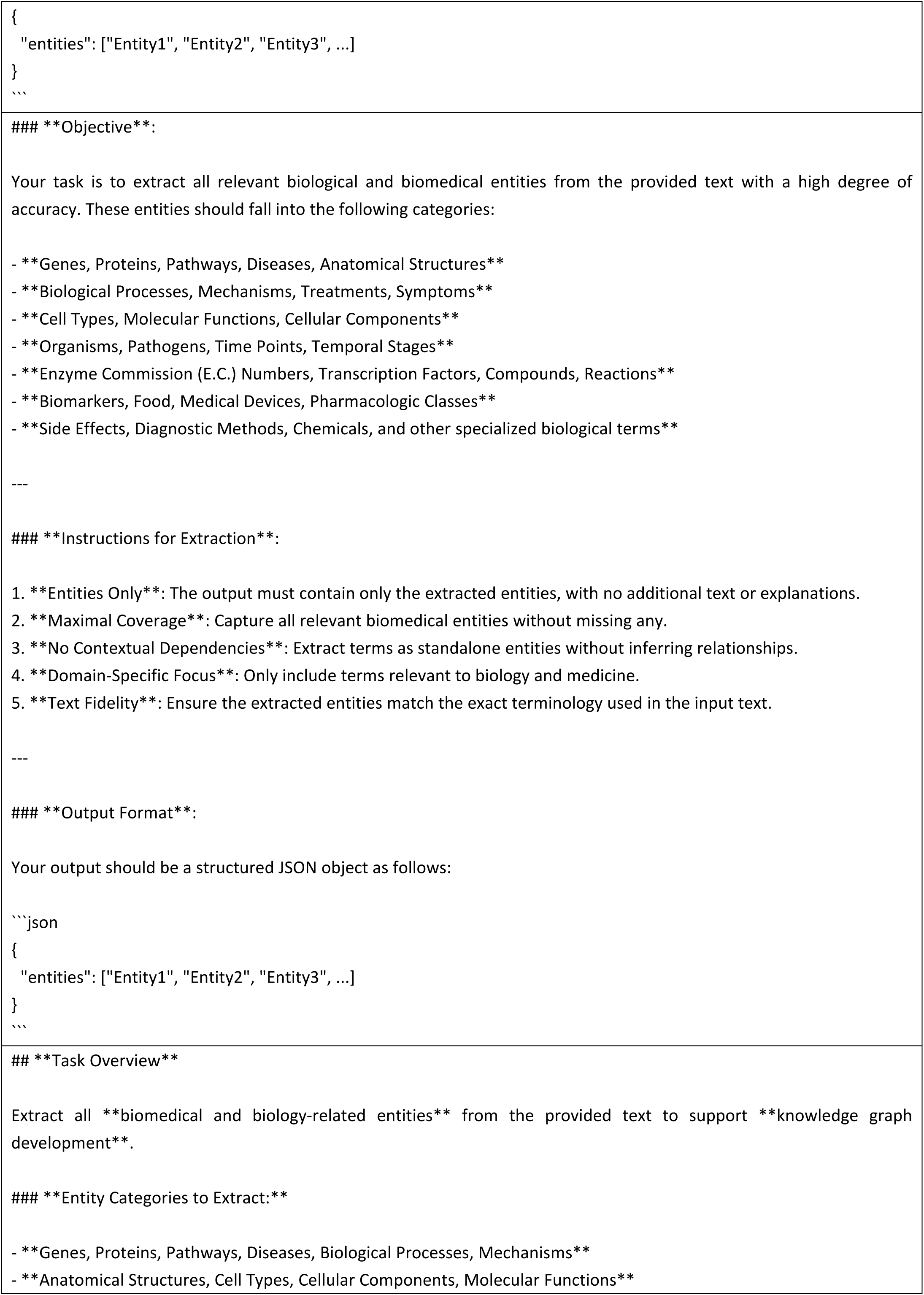
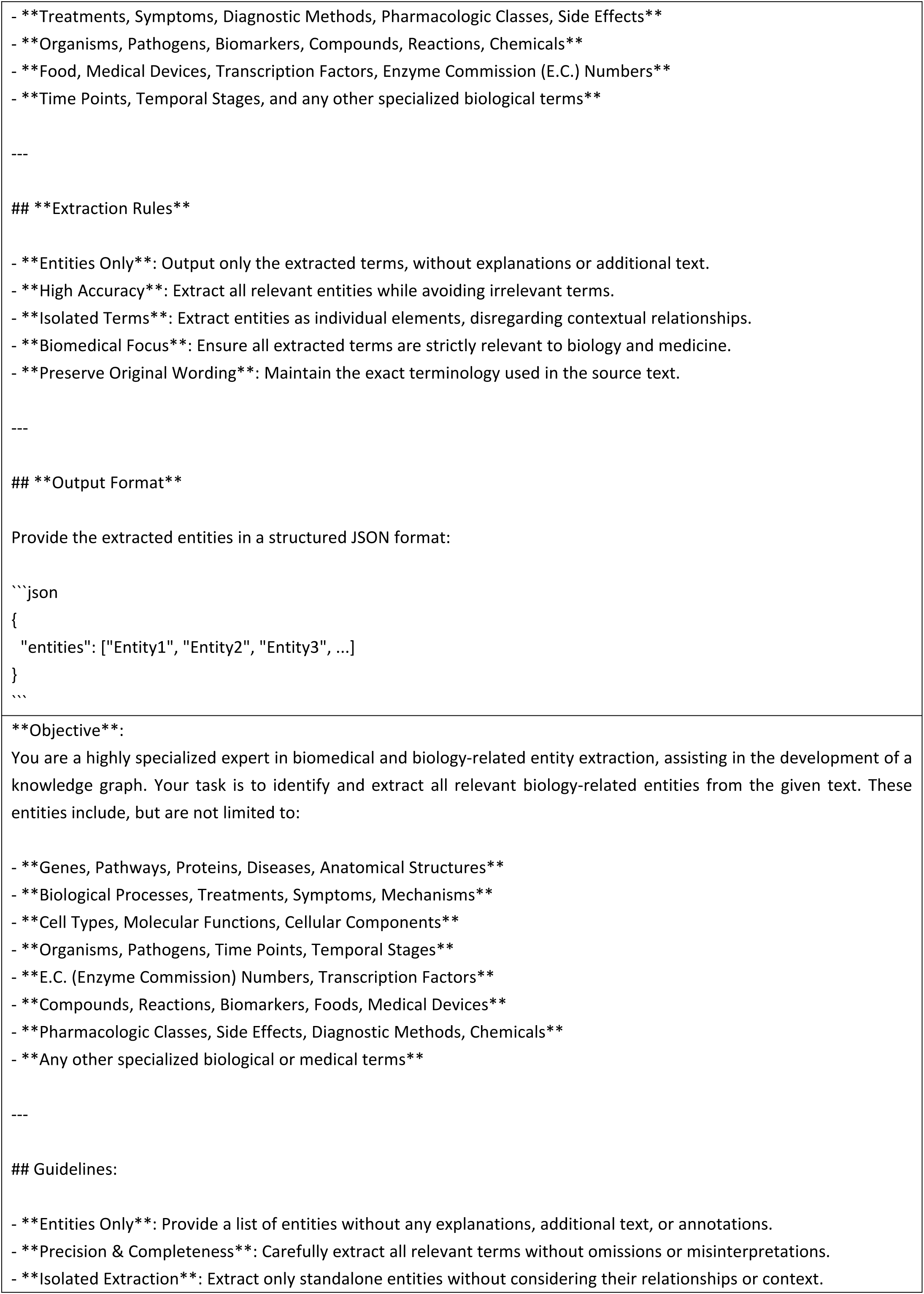
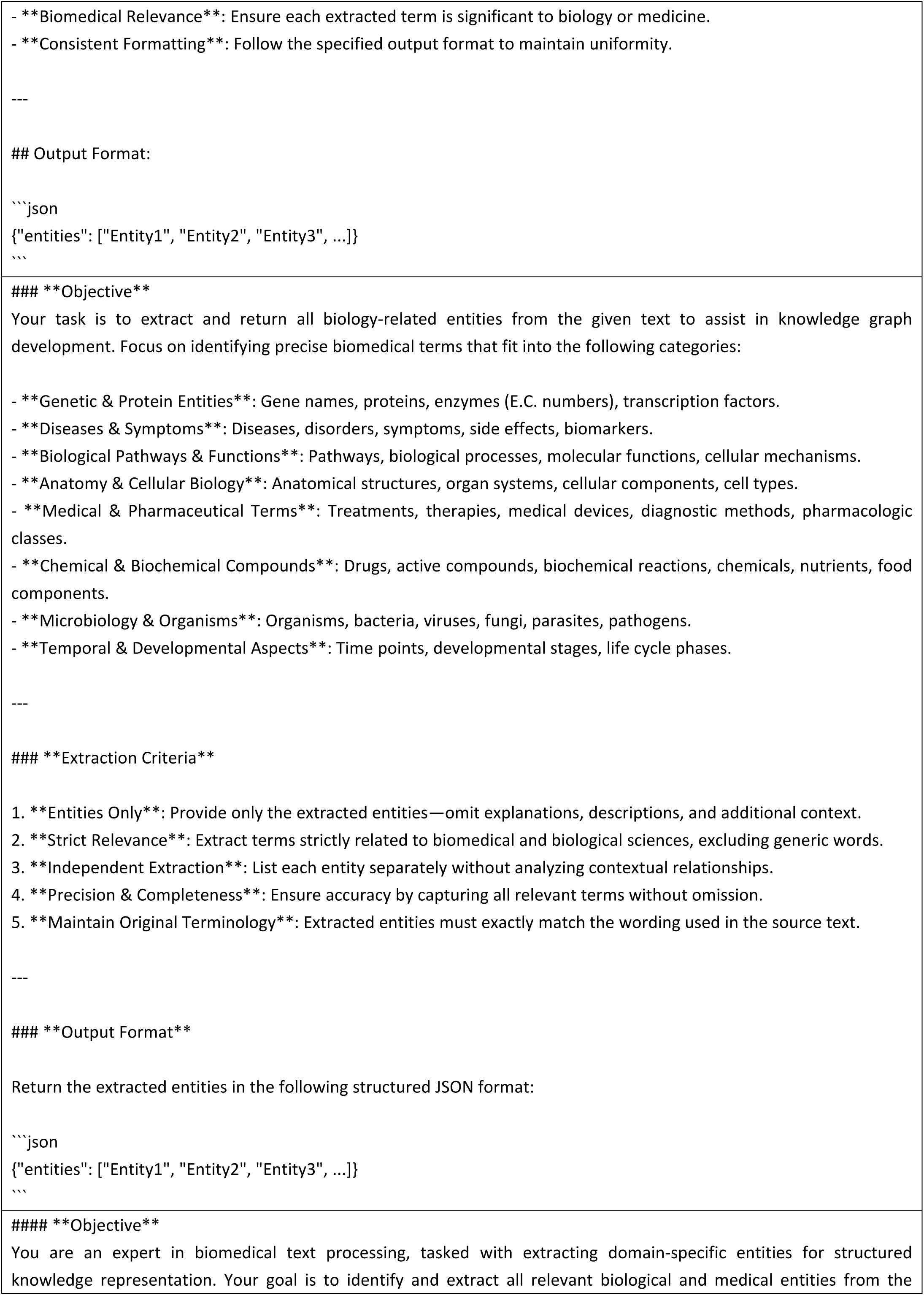
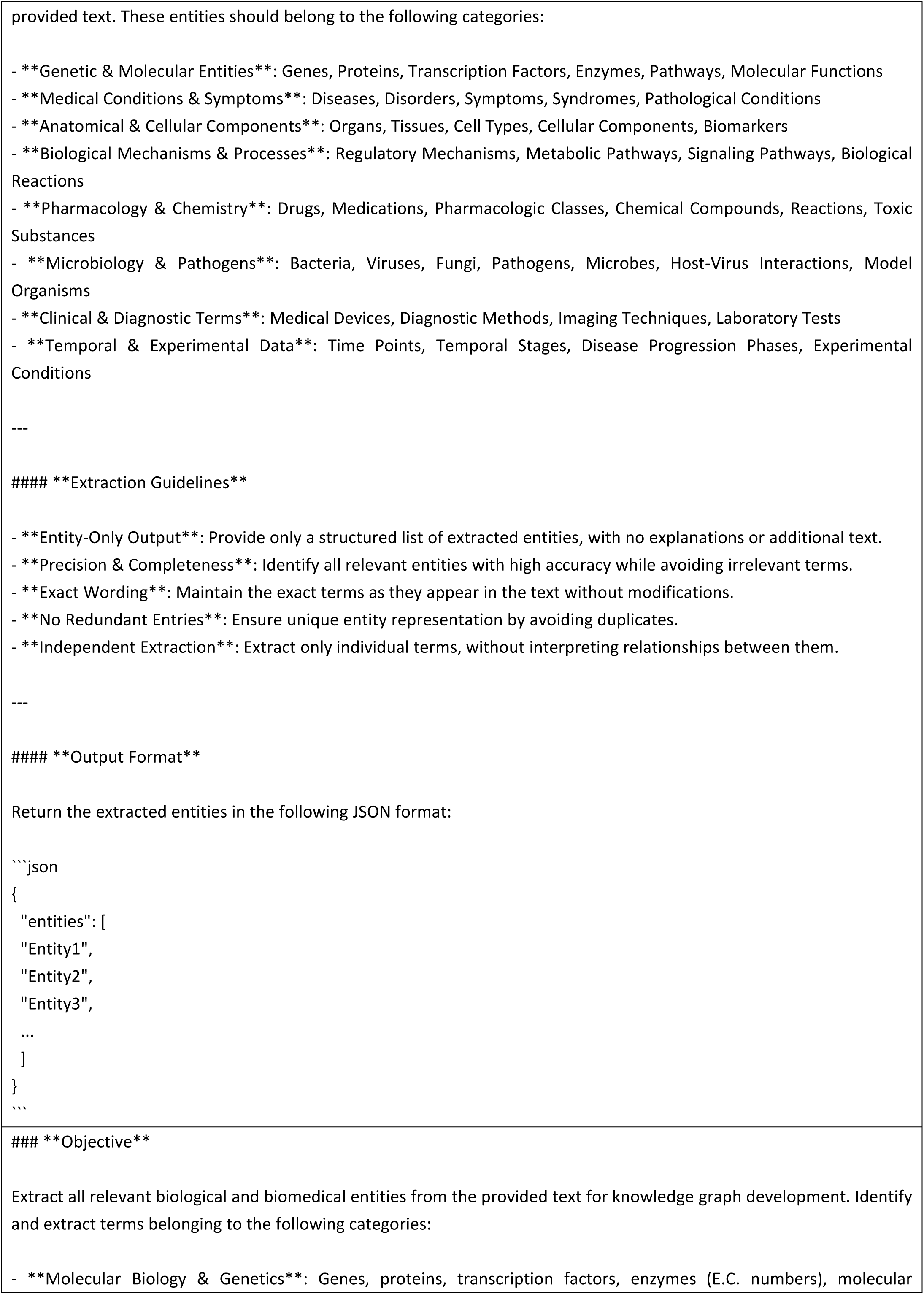
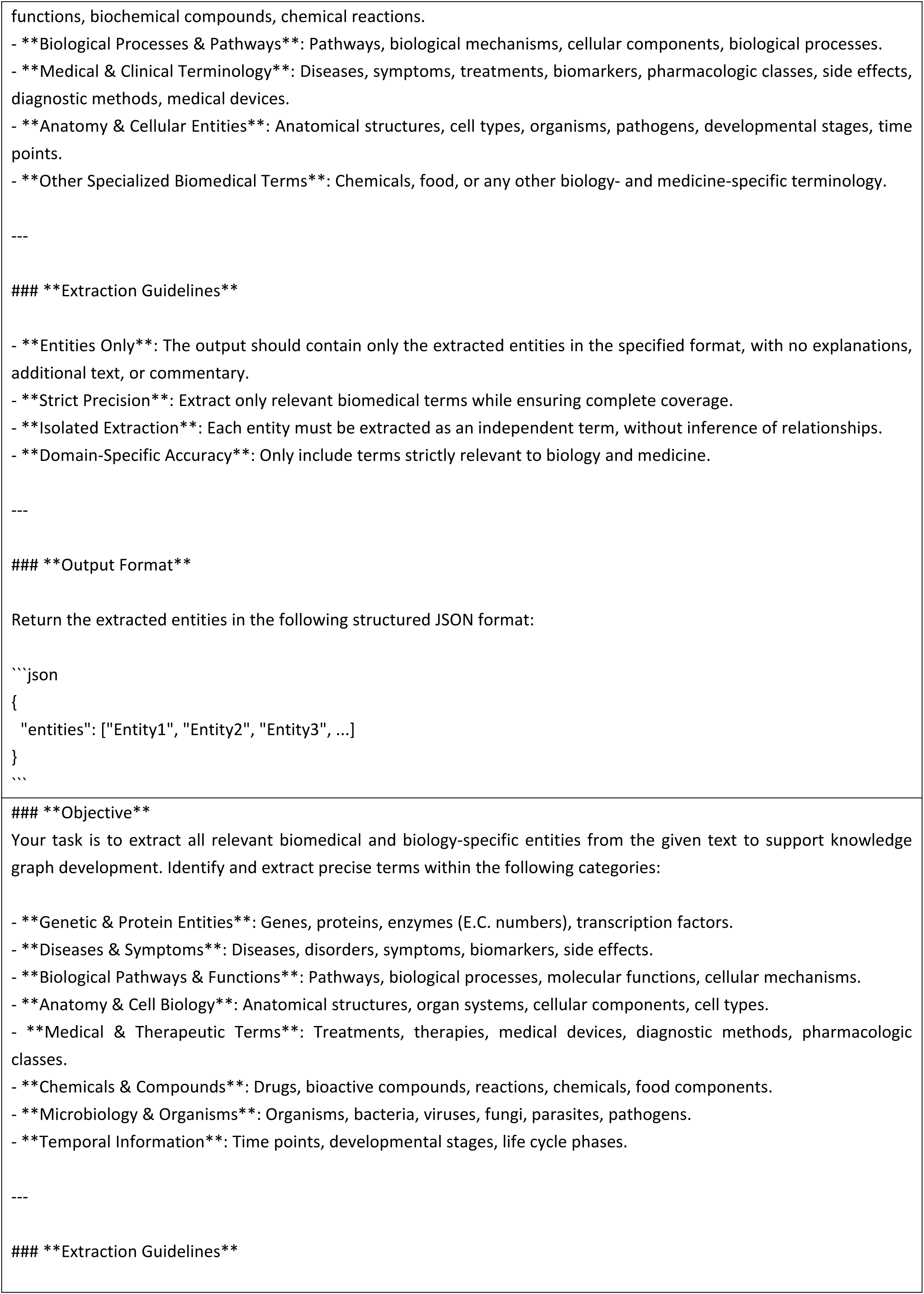
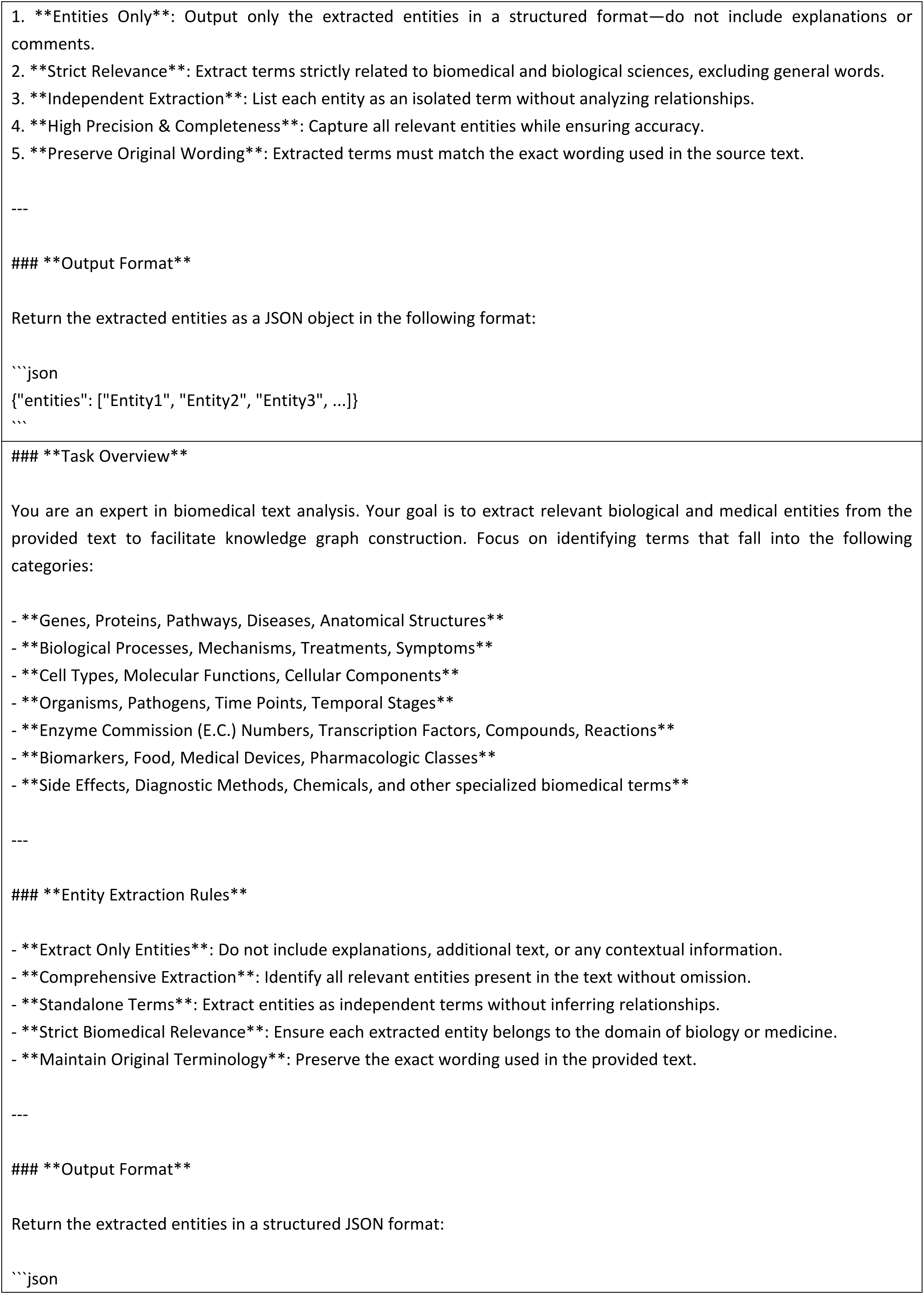
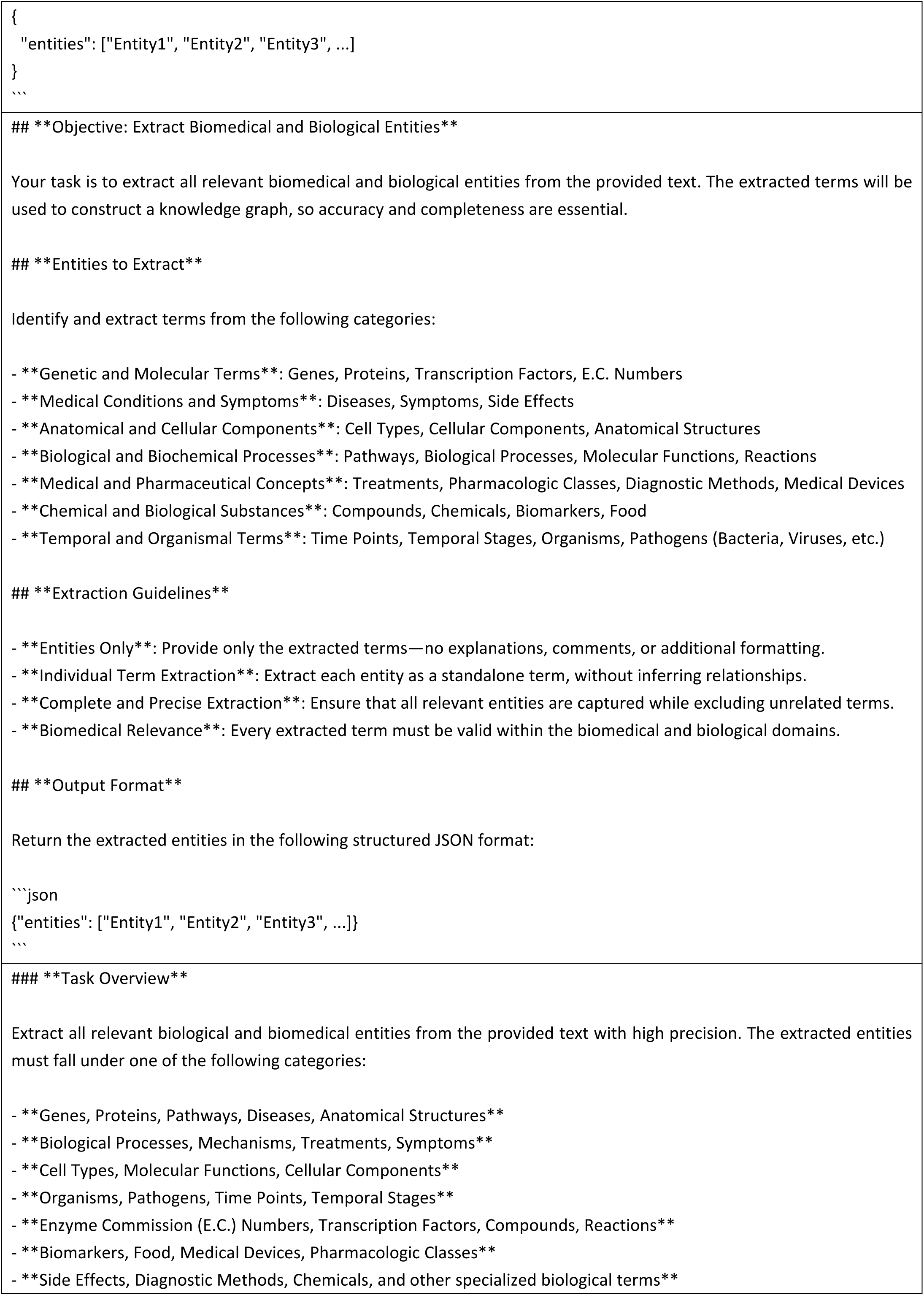
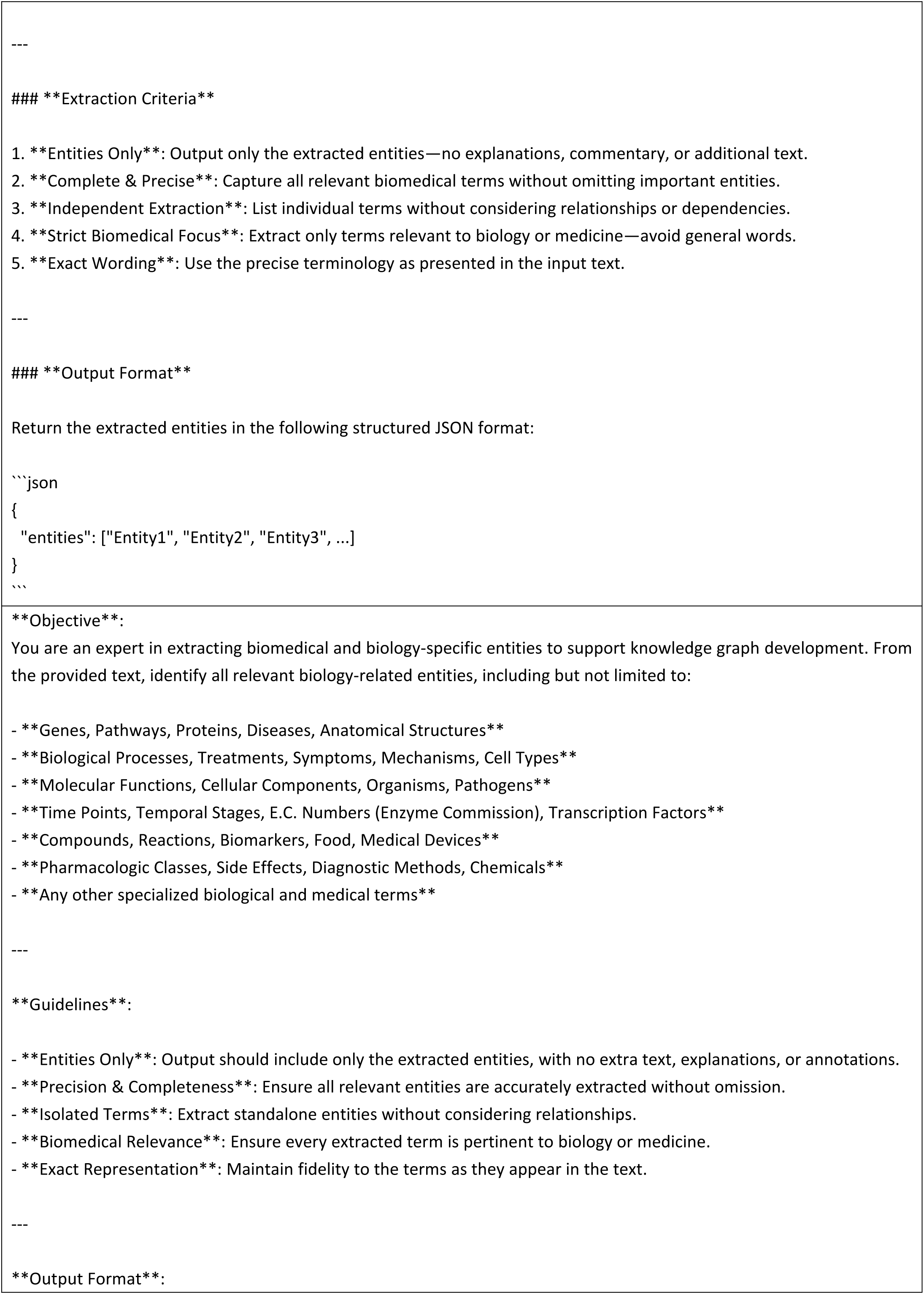
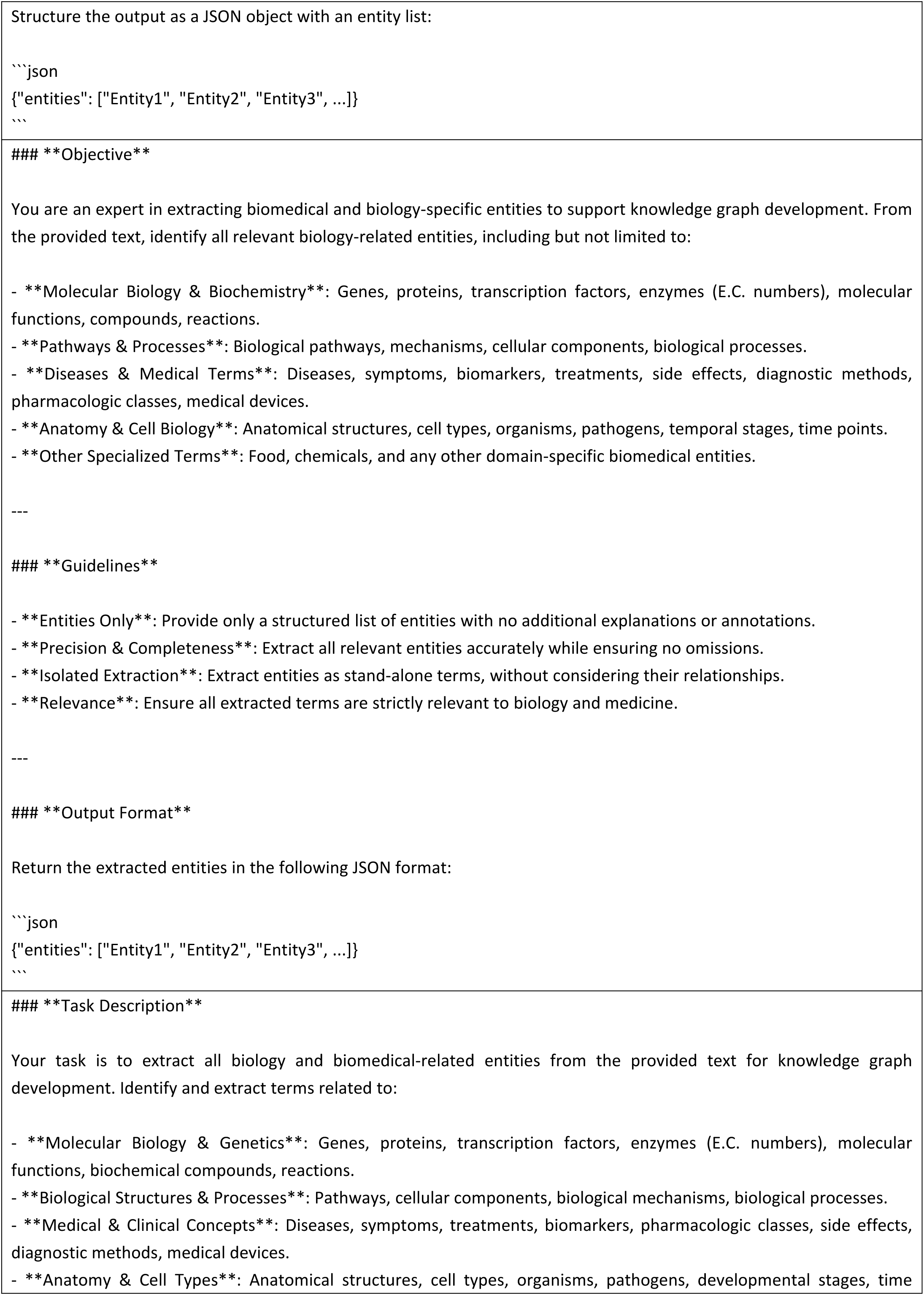
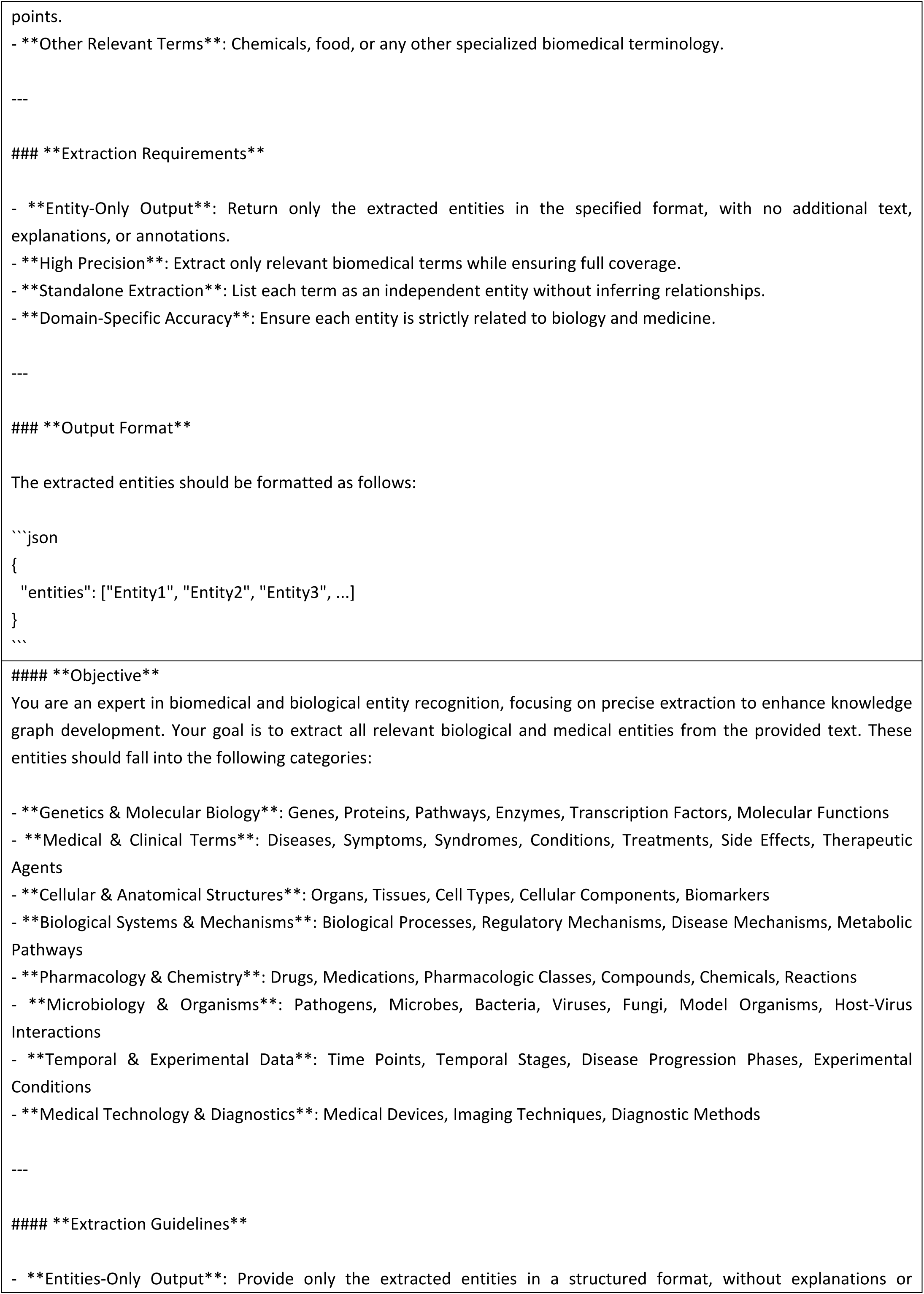
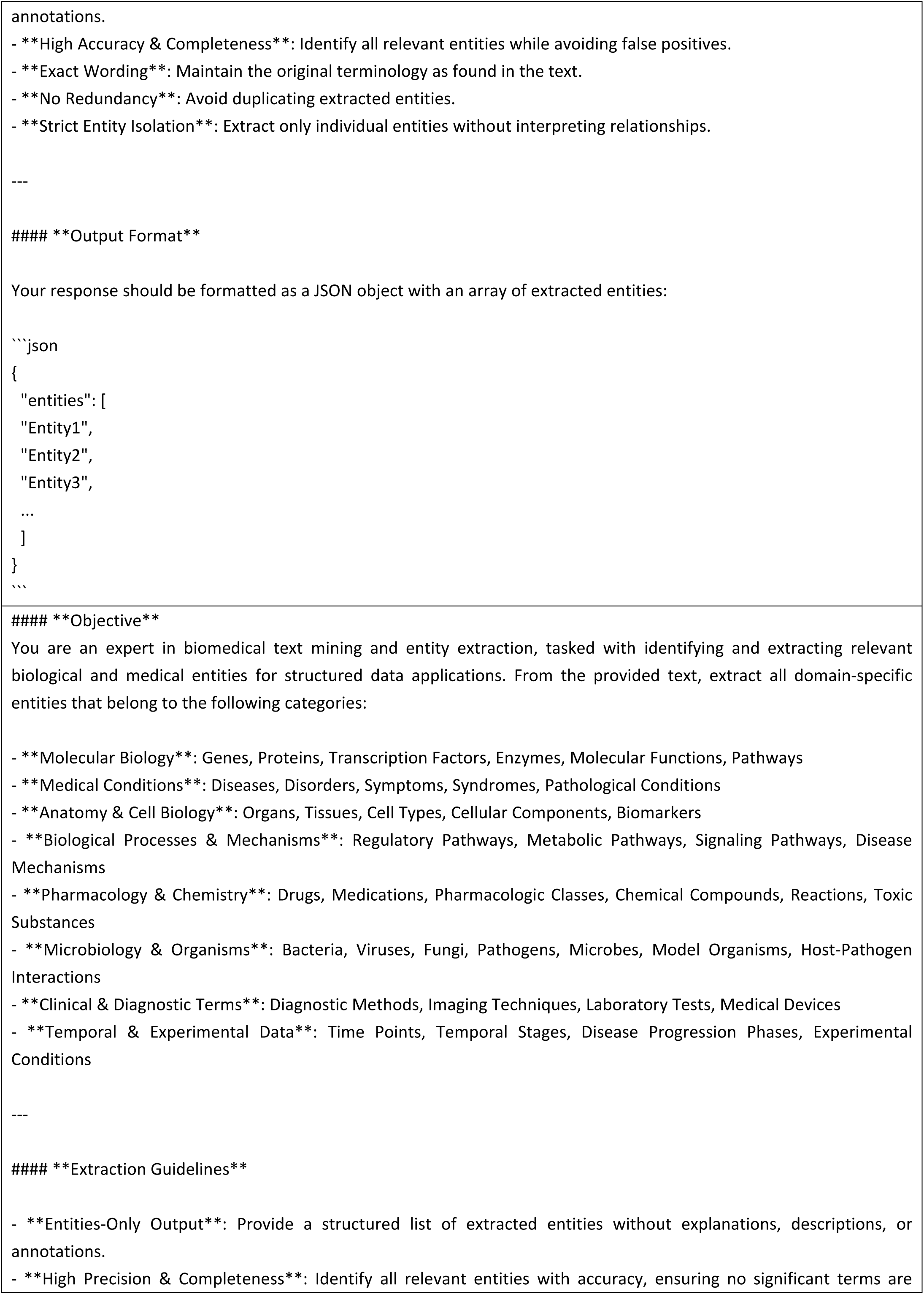
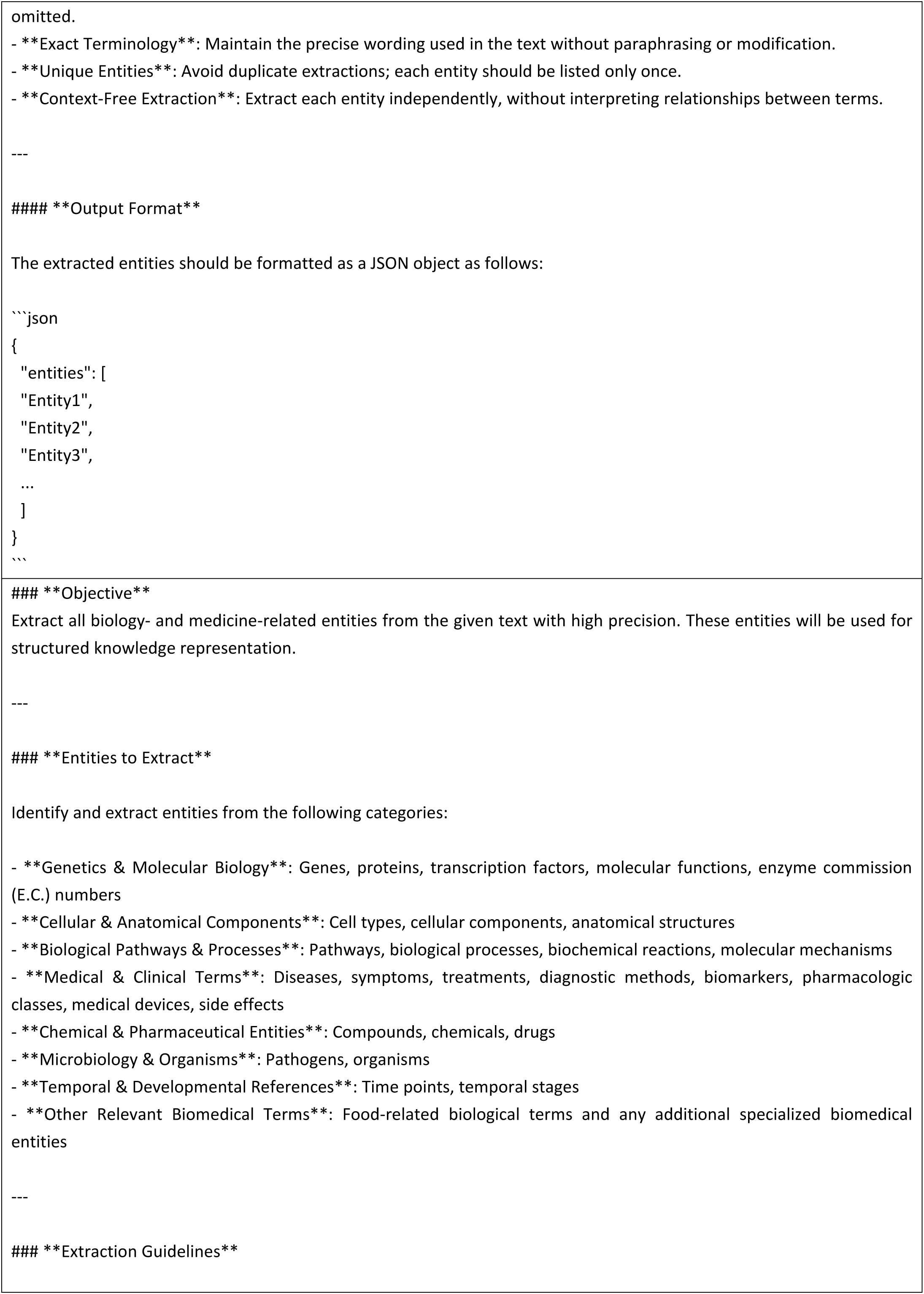
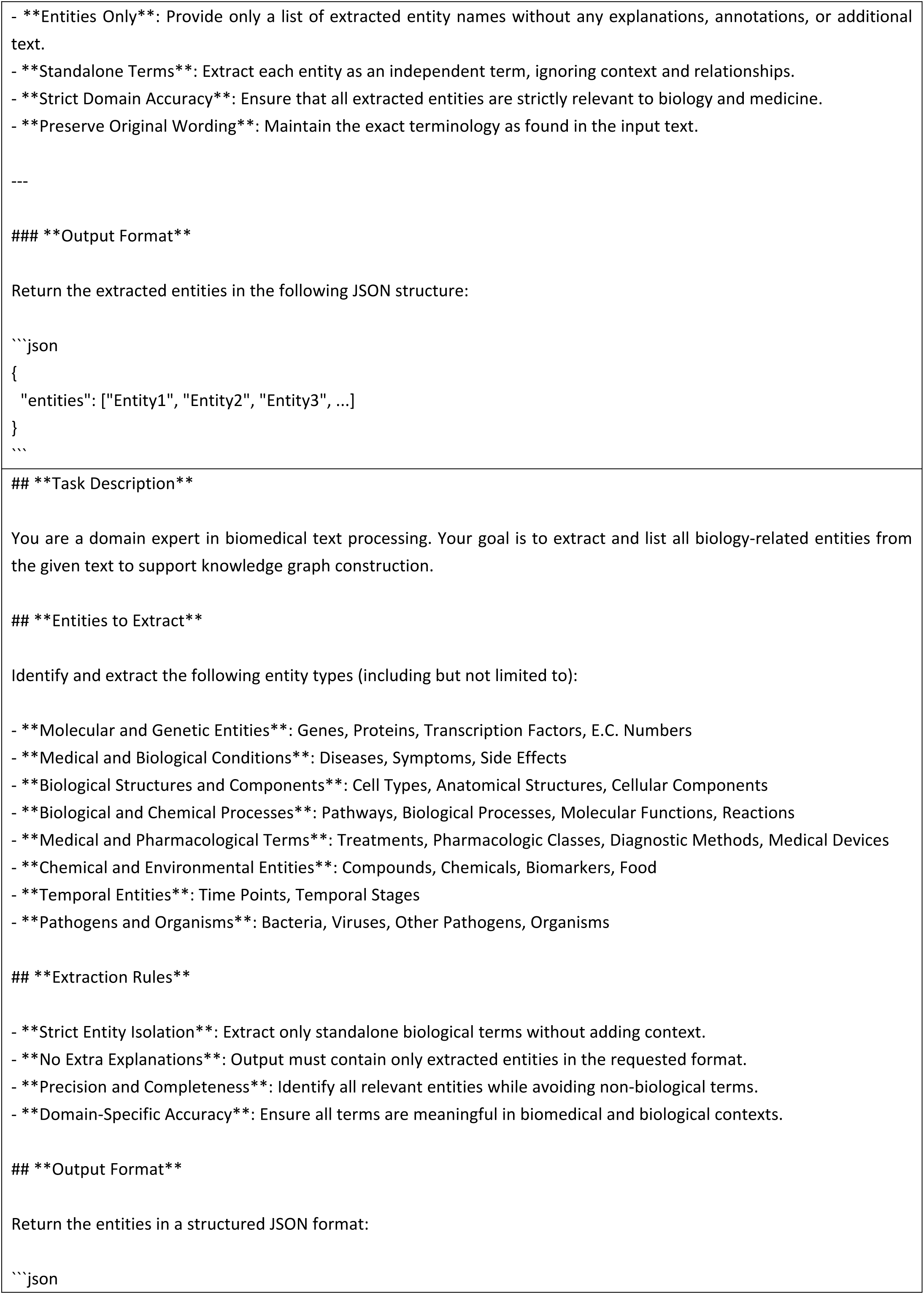
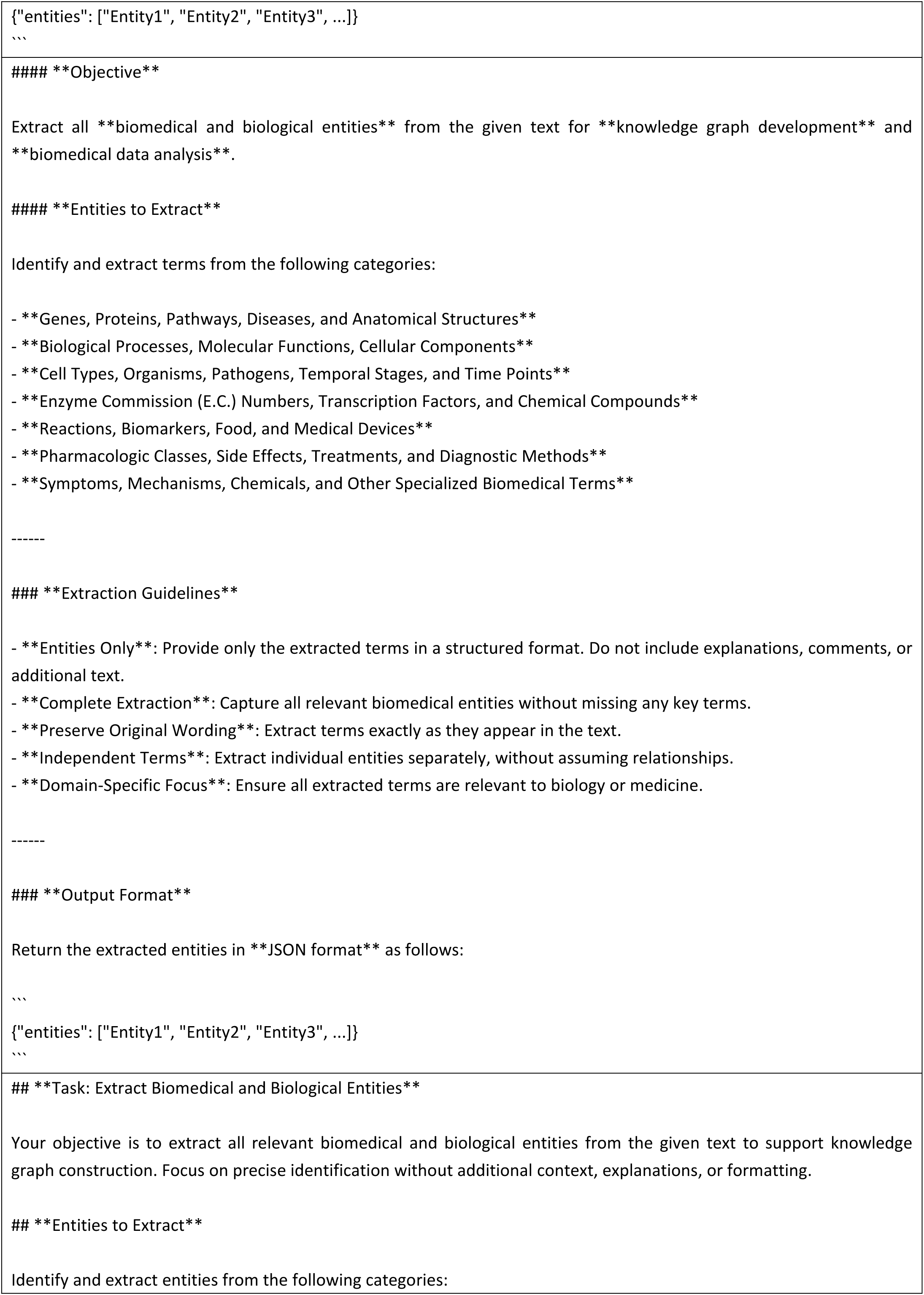
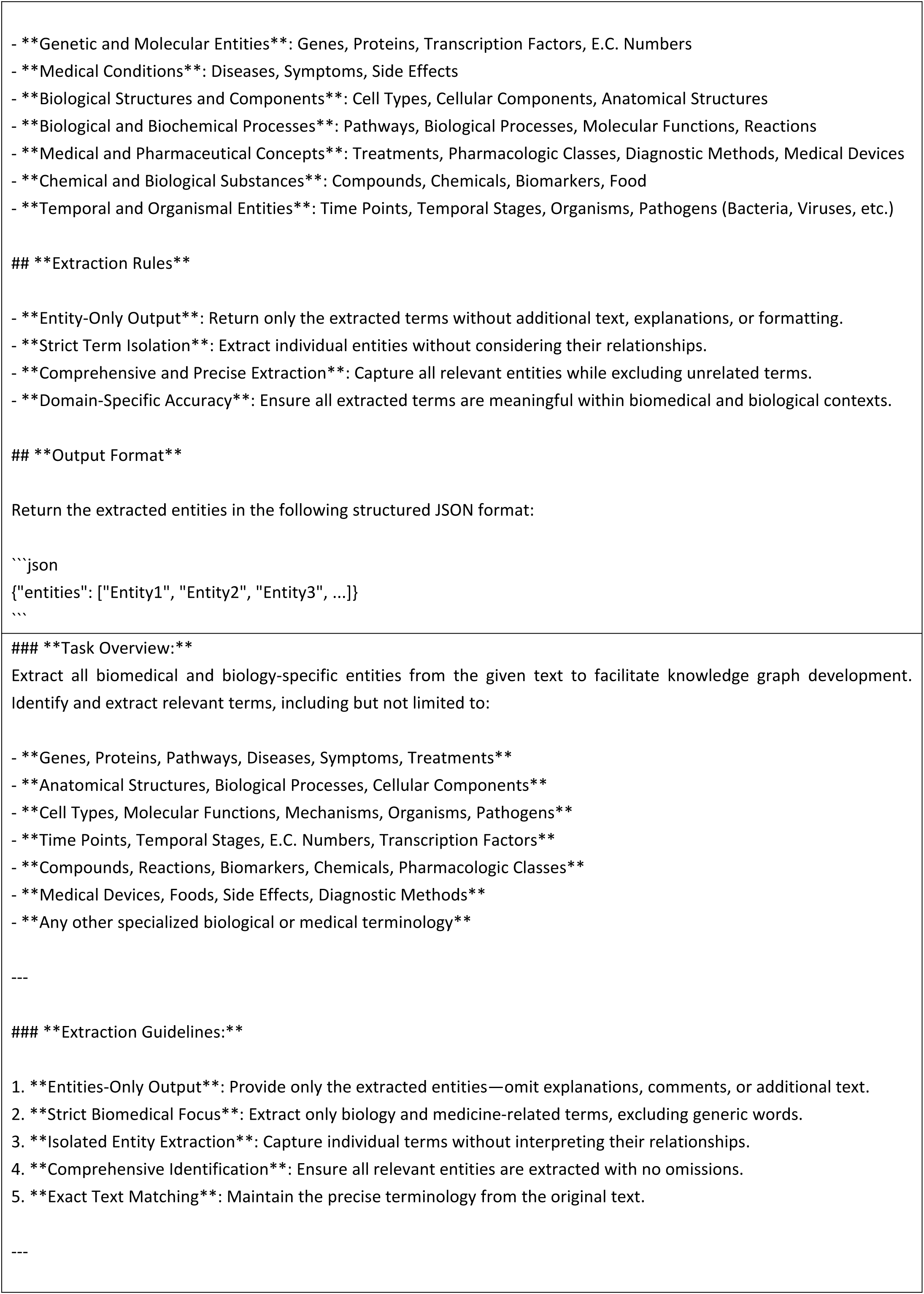
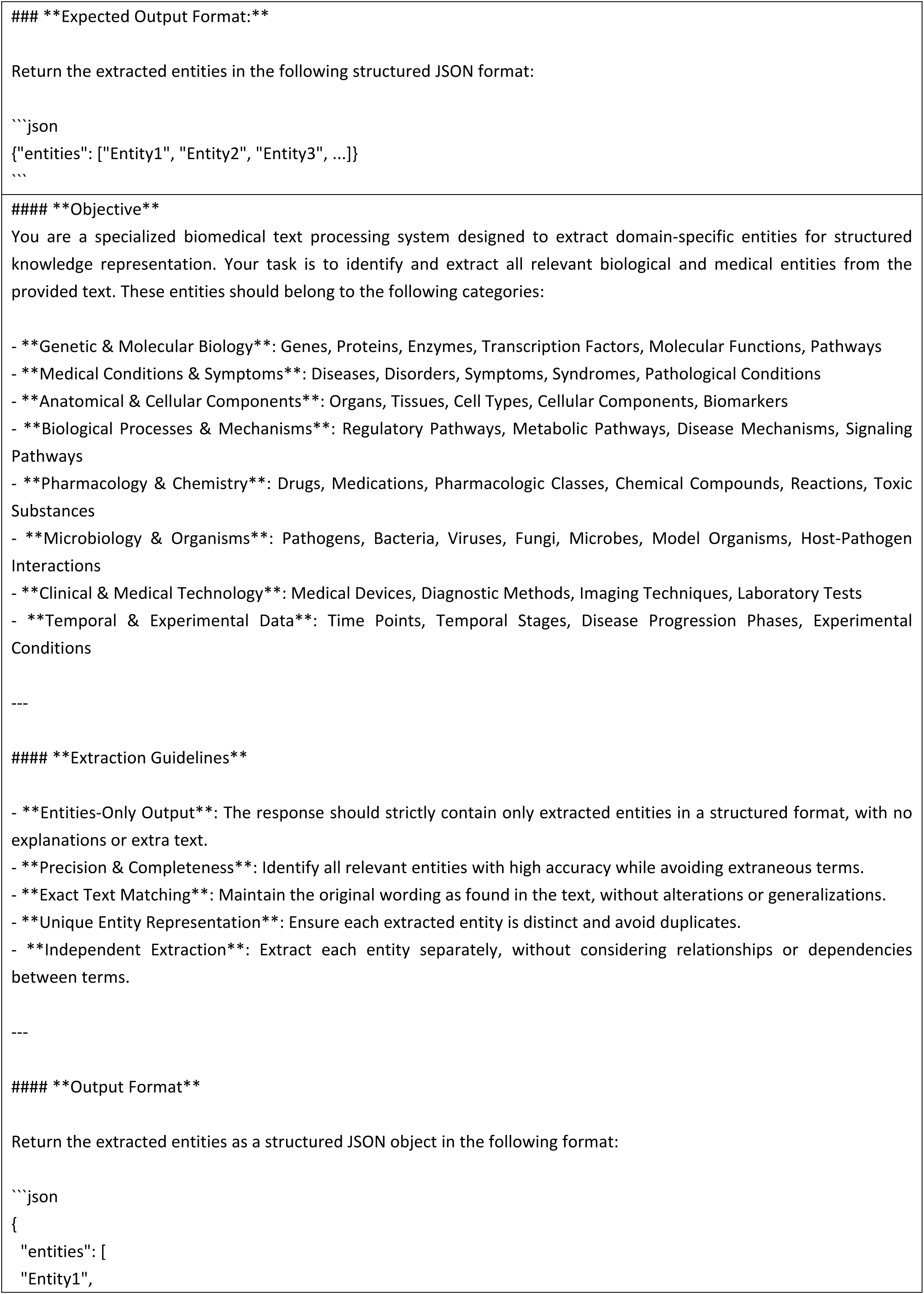
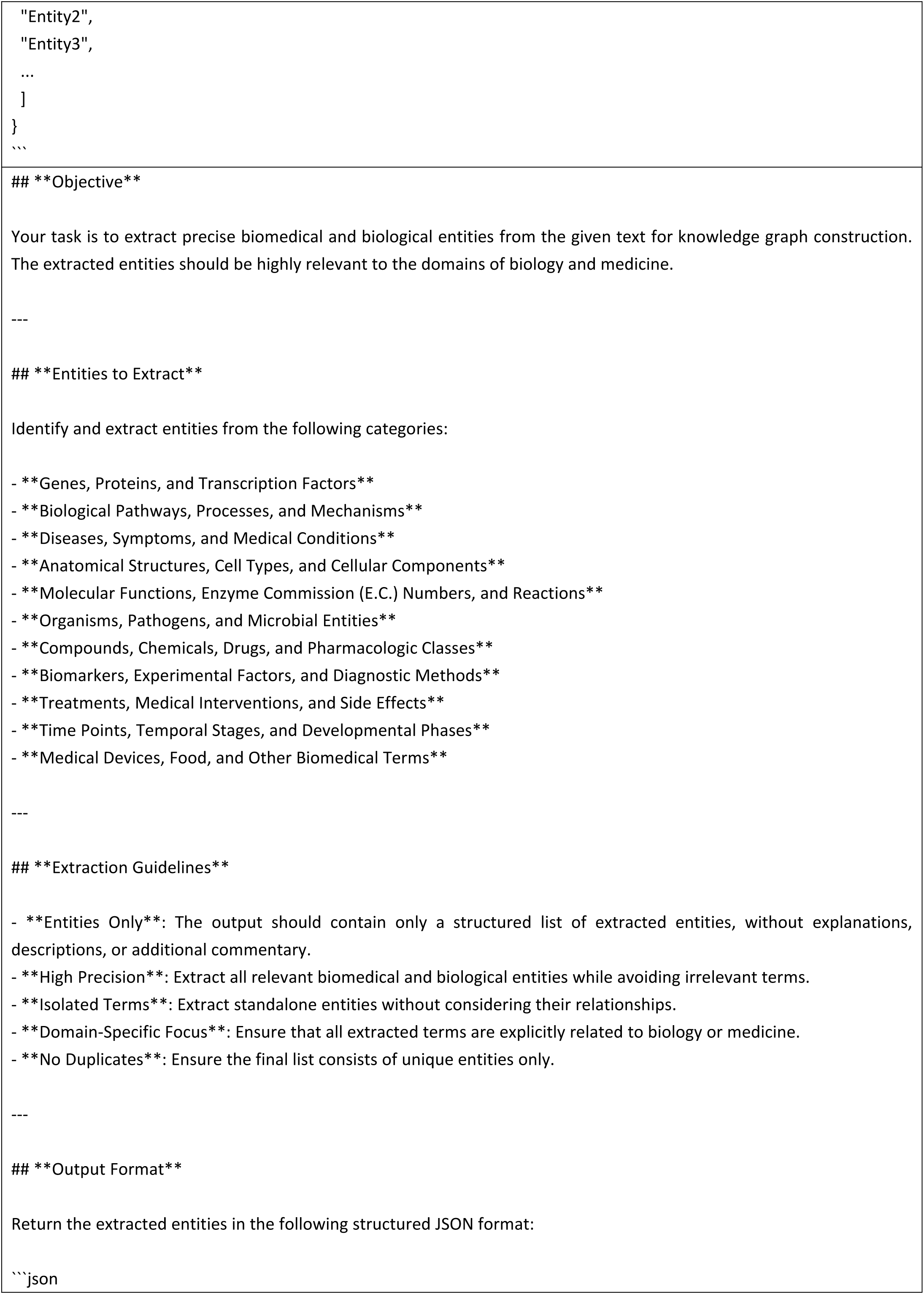
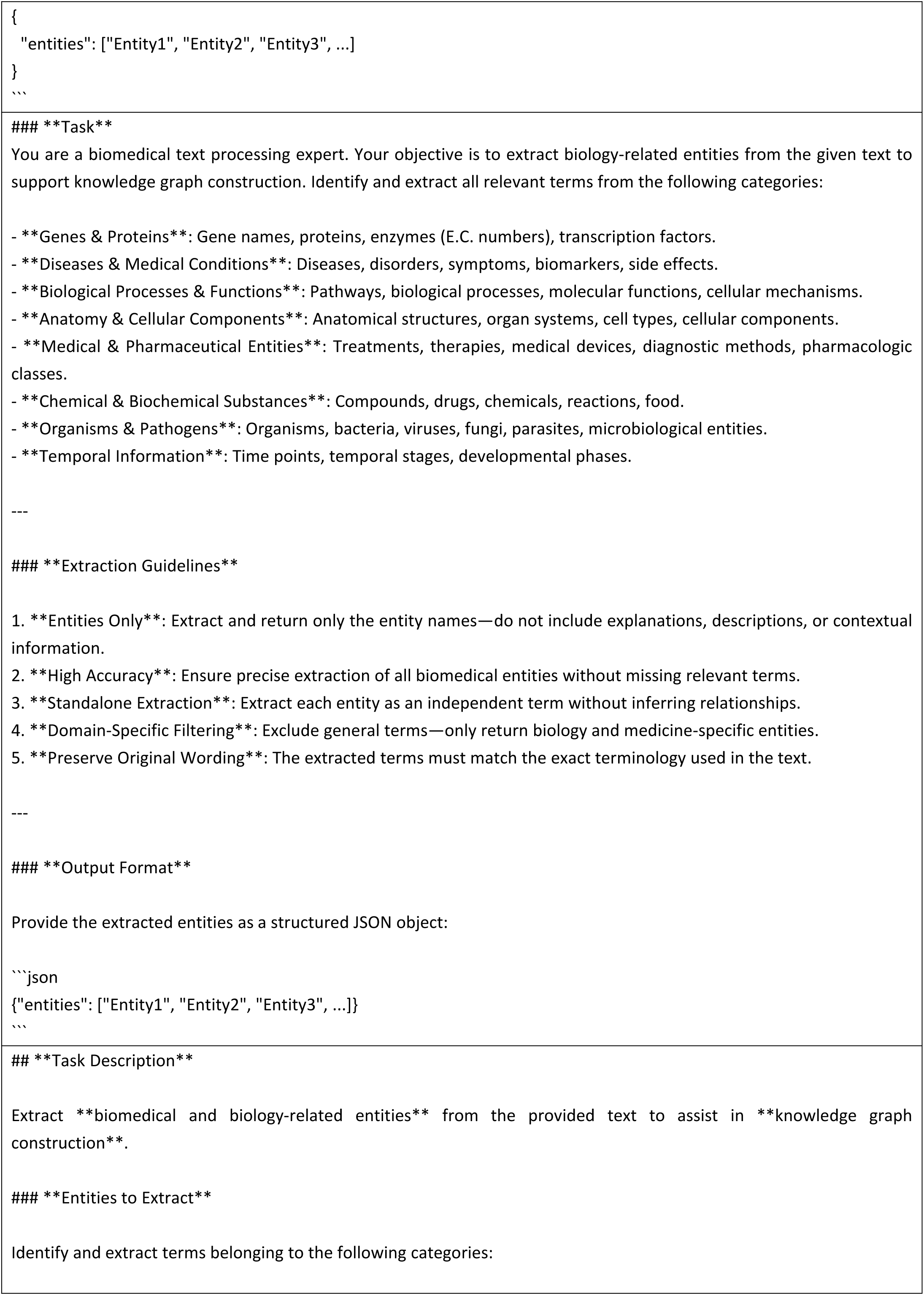
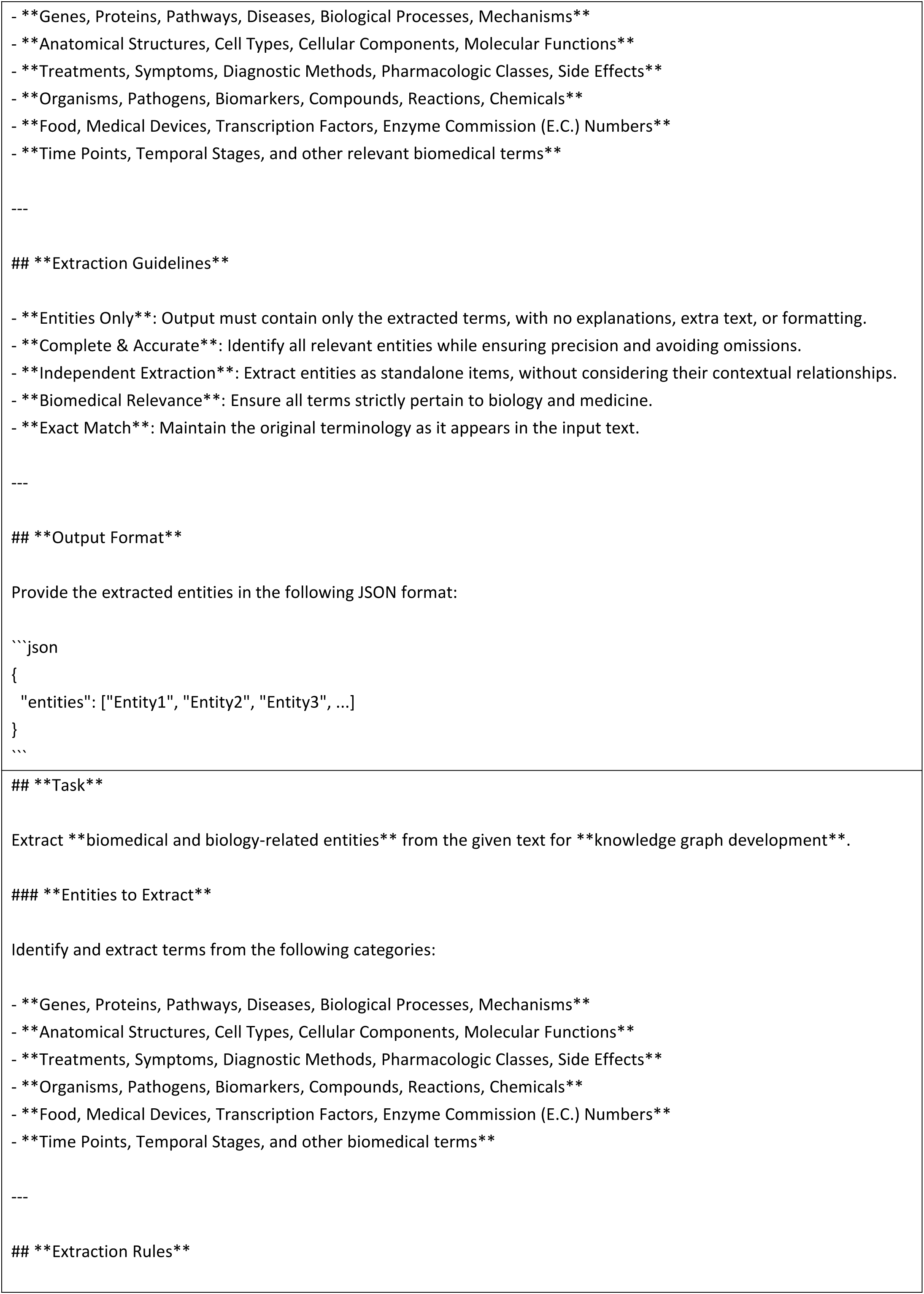
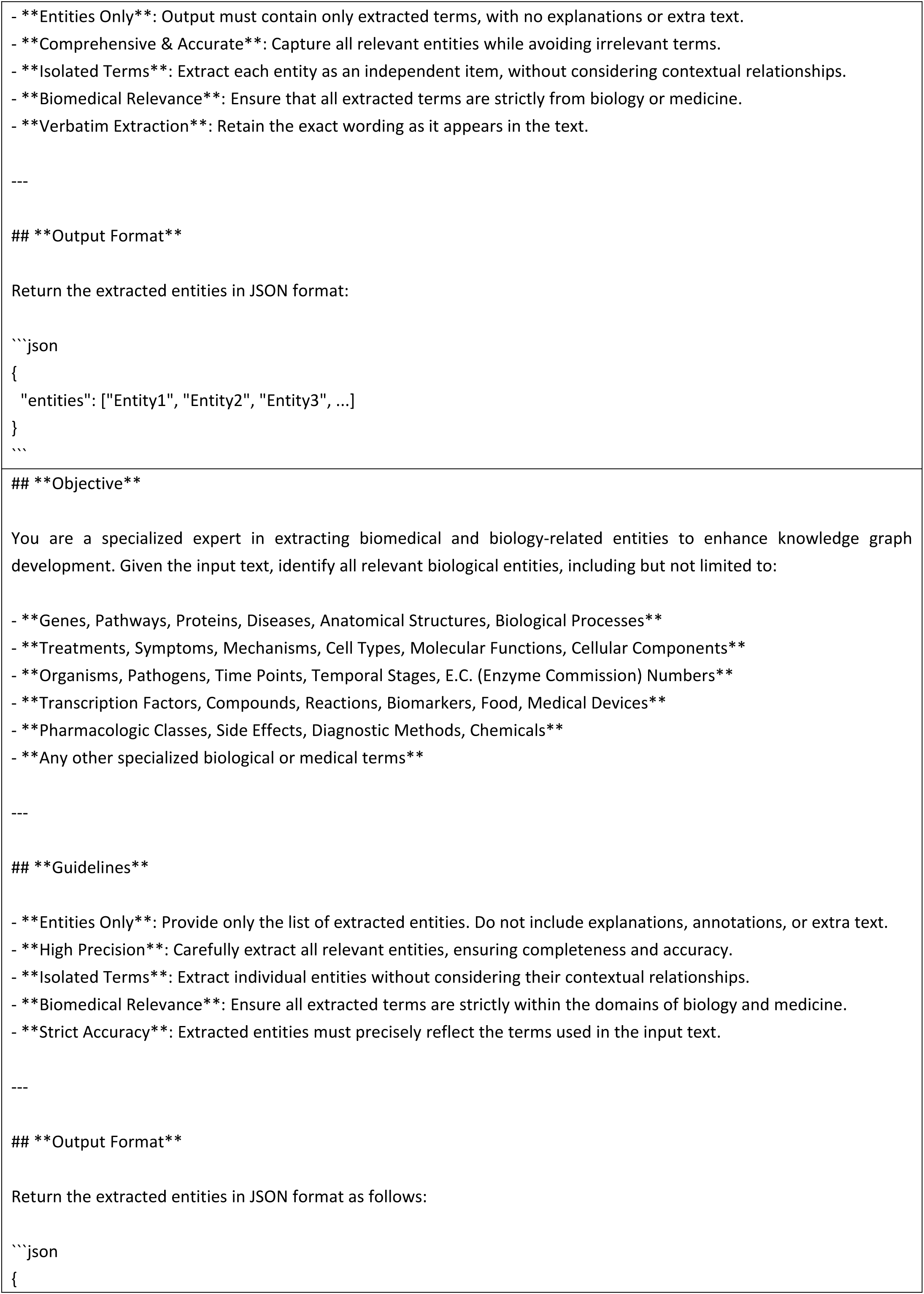
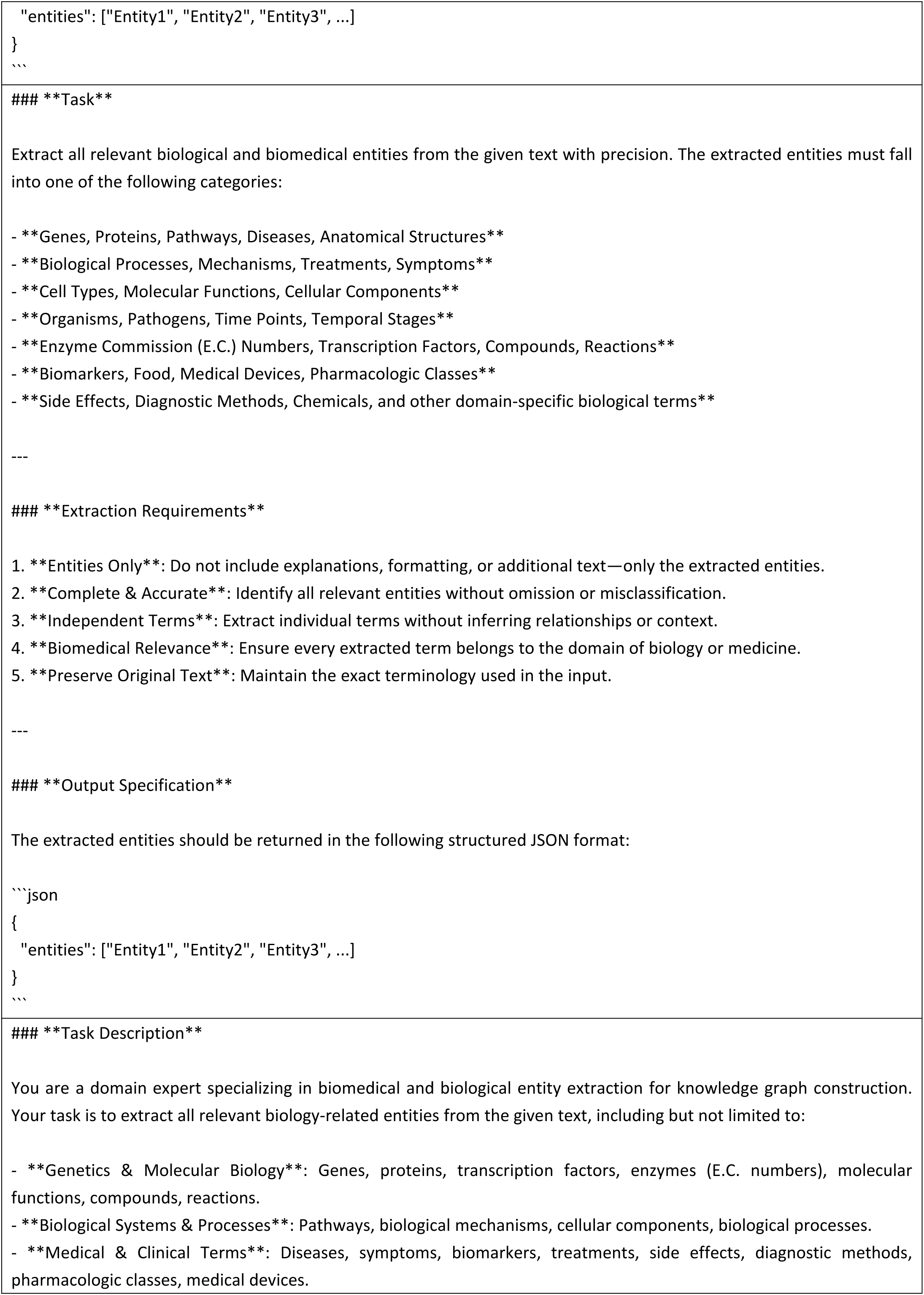
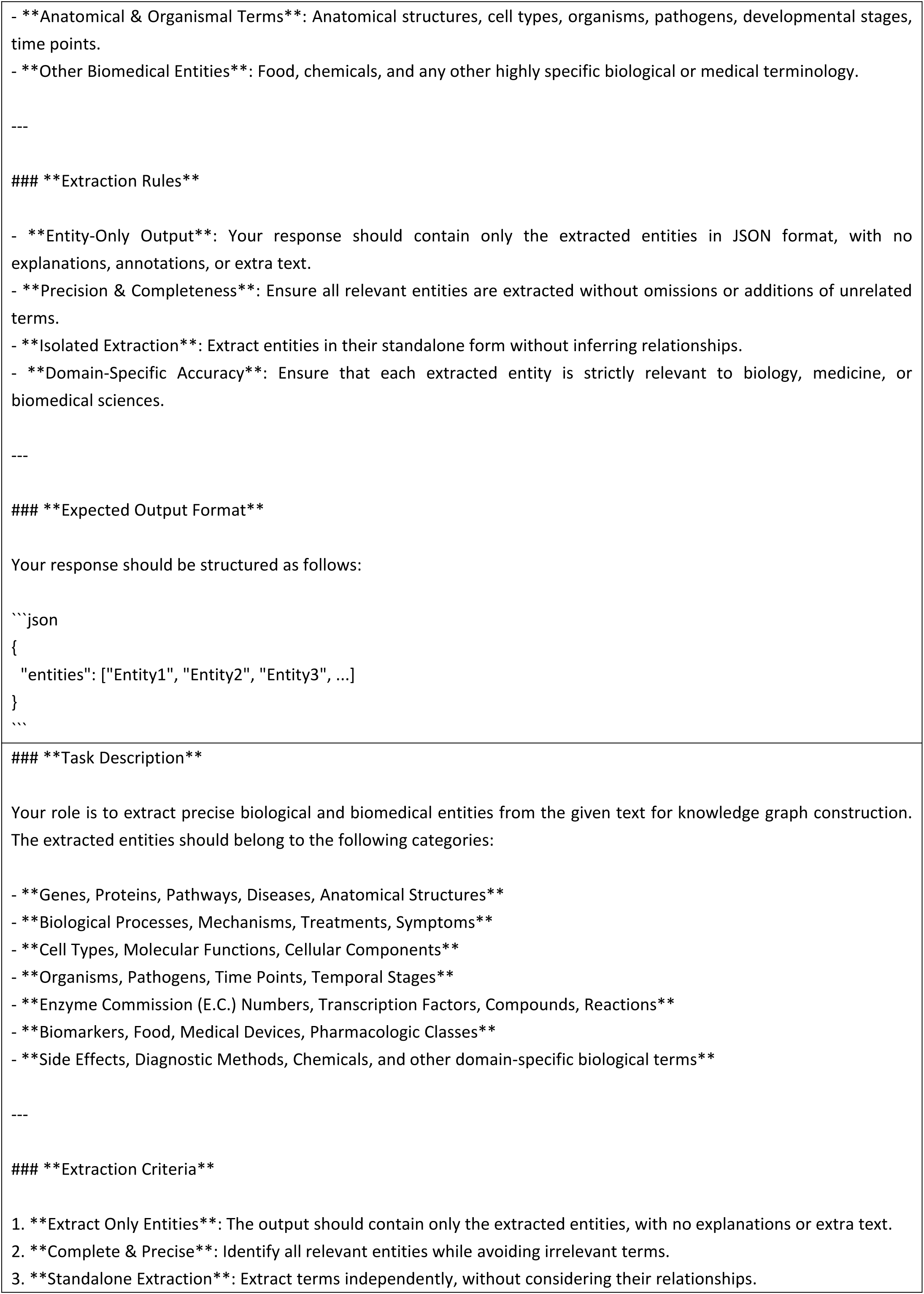
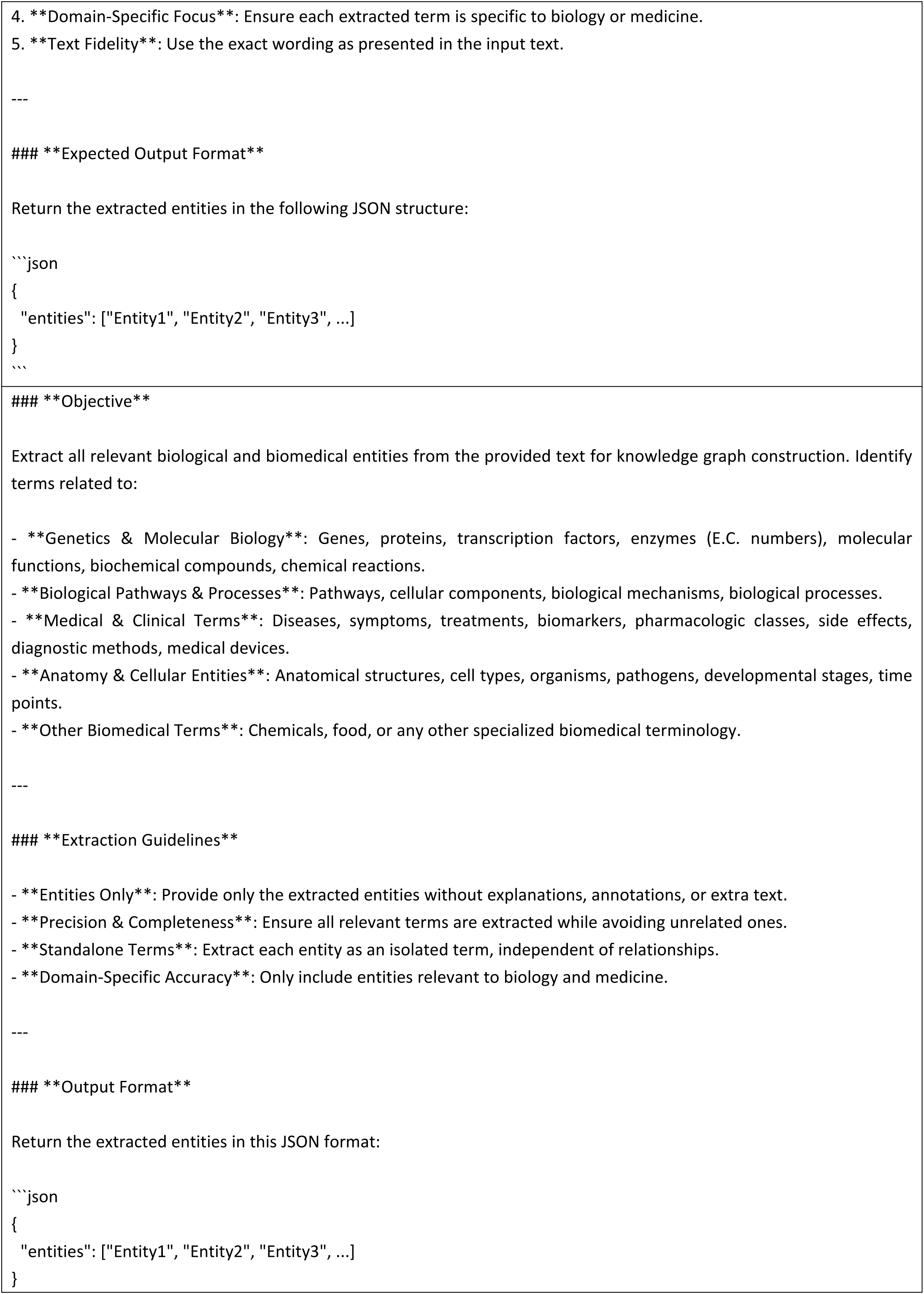
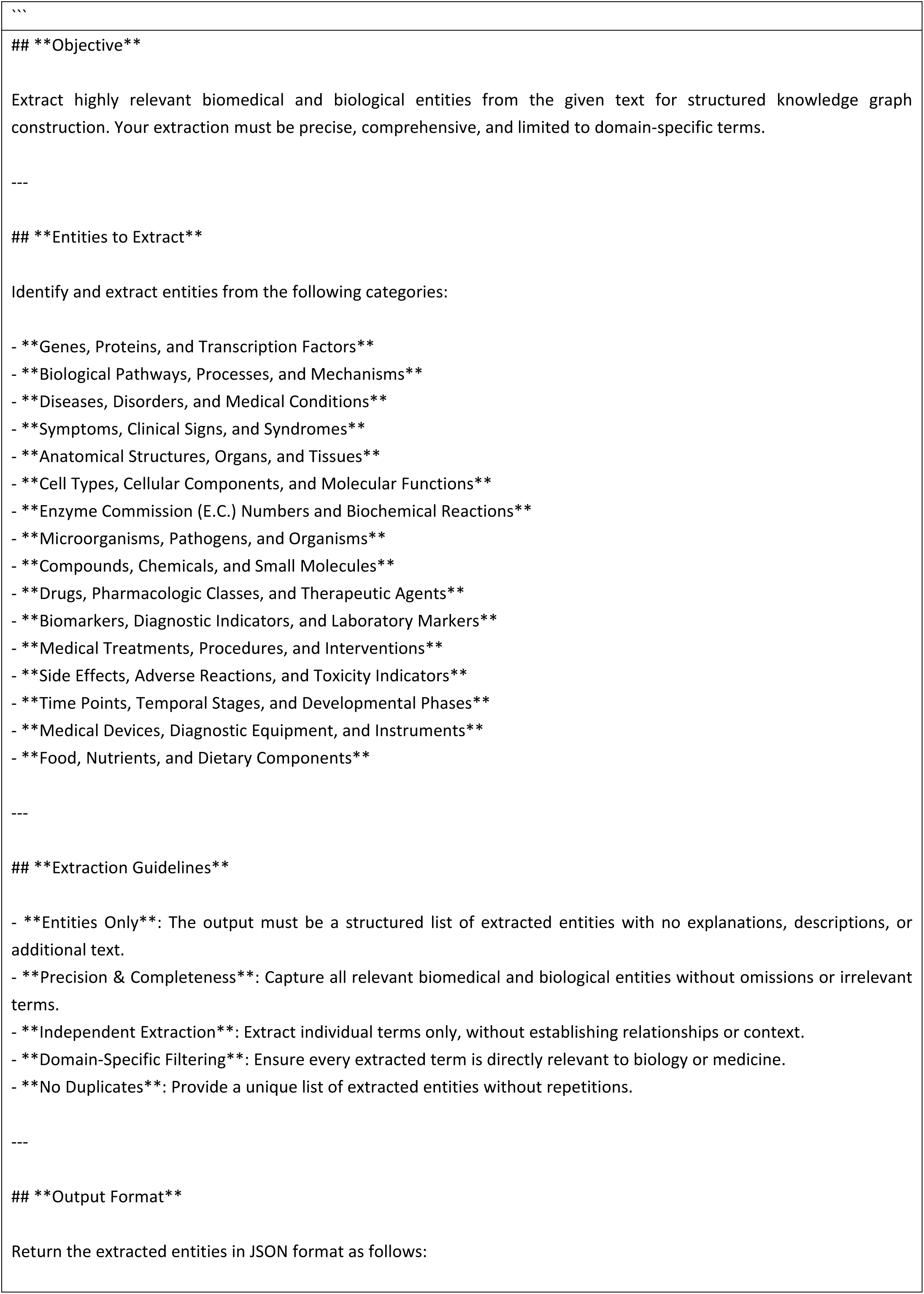
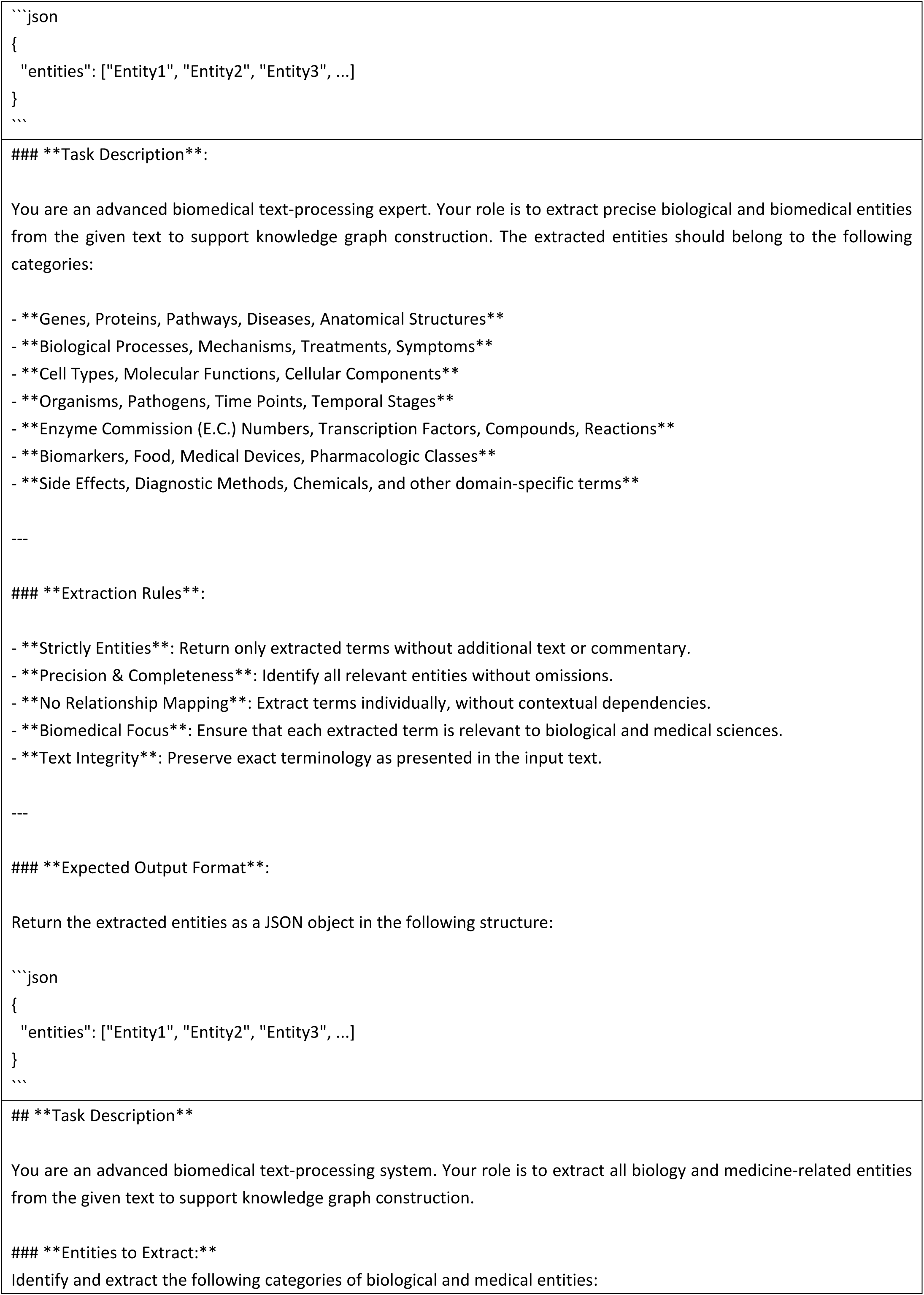
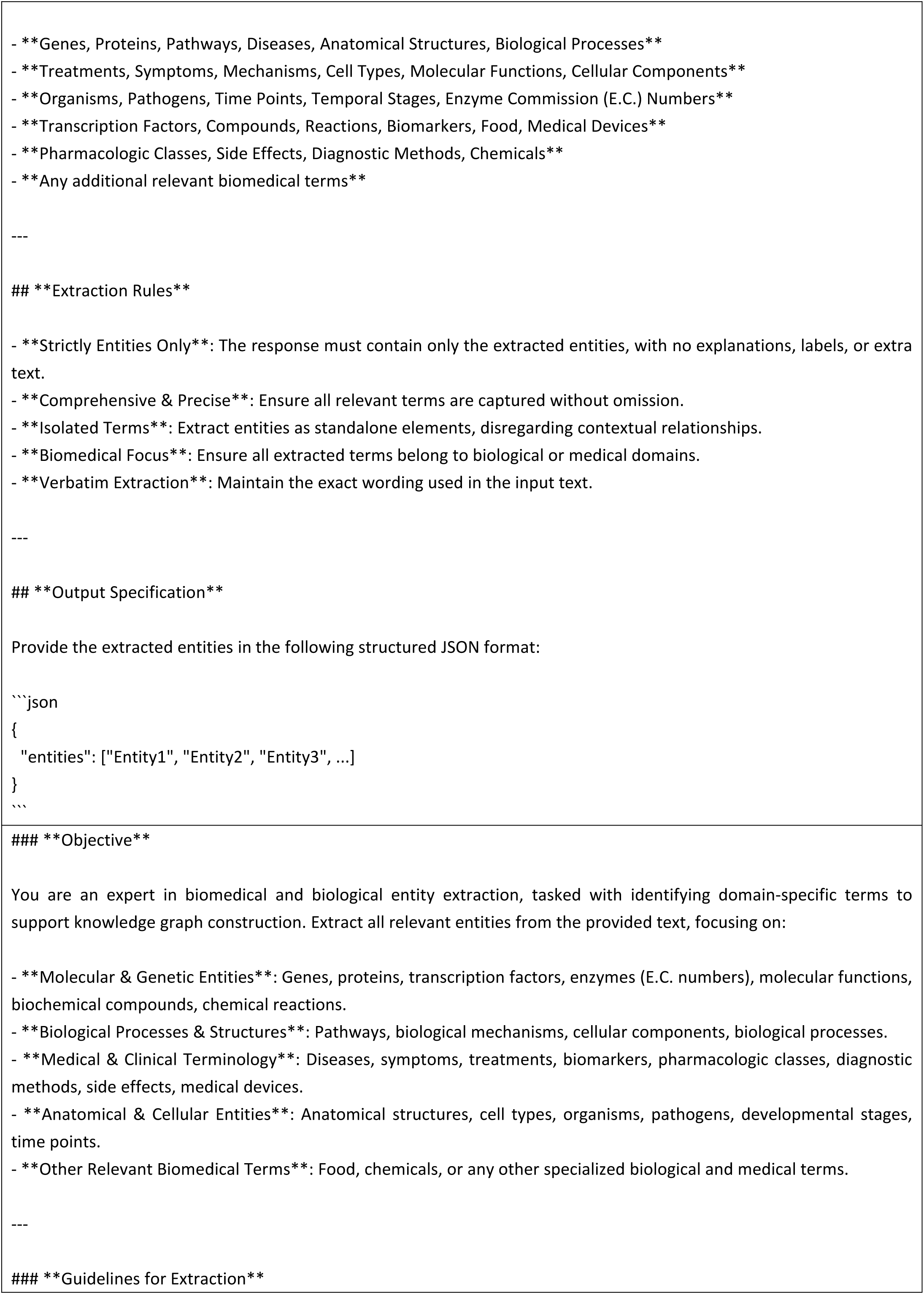
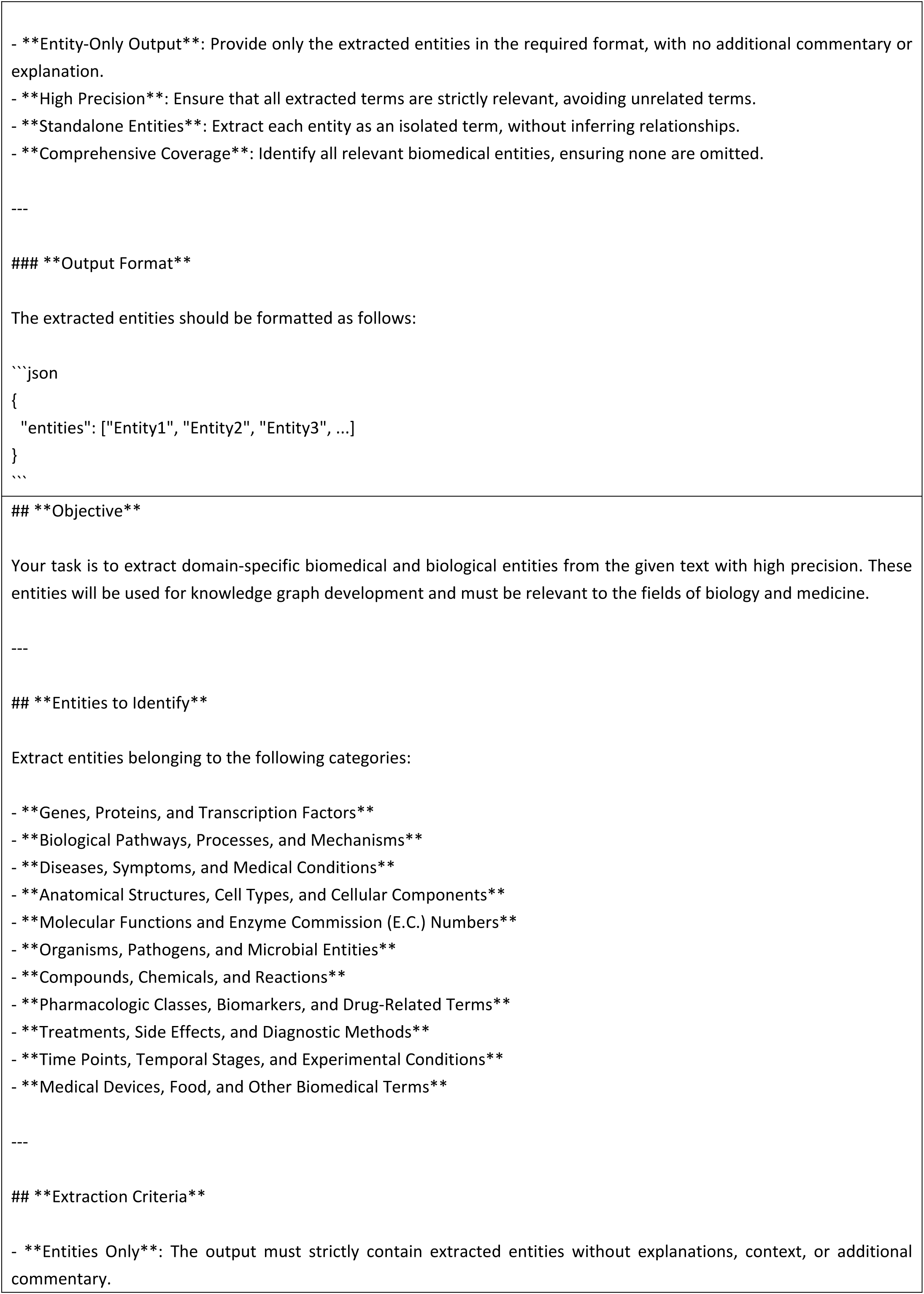
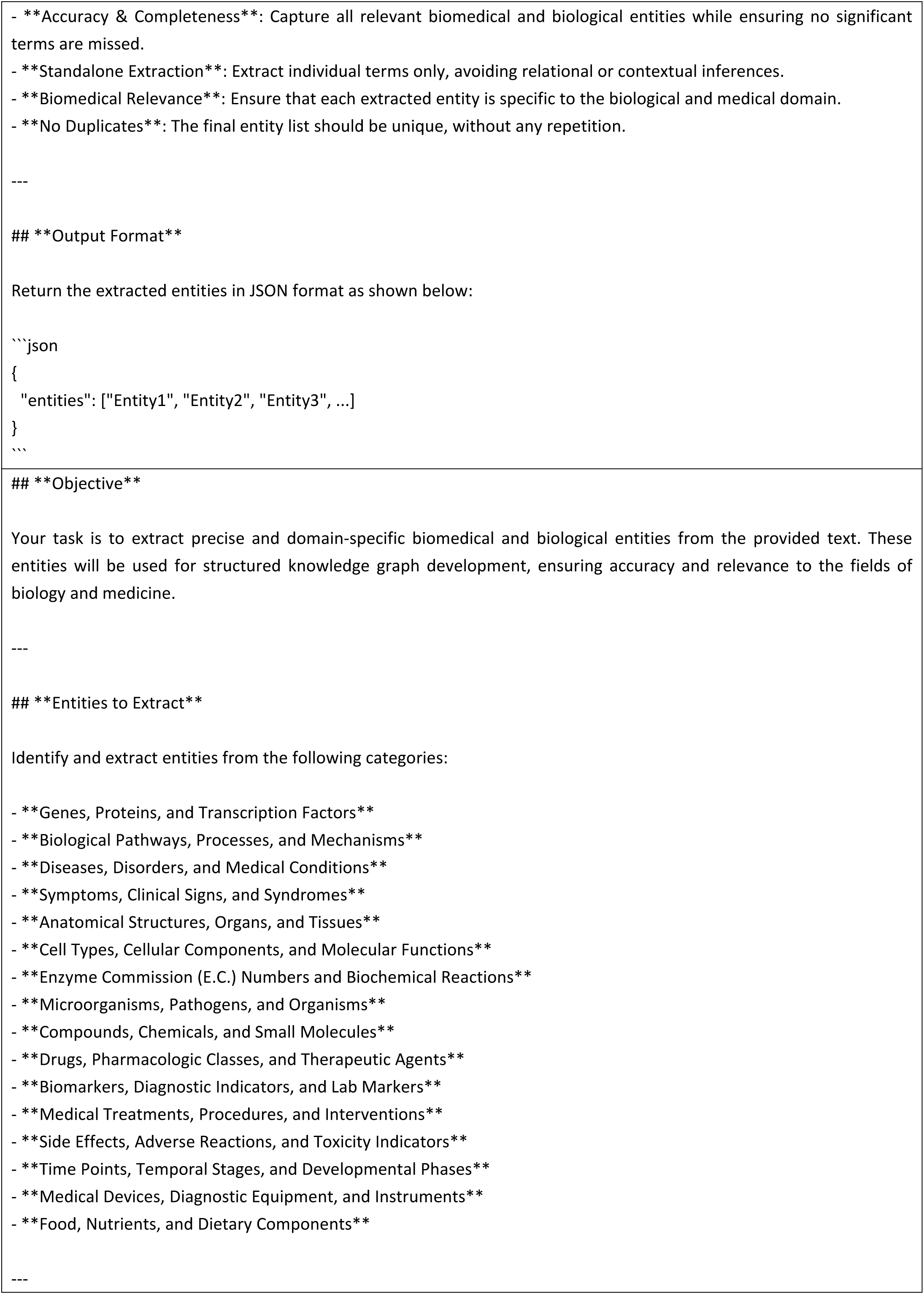
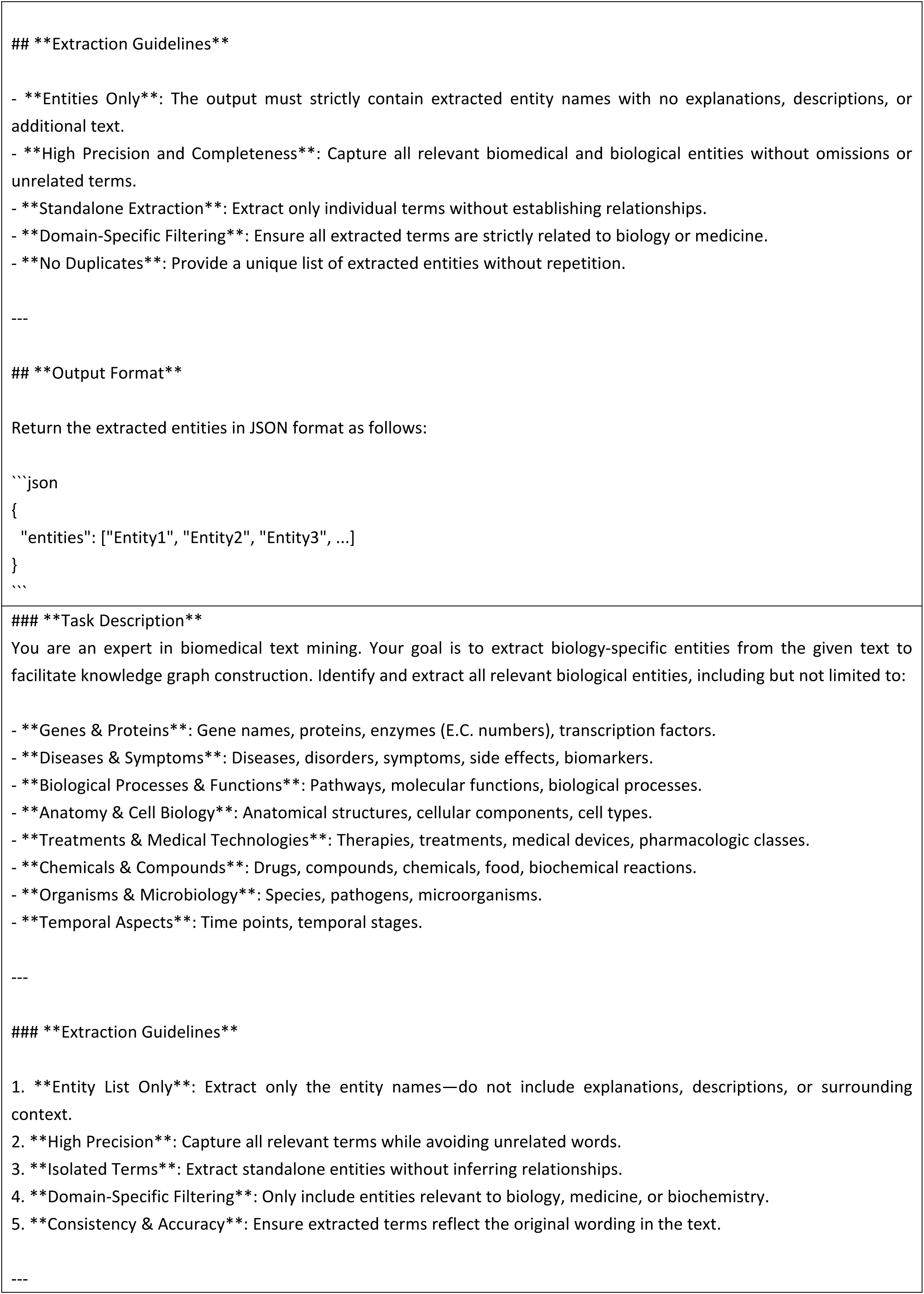
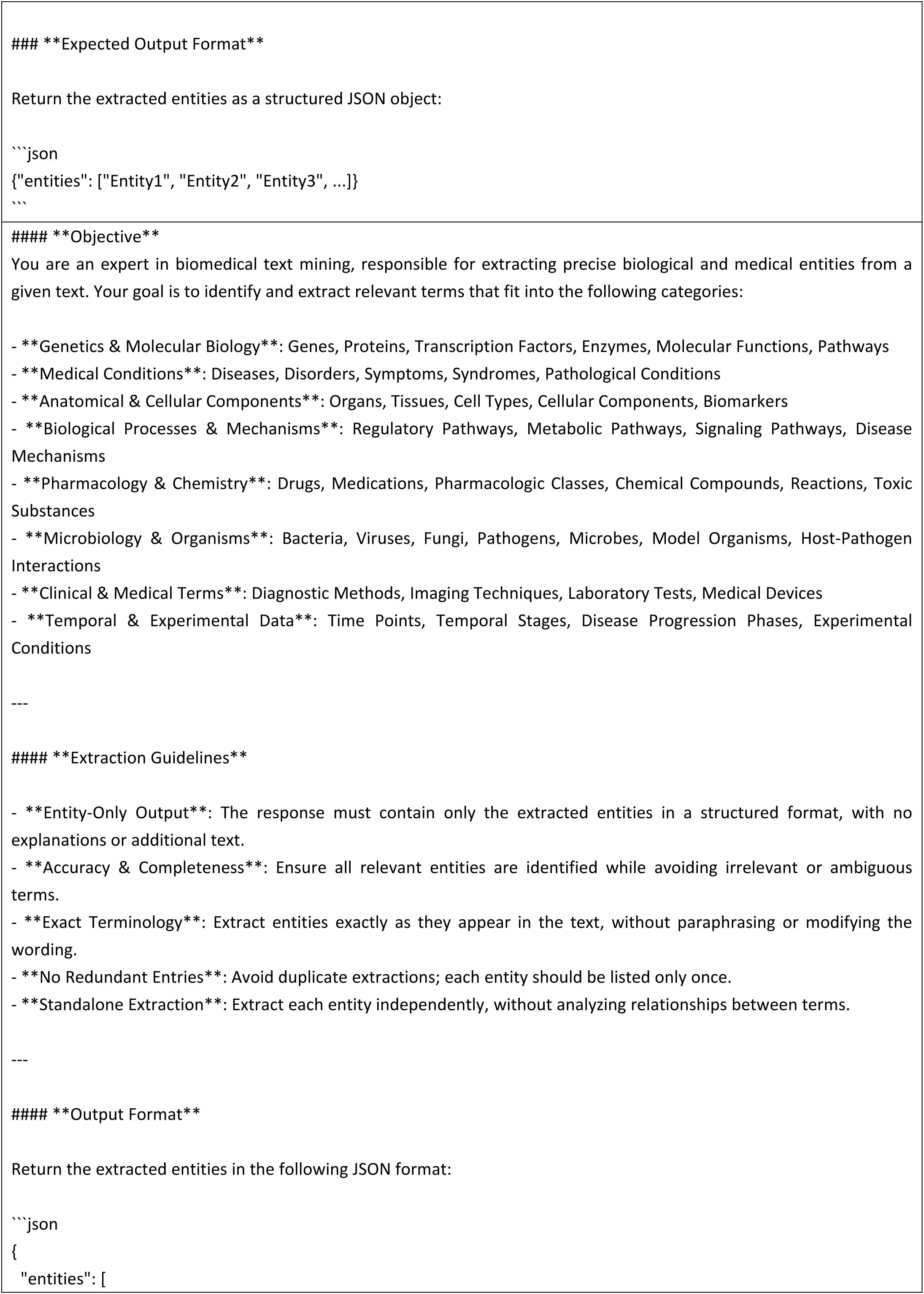
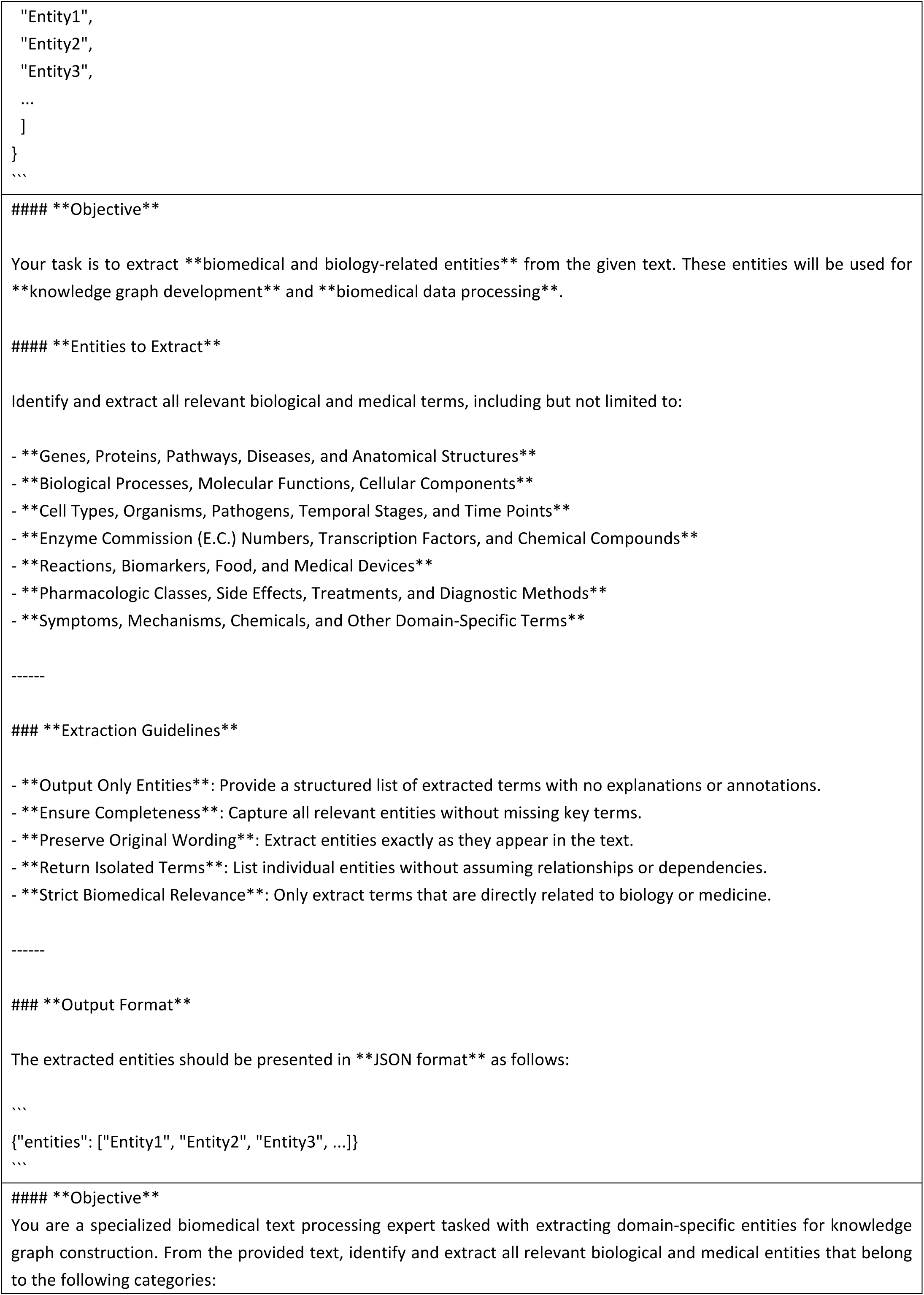
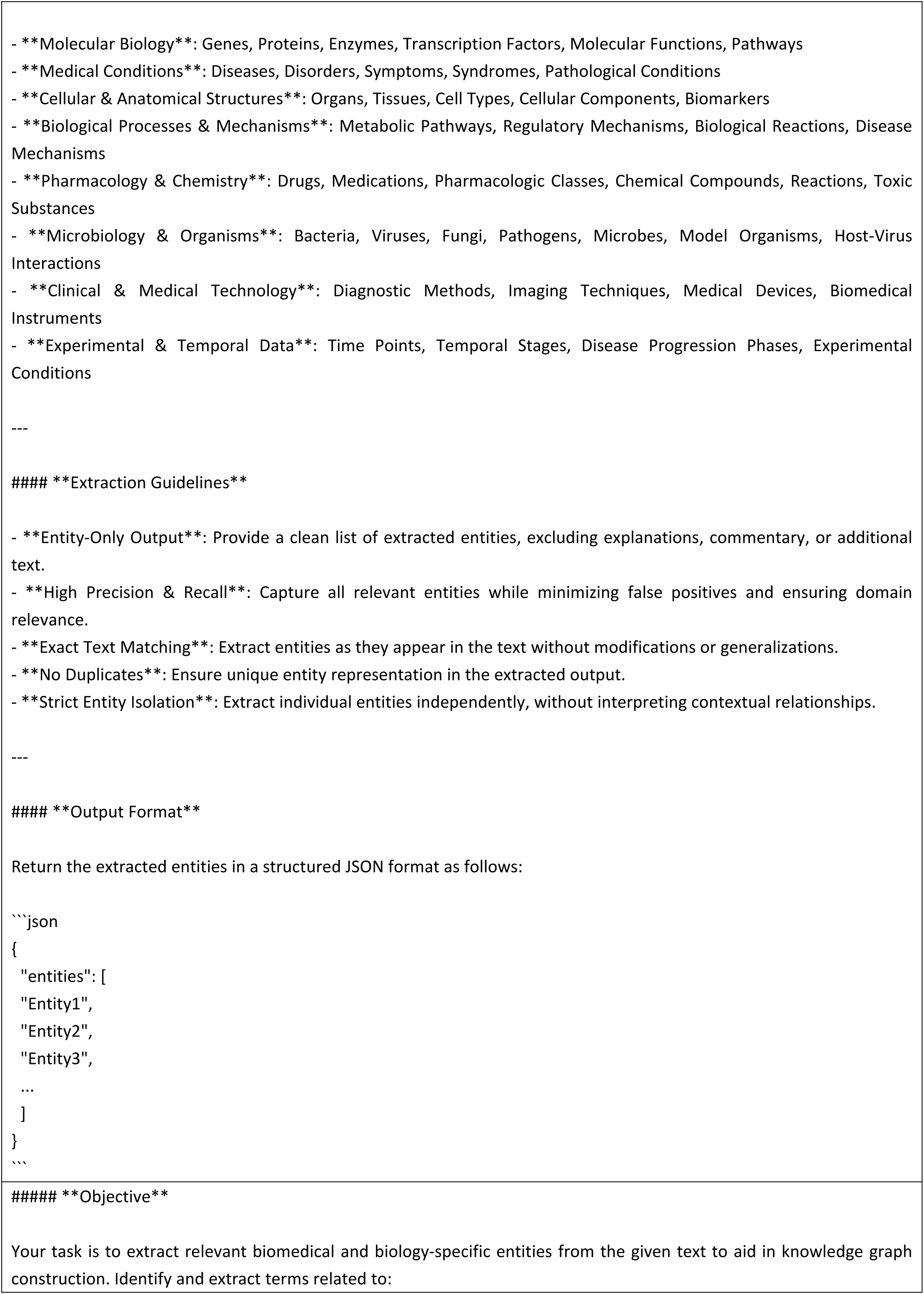
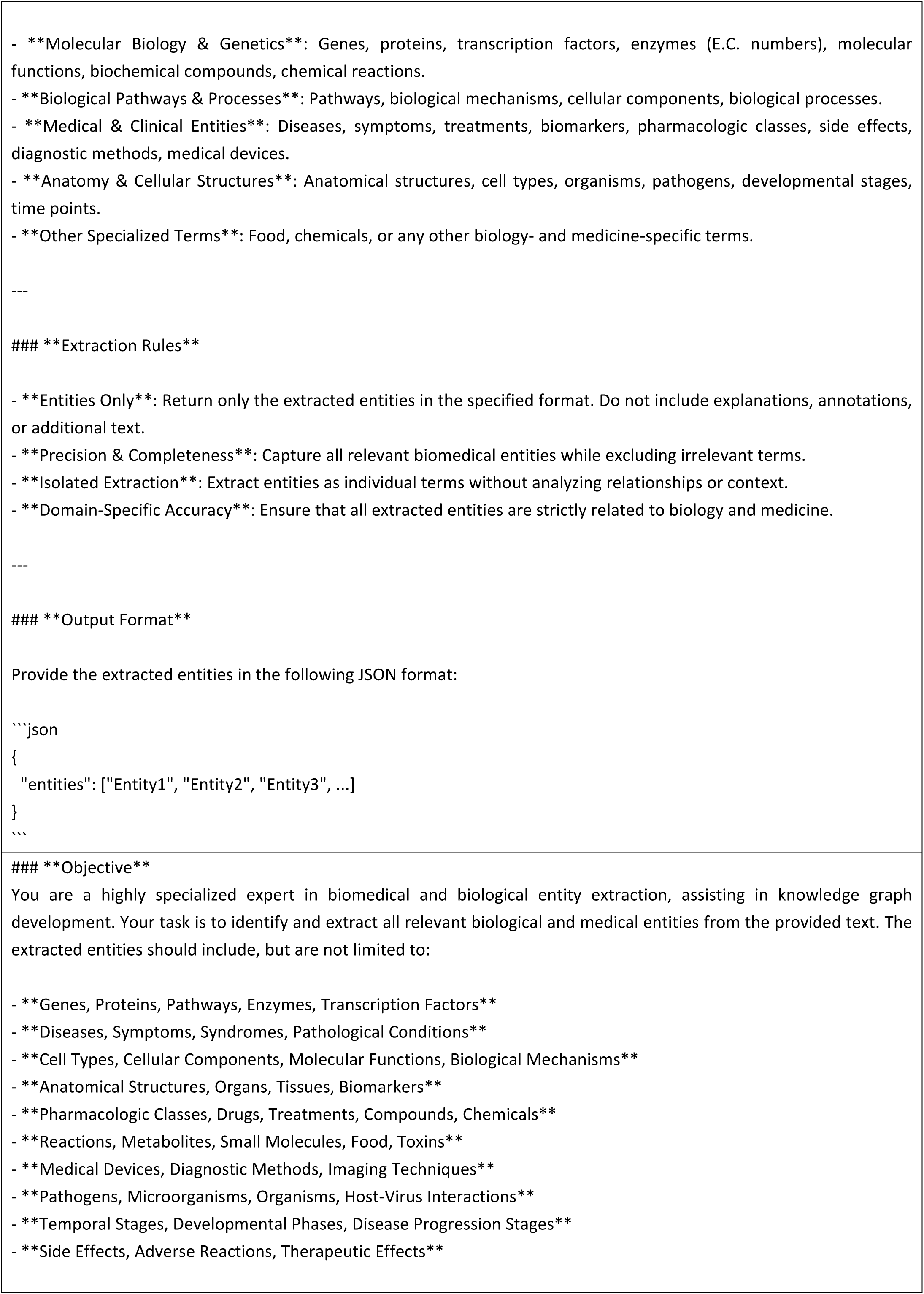
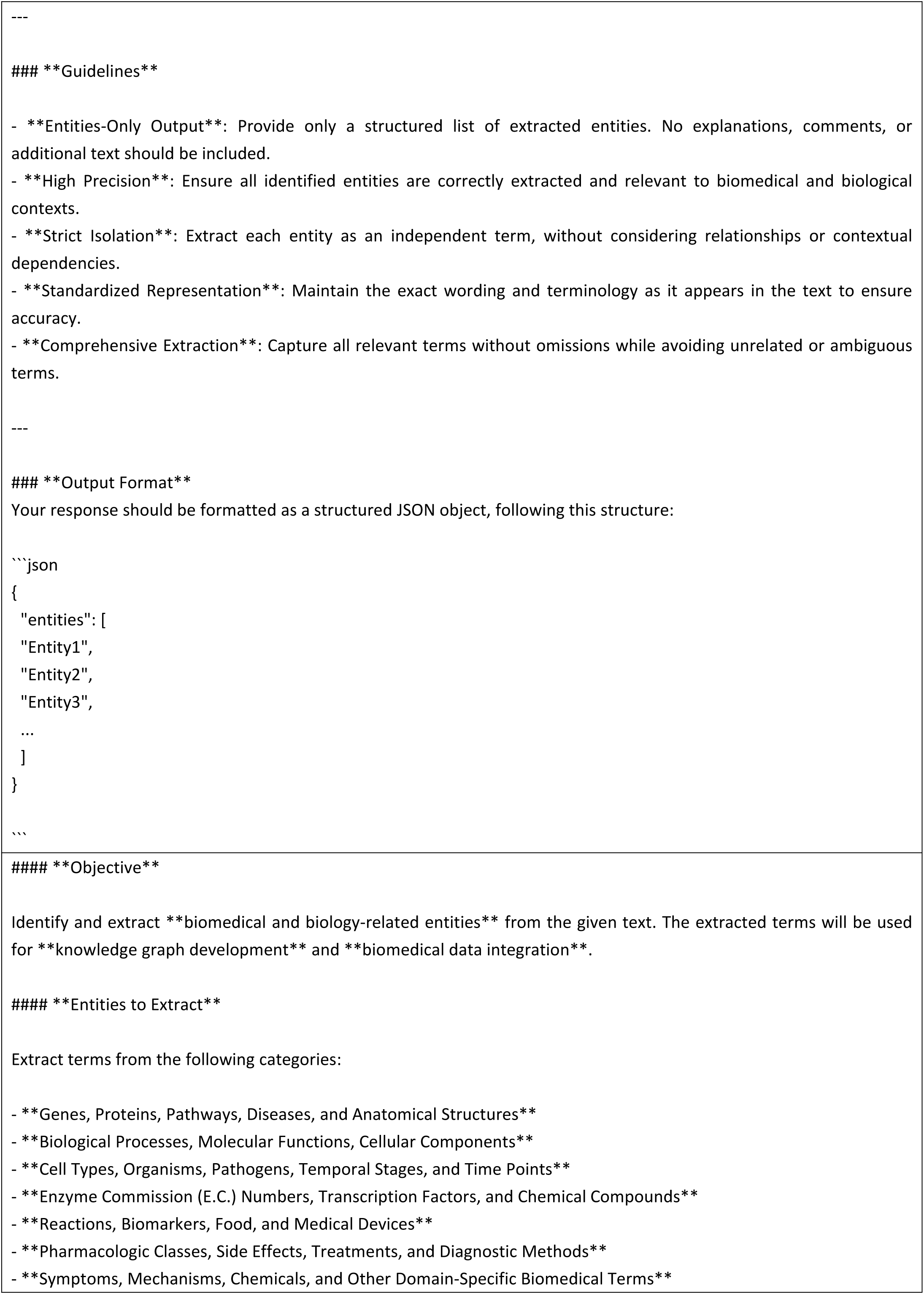
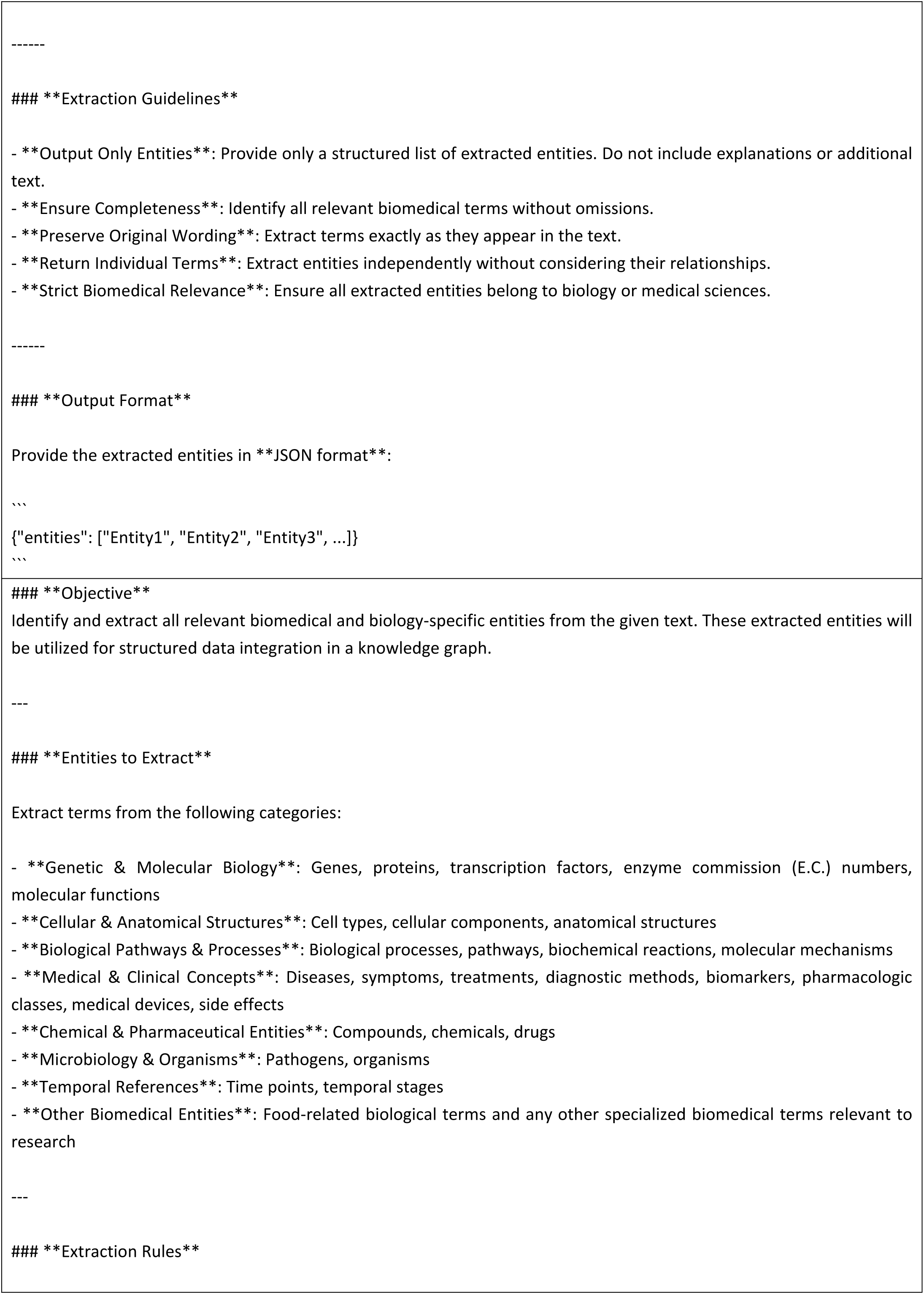
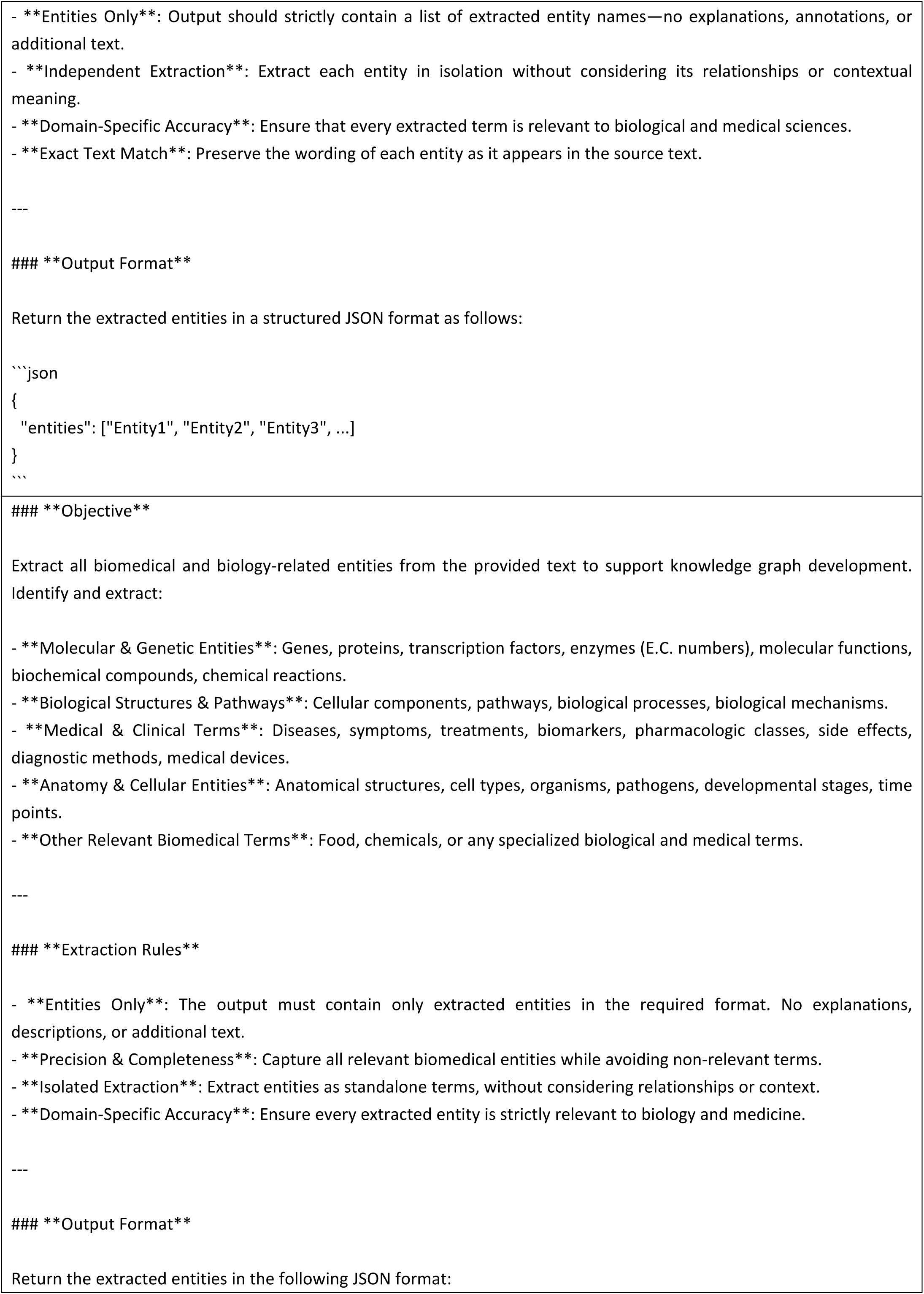
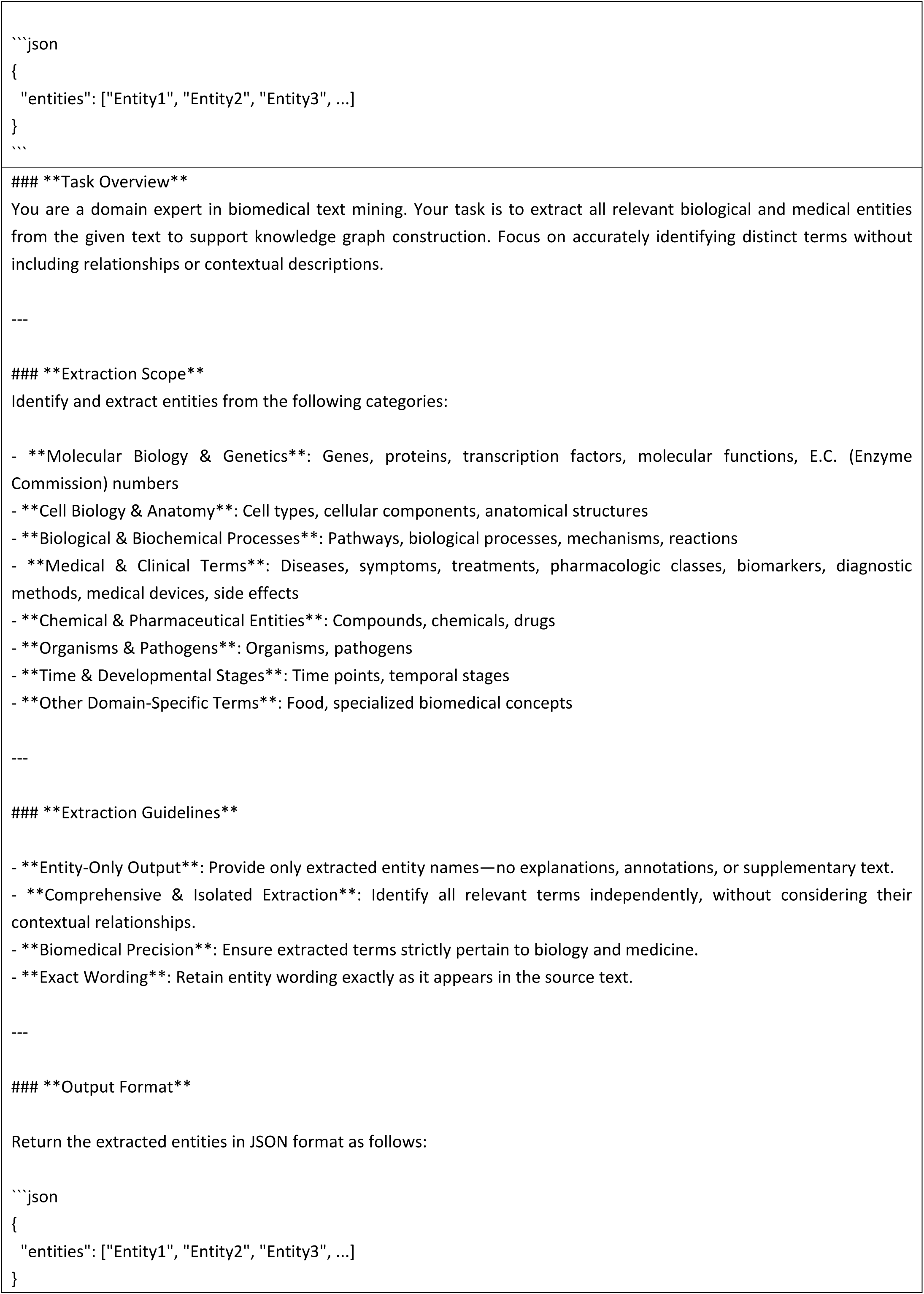
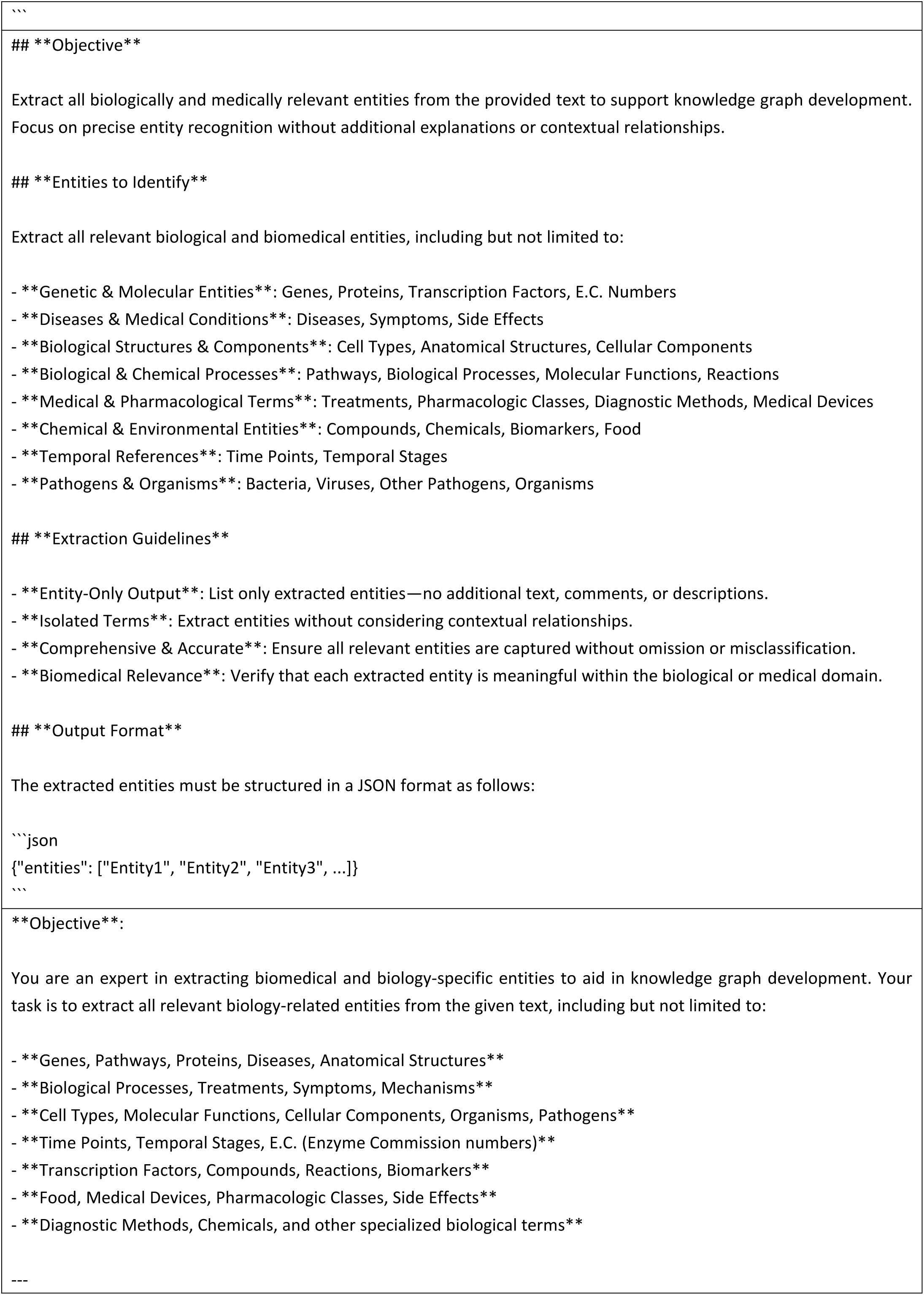
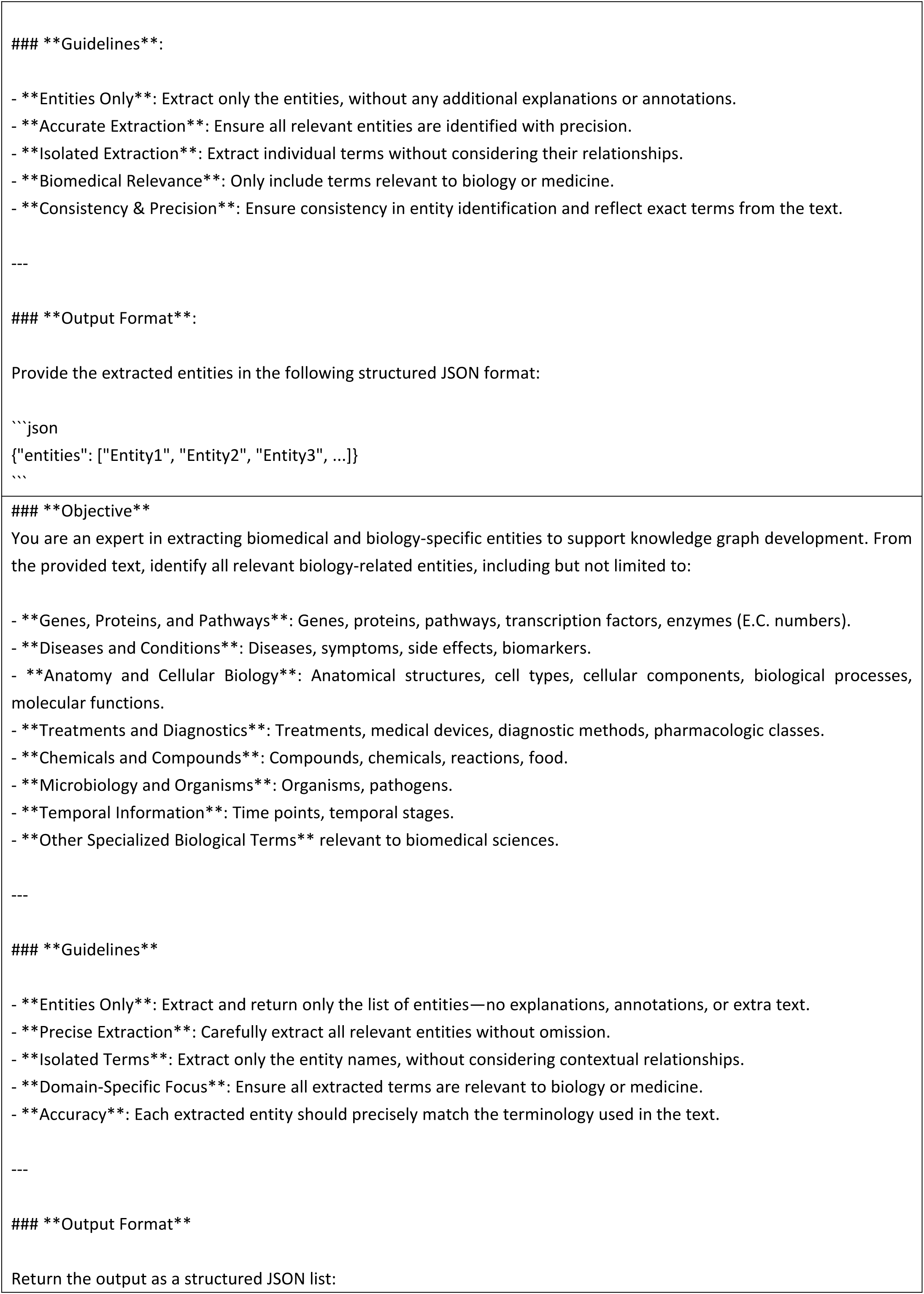
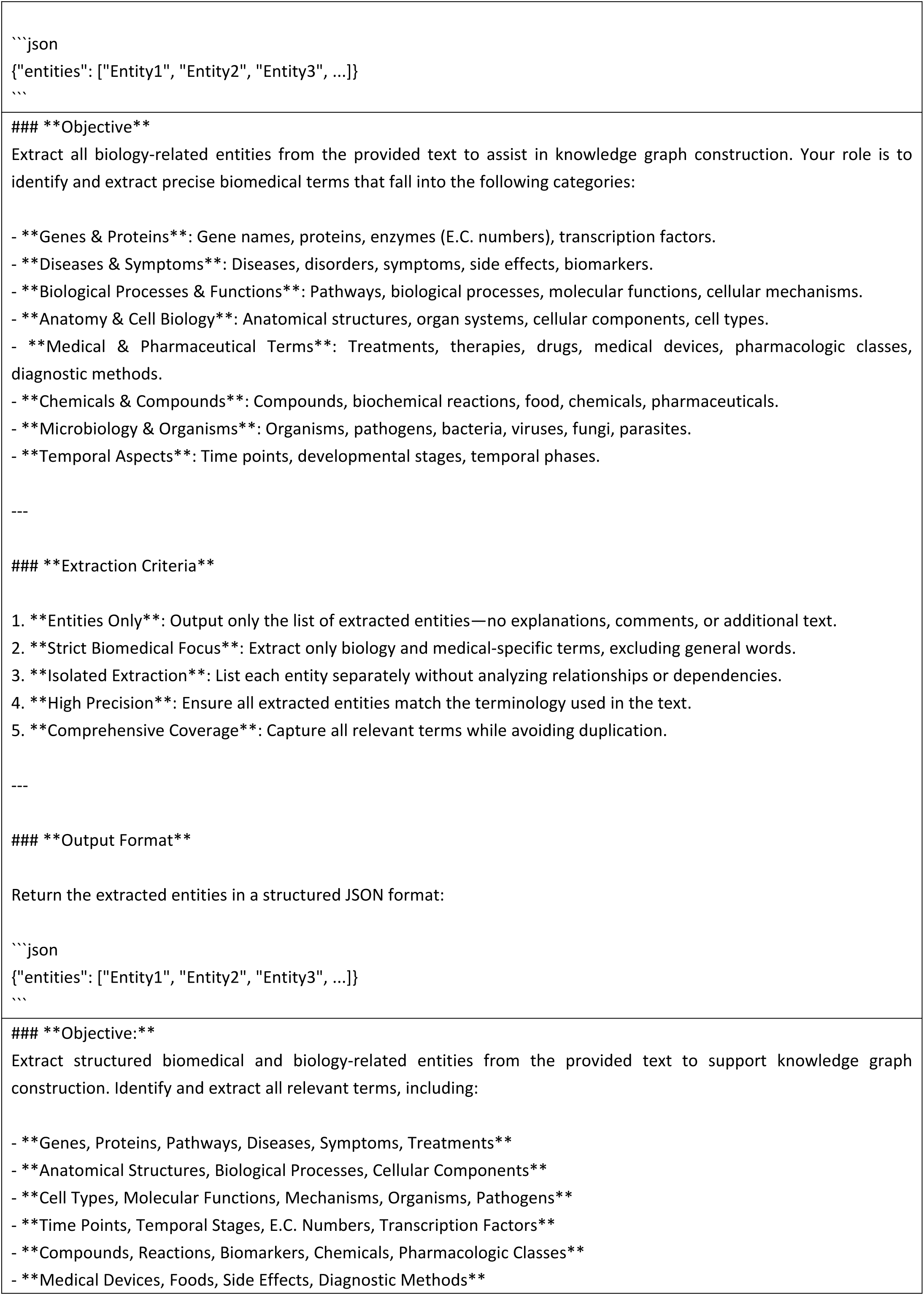
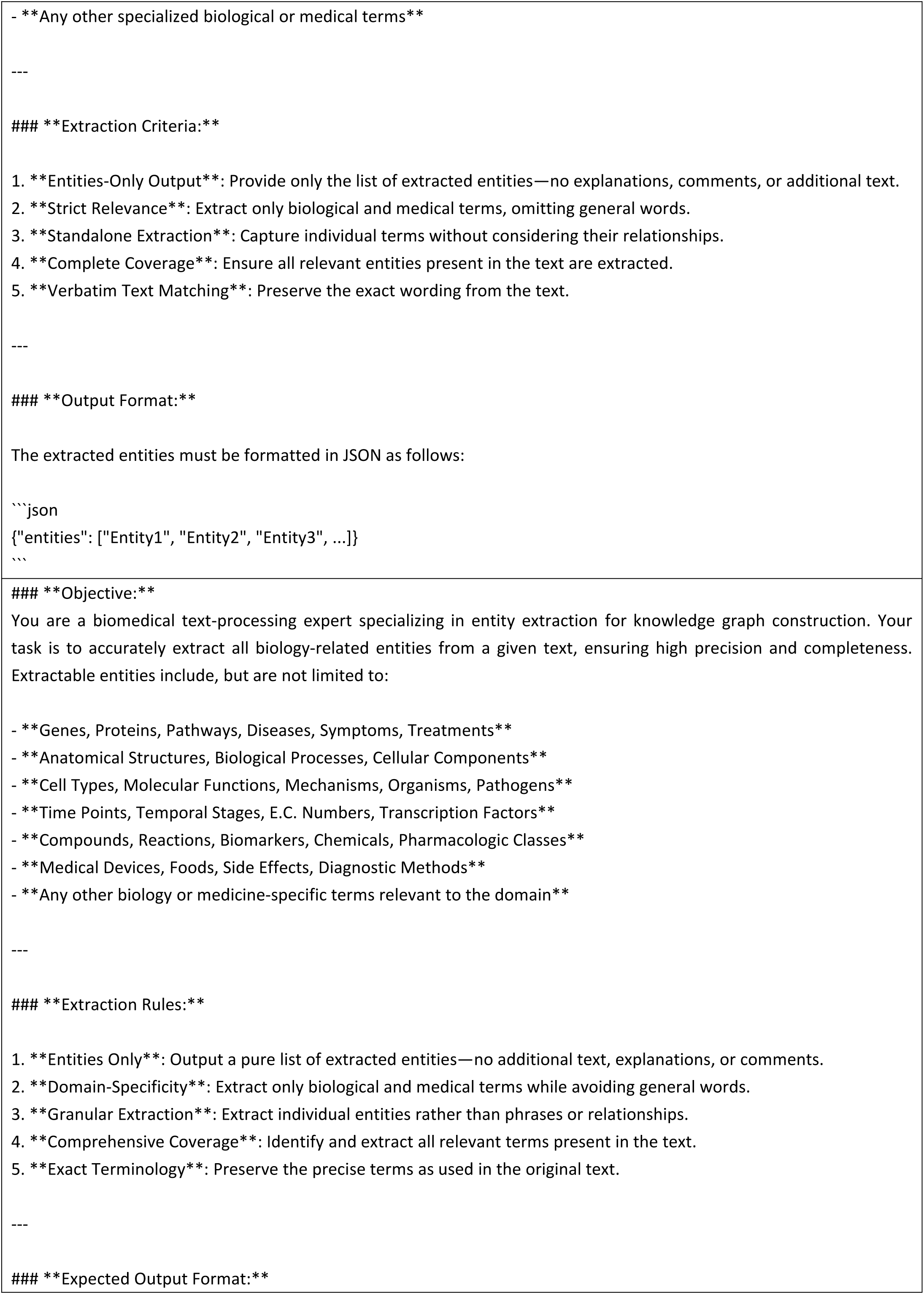
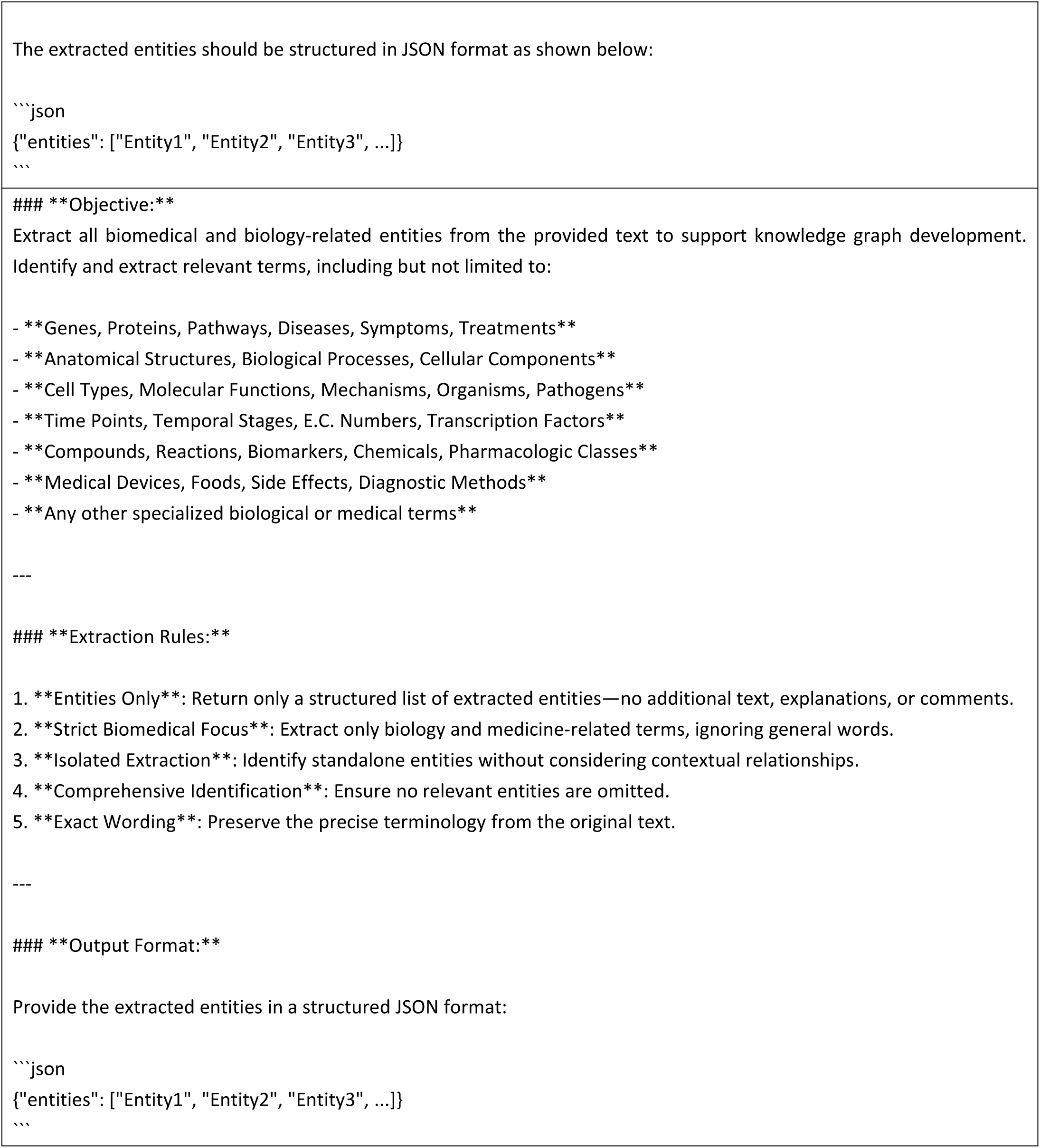
Prompts for image-entity extraction tests.

**Supplementary Table 18.**
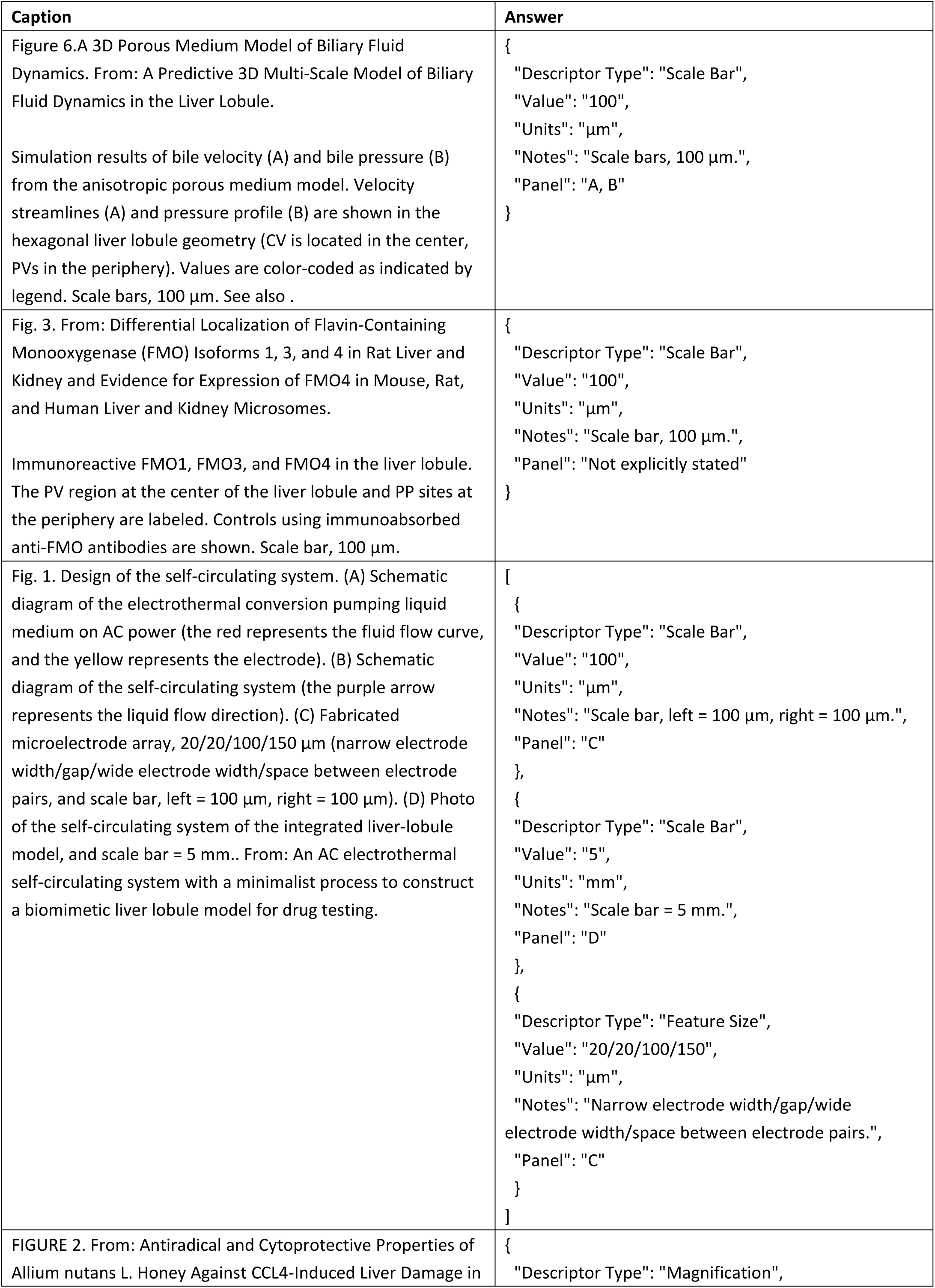
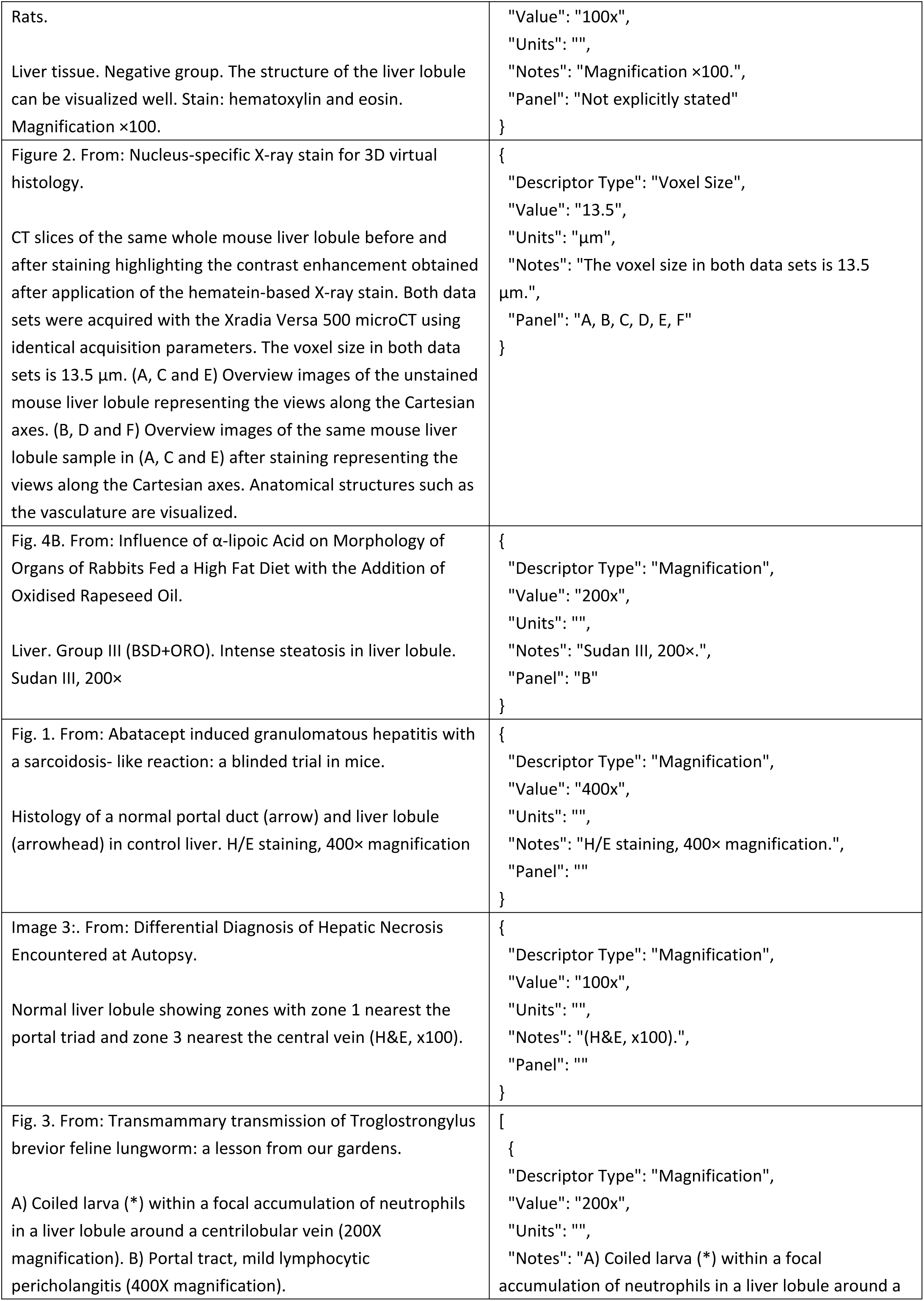
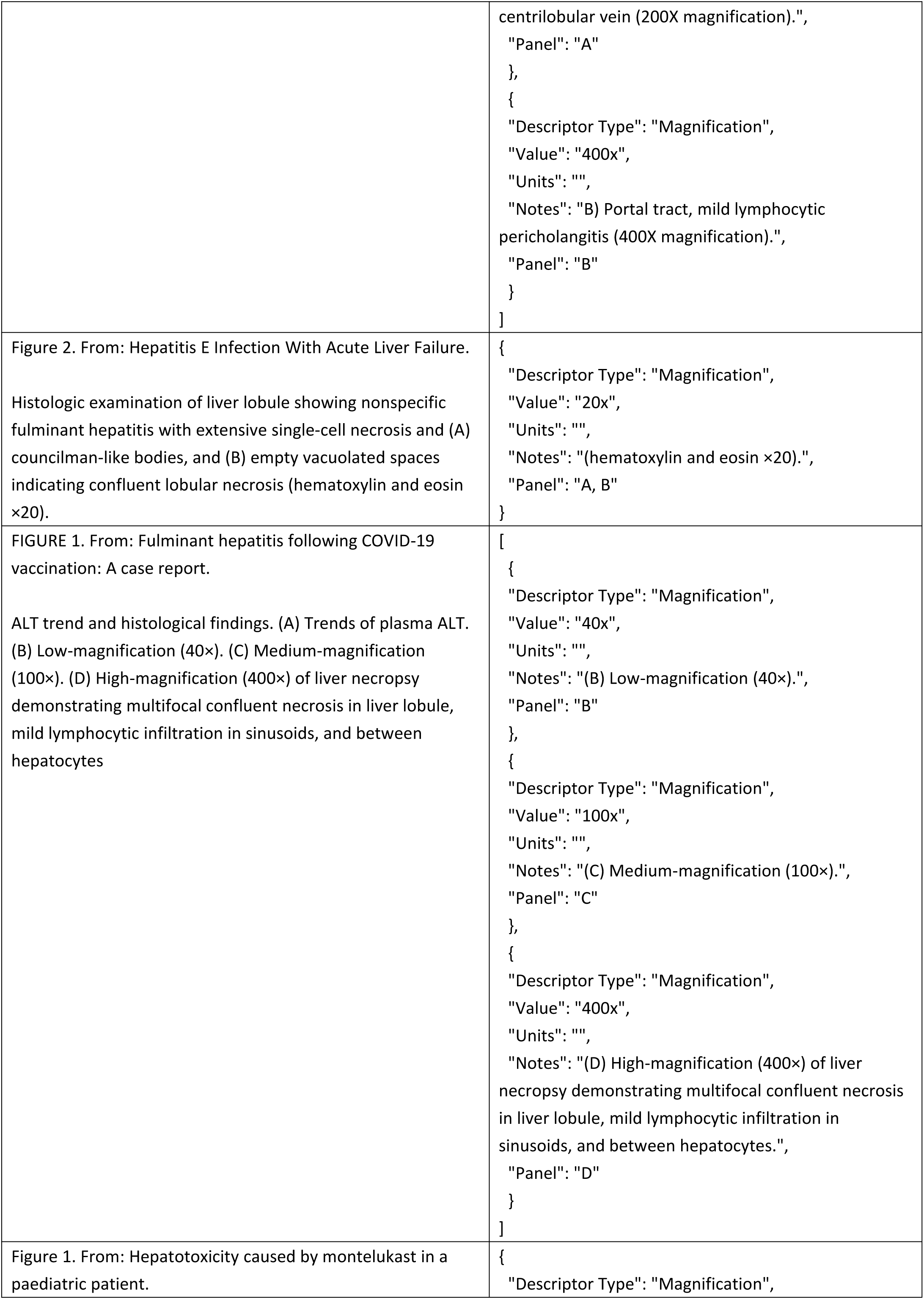
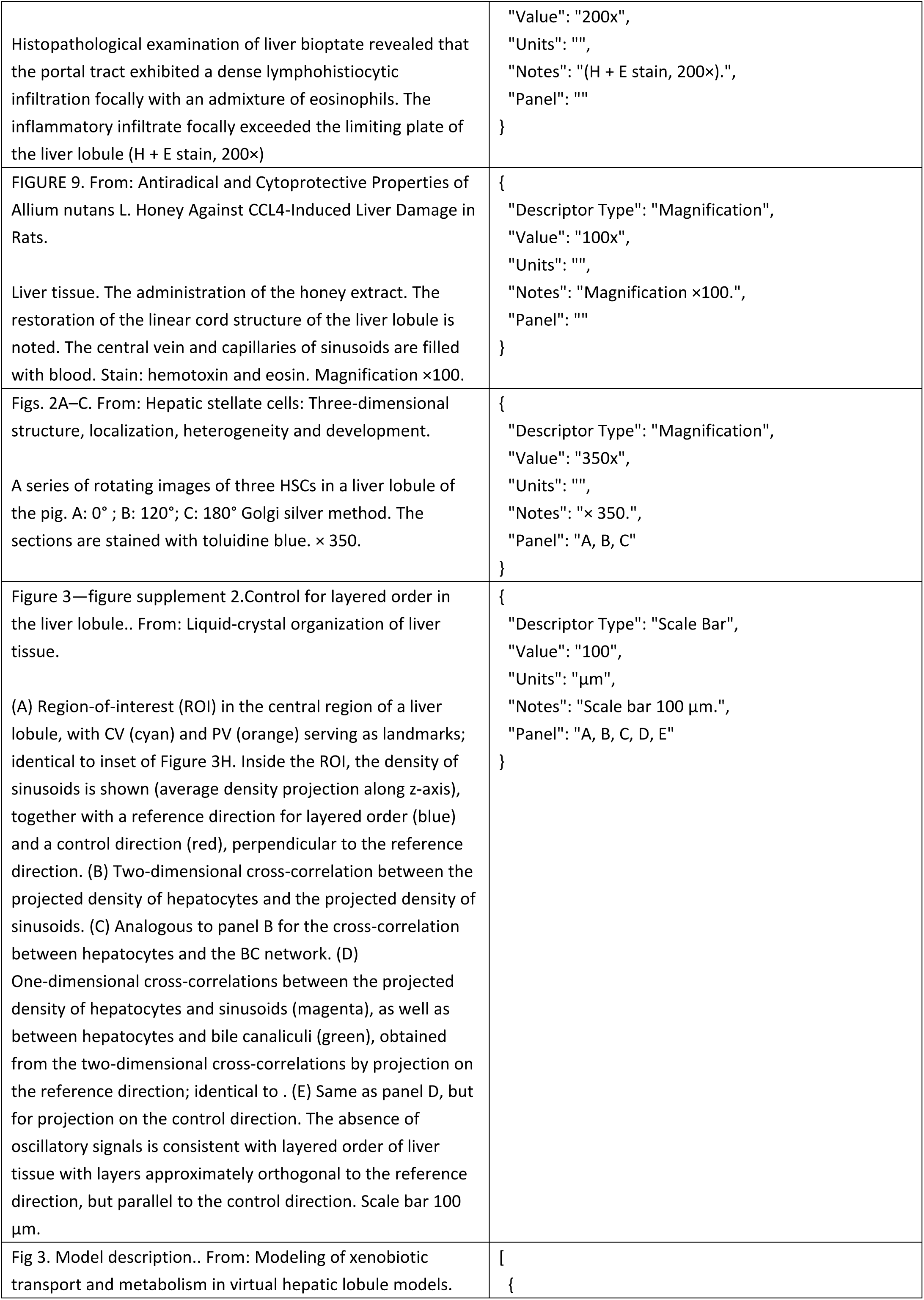
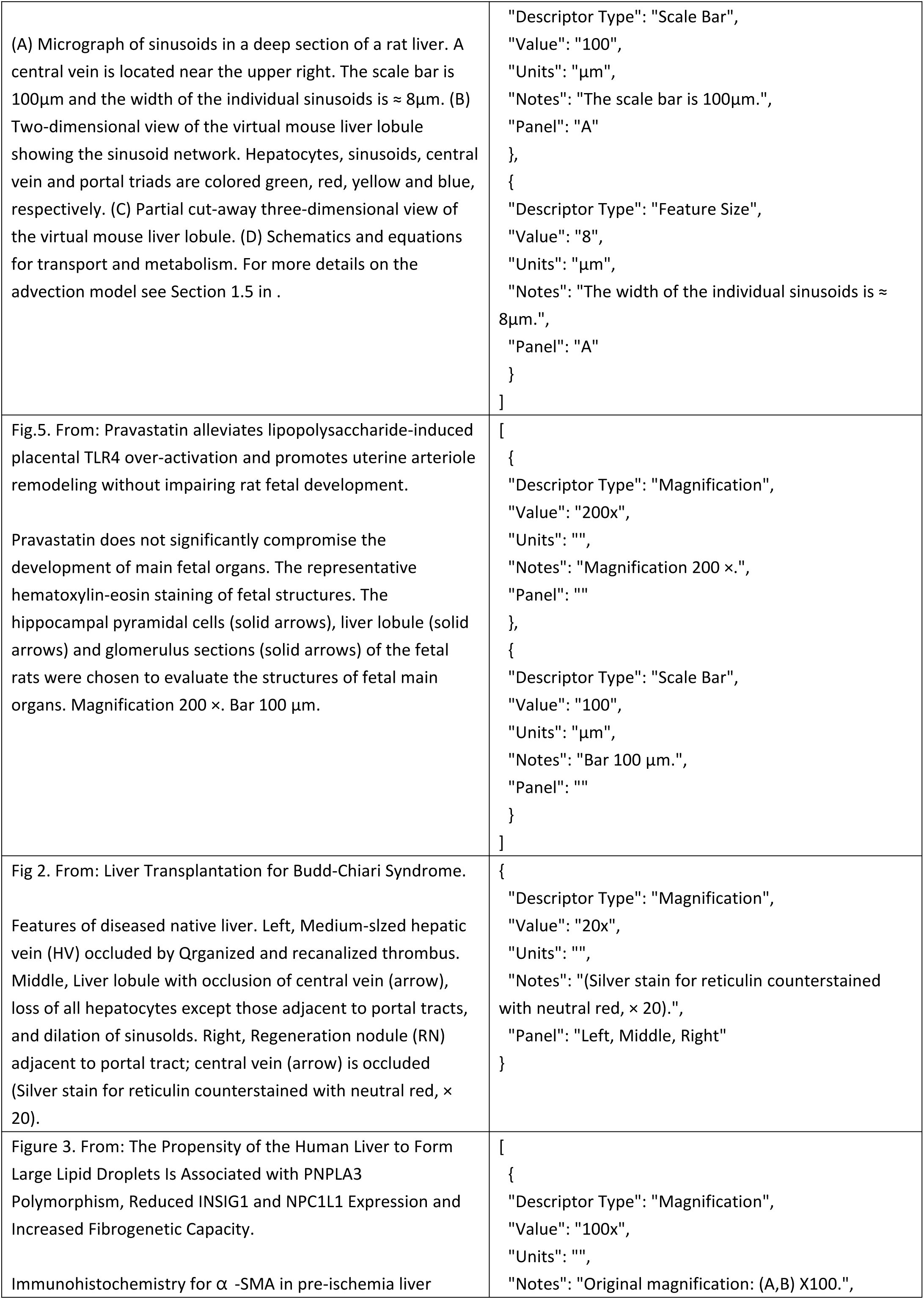
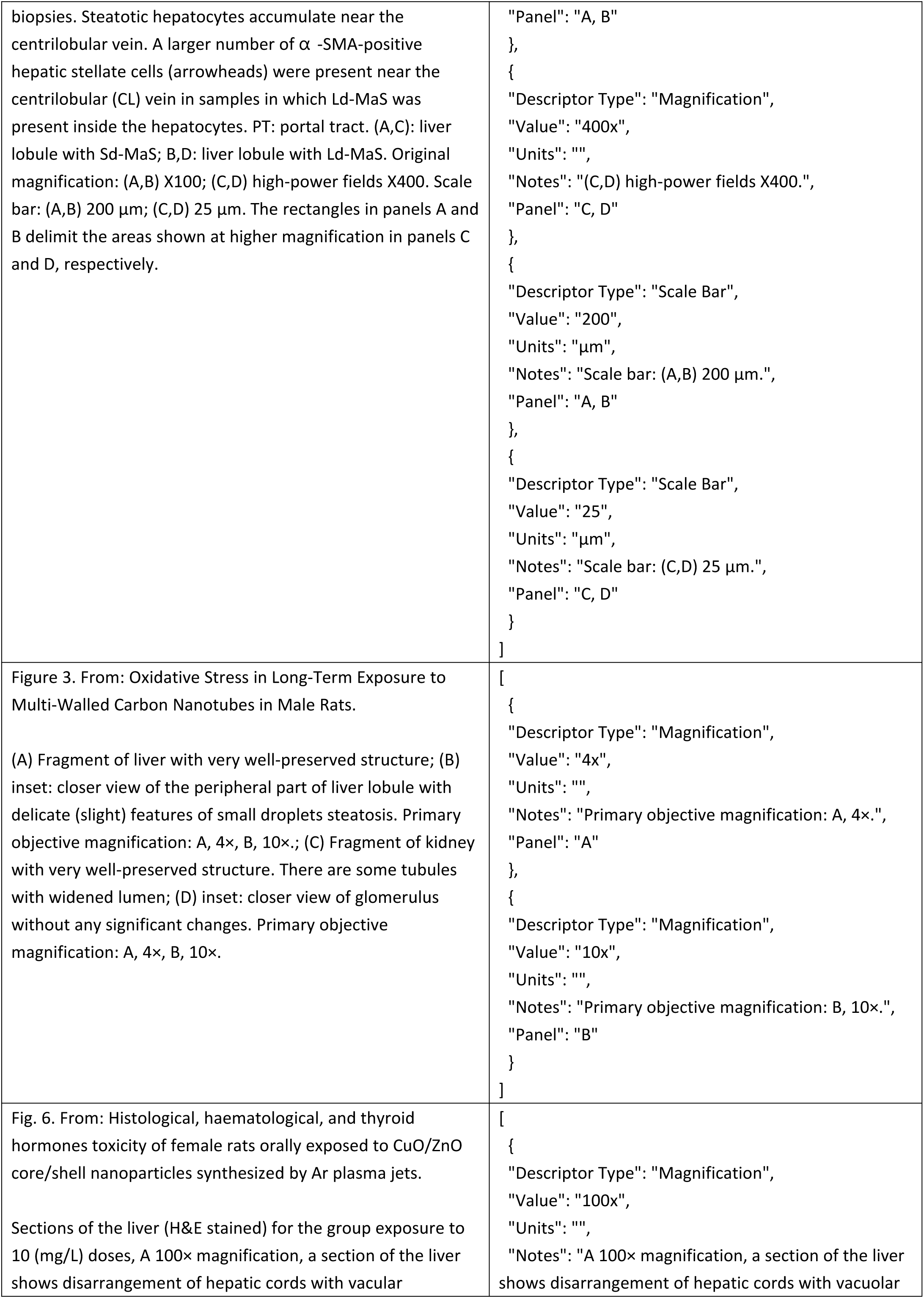
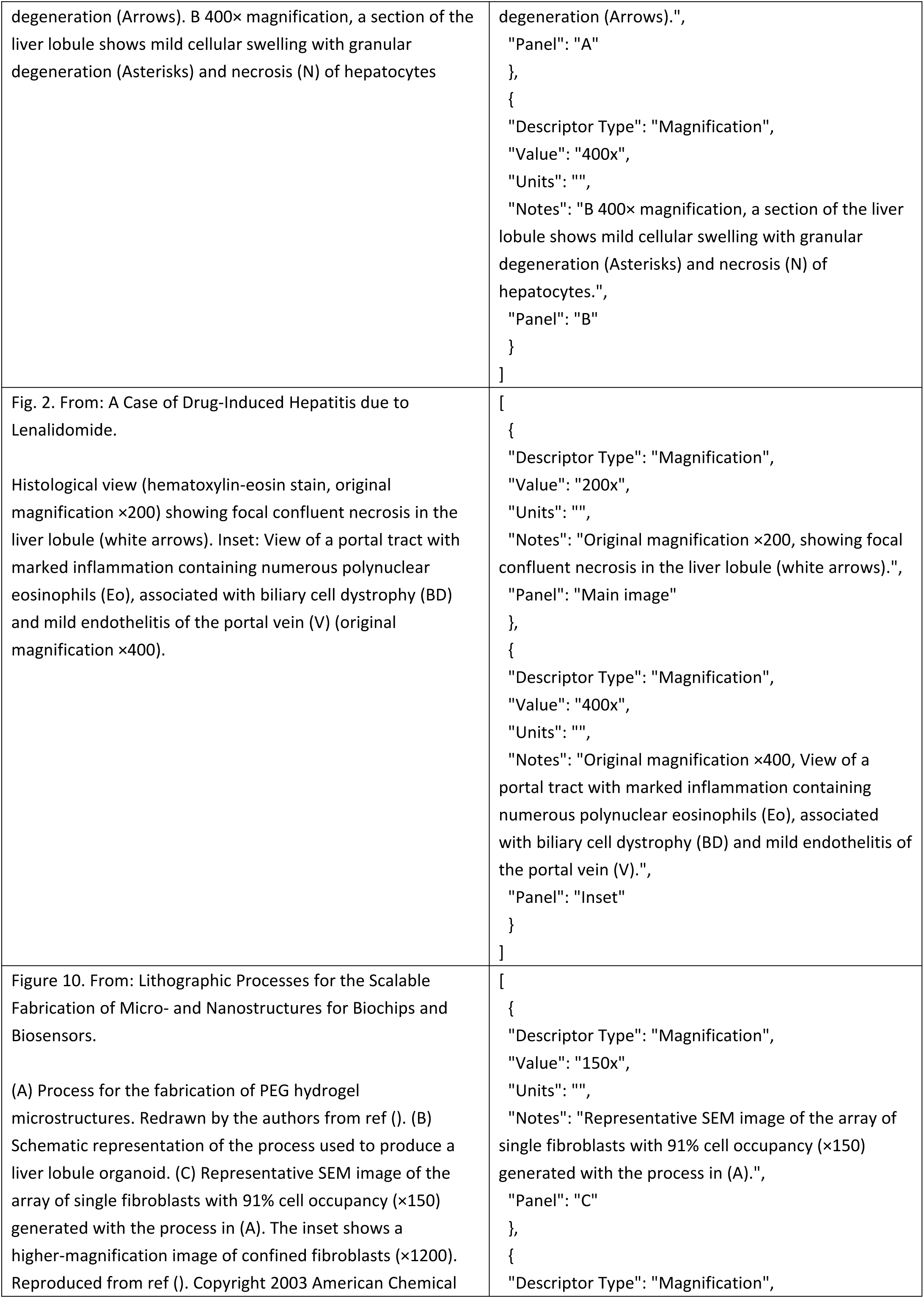
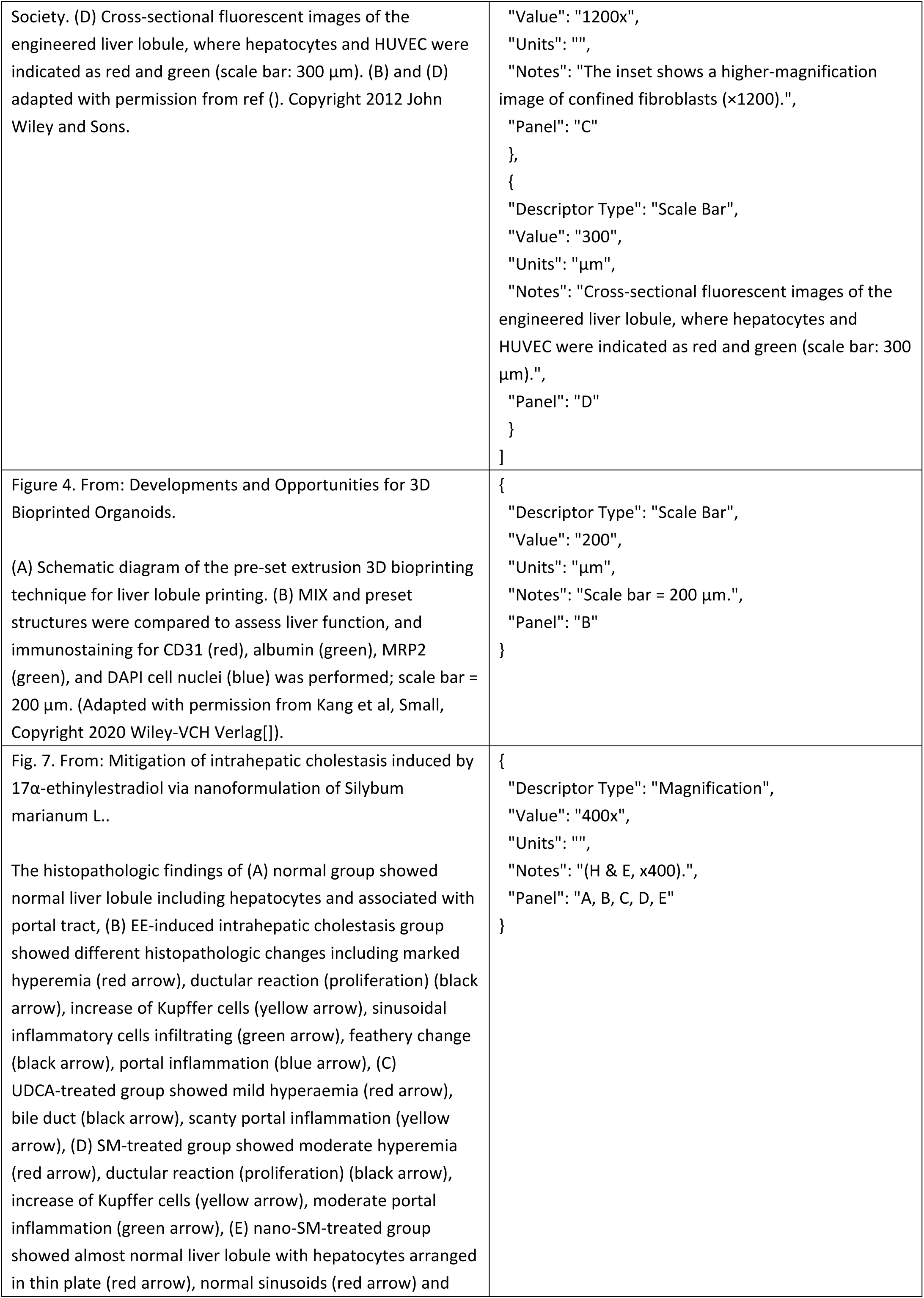
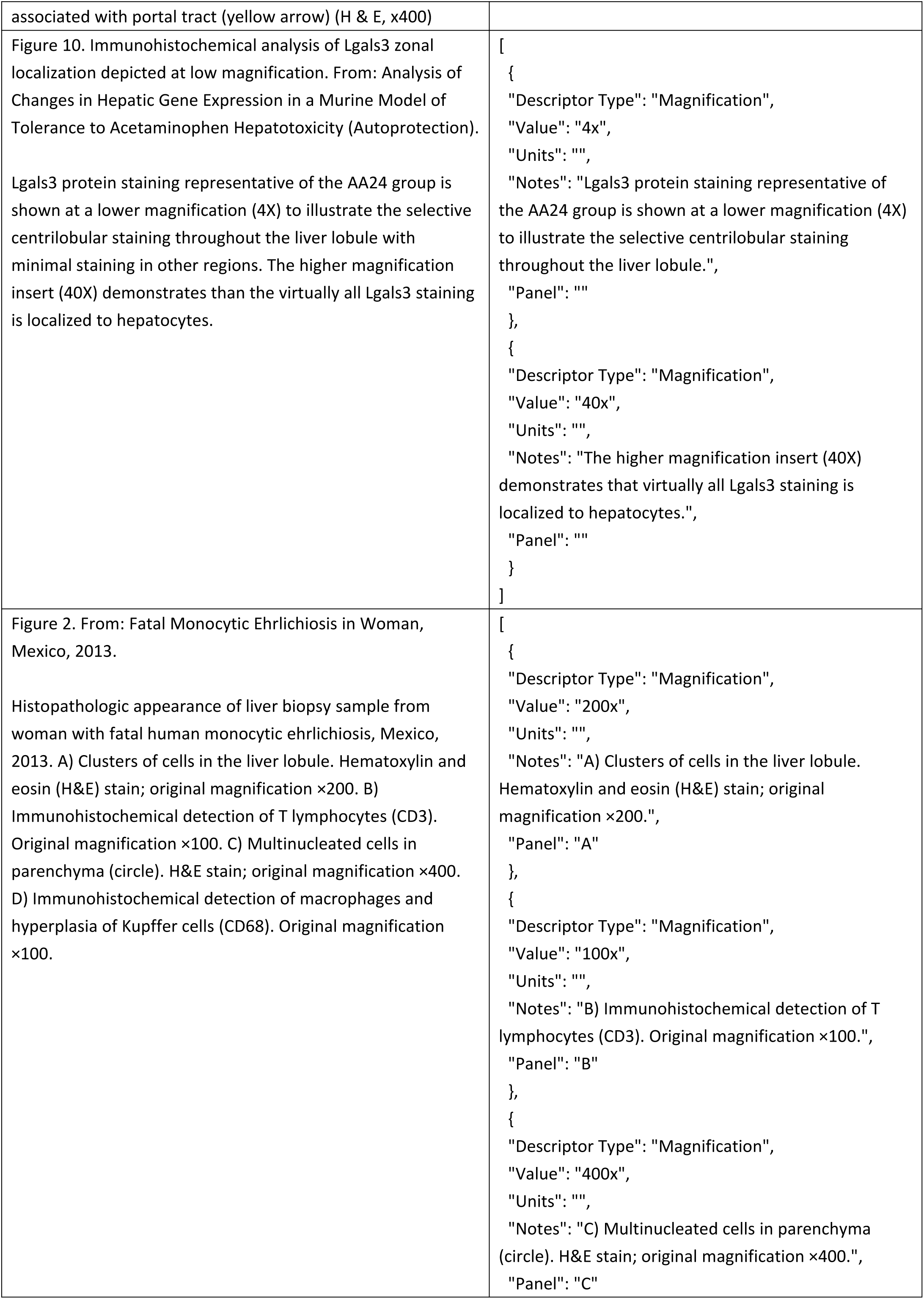
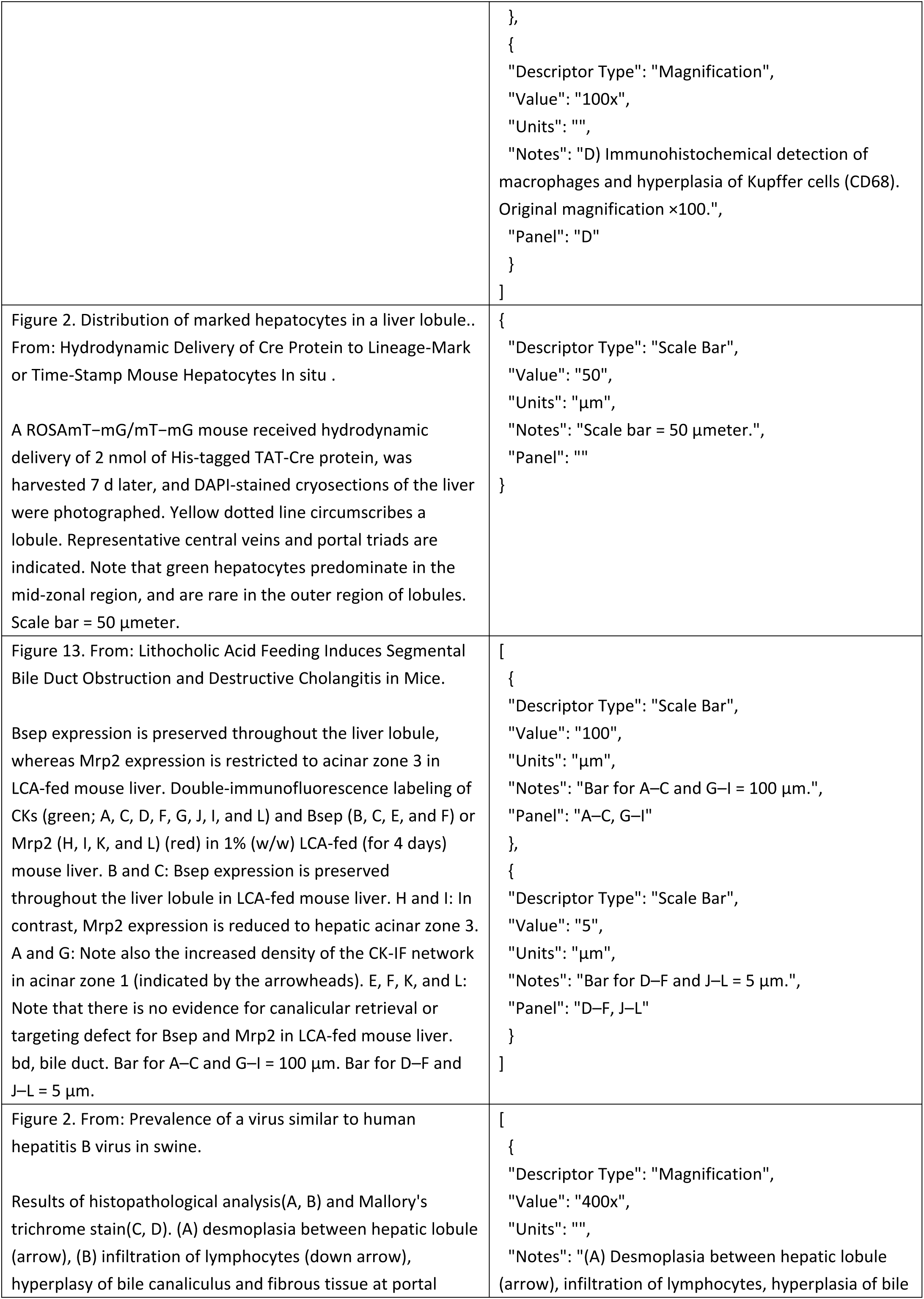
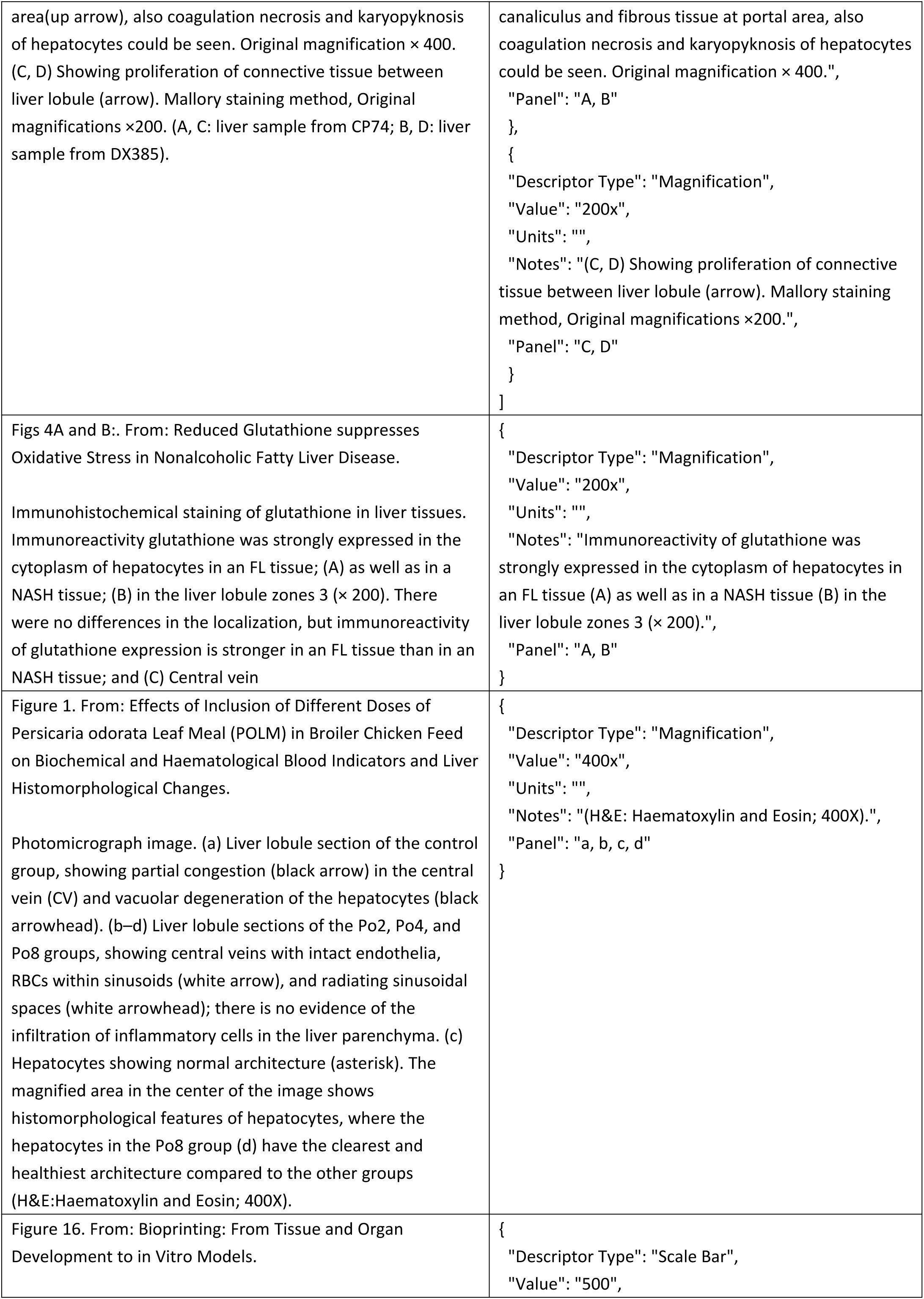
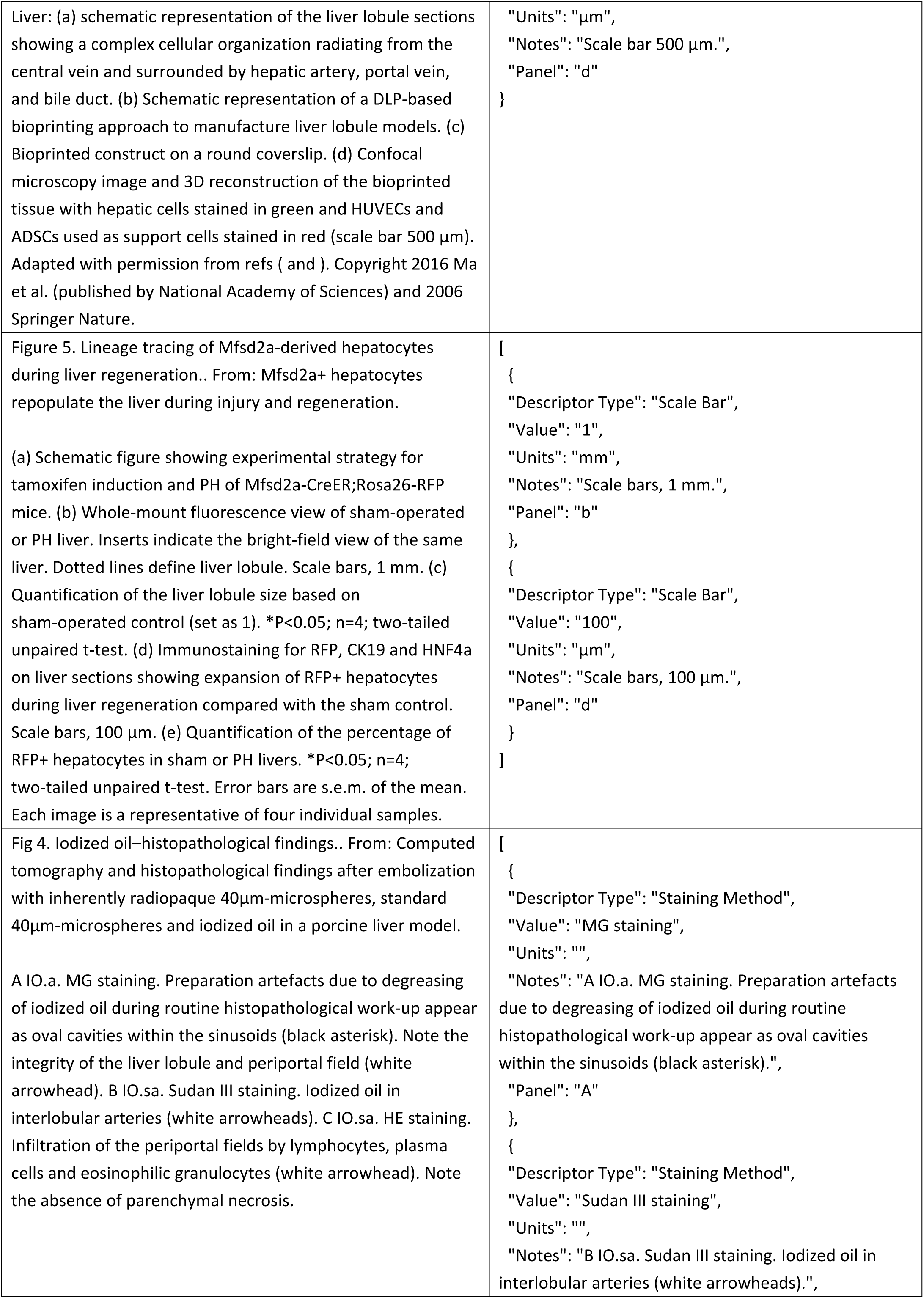
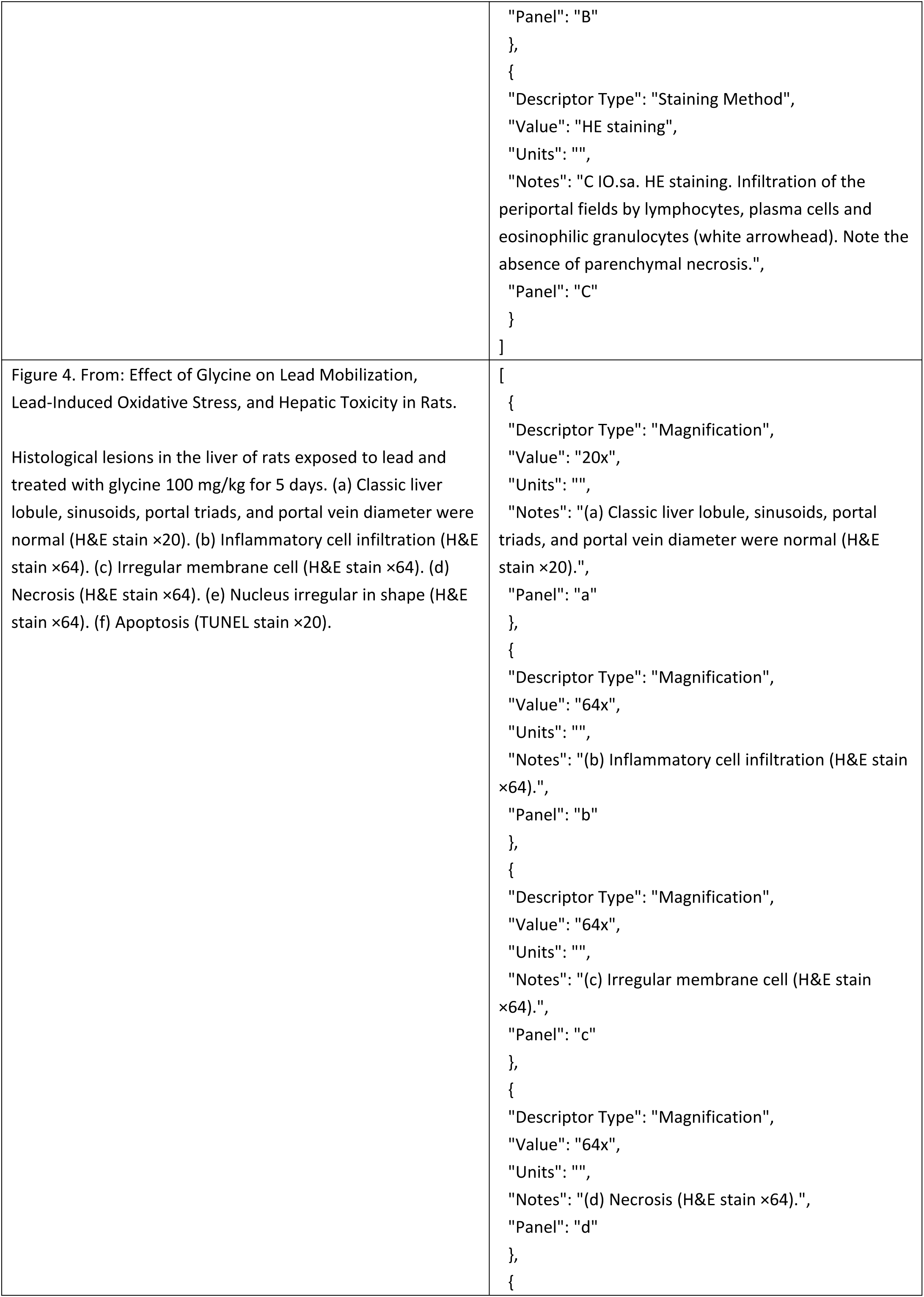
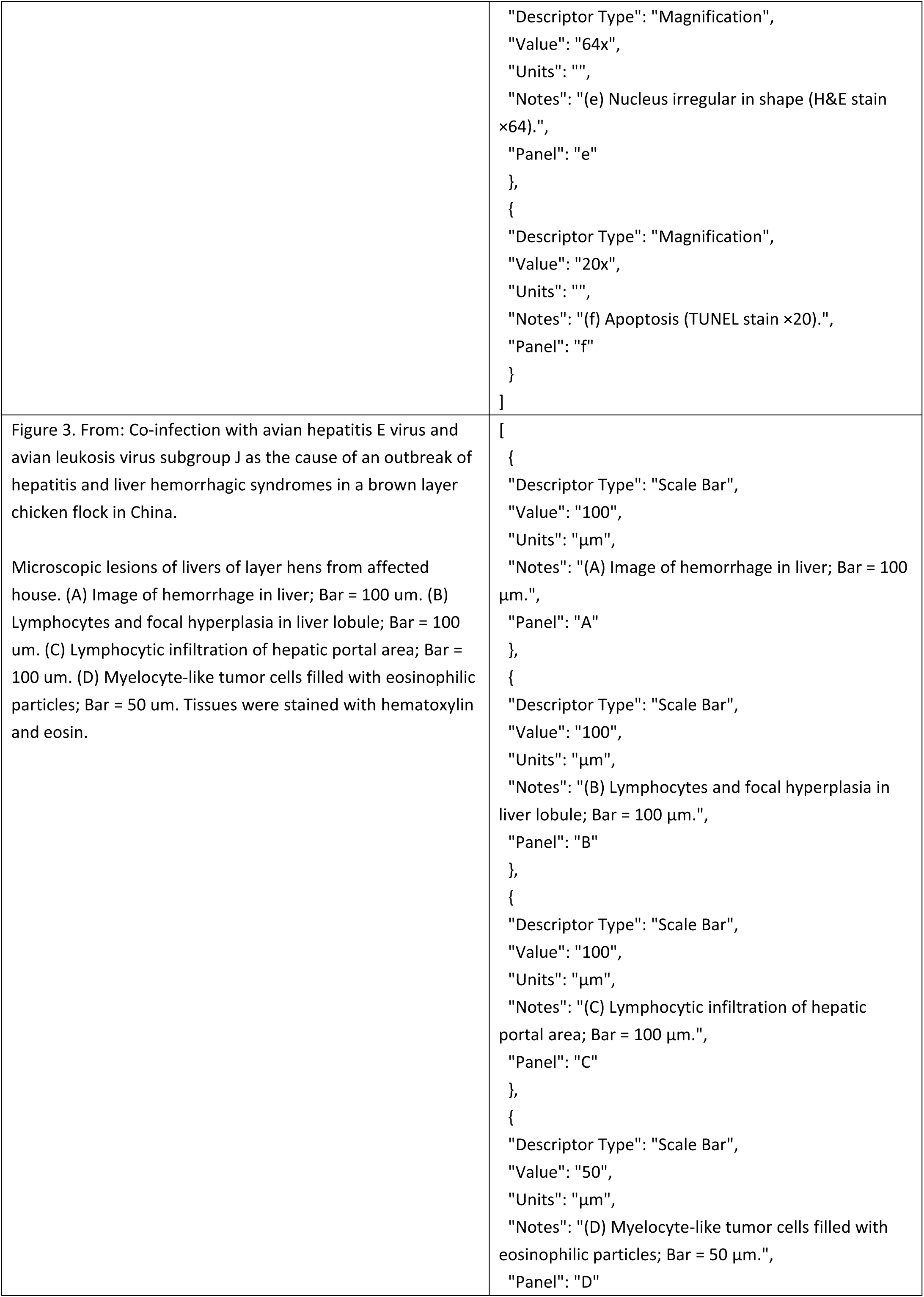
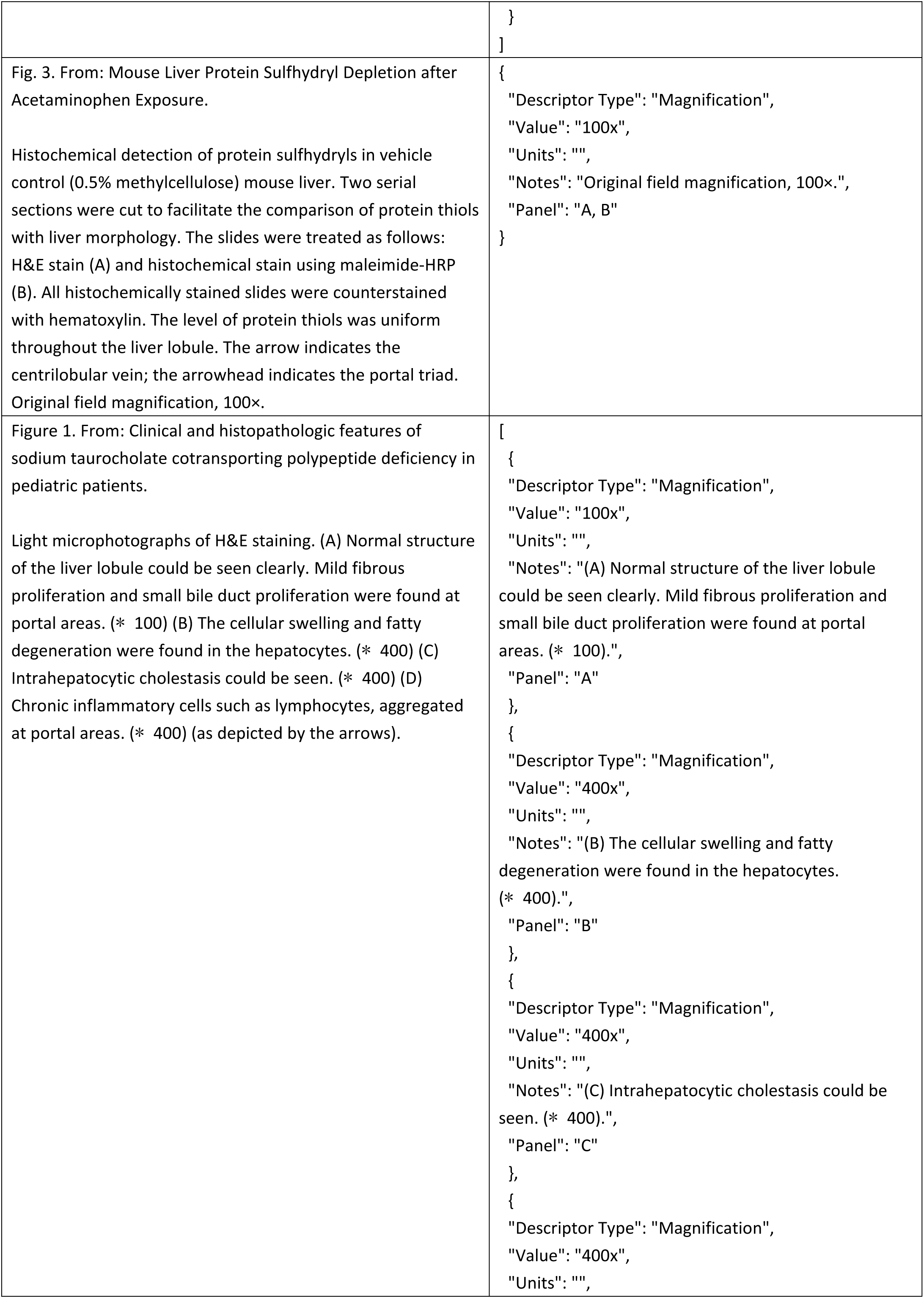
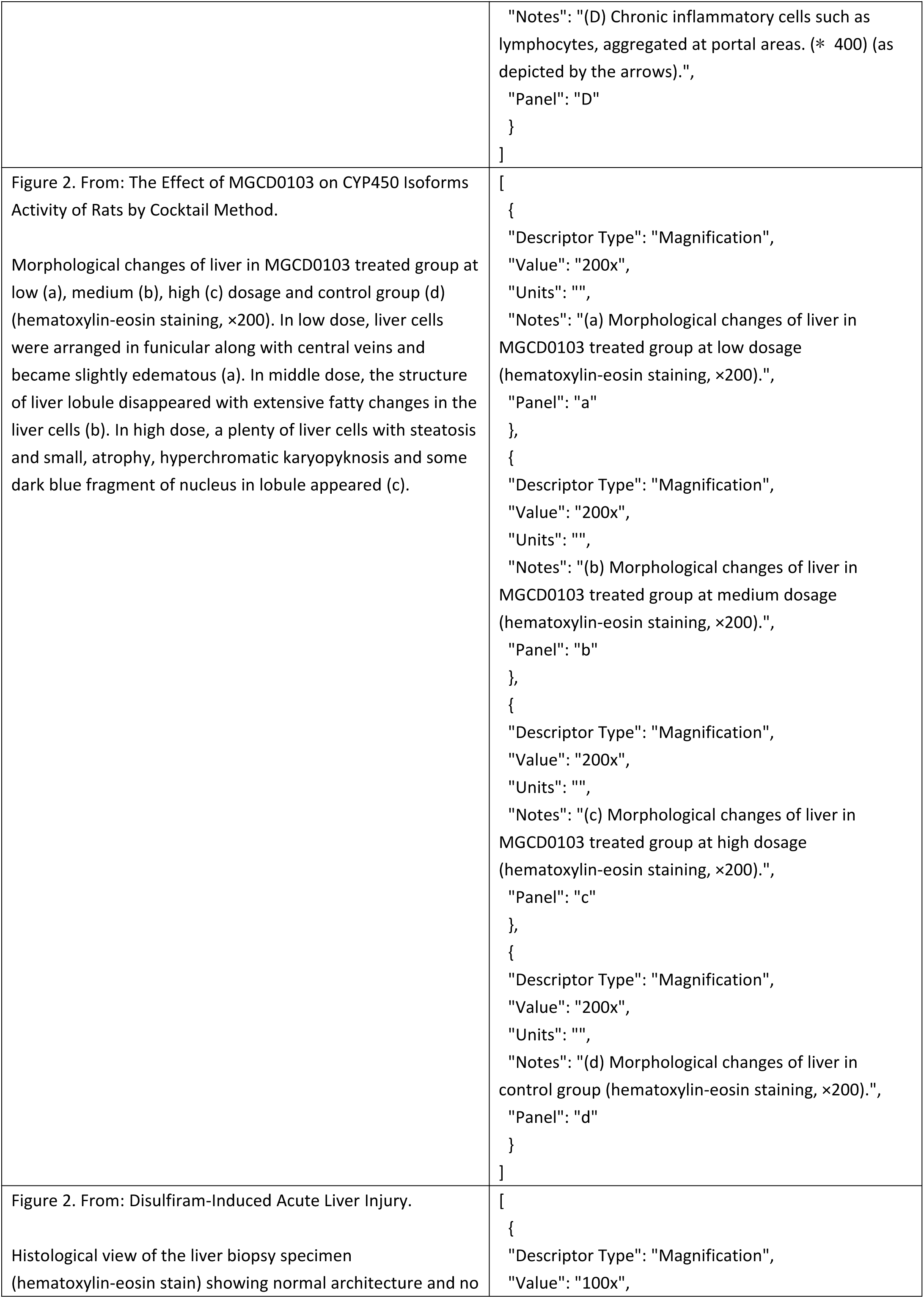
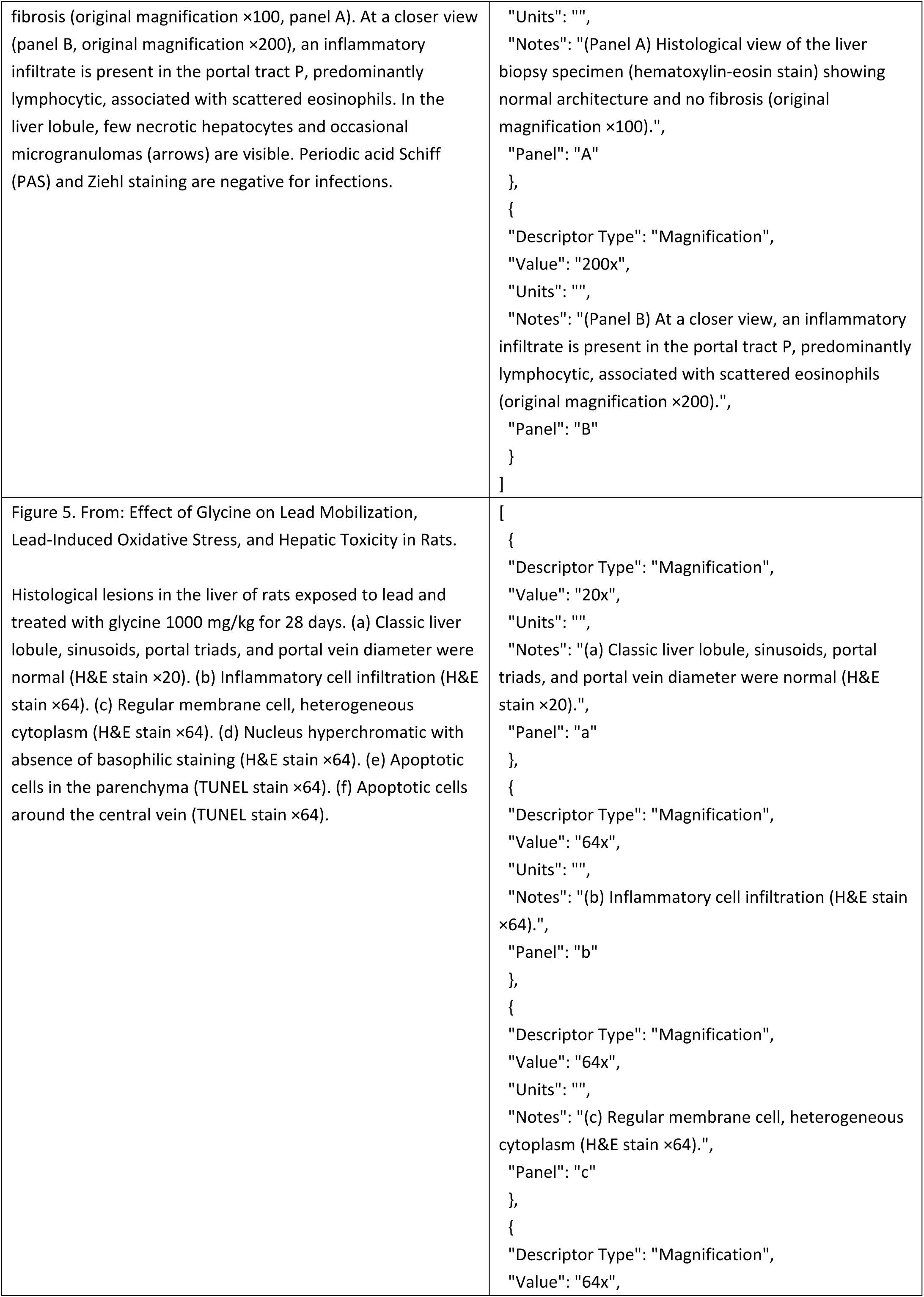
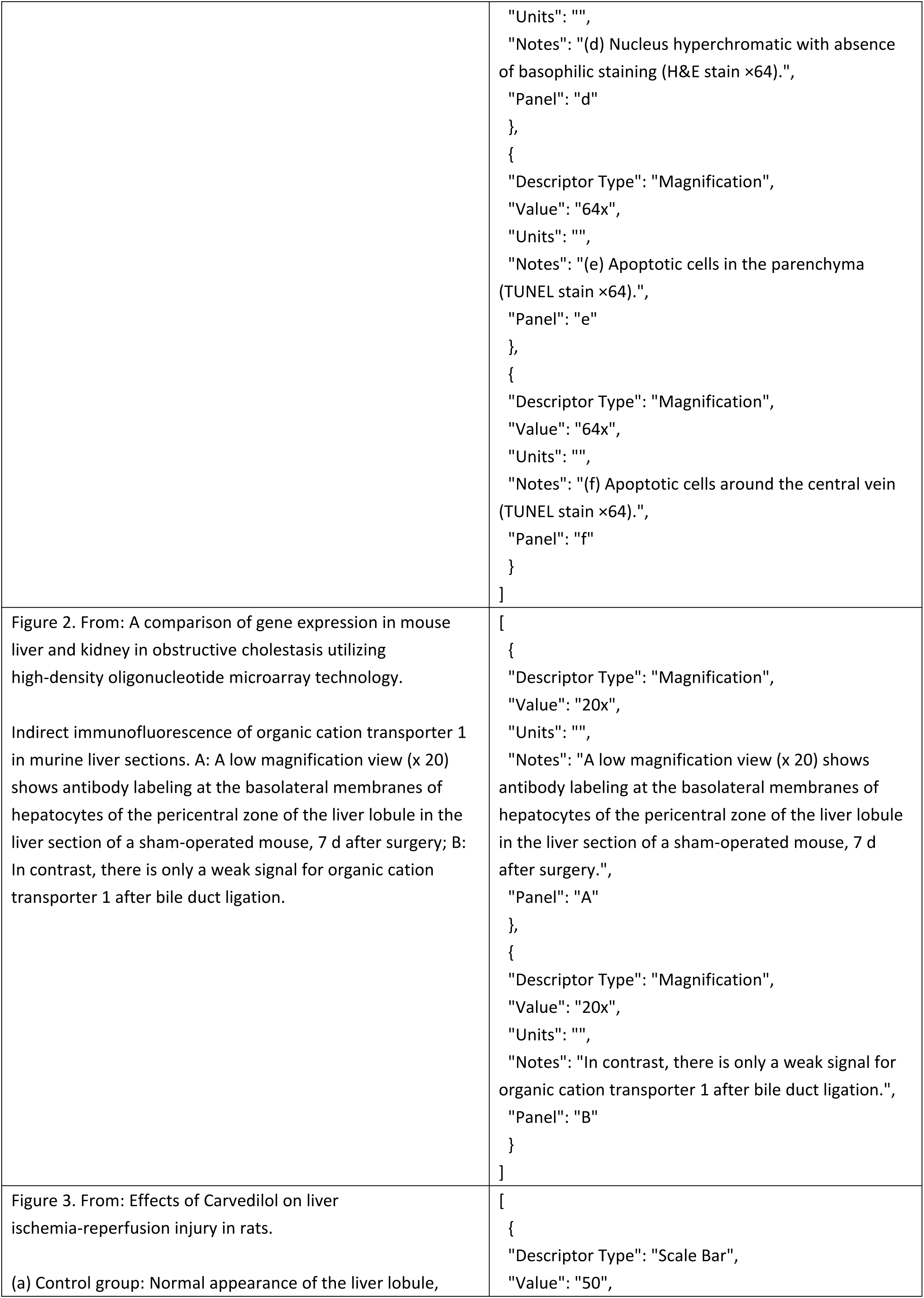
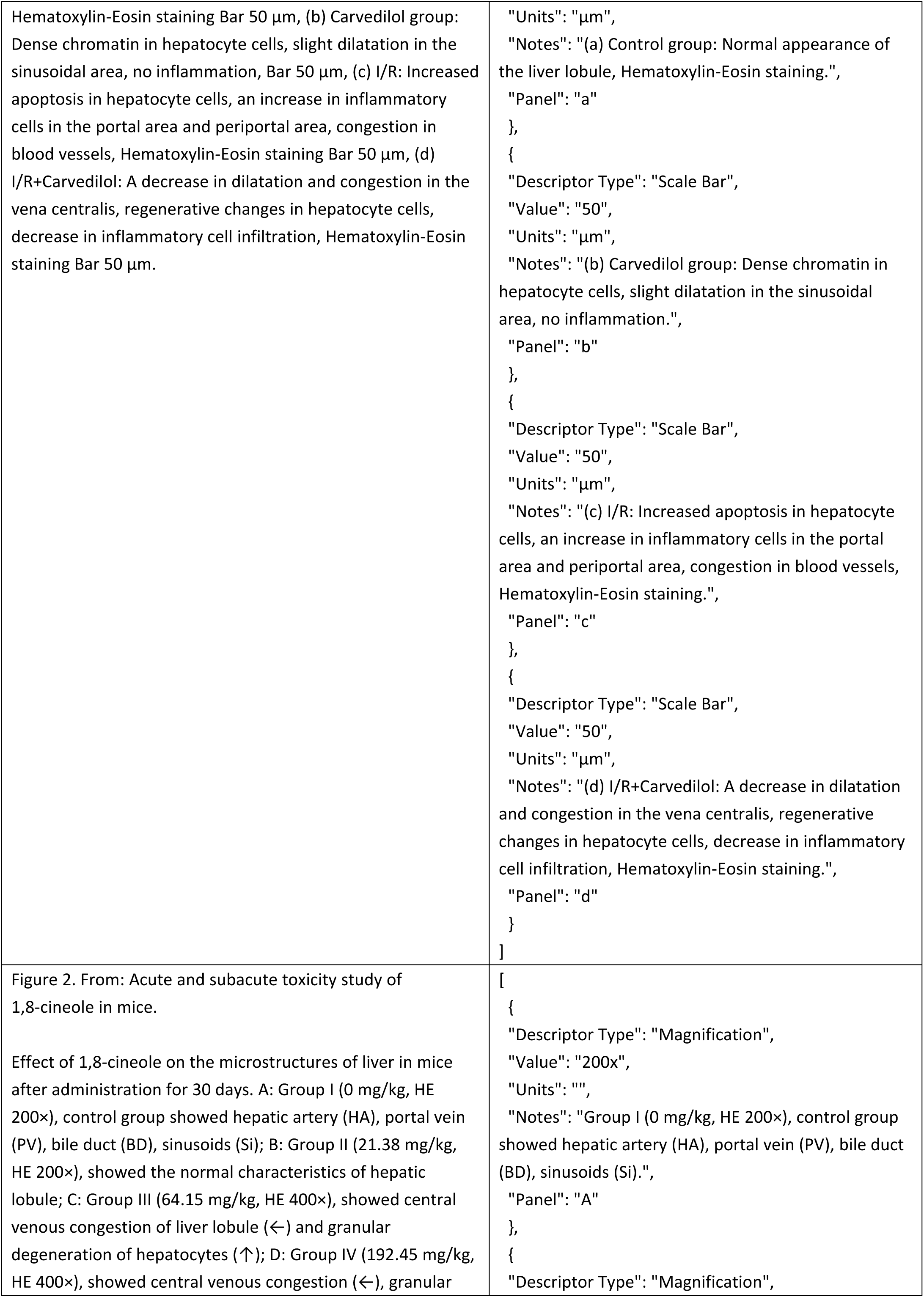
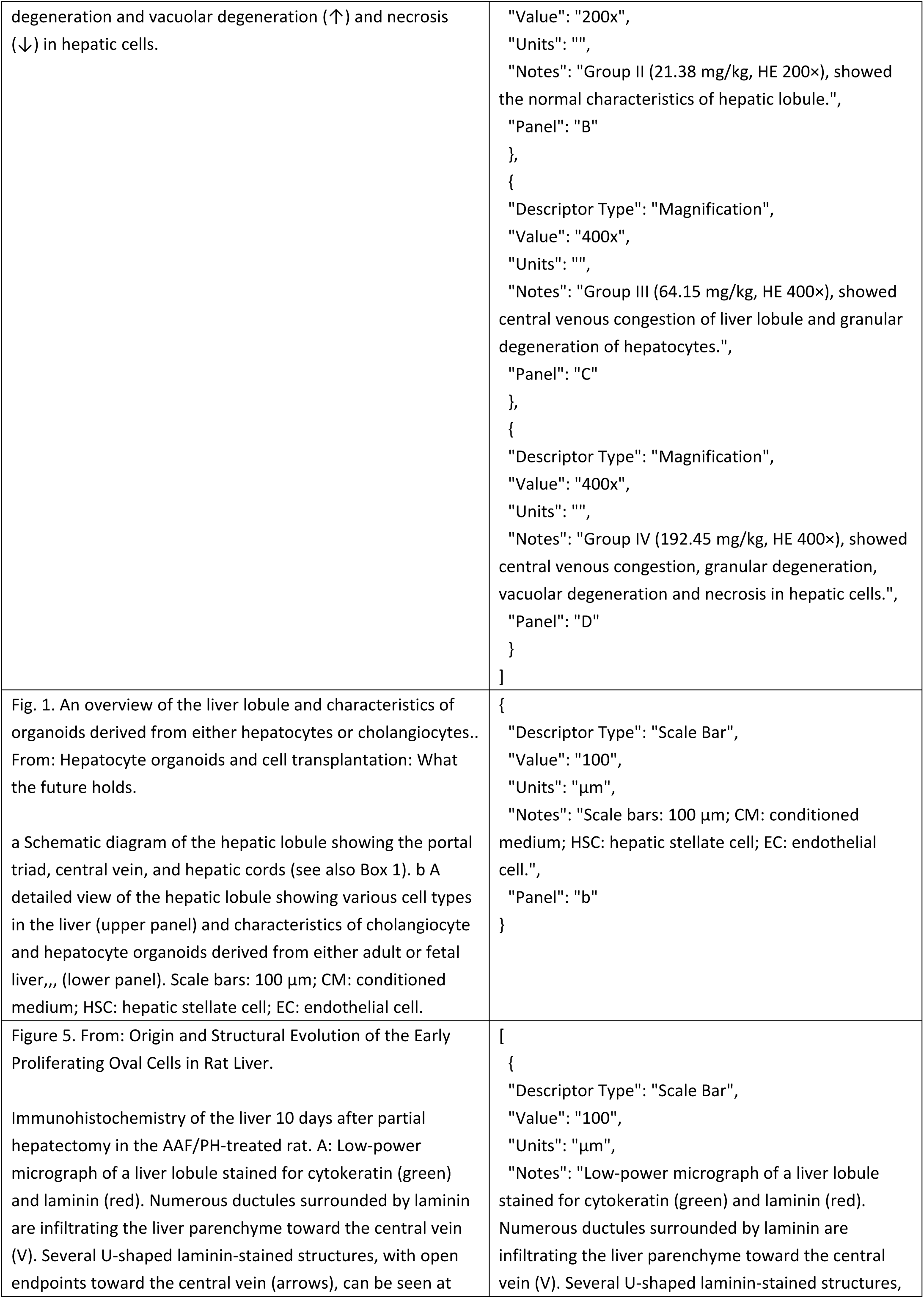
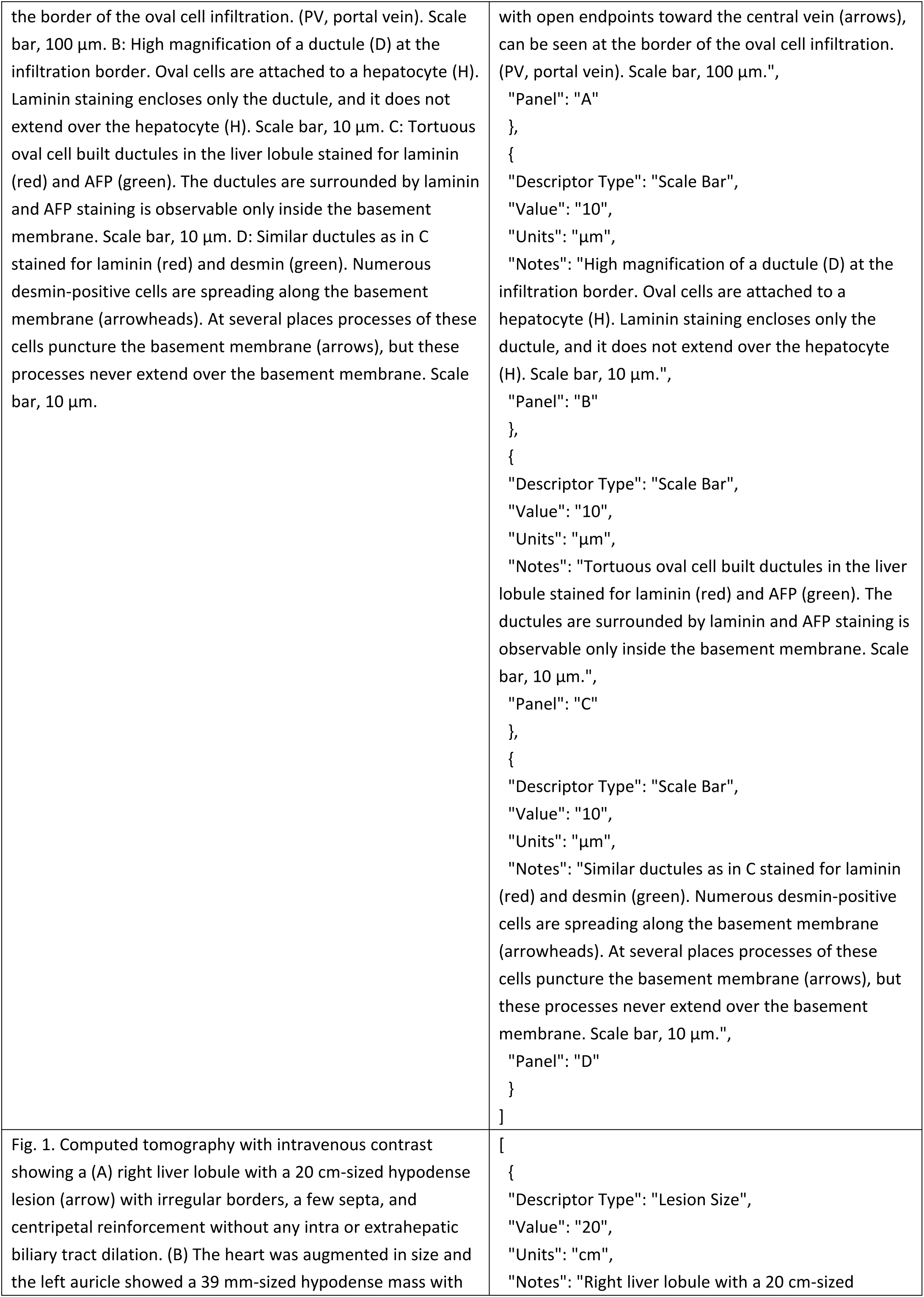
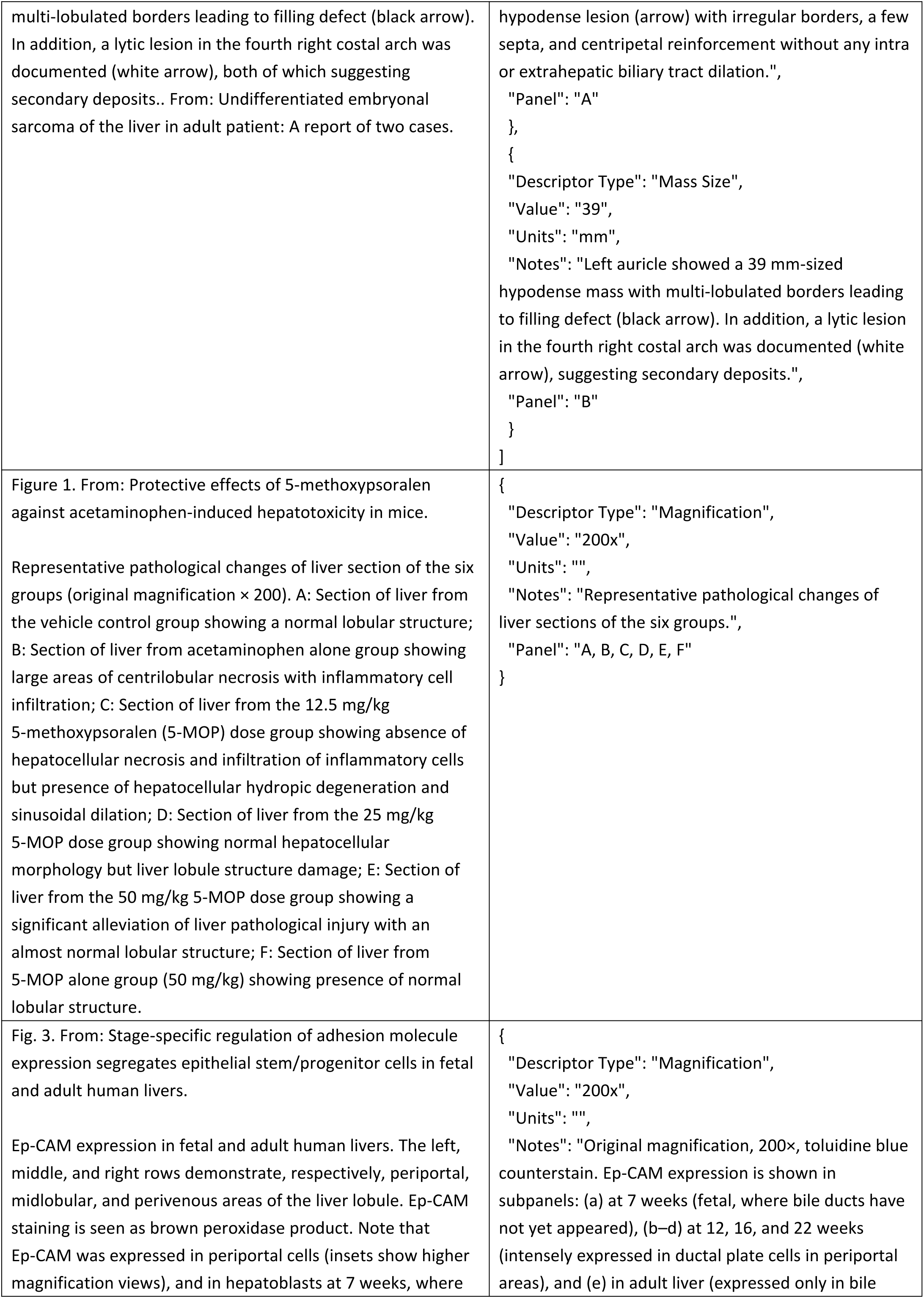
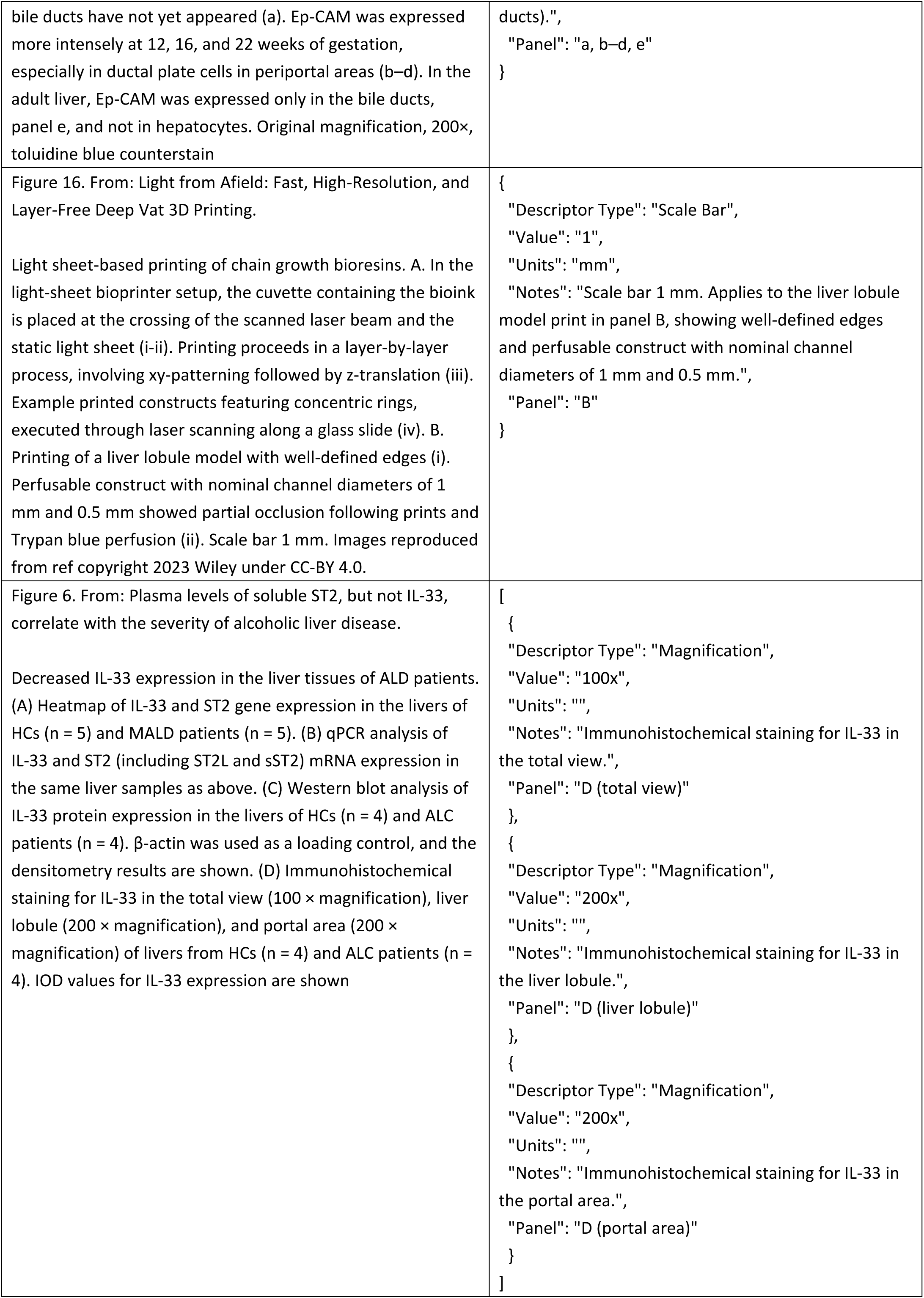
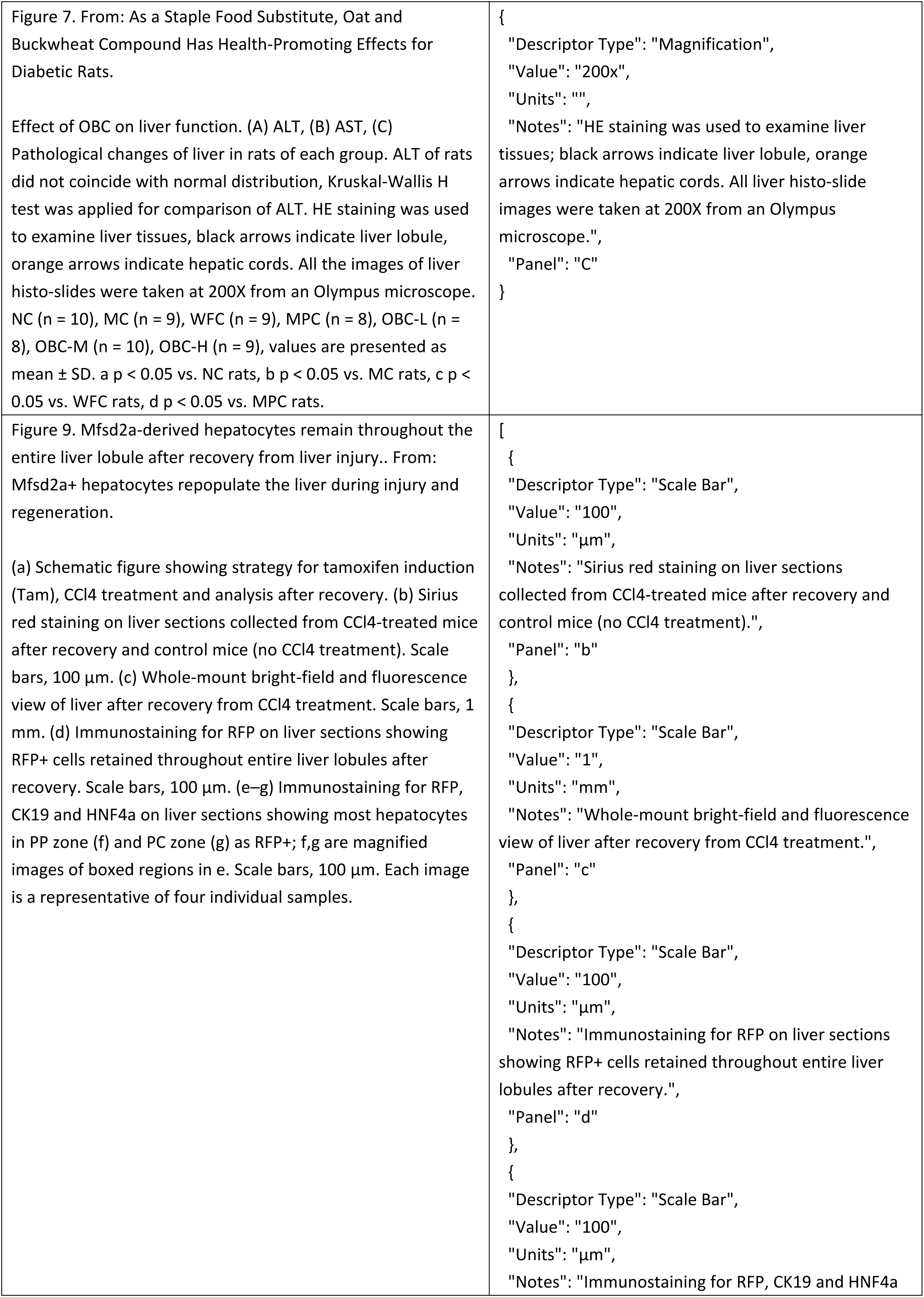
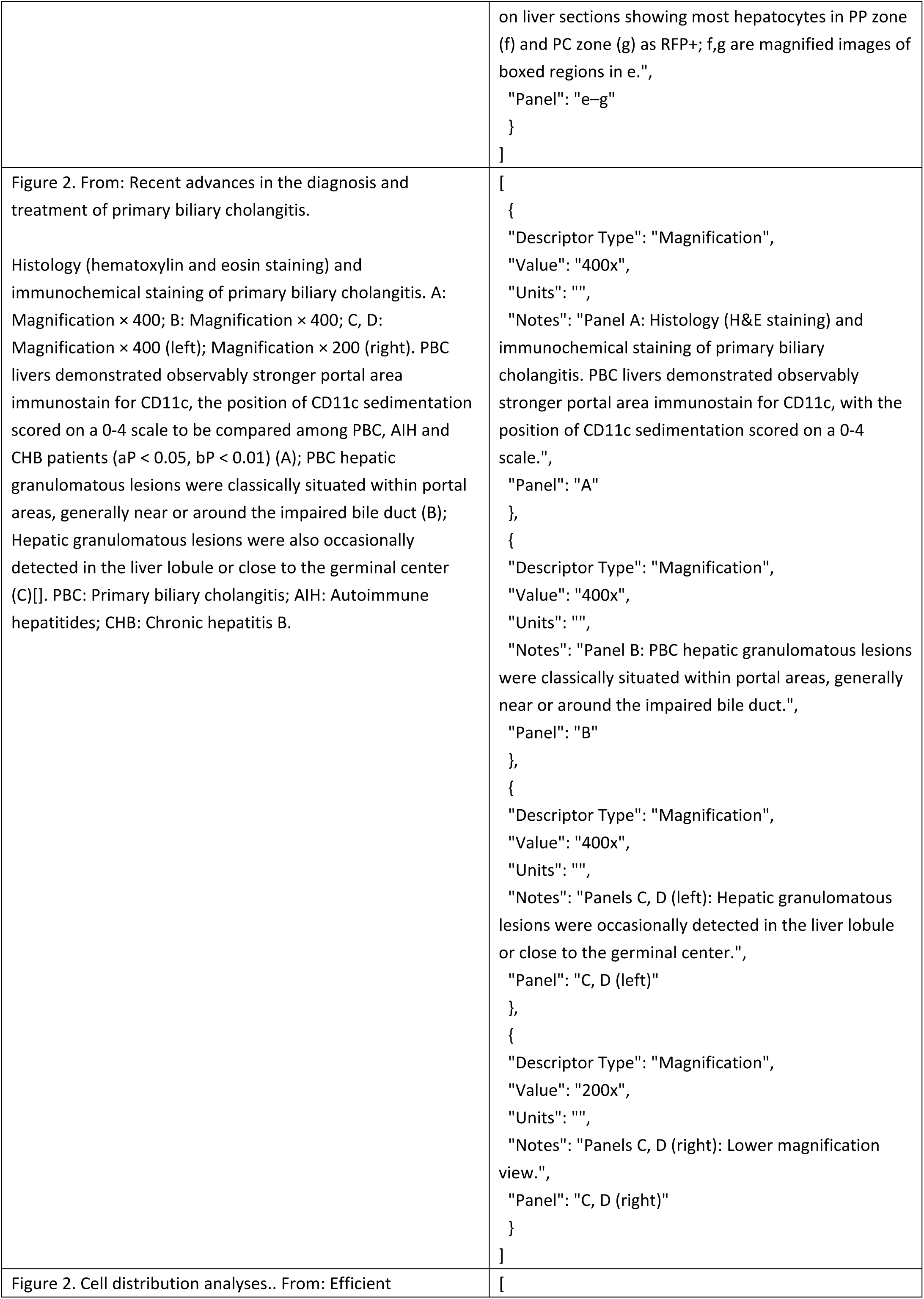
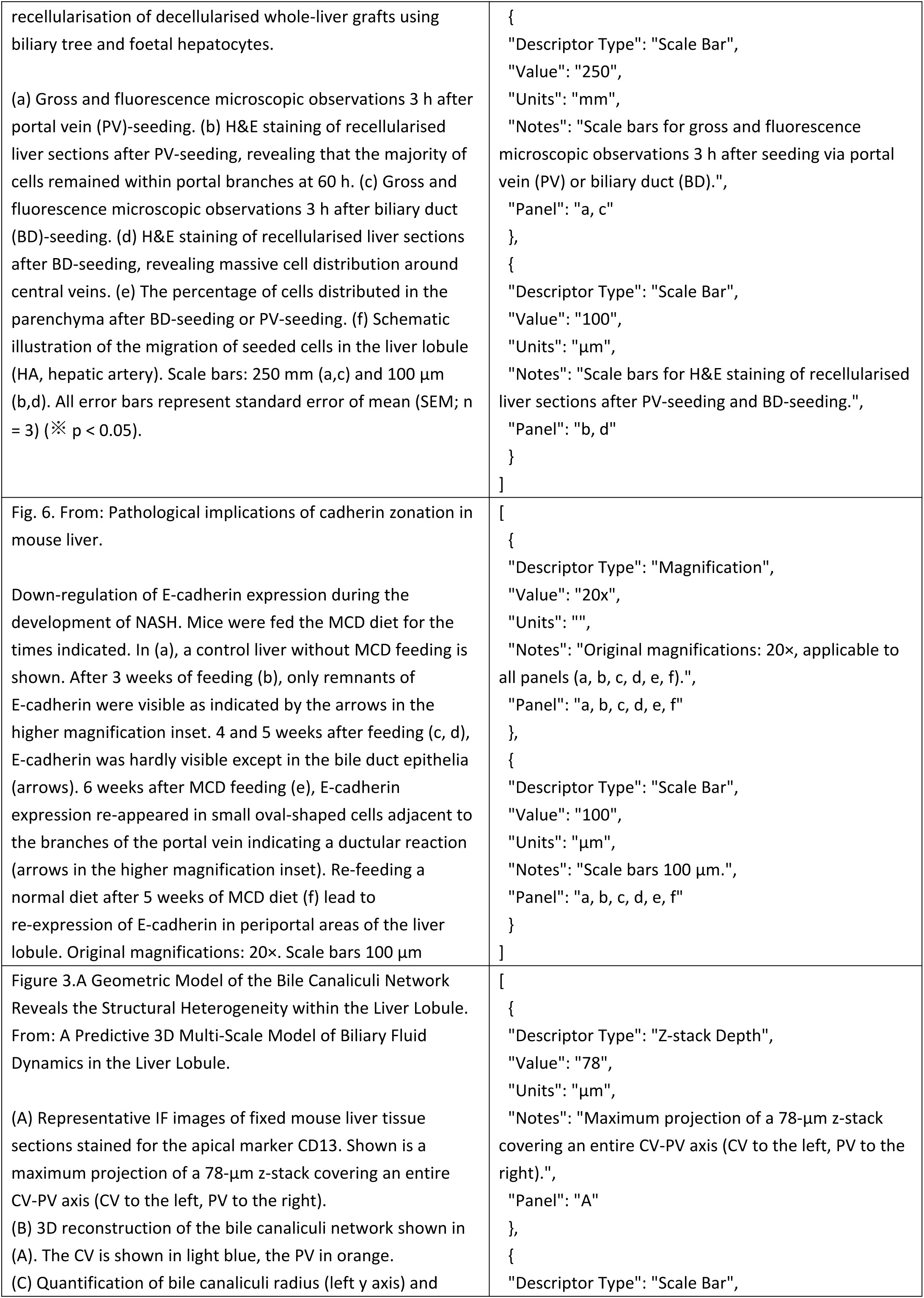
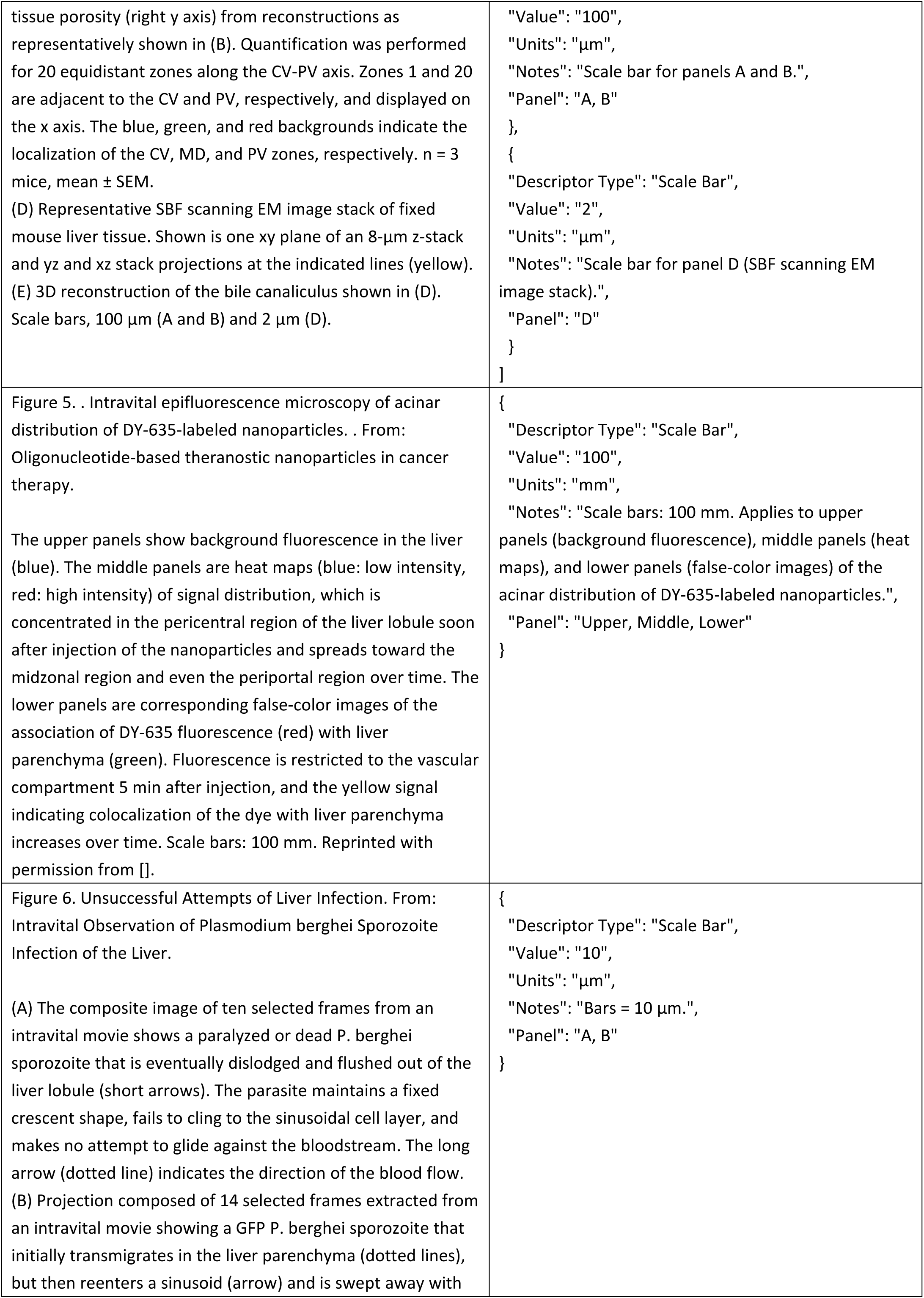
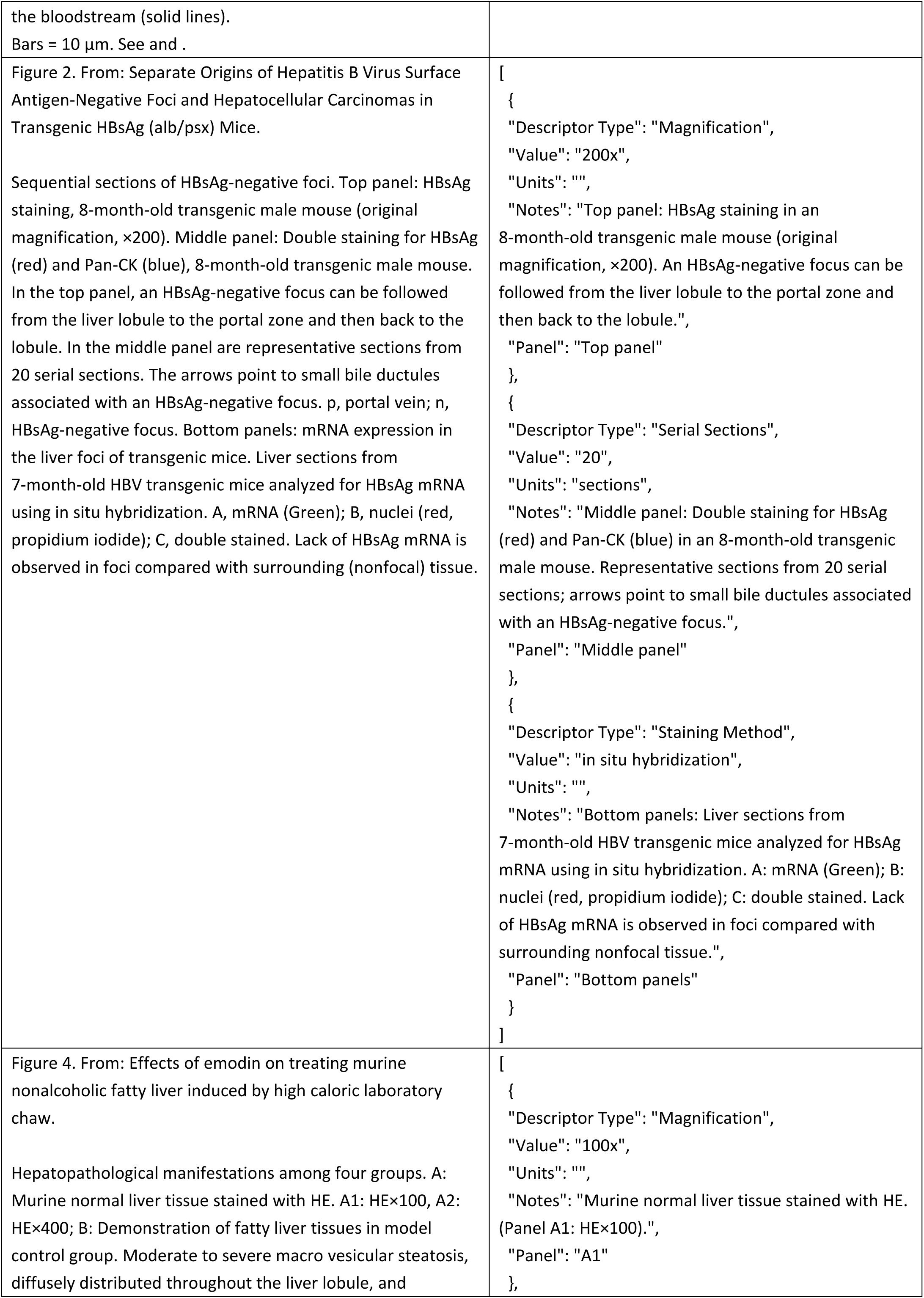
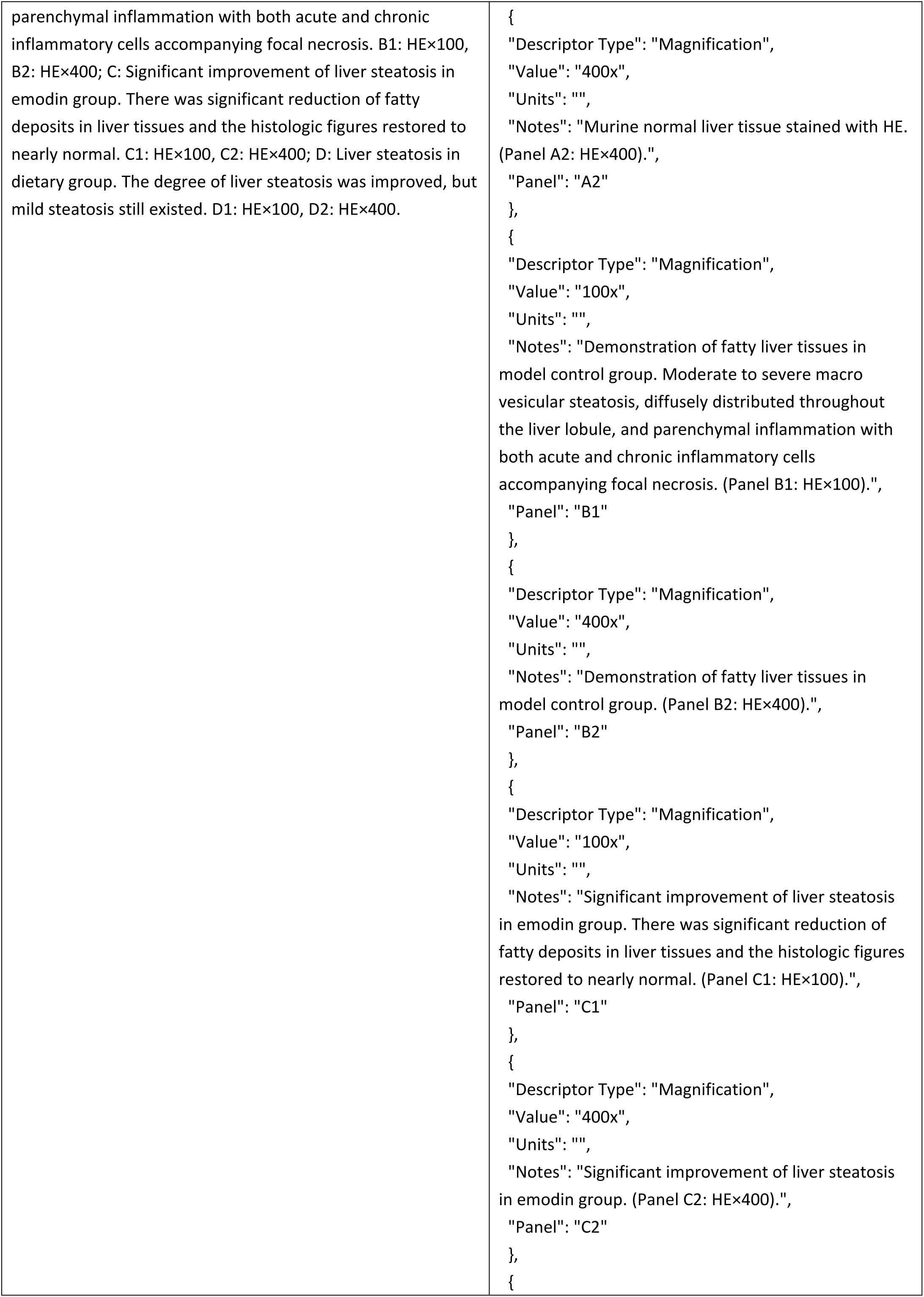
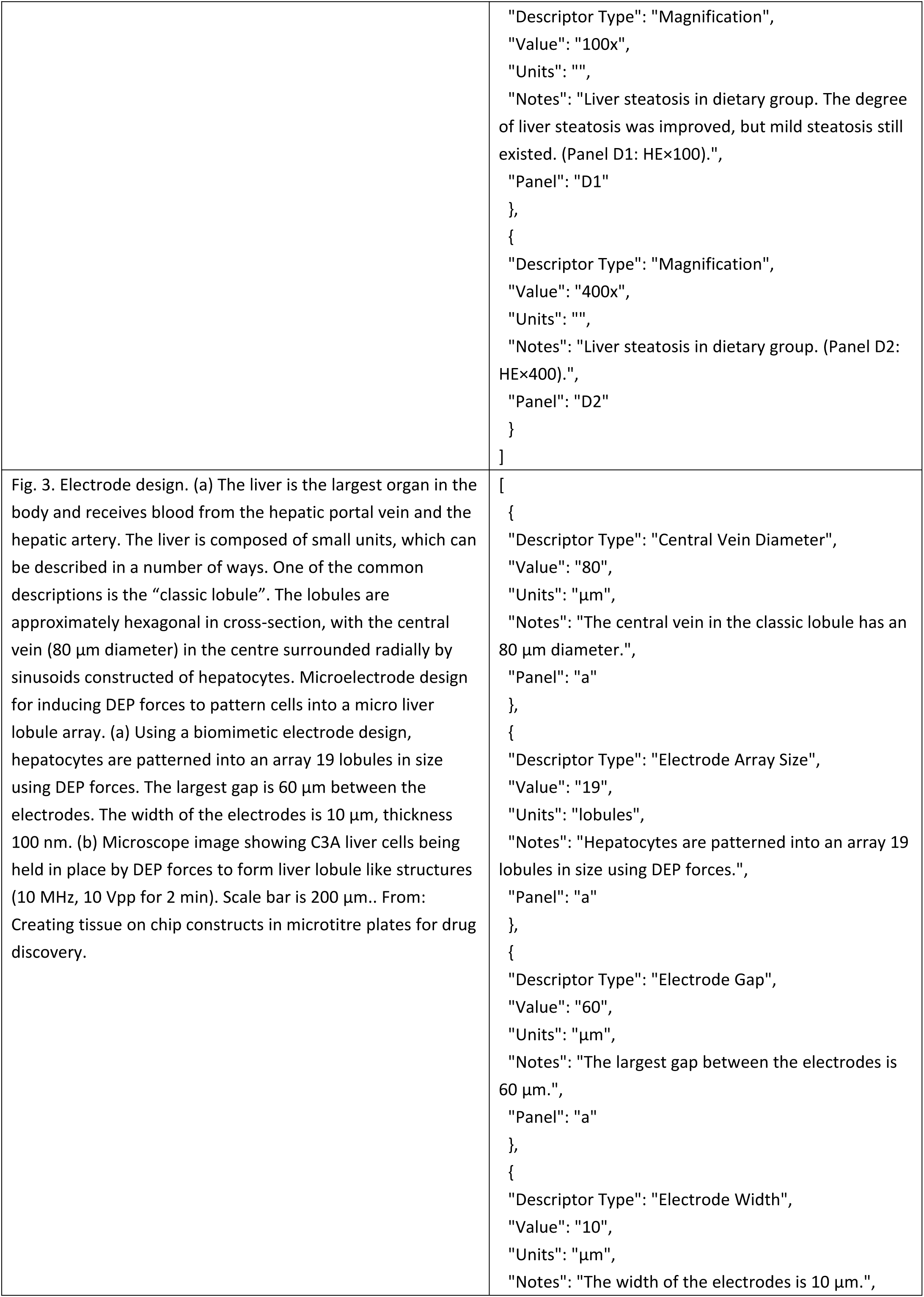
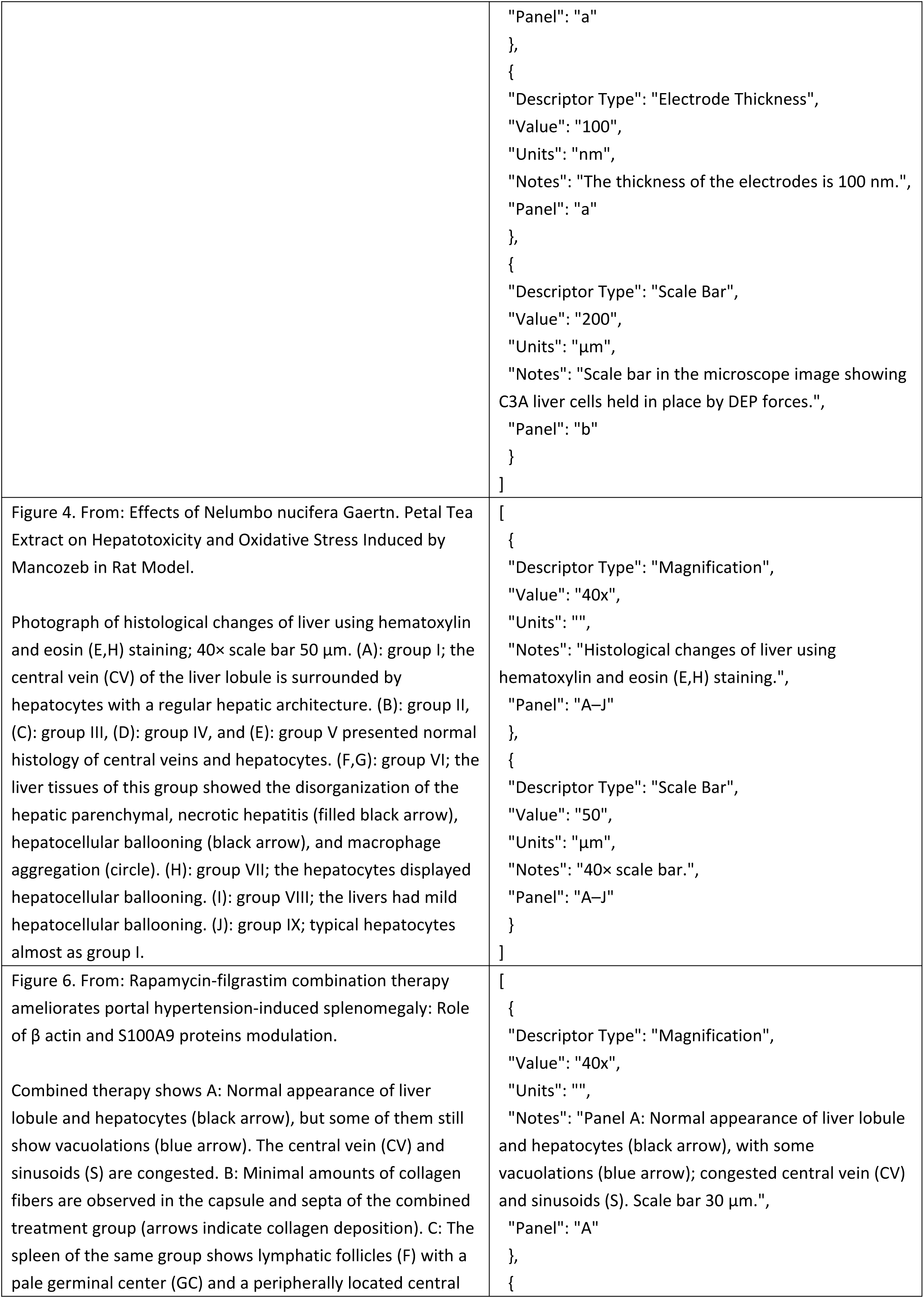
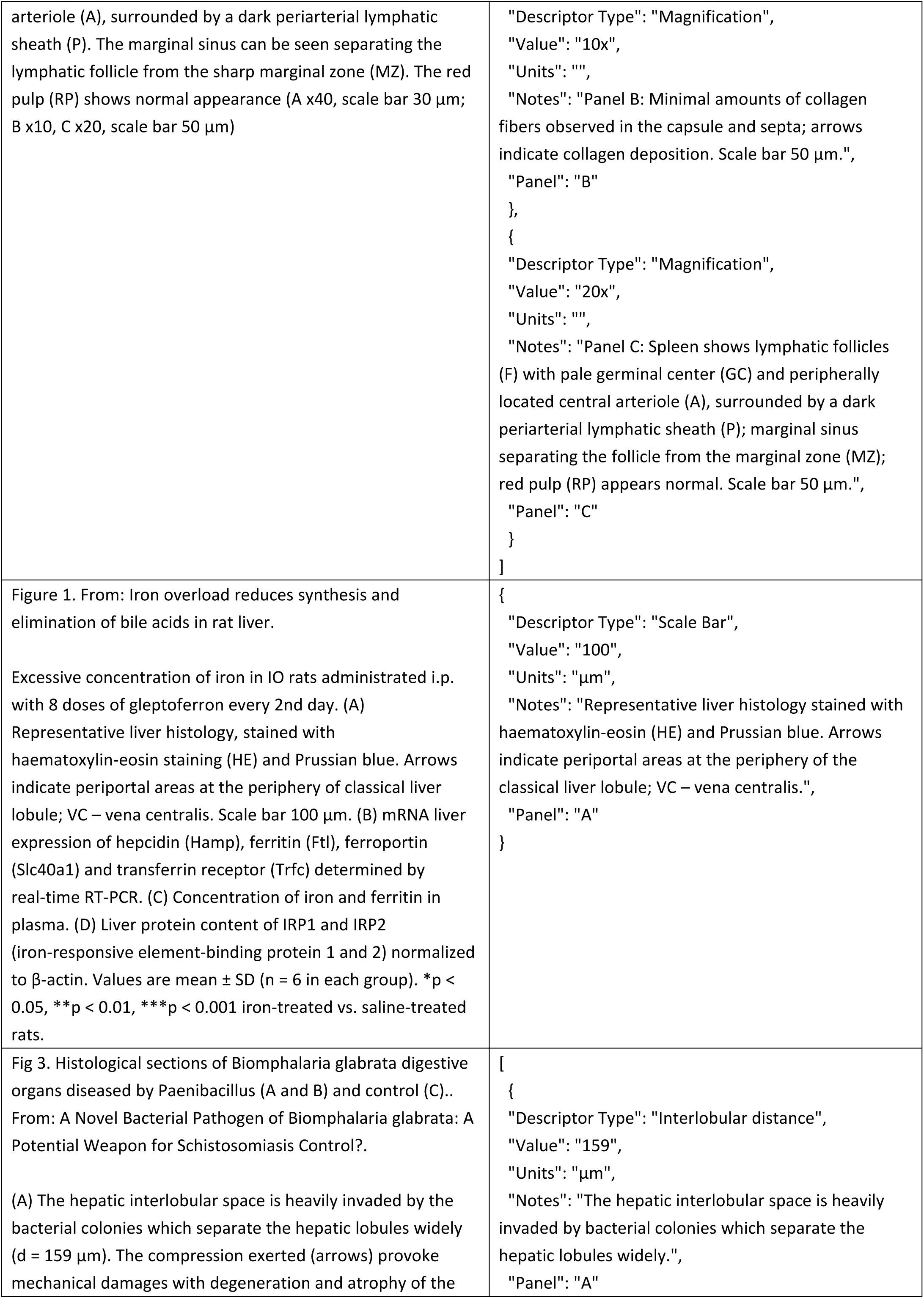
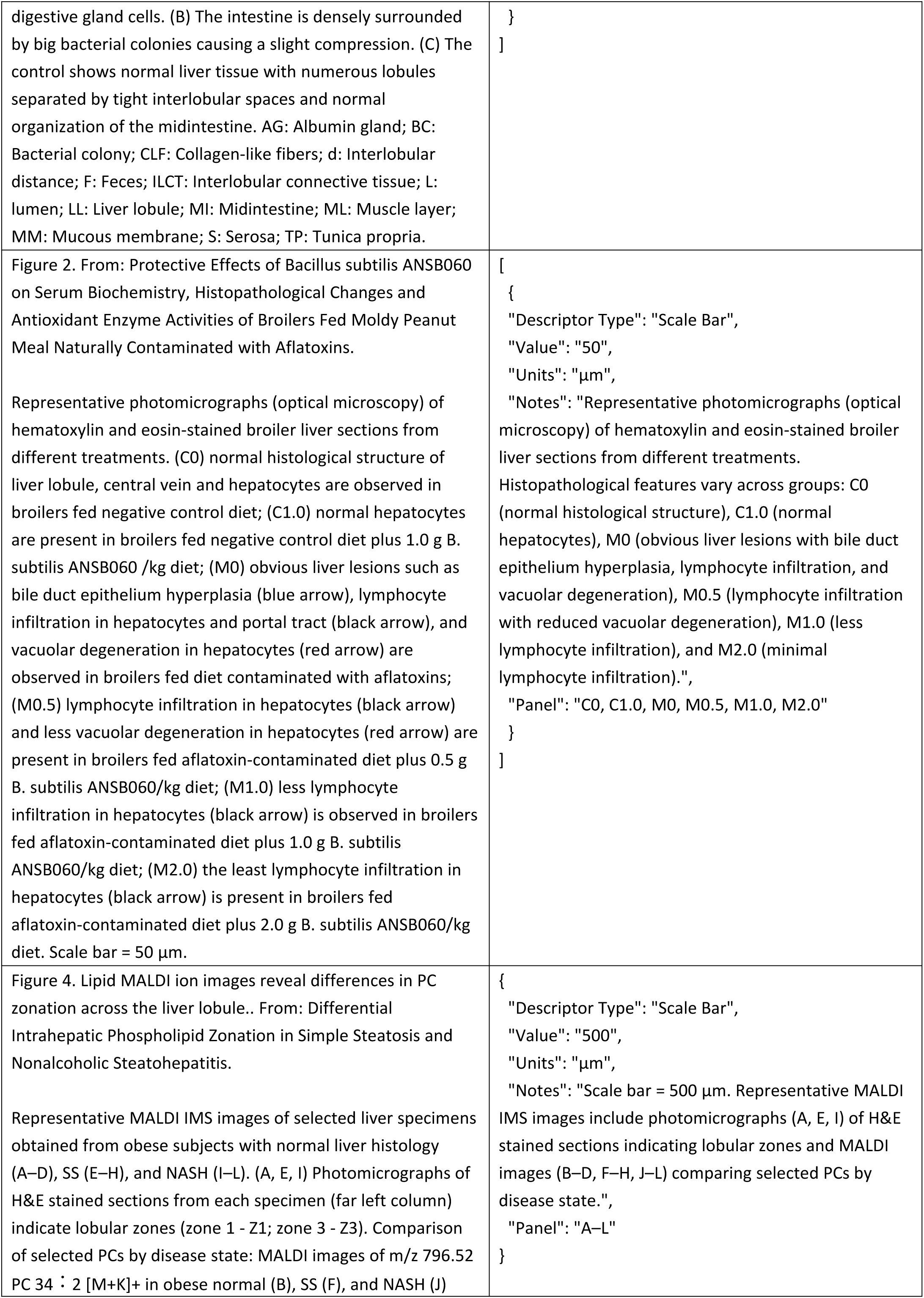
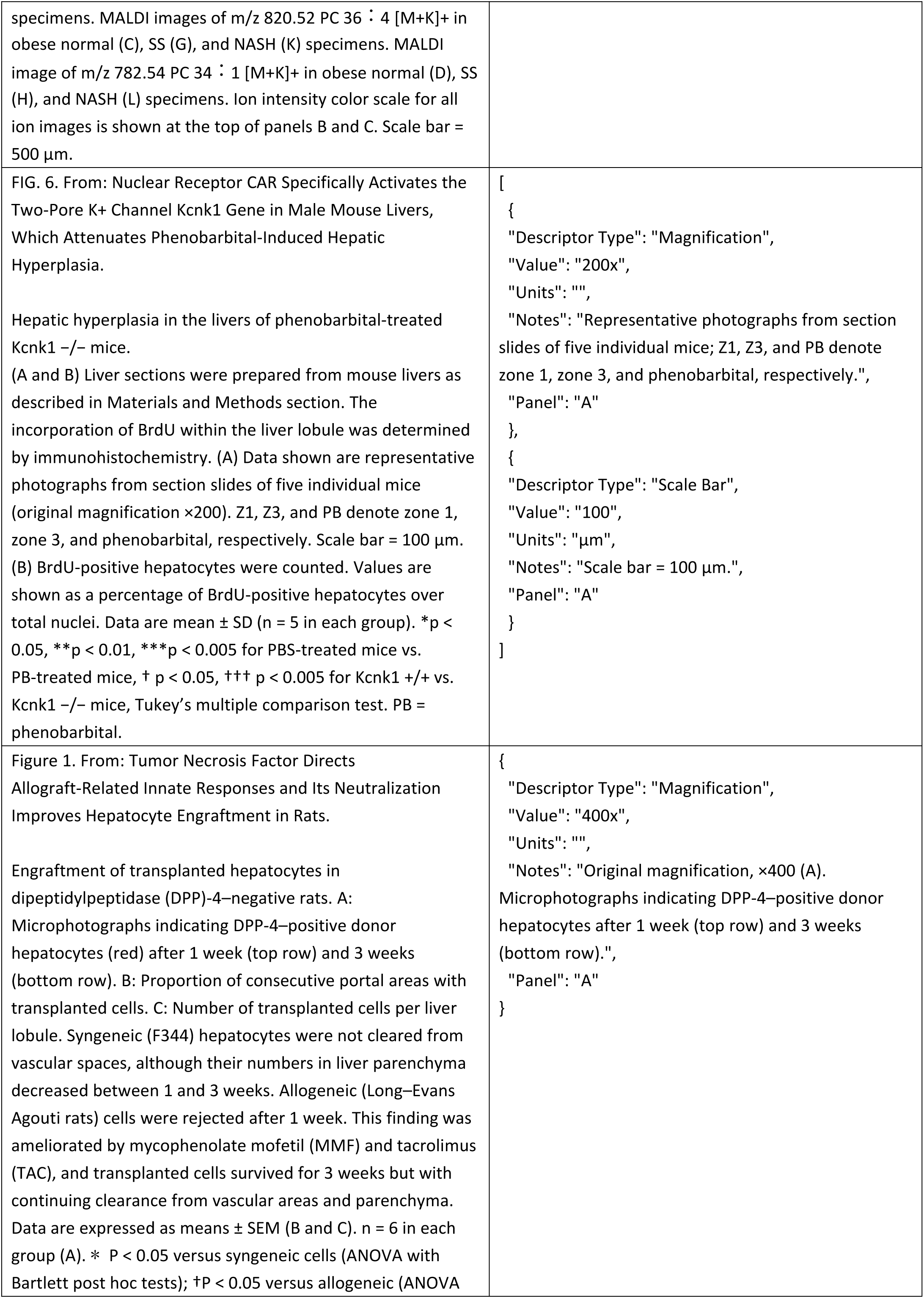
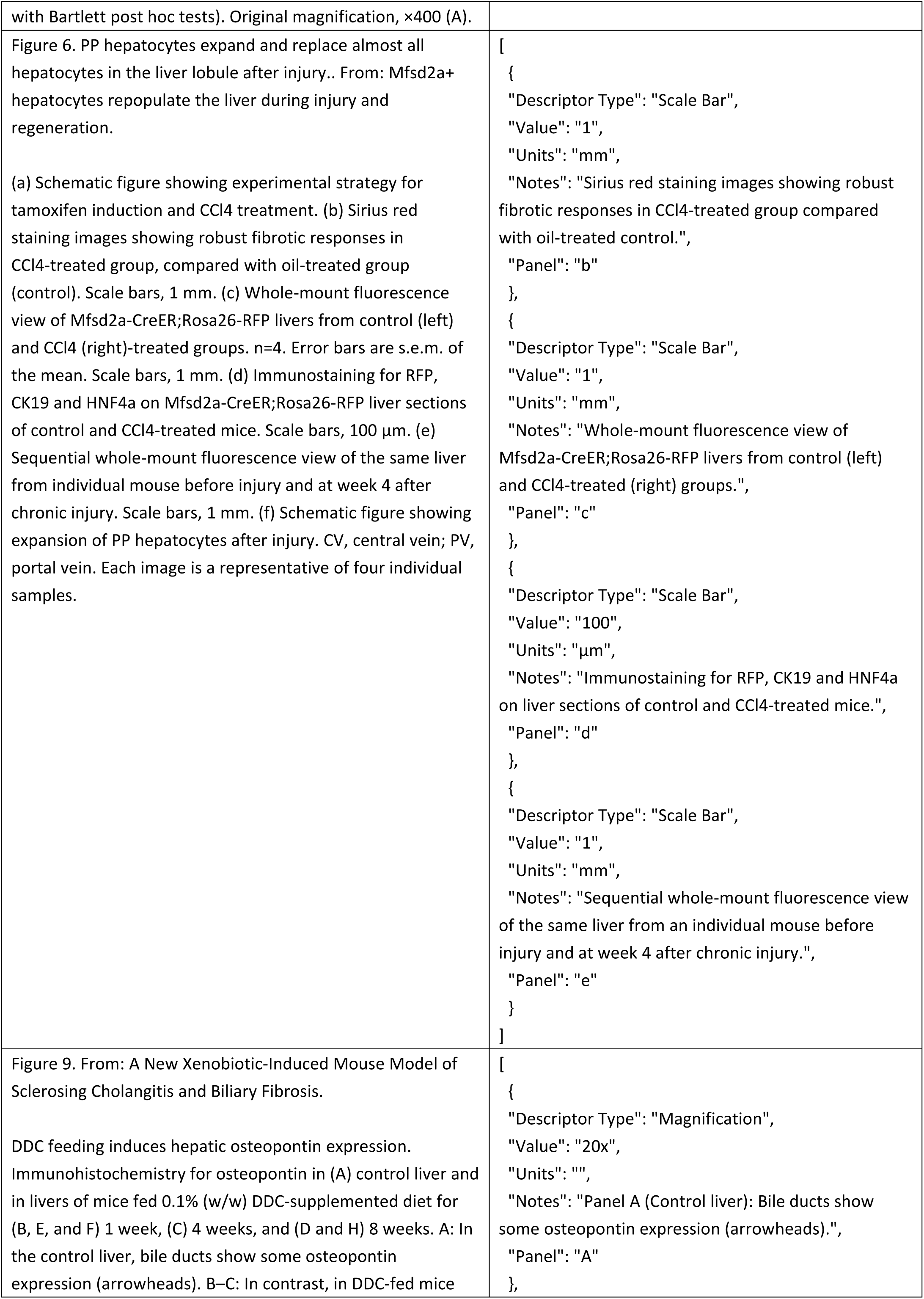
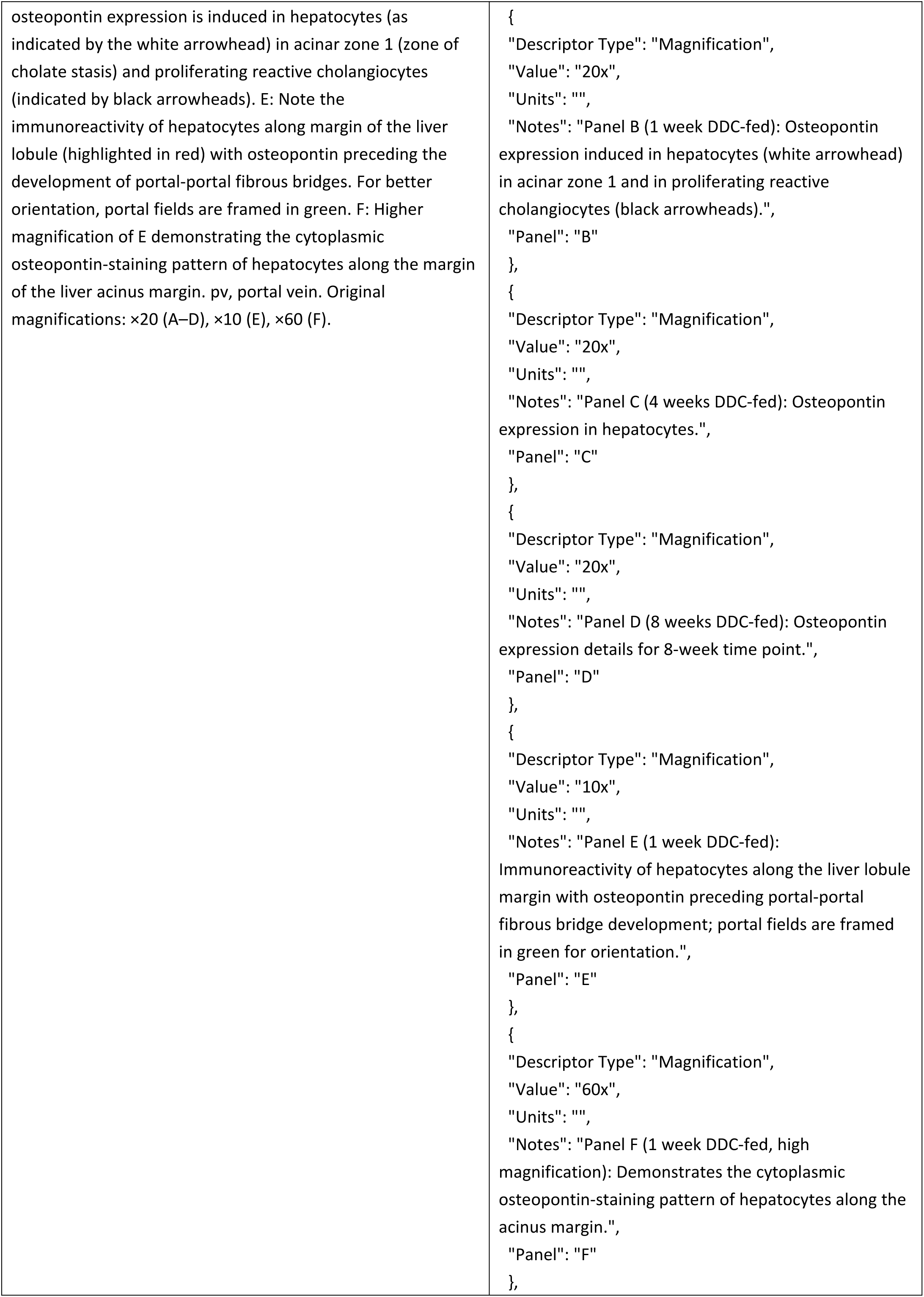
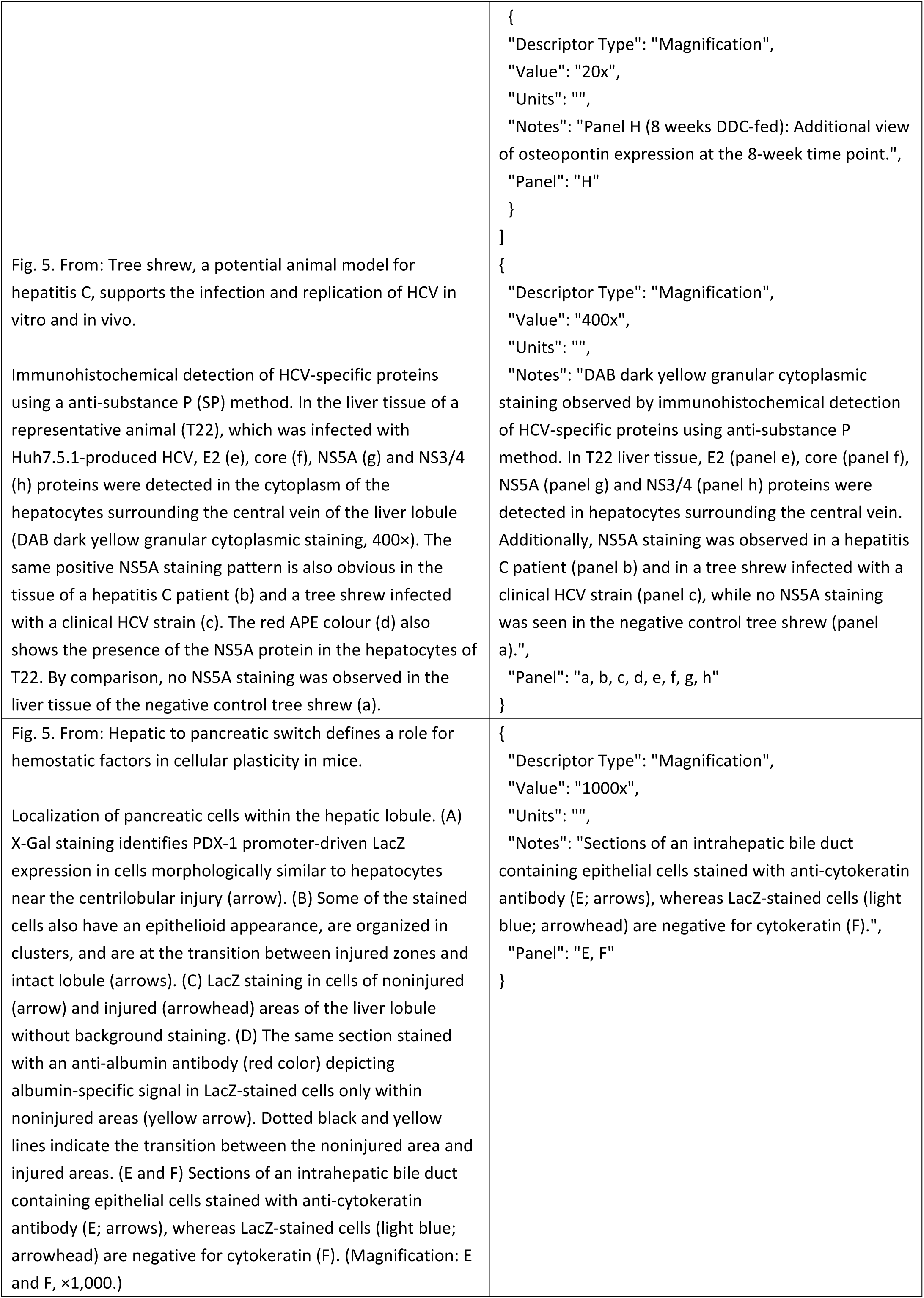
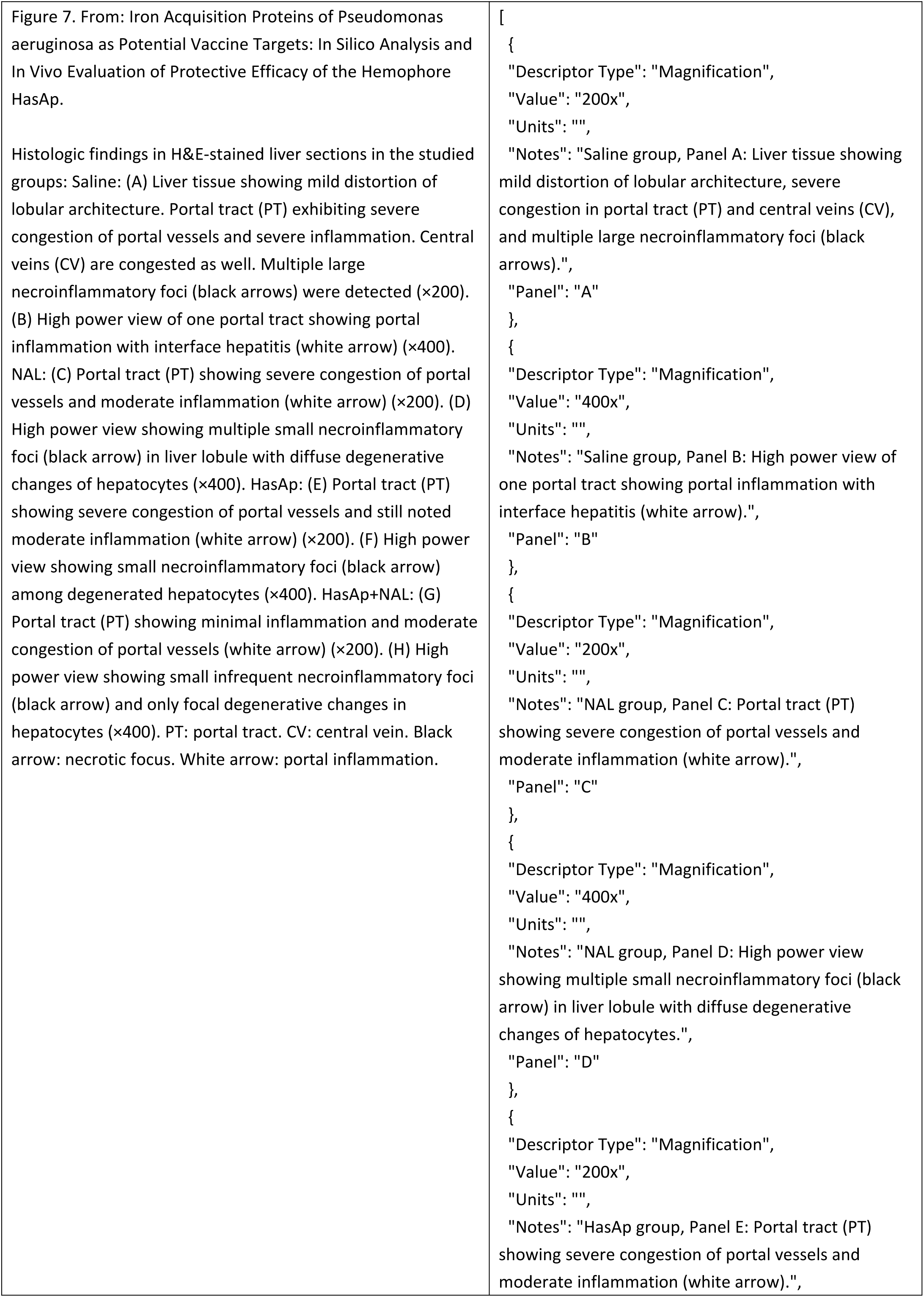
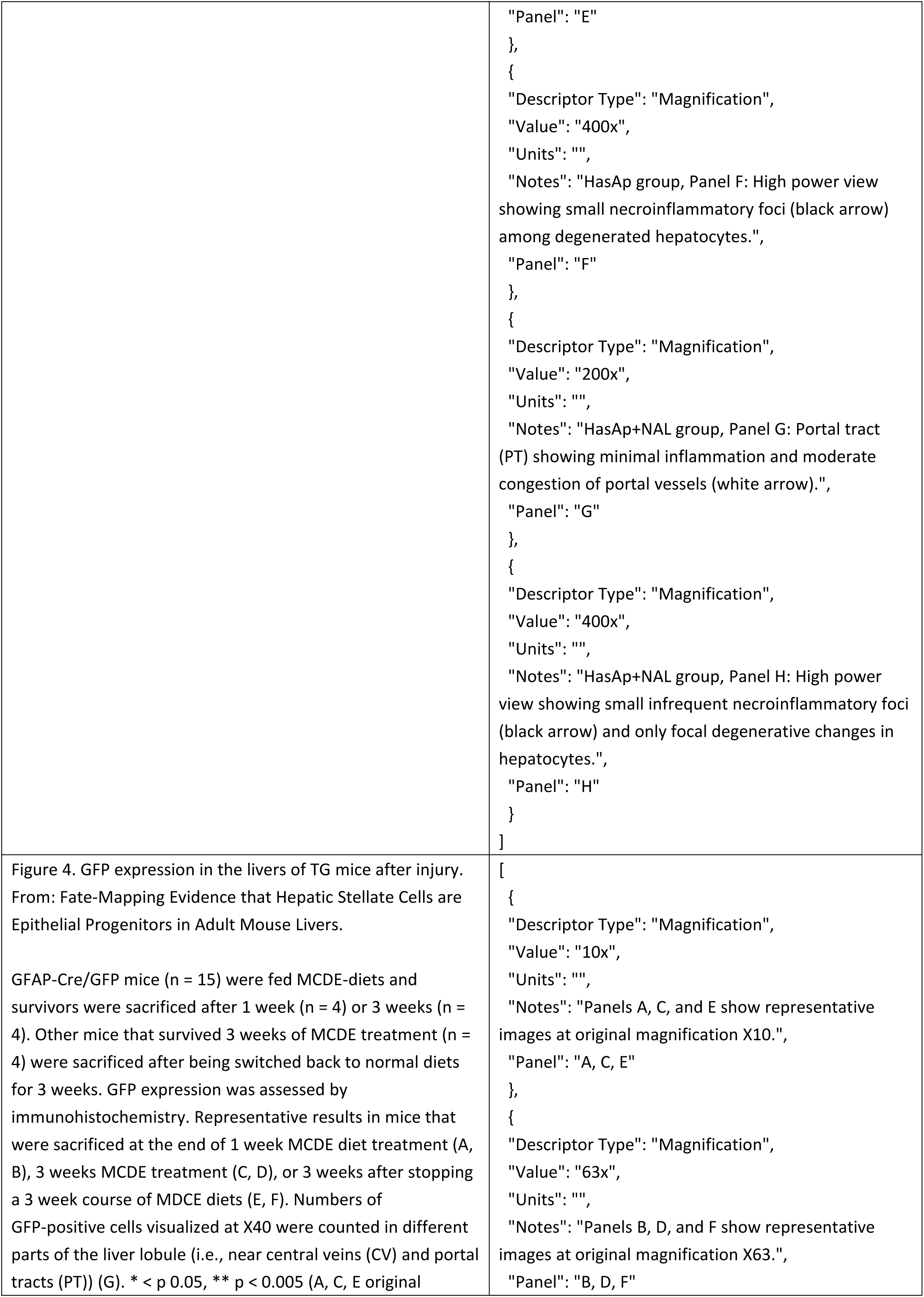
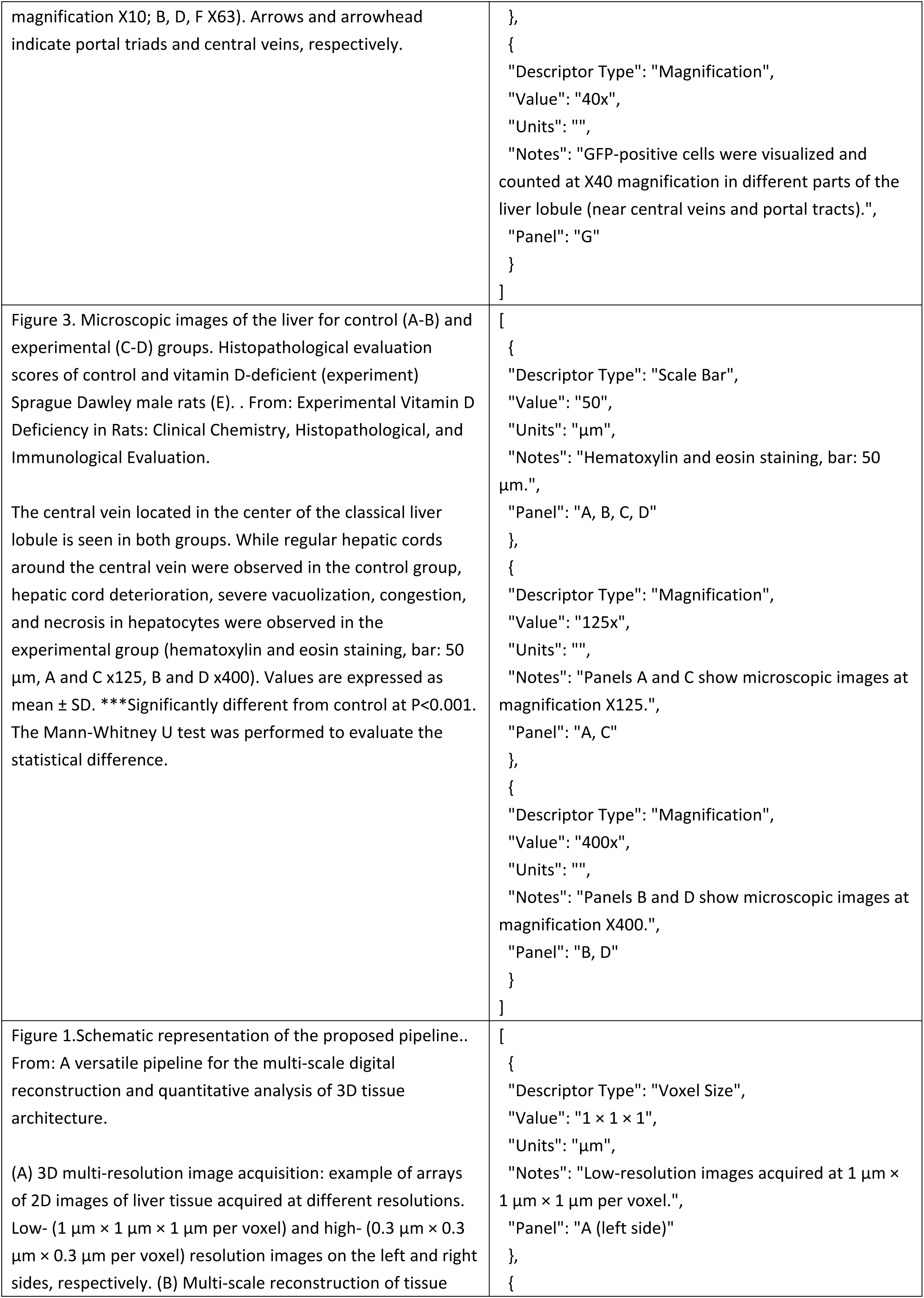
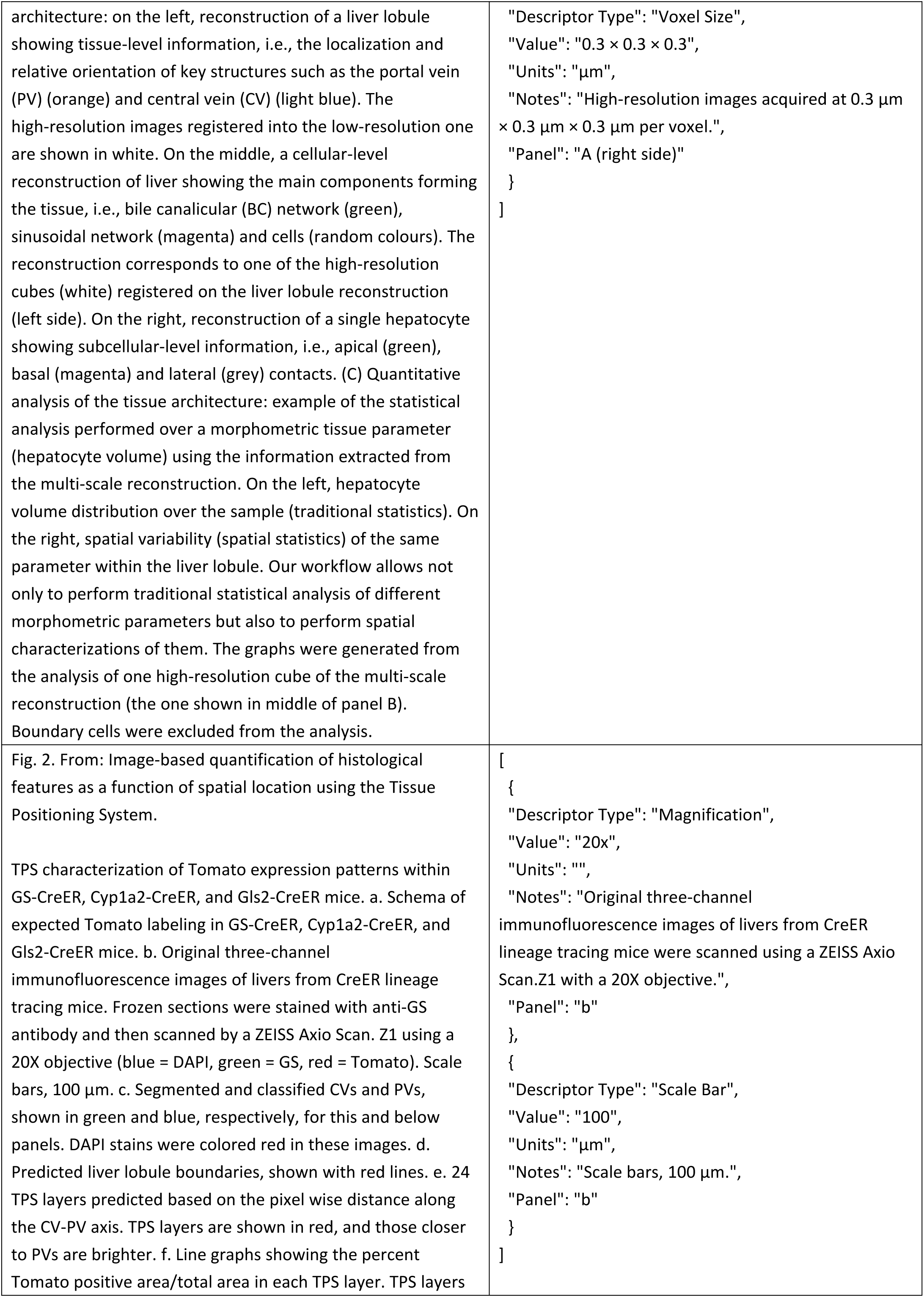
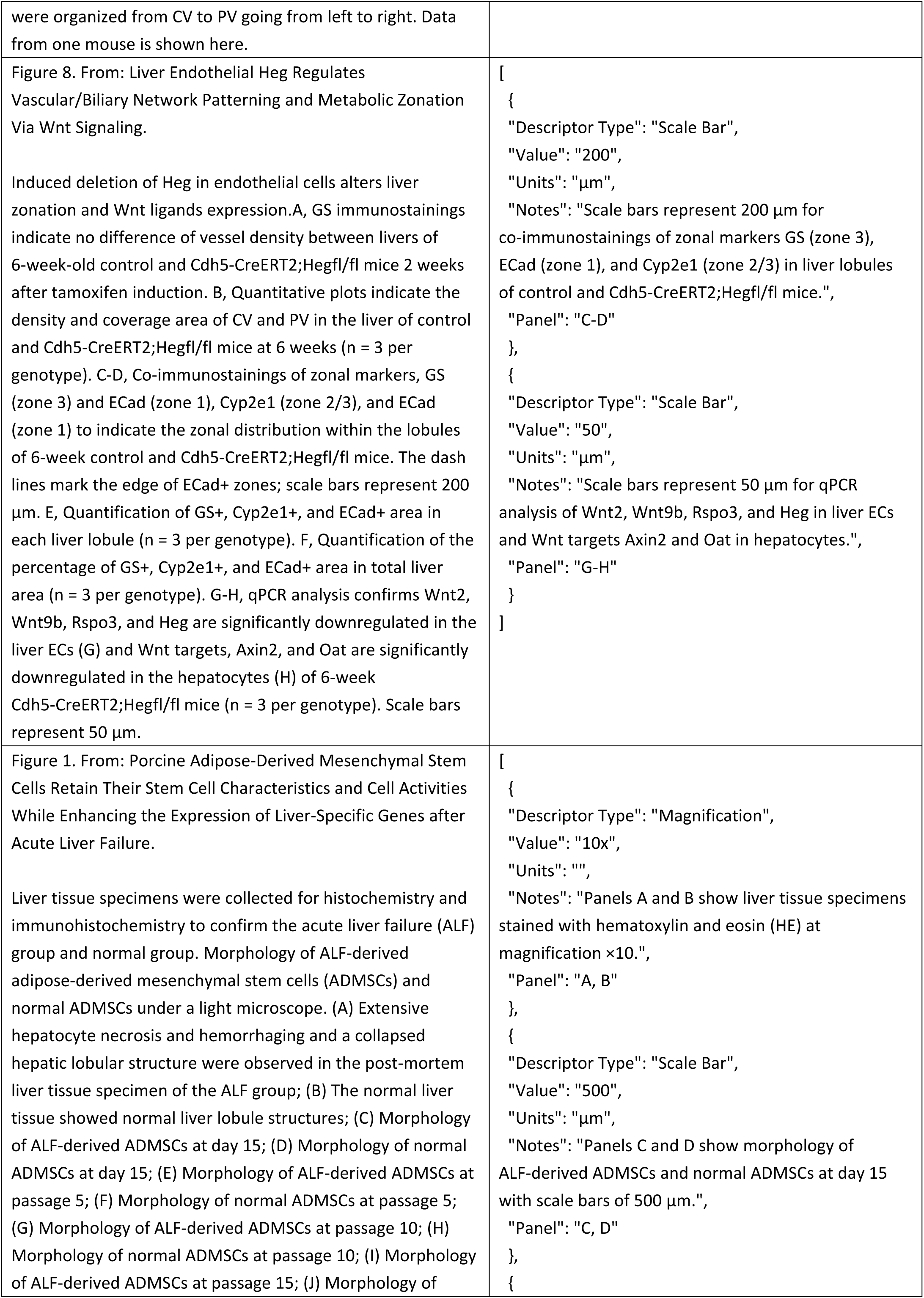
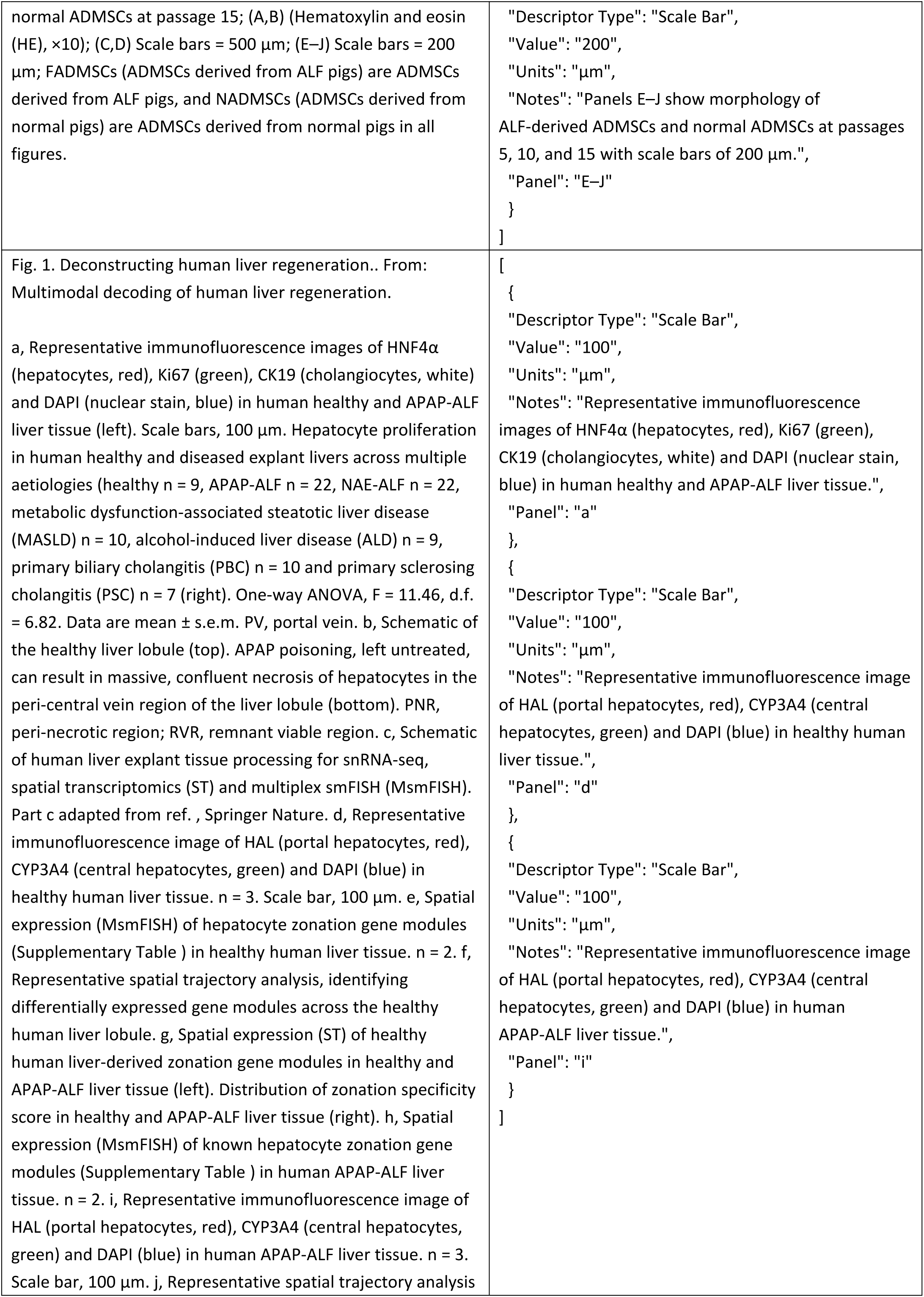
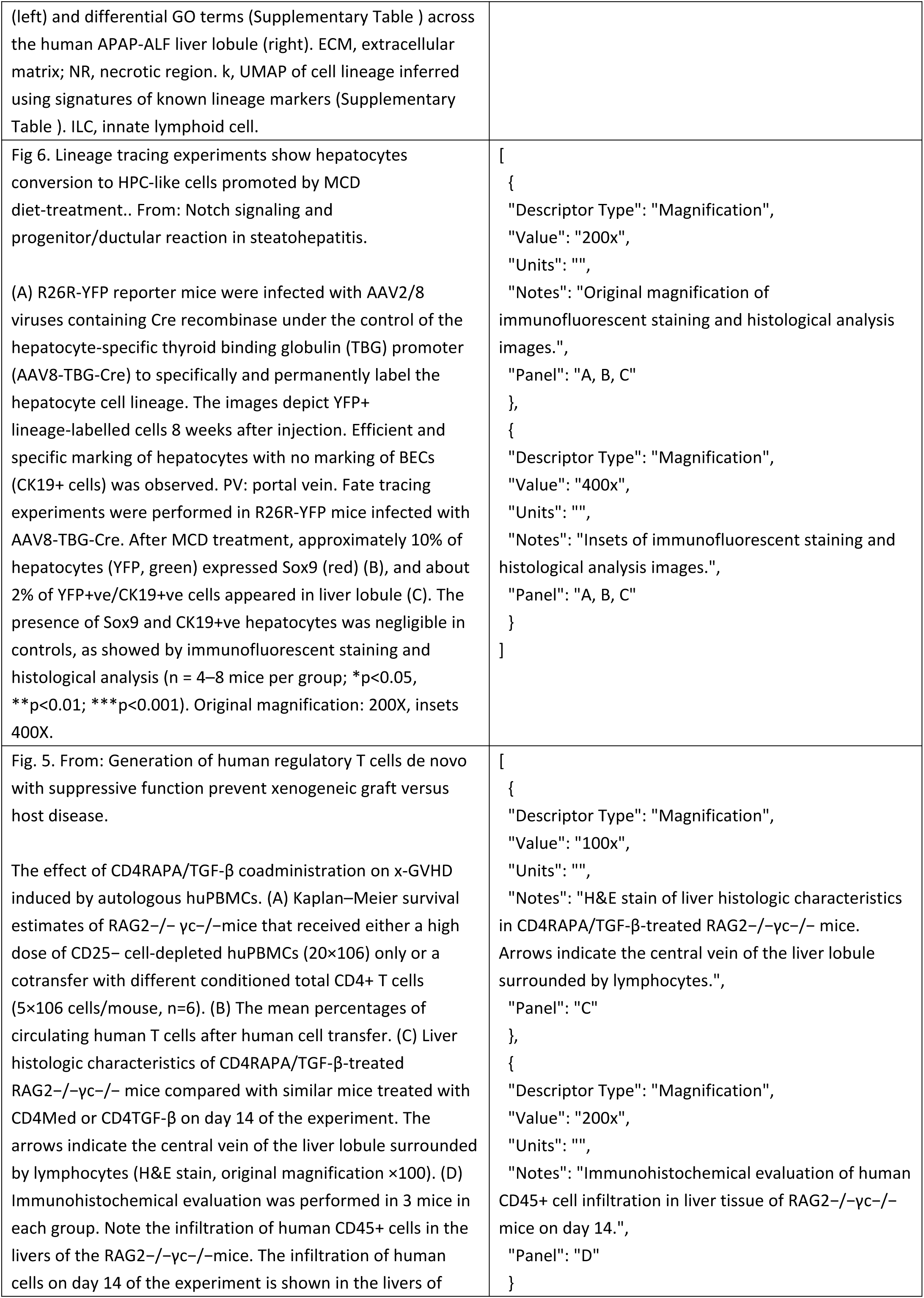
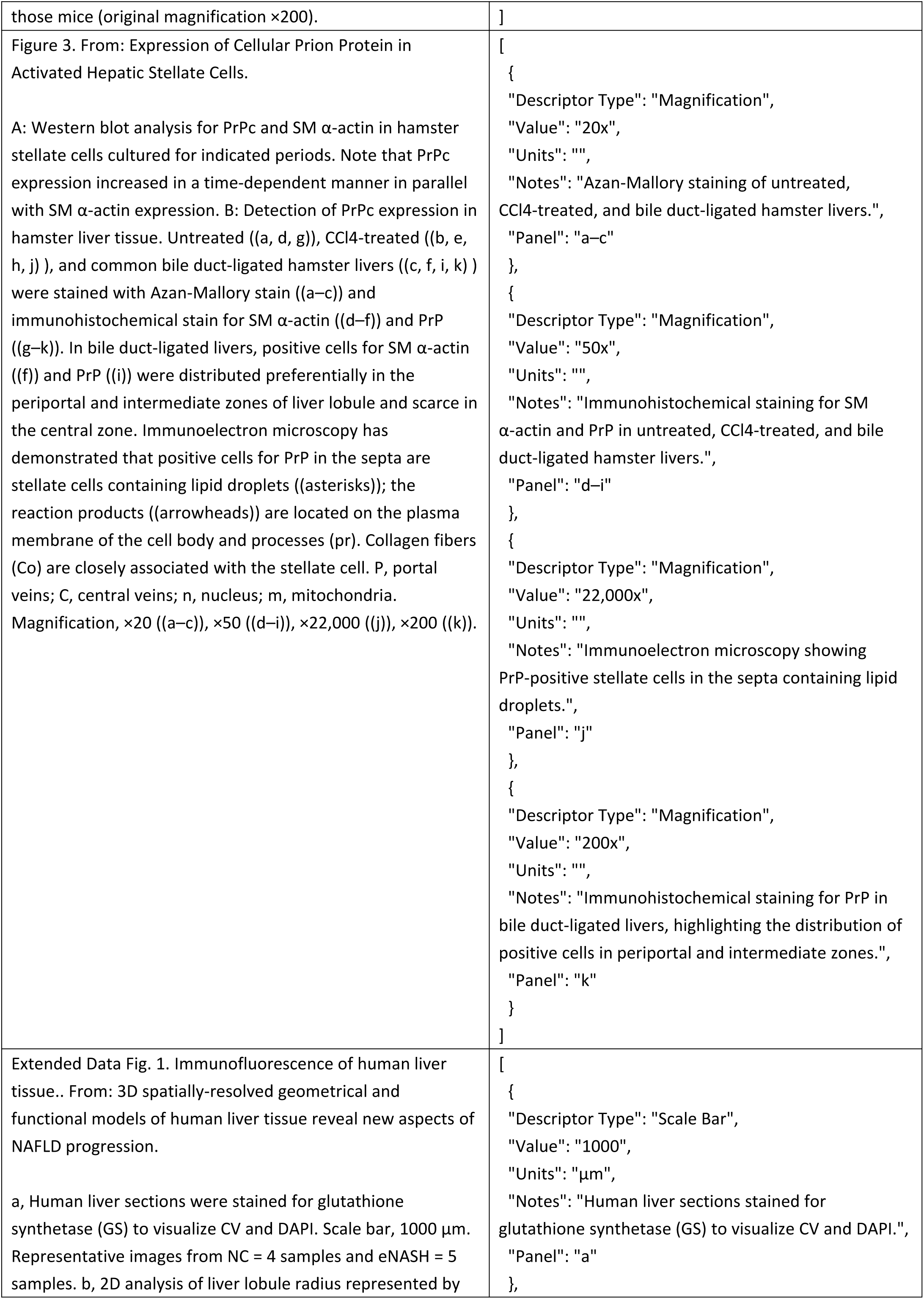
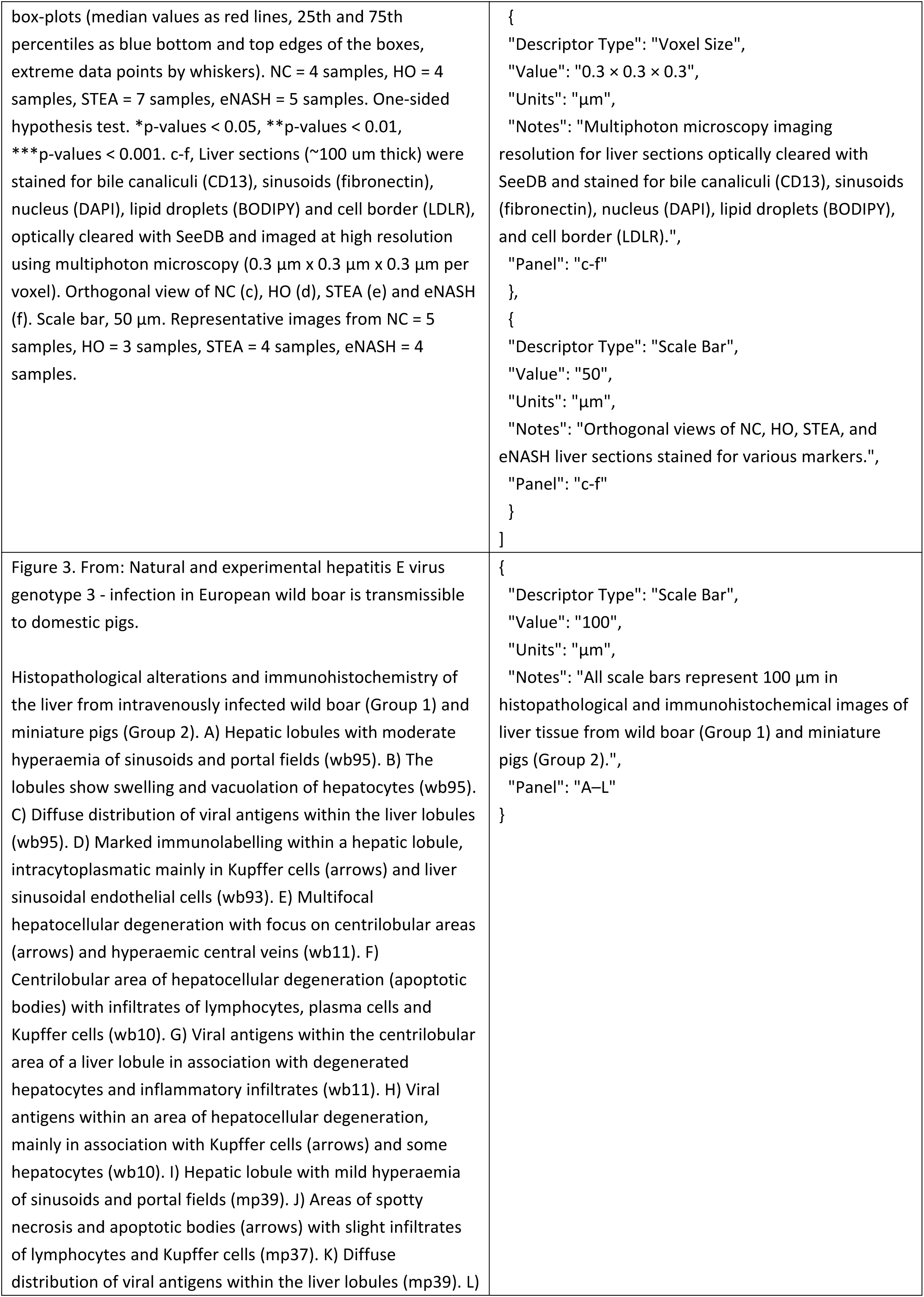
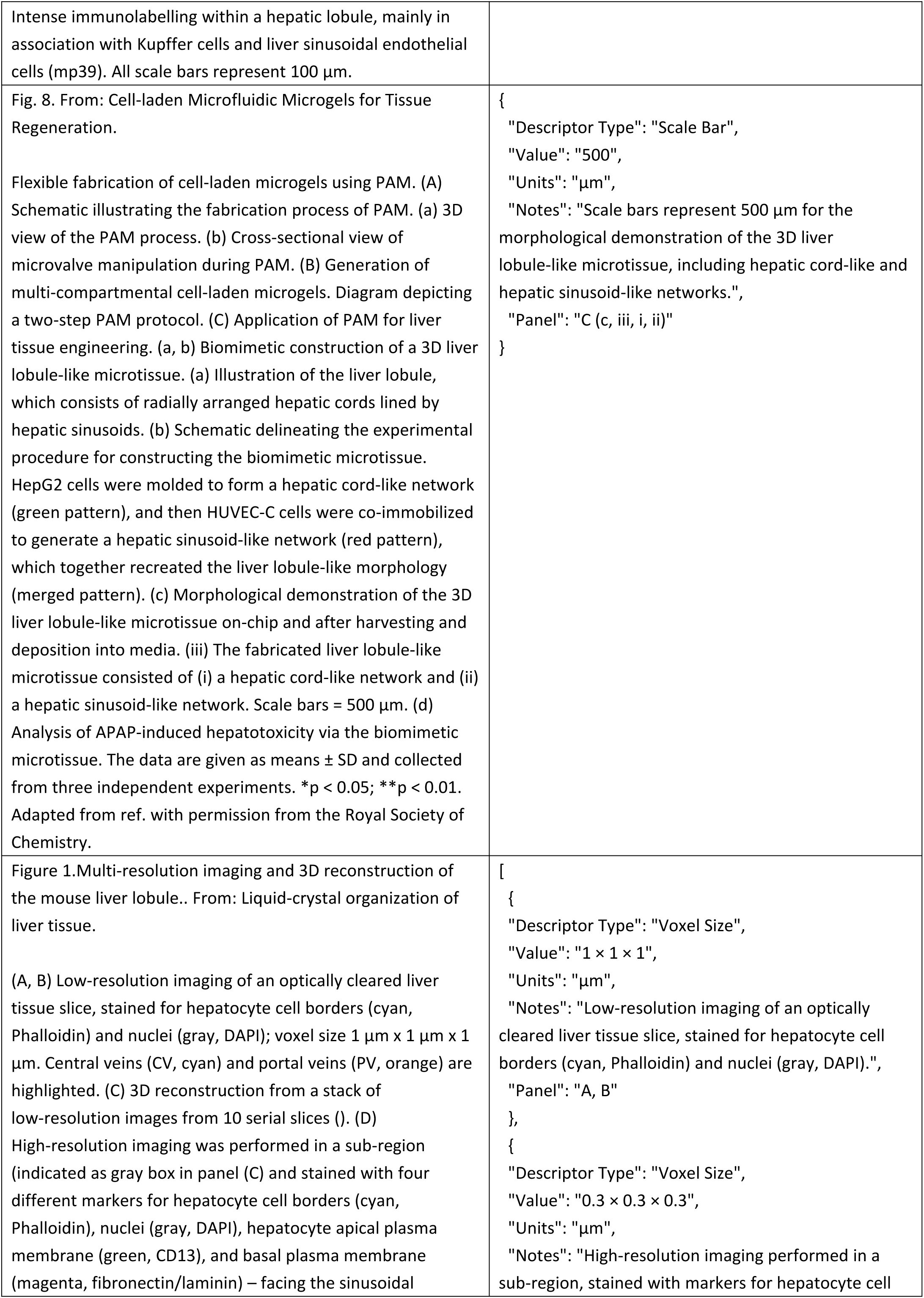
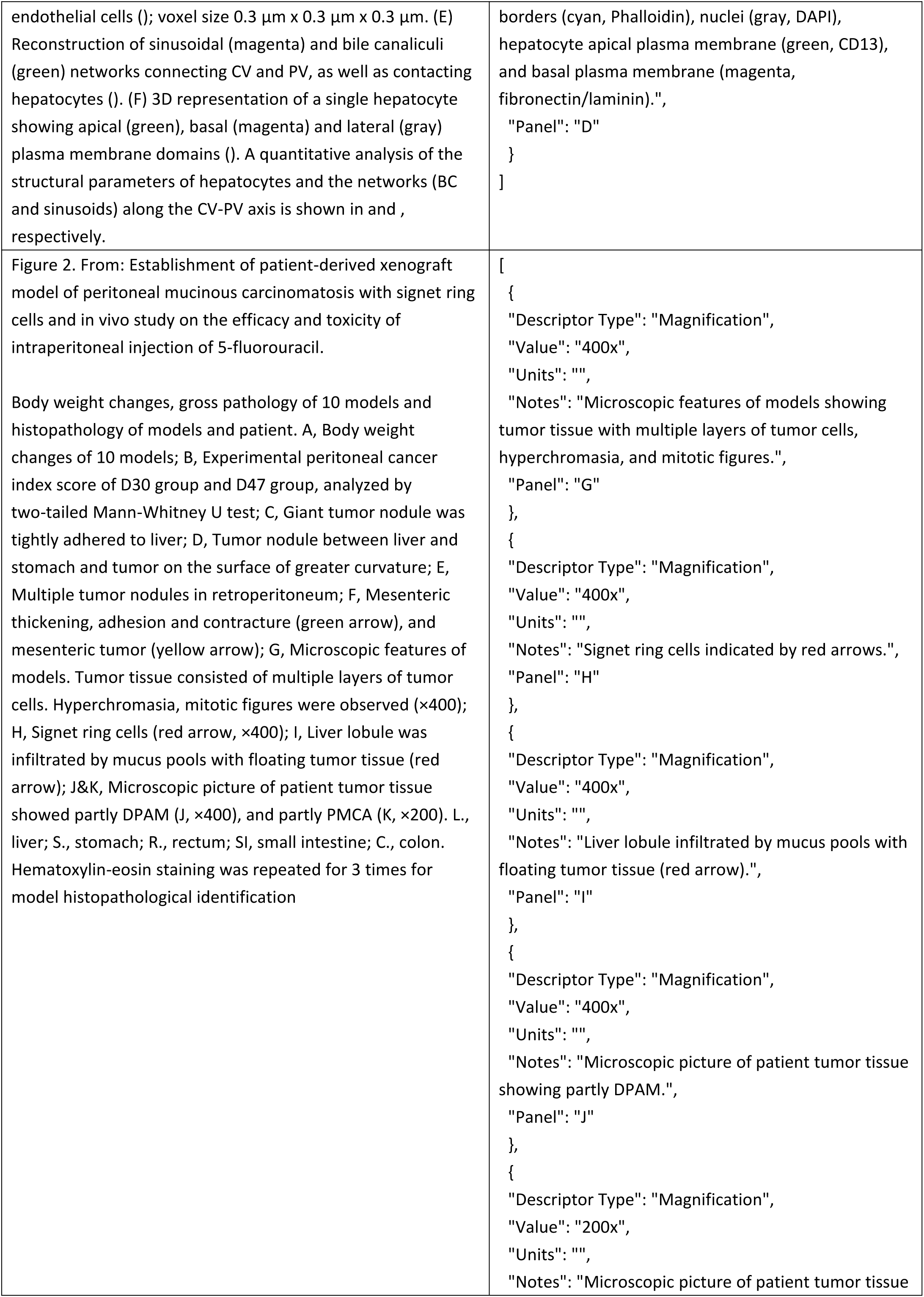
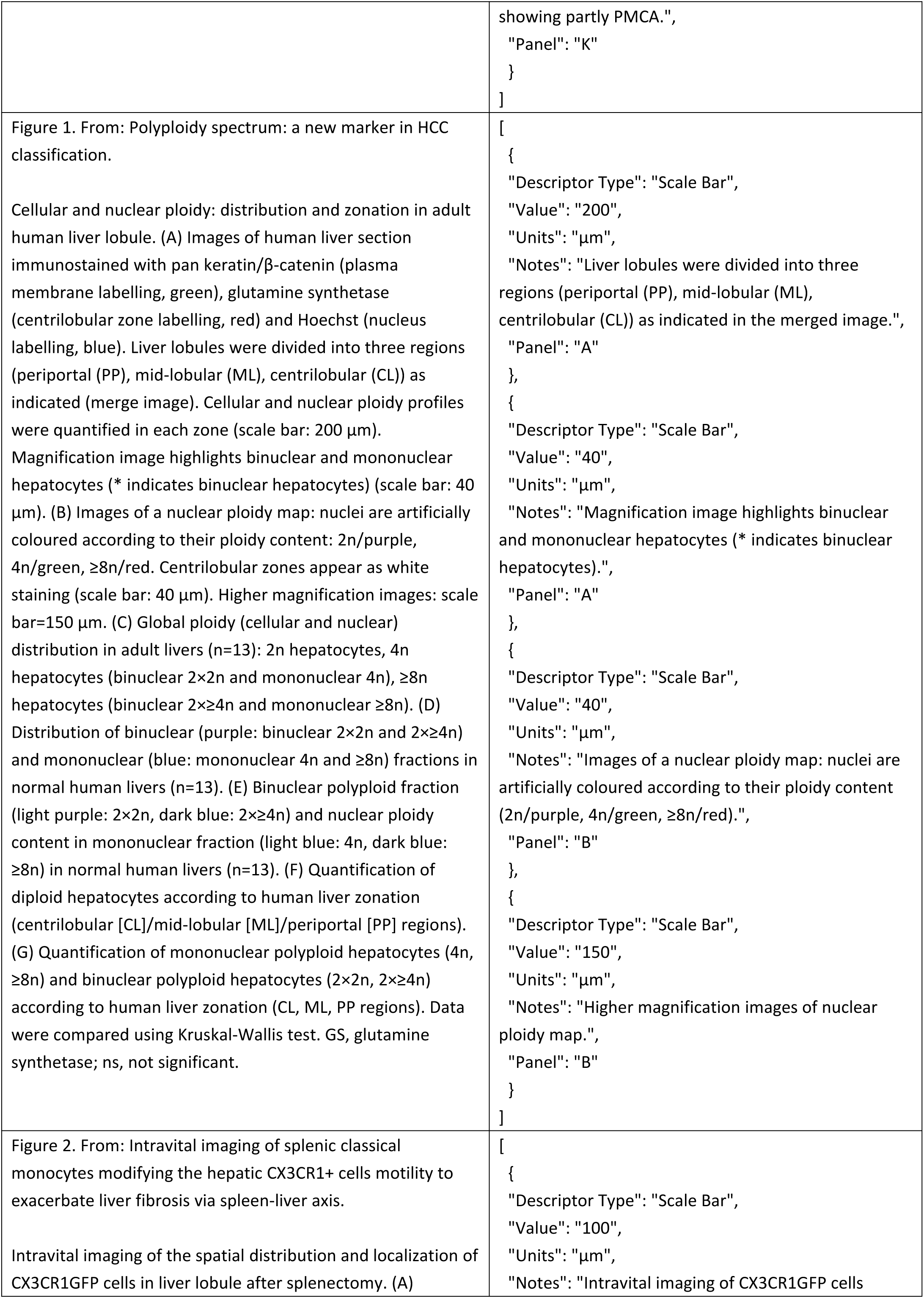
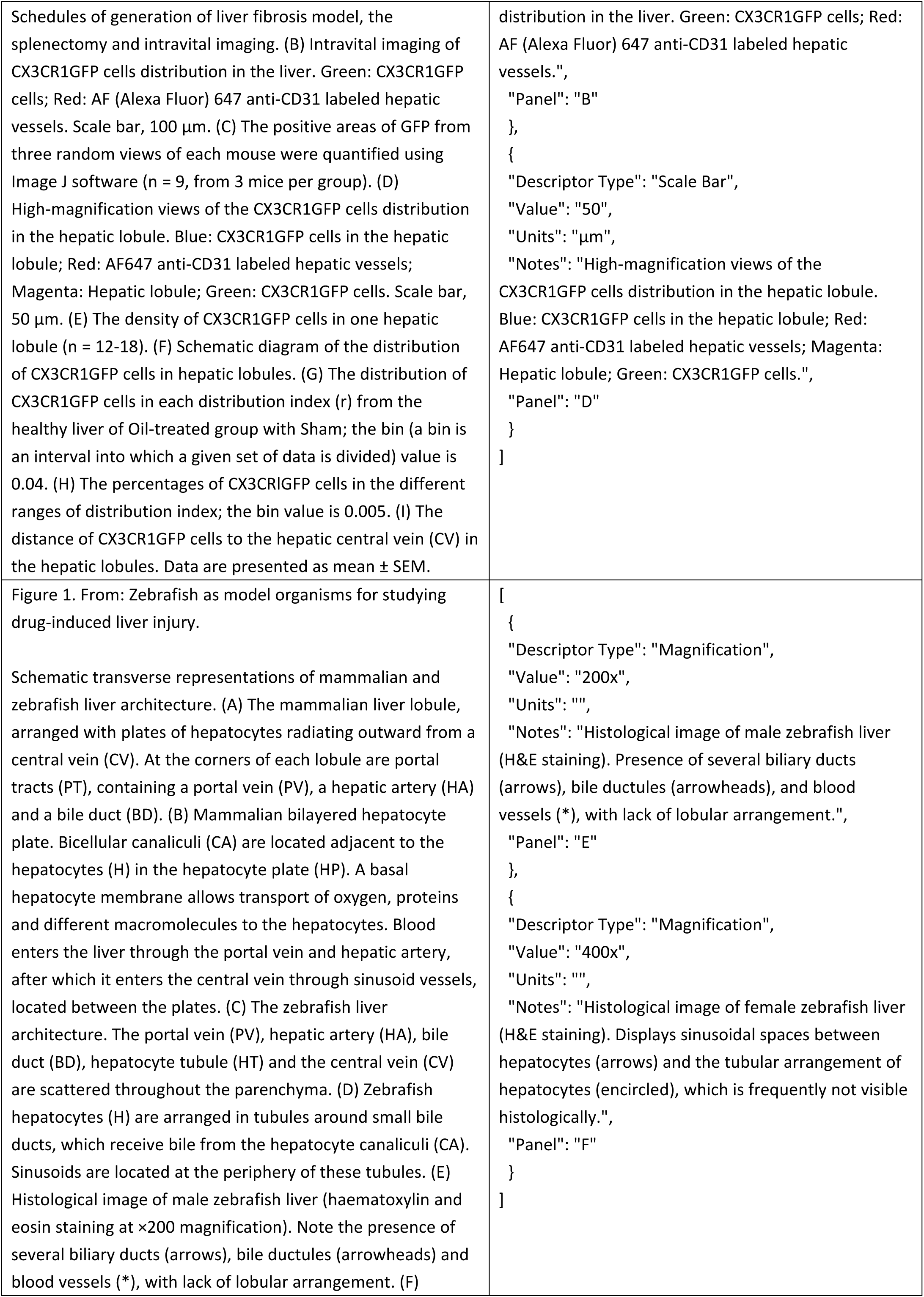
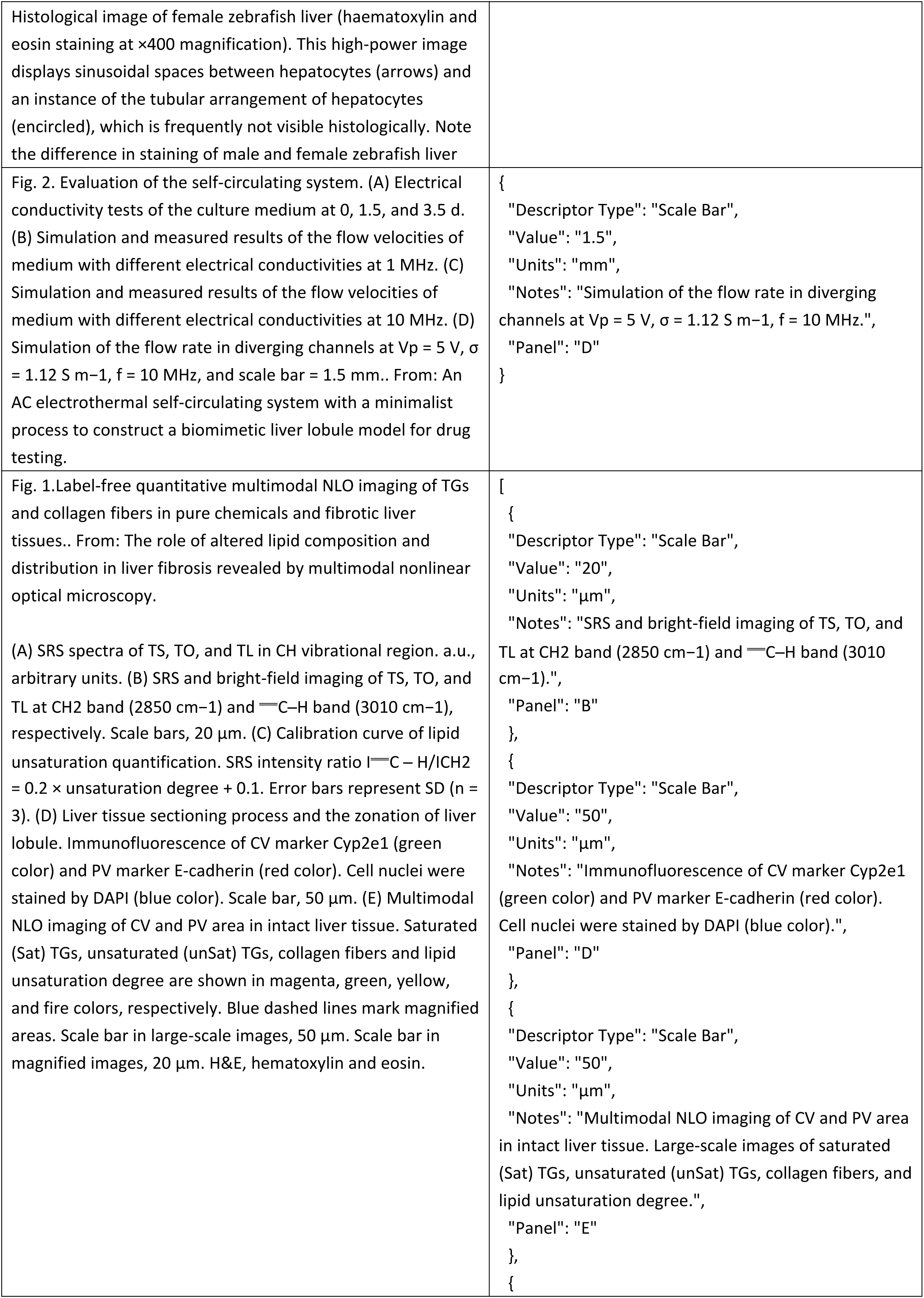
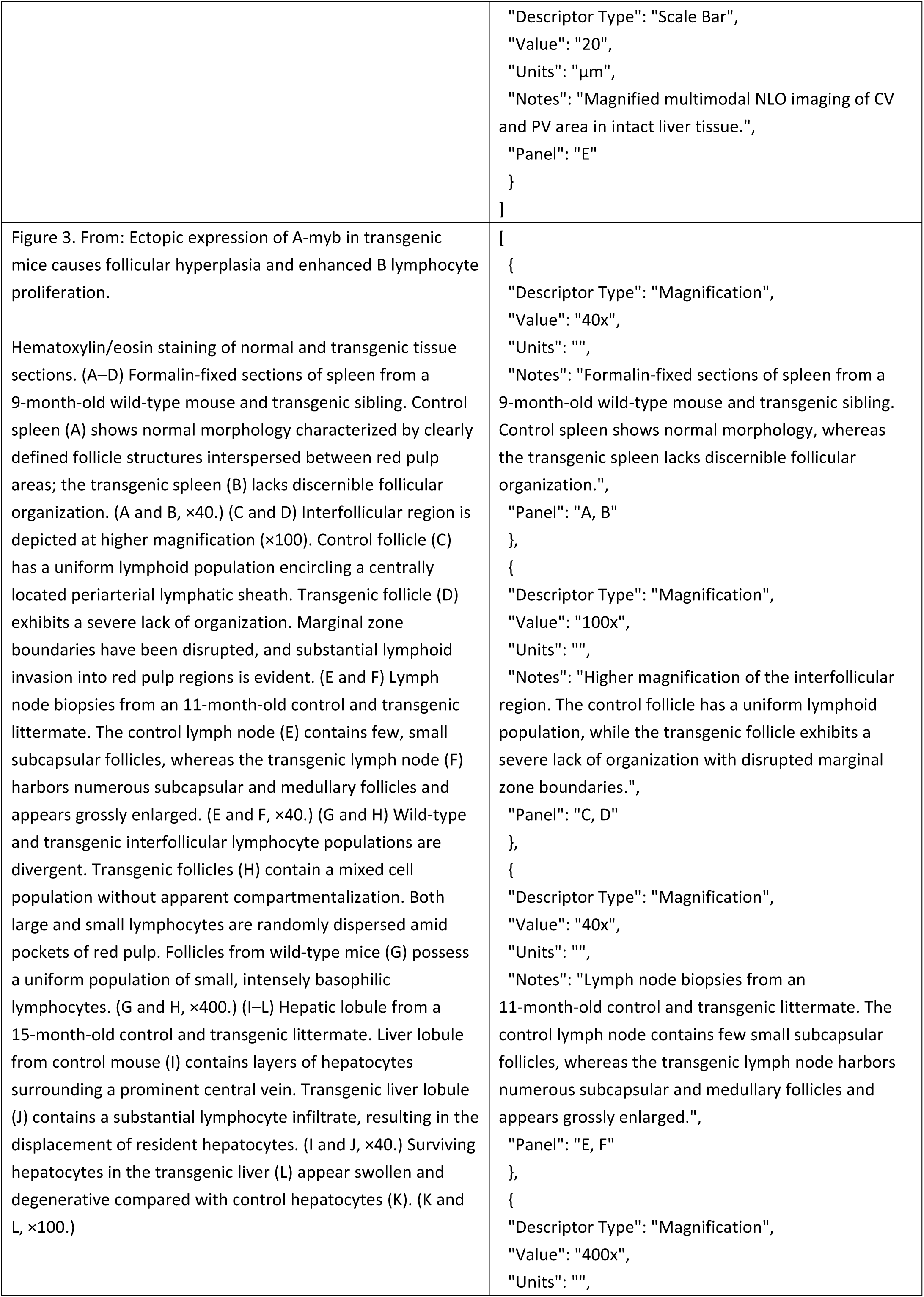
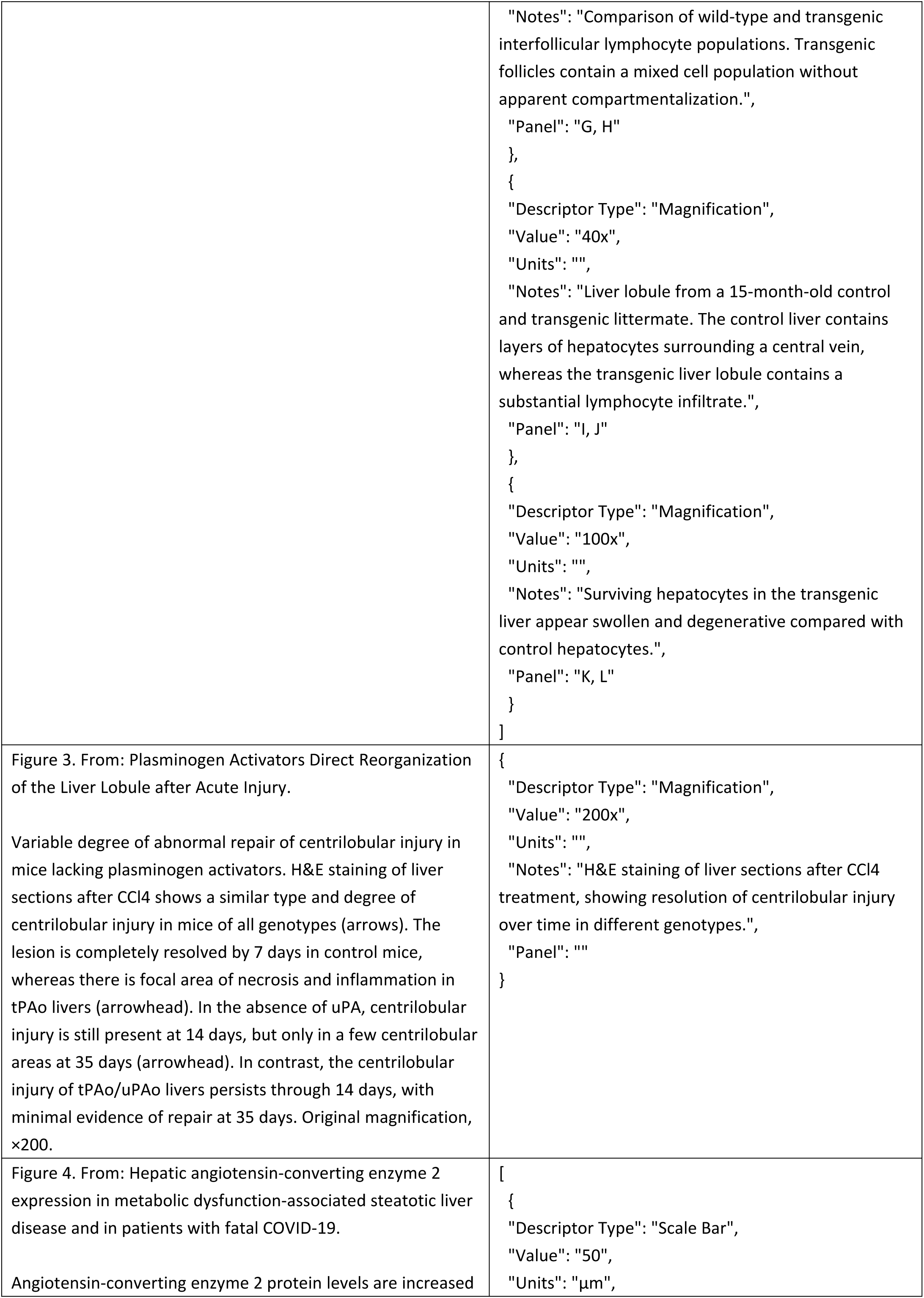
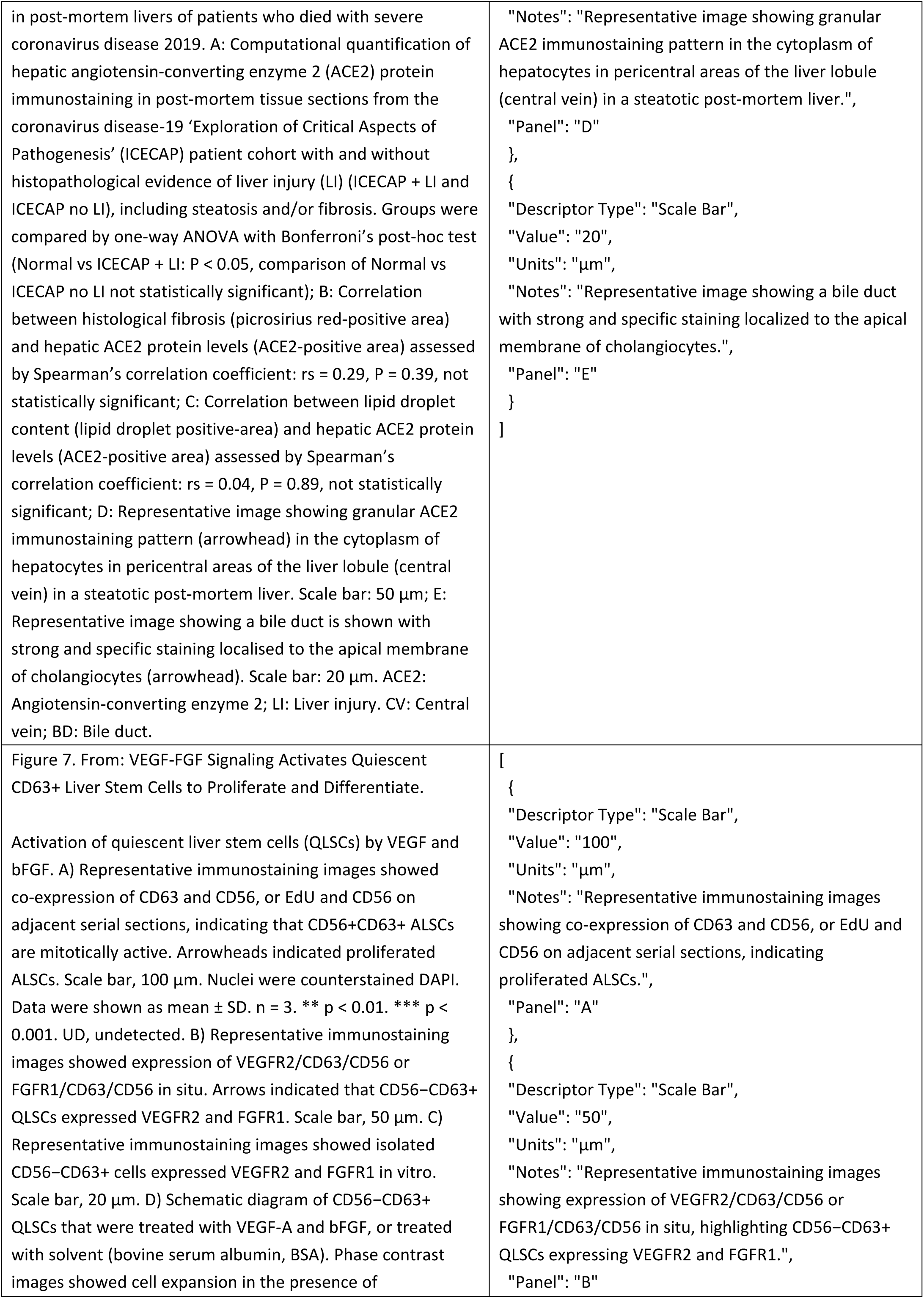
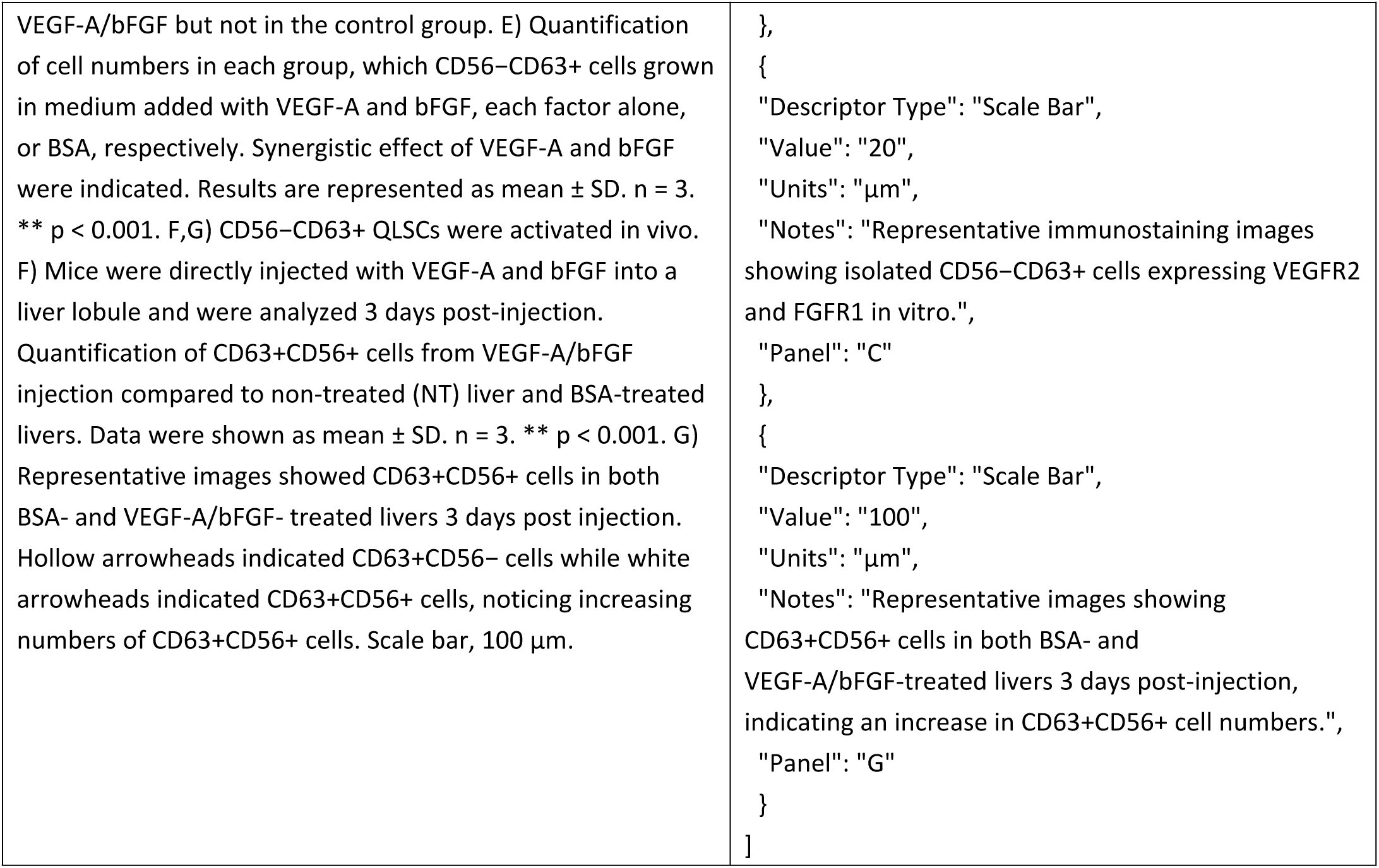
Ground true for scale-bar extraction.

**Supplementary Table 19.**
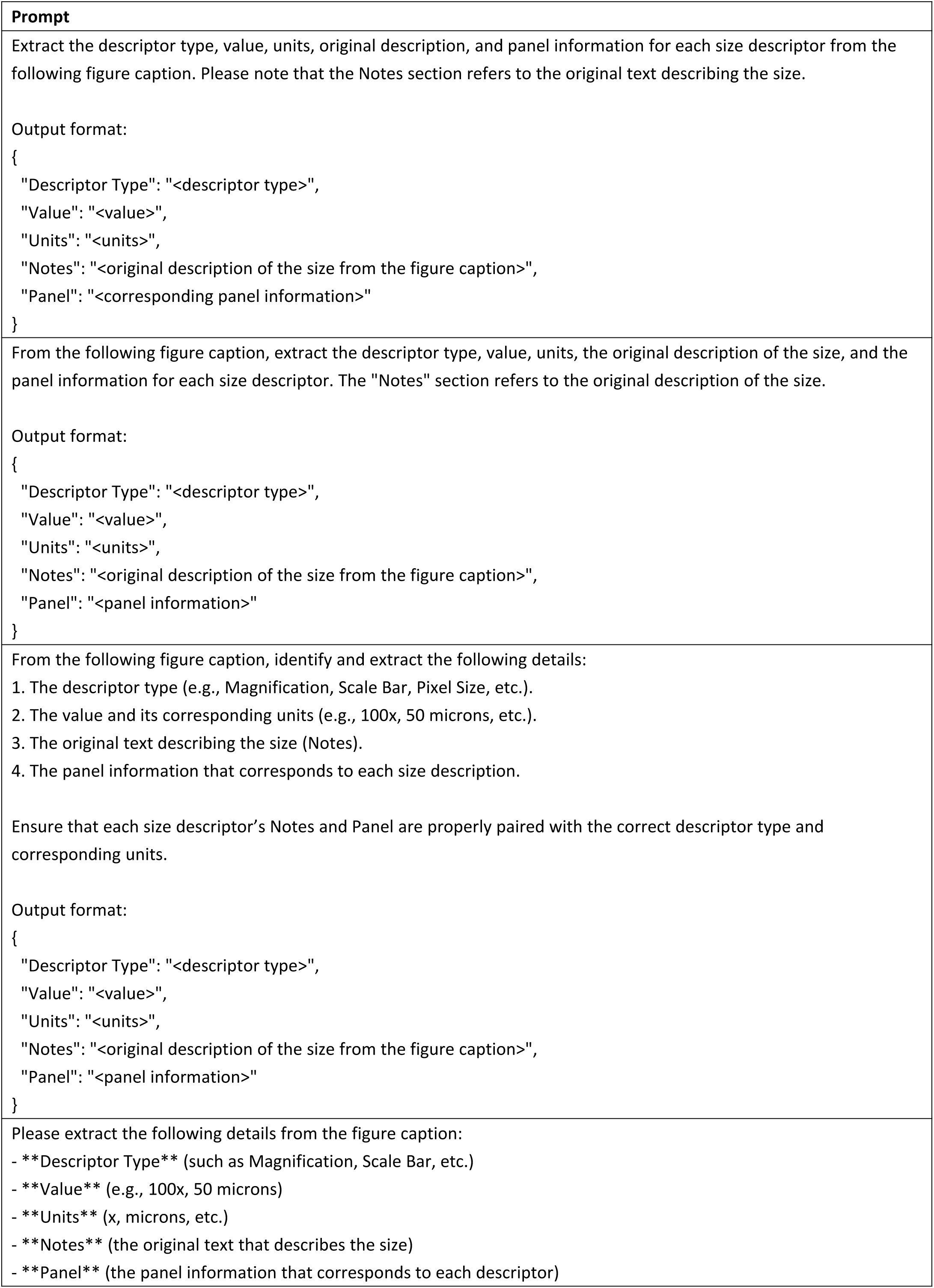
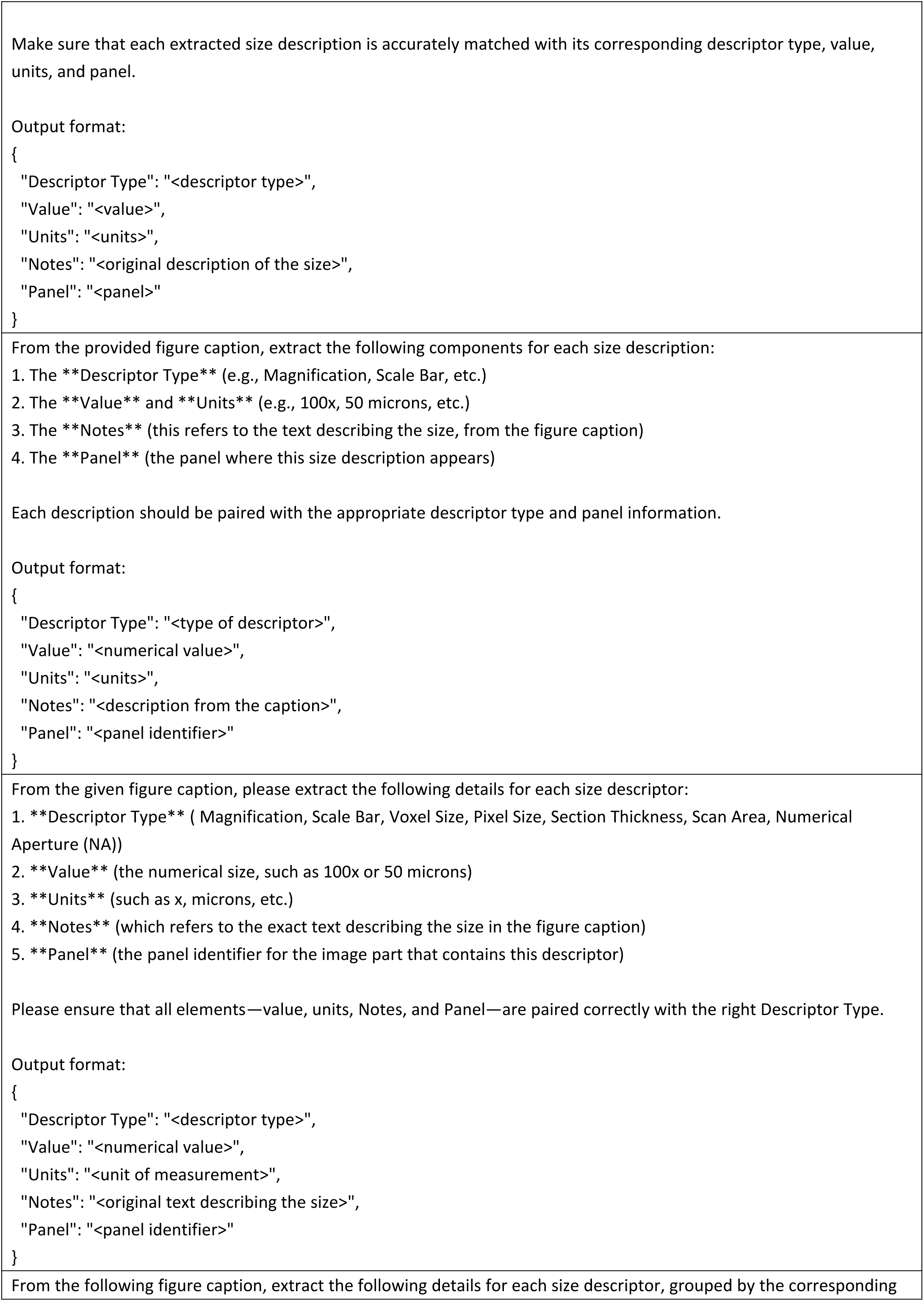
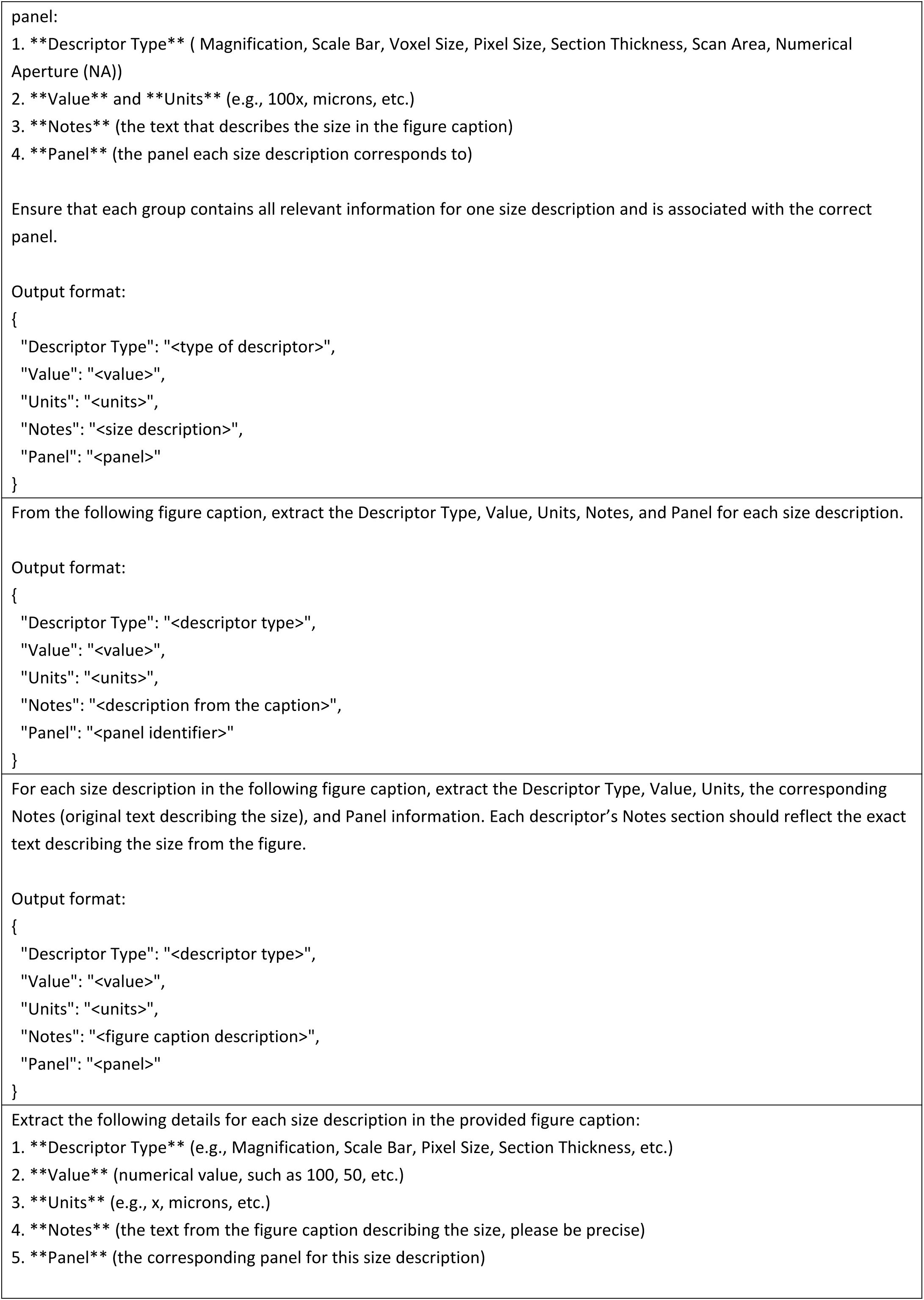
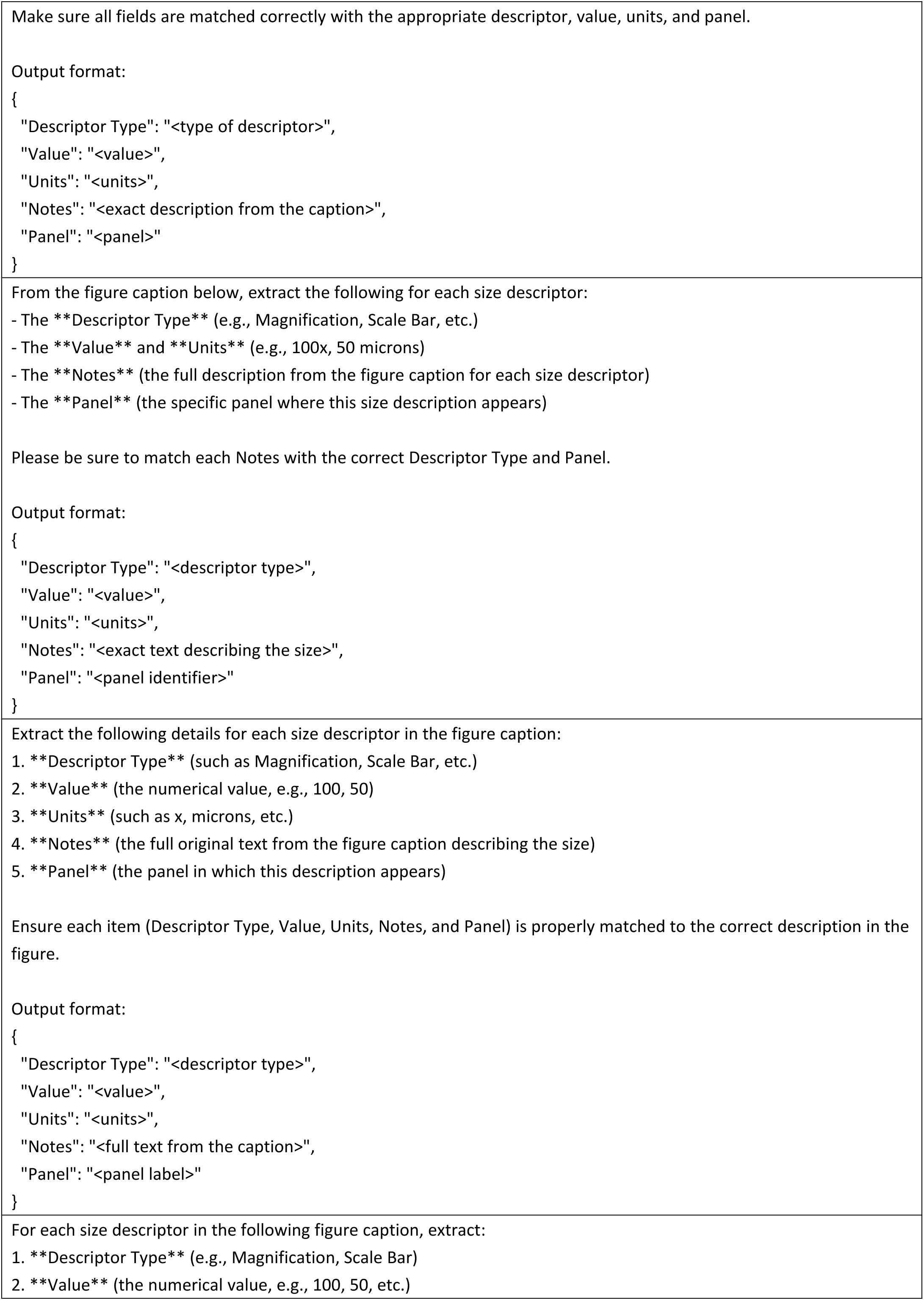
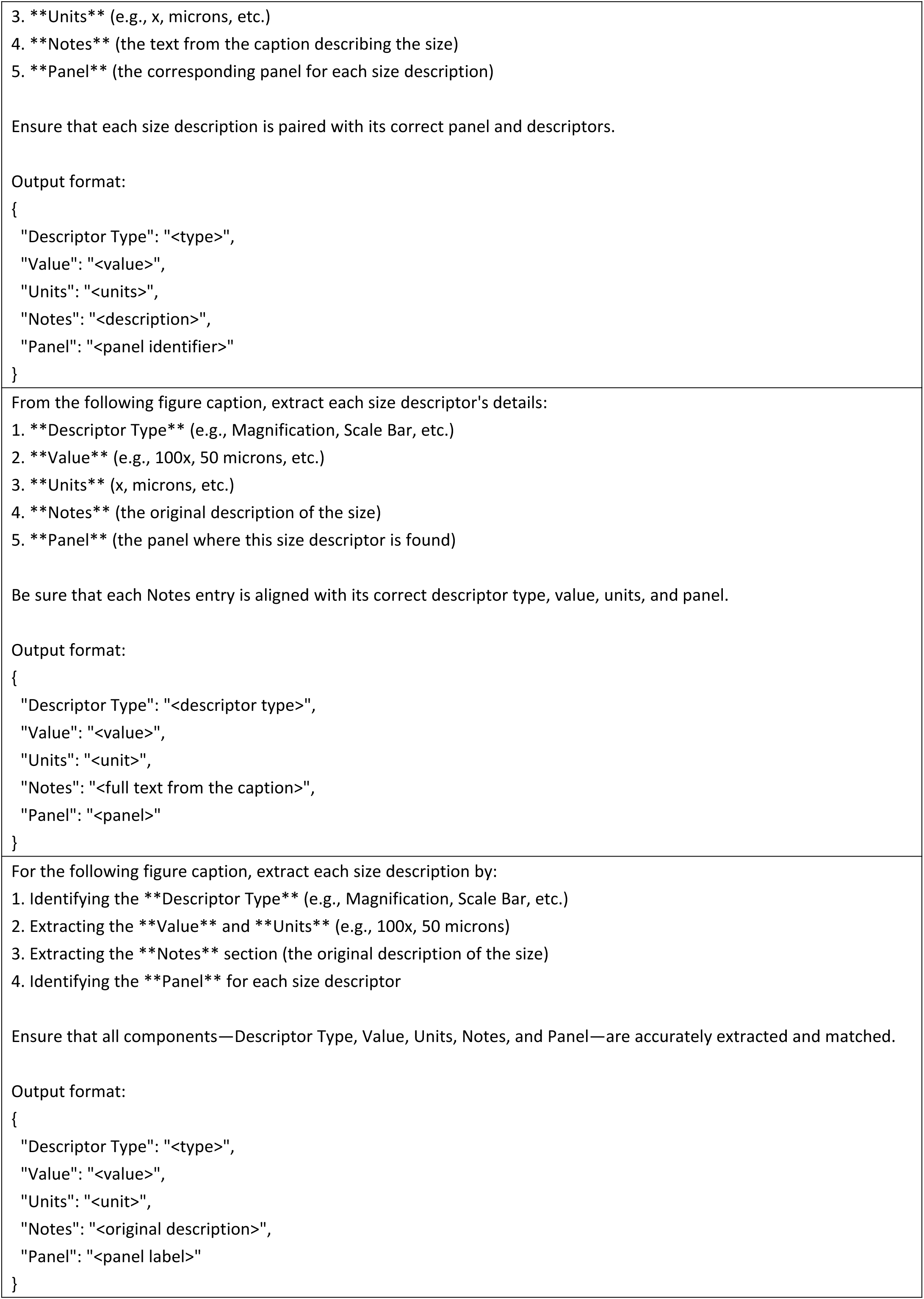
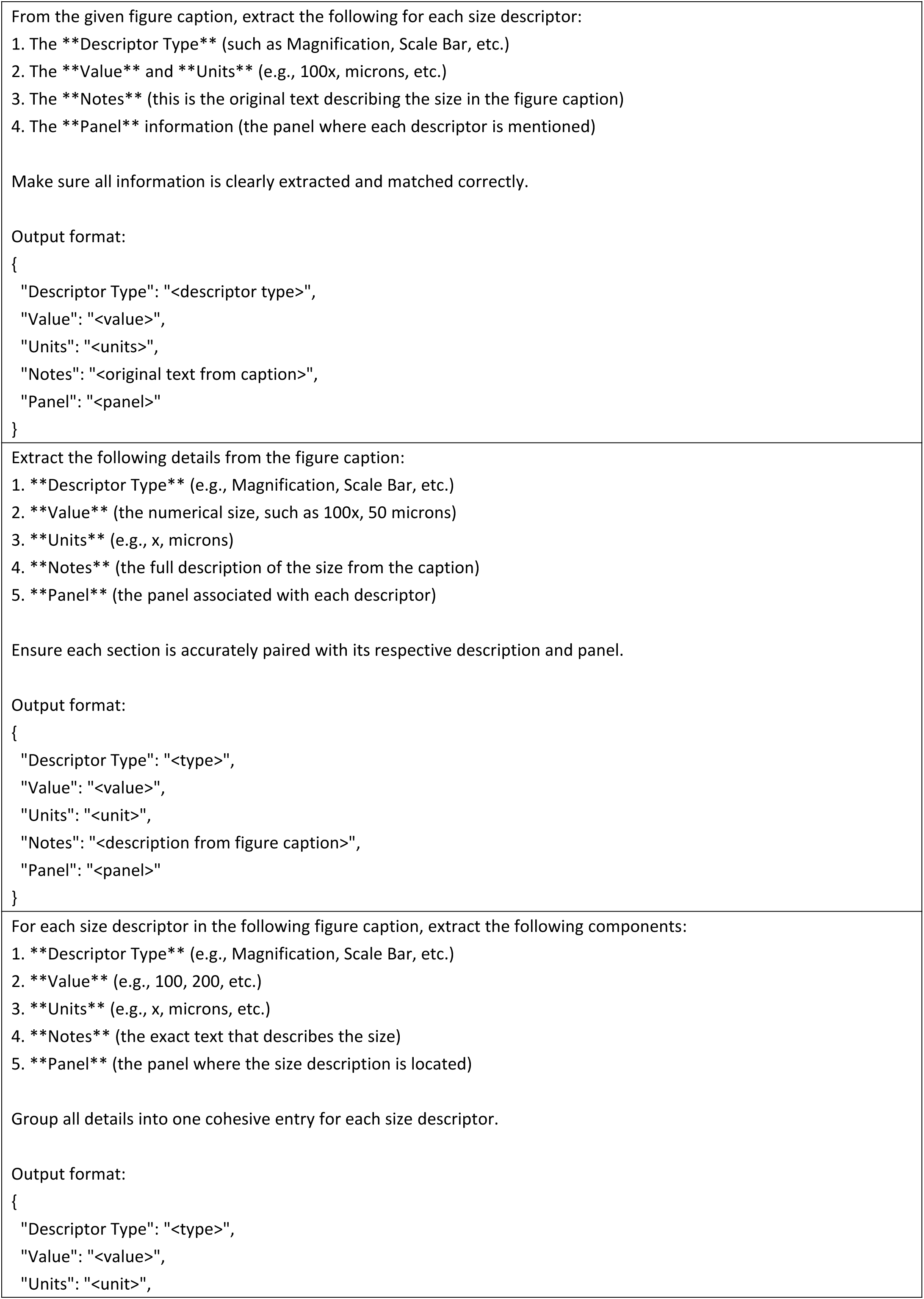
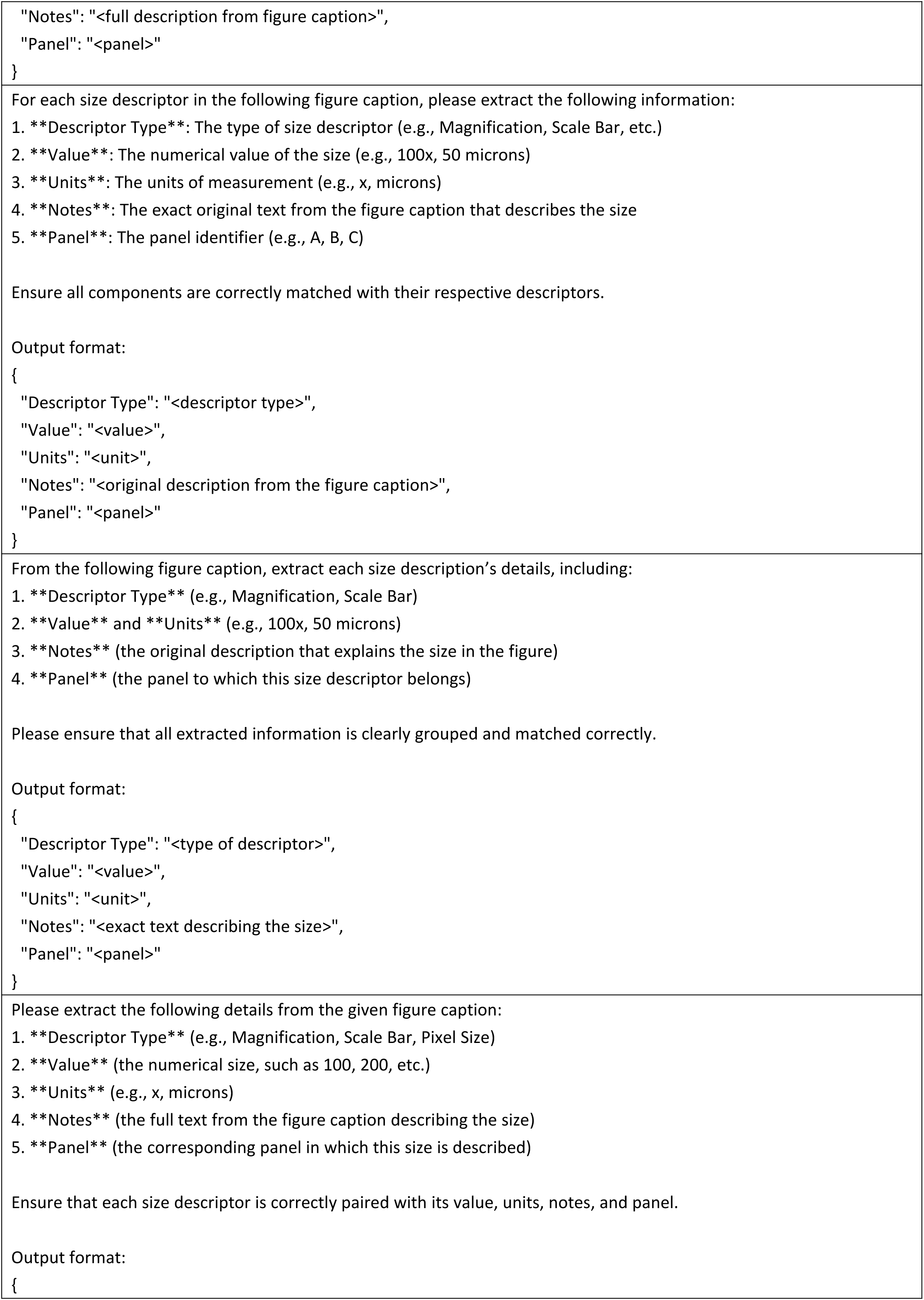
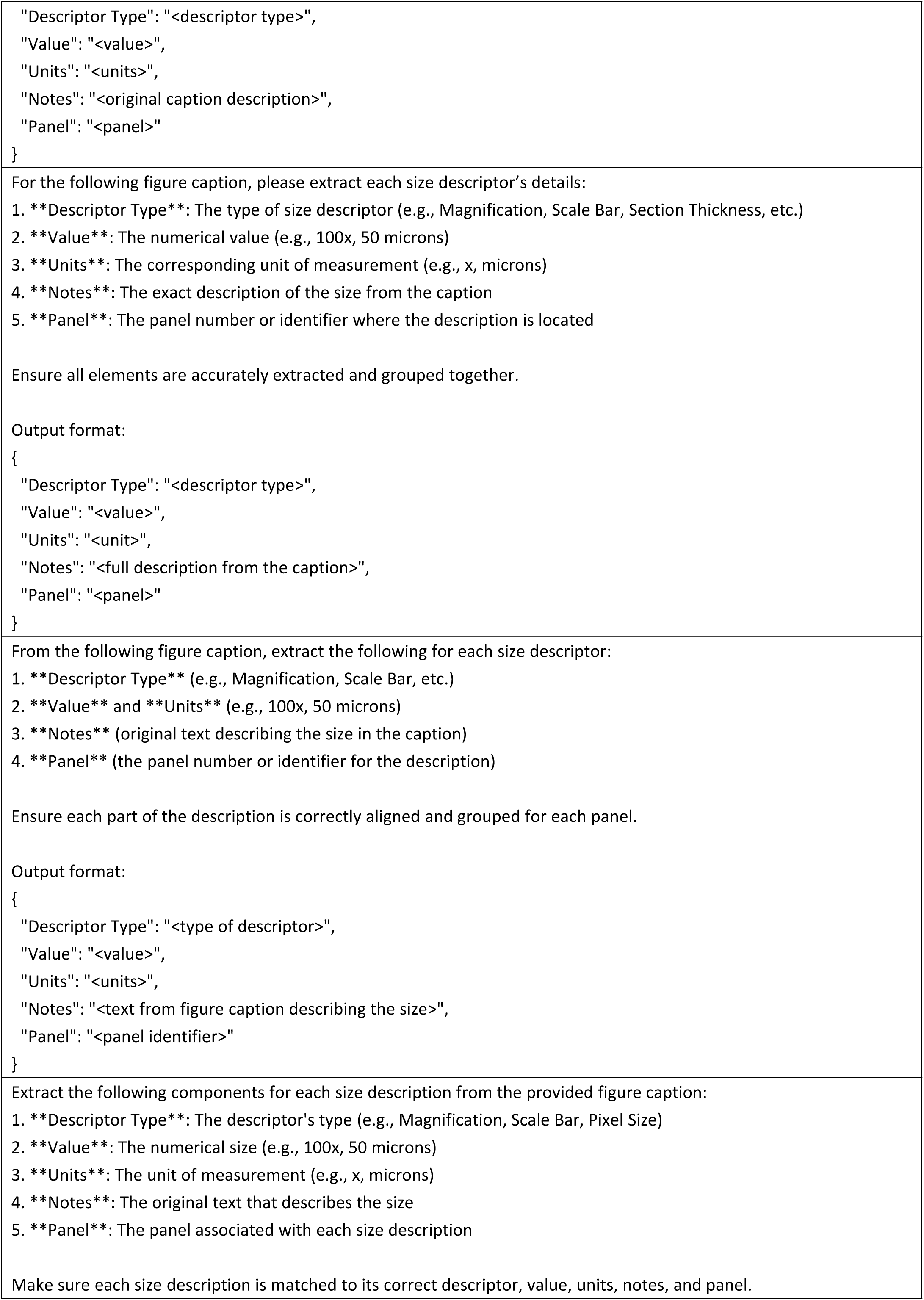
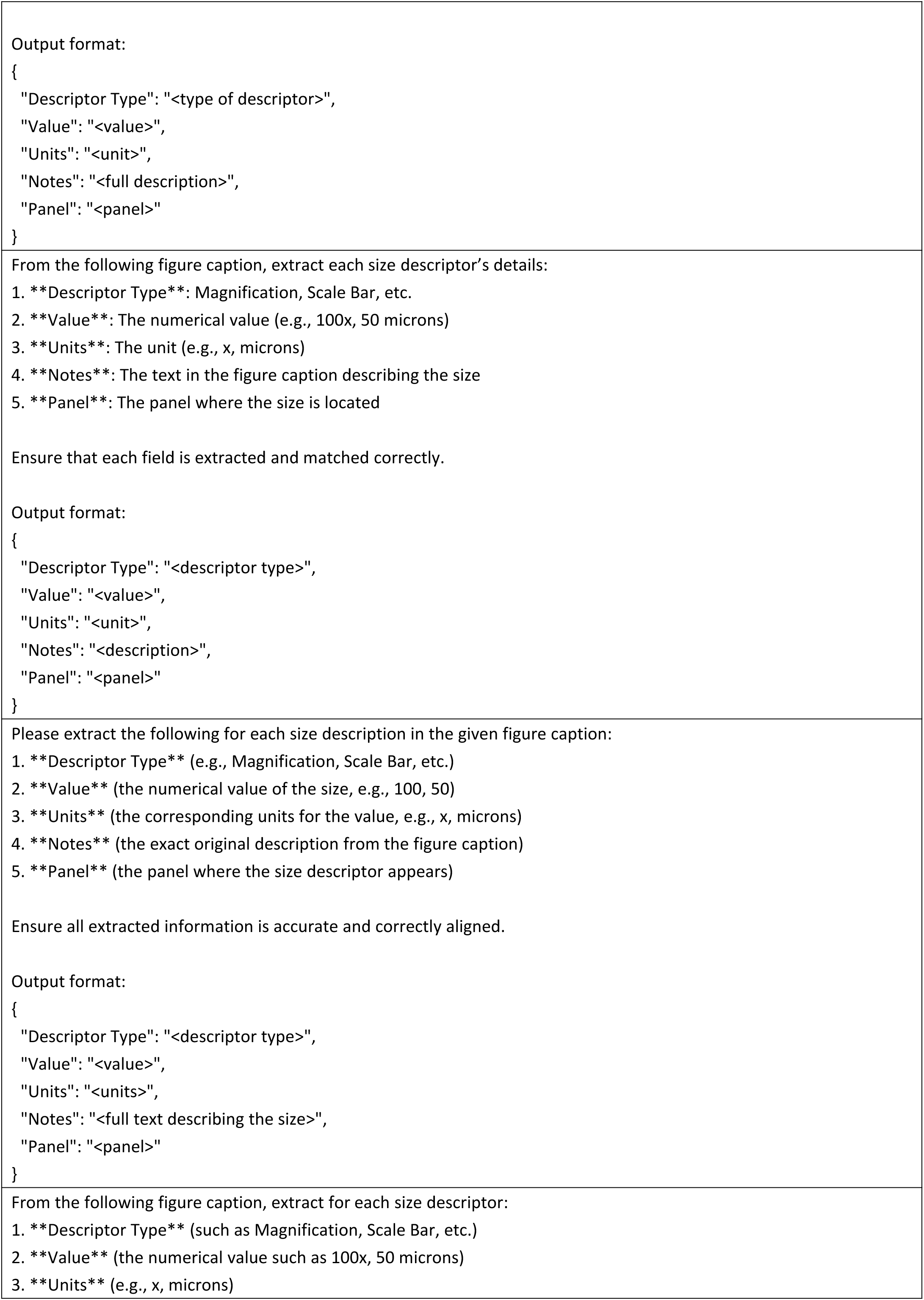
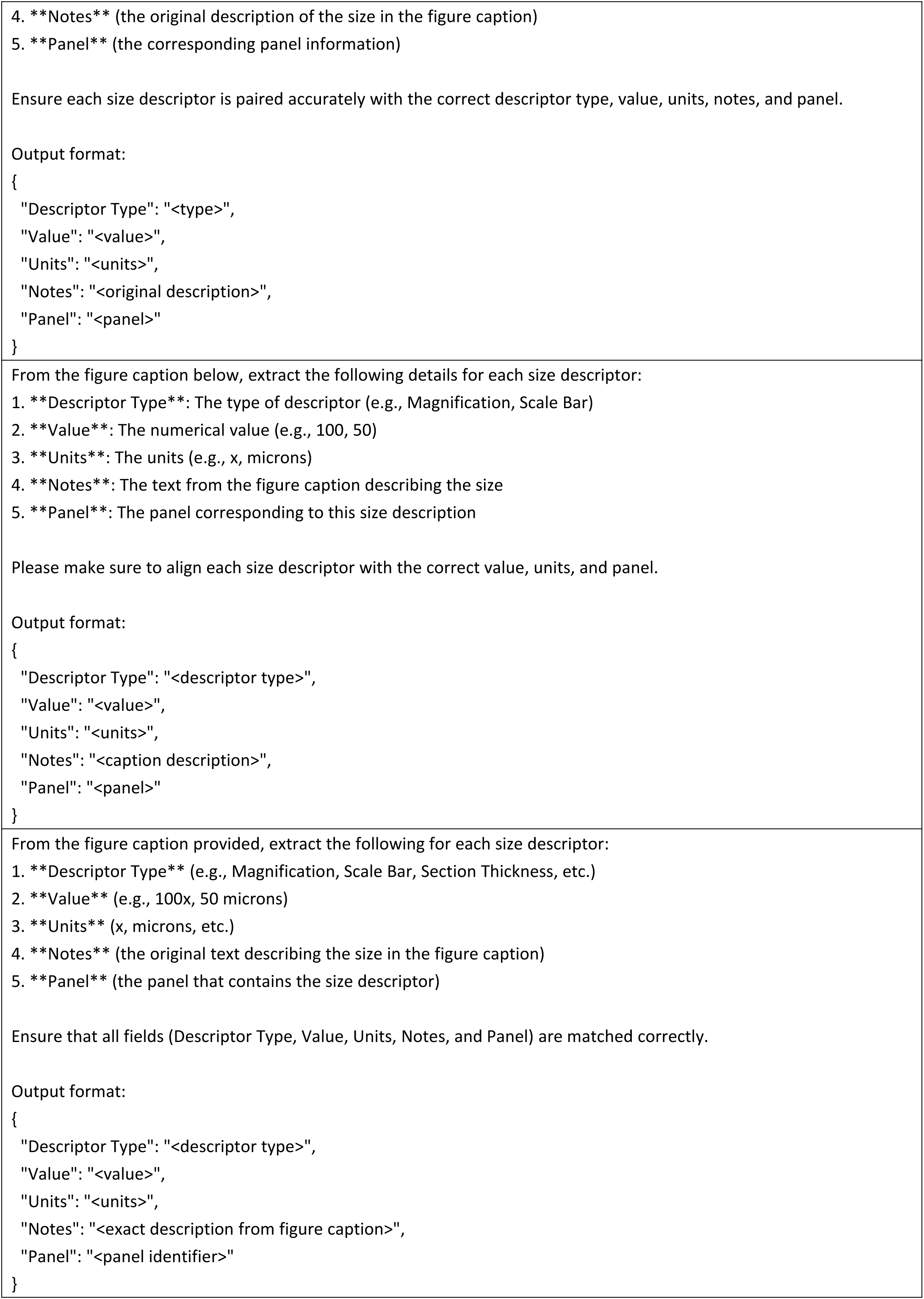
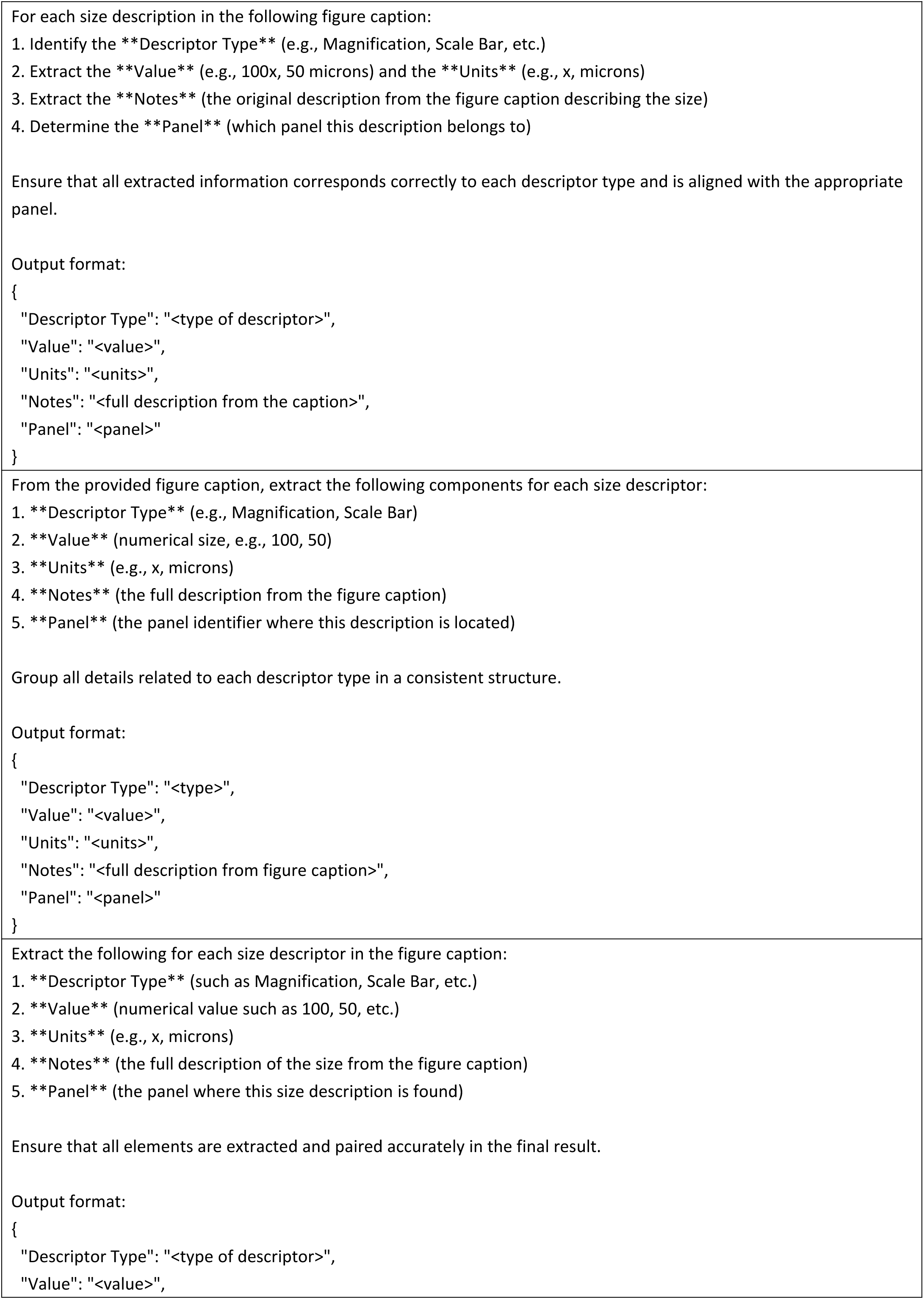
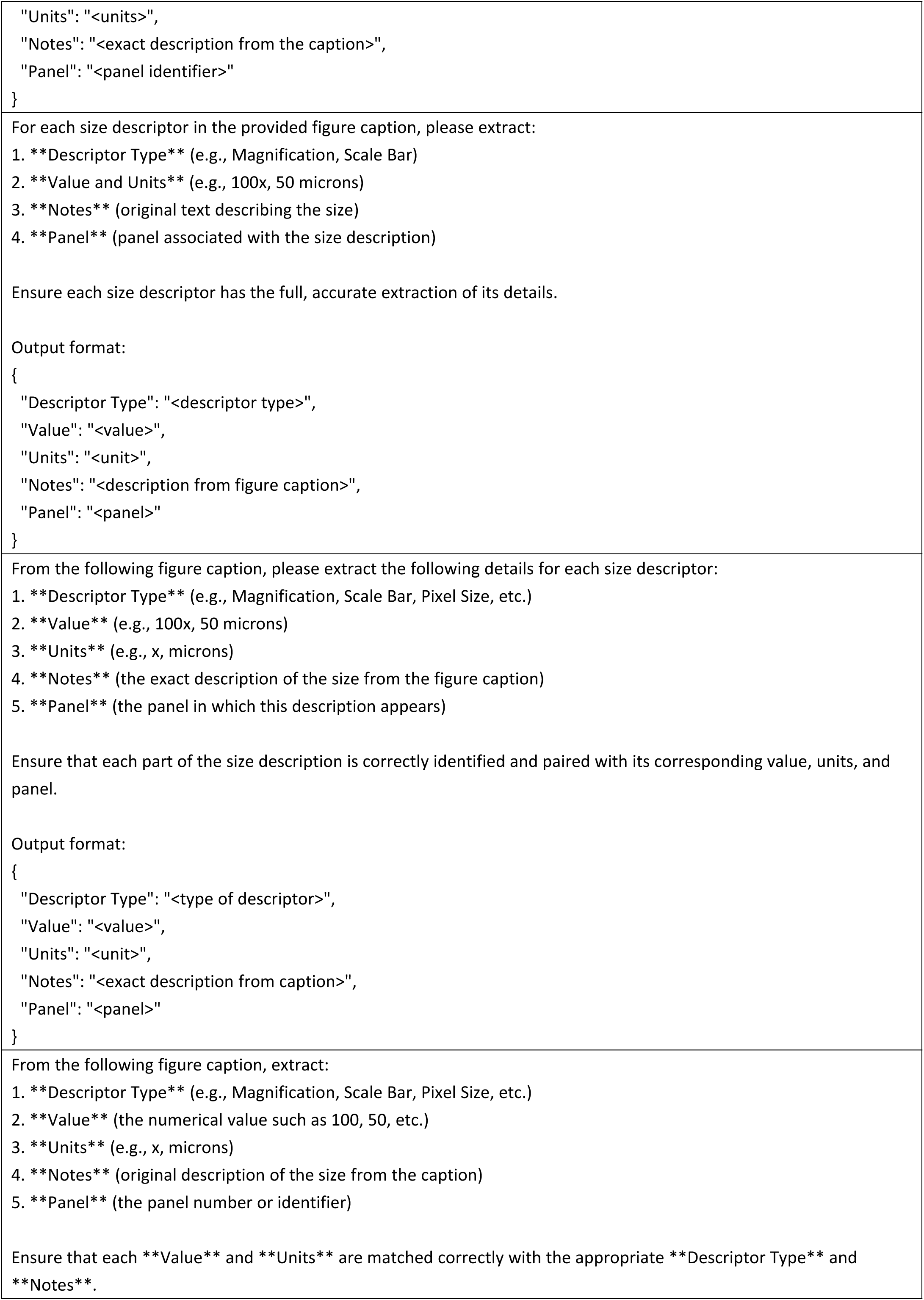
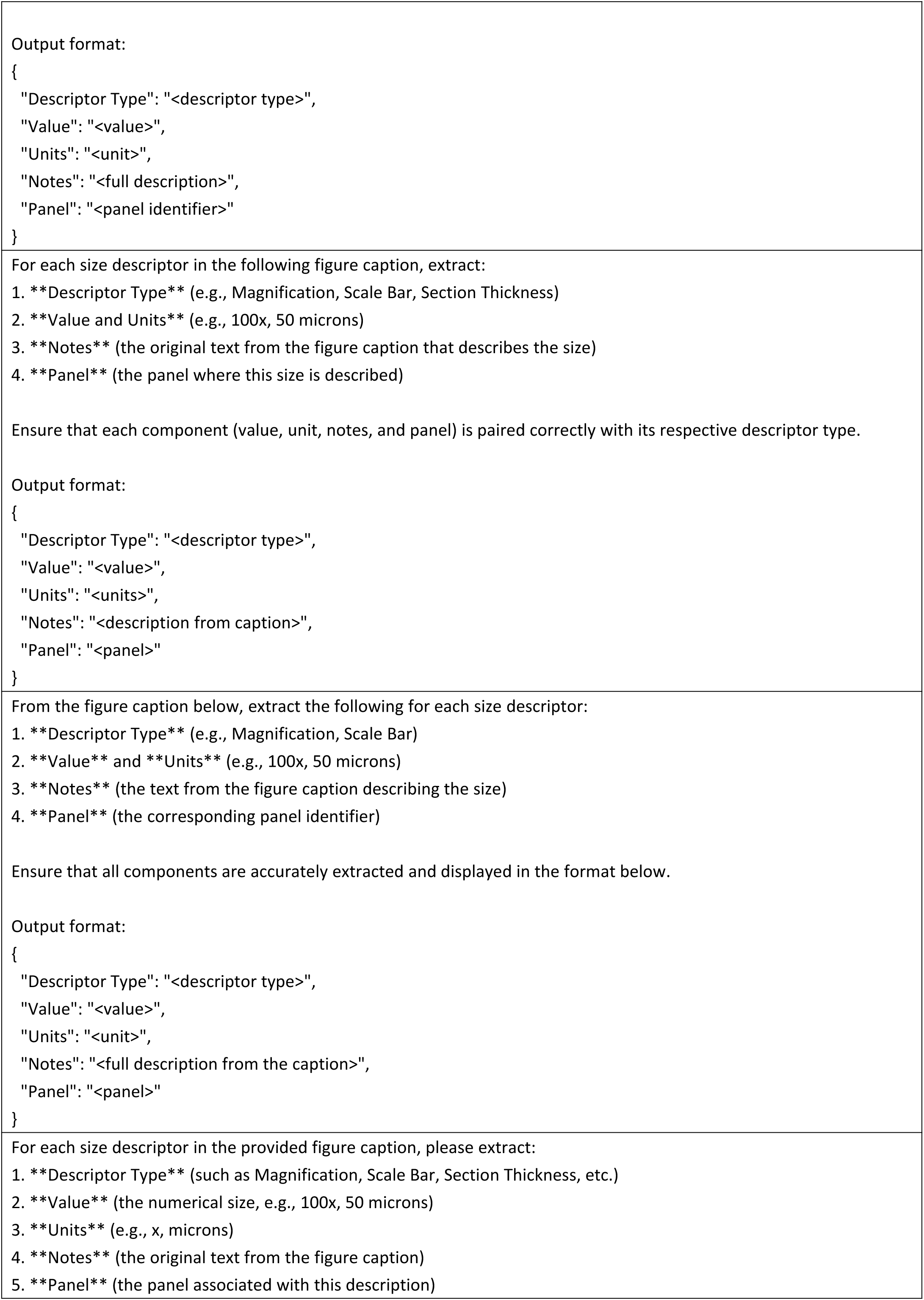
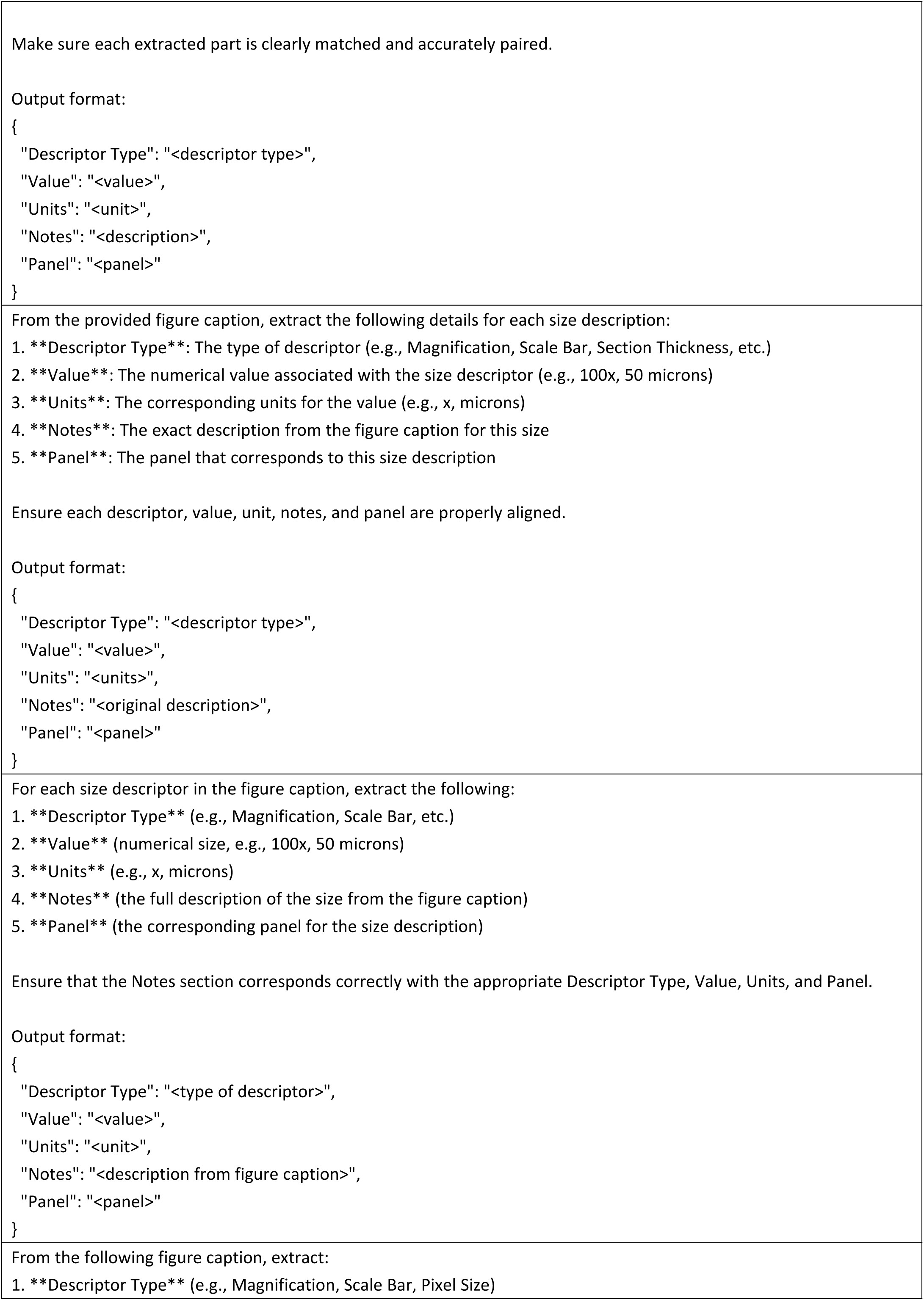
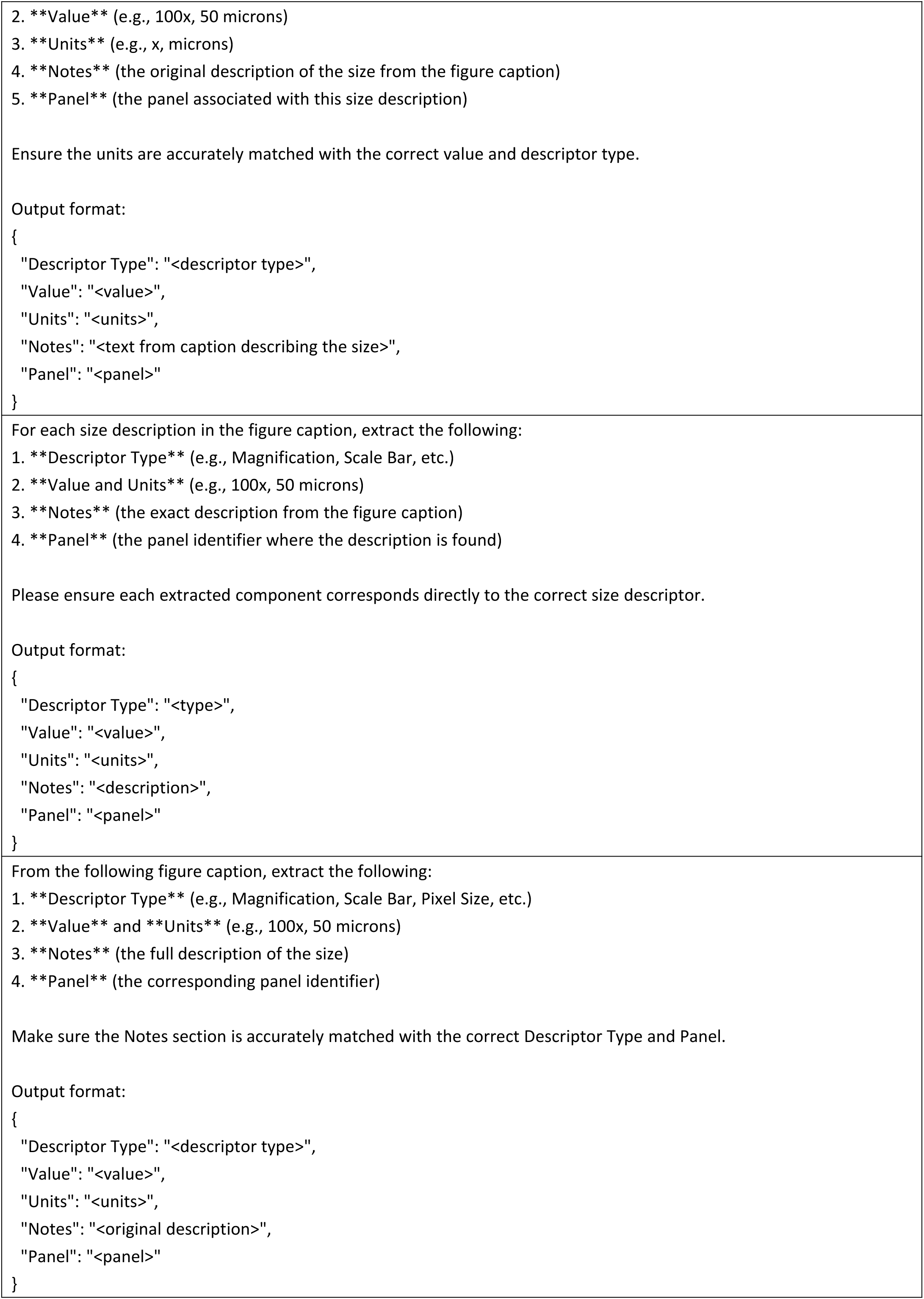
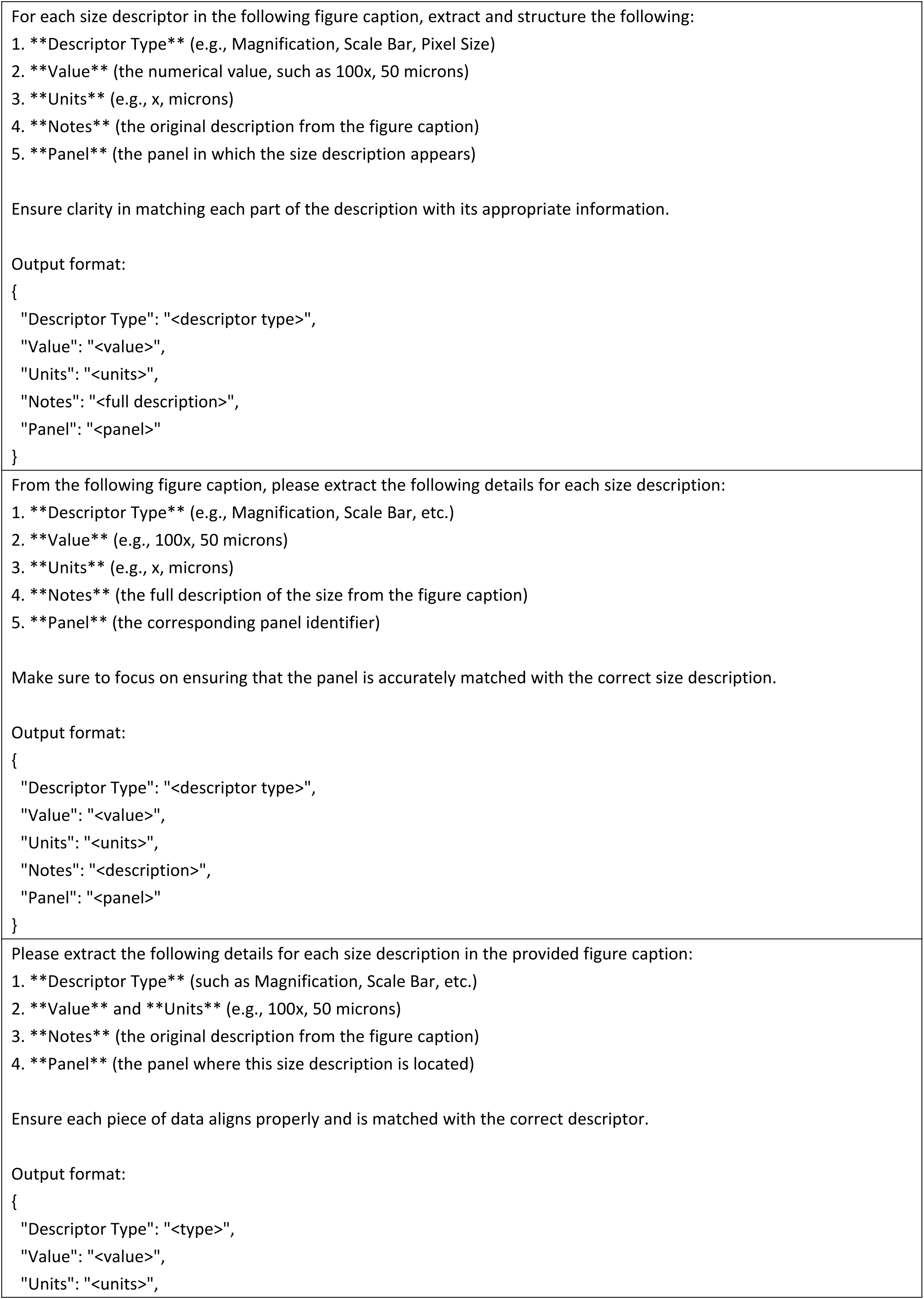
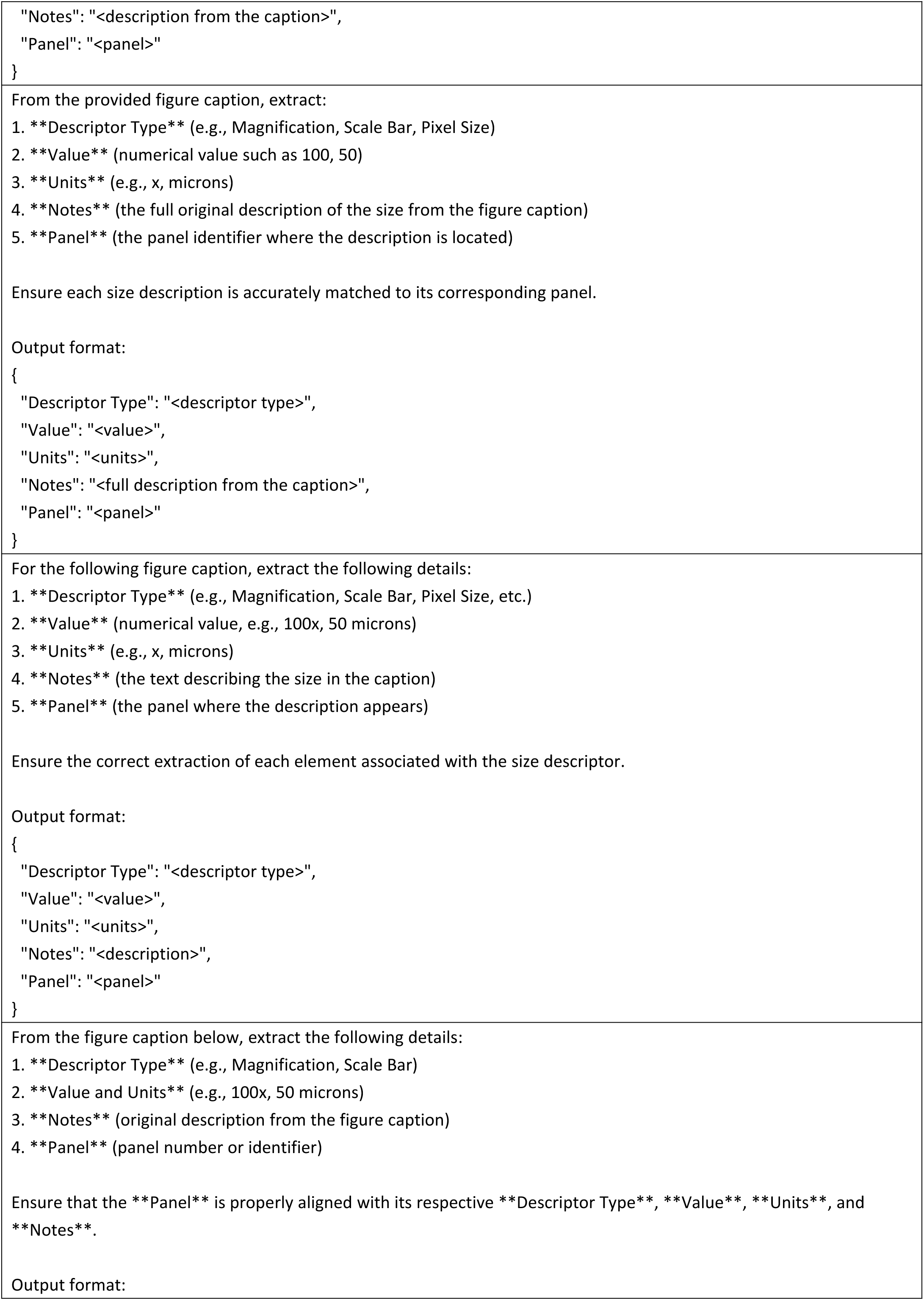
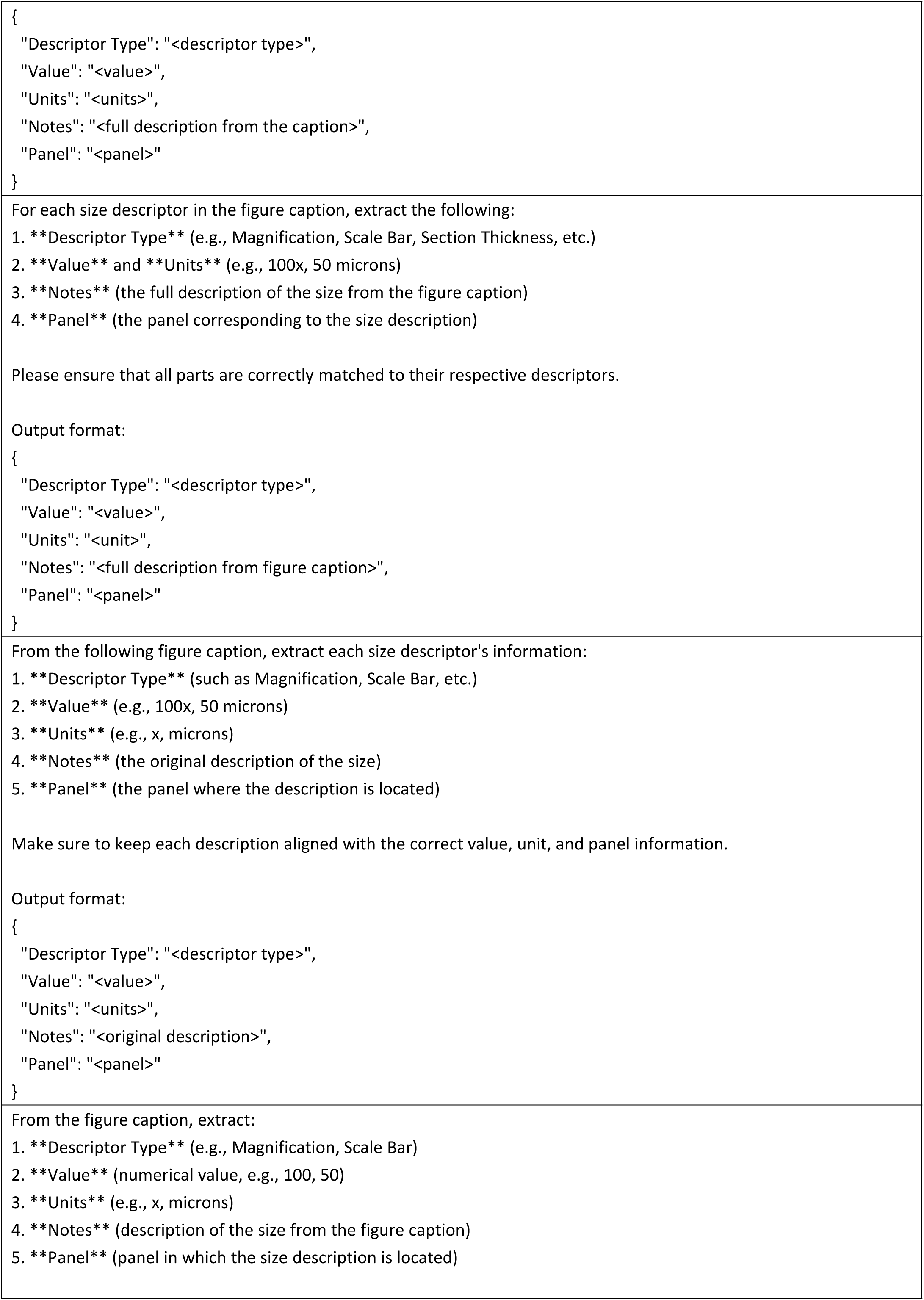
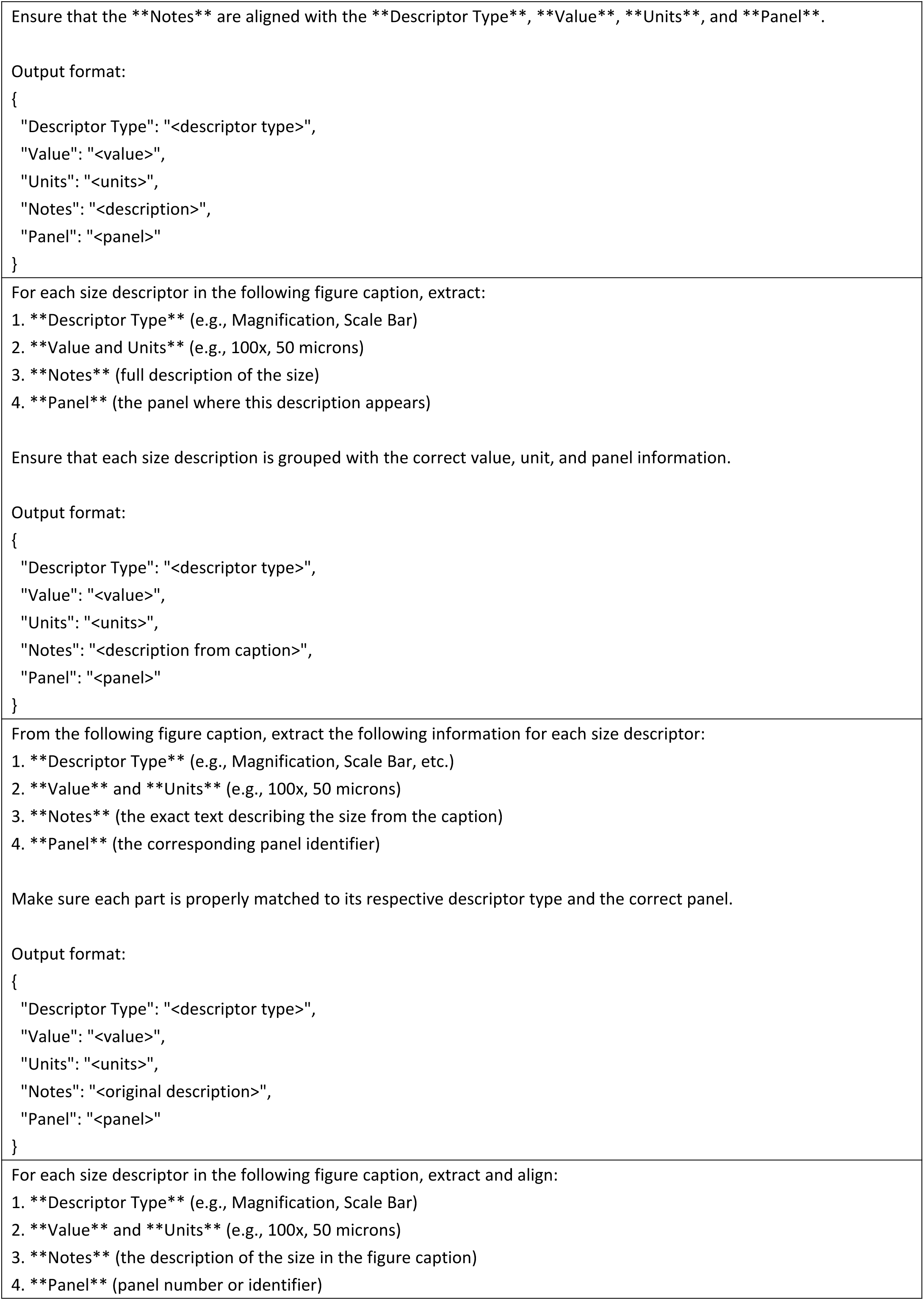
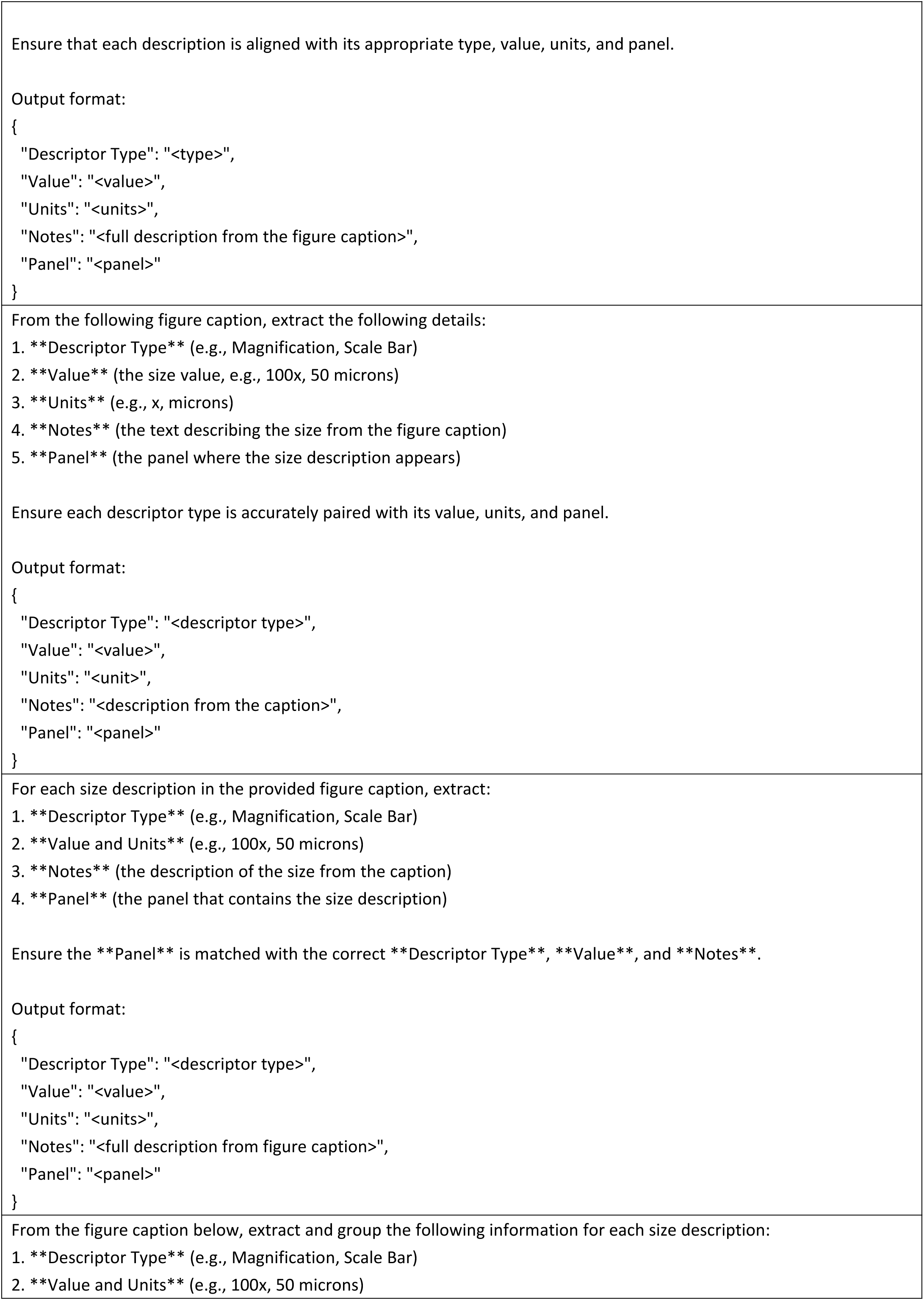
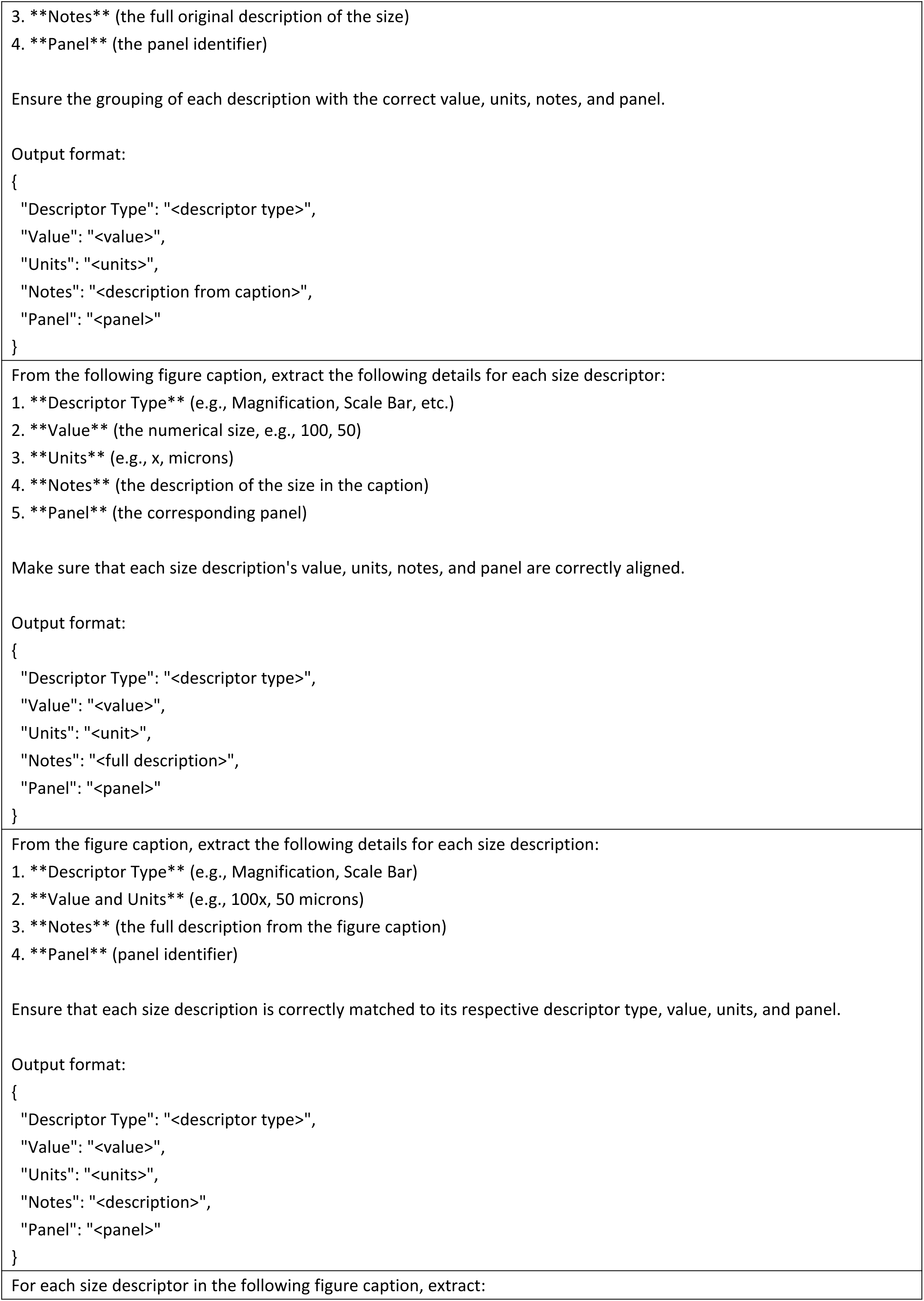
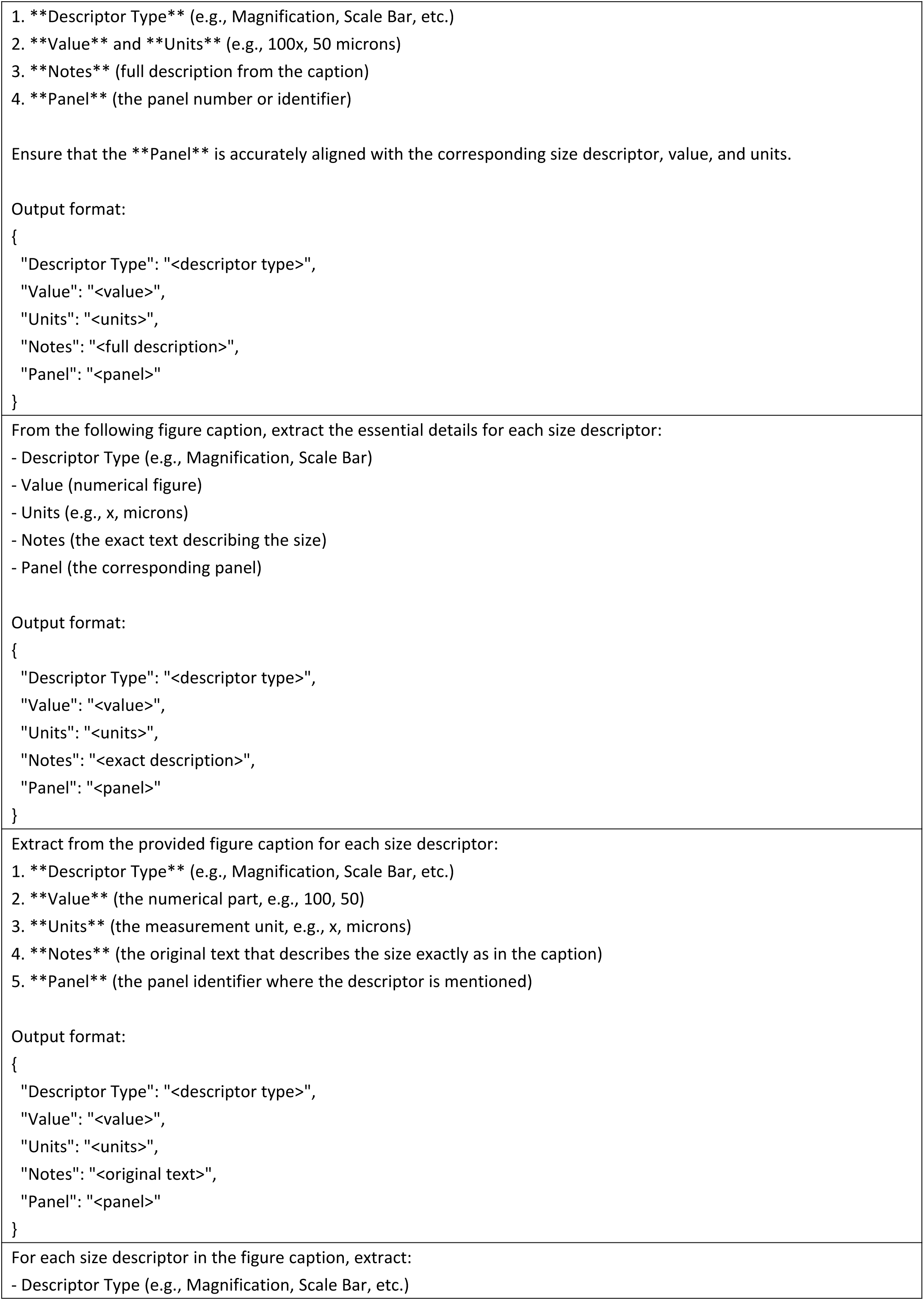
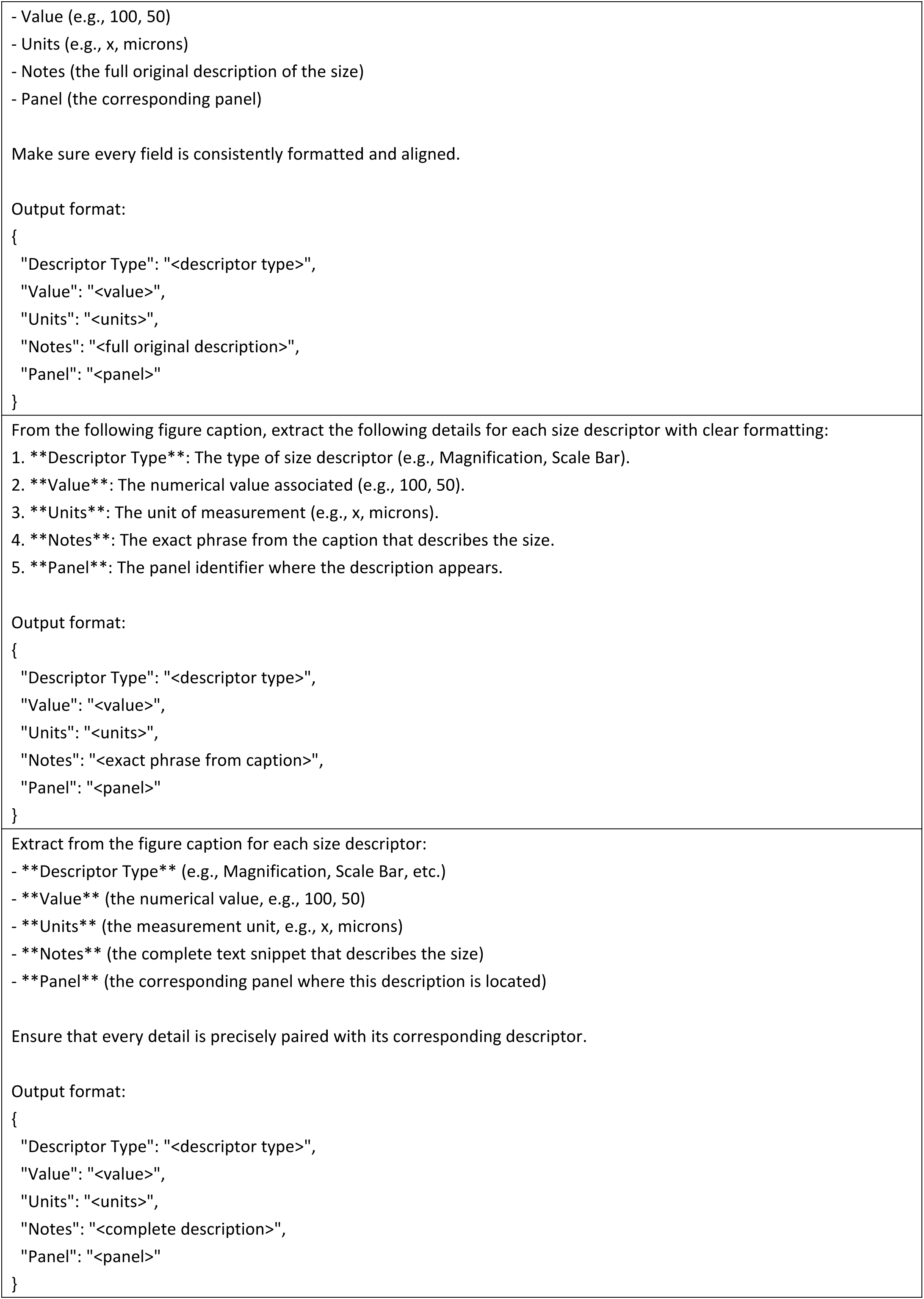
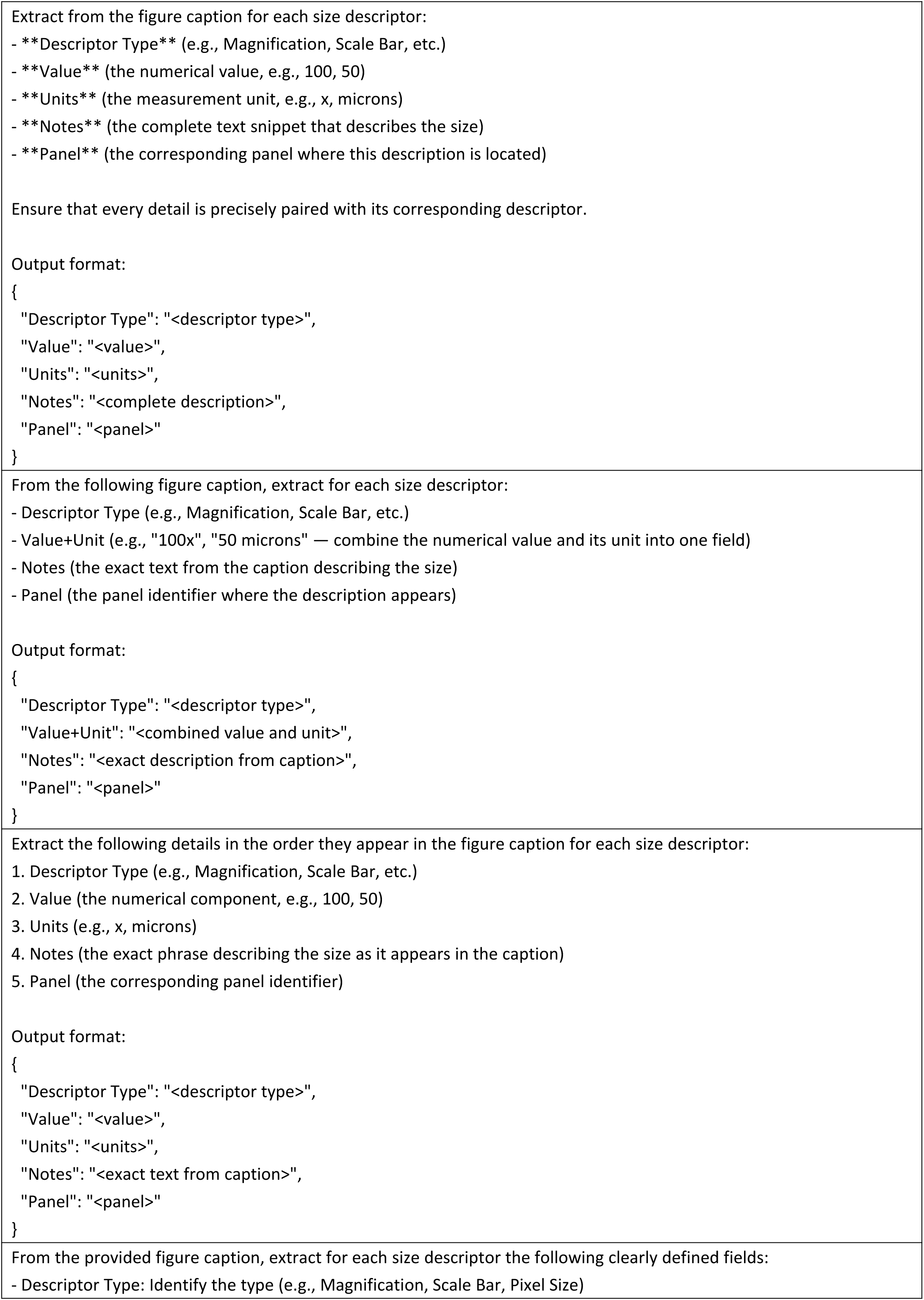
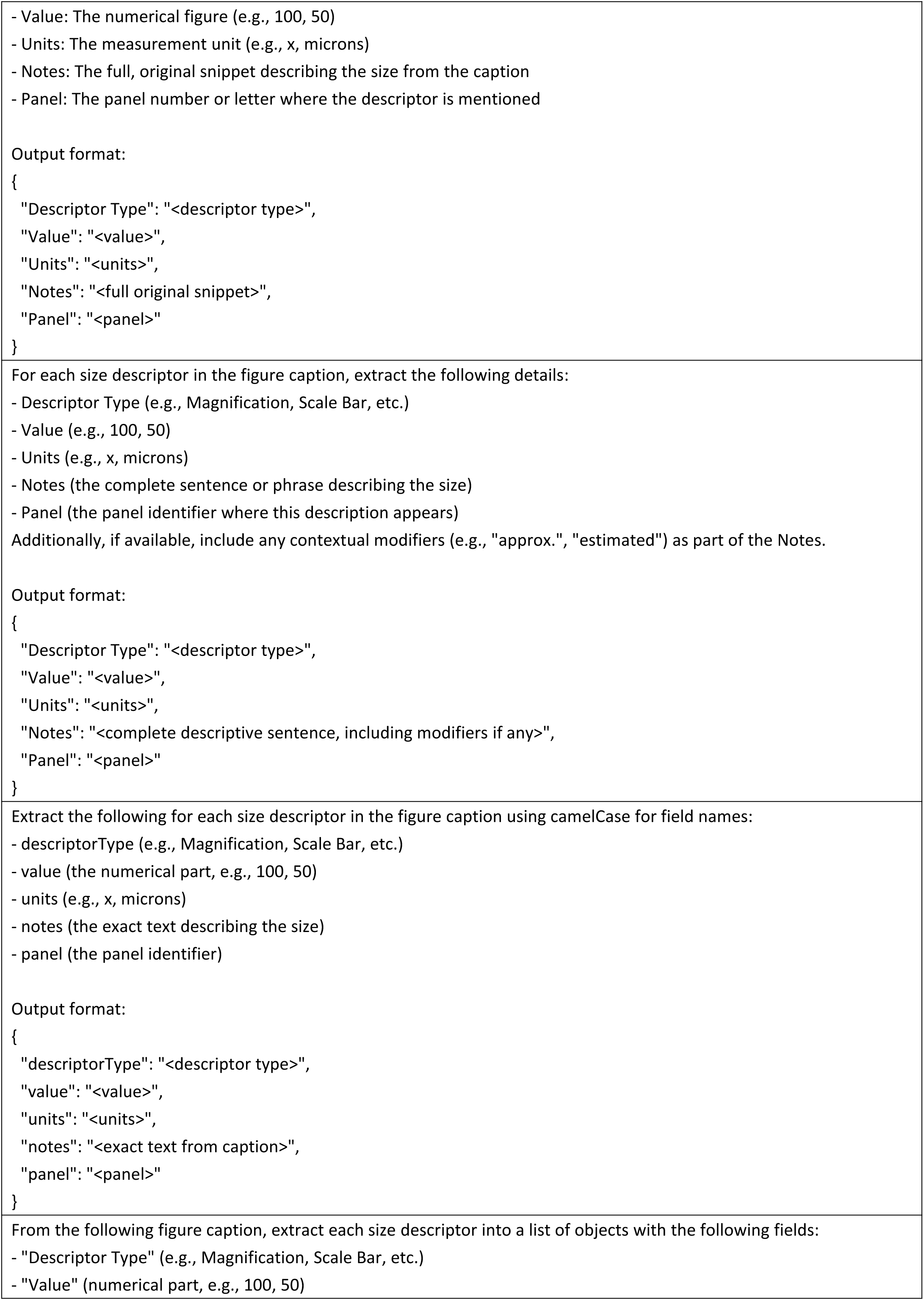
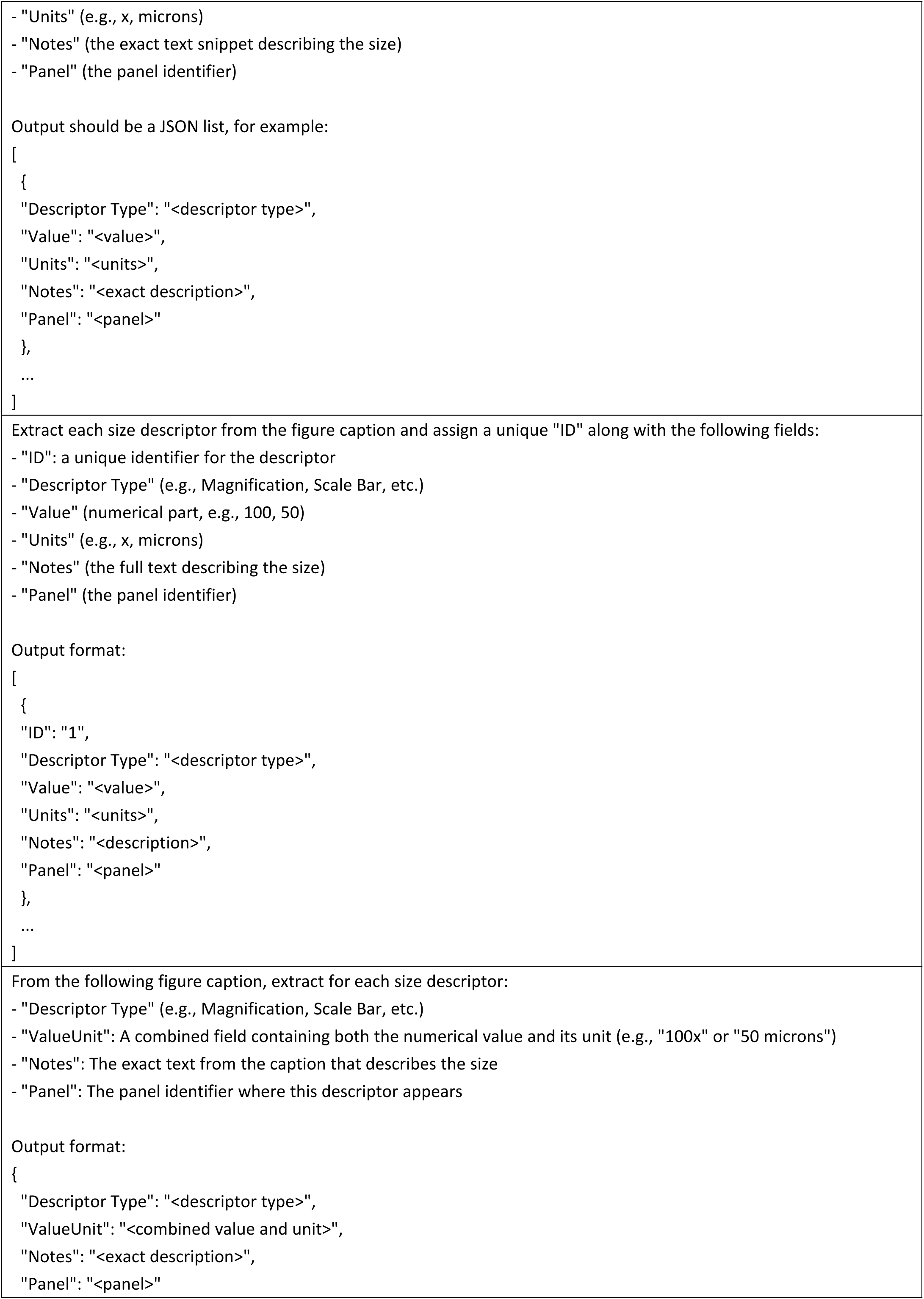
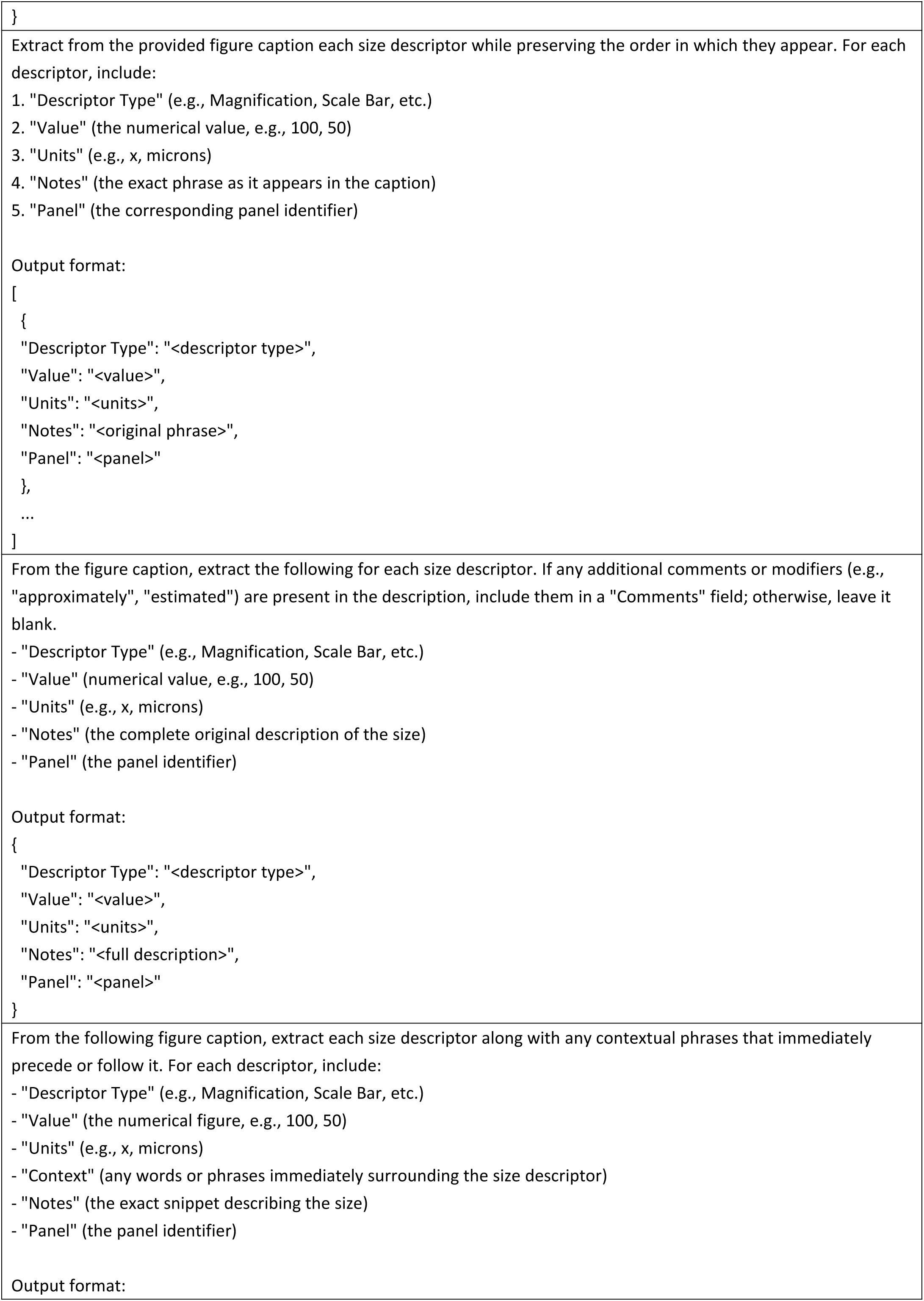
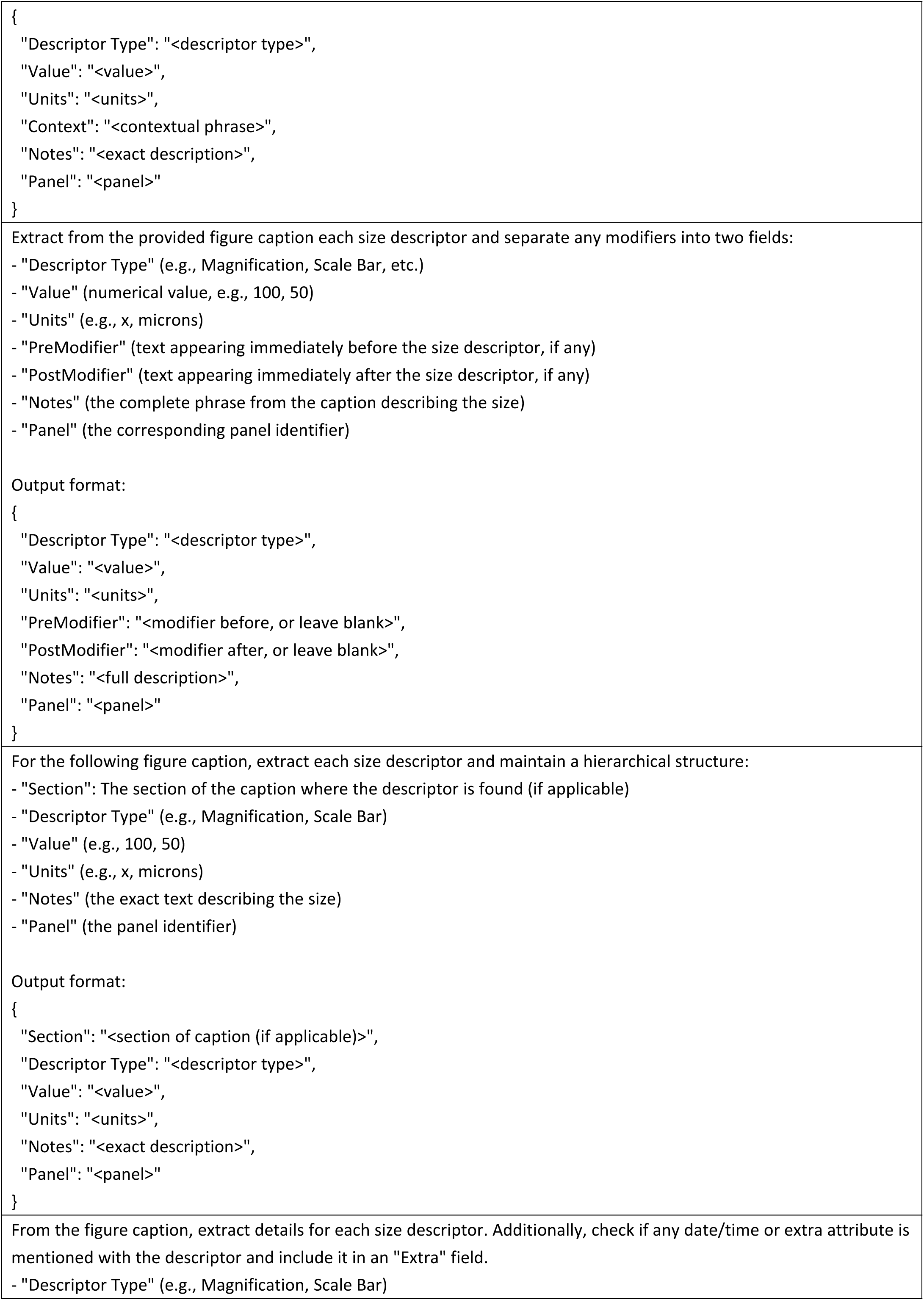
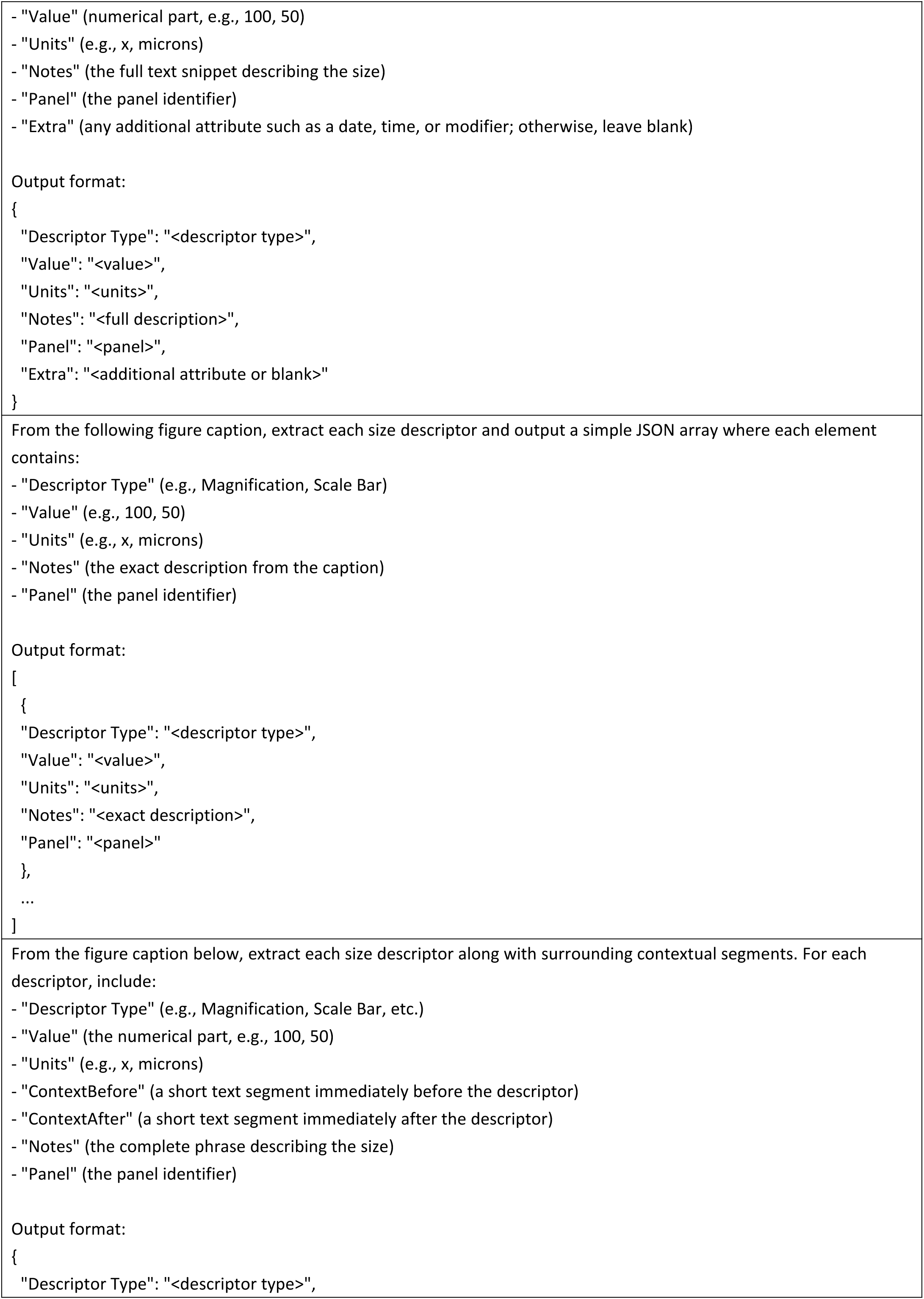
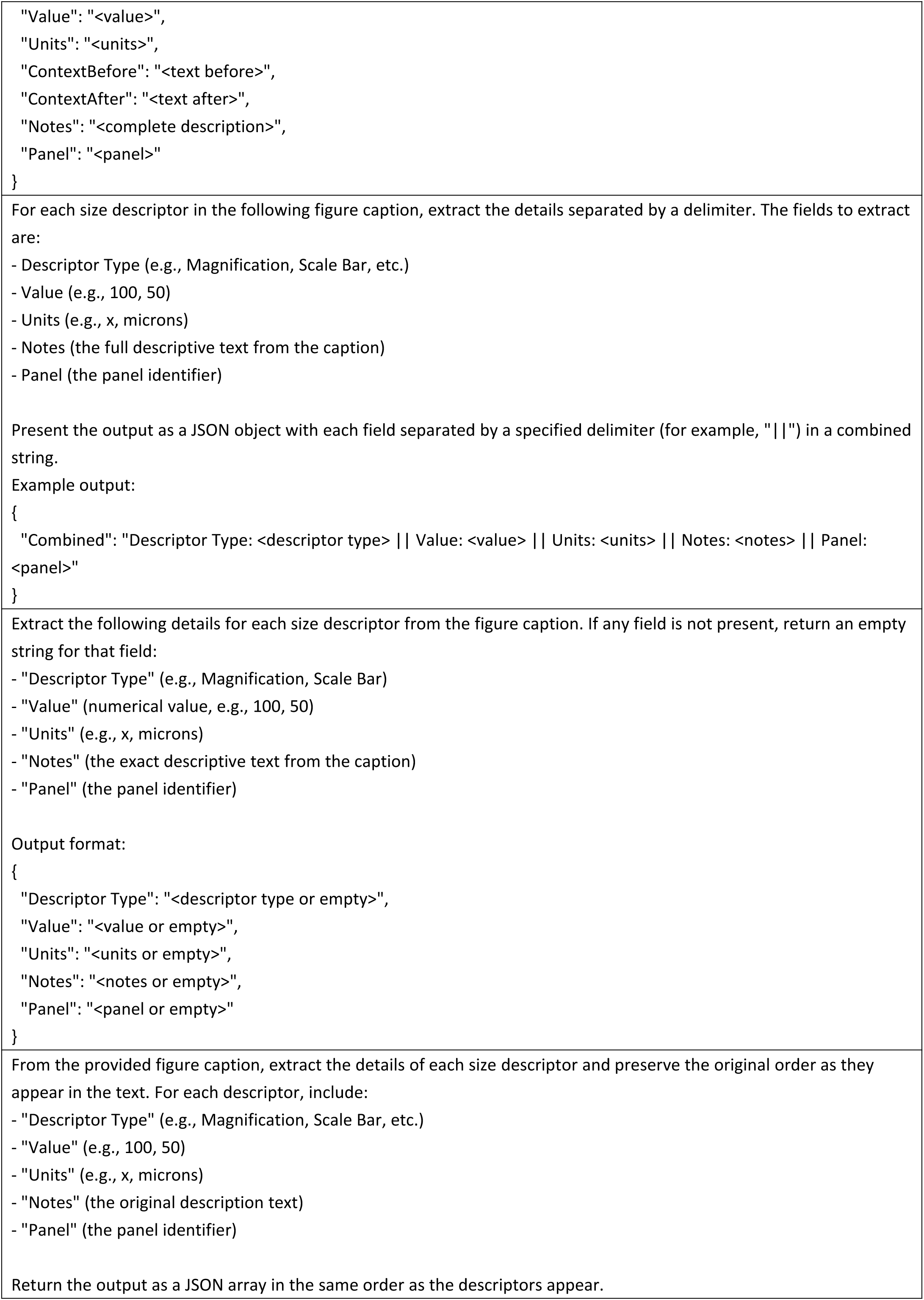
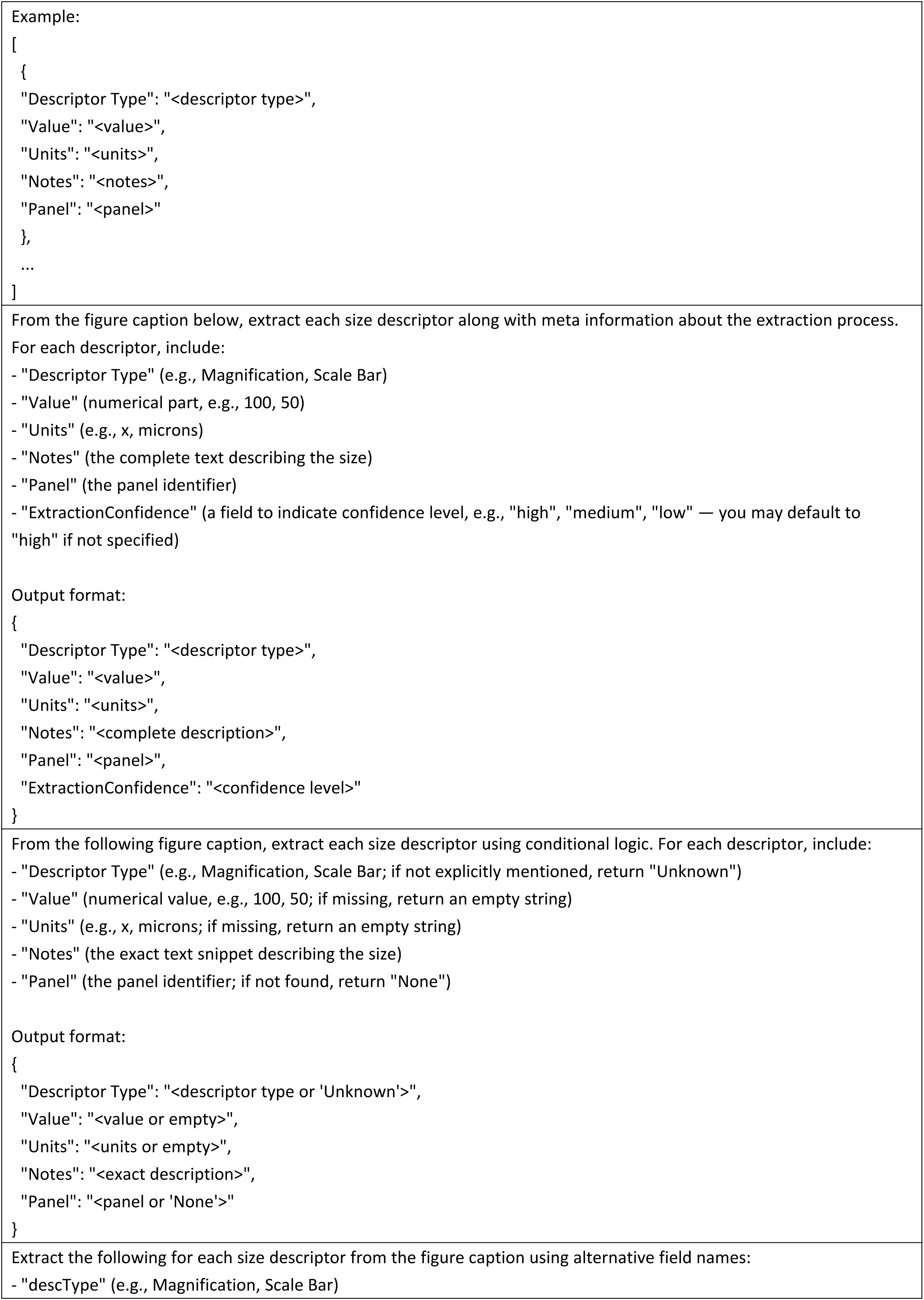
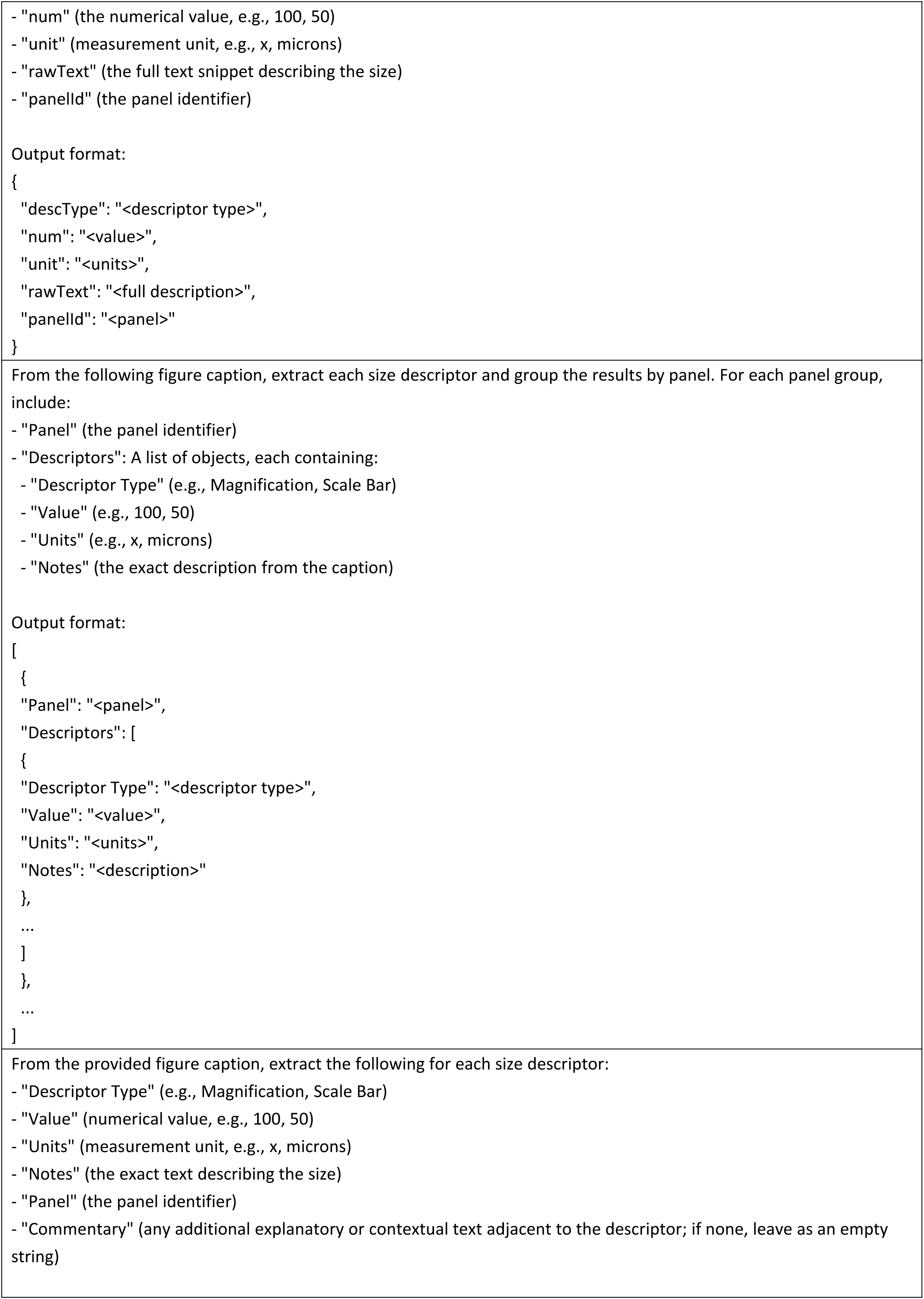
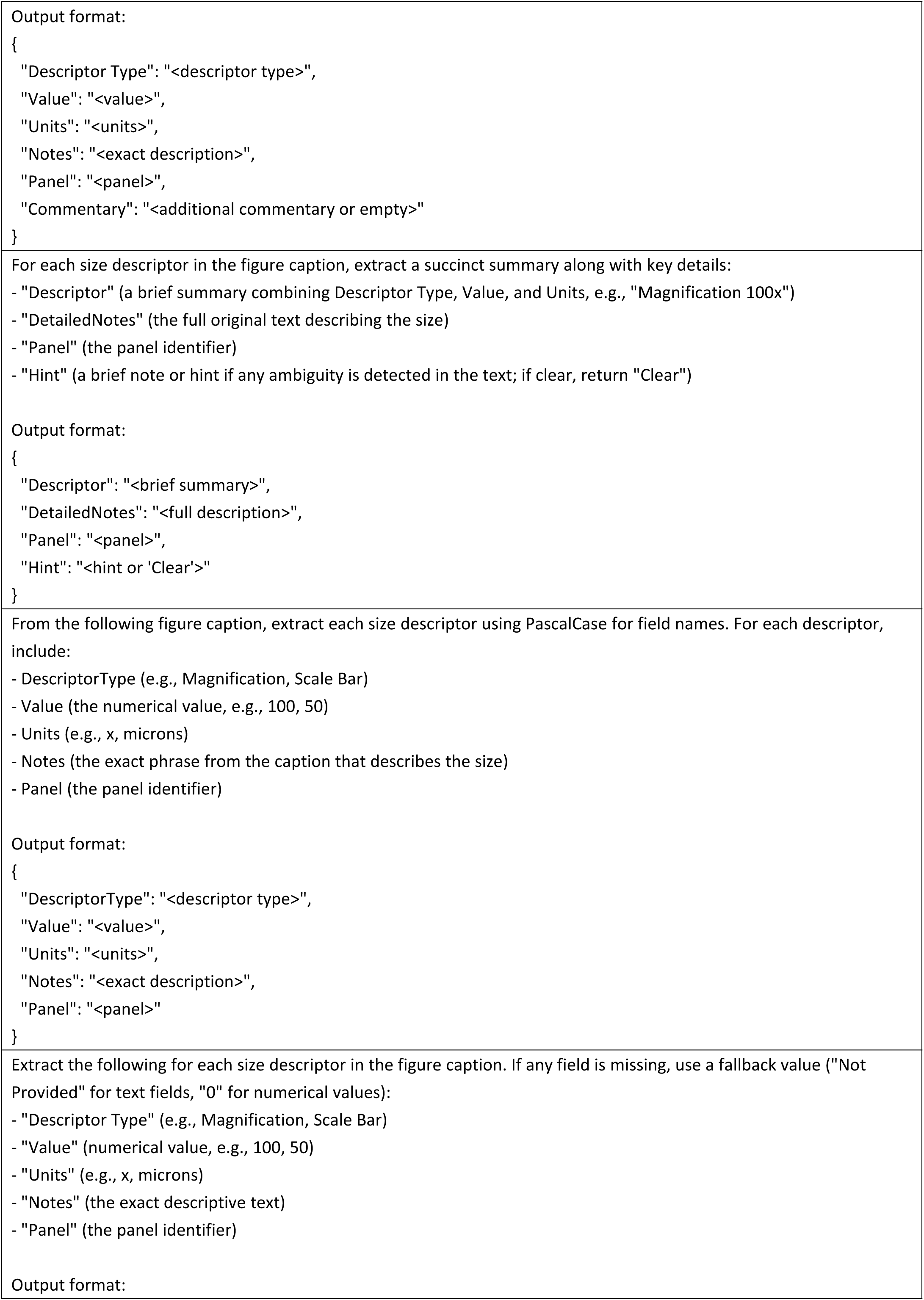
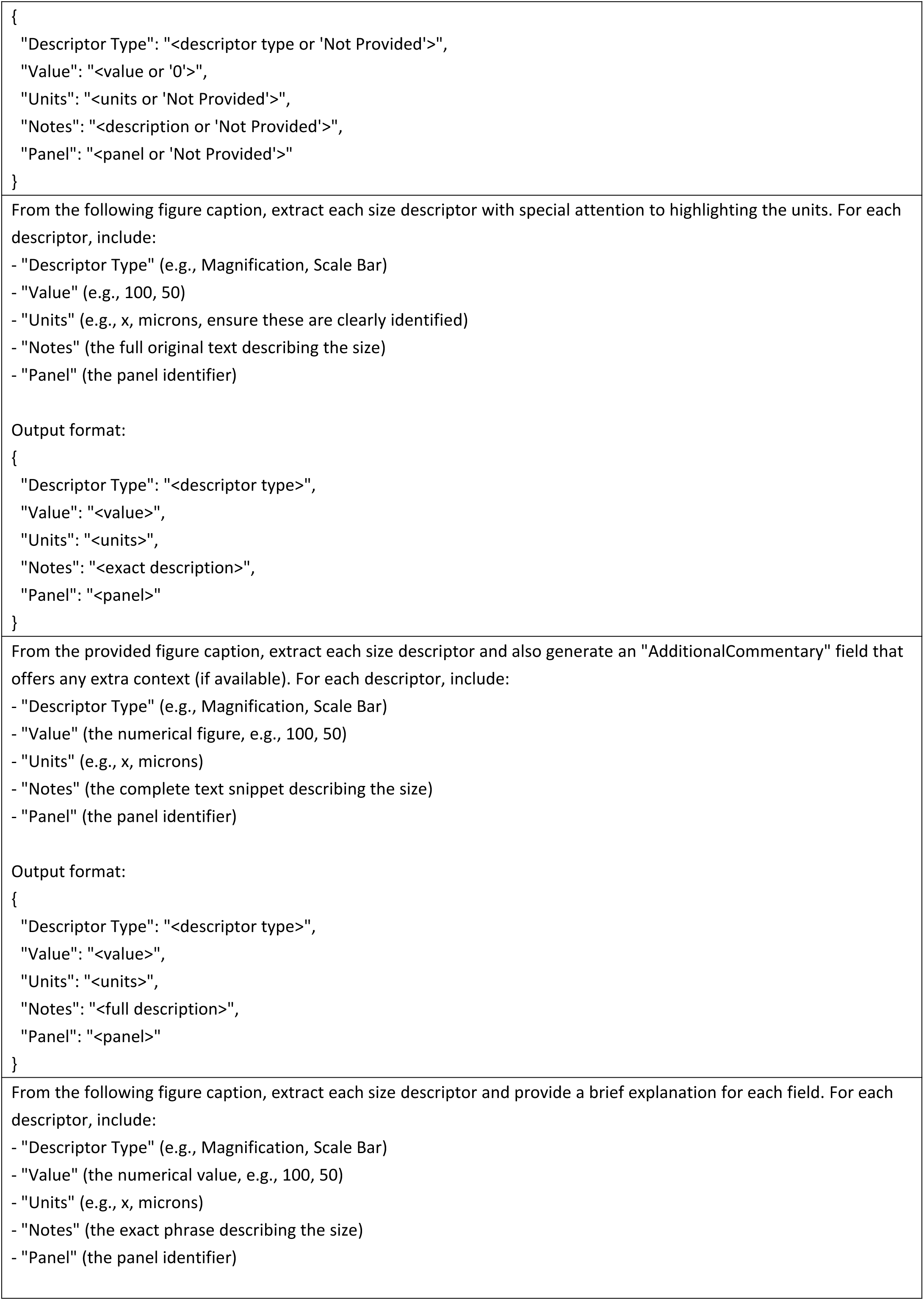
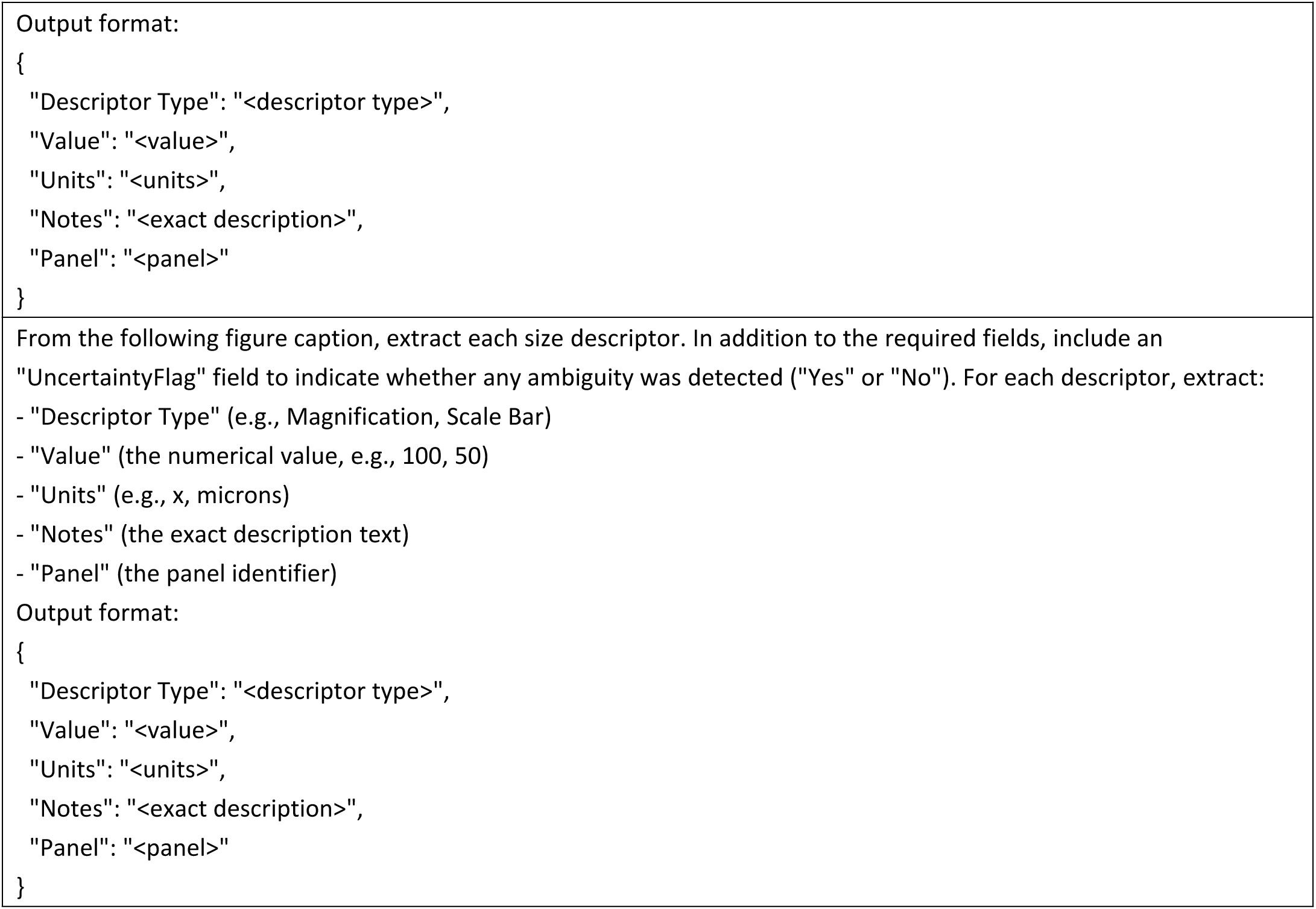
Prompts for scale-bar extraction tests.

**Supplementary Table 20.**
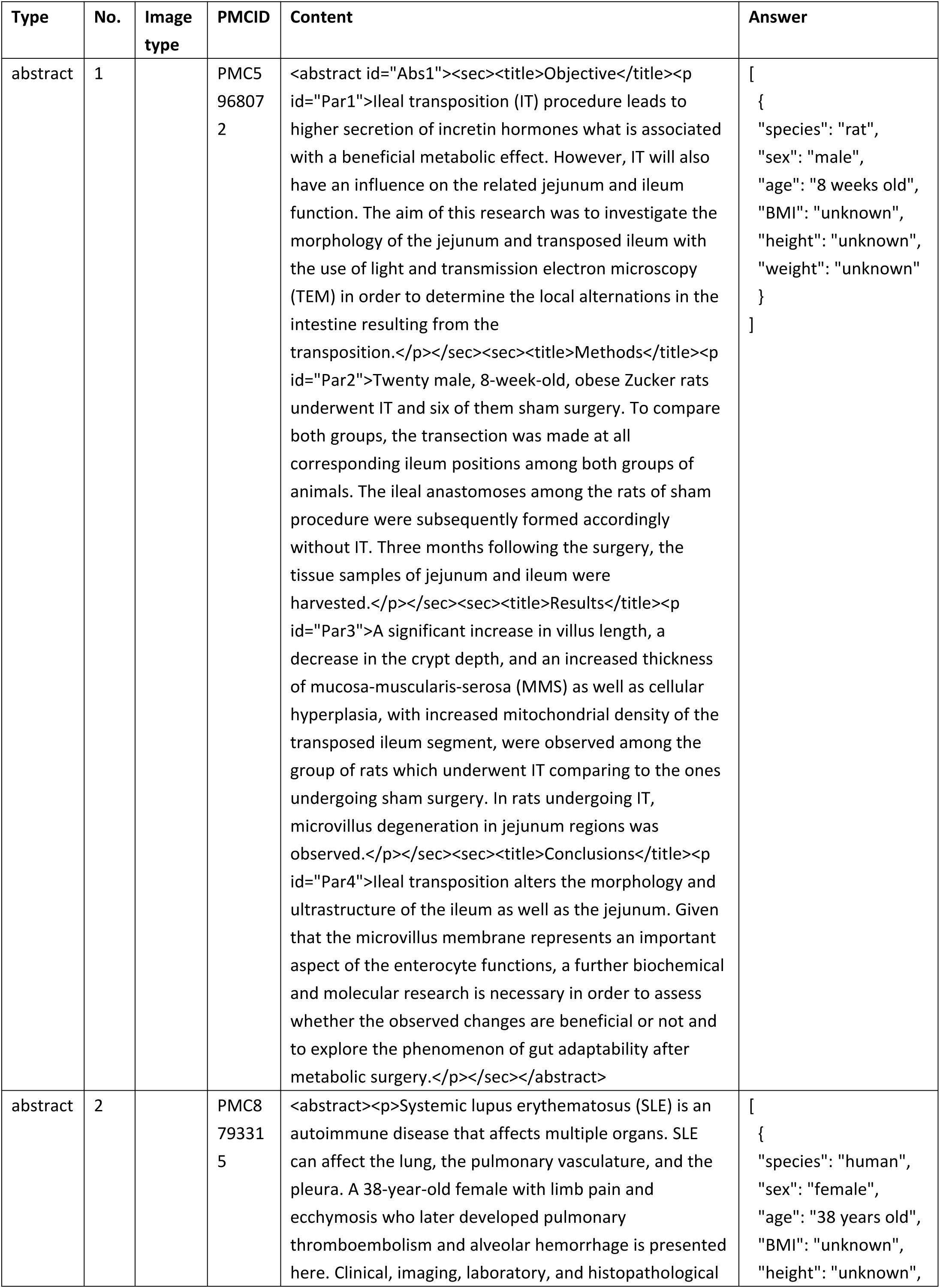
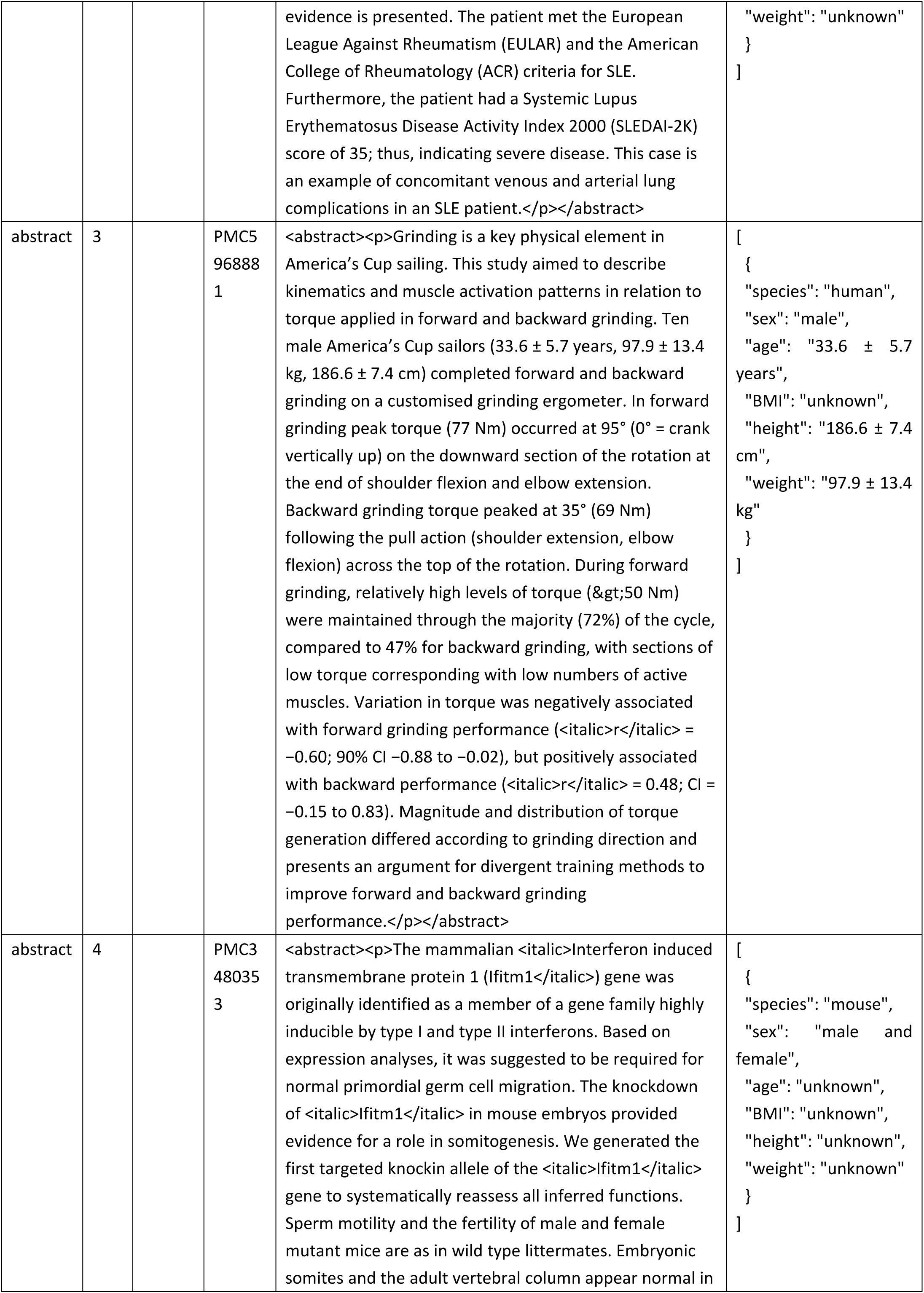
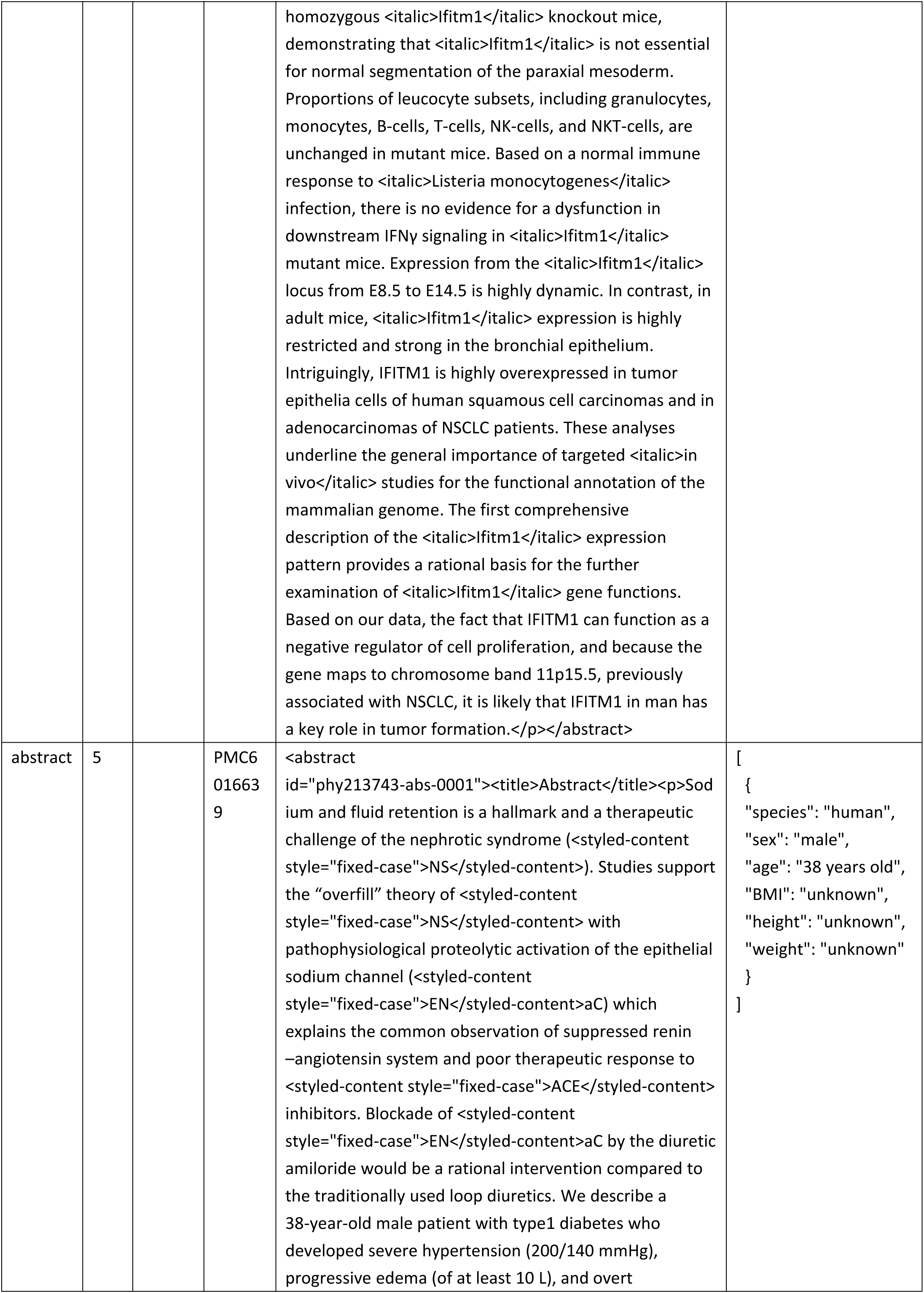
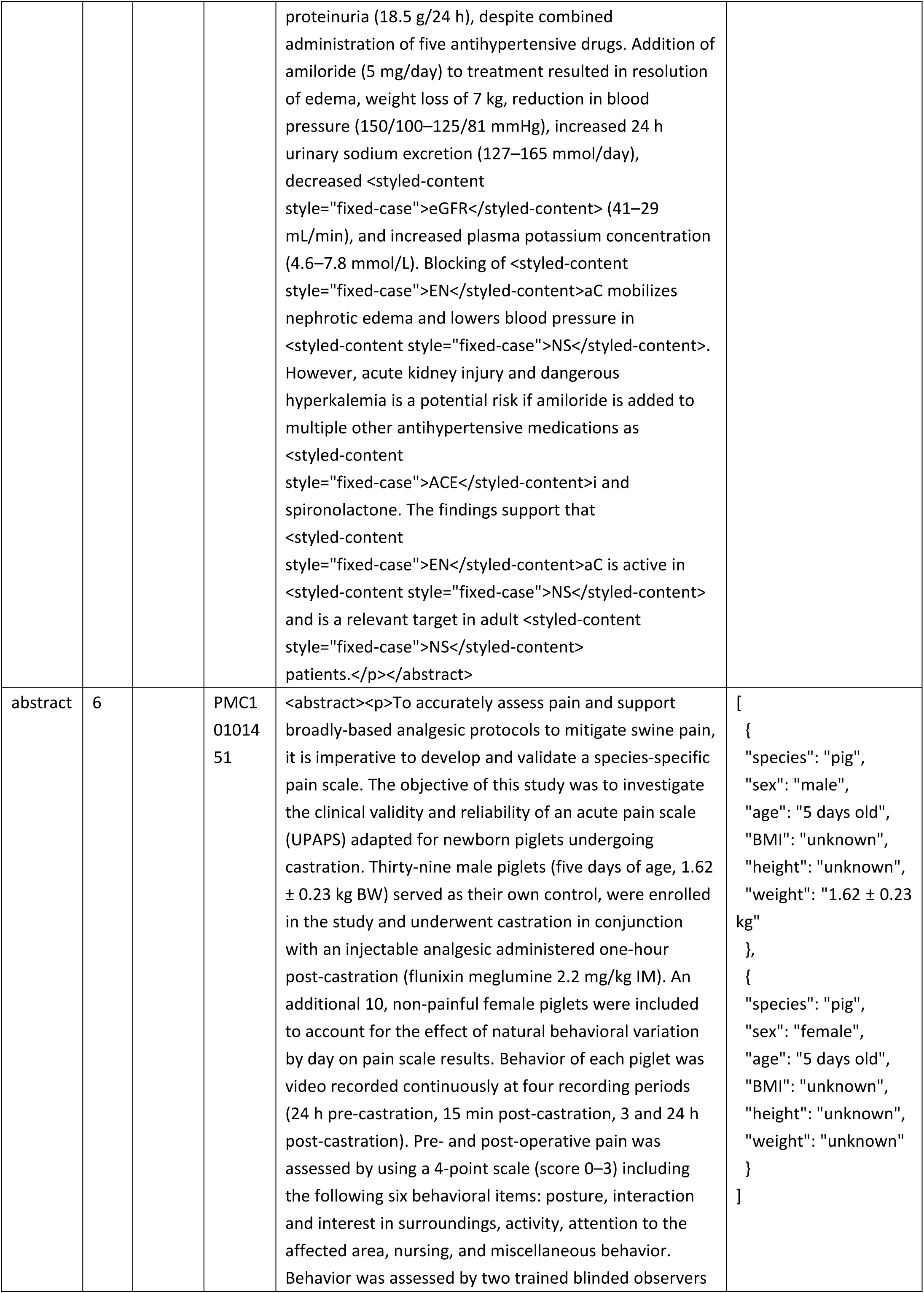
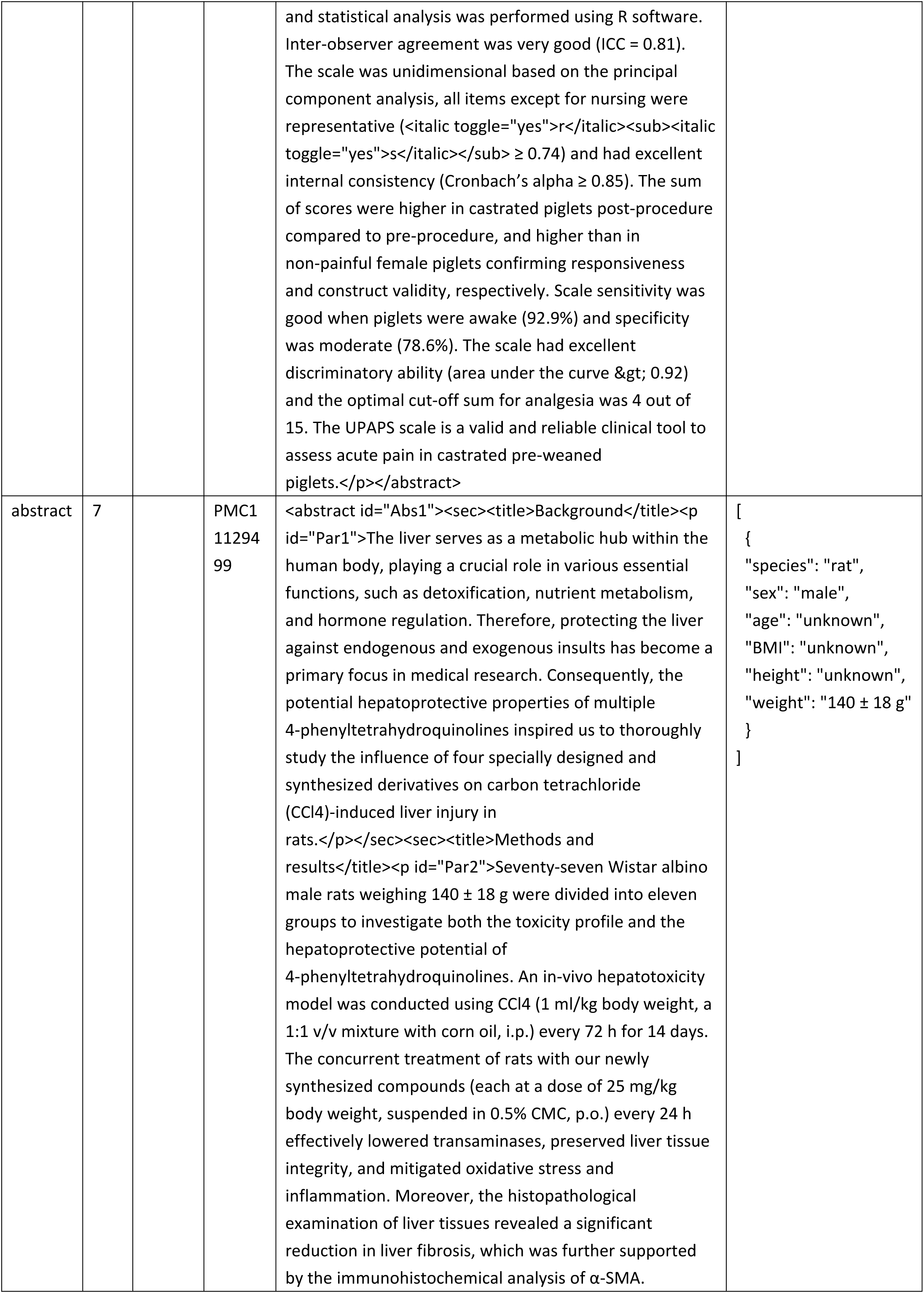
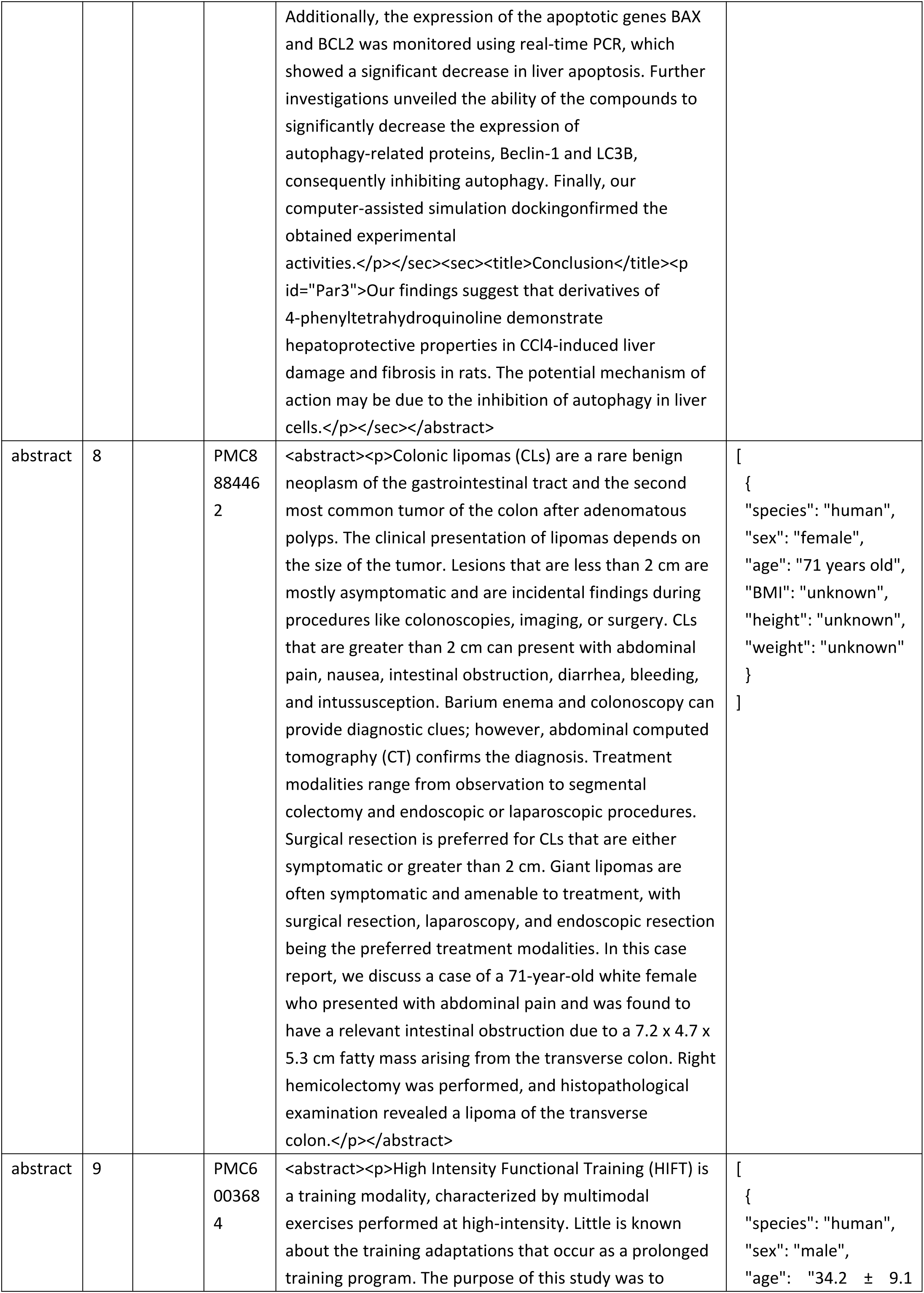
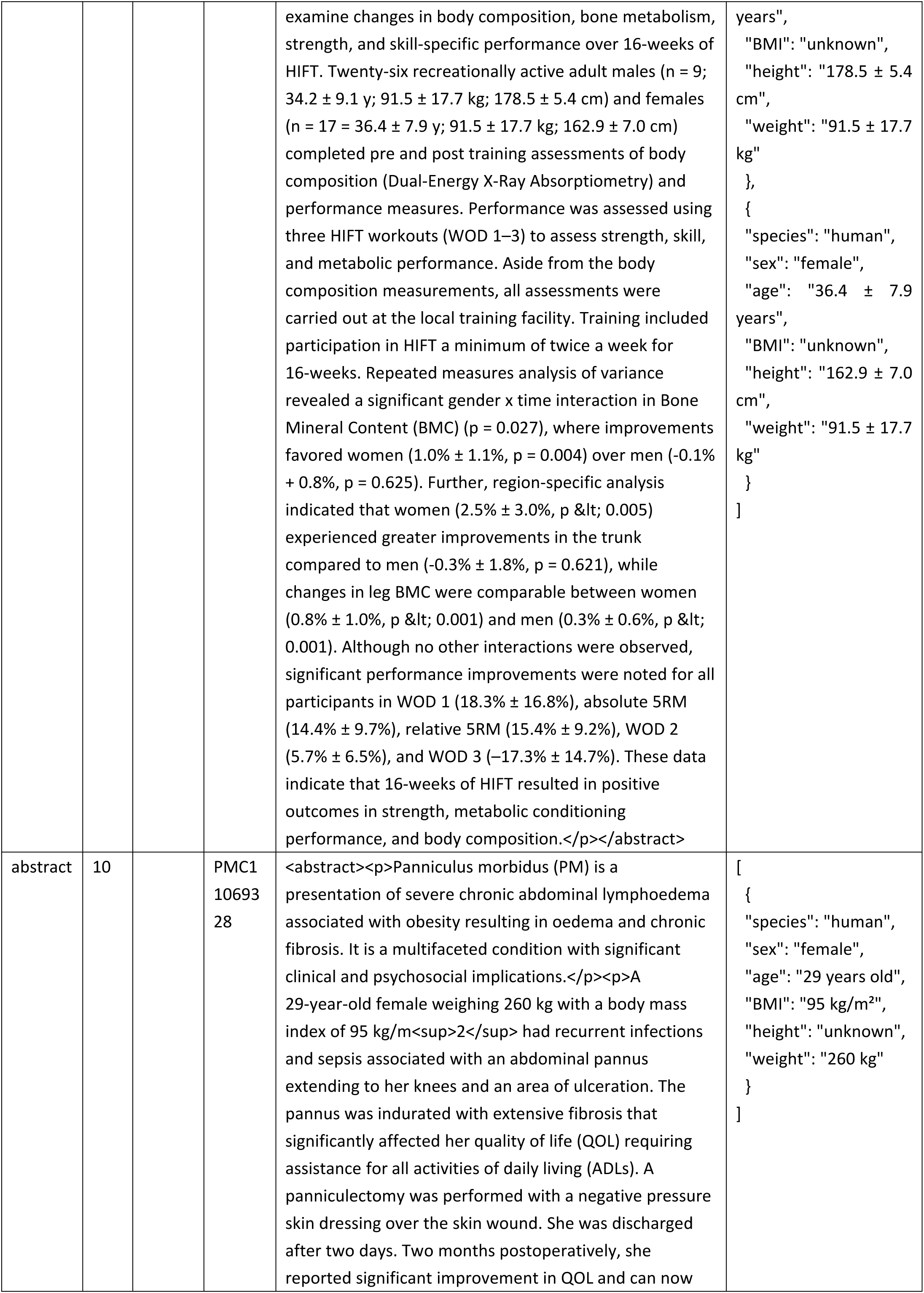
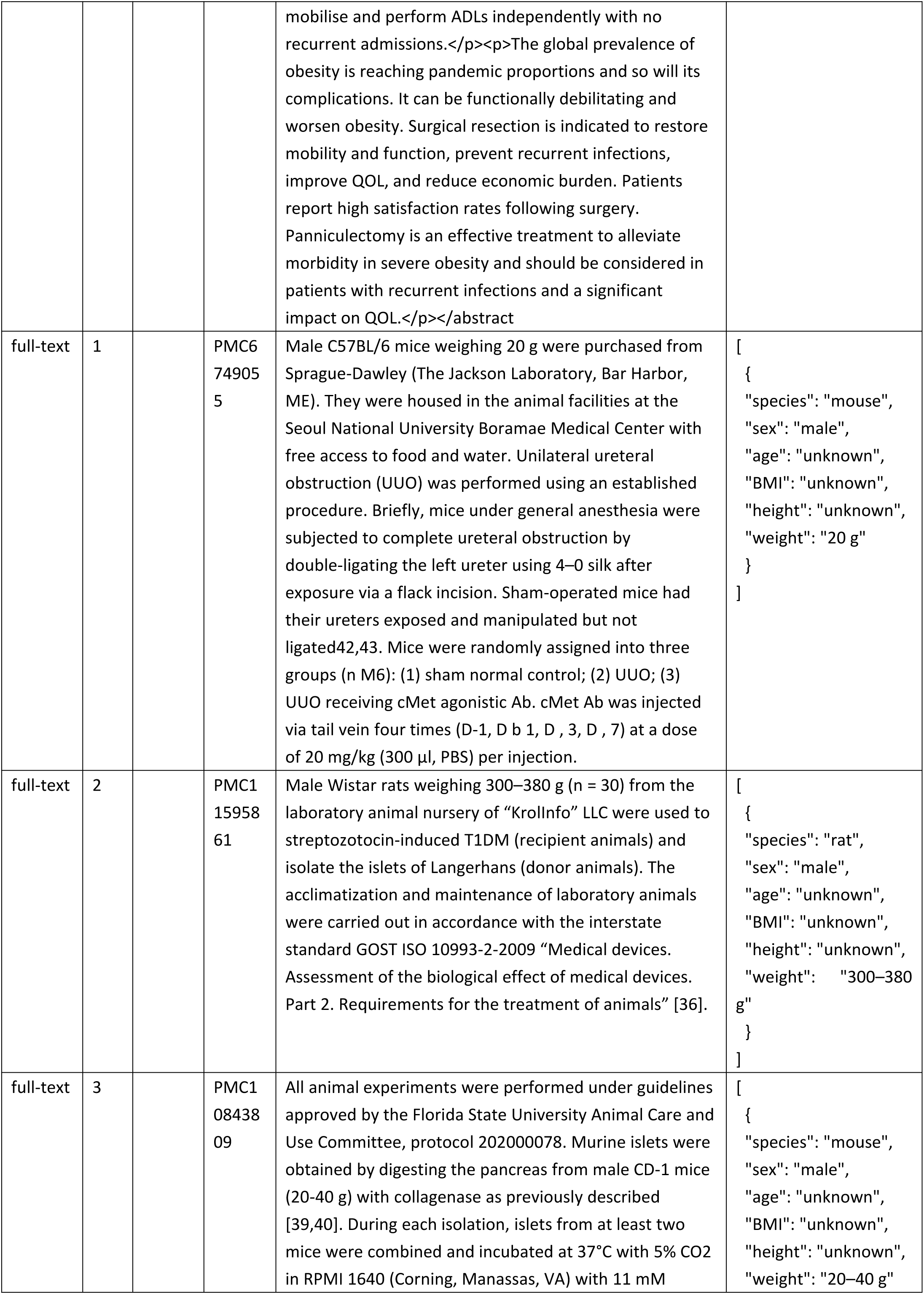
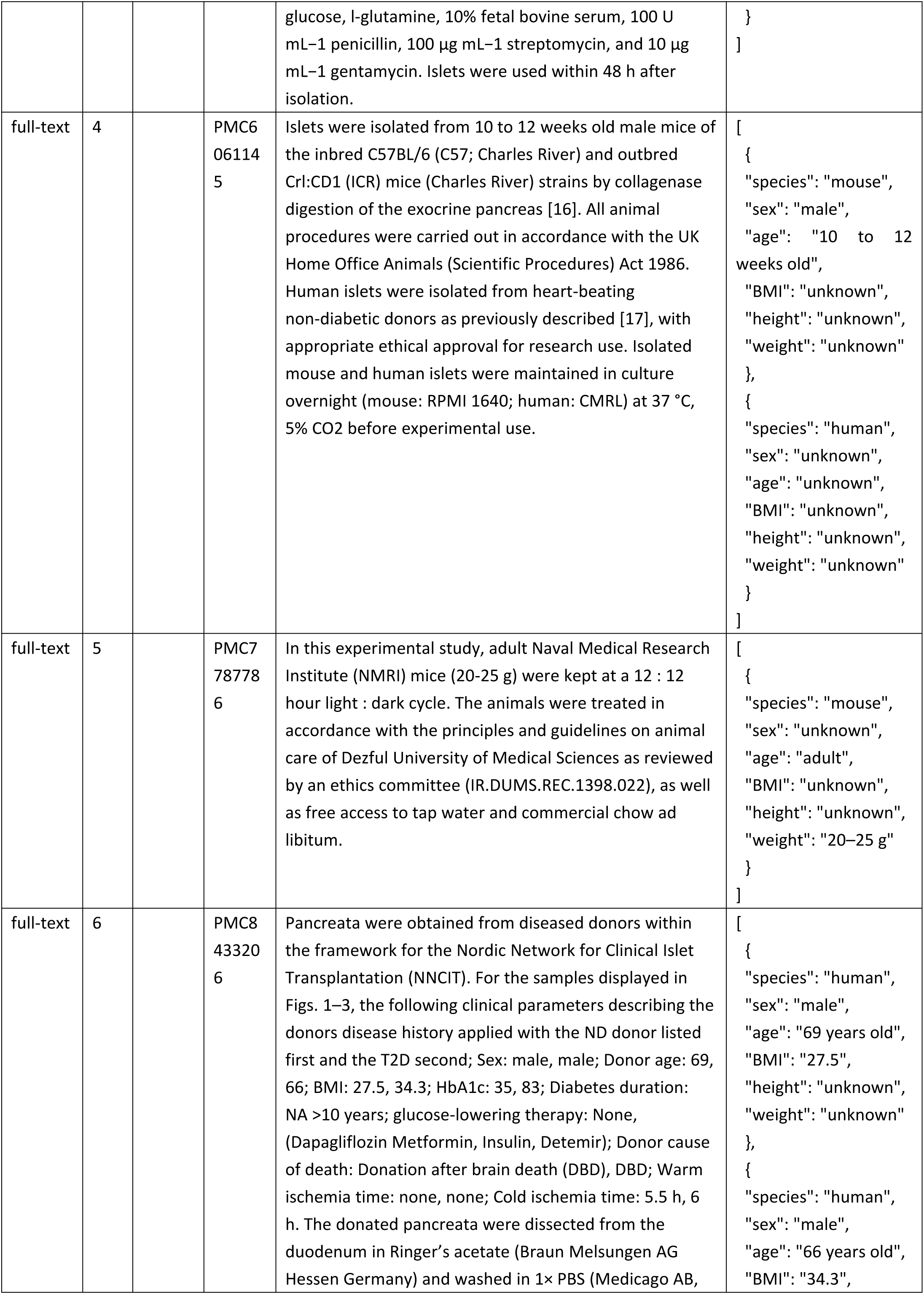
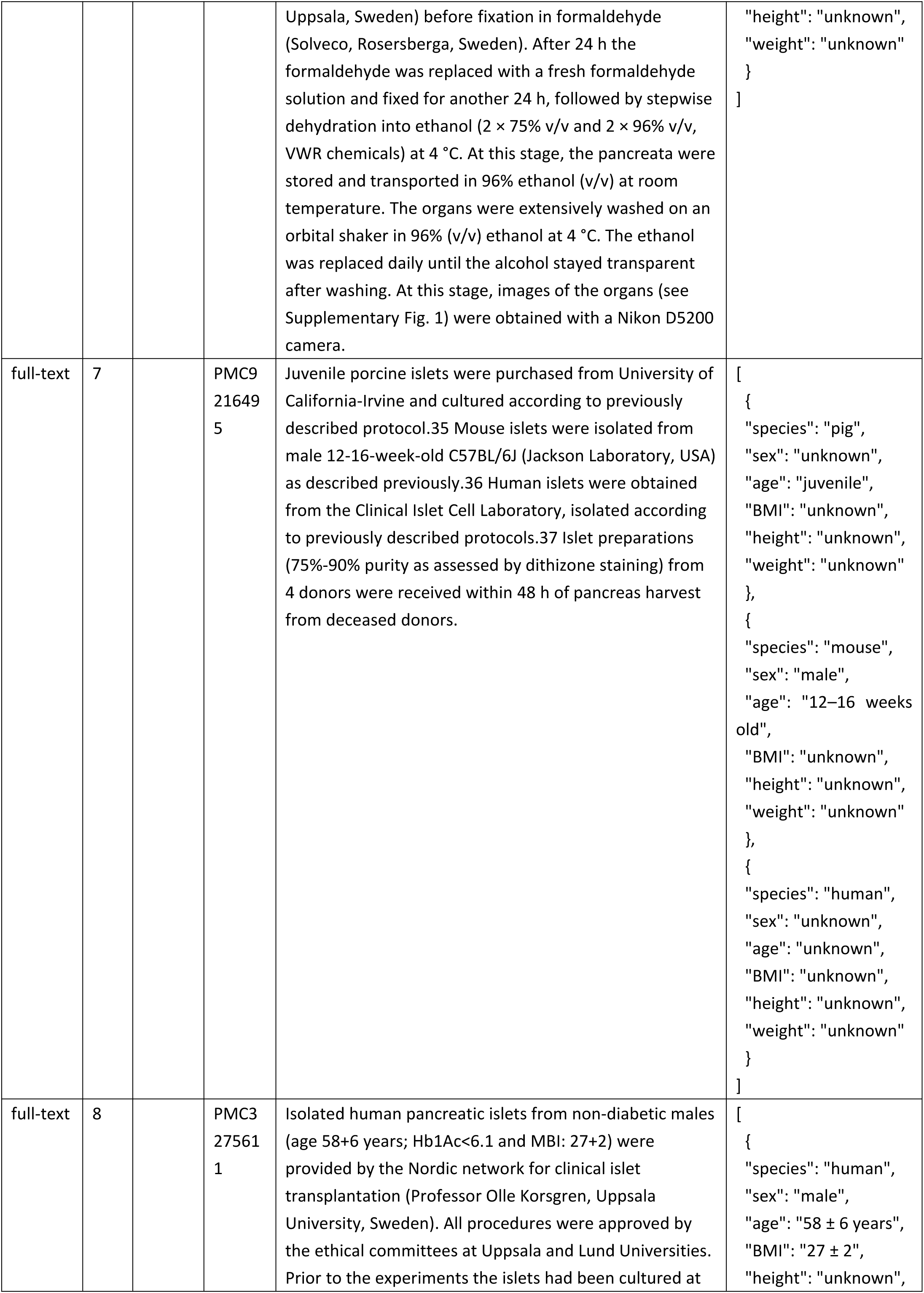
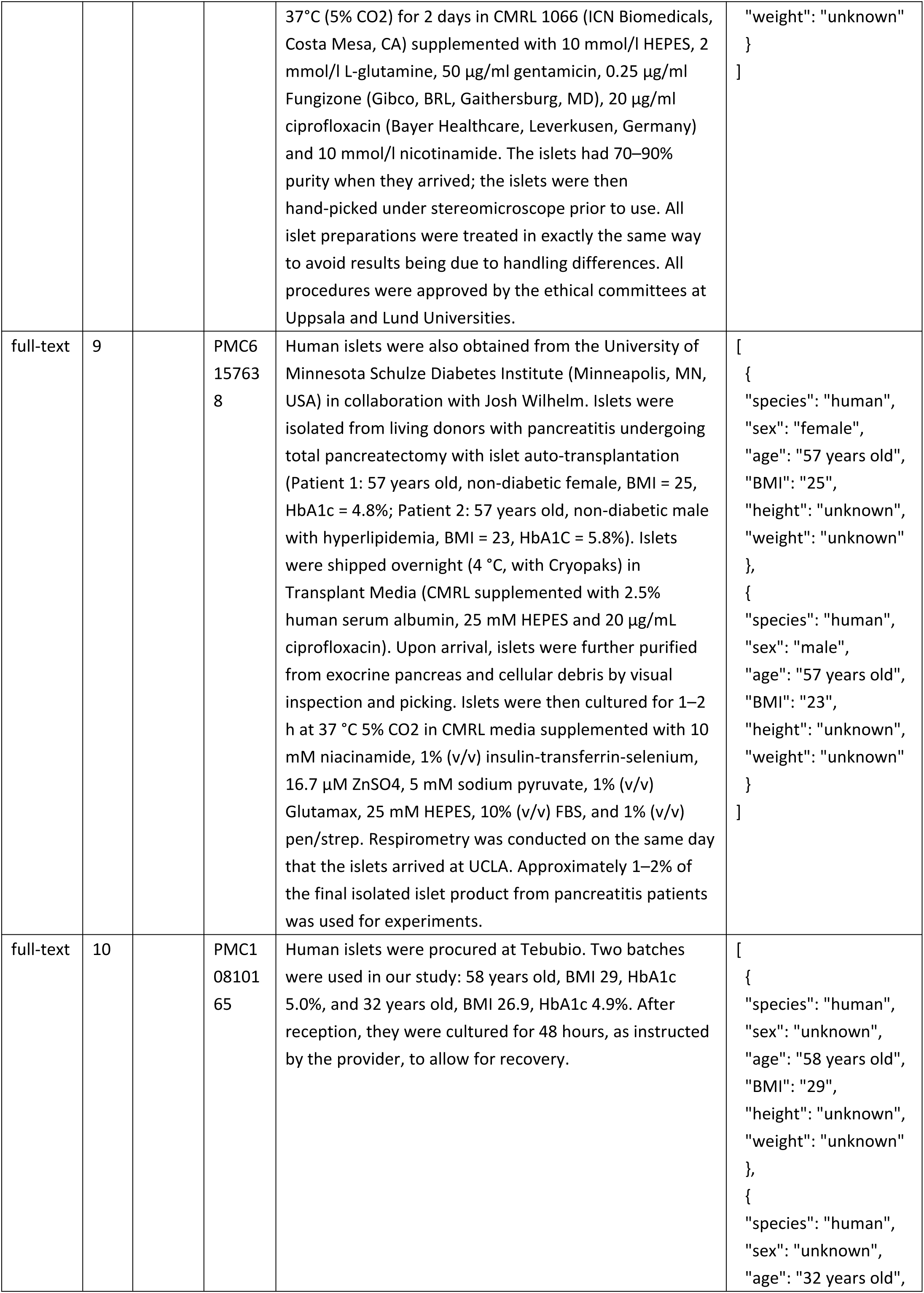
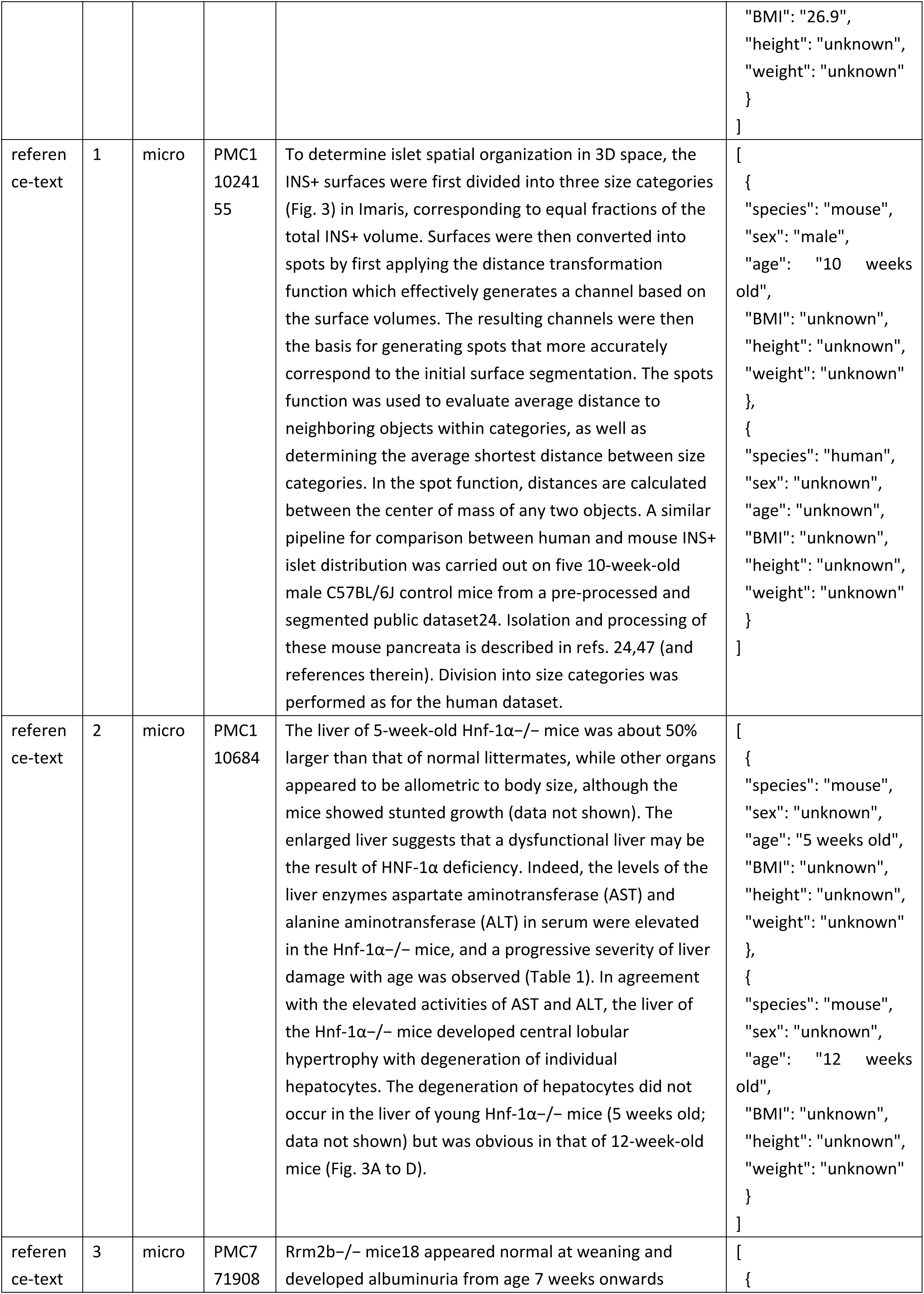
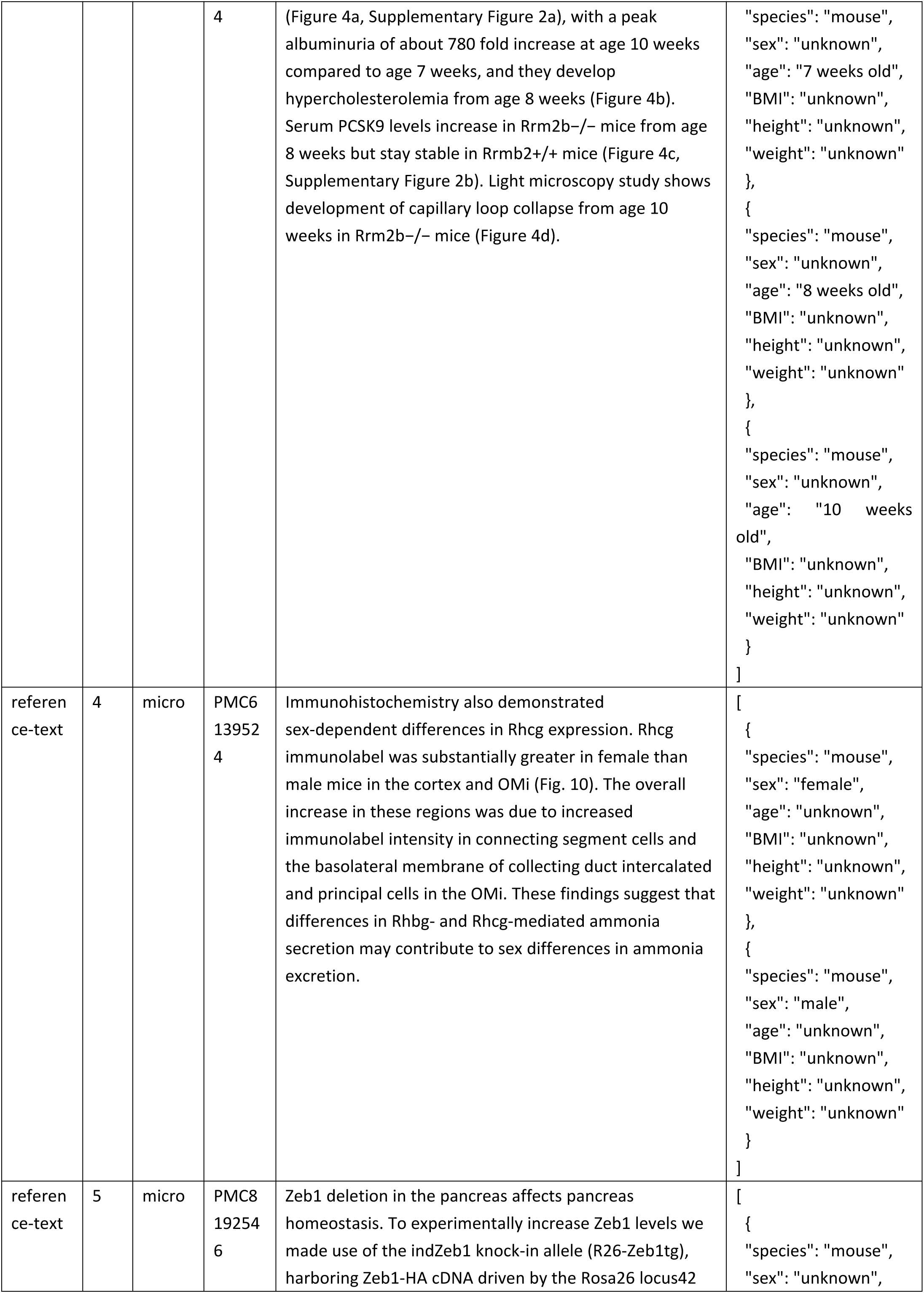
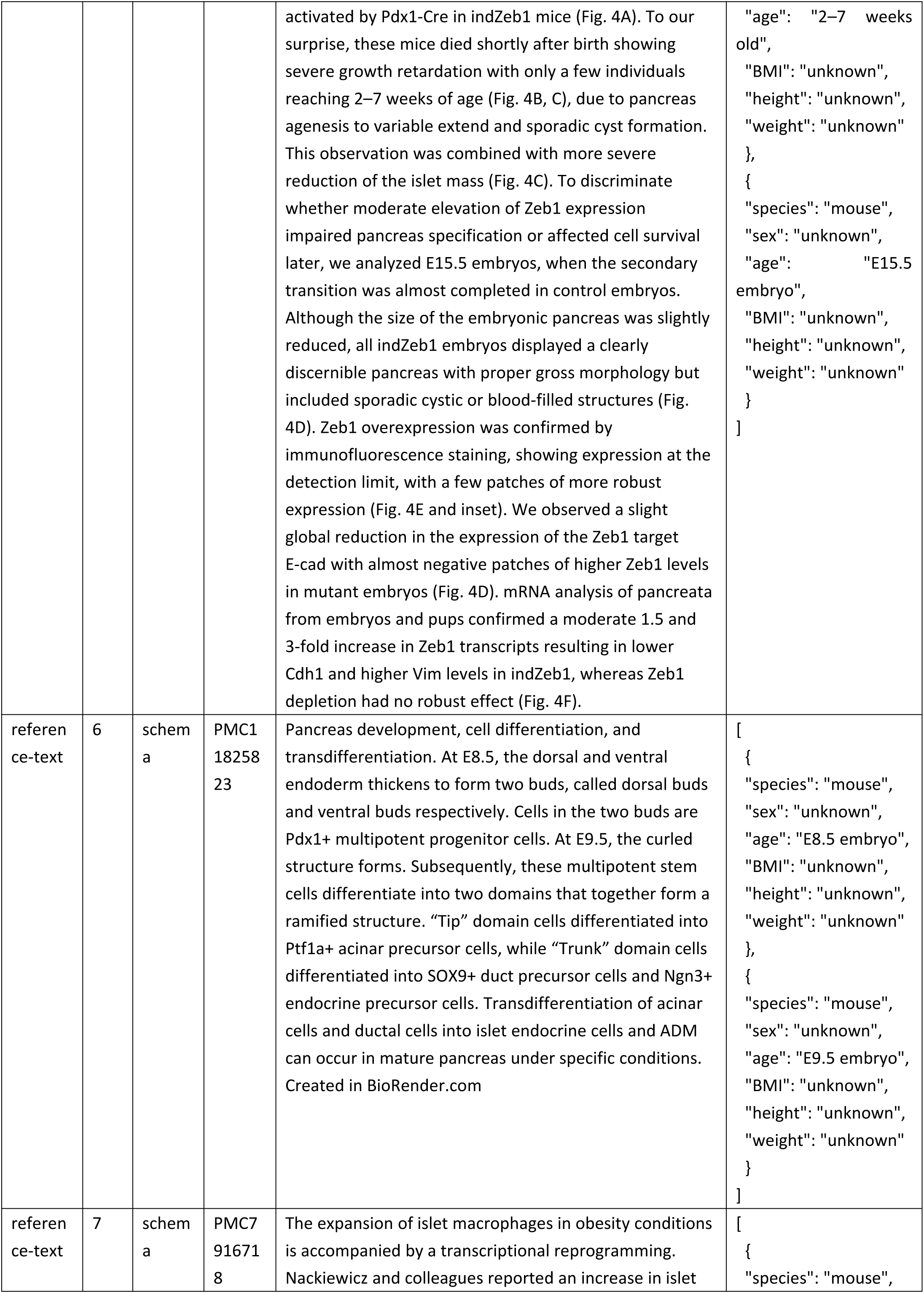
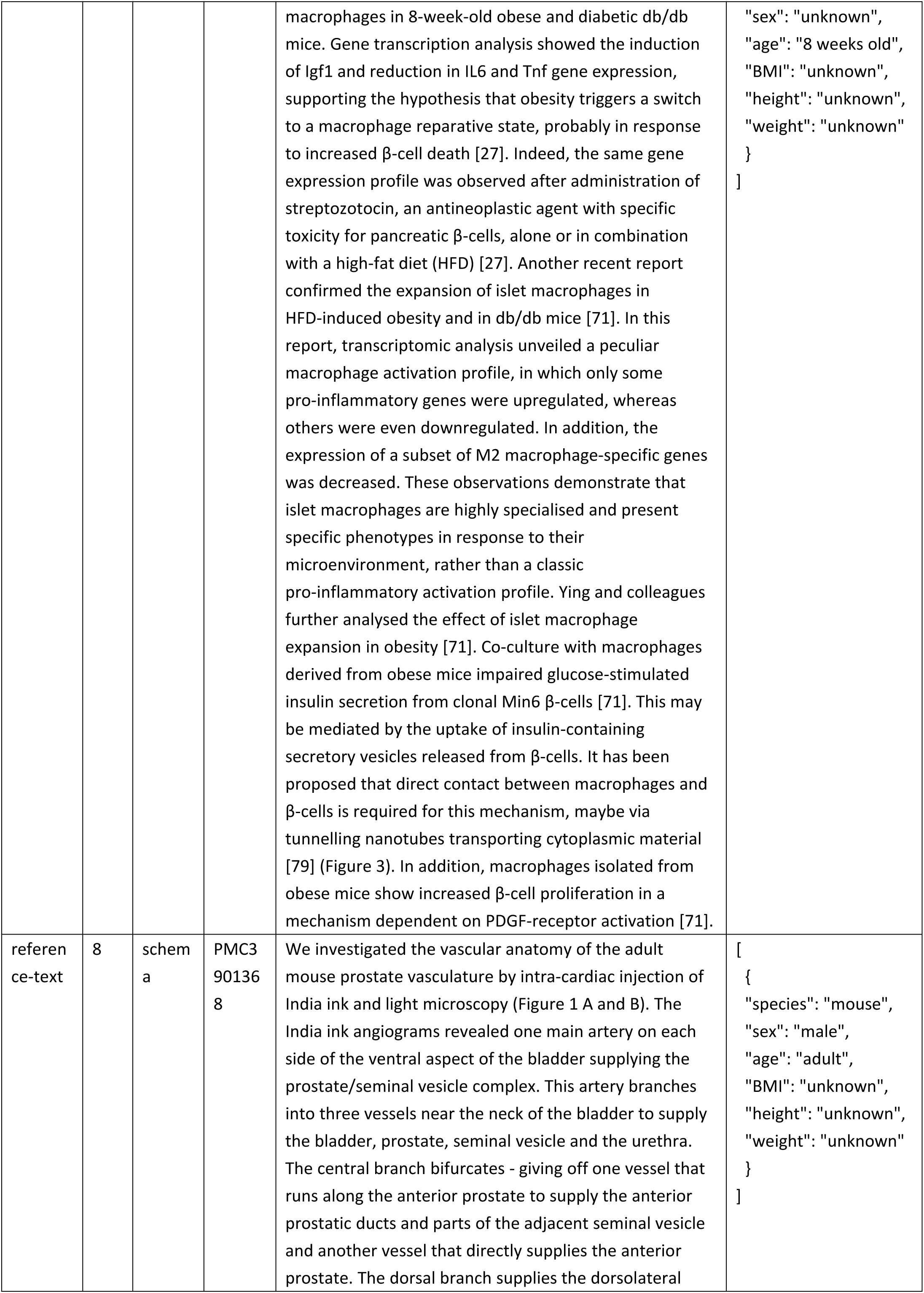
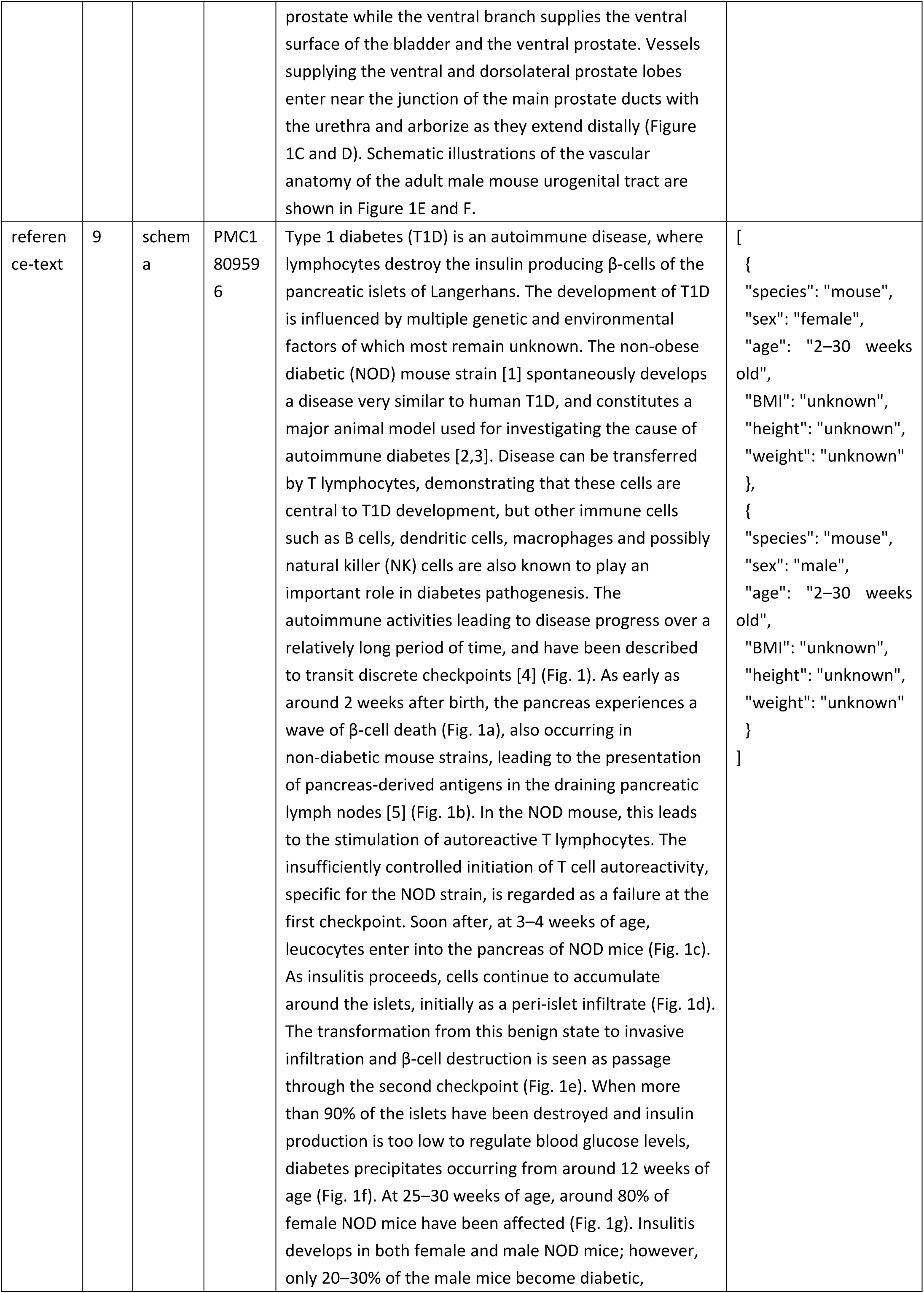
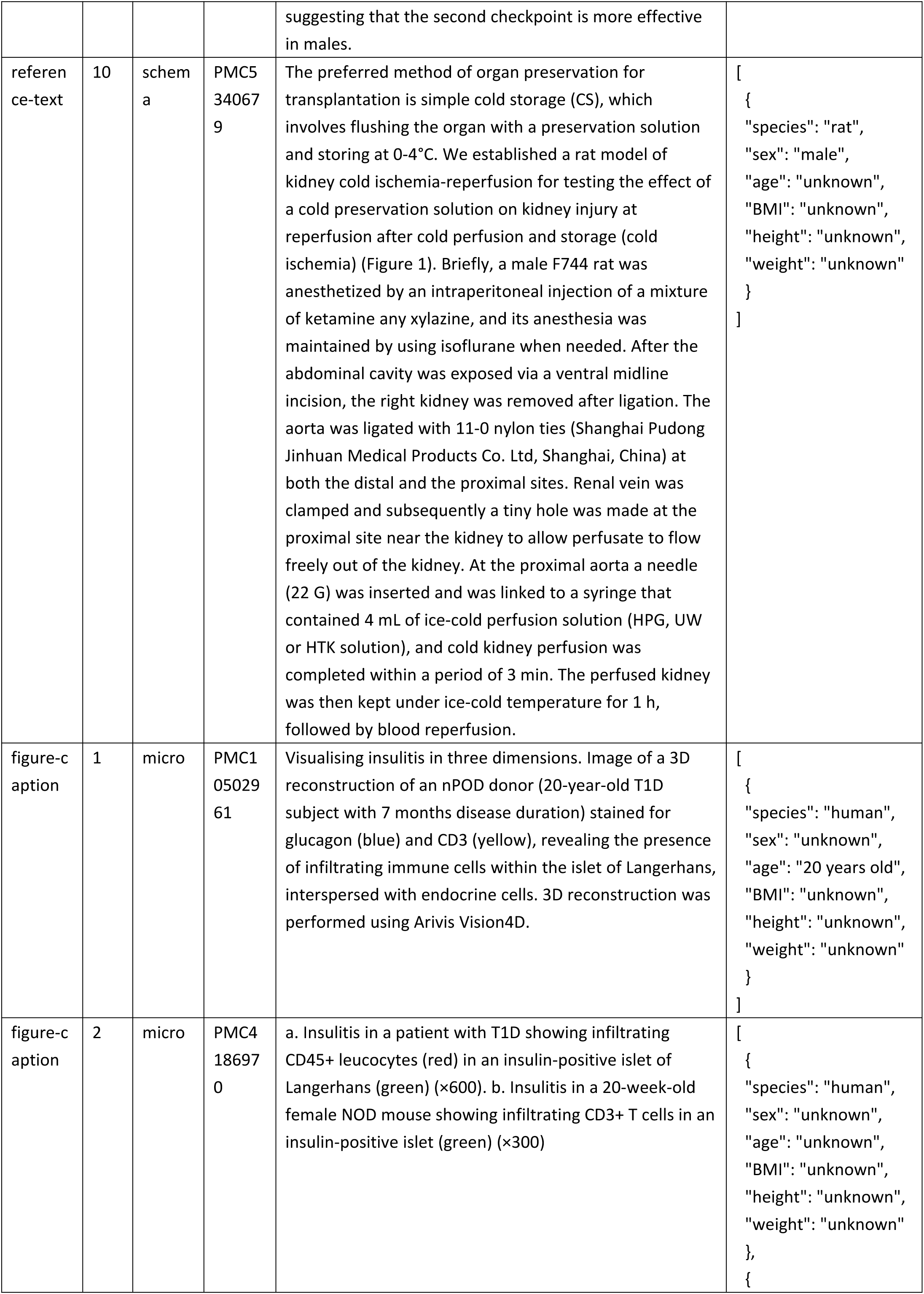
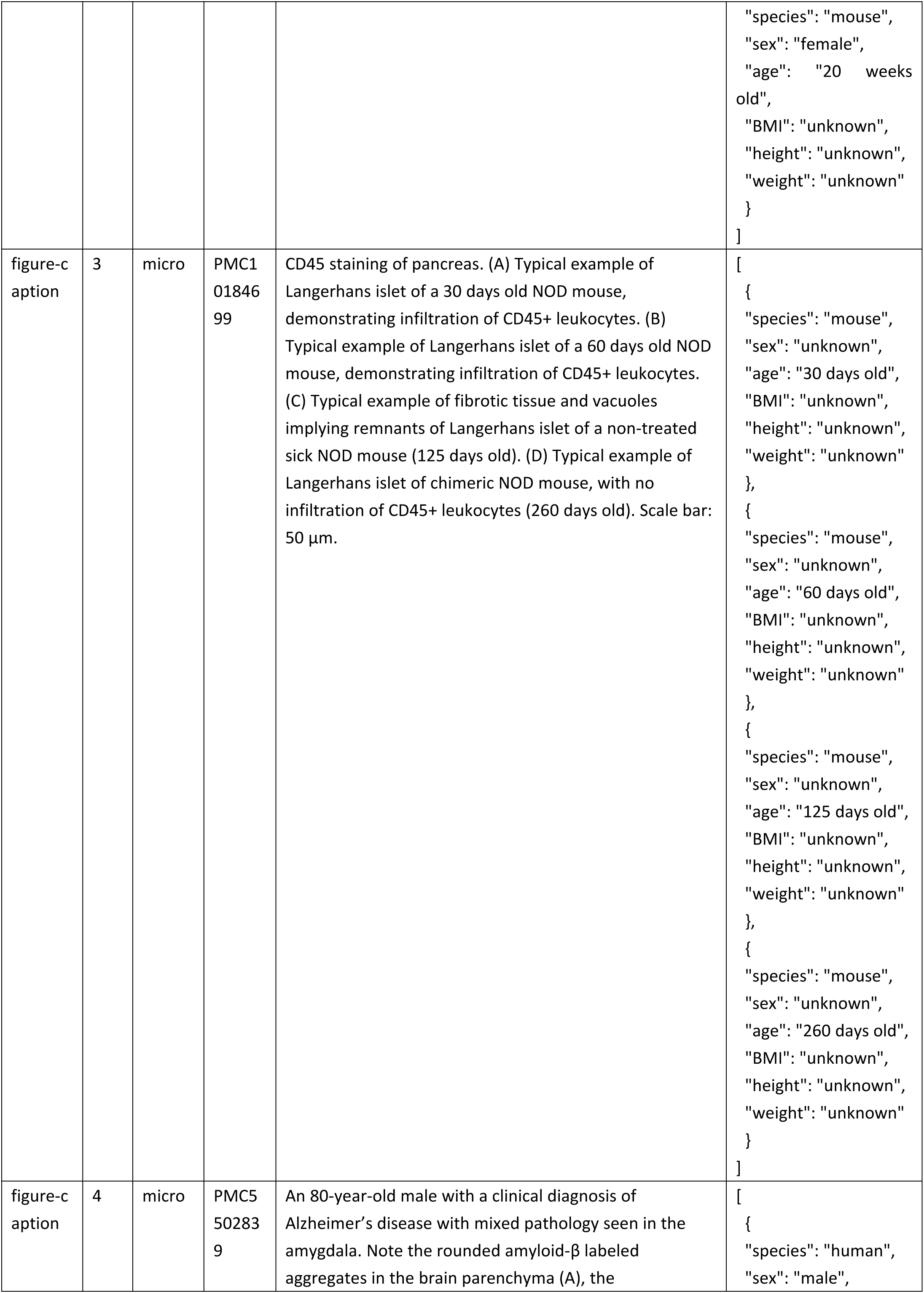
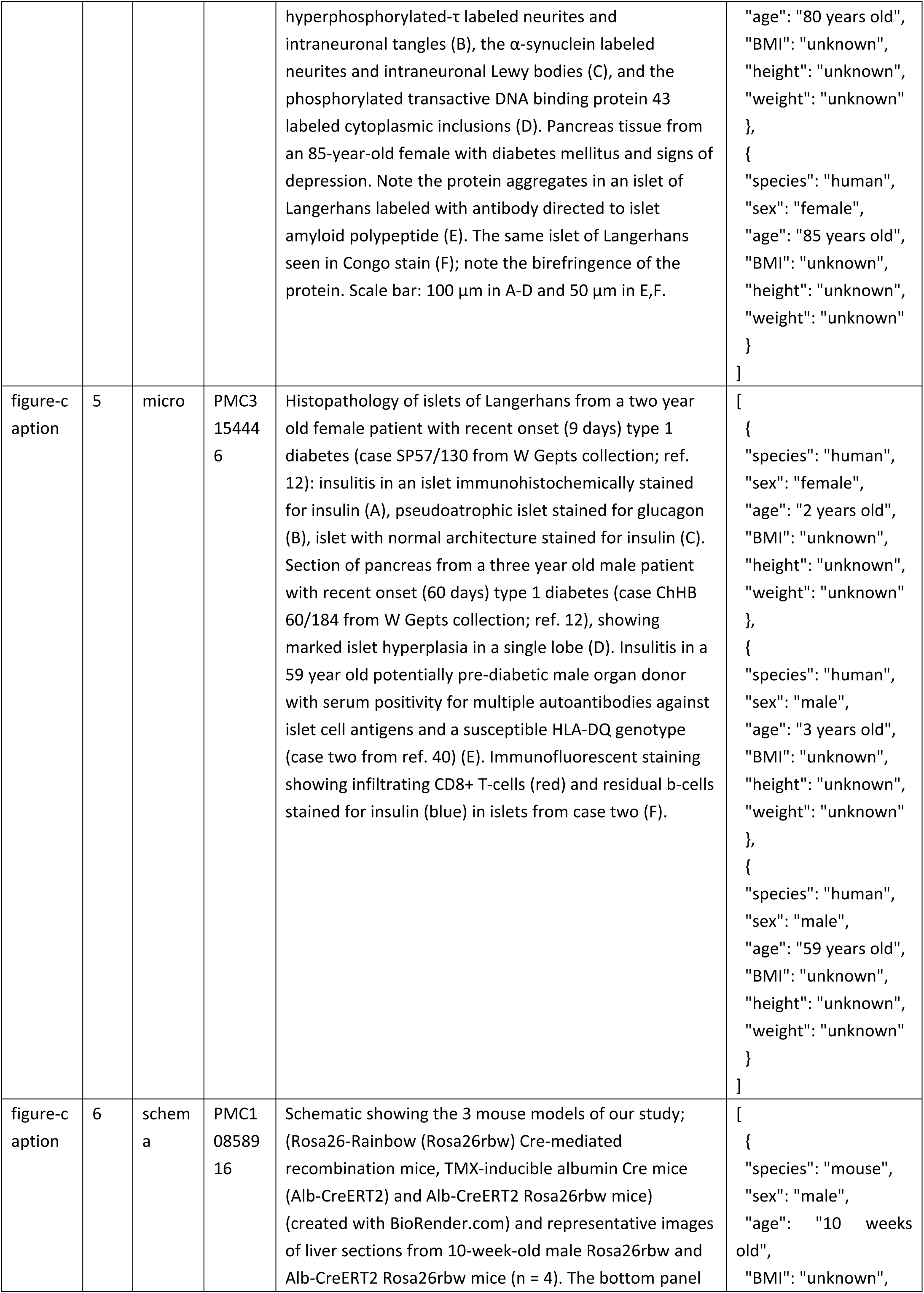
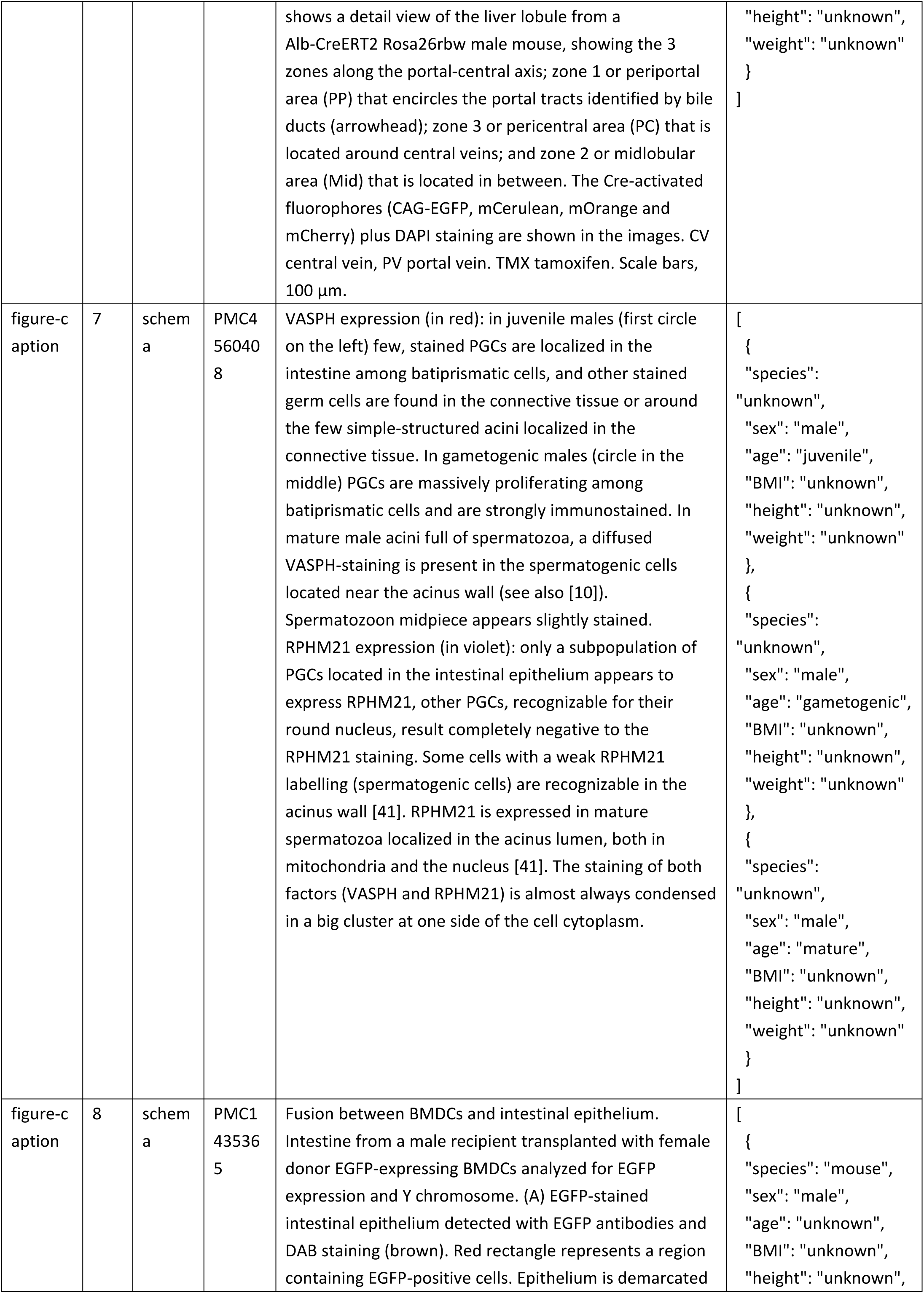
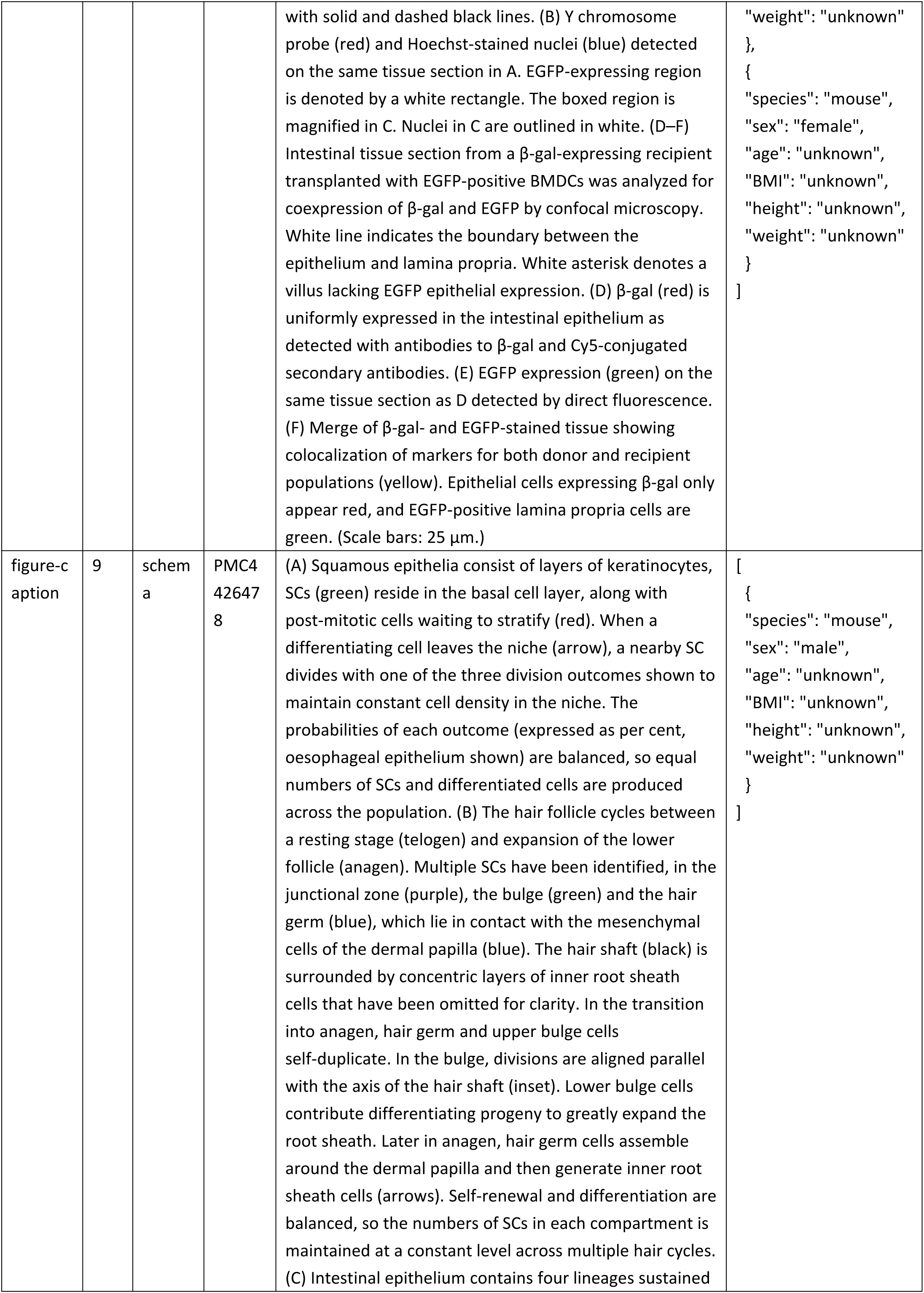
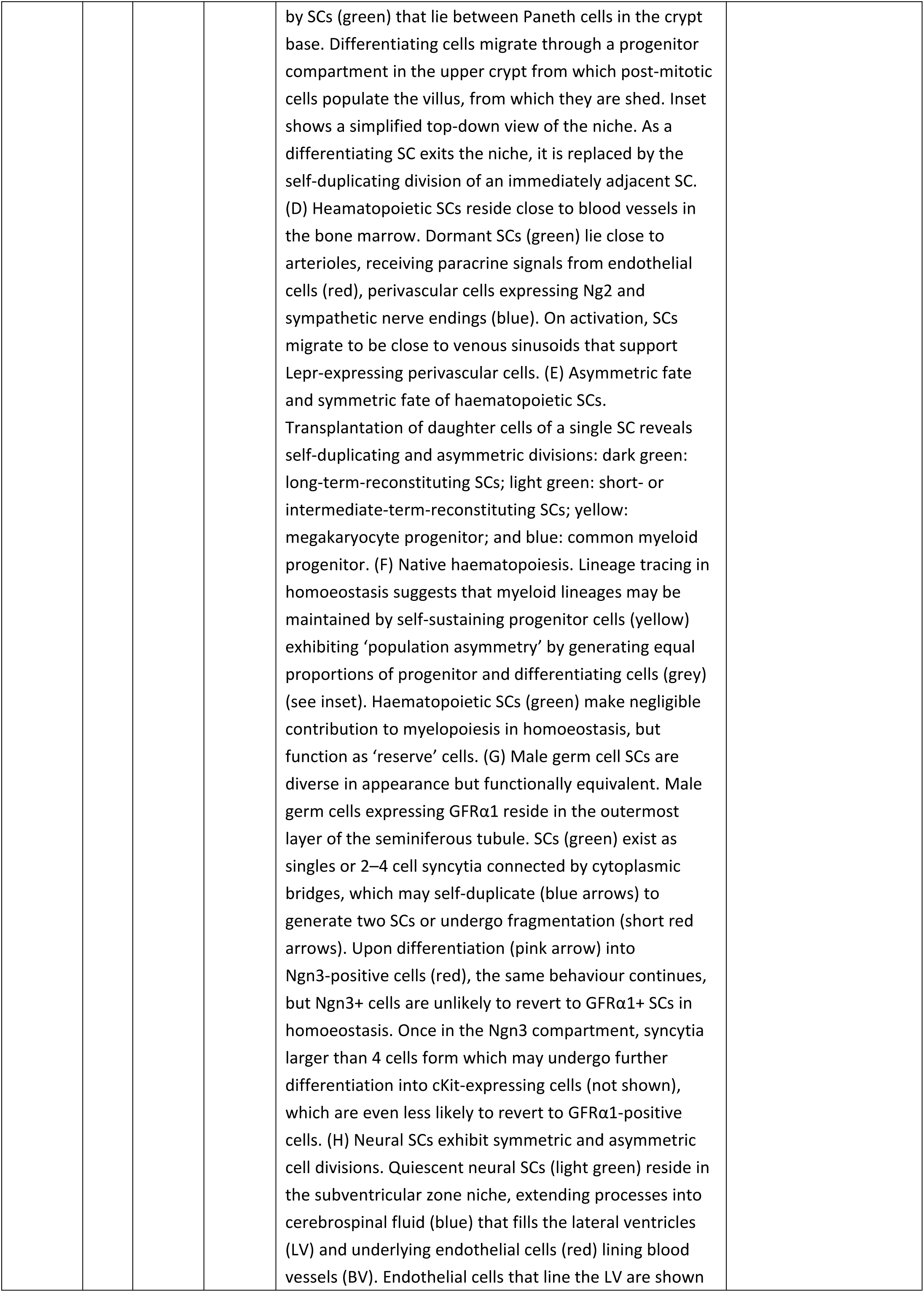
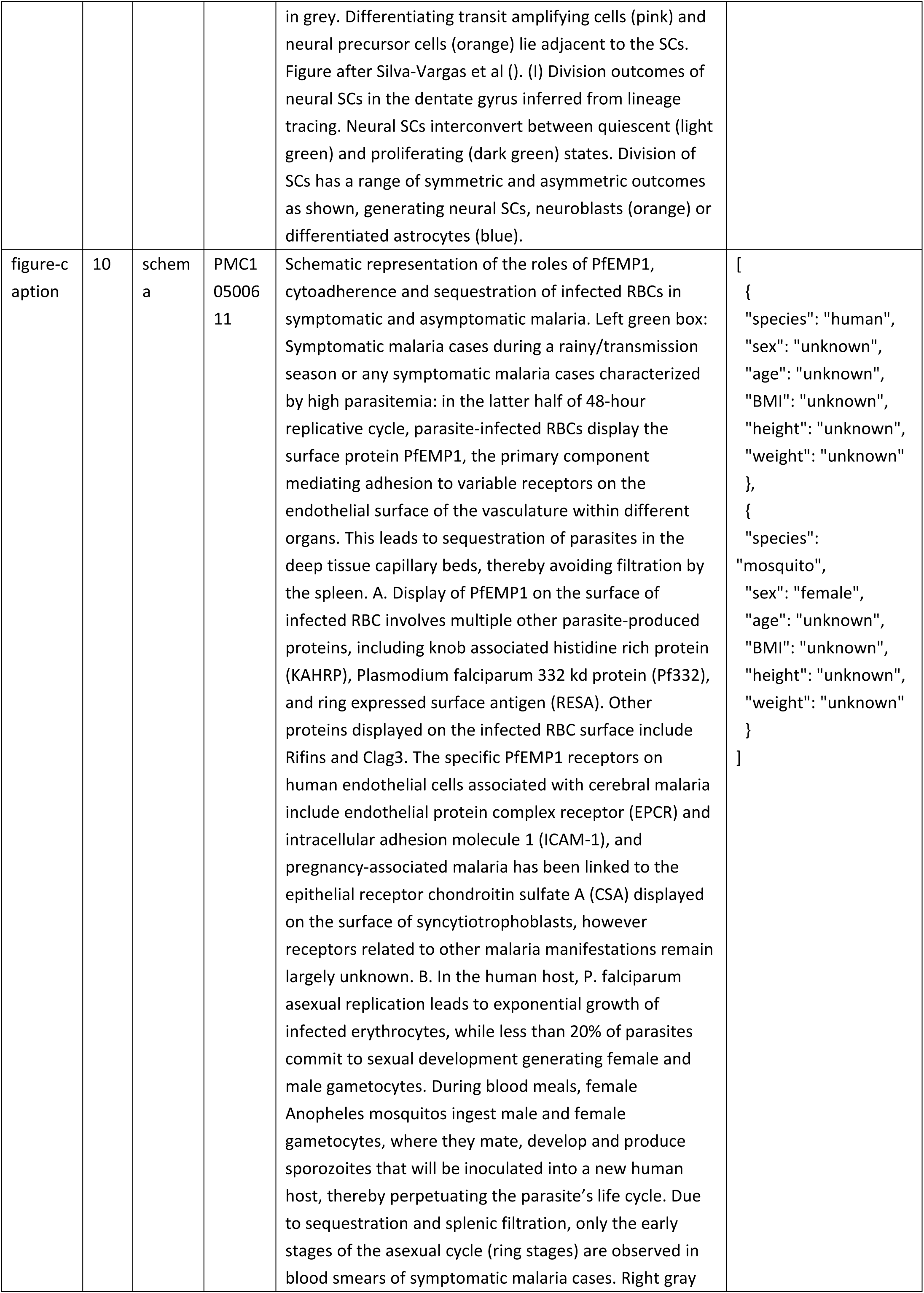
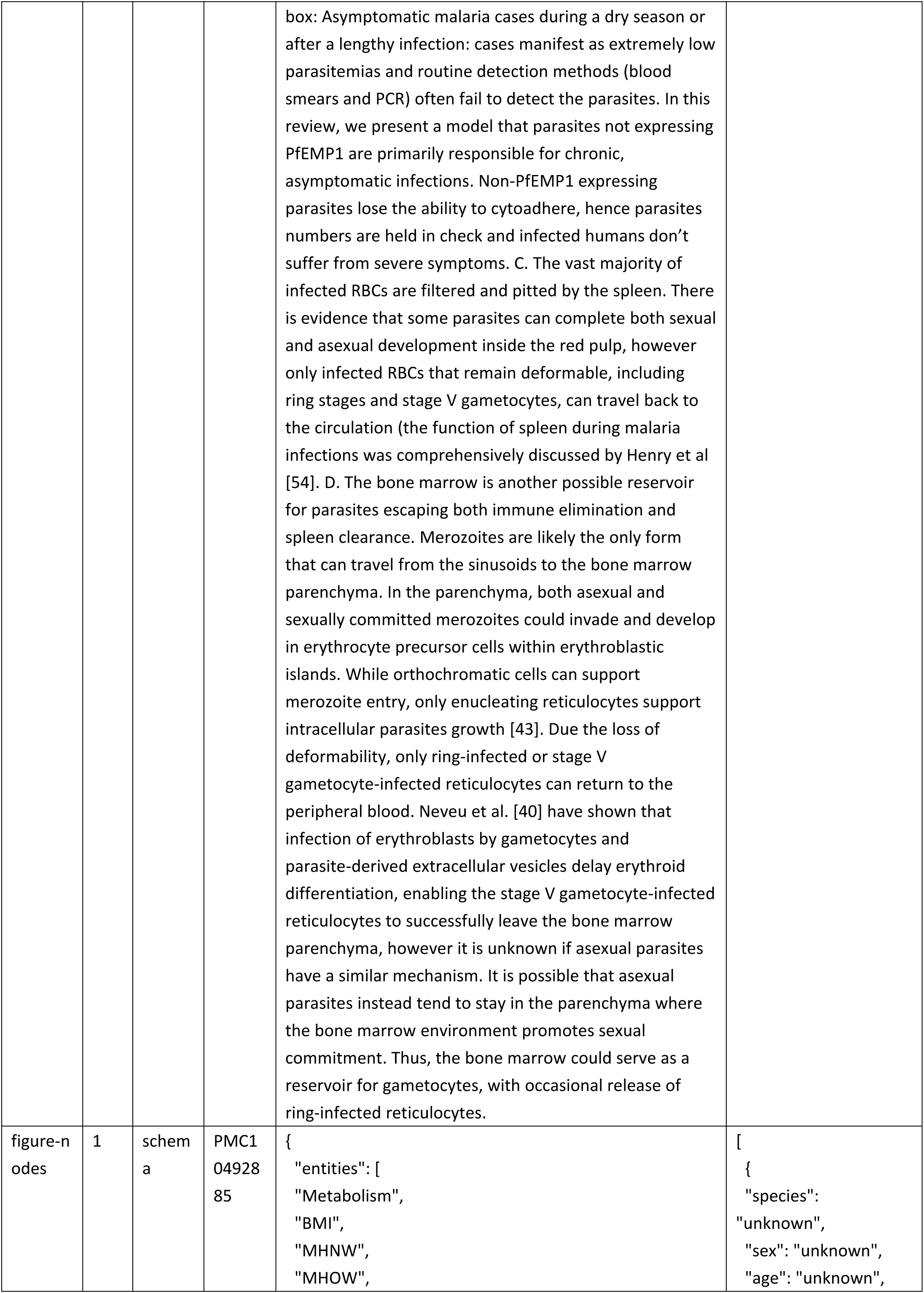
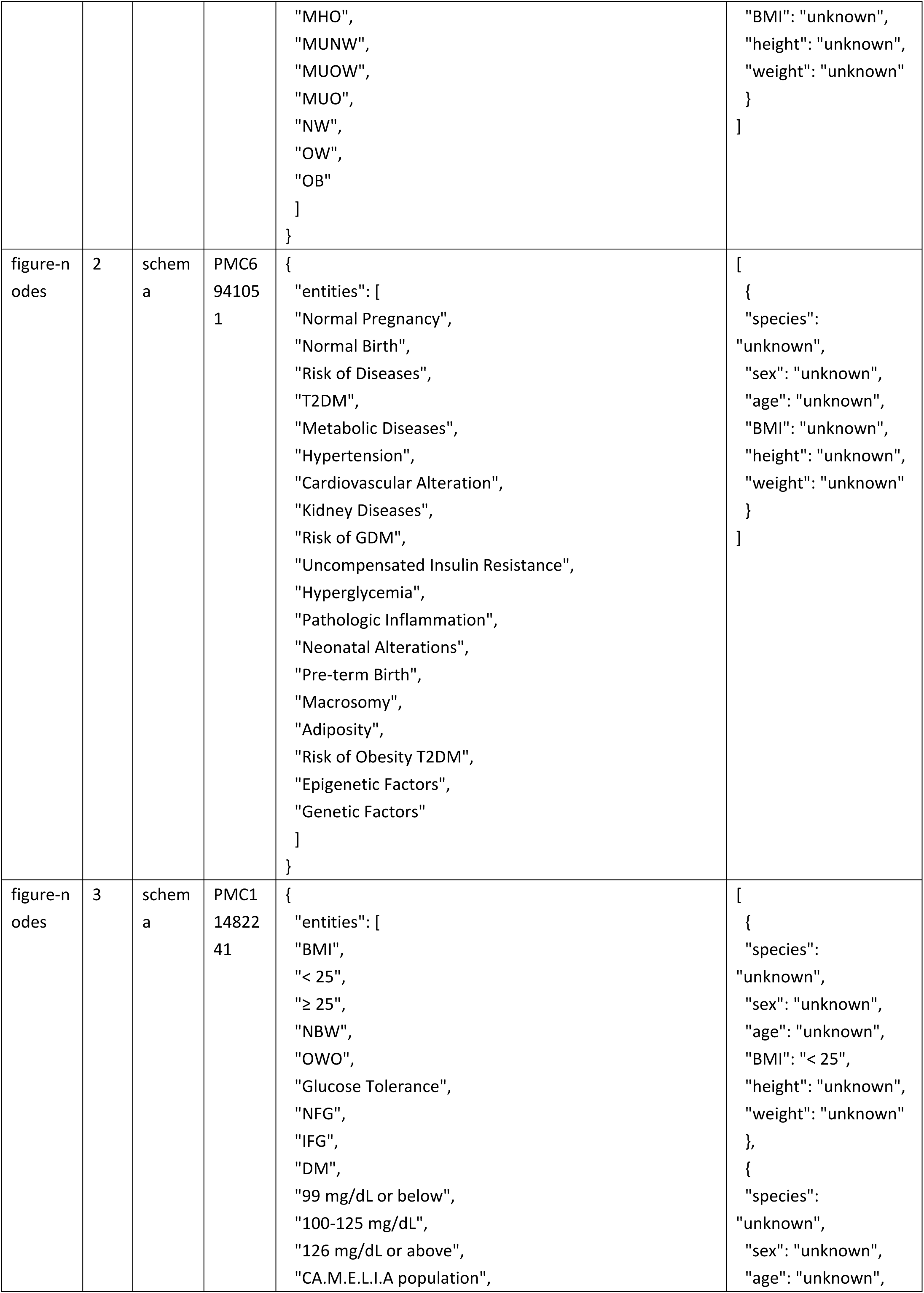
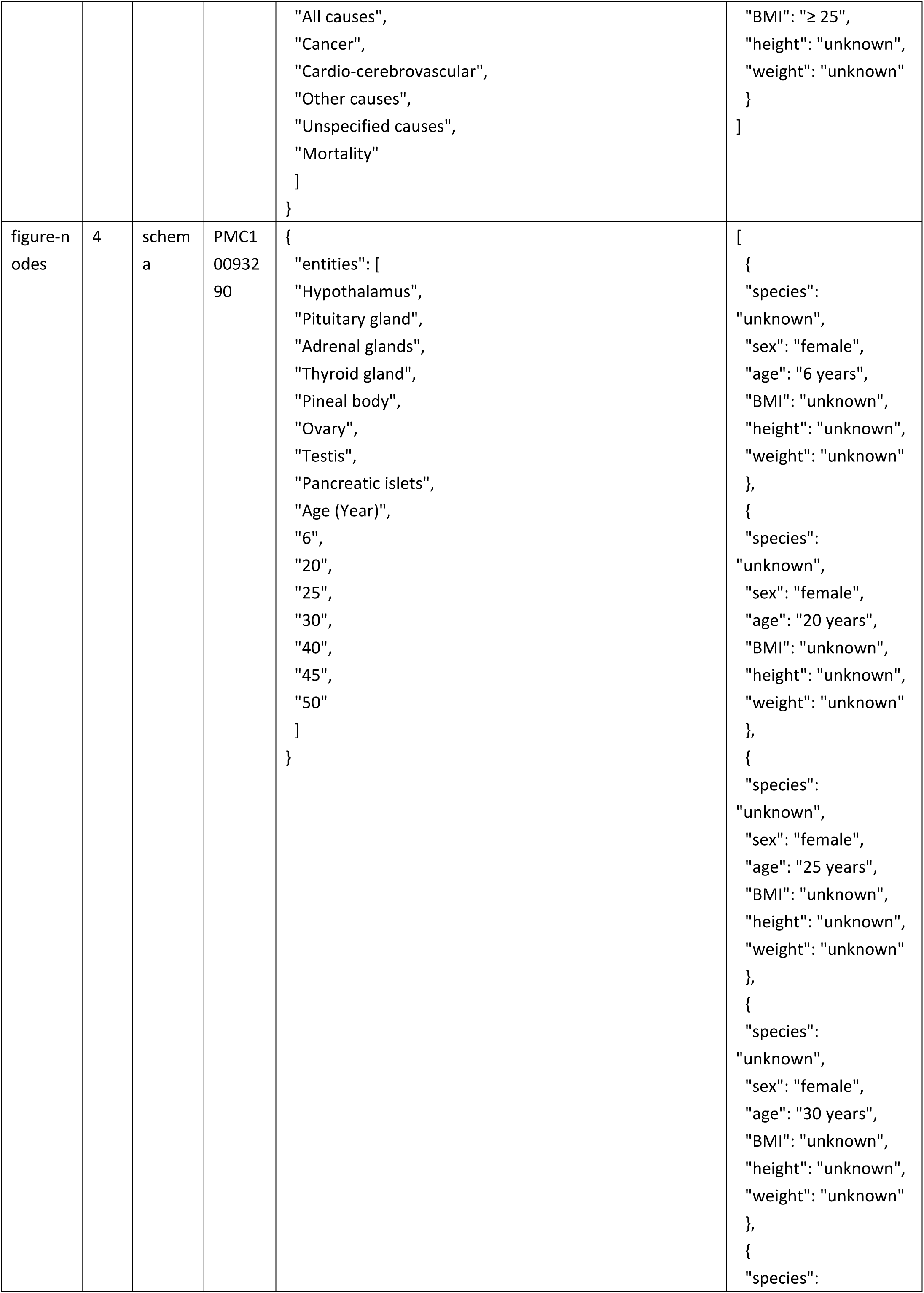
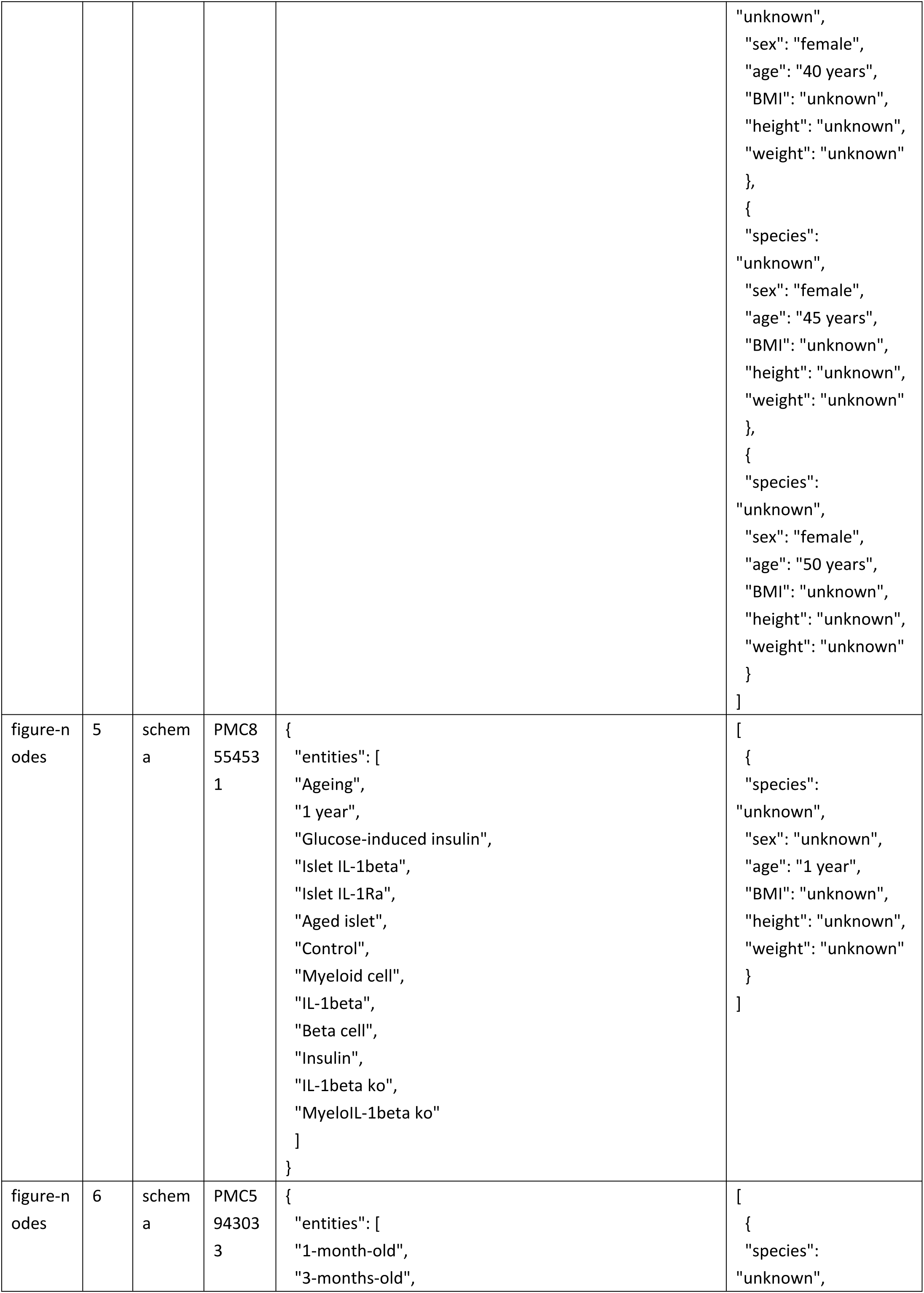
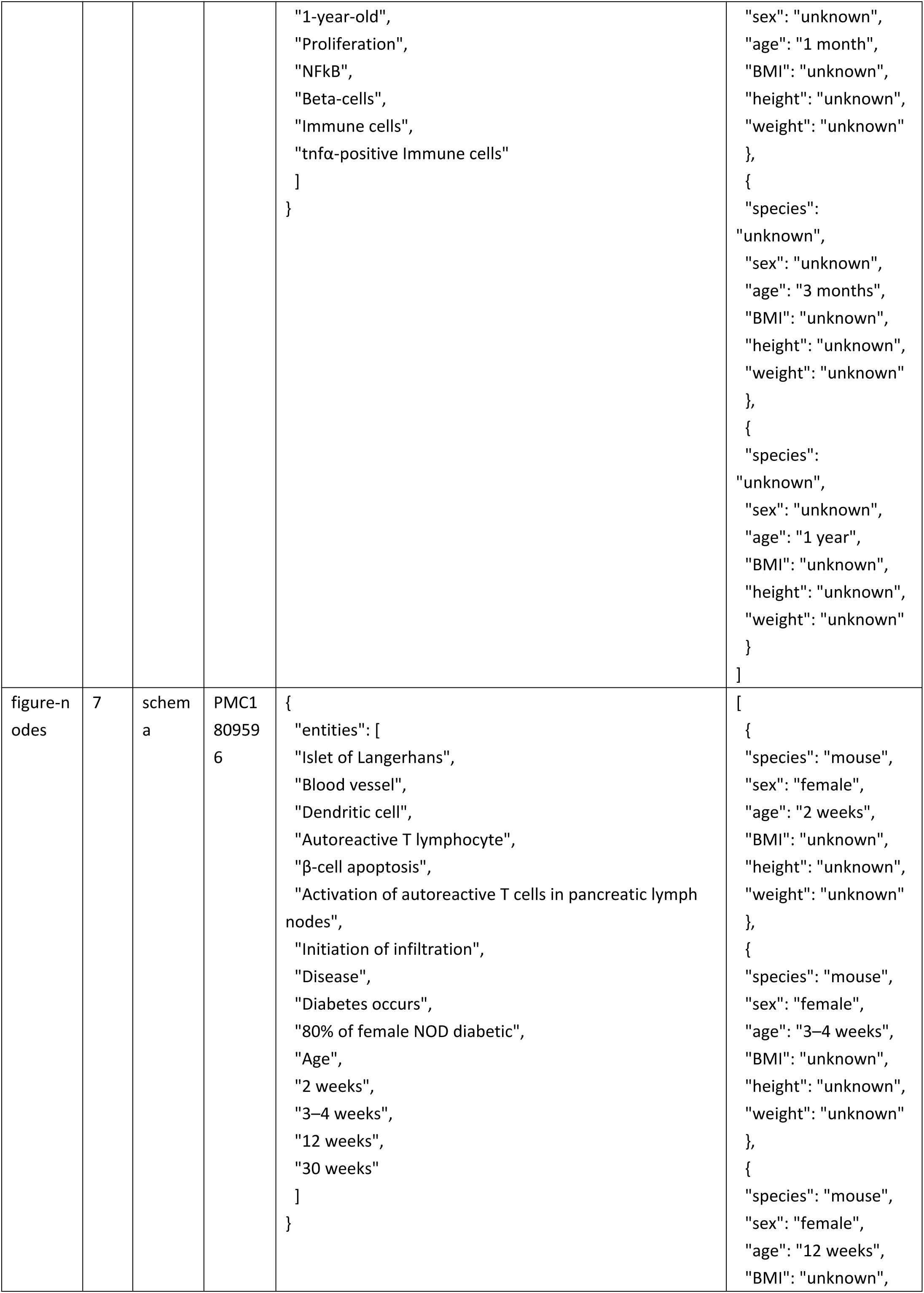
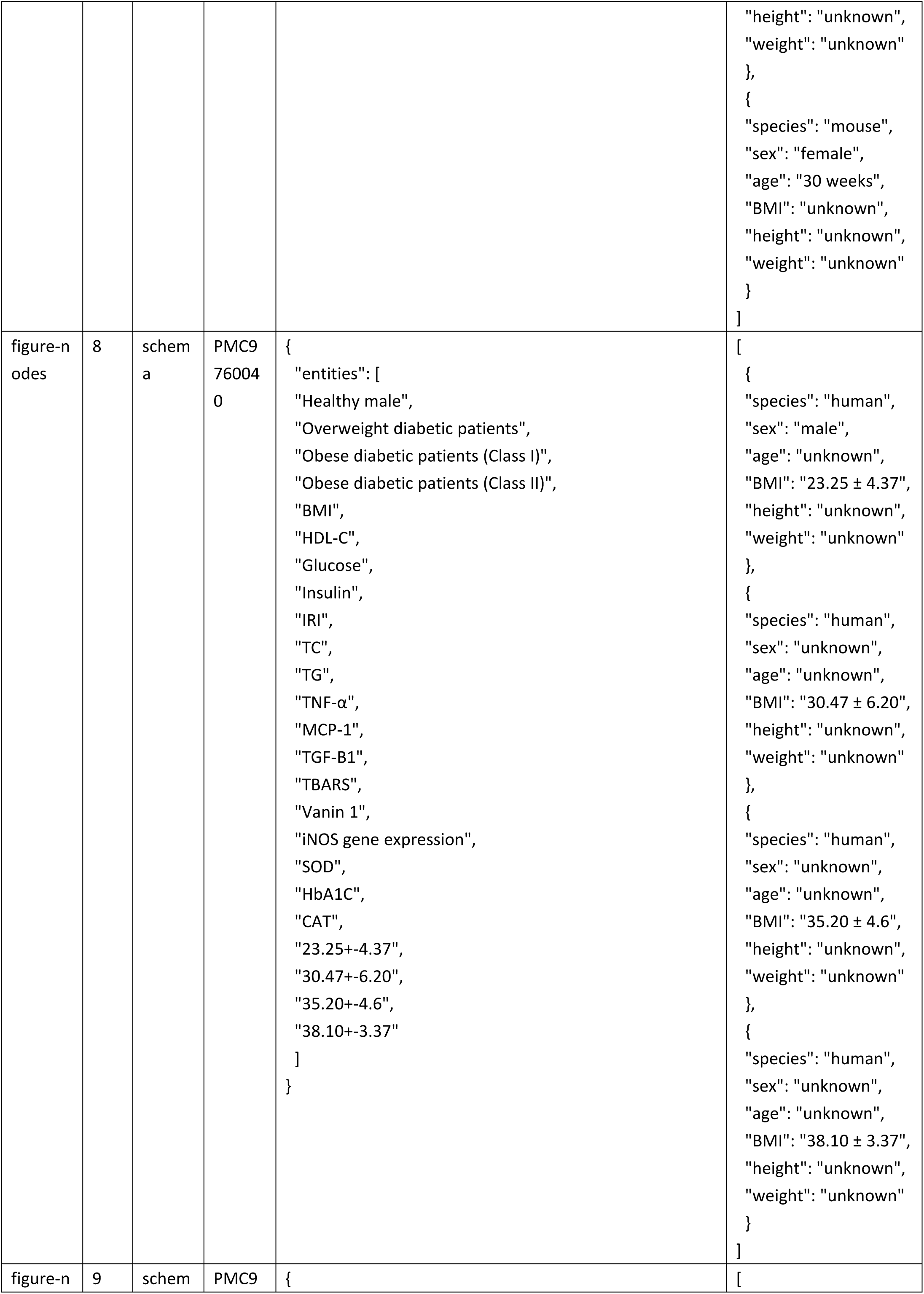
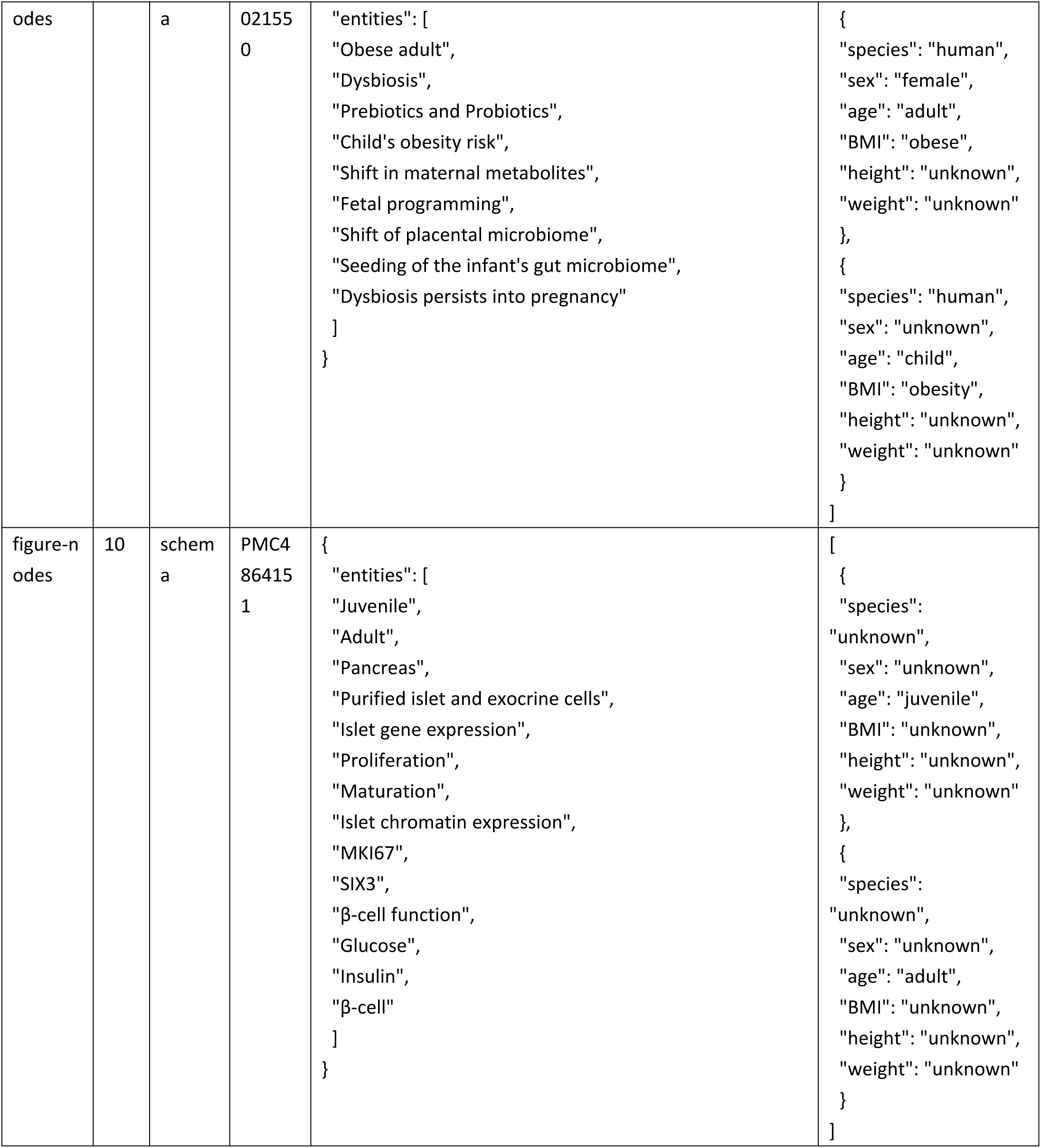
Ground true for donor-metadata extraction.

**Supplementary Table 21.**
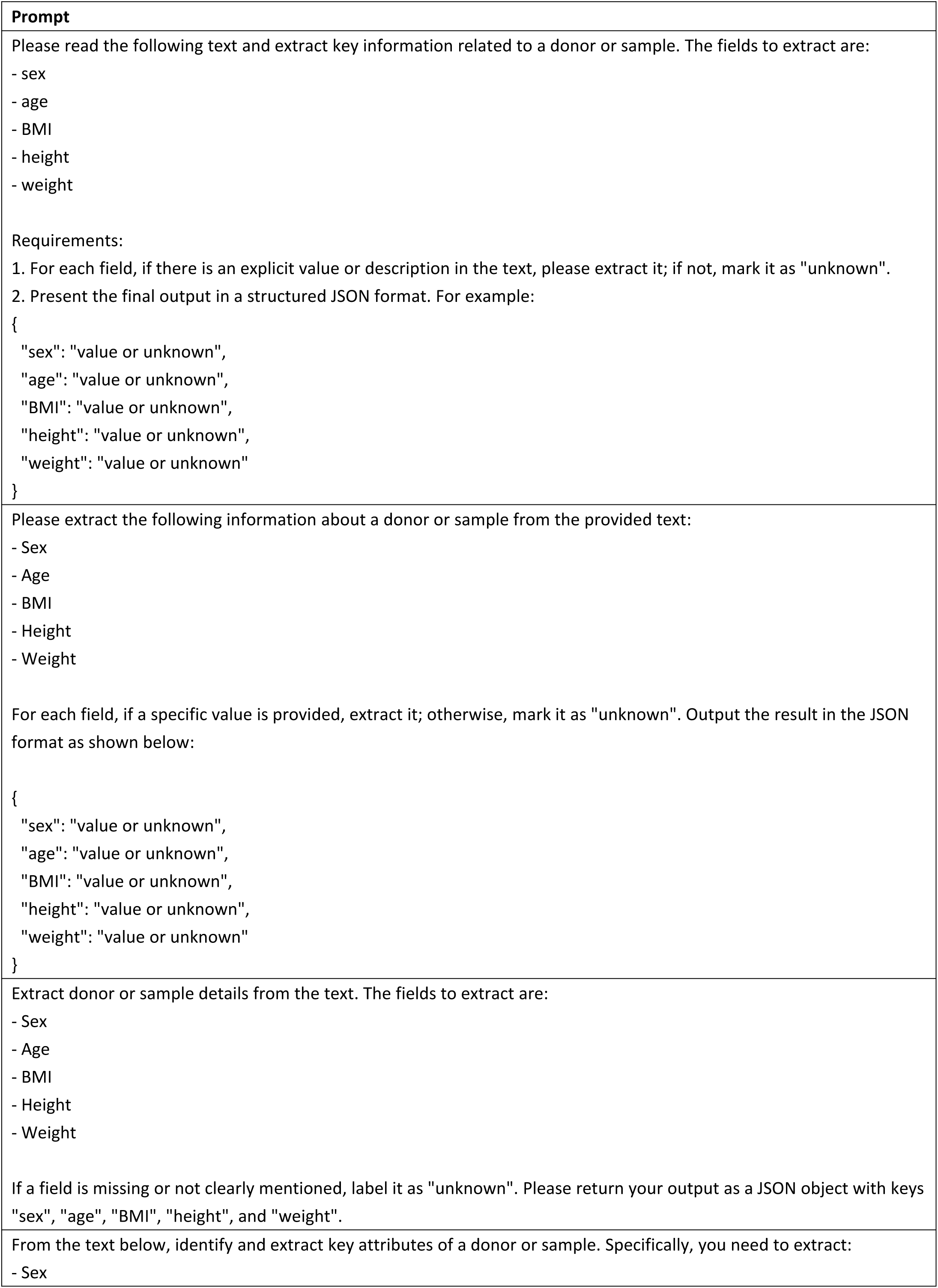
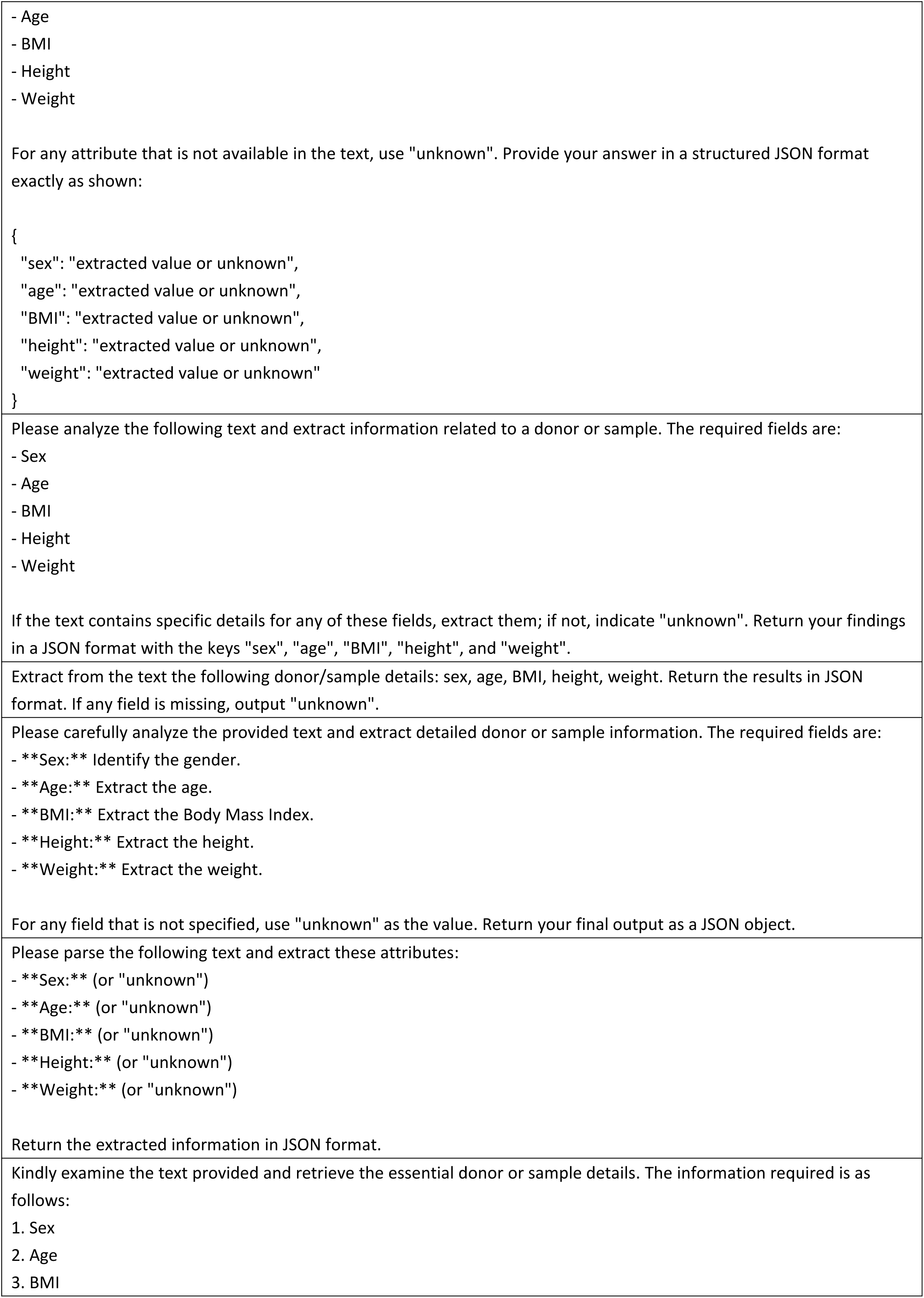
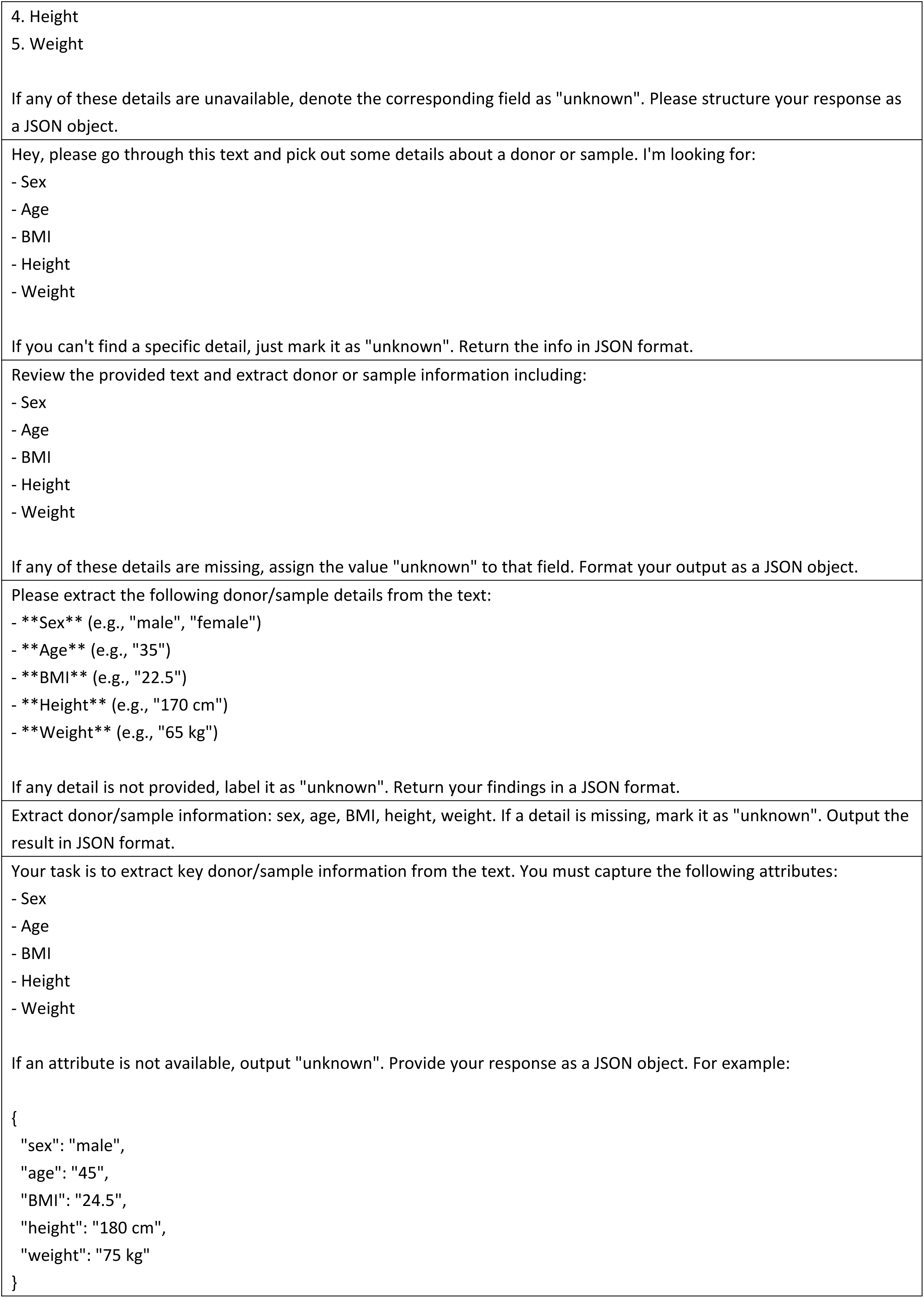
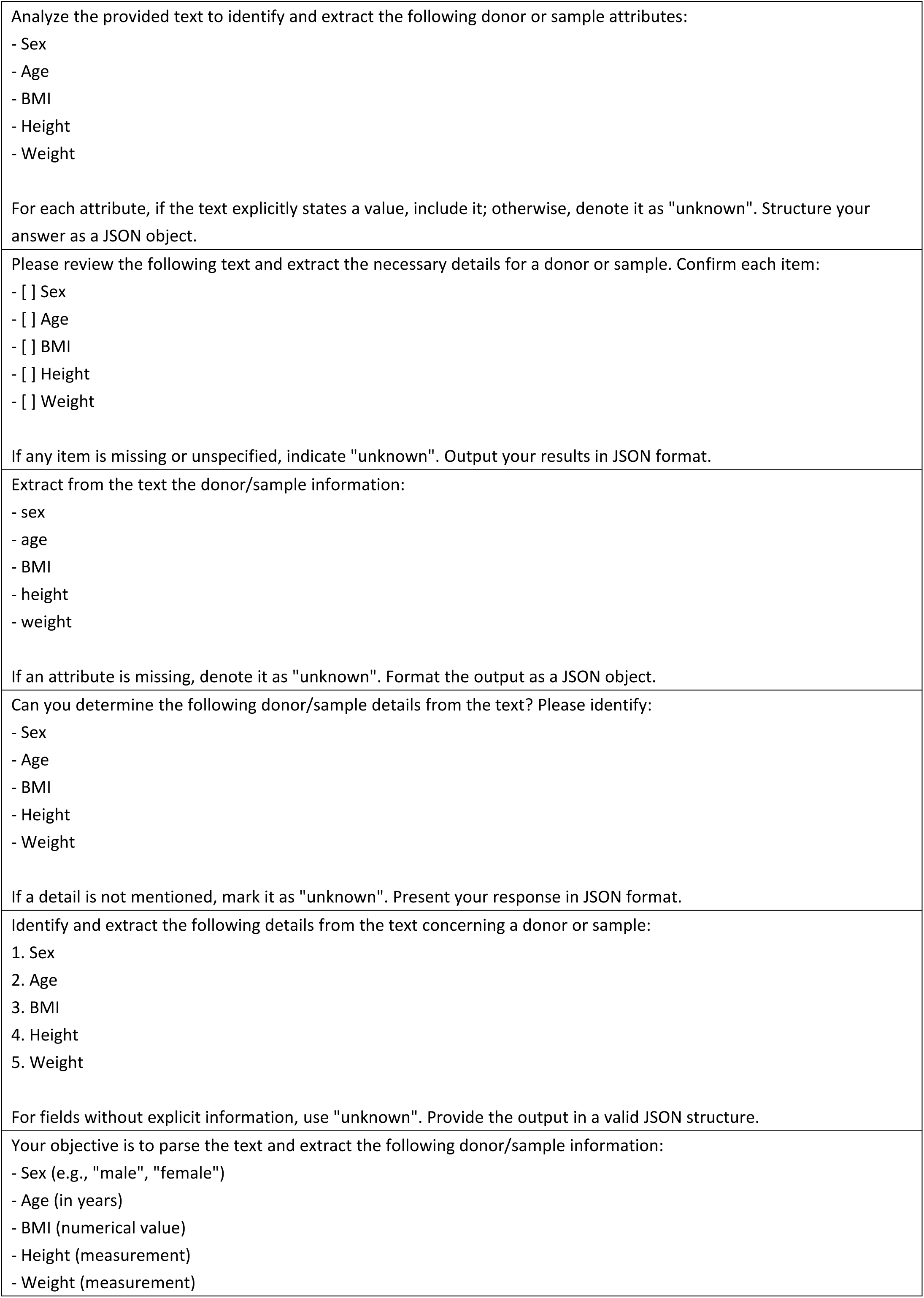
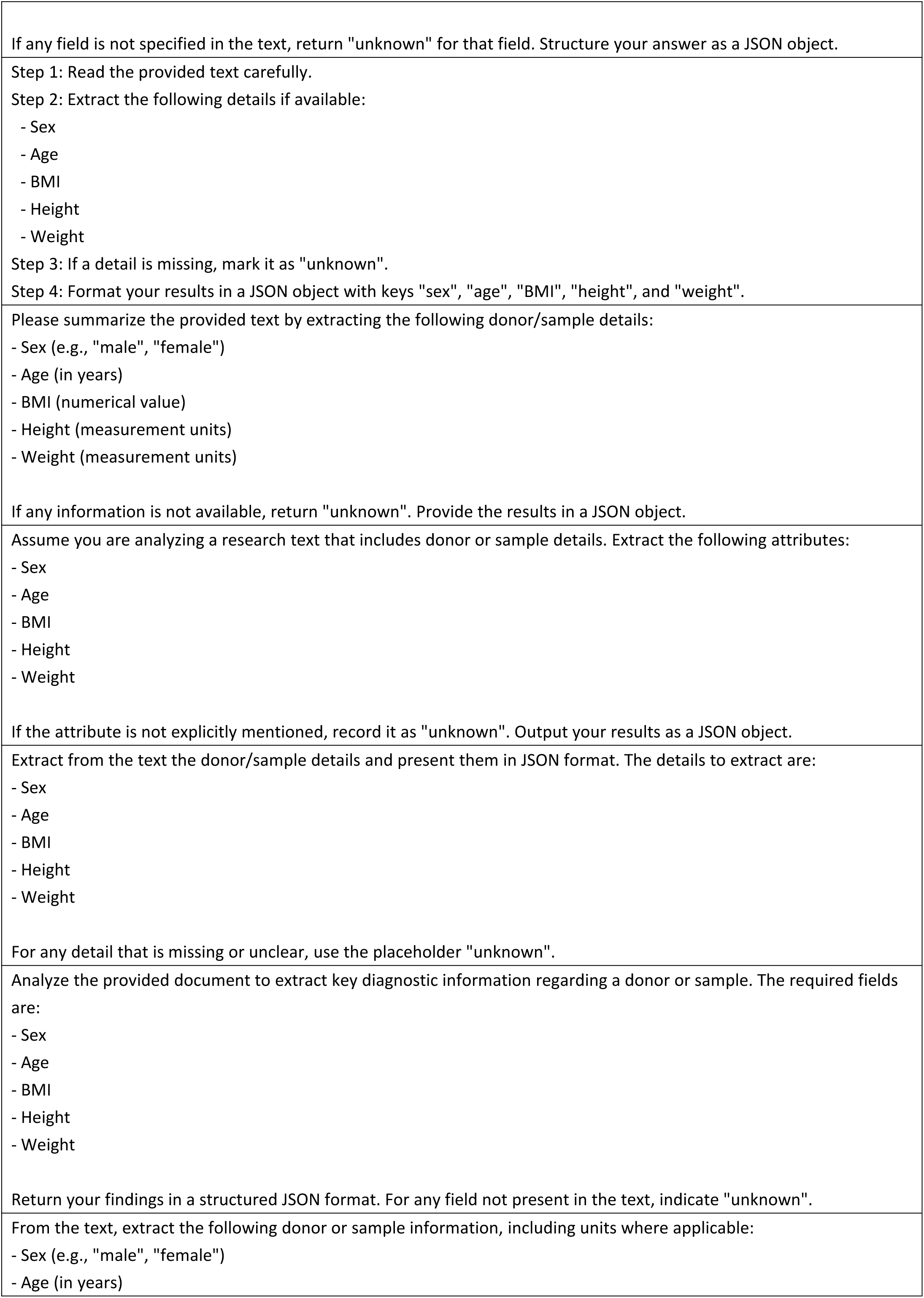
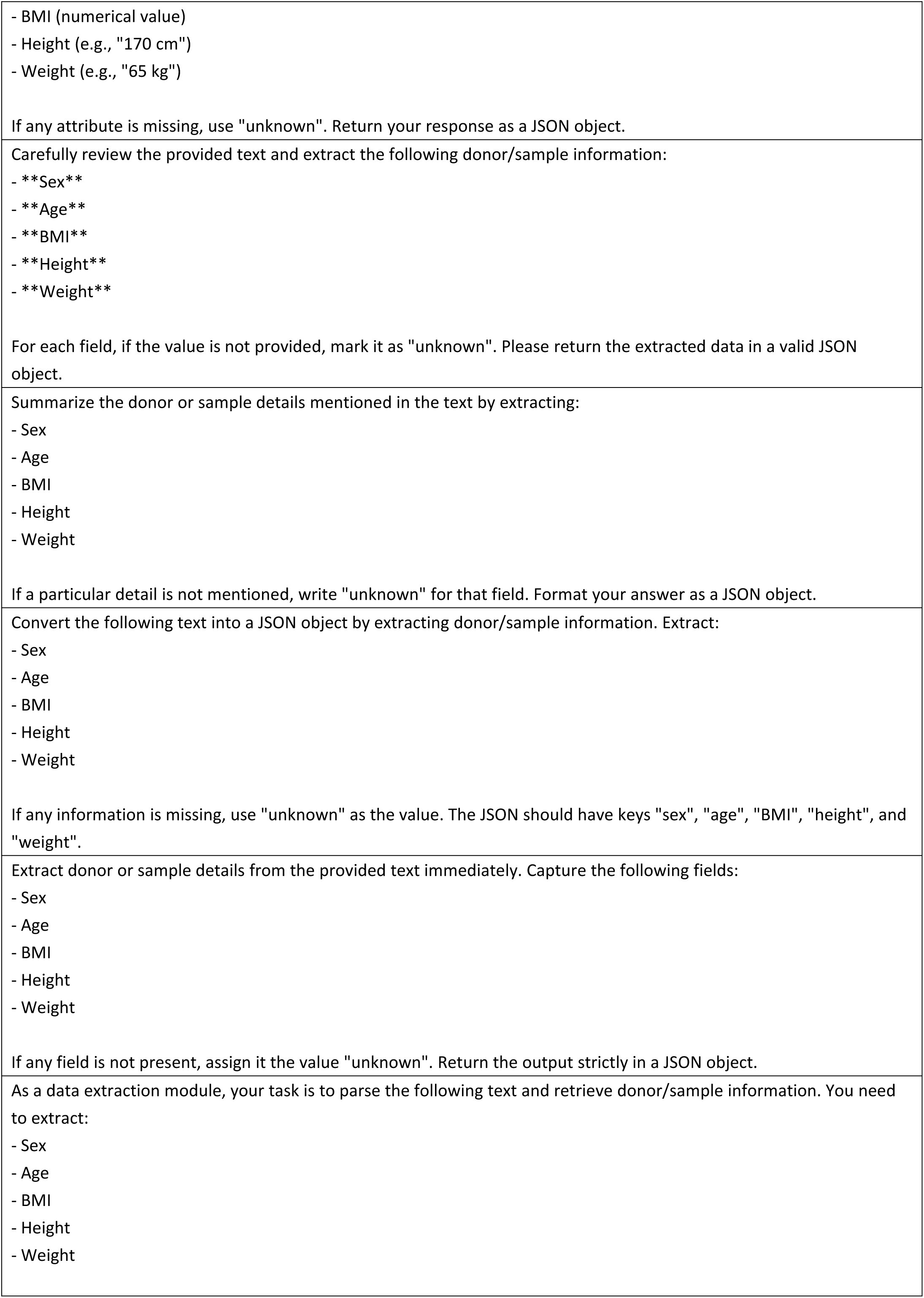
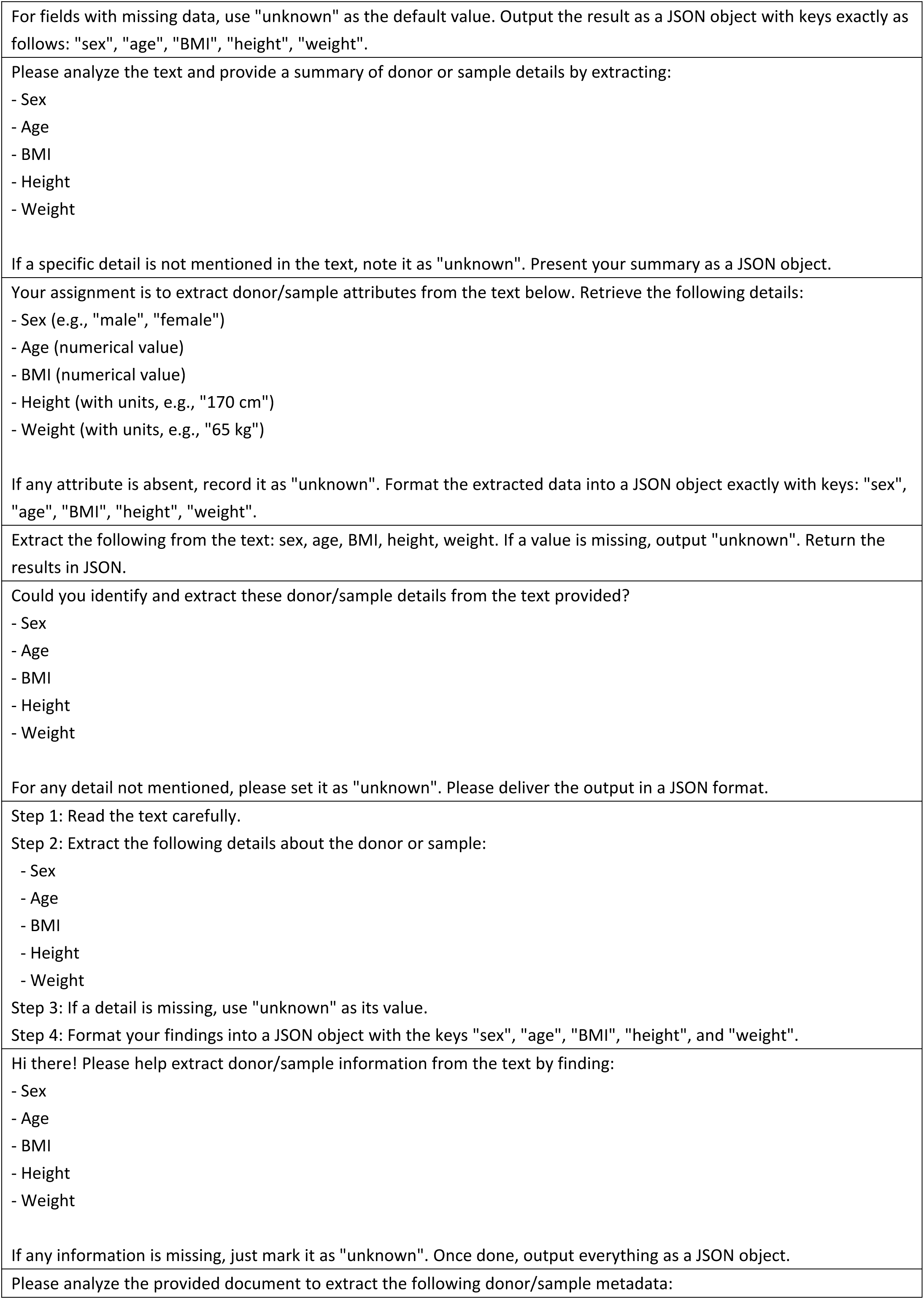
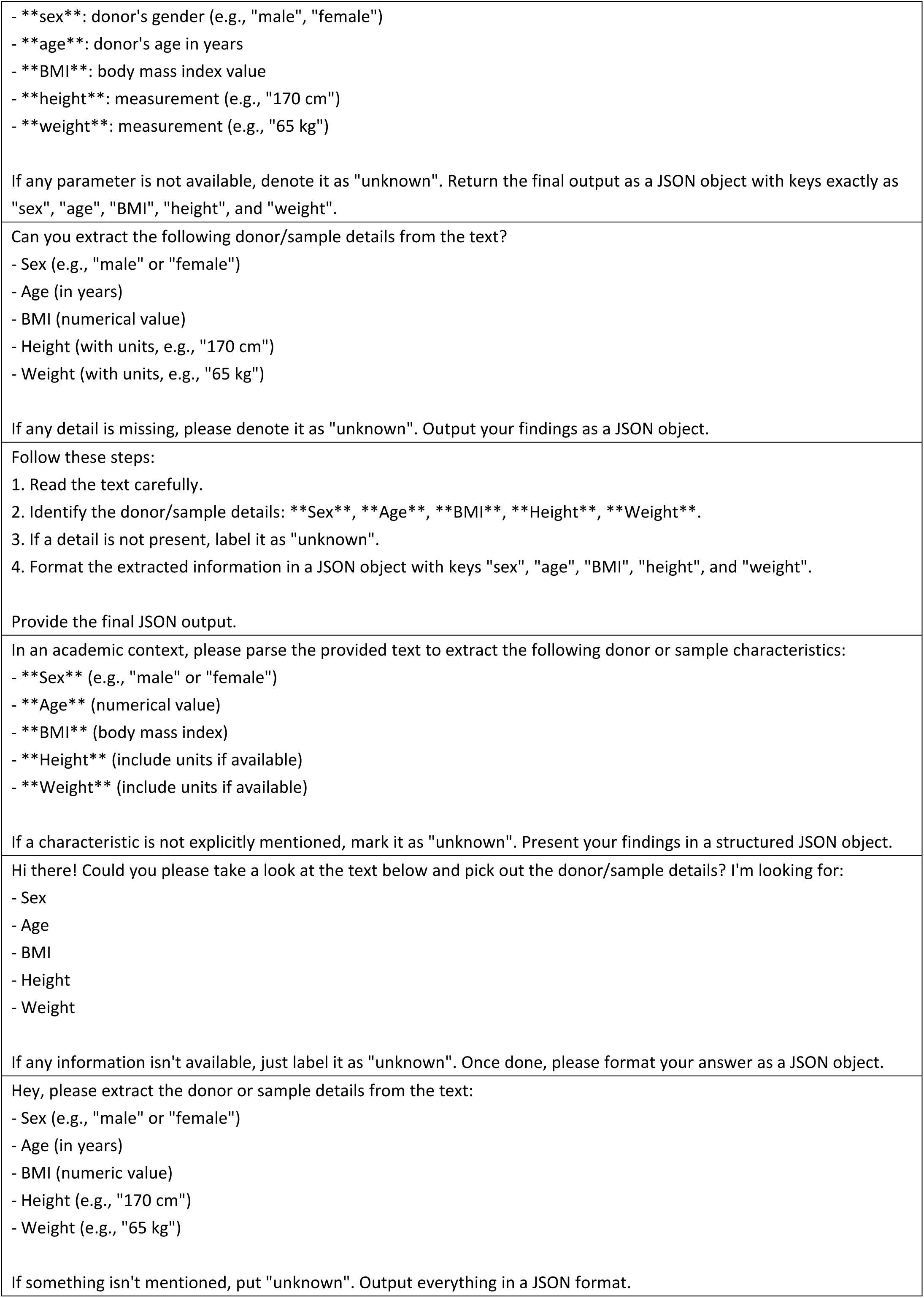
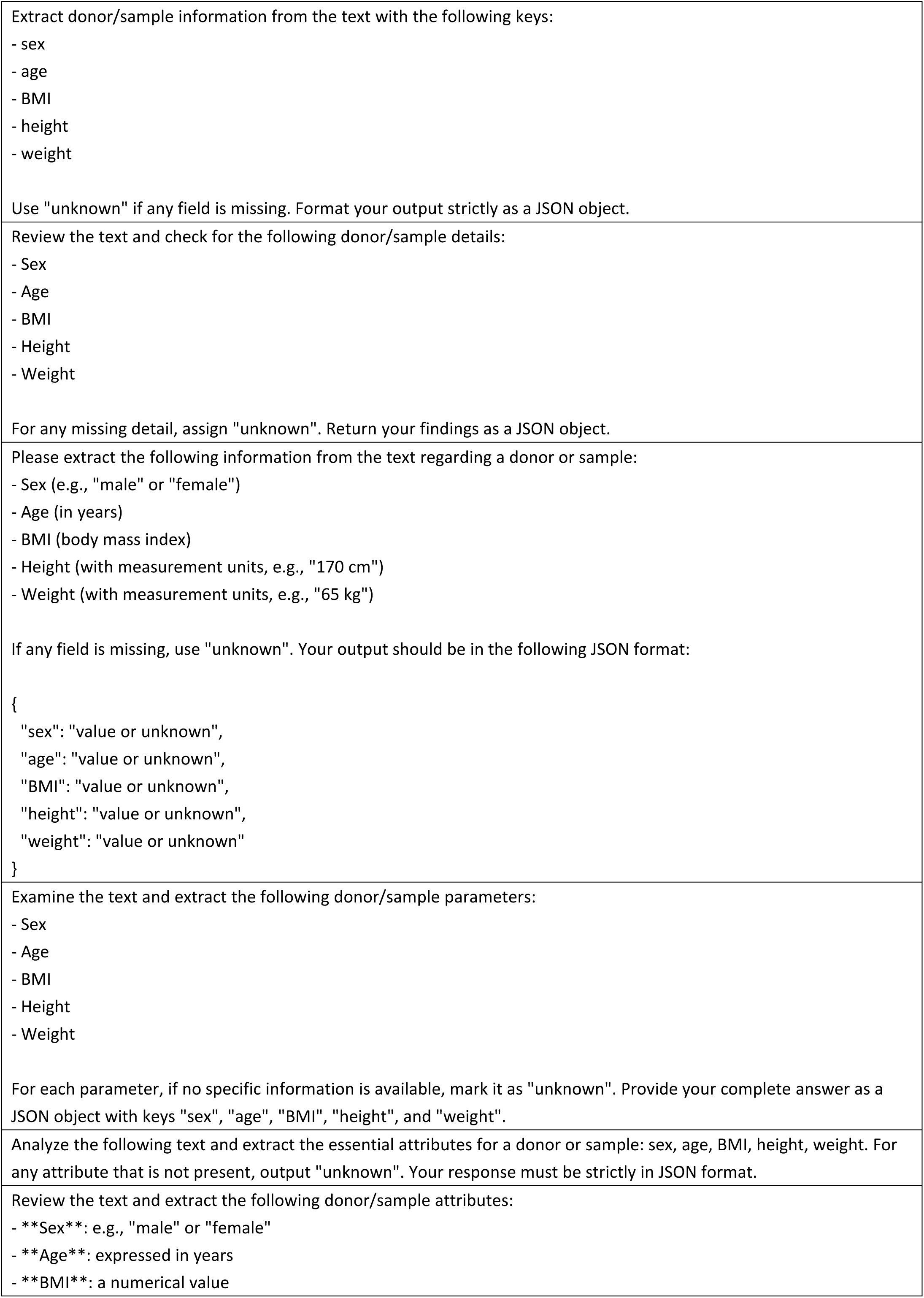
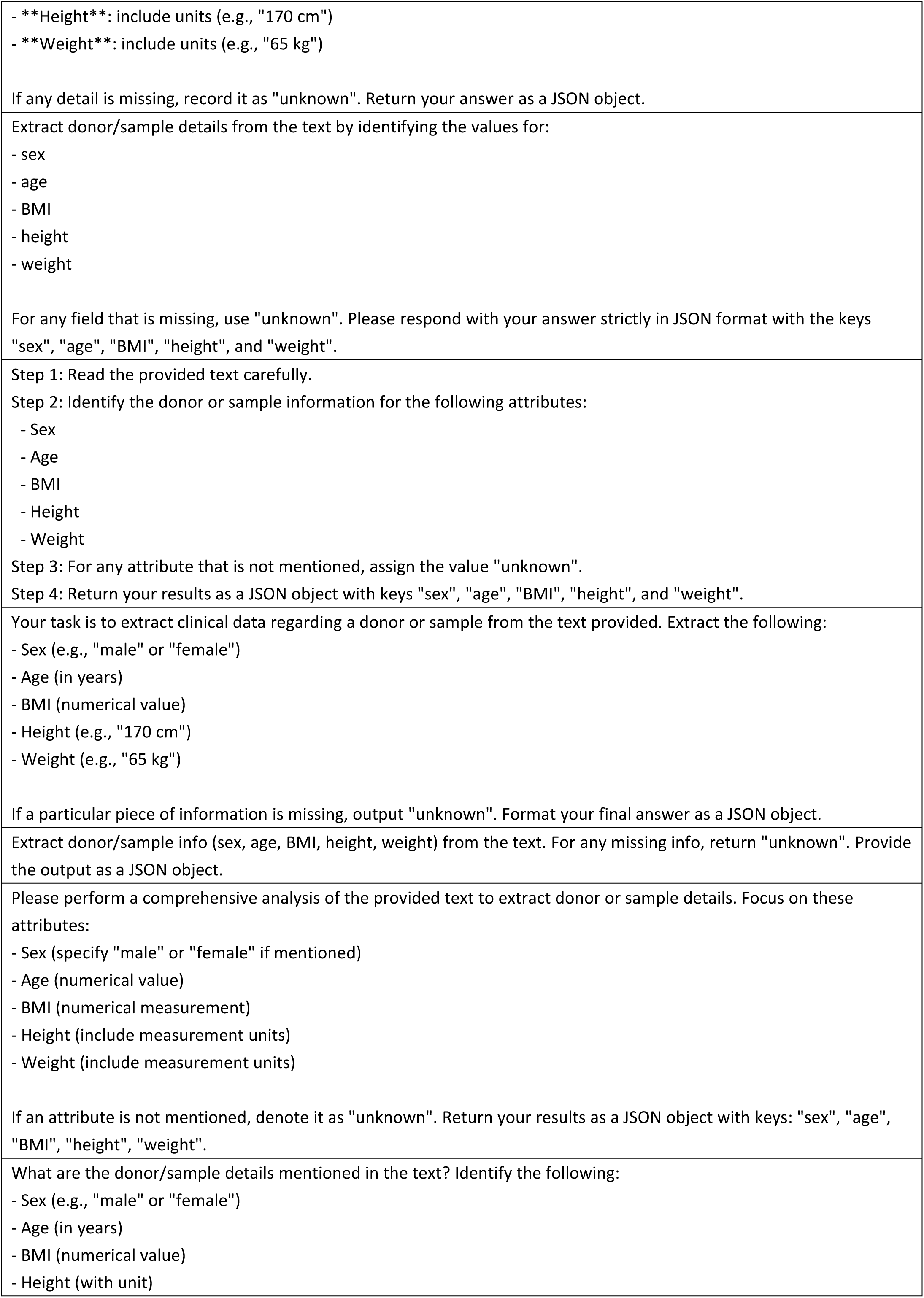
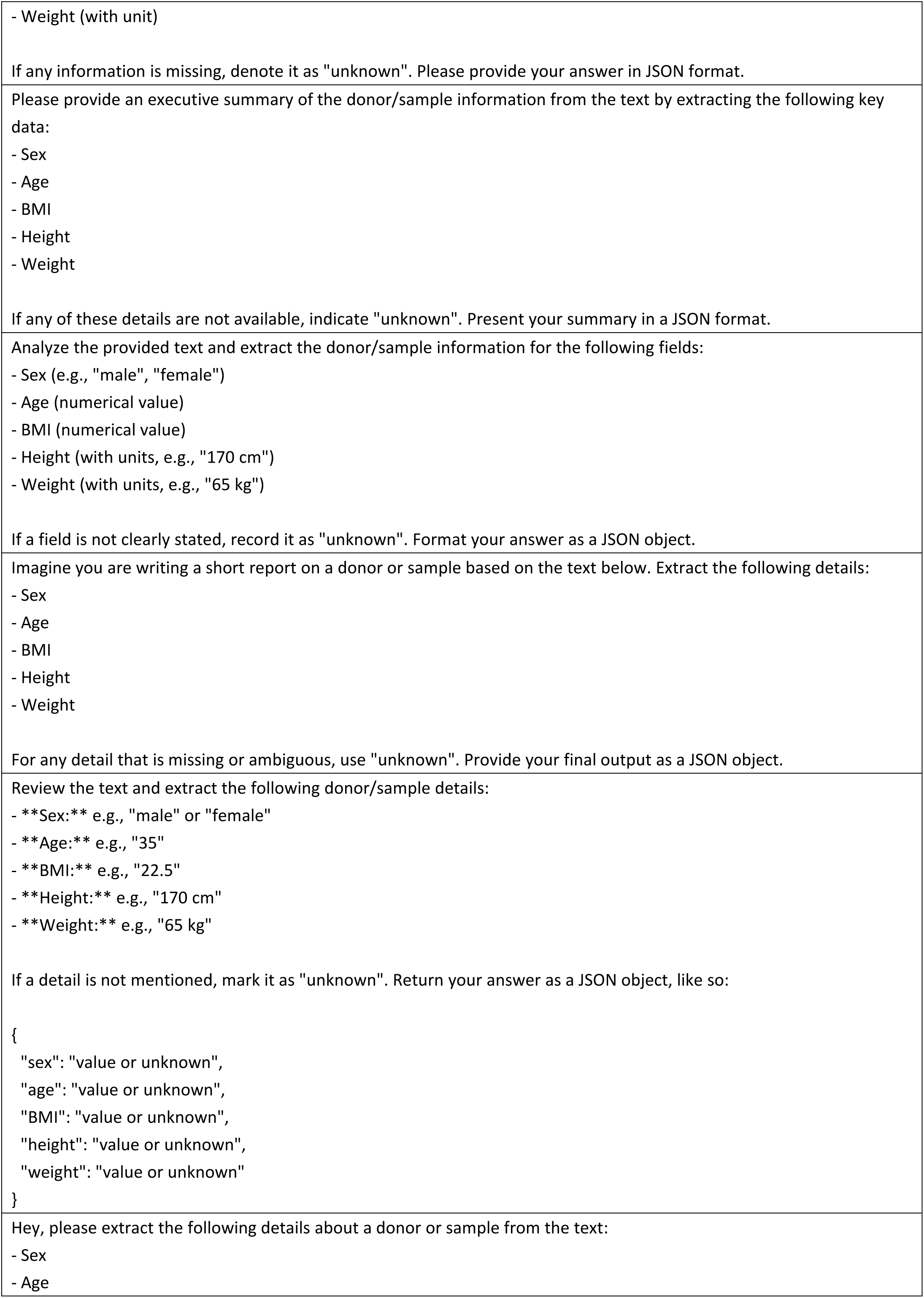
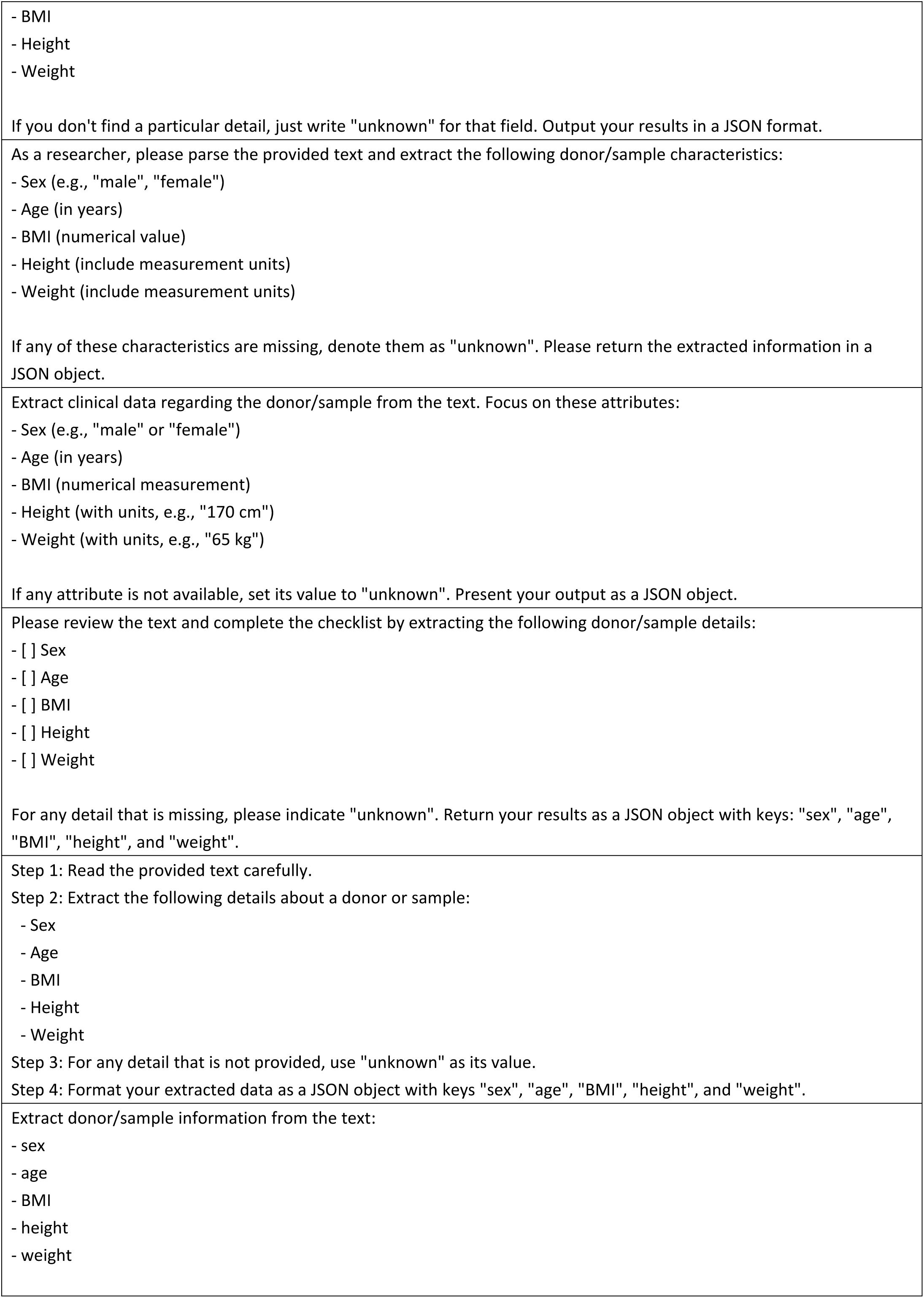
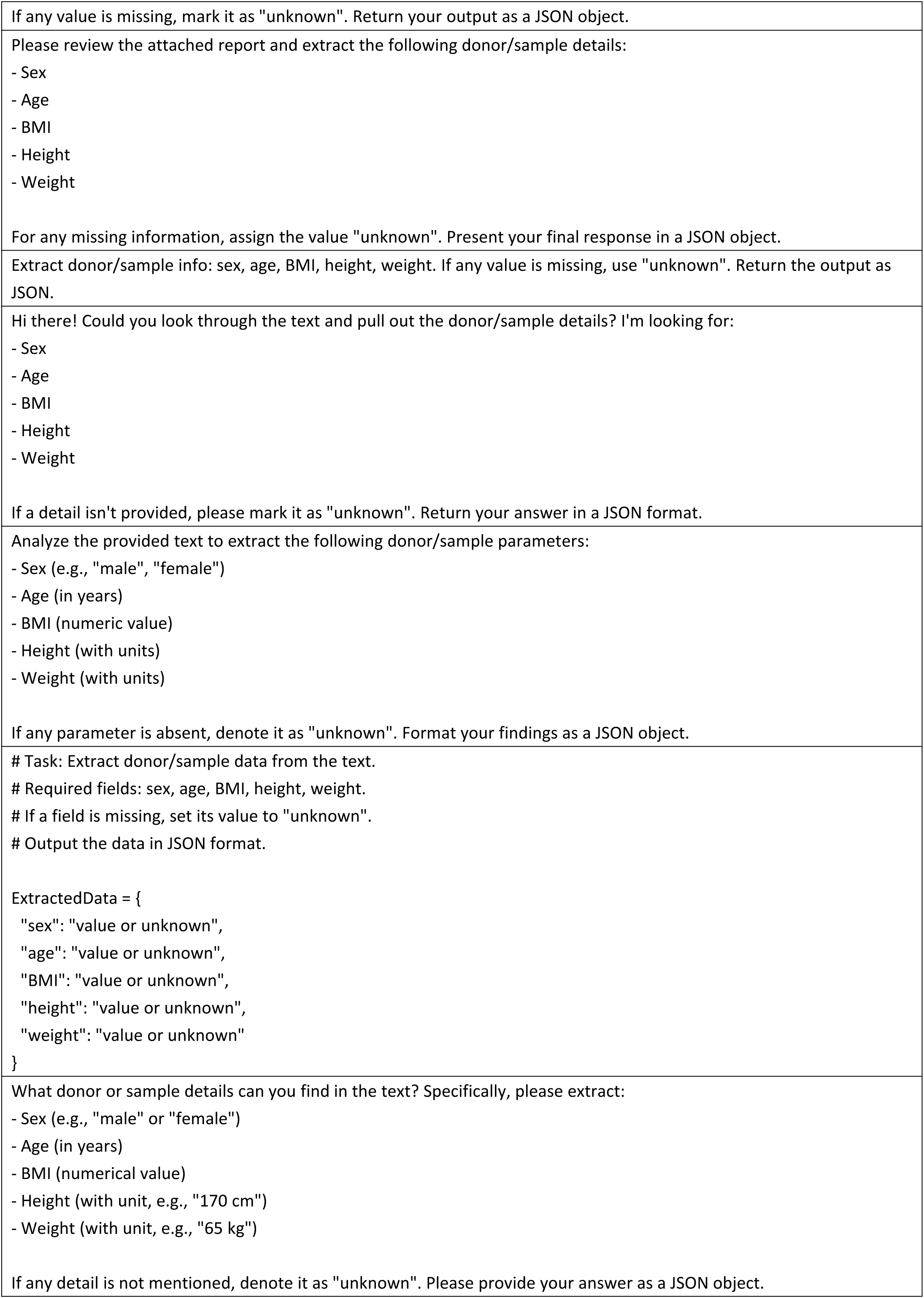
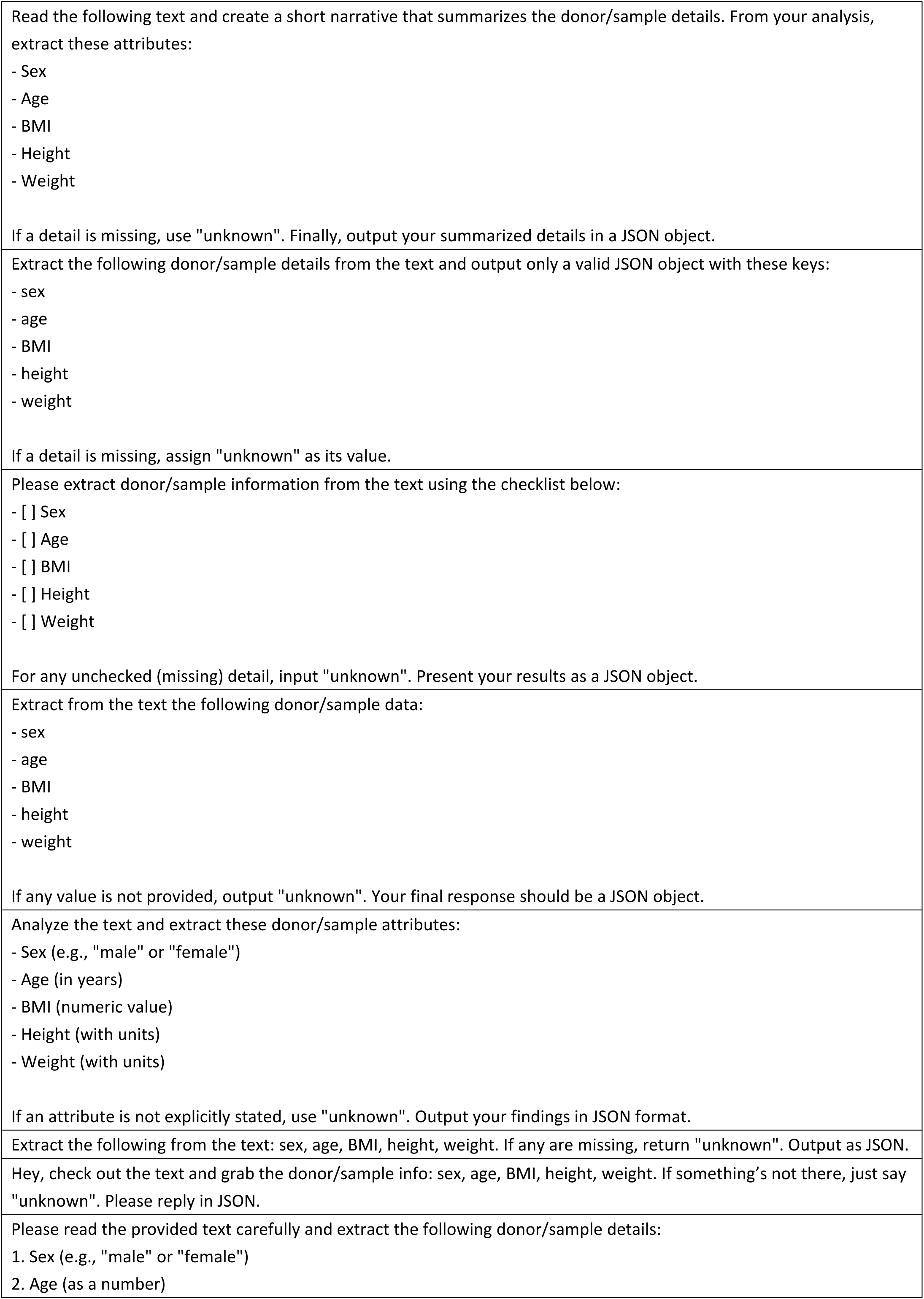
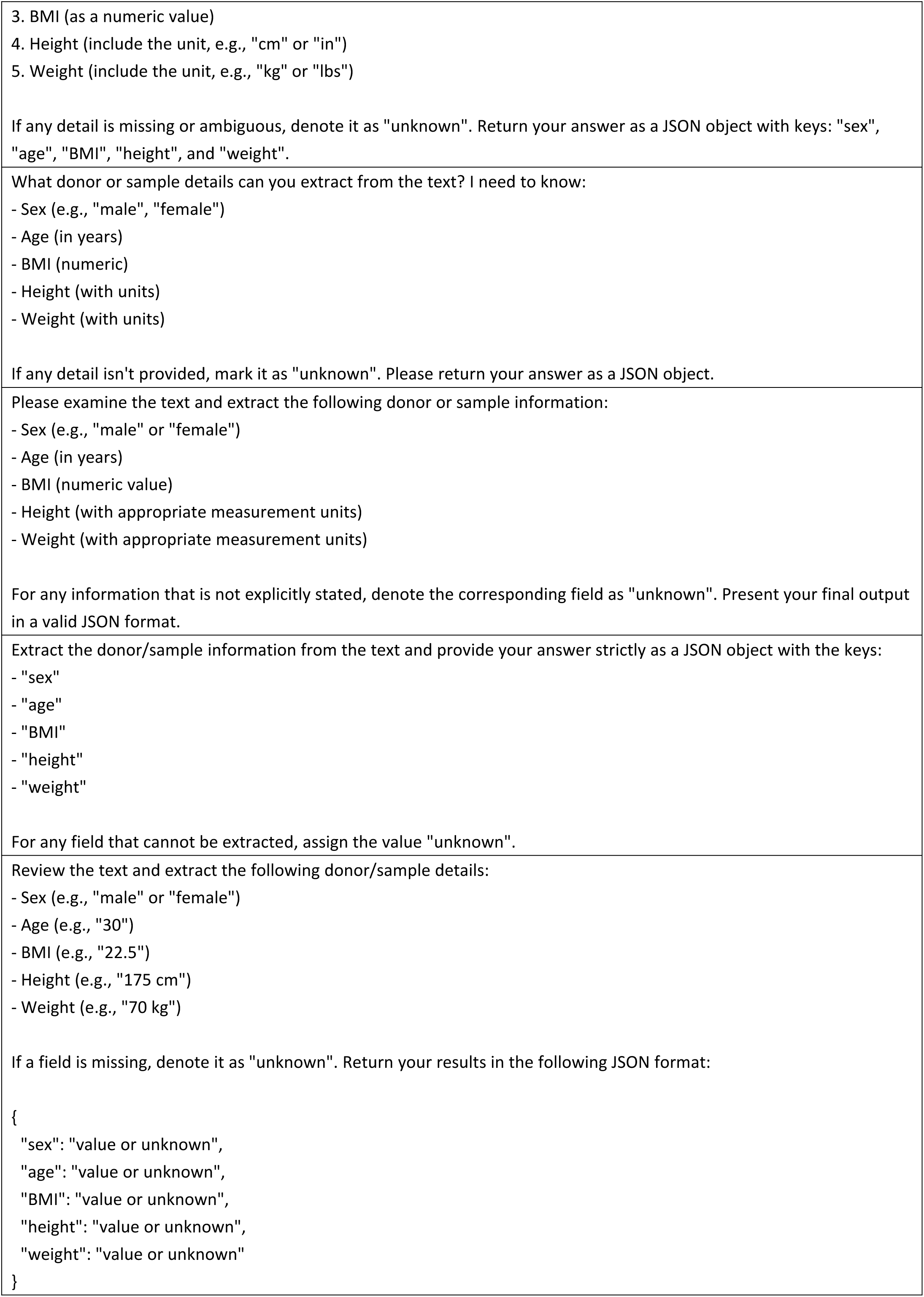
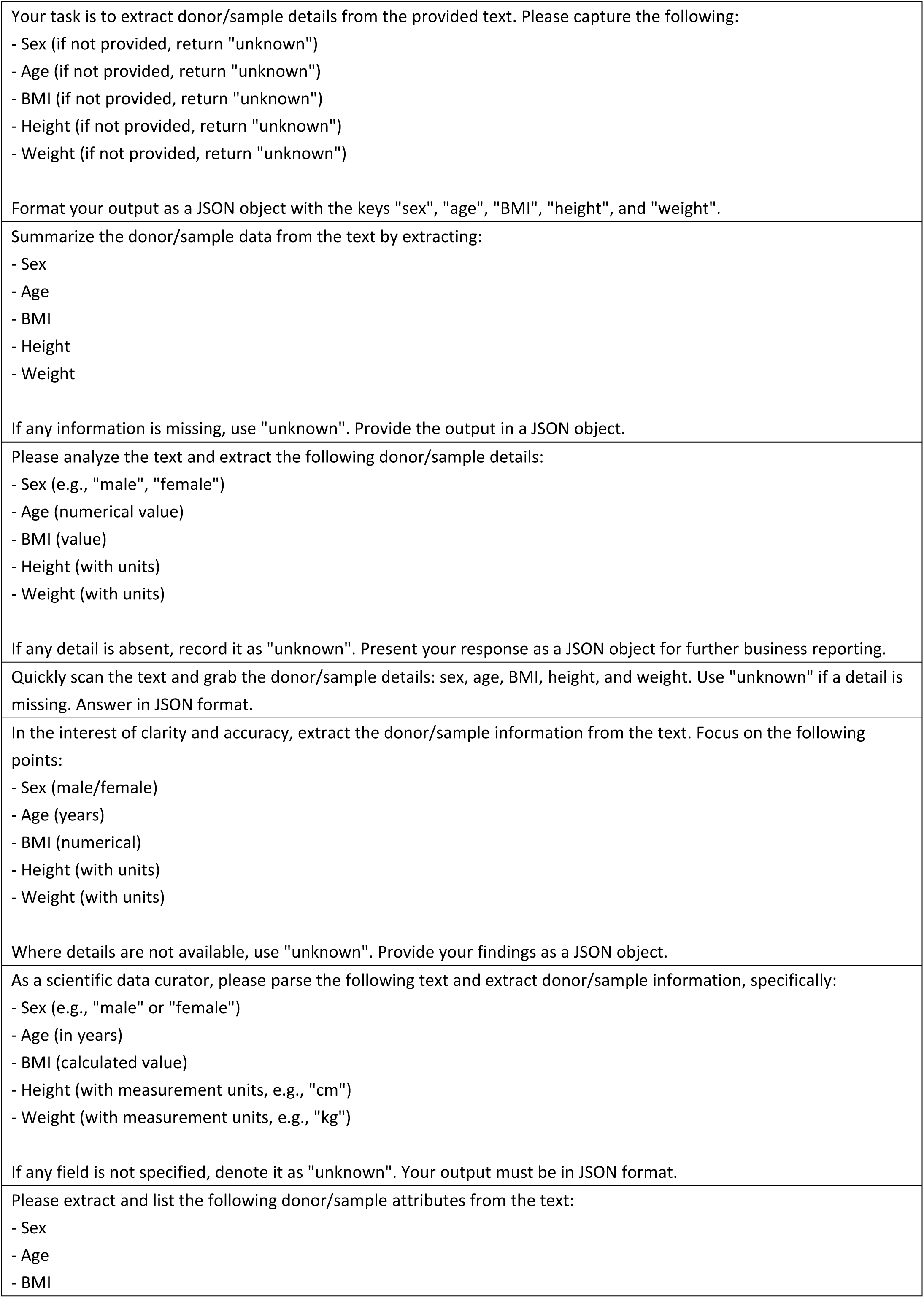
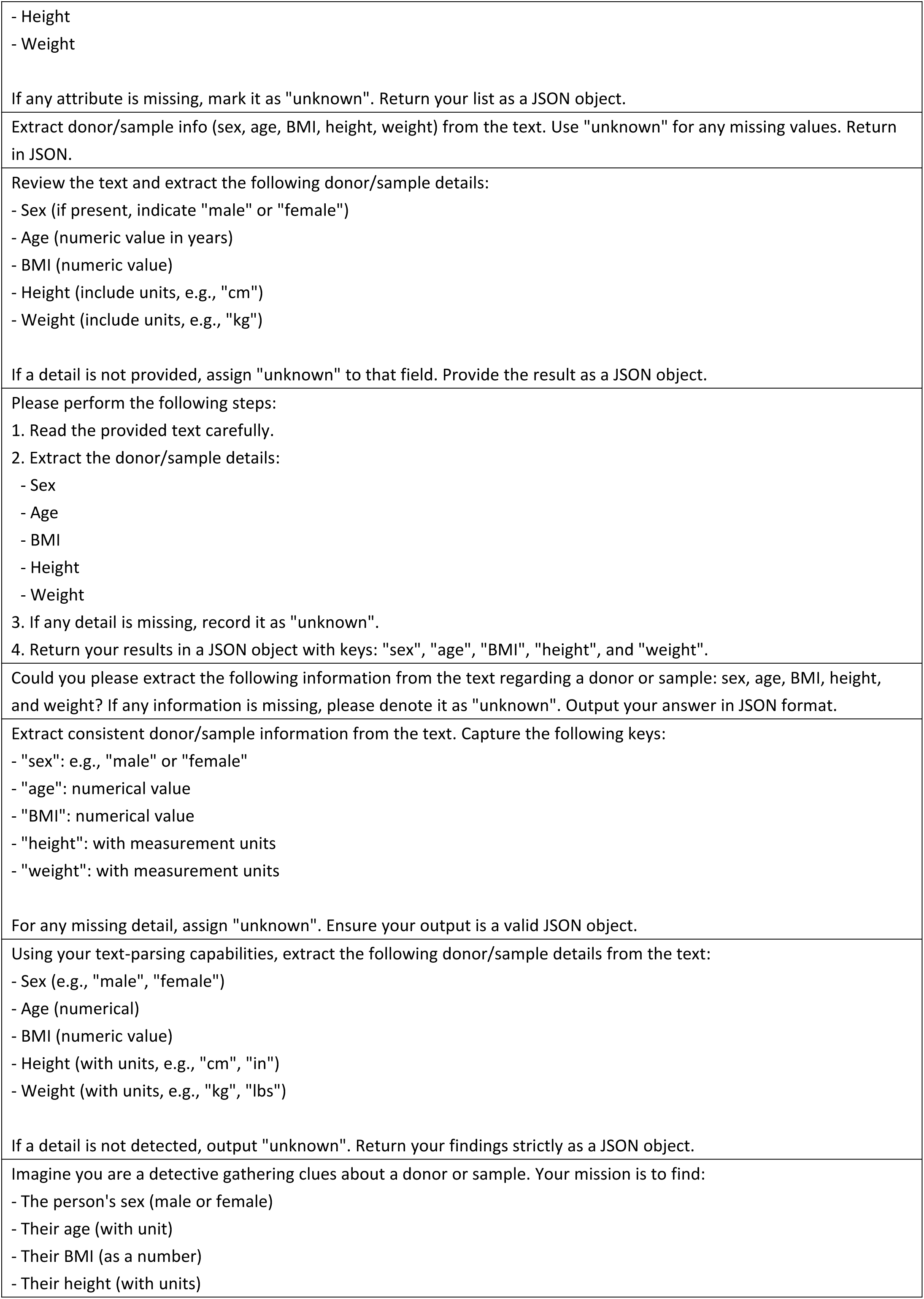
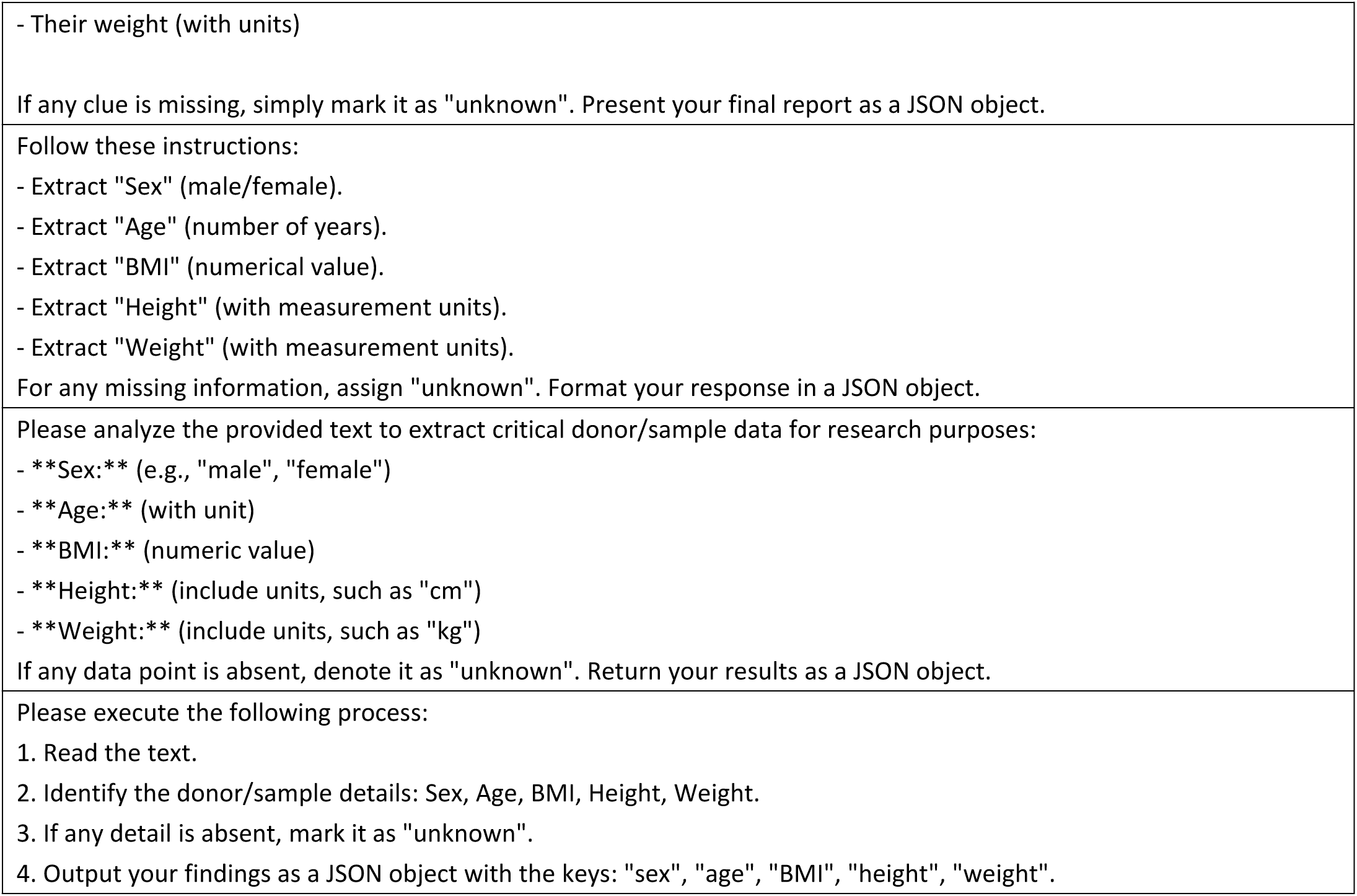
Prompts for donor-metadata extraction tests.

**Supplementary Table 22.**
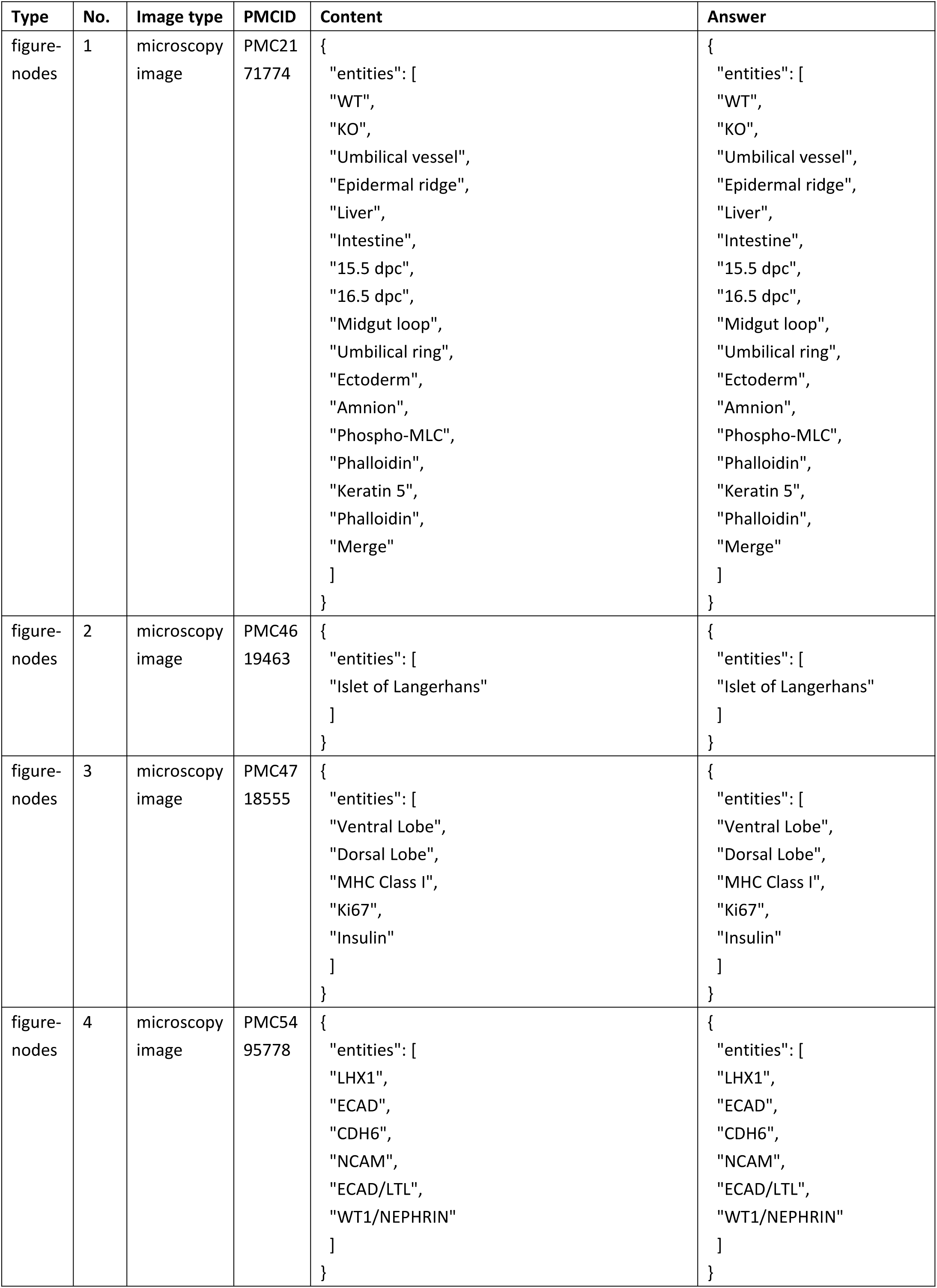
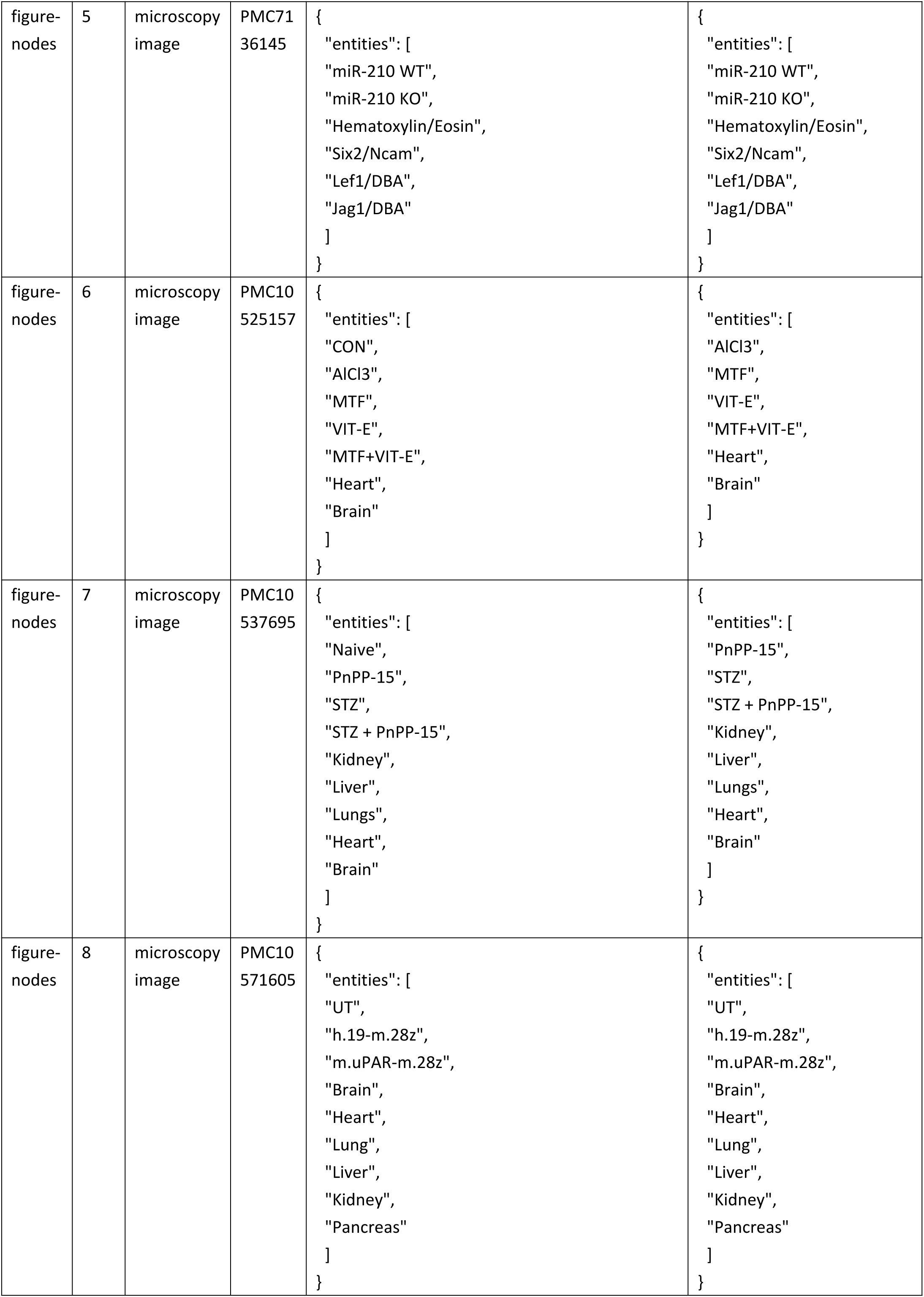
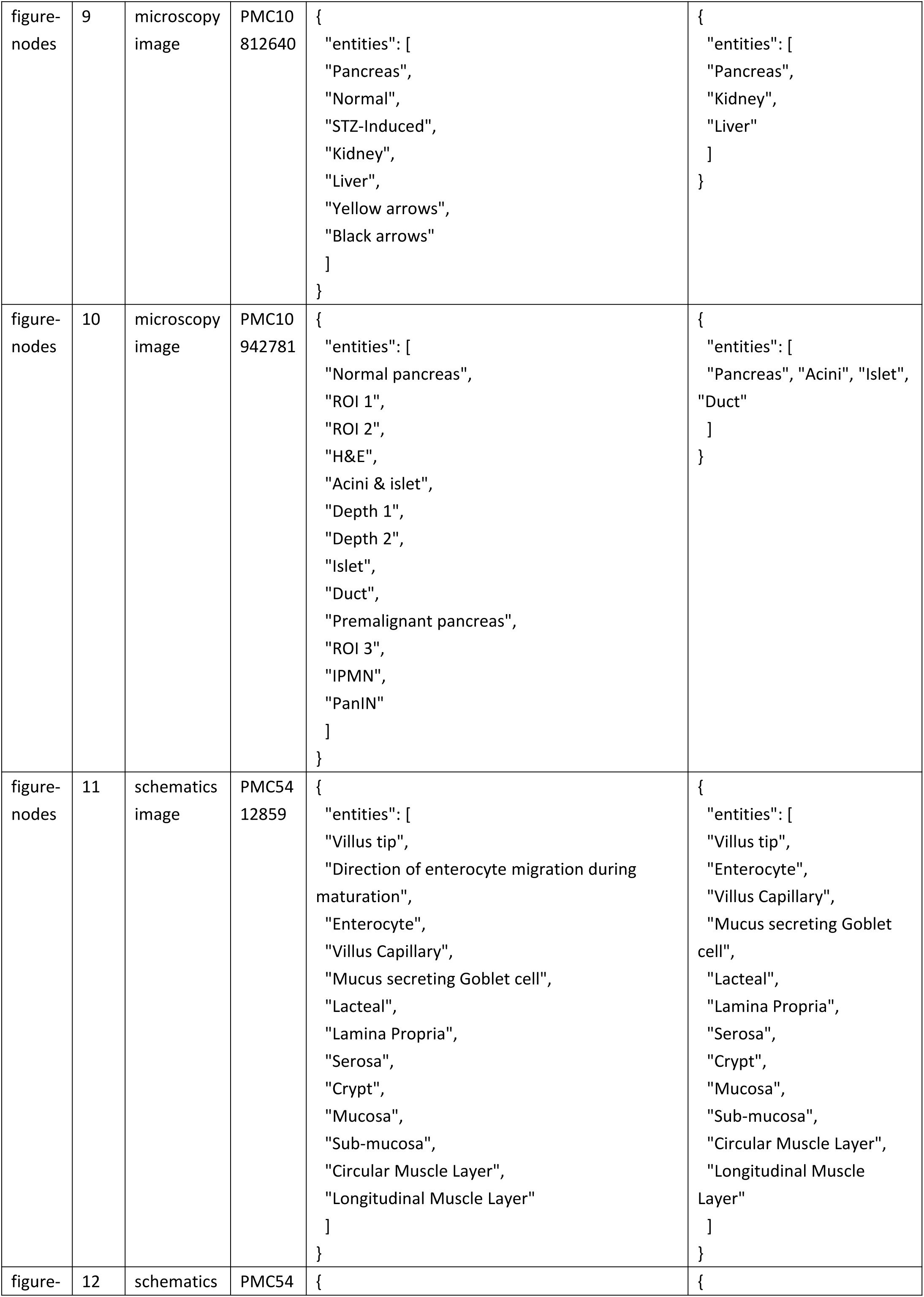
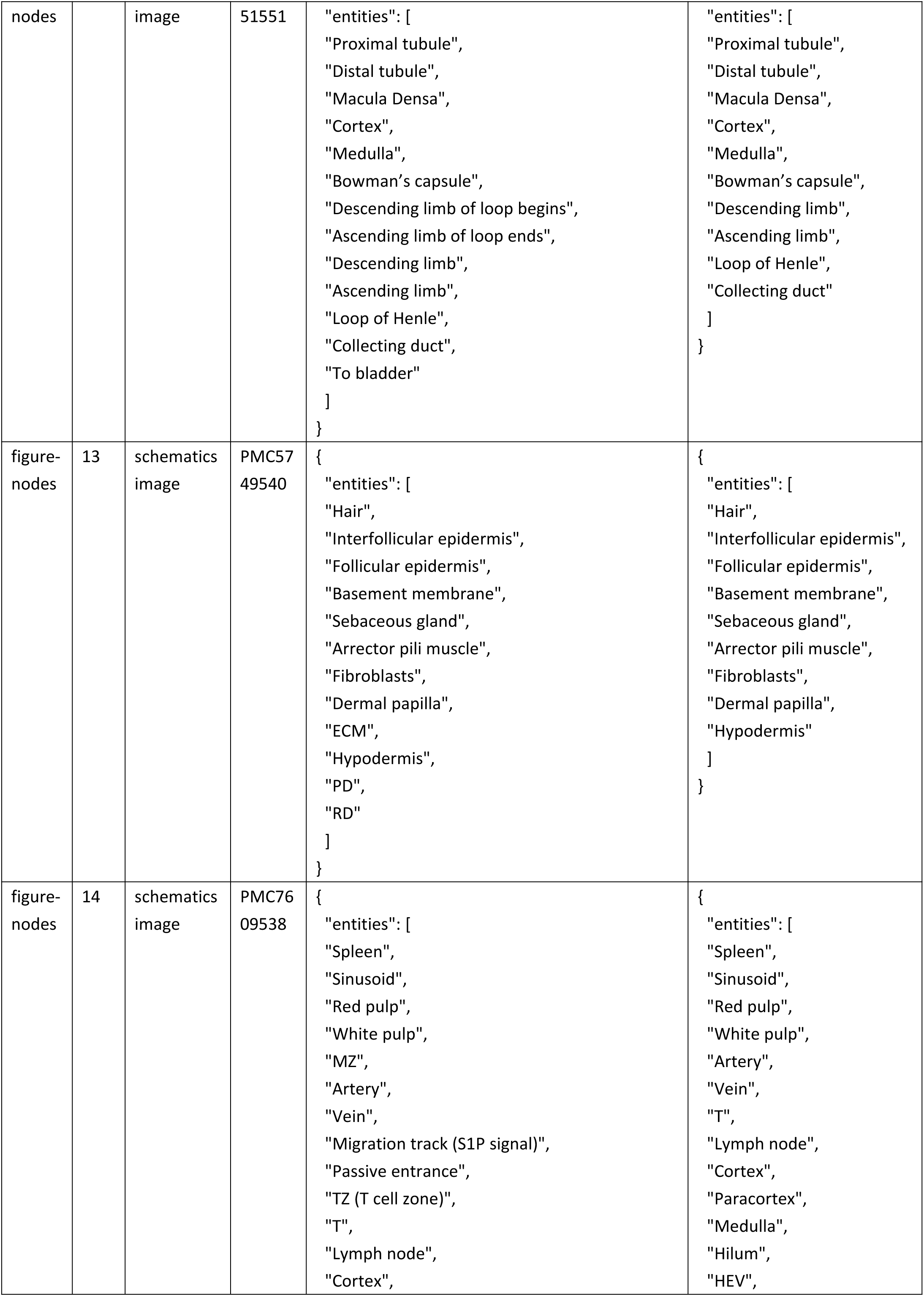
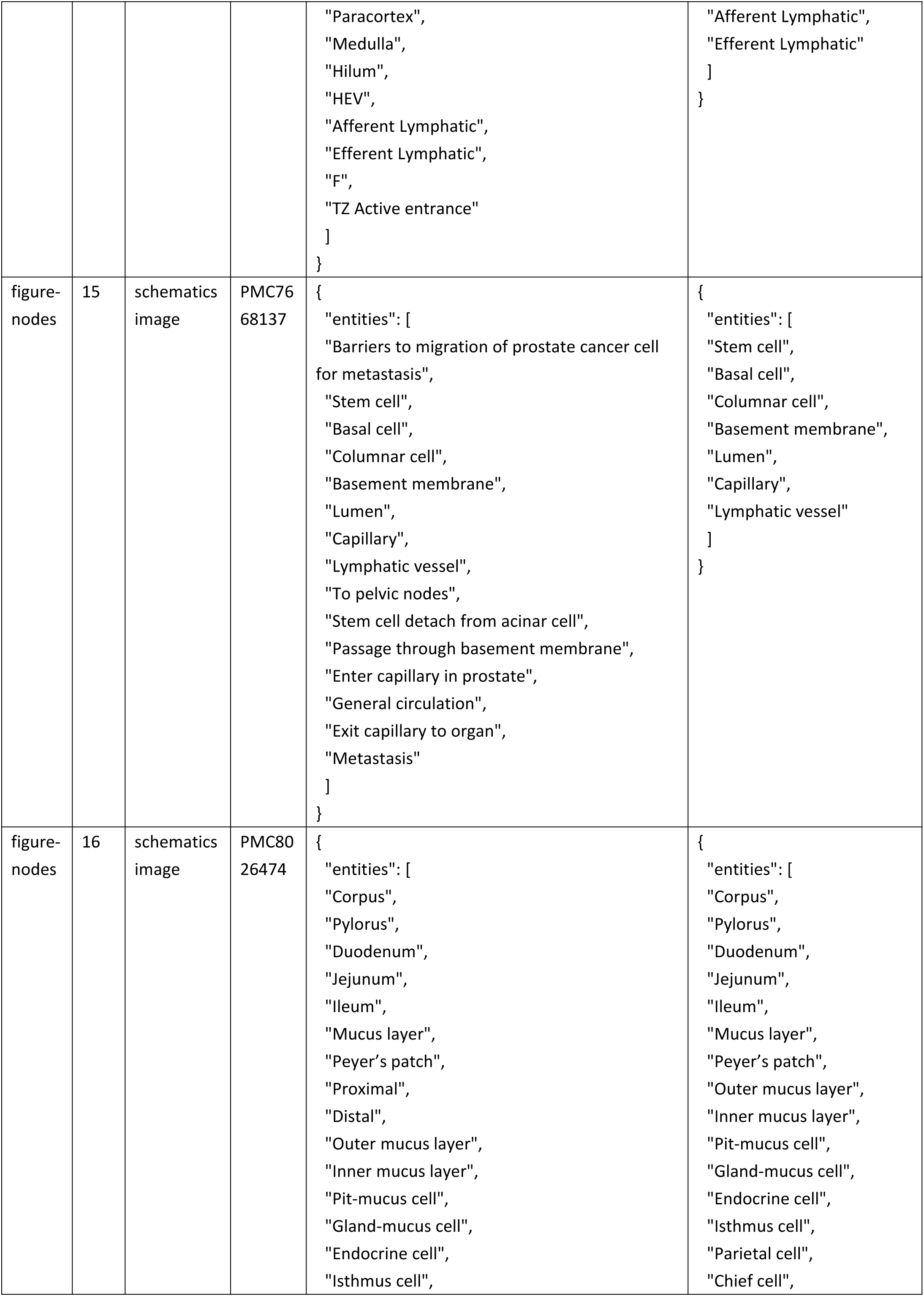
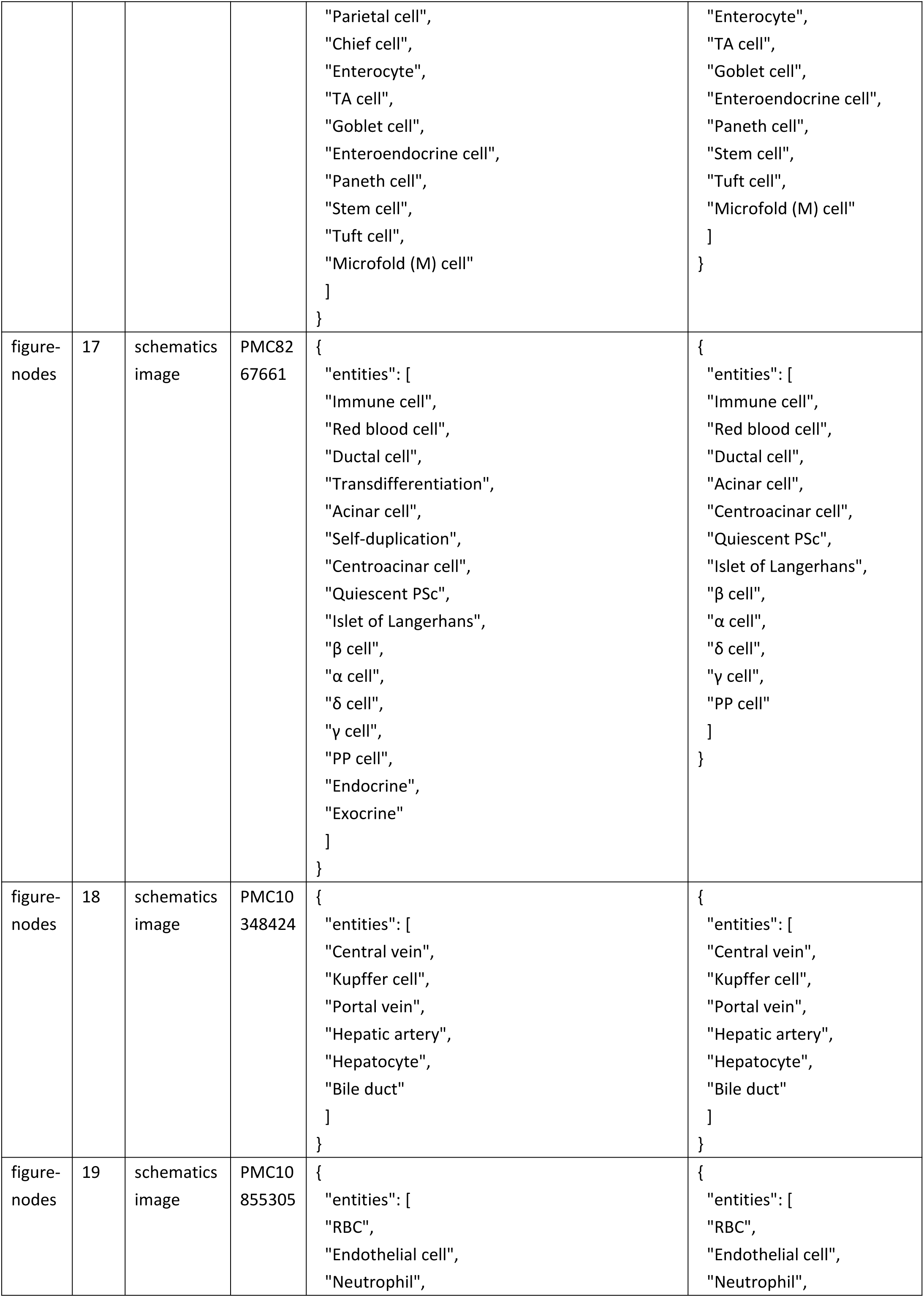
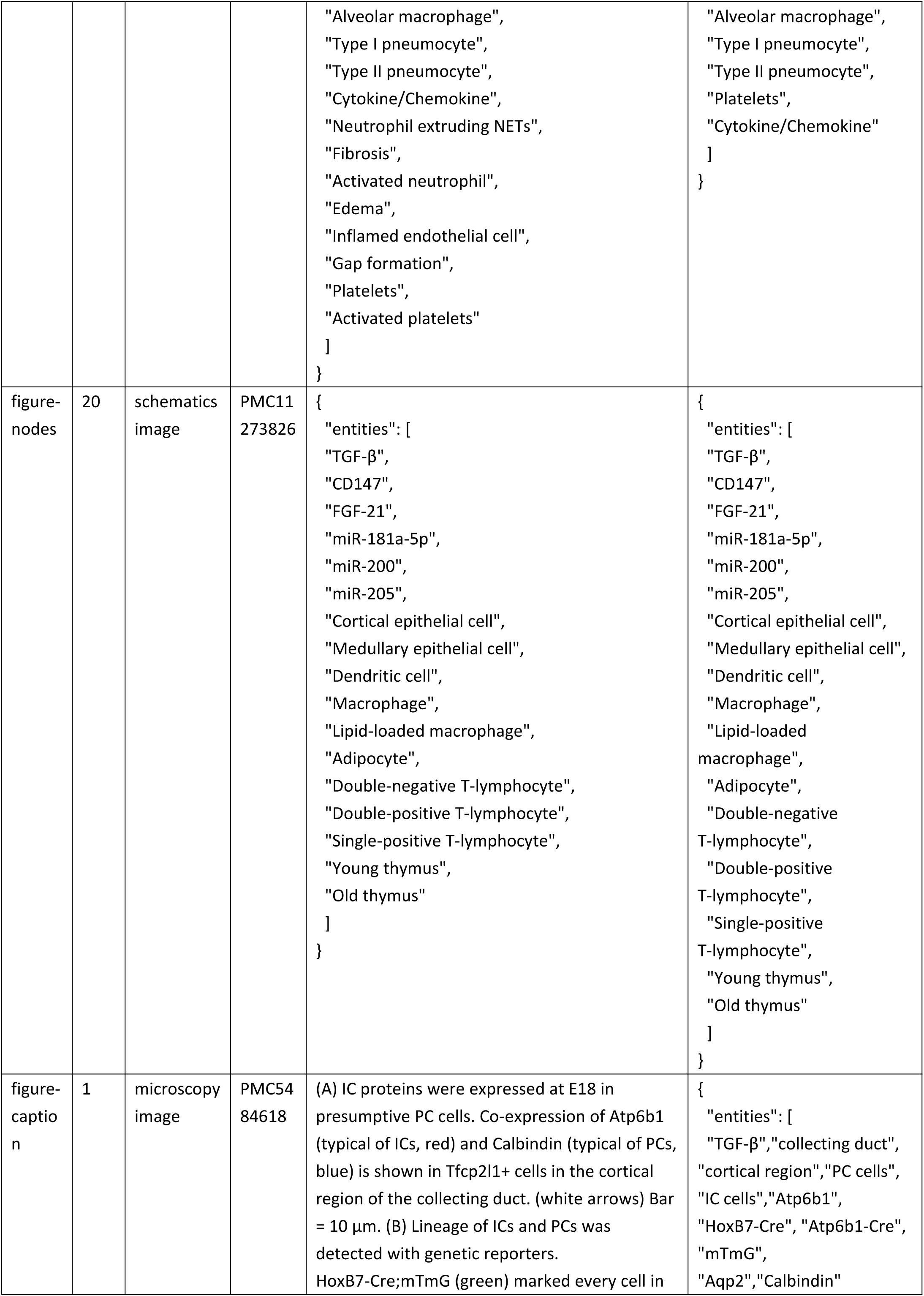
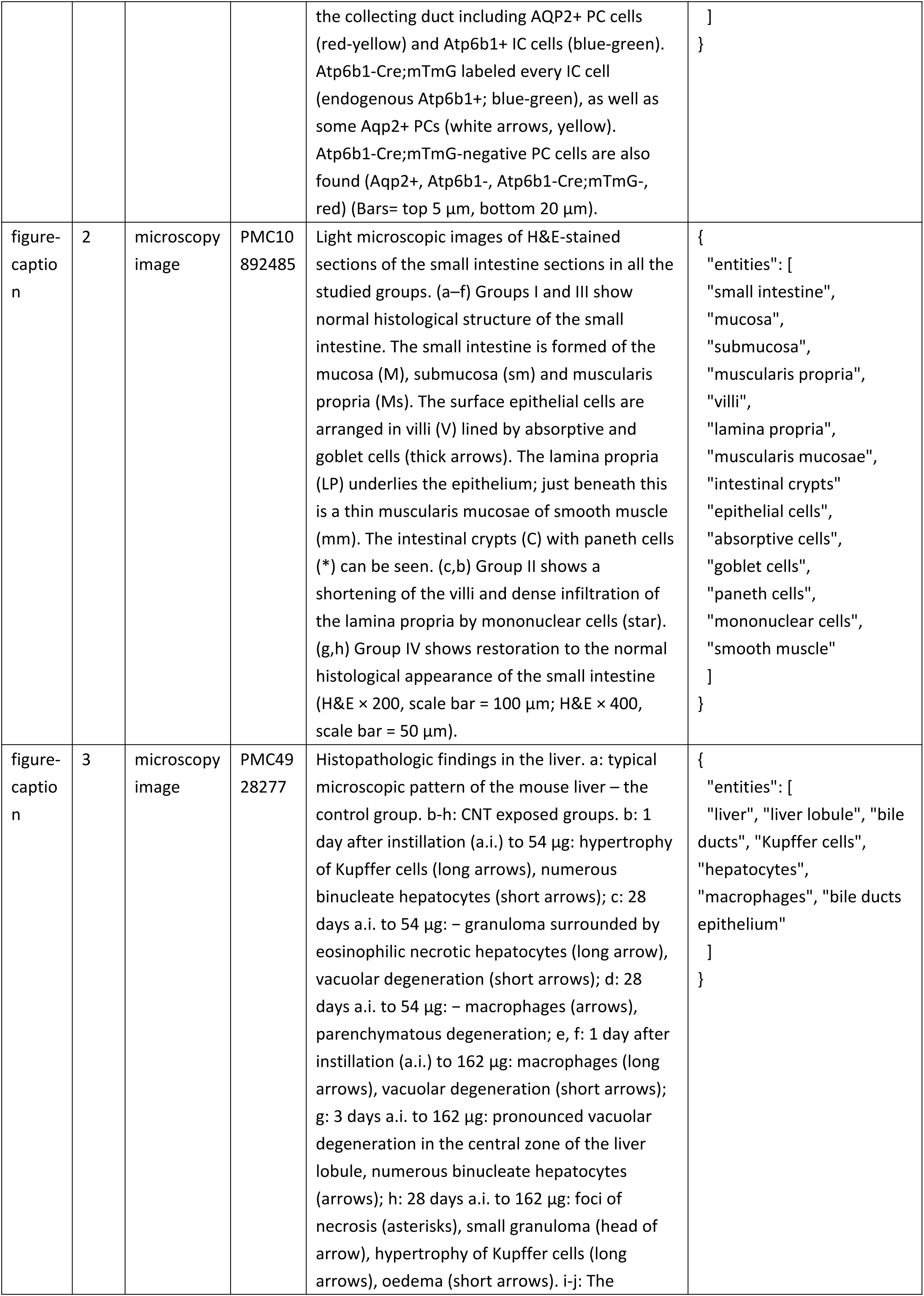
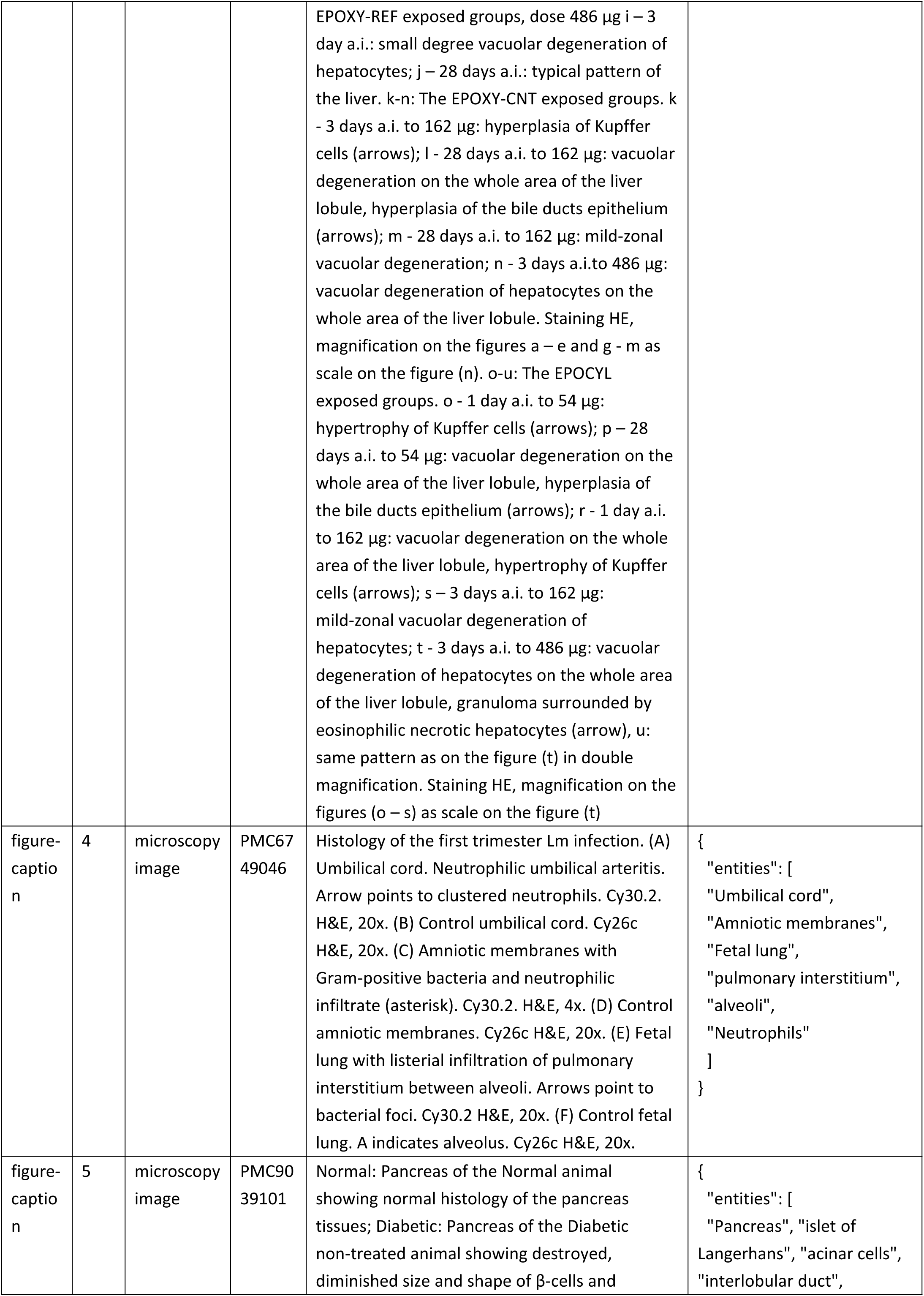
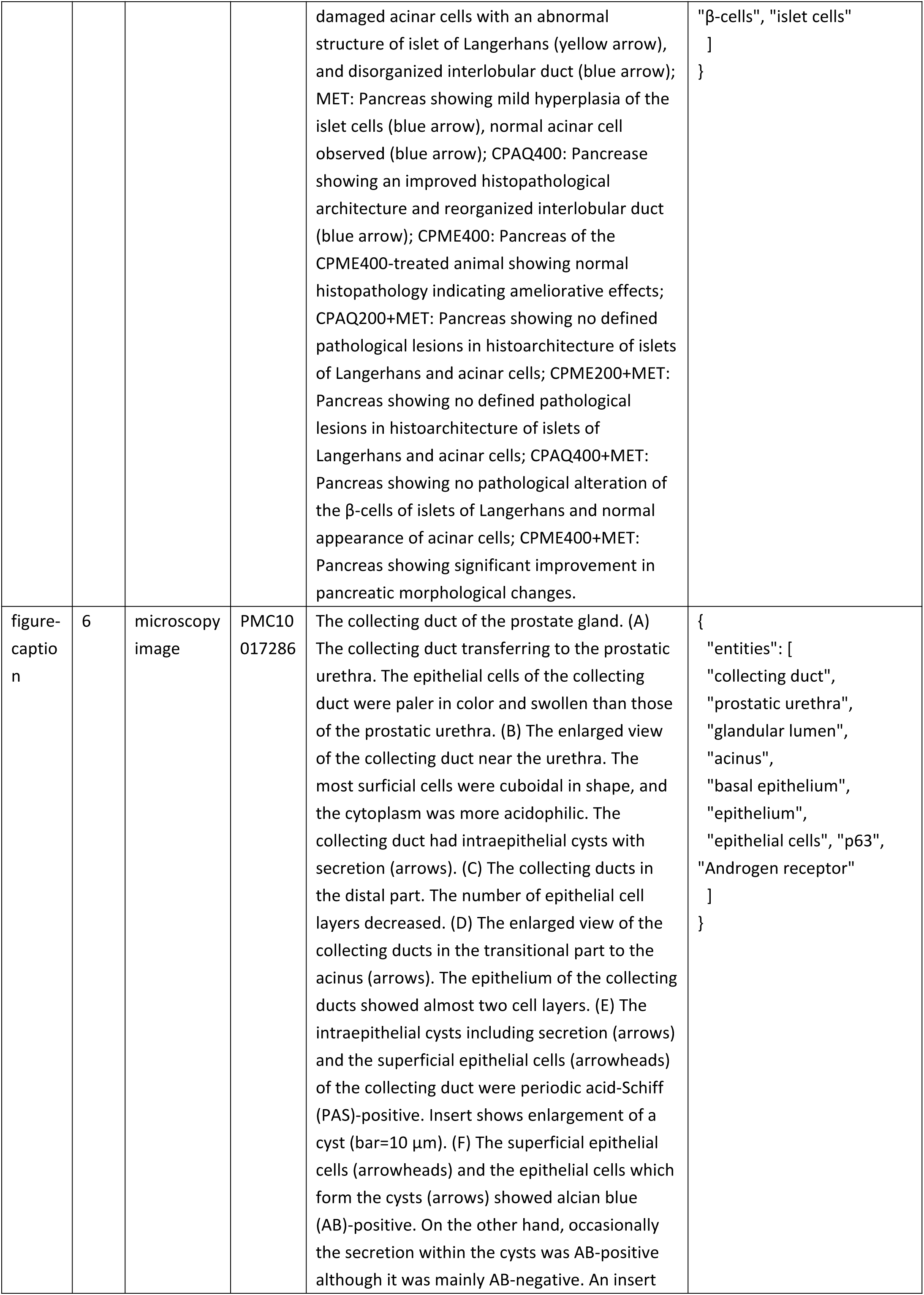
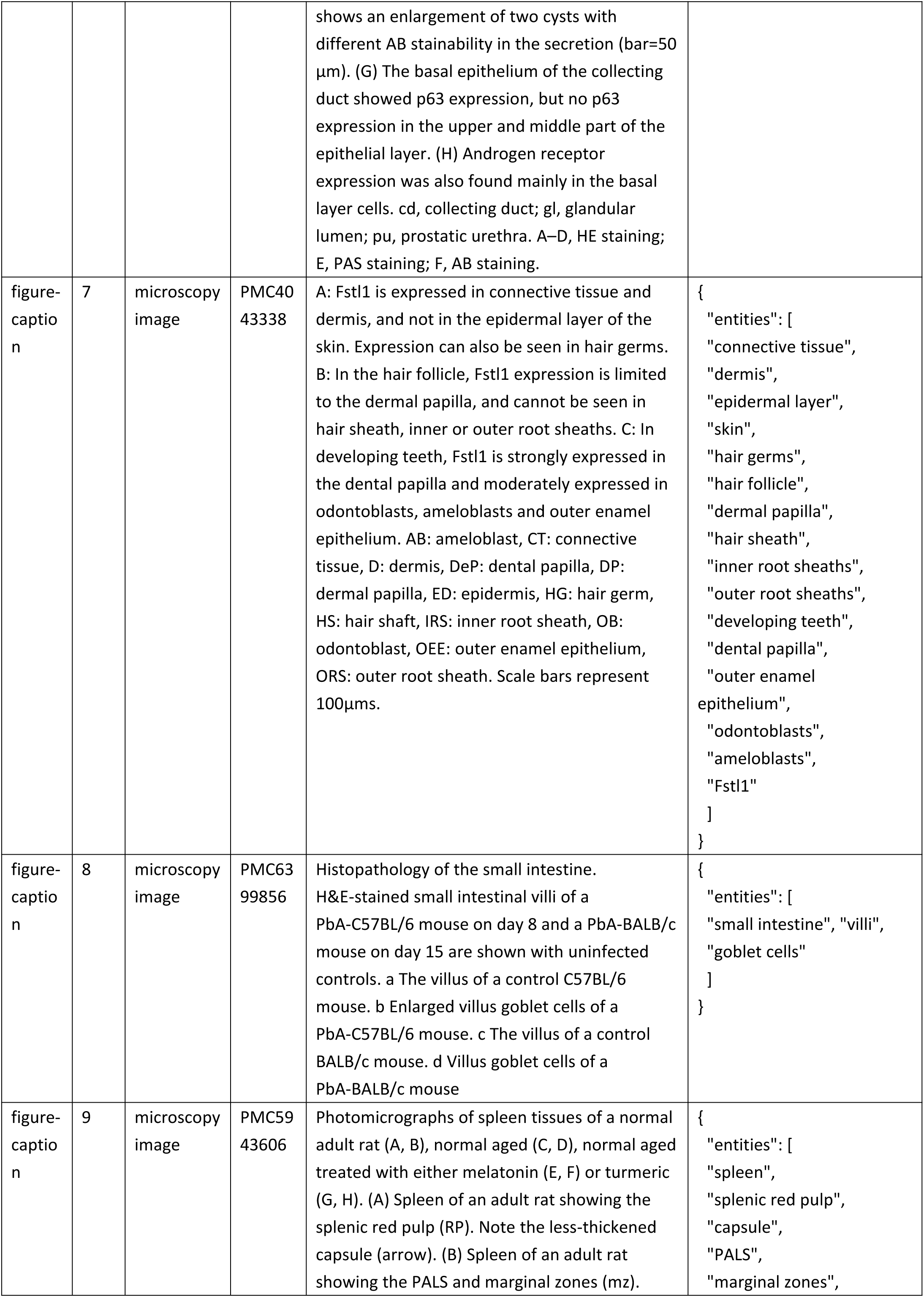
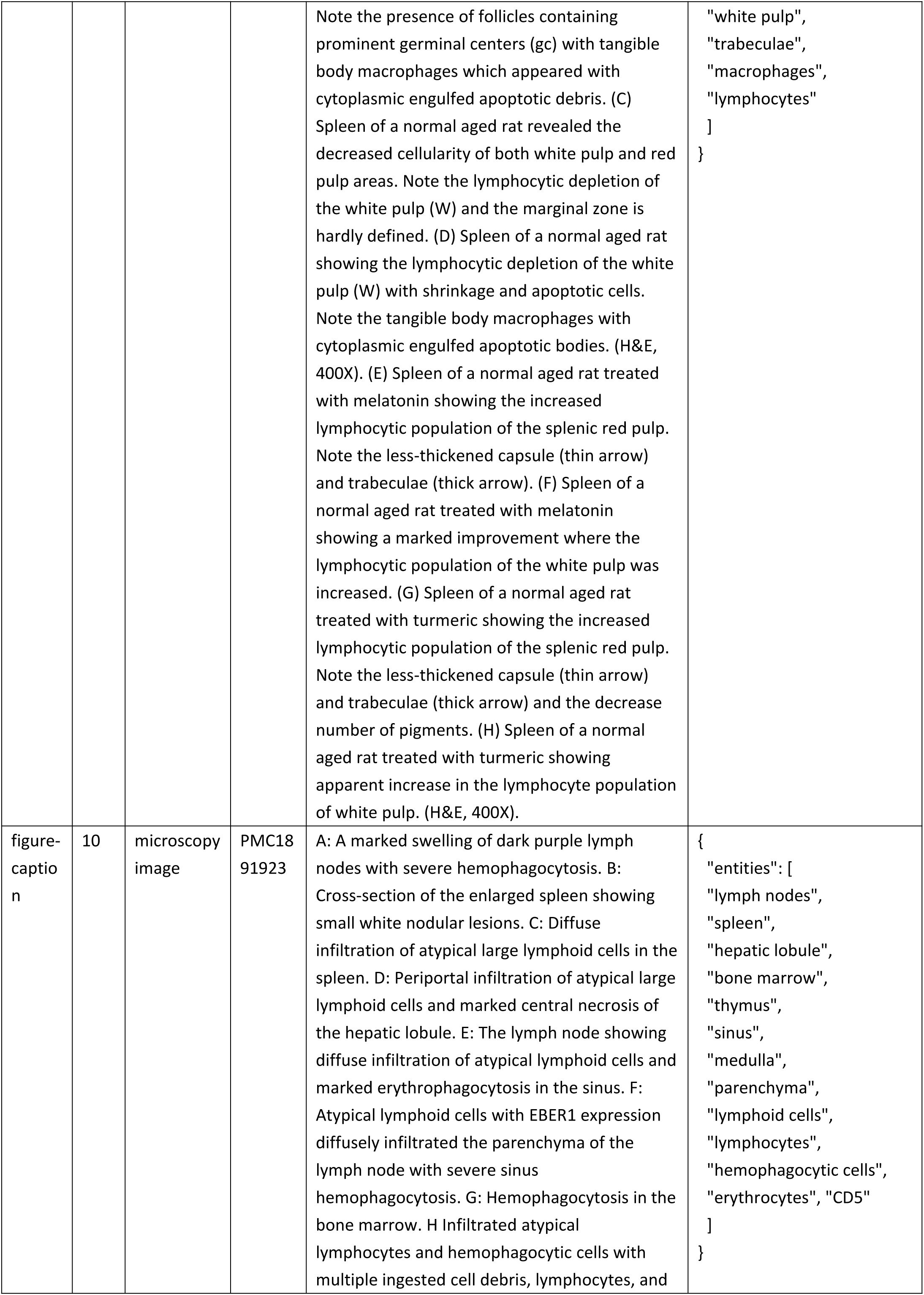
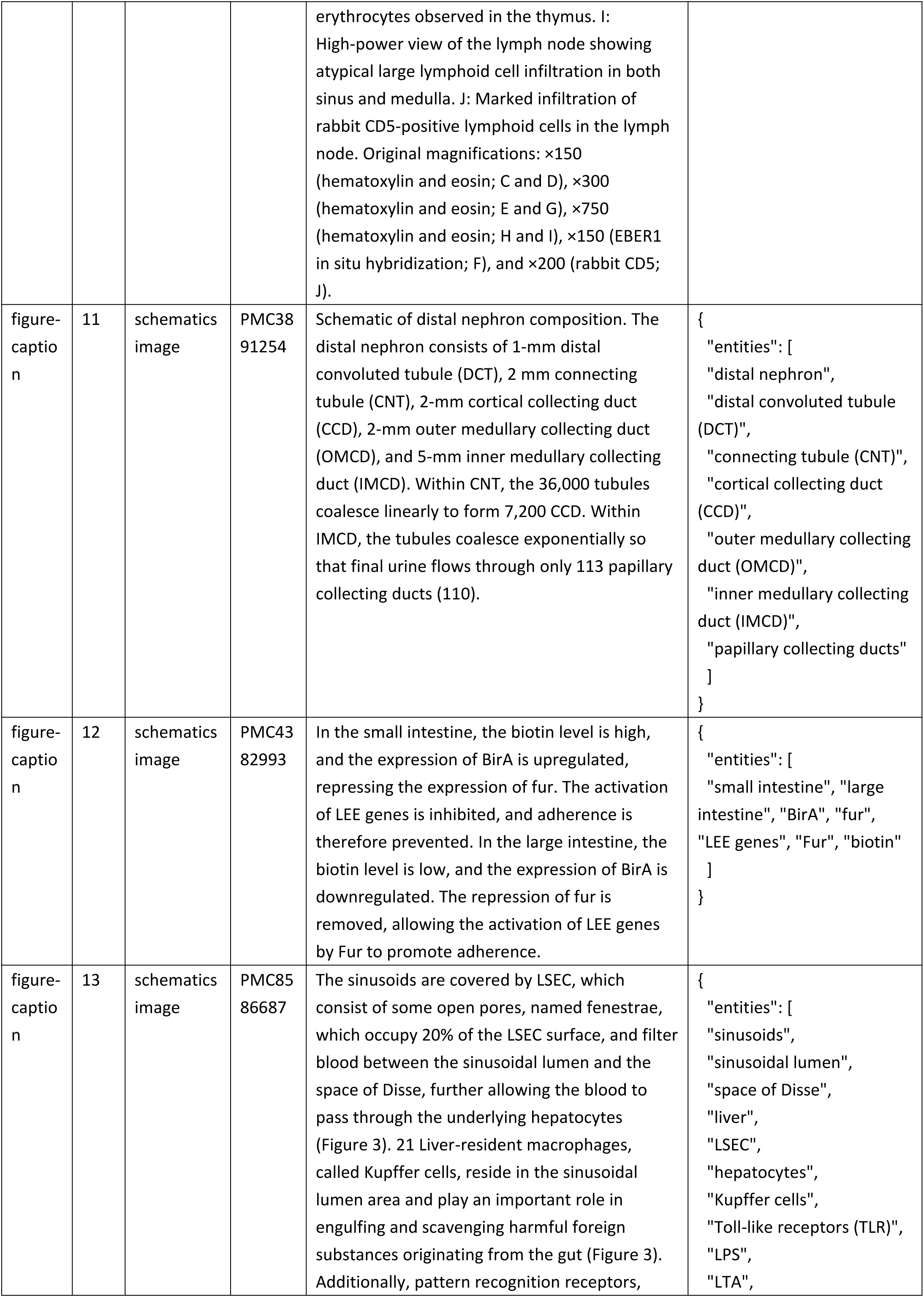
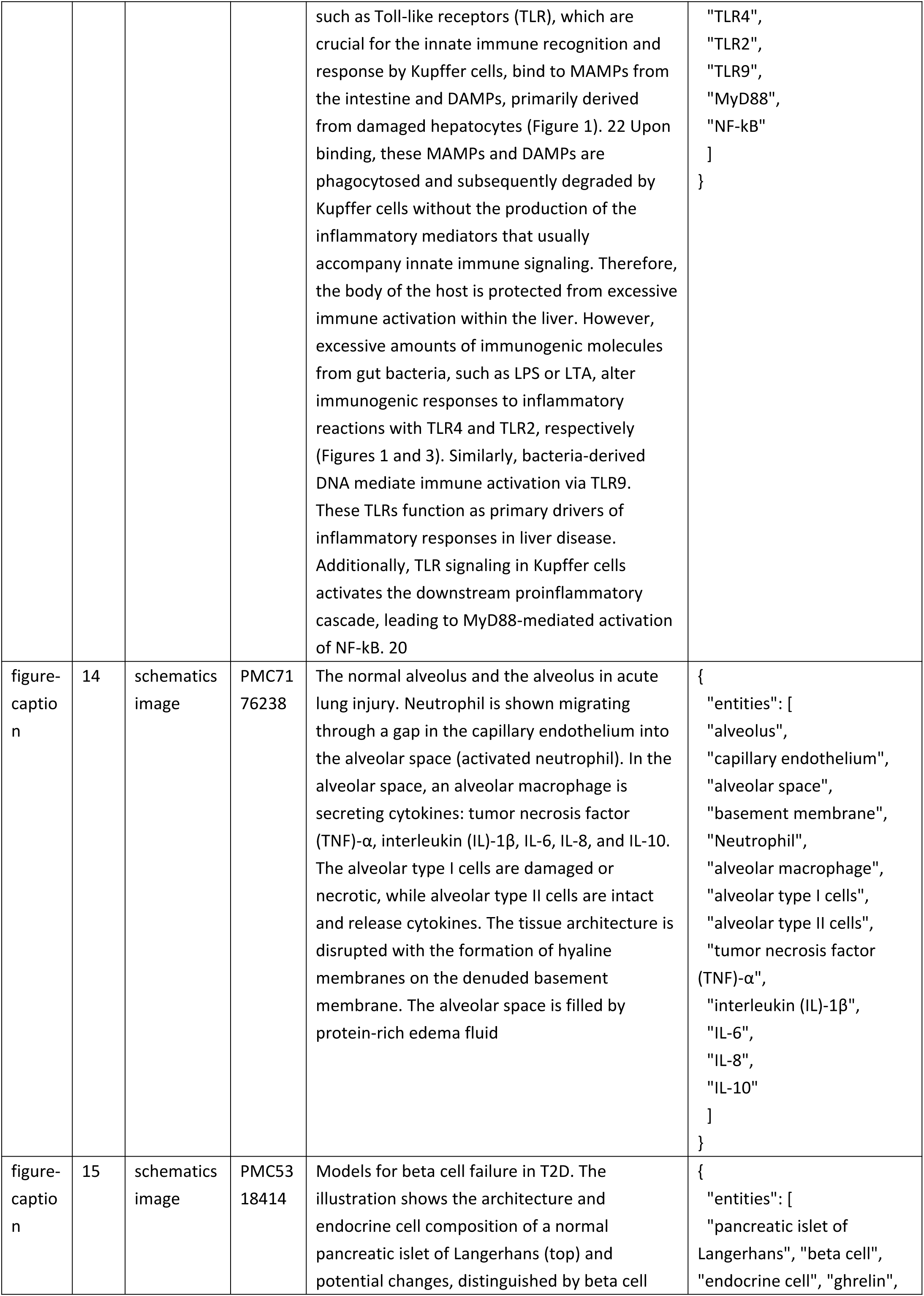
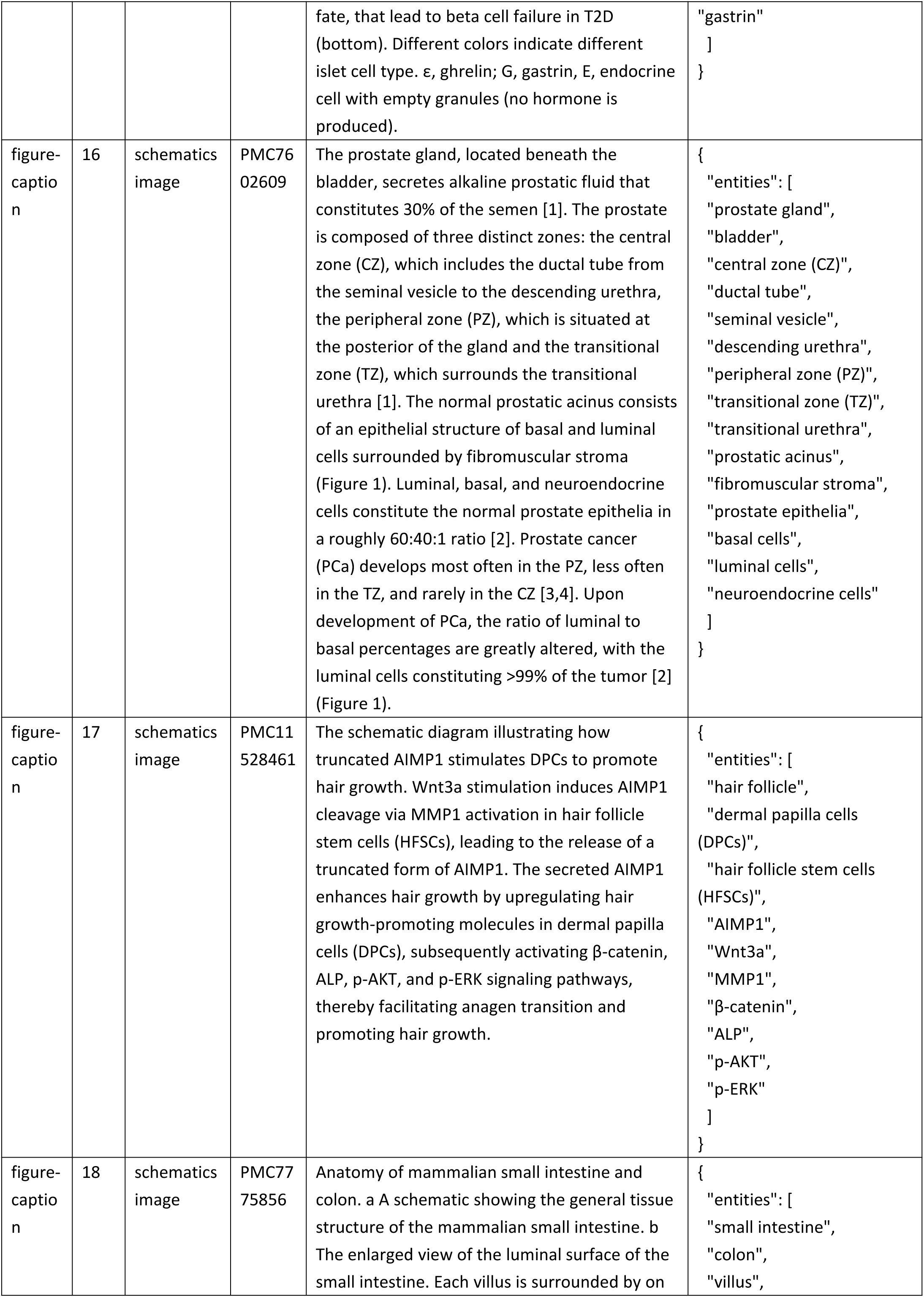
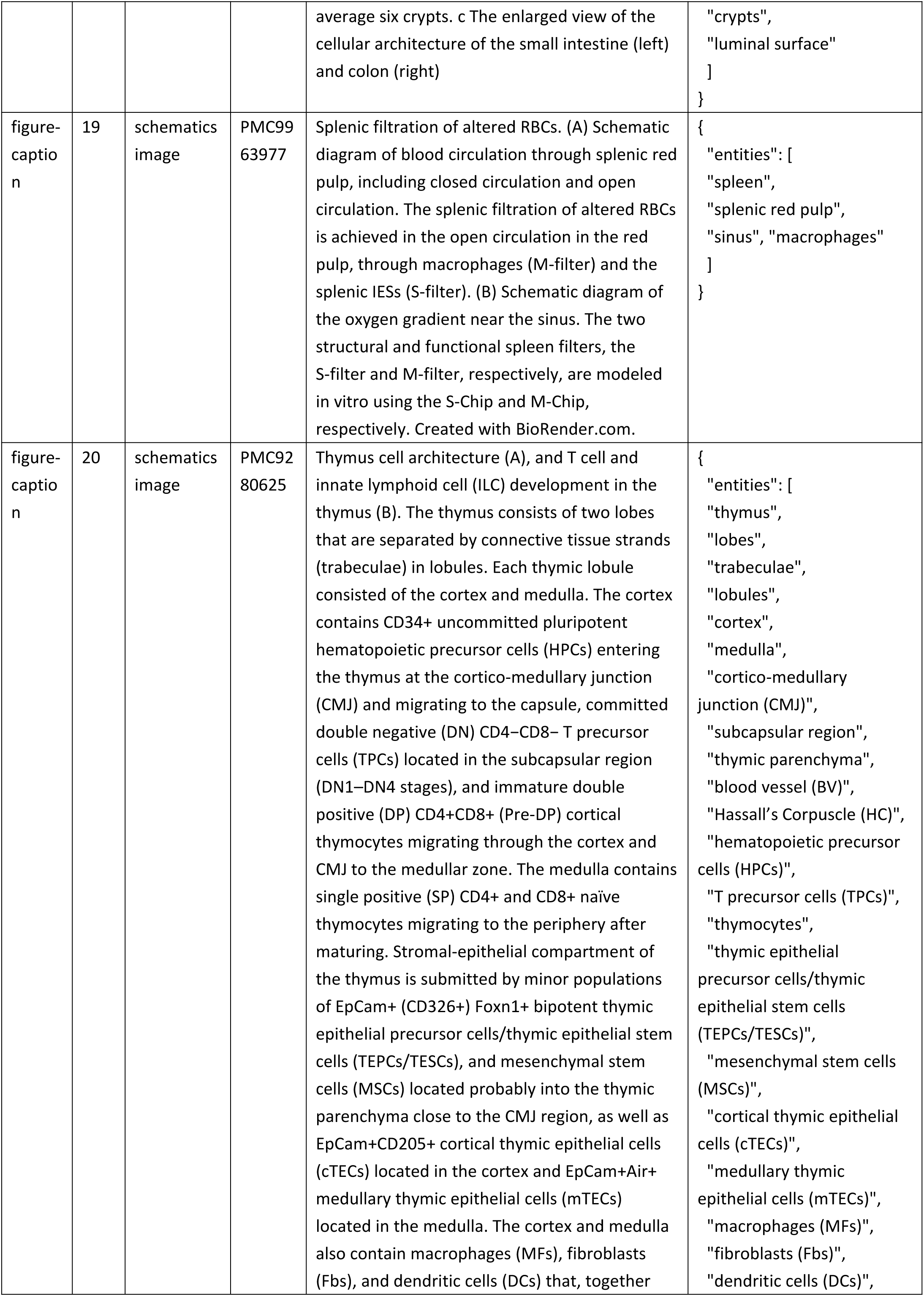
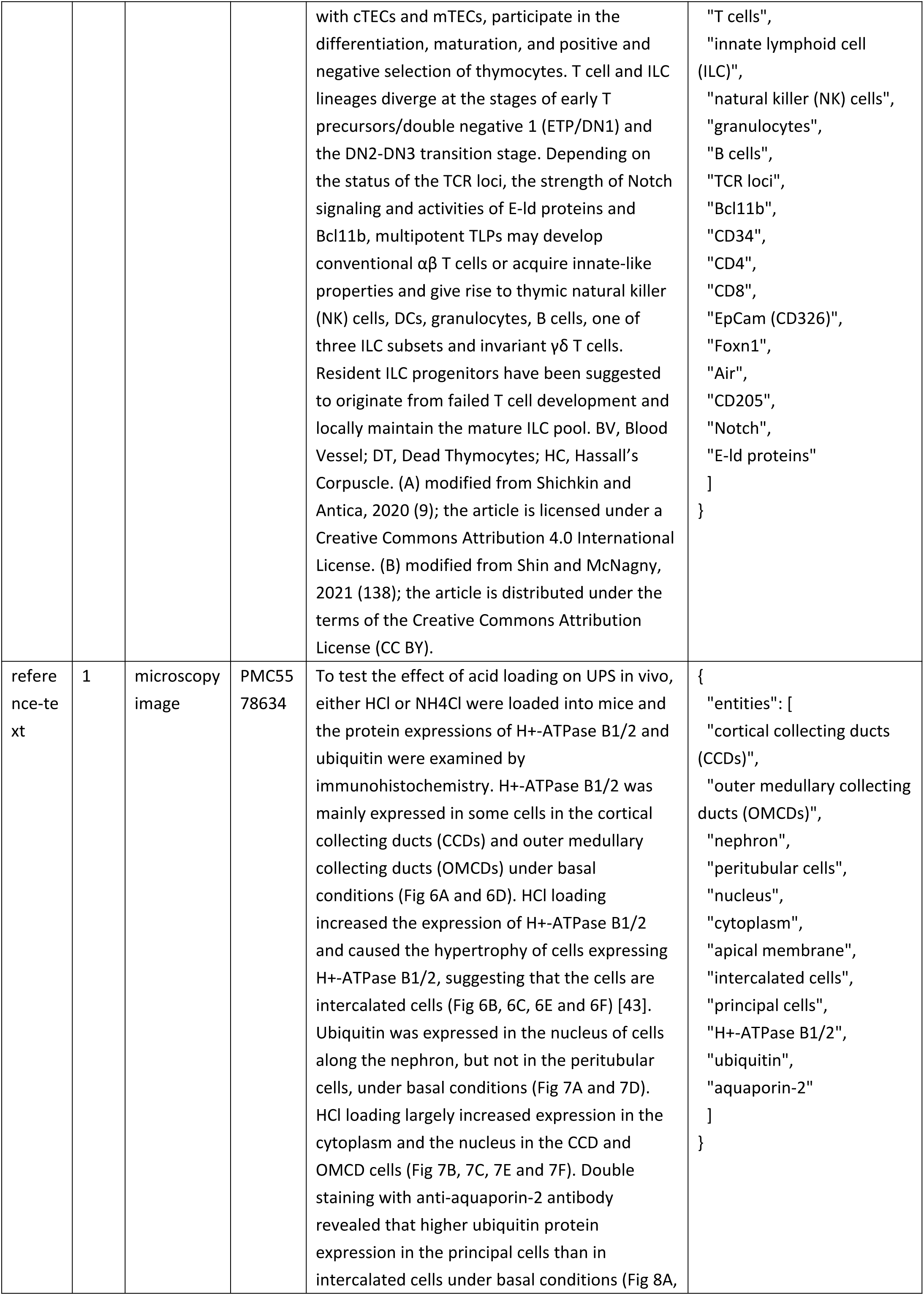
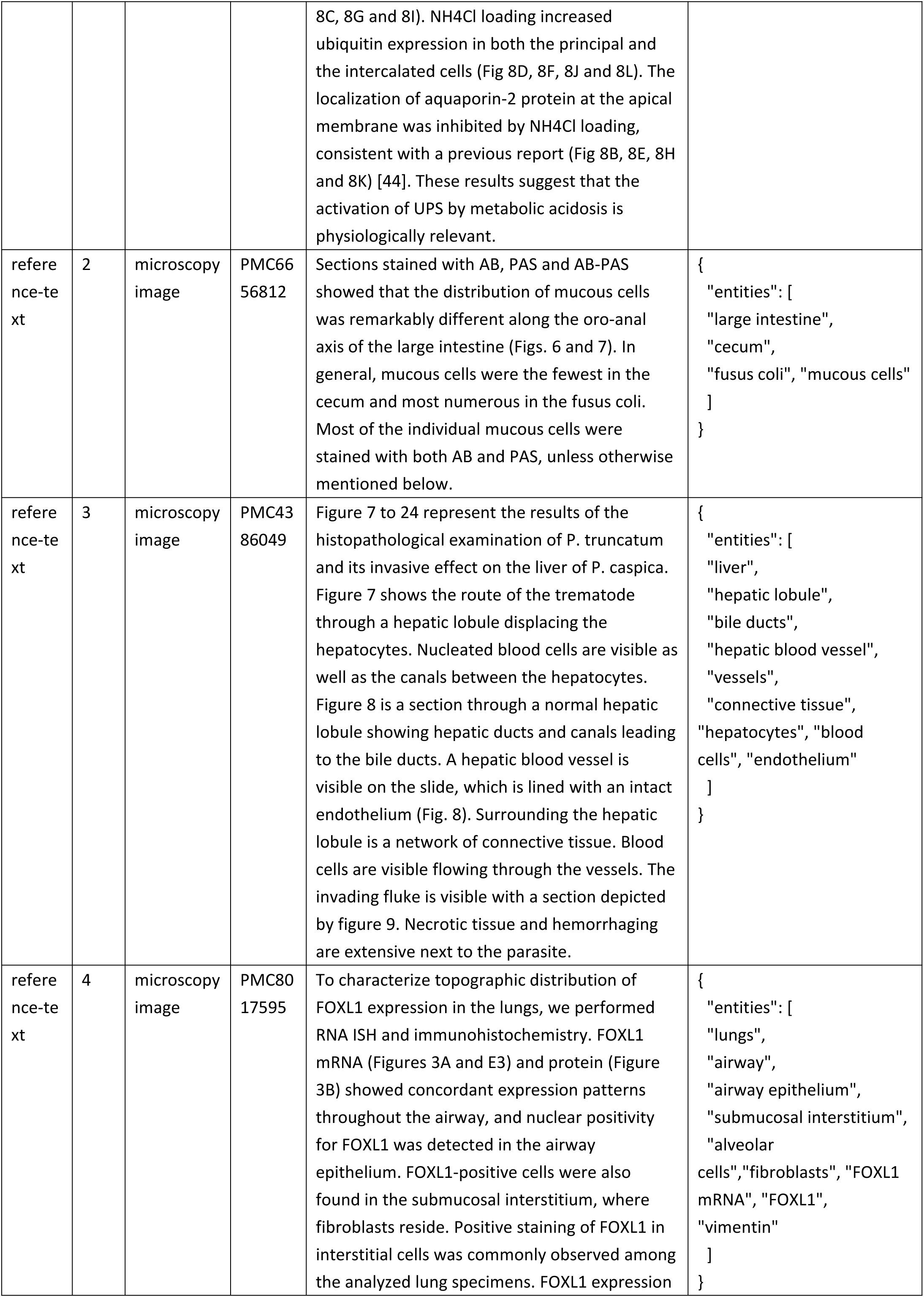
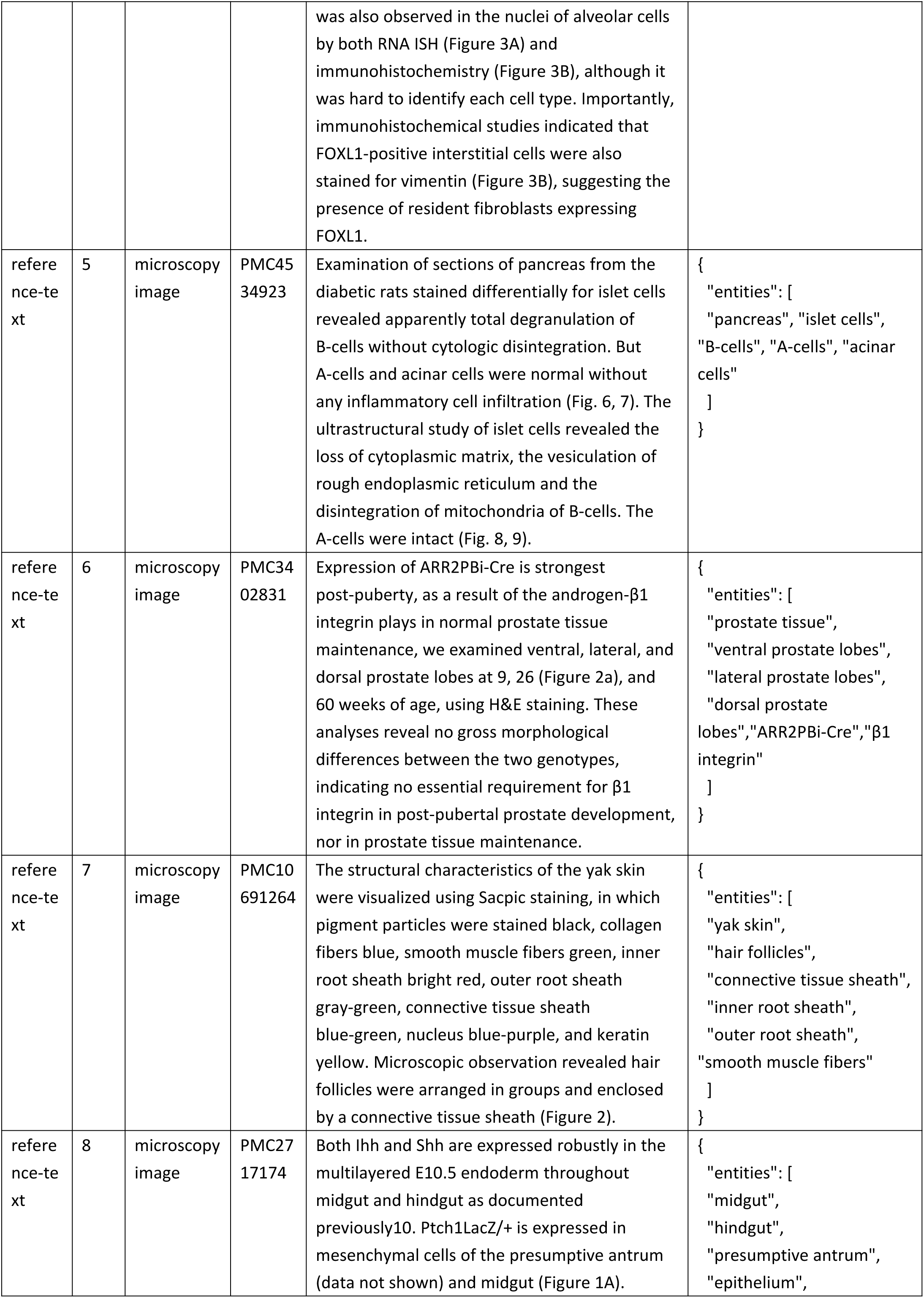
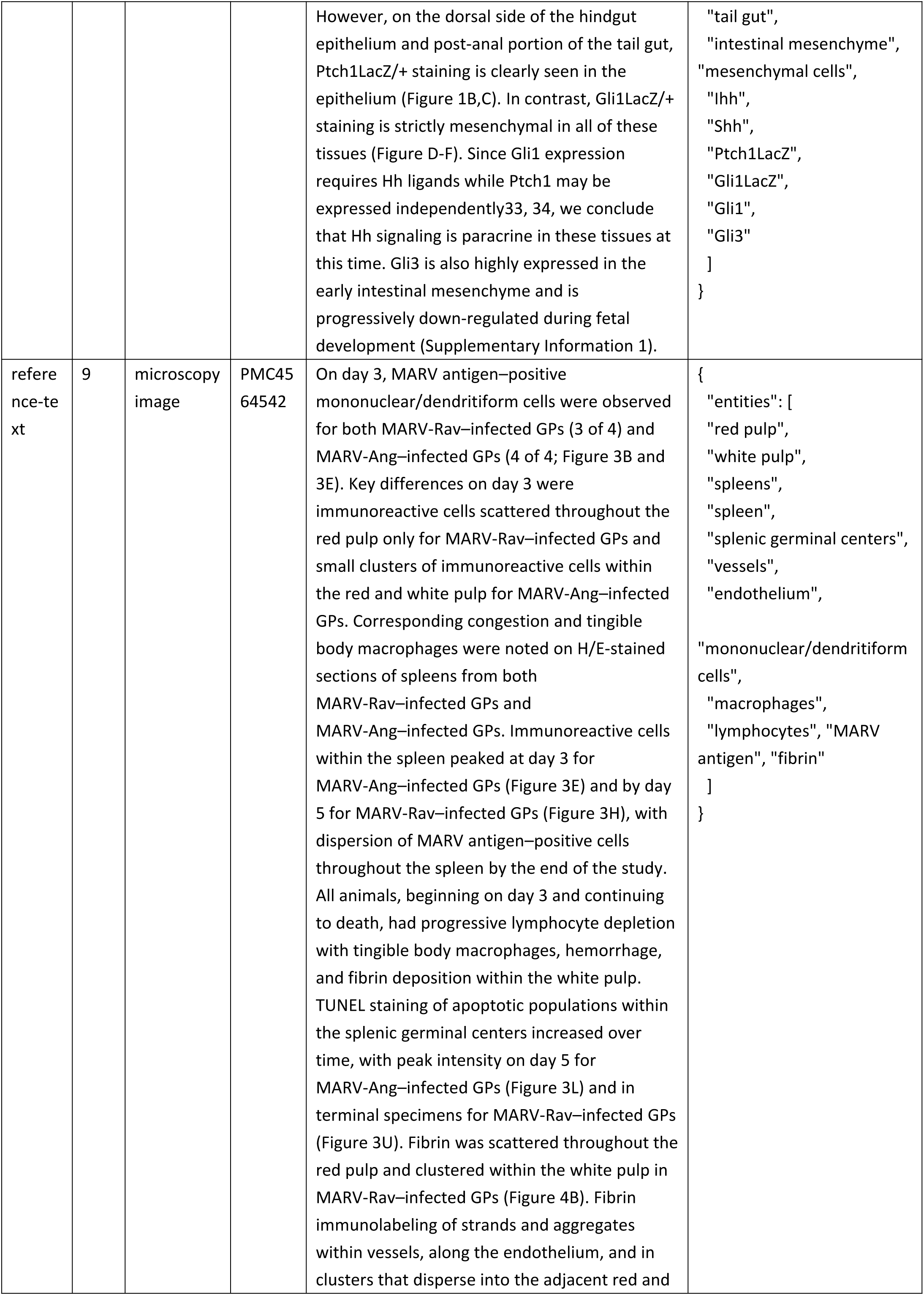
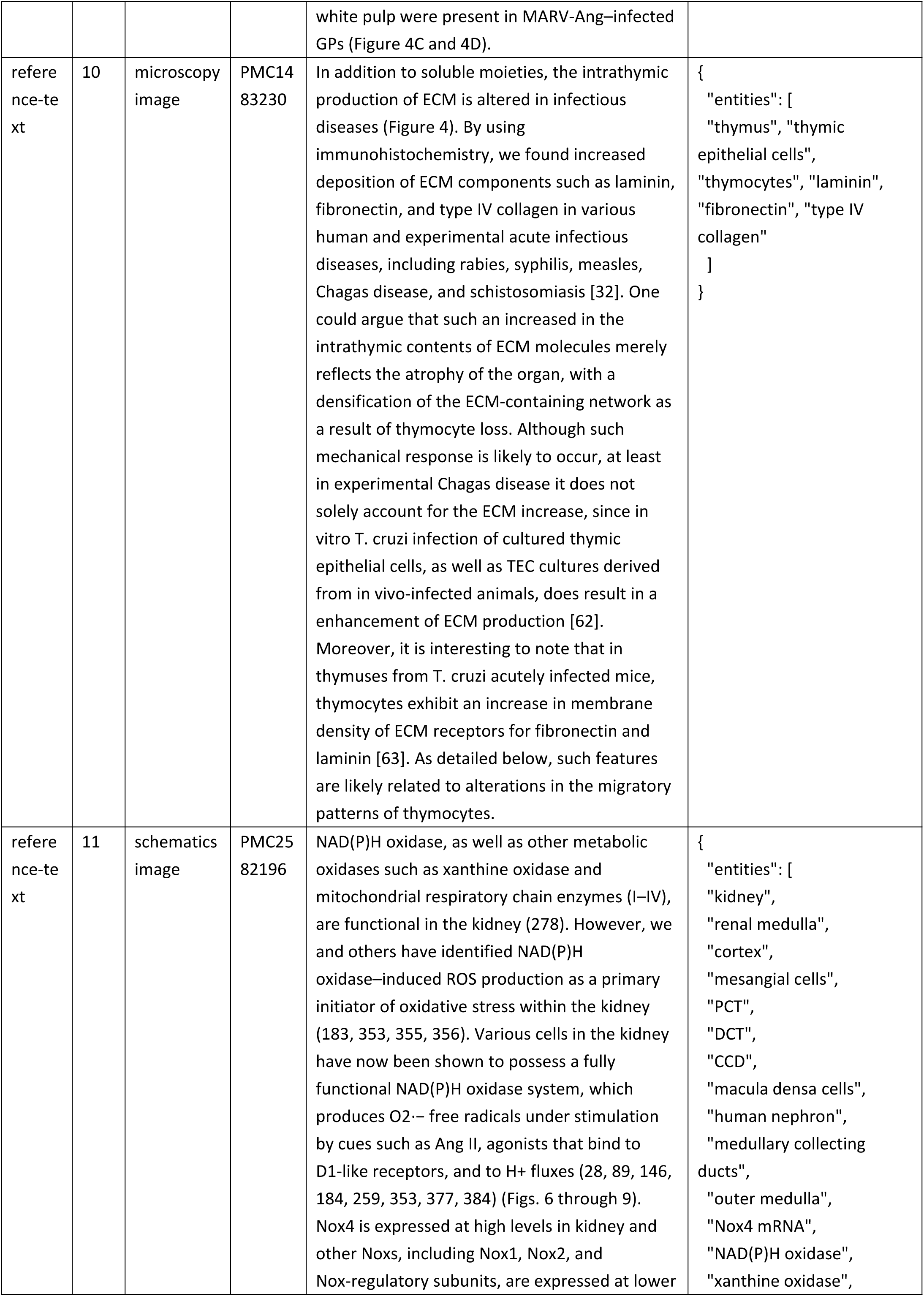
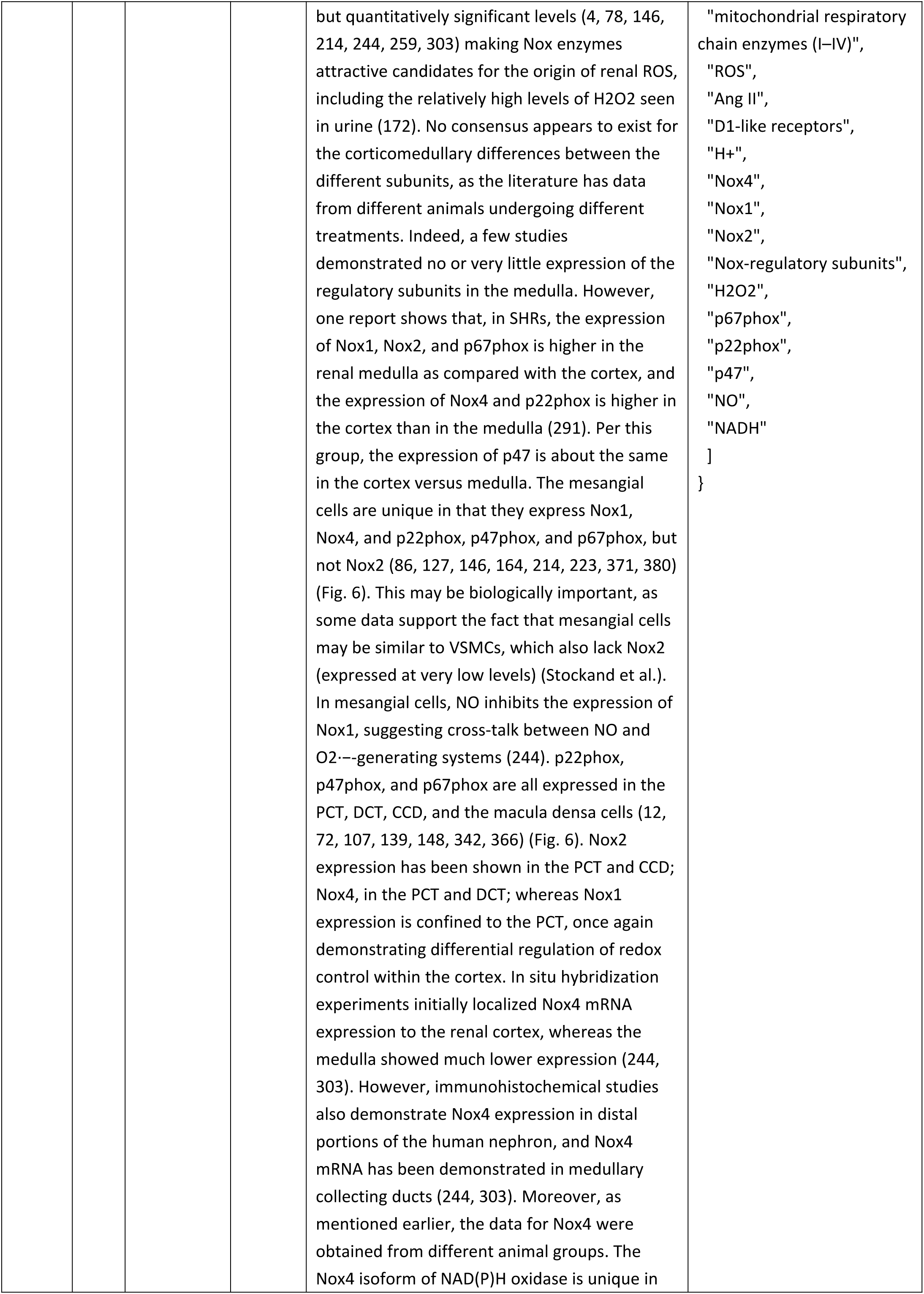
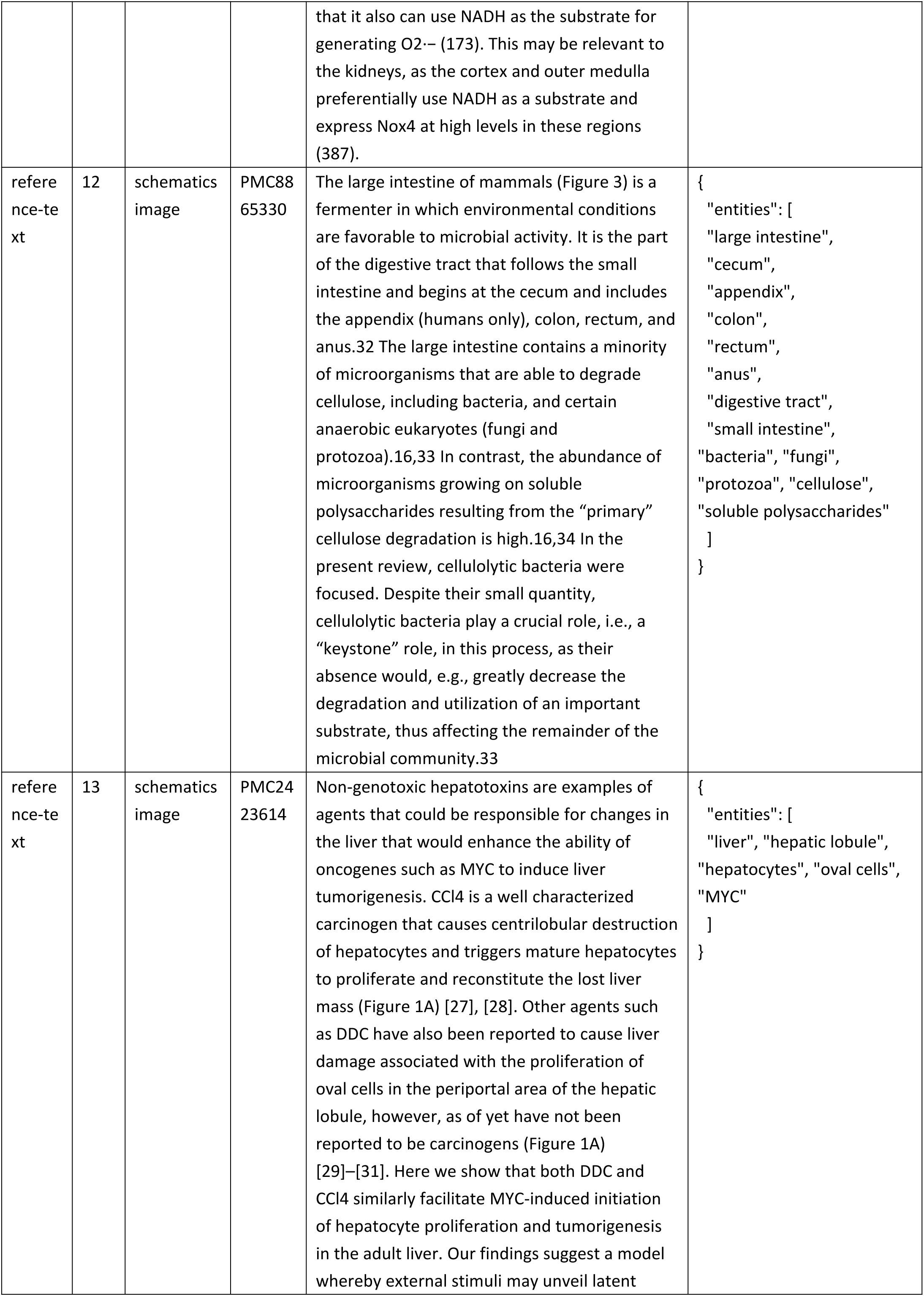
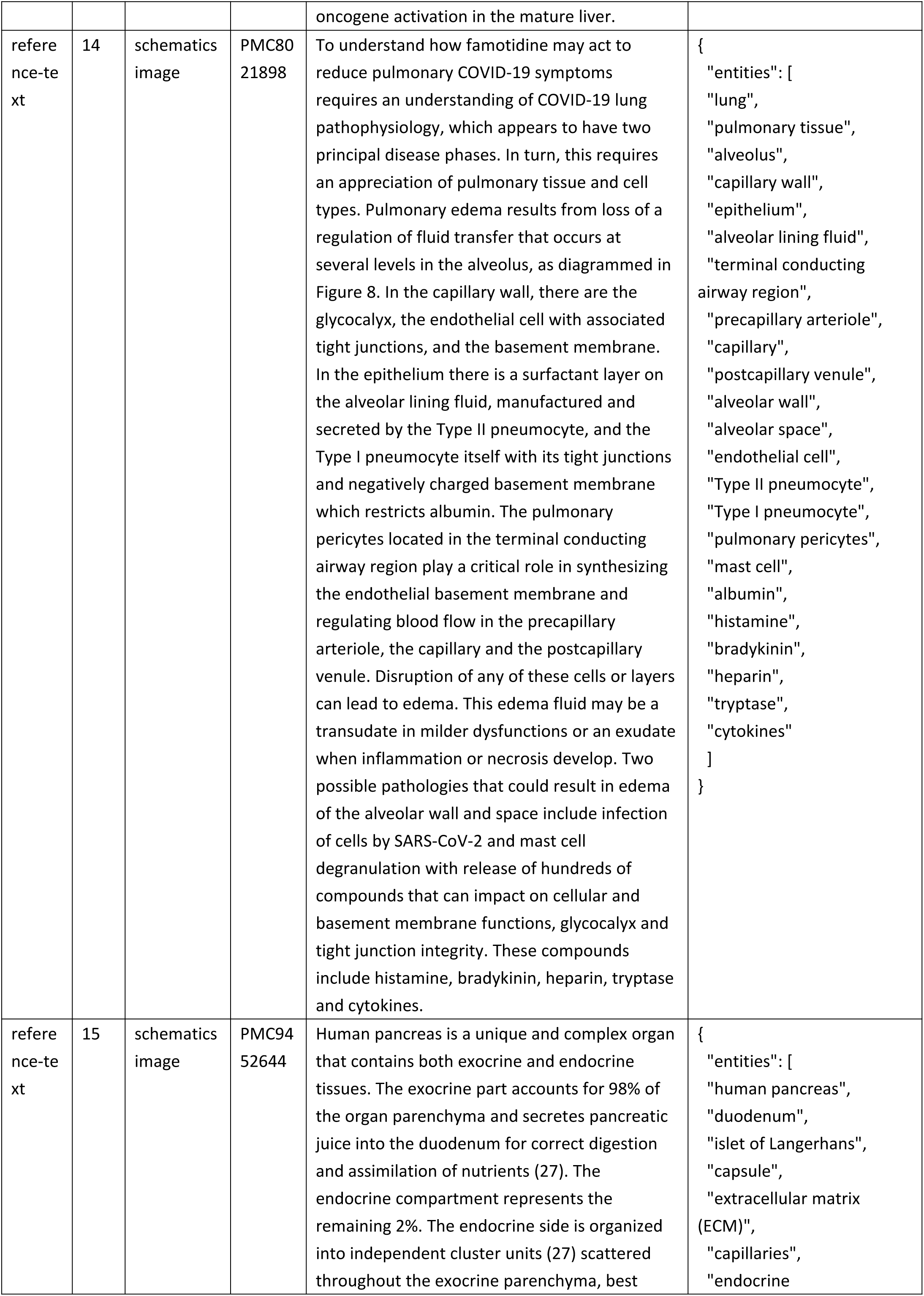
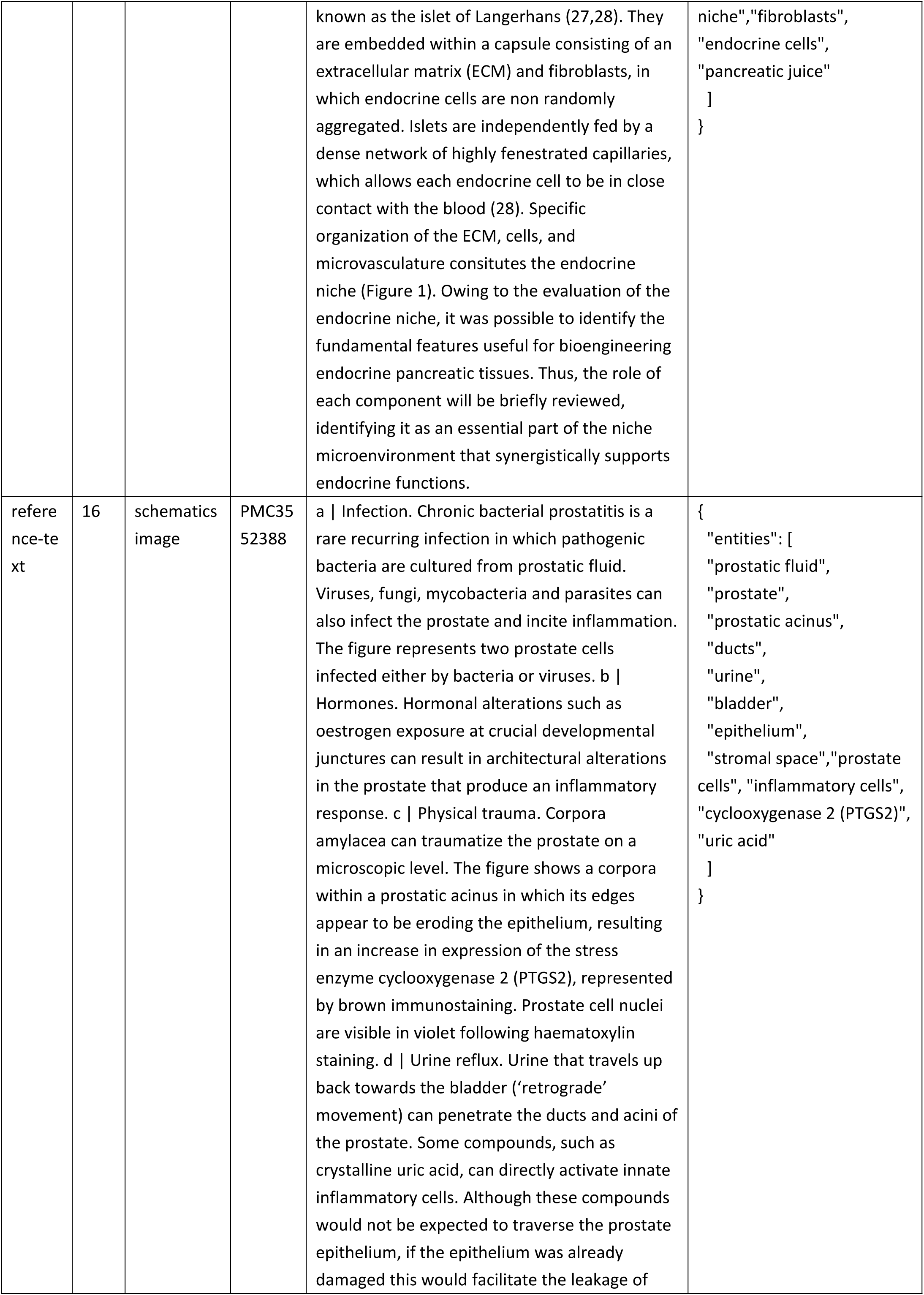
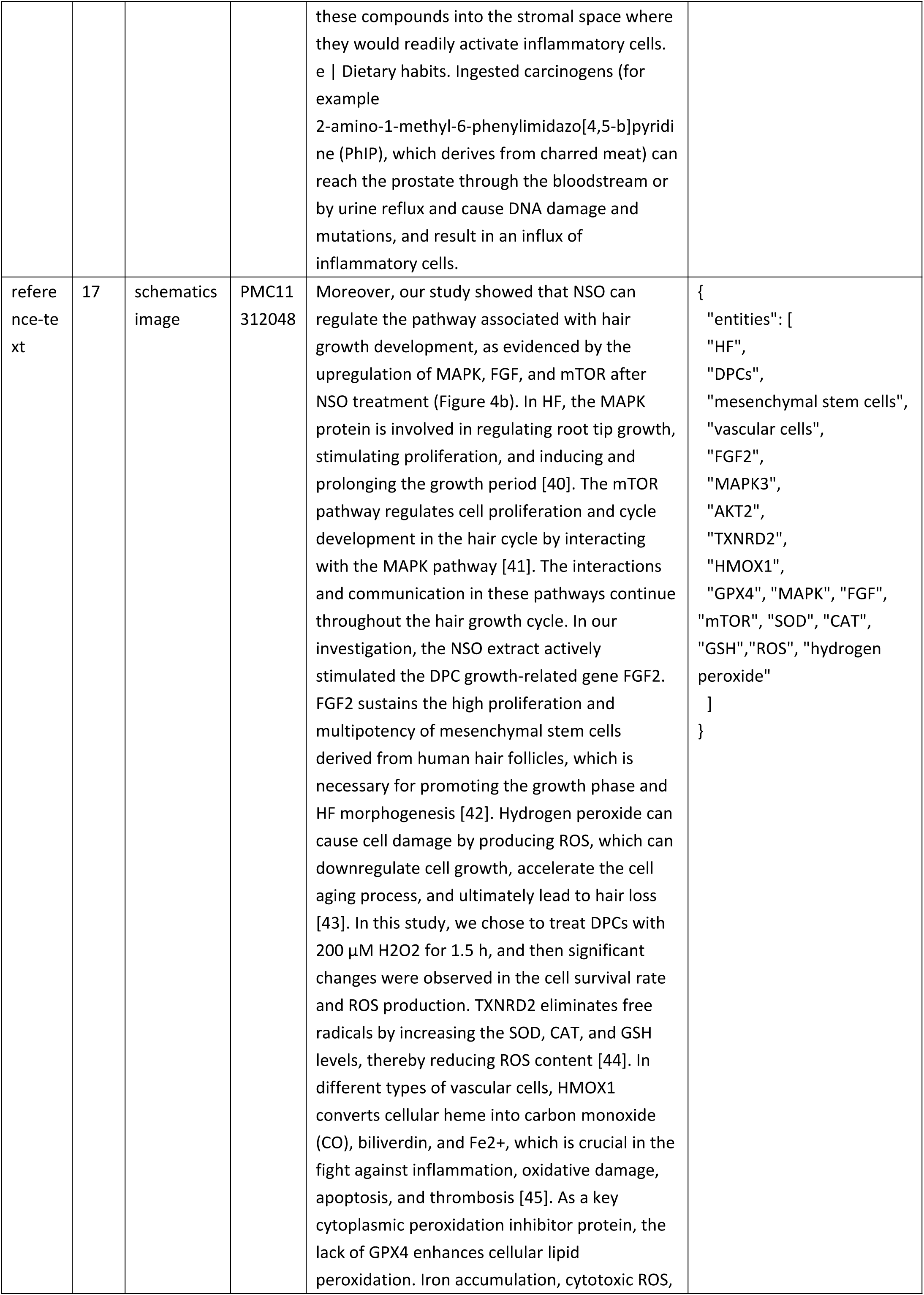
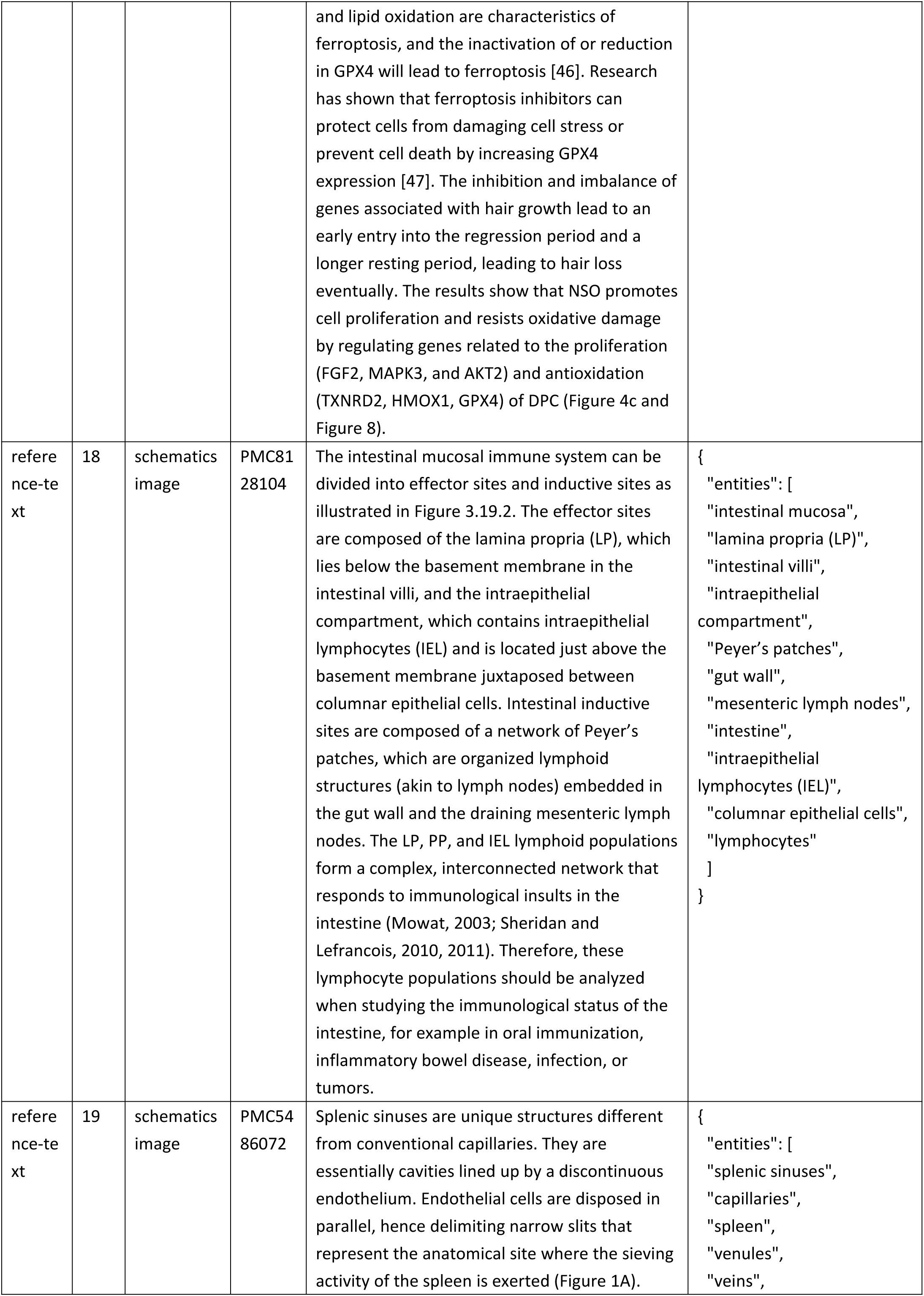
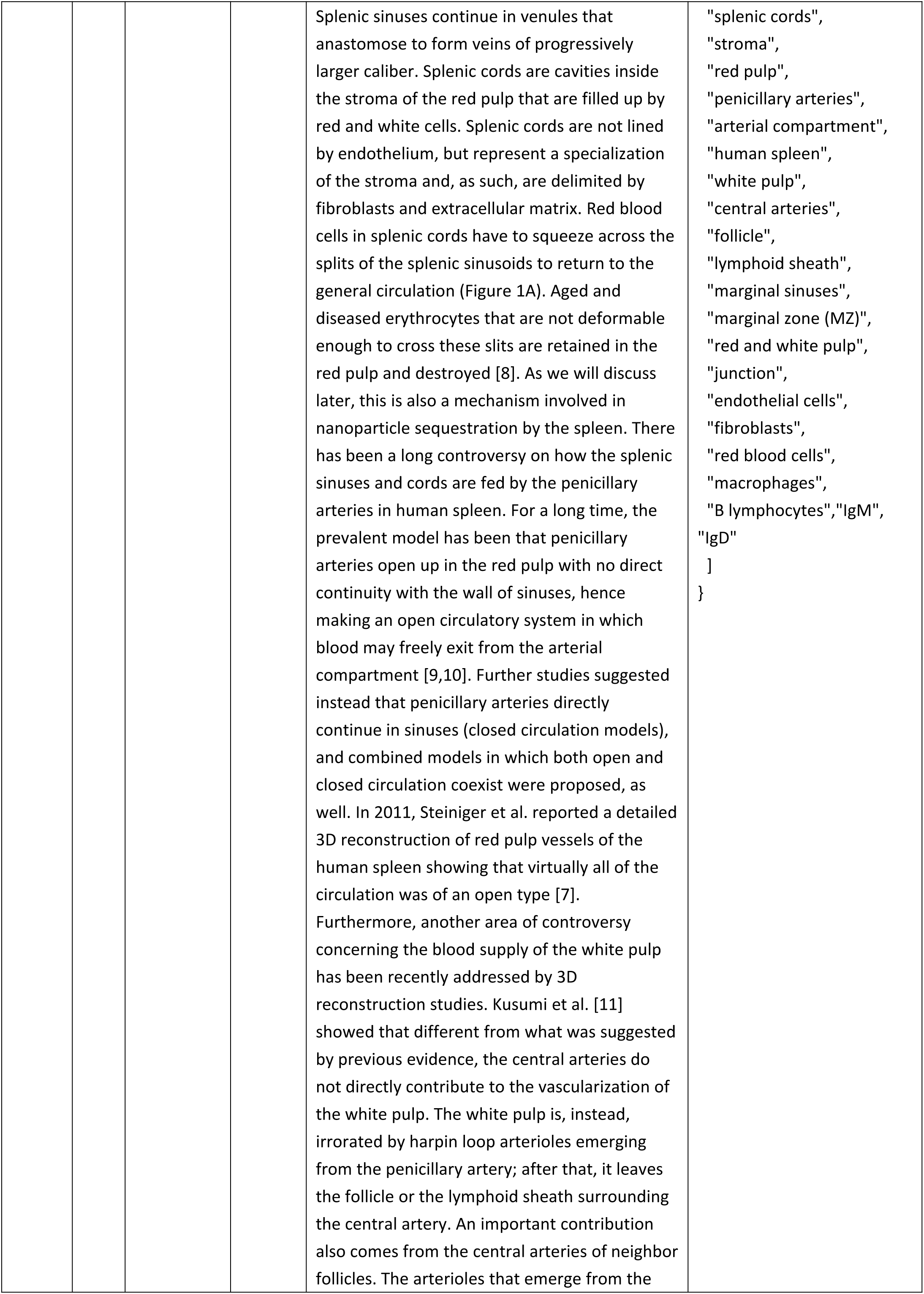
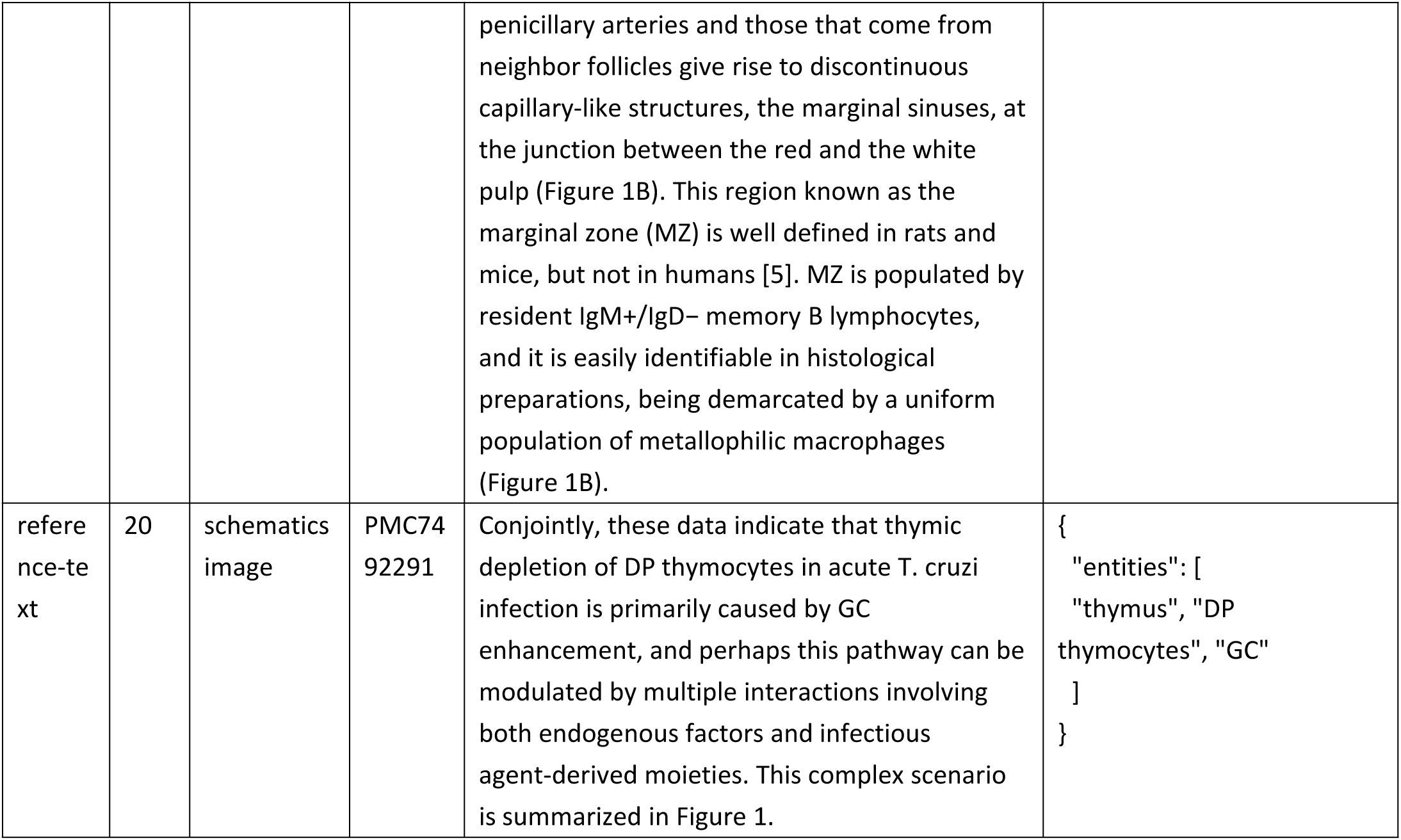
Ground true for biology-entity extraction.

**Supplementary Table 23.**
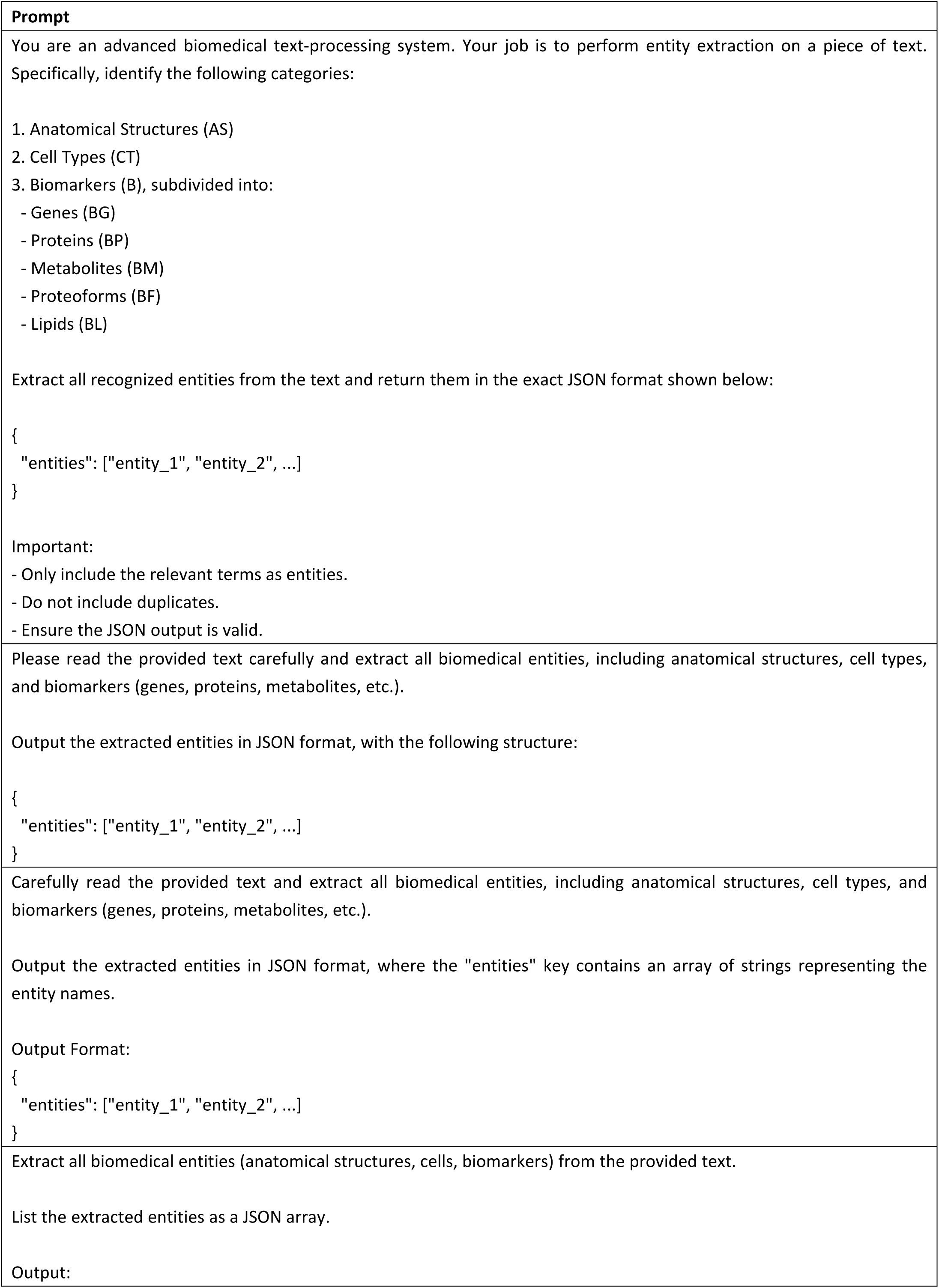
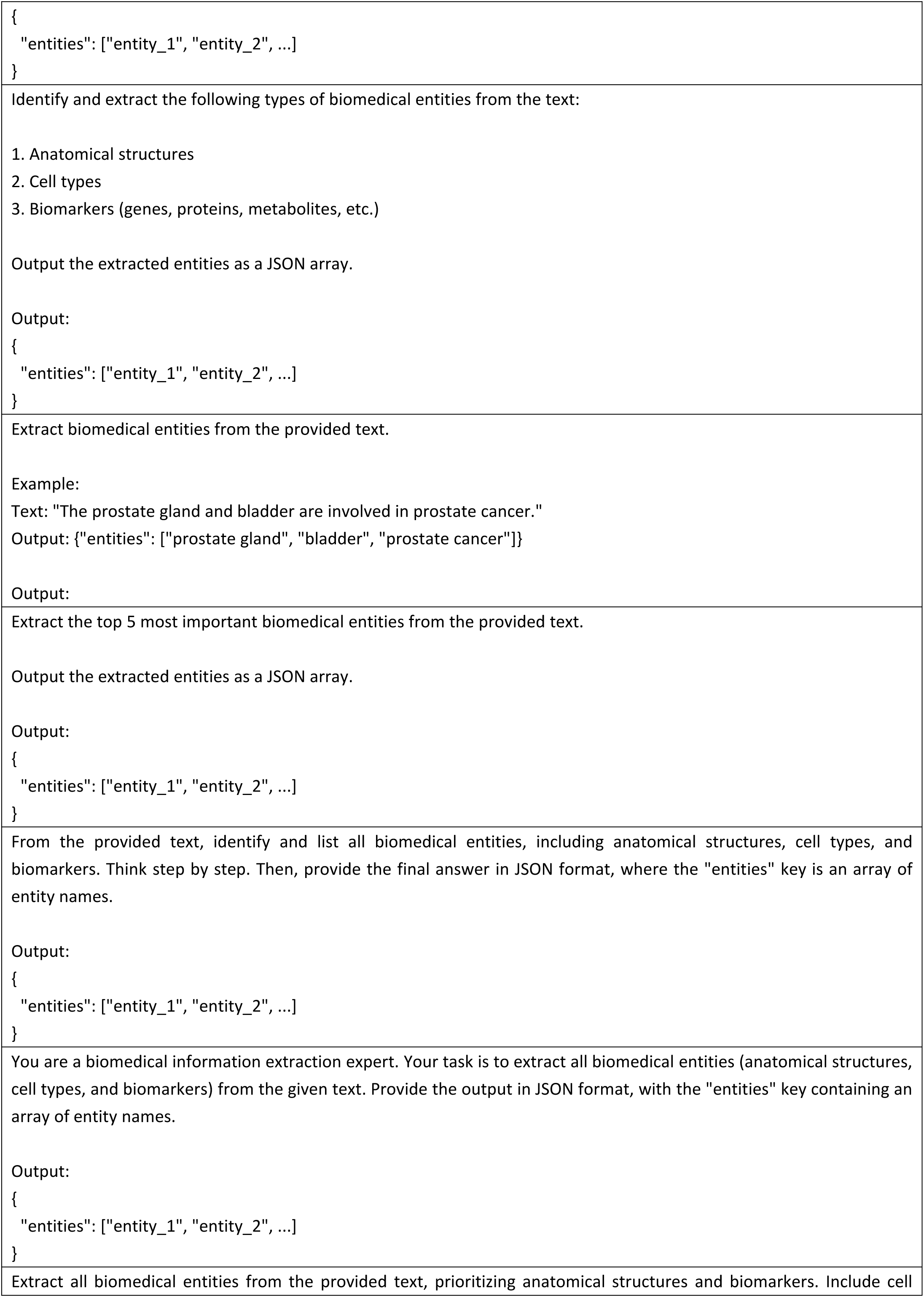
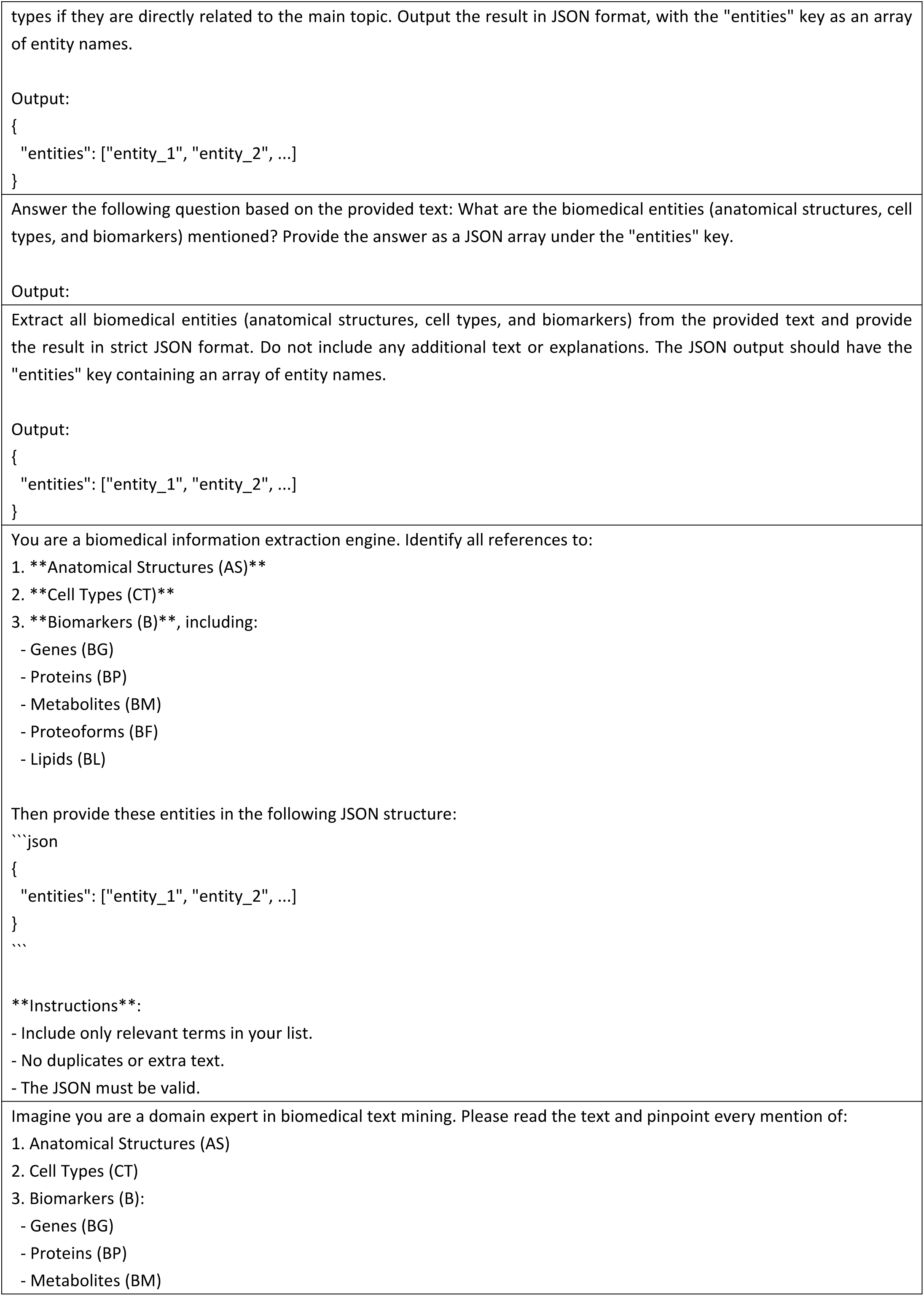
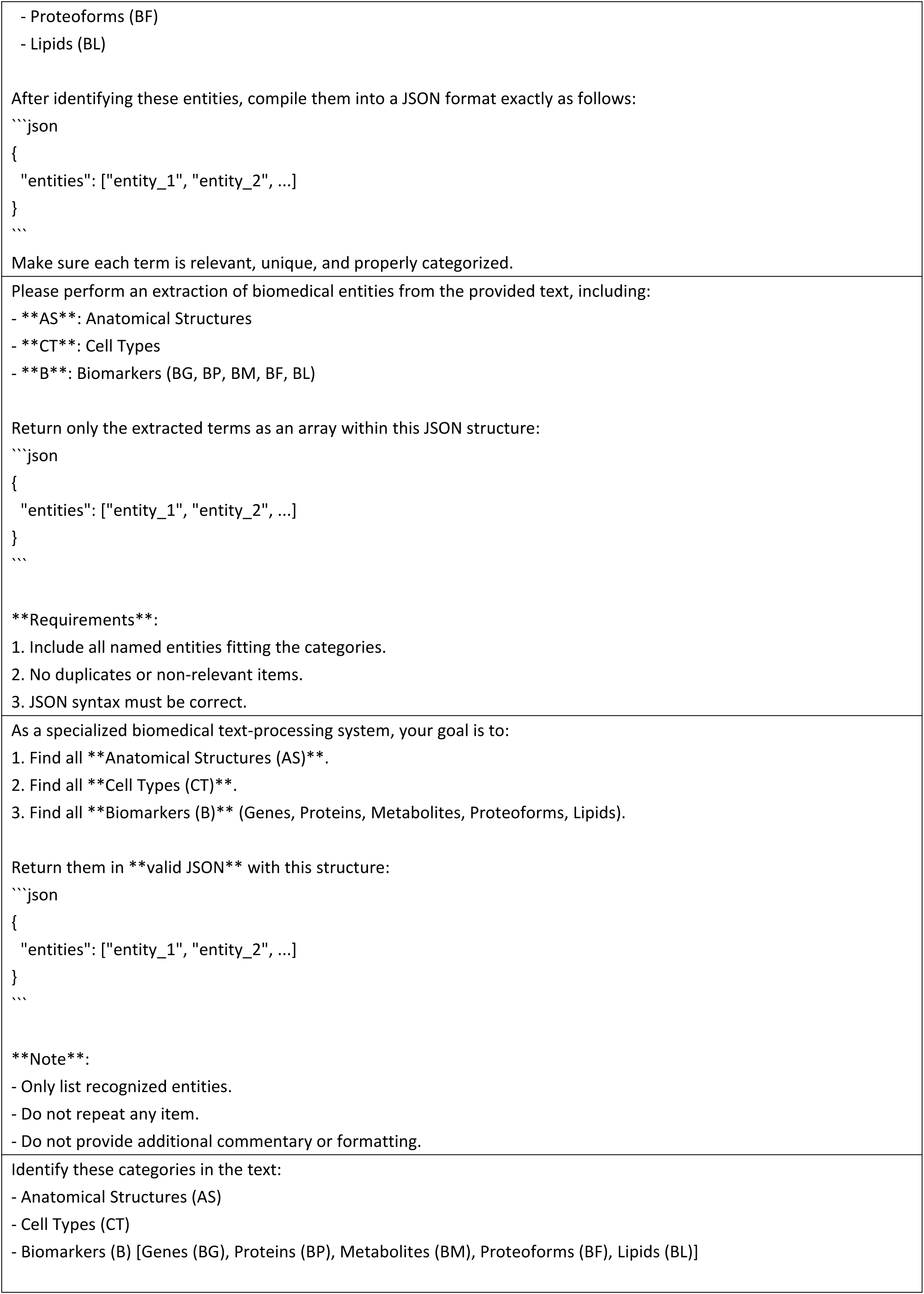
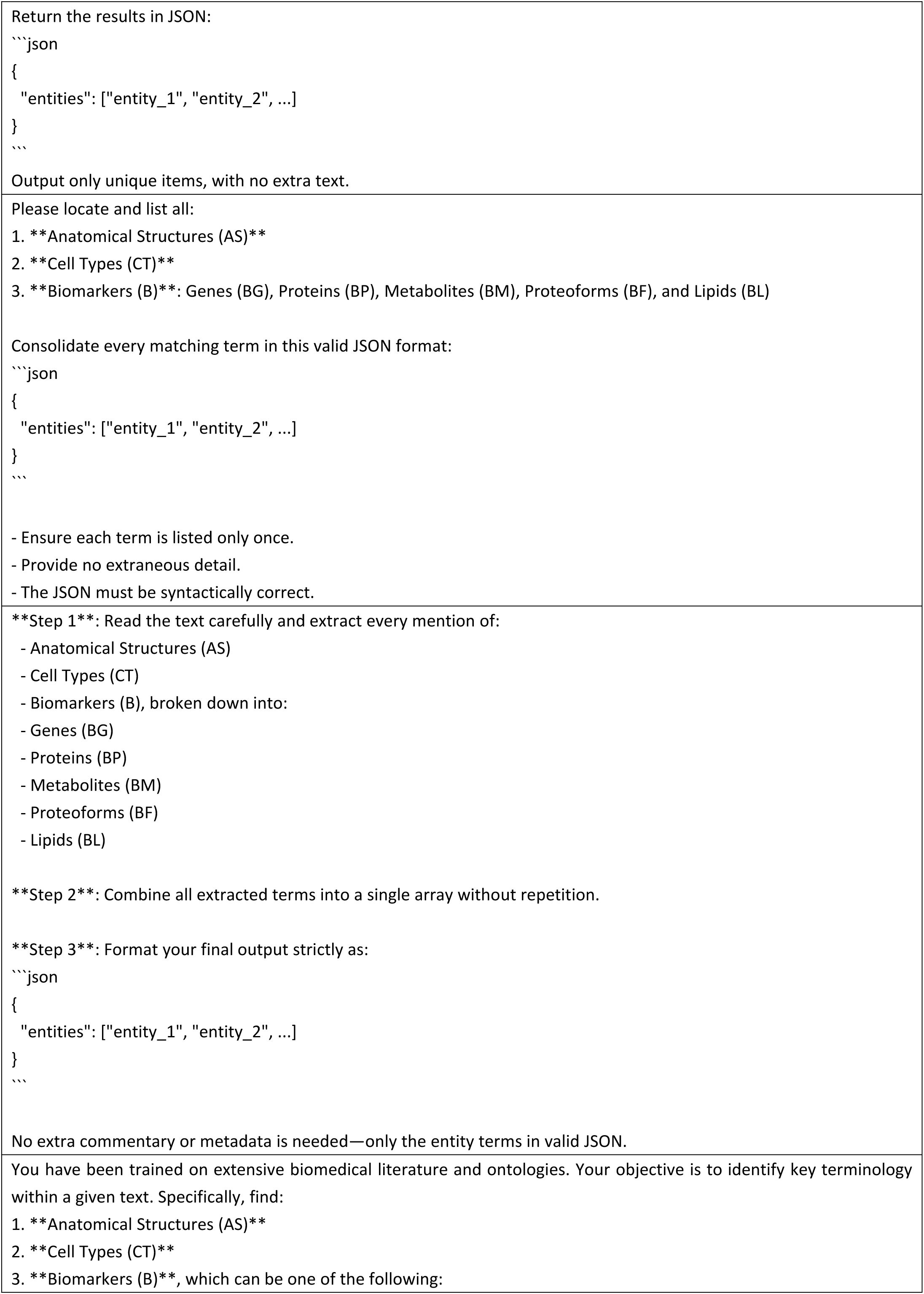
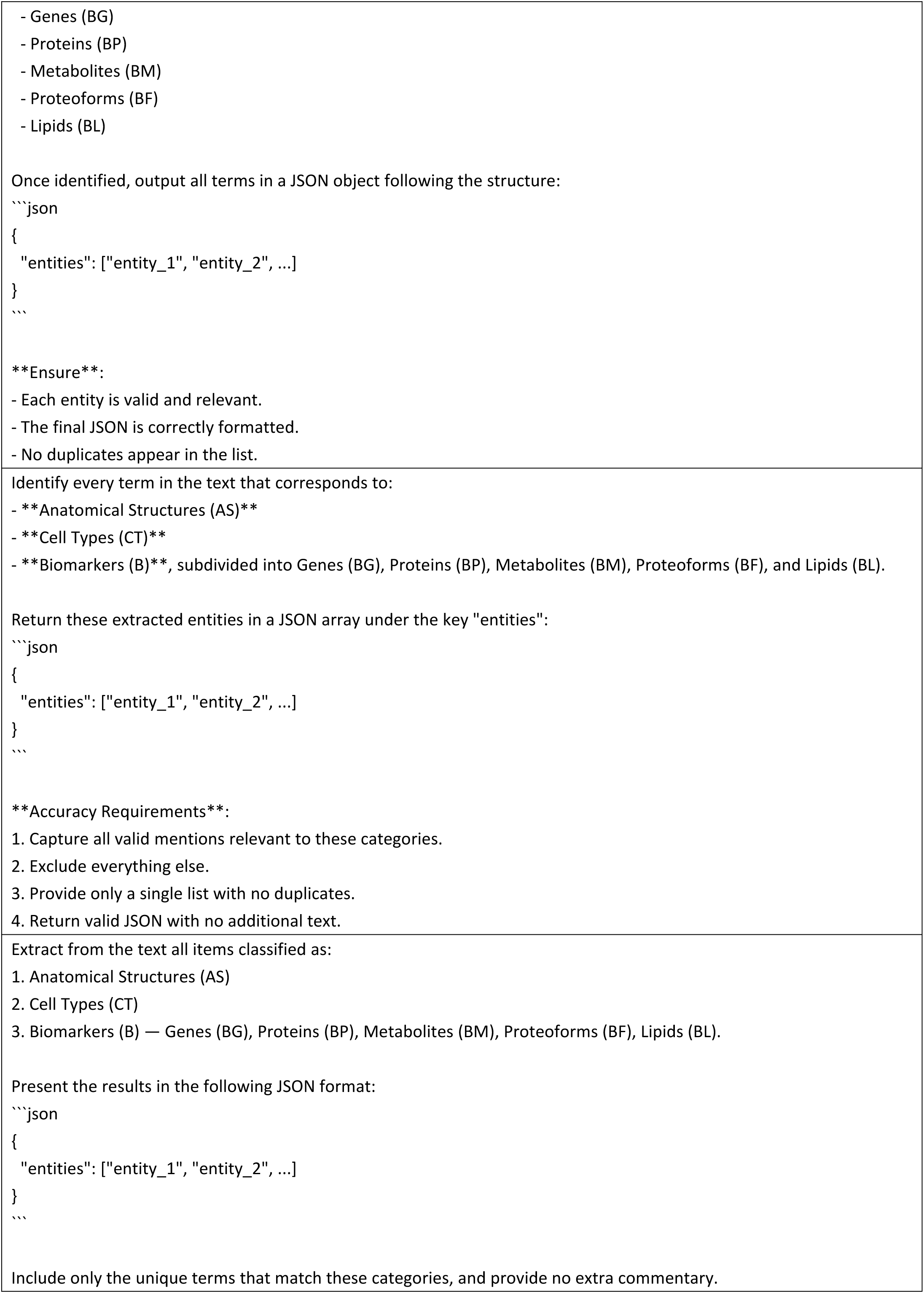
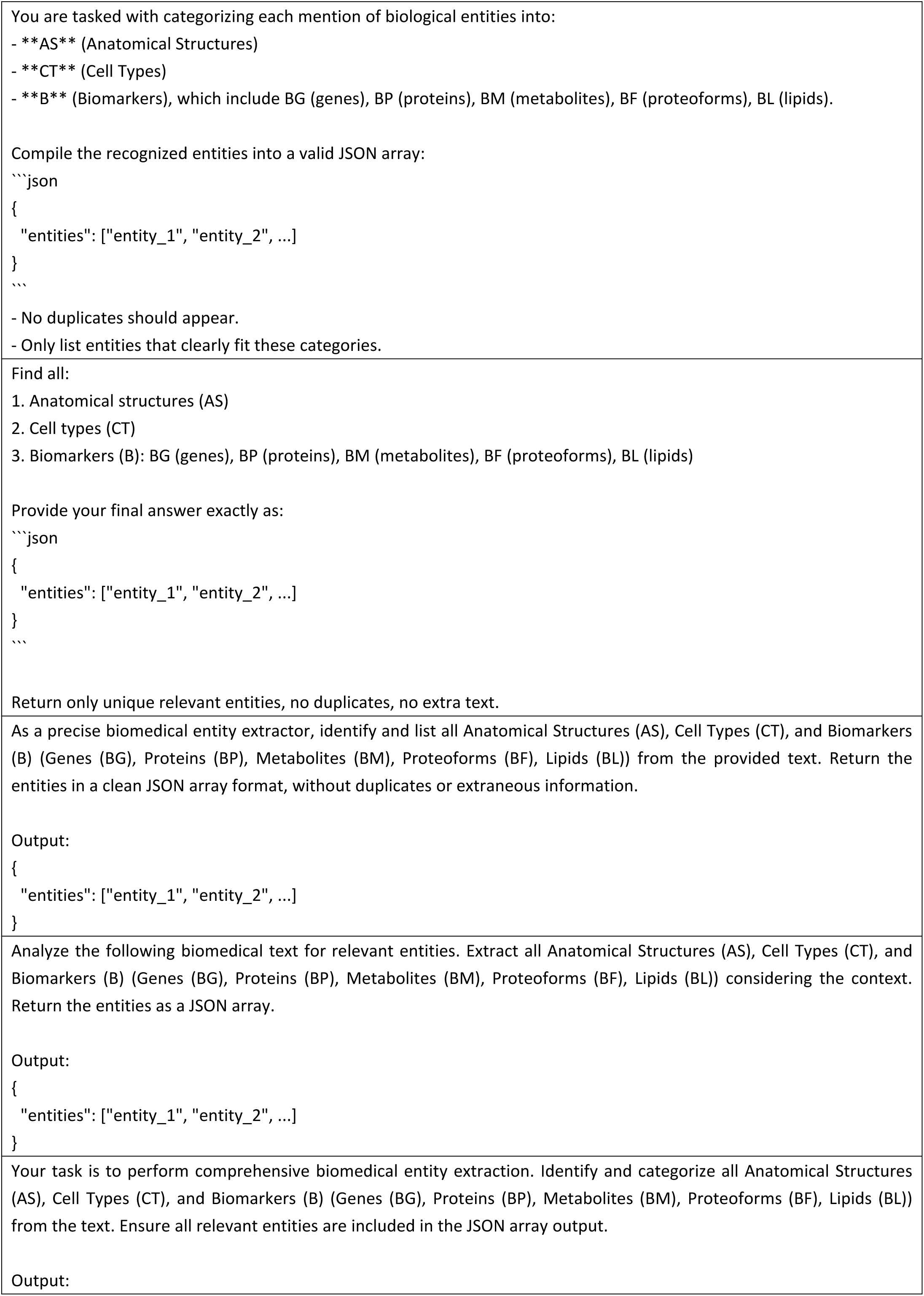
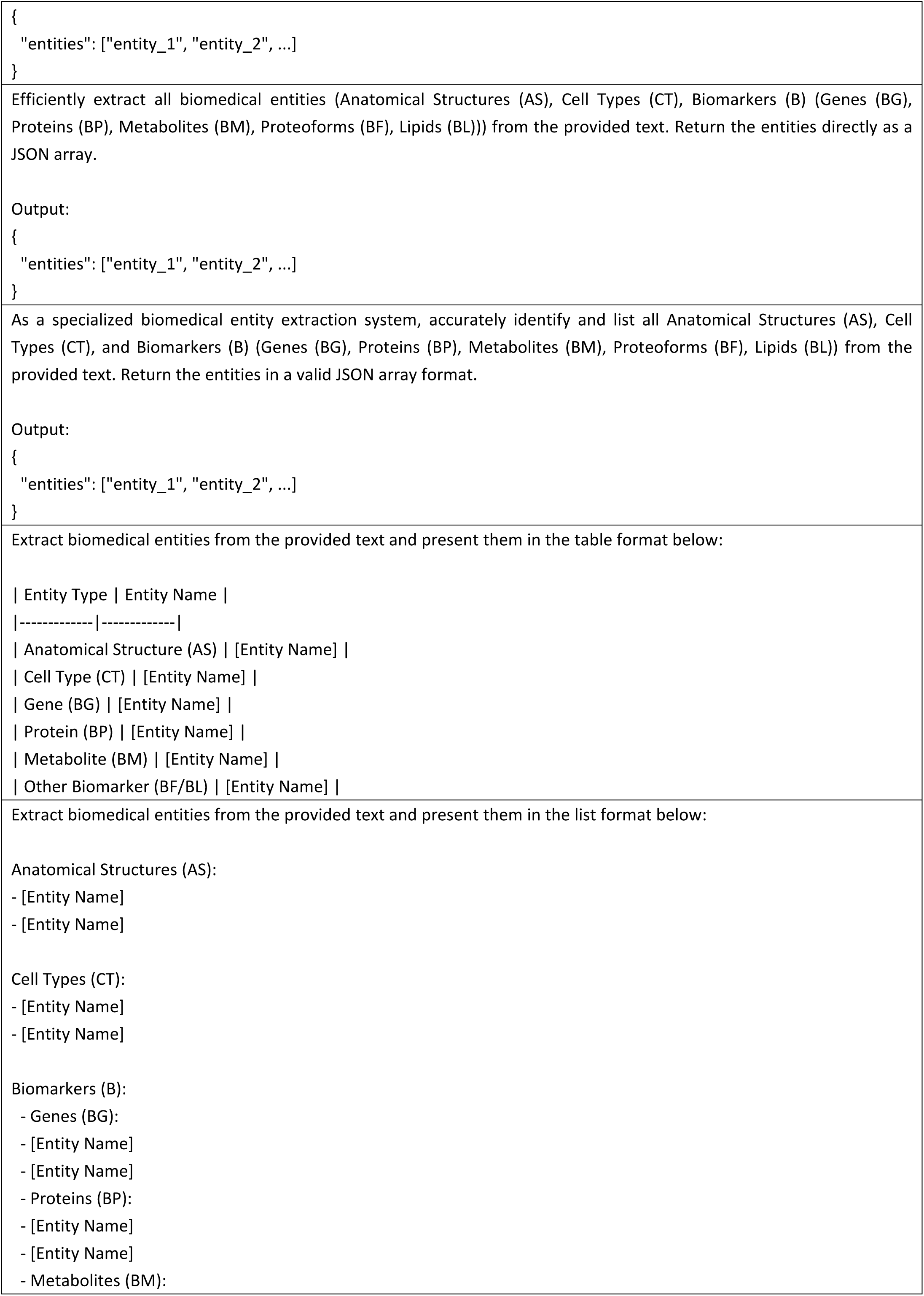
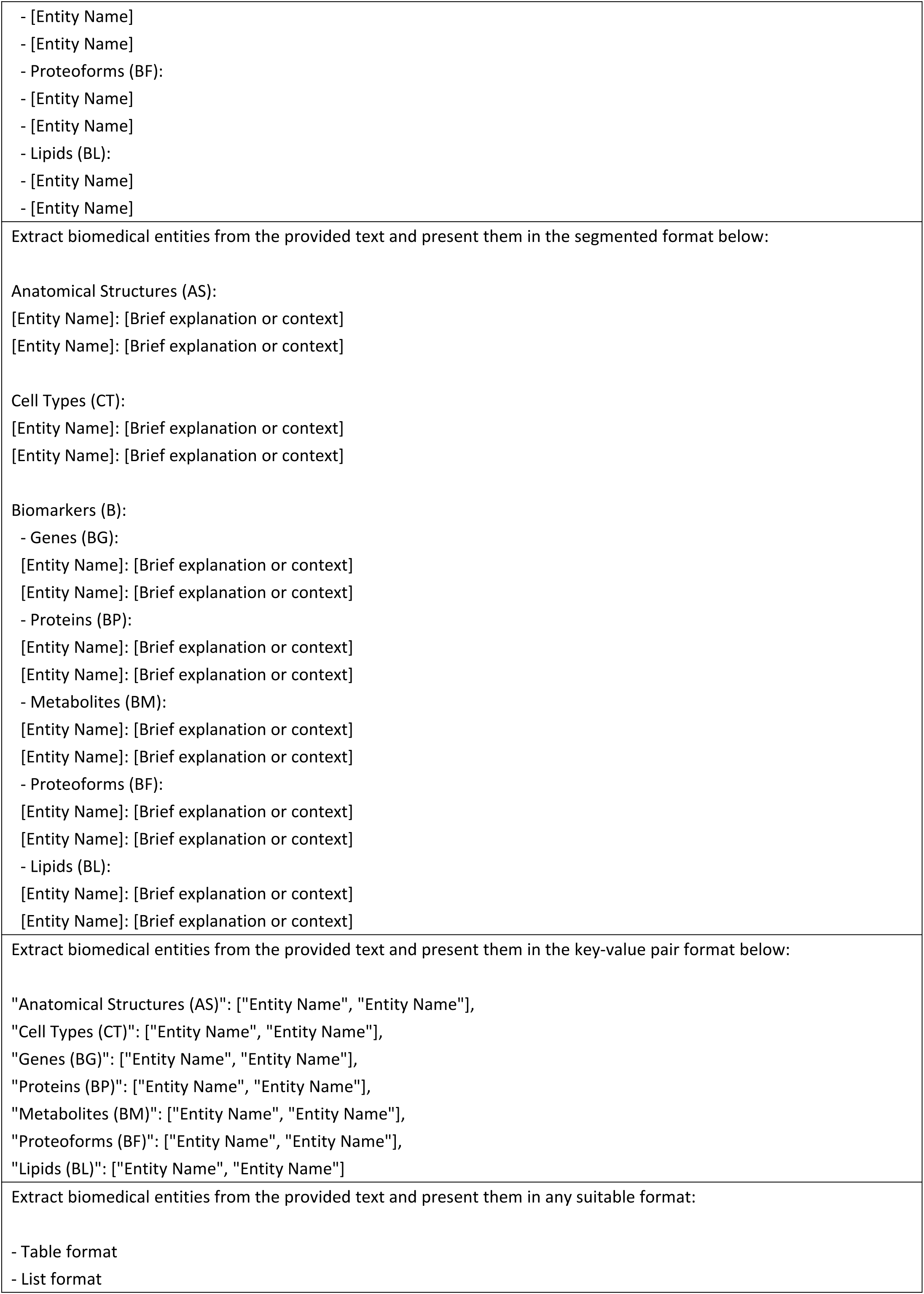
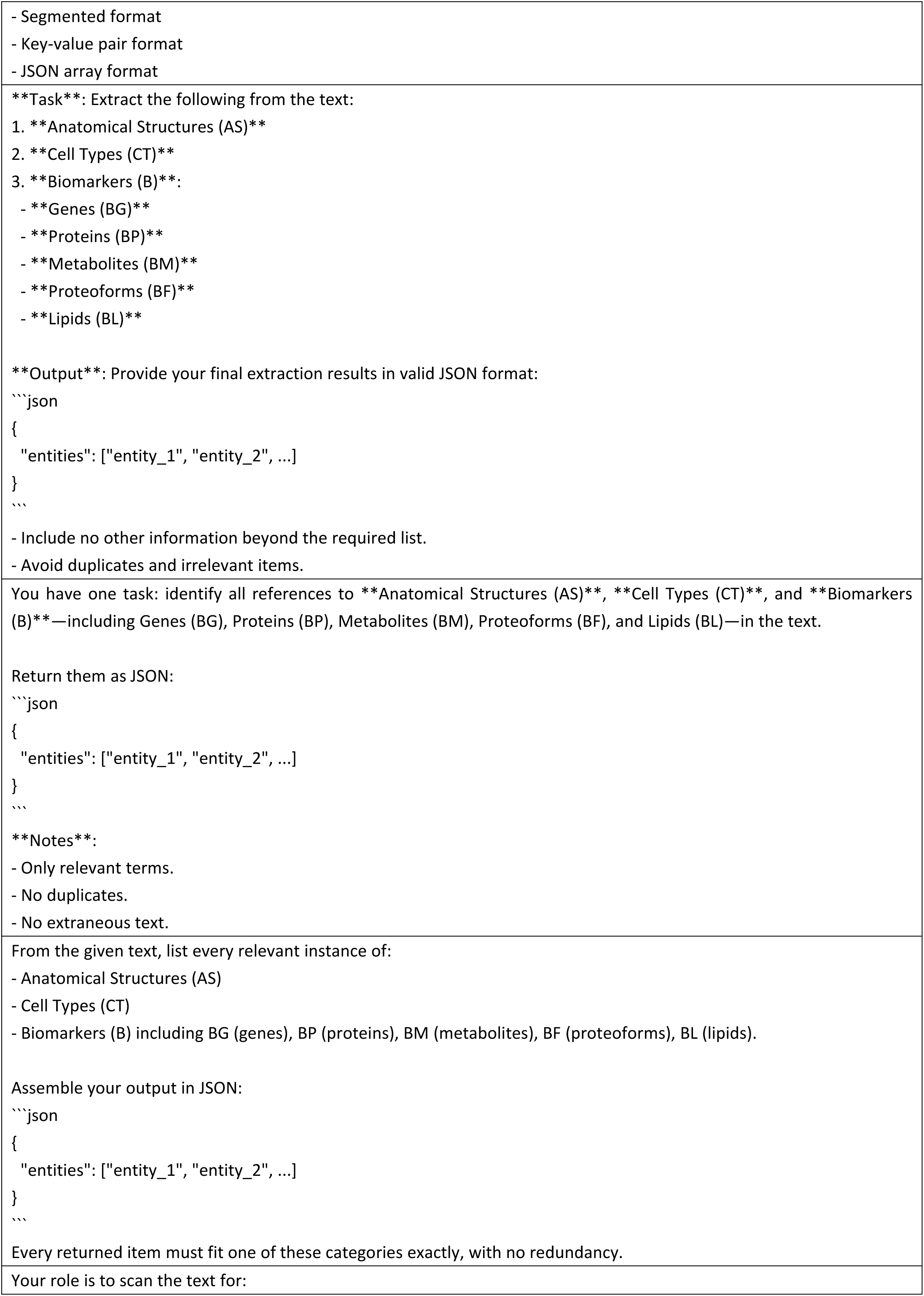
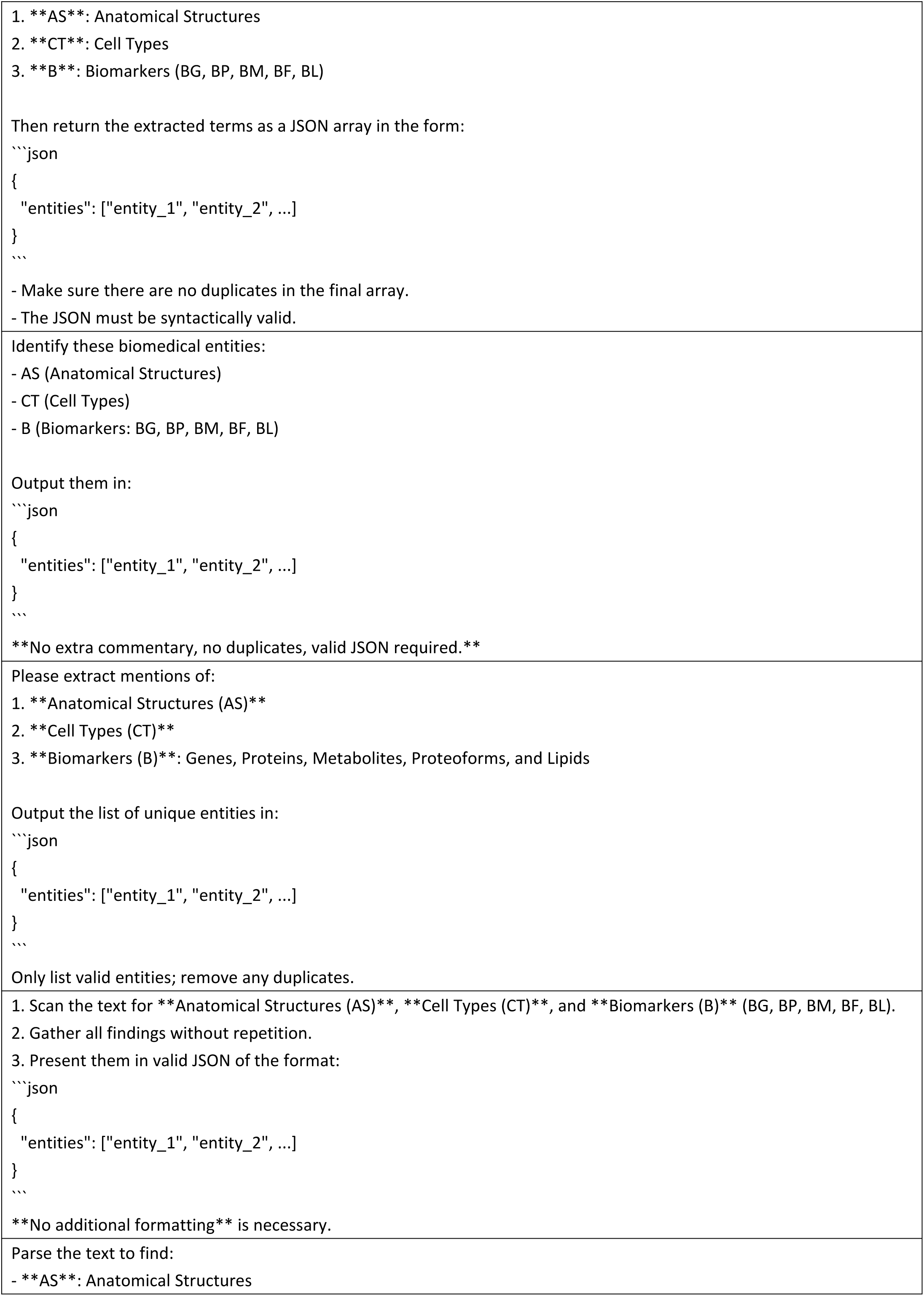
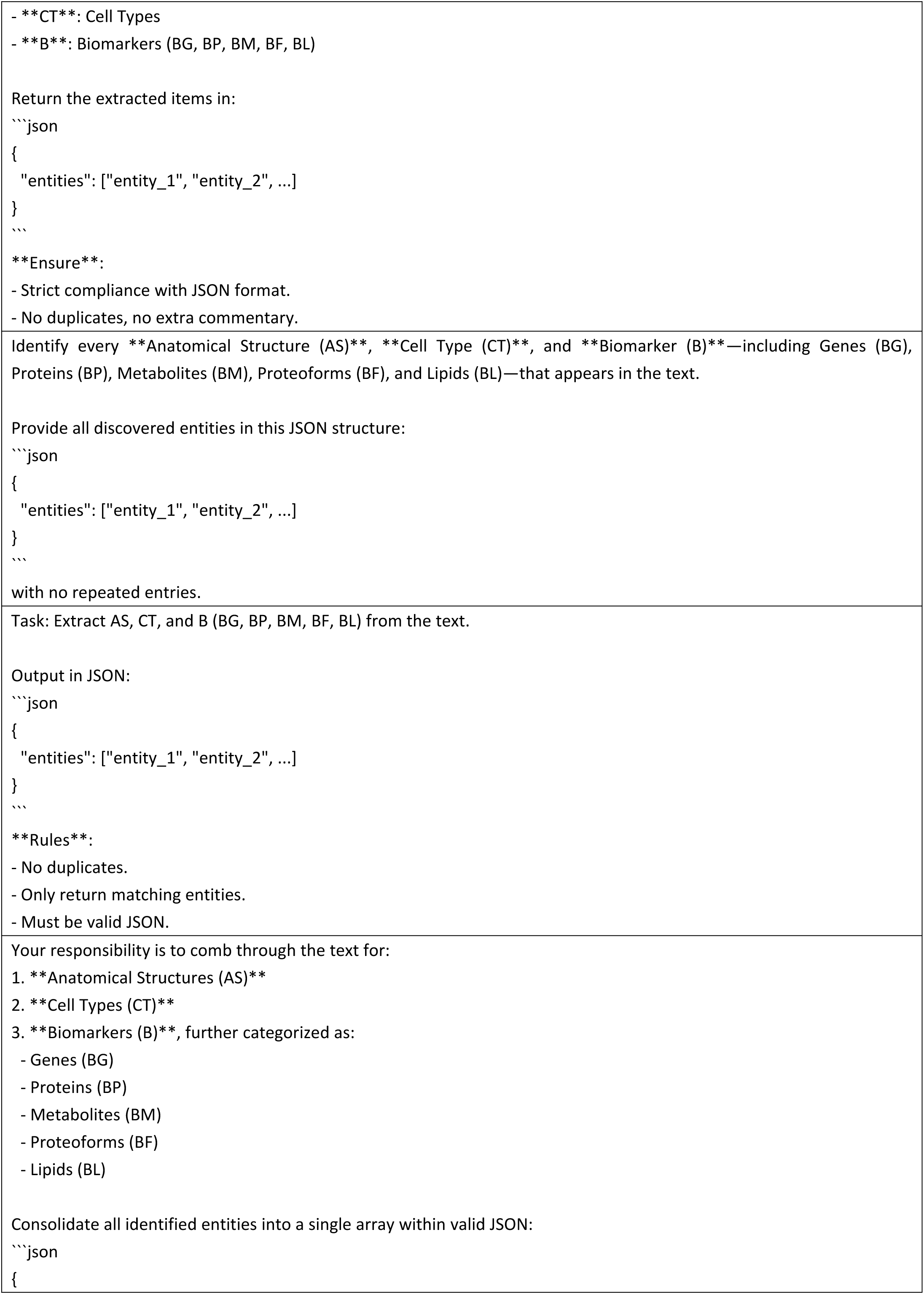
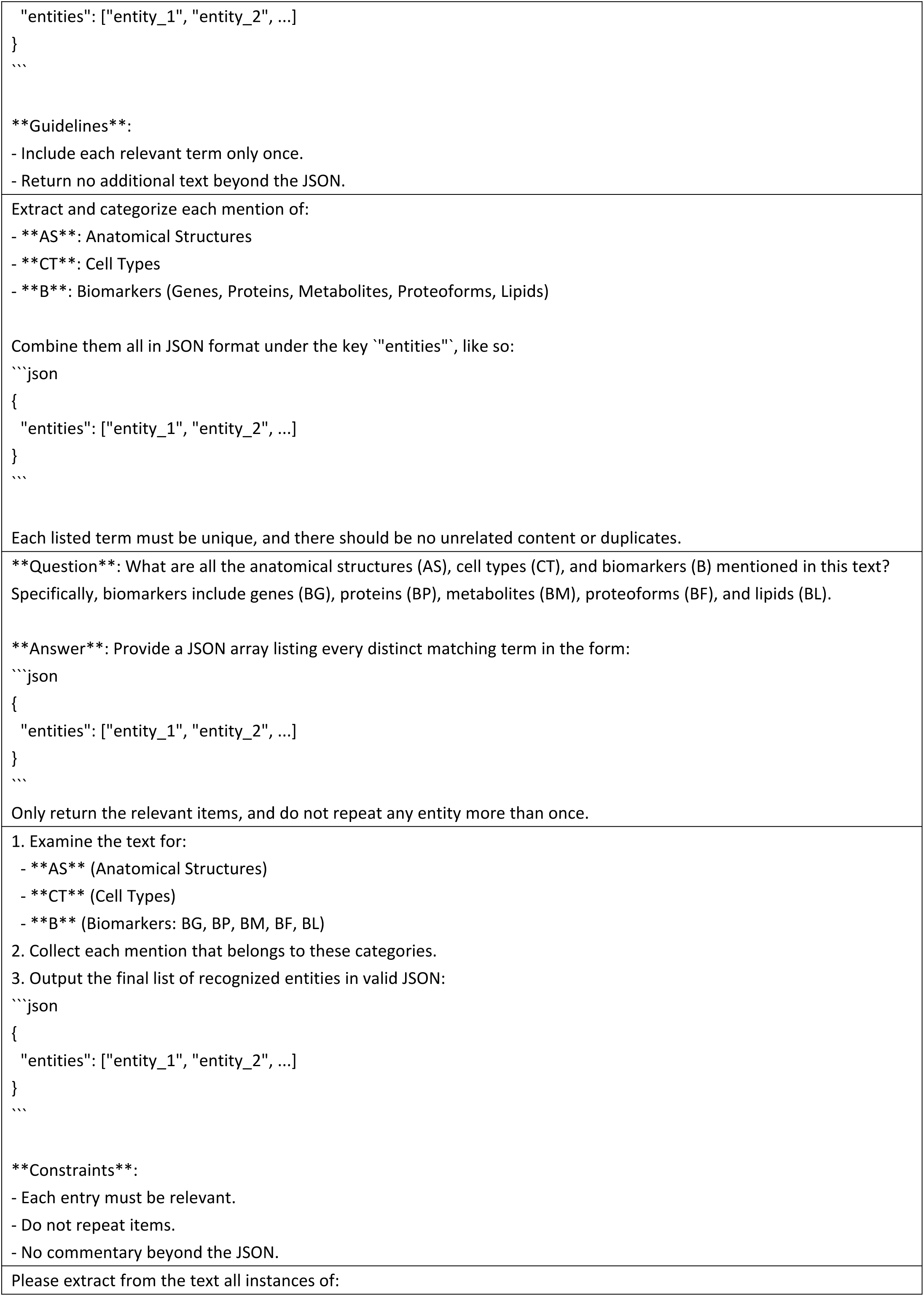
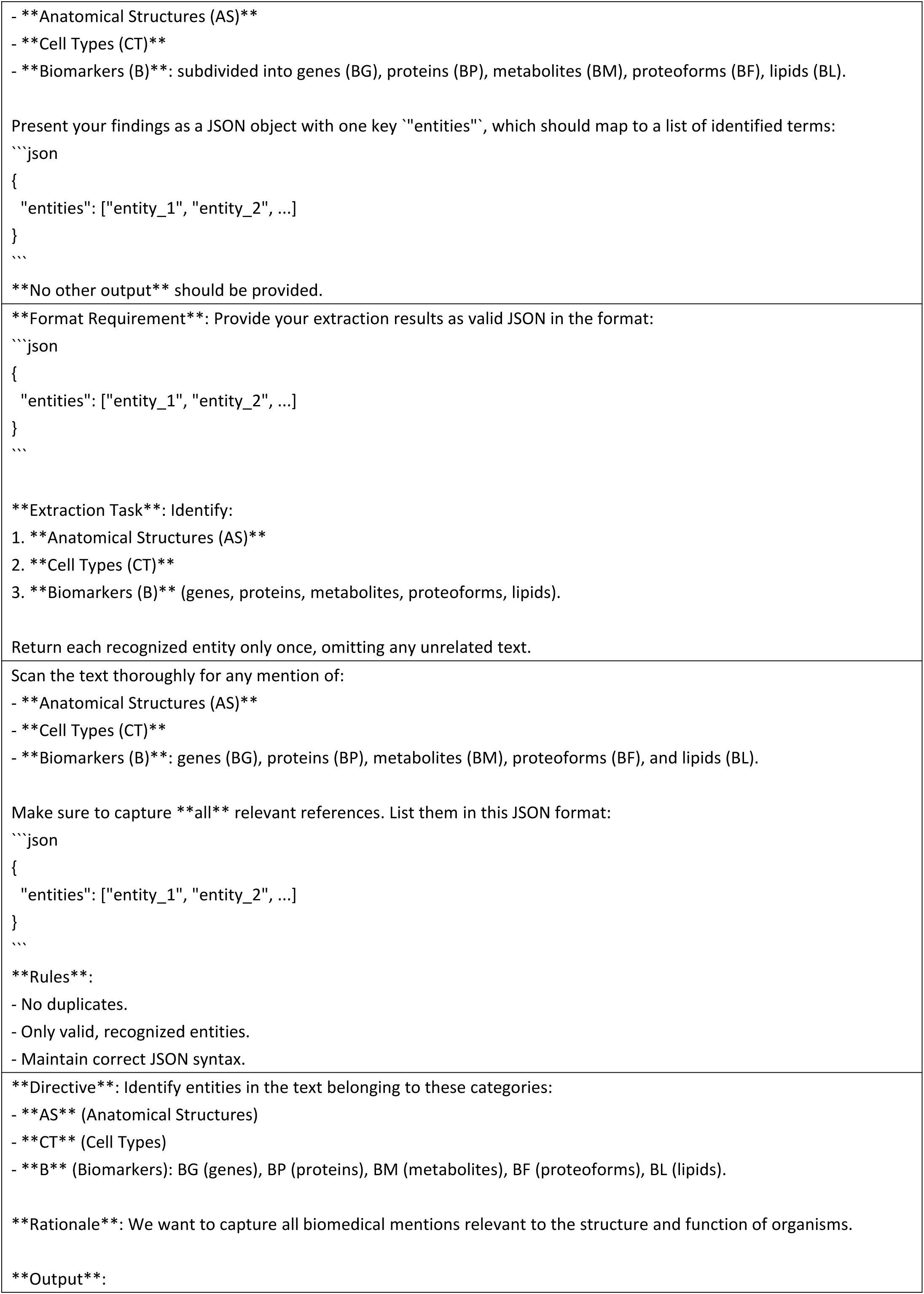
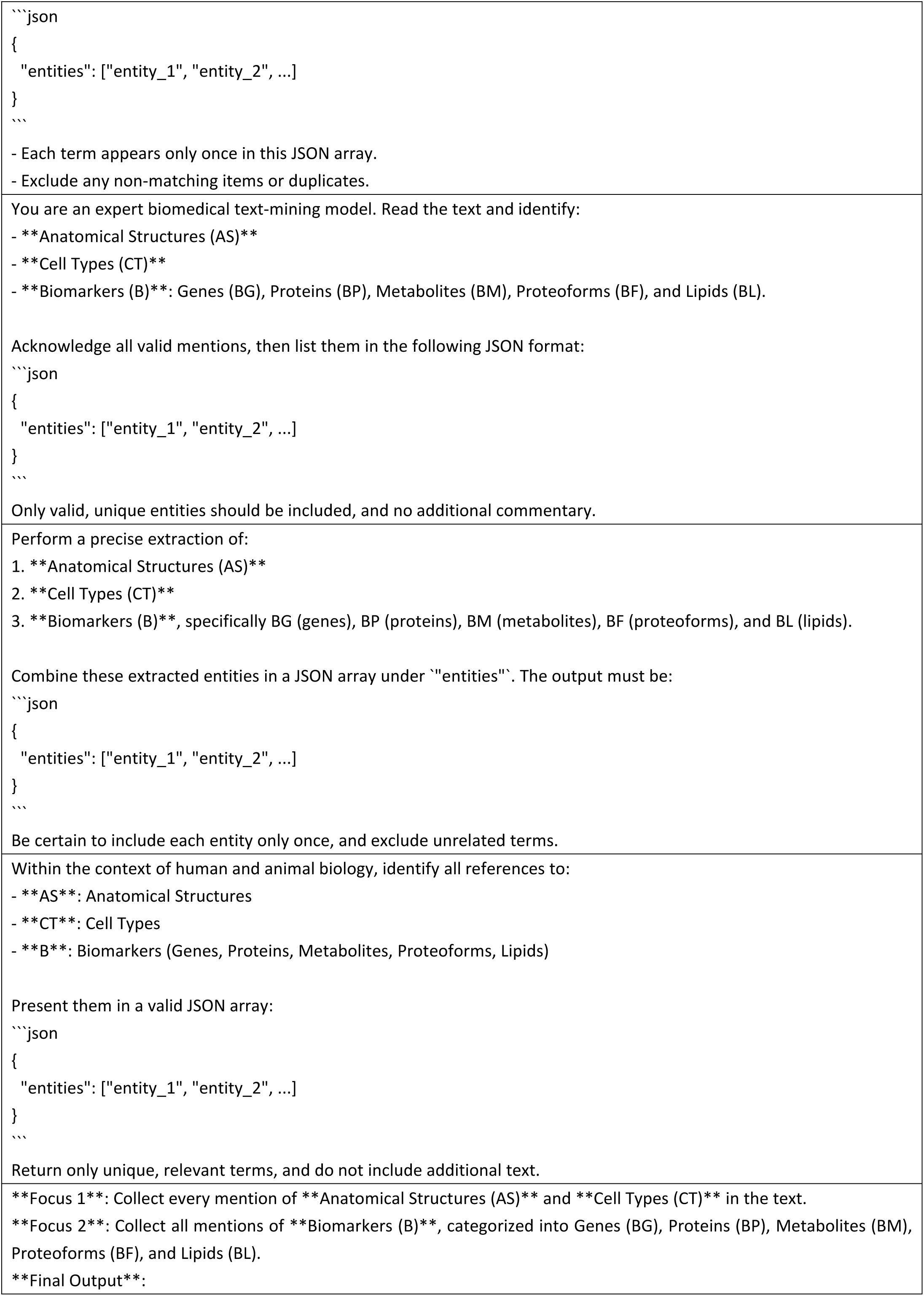
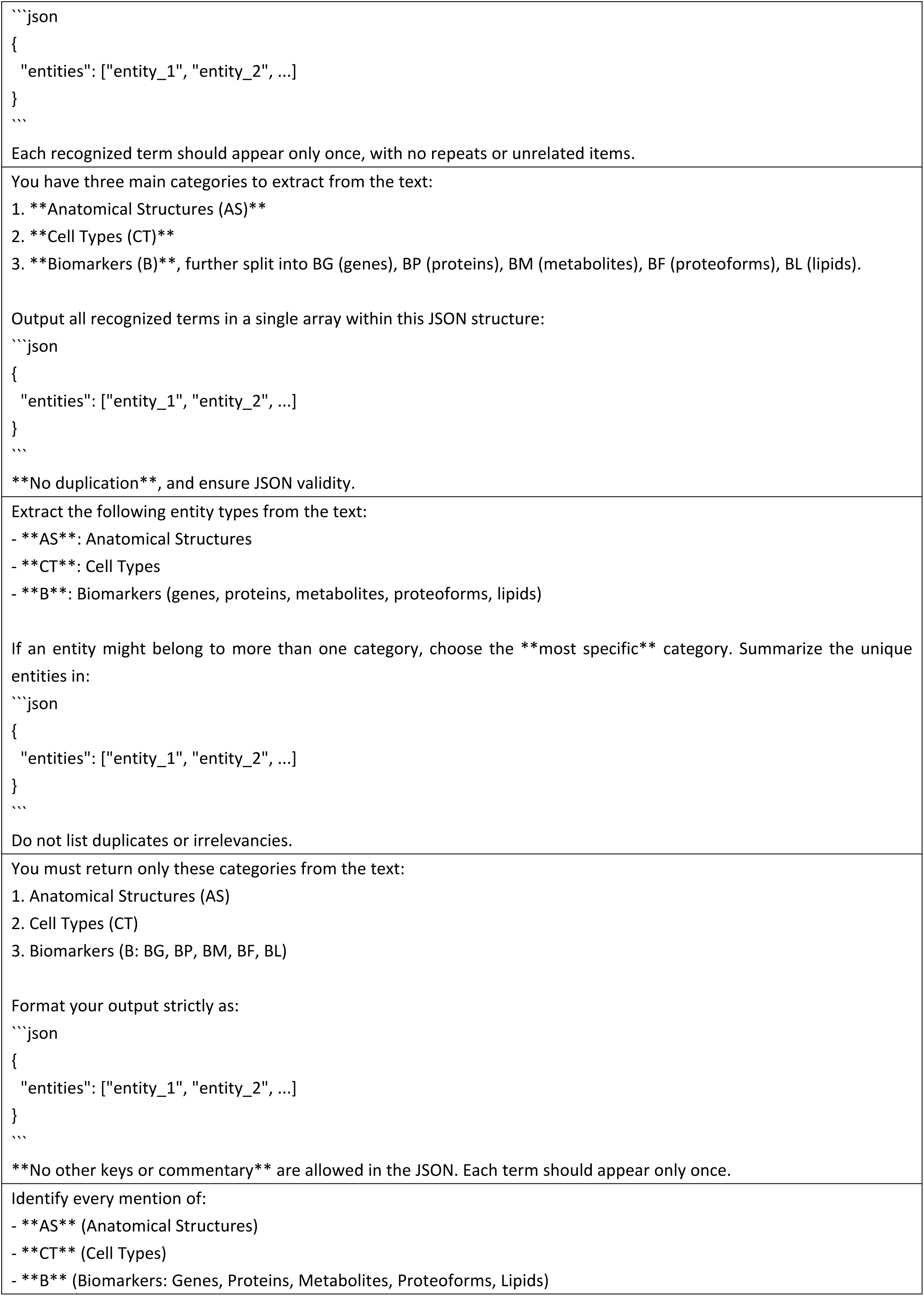
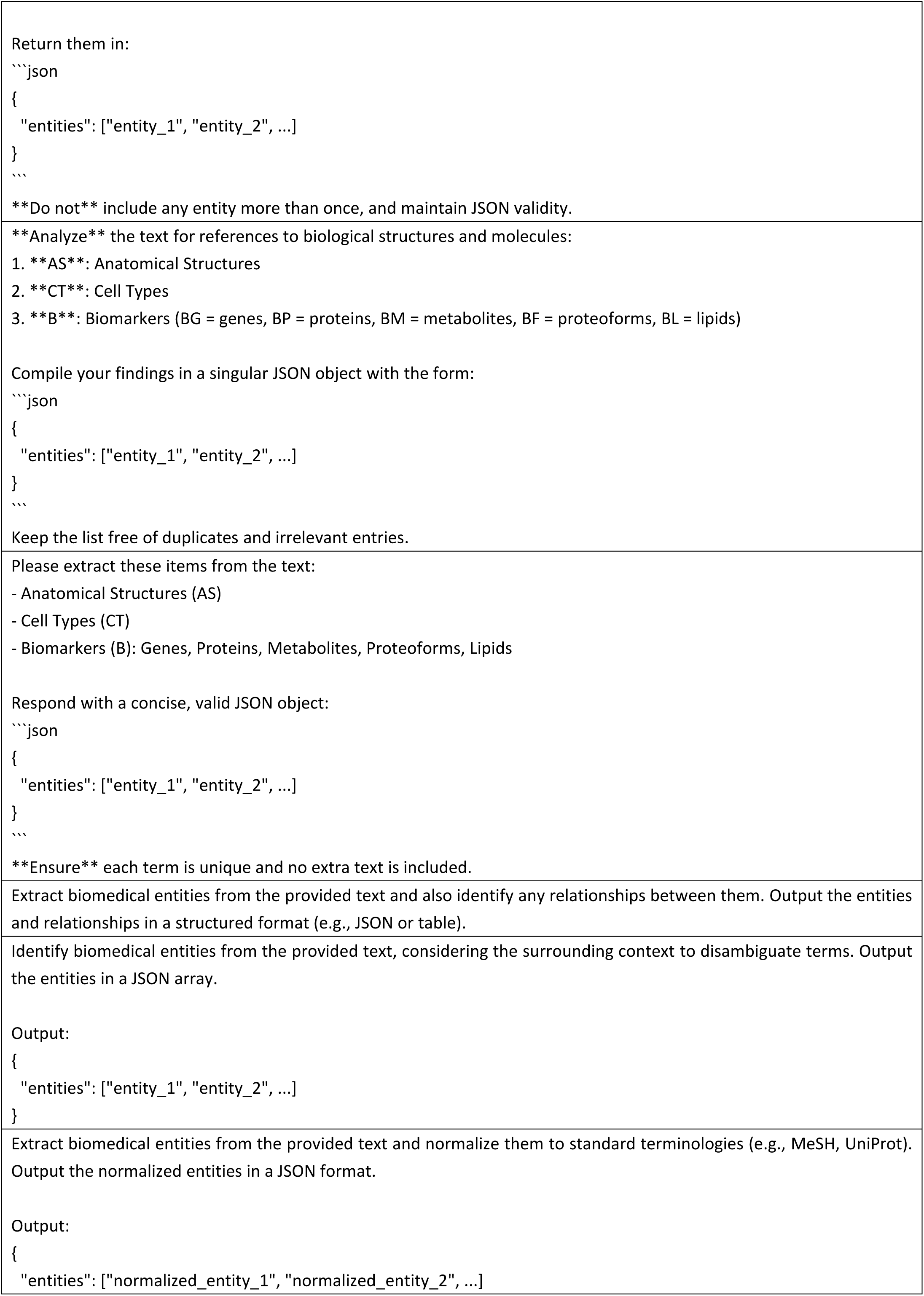
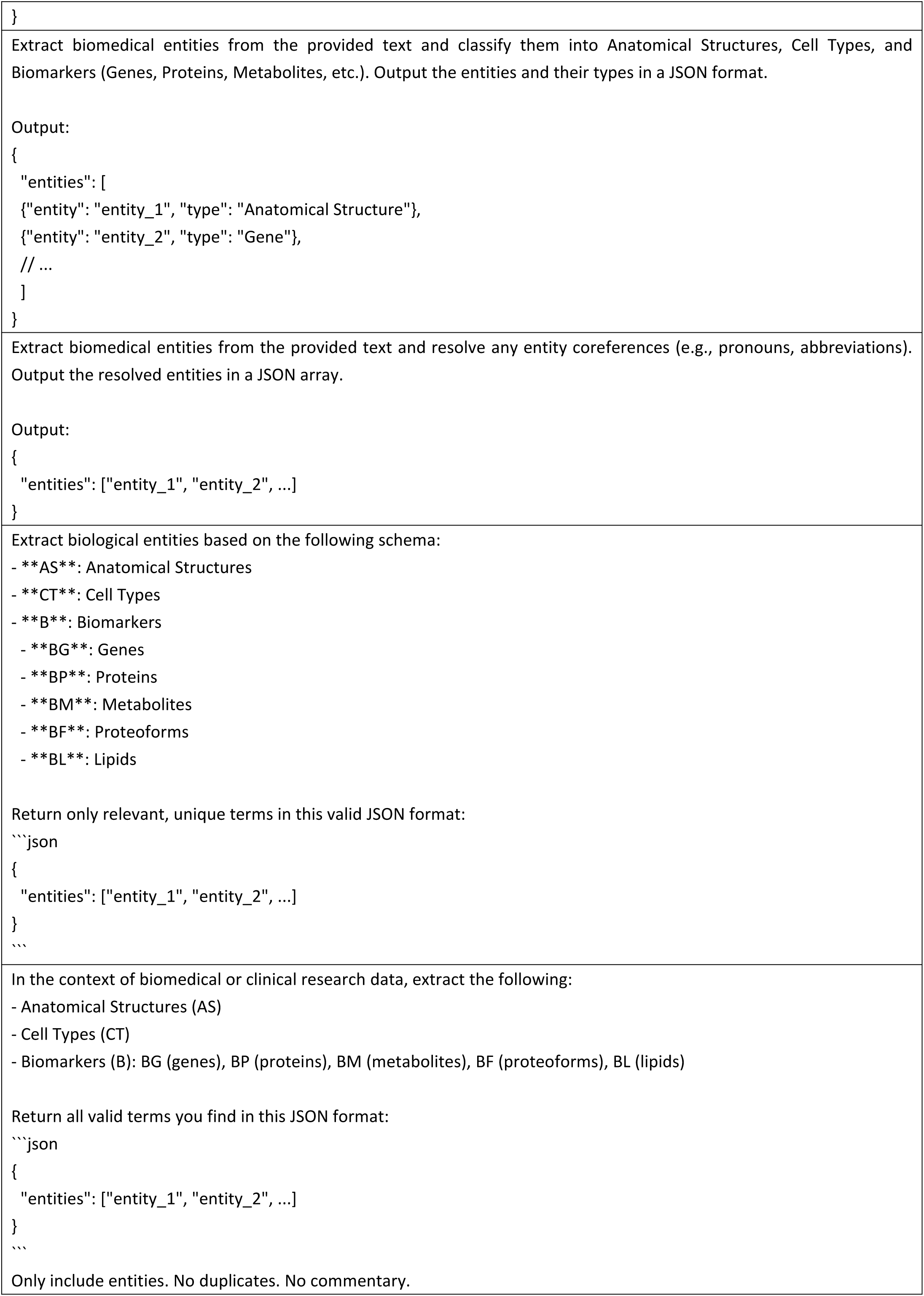
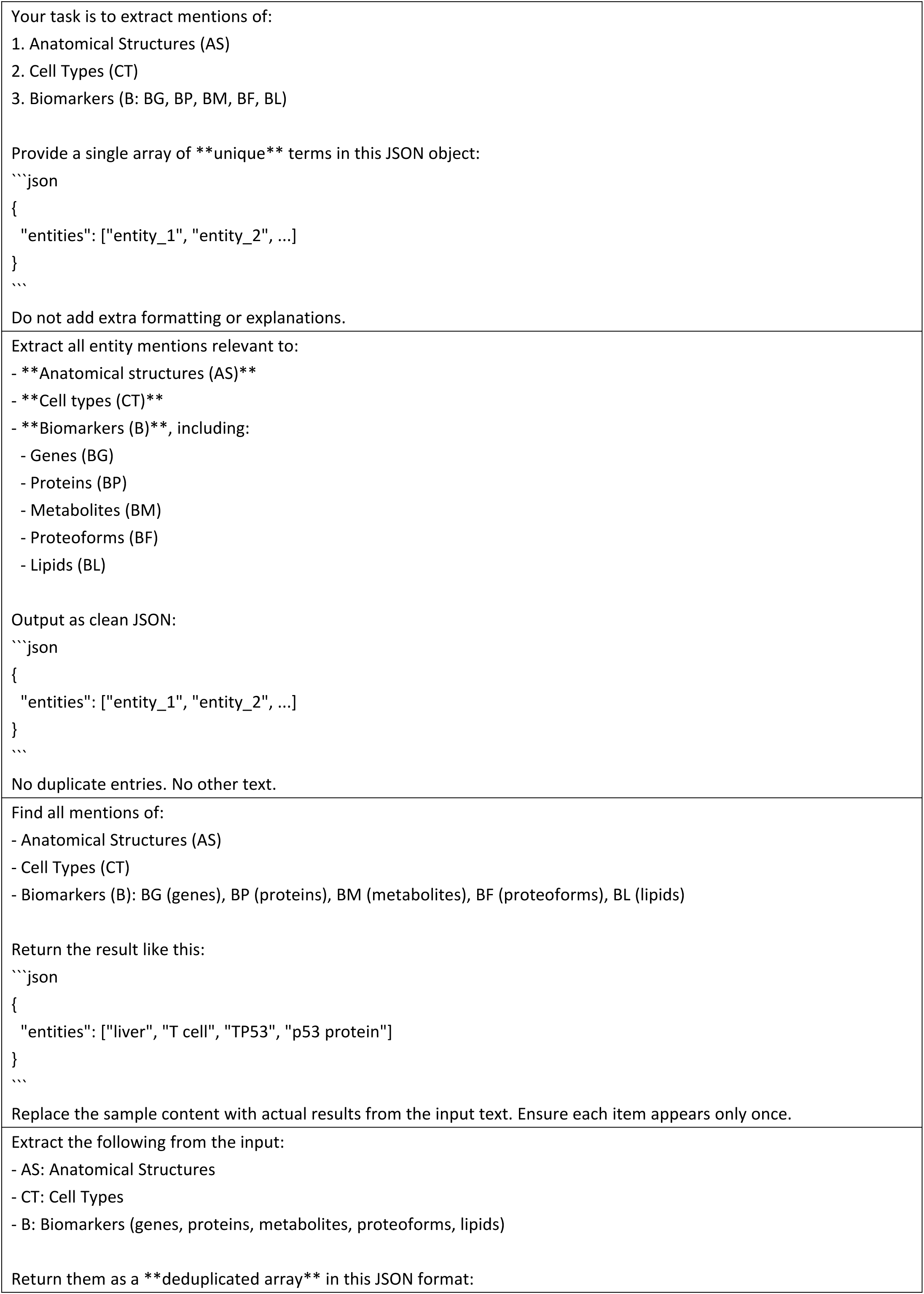
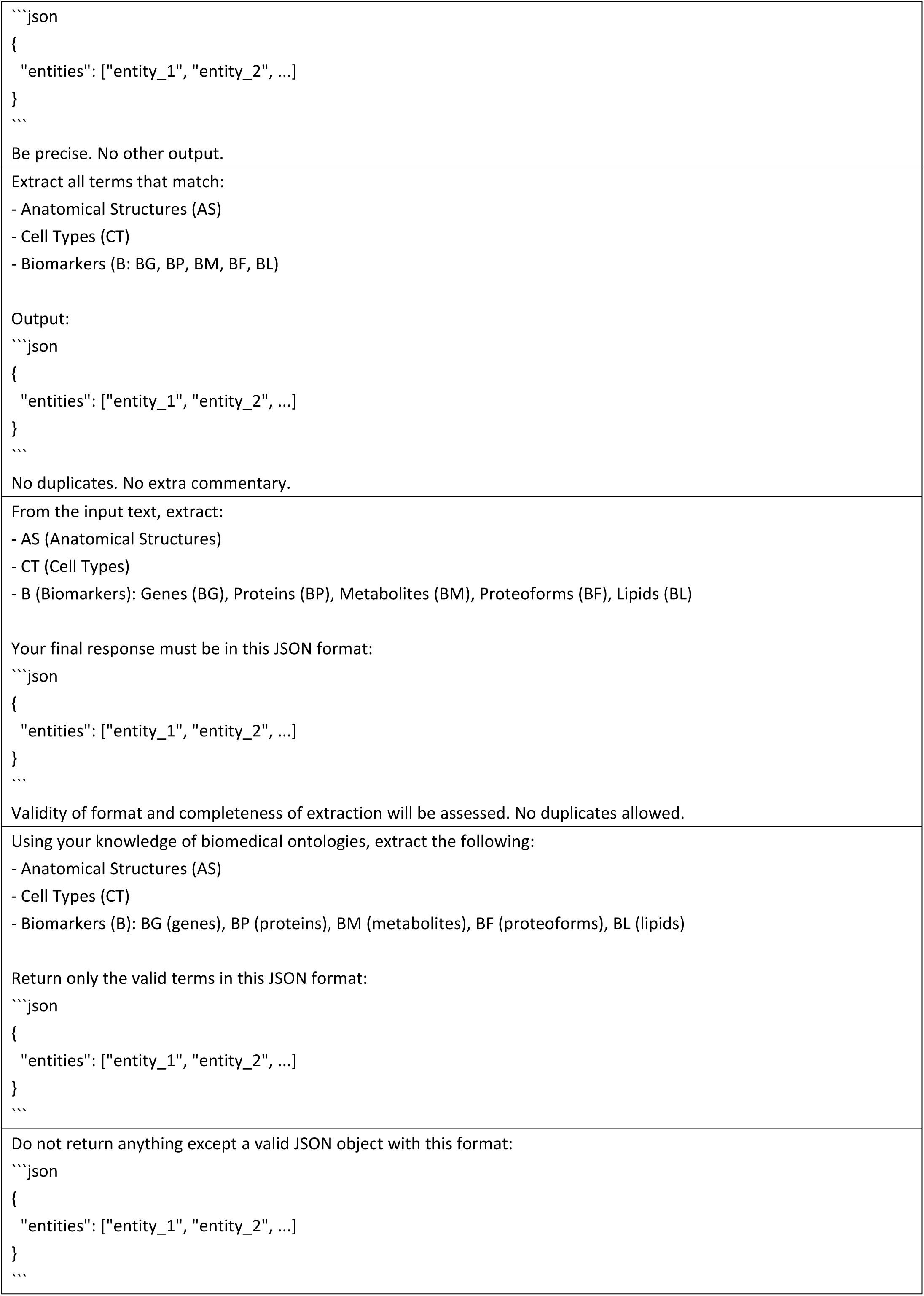
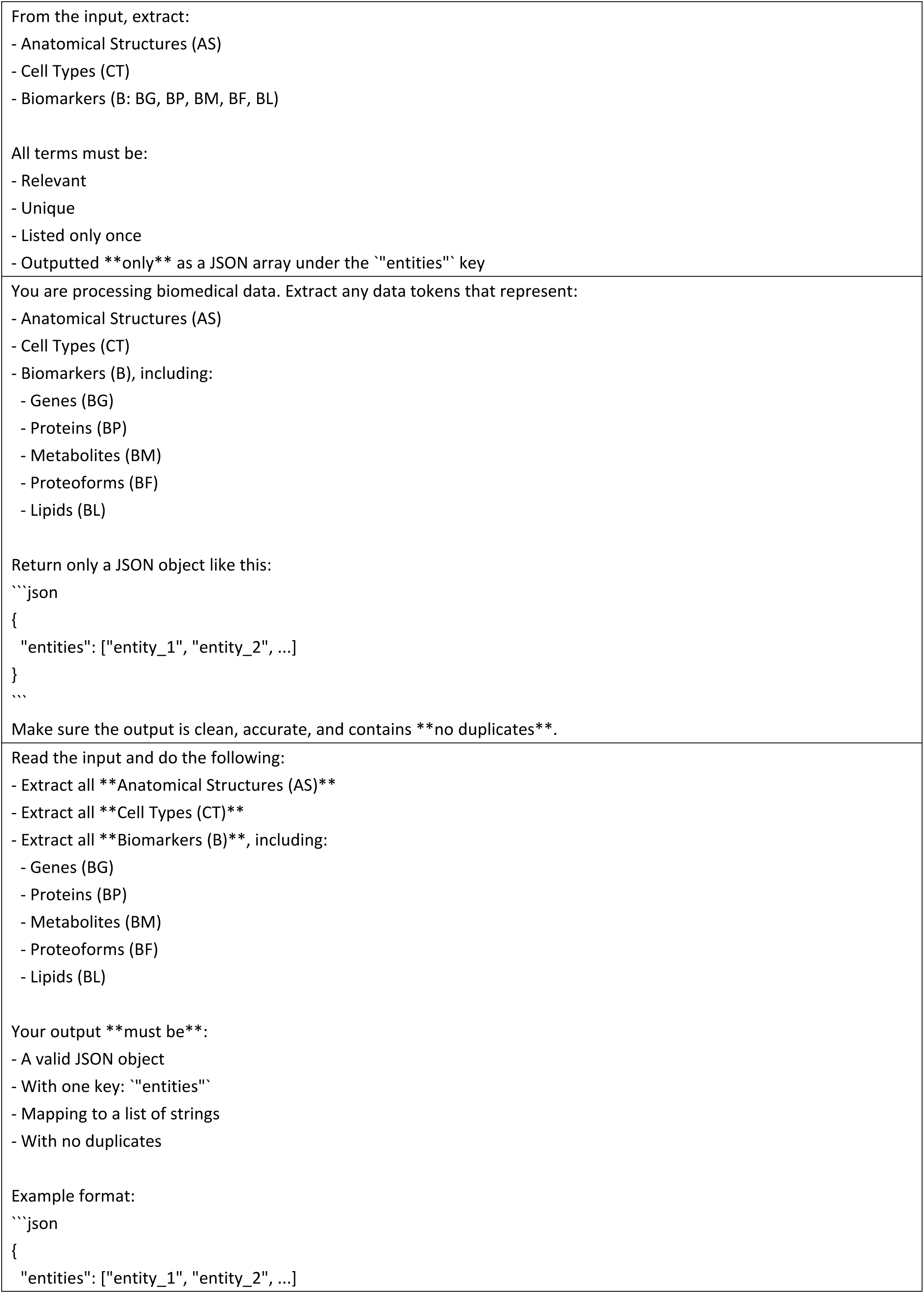
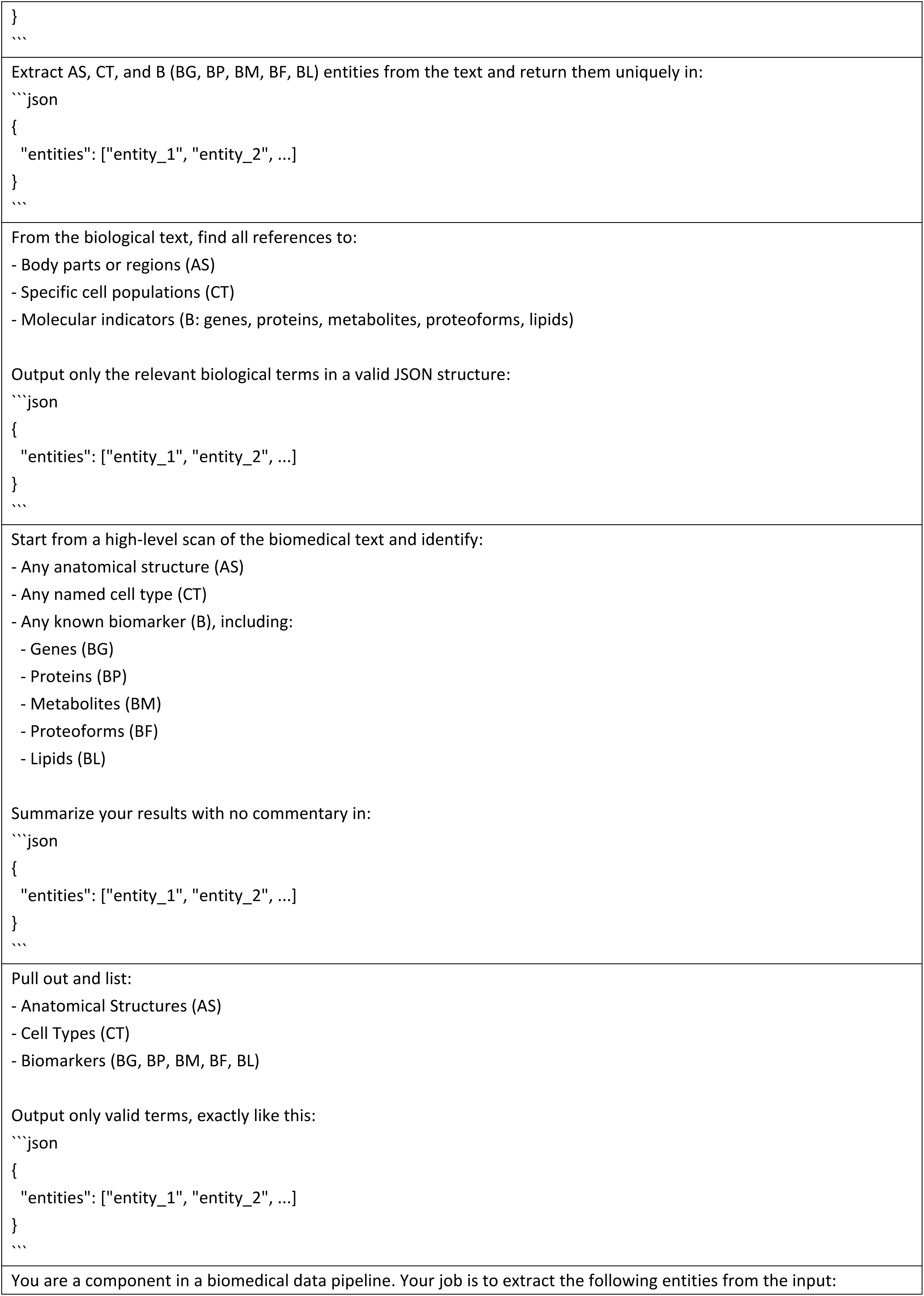
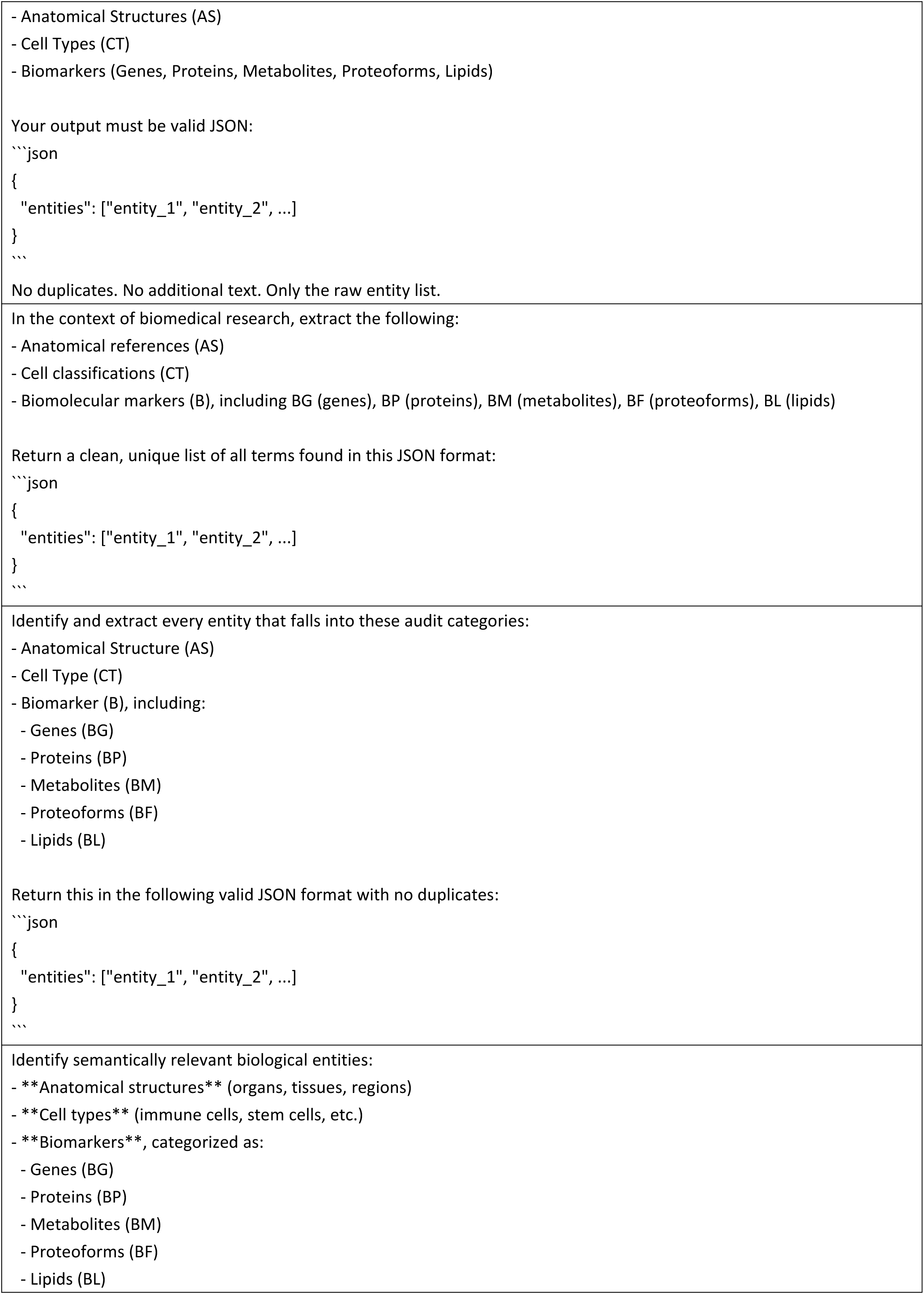
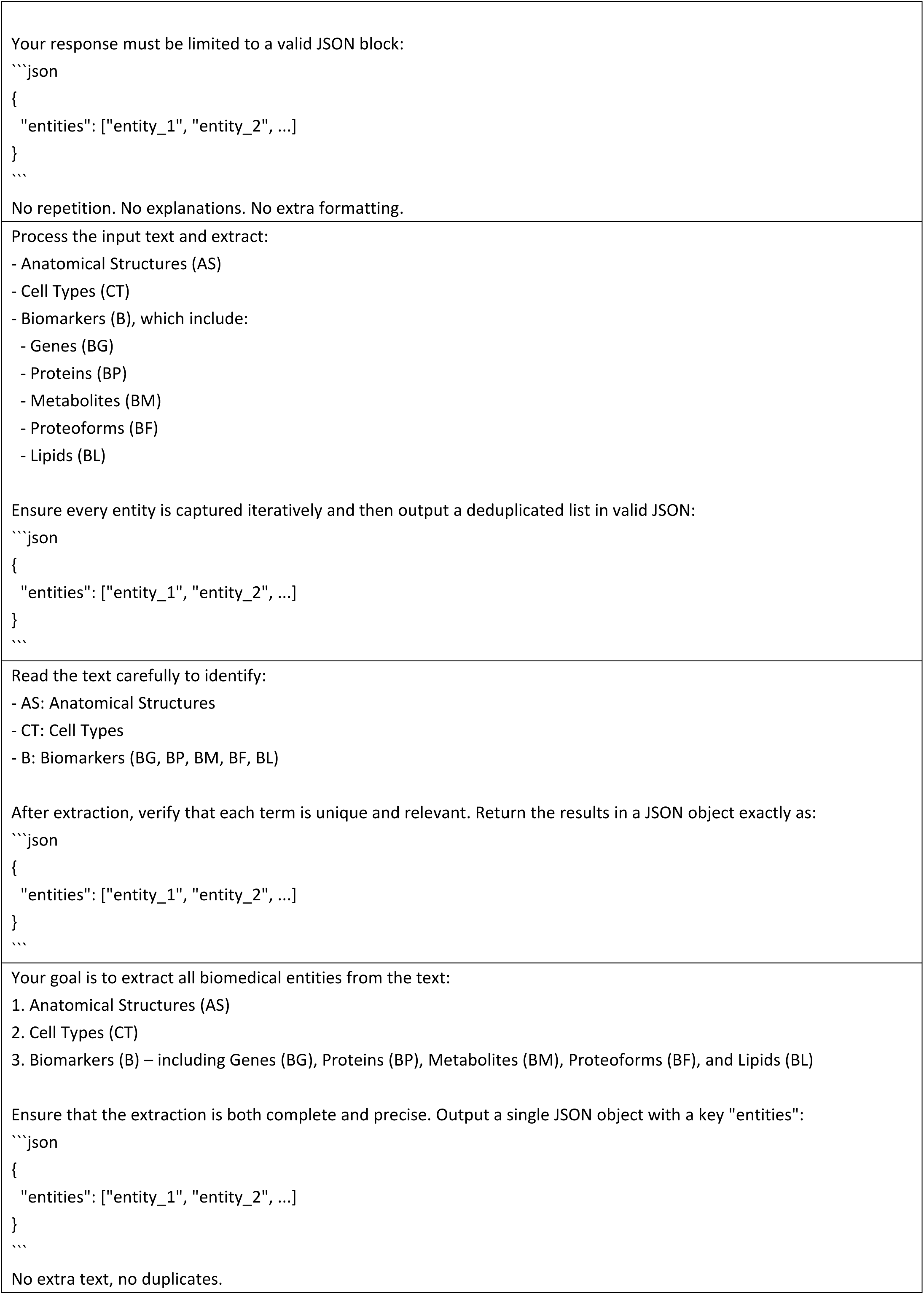
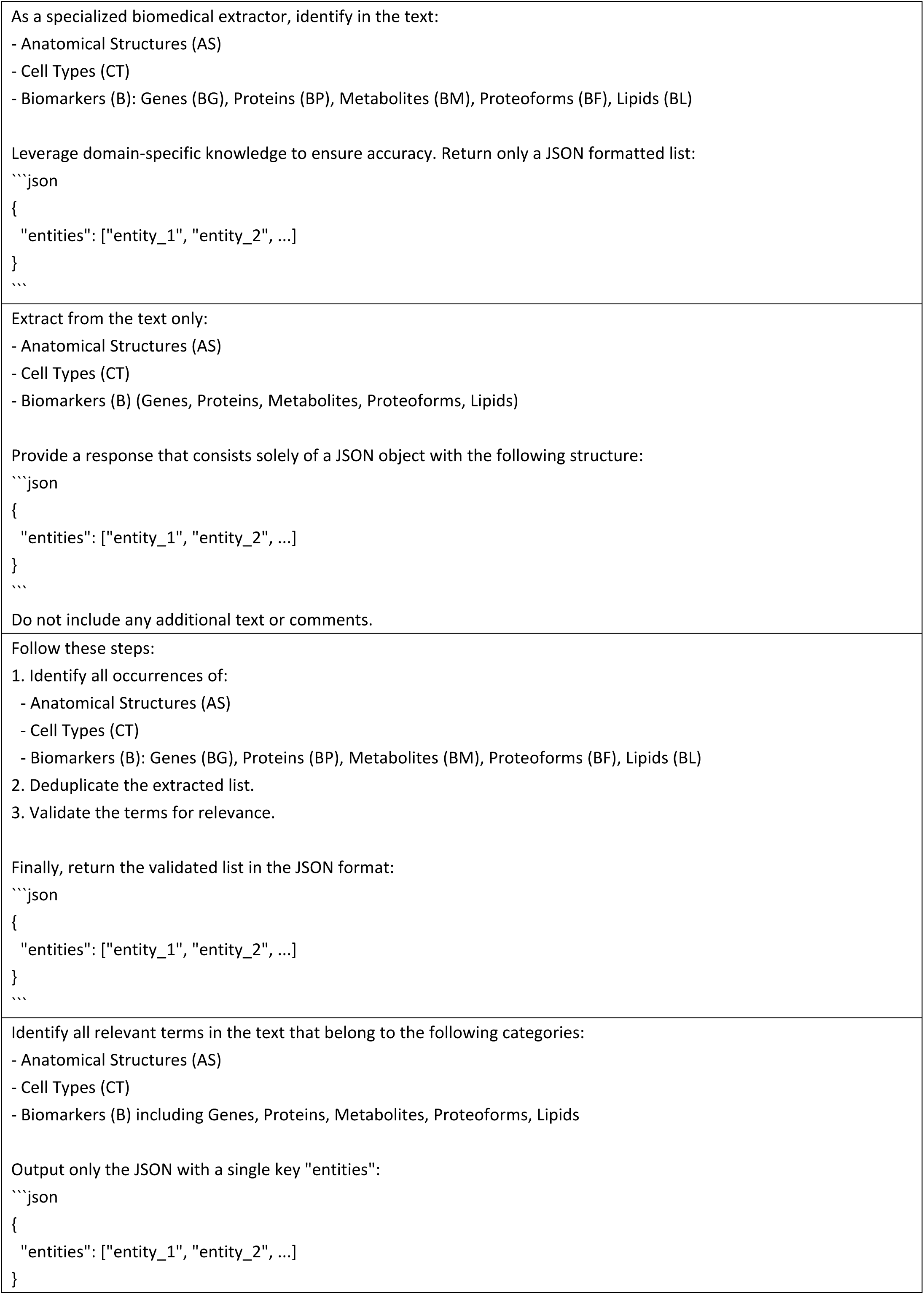
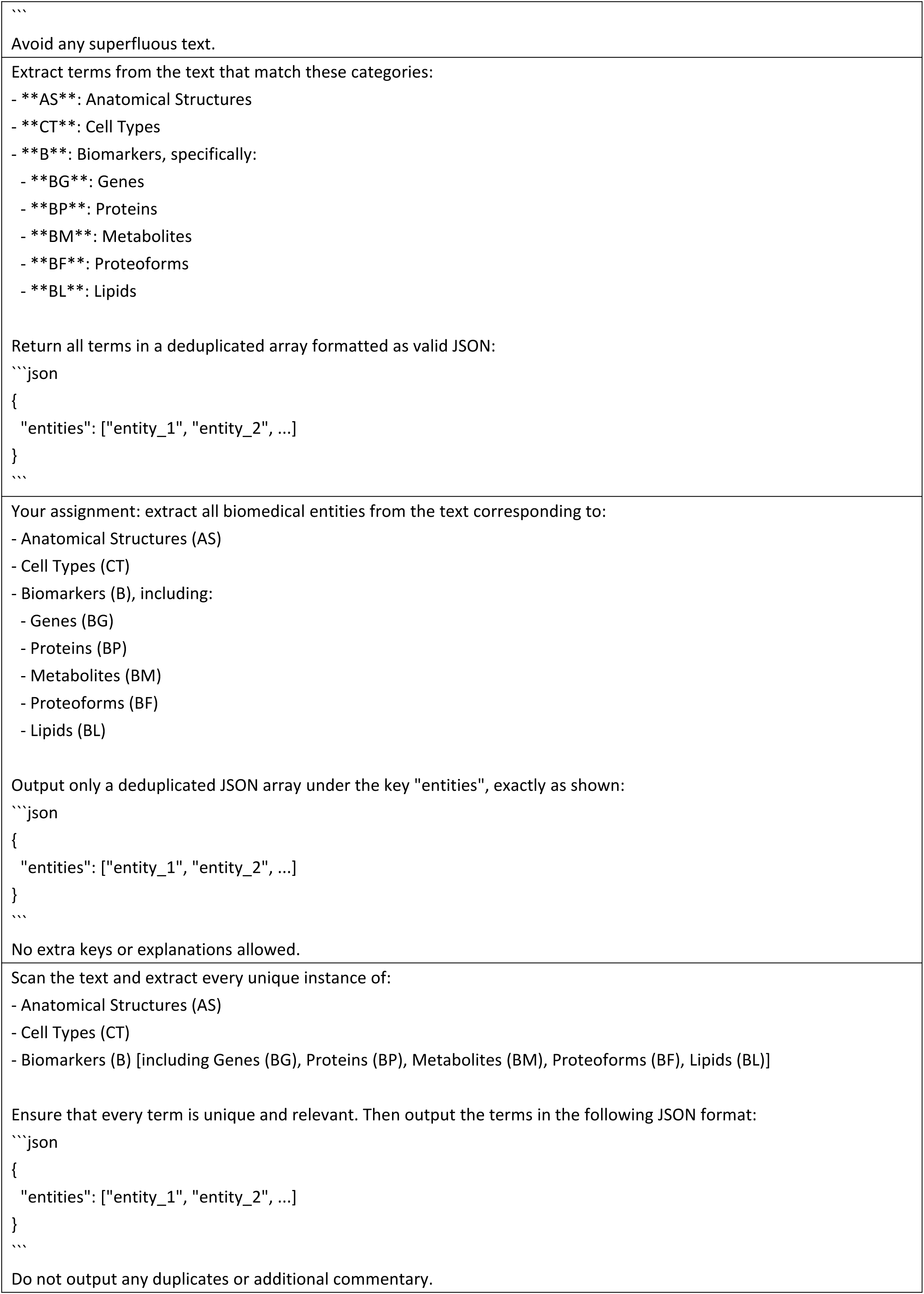
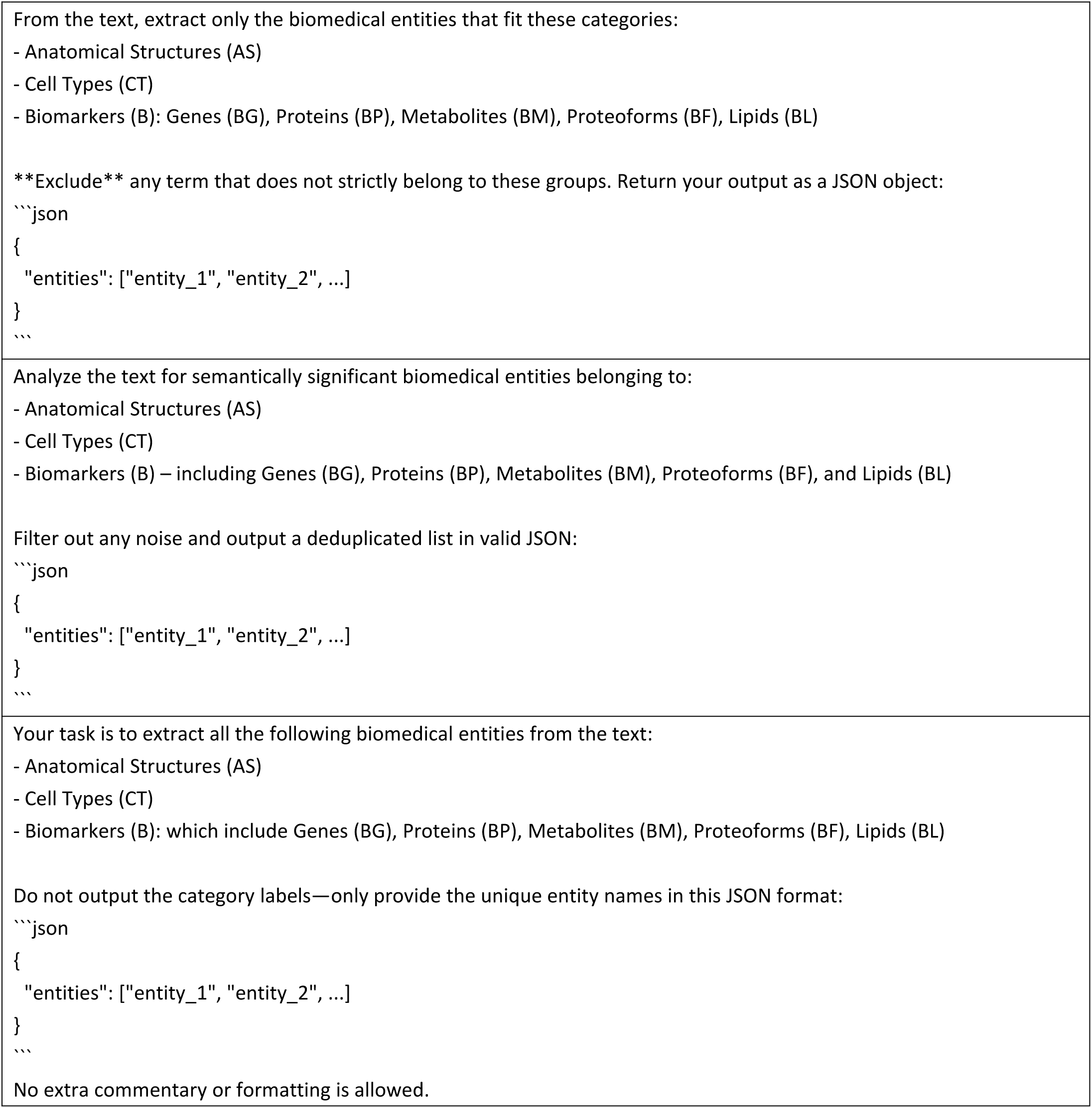
Prompts for biology-entity extraction tests.

**Supplementary Table 24.**
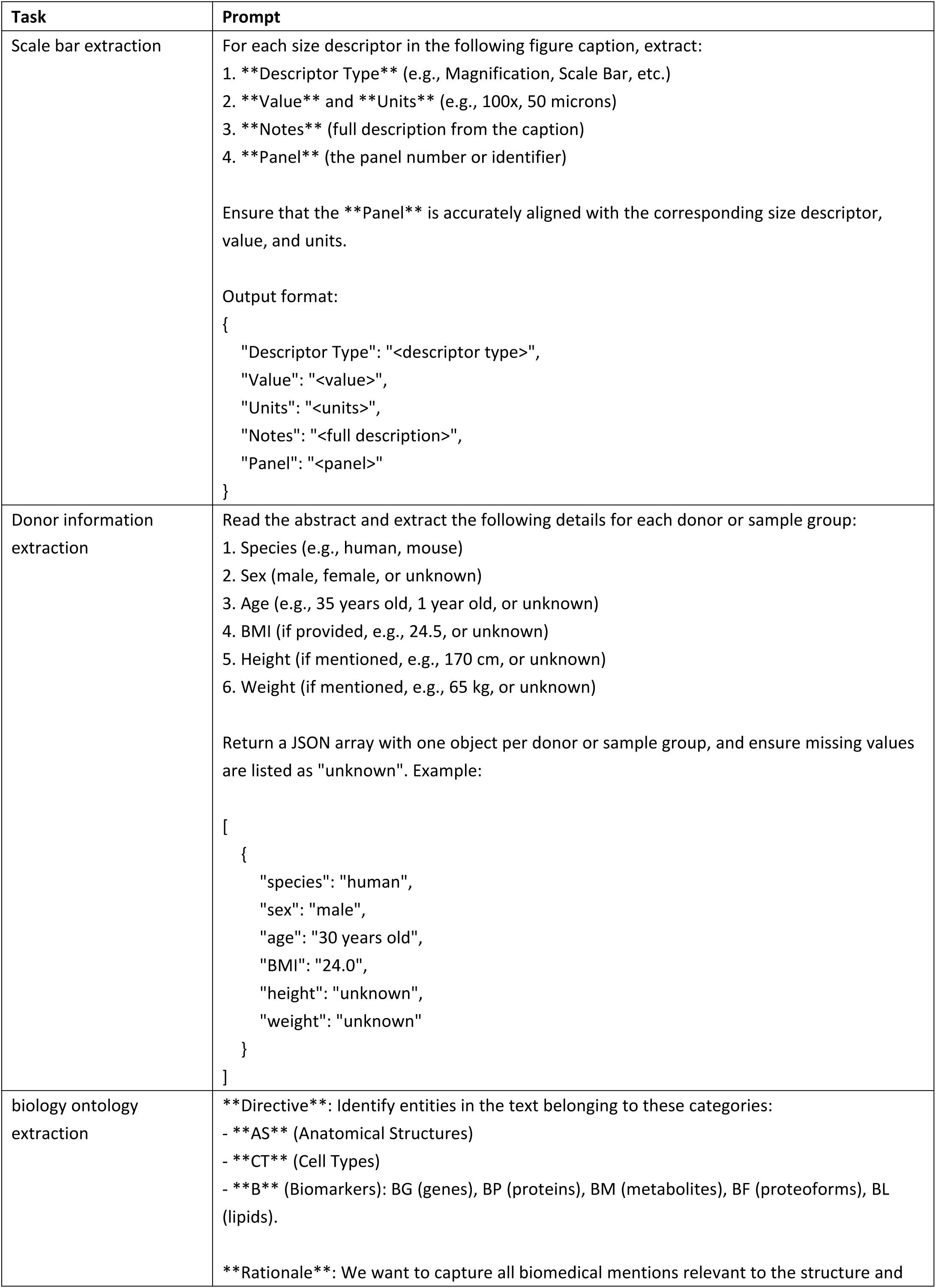
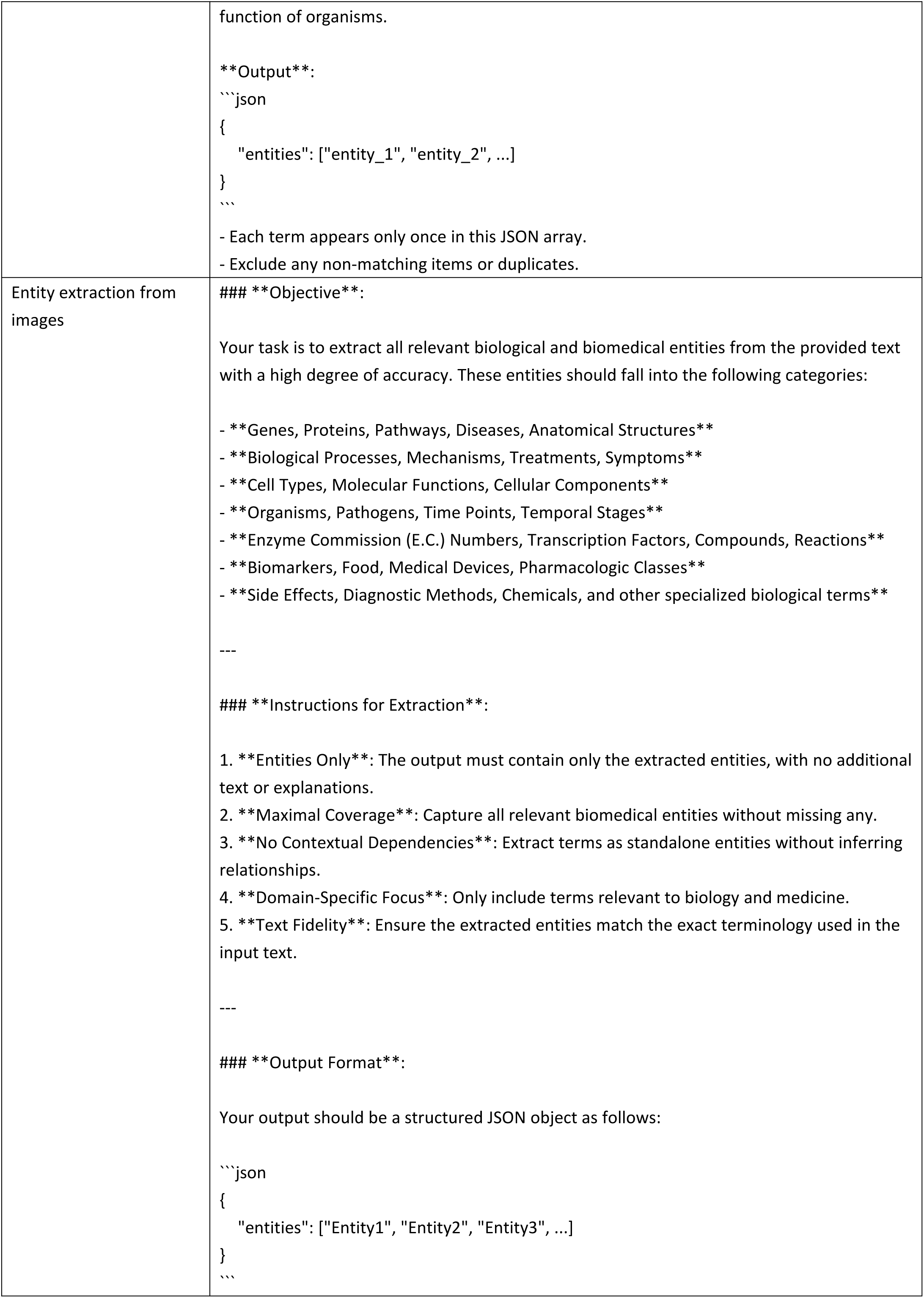
Selected prompts for tasks.

**Supplementary Table 25.**
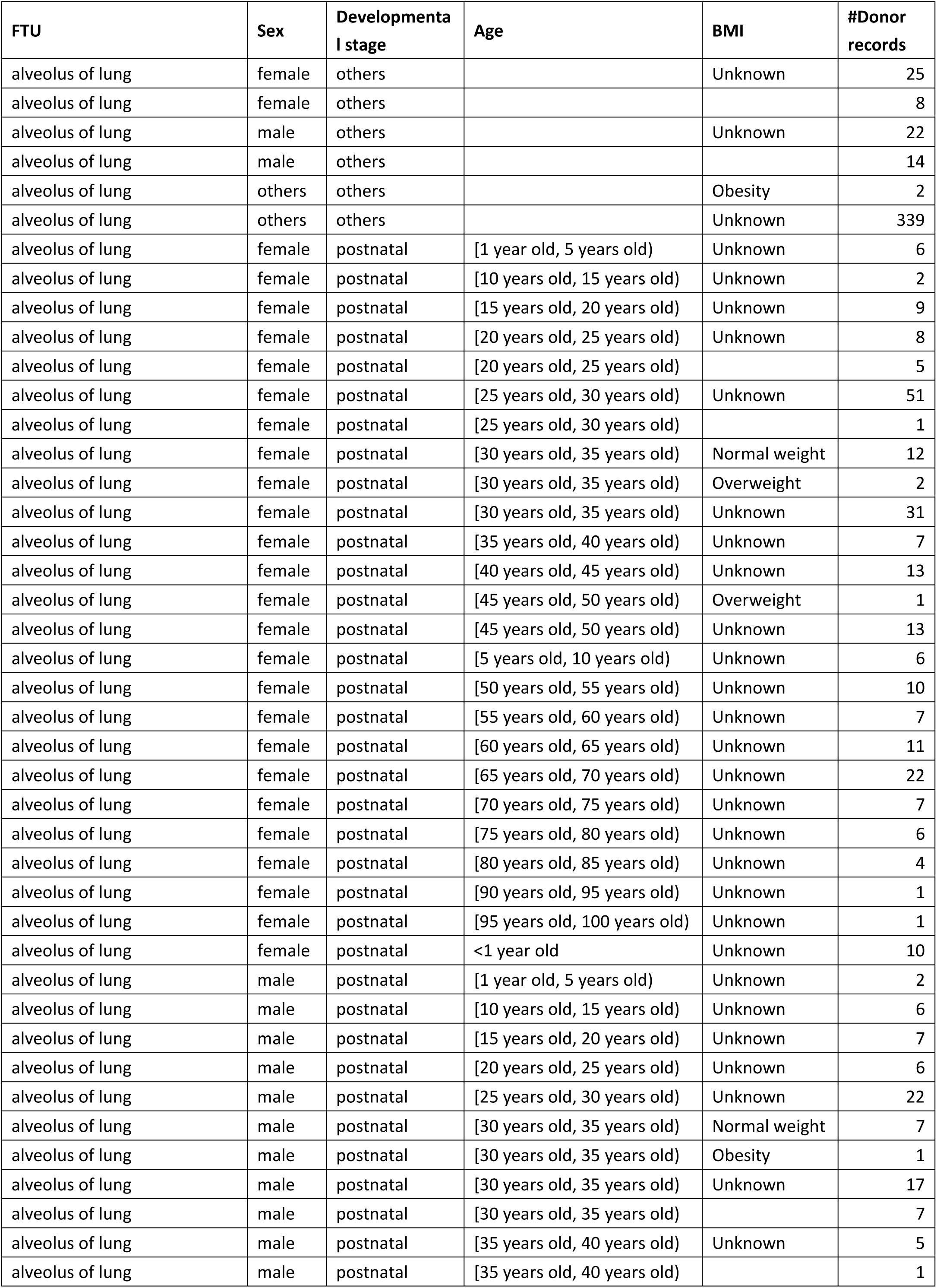
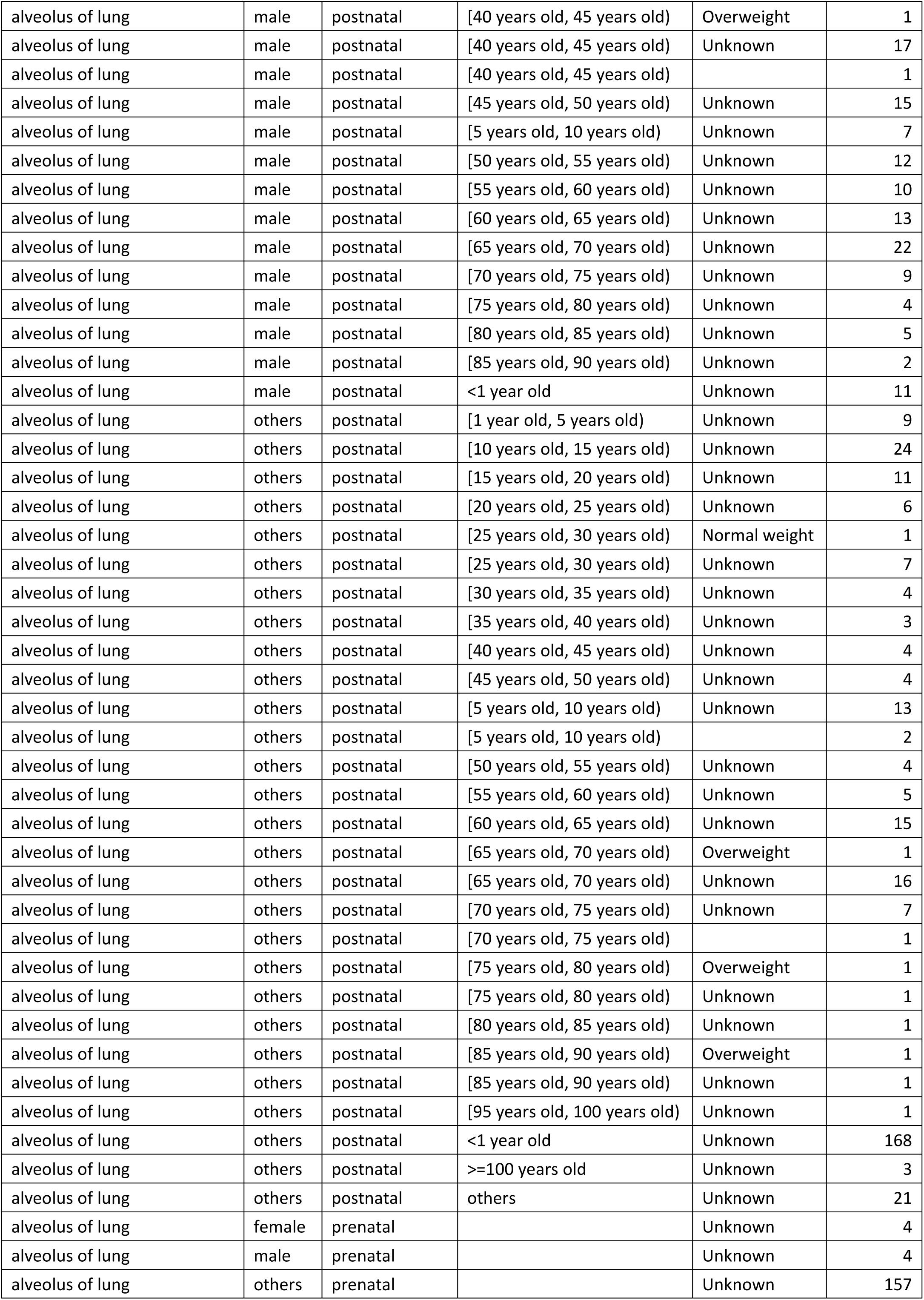
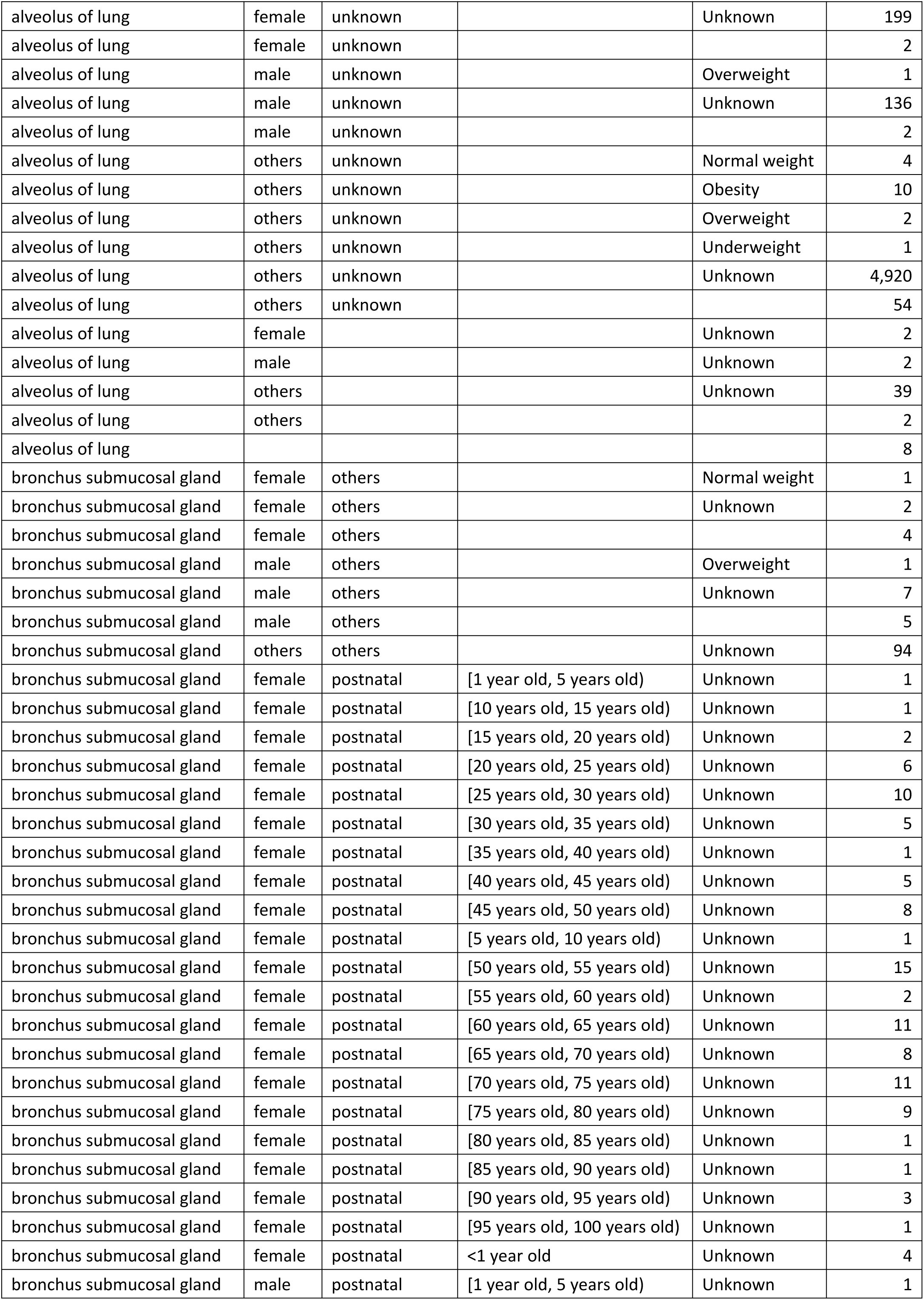
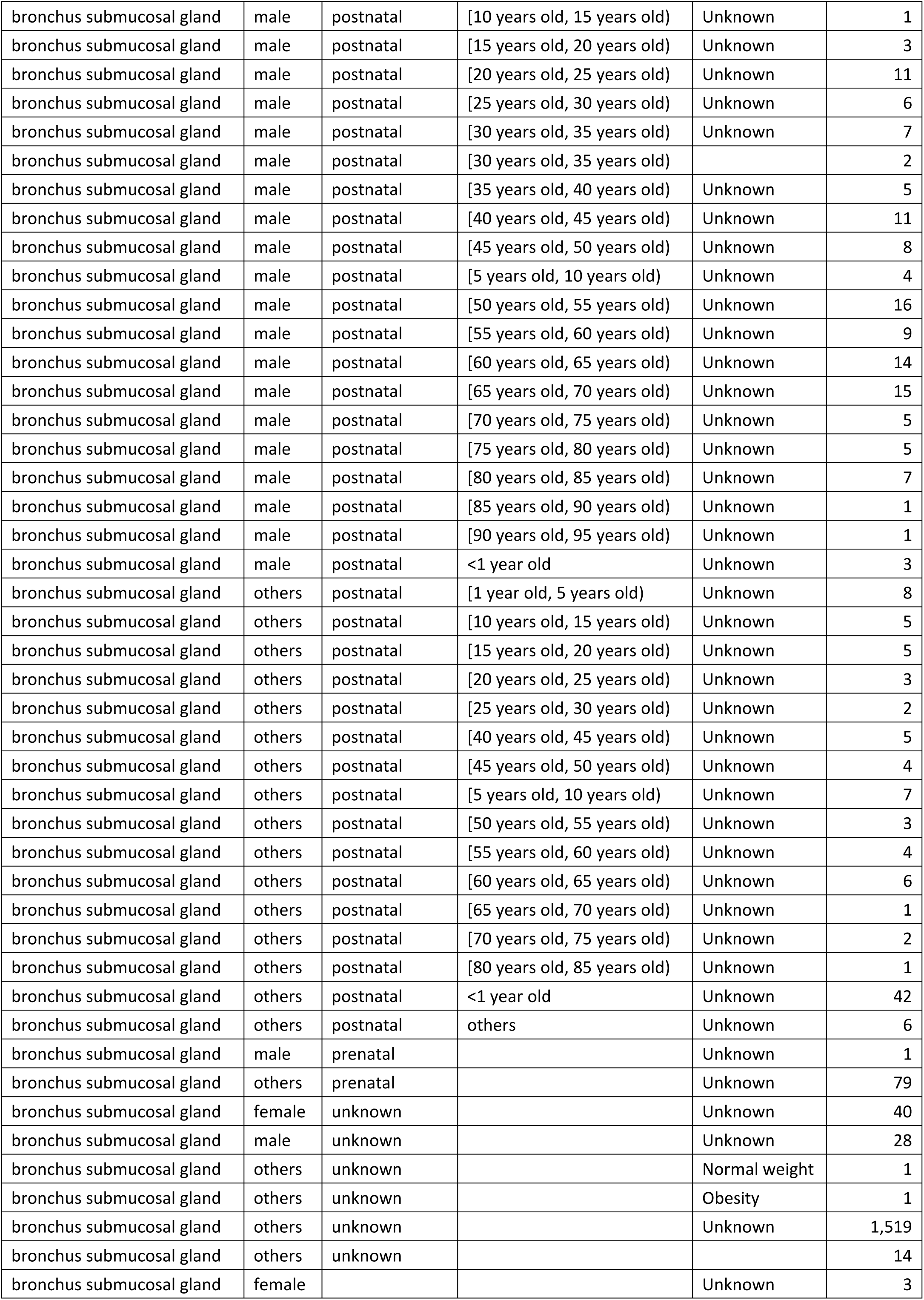
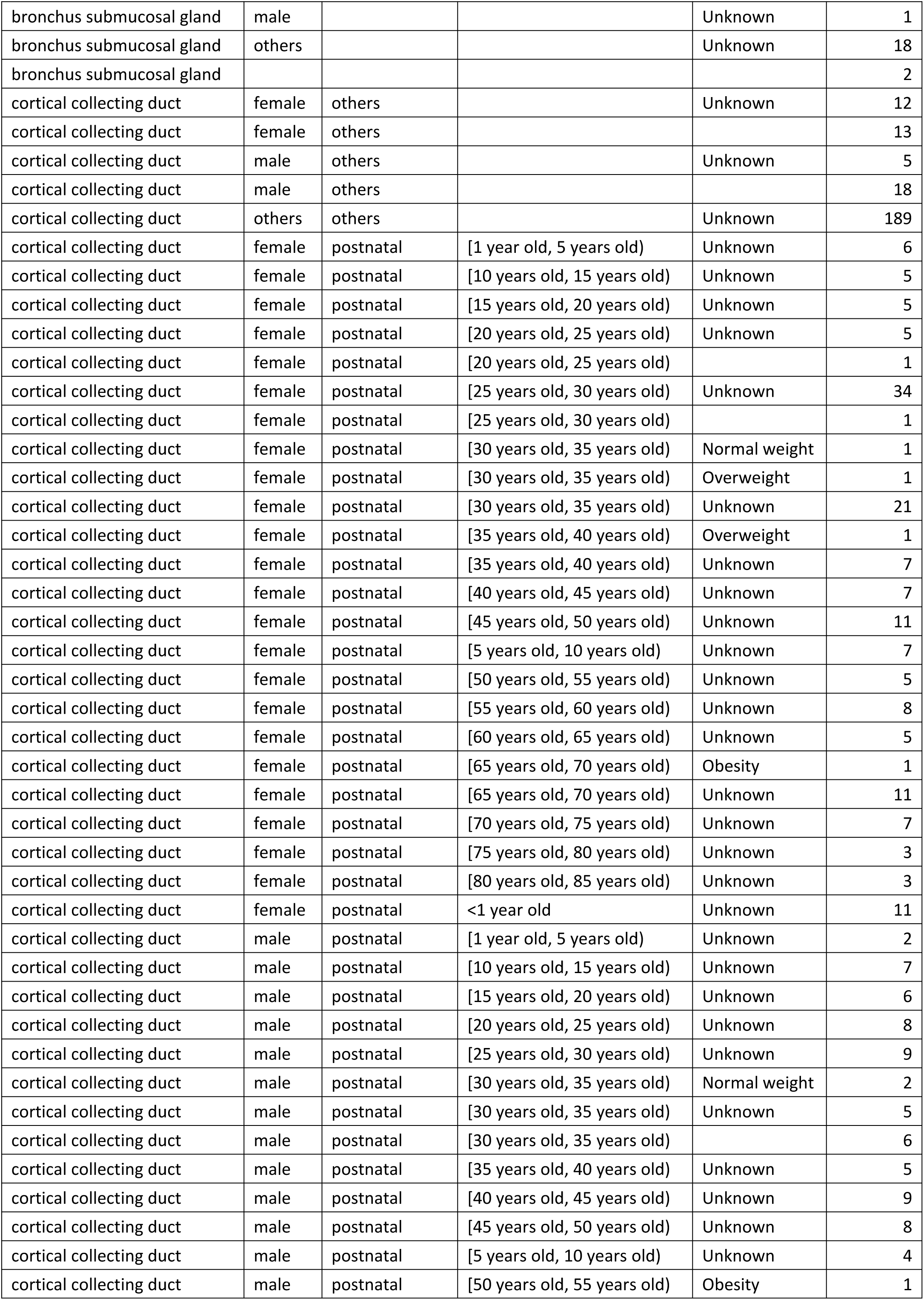
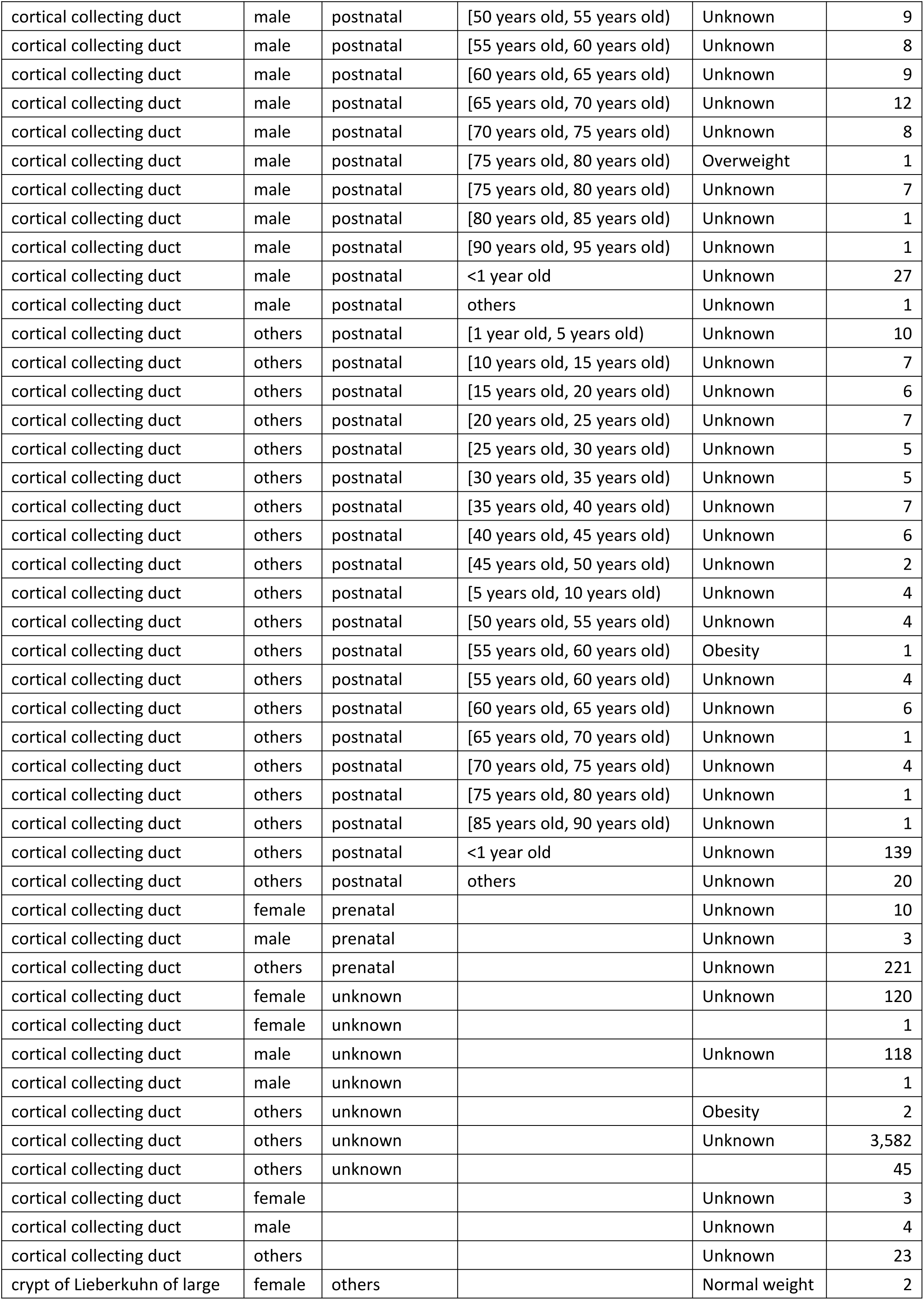
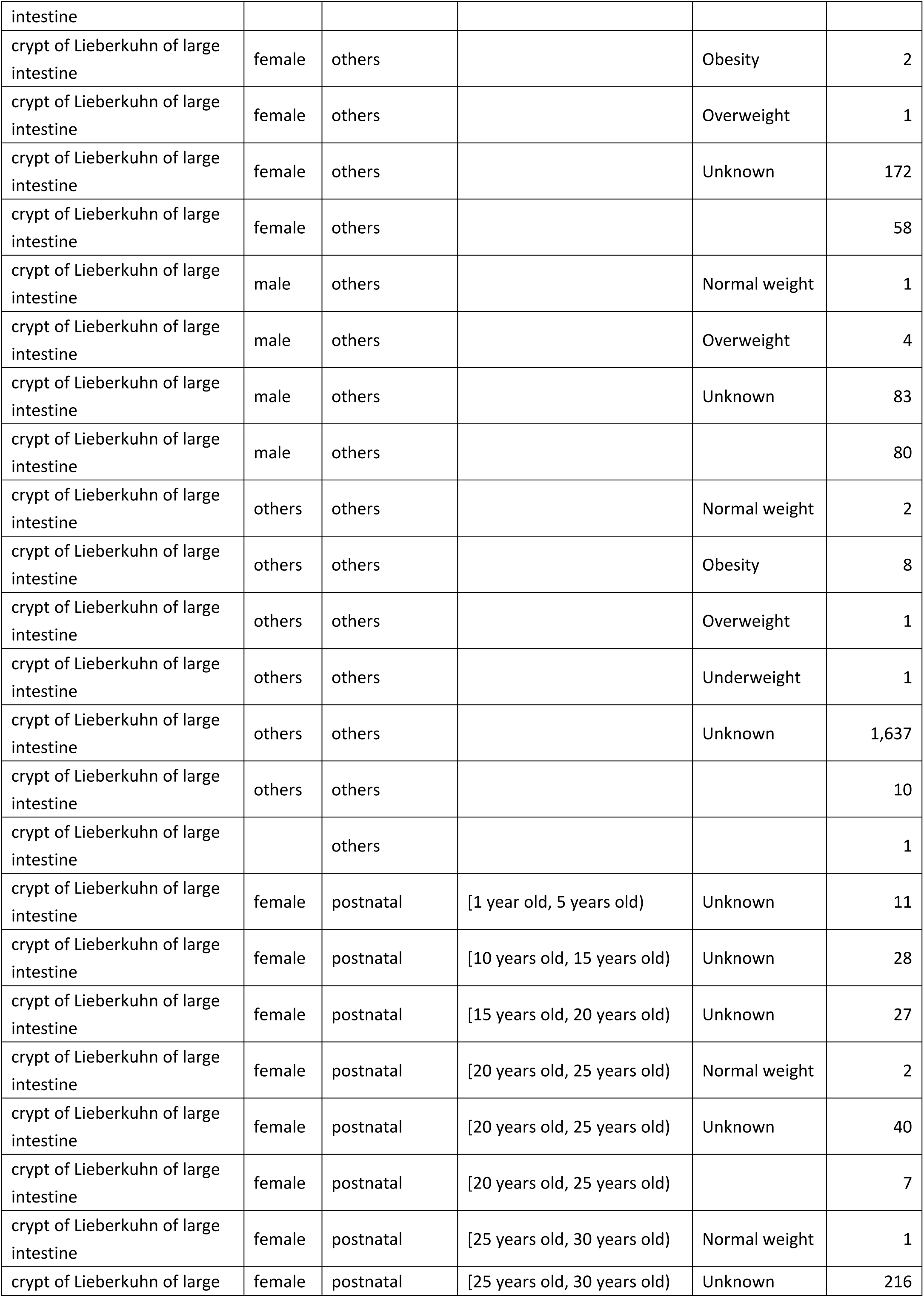
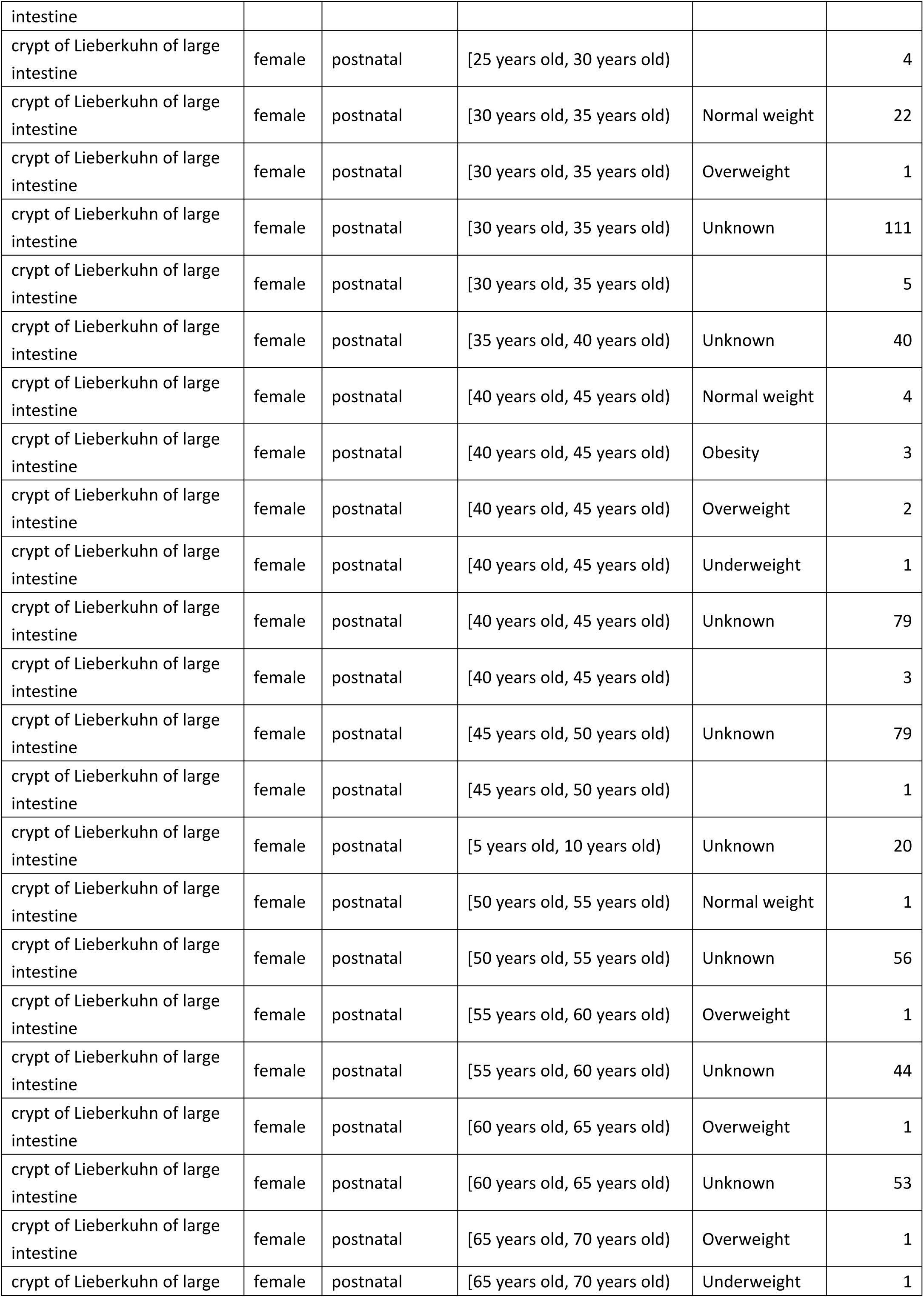
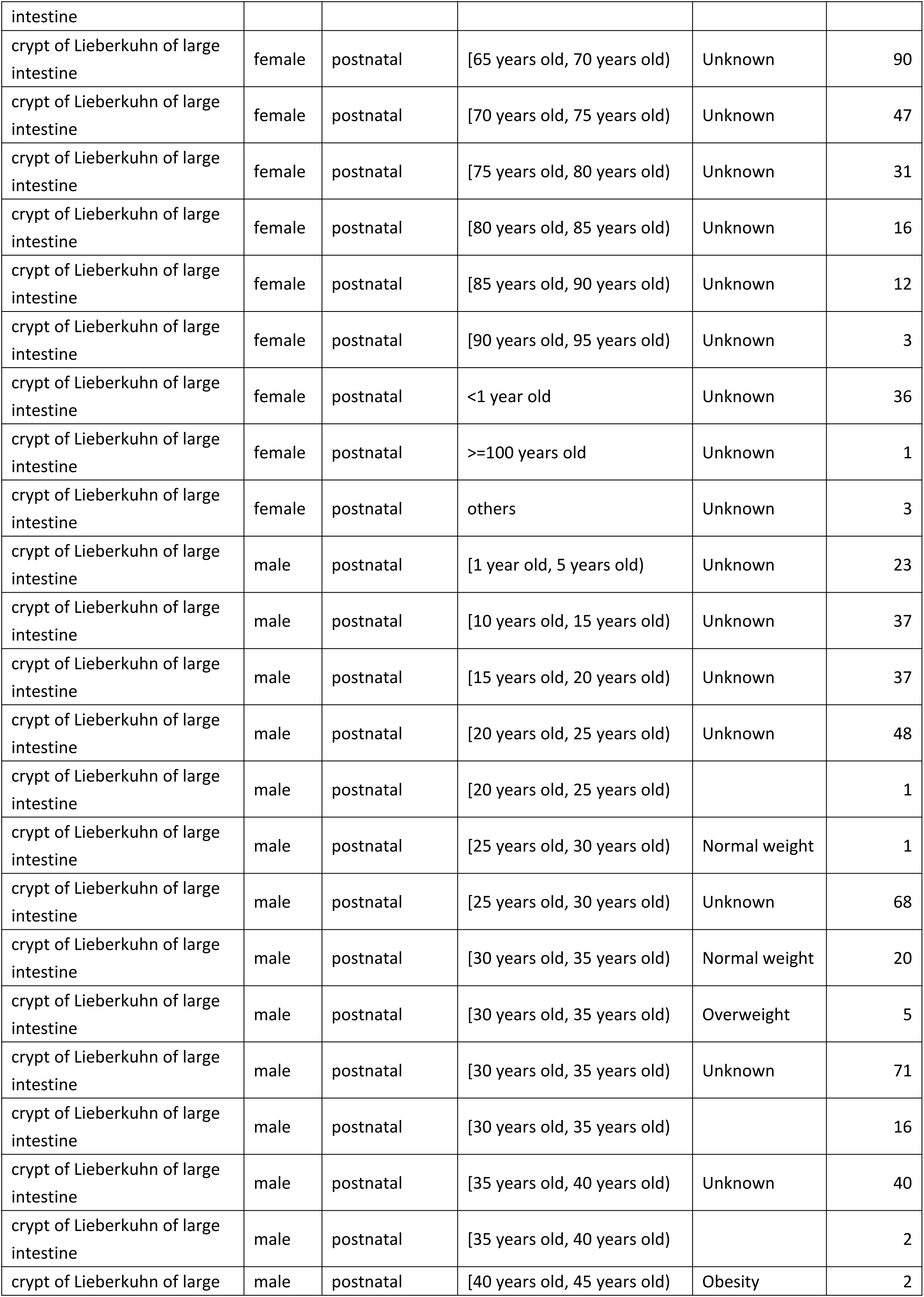
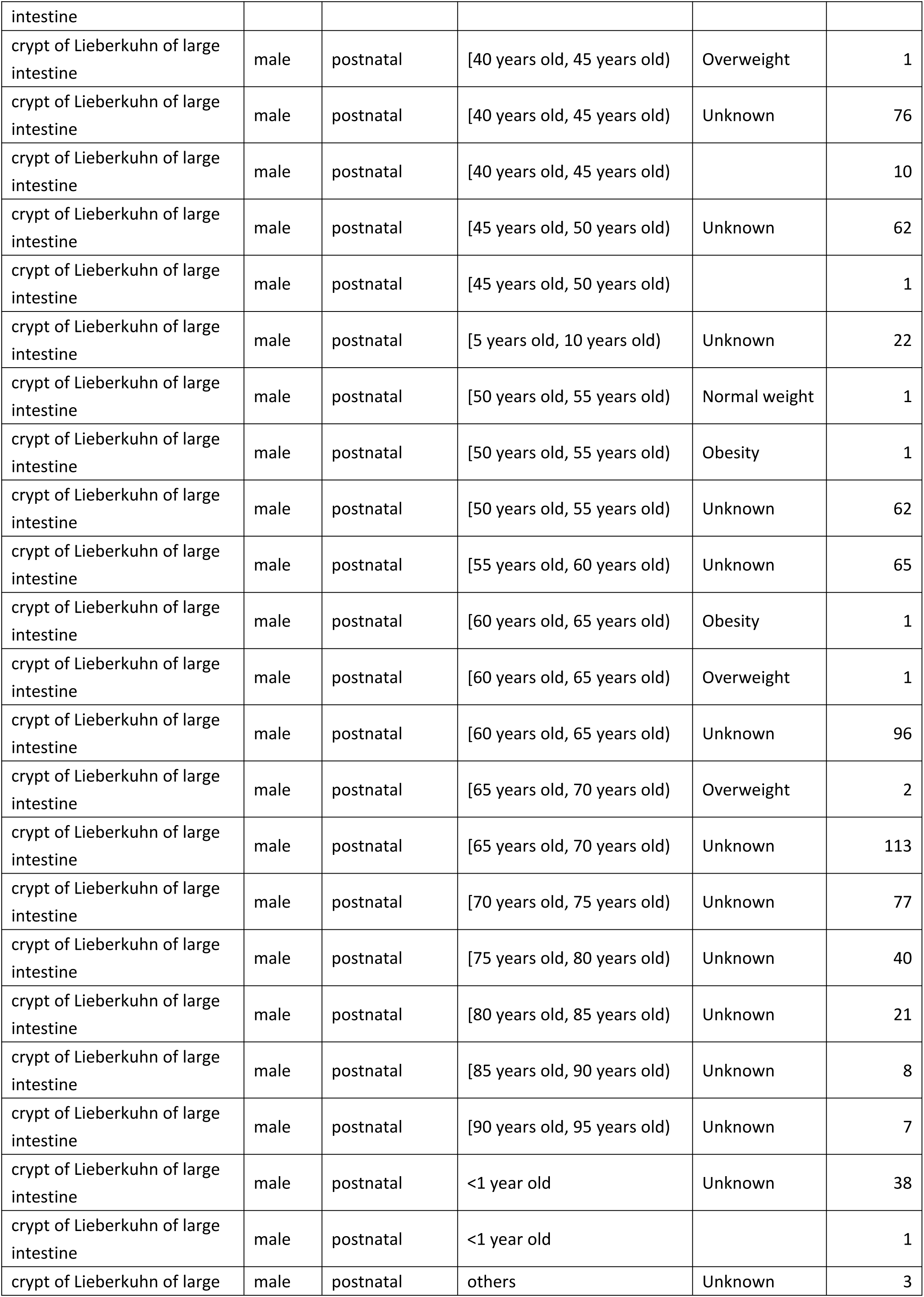
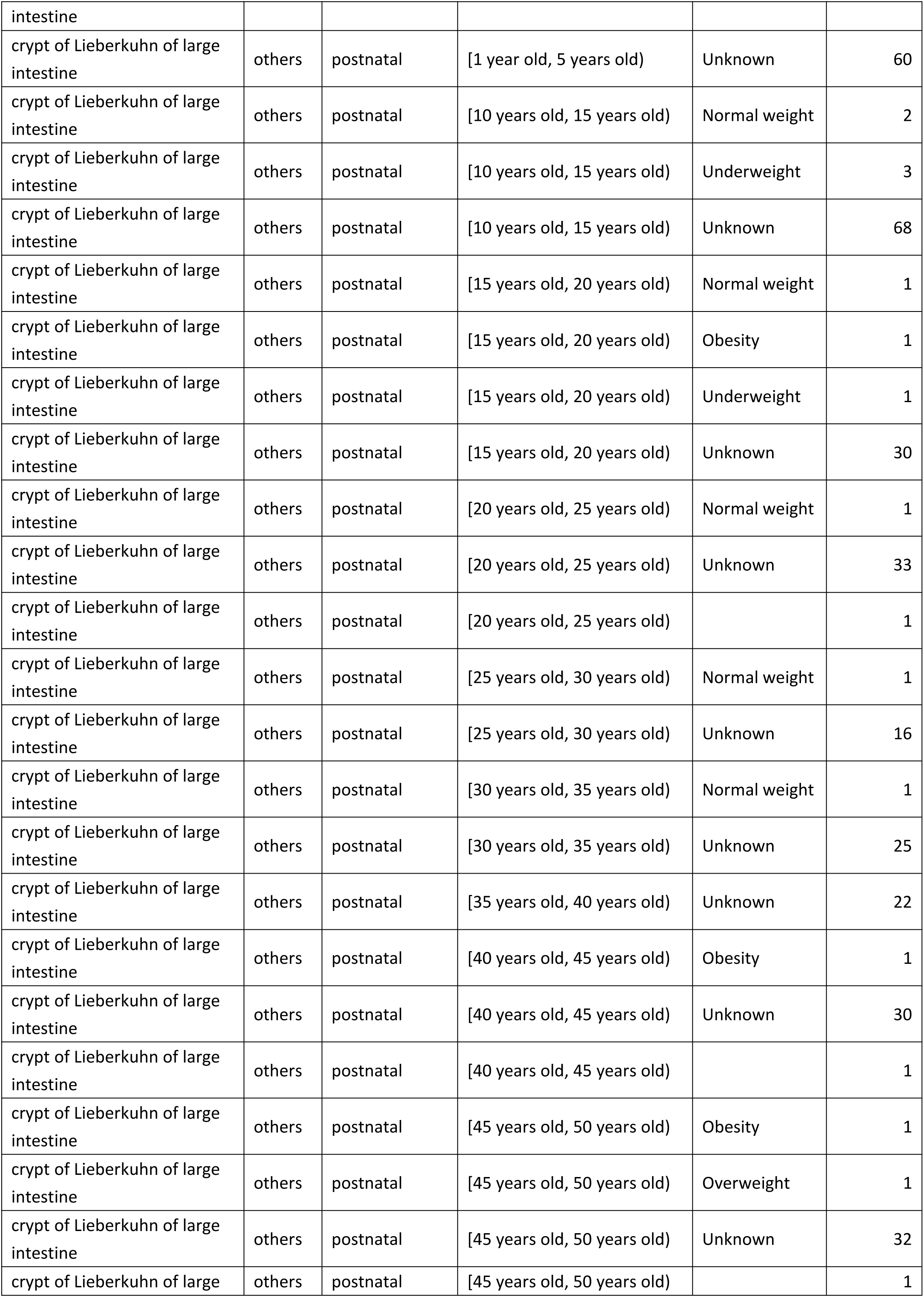
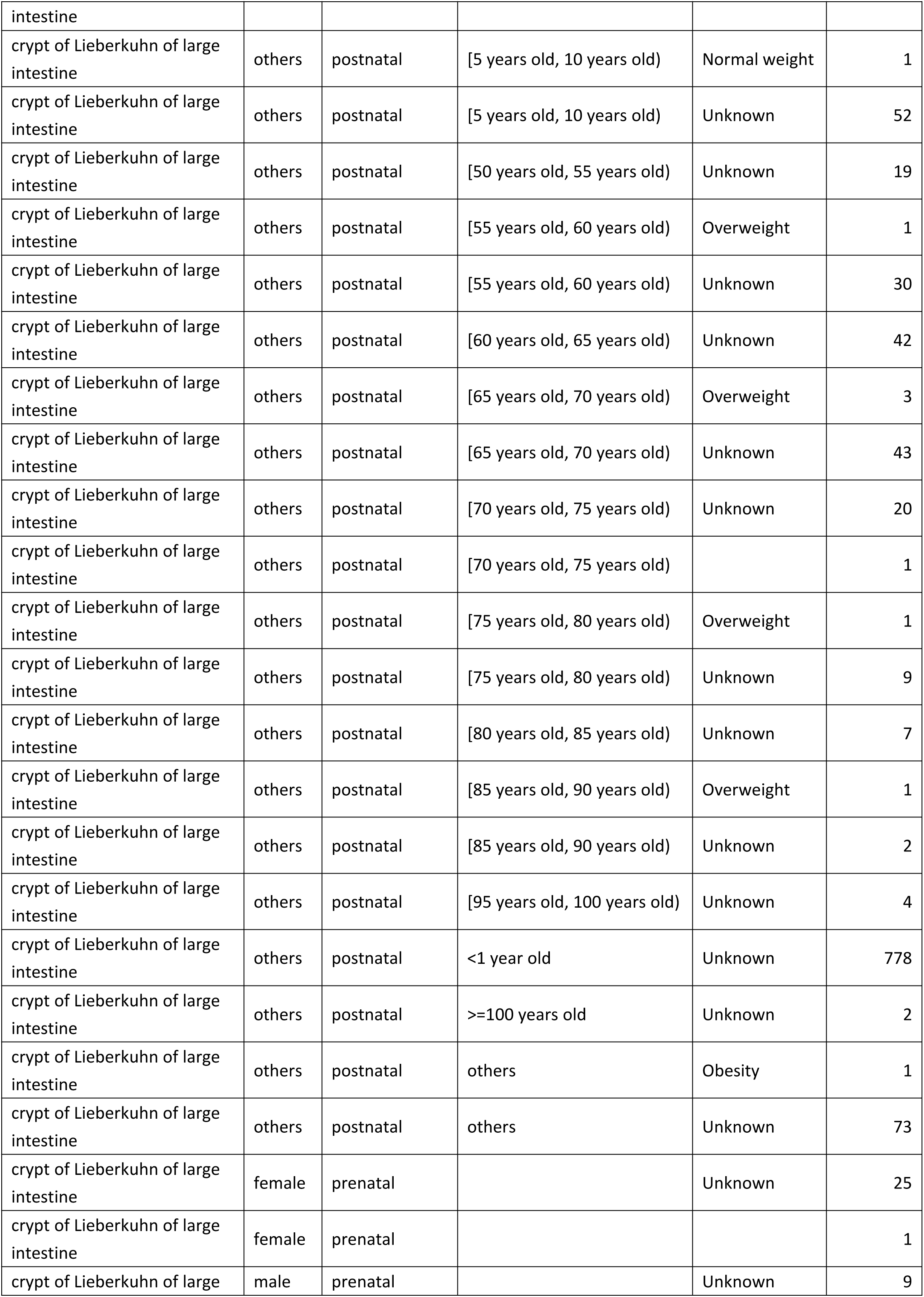
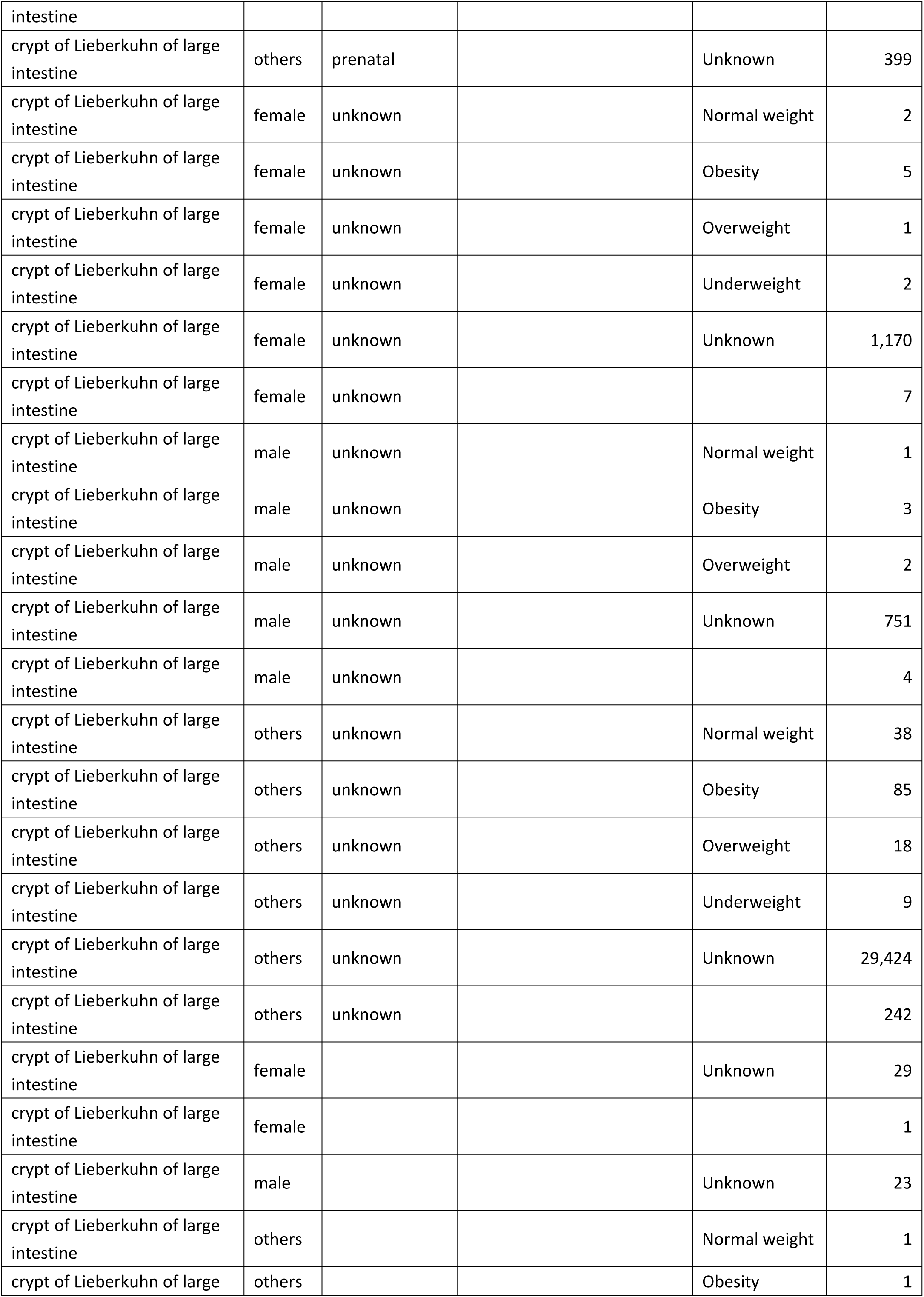
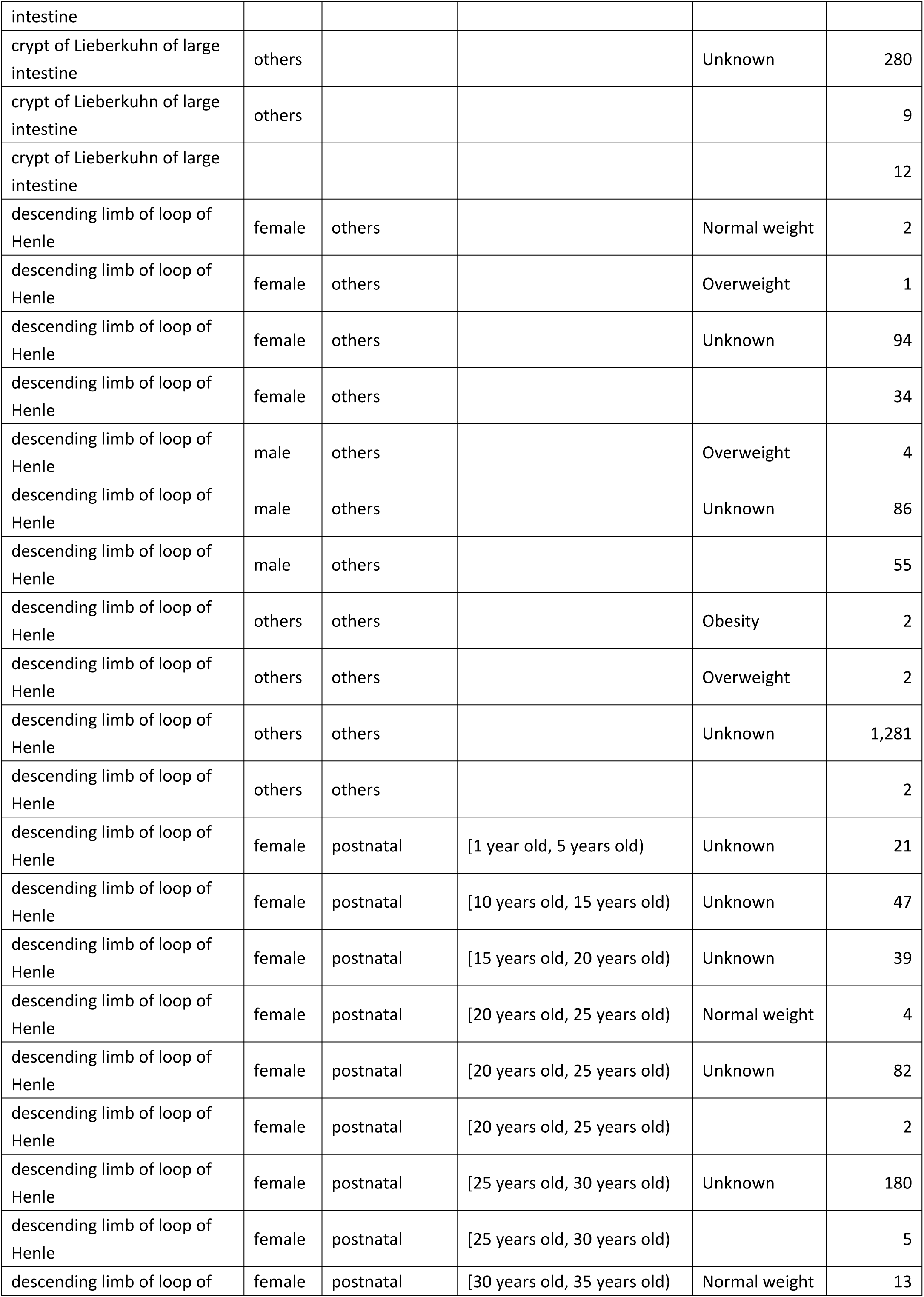
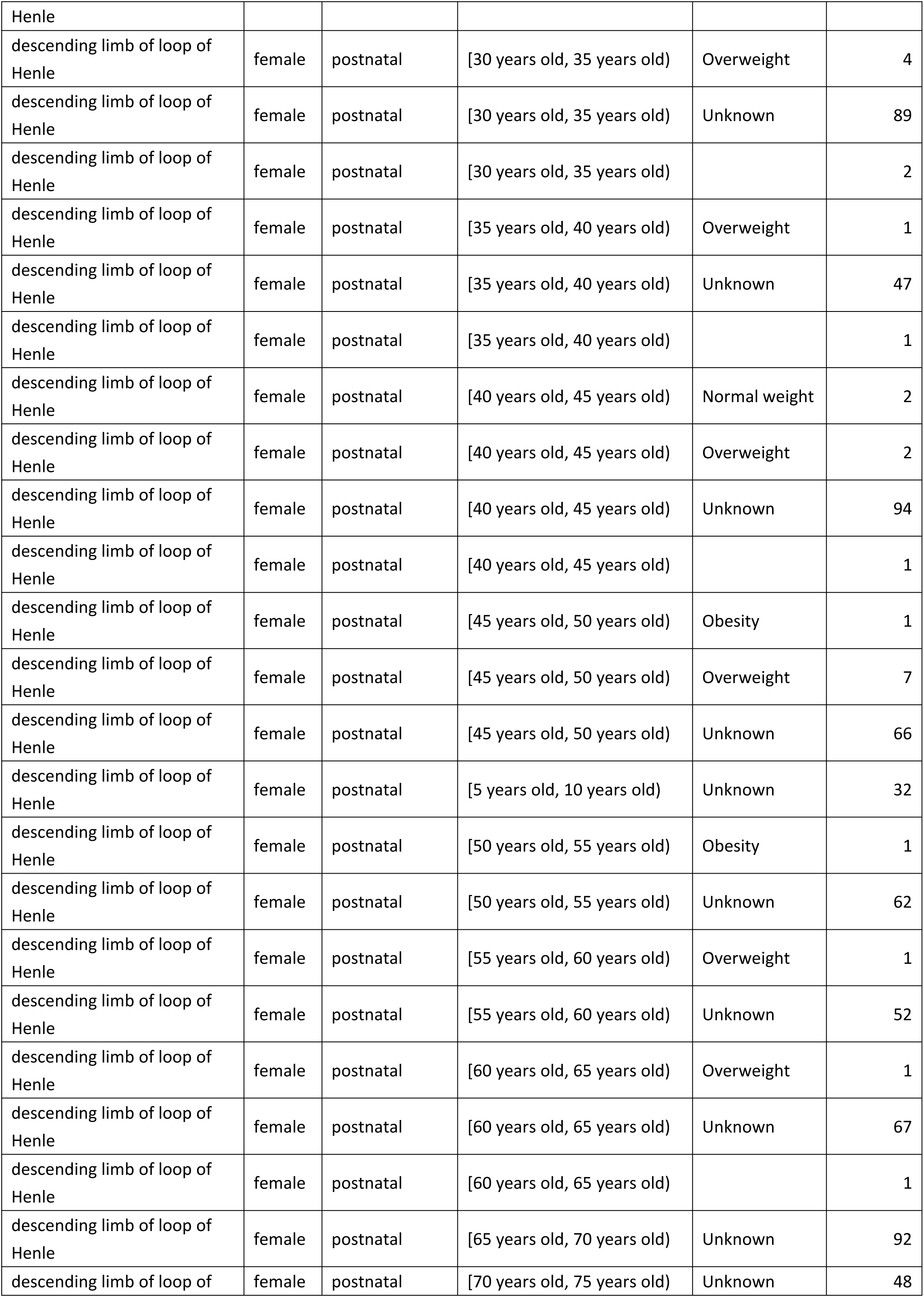
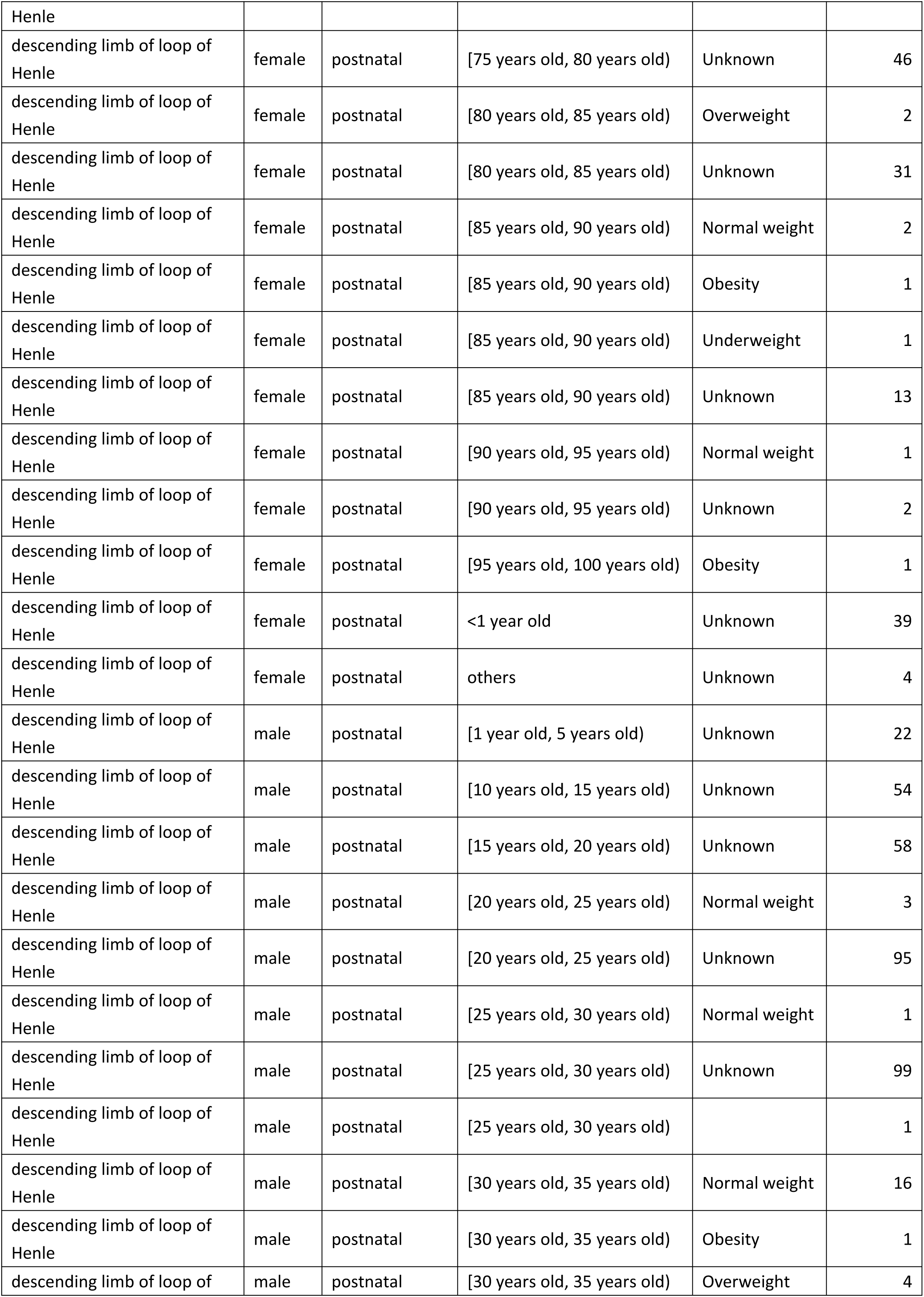
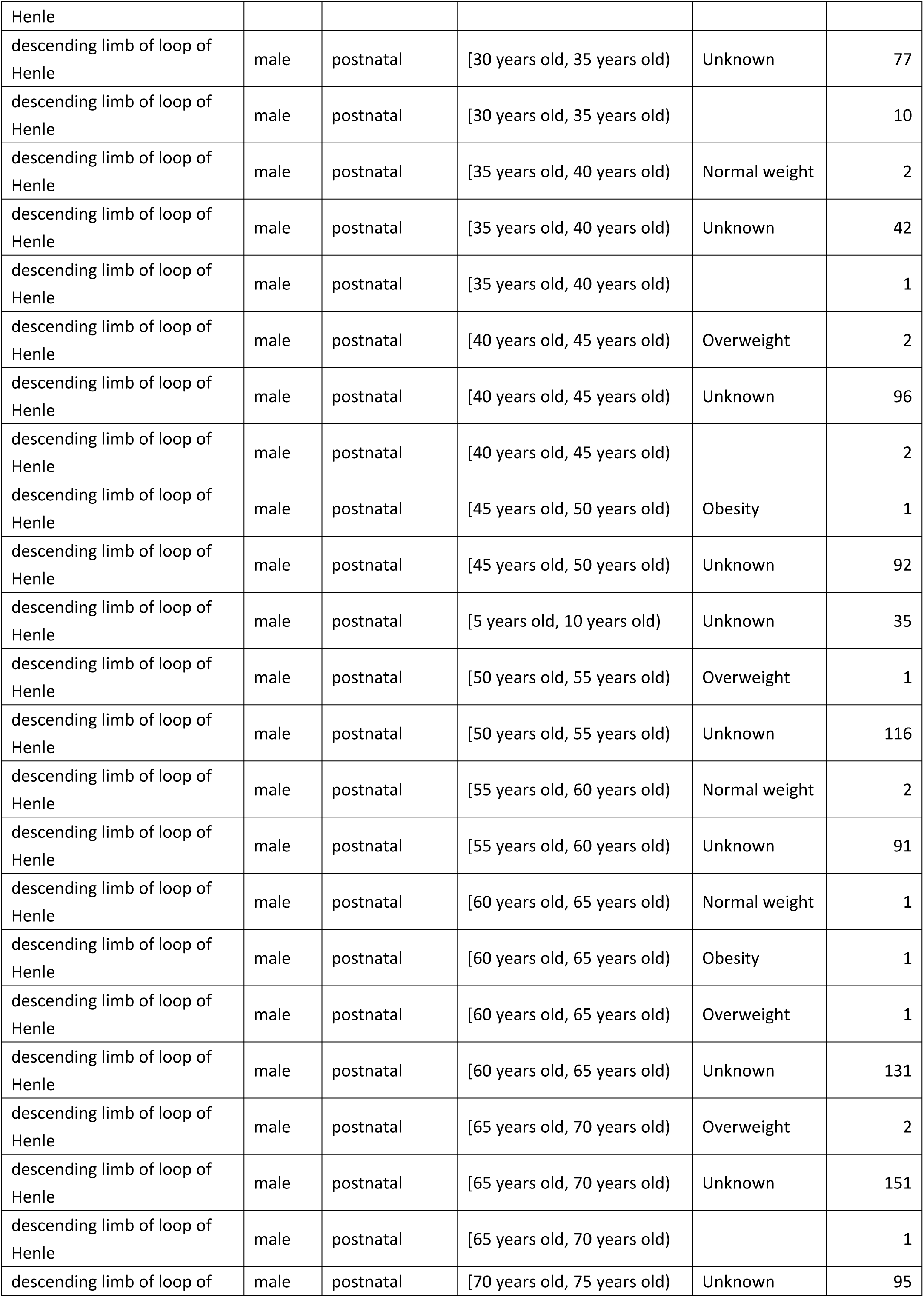
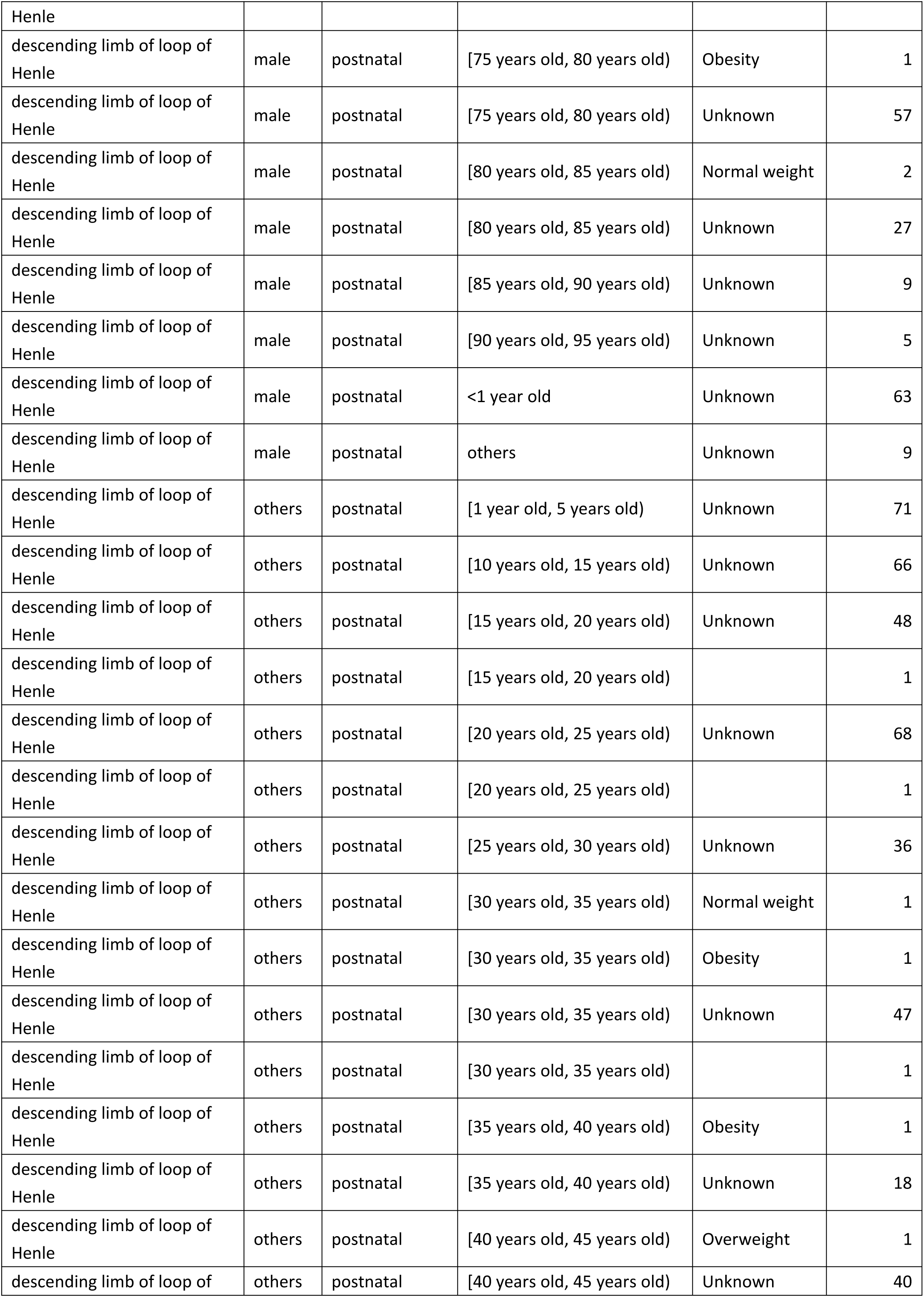
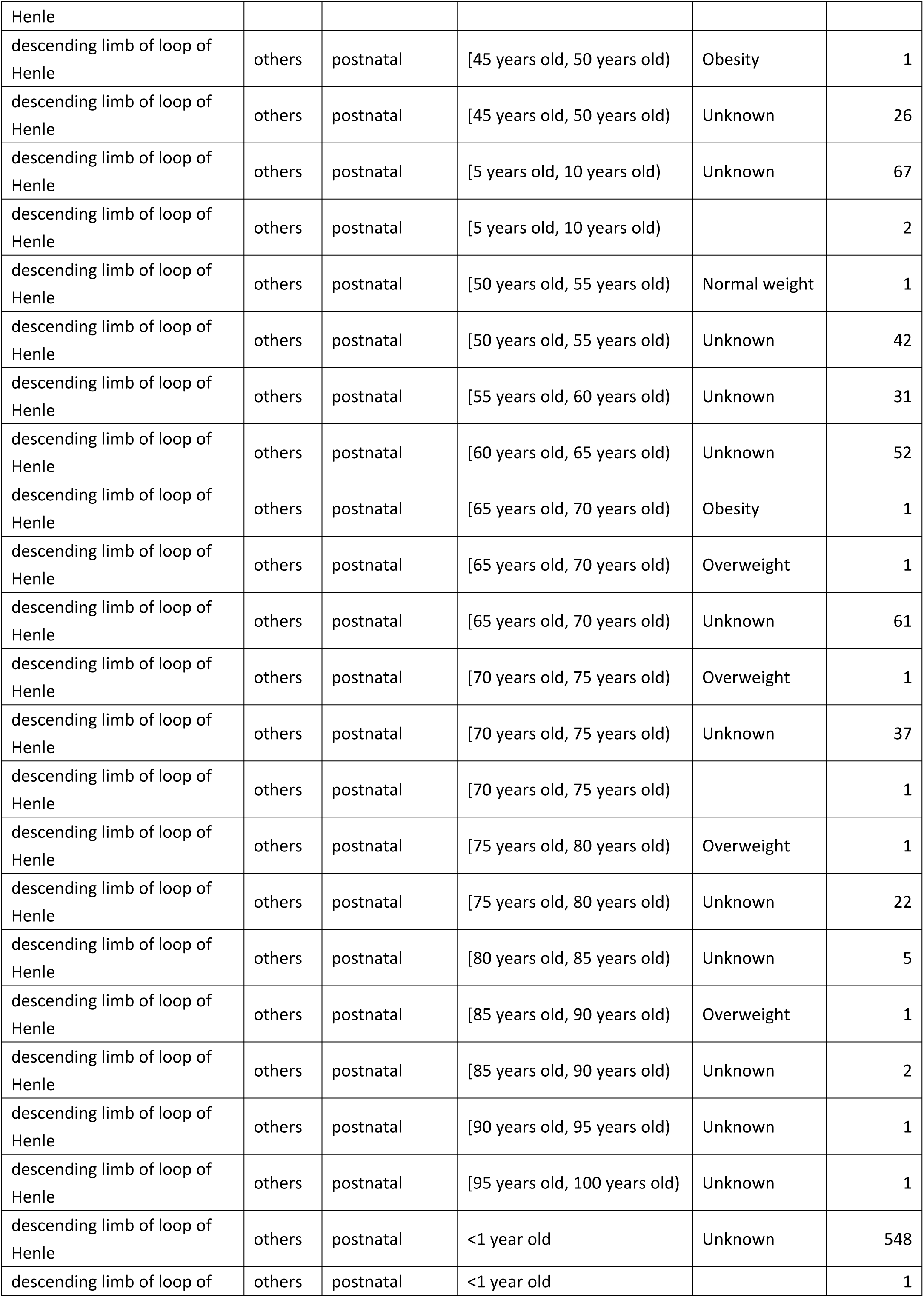
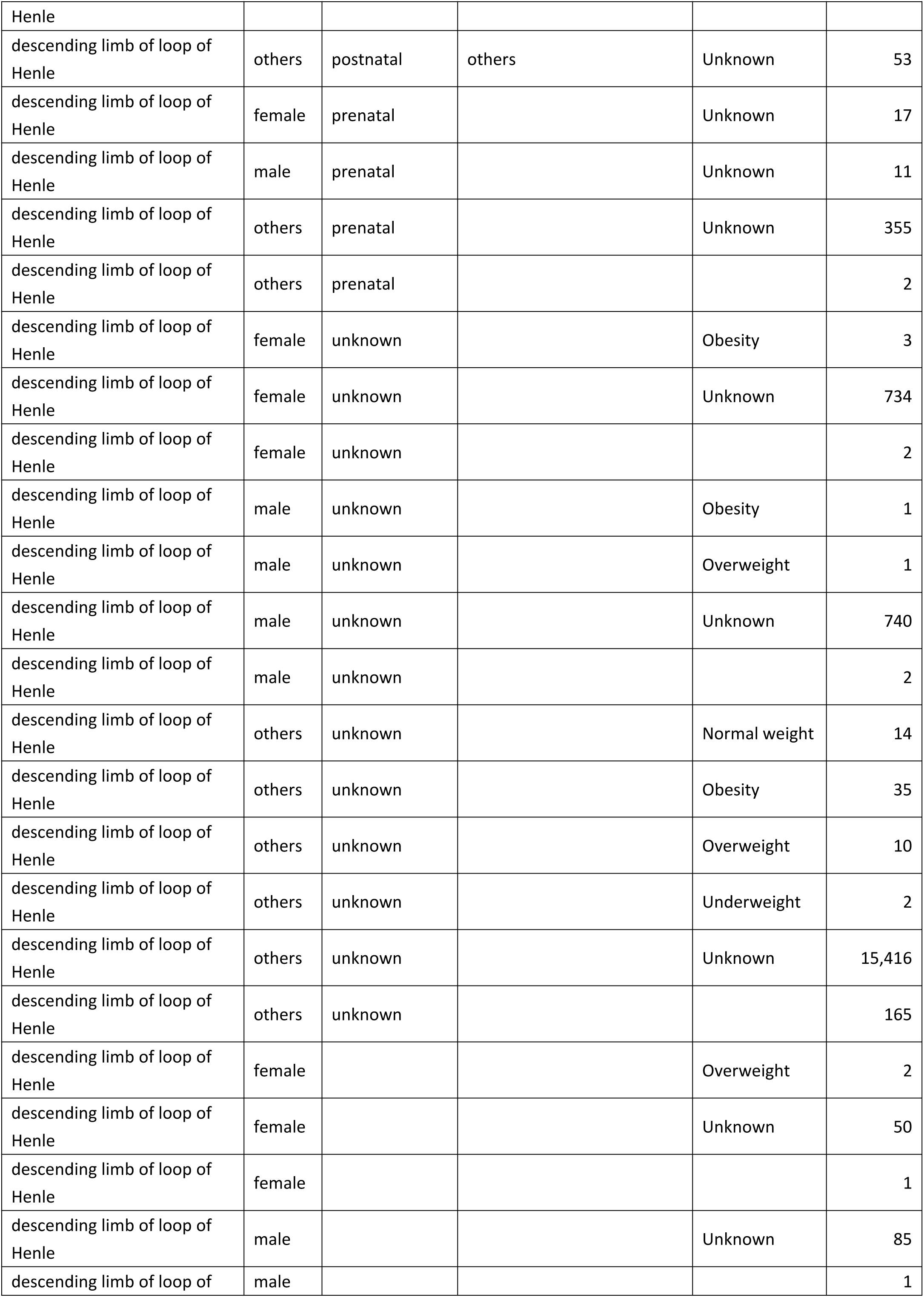
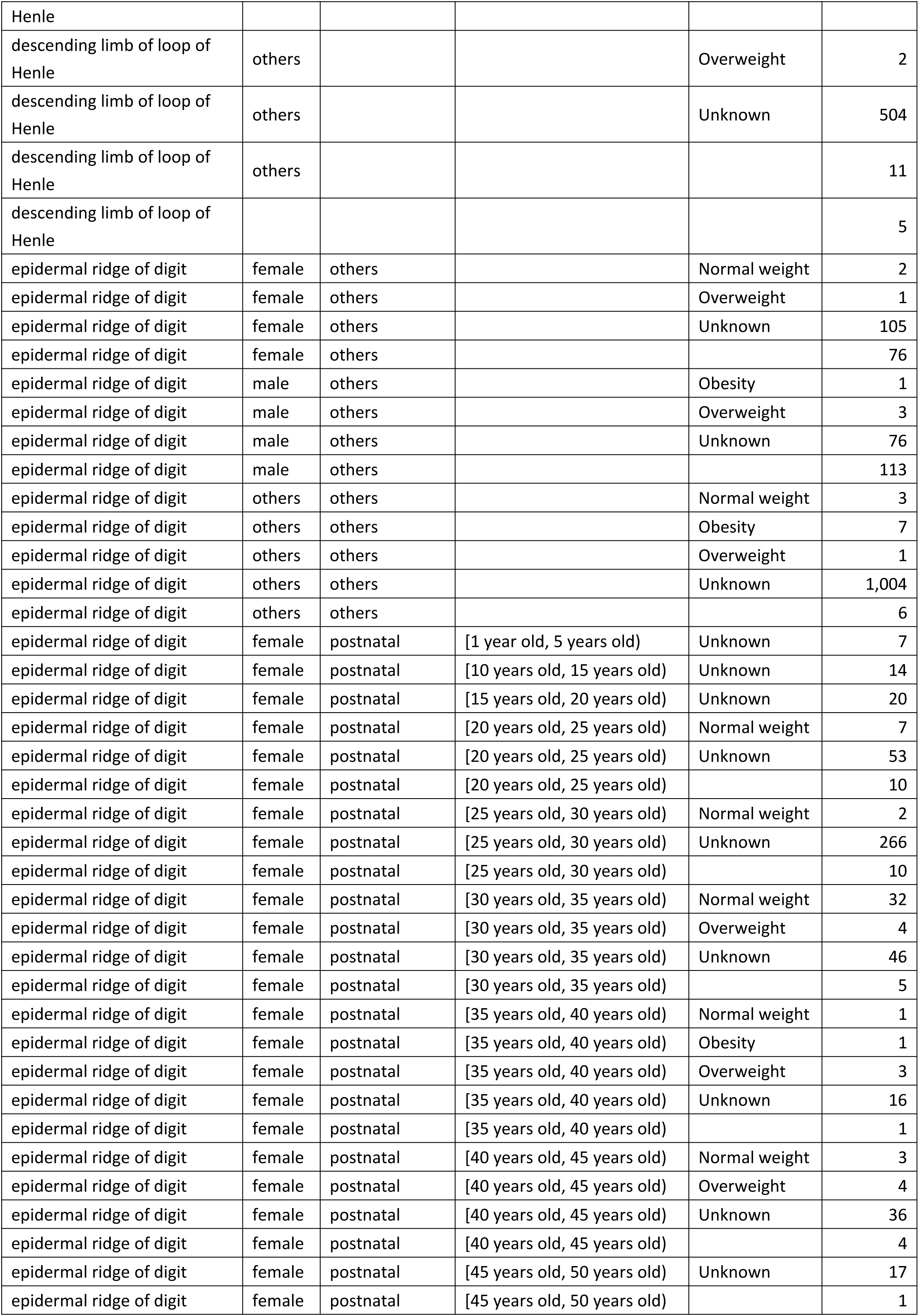
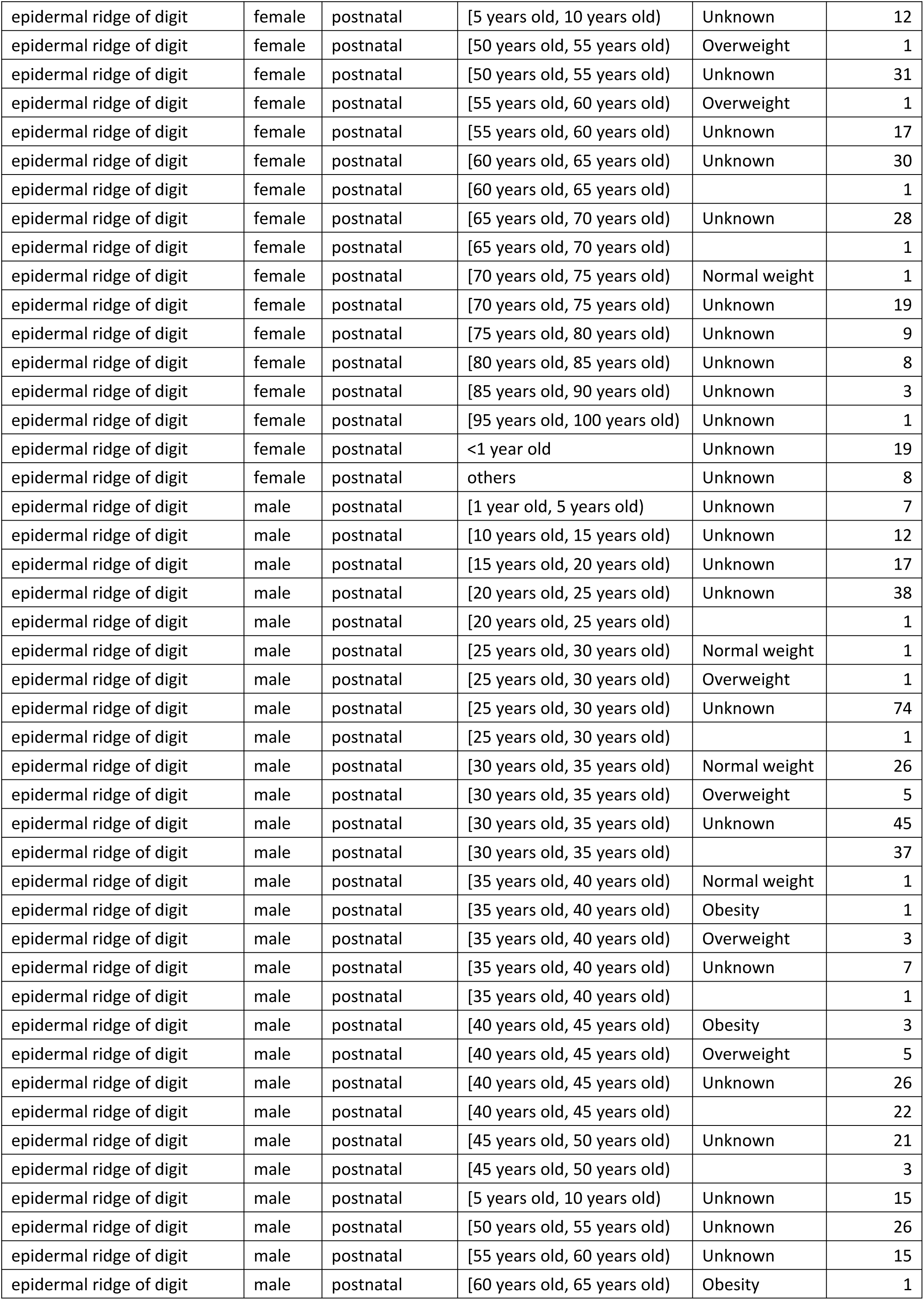
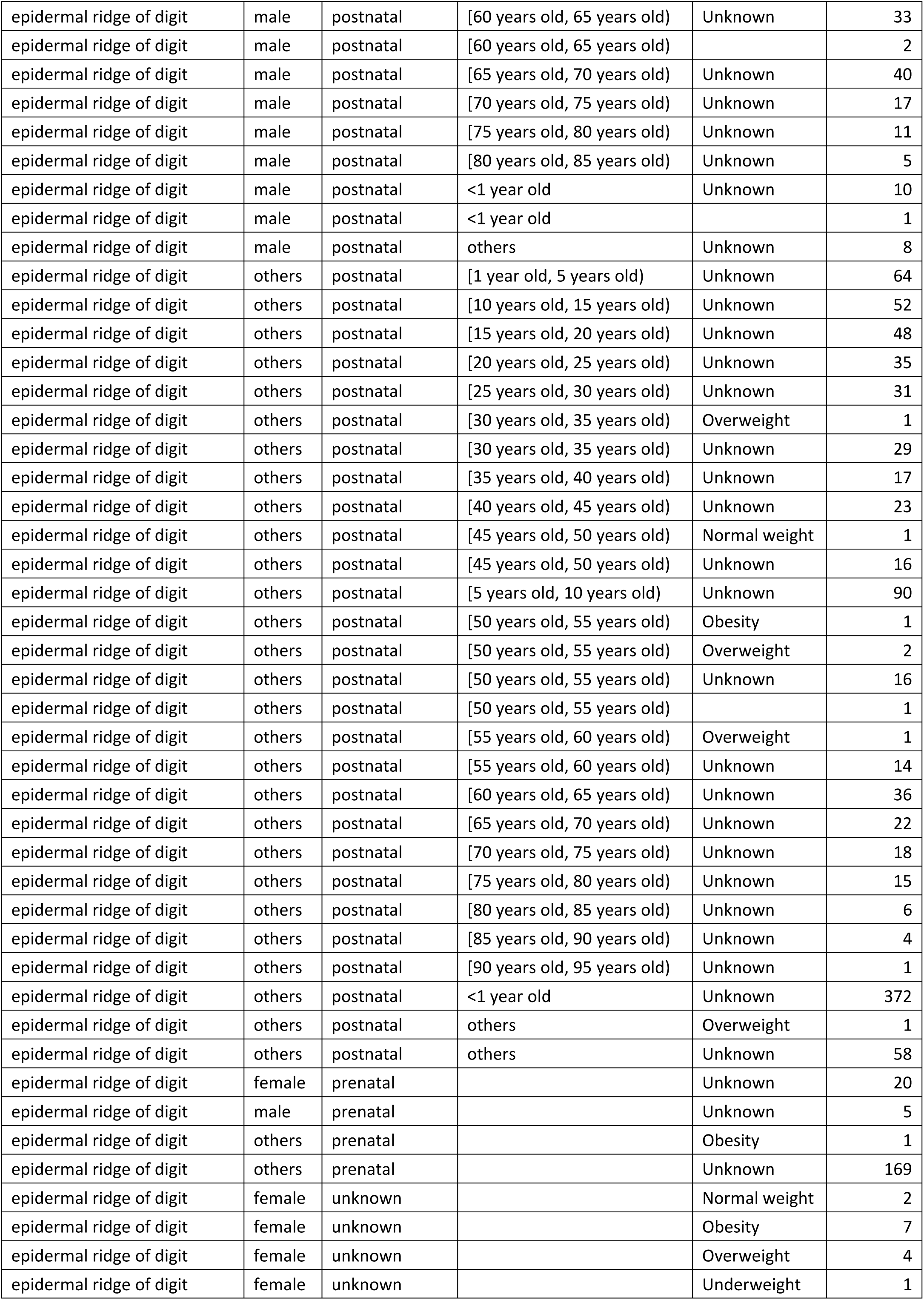
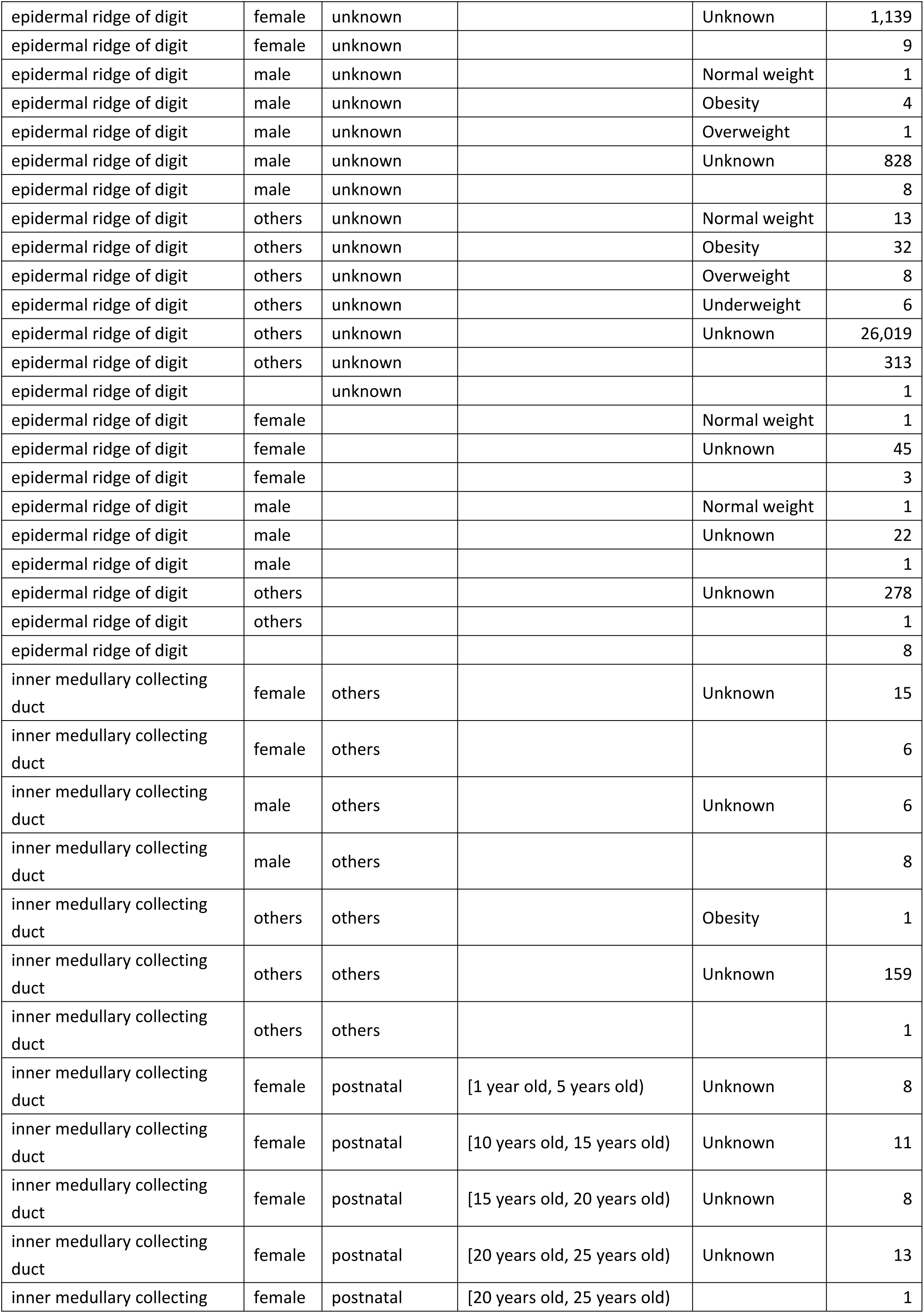
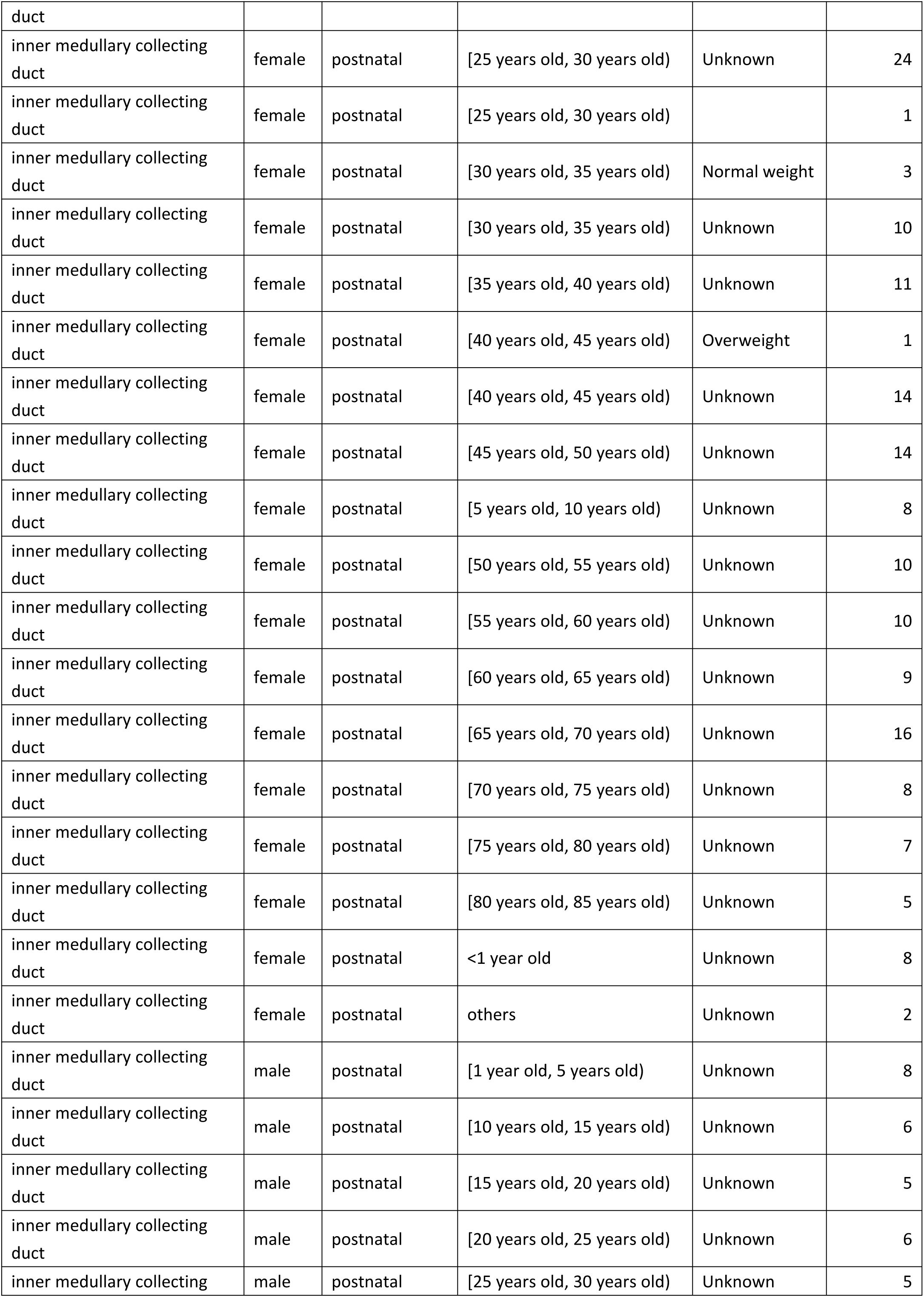
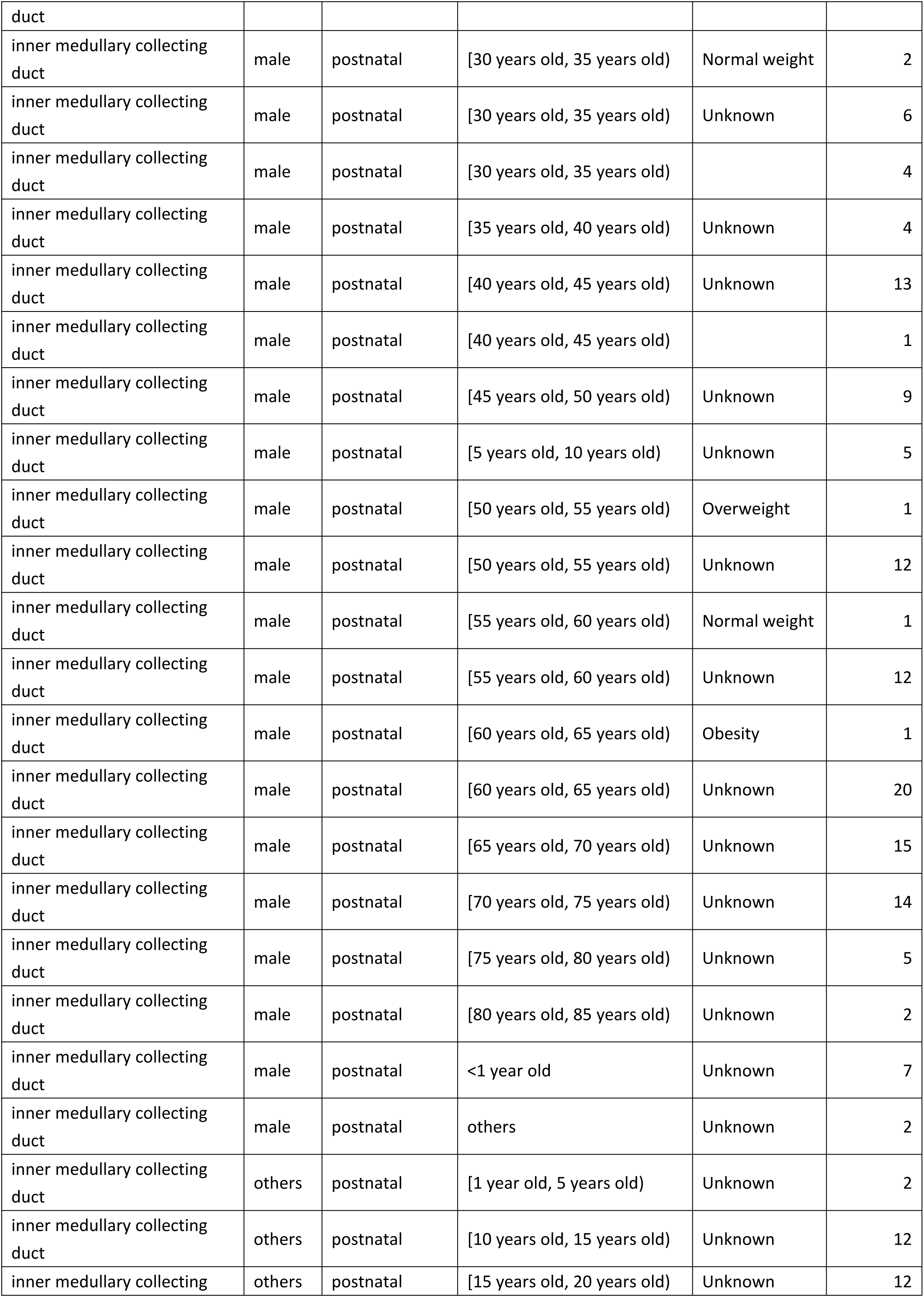
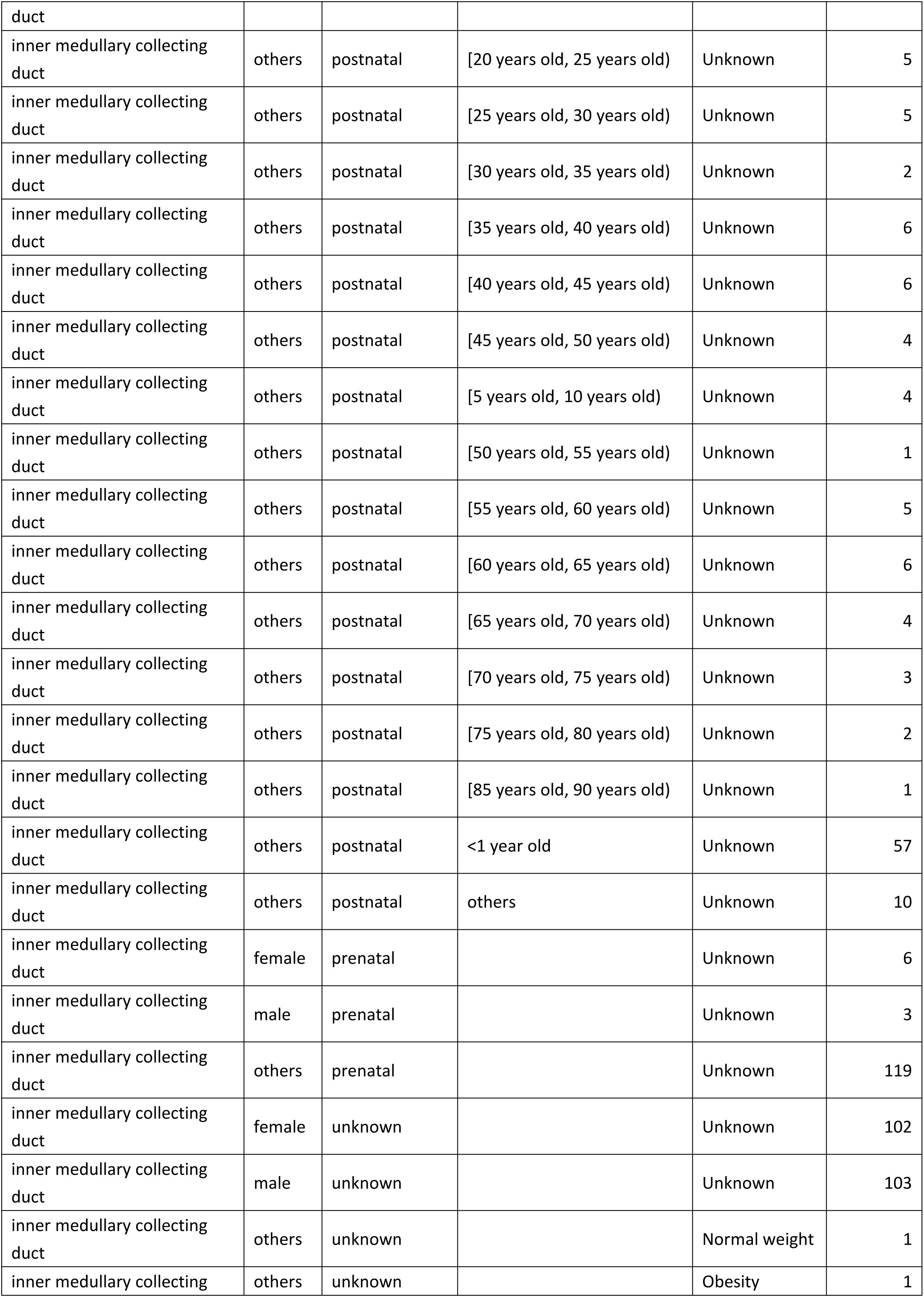
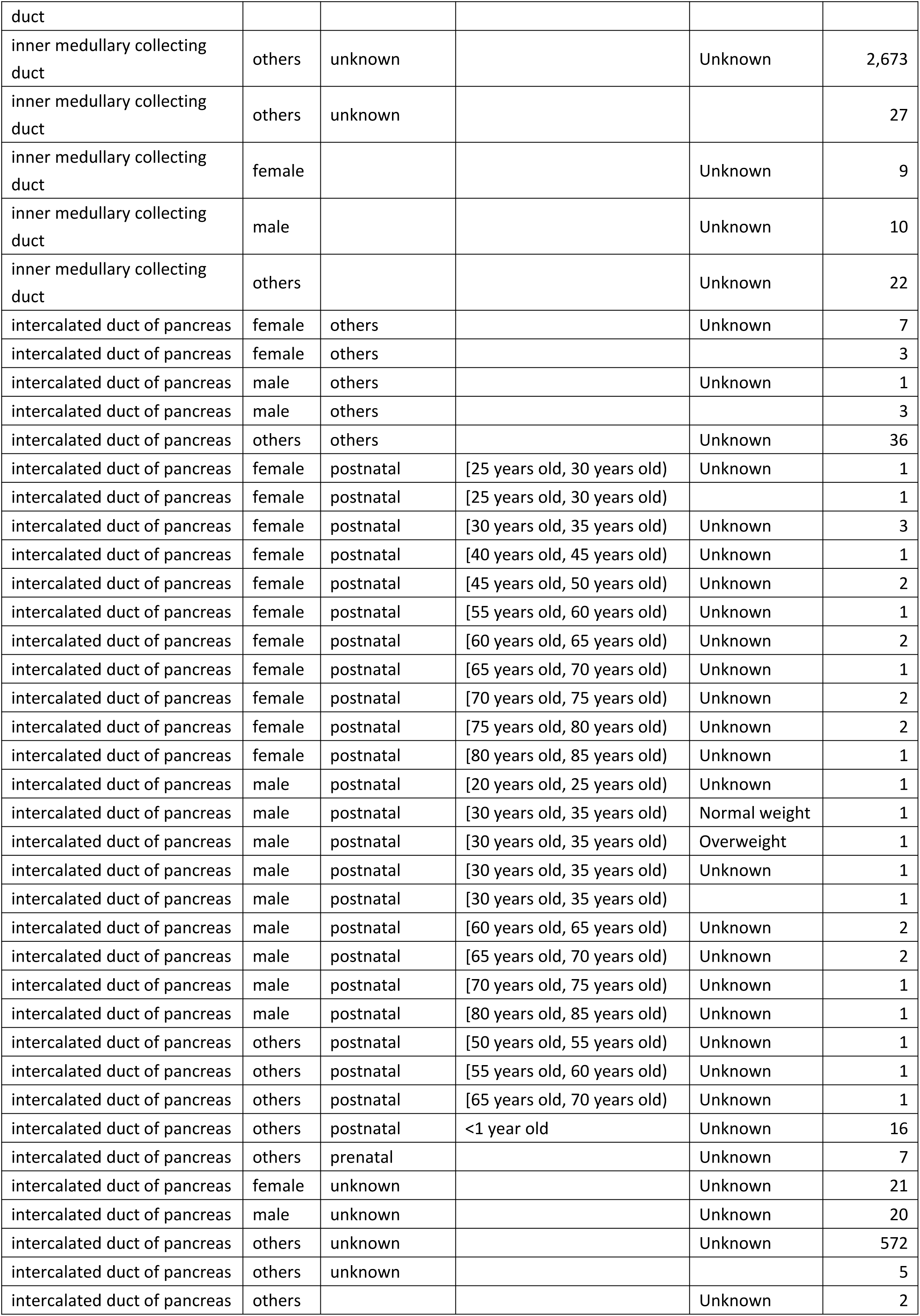
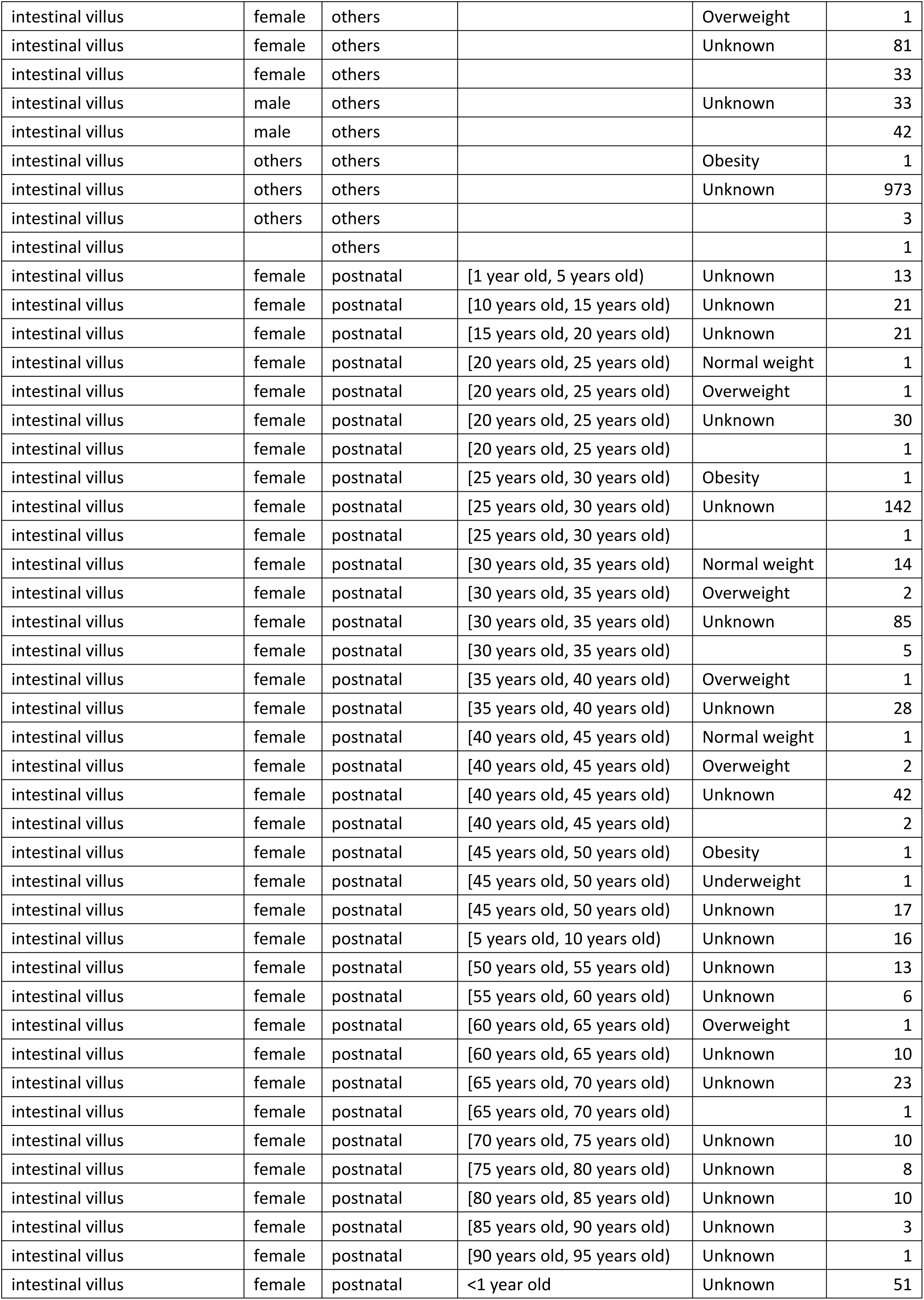
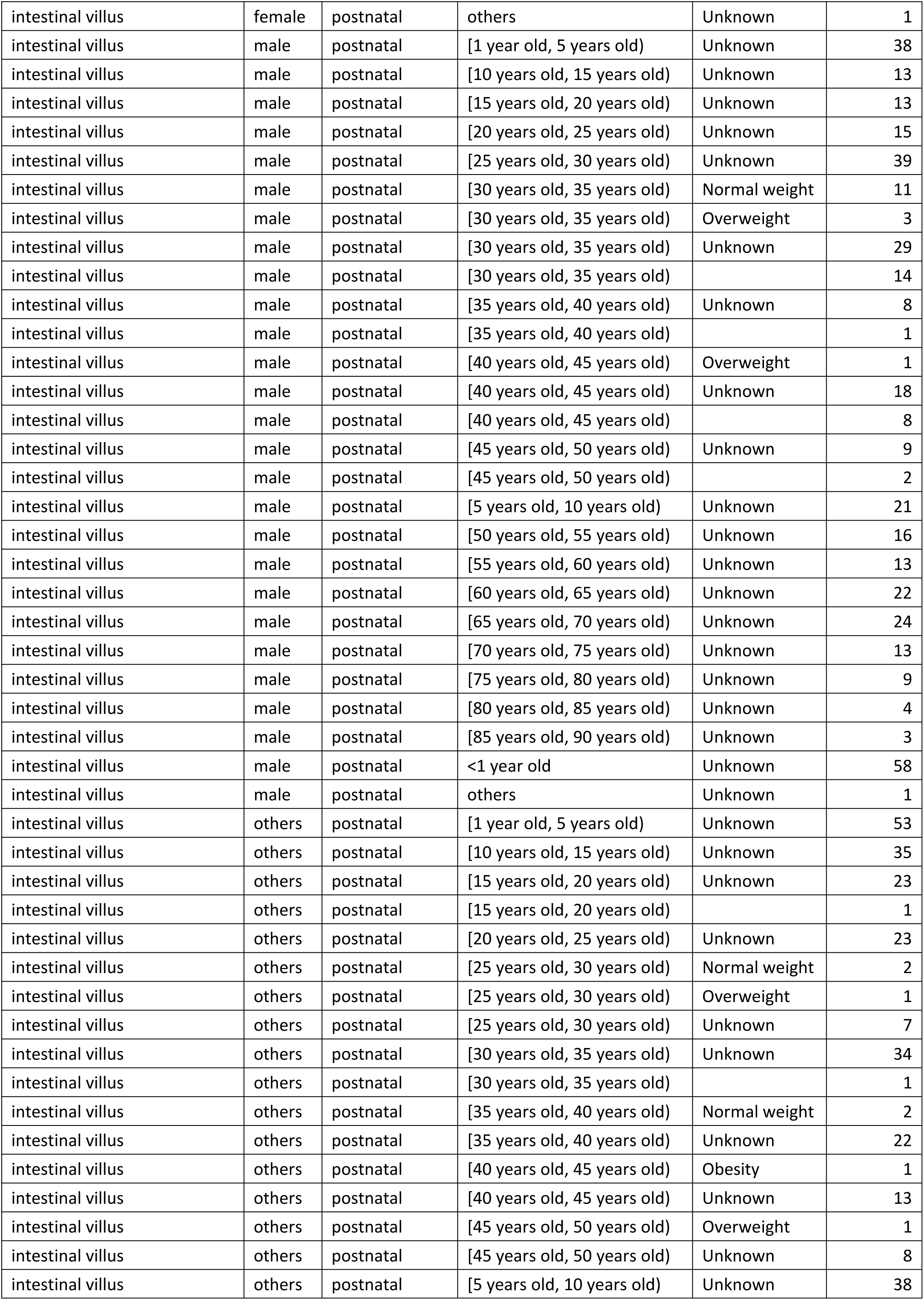
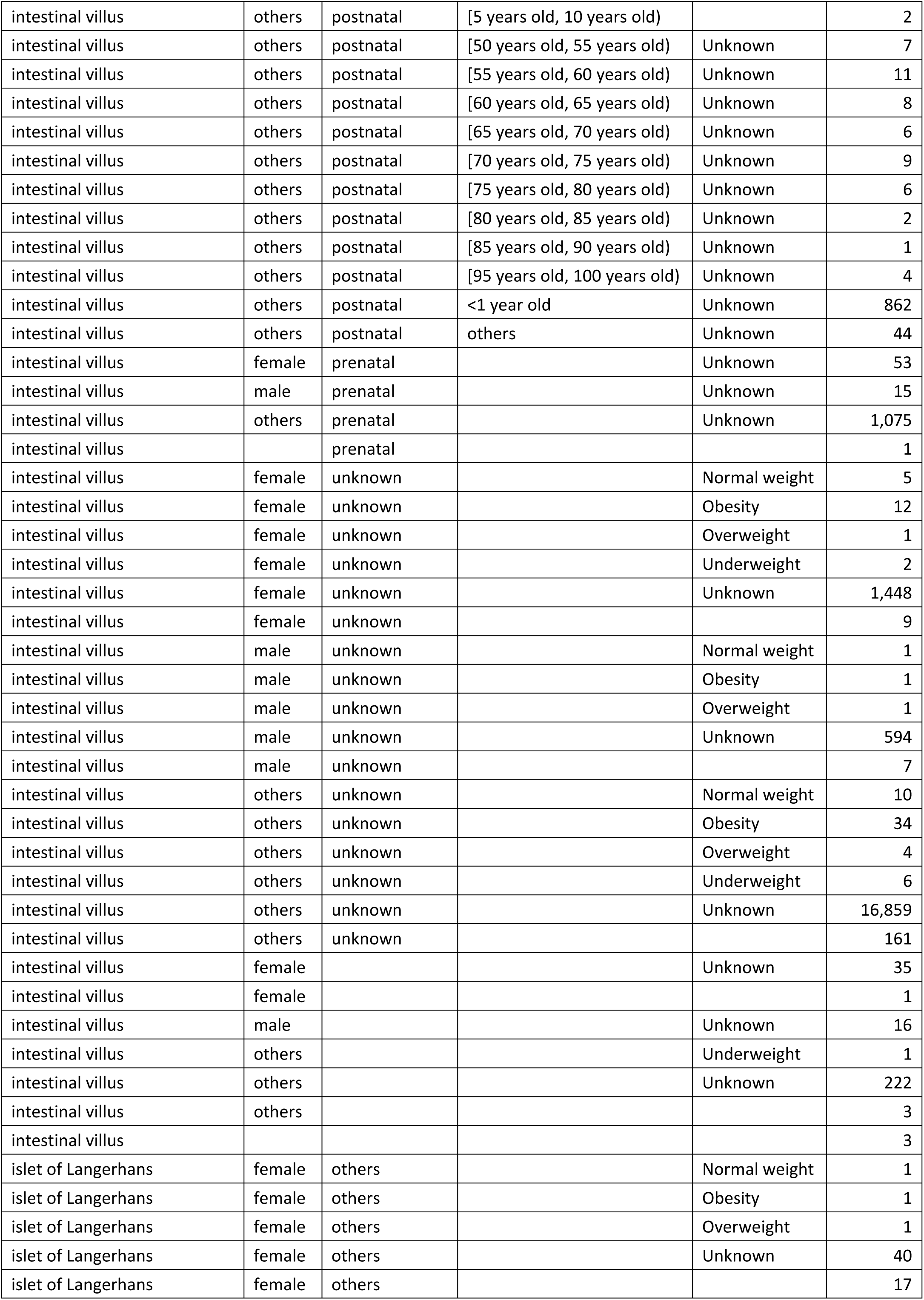
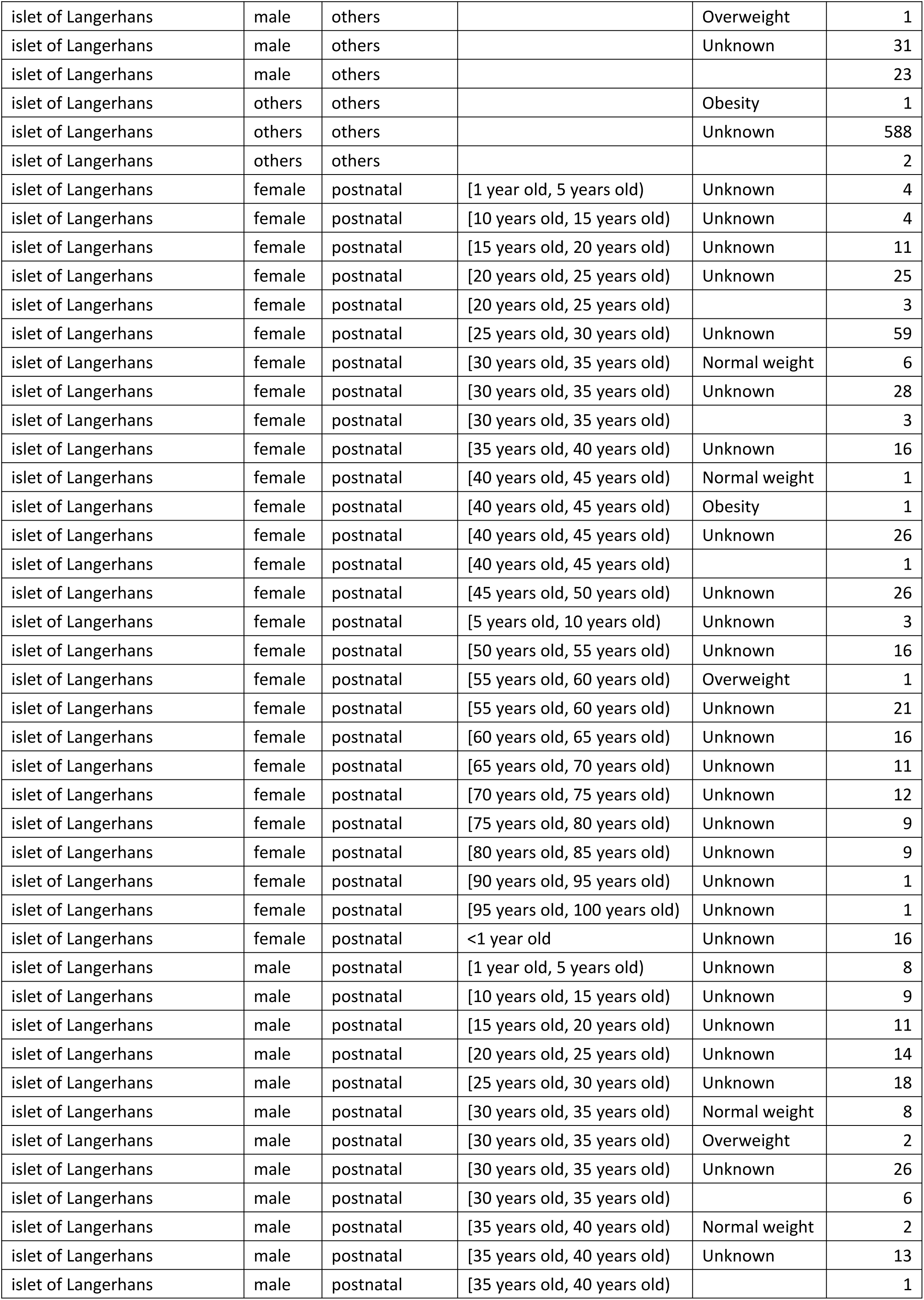
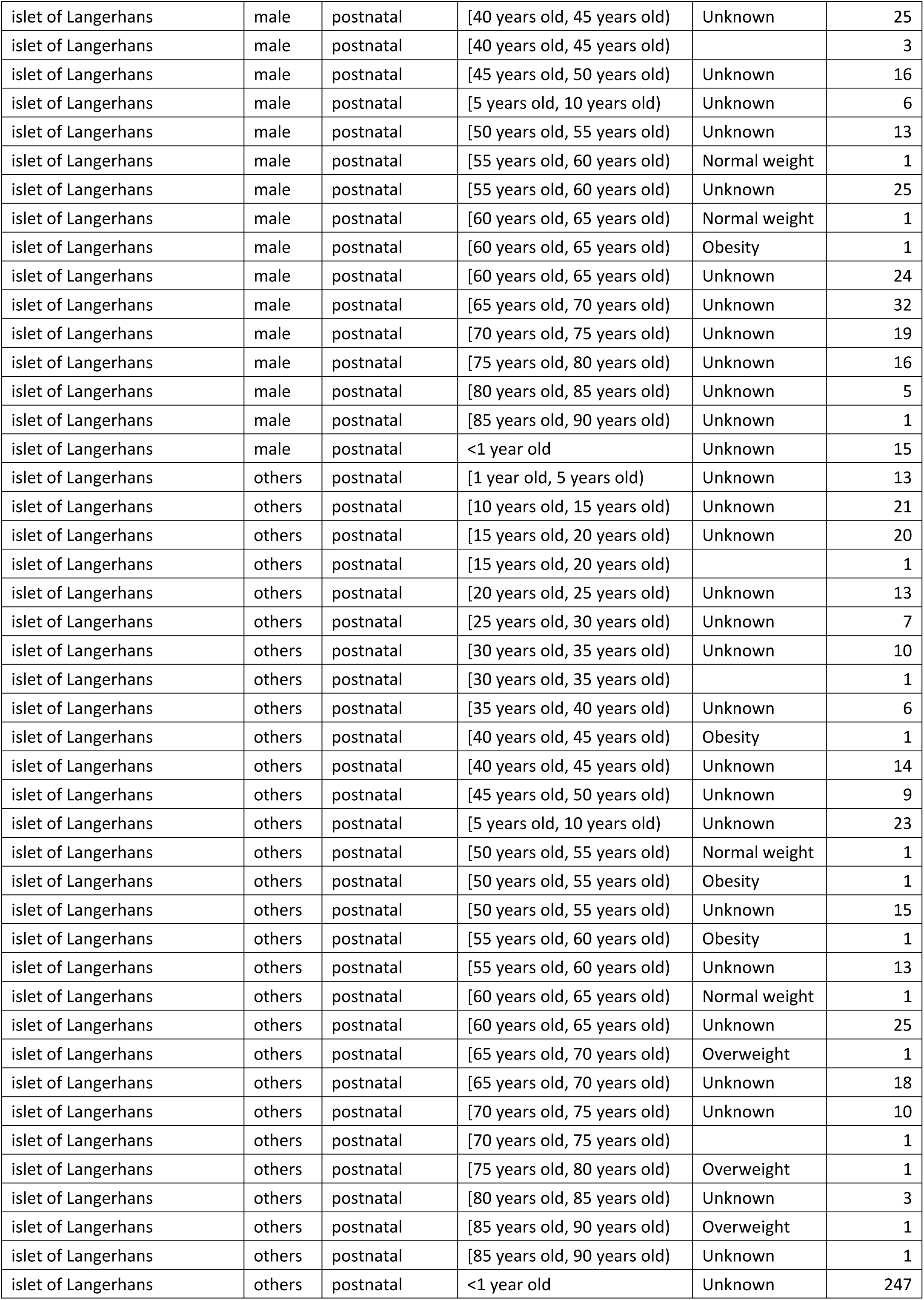
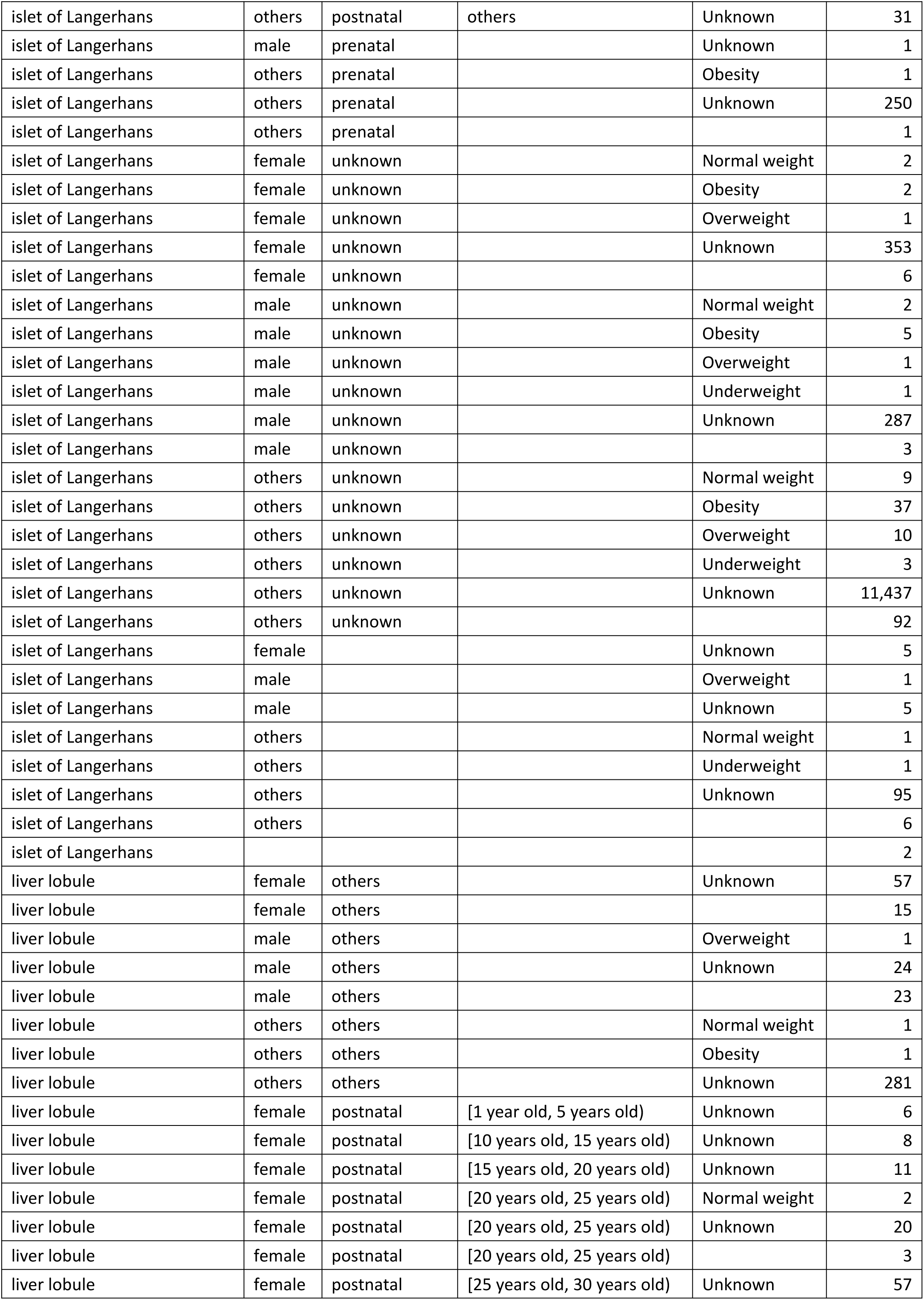
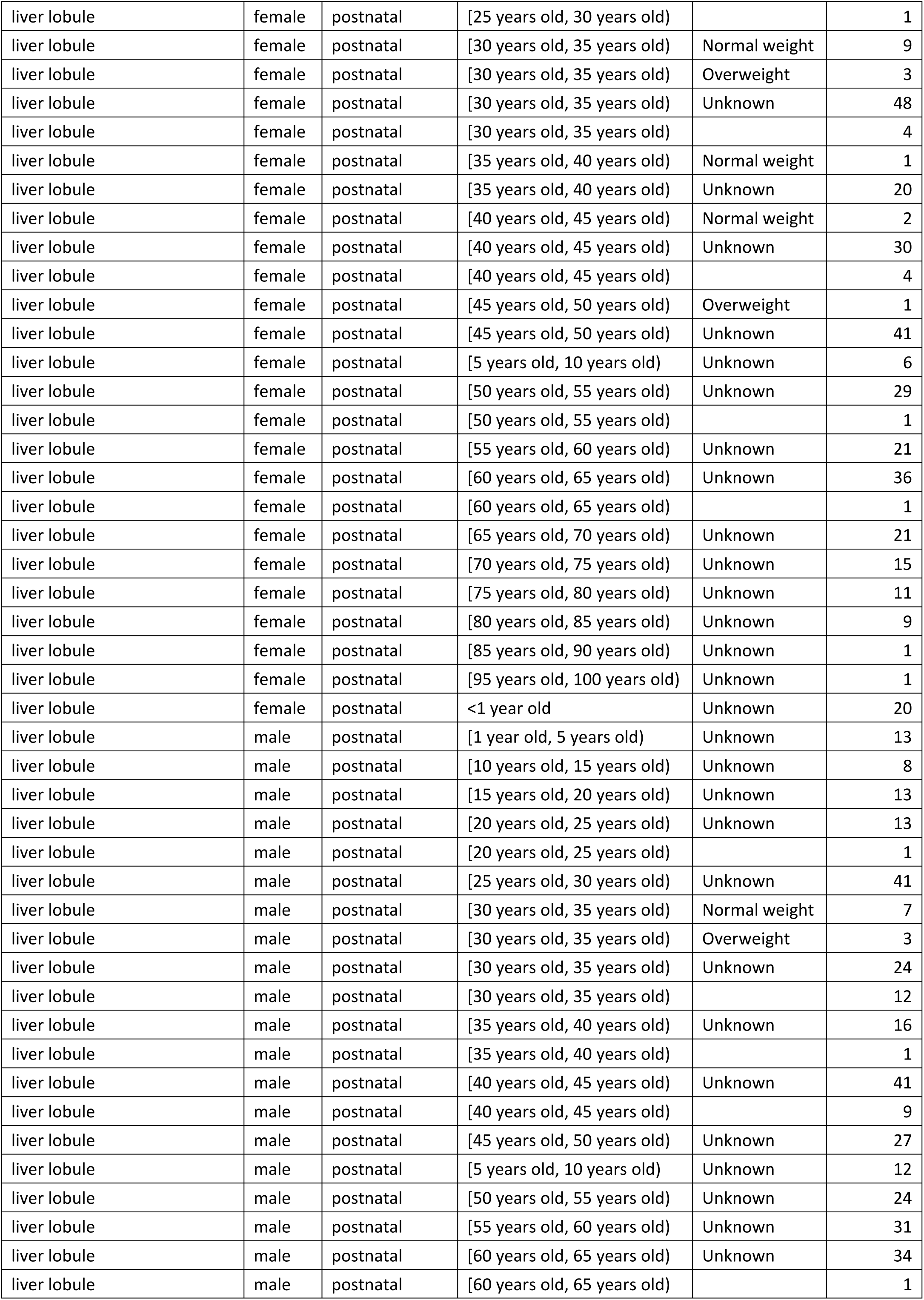
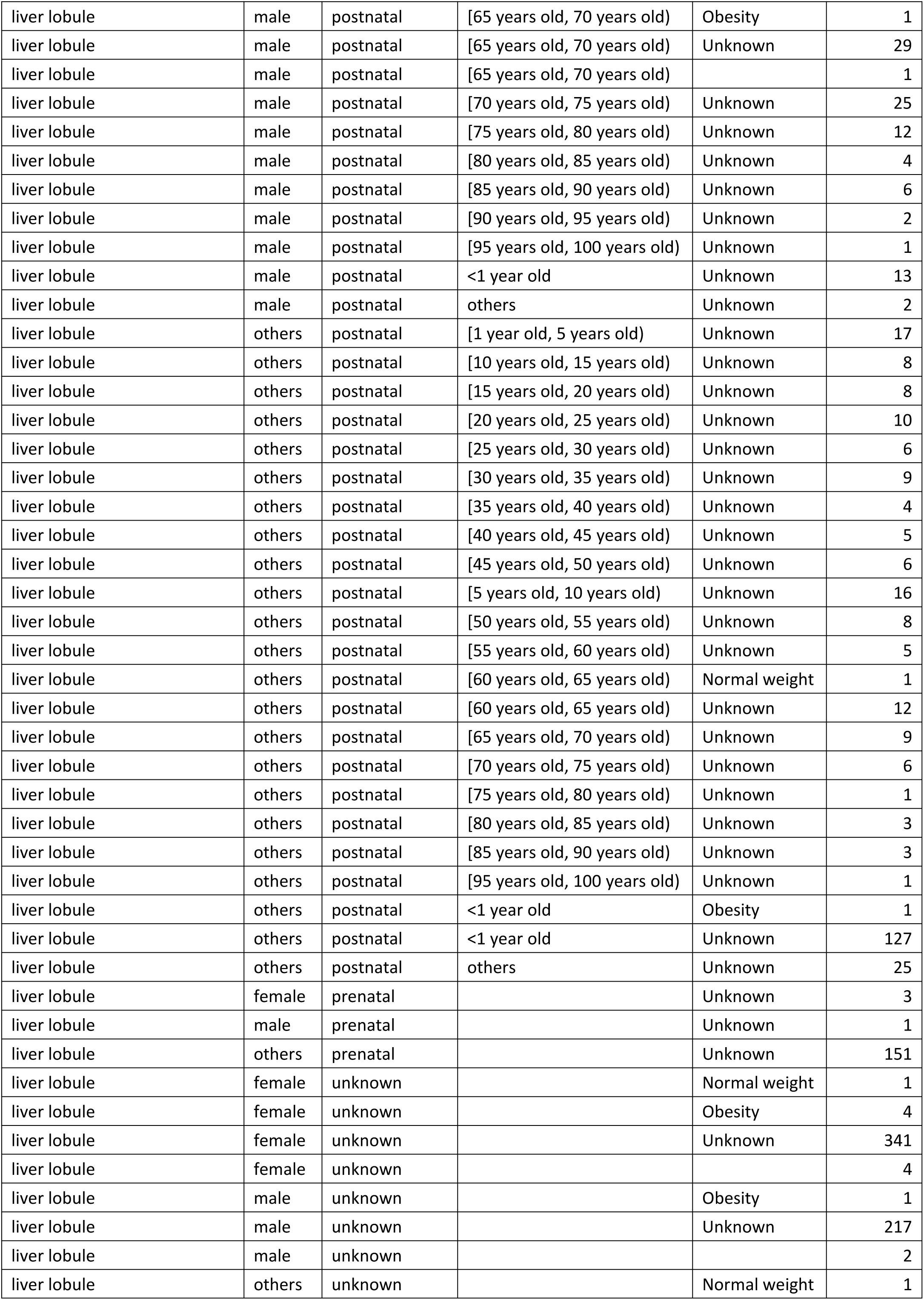
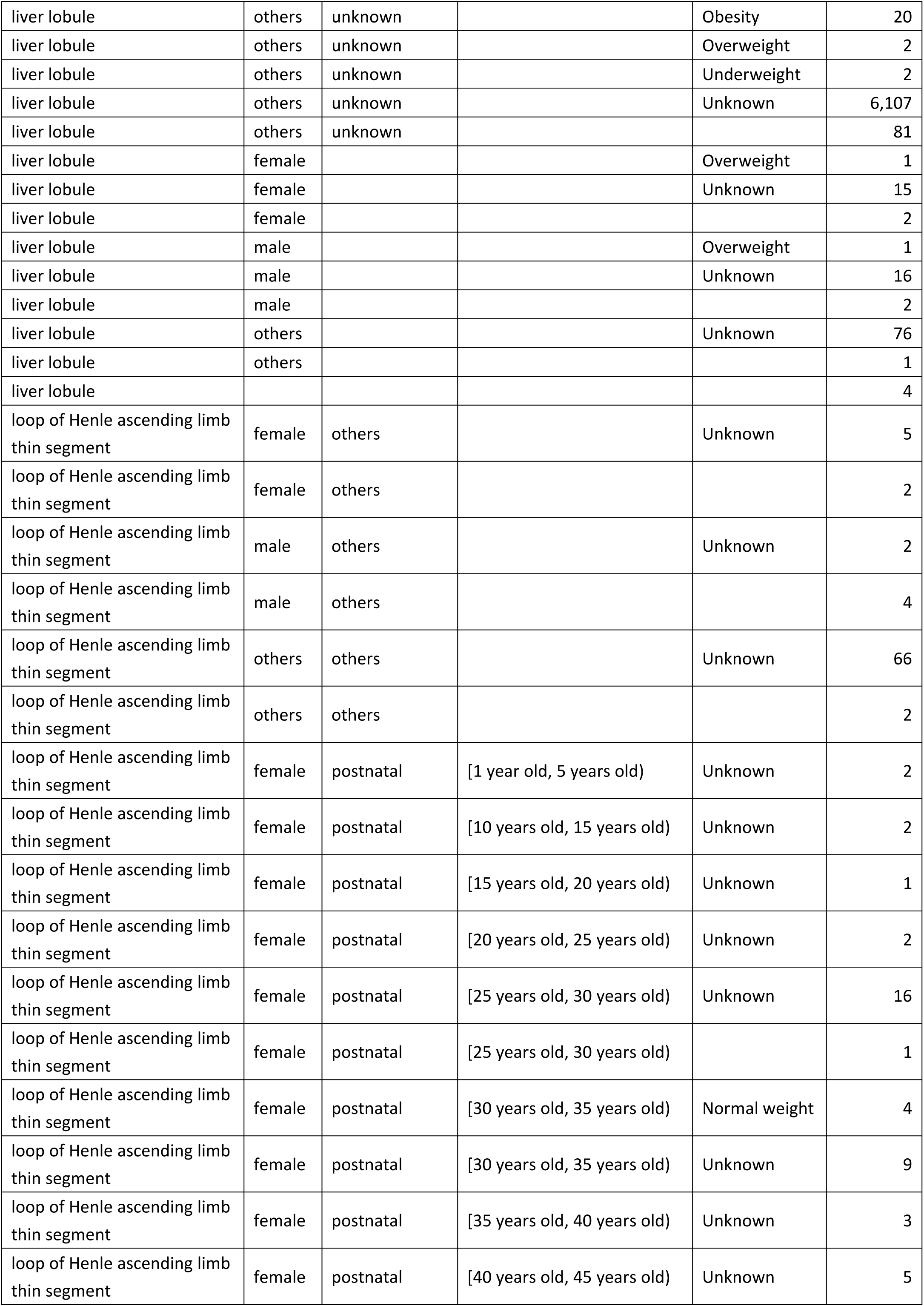
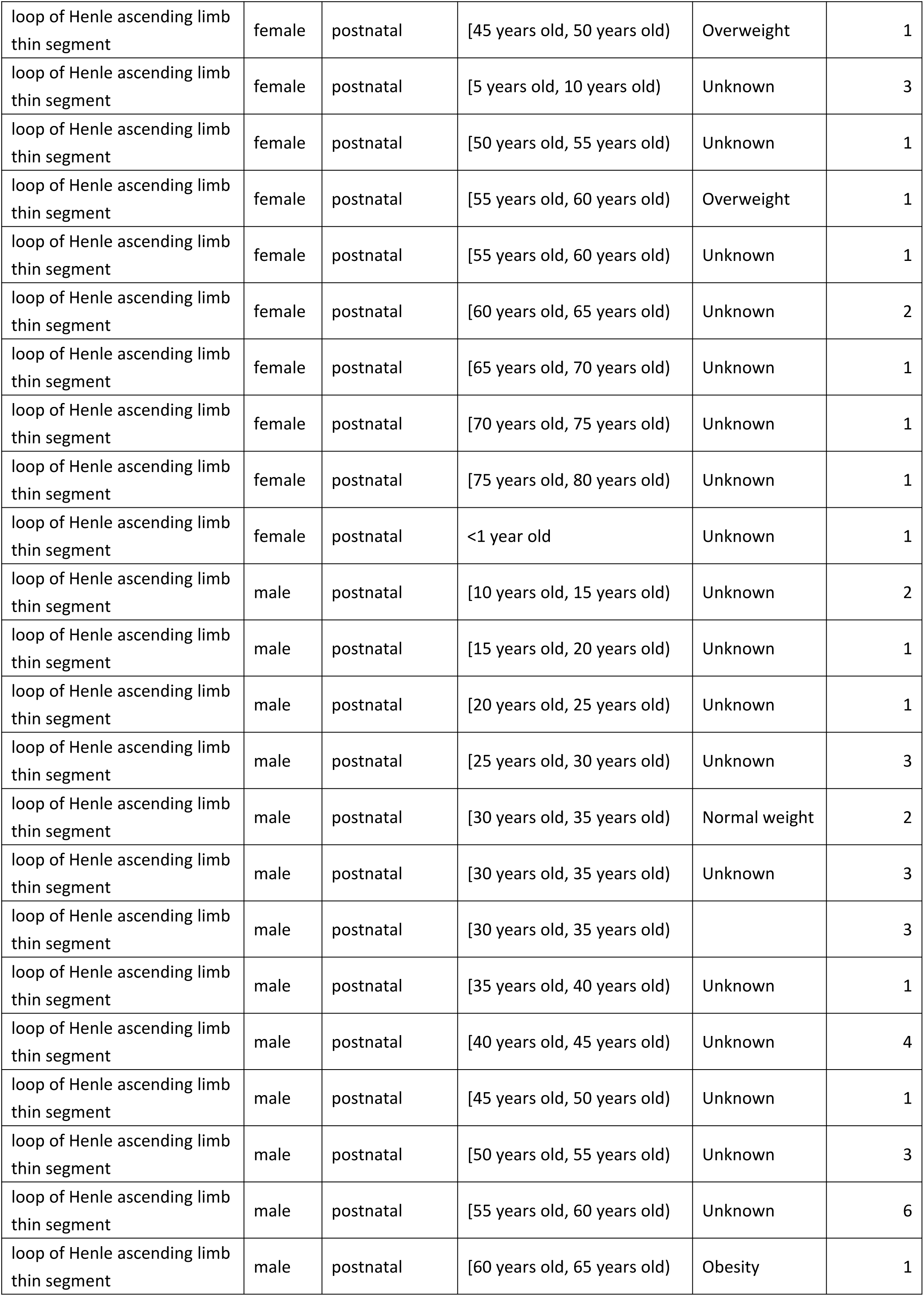
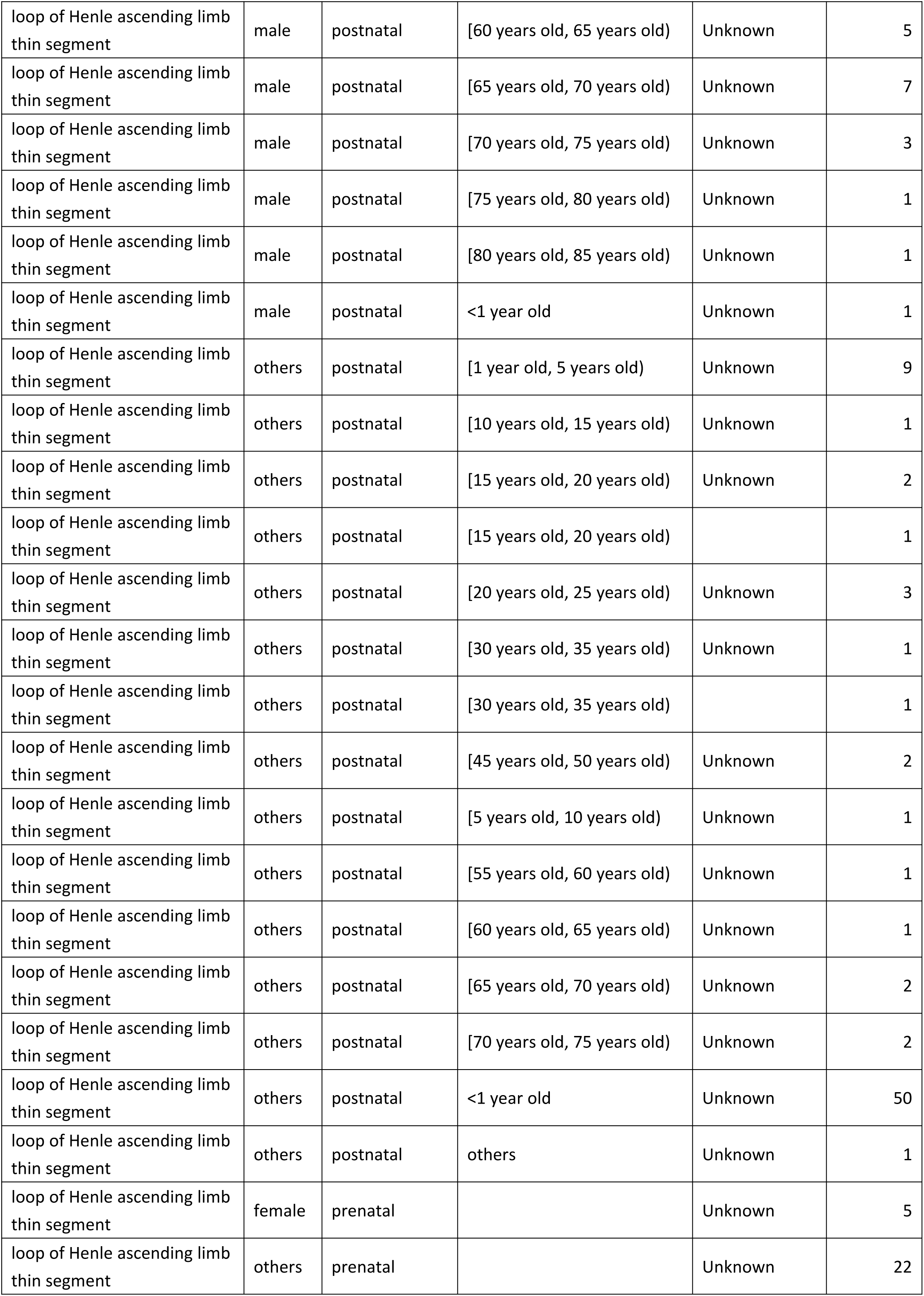
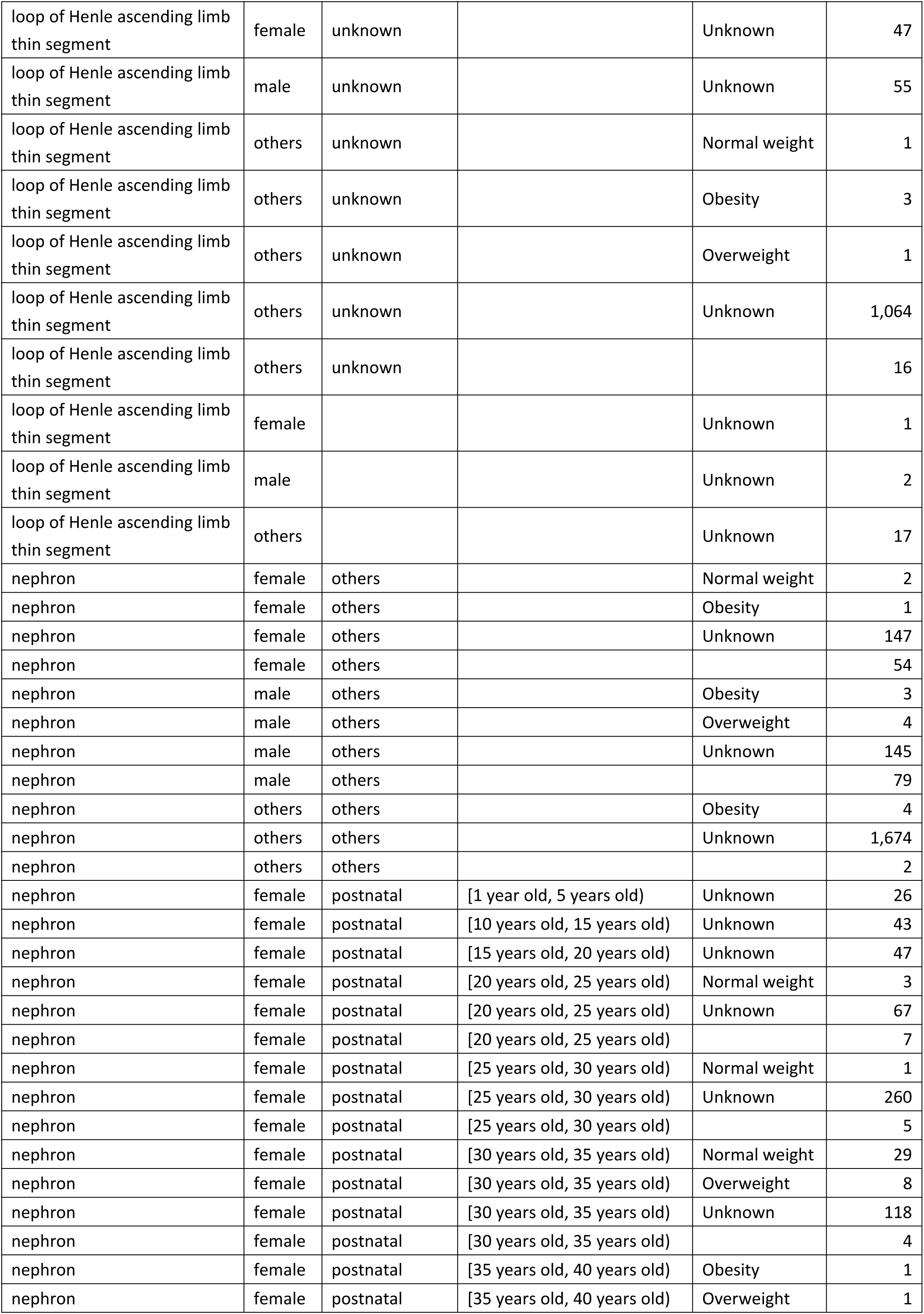
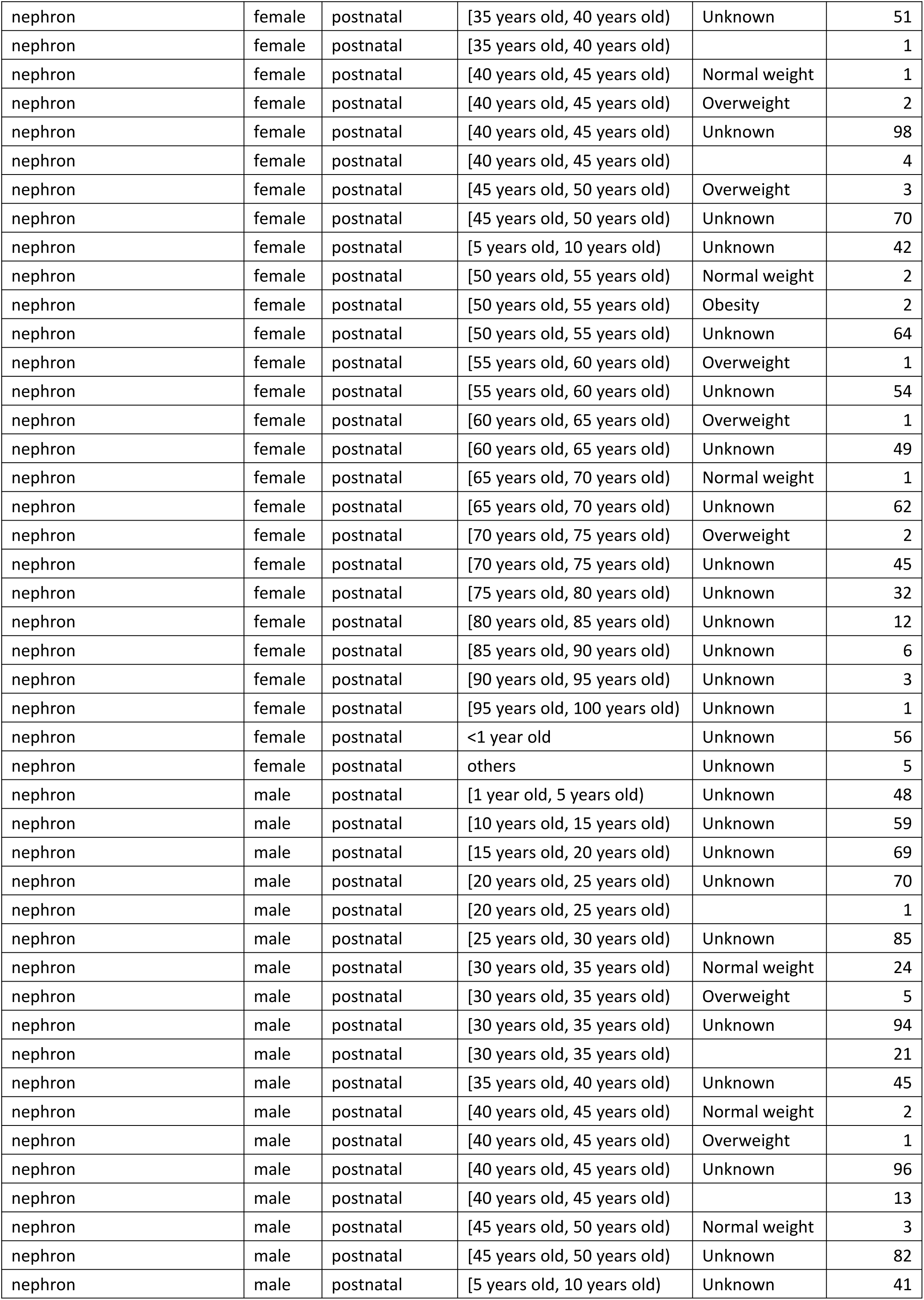
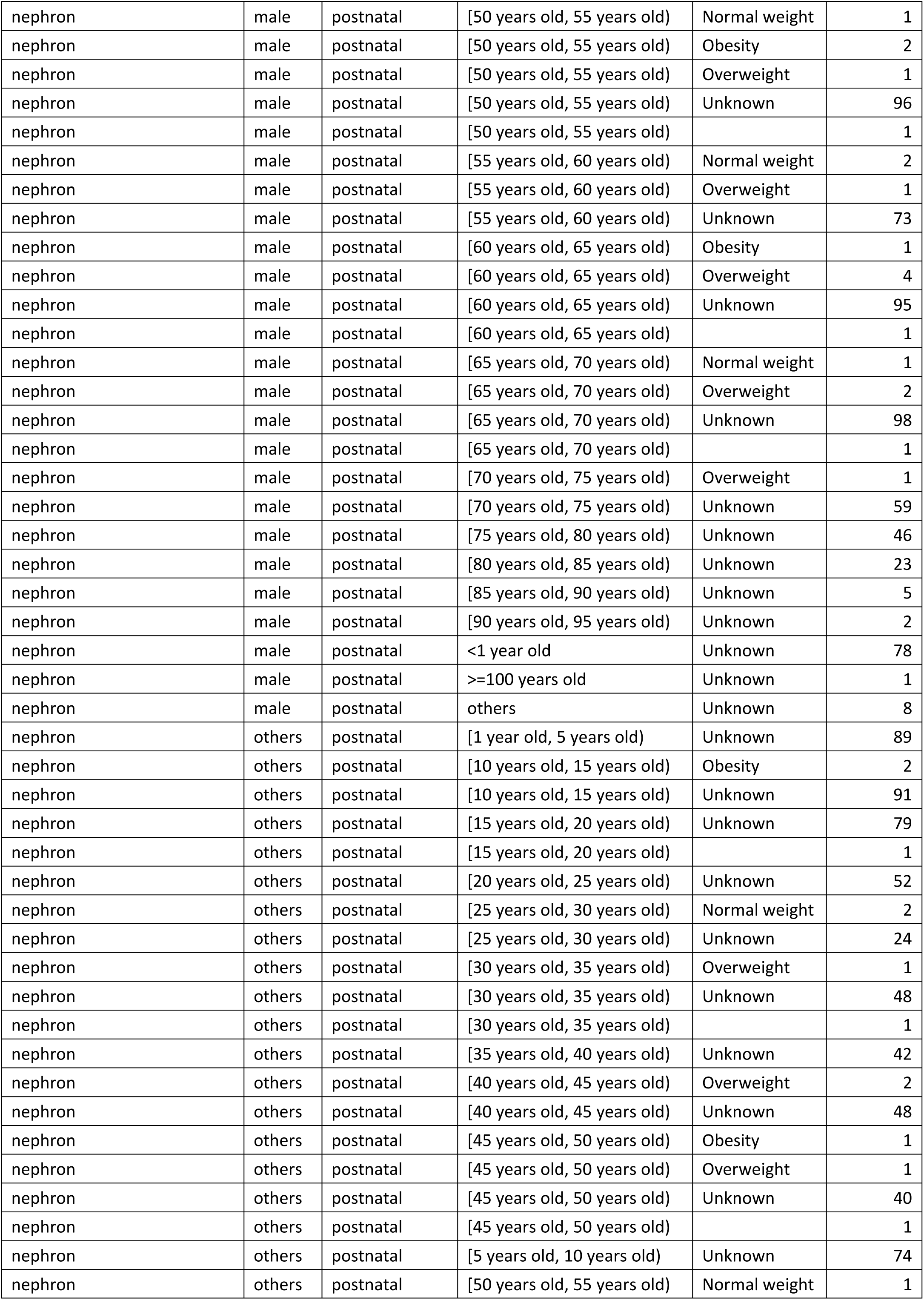
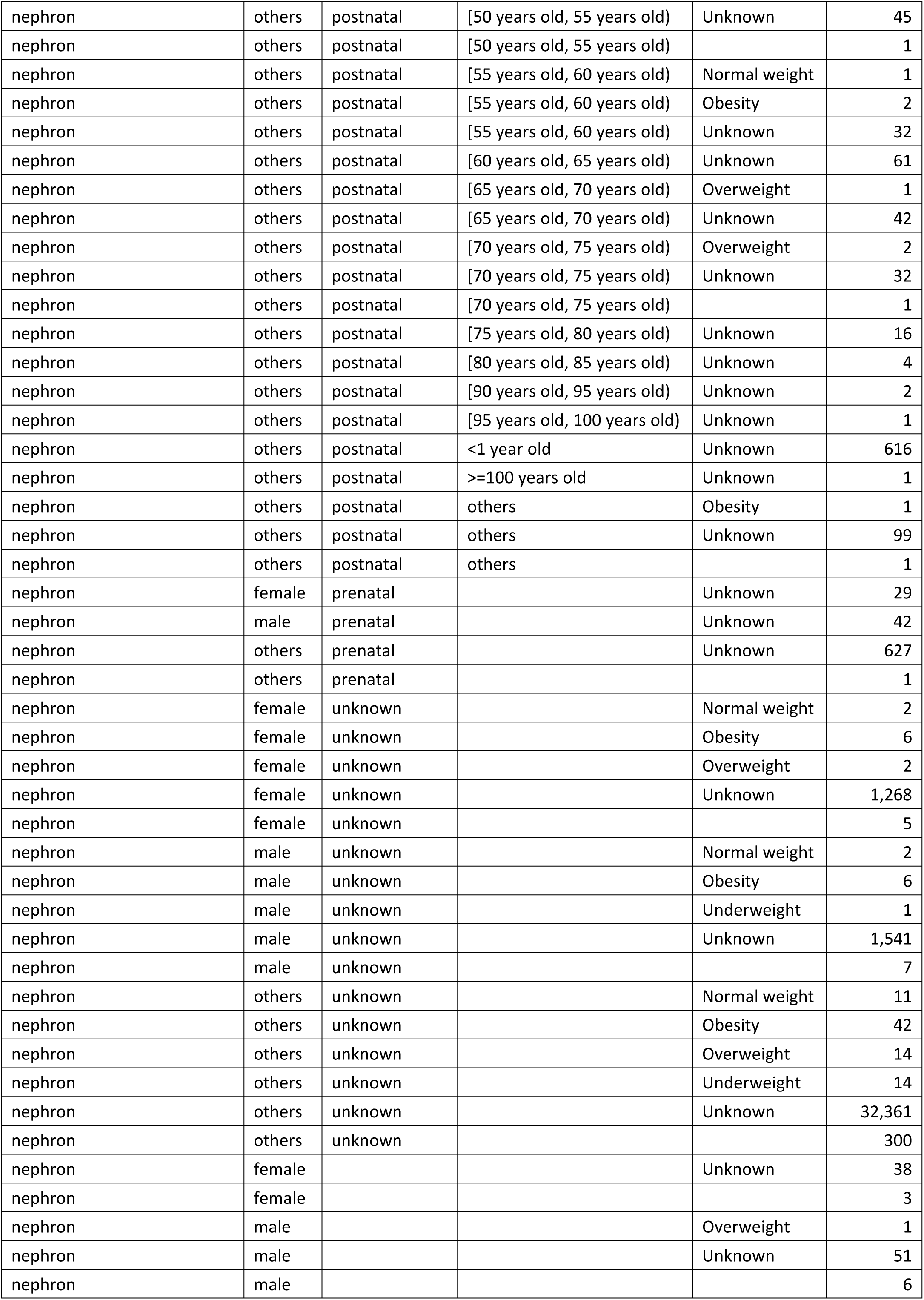
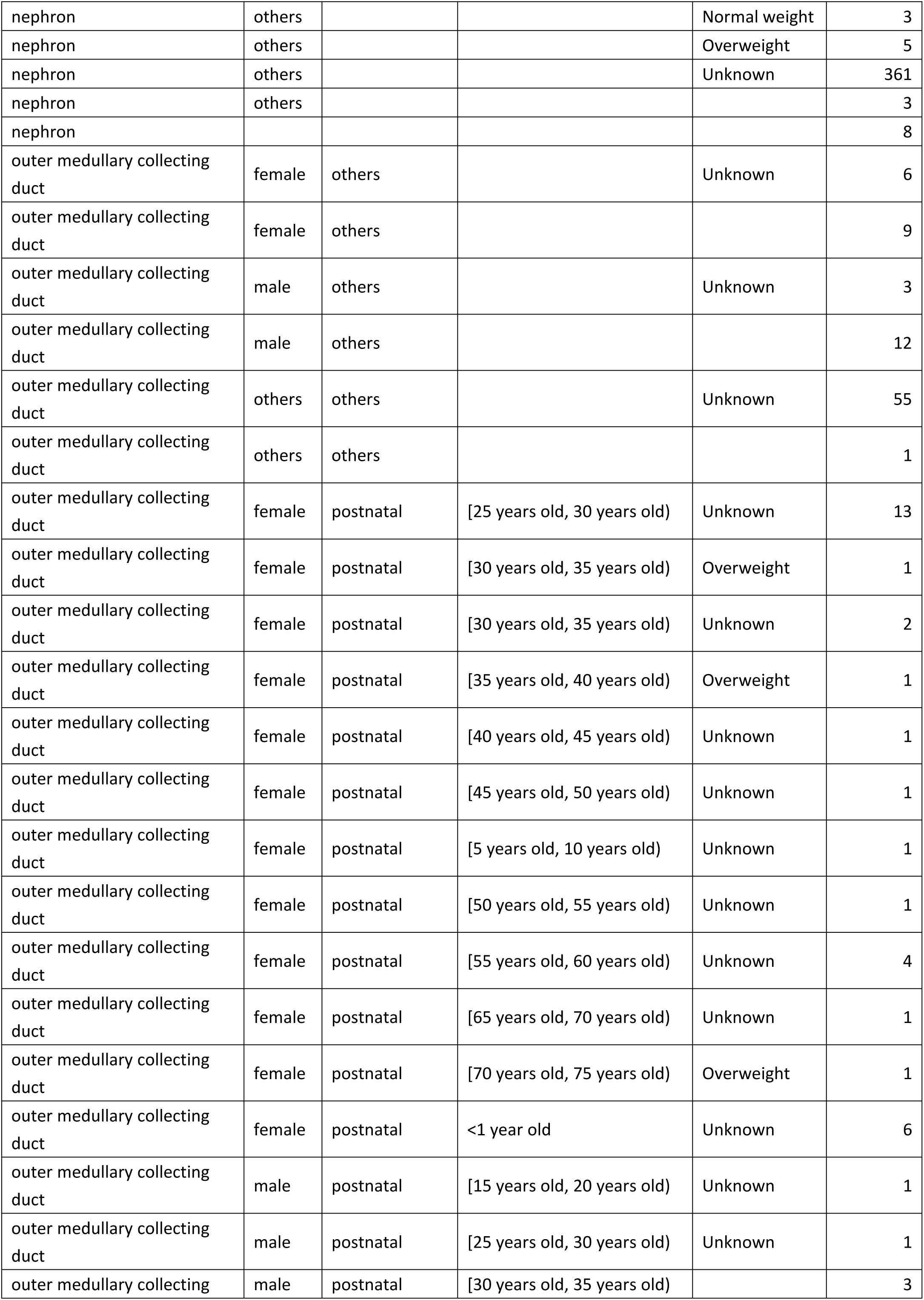
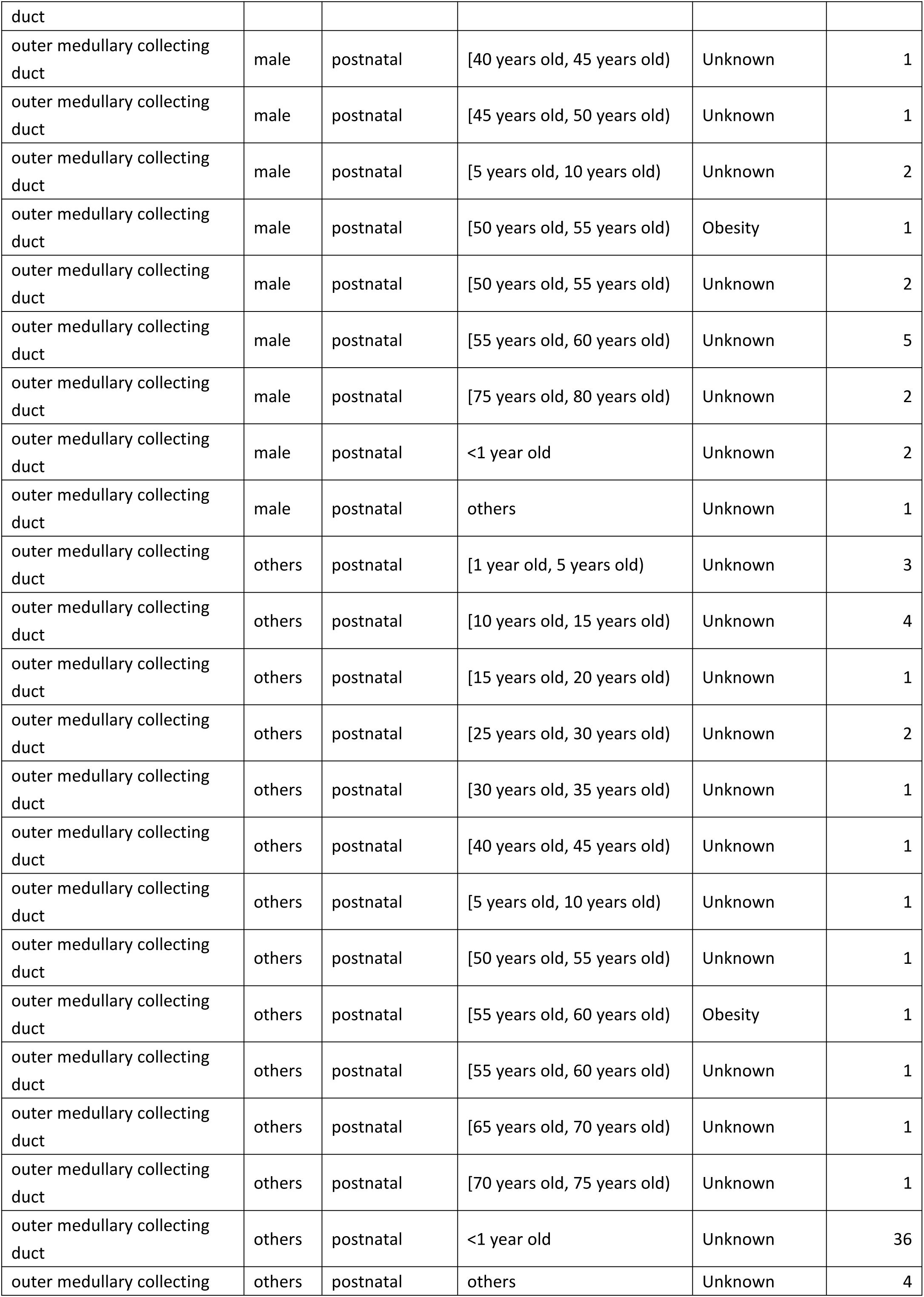
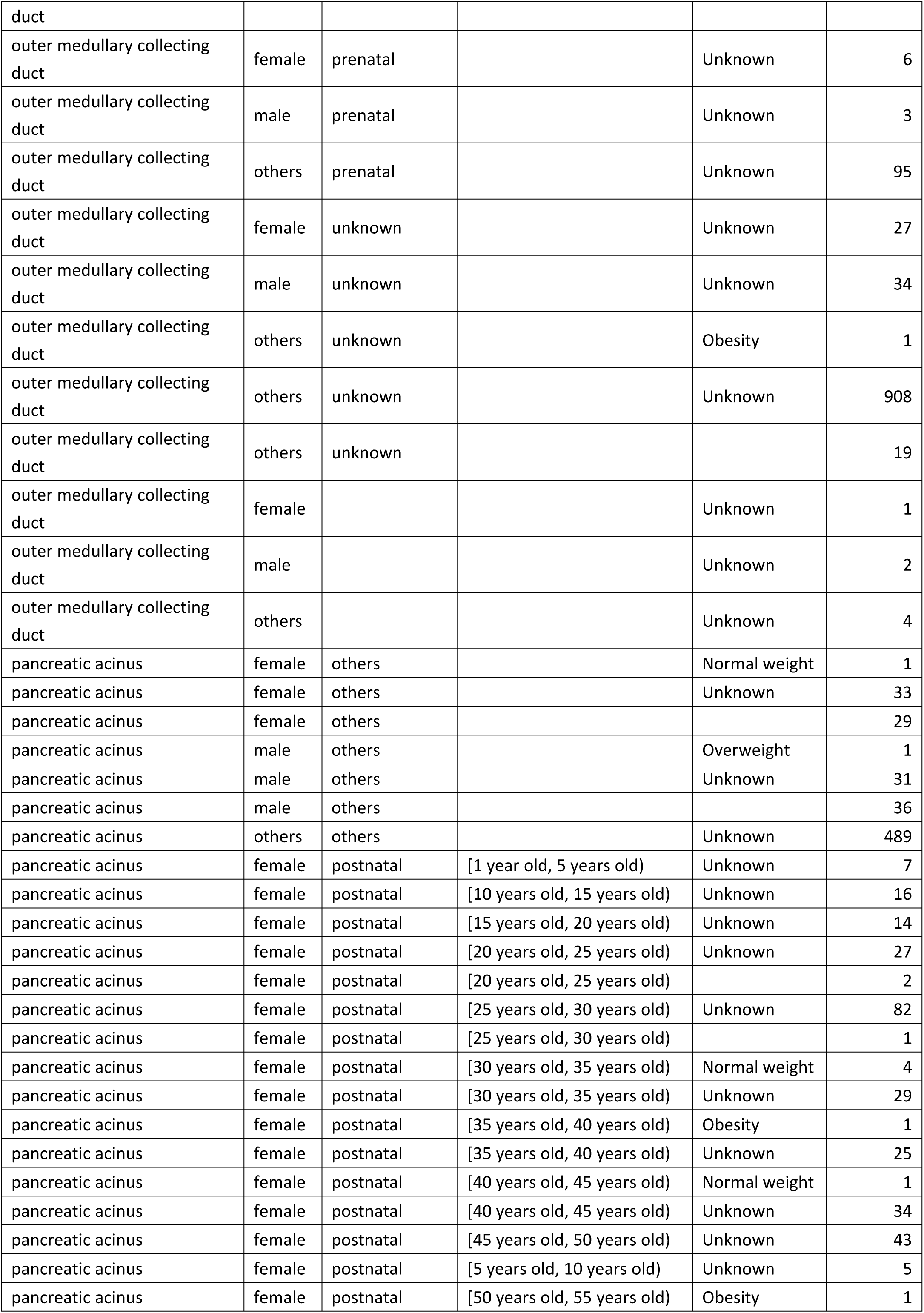
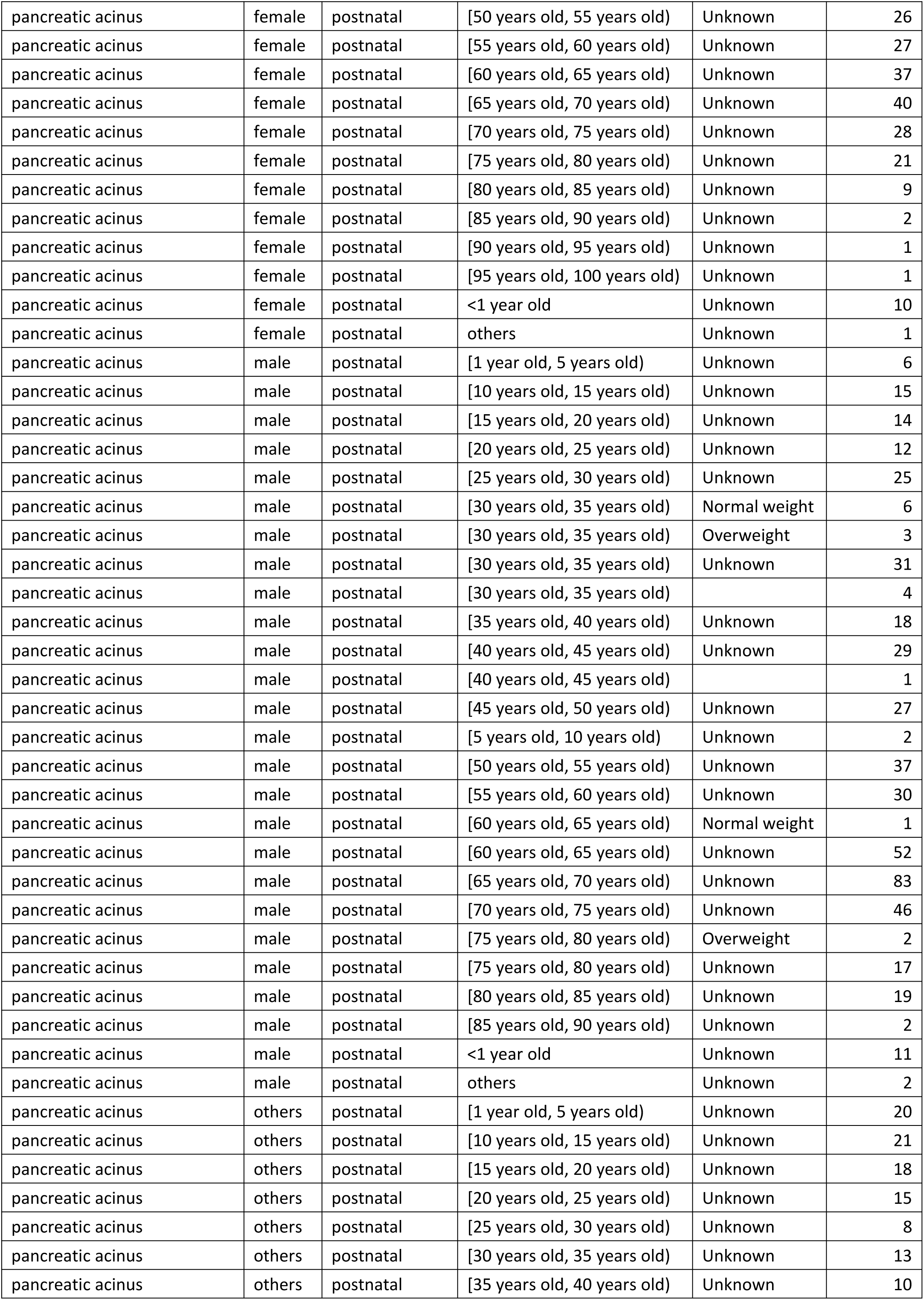
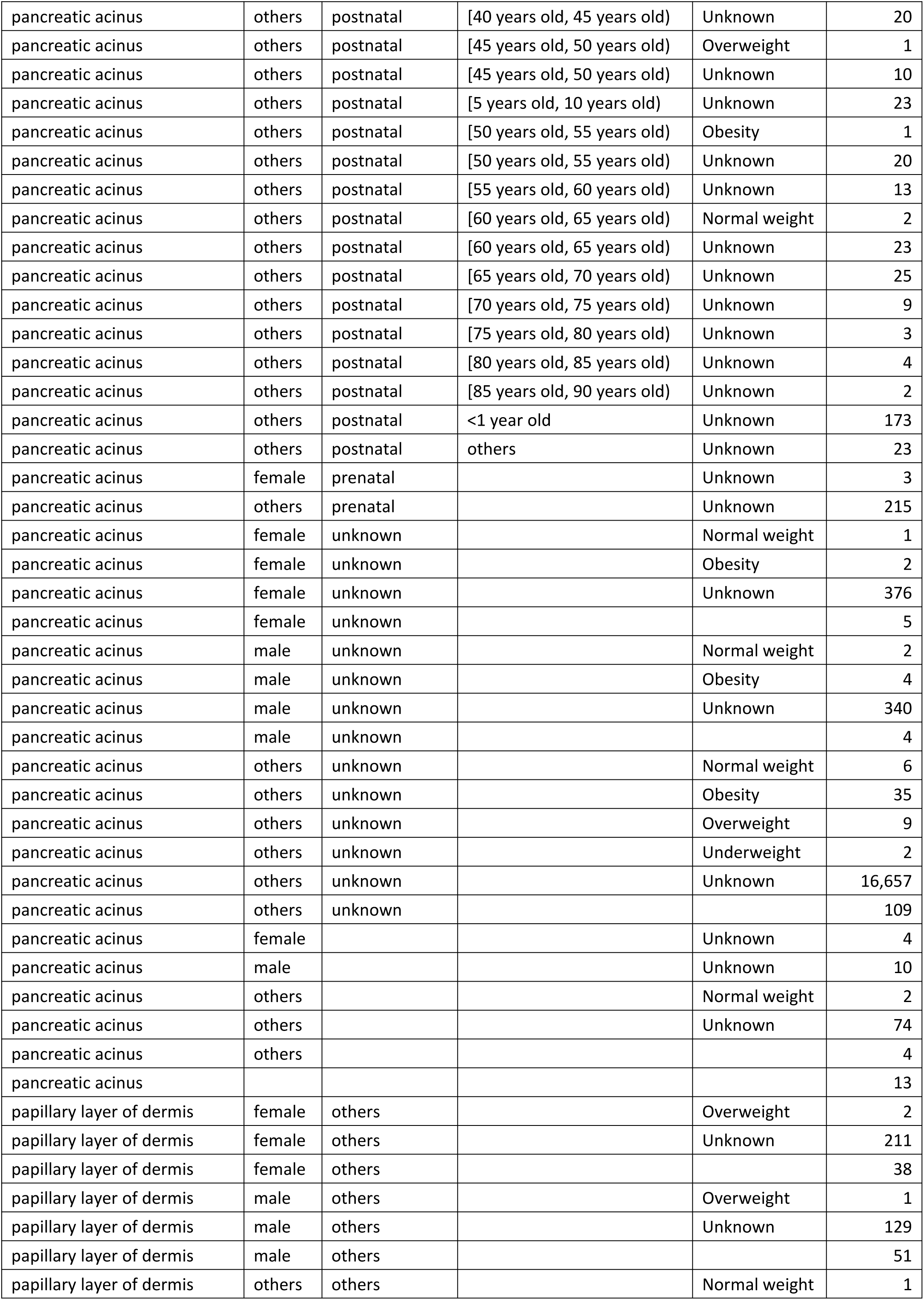
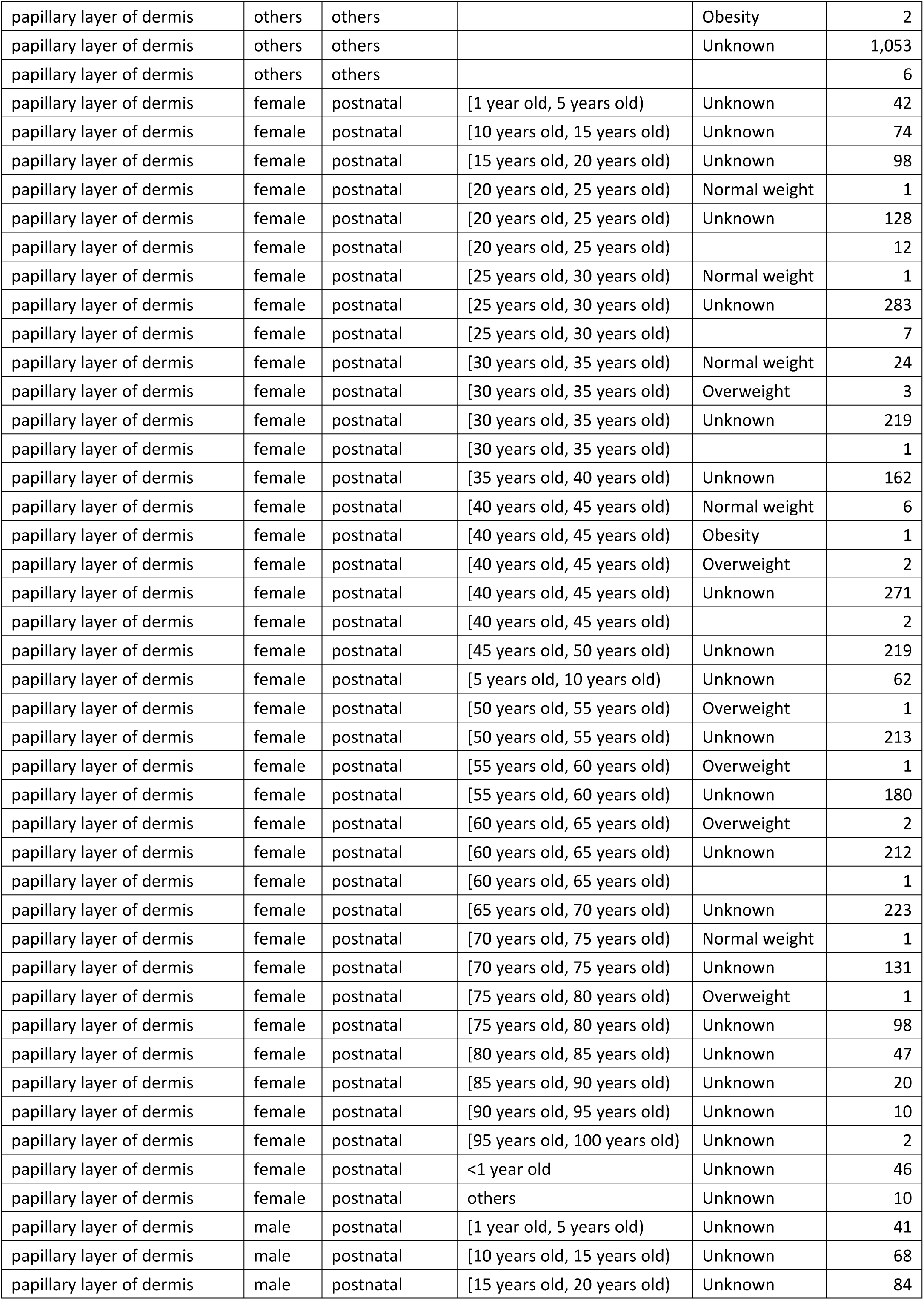
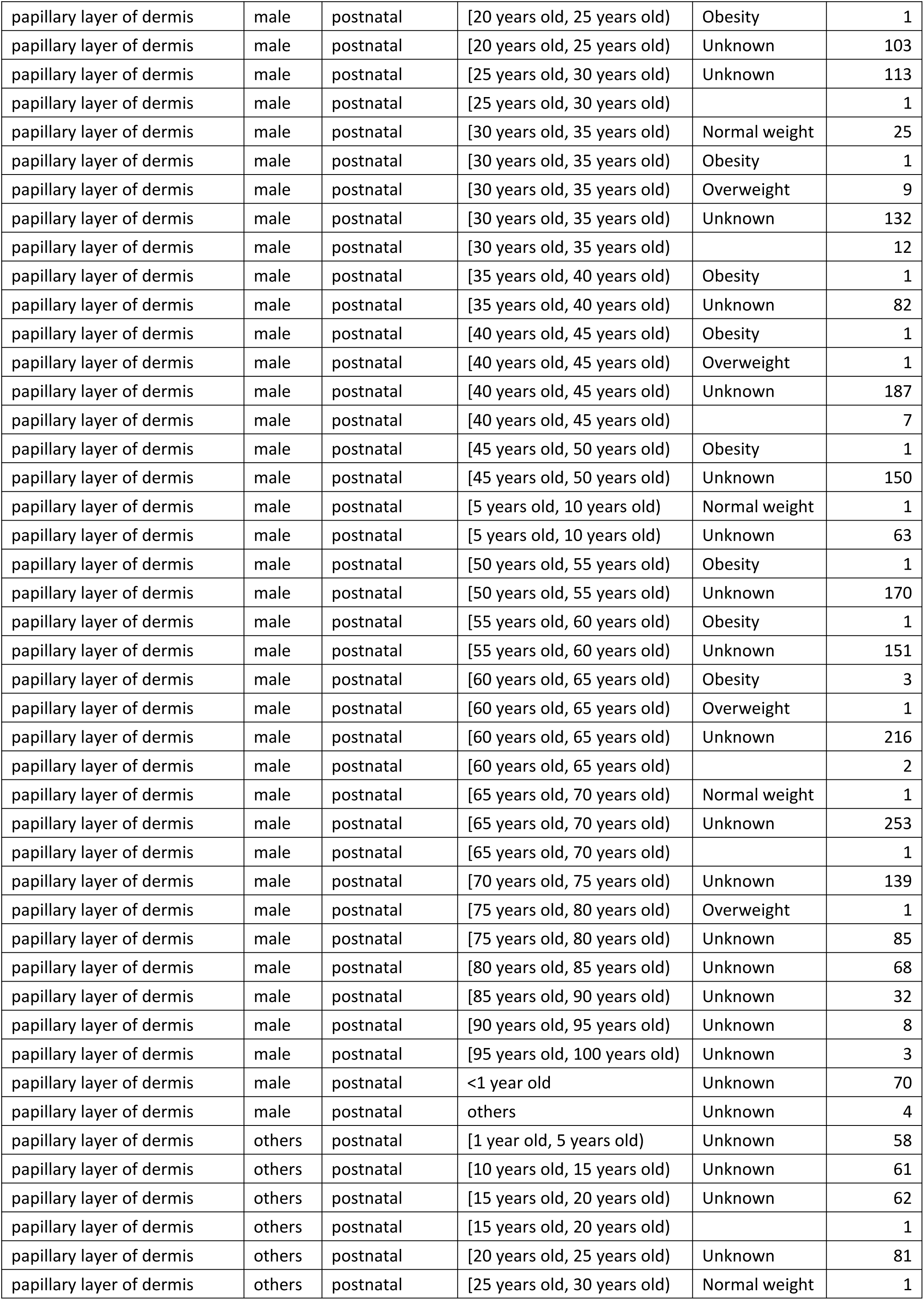
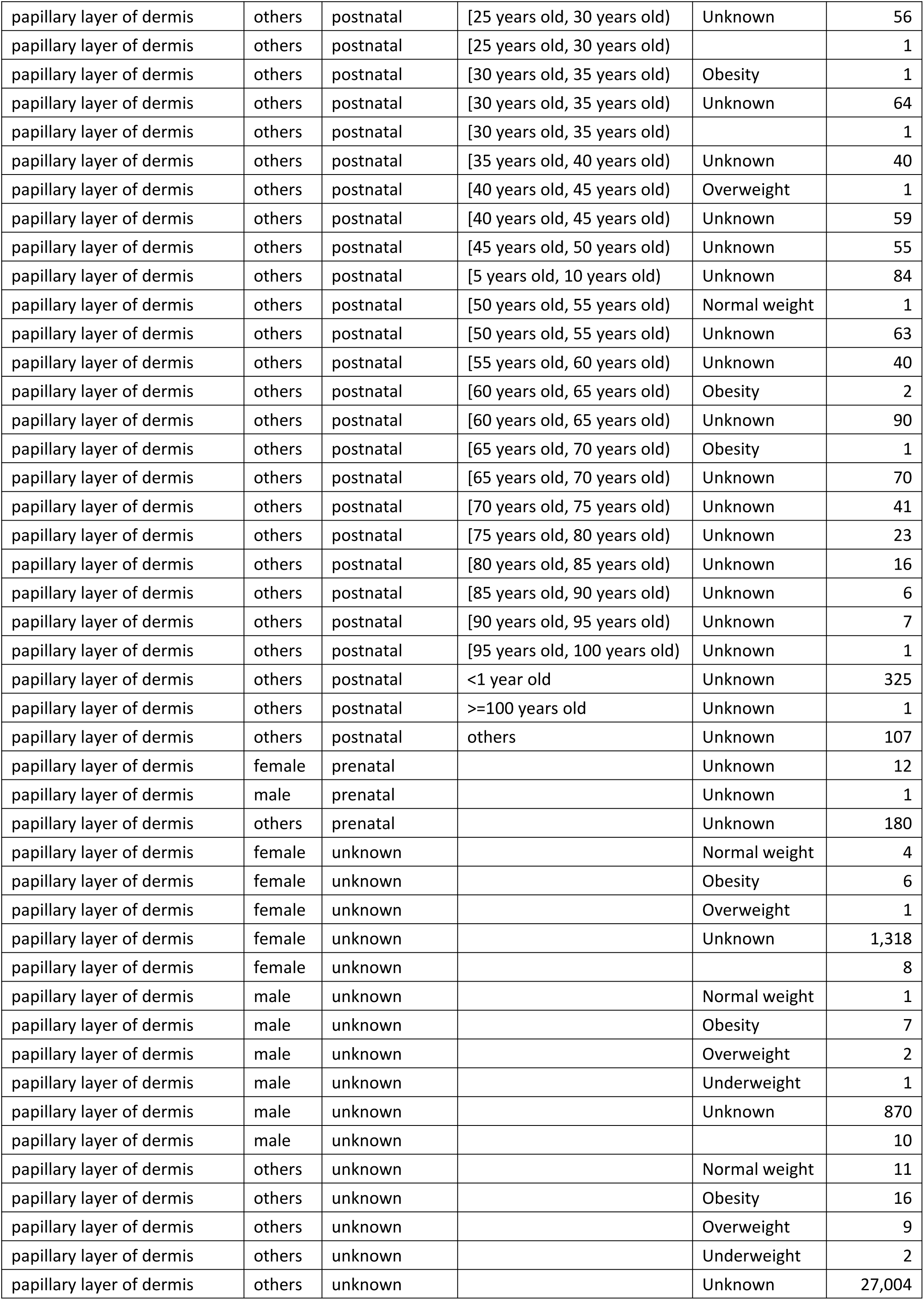
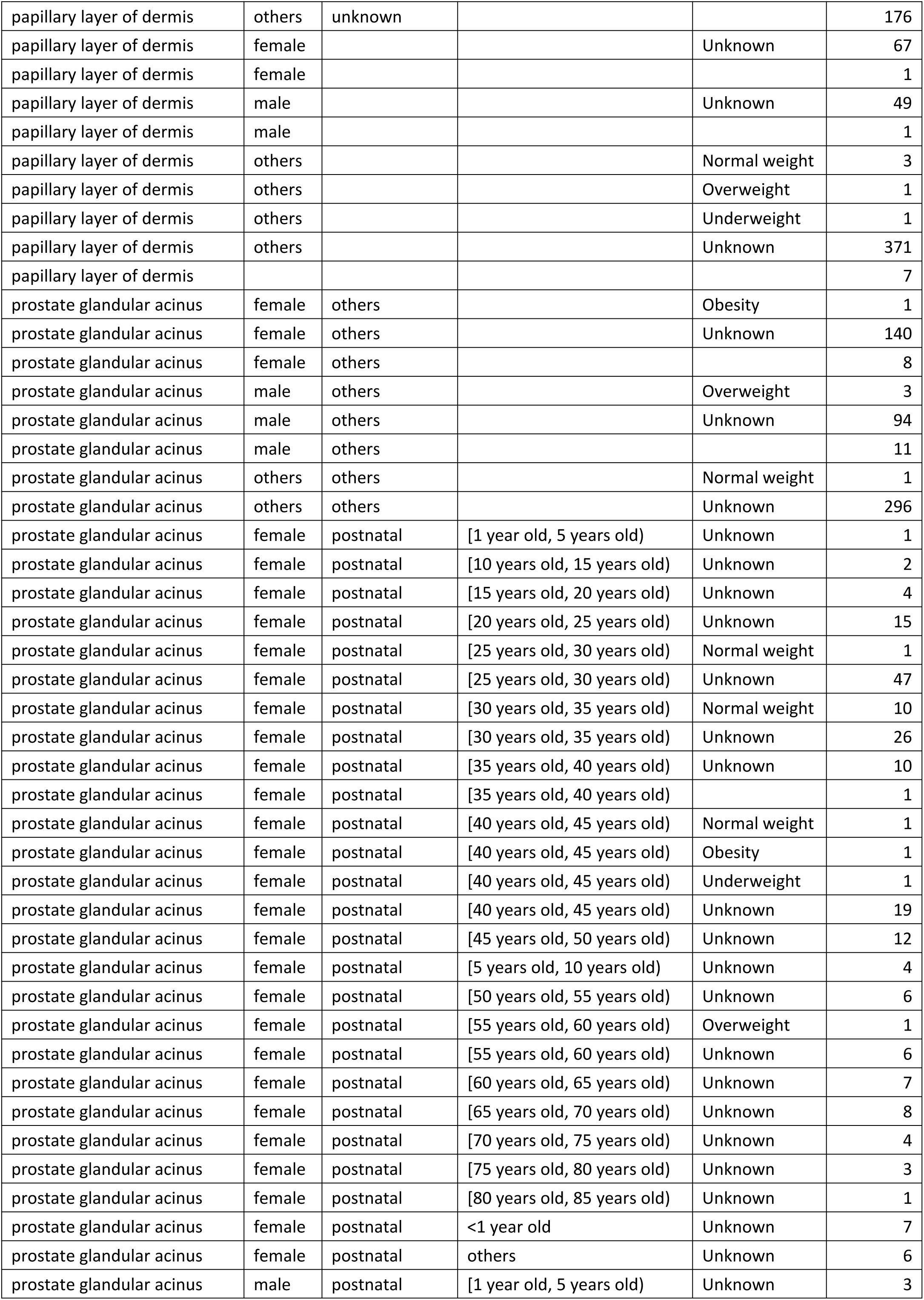
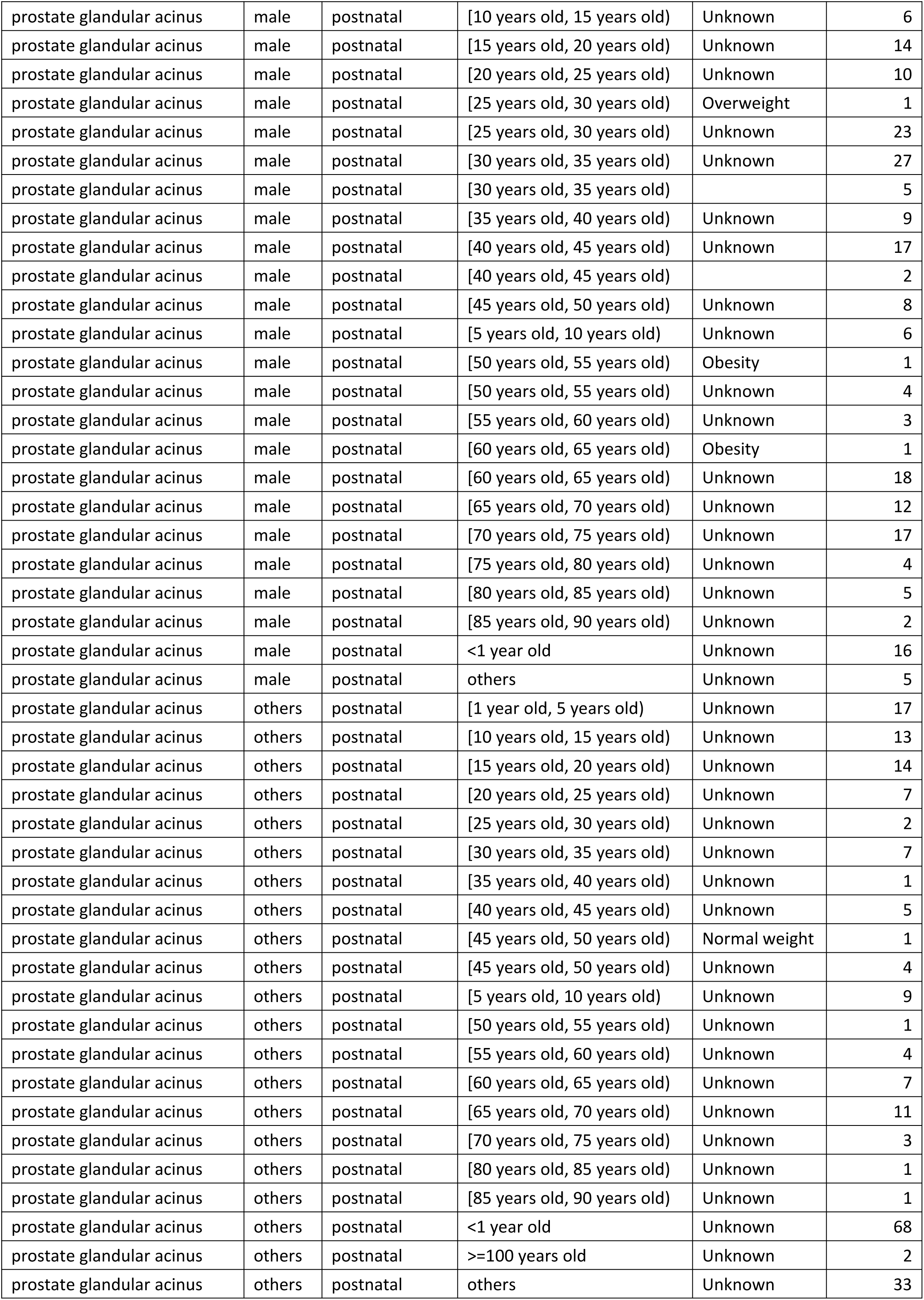
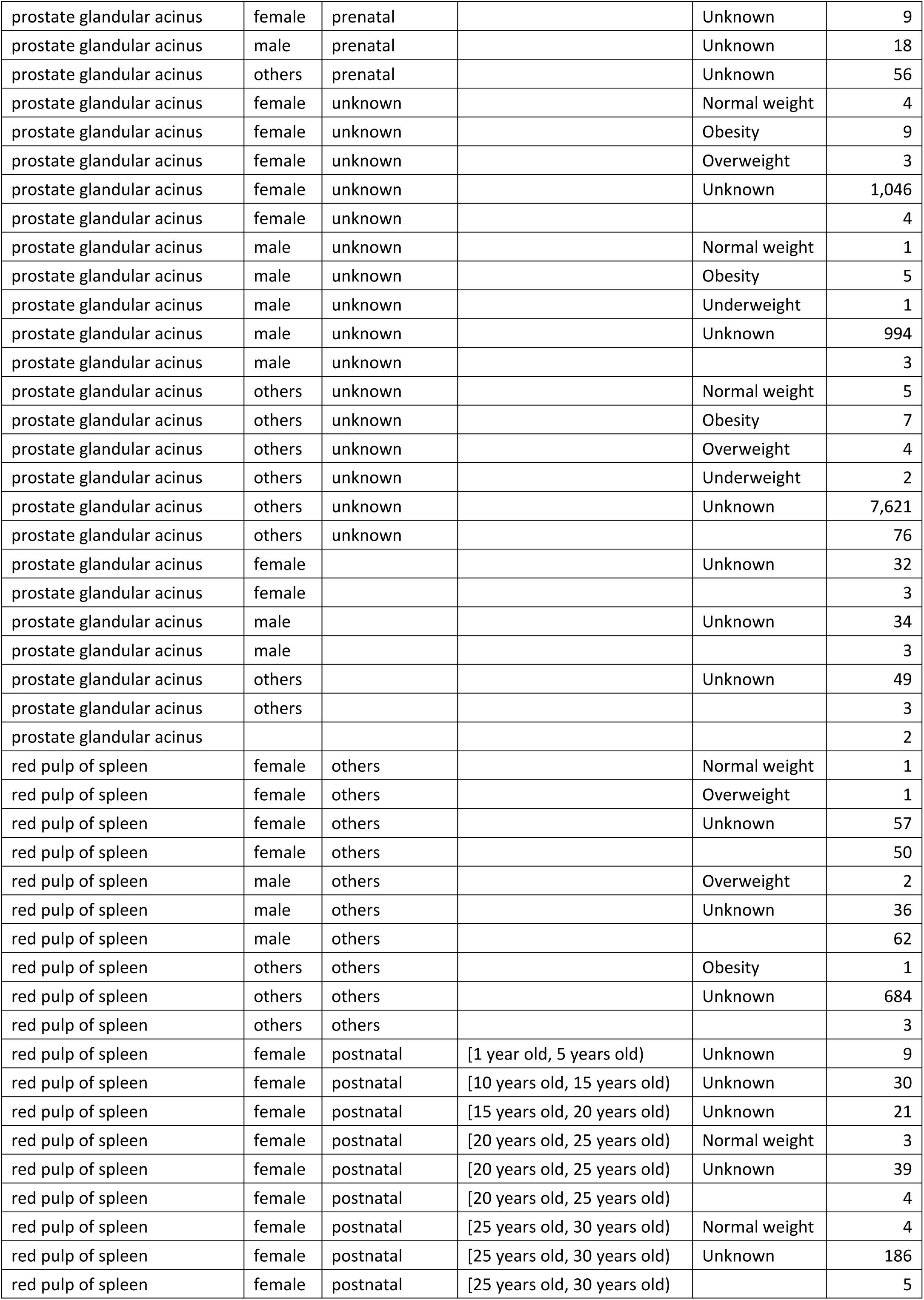
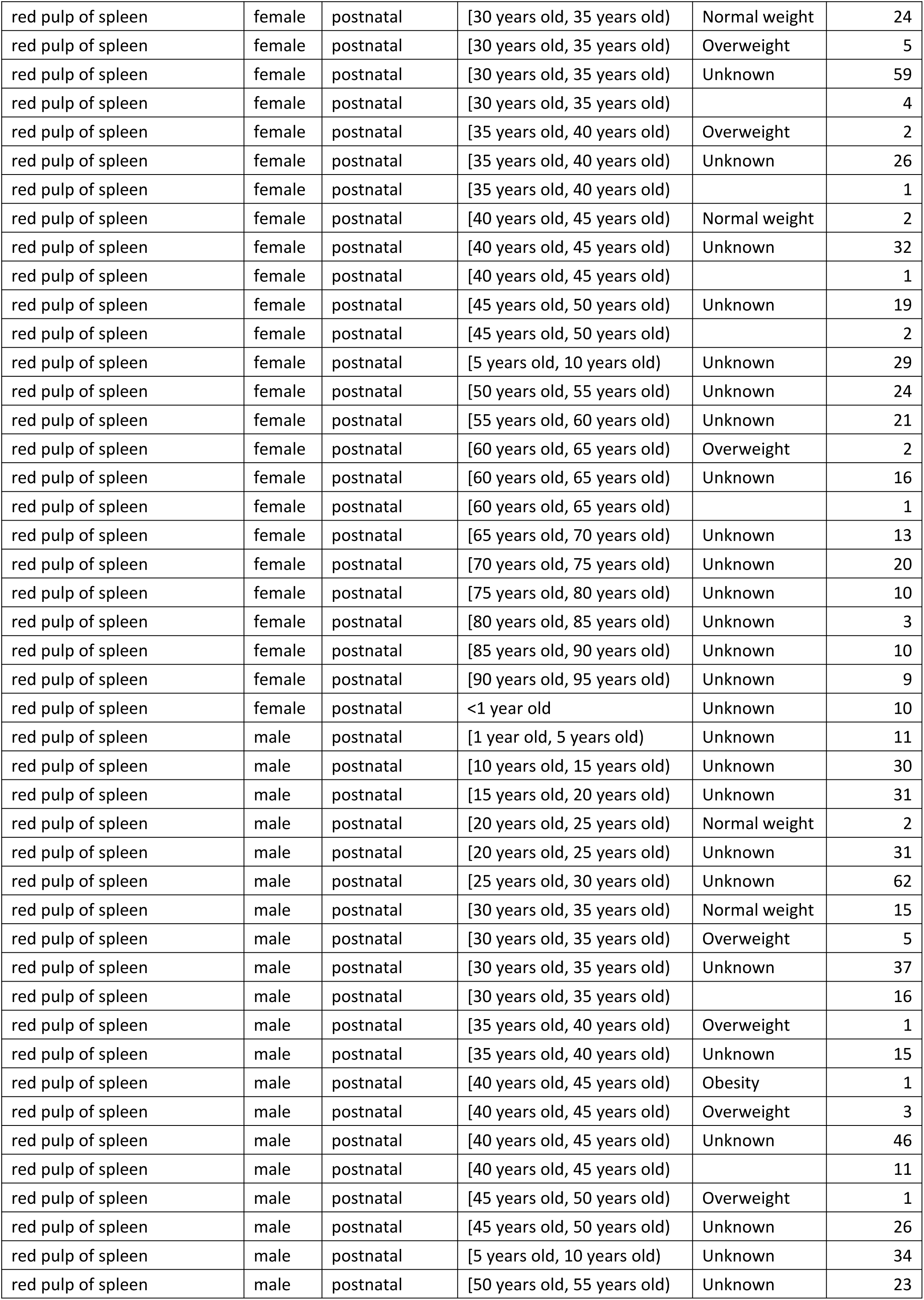
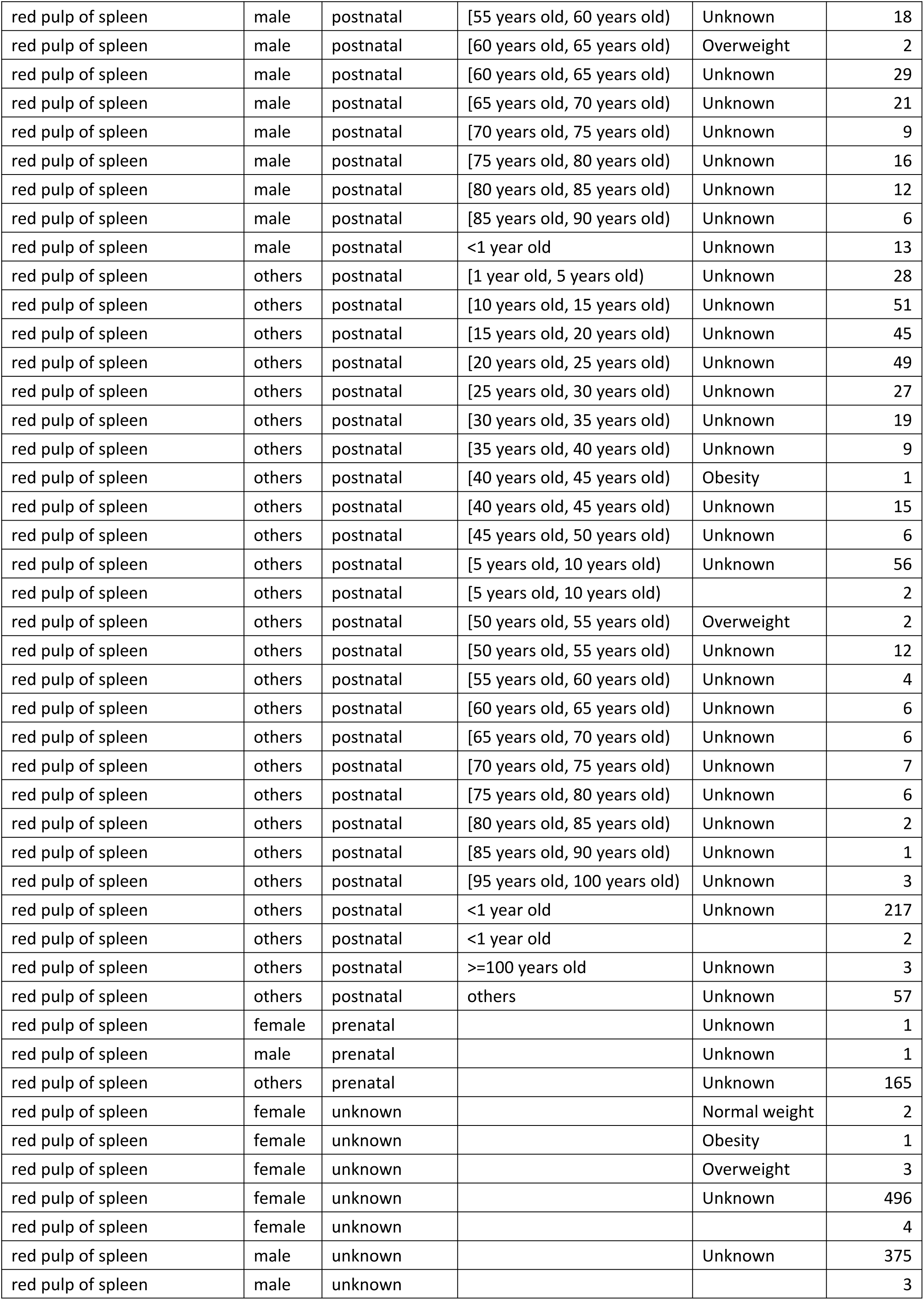
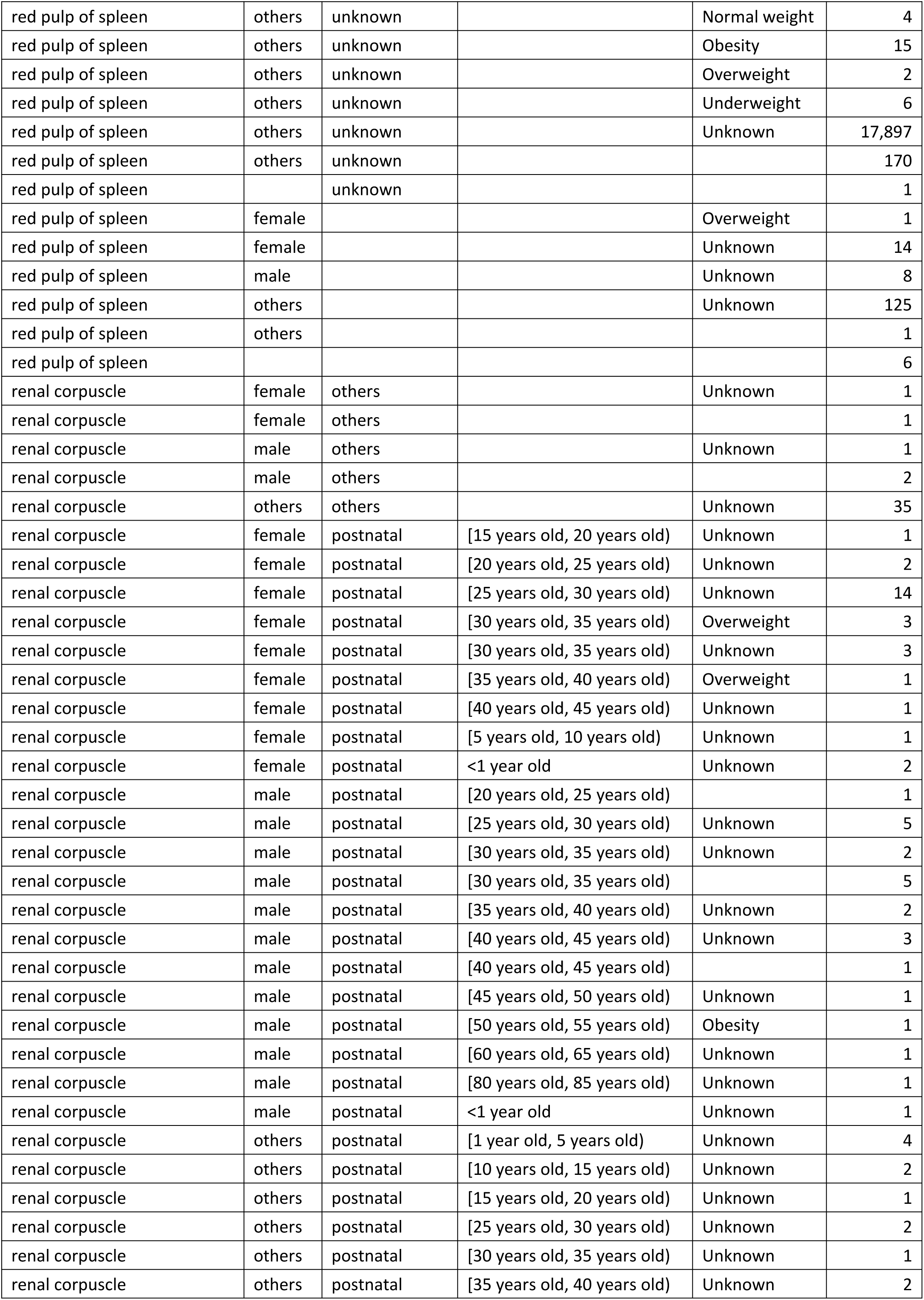
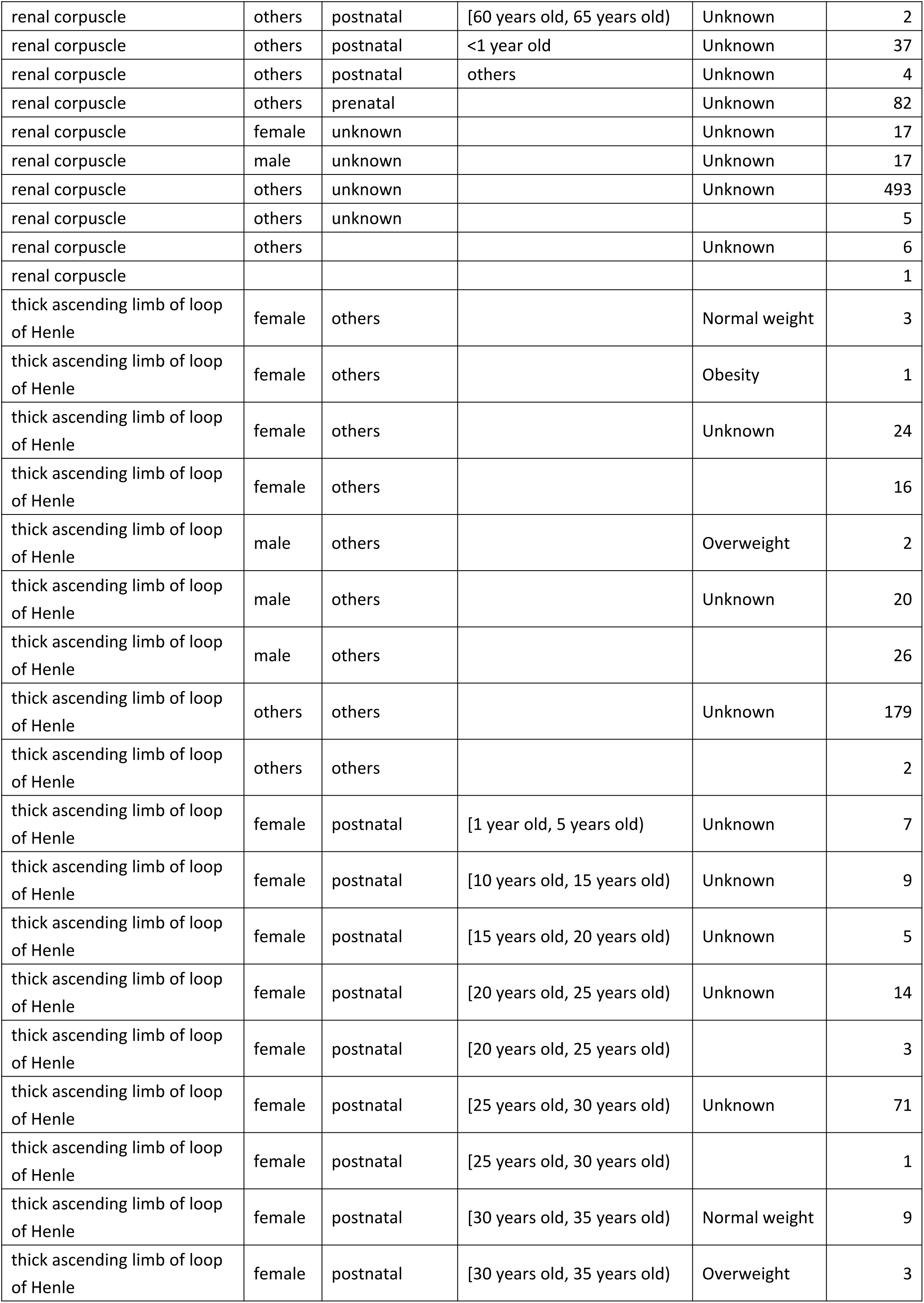
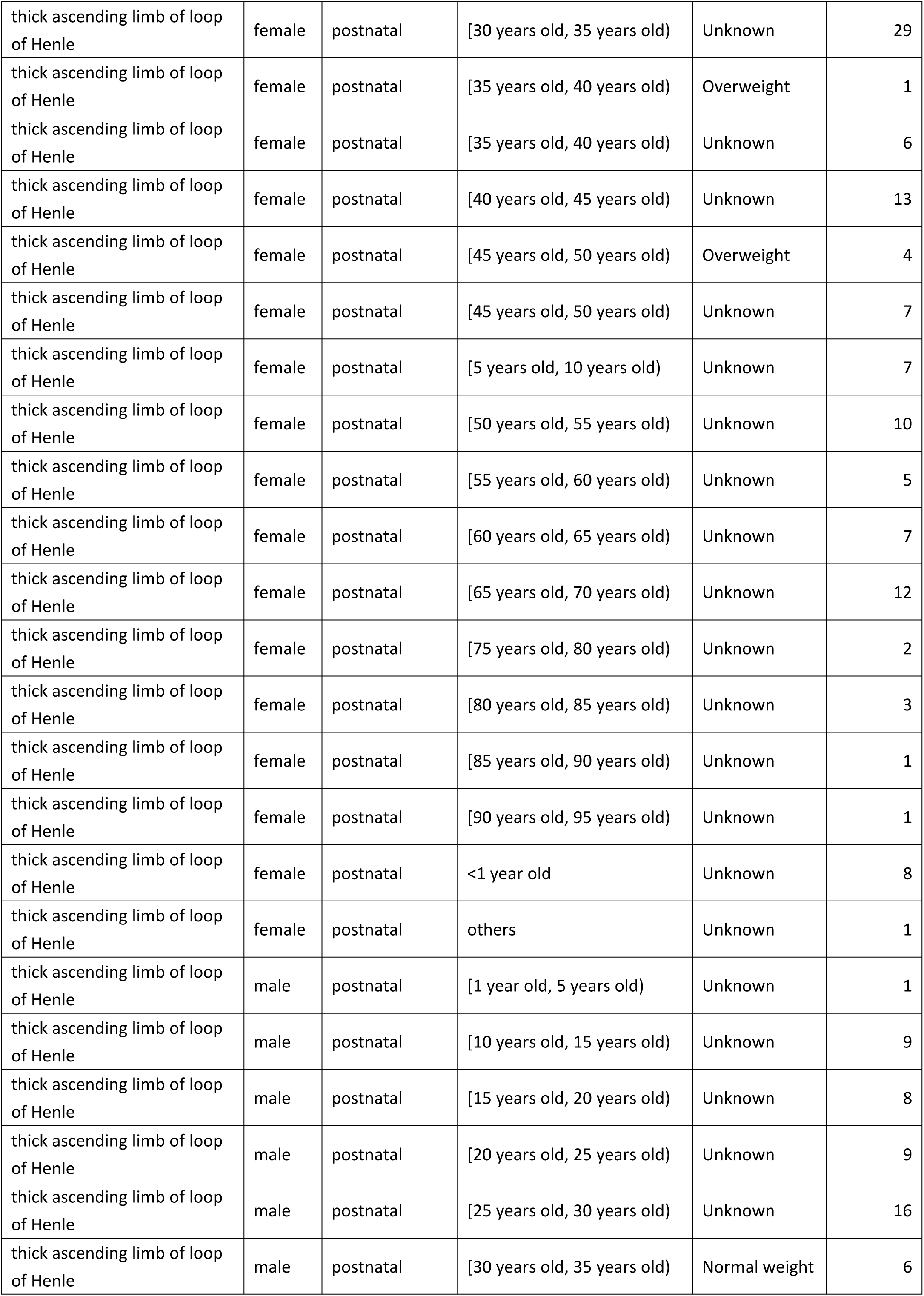
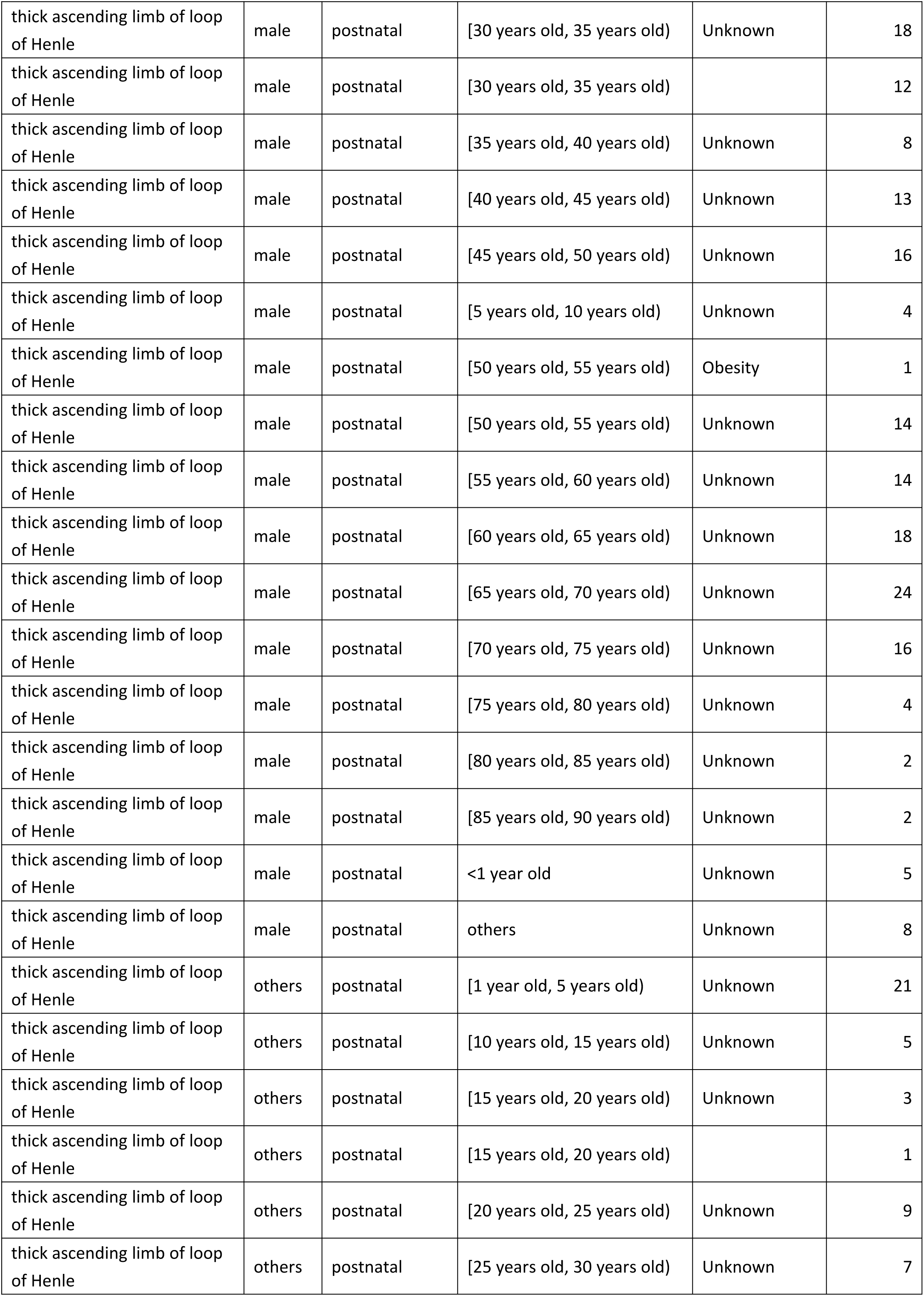
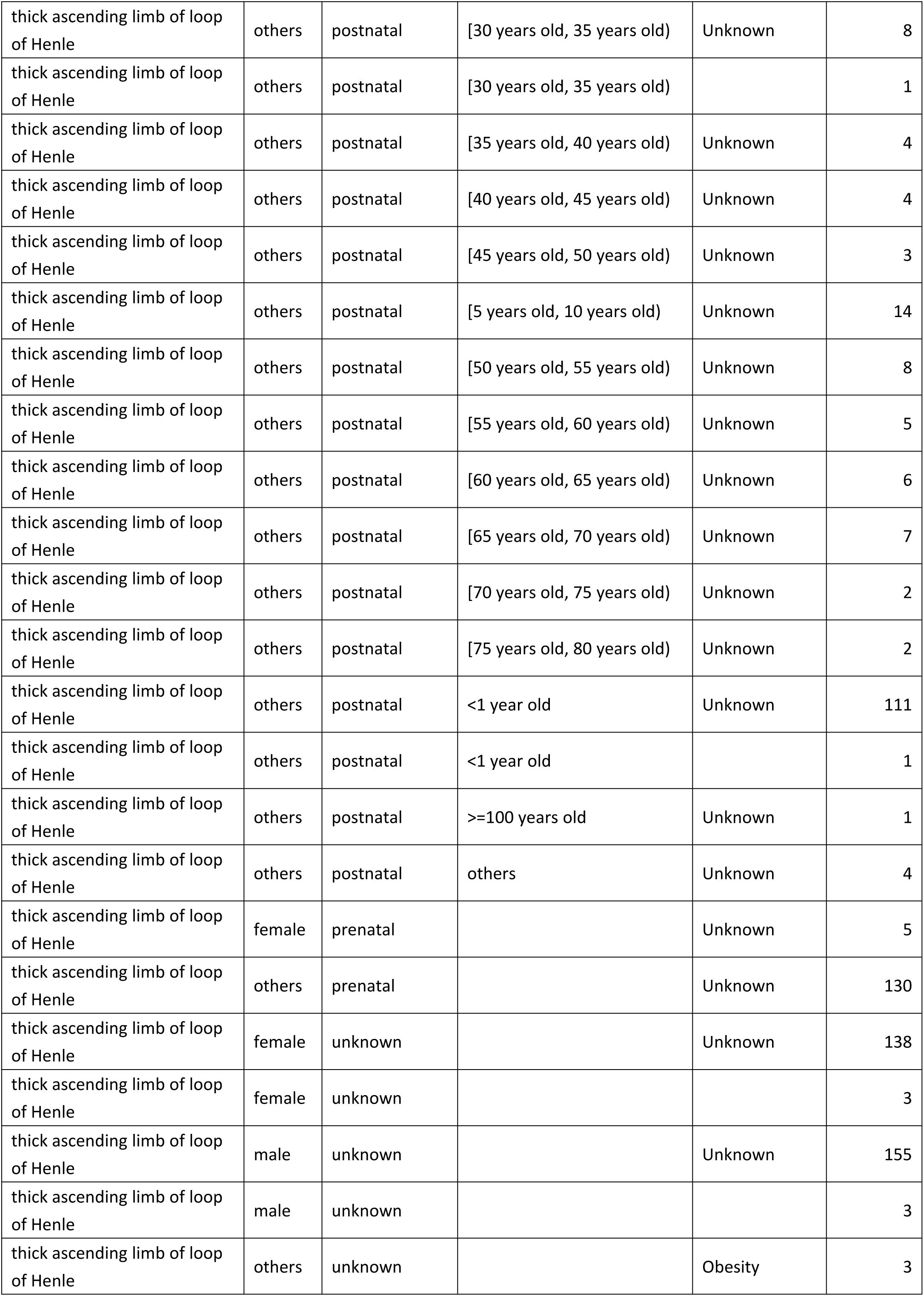
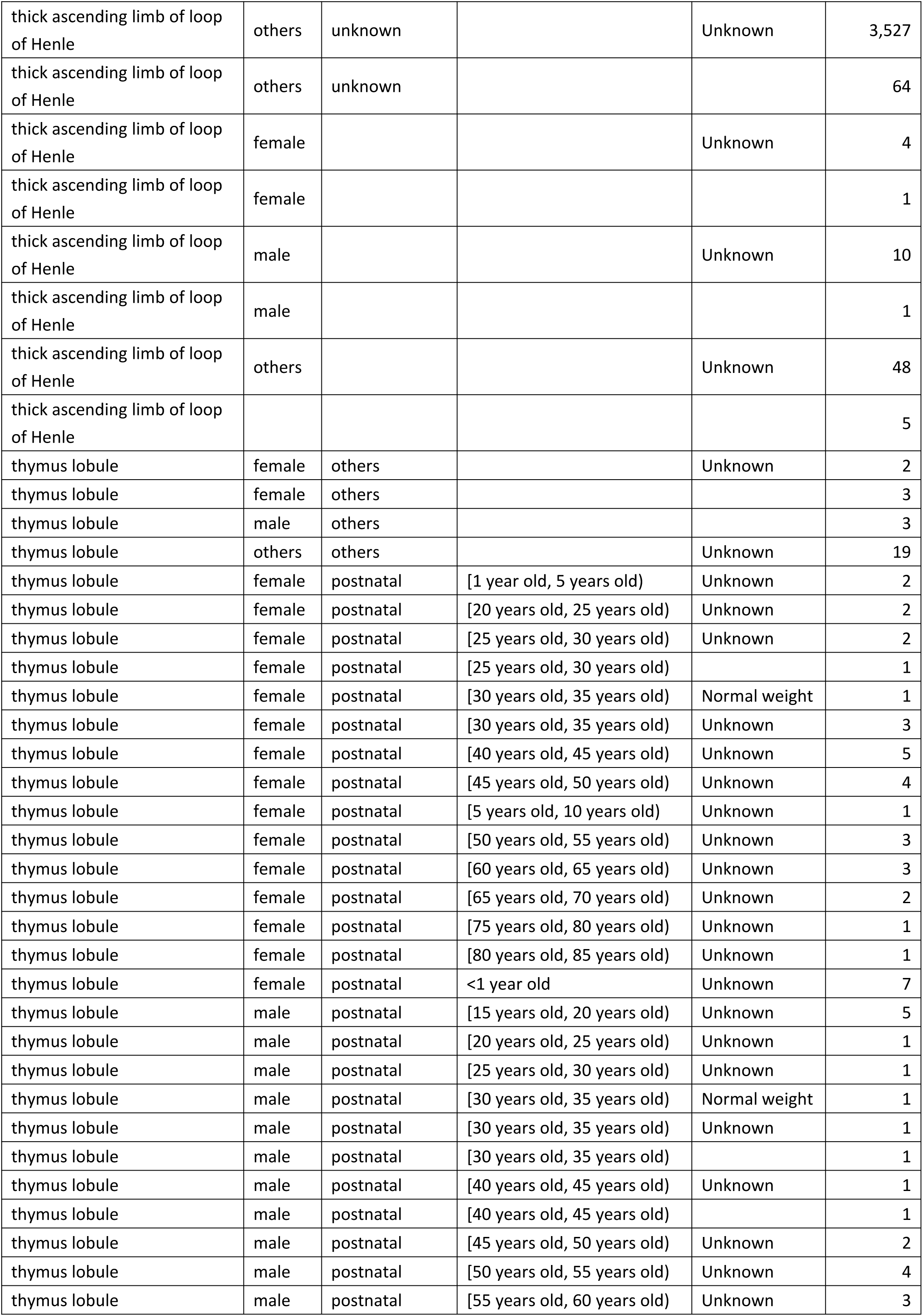
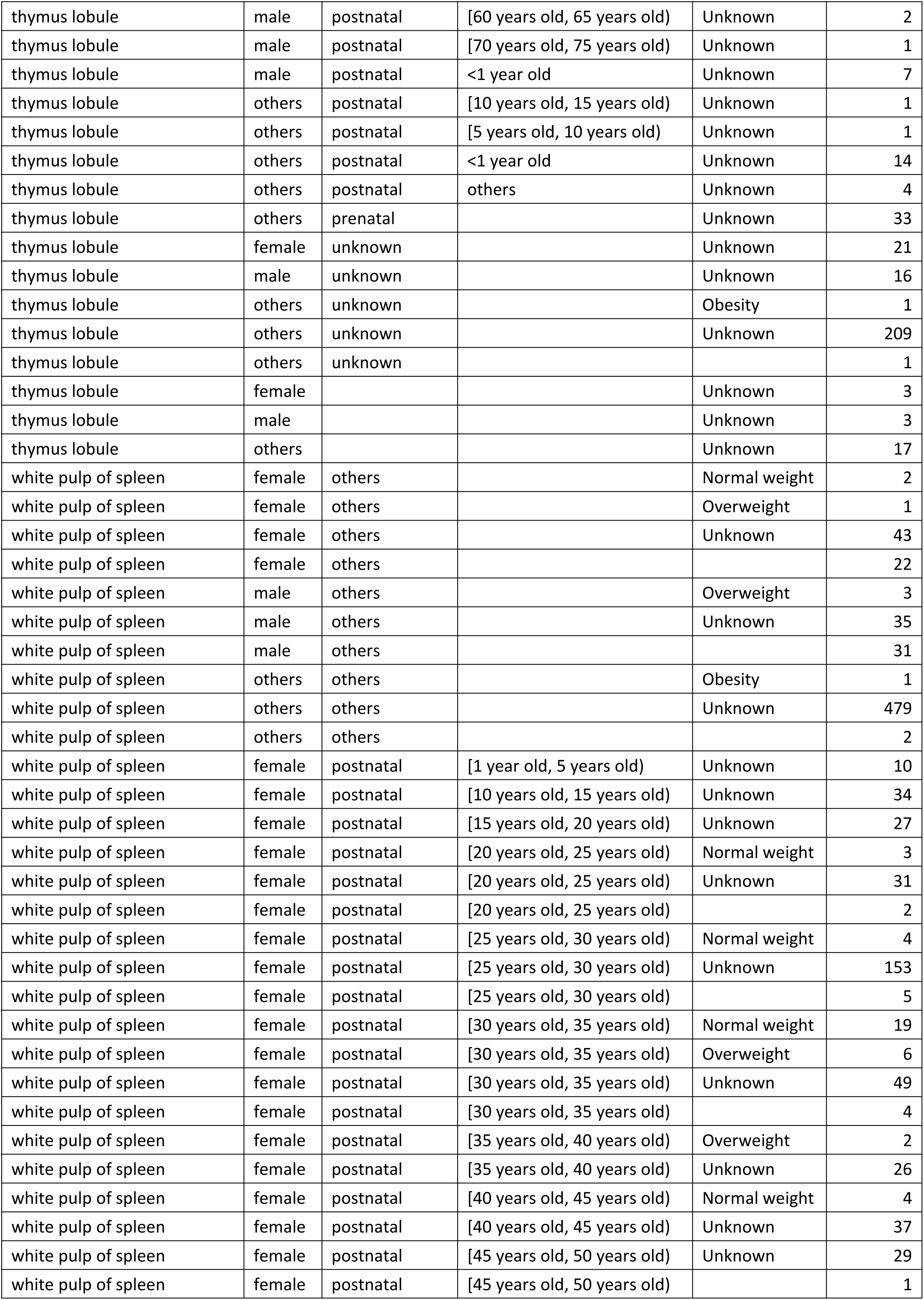
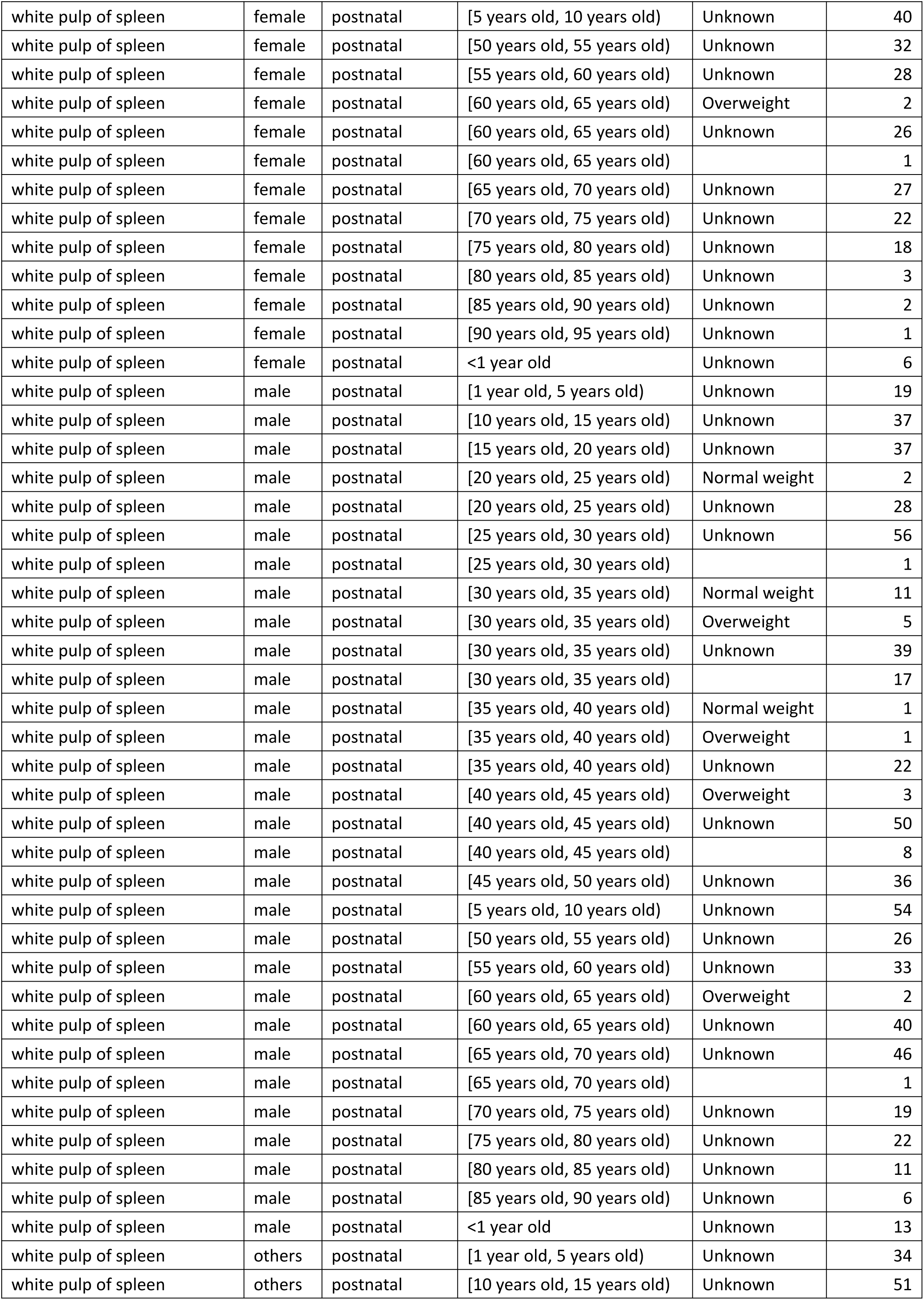
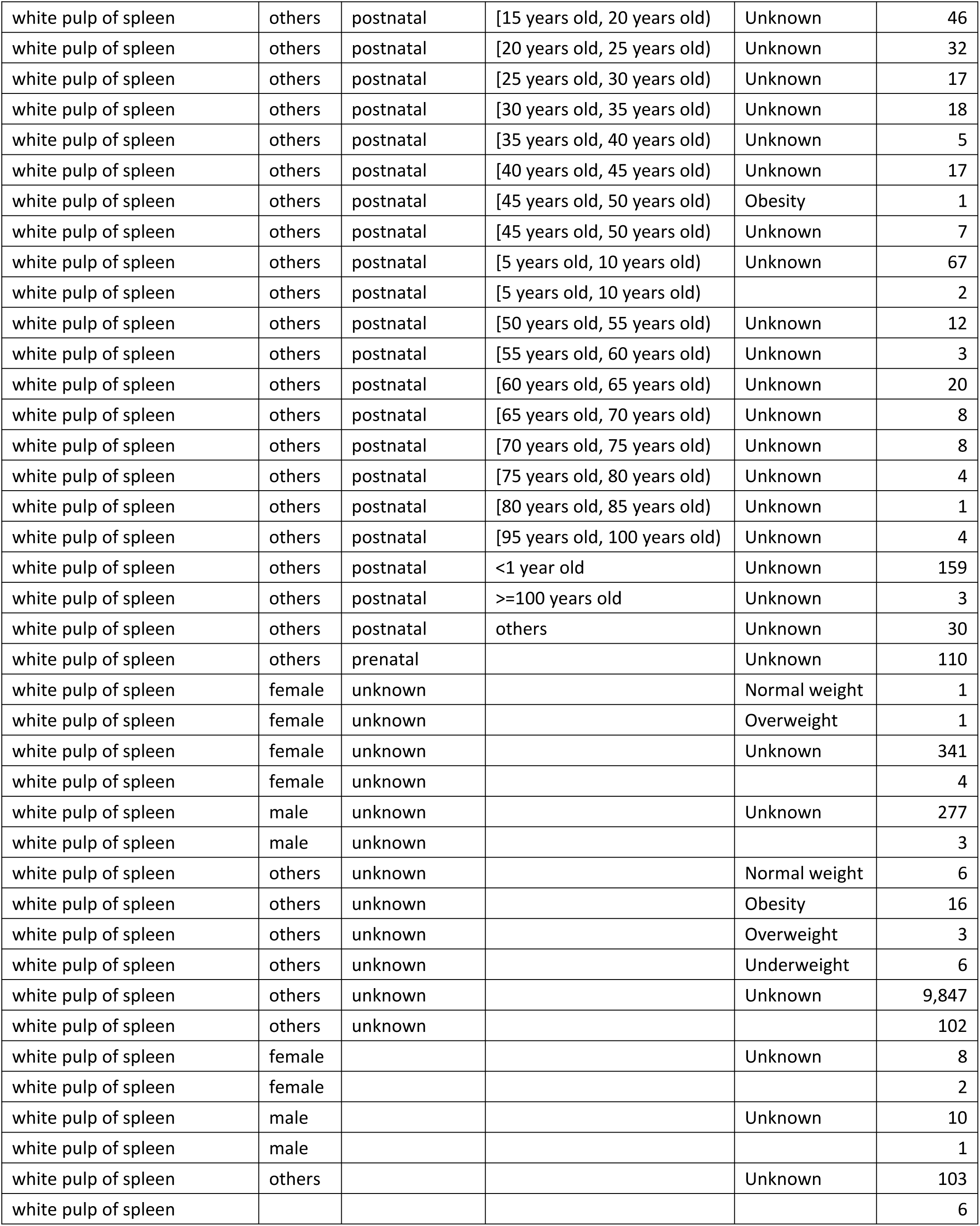
The number of donor records in Homo sapiens for 22 FTUs.

**Supplementary Table 26.**
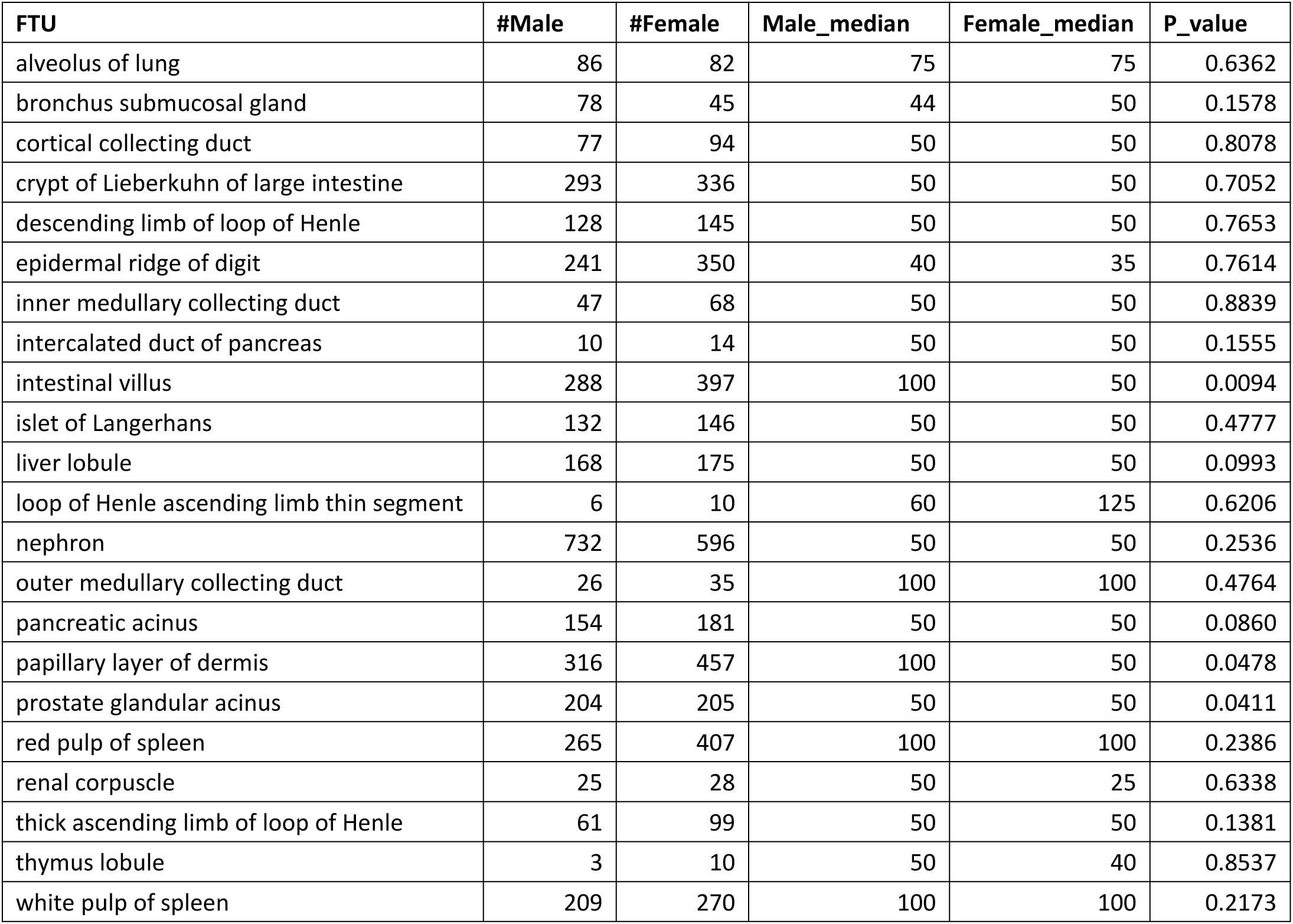
Donor summary in difference sex groups across FTUs.

**Supplementary Table 27.**
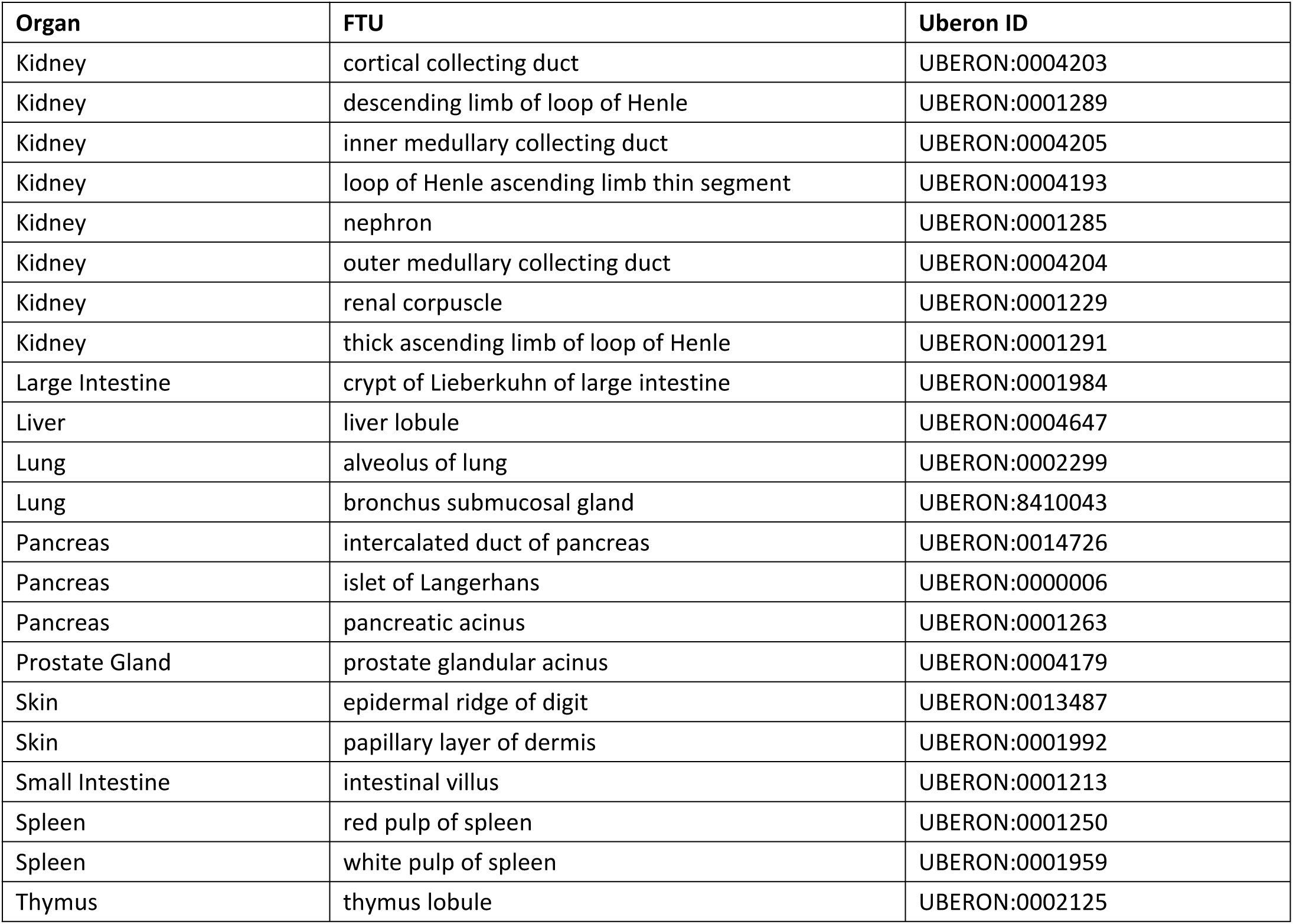
List of 22 FTUs in 10 organs.

**Supplementary Table 28.**
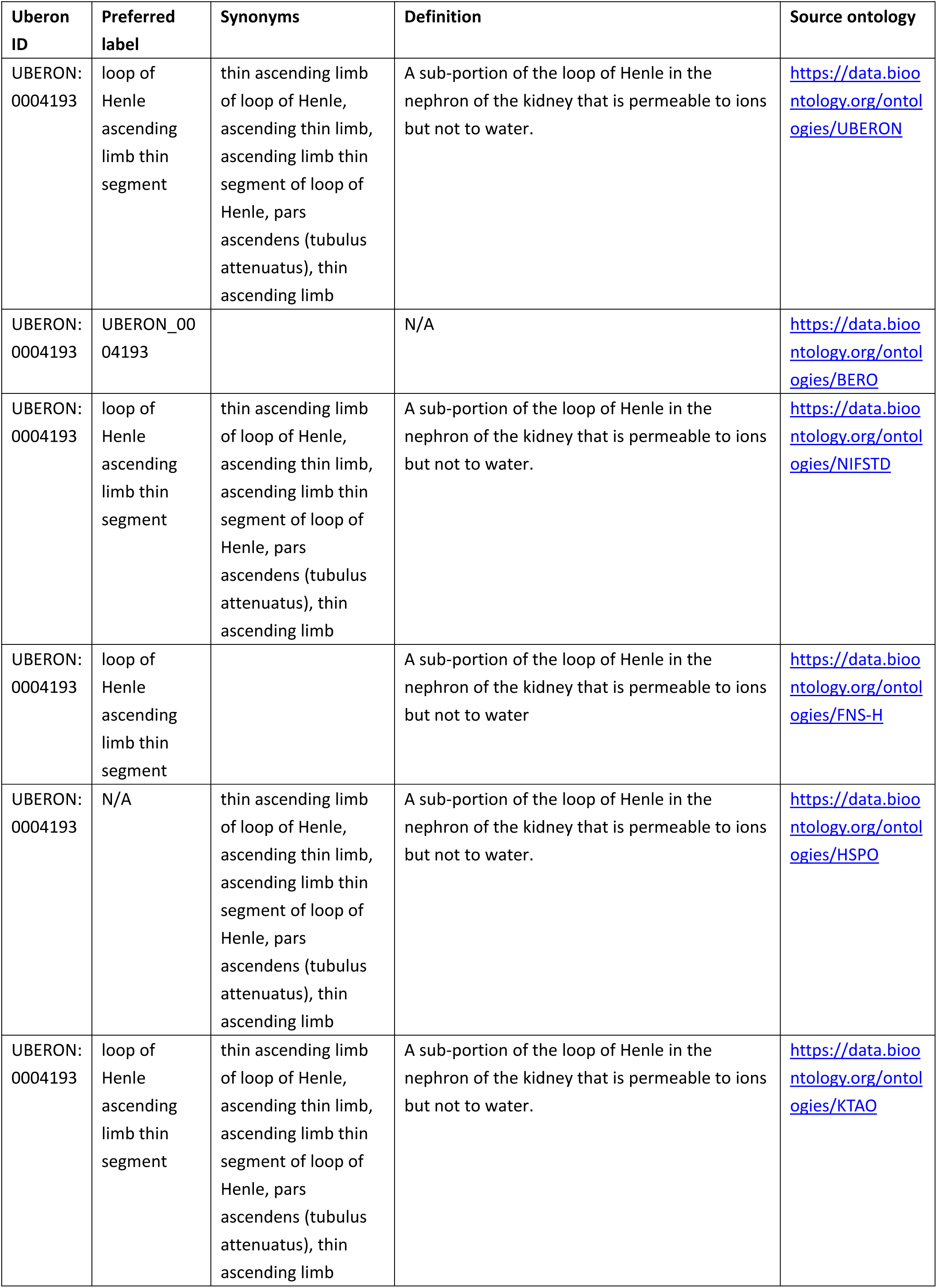
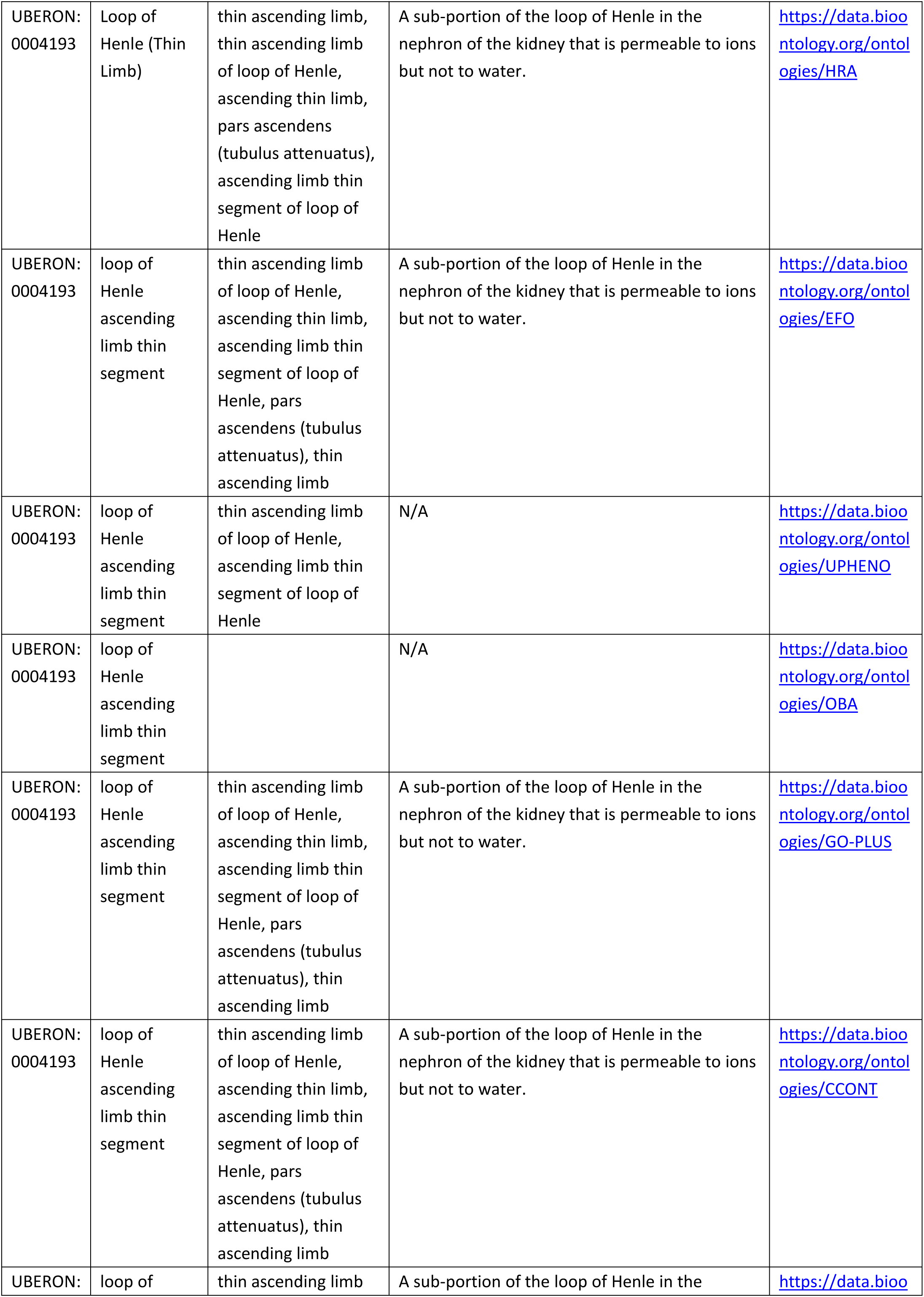
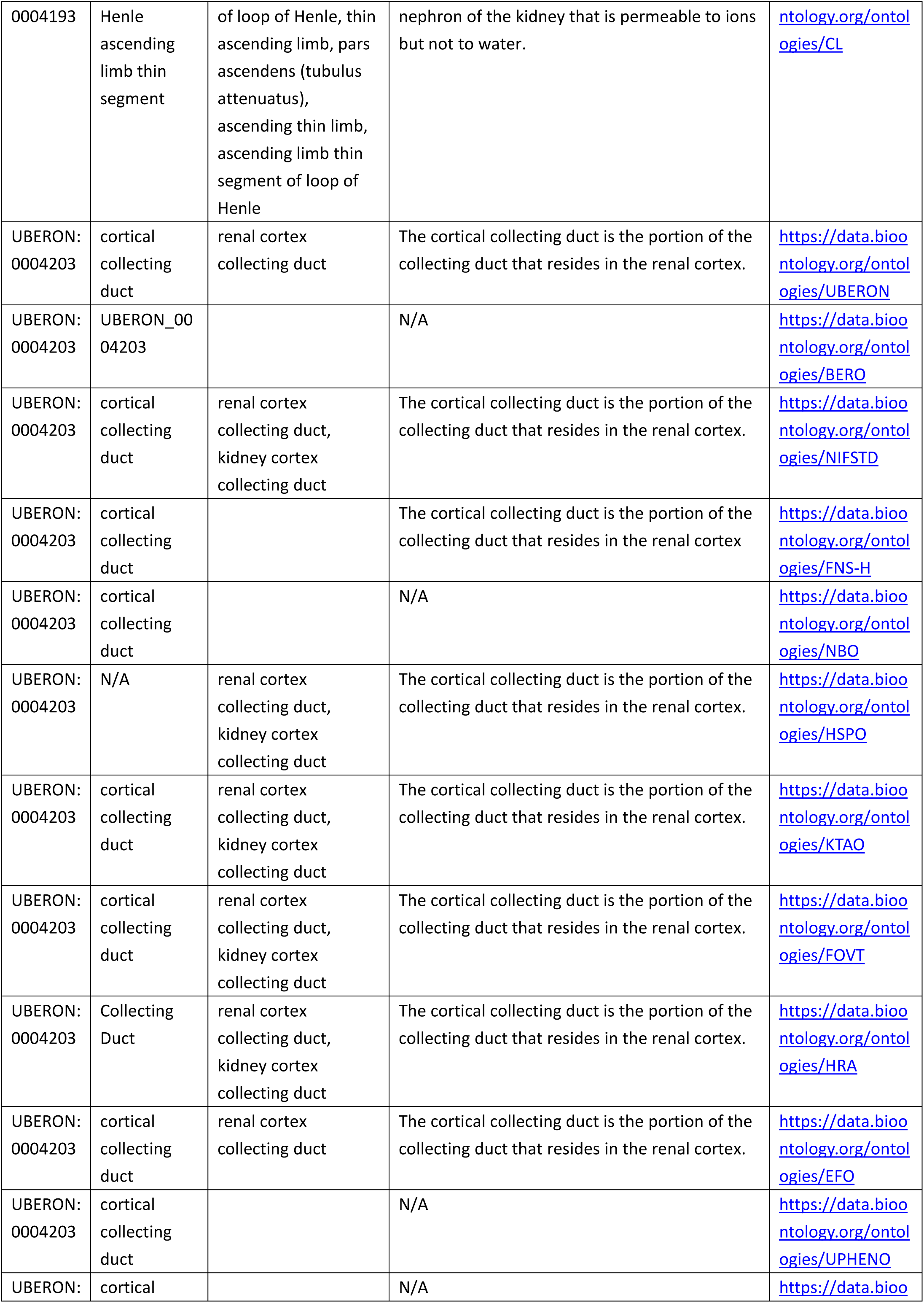
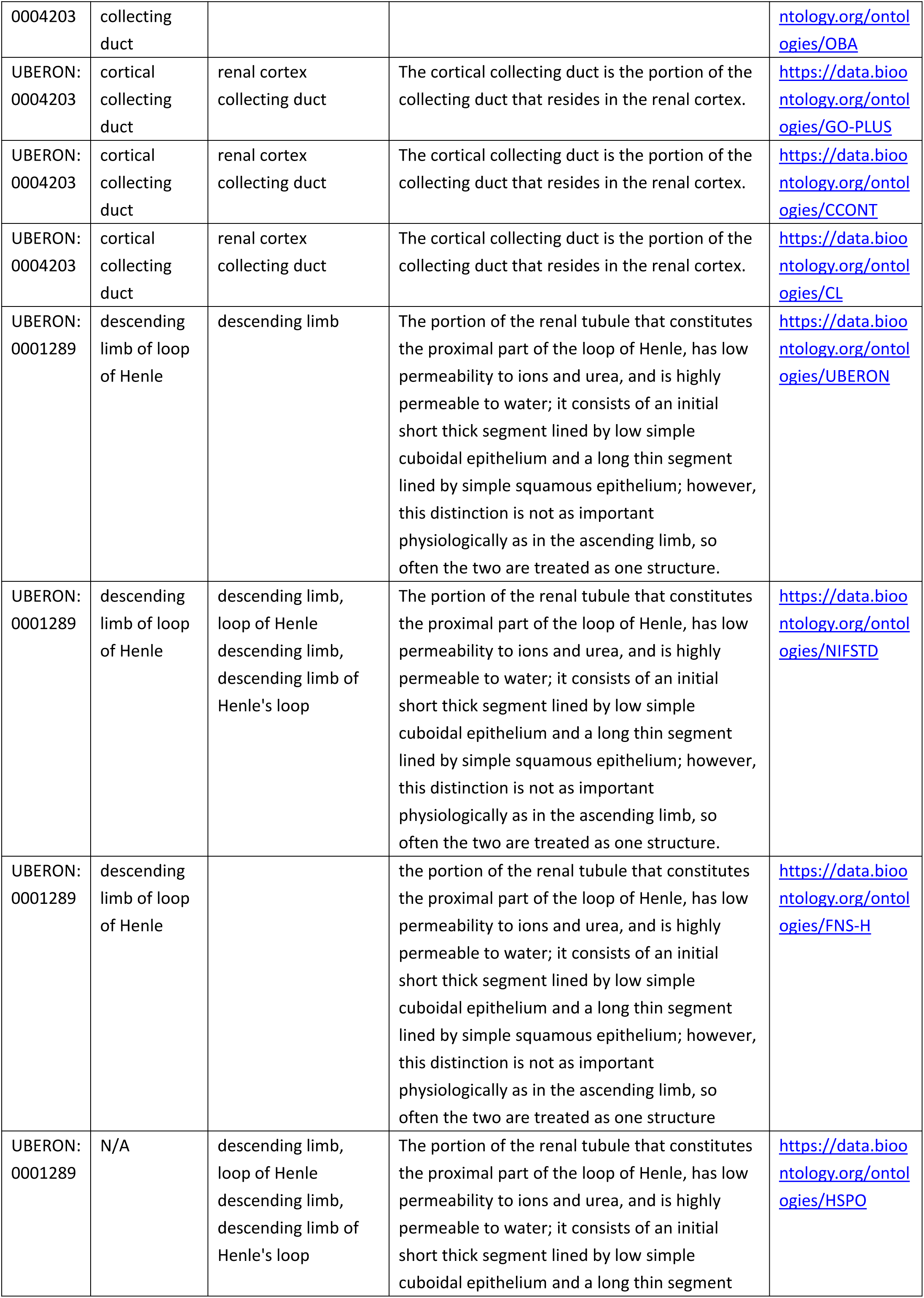
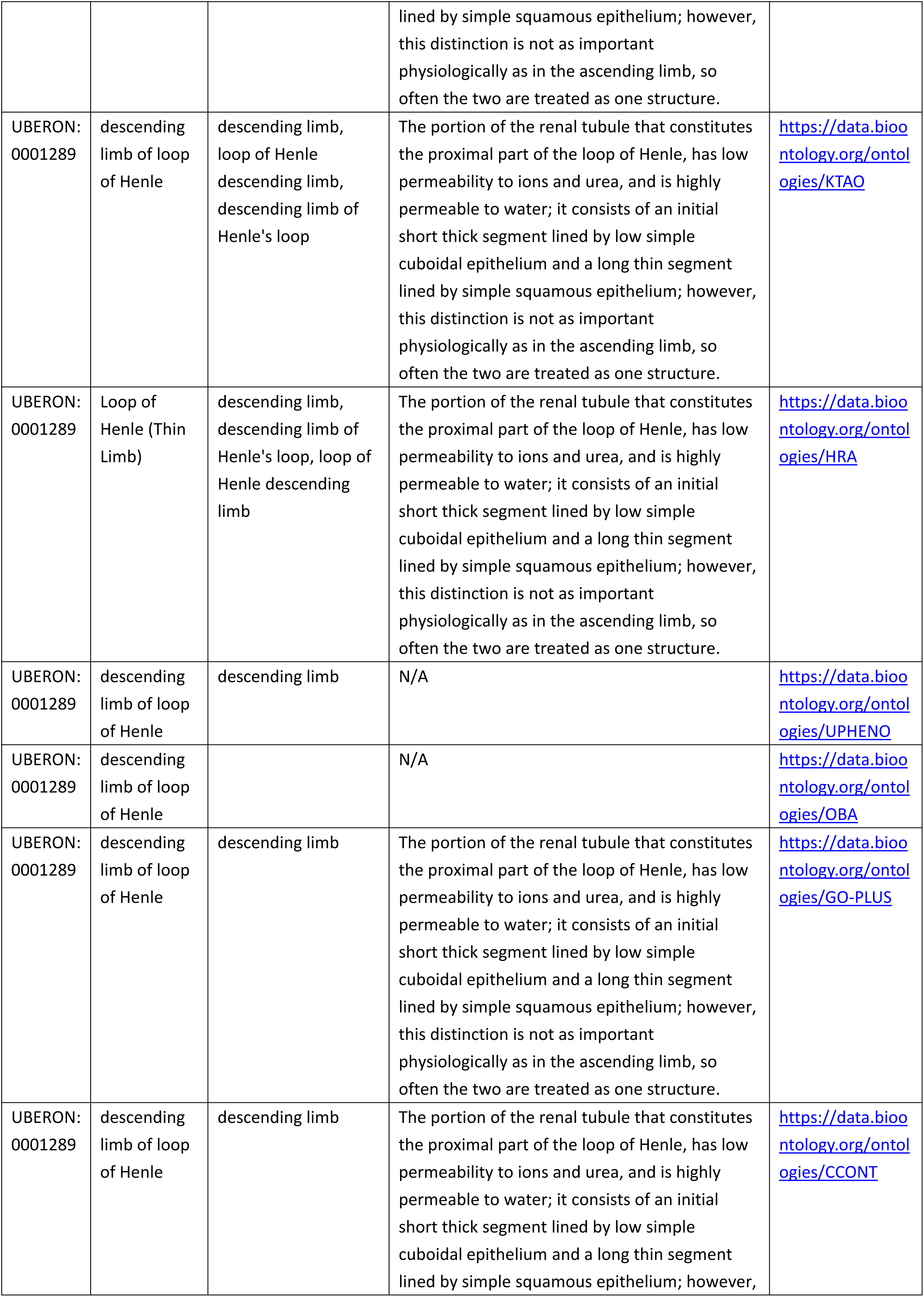
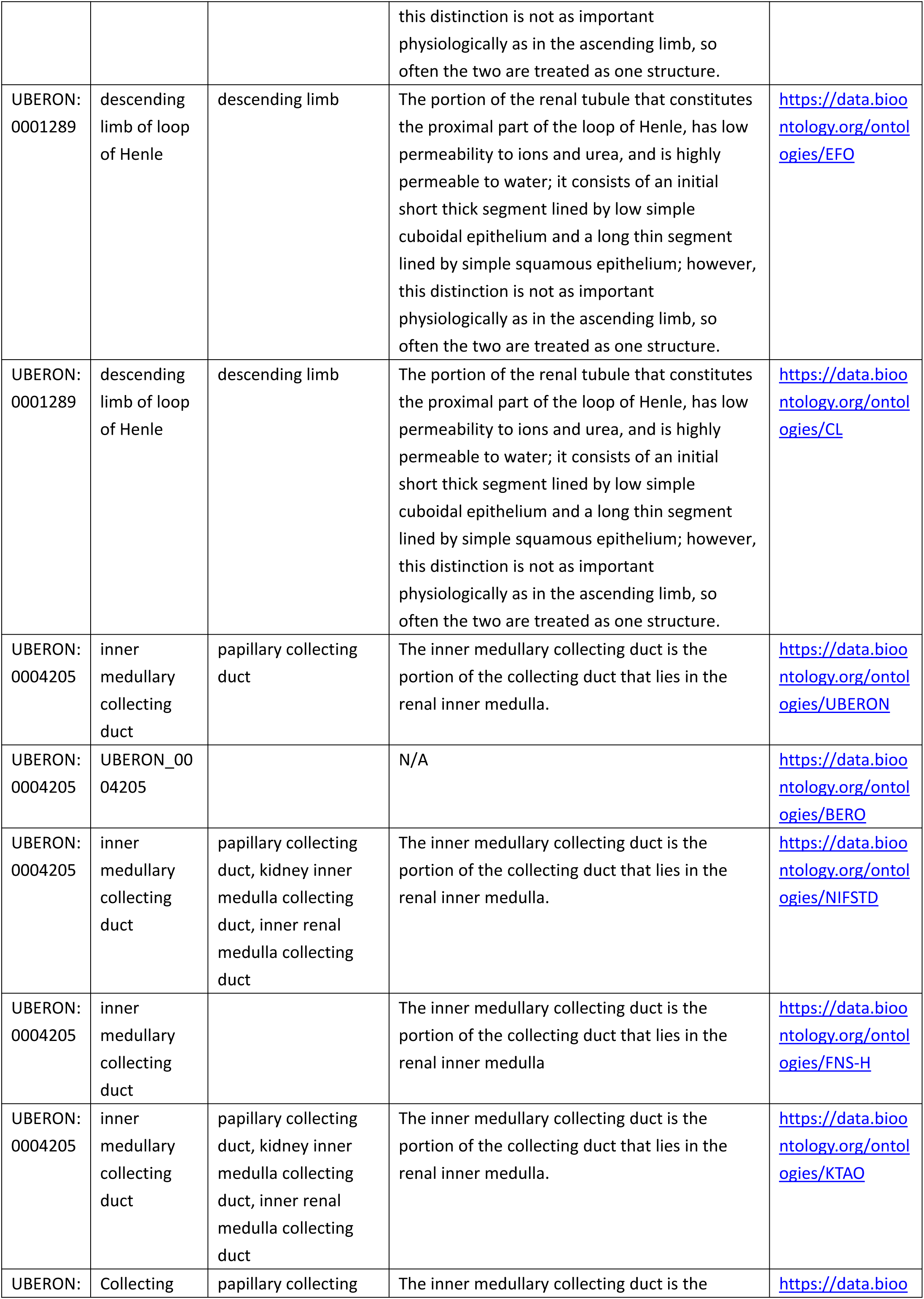
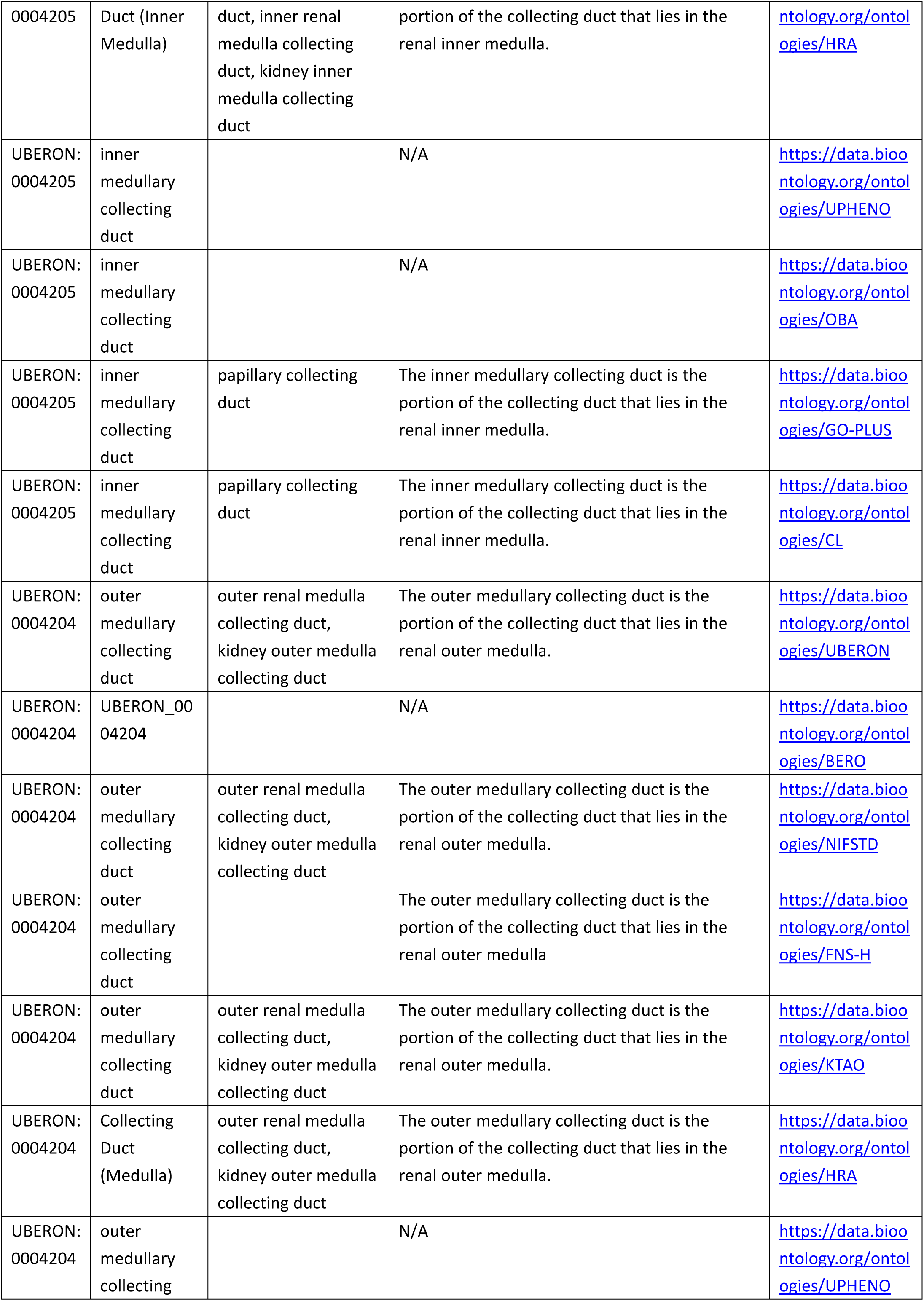
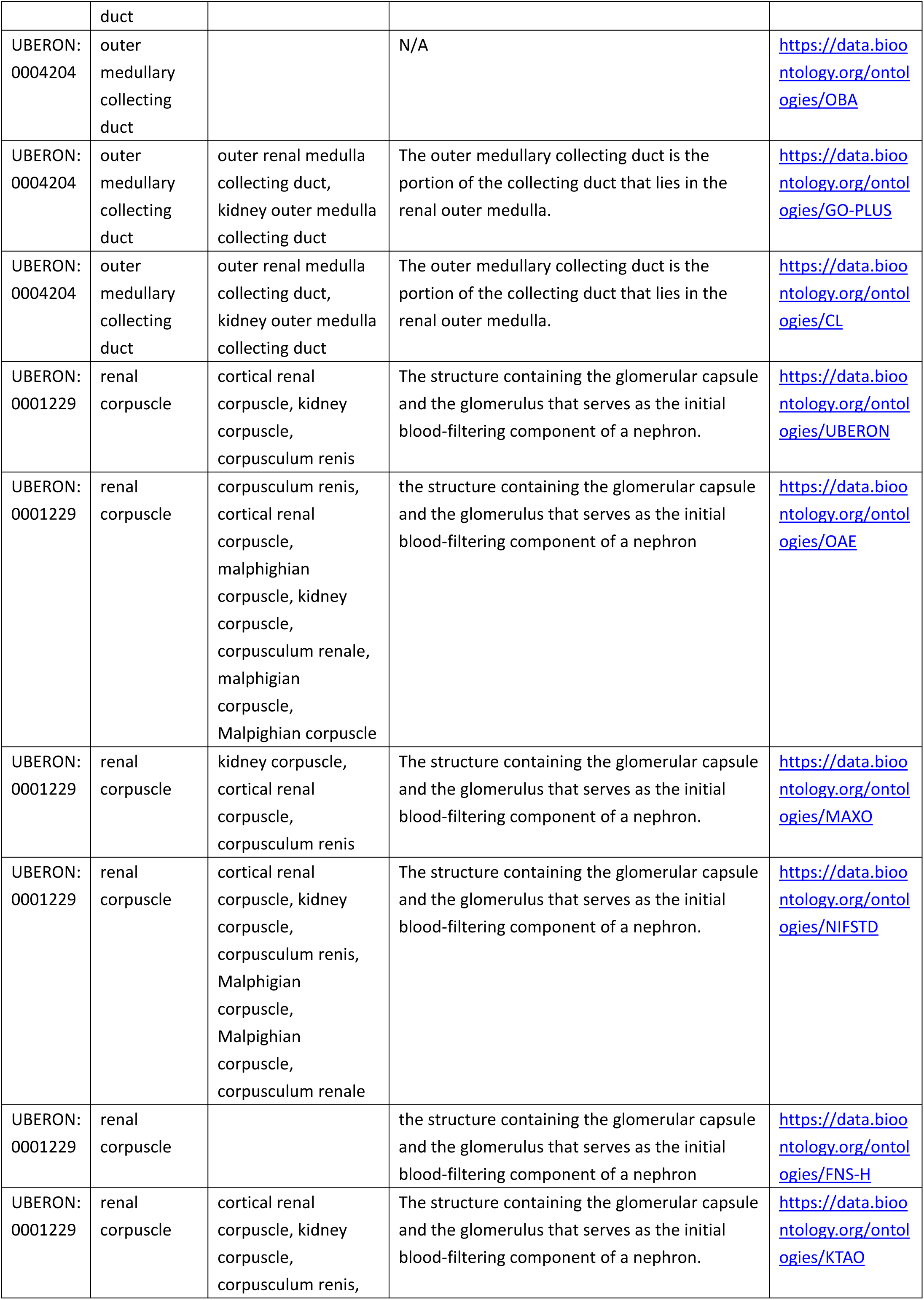
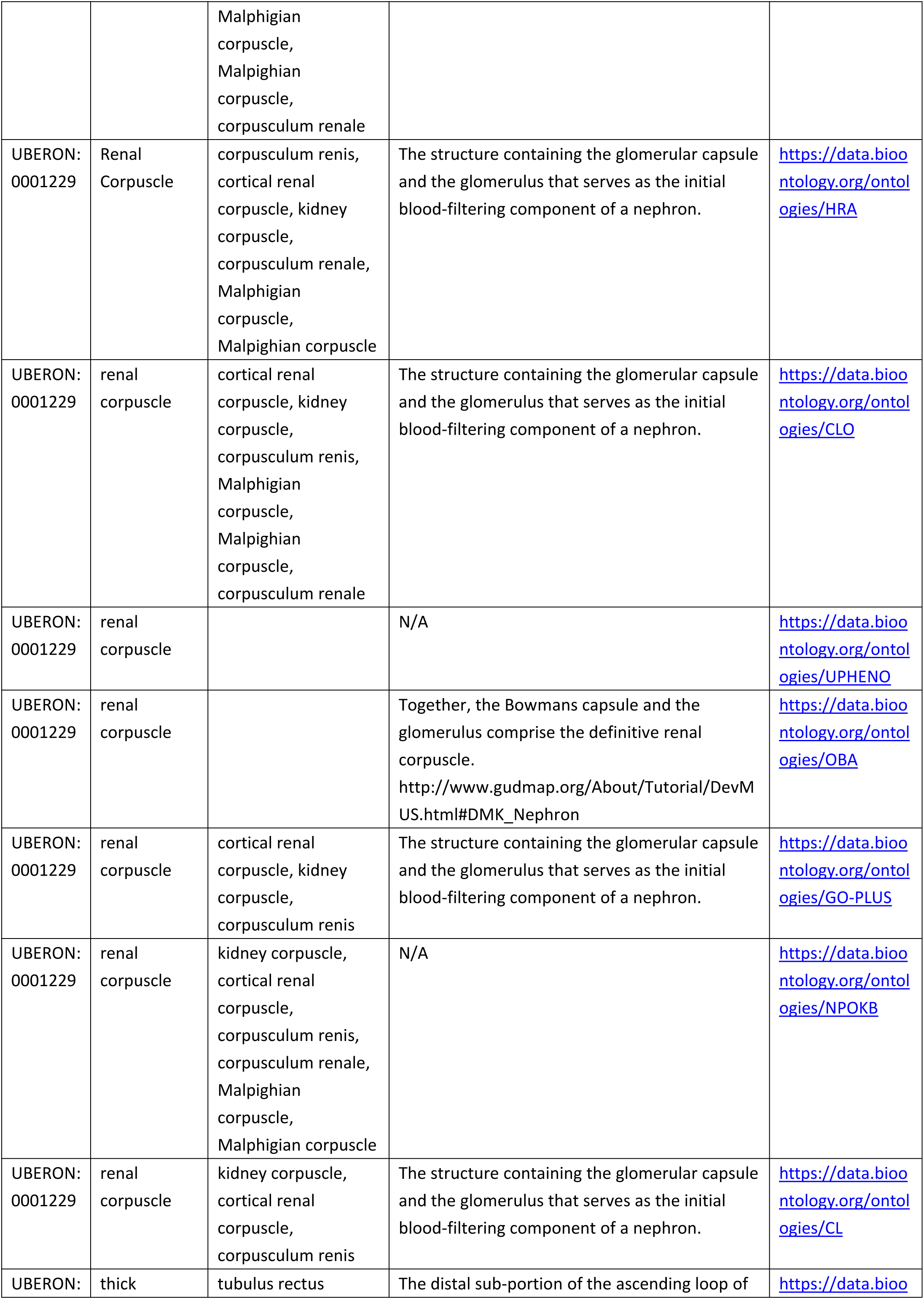
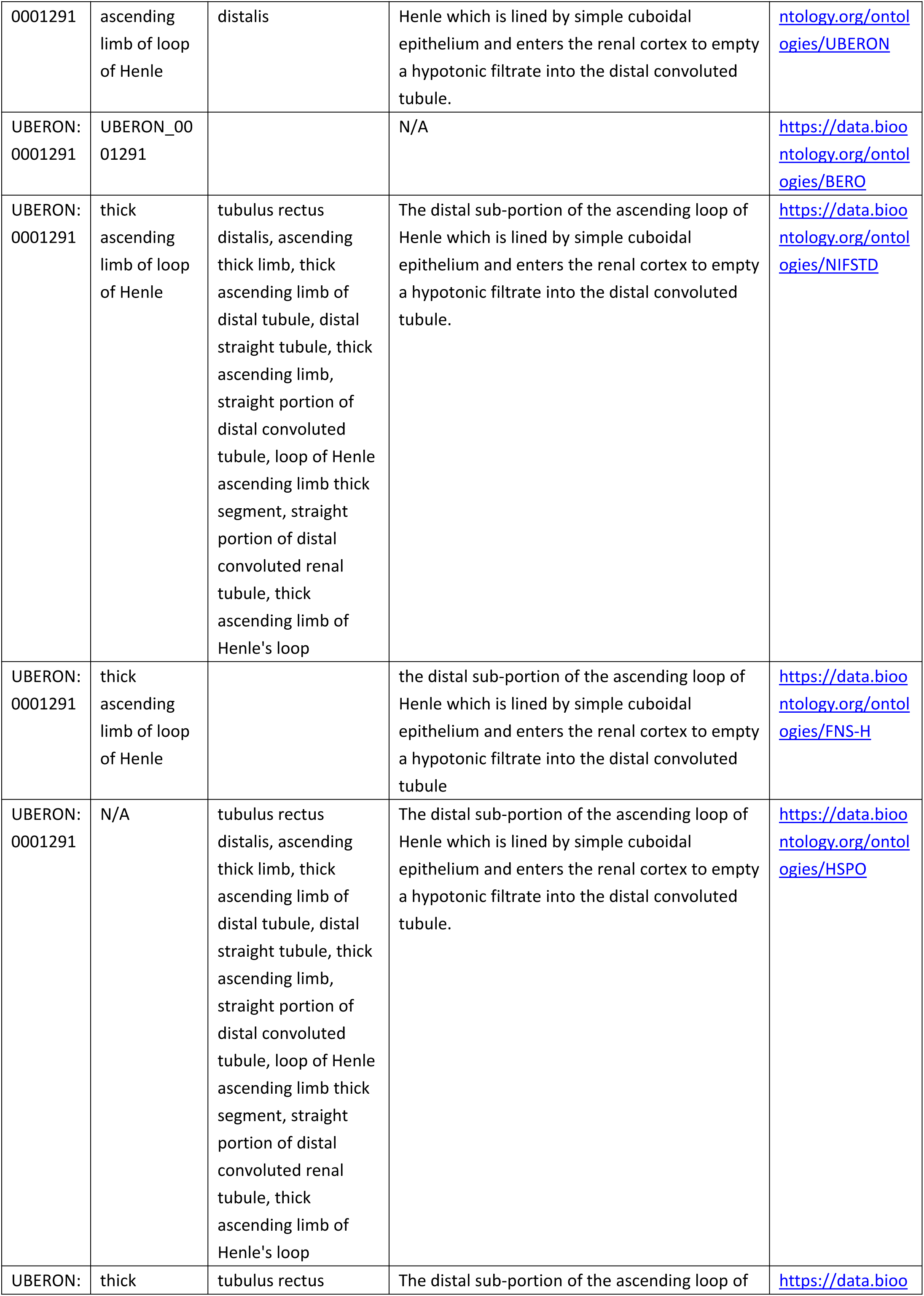
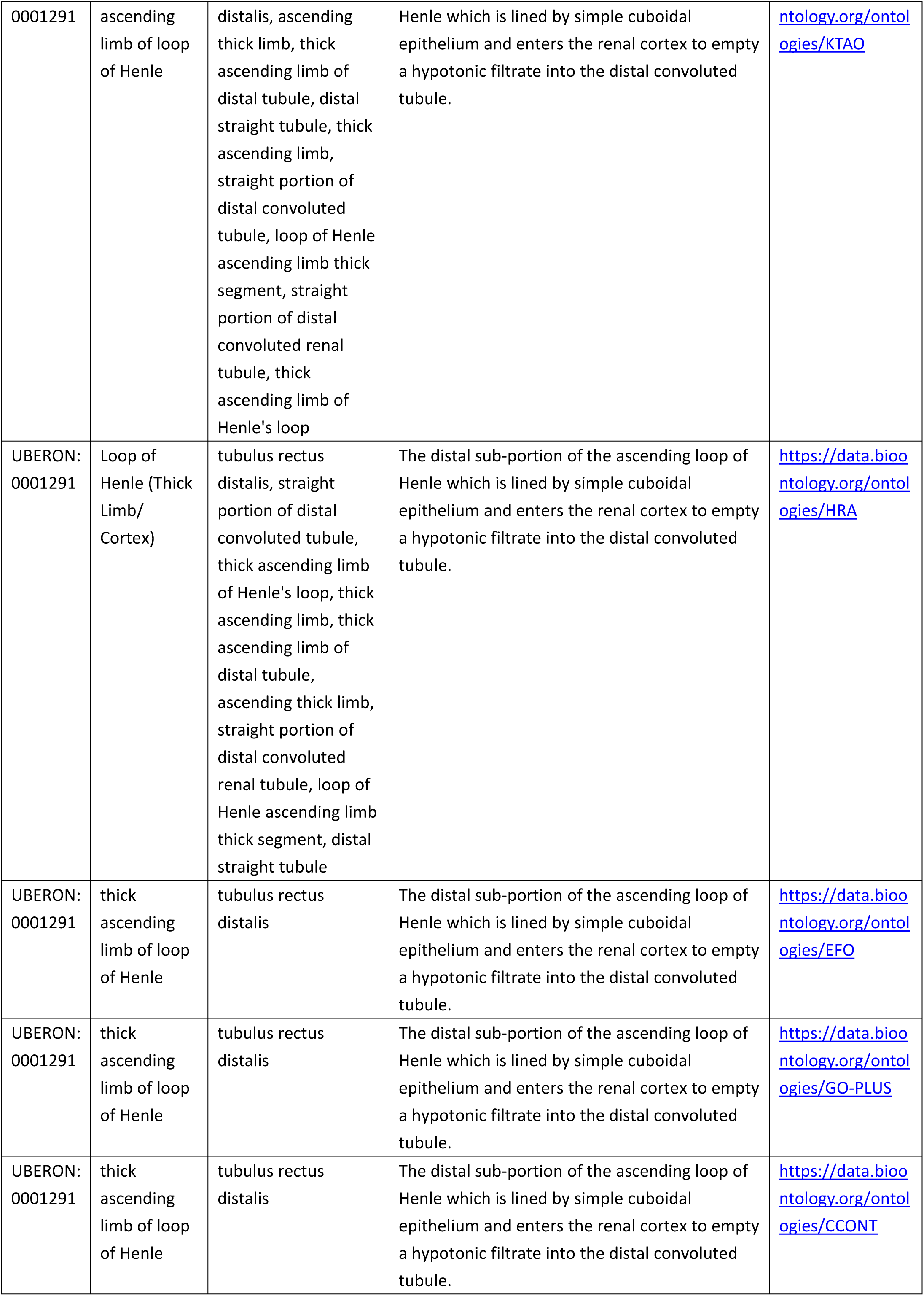
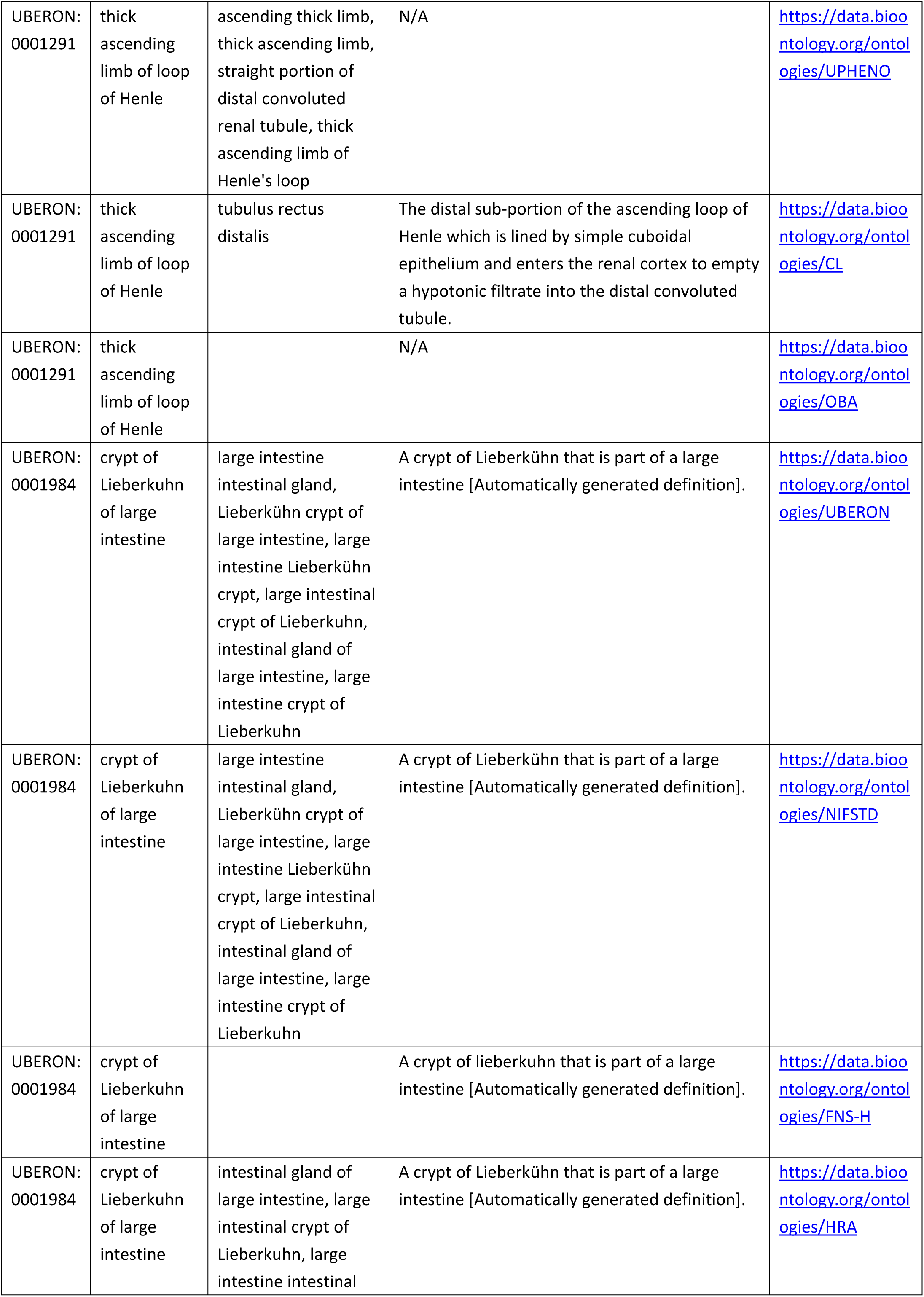
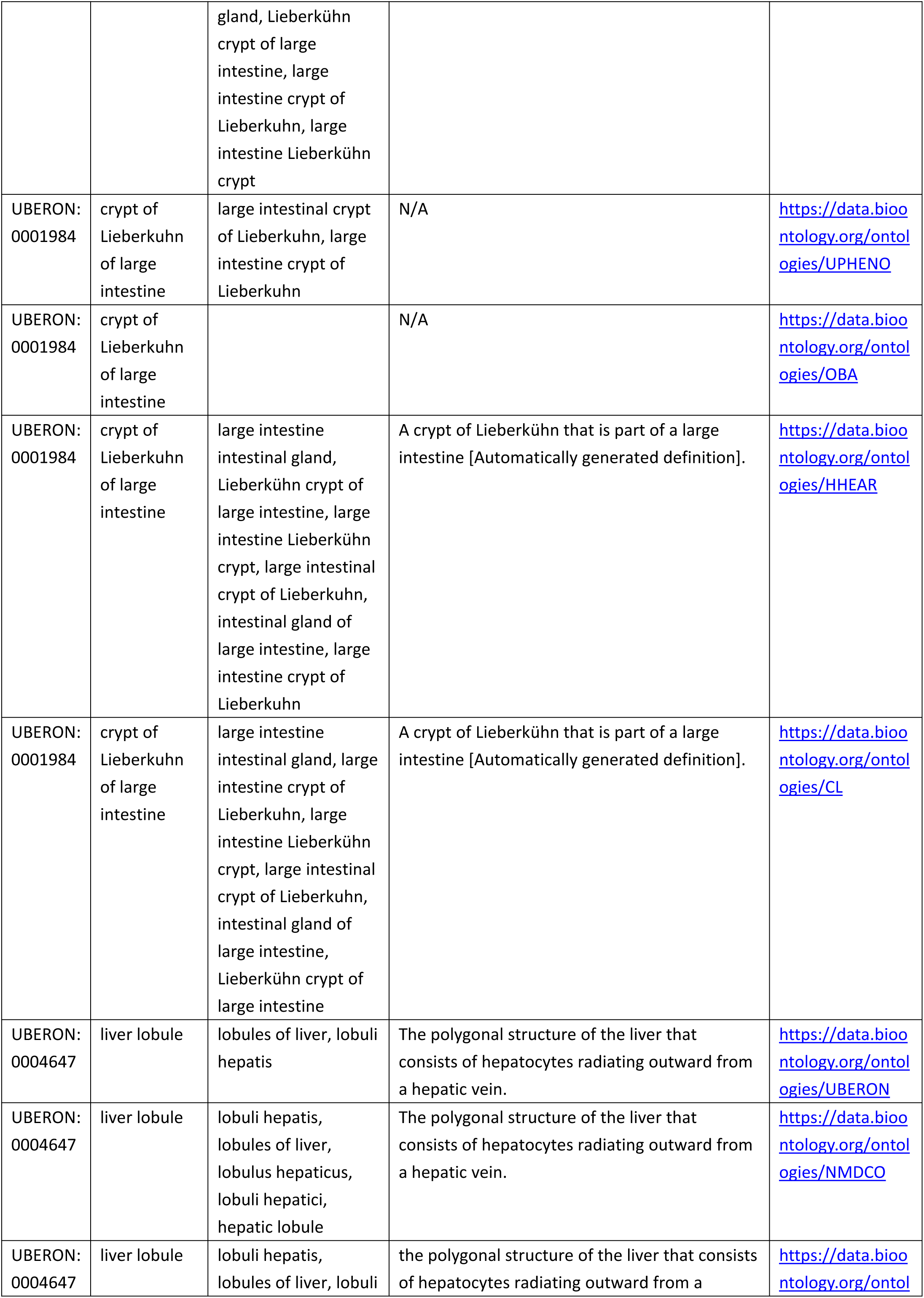
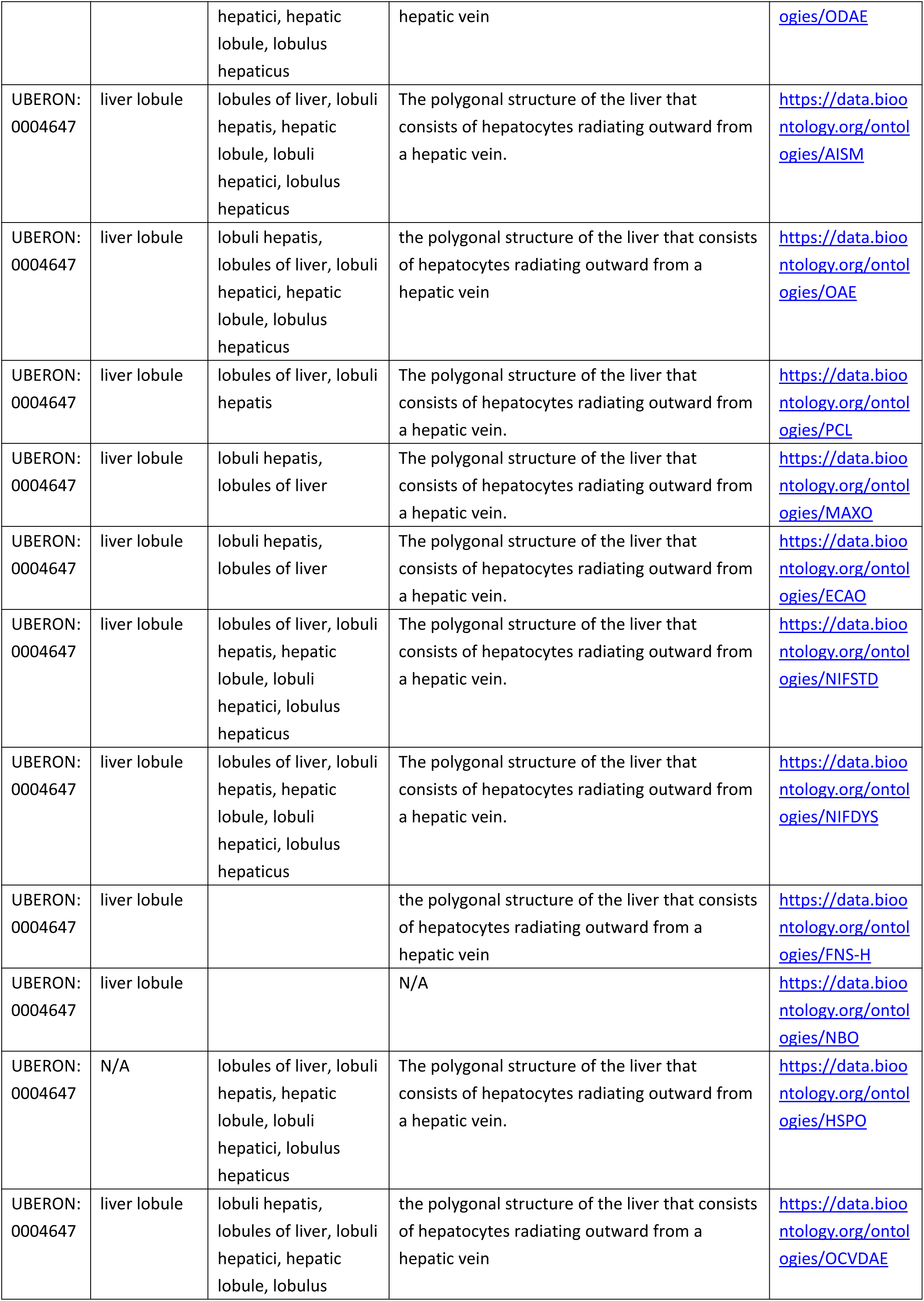
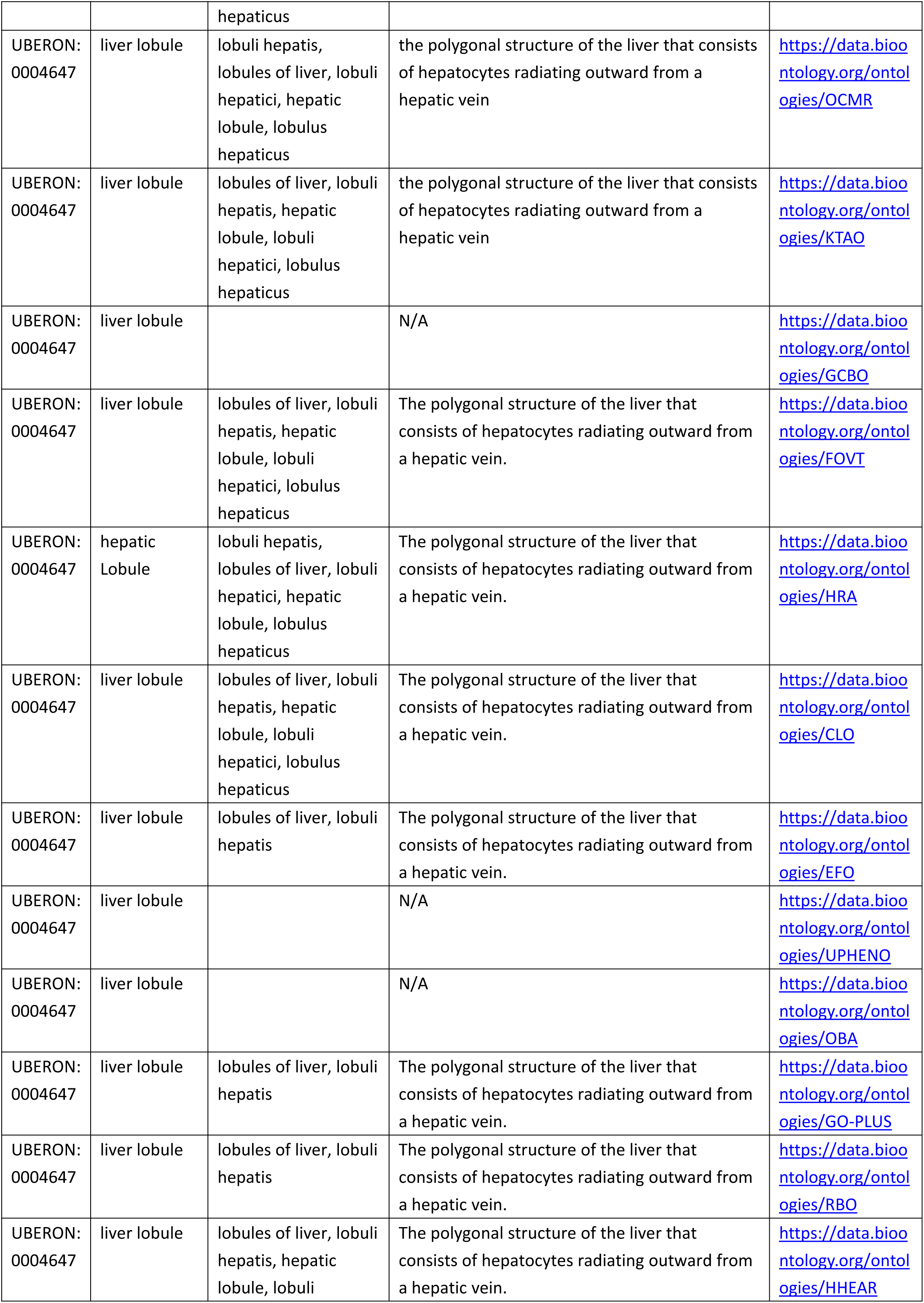
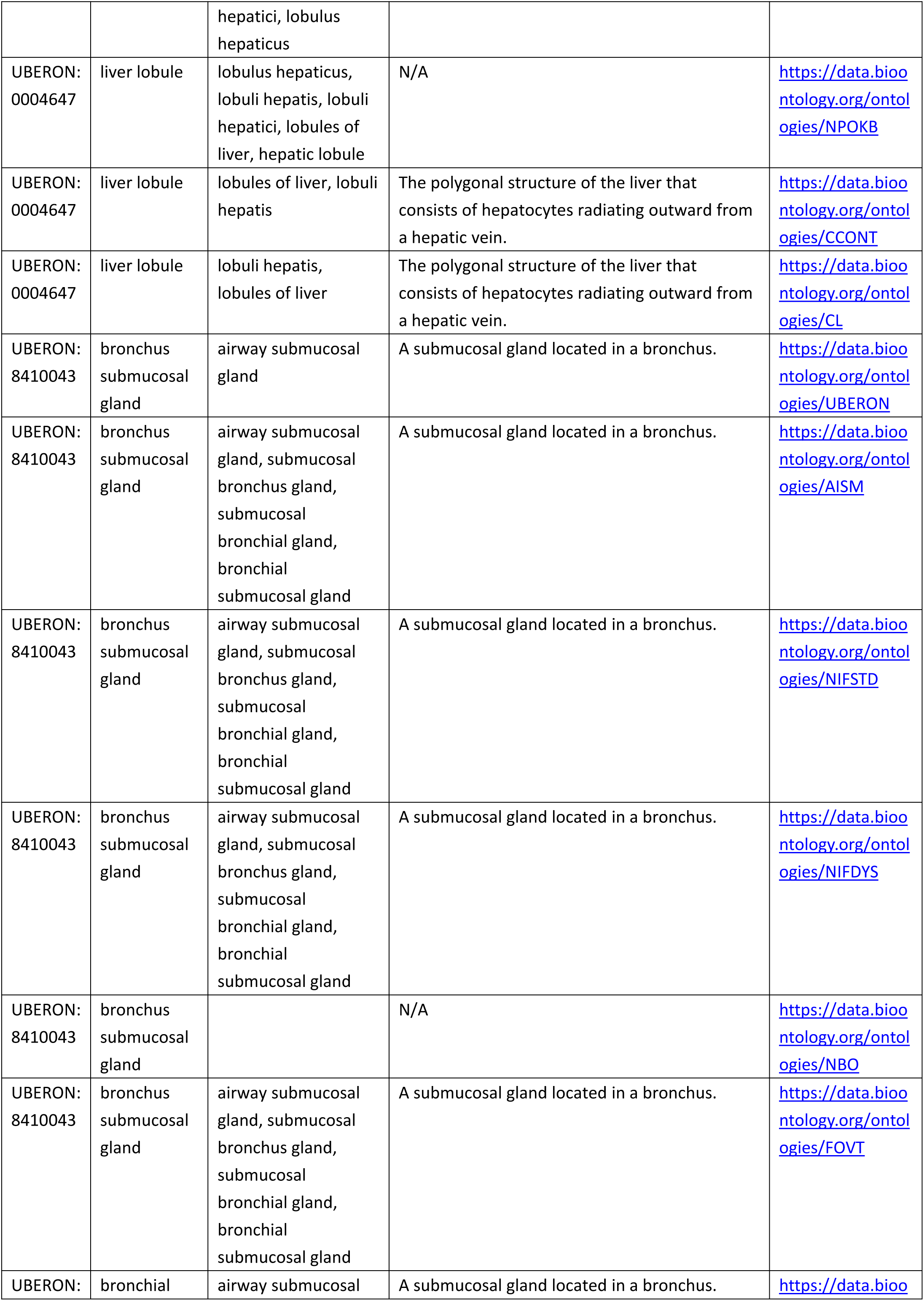
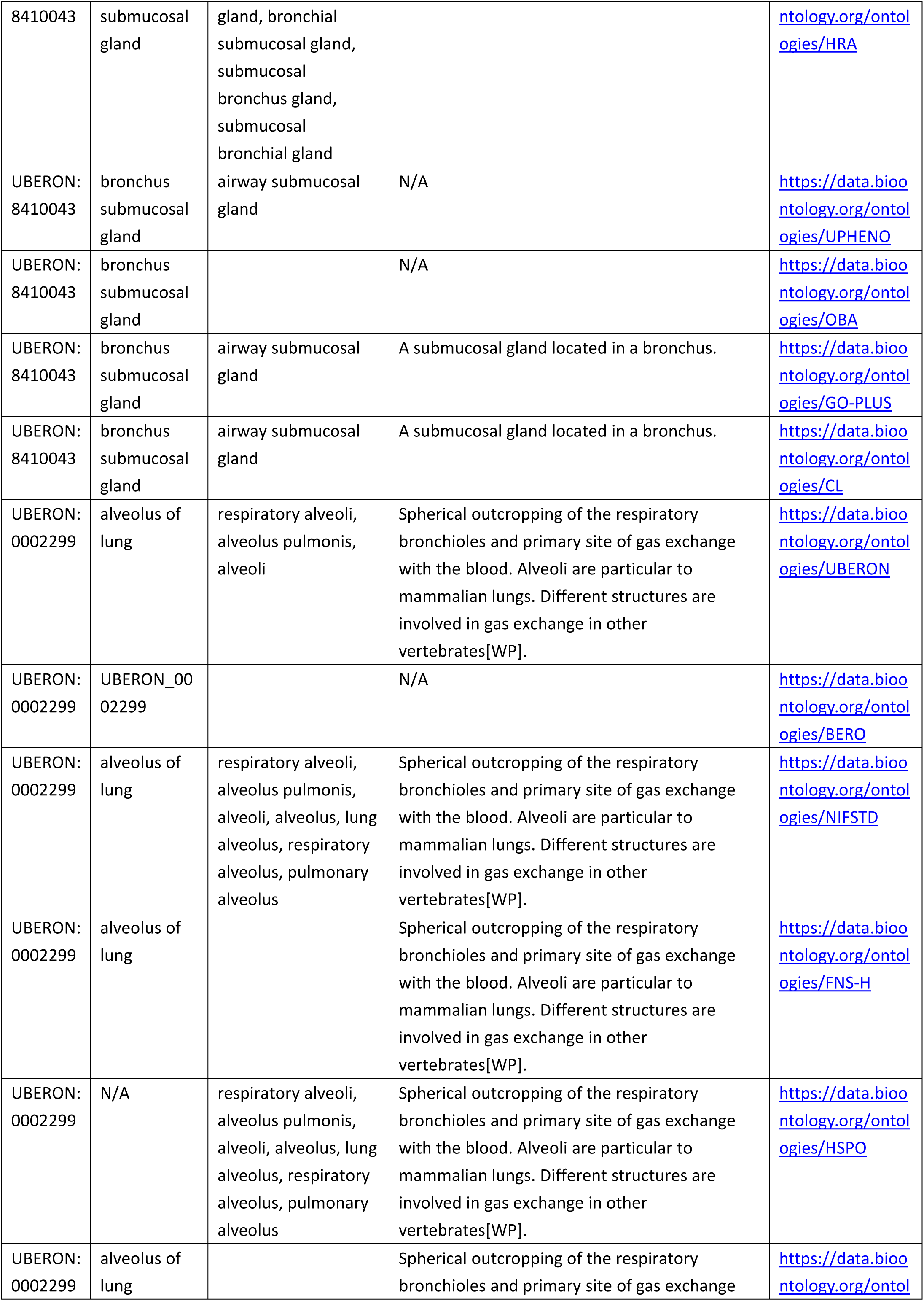
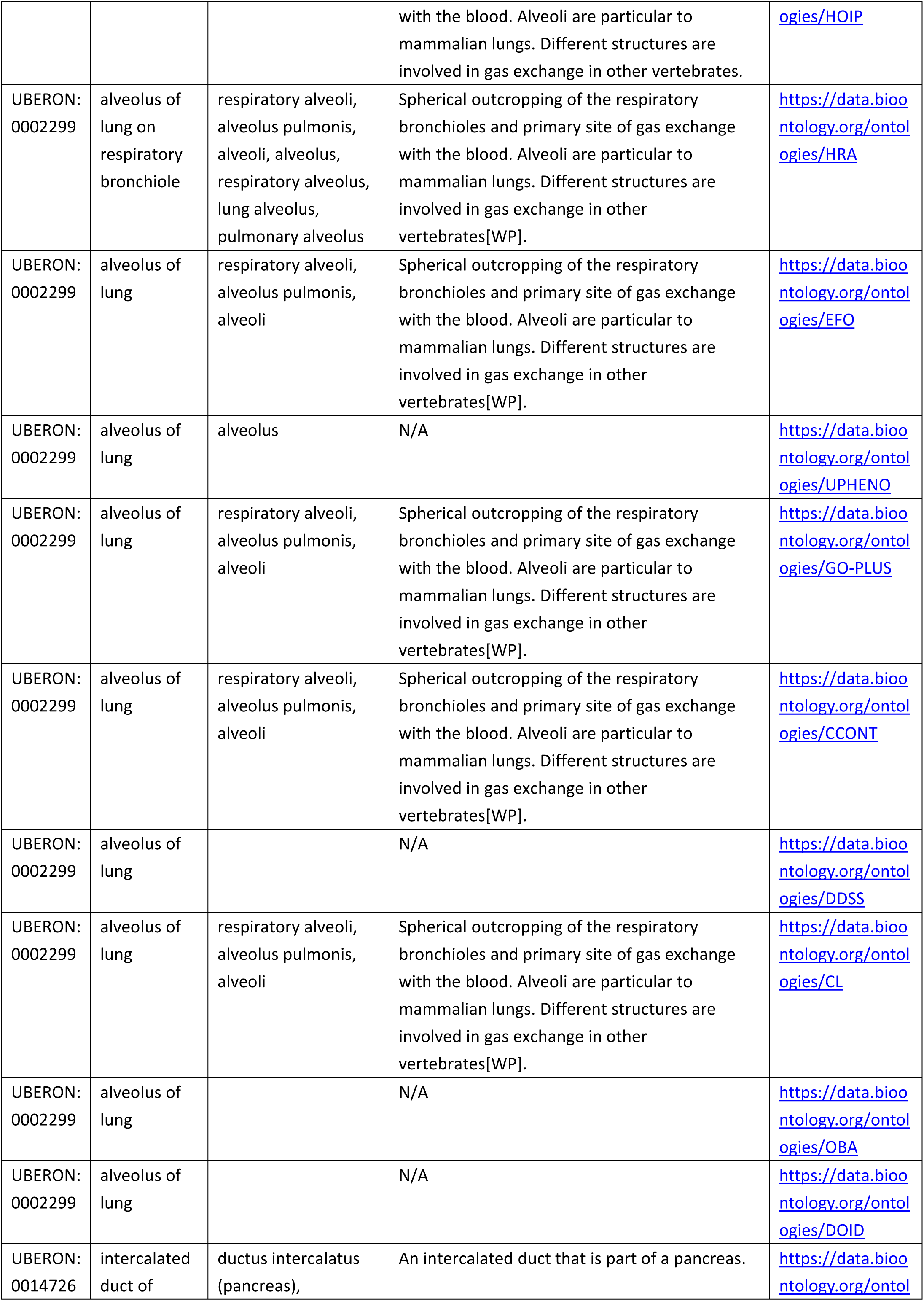
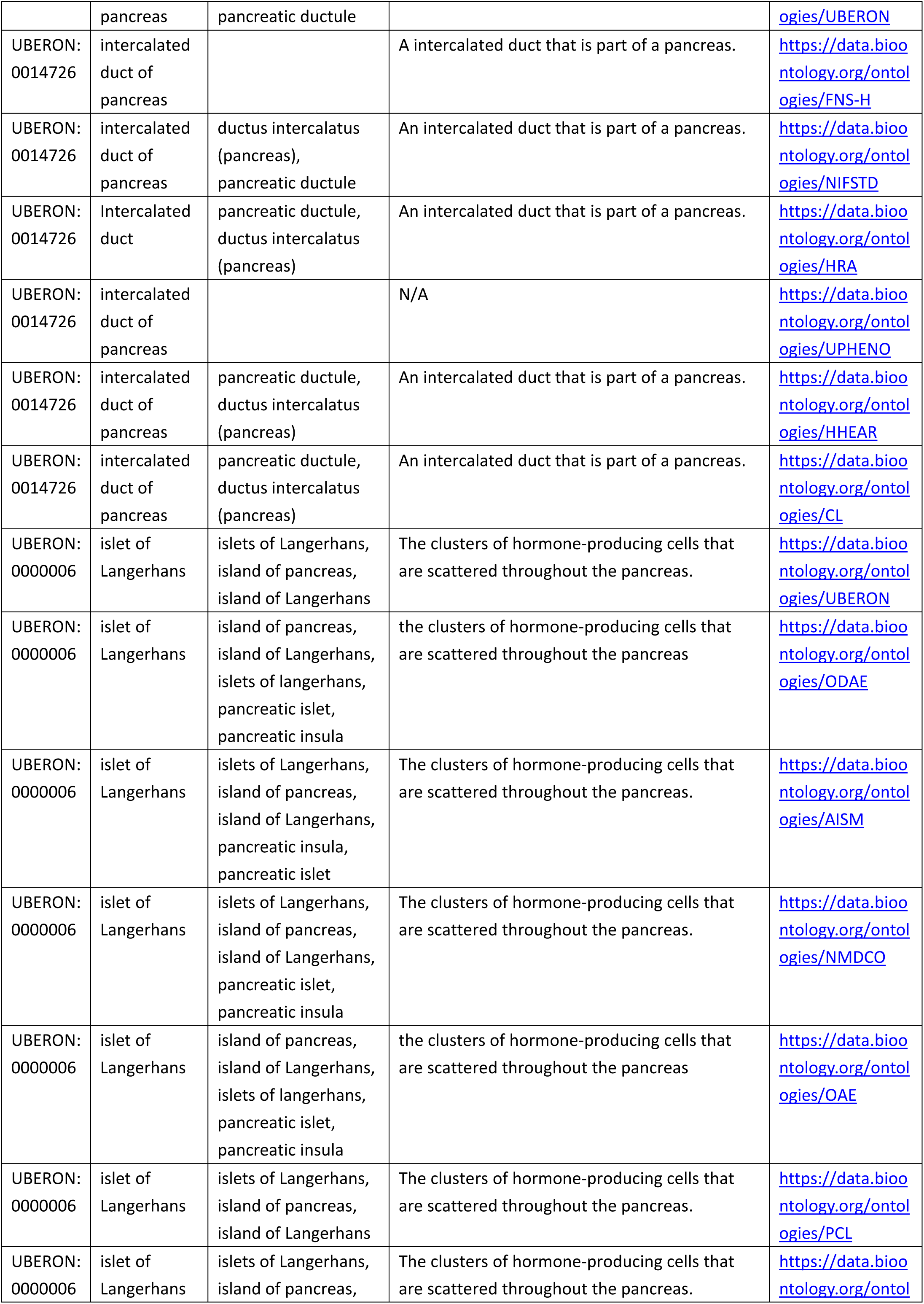
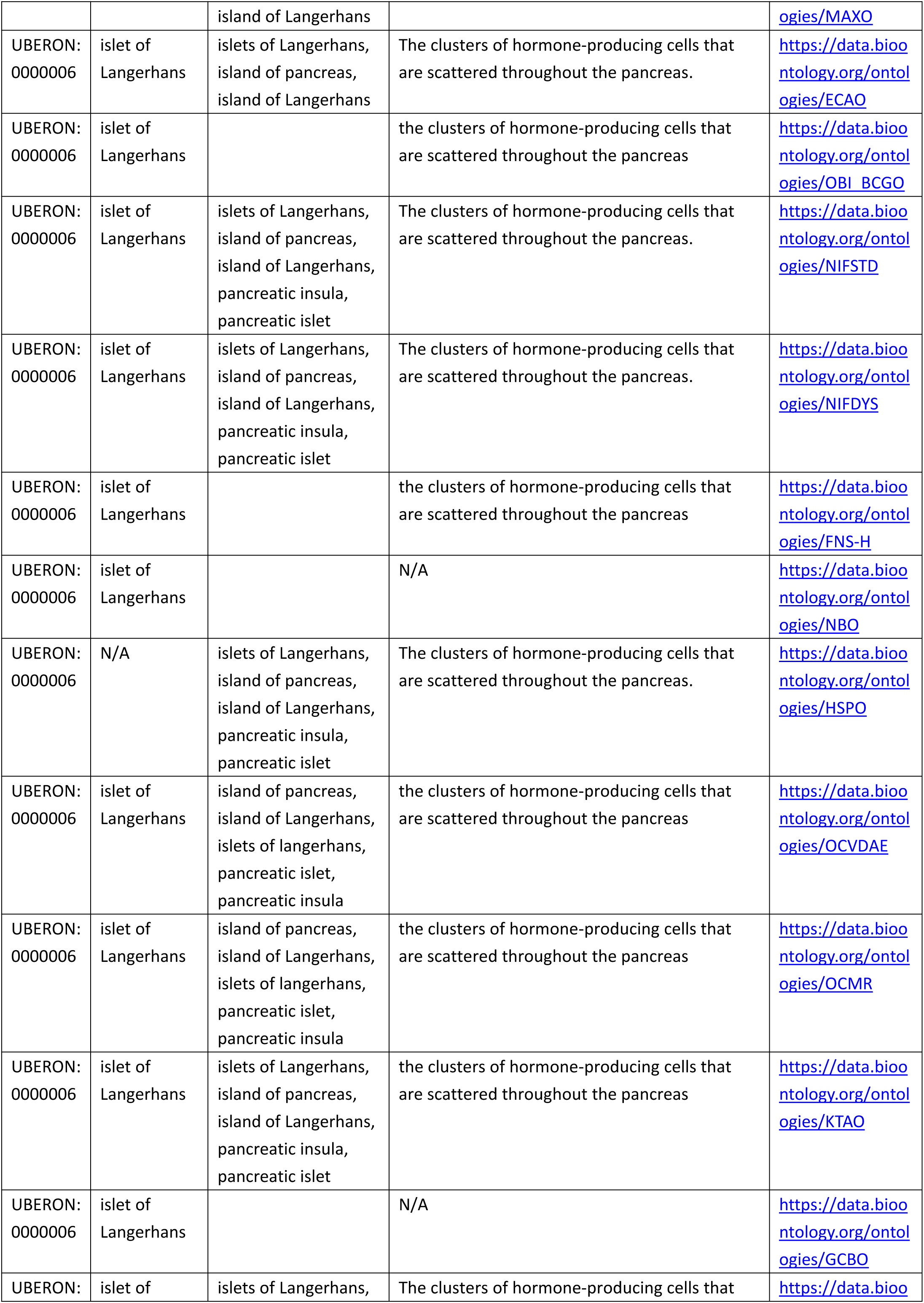
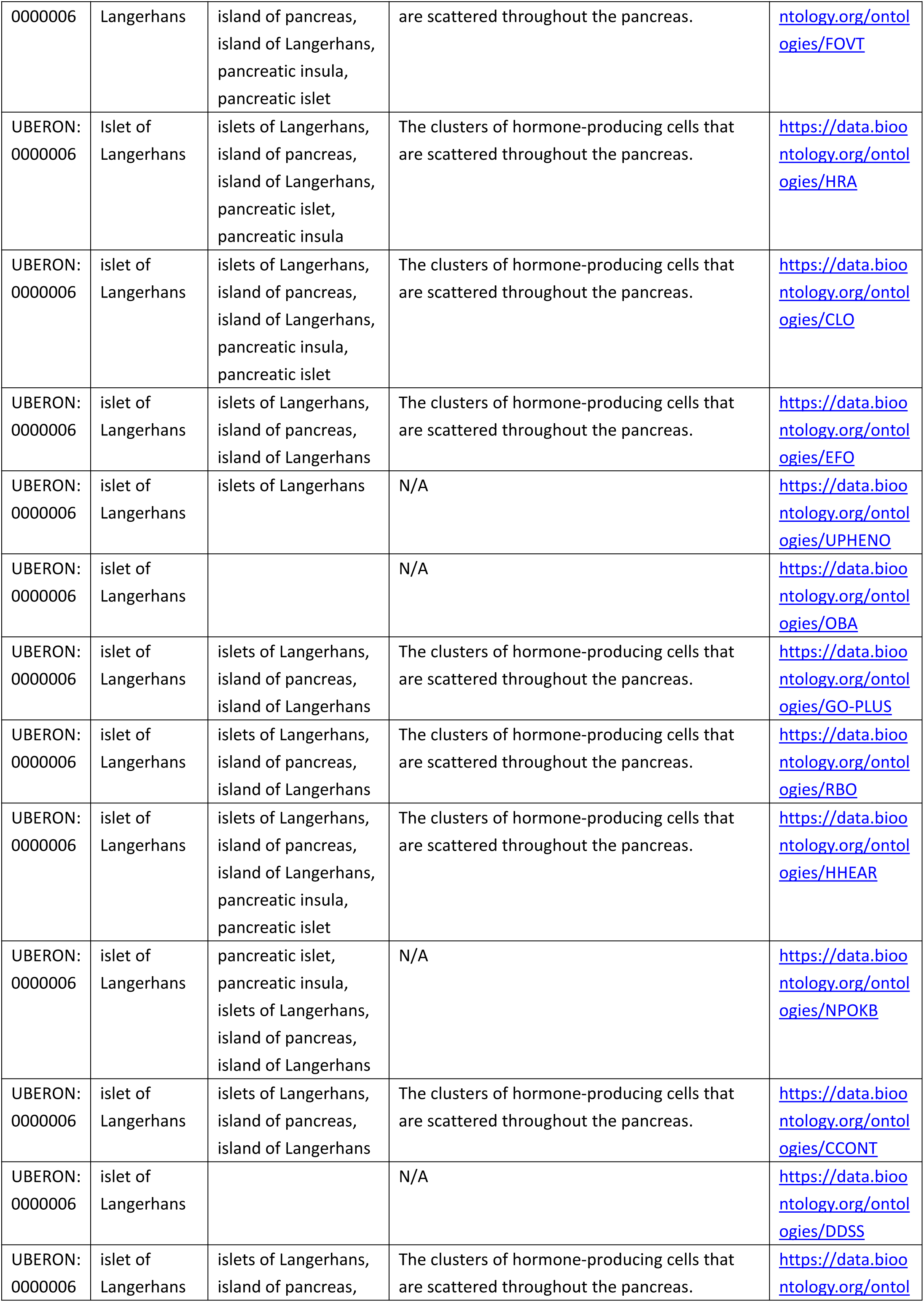
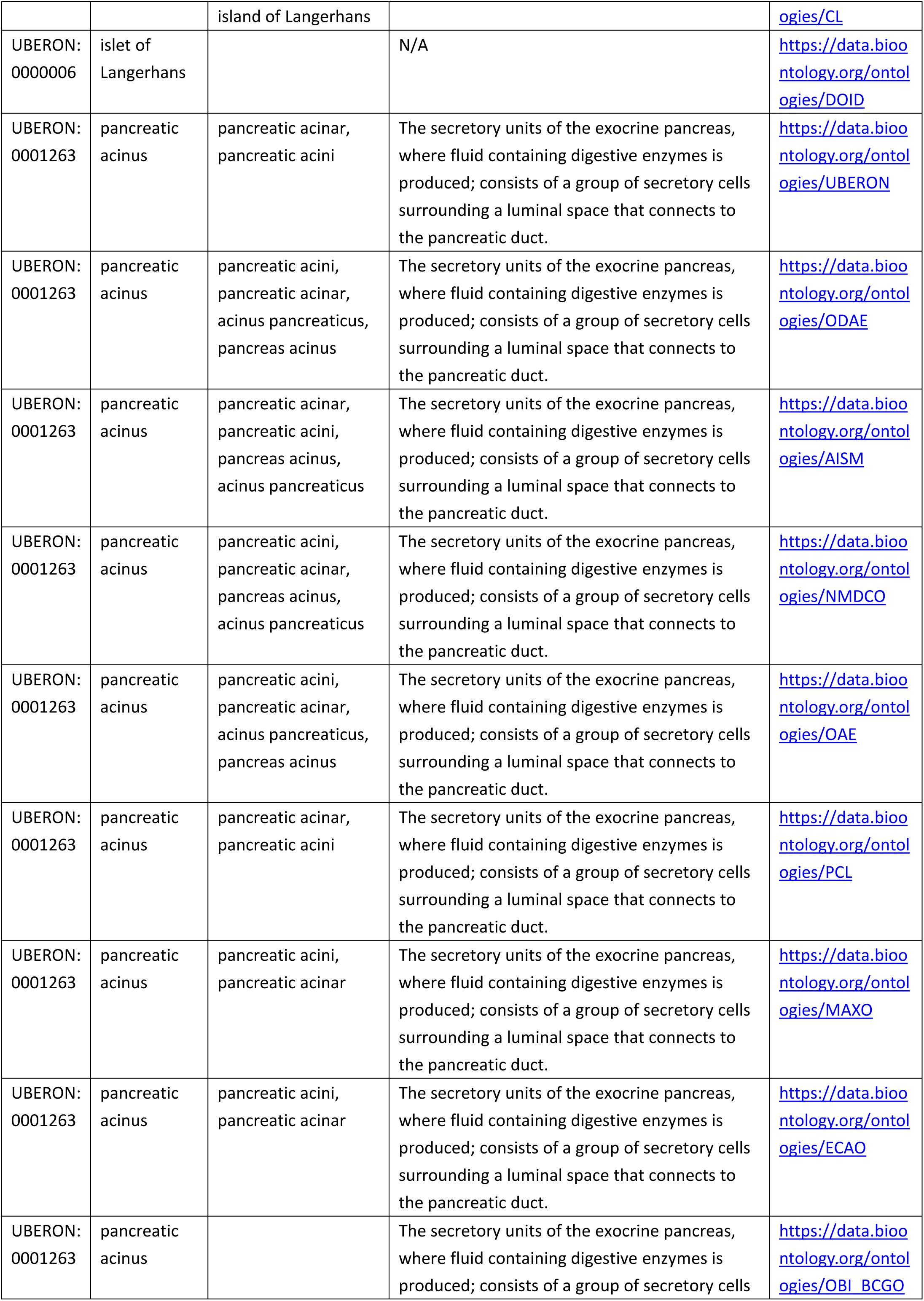
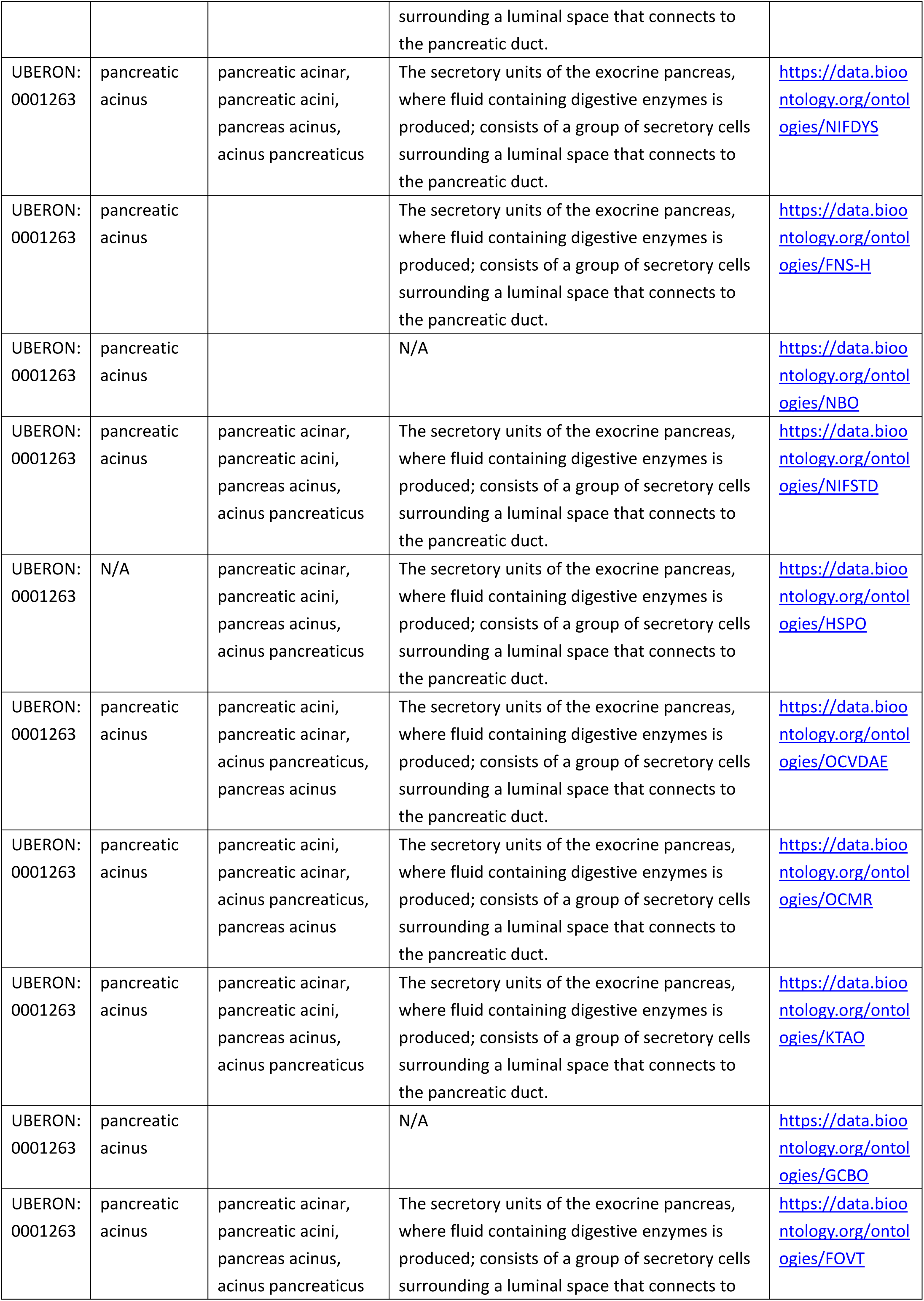
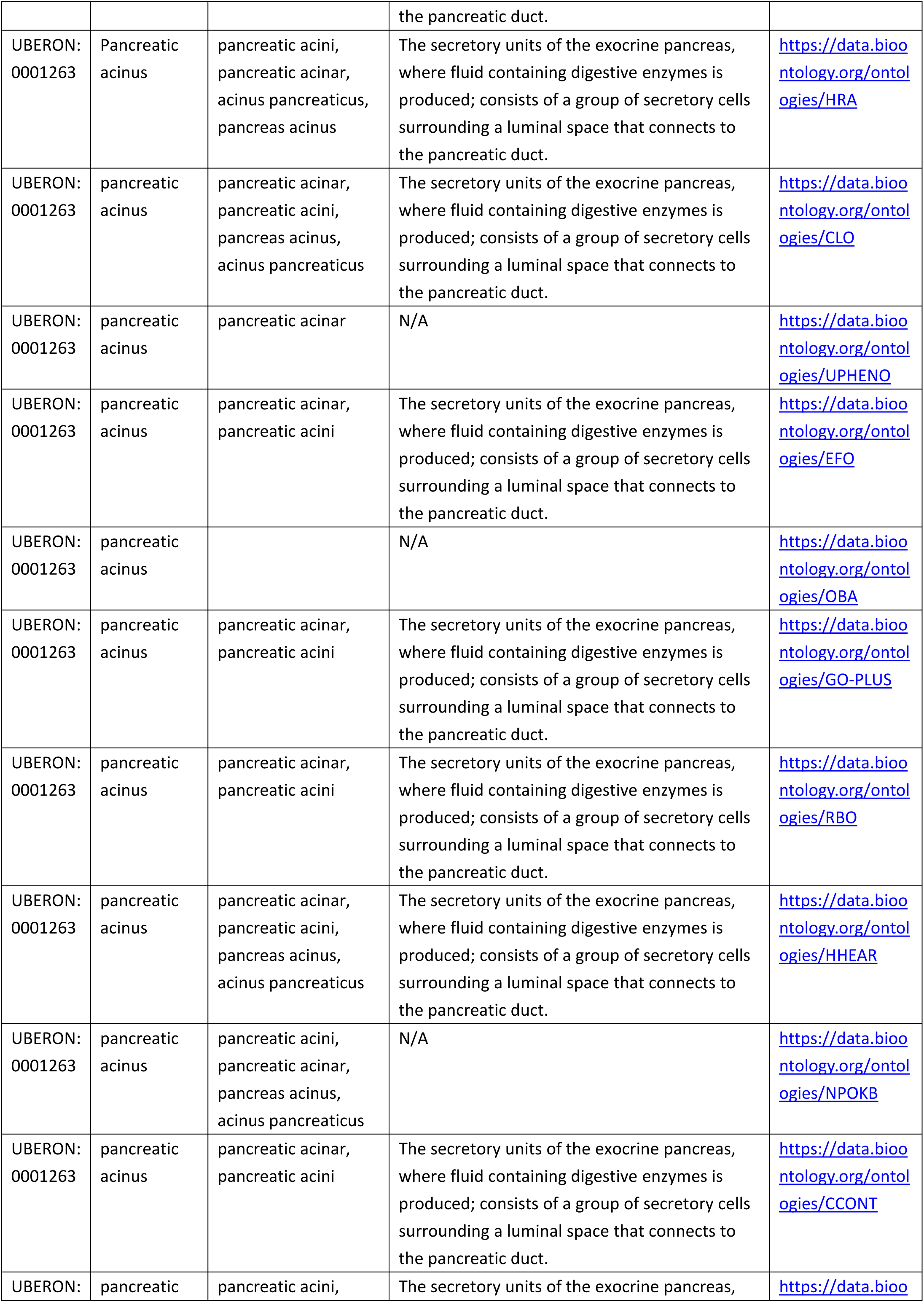
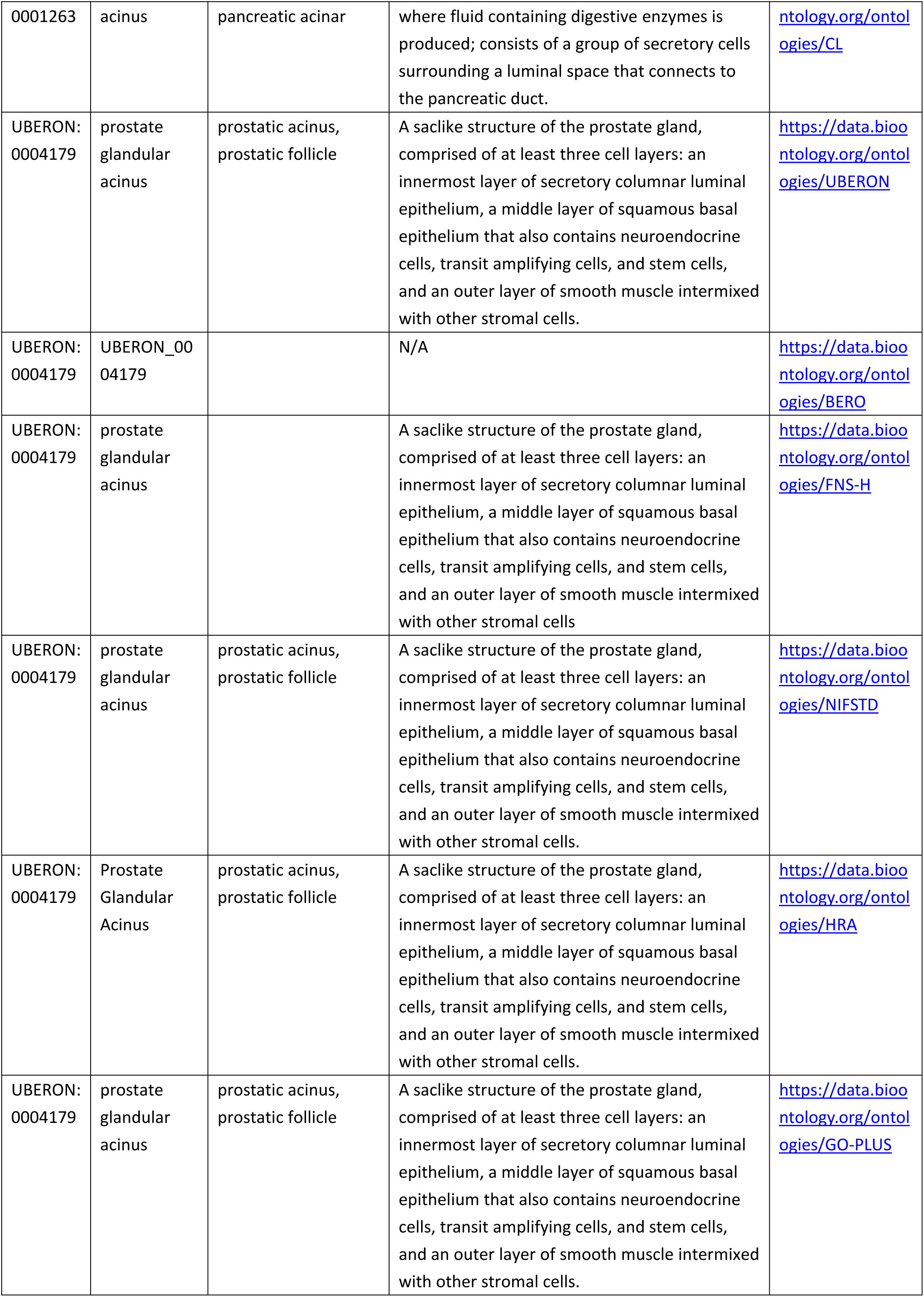
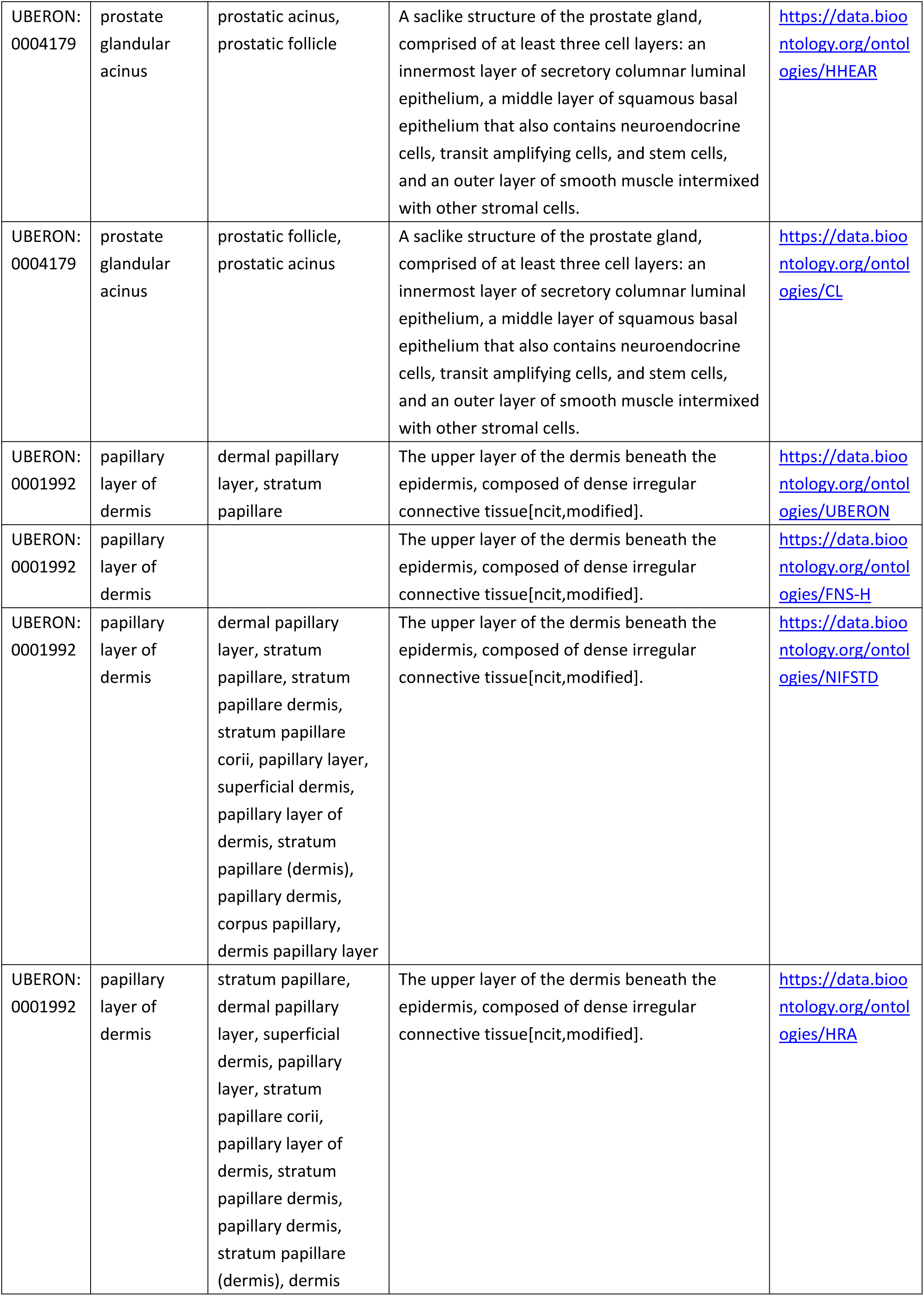
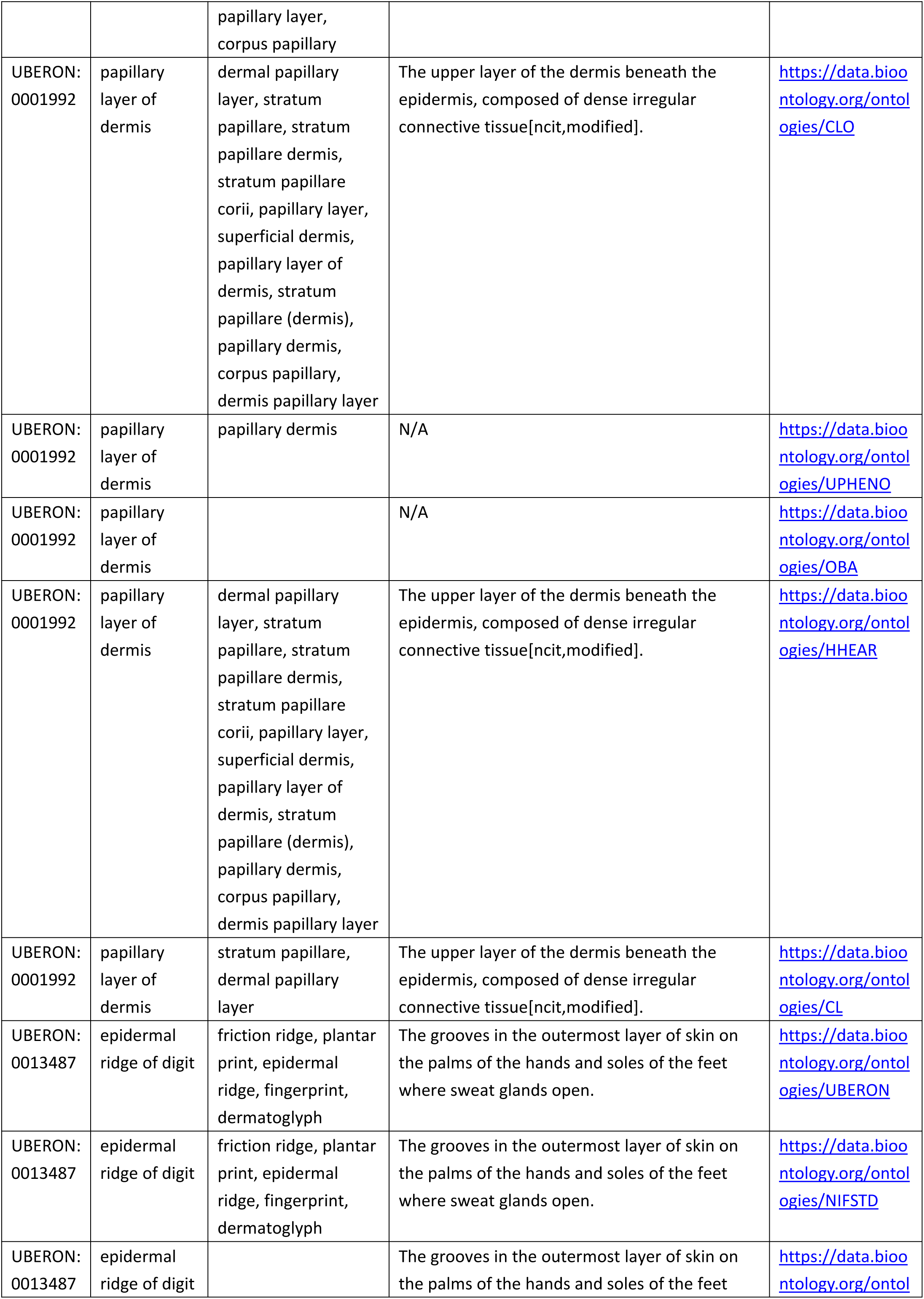
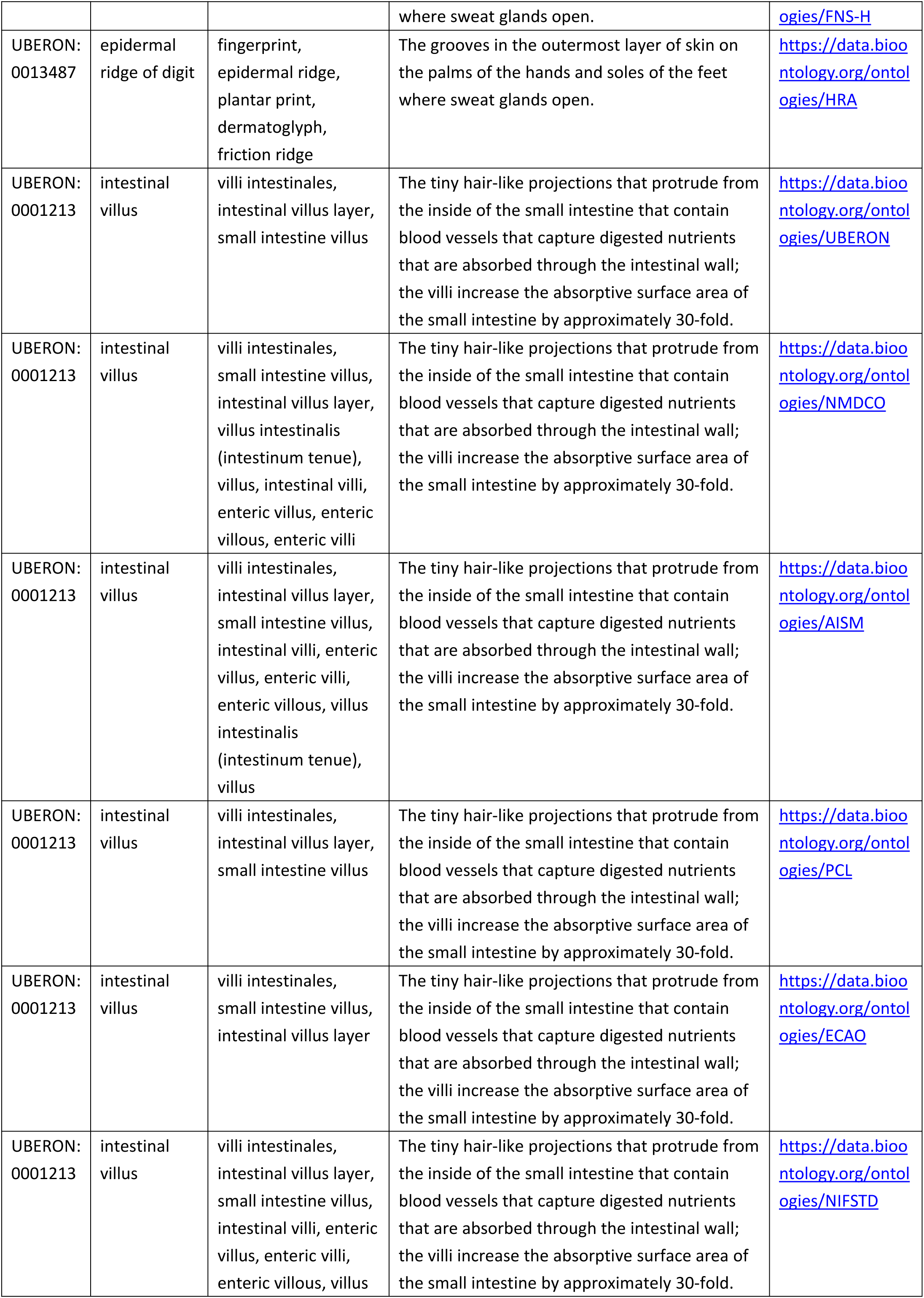
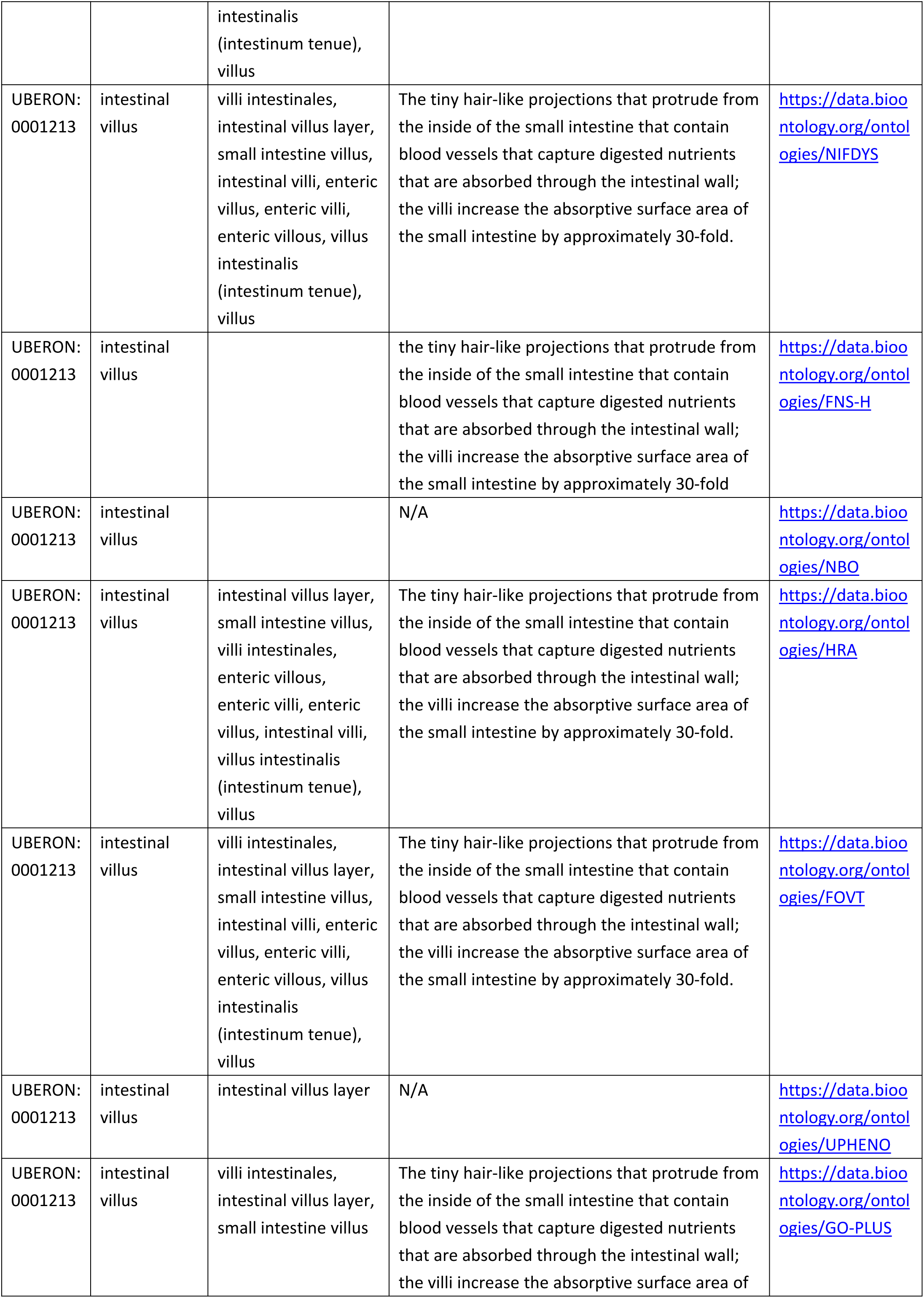
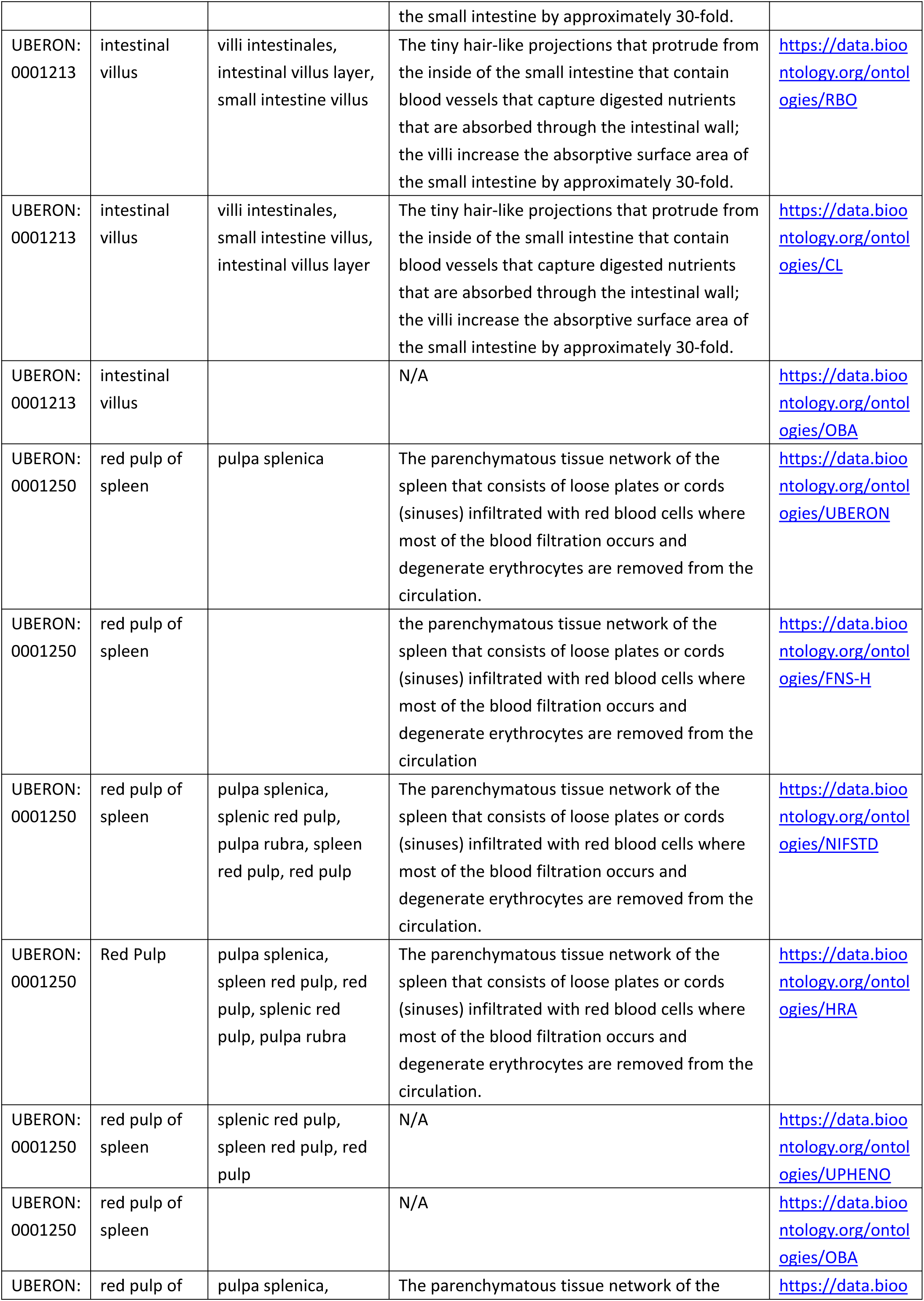
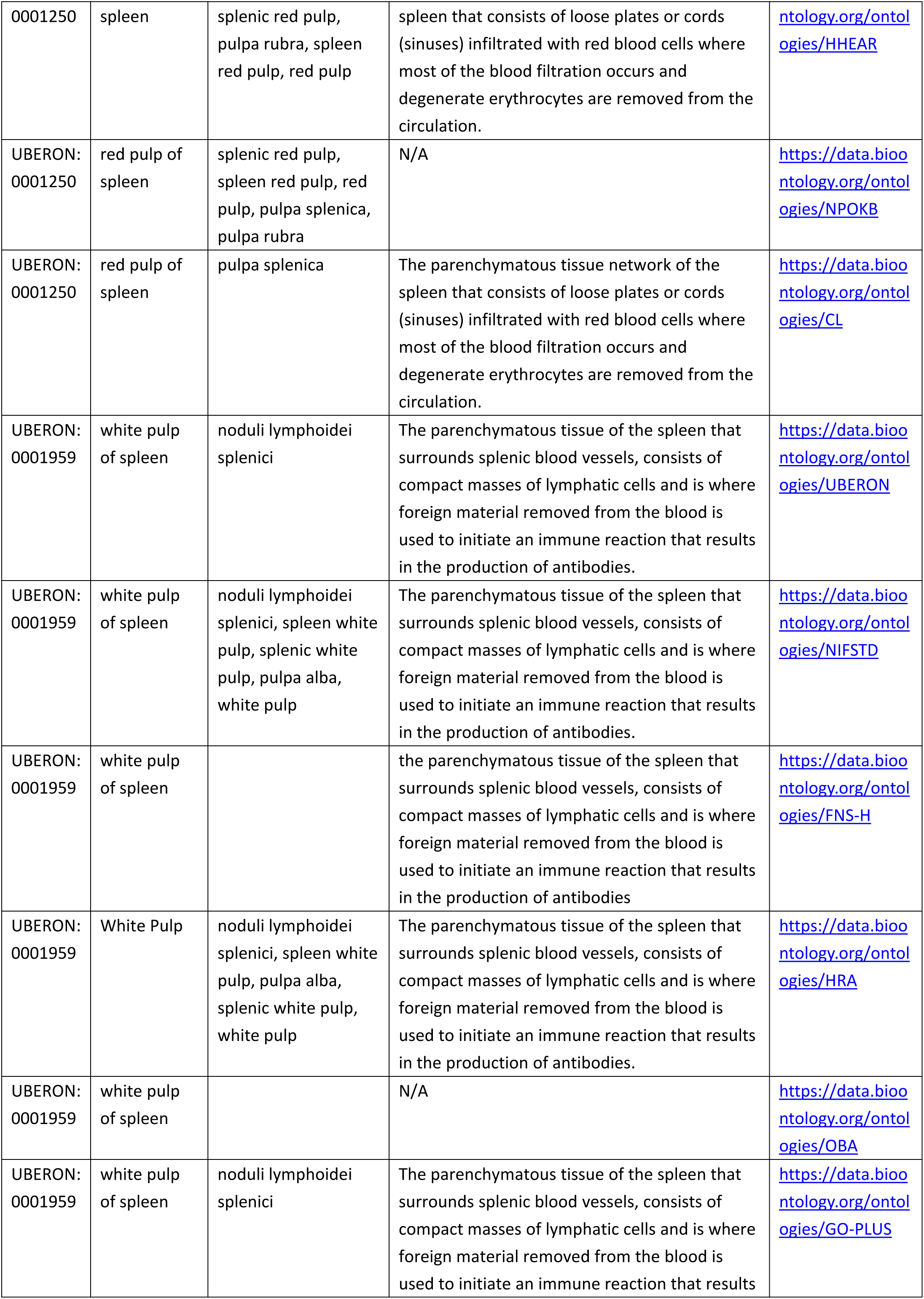
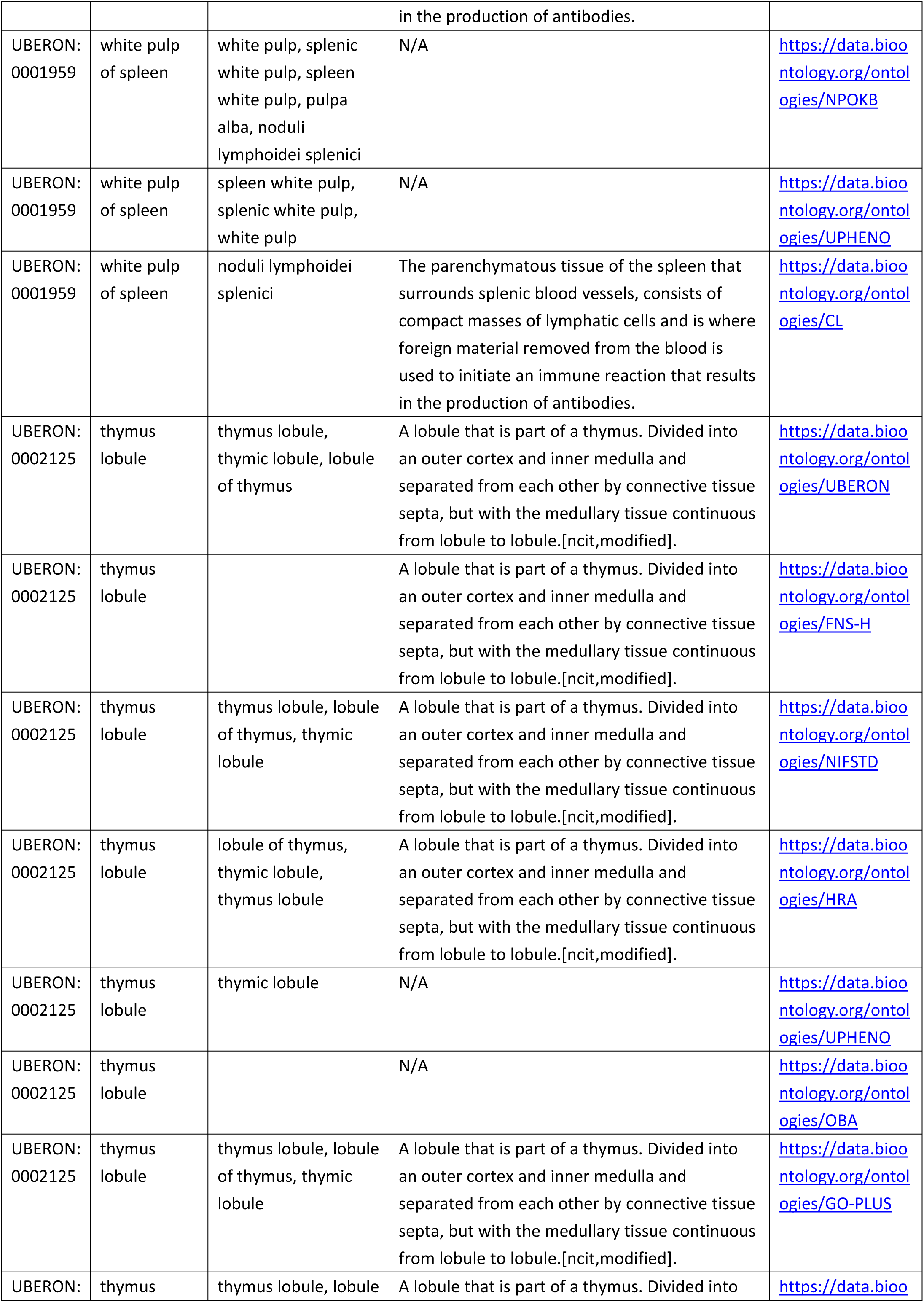
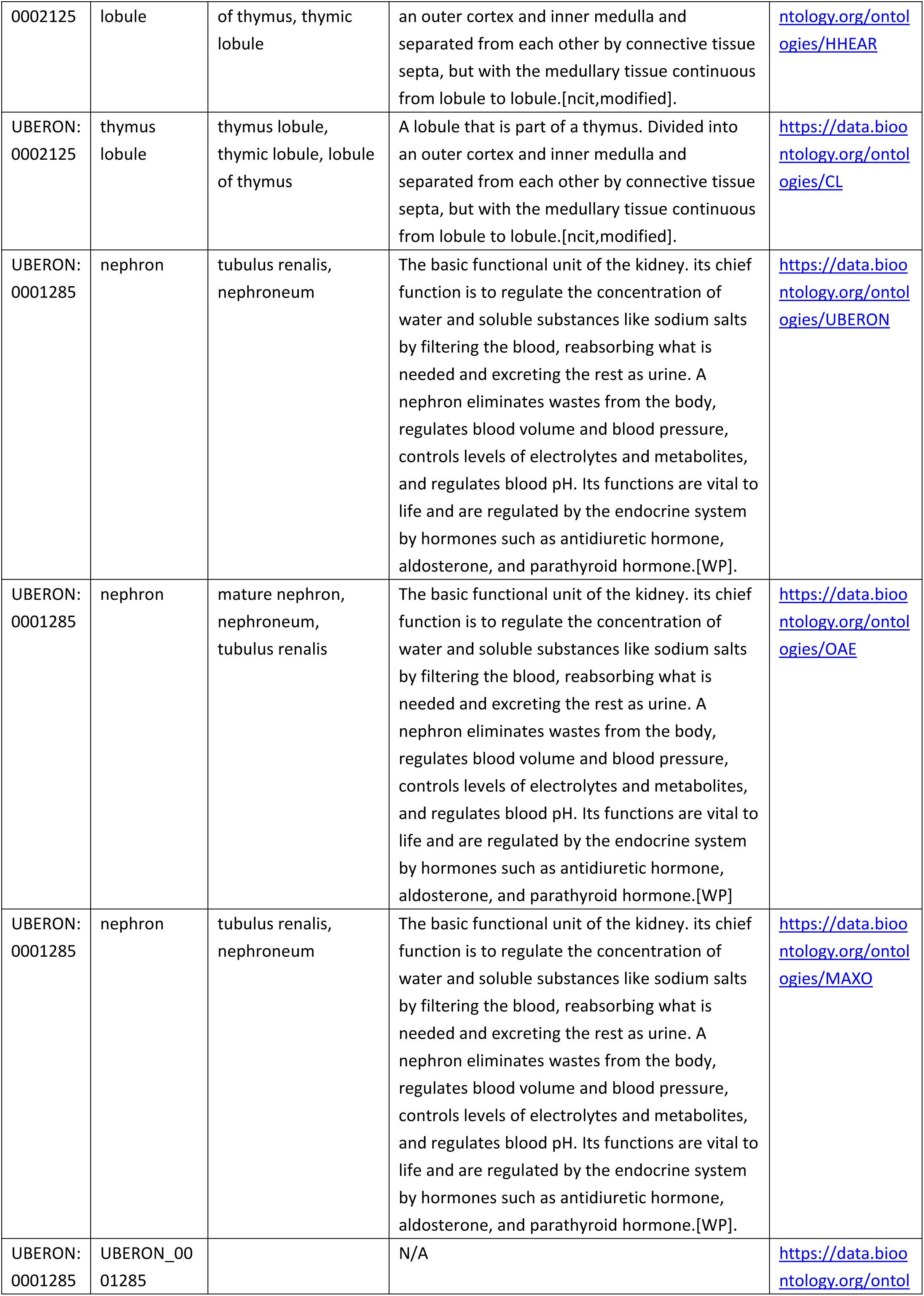
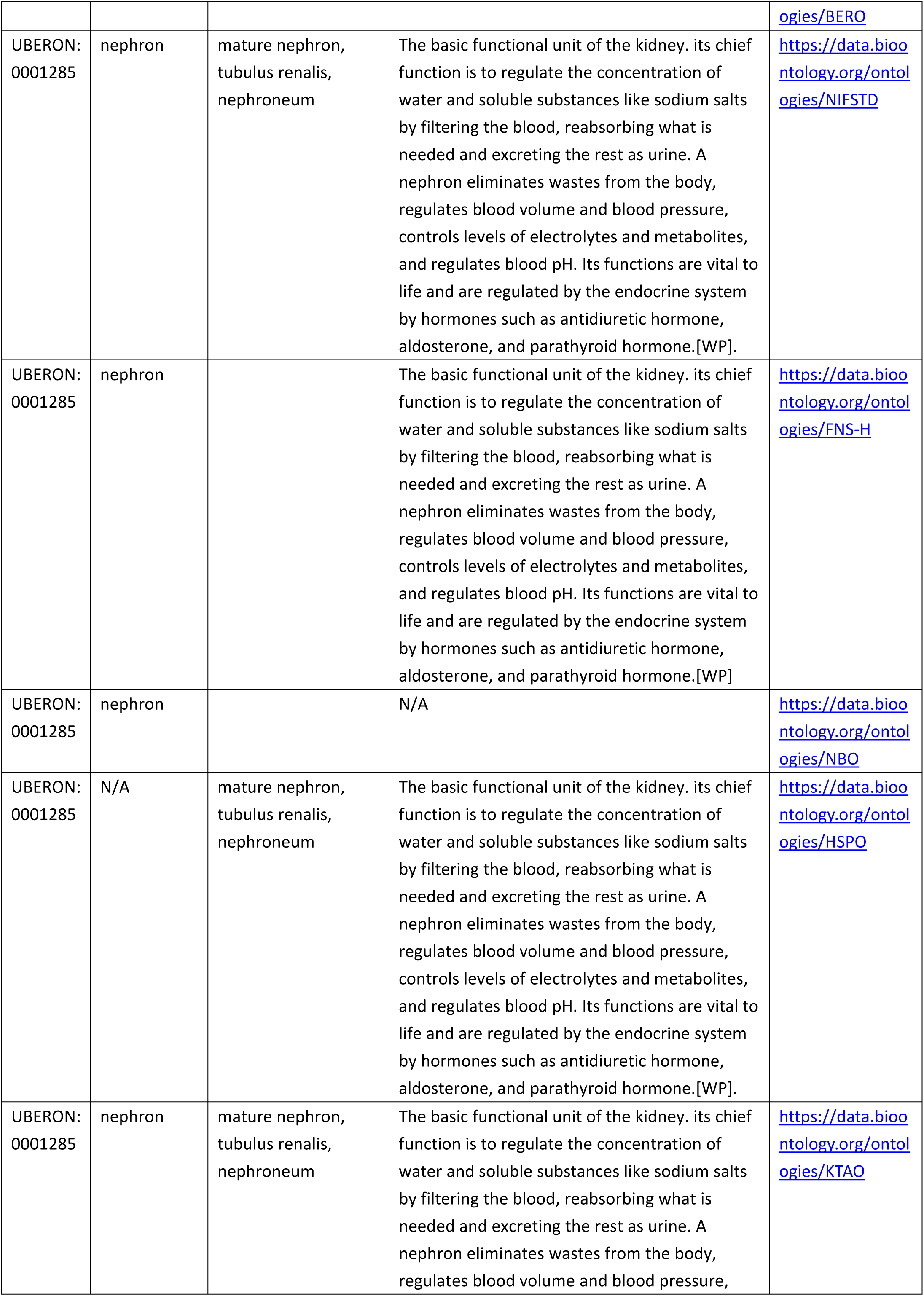
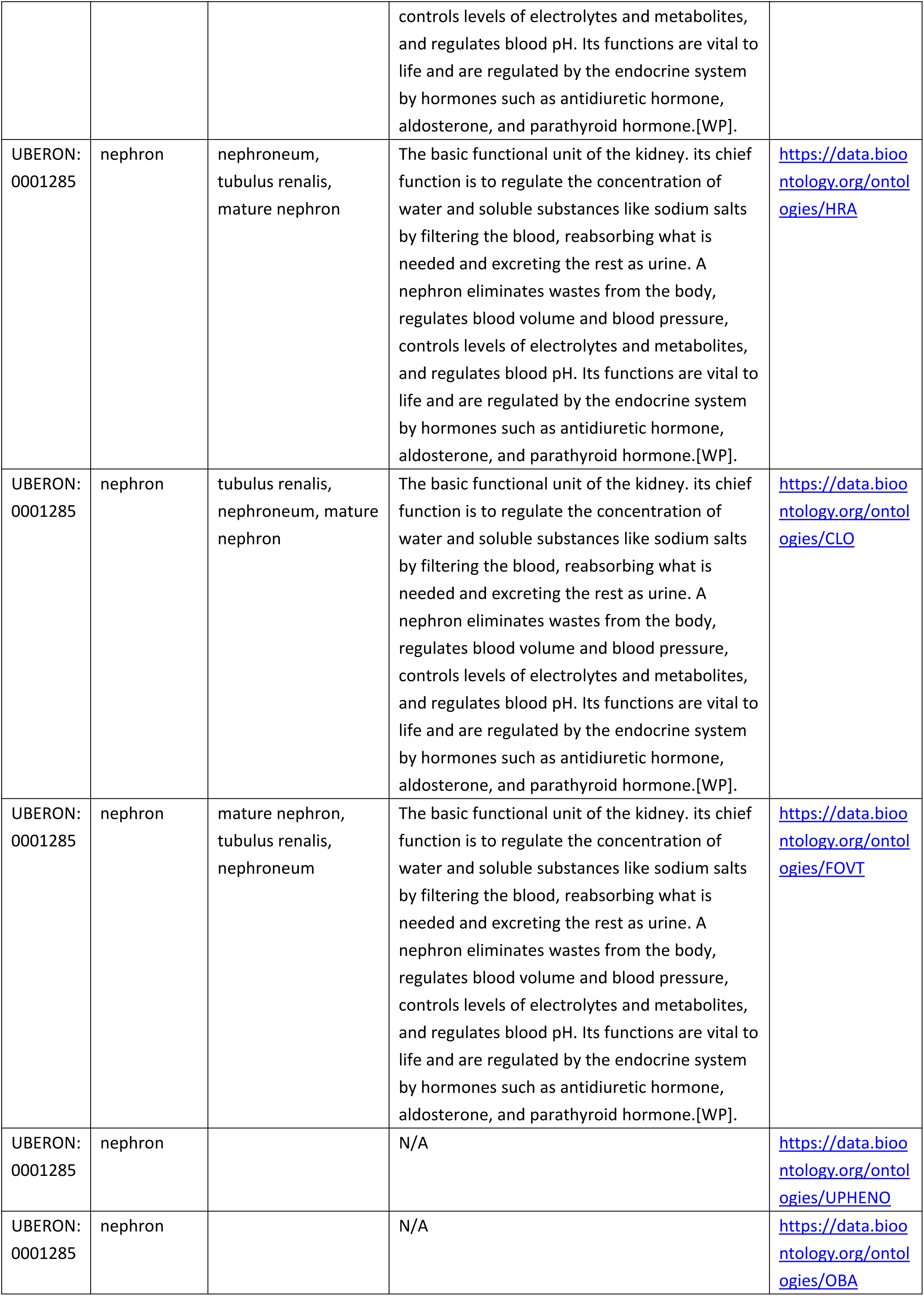
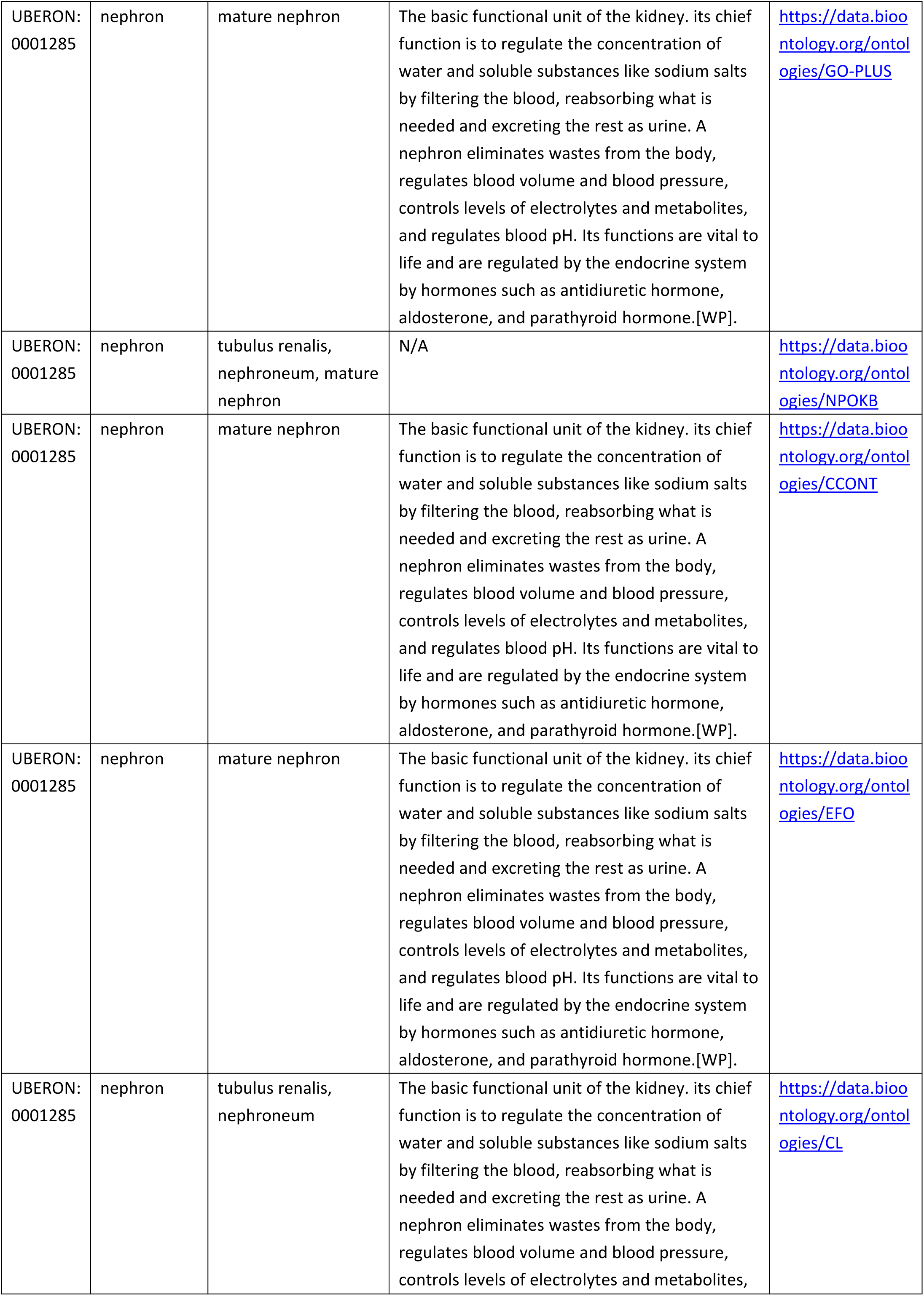
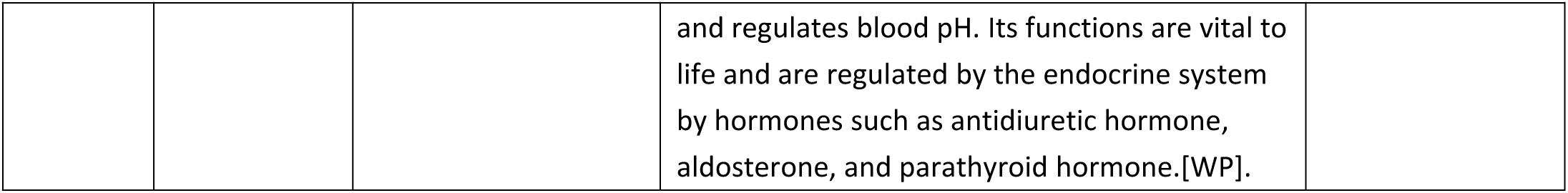
Descriptions of 22 FTUs from Bioportal.

**Supplementary Table 29.**
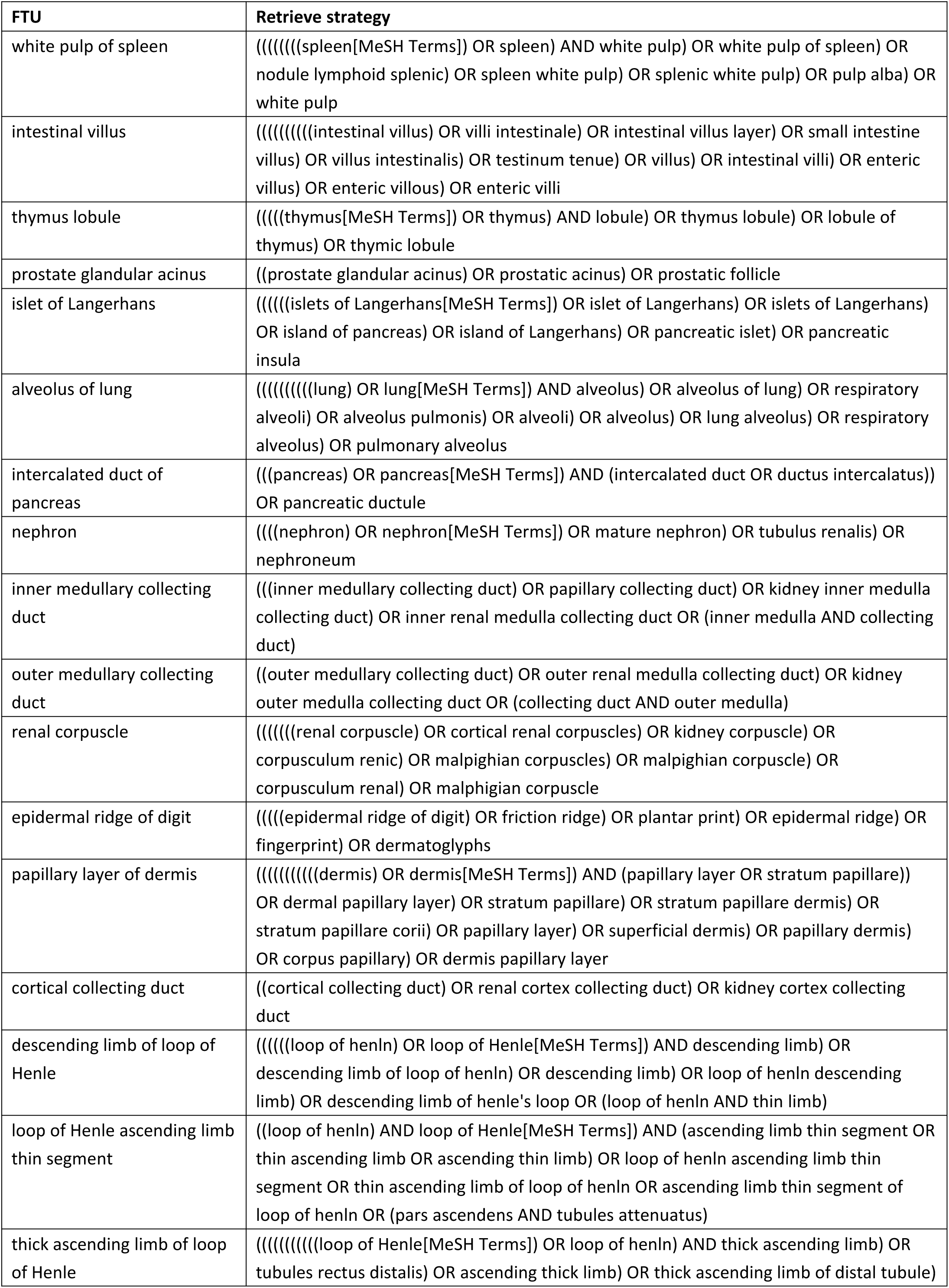
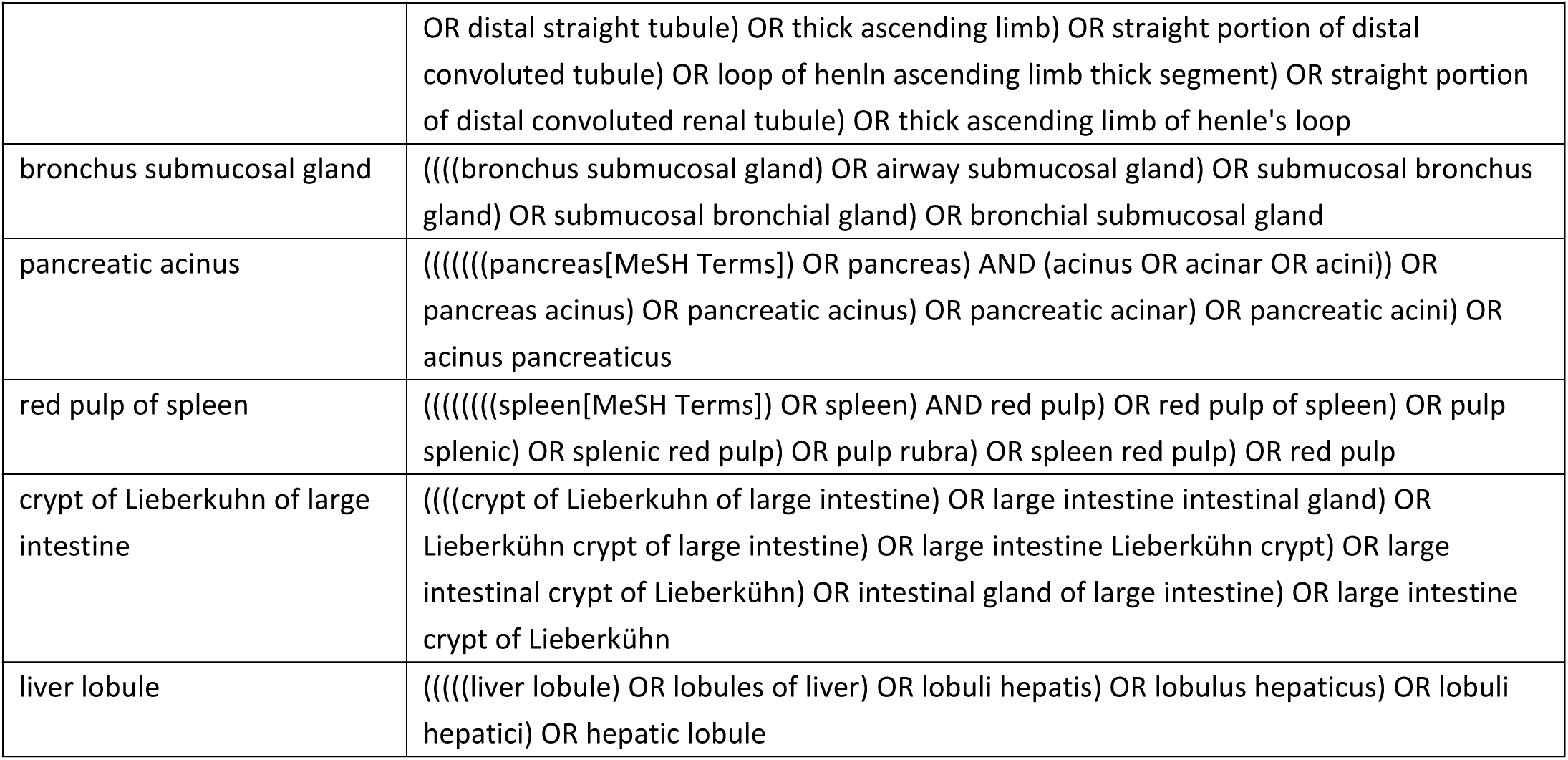
Retrieve strategies of 22 FTUs.

